# Melanopsin regulates axonal translation underlying retinohypothalamic circuit assembly

**DOI:** 10.64898/2026.04.21.716817

**Authors:** R. Rashmi, A.T. Belew, C. Zhang, C.J. Hernández, T. Alexander, R. Pomerat, L.P. Abadir, S.P. D’Souza, N.M. El-Sayed, C.M. Speer

## Abstract

Intrinsically photosensitive retinal ganglion cells (ipRGCs) influence visual system development via melanopsin before photoreceptor-mediated vision, but how melanopsin signaling contributes to ipRGC circuit assembly remains unknown. Here we show that melanopsin coordinates retinohypothalamic tract development by regulating local translation in developing ipRGC axons. Loss of melanopsin selectively disrupted local translation in axons without affecting somatic translation. The affected transcripts encoded cytoskeletal regulators, adhesion molecules, and trafficking proteins, and activity-dependent changes in translation were restricted to the period before eye-opening. Consistent with impaired axonal growth and synaptogenesis, *Opn4* knockout mice showed reduced ipsilateral suprachiasmatic nucleus innervation and fewer retinohypothalamic synapses, while nanoscale synaptic molecular organization and microglial engulfment were unaffected. Reduced visual drive in *Opn4* knockouts further altered developmental gene expression programs across the retina, suprachiasmatic nucleus, and lateral geniculate nucleus, with region-specific differences in expression timing. These findings identify melanopsin as a regulator of local axonal translation during early circuit development, linking sensory phototransduction to translational control mechanisms that guide retinohypothalamic tract assembly and postsynaptic target maturation.

## Introduction

Intrinsically photosensitive retinal ganglion cells (ipRGCs) provide the sole photic input to the rodent brain in the first postnatal week ^1–11^. During this period, ipRGCs influence multiple aspects of visual system maturation, establishing an early role in activity-dependent visual system plasticity before conventional photoreceptor-driven vision ^12–18^. Within this same developmental window, ipRGC axons form the retinohypothalamic tract (RHT), innervating the suprachiasmatic nucleus (SCN) ^15,19–21^. Melanopsin is required for light-induced *Fos* expression in the neonatal SCN, and dark-rearing reduces RHT synaptic input before eye-opening ^3,10,22^, suggesting that melanopsin-dependent activity influences RHT-SCN connectivity. However, the molecular mechanisms by which melanopsin-dependent activity regulates the development of ipRGC projections remain unknown.

Local translation in developing axons represents a candidate mechanism linking melanopsin-driven activity to circuit refinement. Retinal ganglion cells (RGCs) translate mRNAs locally in axons, with transcriptomes and translatomes that change during axon outgrowth and target innervation ^23,24^. Local translation contributes to axon branching, guidance, and synaptogenesis ^25^. Melanopsin activates mTOR signaling in ipRGCs ^26^, a potent regulator of local translation ^27–34^, raising the possibility that early light-driven activity shapes axon development through translational control. However, whether sensory activity regulates local translation in developing RGC axons remains unknown.

The RHT is formed primarily by M1-type ipRGCs, which are genetically accessible using Opn4-Cre knock-in/knock-out mice ^15,35–40^, enabling direct tests of melanopsin’s role in regulating axonal translation during circuit development. Here we examined how melanopsin influences ipRGC development across the eye-opening transition using complementary transcriptomic, translatomic, and structural approaches. We identified temporally restricted, compartment-specific effects of melanopsin loss on ribosome-associated mRNAs within ipRGC axon terminals. These subcellular changes were accompanied by selective defects in RHT targeting and synapse formation, while synaptic nanoscale molecular organization and microglial engulfment were normal.

Reduced functional drive from ipRGCs was further reflected in region-specific shifts in developmental gene expression timing across retinorecipient targets. Together, these findings link early melanopsin-dependent activity to axonal translational control mechanisms that support presynaptic retinohypothalamic circuit assembly and postsynaptic target maturation.

## Results

### Axon-TRAP reveals compartment-specific ipRGC translatomes

To examine how melanopsin affects mRNA translation, specifically within ipRGC somatic and axonal compartments, we developed a translating ribosome affinity purification (TRAP) strategy to separately profile ribosome-associated mRNAs from ipRGC somata in the retina and axon terminals in the SCN **(Figure 1A)**. We crossed Opn4-Cre mice with Rpl22-HA reporters to generate WT (no Cre), Opn4-Het, and Opn4-KO mice expressing hemagglutinin (HA)-tagged ribosomes selectively in ipRGCs, with intact or absent melanopsin signaling **(Figure 1A-C)**. WT samples lacking Cre served as immunoprecipitation (IP) controls, creating a stringent filter of non-specifically isolated mRNAs from all downstream analyses.

**Figure 1:**
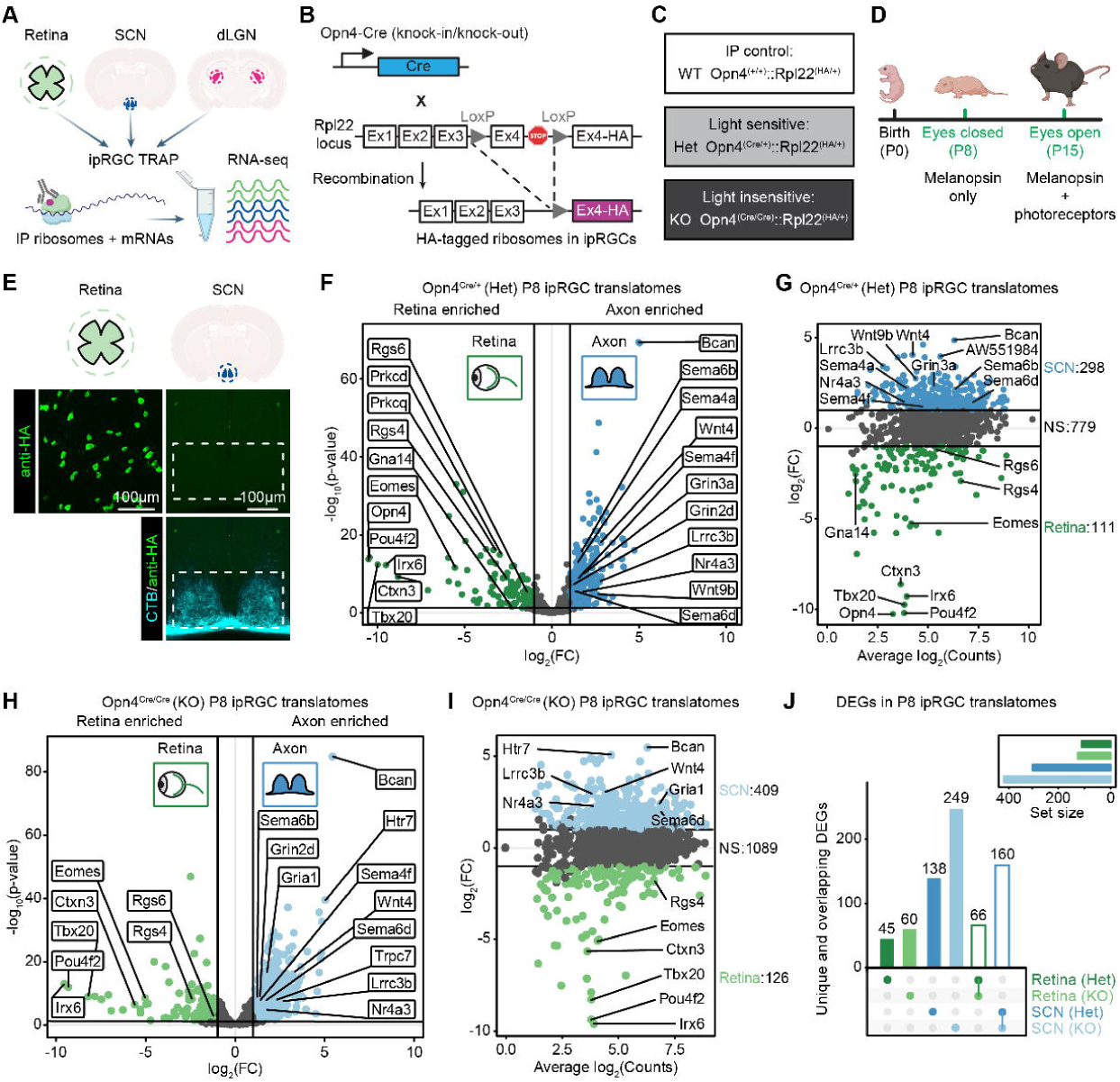
Axon-TRAP profiling of ipRGC subcompartment translatomes. **(A)** Experimental design showing ipRGC-TRAP approach. Retina, SCN, and dLGN tissues were collected for immunoprecipitation of HA-tagged ribosomes, followed by RNA-seq analysis in Opn4 WT, Het, and KO mice. **(B)** Knock-in/knock-out genetic strategy for expressing HA-tagged ribosomes in ipRGCs. **(C)** Three genotypes used: WT (Opn4^+/+^::Rpl22^HA/+)^ lacks Cre and serve as IP controls. Het (Opn4^Cre/+^::Rpl22^HA/+^) expresses tagged ribosomes and retains light sensitivity. KO (Opn4^Cre/Cre^::Rpl22^HA/+^) expresses tagged ribosomes but is light-insensitive. **(D)** Experimental timeline showing tissue collection at P8 (before eye-opening) and P15 (after eye-opening). **(E)** Confocal microscopy showing anti-HA immunolabeling in retina (top left panel) and SCN (top right panel) with CTB labeling of RHT axons overlaid (cyan, bottom right panel). **(F)** Differential expression analysis of P8 heterozygous ipRGC translatomes showing retina-enriched (green) and axon-enriched (blue) transcripts with labeled genes. **(G)** MA plot showing fold change versus expression for P8 Het ipRGC translatomes with SCN-enriched (blue, n=298) and retina-enriched (green, n=111) genes. **(H)** Volcano plot of P8 KO ipRGC translatomes showing retina-enriched (green) and axon-enriched (blue) transcripts with labeled genes. **(I)** MA plot showing fold change versus expression for P8 KO ipRGC translatomes with SCN-enriched (blue, n=409) and retina-enriched (green, n=126) genes. **(J)** Upset plot comparing differentially expressed genes (DEGs) across genotypes and compartments showing unique and overlapping gene sets.

We validated HA expression and specificity by crossing our ipRGC-TRAP line to Ai9 reporters and immunostaining for HA and tdTomato **(Figure S1A)**. In the retina, tdTomato-positive cells (tdT+) were observed at densities consistent with previous reports ^39,41^ **(Figure S1B)**, with ∼72% of tdT cells also expressing HA. In the SCN, we observed tdT+ ipRGC axons and no HA-labeled cell bodies **(Figure S1A)**. In contrast, the dLGN contained HA-positive somata, indicating ectopic expression that precluded axon-specific TRAP analysis in the thalamus **(Figure S1A)**. Western blotting and TapeStation analysis confirmed successful immunoprecipitation and high RNA integrity (RIN >7.0) in Cre-expressing samples **(Figure S1C-F)**.

To investigate melanopsin-dependent local translation, we collected retina and SCN tissue at P8 (before eye-opening) and P15 (after eye-opening), capturing the transition from melanopsin-only to rod/cone-driven photic input **(Figure 1D)**. At P8, background-corrected Het samples showed distinct ribosome-associated mRNA populations in retina versus SCN **(Figure 1E-G)**. Somatic translatomes were enriched for transcription factors and G protein-coupled receptors expressed in developing ipRGCs ^42^, while axonal translatomes were enriched for genes involved in axon growth and synaptic function. Similar compartmental segregation was observed in KO mice **(Figure 1H-J)**.

### Melanopsin selectively regulates axonal translation before eye-opening

We next compared Het and KO translatomes to identify melanopsin-dependent effects **(Figure 2A)**. At P8, retinal translatomes were nearly identical between genotypes apart from *Opn4* itself, indicating that melanopsin loss does not substantially alter ribosome-mRNA association in ipRGC cell bodies or dendrites **(Figure 2B)**. In contrast, the P8 axonal translatome showed striking melanopsin dependence. Dozens of transcripts were enriched in Het axons but depleted in KO axons **(Figure 2C)**. Gene ontology analysis highlighted processes related to axon growth, cargo trafficking, and synapse development **(Figure 2D and S2)**. Depleted transcripts encoded cytoskeletal regulators (Map1b, Map4, Dpysl2/Crmp2, Dpysl3/Crmp4, Fign, Gsk3b, Actb, Cfl2, Marcks), cell adhesion molecules (Ncam1, Nlgn3, Cadm1), trafficking proteins (Sec22b, Scamp1), and growth factors (Nrn1, Fgf9, Wnt9b). These findings indicate that melanopsin promotes ribosomal engagement with growth-associated mRNAs in ipRGC axons prior to eye-opening.

**Figure 2:**
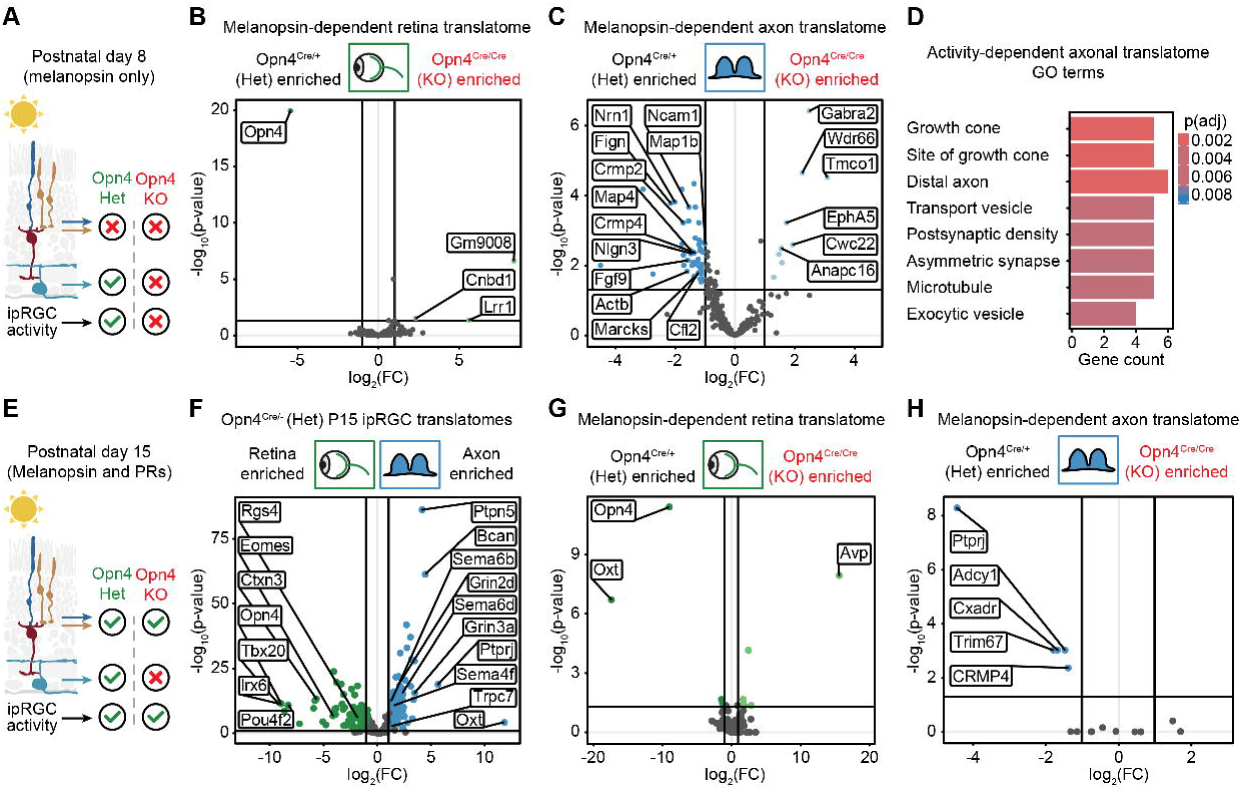
Melanopsin regulates the axonal translatome before eye-opening. **(A)** Schematic of visual input at P8 when melanopsin is the sole source of light-driven activity. **(B)** Volcano plot of melanopsin-dependent retina translatomes at P8 showing Het enriched versus KO enriched genes. **(C)** Volcano plot of melanopsin-dependent axon translatomes in the SCN at P8 showing Het enriched versus KO enriched genes. **(D)** Gene Ontology enrichment analysis for activity-dependent axonal translatome showing significant GO terms. **(E)** Schematic of visual input at P15 when rods and cones are active. **(F)** Volcano plot of P15 heterozygous ipRGC translatomes comparing retina-enriched (green) and axon-enriched (blue) transcripts. **(G)** Volcano plot of melanopsin-dependent retina translatomes at P15 showing Het enriched versus KO enriched genes. **(H)** Volcano plot of melanopsin-dependent axon translatomes in the SCN at P15 showing Het enriched versus KO enriched genes.

We next determined whether axon-specific regulation persists after eye-opening at P15, when rod and cone photoreceptor inputs provide visual drive to ipRGCs independent of melanopsin **(Figure 2E)**. Similar to P8, the P15 Het retinal translatome was enriched for known ipRGC transcription factors while the axonal translatome contained transcripts associated with synaptic function (**Figure 2F, S2, and S3**). However, retinal and axonal translatomes were similar between Het and KO mice **(Figure 2G-H)**, suggesting that photoreceptor-driven activity compensates for melanopsin loss at later stages.

Developmental comparisons further supported this conclusion. Retinal translatomes changed from P8 to P15 in both genotypes, reflecting ongoing retinal maturation **(Figure S4A-B)**. In the SCN, Het axonal translatomes remained stable across ages **(Figure S4C)**, whereas KO axons exhibited a developmental shift characterized by transcript enrichment at P8 relative to P15 **(Figure S4D)**. Together, these data reveal an unexpected specificity in melanopsin-dependent translational regulation in ipRGCs.

Before eye-opening, melanopsin is dispensable for translation in ipRGC cell bodies and dendrites but required for maintaining ribosomal engagement with growth-associated transcripts in RHT axon terminals. After eye-opening, melanopsin loss produces minimal translational changes across ipRGC subcompartments, indicating a temporally restricted window of activity-dependent local translation coinciding with early RHT circuit assembly.

Because the dLGN showed ectopic HA expression in postsynaptic neurons **(Figure S1)**, we were unable to isolate non-M1-type ipRGC axonal translatomes in the thalamus.

Nevertheless, TRAP analysis of the dLGN revealed a developmental shift from P8 to P15 in both Het and KO mice **(Figure S5A-B)**, corresponding to when RGC spike output transitions from spontaneous retinal waves to visually-driven activity ^43^.

Comparing Het versus KO translatomes at each age revealed melanopsin-dependent enrichment of a subset of transcripts in KO dLGNs **(Figure S5C-D)**. However, the number of affected transcripts was small compared to developmental changes in either genotype, suggesting a minor role for melanopsin in regulating mRNA translation in postsynaptic dLGN neurons.

### Melanopsin is required for ipsilateral RHT innervation

The depletion of growth-associated transcripts in P8 KO axons suggests anatomical defects in RHT development. To test this, we performed dual-eye fluorescent anterograde tracing by injecting cholera toxin subunit B (CTB) conjugated to spectrally distinct fluorophores into the left and right eyes of WT and Opn4-KO mice at P7. Brains were collected at P8 and coronal sections through the SCN [Bregma -0.46 to -0.58 ^44^] were imaged to analyze the distribution of eye-specific RHT projections. To quantify genotype differences, we cropped individual confocal SCN images dorsal to the optic chiasm, centering them along the midline, and computed the normalized average image intensity across biological replicates (N=10 per genotype; see Materials and Methods).

In WT mice, CTB mean intensity was symmetric across the two SCN hemispheres, with robust contralateral and ipsilateral innervation from each eye **(Figure 3A)**. In Opn4-KO mice, RHT axons innervated the contralateral SCN and appeared enriched in the outer core relative to WT, though this difference was not significant **(Figure 3B-C)**. In contrast, ipsilateral innervation was significantly reduced in Opn4-KO mice compared to controls. This defect was bilateral, with both eyes showing reduced ipsilateral projections (**Figure 3C**). Subtracting Opn4-KO and WT average images confirmed the two-dimensional spatial pattern, with relative axon enrichment in the contralateral outer core and depletion of ipsilateral input in KO mice compared to controls. Consistent with our TRAP analysis showing depletion of growth-associated transcripts, these results demonstrate that melanopsin is necessary for the normal development of ipsilateral ipRGC projections to the SCN.

**Figure 3:**
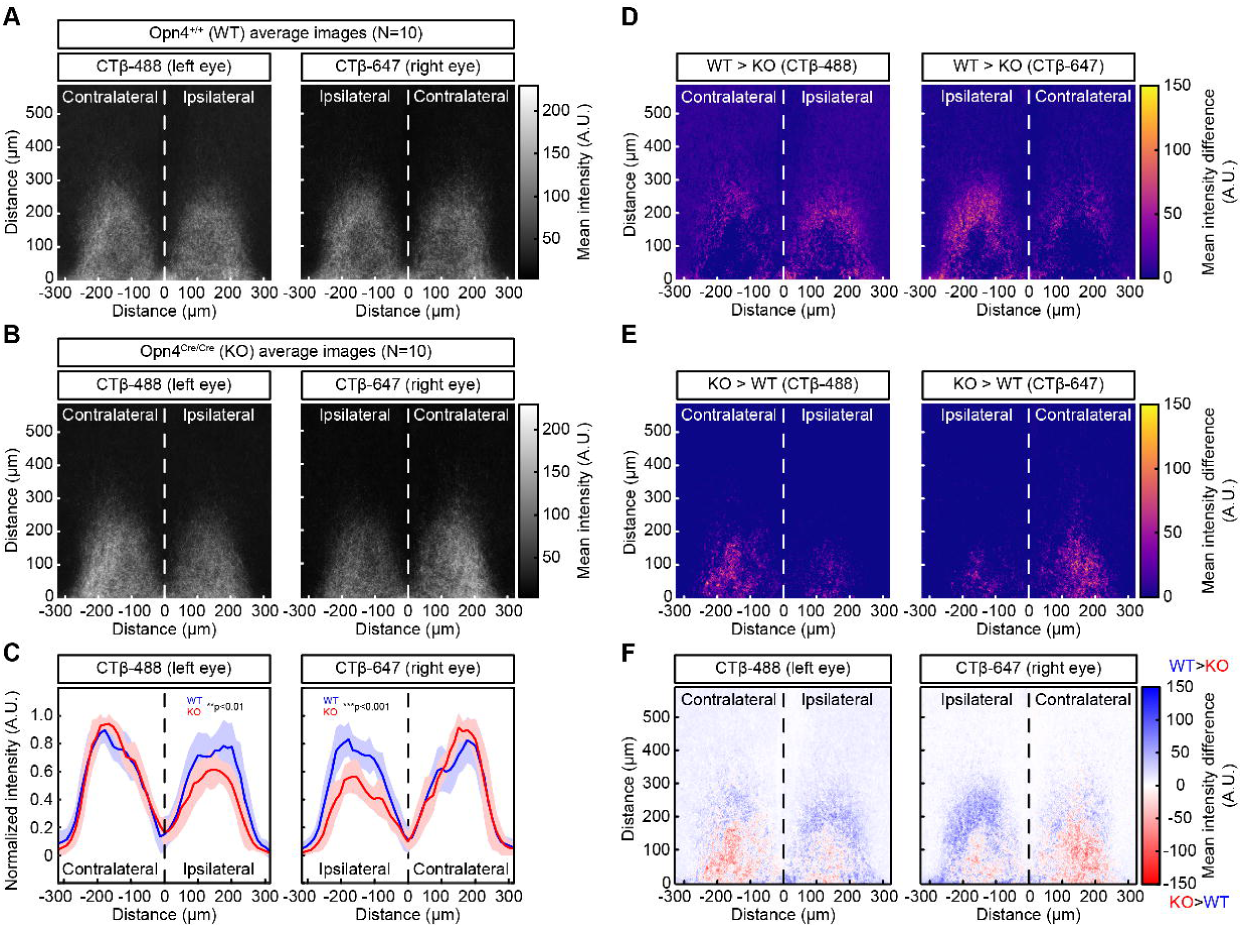
Melanopsin is required for innervation of the ipsilateral SCN. **(A)** RHT innervation in WT mice at P8 showing CTB-Alexa Fluor 488 (left eye, green) and CTB-Alexa Fluor 647 (right eye, magenta) projections. Confocal images were aligned and averaged to show mean intensity (N=10 biological replicates). **(B)** Average intensity images P8 Opn4-KO mice (N=10 biological replicates). Average images are shown on a global intensity scale computed across both eye-specific channels and genotypes to enable direct visual comparison. **(C)** Quantification of normalized CTB intensity across the mediolateral extent of the contralateral and ipsilateral SCN for both eye-specific channels. KO mice show significant reduction in ipsilateral innervation from the left and right eyes. Solid lines and shaded regions show the mean ± S.D. computed from 20-pixel bins (∼12.6 µm). Left eye ipsilateral SCN (0 : 300) — WT mean ± S.D. (0.48 ± 0.08), KO mean ± S.D. (0.35 ± 0.07), **P<0.01, Cohen’s d 1.6; Right eye ipsilateral SCN (0 : -300) — WT mean ± S.D. (0.47 ± 0.07), KO mean ± S.D. (0.31 ± 0.07), ***p<0.001, Cohen’s d 2.5; Mann-Whitney U test. **(D-E)** Spatial difference heatmaps for left eye (left panels) and right eye (right panels) RHT inputs showing SCN regions where WT signal intensity exceeds KO intensity (D) or vice versa (E). **(F)** Combined intensity difference map where blue colors indicate WT > KO (positive values), white indicates equal intensity (0), and red colors indicate KO > WT (negative values). Dashed lines show SCN midline in all figure panels.

### Melanopsin regulates synapse number but not nanoscale organization

Given the reduction in ipsilateral SCN innervation and the axonal depletion of synapse-associated transcripts in Opn4-KO mice, we next asked whether melanopsin regulates ipRGC synapse development in the SCN. We performed volumetric super-resolution microscopy using a custom STORM pipeline (**Figure 4A**) ^45^. We injected both eyes with CTB-Alexa Fluor 488 at P7 and collected SCN tissue from Opn4-Het and Opn4-KO mice at P8 (N=6 biological replicates per genotype) (**Figure 4B**). We immunostained the SCN for Bassoon (active zones), Homer1/2/3 (postsynaptic scaffolds), and VGluT2 (vesicle pools in RHT synapses) ^21,46,47^. RHT synapses were identified by spatial apposition of paired Bassoon and Homer1/2/3 signals with an adjacent VGluT2 cluster (**Figure 4C**).

**Figure 4:**
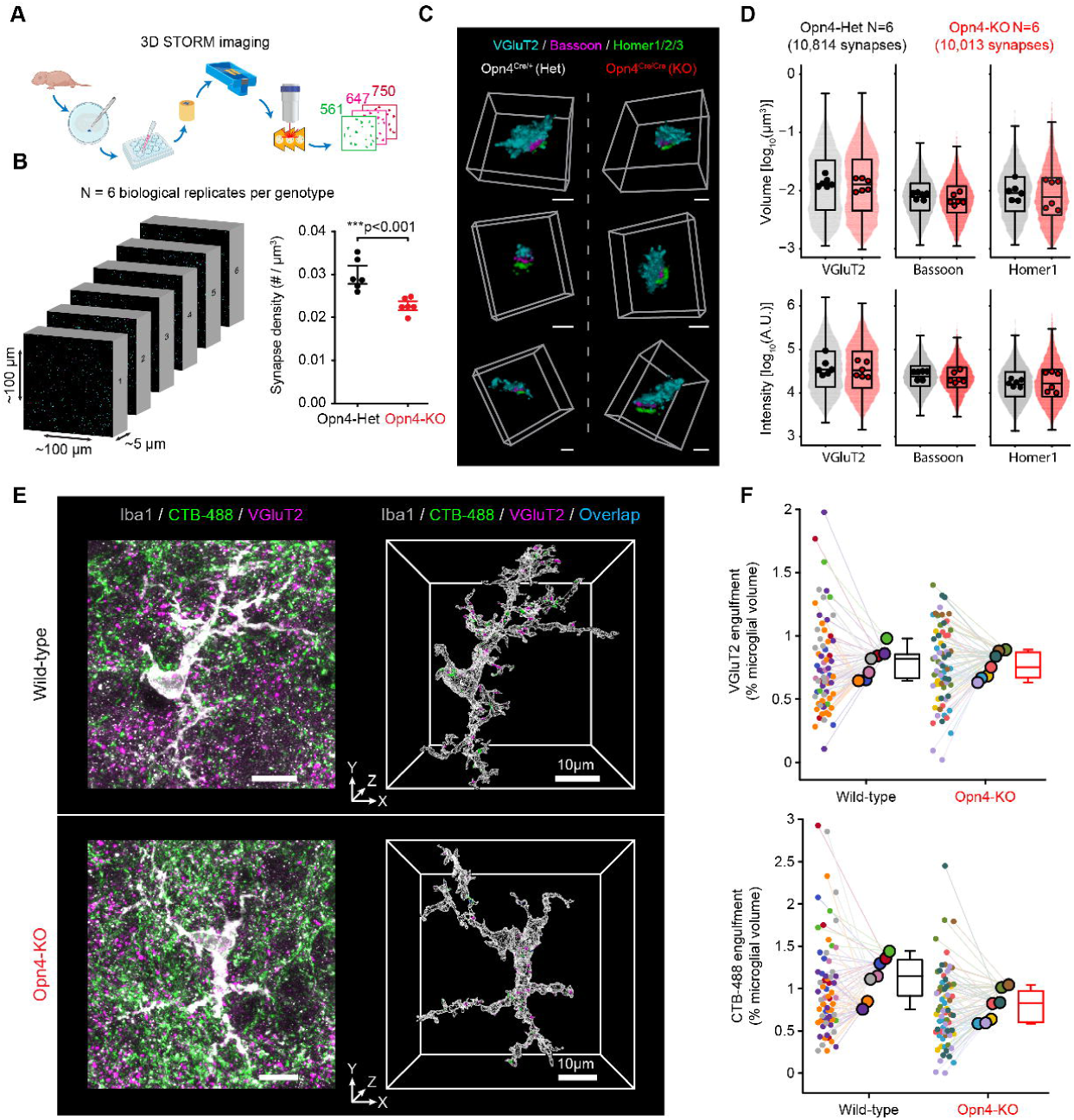
Melanopsin regulates RHT synaptogenesis prior to eye-opening. **(A)** Experimental workflow showing volumetric STORM imaging approach by ultrasectioning and serial reconstruction. **(B)** STORM analysis was performed in N=6 biological replicates per genotype (∼50K µm^3^ volume each). Quantification shows significantly reduced synapse density Opn4-KO mice (red dots) compared to Opn4-Het controls (black dots). Het mean ± S.D. (0.03 ± 0.004); KO mean ± S.D. (0.023 ± 0.002); Cohen’s d 2.5; ***p<0.001 student’s t-test. **(C)** Representative 3D reconstructions of individual synapses immunolabeled for VGluT2 (cyan, presynaptic marker), Bassoon (magenta, active zone protein), and Homer1/2/3 (green, postsynaptic density marker) in Opn4^Cre/+^ (Het, left panels) and Opn4^Cre/Cre^ (KO, right panels). Scale bars = 1 μm. **(D)** Quantitative analysis of synaptic protein distributions showing volume (top) and intensity (bottom) measurements for VGluT2, Bassoon, and Homer1 across N=6 Het (10,814 synapses, black) and N=6 KO (10,013 synapses, red) mice. Box-and-whisker plots with violin overlays show median, quartiles, and distribution within 1.5x the interquartile range. No significant differences observed in volume or intensity measurements between genotypes. **(E)** Engulfment of VGluT2 (magenta) and CTB-Alexa Fluor 488 puncta (green) by microglia (grey) in the SCN. Blue puncta in 3D renderings represent colocalized VGluT2/CTB punctae. Images show maximum intensity Z-projections (left) and 3D renderings (right) of wild-type (14.31 μm Z-stack) and Opn4-KO (12.96 μm Z-stack) tissue. **(F)** Superplots showing VGluT2 (top) and CTB-Alexa Fluor 488 (bottom) engulfment as a percentage of microglial volume. Small circles represent measurements from individual microglia, colored by source animal. N = 7 animals per group; Wild-type: 66 cells; Opn4-KO: 69 cells. Large circles represent individual animal means, connected to their respective cells by thin lines. Box-and-whisker plots show median, quartiles, and distribution within 1.5x the interquartile range. Statistical comparison was performed using a linear mixed model with genotype as a fixed effect and animal number as a random effect.

We cut coronal sections through the center of the SCN, as in our RHT tracing experiments, and collected individual STORM image blocks (∼50,000 µm^3^ each) containing the SCN core. To assess the effect of melanopsin loss on RHT synaptogenesis, we measured synapse density within each block by dividing the number of paired VGluT2 clusters by the block volume, accounting for edge artifacts. KO mice had significantly lower synapse density than Het mice (**Figure 4B**), consistent with both the molecular depletion of synapse-associated transcripts and anatomical reduction in ipsilateral innervation. However, the synapses that did form appeared structurally normal. Across more than 10,000 synapses per genotype, we found no differences in the mean volume for VGluT2, Bassoon, or Homer1/2/3 clusters (**Figure 4D, upper panels**). Similarly, we found no difference in intensity for any of these markers, a proxy for relative protein abundance (**Figure 4D, lower panels**). Thus, melanopsin is necessary for RHT synapse formation in the SCN before eye-opening but does not affect vesicle pool size or the volume and intensity of Bassoon and Homer1/2/3 signals in individual contacts.

To investigate whether the reduction in synapse density was caused by increased microglial pruning, we performed an engulfment assay to measure VGluT2 and CTB-Alexa Fluor 488 signals within microglia in the SCN **(Figure 4E)**. The engulfment percentage for both markers was similar between genotypes, indicating that melanopsin loss does not increase microglial engulfment leading to reduced synapse density **(Figure 4F)**. This suggests the reduced synapse density reflects impaired synapse formation rather than enhanced synapse elimination.

### Melanopsin loss alters developmental gene expression across visual structures

In Opn4-KO mice, reduced RHT axon innervation and synapse density diminish ipRGC input to the SCN during a critical period of circuit maturation. These structural deficits raise the question of whether melanopsin loss also reshapes the gene expression landscape of retinorecipient targets. To assess this, we performed bulk RNA sequencing on retina, SCN, and dLGN from wild-type (WT), Het (Opn4^Cre/+^), and KO (Opn4^Cre/Cre^) mice at P8 **(Figure 5A)**. Principal component analysis revealed clear clustering by tissue type **(Figure 5B)**, with tissue-specific differential expression profiles **(Figure 5C)**. Gene ontology analysis identified changes in transcripts associated with neurogenesis, synaptic transmission, mitochondrial function, and extracellular matrix organization **(Figure S6)**. The largest transcriptomic changes were observed between WT and Opn4-KO mice, while heterozygotes exhibited an intermediate profile, suggesting a graded effect of melanopsin signaling **(Figure S7**).

**Figure 5:**
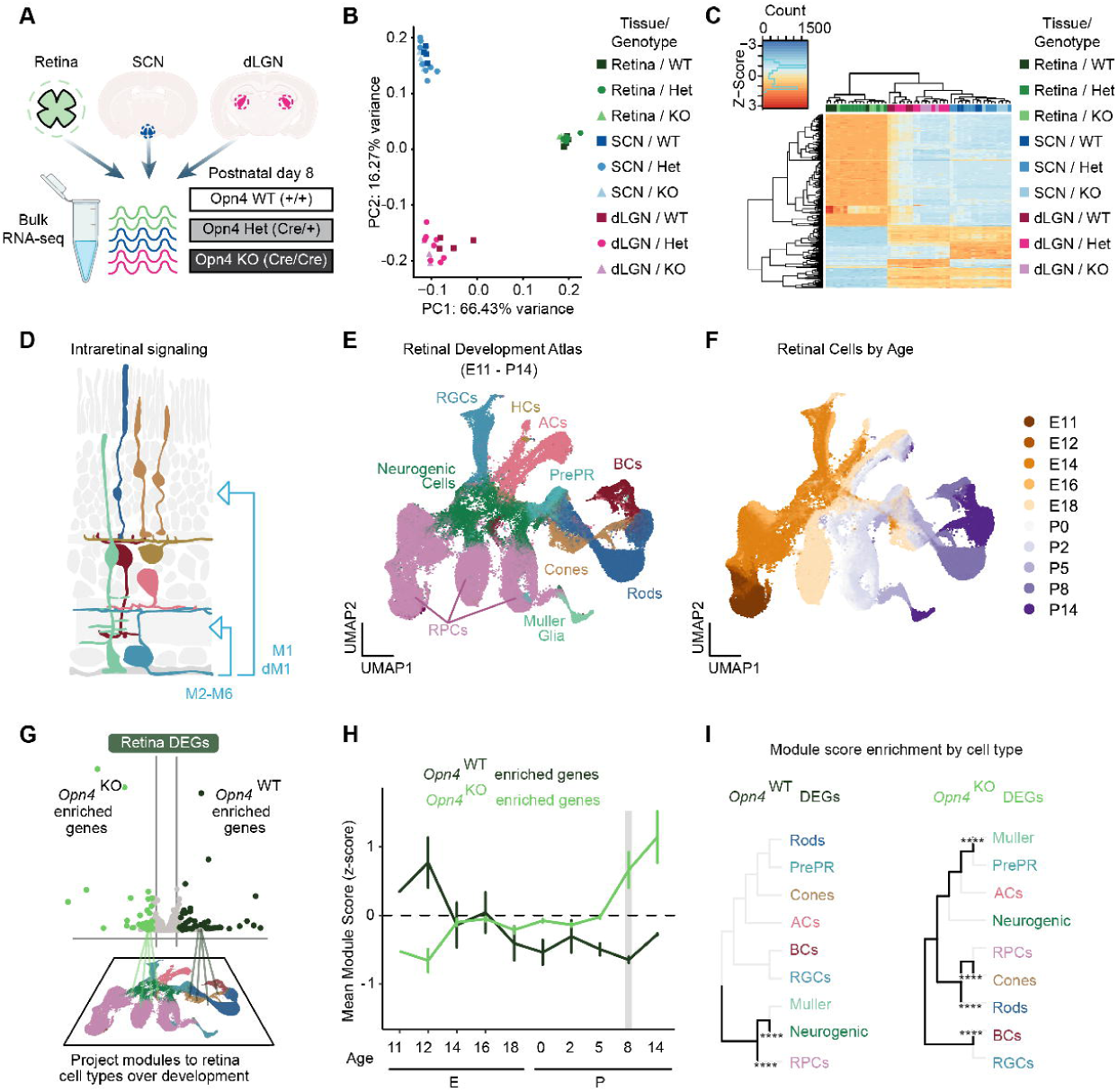
Melanopsin regulates gene expression in the retina and central targets before eye-opening. **(A)** Schematic of bulk RNA-seq analysis of retina, suprachiasmatic nucleus (SCN), and dorsal lateral geniculate nucleus (dLGN) from Opn4 wild-type (WT), Heterozygous (Opn4^Cre/+^), and knockout mice (Opn4^Cre/Cre^) at P8. **(B)** Principal component analysis shows discrete transcriptome clustering by tissue type. **(C)** Hierarchical clustering heatmap of top differentially expressed genes between melanopsin WT, Het, and KO mice at P8 (Z-score normalized). **(D)** Schematic of ipRGC communication within the retina. **(E)** UMAP projection of Clark et al., 2019 retinal development atlas color coded by major cell class. **(F)** UMAP visualization of retinal cells colored by developmental age. **(G)** Module mapping of Opn4 WT-enriched (dark green) and Opn4 KO-enriched (light green) genes to the developmental atlas. **(H)** Mean module scores averaged across developmental age (E = embryonic, P = postnatal). **(I)** Cell type-specific enrichment analysis showing module scores for Opn4-WT and Opn4-KO differentially expressed genes across retinal cell types, with hierarchical clustering indicating cell type relationships. ****p < 0.0001, Mann-Whitney U test with Benjamini-Hochberg correction comparing mean module score between cell class and label-shuffled dataset.

To identify affected retinal cell types **(Figure 5D)**, we mapped differentially-expressed genes onto a single-cell RNA-seq atlas of retinal development ^48^ **(Figure 5E-F)**. Module mapping of P8 bulk RNA-seq genes showed that most genes were identified in the atlas, with WT-enriched and KO-enriched gene modules showing opposing developmental trajectories **(Figure 5G-H)**. WT modules were associated with retinal progenitors and neurogenic populations, while KO modules correlated with late-born cell types including rods, cones, bipolar cells, and Müller glia **(Figure 5I)**.

Comparison with a previous transcriptomic analysis of P4 Opn4-KO retina revealed minimal gene overlap, with only four shared transcripts including *Opn4* ^18^ **(Figure S8A)**. Despite this, both studies showed similar temporal enrichment patterns **(Figure S8B-C)** and nearly identical cell-type associations **(Figure S8D)**, suggesting that melanopsin influences developmental timing of retinal gene expression rather than regulating a fixed gene set.

We applied the same analysis to the SCN using a hypothalamic single-cell atlas spanning E15 to P65 ^49^ **(Figure 6A-F).** Focusing on SCN-annotated (i-S1-7) and SCN-related (i-S8) populations **(Figure 6B-C)**, module mapping again revealed opposing genotype-dependent trajectories **(Figure 6D-F)**. WT-enriched genes predominated at earlier developmental stages, whereas KO-enriched genes were associated with more mature profiles **(Figure 6E)**. Cell-type analysis linked WT modules to i-S1 (SCN-Avp/C1ql3) and i-S8 (BNST-Lhx1/Satb2) neurons, while KO modules were enriched in i-S1 and i-S4 (SCN-Vip/C1ql3) populations **(Figure 6F)**.

**Figure 6:**
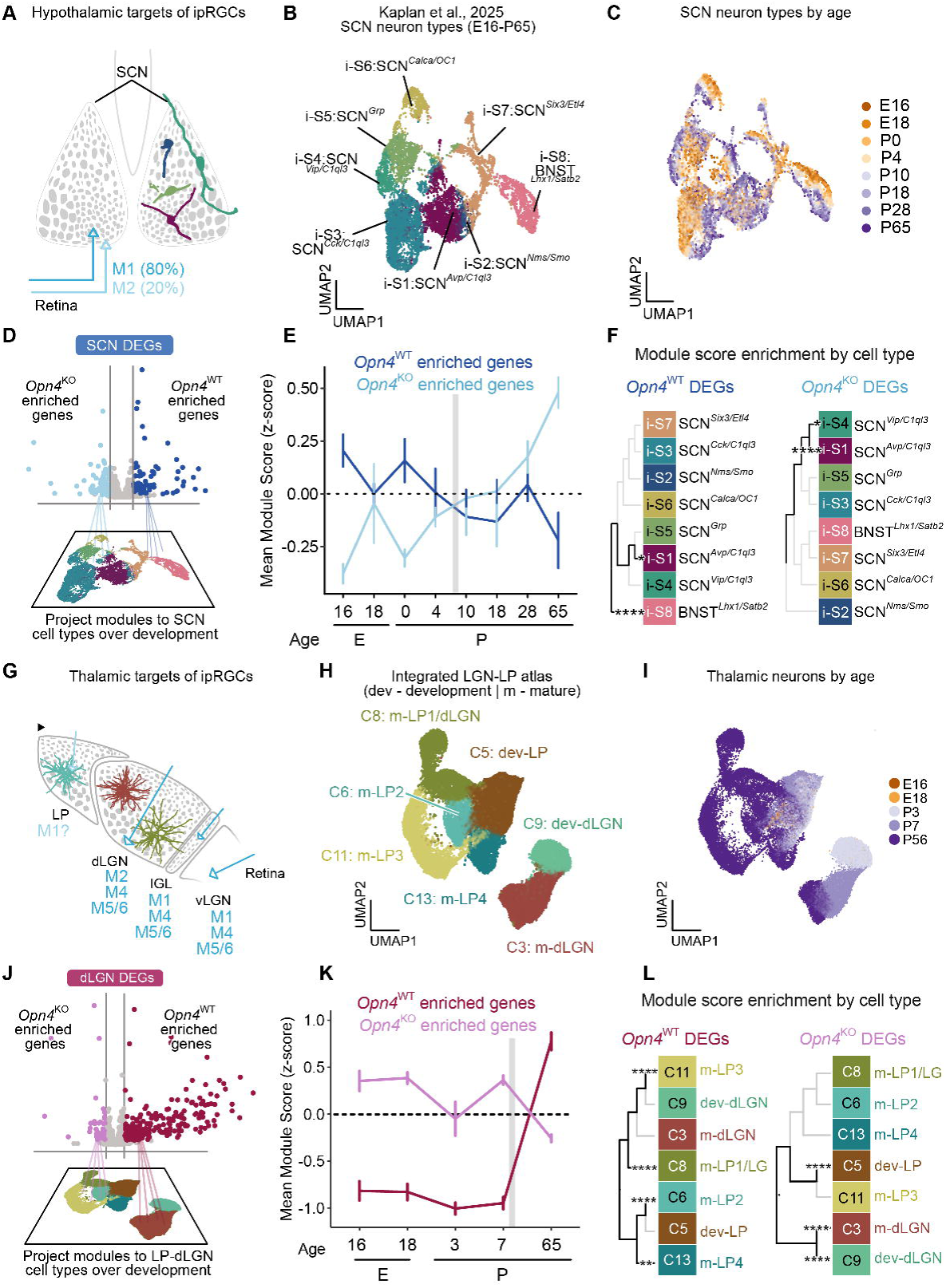
Melanopsin-dependent gene expression in hypothalamic and thalamic targets receiving ipRGC input. **(A)** Schematic of ipRGC subtype connectivity with the SCN. **(B)** UMAP projection of SCN neuron types from Kaplan et al. 2025 spanning embryonic day 16 to postnatal day 65 (E16-P65), showing distinct subpopulations with molecular markers indicated. **(C)** UMAP visualization of SCN neurons colored by developmental age from E16 to P65. **(D)** Module mapping of Opn4 KO-enriched genes (light blue) and Opn4 WT-enriched genes (dark blue) to the developmental SCN atlas. **(E)** Temporal dynamics of mean module scores for Opn4 WT-enriched (dark blue) and Opn4 KO-enriched (light blue) gene sets across embryonic (E) and postnatal development (P) in the SCN. **(F)** Cell type-specific enrichment analysis showing module scores for Opn4-WT and Opn4-KO enriched genes across SCN neuronal subtypes. *p < 0.05, ****p < 0.0001, Mann-Whitney U test with Benjamini-Hochberg correction. **(G)** Schematic of thalamic targets of ipRGCs showing projections to the dorsal lateral geniculate nucleus (dLGN), intergeniculate leaflet (IGL), and ventral LGN (vLGN). **(H)** UMAP projection of an integrated LGN-LP embedding of Lo Giudice et al., 2024 (development) and Allen Brain Cell Atlas 2024 (mature) data identifying major cell clusters. **(I)** UMAP visualization of thalamic neurons colored by developmental age from E16 to P56. **(J)** Module mapping to the integrated dLGN atlas. **(K)** Temporal dynamics of mean module scores for Opn4 WT-enriched (magenta) and Opn4 KO-enriched (light magenta) gene sets across embryonic and postnatal development in the dLGN. **(L)** Cell type-specific enrichment analysis showing module scores for Opn4-WT and Opn4-KO enriched genes across dLGN neuronal subtypes. **p < 0.01, ****p < 0.0001, Mann-Whitney U test with Benjamini-Hochberg correction.

To determine whether these cell types receive ipRGC input, we performed label transfer using a published SCN scRNA-seq dataset annotated for photic responsiveness ^50^.

After reidentifying SCN neurons based on Six6 and Lhx1 expression and regressing out rhythmic genes **(Figure S9A-C)**, we integrated the Wen and Kaplan datasets using CCA-based alignment restricted to adult cell types **(Figure S9D-E)**. This analysis confirmed that affected populations correspond to light-responsive SCN neurons receiving direct ipRGC input **(Figure S9F)**. Specifically, i-S1 neurons mapped primarily to C2 (Avp^+^Nms^+^C1ql3^+^) cells of the Wen dataset, i-S4 neurons corresponded to C3 (Vip^+^C1ql3^+^) cells, consistent with shell and core SCN populations ^50^.

We next examined the visual thalamus **(Figure 6G-L)** using an integrated atlas combining developmental single-nucleus RNA-seq ^51^ with adult scRNA-seq data from the Allen Brain Cell Atlas ^52^ **(Figure 6H-I)**. In contrast to retina and SCN, WT-enriched genes in the dLGN were associated with mature cell states, whereas KO-enriched correlated with immature profiles **(Figure 6K-L)**, indicating region-specific effects of melanopsin on developmental gene expression trajectories.

## Discussion

Melanopsin-expressing ipRGCs shape visual system development before eye-opening, yet whether melanopsin regulates the formation of ipRGC circuits themselves has remained unresolved. Here, we find that melanopsin coordinates multiple aspects of retinohypothalamic tract (RHT) development during the first postnatal week in mice.

Using axon-TRAP, we show that melanopsin loss selectively reduces ribosome association with transcripts encoding axon growth and synaptic proteins in ipRGC axon terminals, without affecting the somatic translatome. This compartment-specific regulation is restricted to the pre-visual period and disappears after eye-opening when rod and cone inputs become functional. Structurally, Opn4-KO mice show reduced ipsilateral RHT innervation and fewer RHT synapses in the SCN, with no increase in microglial engulfment. RHT synapses that do form display normal nanoscale organization, suggesting that melanopsin-driven activity influences synaptogenesis rather than the molecular architecture of individual contacts. Together, these findings identify melanopsin as a regulator of axonal ribosome occupancy during a critical window of circadian circuit development.

### Melanopsin regulates a cytoskeletal developmental program

Melanopsin regulates the developing ipRGC axonal translatome with striking spatial and temporal specificity. Ribosome-associated transcripts are altered in axons but not in cell bodies or dendrites, and these effects are confined to P8, prior to eye-opening. The affected transcripts include a functionally coherent set of cytoskeletal regulators, implicating melanopsin-dependent local translation in axon growth, branching, and structural remodeling during early postnatal development.

The microtubule regulatory network depleted in Opn4-KO axons includes stabilizing proteins (Map1b, Crmp2, Crmp4) as well as their negative regulator (Gsk3b), suggesting melanopsin-dependent regulation of microtubule dynamics during axon growth and branching. Map1b, Crmp2, and Crmp4 promote microtubule assembly underlying axon elongation and branching ^53–59^. Phosphorylation of Map1b and Crmp2 by Gsk3b modulates their interactions with microtubules, linking extracellular signaling and cytoskeletal remodeling in developing RGCs ^58,60–65^. Fidgetin, which severs microtubules to regulate axon growth and targeting ^66,67^, may provide additional spatial control over microtubule remodeling within ipRGC axons.

Complementing these microtubule regulators, transcripts encoding the actin regulatory proteins Actb, Cfl2, and Marcks, were also depleted in Opn4-KO axons, implicating impaired actin dynamics underlying growth cone motility and axon branching. β-actin is locally translated at axonal hotspots to drive growth cone turning and branch formation in RGCs ^68–70^, while the actin severing protein cofilin 2 is also locally synthesized in RGC growth cones in response to extracellular cues ^71^. Marcks further contributes to axon development by mediating vesicular cargo delivery to the membrane ^72,73^.

This cytoskeletal machinery is further influenced by growth promoting signals and adhesion molecules that were also enriched in Het axons relative to KOs, including Ncam1, Nlgn3, Wnt9b, Fgf9, and Nrn1. Interestingly, Nrn1 expression in the brain is light-dependent, and RGC depolarization promotes calcium-dependent insertion of Nrn1 into the axonal membrane ^74,75^. Our findings suggest that Nrn1 may be locally translated in an activity-dependent manner, linking melanopsin signaling to axon growth and synapse formation. Together, these data support a model in which melanopsin-dependent local translation regulates an integrated developmental program for ipRGC axon growth and target innervation during early postnatal development.

### Temporal specificity of melanopsin-dependent translation

The presence of melanopsin-dependent differences in axonal translatomes at P8, but not P15, coincides with the pre-visual period when melanopsin provides the only source of photic input to the brain. At P8, dozens of transcripts were enriched in Het axons compared to KOs, consistent with activity-dependent local translation. Neural activity regulates local translation elsewhere in the brain by modulating polysome recruitment , mRNA trafficking, and translation at active synapses ^76–85^. In this context, transcript enrichment in Het axons at P8 is consistent with activity-dependent local translation associated with melanopsin signaling.

Interestingly, KO axons also showed transcript enrichment at P8. Developmental comparisons confirmed that KO axons contained more ribosome-associated transcripts at P8 than at P15. The genes enriched in this KO temporal comparison were largely distinct from those identified in the Het versus KO comparison at P8, with only minimal overlap between gene sets. The enrichment in KO axons may reflect aberrant translational regulation in the absence of melanopsin signaling, including altered ribosome stalling or mRNA trafficking and stability.

Approximately 15% of transcripts isolated from the neonatal retina may reflect stalled ribosomes ^23^. However, we observed no differences in retinal translatomes between Het and KO mice at either age, suggesting that ribosome stalling in ipRGC cell bodies and dendrites is activity-independent. Alternatively, enrichment in KO axons may reflect altered delivery or retention of mRNAs within axons. In adult RGCs, continued axonal mRNA transport is required to sustain local translation ^86^, but whether mRNA abundance correlates with translational output during development remains poorly understood. Future mRNA tagging and pulse-chase experiments will be necessary to distinguish whether melanopsin regulates the trafficking and stability of axonal mRNAs, their conversion into proteins, or both.

### Structural defects in RHT development

Local translation defects in Opn4-KO mice correlate with reduced ipsilateral innervation bilaterally within the SCN. This reduction is notable given the known projection pattern of ipRGC axons. A minor population of ipRGCs in the ventrotemporal retina projects purely ipsilaterally, while most M1-type ipRGCs project contralaterally, branching to form collateral inputs to the opposite SCN ^47^. These branching patterns normally produce roughly symmetric bilateral innervation from both eyes ^36,47,87^. In Opn4-KO mice, we observed reduced ipsilateral innervation together with a trend toward increased CTB signal at the lateral edge of the contralateral SCN. This pattern suggests abnormal development of contralaterally projecting ipRGC axons, potentially reflecting impaired branch formation or stabilization. These anatomical defects are consistent with the depletion of locally-translated growth regulators in ipRGC axons leading to the disruption of coordinated axon growth.

Melanopsin loss also reduces synaptic transmission from ipRGCs and alters retinal wave properties prior to eye-opening ^14,16,88,89^. However, retinal waves appear unlikely to be the primary driver of RHT development defects, as mice with aberrant retinal wave properties exhibit largely normal binocular SCN innervation ^90^. Moreover, M1-type ipRGCs, the major RHT contributors, are weakly recruited into retinal waves and rely more heavily on melanopsin-driven activity during this developmental period ^5^.

### Reduced synapse formation with preserved nanoscale architecture

Volumetric STORM imaging revealed a significant reduction in RHT synapse density in the SCN of Opn4-KO mice at P8. This finding is consistent with prior electrophysiological studies showing that dark rearing reduces both the frequency and release probability of RHT inputs ^22^. The reduction in synapse number does not appear to result from increased microglial pruning, as engulfment of VGluT2 and CTB-Alexa Fluor 488 by microglia was unchanged between WT controls and KOs. This aligns with recent findings that microglial ablation from birth does not impact major features of circuit development ^91,92^. Nevertheless, we cannot rule out melanopsin-dependent synapse pruning by astrocytes, as shown in the developing dLGN ^93^.

Despite reduced synapse density, individual RHT synapses in Opn4-KO mice exhibited normal VGluT2, Bassoon, and Homer1/2/3 cluster volumes and intensities, indicating preserved nanoscale organization. This contrasts with the developing dLGN, where retinal activity regulates presynaptic vesicle pool size and protein content at individual RGC synapses ^94,95^. This distinction may reflect the unique development of ipRGC synapses in the SCN, which are smaller, contain fewer mitochondria, and engage distinct postsynaptic signaling programs ^96^. However, other synaptic components that we did not assess here could still be affected by the loss of melanopsin.

### Target-specific effects on gene expression

Beyond axonal translation, melanopsin loss induced target-specific effects on developmental gene expression across the visual system, suggesting that early ipRGC activity influences both pre- and postsynaptic cellular development. While ipRGC input promotes maturation in the visual thalamus, it appears to delay maturation in the developing retina and SCN. This divergence may reflect distinct contributions from ipRGC subtypes. M1-type ipRGCs mediate light-dependent retinal development ^17,18^ and provide most SCN input, whereas non-M1 types innervate the visual thalamus.

Consistent with this model, our sequencing results show that melanopsin loss preferentially affects VIP- and AVP-expressing SCN neurons that receive direct ipRGC input ^50^, suggesting that reduced neurotransmission contributes to altered developmental trajectories in postsynaptic targets.

### Implications for activity-dependent circuit development

Our findings place ipRGCs among other systems that use sensory-regulated local translation to guide circuit plasticity. Contextual fear conditioning increases ribosome-mRNA binding in CA1 neurites ^97^ and further modulates local translation in astrocytic endfeet at tripartite synapses ^98^. Similarly, auditory fear conditioning increases axonal translation of mitochondrial and ribosomal protein coding transcripts in auditory cortical projections neurons during memory consolidation ^99^. Developing ipRGC axons may share parallels with olfactory sensory neuron axons, which exhibit developmental regulation of local odorant receptor mRNA translation proposed to guide axon targeting^100^. Although mechanistically distinct, these examples suggest that activity-dependent local translation may represent a general mechanism through which sensory input influences axon growth, synapse formation, and plasticity in the developing brain.

## Conclusions

Our results demonstrate that ipRGCs express compartment-specific translatomes regulated by melanopsin-dependent activity during a discrete developmental window before eye-opening. The melanopsin-dependent axonal translatome reveals a coordinated program of local translation supporting cytoskeletal dynamics, axon growth, and synapse formation. Melanopsin is required for both RHT innervation and synapse formation during SCN development, but not for nanoscale synaptic organization. These findings support a model in which early light input regulates local axon translation during circadian circuit assembly and raise the possibility that activity-dependent local translation is a general mechanism through which sensory input shapes neural connectivity during early brain development.

## Materials and Methods

### Mouse lines

The Opn4-Cre transgenic mouse line (JAX stock #035925) ^101^, Ribotag mice (#029977, B6J.129(Cg) Rpl22<tm1.1Psam>/SjJ) ^102^ and Ai9(RCL-tdT) (B6.Cg-*Gt(ROSA)26Sortm9(CAG-tdTomato)Hze*/J) (JAX stock #007909) ^103^ were obtained from Jackson. All experimental procedures were performed in accordance with an animal study protocol approved by the Institutional Animal Care and Use Committee (IACUC) at the University of Maryland. The Rpl22^HA^ transgenic line was crossed with the Opn4^Cre^ line to generate mice expressing HA epitope-tagged ribosomes only in the presence of Cre recombinase. Three genotypes were generated for the TRAP study. Opn4^Cre/Cre^::Rpl22^HA/+^ are melanopsin knockouts (KO). Opn4^Cre/+^::Rpl22^HA/+^ retain melanopsin expression (Het). Opn4^+/+^::Rpl22^HA/+^ are wild type immunoprecipitation controls (WT). There were no gross phenotypic differences (e.g. body weight, activity, etc.) between *Opn4* genotypes.

PCR was performed using published primers. Primers for *Opn4* genotyping were FP:GGGTTCTGAGAGTGAAGTGG, RP:AAGAGGCCTTGAGTTCTCC ^101^. Primers for Opn4^Cre^ genotyping were FP:GCCAGCTAAACATGCTTCATC and RP:ATTGCCCCTGTTTCACTATCC. Primers for Rpl22^HA^ genotyping were FP:GGGAGGCTTGCTGGATATG RP:TTCCAGACACAGGCTAAGTACAC (The Jackson Laboratory). Primers for Ai9 were FP:GGGAGGCTTGCTGGATATG and RP: TTCCAGACACAGGCTAAGTACAC (The Jackson Laboratory). Some samples were also genotyped by Transnetyx (Cordova, TN). Cre-negative samples served as negative controls for non-specific binding in TRAP experiments. All samples were Rpl22^HA/+^ as in published axon-TRAP protocols ^104^. Male mice were used for axon-TRAP-RiboTag, and mice of both sexes were used interchangeably in all other experiments. Animals were maintained on a 12-hour light / 12-hour dark cycle, and all procedures were performed during the light phase between 12:00 PM and 5:00 PM.

### Bulk RNA isolation

P8 brains were sectioned at 750 µm on a vibratome (Leica) and central targets (SCN and dLGN) were manually dissected and frozen in low-bind Eppendorf tubes in liquid nitrogen. Retinas were dissected and frozen separately in lo-bind tubes.

Tissues were stored at −80°C until processing. Homogenization was performed using a pellet pestle in lysis buffer (20 mM HEPES-KOH, 5 mM MgCl_2_, 150 mM KCl, 1 mM DTT, SUPERase In, and Complete EDTA-free Protease Inhibitor Cocktail).

Homogenates were centrifuged at 16,000 x g for 10 minutes at 4 °C. The supernatant was incubated with 500 µL ice-cold Trizol (Invitrogen, 15596026) for 5 minutes at room temperature. After adding 100 µL (5:1 ratio) of chloroform, the lysate tubes were incubated for 2-3 minutes, vortexed, and centrifuged at 12,000 x g for 15 minutes at 4°C. The aqueous phase was mixed with 250 µL isopropanol and centrifuged at 12,000 x g for 10 minutes at 4° C. The pellet was washed with 500 µL 75% ethanol and centrifuged at 7,500 x g for 5 minutes at 4°C. RNA was purified using DNase I (Qiagen) according to manufacturer instructions and eluted in 15 µL RNase-free water. RNA quantity was measured using a Qubit 3 fluorometer High Sensitivity RNA Assay Kit (Invitrogen, Q32852). RNA integrity was assessed on an Agilent TapeStation 4150 (Agilent Technologies) using RNA and HS RNA assay kits. Samples were stored at -80°C before sequencing.

### Library preparation and bulk RNA sequencing

Library preparation and sequencing were performed at the Brain and Behavior Institute - Advanced Genomic Technologies Core (BBI-AGTC). Libraries were prepared using Illumina Stranded mRNA with IDT for Illumina DNA/RNA UD Indexes Sets A. Starting material was 275 ng of total RNA in 25 µL RNase-free water. Fragmentation time was 8 minutes with 12 PCR amplification cycles. Library quality was assessed on an Agilent TapeStation 4150 using a D1000 Assay kit. Following PolyA selection, library concentration was determined by qPCR on a Roche LightCycler 96. Libraries were pooled (10 µL each at 2 nm) and sequenced in a single batch run on an Illumina NextSeq 1000 using a 58 bp paired-end read protocol with dual indexing. Raw data were provided in FASTQ format.

### Bulk RNA sequencing read preprocessing

Fastq files were trimmed using Trimmomatic ^105^ to remove Illumina and Epicenter adapters and perform sliding window removal of bases with quality scores below 25. Trimmed reads were aligned to the Genome Reference Consortium Mouse Build 38 using HISAT2 ^106^. Gene counts were obtained using HTSeq ^107,108^.

### Bioinformatic analysis of bulk RNA-seq data

Analyses were performed in R version 4.2.1 (R Core Team, 2021) using custom functions and the hpgltools package (https://github.com/elsayed-lab/hpgltools). Lowly expressed genes were filtered to retina only those with > 10 reads across all samples (14,107 genes remaining after filtering). Count data were converted to counts per million (CPM), quantile normalized, and log2 transformed before principal component analysis (PCA). Differential expression (DE) analysis was conducted using DESeq2 with raw counts as input ^109^. Significance thresholds were log2 fold change > 1 and adjusted p-value ≤ 0.05. Gene set enrichment analysis was performed using gProfiler2 ^110^ with Gene Ontology (GO), KEGG, and REACTOME databases using an adjusted p-value cutoff of ≤ 0.05.

### Module score assignment across developmental datasets

To uncover the cell type specificity and developmental effects of melanopsin during development of the retina, SCN, and dLGN, we projected differentially enriched genes from bulk RNA sequencing onto single cell transcriptomic atlases by computing module scores for each gene set per cell. We implemented the Seurat AddModuleScore function, which aggregates expression of module genes into discrete bins and randomly selects control genes with similar aggregate expression (repeated n *=* 10 times). The module score represents enrichment of module genes compared to randomly sampled genes of comparable expression distribution. To make results comparable between modules, we computed the z-score across all cell types and ages for visualization and cell type-specific permutation testing.

### Tissue collection and immunostaining for TRAP validation

P8 Opn4^Cre/Cre^::Rpl22^HA/+:^Ai9^+/-^ mice were anaesthetized by intraperitoneal injection of 100 µL ketamine/xylazine. Animals were transcardially perfused with 5-10 mL of 0.9% sterile saline followed by 5-10 mL of 4% EM-grade paraformaldehyde (PFA, Electron Microscopy Sciences) diluted in 1X PBS. Brains were postfixed overnight in 4% PFA at 4°C. Retinas were dissected, postfixed in in 4% PFA for 30 minutes at room temperature, and washed for 20 minutes in 1X PBS at room temperature in preparation for whole-mount immunohistochemical labeling. Brains were cryoprotected overnight in 30% sucrose in 1X PBS at 4°C and embedding in OCT (optical cutting temperature) compound before freezing at -20°C. Brains were sectioned at 60 µm on a cryostat (Microm HM550). Retinas and brains were blocked for 2 hours at room temperature in 10% normal donkey serum (Jackson Immunoresearch) with 0.3% Triton-X 100. Retinas and brain sections were incubated for 48 hours at 4°C with primary antibodies (rat anti-HA, Roche 11-867-423, 1:500 dilution; chicken anti-mCherry, Kerafast, EMU110, 1:500 dilution), followed by extensive washing in 1X PBS at room temperature (6 x 20-minute exchanges). Samples were incubated in secondary antibodies (Alexa Fluor 488 AffiniPure Donkey Anti-Rat IgG (Jackson ImmunoResearch Laboratories, 712-545-150, 1:500 dilution; Alexa Fluor 594 AffiniPure Donkey Anti-Chicken IgY (Jackson ImmunoResearch Laboratories, 712-545-150, 1:500 dilution) for 1 hour at room temperature. Tissues were washed in 1X PBS 6x for 20 minutes each exchange. Samples were mounted with Vectashield (Vector Laboratories, H-1000-10). Images were acquired on a Zeiss LSM 980 Airyscan 2 and analyzed with Fiji ^111^.

### RHT tract tracing for TRAP tissue isolation

We collected tissue at P8 (before eye-opening, when activity is photoreceptor-independent and governed by melanopsin) and P15 (after eye-opening, when rods and cones are also active). Twenty-four hours before tissue collection, bilateral intraocular injections were performed. At P7 or P14, mice were anesthetized with isoflurane. For P7 pups, sterile surgical spring scissors were used to gently part the eyelid and expose the corneoscleral junction. A small hole was made with a sterile 34-gauge needle, and 0.5 μL of CTB-Alexa Fluor 488 (1 μg/μL, ThermoFisher Scientific C34775) in 0.9% sterile saline was injected intravitreally using a Hamilton syringe fit with a 34-gauge, 1-inch blunt end needle (Hamilton part number 207434) attached to a micromanipulator (MM-3, Narishige Group).

### Tissue collection for axon-TRAP RNA-seq

At P8 or P15, animals were anesthetized with isoflurane and decapitated. Eyes and brain were immediately submerged in ice cold 200 µM cycloheximide diluted in 1X PBS to prevent ribosome runoff. Brains were sectioned at 750 µm on a vibratome in ice cold 200 µM cycloheximide solution. SCN, dLGN, and retinas were manually dissected, collected into separate low-bind Eppendorf tubes, frozen in liquid nitrogen within 30 minutes of euthanasia, and stored at −80°C.

### Immunoblotting

Mouse retinas (50 mg) were homogenized using a pellet pestle. Following centrifugation at 16,000 x g for 10 minutes, supernatants were collected. An aliquot (50 µL) was reserved as input. Precleared lysates were incubated with 5 µL Rabbit anti-HA-antibody (Abcam, ab9110) overnight at 4°C, then with 50 µL Dynabeads protein A/G for four hours at 4°C. Immunoprecipitates were denatured by boiling in Laemmli buffer, fractionated by SDS-PAGE, transferred to nitrocellulose membranes, and blocked with 5% BSA in TBST for 1 hour at room temperature. Blots were incubated overnight at 4°C with rat anti-HA (Roche, 11-867-423) and rabbit-anti-GAPDH antibodies (ThermoFisher, MA515738). After four washes in 1X TBST at room temperature, blots were incubated with secondary antibodies (Alexa Fluor 546 goat anti-rat, Invitrogen A11081; Alexa Fluor 647 goat anti-rabbit, ThermoFisher A32733) for 1 hour at room temperature. Bands were visualized on a Typhoon TRIO Variable Imager (GE Healthcare) and analyzed using Image Studio Lite v.5.x (LI-COR Biosciences).

### Translating ribosome affinity purification (TRAP)

Immunoprecipitation of ribosome-associated mRNAs was performed according to published protocols ^112^. Tissues from animals of the same age and genotype were pooled to create a single biological replicate sample for each tissue type (10-12 retinas, dLGNs, and SCNs). Pooled biological replicate samples were collected from: P8 — wild-type retina (N=5), SCN (N=3), dLGN (N=5); *Opn4* heterozygous retina (N=3), SCN (N=3), dLGN (N=3); *Opn4* knockout retina (N=3), SCN (N=3), dLGN (N=3); P15 — wild-type retina (N=5), SCN (N=2), dLGN (N=5); *Opn4* heterozygous retina (N=4), SCN (N=3), dLGN (N=4); *Opn4* knockout retina (N=3), SCN (N=3), dLGN (N=3). Final Samples were homogenized 3-4 times at 10-15 second intervals using a pellet pestle in lysis buffer (20 mM HEPES-KOH, 5 mM MgCl_2_, 150 mM KCl, 1 mM DTT, SUPERase In, 0.5 % NP-40, EDTA-free Protease Inhibitor, and 100 µg/mL cycloheximide) on ice. Homogenates were centrifuged at 16,000 x g for 10 minutes at 4° C. Supernatants were transferred to new tubes and pre-cleared with 40 µL Protein G beads (Thermo Scientific Pierce, 88803). Pre-cleared lysates were incubated with polyclonal anti-HA antibody (Abcam, ab9110) (5 µL for retina samples and 2.5 µL for dLGN/SCN samples) overnight at 4° C with rotation. Dynabeads Protein G (60 µL, Life Technologies, 10004D) were added and incubated for 4 hours at 4° C with rotation. Beads were washed in ice-cold high salt buffer (20 mM HEPES pH 7.4, 350 mM KCl, 10 mM MgCl_2_, 0.5 % NP-40, 1 mM DTT, 100 μg/mL cycloheximide, EDTA-free protease inhibitors, and 100 U/mL SUPERase IN). After four washes, beads were resuspended in 100 µL cycloheximide lysis buffer and incubated with 500 µL Trizol (Invitrogen, 15596026) for 5 minutes at room temperature. Samples were centrifuged (12,500 x g) for 1 minute at room temperature and placed on a magnetic stand for 30 seconds. The supernatant was transferred to a new tube, 100 µL of chloroform was added and the samples were incubated for 2-3 minutes, vortexed, and centrifuged at 12,000 x g for 15 minutes at 4°C. The aqueous phase was mixed with 250 µL isopropanol and centrifuged at 12,000 x g for 10 minutes at 4°C. The pellet was washed with 500 µL 75% ethanol and centrifuged at 7,500 x g for 5 minutes at 4°C. RNA was purified using DNase I (Qiagen) according to manufacturer instructions and eluted in 15 µL RNase-free water. RNA quantity was measured using a Qubit 3 fluorometer High Sensitivity RNA Assay Kit (Invitrogen, Q32852). RNA integrity was assessed on an Agilent TapeStation 4150 (Agilent Technologies) using HS RNA assay kits. Samples were stored at -80°C before sequencing.

### Library preparation and sequencing of TRAP RNA

Library preparation was performed at the BBI-AGTC using the TakaraBio SMARTer Stranded Total RNA-Seq Kit v3 - Pico Input Mammalian Kit (Takara, 634486) with SMARTer RNA Unique Dual Index Kit – 96U (8bp) (Takara, 634452). Libraries were prepared from 1-6 ng total RNA with 2-minute fragmentation time, 5 PCR1 amplification cycles, and 12-14 PCR2 cycles depending on RNA concentration.

Library quality was assessed on an Agilent TapeStation 4150 using a HSD1000 Assay kit. Libraries were pooled and sequenced on an Illumina NextSeq 1000 using a 60 bp paired-end protocol. A single pool was prepared by combining 10 µL of each sample at 2 nM. FASTQ files were generated using BCL Convert.

### Axon-TRAP-RiboTag sequence processing

Fastqc ^113^ and fastp ^114^ were used to query sequence quality and Trimmomatic ^105^ was used to remove low-quality sequence. The remaining reads were marked by UMI-tools ^115^ with the default regular expression sensitive to the Takara unique molecular identifiers. Marked reads were aligned against release 112 of the ensembl mouse genome ^116^ with hisat2 ^106^ with the very-sensitive parameter set. The resulting alignments were compressed, sorted, and indexed via samtools ^117^ and deduplicated via UMI-tools. Reads per gene were quantified by featureCounts ^118^ using the pre and post-deduplication alignments with default parameters; transcripts were similarly quantified via salmon ^119^. All raw reads for this project are available at the NCBI Short Read Archive (SRA) under BioProject accession PRJNA1427139.

### Global data assessment, visualization, and differential translation analyses

Processed data were assessed and visualized using the hpgltools ^120^ package: normalized data were visualized using log2 transformed counts per million reads following low-count per gene filtering. Various visualizations, including density plots, depth boxplots, CV/condition, hierarchical clustering using Pearson’s correlation coefficient and Euclidean distance, variance partition ^121^, and principal component analyses were performed to observe sample relationships. Batch effects and/or surrogate variables were queried via a combination of normalization strategies with and without sva ^122^. Differential expression analyses were performed in multiple rounds using a single pipeline, which performed all pairwise comparisons using the Bioconductor packages: DESeq2 ^109^, EBSeq ^123^, edgeR ^124^, limma ^125^, NOISeq ^126^, variancePartition/Dream ^127^, and a basic analysis using simple log2 CPM comparisons; when applicable the surrogate estimates provided by sva were appended to the statistical model (e.g. not for EBSeq, basic, NOISeq, nor Dream). DESeq2 values were used in subsequent analyses. The first round of differential expression compared cre-negative wild-type samples against heterozygous and knockout samples from the same tissue. Genes not detected above the non-specific (wild-type) background in the experimental (heterozygous and knockouts) samples were excluded from subsequent analyses. A second round of differential expression was performed with contrasts to compare expression by tissue type, genotype, and developmental age. The resulting significantly differentially expressed gene sets (defined as |log2FC| >= 1.0 and adjusted p-value <= 0.05) as well as the full set of log2FC values were passed to a combination of gProfiler2 ^110^ and clusterProfiler ^128^ to perform complementary enrichment and GSEA searches for statistically significant categories among GO, KEGG, miRNA, mSigDB (C2), reactome, TFdb, etc. All analyses performed using the pre/post deduplication data are available as a container recipe at doi:10.5281/zenodo.18985313.

### Two-color binocular RHT tract tracing

At P7, mice were anesthetized with isoflurane. Sterile surgical spring scissors were used to gently part the eyelid and expose the corneoscleral junction. A small hole was made with a sterile 34-gauge needle, and 1 µL of CTB-Alexa Fluor 488 (1 μg/μL, ThermoFisher Scientific C34775) in 0.9% sterile saline was injected intravitreally into the left eye using a Hamilton syringe fit with a 34-gauge, 1-inch blunt end needle (Hamilton part number 207434) attached to a micromanipulator (MM-3, Narishige Group). The right eye was injected with 1 µL of CTB-Alexa Fluor 647 (2 µg/µL, Invitrogen, C34778). Twenty-four hours later (P8), pups were anesthetized by intraperitoneal injection of ketamine/xylazine in 0.9% sterile saline. Animals were transcardially perfused with 7 mL of 0.9% sterile saline, followed by 7 mL of 4% PFA in 1x PBS (Electron Microscopy Sciences, pH 7.4). Brains were dissected and postfixed overnight in 4% PFA at 4°C. After three 10-minute washes in 1x PBS, brains were cryoprotected in 30% sucrose at 4°C for 48 hours. Tissues were embedded in 2:1 OCT:30% sucrose, frozen at -80°C, and cryosectioned coronally at 60 µm. Sections containing the SCN (Bregma -0.46 to -0.58 mm) were collected in 1x PBS and screened under a fluorescence stereoscope. Selected sections were mounted with Vectashield (Vector Laboratories, H-1000-10). Retinal integrity was verified for each injection to ensure no injection-related damage.

### Two-color binocular RHT Imaging

Brain slices were imaged on a Leica Stellaris 8 FALCON laser scanning confocal microscope in the Department of Cell Biology and Molecular Genetics Imaging Core at the University of Maryland, College Park. Sections containing the SCN were imaged individually using a Leica 20x/0.75 NA objective with 0.9x zoom, acquiring a single optical plane from each slice.

### Two-color binocular RHT image analysis

Red and green channels were extracted and normalized separately for each genotype group using linear histogram stretching with 1% and 99% intensity saturation limits to normalize within-group intensity variation while preserving between-group differences. One-dimensional intensity profiles were computed by averaging pixel intensities along the Y-axis within spatial bins of 80 pixels (50.51 µm) for alignment and 20 pixels (12.63 µm) for visualization. Images were aligned to the midline using a valley-detection algorithm that identified the intensity minimum within a 400–600-pixel search window. The mean valley position across all animals was computed and individual images were shifted horizontally to align their valleys to this common reference point. To assess spatial distribution patterns independent of absolute intensity, profiles were additionally normalized to the 0-1 range within each animal. Average intensity images were generated by computing pixel-wise means across aligned images within each group.

Difference maps were calculated by subtracting KO from WT group averages. Spatial coordinates were converted from pixels to microns (0.6313 µm/pixel) with the X-axis centered at the aligned midline.

To compare intensity distributions between WT (N = 10) and KO (N = 10) groups, aligned profiles were divided into negative (ipsilateral, x < 0) and positive (contralateral, x > 0) halves. For each animal, a single summary statistic was computed as the mean intensity across all bins within each half, ensuring each biological replicate contributed one value. Group differences were assessed using the Mann-Whitney U test. Effect sizes were quantified using Cohen’s d with conventional thresholds (small d = 0.2, medium d = 0.5, and large d = 0.8). Post-hoc power was estimated using the non-central t-distribution. Results were considered robust when both p < 0.05 and power > 0.8. All analyses were performed in MATLAB (version 2025a) using a custom pipeline developed with assistance from Claude 4.5 Sonnet, Anthropic.

### Microglial engulfment assay

WT and Opn4-KO mice of both sexes received binocular CTB-Alexa Fluor 488 injections at P7 (0.5 µL, 3 mg/mL). At P8, animals were transcardially perfused with 5 mL of 0.9% sterile saline followed by 10 mL of 4% PFA in 1x PBS at room temperature. Brains were collected (N=7 biological replicates from each genotype) and immersion fixed overnight (∼16 hours) in 4% PFA in 1x PBS at 4°C. Brains were washed 2 x 20 minutes in 1X PBS at room temperature and sectioned on a vibratome (unembedded) at 50 µm. Tissue sections containing the SCN were blocked in 10% normal donkey serum (Jackson ImmunoResearch) in 1X PBS with 0.3% Triton-X 100 + 0.02% sodium azide for ∼4 hours at room temperature on an orbital shaker. Sections were then incubated in this same blocking solution for ∼64 hours with added primary antibodies: rabbit anti-Iba1 (Cat # 019-19741; Fujifilm Wako; 1:100 dilution) + guinea pig anti-VGluT2 (AB251-I; Millipore; 1:200 dilution). Sections were washed 6 x 20 minutes each in 1X PBS at room temperature on an orbital shaker and incubated in blocking solution plus secondary antibodies (Jackson ImmunoResearch) overnight at 4°C with gentle agitation: donkey anti-rabbit Atto565 (1:100; 2.8 dyes/antibody; 711-005-152) and donkey anti-guinea pig Atto647N (1:100; 4.5 dyes/antibody; 706-005-148). Secondary antibodies were custom conjugated as described previously ^45^. Sections were washed 6 x 20 minutes each in 1X PBS at room temperature on an orbital shaker, mounted, and coverslipped (#1.5) with Vectashield (Vector Laboratories). Slides were sealed with nail polish and stored flat in a slide folder at 4°C prior to imaging. Three color image stacks were acquired through a 63x 1.4NA oil immersion objective on a Zeiss LSM 980 Airyscan 2 Laser Scanning Confocal microscope in the UMD Imaging Core. Three color confocal z-stacks were processed using custom MATLAB scripts (MathWorks, R2025a). Each channel was normalized independently using linear contrast stretching, mapping pixel intensities from the 1st to 99.5th percentile to the 0–255 range. A 3D Gaussian filter (σ = 1.5 pixels) was applied, and adaptive percentile-based thresholding was used to account for brightness variation across images. Pixels above the 97th (green channel, Iba1), 99th (red channel, VGluT2), and 99th (blue channel, CTB-Alexa Fluor 488) percentiles were classified as foreground. Connected components were identified in 3D and volume filtered to exclude microglial signal less than 50 µm^3^ and VGluT2 and CTB(+) puncta less than 0.1 µm^3^. Engulfed puncta were defined as VGluT2(+) or CTB(+) voxels spatially overlapping with the thresholded microglial signal. Engulfment was quantified as the percentage of microglial volume occupied by engulfed puncta. Data were collected from 7 animals per group (WT: 66 total cells; Opn4-KO: 69 total cells). To account for the hierarchical data structure (multiple imaging fields nested within animals), a linear mixed-effects model was used with genotype as a fixed effect and animal number as a random effect. Analysis was performed in MATLAB (version 2025a) using a custom pipeline developed with assistance from Claude 4.5 Sonnet, Anthropic. 3D engulfment models were generated in ParaView ^129^.

### Tract tracing for STORM experiments

Intraocular injections were performed one day before tissue collection. P7 mice were anesthetized with isoflurane, and the eyelid was parted with sterile surgical spring scissors to expose the corneoscleral junction. A small hole was made in the eye with a sterile 34-gauge needle and 0.5 µL of CTB-Alexa Fluor 488 (ThermoFisher Scientific, C34775) in 0.9% sterile saline was injected intravitreally into each eye using a pulled-glass micropipette coupled to a Picospritzer (Parker Hannifin).

### Tissue collection for STORM

Animals were deeply anesthetized with ketamine/xylazine and transcardially perfused with 5-10 mL of 37°C 0.9% sterile saline followed by 10 mL room temperature 4% EM-Grade PFA (Electron Microscopy Sciences) in 0.9% saline. Brains were embedded in 2.5% agarose and sectioned coronally at 100 µm on a vibratome. From the full anterior-posterior SCN series (approximately 3-4 sections), the central two sections (Bregma -0.46 to -0.58) were selected for staining in all biological replicates ^44^. Selected sections were postfixed in 4% PFA for 30 minutes at room temperature and washed for 30 minutes in 1X PBS. The SCN was identified by CTB-Alexa Fluor 488 signal using a fluorescence dissecting microscope. A circular tissue punch (500 μm diameter) containing the SCN was microdissected from each section using a blunt-end needle.

### Immunostaining for STORM imaging

Samples were prepared for serial-section single-molecule localization imaging as previously described ^45^. SCN tissue punches were blocked in 10% normal donkey serum (Jackson ImmunoResearch, 017-000-121) with 0.3% Triton X-100 and 0.02% sodium azide (Sigma-Aldrich Inc.) in 1X PBS for 2-3 hours at room temperature.

Tissues were incubated in primary antibodies for approximately 72 hours at 4°C on an orbital shaker. Primary antibodies were rabbit anti-Homer1/2/3 (Synaptic Systems, 160103, 1:100 dilution) for postsynaptic densities, mouse anti-Bassoon (Abcam, AB82958, 1:100 dilution) for presynaptic active zones, and guinea pig anti-VGluT2 (Millipore, AB251-I, 1:100 dilution) for presynaptic vesicles. Tissues were washed in 1X PBS for 6 x 20 minutes at room temperature and incubated in secondary antibodies overnight for approximately 36 hours at 4°C on an orbital shaker.

Secondary antibodies were donkey anti-rabbit IgG (Jackson ImmunoResearch, 711-005-152, 1:100 dilution) conjugated with Alexa Fluor 647 and Alexa Fluor 405 (ThermoFisher, A20006, A30000), donkey anti-mouse IgG (Jackson ImmunoResearch, 715-005-150, 1:100 dilution) conjugated with Dy749P1 and Alexa Fluor 405 (Dyomics, 749P1-01; ThermoFisher, A30000), and donkey anti-guinea pig IgG (Jackson ImmunoResearch, 706-005-148, 1:100 dilution) conjugated with Cy3b (Cytiva, PA63101). Tissues were washed 6 x 20 minutes in 1X PBS at room temperature after secondary antibody incubation.

### Postfixation, dehydration, and embedding in epoxy resin

Tissue embedding was performed as previously described (Vatan et al., 2021). Tissues were postfixed with 3% PFA + 0.1% GA (Electron Microscopy Sciences) in 1X PBS for 2 hours at room temperature and washed in 1X PBS for 20 minutes. For ultrasectioning, tissues were dehydrated in a graded ethanol series (50%, 70%, 90%, 100%, 100%) for 15 minutes each at room temperature. Tissues were then immersed in a series of epoxy resin/100% ethanol exchanges (Electron Microscopy Sciences) with increasing resin concentration (25%, 50%, 75%, 100%, 100% resin) for 2 hours each. Tissues were transferred to BEEM capsules (Electron Microscopy Sciences) filled with 100% resin and polymerized for 16 hours at 70°C.

### Ultrasectioning

Plasticized tissue sections were cut at 70nm using a Leica UC7 ultramicrotome with a Histo Jumbo diamond knife (DiATOME). Chloroform vapor was used to reduce section compression. For each sample, approximately 100 sections were collected coverslips coated with 0.5% gelatin and 0.05% chromium potassium (Sigma-Aldrich Inc.), dried at 60°C for 25 minutes, and protected from light prior to imaging.

### Imaging chamber preparation

Coverslips were chemically etched in 10% sodium ethoxide for 5 minutes at room temperature to remove epoxy resin and expose dyes to the imaging buffer.

Coverslips were rinsed with ethanol and deionized water. Fiducial beads for flat-field and chromatic corrections were prepared by mixing 715/755 nm and 540/560 nm carboxylate-modified microspheres (Invitrogen, F8799, F8809, 1:8 ratio) to create a high-density bead solution, then diluting 1:750 with Dulbecco’s PBS to create a low-density bead solution. Both high- and low-density bead solutions were spotted on coverslips (approximately 0.7 µL each). Excess beads were rinsed with deionized water for 1-2 minutes. Coverslips were attached to glass slides with double-sided tape to form imaging chambers. Chambers were filled with STORM imaging buffer (10% glucose, 17.5 μM glucose oxidase, 708 nM catalase, 10 mM MEA, 10 mM NaCl, and 200 mM Tris) and sealed with epoxy.

### Imaging setup

Imaging was performed on a custom single-molecule super-resolution imaging system. The microscope contained low (4x/10x air) and high (60x 1.4NA oil immersion) magnitude objectives mounted on a Nikon Ti-U frame with back optics arranged for oblique angle illumination. Continuous-wave lasers at 488 nm (Coherent), 561 nm (MPB), 647 nm (MPB), and 750 nm (MPB) excited Alexa Fluor 488, Cy3B, Alexa Fluor 647, and Dy749P1 dyes respectively. A 405 nm laser (Coherent) reactivated Dy749P1 and Alexa Fluor 647 photoswitching. The microscope was fitted with a custom pentaband/pentanotch dichroic filter set and motorized emission filter wheel. An IR laser-based focus lock system maintained optimal focus during automatic acquisition. Images were collected on an 896 x 896-pixel region of an sCMOS camera (ORCA-Flash4.0 V3, Hamamatsu Photonics) with 155 nm pixel size.

### Automated image acquisition

Fiducials and tissue sections were imaged at low magnification (4x) to create a mosaic overview. Beads and sections were then imaged at high magnification (60x) to select regions of interest (ROIs) in Cy3B and Alexa Fluor 488 channels. Laser intensities and incident angle were adjusted to optimize photoswitching for STORM imaging and utilize the full camera dynamic range for conventional imaging. Low-density bead images were acquired in 16 partially overlapping ROIs. 715/755nm beads were excited with 750 nm light and imaged through Dy749P1 and Alexa Fluor 647 filters. 540/560nm beads were excited with 488 nm light and imaged through Alexa Fluor 647, Cy3B, and Alexa Fluor 488 filters. These images were used to generate a non-linear warping transform for chromatic aberration correction. ROIs within each section were then imaged at conventional resolution in all four channels sequentially.

Following conventional image acquisition, a partially overlapping series of 9 images was collected in the high-density bead field for all channels to enable flat-field correction of non-uniform illumination. Additional bead images were acquired in a different low-density bead ROI to confirm chromatic offset stability. All ROIs were then imaged by STORM for Dy749P1 and Alexa Fluor 647 using a custom progression of increasing 405 nm laser intensity. For each ROI, 8,000 frames of Dy749P1 (60 Hz) were acquired followed by 12,000 frames of Alexa Fluor 647 (100 Hz). In a second pass, the same ROIs were imaged for Cy3B and Alexa Fluor 488, each for 8,000 frames (60 Hz). ROIs were positioned in the center of CTB-stained regions in the SCN (core).

### Image processing

Single-molecule localization was performed using a DAOSTORM algorithm ^130,131^. Molecule lists were rendered as 8-bit images with 15.5 nm pixel size where each molecule is plotted as an intensity distribution reflecting its localization precision. Low-density fiducial images were used for chromatic aberration correction. We localized 715/755 beads in Dy749P1 and Alexa Fluor 647 channels, and 540/560 beads in Alexa Fluor 647, Cy3B, and Alexa Fluor 488 channels. A third-order polynomial transform map was generated by matching bead positions across channels to the Alexa Fluor 647 channel. Average residual error was <15 nm for all channels. Transform maps were applied to both conventional and STORM images. Conventional images were upscaled 10x to match the STORM image size. Serial section alignment was performed as previously described ^45^. STORM images were first aligned to corresponding conventional images by correlation. For 3D image stacks, Alexa Fluor 488 images were intensity-normalized and used to generate rigid and elastic transformation matrices for all channels. The final stack was rotated and cropped to exclude incompletely imaged edges. SCN images were further cropped to exclude regions without dense CTB labeling.

### Cell body filter

Aligned STORM images showed non-specific cell body labeling in Dy749P1 and Alexa Fluor 647 channels. To restrict synapse identification to neuropil, we identified cell bodies by Dy749P1 signal and excluded these regions. STORM images were convolved with a Gaussian (σ = 140nm) and binarized using the lower threshold of a two-level Otsu method. Connected components larger than 10^11^ voxels were identified and masked in all channels. Because cell body clusters are orders of magnitude larger than synaptic clusters, this algorithm was robust across a range of size thresholds.

### Synapse identification and quantification

To correct for minor intensity variance across sections, pixel intensity histograms were normalized to the average histogram and rescaled to the full 8-bit range. Using a two-level Otsu method, conventional images were classified into background, non-synaptic labeling, and synaptic structures. Conventional images were binarized by the lower threshold to generate masks for STORM images. STORM images were convolved with a Gaussian (σ = 77.5 nm) and thresholded using the higher two-level Otsu threshold. Connected components were identified in three dimensions using MATLAB ‘conncomp’. Watershedding was applied to split improperly connected large clusters. Clusters spanning fewer than two sections were excluded. Edge synapses were removed by excluding synapses without blank image data on all adjacent sides. To distinguish non-specific labeling from synaptic signals, cluster volume and signal density (ratio of within-cluster pixels with positive signal) were quantified. Two populations were identified in 2D histograms of these parameters. The population with higher volumes and signal densities representing synaptic structures was manually selected. Repeated measurements of the manual cluster selection showed < 1% between-measurement variance.

Pre- and postsynaptic cluster pairing was determined by measuring centroid-to-centroid distances between Dy749P1 and Alexa Fluor 647 clusters and quantifying opposing channel signal intensity within a 140 nm shell around each cluster. Paired clusters with closely positioned centroids and high opposing signal densities were identified using the OPTICS algorithm. Retinal inputs were identified by pairing Bassoon with VGluT2 (Cy3B) clusters using the same method. We identified 10,814 synapses from Opn4-Het mice and 10,013 from Opn4-KO mice (N = 6 biological replicates each at P8). Cluster volume reflected total connected voxel volume and total signal intensity was the sum of voxel intensities within connected voxels.

### Statistical analysis of STORM images

Statistical analysis was performed using SPSS. Plots were generated with SPSS or R (ggplot2). A linear mixed model compared cluster volumes and total signal intensity with genotype as the fixed main factor and biological replicate IDs as nested random factors. Pairwise comparisons among main factor groups used post-hoc Bonferroni tests.

Synapse density was analyzed using Student’s t-test. In violin plots, each violin shows the distribution of grouped data from all biological replicates for each condition. Black (Het) or red (Opn4-KO) dots represent median values for each biological replicate. Horizontal black lines indicate group medians. Significance is indicated as ***P < 0.001.

## Supporting information

Table_S1_bulk_RNA_seq_significant_genes

Table_S2_TRAP_seq_tissue_contrasts_significant_genes

Table_S3_TRAP_seq_genotype_contrasts_significant_genes

Table_S4_TRAP_seq_age_contrasts_significant_genes

## Acknowledgements

We thank Drs. Toshiaki Shigeoka and Hosung Jung for correspondence on axon-TRAP bioinformatics analysis. We thank Dr. Elizabeth Quinlan for generous use of secondary antibodies for western blotting. Certain cartoon figure elements were created with BioRender.com. Purchase of the Zeiss LSM 980 Airyscan 2 was supported by Award Number 1S10OD025223-01A1 from the National Institute of Health. Purchase of the Leica Stellaris 8 was supported by Award Number 1S10ODO34260 from the National Institute of Health. All library preparation and sequencing was conducted at the Brain and Behavior Institute - Advanced Genomic Technologies Core (BBI-AGTC), which is supported by the BBI and the University of Maryland, College Park. We are especially grateful to April Hussey for helpful discussions and technical expertise concerning axon-TRAP sample processing. This study was supported by a Brain and Behavior Institute Seed Grant (UMD to N.M.E. and C.M.S.), a Brain and Behavior Research Foundation Young Investigator Grant (C.M.S.), and an NIH Director’s New Innovator Award DP2MH125812 (C.M.S.).

---

**Figure S1.**
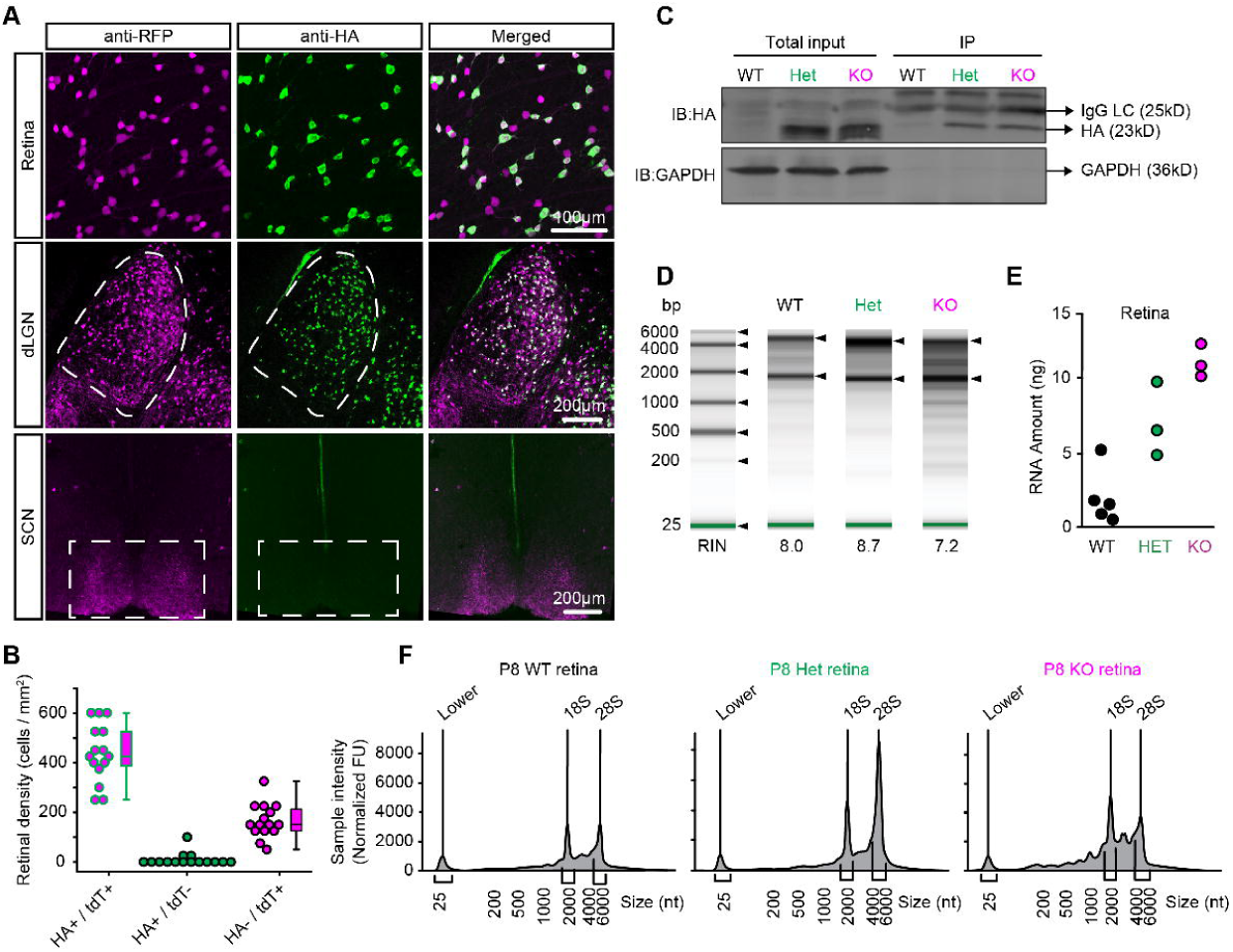
Characterization of ipRGC-TRAP labeling and mRNA isolation. (**A**) Immunofluorescence imaging of tdT and HA labels in P8 retina, dorsal lateral geniculate nucleus (dLGN), and suprachiasmatic nucleus (SCN). Anti-RFP (magenta), anti-HA (green), and merged channels show colocalization of HA and RFP signals. Dashed lines delineate the dLGN and SCN regions dissected for TRAP. **(B)** Quantification of P8 retinal density for HA^+^/tdT^+^, HA^+^/tdt^-^, and HA^-^/tdT^+^ cells. Box plots show median, quartiles, and range. Individual imaging field are shown as single data points collected from N=3 biological replicates. tdT+ RGCs: 609 ± 119 cells / mm^2^ (mean ± S.D.); HA+ / tdT+ RGCs: 440 ± 117 (mean ± S.D.). **(C)** Western blotting of P8 retinal tissue shows HA-tagged protein expression in heterozygous (Het) and knockout (KO) retinal tissue, but not WT TRAP controls. Blots show anti-HA signal (23 kD) in total input and immunoprecipitated samples, with GAPDH (36 kD) as a loading and IP control. **(D)** High sensitivity RNA ScreenTape analysis of P8 WT, Het, and KO retinal samples. RNA Integrity Numbers (RIN) are indicated below each lane. **(E)** Total RNA yield from P8 retinal tissue across genotypes where individual data points show the biological replicates used in axon-TRAP analysis (N=5 WT, N=3 Het, N=3 KO). **(F)** Electropherograms showing RNA quality profiles for P8 retinal samples from WT, Het, and KO mice. Samples correspond to those shown in (D).

**Figure S2.**
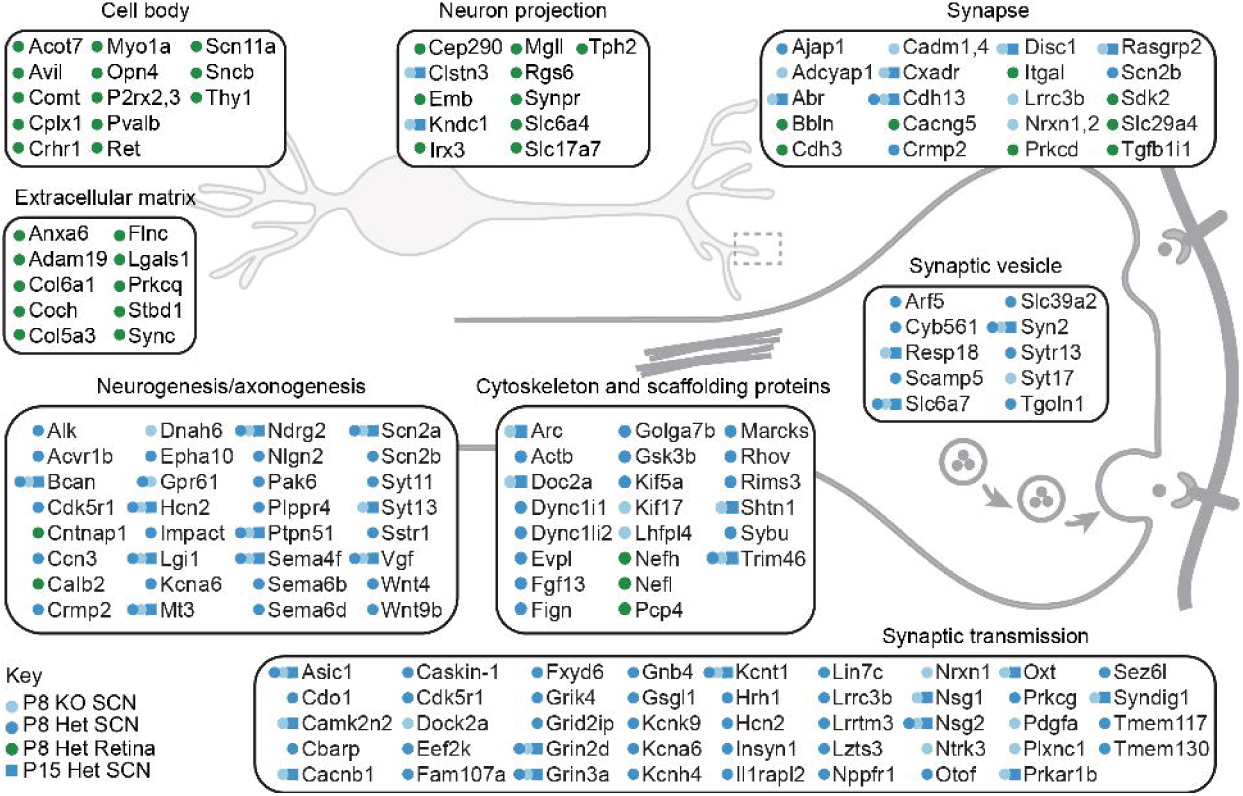
Gene ontology categories enriched in retinohypothalamic compartment-specific gene expression. Schematic diagram showing anatomical compartments with associated Gene Ontology enrichment results comparing retina versus SCN at P8 and P15 in Opn4-Het and KO animals. Color-coded key highlights compartment enrichment: P8 KO SCN (light blue circles), P8 Het SCN (darker blue circles), P8 Het Retina (green circles), and P15 Het SCN (blue squares).

**Figure S3.**
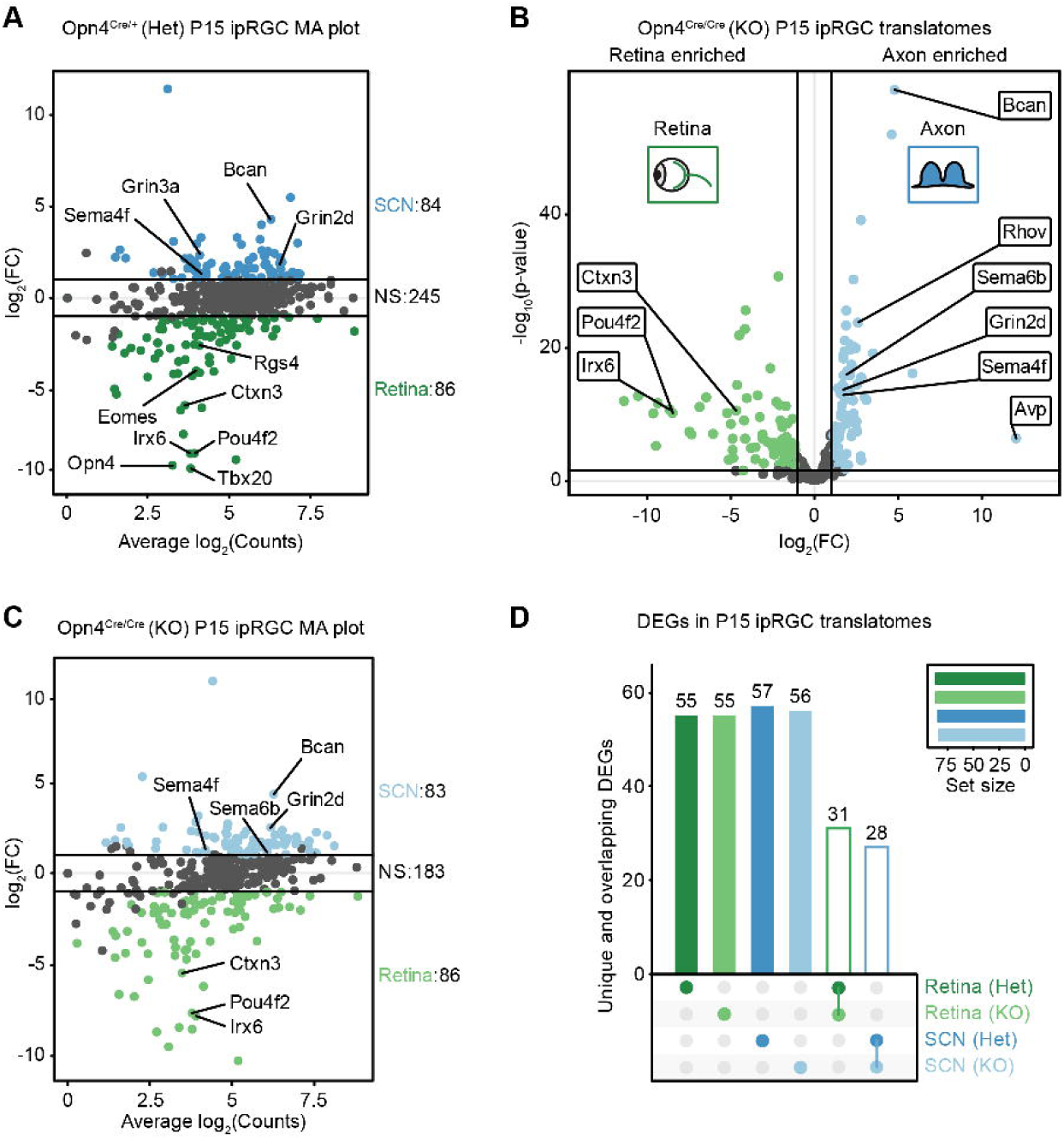
Postnatal day 15 ipRGC axon-TRAP analysis reveals persistent compartment-specific gene expression. **(A)** MA plot of Opn4^Cre/+^ (Het) P15 ipRGC translatomes showing the relationship between average expression and differential expression for retina (green, n=86 genes) versus axon (blue, n=84 genes) compartments. **(B)** Volcano plot of Opn4-KO P15 ipRGC translatomes comparing retina-enriched (green) and axon-enriched (blue) compartments. **(C)** MA plot of Opn4-KO P15 ipRGC translatomes showing SCN-enriched (blue, n=83 genes) versus retina-enriched (green, n=86 genes) transcripts. **(D)** Upset plot showing unique and overlapping differentially expressed genes (DEGs) in P15 ipRGC translatomes across genotypes and compartments. Connected dots and open bars indicate intersections between groups.

**Figure S4.**
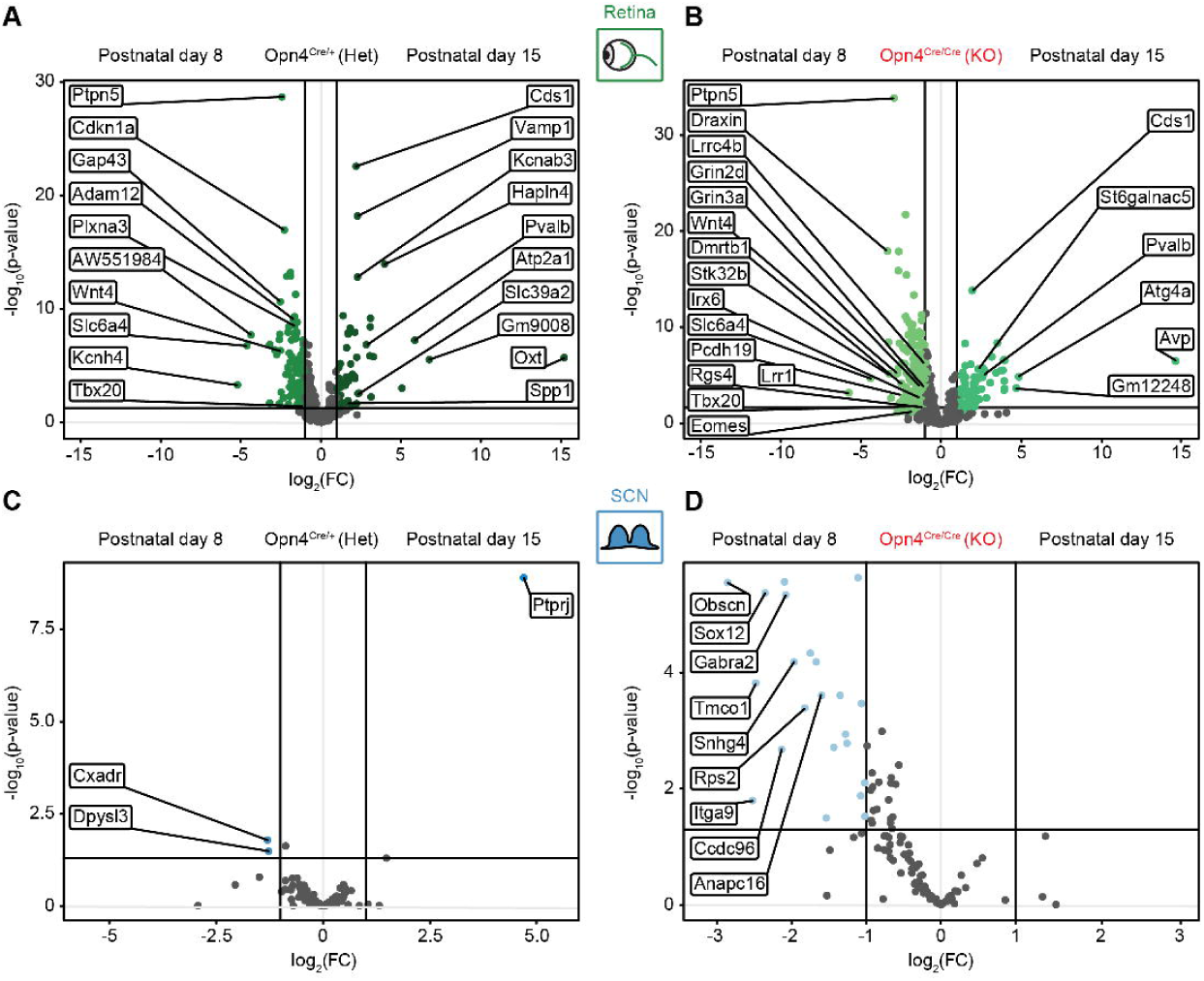
Temporal dynamics of compartment-specific gene expression in ipRGC soma and axon terminals before (P8) and after (P15) eye-opening. **(A-B)** Volcano plots showing differential gene expression within Het (A) and KO (B) retinal translatomes comparing postnatal day 8 and day 15. **(C-D)** Volcano plots showing differential gene expression within Het (C) and KO (D) axonal translatomes in the SCN comparing postnatal day 8 and day 15. Gray dots in all panels represent non-significant genes.

**Figure S5.**
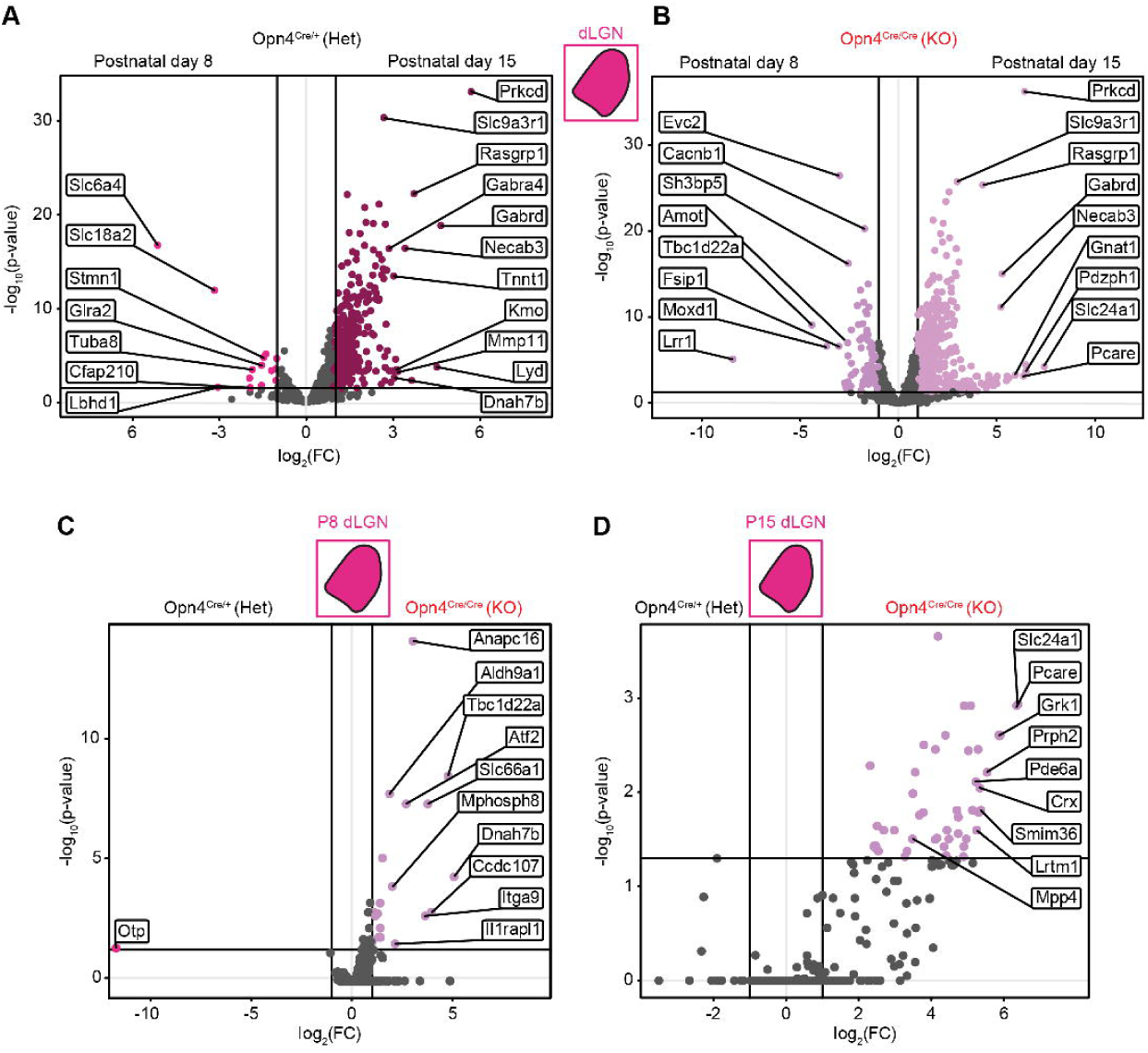
Translatome dynamics in the developing dorsal lateral geniculate nucleus following loss of melanopsin signaling. **(A-B)** Volcano plots showing differential gene expression within Het (A) and KO (B) dLGN translatomes comparing postnatal day 8 and day 15. **(C)** Volcano plot showing melanopsin dependent dLGN translatome at P8 (Het versus KO). **(D)** Volcano plot showing melanopsin dependent dLGN translatome at P15 (Het versus KO). Gray dots in all panels represent non-significant genes.

**Figure S6:**
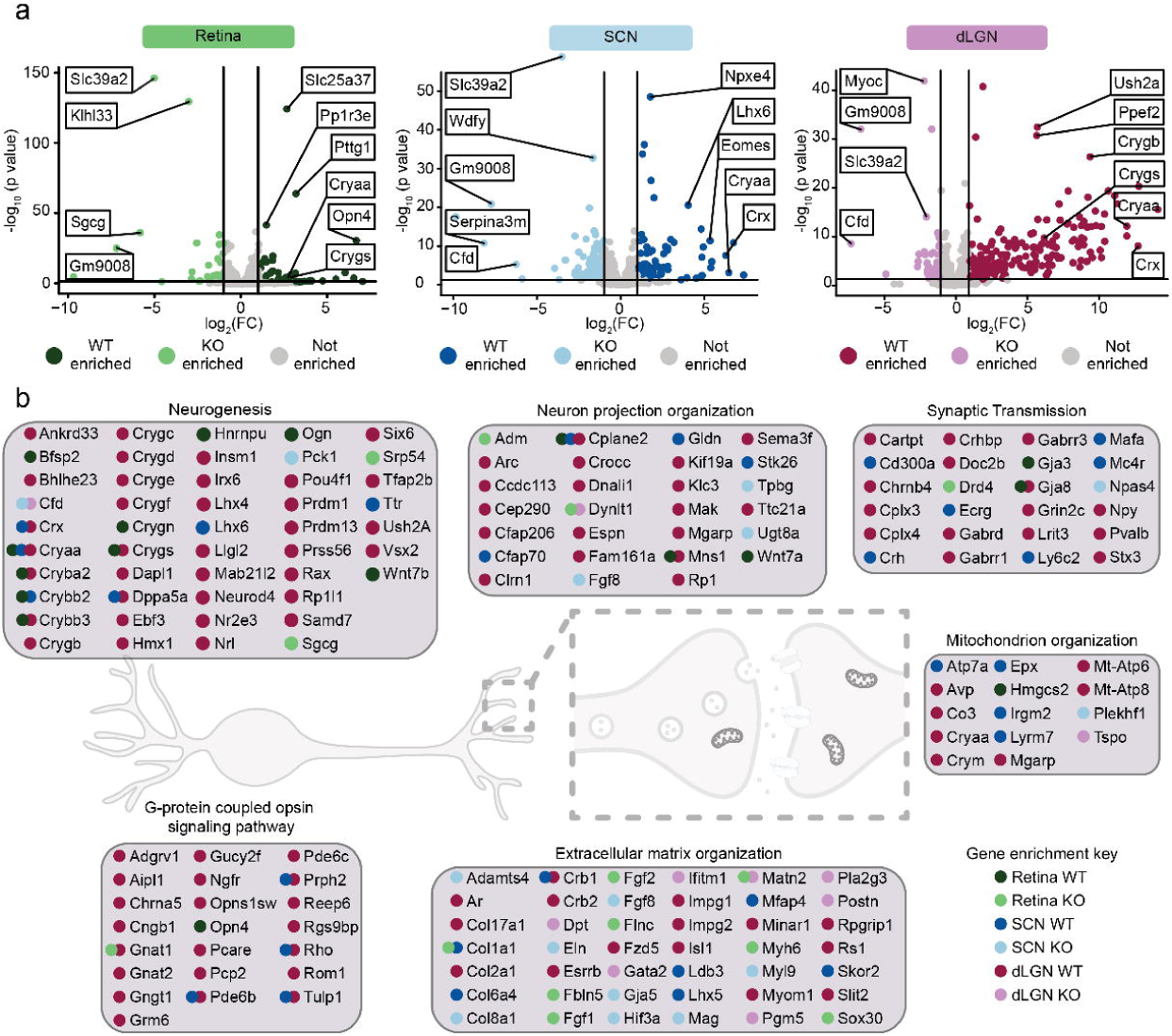
Gene ontology enrichment analysis of melanopsin-dependent differentially expressed genes in retina, SCN, and dLGN. (A) Volcano plots showing differential gene expression in retina (left, green), SCN (middle, blue), and dLGN (right, magenta) tissues comparing Opn4 WT-enriched versus KO-enriched genes at postnatal day 8. Dark colors indicate WT-enriched genes, light colors indicate KO-enriched genes, and gray indicates non-significant genes. (B) Detailed gene lists for major Gene Ontology categories organized by neuronal function. Colored dots indicate tissue enrichment: retina WT=dark green, retina KO=light green, SCN WT=dark blue, SCN KO=light blue, dLGN WT=dark magenta, dLGN KO=light magenta.

**Figure S7:**
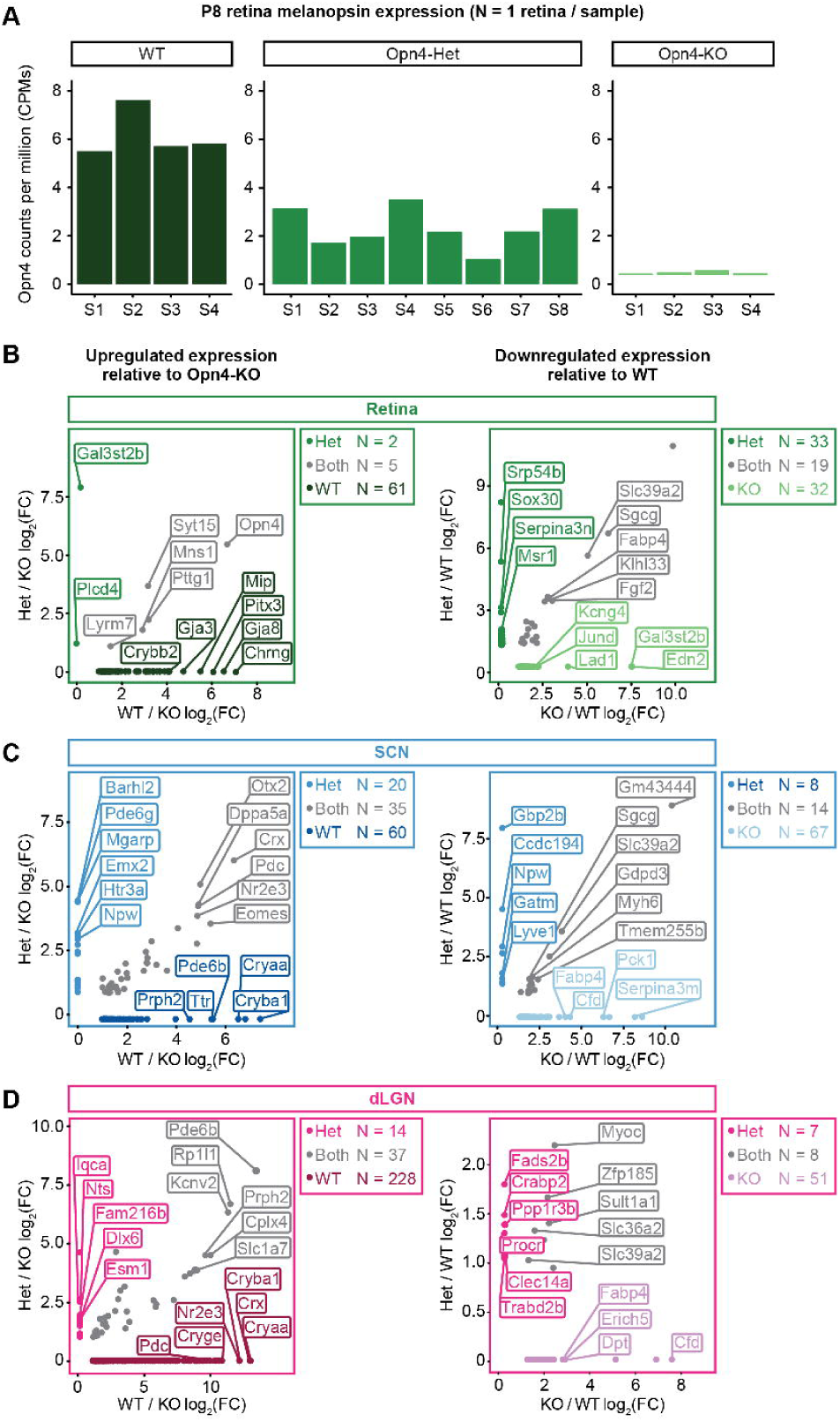
Melanopsin expression levels and differential gene expression across Opn4 genotypes in retina, SCN, and dLGN. (A) Bar graphs showing Opn4 expression levels (counts per million CPMs) from individual retinal samples at P8 across three genotypes: WT (samples S1-S4, consistently high expression), Opn4-Het (samples S1-S8, reduced expression), and Opn4-KO (samples S1-S4, near-zero expression). (B) Scatter plots for the retina showing upregulated expression relative to Opn4-KO tissue (left panel) for Het (y-axis) and WT (x-axis) samples. Downregulated expression relative to WT (right panel) is shown for Het (y-axis) and KO (x-axis) samples. (C) Similar analyses for SCN tissue showing upregulated genes relative to Opn4-KO (left panel) and downregulated genes relative to WT (right panel). (D) Similar analysis for dLGN tissue showing upregulated (left panel) and downregulated genes (right panel).

**Figure S8:**
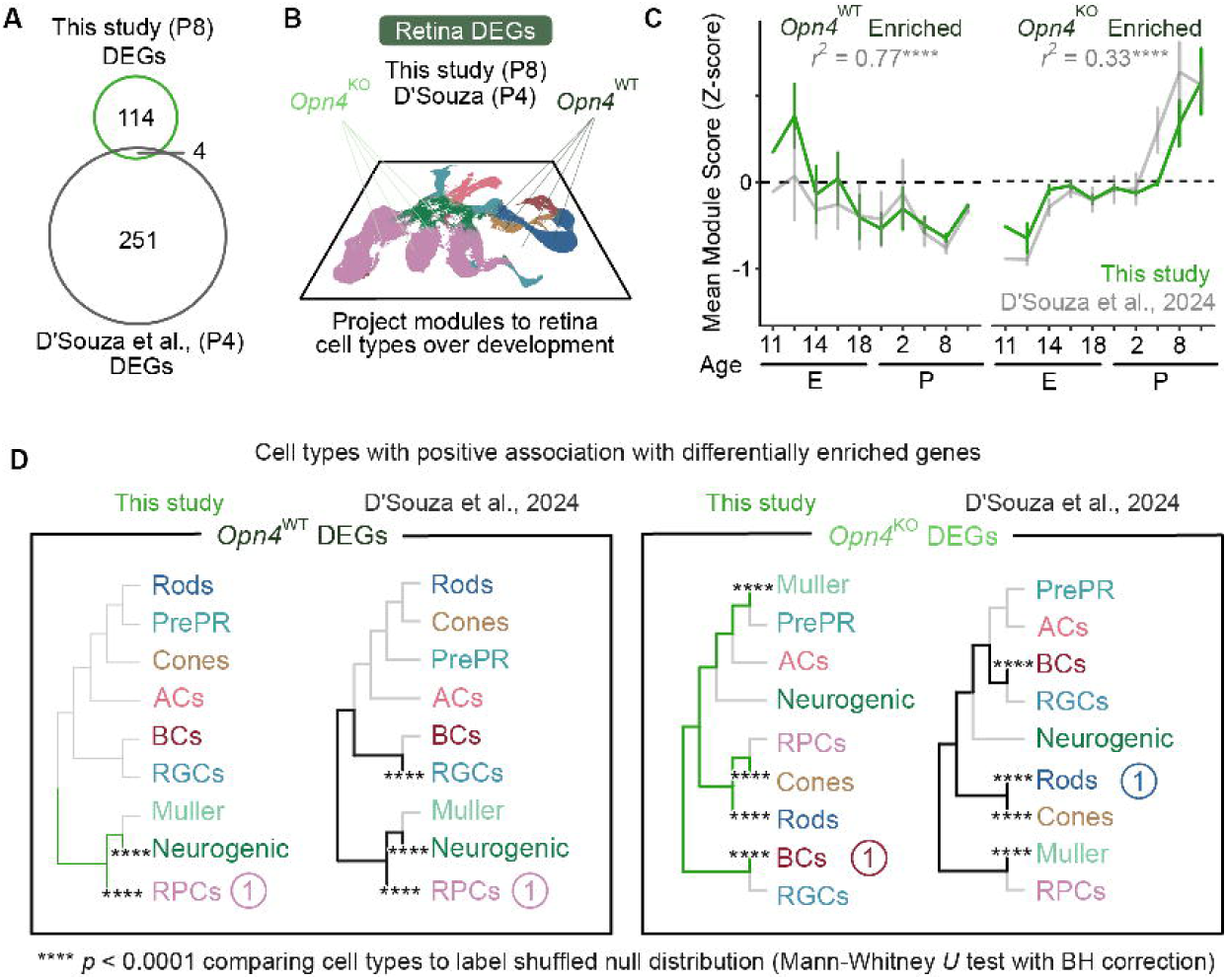
Comparison of melanopsin-dependent retinal gene expression across independent developmental datasets at P4 and P8. (A) Venn diagram showing overlap of differentially expressed genes between this study at P8 and D’Souza et al. at P4. (B) Module mapping for both datasets onto the Clark et al. 2019 retinal atlas. (C) Temporal analysis showing mean module score for Opn4 WT-enriched genes (gray line, this study) and Opn4 KO-enriched genes (green line, this study) averaged across developmental ages, compared with D’Souza et al. data (gray dashed line). ****p < 0.0001. (D) Module enriched cell types in both studies. Left panels show WT enrichment patterns, right panels show KO enrichment patterns. (1) indicates the cell class with the highest module score enrichment. ****p < 0.0001, Mann-Whitney U test with Benjamini-Hochberg correction comparing mean module score between cell class and a label shuffled data set.

**Figure S9:**
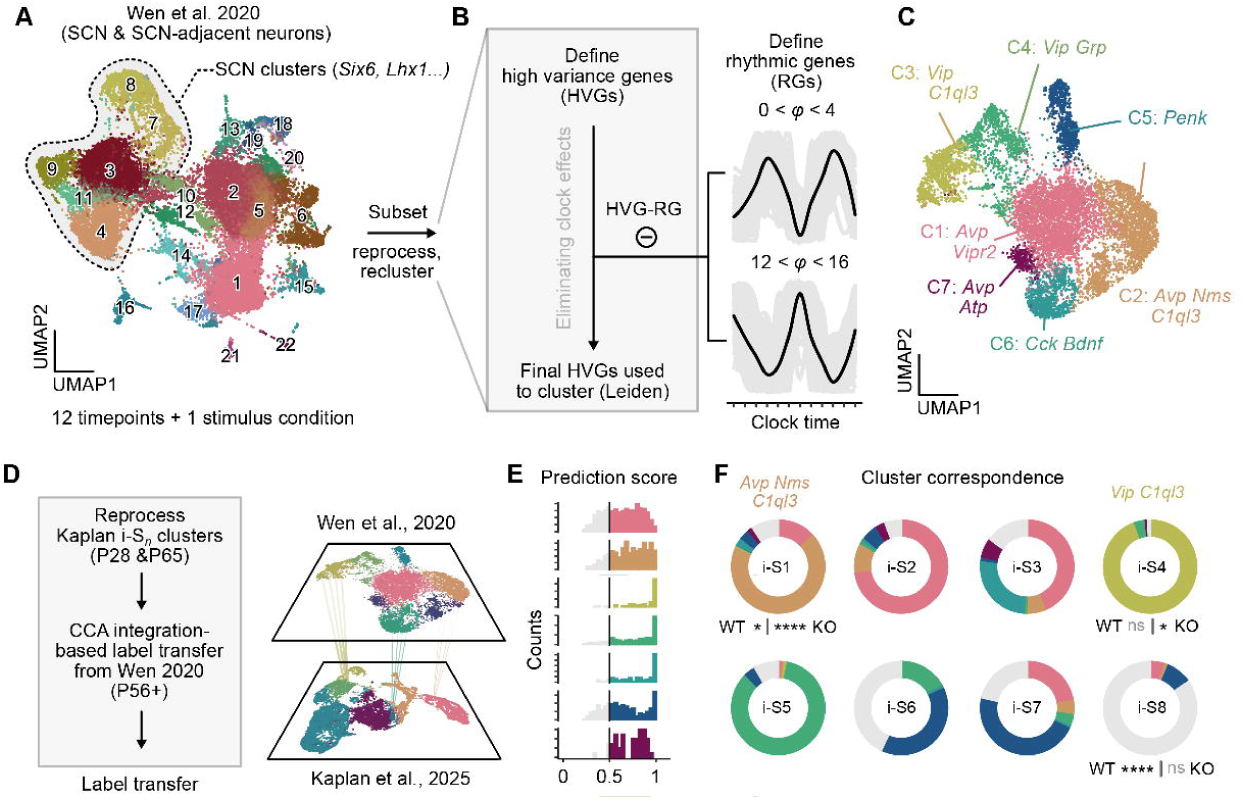
SCN cell types impacted by melanopsin deletion. (A) UMAP embedding of cells from Wen et al., 2020. Dashed line indicates SCN-specific clusters with cluster numbers labeled 1-22. (B) Reanalysis procedure to cluster SCN neurons, regressing out high variance genes displaying strong circadian rhythmicity (defined by JTK analysis). (C) UMAP projection of final cluster annotations showing 7 distinct clusters labeled C1-C7 with representative marker genes: three Avp clusters (C1, 2, 7), two Vip clusters (C3, 4), and single Cck and Penk clusters. (D) Processing workflow CCA integration-based label transfer from Wen et al. 2020 to Kaplan et al. 2025. (E) Histograms showing prediction scores of label transfer confidence for all seven clusters (threshold 0.5). (F) Donut plots showing cluster correspondence between Kaplan et al. 2025 SCN nomenclature (inner labels: i-S1 through i-S8) and Wen et al. 2020 clusters (C1-C7 colors in pie segments and molecular labels above). *p < 0.05, ****p < 0.0001, Mann-Whitney U test with Benjamini-Hochberg correction.

