## Supplementary material for "Melanopsin regulates axonal translation underlying retinohypothalamic circuit assembly": Table_S1_bulk_RNA_seq_significant_genes

**Contrast numerator: WT_dLGN. Contrast denominator: KO_dLGN.**

| ensembl_gene_id | external_gene_name | Significance | description | baseMean | logFC | adj.P.Val |
|---|---|---|---|---|---|---|
| ENSMUSG00000061780 | Cfd | Enriched in KO dLGN | complement factor D (adipsin) [Source:MGI Symbol;Acc:MGI:87931] | 65.58 | -7.33 | 2.494E-09 |
| ENSMUSG00000072476 | Gm9008 | Enriched in KO dLGN | predicted pseudogene 9008 [Source:MGI Symbol;Acc:MGI:3644000] | 170.6 | -6.652 | 7.736E-33 |
| ENSMUSG00000105243 | Gm43444 | Enriched in KO dLGN | predicted gene 43444 [Source:MGI Symbol;Acc:MGI:5663581] | 86.84 | -4.872 | 0.004341 |
| ENSMUSG00000026574 | Dpt | Enriched in KO dLGN | dermatopontin [Source:MGI Symbol;Acc:MGI:1928392] | 101.7 | -2.656 | 0.000126 |
| ENSMUSG00000044726 | Erich5 | Enriched in KO dLGN | glutamate rich 5 [Source:MGI Symbol;Acc:MGI:2447772] | 46.06 | -2.61 | 0.00001046 |
| ENSMUSG00000062515 | Fabp4 | Enriched in KO dLGN | fatty acid binding protein 4, adipocyte [Source:MGI Symbol;Acc:MGI:88038] | 255.4 | -2.55 | 3.422E-07 |
| ENSMUSG00000026697 | Myoc | Enriched in KO dLGN | myocilin [Source:MGI Symbol;Acc:MGI:1202864] | 1087 | -2.192 | 1.018E-42 |
| ENSMUSG00000027559 | Car3 | Enriched in KO dLGN | carbonic anhydrase 3 [Source:MGI Symbol;Acc:MGI:88270] | 216.9 | -2.168 | 1.017E-07 |
| ENSMUSG00000047420 | Fam180a | Enriched in KO dLGN | family with sequence similarity 180, member A [Source:MGI Symbol;Acc:MGI:3039626] | 113.5 | -2.118 | 4.476E-07 |
| ENSMUSG00000072572 | Slc39a2 | Enriched in KO dLGN | solute carrier family 39 (zinc transporter), member 2 [Source:MGI Symbol;Acc:MGI:2684326] | 587.9 | -2.029 | 7.447E-15 |
| ENSMUSG00000022074 | Tnfrsf10b | Enriched in KO dLGN | tumor necrosis factor receptor superfamily, member 10b [Source:MGI Symbol;Acc:MGI:1341090] | 71.96 | -1.982 | 0.00002125 |
| ENSMUSG00000020264 | Slc36a2 | Enriched in KO dLGN | solute carrier family 36 (proton/amino acid symporter), member 2 [Source:MGI Symbol;Acc:MGI:1891430] | 52.84 | -1.952 | 0.0006368 |
| ENSMUSG00000020884 | Asgr1 | Enriched in KO dLGN | asialoglycoprotein receptor 1 [Source:MGI Symbol;Acc:MGI:88081] | 57.45 | -1.945 | 0.003303 |
| ENSMUSG00000030711 | Sult1a1 | Enriched in KO dLGN | sulfotransferase family 1A, phenol-preferring, member 1 [Source:MGI Symbol;Acc:MGI:102896] | 117.9 | -1.88 | 0.0001007 |
| ENSMUSG00000102439 | Flg | Enriched in KO dLGN | filaggrin [Source:MGI Symbol;Acc:MGI:95553] | 47.62 | -1.853 | 0.0003537 |
| ENSMUSG00000054641 | Mmrn1 | Enriched in KO dLGN | multimerin 1 [Source:MGI Symbol;Acc:MGI:1918195] | 51.59 | -1.837 | 0.0317 |
| ENSMUSG00000073406 | H2-Bl | Enriched in KO dLGN | histocompatibility 2, blastocyst [Source:MGI Symbol;Acc:MGI:892004] | 617.1 | -1.805 | 0.005043 |
| ENSMUSG00000079396 | Gm3411 | Enriched in KO dLGN | predicted gene 3411 [Source:MGI Symbol;Acc:MGI:3781589] | 63.58 | -1.732 | 0.0006305 |
| ENSMUSG00000008845 | Cd163 | Enriched in KO dLGN | CD163 antigen [Source:MGI Symbol;Acc:MGI:2135946] | 213.8 | -1.694 | 2.987E-07 |
| ENSMUSG00000055235 | Wdr86 | Enriched in KO dLGN | WD repeat domain 86 [Source:MGI Symbol;Acc:MGI:1915466] | 188 | -1.69 | 0.00815 |
| ENSMUSG00000021091 | Serpina3n | Enriched in KO dLGN | serine (or cysteine) peptidase inhibitor, clade A, member 3N [Source:MGI Symbol;Acc:MGI:105045] | 1451 | -1.654 | 7.736E-33 |
| ENSMUSG00000023046 | Igfbp6 | Enriched in KO dLGN | insulin-like growth factor binding protein 6 [Source:MGI Symbol;Acc:MGI:96441] | 125.6 | -1.602 | 0.0004609 |
| ENSMUSG00000031351 | Zfp185 | Enriched in KO dLGN | zinc finger protein 185 [Source:MGI Symbol;Acc:MGI:108095] | 48.85 | -1.578 | 0.002459 |
| ENSMUSG00000024650 | Slc22a6 | Enriched in KO dLGN | solute carrier family 22 (organic anion transporter), member 6 [Source:MGI Symbol;Acc:MGI:892001] | 371.7 | -1.536 | 1.554E-07 |
| ENSMUSG00000096255 | Dynlt1b | Enriched in KO dLGN | dynein light chain Tctex-type 1B [Source:MGI Symbol;Acc:MGI:98643] | 104.6 | -1.436 | 0.007071 |
| ENSMUSG00000031070 | Mrgprf | Enriched in KO dLGN | MAS-related GPR, member F [Source:MGI Symbol;Acc:MGI:2384823] | 81.14 | -1.342 | 0.03124 |
| ENSMUSG00000074971 | Fibin | Enriched in KO dLGN | fin bud initiation factor homolog (zebrafish) [Source:MGI Symbol;Acc:MGI:1914856] | 77.92 | -1.339 | 0.01325 |
| ENSMUSG00000041736 | Tspo | Enriched in KO dLGN | translocator protein [Source:MGI Symbol;Acc:MGI:88222] | 86.27 | -1.338 | 0.004915 |
| ENSMUSG00000025150 | Cbr2 | Enriched in KO dLGN | carbonyl reductase 2 [Source:MGI Symbol;Acc:MGI:107200] | 64.52 | -1.327 | 0.04467 |
| ENSMUSG00000048572 | Tmem252 | Enriched in KO dLGN | transmembrane protein 252 [Source:MGI Symbol;Acc:MGI:3583948] | 143.9 | -1.322 | 0.001153 |
| ENSMUSG00000029843 | Slc13a4 | Enriched in KO dLGN | solute carrier family 13 (sodium/sulfate symporters), member 4 [Source:MGI Symbol;Acc:MGI:2442367] | 465.6 | -1.307 | 0.0001361 |
| ENSMUSG00000093985 | Gm10406 | Enriched in KO dLGN | predicted gene 10406 [Source:MGI Symbol;Acc:MGI:3711272] | 206 | -1.286 | 0.0000169 |
| ENSMUSG00000030787 | Lyve1 | Enriched in KO dLGN | lymphatic vessel endothelial hyaluronan receptor 1 [Source:MGI Symbol;Acc:MGI:2136348] | 280.2 | -1.251 | 0.006323 |
| ENSMUSG00000022324 | Matn2 | Enriched in KO dLGN | matrilin 2 [Source:MGI Symbol;Acc:MGI:109613] | 337.6 | -1.232 | 1.545E-11 |
| ENSMUSG00000038216 | Pnmt | Enriched in KO dLGN | phenylethanolamine-N-methyltransferase [Source:MGI Symbol;Acc:MGI:97724] | 82.71 | -1.225 | 0.01847 |
| ENSMUSG00000021943 | Gdf10 | Enriched in KO dLGN | growth differentiation factor 10 [Source:MGI Symbol;Acc:MGI:95684] | 423.7 | -1.216 | 4.235E-07 |
| ENSMUSG00000022025 | Cnmd | Enriched in KO dLGN | chondromodulin [Source:MGI Symbol;Acc:MGI:1341171] | 165.3 | -1.213 | 0.0008145 |
| ENSMUSG00000048489 | Depp1 | Enriched in KO dLGN | DEPP1 autophagy regulator [Source:MGI Symbol;Acc:MGI:1918730] | 73.85 | -1.205 | 0.01188 |
| ENSMUSG00000015053 | Gata2 | Enriched in KO dLGN | GATA binding protein 2 [Source:MGI Symbol;Acc:MGI:95662] | 166 | -1.172 | 0.001088 |
| ENSMUSG00000015090 | Ptgds | Enriched in KO dLGN | prostaglandin D2 synthase (brain) [Source:MGI Symbol;Acc:MGI:99261] | 9324 | -1.169 | 0.04807 |
| ENSMUSG00000051439 | Cd14 | Enriched in KO dLGN | CD14 antigen [Source:MGI Symbol;Acc:MGI:88318] | 72.9 | -1.168 | 0.009249 |
| ENSMUSG00000026656 | Fcgr2b | Enriched in KO dLGN | Fc receptor, IgG, low affinity IIb [Source:MGI Symbol;Acc:MGI:95499] | 154.4 | -1.156 | 0.00613 |
| ENSMUSG00000036814 | Slc6a20a | Enriched in KO dLGN | solute carrier family 6 (neurotransmitter transporter), member 20A [Source:MGI Symbol;Acc:MGI:2143217] | 584.1 | -1.152 | 0.00009072 |
| ENSMUSG00000066058 | Cldn19 | Enriched in KO dLGN | claudin 19 [Source:MGI Symbol;Acc:MGI:3033992] | 220.4 | -1.141 | 0.00006986 |
| ENSMUSG00000041731 | Pgm5 | Enriched in KO dLGN | phosphoglucomutase 5 [Source:MGI Symbol;Acc:MGI:1925668] | 138.8 | -1.137 | 0.002891 |
| ENSMUSG00000032419 | Tbx18 | Enriched in KO dLGN | T-box18 [Source:MGI Symbol;Acc:MGI:1923615] | 456.9 | -1.129 | 0.0006722 |
| ENSMUSG00000038457 | Tmem255b | Enriched in KO dLGN | transmembrane protein 255B [Source:MGI Symbol;Acc:MGI:2685533] | 399.6 | -1.127 | 1.078E-08 |
| ENSMUSG00000034579 | Pla2g3 | Enriched in KO dLGN | phospholipase A2, group III [Source:MGI Symbol;Acc:MGI:2444945] | 3333 | -1.116 | 1.493E-10 |
| ENSMUSG00000073940 | Hbb-bt | Enriched in KO dLGN | hemoglobin, beta adult t chain [Source:MGI Symbol;Acc:MGI:5474850] | 401 | -1.111 | 4.447E-09 |
| ENSMUSG00000029546 | Uncx | Enriched in KO dLGN | UNC homeobox [Source:MGI Symbol;Acc:MGI:108013] | 114.6 | -1.1 | 0.03124 |
| ENSMUSG00000030108 | Slc6a13 | Enriched in KO dLGN | solute carrier family 6 (neurotransmitter transporter, GABA), member 13 [Source:MGI Symbol;Acc:MGI:95629] | 616.1 | -1.097 | 0.00282 |
| ENSMUSG00000027750 | Postn | Enriched in KO dLGN | periostin, osteoblast specific factor [Source:MGI Symbol;Acc:MGI:1926321] | 467.9 | -1.085 | 0.001469 |
| ENSMUSG00000113255 | Tes3-ps | Enriched in KO dLGN | testis derived transcript 3, pseudogene [Source:MGI Symbol;Acc:MGI:3582925] | 87.89 | -1.074 | 0.02434 |
| ENSMUSG00000073643 | Wdfy1 | Enriched in KO dLGN | WD repeat and FYVE domain containing 1 [Source:MGI Symbol;Acc:MGI:1916618] | 1689 | -1.057 | 2.929E-07 |
| ENSMUSG00000031750 | Il34 | Enriched in KO dLGN | interleukin 34 [Source:MGI Symbol;Acc:MGI:1923777] | 248.7 | -1.047 | 0.001249 |
| ENSMUSG00000025784 | Clec3b | Enriched in KO dLGN | C-type lectin domain family 3, member b [Source:MGI Symbol;Acc:MGI:104540] | 834.1 | -1.038 | 0.0008764 |
| ENSMUSG00000025491 | Ifitm1 | Enriched in KO dLGN | interferon induced transmembrane protein 1 [Source:MGI Symbol;Acc:MGI:1915963] | 105.9 | -1.023 | 0.02881 |
| ENSMUSG00000043424 | Eif3j2 | Enriched in KO dLGN | eukaryotic translation initiation factor 3, subunit J2 [Source:MGI Symbol;Acc:MGI:3704486] | 240.8 | -1.02 | 0.03389 |
| ENSMUSG00000035686 | Thrsp | Enriched in KO dLGN | thyroid hormone responsive [Source:MGI Symbol;Acc:MGI:109126] | 191.7 | -1.004 | 0.008242 |
| ENSMUSG00000021996 | Esd | Enriched in WT dLGN | esterase D/formylglutathione hydrolase [Source:MGI Symbol;Acc:MGI:95421] | 1867 | 1.013 | 3.801E-17 |
| ENSMUSG00000020887 | A230052G05Rik | Enriched in WT dLGN | RIKEN cDNA A230052G05 gene [Source:MGI Symbol;Acc:MGI:3045239] | 88.99 | 1.022 | 0.03137 |
| ENSMUSG00000024049 | Myom1 | Enriched in WT dLGN | myomesin 1 [Source:MGI Symbol;Acc:MGI:1341430] | 299.5 | 1.027 | 0.003351 |
| ENSMUSG00000053219 | Raet1e | Enriched in WT dLGN | retinoic acid early transcript 1E [Source:MGI Symbol;Acc:MGI:2675273] | 360.5 | 1.052 | 0.000009073 |
| ENSMUSG00000034684 | Sema3f | Enriched in WT dLGN | sema domain, immunoglobulin domain (Ig), short basic domain, secreted, (semaphorin) 3F [Source:MGI Symbol;Acc:MGI:109634 | 629.3 | 1.061 | 0.00002199 |
| ENSMUSG00000036192 | Rorb | Enriched in WT dLGN | RAR-related orphan receptor beta [Source:MGI Symbol;Acc:MGI:1343464] | 2871 | 1.071 | 0.000001417 |
| ENSMUSG00000032251 | Irak1bp1 | Enriched in WT dLGN | interleukin-1 receptor-associated kinase 1 binding protein 1 [Source:MGI Symbol;Acc:MGI:1929475] | 882.6 | 1.074 | 5.915E-07 |
| ENSMUSG00000091345 | Col6a5 | Enriched in WT dLGN | collagen, type VI, alpha 5 [Source:MGI Symbol;Acc:MGI:3648134] | 65 | 1.081 | 0.03222 |
| ENSMUSG00000035299 | Mid1 | Enriched in WT dLGN | midline 1 [Source:MGI Symbol;Acc:MGI:1100537] | 240.3 | 1.082 | 0.000408 |
| ENSMUSG00000040543 | Pitpnm3 | Enriched in WT dLGN | PITPNM family member 3 [Source:MGI Symbol;Acc:MGI:2685726] | 2356 | 1.092 | 0.000005141 |
| ENSMUSG00000114995 | Gm49284 | Enriched in WT dLGN | predicted gene, 49284 [Source:MGI Symbol;Acc:MGI:6118771] | 115.4 | 1.095 | 0.01542 |
| ENSMUSG00000037627 | Rgs22 | Enriched in WT dLGN | regulator of G-protein signalling 22 [Source:MGI Symbol;Acc:MGI:3613651] | 129.2 | 1.113 | 0.03352 |
| ENSMUSG00000017692 | Rhbdl3 | Enriched in WT dLGN | rhomboid like 3 [Source:MGI Symbol;Acc:MGI:2179276] | 414.3 | 1.137 | 0.001734 |
| ENSMUSG00000031558 | Slit2 | Enriched in WT dLGN | slit guidance ligand 2 [Source:MGI Symbol;Acc:MGI:1315205] | 706 | 1.145 | 0.0004768 |
| ENSMUSG00000020806 | Rhbdf2 | Enriched in WT dLGN | rhomboid 5 homolog 2 [Source:MGI Symbol;Acc:MGI:2442473] | 148.2 | 1.152 | 0.01141 |
| ENSMUSG00000096039 | D830030K20Rik | Enriched in WT dLGN | RIKEN cDNA D830030K20 gene [Source:MGI Symbol;Acc:MGI:2443830] | 81.94 | 1.157 | 0.0405 |
| ENSMUSG00000064354 | mt-Co2 | Enriched in WT dLGN | mitochondrially encoded cytochrome c oxidase II [Source:MGI Symbol;Acc:MGI:102503] | 188.7 | 1.183 | 0.0007845 |
| ENSMUSG00000040164 | Kcns1 | Enriched in WT dLGN | K+ voltage-gated channel, subfamily S, 1 [Source:MGI Symbol;Acc:MGI:1197019] | 219.5 | 1.184 | 0.0006672 |
| ENSMUSG00000052407 | Ccdc171 | Enriched in WT dLGN | coiled-coil domain containing 171 [Source:MGI Symbol;Acc:MGI:1922152] | 74.14 | 1.184 | 0.01158 |
| ENSMUSG00000032221 | Mns1 | Enriched in WT dLGN | meiosis-specific nuclear structural protein 1 [Source:MGI Symbol;Acc:MGI:107933] | 204.4 | 1.186 | 0.00001308 |
| ENSMUSG00000019971 | Cep290 | Enriched in WT dLGN | centrosomal protein 290 [Source:MGI Symbol;Acc:MGI:2384917] | 585.8 | 1.189 | 0.001519 |
| ENSMUSG00000095440 | Fignl2 | Enriched in WT dLGN | fidgetin-like 2 [Source:MGI Symbol;Acc:MGI:3646919] | 138.2 | 1.19 | 0.007953 |
| ENSMUSG00000012126 | Ubxn11 | Enriched in WT dLGN | UBX domain protein 11 [Source:MGI Symbol;Acc:MGI:1914836] | 217.1 | 1.193 | 0.0166 |
| ENSMUSG00000033405 | Nudt15 | Enriched in WT dLGN | nudix (nucleoside diphosphate linked moiety X)-type motif 15 [Source:MGI Symbol;Acc:MGI:2443366] | 81.86 | 1.194 | 0.01807 |
| ENSMUSG00000019996 | Map7 | Enriched in WT dLGN | microtubule-associated protein 7 [Source:MGI Symbol;Acc:MGI:1328328] | 1645 | 1.196 | 0.000006597 |
| ENSMUSG00000050786 | Ccdc126 | Enriched in WT dLGN | coiled-coil domain containing 126 [Source:MGI Symbol;Acc:MGI:1889376] | 496.1 | 1.196 | 0.005121 |
| ENSMUSG00000027210 | Meis2 | Enriched in WT dLGN | Meis homeobox 2 [Source:MGI Symbol;Acc:MGI:108564] | 1133 | 1.203 | 3.218E-09 |
| ENSMUSG00000029335 | Bmp3 | Enriched in WT dLGN | bone morphogenetic protein 3 [Source:MGI Symbol;Acc:MGI:88179] | 221.6 | 1.213 | 0.002497 |
| ENSMUSG00000026896 | Ifih1 | Enriched in WT dLGN | interferon induced with helicase C domain 1 [Source:MGI Symbol;Acc:MGI:1918836] | 63.17 | 1.215 | 0.01192 |
| ENSMUSG00000028919 | Arhgef19 | Enriched in WT dLGN | Rho guanine nucleotide exchange factor (GEF) 19 [Source:MGI Symbol;Acc:MGI:1925912] | 125.5 | 1.215 | 0.006555 |
| ENSMUSG00000021255 | Esrrb | Enriched in WT dLGN | estrogen related receptor, beta [Source:MGI Symbol;Acc:MGI:1346832] | 185.2 | 1.216 | 0.0003039 |
| ENSMUSG00000026247 | Ecel1 | Enriched in WT dLGN | endothelin converting enzyme-like 1 [Source:MGI Symbol;Acc:MGI:1343461] | 758.3 | 1.226 | 0.0004744 |
| ENSMUSG00000074252 | Gm10654 | Enriched in WT dLGN | predicted gene 10654 [Source:MGI Symbol;Acc:MGI:3643366] | 70.24 | 1.227 | 0.04938 |
| ENSMUSG00000021904 | Sema3g | Enriched in WT dLGN | sema domain, immunoglobulin domain (Ig), short basic domain, secreted, (semaphorin) 3G [Source:MGI Symbol;Acc:MGI:304124 | 346.1 | 1.23 | 0.00003546 |
| ENSMUSG00000033966 | Cdkl4 | Enriched in WT dLGN | cyclin-dependent kinase-like 4 [Source:MGI Symbol;Acc:MGI:3587025] | 207.1 | 1.237 | 0.03134 |
| ENSMUSG00000046532 | Ar | Enriched in WT dLGN | androgen receptor [Source:MGI Symbol;Acc:MGI:88064] | 86.73 | 1.243 | 0.03352 |
| ENSMUSG00000037995 | Igsf9 | Enriched in WT dLGN | immunoglobulin superfamily, member 9 [Source:MGI Symbol;Acc:MGI:2135283] | 265.6 | 1.246 | 0.00613 |
| ENSMUSG00000058498 | Rnf207 | Enriched in WT dLGN | ring finger protein 207 [Source:MGI Symbol;Acc:MGI:2684989] | 154.5 | 1.249 | 0.0001286 |
| ENSMUSG00000064358 | mt-Co3 | Enriched in WT dLGN | mitochondrially encoded cytochrome c oxidase III [Source:MGI Symbol;Acc:MGI:102502] | 85.41 | 1.249 | 0.03298 |
| ENSMUSG00000035513 | Ntng2 | Enriched in WT dLGN | netrin G2 [Source:MGI Symbol;Acc:MGI:2159341] | 600.5 | 1.25 | 0.001143 |
| ENSMUSG00000070880 | Gad1 | Enriched in WT dLGN | glutamate decarboxylase 1 [Source:MGI Symbol;Acc:MGI:95632] | 9591 | 1.255 | 0.02247 |
| ENSMUSG00000042282 | Gucy2f | Enriched in WT dLGN | guanylate cyclase 2f [Source:MGI Symbol;Acc:MGI:105119] | 119.8 | 1.277 | 0.002078 |
| ENSMUSG00000028883 | Sema3a | Enriched in WT dLGN | sema domain, immunoglobulin domain (Ig), short basic domain, secreted, (semaphorin) 3A [Source:MGI Symbol;Acc:MGI:107558 | 201.8 | 1.297 | 0.000002407 |
| ENSMUSG00000046861 | Hectd3 | Enriched in WT dLGN | HECT domain E3 ubiquitin protein ligase 3 [Source:MGI Symbol;Acc:MGI:1923858] | 1088 | 1.298 | 0.000004062 |
| ENSMUSG00000020912 | Krt12 | Enriched in WT dLGN | keratin 12 [Source:MGI Symbol;Acc:MGI:96687] | 57.39 | 1.299 | 0.03538 |
| ENSMUSG00000020481 | Ankrd36 | Enriched in WT dLGN | ankyrin repeat domain 36 [Source:MGI Symbol;Acc:MGI:1923639] | 128.2 | 1.303 | 0.0001027 |
| ENSMUSG00000036395 | Glb1l2 | Enriched in WT dLGN | galactosidase, beta 1-like 2 [Source:MGI Symbol;Acc:MGI:2388283] | 94.46 | 1.316 | 0.02963 |
| ENSMUSG00000039313 | Minar1 | Enriched in WT dLGN | membrane integral NOTCH2 associated receptor 1 [Source:MGI Symbol;Acc:MGI:2667167] | 664.8 | 1.357 | 0.00007734 |
| ENSMUSG00000029561 | Oasl2 | Enriched in WT dLGN | 2'-5' oligoadenylate synthetase-like 2 [Source:MGI Symbol;Acc:MGI:1344390] | 132 | 1.368 | 0.03468 |
| ENSMUSG00000069170 | Adgrv1 | Enriched in WT dLGN | adhesion G protein-coupled receptor V1 [Source:MGI Symbol;Acc:MGI:1274784] | 217.7 | 1.371 | 0.0004084 |
| ENSMUSG00000043068 | Fam89a | Enriched in WT dLGN | family with sequence similarity 89, member A [Source:MGI Symbol;Acc:MGI:1916877] | 86.88 | 1.376 | 0.03955 |
| ENSMUSG00000021009 | Ptpn21 | Enriched in WT dLGN | protein tyrosine phosphatase, non-receptor type 21 [Source:MGI Symbol;Acc:MGI:1344406] | 515.5 | 1.391 | 0.00008276 |
| ENSMUSG00000028294 | Cfap206 | Enriched in WT dLGN | cilia and flagella associated protein 206 [Source:MGI Symbol;Acc:MGI:1916579] | 80.36 | 1.399 | 0.0299 |
| ENSMUSG00000053846 | Lipg | Enriched in WT dLGN | lipase, endothelial [Source:MGI Symbol;Acc:MGI:1341803] | 50.8 | 1.411 | 0.01811 |
| ENSMUSG00000055865 | Tafa3 | Enriched in WT dLGN | TAFA chemokine like family member 3 [Source:MGI Symbol;Acc:MGI:3046463] | 282.2 | 1.419 | 0.0002835 |
| ENSMUSG00000022602 | Arc | Enriched in WT dLGN | activity regulated cytoskeletal-associated protein [Source:MGI Symbol;Acc:MGI:88067] | 358.2 | 1.42 | 0.001358 |
| ENSMUSG00000026610 | Esrrg | Enriched in WT dLGN | estrogen-related receptor gamma [Source:MGI Symbol;Acc:MGI:1347056] | 3135 | 1.432 | 0.002078 |
| ENSMUSG00000073733 | Cplane2 | Enriched in WT dLGN | ciliogenesis and planar polarity effector 2 [Source:MGI Symbol;Acc:MGI:1923416] | 63.36 | 1.439 | 0.003433 |
| ENSMUSG00000079941 | Cox5b-ps | Enriched in WT dLGN | cytochrome c oxidase subunit 5B, pseudogene [Source:MGI Symbol;Acc:MGI:3649411] | 678.1 | 1.445 | 3.354E-31 |
| ENSMUSG00000073530 | Pappa2 | Enriched in WT dLGN | pappalysin 2 [Source:MGI Symbol;Acc:MGI:3051647] | 85.4 | 1.457 | 0.02319 |
| ENSMUSG00000000120 | Ngfr | Enriched in WT dLGN | nerve growth factor receptor (TNFR superfamily, member 16) [Source:MGI Symbol;Acc:MGI:97323] | 614.8 | 1.461 | 0.00001683 |
| ENSMUSG00000078963 | Hsbp1l1 | Enriched in WT dLGN | heat shock factor binding protein 1-like 1 [Source:MGI Symbol;Acc:MGI:1913505] | 162.8 | 1.464 | 0.0002156 |
| ENSMUSG00000040860 | Crocc | Enriched in WT dLGN | ciliary rootlet coiled-coil, rootletin [Source:MGI Symbol;Acc:MGI:3529431] | 1247 | 1.505 | 0.01542 |
| ENSMUSG00000022860 | Chodl | Enriched in WT dLGN | chondrolectin [Source:MGI Symbol;Acc:MGI:2179069] | 83.01 | 1.508 | 0.00446 |
| ENSMUSG00000020734 | Grin2c | Enriched in WT dLGN | glutamate receptor, ionotropic, NMDA2C (epsilon 3) [Source:MGI Symbol;Acc:MGI:95822] | 103.1 | 1.519 | 0.04354 |
| ENSMUSG00000038135 | Crygn | Enriched in WT dLGN | crystallin, gamma N [Source:MGI Symbol;Acc:MGI:2449167] | 58.25 | 1.524 | 0.02123 |

| | | | | | | |
|---|---|---|---|---|---|---|
| ENSMUSG00000049811 | Fam161a | Enriched in WT dLGN | family with sequence similarity 161, member A [Source:MGI Symbol;Acc:MGI:1921123] | 501.7 | 1.531 | 0.04315 |
| ENSMUSG00000035504 | Reep6 | Enriched in WT dLGN | receptor accessory protein 6 [Source:MGI Symbol;Acc:MGI:1917585] | 829.2 | 1.537 | 0.01715 |
| ENSMUSG00000020848 | Doc2b | Enriched in WT dLGN | double C2, beta [Source:MGI Symbol;Acc:MGI:1100497] | 1999 | 1.558 | 0.002876 |
| ENSMUSG00000020415 | Pttg1 | Enriched in WT dLGN | pituitary tumor-transforming gene 1 [Source:MGI Symbol;Acc:MGI:1353578] | 381.6 | 1.56 | 2.203E-14 |
| ENSMUSG00000064357 | mt-Atp6 | Enriched in WT dLGN | mitochondrially encoded ATP synthase 6 [Source:MGI Symbol;Acc:MGI:99927] | 57.8 | 1.587 | 0.008751 |
| ENSMUSG00000041488 | Stx3 | Enriched in WT dLGN | syntaxin 3 [Source:MGI Symbol;Acc:MGI:103077] | 1821 | 1.662 | 0.01784 |
| ENSMUSG00000028906 | Epb41 | Enriched in WT dLGN | erythrocyte membrane protein band 4.1 [Source:MGI Symbol;Acc:MGI:95401] | 3569 | 1.665 | 0.008777 |
| ENSMUSG00000043456 | Zfp536 | Enriched in WT dLGN | zinc finger protein 536 [Source:MGI Symbol;Acc:MGI:1926102] | 762.2 | 1.675 | 0.003352 |
| ENSMUSG00000018862 | Otop3 | Enriched in WT dLGN | otopetrin 3 [Source:MGI Symbol;Acc:MGI:1916852] | 68.57 | 1.69 | 0.02194 |
| ENSMUSG00000032514 | Ttc21a | Enriched in WT dLGN | tetratricopeptide repeat domain 21A [Source:MGI Symbol;Acc:MGI:1921302] | 53.53 | 1.69 | 0.006342 |
| ENSMUSG00000030898 | Cckbr | Enriched in WT dLGN | cholecystokinin B receptor [Source:MGI Symbol;Acc:MGI:99479] | 109.5 | 1.691 | 0.0001584 |
| ENSMUSG00000029343 | Crybb1 | Enriched in WT dLGN | crystallin, beta B1 [Source:MGI Symbol;Acc:MGI:104992] | 480.1 | 1.696 | 0.00003031 |
| ENSMUSG00000049928 | Glp2r | Enriched in WT dLGN | glucagon-like peptide 2 receptor [Source:MGI Symbol;Acc:MGI:2136733] | 57.09 | 1.698 | 0.0008342 |
| ENSMUSG00000028280 | Gabrr1 | Enriched in WT dLGN | gamma-aminobutyric acid (GABA) C receptor, subunit rho 1 [Source:MGI Symbol;Acc:MGI:95625] | 141.8 | 1.706 | 0.01635 |
| ENSMUSG00000014198 | Zfp385c | Enriched in WT dLGN | zinc finger protein 385C [Source:MGI Symbol;Acc:MGI:3608347] | 103.3 | 1.709 | 0.000005141 |
| ENSMUSG00000028736 | Pax7 | Enriched in WT dLGN | paired box 7 [Source:MGI Symbol;Acc:MGI:97491] | 75.77 | 1.719 | 0.001997 |
| ENSMUSG00000035295 | Wdr38 | Enriched in WT dLGN | WD repeat domain 38 [Source:MGI Symbol;Acc:MGI:1923896] | 124.7 | 1.751 | 0.0001509 |
| ENSMUSG00000025359 | Pmel | Enriched in WT dLGN | premelanosome protein [Source:MGI Symbol;Acc:MGI:98301] | 69.08 | 1.753 | 0.03408 |
| ENSMUSG00000036598 | Ccdc113 | Enriched in WT dLGN | coiled-coil domain containing 113 [Source:MGI Symbol;Acc:MGI:3606076] | 76.78 | 1.774 | 0.00008276 |
| ENSMUSG00000038963 | Slco4a1 | Enriched in WT dLGN | solute carrier organic anion transporter family, member 4a1 [Source:MGI Symbol;Acc:MGI:1351866] | 224.4 | 1.782 | 0.00001122 |
| ENSMUSG00000021680 | Crhbp | Enriched in WT dLGN | corticotropin releasing hormone binding protein [Source:MGI Symbol;Acc:MGI:88497] | 76.8 | 1.788 | 0.02716 |
| ENSMUSG00000020212 | Mdm1 | Enriched in WT dLGN | transformed mouse 3T3 cell double minute 1 [Source:MGI Symbol;Acc:MGI:96951] | 841.5 | 1.789 | 8.919E-09 |
| ENSMUSG00000031636 | Pdlim3 | Enriched in WT dLGN | PDZ and LIM domain 3 [Source:MGI Symbol;Acc:MGI:1859274] | 77.07 | 1.793 | 0.01196 |
| ENSMUSG00000046110 | Serinc4 | Enriched in WT dLGN | serine incorporator 4 [Source:MGI Symbol;Acc:MGI:2441842] | 108.6 | 1.793 | 0.0002042 |
| ENSMUSG00000029054 | Gabrd | Enriched in WT dLGN | gamma-aminobutyric acid (GABA) A receptor, subunit delta [Source:MGI Symbol;Acc:MGI:95622] | 85.7 | 1.802 | 0.02908 |
| ENSMUSG00000035594 | Chrna5 | Enriched in WT dLGN | cholinergic receptor, nicotinic, alpha polypeptide 5 [Source:MGI Symbol;Acc:MGI:87889] | 110.2 | 1.802 | 0.000000502 |
| ENSMUSG00000004366 | Sst | Enriched in WT dLGN | somatostatin [Source:MGI Symbol;Acc:MGI:98326] | 3157 | 1.806 | 0.01794 |
| ENSMUSG00000091636 | Akain1 | Enriched in WT dLGN | A kinase (PRKA) anchor inhibitor 1 [Source:MGI Symbol;Acc:MGI:2444600] | 60.01 | 1.813 | 0.0008058 |
| ENSMUSG00000032420 | Nt5e | Enriched in WT dLGN | 5' nucleotidase, ecto [Source:MGI Symbol;Acc:MGI:99782] | 428.4 | 1.84 | 0.03482 |
| ENSMUSG00000022876 | Samsn1 | Enriched in WT dLGN | SAM domain, SH3 domain and nuclear localization signals, 1 [Source:MGI Symbol;Acc:MGI:1914992] | 155.2 | 1.846 | 0.003604 |
| ENSMUSG00000019982 | Myb | Enriched in WT dLGN | myeloblastosis oncogene [Source:MGI Symbol;Acc:MGI:97249] | 117.2 | 1.849 | 0.0005426 |
| ENSMUSG00000010021 | Kif19a | Enriched in WT dLGN | kinesin family member 19A [Source:MGI Symbol;Acc:MGI:2447024] | 738.6 | 1.881 | 0.02155 |
| ENSMUSG00000021903 | Galnt15 | Enriched in WT dLGN | polypeptide N-acetylgalactosaminyltransferase 15 [Source:MGI Symbol;Acc:MGI:1926004] | 84.6 | 1.884 | 0.000009963 |
| ENSMUSG00000045005 | Fzd5 | Enriched in WT dLGN | frizzled class receptor 5 [Source:MGI Symbol;Acc:MGI:108571] | 202.1 | 1.911 | 0.0005747 |
| ENSMUSG00000029819 | Npy | Enriched in WT dLGN | neuropeptide Y [Source:MGI Symbol;Acc:MGI:97374] | 2719 | 1.916 | 0.0001509 |
| ENSMUSG00000026830 | Ermn | Enriched in WT dLGN | ermin, ERM-like protein [Source:MGI Symbol;Acc:MGI:1925017] | 49.08 | 1.927 | 0.008094 |
| ENSMUSG00000032346 | Ooep | Enriched in WT dLGN | oocyte expressed protein [Source:MGI Symbol;Acc:MGI:1915218] | 114.1 | 1.93 | 0.000003004 |
| ENSMUSG00000100486 | Gm4131 | Enriched in WT dLGN | predicted gene 4131 [Source:MGI Symbol;Acc:MGI:3782307] | 700.1 | 1.96 | 1.423E-41 |
| ENSMUSG00000084989 | Crocc2 | Enriched in WT dLGN | ciliary rootlet coiled-coil, rootletin family member 2 [Source:MGI Symbol;Acc:MGI:3045962] | 44.72 | 1.961 | 0.04462 |
| ENSMUSG00000030230 | Plcz1 | Enriched in WT dLGN | phospholipase C, zeta 1 [Source:MGI Symbol;Acc:MGI:2150308] | 66.28 | 1.977 | 0.006493 |
| ENSMUSG00000041460 | Cacna2d4 | Enriched in WT dLGN | calcium channel, voltage-dependent, alpha 2/delta subunit 4 [Source:MGI Symbol;Acc:MGI:2442632] | 1003 | 1.988 | 0.008094 |
| ENSMUSG00000035403 | Crb2 | Enriched in WT dLGN | crumbs family member 2 [Source:MGI Symbol;Acc:MGI:2679260] | 222.1 | 2.005 | 0.00001229 |
| ENSMUSG00000038805 | Six3 | Enriched in WT dLGN | sine oculis-related homeobox 3 [Source:MGI Symbol;Acc:MGI:102764] | 2831 | 2.061 | 0.002809 |
| ENSMUSG00000048038 | Ccdc187 | Enriched in WT dLGN | coiled-coil domain containing 187 [Source:MGI Symbol;Acc:MGI:3045295] | 137.7 | 2.062 | 0.00001706 |
| ENSMUSG00000041046 | Ramp3 | Enriched in WT dLGN | receptor (calcitonin) activity modifying protein 3 [Source:MGI Symbol;Acc:MGI:1860292] | 2034 | 2.065 | 0.003094 |
| ENSMUSG00000005892 | Trh | Enriched in WT dLGN | thyrotropin releasing hormone [Source:MGI Symbol;Acc:MGI:98823] | 687.9 | 2.095 | 0.01512 |
| ENSMUSG00000021384 | Susd3 | Enriched in WT dLGN | sushi domain containing 3 [Source:MGI Symbol;Acc:MGI:1913579] | 113.7 | 2.101 | 0.00008228 |
| ENSMUSG00000022483 | Col2a1 | Enriched in WT dLGN | collagen, type II, alpha 1 [Source:MGI Symbol;Acc:MGI:88452] | 555.6 | 2.11 | 0.01811 |
| ENSMUSG00000046593 | Tmem215 | Enriched in WT dLGN | transmembrane protein 215 [Source:MGI Symbol;Acc:MGI:2444167] | 272.5 | 2.148 | 0.0405 |
| ENSMUSG00000048562 | Sp8 | Enriched in WT dLGN | trans-acting transcription factor 8 [Source:MGI Symbol;Acc:MGI:2443471] | 56.07 | 2.184 | 0.005776 |
| ENSMUSG00000086228 | Ubap1l | Enriched in WT dLGN | ubiquitin-associated protein 1-like [Source:MGI Symbol;Acc:MGI:2685360] | 636.1 | 2.184 | 0.01496 |
| ENSMUSG00000056888 | Glipr1 | Enriched in WT dLGN | GLI pathogenesis-related 1 (glioma) [Source:MGI Symbol;Acc:MGI:1920940] | 68.84 | 2.187 | 0.000463 |
| ENSMUSG00000000093 | Tbx2 | Enriched in WT dLGN | T-box 2 [Source:MGI Symbol;Acc:MGI:98494] | 461.2 | 2.197 | 3.037E-08 |
| ENSMUSG00000091402 | Rd3l | Enriched in WT dLGN | retinal degeneration 3-like [Source:MGI Symbol;Acc:MGI:2675860] | 248.7 | 2.198 | 6.258E-08 |
| ENSMUSG00000025161 | Slc16a3 | Enriched in WT dLGN | solute carrier family 16 (monocarboxylic acid transporters), member 3 [Source:MGI Symbol;Acc:MGI:1933438] | 198.8 | 2.247 | 0.000009073 |
| ENSMUSG00000048617 | Rtbdn | Enriched in WT dLGN | retbindin [Source:MGI Symbol;Acc:MGI:2443686] | 278.8 | 2.258 | 0.000003461 |
| ENSMUSG00000025064 | Col17a1 | Enriched in WT dLGN | collagen, type XVII, alpha 1 [Source:MGI Symbol;Acc:MGI:88450] | 86.57 | 2.271 | 0.0008563 |
| ENSMUSG00000054580 | Pla2r1 | Enriched in WT dLGN | phospholipase A2 receptor 1 [Source:MGI Symbol;Acc:MGI:102468] | 99.53 | 2.368 | 0.0001513 |
| ENSMUSG00000042707 | Dnali1 | Enriched in WT dLGN | dynein, axonemal, light intermediate polypeptide 1 [Source:MGI Symbol;Acc:MGI:1922813] | 57.63 | 2.371 | 0.04202 |
| ENSMUSG00000005716 | Pvalb | Enriched in WT dLGN | parvalbumin [Source:MGI Symbol;Acc:MGI:97821] | 1974 | 2.374 | 0.01439 |
| ENSMUSG00000040258 | Nxph4 | Enriched in WT dLGN | neurexophilin 4 [Source:MGI Symbol;Acc:MGI:1336197] | 225.4 | 2.407 | 0.01811 |
| ENSMUSG00000028977 | Casz1 | Enriched in WT dLGN | castor zinc finger 1 [Source:MGI Symbol;Acc:MGI:1196251] | 721 | 2.492 | 0.001006 |
| ENSMUSG00000070337 | Gpr179 | Enriched in WT dLGN | G protein-coupled receptor 179 [Source:MGI Symbol;Acc:MGI:2443409] | 245.8 | 2.527 | 0.000001692 |
| ENSMUSG00000108841 | Frmpd2 | Enriched in WT dLGN | FERM and PDZ domain containing 2 [Source:MGI Symbol;Acc:MGI:2685472] | 311.6 | 2.538 | 4.447E-09 |
| ENSMUSG00000029086 | Prom1 | Enriched in WT dLGN | prominin 1 [Source:MGI Symbol;Acc:MGI:1100886] | 1695 | 2.552 | 0.003284 |
| ENSMUSG00000021396 | Nxnl2 | Enriched in WT dLGN | nucleoredoxin-like 2 [Source:MGI Symbol;Acc:MGI:1922374] | 254.9 | 2.562 | 0.02285 |
| ENSMUSG00000021337 | Scgn | Enriched in WT dLGN | secretagogin, EF-hand calcium binding protein [Source:MGI Symbol;Acc:MGI:2384873] | 113.1 | 2.567 | 0.0007077 |
| ENSMUSG00000062859 | Tcp11 | Enriched in WT dLGN | t-complex protein 11 [Source:MGI Symbol;Acc:MGI:98544] | 57.91 | 2.582 | 0.0005036 |
| ENSMUSG00000075410 | Prcd | Enriched in WT dLGN | photoreceptor disc component [Source:MGI Symbol;Acc:MGI:3649529] | 261 | 2.582 | 2.728E-07 |
| ENSMUSG00000044254 | Pcsk9 | Enriched in WT dLGN | proprotein convertase subtilisin/kexin type 9 [Source:MGI Symbol;Acc:MGI:2140260] | 64.47 | 2.602 | 9.209E-10 |
| ENSMUSG00000002930 | Ppp1r17 | Enriched in WT dLGN | protein phosphatase 1, regulatory subunit 17 [Source:MGI Symbol;Acc:MGI:1333876] | 80.95 | 2.71 | 0.000001253 |
| ENSMUSG00000033080 | Vsx1 | Enriched in WT dLGN | visual system homeobox 1 [Source:MGI Symbol;Acc:MGI:1890816] | 179.5 | 2.769 | 0.009817 |
| ENSMUSG00000037727 | Avp | Enriched in WT dLGN | arginine vasopressin [Source:MGI Symbol;Acc:MGI:88121] | 43.05 | 2.771 | 0.0002811 |
| ENSMUSG00000021803 | Cdhr1 | Enriched in WT dLGN | cadherin-related family member 1 [Source:MGI Symbol;Acc:MGI:2157782] | 2402 | 2.839 | 0.00002707 |
| ENSMUSG00000039714 | Cplx3 | Enriched in WT dLGN | complexin 3 [Source:MGI Symbol;Acc:MGI:2384571] | 648.2 | 2.878 | 0.001469 |
| ENSMUSG00000075256 | Cerkl | Enriched in WT dLGN | ceramide kinase-like [Source:MGI Symbol;Acc:MGI:3037816] | 146.9 | 2.893 | 0.000008331 |
| ENSMUSG00000021363 | Mak | Enriched in WT dLGN | male germ cell-associated kinase [Source:MGI Symbol;Acc:MGI:96913] | 357.3 | 2.903 | 0.0001007 |
| ENSMUSG00000024987 | Cyp26a1 | Enriched in WT dLGN | cytochrome P450, family 26, subfamily a, polypeptide 1 [Source:MGI Symbol;Acc:MGI:1096359] | 41.44 | 2.941 | 0.000408 |
| ENSMUSG00000041193 | Pla2g5 | Enriched in WT dLGN | phospholipase A2, group V [Source:MGI Symbol;Acc:MGI:101899] | 109.2 | 3.027 | 0.0002104 |
| ENSMUSG00000038151 | Prdm1 | Enriched in WT dLGN | PR domain containing 1, with ZNF domain [Source:MGI Symbol;Acc:MGI:99655] | 331.6 | 3.07 | 1.218E-11 |
| ENSMUSG00000021647 | Cartpt | Enriched in WT dLGN | CART prepropeptide [Source:MGI Symbol;Acc:MGI:1351330] | 84.34 | 3.111 | 0.00001459 |
| ENSMUSG00000066975 | Cryba4 | Enriched in WT dLGN | crystallin, beta A4 [Source:MGI Symbol;Acc:MGI:102716] | 252.4 | 3.216 | 2.44E-12 |
| ENSMUSG00000048349 | Pou4f1 | Enriched in WT dLGN | POU domain, class 4, transcription factor 1 [Source:MGI Symbol;Acc:MGI:102525] | 87.43 | 3.217 | 0.0001824 |
| ENSMUSG00000063681 | Crb1 | Enriched in WT dLGN | crumbs family member 1, photoreceptor morphogenesis associated [Source:MGI Symbol;Acc:MGI:2136343] | 512.5 | 3.229 | 0.00002163 |
| ENSMUSG00000038115 | Ano2 | Enriched in WT dLGN | anoctamin 2 [Source:MGI Symbol;Acc:MGI:2387214] | 321.5 | 3.246 | 0.0003581 |
| ENSMUSG00000035200 | Chrnb4 | Enriched in WT dLGN | cholinergic receptor, nicotinic, beta polypeptide 4 [Source:MGI Symbol;Acc:MGI:87892] | 108.8 | 3.249 | 5.321E-07 |
| ENSMUSG00000031738 | Irx6 | Enriched in WT dLGN | Iroquois homeobox 6 [Source:MGI Symbol;Acc:MGI:1927642] | 58.39 | 3.331 | 0.0009597 |
| ENSMUSG00000040714 | Klc3 | Enriched in WT dLGN | kinesin light chain 3 [Source:MGI Symbol;Acc:MGI:1277971] | 329.4 | 3.331 | 0.000203 |
| ENSMUSG00000020080 | Hkdc1 | Enriched in WT dLGN | hexokinase domain containing 1 [Source:MGI Symbol;Acc:MGI:2384910] | 109.5 | 3.348 | 0.02626 |
| ENSMUSG00000068154 | Insm1 | Enriched in WT dLGN | insulinoma-associated 1 [Source:MGI Symbol;Acc:MGI:1859980] | 792.6 | 3.382 | 0.0002055 |
| ENSMUSG00000021123 | Rdh12 | Enriched in WT dLGN | retinol dehydrogenase 12 [Source:MGI Symbol;Acc:MGI:1925224] | 289.9 | 3.43 | 0.00005949 |
| ENSMUSG00000060461 | Dppa5a | Enriched in WT dLGN | developmental pluripotency associated 5A [Source:MGI Symbol;Acc:MGI:101800] | 84.7 | 3.435 | 3.829E-10 |
| ENSMUSG00000021359 | Tfap2a | Enriched in WT dLGN | transcription factor AP-2, alpha [Source:MGI Symbol;Acc:MGI:104671] | 156.8 | 3.458 | 0.003402 |
| ENSMUSG00000028943 | Espn | Enriched in WT dLGN | espin [Source:MGI Symbol;Acc:MGI:1861630] | 99.89 | 3.467 | 4.846E-07 |
| ENSMUSG00000020782 | Llgl2 | Enriched in WT dLGN | LLGL2 scribble cell polarity complex component [Source:MGI Symbol;Acc:MGI:1918843] | 247.7 | 3.485 | 0.0002976 |
| ENSMUSG00000020890 | Gucy2e | Enriched in WT dLGN | guanylate cyclase 2e [Source:MGI Symbol;Acc:MGI:105123] | 407.1 | 3.489 | 6.531E-07 |
| ENSMUSG00000028125 | Abca4 | Enriched in WT dLGN | ATP-binding cassette, sub-family A (ABC1), member 4 [Source:MGI Symbol;Acc:MGI:109424] | 528 | 3.5 | 0.00003405 |
| ENSMUSG00000038811 | Gngt2 | Enriched in WT dLGN | guanine nucleotide binding protein (G protein), gamma transducing activity polypeptide 2 [Source:MGI Symbol;Acc:MGI:893584] | 355.5 | 3.519 | 0.004869 |
| ENSMUSG00000027270 | Lamp5 | Enriched in WT dLGN | lysosomal-associated membrane protein family, member 5 [Source:MGI Symbol;Acc:MGI:1923411] | 108.3 | 3.567 | 0.000001104 |
| ENSMUSG00000042258 | Isl1 | Enriched in WT dLGN | ISL1 transcription factor, LIM/homeodomain [Source:MGI Symbol;Acc:MGI:101791] | 640.2 | 3.592 | 0.000001749 |
| ENSMUSG00000044429 | Cryga | Enriched in WT dLGN | crystallin, gamma A [Source:MGI Symbol;Acc:MGI:88521] | 66.43 | 3.655 | 4.235E-07 |
| ENSMUSG00000043850 | Clrn1 | Enriched in WT dLGN | clarin 1 [Source:MGI Symbol;Acc:MGI:2388124] | 58.84 | 3.74 | 1.078E-08 |
| ENSMUSG00000010476 | Ebf3 | Enriched in WT dLGN | early B cell factor 3 [Source:MGI Symbol;Acc:MGI:894289] | 164.2 | 3.795 | 2.148E-09 |
| ENSMUSG00000075330 | A930003A15Rik | Enriched in WT dLGN | RIKEN cDNA A930003A15 gene [Source:MGI Symbol;Acc:MGI:1915412] | 82.05 | 3.988 | 0.00005135 |
| ENSMUSG00000074991 | Gabrr3 | Enriched in WT dLGN | gamma-aminobutyric acid (GABA) receptor, rho 3 [Source:MGI Symbol;Acc:MGI:3588203] | 77.34 | 4.133 | 1.031E-09 |
| ENSMUSG00000032343 | Impg1 | Enriched in WT dLGN | interphotoreceptor matrix proteoglycan 1 [Source:MGI Symbol;Acc:MGI:1926876] | 226.2 | 4.182 | 0.000006515 |
| ENSMUSG00000057132 | Rpgrip1 | Enriched in WT dLGN | retinitis pigmentosa GTPase regulator interacting protein 1 [Source:MGI Symbol;Acc:MGI:1932134] | 1278 | 4.192 | 3.46E-10 |
| ENSMUSG00000079550 | Mpp4 | Enriched in WT dLGN | membrane protein, palmitoylated 4 (MAGUK p55 subfamily member 4) [Source:MGI Symbol;Acc:MGI:2386681] | 424.7 | 4.241 | 1.925E-08 |
| ENSMUSG00000056947 | Mab21l1 | Enriched in WT dLGN | mab-21-like 1 [Source:MGI Symbol;Acc:MGI:1333773] | 395.9 | 4.284 | 1.078E-08 |
| ENSMUSG00000064330 | Pde6h | Enriched in WT dLGN | phosphodiesterase 6H, cGMP-specific, cone, gamma [Source:MGI Symbol;Acc:MGI:1925850] | 117.3 | 4.355 | 3.209E-09 |
| ENSMUSG00000071648 | Rom1 | Enriched in WT dLGN | rod outer segment membrane protein 1 [Source:MGI Symbol;Acc:MGI:97998] | 1866 | 4.444 | 0.000007859 |
| ENSMUSG00000056494 | Cngb3 | Enriched in WT dLGN | cyclic nucleotide gated channel beta 3 [Source:MGI Symbol;Acc:MGI:1353562] | 54.78 | 4.512 | 0.000002591 |
| ENSMUSG00000030905 | Crym | Enriched in WT dLGN | crystallin, mu [Source:MGI Symbol;Acc:MGI:102675] | 928.5 | 4.614 | 0.00006613 |
| ENSMUSG00000029352 | Crybb3 | Enriched in WT dLGN | crystallin, beta B3 [Source:MGI Symbol;Acc:MGI:102717] | 457.1 | 4.697 | 1.654E-09 |
| ENSMUSG00000030206 | Gsg1 | Enriched in WT dLGN | germ cell associated 1 [Source:MGI Symbol;Acc:MGI:1194499] | 44.86 | 4.733 | 0.0001602 |
| ENSMUSG00000057777 | Mab21l2 | Enriched in WT dLGN | mab-21-like 2 [Source:MGI Symbol;Acc:MGI:1346022] | 136.6 | 4.965 | 0.0002283 |
| ENSMUSG00000022237 | Ankrd33b | Enriched in WT dLGN | ankyrin repeat domain 33B [Source:MGI Symbol;Acc:MGI:1917904] | 765.5 | 4.969 | 1.079E-08 |
| ENSMUSG00000037446 | Tulp1 | Enriched in WT dLGN | tubby like protein 1 [Source:MGI Symbol;Acc:MGI:109571] | 1768 | 4.979 | 9.107E-07 |
| ENSMUSG00000031142 | Cacna1f | Enriched in WT dLGN | calcium channel, voltage-dependent, alpha 1F subunit [Source:MGI Symbol;Acc:MGI:1859639] | 114.6 | 5.192 | 0.0001761 |
| ENSMUSG00000045776 | Lrtm1 | Enriched in WT dLGN | leucine-rich repeats and transmembrane domains 1 [Source:MGI Symbol;Acc:MGI:2442106] | 29.22 | 5.252 | 0.00003346 |
| ENSMUSG00000024992 | Pde6c | Enriched in WT dLGN | phosphodiesterase 6C, cGMP specific, cone, alpha prime [Source:MGI Symbol;Acc:MGI:105956] | 112.3 | 5.341 | 2.445E-11 |
| ENSMUSG00000110344 | Gm45716 | Enriched in WT dLGN | predicted gene 45716 [Source:MGI Symbol;Acc:MGI:5804831] | 63.87 | 5.355 | 0.001899 |
| ENSMUSG00000067220 | Cnga1 | Enriched in WT dLGN | cyclic nucleotide gated channel alpha 1 [Source:MGI Symbol;Acc:MGI:88436] | 505.5 | 5.5 | 1.314E-08 |
| ENSMUSG00000070683 | Lactbl1 | Enriched in WT dLGN | lactamase, beta-like 1 [Source:MGI Symbol;Acc:MGI:2448566] | 90.85 | 5.512 | 1.558E-11 |
| ENSMUSG00000086322 | E130218I03Rik | Enriched in WT dLGN | RIKEN cDNA E130218I03 gene [Source:MGI Symbol;Acc:MGI:3528958] | 500.5 | 5.549 | 3.802E-07 |
| ENSMUSG00000004630 | Pcp2 | Enriched in WT dLGN | Purkinje cell protein 2 (L7) [Source:MGI Symbol;Acc:MGI:97508] | 81.77 | 5.608 | 0.000523 |
| ENSMUSG00000005649 | Cabp5 | Enriched in WT dLGN | calcium binding protein 5 [Source:MGI Symbol;Acc:MGI:1352746] | 79.94 | 5.616 | 0.0001281 |

| | | | | | | |
|---|---|---|---|---|---|---|
| ENSMUSG00000040632 | Nrl | Enriched in WT dLGN | neural retina leucine zipper gene [Source:MGI Symbol;Acc:MGI:102567] | 2290 | 5.635 | 1.543E-08 |
| ENSMUSG00000034837 | Gnat1 | Enriched in WT dLGN | guanine nucleotide binding protein, alpha transducing 1 [Source:MGI Symbol;Acc:MGI:95778] | 274.3 | 5.639 | 7.537E-07 |
| ENSMUSG00000036480 | Prss56 | Enriched in WT dLGN | protease, serine 56 [Source:MGI Symbol;Acc:MGI:1916703] | 249.2 | 5.689 | 0.00001256 |
| ENSMUSG00000029410 | Ppef2 | Enriched in WT dLGN | protein phosphatase, EF hand calcium-binding domain 2 [Source:MGI Symbol;Acc:MGI:1342304] | 362.2 | 5.747 | 1.565E-31 |
| ENSMUSG00000026609 | Ush2a | Enriched in WT dLGN | usherin [Source:MGI Symbol;Acc:MGI:1341292] | 245.4 | 5.788 | 3.014E-33 |
| ENSMUSG00000034452 | Slc24a1 | Enriched in WT dLGN | solute carrier family 24 (sodium/potassium/calcium exchanger), member 1 [Source:MGI Symbol;Acc:MGI:2384871] | 138.5 | 5.821 | 2.555E-14 |
| ENSMUSG00000026989 | Dapl1 | Enriched in WT dLGN | death associated protein-like 1 [Source:MGI Symbol;Acc:MGI:1923997] | 38.68 | 5.988 | 5.202E-08 |
| ENSMUSG00000031789 | Cngb1 | Enriched in WT dLGN | cyclic nucleotide gated channel beta 1 [Source:MGI Symbol;Acc:MGI:2664102] | 970.5 | 6.089 | 3.209E-09 |
| ENSMUSG00000085139 | A730046J19Rik | Enriched in WT dLGN | RIKEN cDNA A730046J19 gene [Source:MGI Symbol;Acc:MGI:2442684] | 87.93 | 6.122 | 0.00007774 |
| ENSMUSG00000033501 | Crygs | Enriched in WT dLGN | crystallin, gamma S [Source:MGI Symbol;Acc:MGI:1298216] | 207.8 | 6.273 | 1.659E-10 |
| ENSMUSG00000064356 | mt-Atp8 | Enriched in WT dLGN | mitochondrially encoded ATP synthase 8 [Source:MGI Symbol;Acc:MGI:99926] | 43.05 | 6.435 | 0.006649 |
| ENSMUSG00000035270 | Impg2 | Enriched in WT dLGN | interphotoreceptor matrix proteoglycan 2 [Source:MGI Symbol;Acc:MGI:3044955] | 946.8 | 6.448 | 3.303E-14 |
| ENSMUSG00000092349 | Smim40 | Enriched in WT dLGN | small integral membrane protein 40 [Source:MGI Symbol;Acc:MGI:3054967] | 183.3 | 6.496 | 0.000001626 |
| ENSMUSG00000026468 | Lhx4 | Enriched in WT dLGN | LIM homeobox protein 4 [Source:MGI Symbol;Acc:MGI:101776] | 200.2 | 6.535 | 3.209E-09 |
| ENSMUSG00000024575 | Pde6a | Enriched in WT dLGN | phosphodiesterase 6A, cGMP-specific, rod, alpha [Source:MGI Symbol;Acc:MGI:97524] | 282.2 | 6.539 | 0.00005565 |
| ENSMUSG00000074365 | Crxos | Enriched in WT dLGN | cone-rod homeobox, opposite strand [Source:MGI Symbol;Acc:MGI:2451355] | 74.43 | 6.562 | 0.001602 |
| ENSMUSG00000058831 | Opn1sw | Enriched in WT dLGN | opsin 1 (cone pigments), short-wave-sensitive (color blindness, tritan) [Source:MGI Symbol;Acc:MGI:99438] | 230.1 | 6.633 | 0.000004264 |
| ENSMUSG00000097050 | Gm9918 | Enriched in WT dLGN | predicted gene 9918 [Source:MGI Symbol;Acc:MGI:3646315] | 107.5 | 6.83 | 0.00006769 |
| ENSMUSG00000042240 | Crybb2 | Enriched in WT dLGN | crystallin, beta B2 [Source:MGI Symbol;Acc:MGI:88519] | 374.3 | 6.851 | 7.432E-09 |
| ENSMUSG00000043418 | Lrit2 | Enriched in WT dLGN | leucine-rich repeat, immunoglobulin-like and transmembrane domains 2 [Source:MGI Symbol;Acc:MGI:2444885] | 170.6 | 6.859 | 0.000000502 |
| ENSMUSG00000045493 | Bhlhe23 | Enriched in WT dLGN | basic helix-loop-helix family, member e23 [Source:MGI Symbol;Acc:MGI:2153710] | 86.21 | 6.873 | 6.595E-10 |
| ENSMUSG00000020907 | Rcvrn | Enriched in WT dLGN | recoverin [Source:MGI Symbol;Acc:MGI:97883] | 558.4 | 7.007 | 9.898E-09 |
| ENSMUSG00000037161 | Mgarp | Enriched in WT dLGN | mitochondria localized glutamic acid rich protein [Source:MGI Symbol;Acc:MGI:1914999] | 632.7 | 7.123 | 5.873E-07 |
| ENSMUSG00000056043 | Rgs9bp | Enriched in WT dLGN | regulator of G-protein signalling 9 binding protein [Source:MGI Symbol;Acc:MGI:2384418] | 281.1 | 7.129 | 6.251E-11 |
| ENSMUSG00000027530 | Fabp12 | Enriched in WT dLGN | fatty acid binding protein 12 [Source:MGI Symbol;Acc:MGI:1922747] | 97.34 | 7.263 | 0.000001326 |
| ENSMUSG00000034829 | Nxnl1 | Enriched in WT dLGN | nucleoredoxin-like 1 [Source:MGI Symbol;Acc:MGI:1924446] | 270.2 | 7.264 | 0.000002348 |
| ENSMUSG00000023439 | Gnb3 | Enriched in WT dLGN | guanine nucleotide binding protein (G protein), beta 3 [Source:MGI Symbol;Acc:MGI:95785] | 770.3 | 7.285 | 1.543E-08 |
| ENSMUSG00000031293 | Rs1 | Enriched in WT dLGN | retinoschisis (X-linked, juvenile) 1 (human) [Source:MGI Symbol;Acc:MGI:1336189] | 932.3 | 7.325 | 1.69E-17 |
| ENSMUSG00000025389 | Mip | Enriched in WT dLGN | major intrinsic protein of lens fiber [Source:MGI Symbol;Acc:MGI:96990] | 109.6 | 7.401 | 3.455E-14 |
| ENSMUSG00000103119 | Gm37583 | Enriched in WT dLGN | predicted gene, 37583 [Source:MGI Symbol;Acc:MGI:5610811] | 152.8 | 7.46 | 0.000001849 |
| ENSMUSG00000056055 | Sag | Enriched in WT dLGN | S-antigen, retina and pineal gland (arrestin) [Source:MGI Symbol;Acc:MGI:98227] | 3480 | 7.89 | 2.332E-13 |
| ENSMUSG00000009108 | Gnat2 | Enriched in WT dLGN | guanine nucleotide binding protein, alpha transducing 2 [Source:MGI Symbol;Acc:MGI:95779] | 97.51 | 7.967 | 1.307E-13 |
| ENSMUSG00000030523 | Trpm1 | Enriched in WT dLGN | transient receptor potential cation channel, subfamily M, member 1 [Source:MGI Symbol;Acc:MGI:1330305] | 117.8 | 7.968 | 3.766E-20 |
| ENSMUSG00000041044 | Lrit1 | Enriched in WT dLGN | leucine-rich repeat, immunoglobulin-like and transmembrane domains 1 [Source:MGI Symbol;Acc:MGI:2385320] | 489 | 8.247 | 1.111E-09 |
| ENSMUSG00000021099 | Six6 | Enriched in WT dLGN | sine oculis-related homeobox 6 [Source:MGI Symbol;Acc:MGI:1341840] | 180.2 | 8.312 | 0.00002041 |
| ENSMUSG00000025386 | Pde6g | Enriched in WT dLGN | phosphodiesterase 6G, cGMP-specific, rod, gamma [Source:MGI Symbol;Acc:MGI:97526] | 1042 | 8.359 | 1.08E-12 |
| ENSMUSG00000048015 | Neurod4 | Enriched in WT dLGN | neurogenic differentiation 4 [Source:MGI Symbol;Acc:MGI:108055] | 749.4 | 8.388 | 3.829E-10 |
| ENSMUSG00000000617 | Grm6 | Enriched in WT dLGN | glutamate receptor, metabotropic 6 [Source:MGI Symbol;Acc:MGI:1351343] | 260.4 | 8.68 | 9.264E-09 |
| ENSMUSG00000051860 | Samd7 | Enriched in WT dLGN | sterile alpha motif domain containing 7 [Source:MGI Symbol;Acc:MGI:1923203] | 460.4 | 8.737 | 9.014E-10 |
| ENSMUSG00000031450 | Grk1 | Enriched in WT dLGN | G protein-coupled receptor kinase 1 [Source:MGI Symbol;Acc:MGI:1345146] | 554.6 | 8.757 | 2.387E-09 |
| ENSMUSG00000025927 | Tfap2b | Enriched in WT dLGN | transcription factor AP-2 beta [Source:MGI Symbol;Acc:MGI:104672] | 524.6 | 8.76 | 7.447E-15 |
| ENSMUSG00000040478 | Prdm13 | Enriched in WT dLGN | PR domain containing 13 [Source:MGI Symbol;Acc:MGI:2448528] | 96.48 | 8.789 | 0.0001524 |
| ENSMUSG00000067438 | Hmx1 | Enriched in WT dLGN | H6 homeobox 1 [Source:MGI Symbol;Acc:MGI:107178] | 82.19 | 8.868 | 0.000006597 |
| ENSMUSG00000093865 | Lrit3 | Enriched in WT dLGN | leucine-rich repeat, immunoglobulin-like and transmembrane domains 3 [Source:MGI Symbol;Acc:MGI:2685267] | 39 | 8.883 | 1.182E-09 |
| ENSMUSG00000025945 | Crygf | Enriched in WT dLGN | crystallin, gamma F [Source:MGI Symbol;Acc:MGI:88526] | 72.24 | 8.892 | 4.787E-13 |
| ENSMUSG00000008932 | Slc1a7 | Enriched in WT dLGN | solute carrier family 1 (glutamate transporter), member 7 [Source:MGI Symbol;Acc:MGI:2444087] | 266.9 | 8.908 | 1.493E-10 |
| ENSMUSG00000040554 | Aipl1 | Enriched in WT dLGN | aryl hydrocarbon receptor-interacting protein-like 1 [Source:MGI Symbol;Acc:MGI:2148800] | 433.3 | 8.933 | 2.148E-09 |
| ENSMUSG00000006546 | Cryba2 | Enriched in WT dLGN | crystallin, beta A2 [Source:MGI Symbol;Acc:MGI:104336] | 205.5 | 9.061 | 2.516E-10 |
| ENSMUSG00000047034 | Ankrd33 | Enriched in WT dLGN | ankyrin repeat domain 33 [Source:MGI Symbol;Acc:MGI:2443398] | 182.9 | 9.065 | 4.447E-09 |
| ENSMUSG00000021239 | Vsx2 | Enriched in WT dLGN | visual system homeobox 2 [Source:MGI Symbol;Acc:MGI:88401] | 1317 | 9.116 | 5.463E-14 |
| ENSMUSG00000044375 | Pcare | Enriched in WT dLGN | photoreceptor cilium actin regulator [Source:MGI Symbol;Acc:MGI:2385061] | 276.9 | 9.275 | 0.00001105 |
| ENSMUSG00000049908 | Gja8 | Enriched in WT dLGN | gap junction protein, alpha 8 [Source:MGI Symbol;Acc:MGI:99953] | 50.39 | 9.317 | 1.08E-12 |
| ENSMUSG00000025952 | Crygc | Enriched in WT dLGN | crystallin, gamma C [Source:MGI Symbol;Acc:MGI:88523] | 146.1 | 9.357 | 7.383E-19 |
| ENSMUSG00000073658 | Crygb | Enriched in WT dLGN | crystallin, gamma B [Source:MGI Symbol;Acc:MGI:88522] | 257.4 | 9.511 | 3.788E-27 |
| ENSMUSG00000024519 | Cplx4 | Enriched in WT dLGN | complexin 4 [Source:MGI Symbol;Acc:MGI:2685803] | 177.1 | 9.527 | 0.000000136 |
| ENSMUSG00000025900 | Rp1 | Enriched in WT dLGN | retinitis pigmentosa 1 (human) [Source:MGI Symbol;Acc:MGI:1341105] | 2273 | 9.6 | 7.52E-13 |
| ENSMUSG00000053773 | Rdh8 | Enriched in WT dLGN | retinol dehydrogenase 8 [Source:MGI Symbol;Acc:MGI:2685028] | 69.84 | 9.669 | 0.000001741 |
| ENSMUSG00000030324 | Rho | Enriched in WT dLGN | rhodopsin [Source:MGI Symbol;Acc:MGI:97914] | 5049 | 9.877 | 4.737E-16 |
| ENSMUSG00000023978 | Prph2 | Enriched in WT dLGN | peripherin 2 [Source:MGI Symbol;Acc:MGI:102791] | 2662 | 9.921 | 4.591E-13 |
| ENSMUSG00000029663 | Gngt1 | Enriched in WT dLGN | guanine nucleotide binding protein (G protein), gamma transducing activity polypeptide 1 [Source:MGI Symbol;Acc:MGI:109165] | 903.3 | 10.22 | 5.341E-08 |
| ENSMUSG00000067299 | Crygd | Enriched in WT dLGN | crystallin, gamma D [Source:MGI Symbol;Acc:MGI:88524] | 244 | 10.3 | 7.2E-13 |
| ENSMUSG00000024518 | Rax | Enriched in WT dLGN | retina and anterior neural fold homeobox [Source:MGI Symbol;Acc:MGI:109632] | 348.4 | 10.31 | 0.000001741 |
| ENSMUSG00000041534 | Rbp3 | Enriched in WT dLGN | retinol binding protein 3, interstitial [Source:MGI Symbol;Acc:MGI:97878] | 4106 | 10.53 | 2.004E-15 |
| ENSMUSG00000006007 | Pdc | Enriched in WT dLGN | phosducin [Source:MGI Symbol;Acc:MGI:98090] | 3344 | 10.54 | 8.405E-10 |
| ENSMUSG00000070870 | Cryge | Enriched in WT dLGN | crystallin, gamma E [Source:MGI Symbol;Acc:MGI:88525] | 184.6 | 10.78 | 3.688E-20 |
| ENSMUSG00000047298 | Kcnv2 | Enriched in WT dLGN | potassium channel, subfamily V, member 2 [Source:MGI Symbol;Acc:MGI:2670981] | 452.7 | 11.24 | 3.439E-19 |
| ENSMUSG00000046049 | Rp1l1 | Enriched in WT dLGN | retinitis pigmentosa 1 homolog like 1 [Source:MGI Symbol;Acc:MGI:2384303] | 752 | 11.41 | 1.509E-17 |
| ENSMUSG00000024041 | Cryaa | Enriched in WT dLGN | crystallin, alpha A [Source:MGI Symbol;Acc:MGI:88515] | 1565 | 12.1 | 6.141E-13 |
| ENSMUSG00000032292 | Nr2e3 | Enriched in WT dLGN | nuclear receptor subfamily 2, group E, member 3 [Source:MGI Symbol;Acc:MGI:1346317] | 2583 | 12.1 | 6.251E-11 |
| ENSMUSG00000041578 | Crx | Enriched in WT dLGN | cone-rod homeobox [Source:MGI Symbol;Acc:MGI:1194883] | 2316 | 12.89 | 7.023E-09 |
| ENSMUSG00000000724 | Cryba1 | Enriched in WT dLGN | crystallin, beta A1 [Source:MGI Symbol;Acc:MGI:88518] | 732 | 12.94 | 4.046E-21 |
| ENSMUSG00000029491 | Pde6b | Enriched in WT dLGN | phosphodiesterase 6B, cGMP, rod receptor, beta polypeptide [Source:MGI Symbol;Acc:MGI:97525] | 1563 | 13.31 | 2.296E-17 |

**Contrast numerator: KO_dLGN. Contrast denominator: Het_dLGN.**

| ensembl_gene_id | external_gene_name | Significance | description | baseMean | logFC | adj.P.Val |
|---|---|---|---|---|---|---|
| ENSMUSG00000029491 | Pde6b | Enriched in Het dLGN | phosphodiesterase 6B, cGMP, rod receptor, beta polypeptide [Source:MGI Symbol;Acc:MGI:97525] | 287.1 | -8.114 | 0.0002779 |
| ENSMUSG00000046049 | Rp1l1 | Enriched in Het dLGN | retinitis pigmentosa 1 homolog like 1 [Source:MGI Symbol;Acc:MGI:2384303] | 163.9 | -6.67 | 0.0003124 |
| ENSMUSG00000047298 | Kcnv2 | Enriched in Het dLGN | potassium channel, subfamily V, member 2 [Source:MGI Symbol;Acc:MGI:2670981] | 106 | -6.313 | 0.005991 |
| ENSMUSG00000031022 | BC051019 | Enriched in Het dLGN | cDNA sequence BC051019 [Source:MGI Symbol;Acc:MGI:1928824] | 16.5 | -4.637 | 0.02627 |
| ENSMUSG00000037727 | Avp | Enriched in Het dLGN | arginine vasopressin [Source:MGI Symbol;Acc:MGI:88121] | 493.3 | -4.632 | 0.02226 |
| ENSMUSG00000023978 | Prph2 | Enriched in Het dLGN | peripherin 2 [Source:MGI Symbol;Acc:MGI:102791] | 560.3 | -4.49 | 0.01876 |
| ENSMUSG00000024519 | Cplx4 | Enriched in Het dLGN | complexin 4 [Source:MGI Symbol;Acc:MGI:2685803] | 46.36 | -4.485 | 0.04252 |
| ENSMUSG00000025927 | Tfap2b | Enriched in Het dLGN | transcription factor AP-2 beta [Source:MGI Symbol;Acc:MGI:104672] | 110.4 | -3.884 | 0.02104 |
| ENSMUSG00000008932 | Slc1a7 | Enriched in Het dLGN | solute carrier family 1 (glutamate transporter), member 7 [Source:MGI Symbol;Acc:MGI:2444087] | 54.39 | -3.853 | 0.02262 |
| ENSMUSG00000000617 | Grm6 | Enriched in Het dLGN | glutamate receptor, metabotropic 6 [Source:MGI Symbol;Acc:MGI:1351343] | 55.11 | -3.824 | 0.03477 |
| ENSMUSG00000025386 | Pde6g | Enriched in Het dLGN | phosphodiesterase 6G, cGMP-specific, rod, gamma [Source:MGI Symbol;Acc:MGI:97526] | 242.3 | -3.713 | 0.0336 |
| ENSMUSG00000030523 | Trpm1 | Enriched in Het dLGN | transient receptor potential cation channel, subfamily M, member 1 [Source:MGI Symbol;Acc:MGI:1330305] | 45.51 | -3.575 | 0.007169 |
| ENSMUSG00000060461 | Dppa5a | Enriched in Het dLGN | developmental pluripotency associated 5A [Source:MGI Symbol;Acc:MGI:101800] | 73.61 | -3.154 | 0.000003632 |
| ENSMUSG00000021647 | Cartpt | Enriched in Het dLGN | CART prepropeptide [Source:MGI Symbol;Acc:MGI:1351330] | 82.61 | -2.988 | 0.00001264 |
| ENSMUSG00000026301 | Iqca | Enriched in Het dLGN | IQ motif containing with AAA domain [Source:MGI Symbol;Acc:MGI:1922168] | 26.27 | -2.666 | 0.01862 |
| ENSMUSG00000024987 | Cyp26a1 | Enriched in Het dLGN | cytochrome P450, family 26, subfamily a, polypeptide 1 [Source:MGI Symbol;Acc:MGI:1096359] | 37.1 | -2.622 | 0.0002004 |
| ENSMUSG00000056043 | Rgs9bp | Enriched in Het dLGN | regulator of G-protein signalling 9 binding protein [Source:MGI Symbol;Acc:MGI:2384418] | 74.59 | -2.58 | 0.006247 |
| ENSMUSG00000004630 | Pcp2 | Enriched in Het dLGN | Purkinje cell protein 2 (L7) [Source:MGI Symbol;Acc:MGI:97508] | 29.57 | -2.573 | 0.04706 |
| ENSMUSG00000019890 | Nts | Enriched in Het dLGN | neurotensin [Source:MGI Symbol;Acc:MGI:1328351] | 138.3 | -2.536 | 0.01034 |
| ENSMUSG00000034452 | Slc24a1 | Enriched in Het dLGN | solute carrier family 24 (sodium/potassium/calcium exchanger), member 1 [Source:MGI Symbol;Acc:MGI:2384871] | 58.63 | -2.408 | 0.01896 |
| ENSMUSG00000026609 | Ush2a | Enriched in Het dLGN | usherin [Source:MGI Symbol;Acc:MGI:1341292] | 63.5 | -2.289 | 0.01364 |
| ENSMUSG00000002930 | Ppp1r17 | Enriched in Het dLGN | protein phosphatase 1, regulatory subunit 17 [Source:MGI Symbol;Acc:MGI:1333876] | 67.8 | -2.112 | 0.02775 |
| ENSMUSG00000042258 | Isl1 | Enriched in Het dLGN | ISL1 transcription factor, LIM/homeodomain [Source:MGI Symbol;Acc:MGI:101791] | 240.3 | -2.096 | 0.003153 |
| ENSMUSG00000045655 | Fam216b | Enriched in Het dLGN | family with sequence similarity 216, member B [Source:MGI Symbol;Acc:MGI:2145738] | 57.89 | -1.968 | 0.03506 |
| ENSMUSG00000048562 | Sp8 | Enriched in Het dLGN | trans-acting transcription factor 8 [Source:MGI Symbol;Acc:MGI:2443471] | 37.66 | -1.905 | 0.004219 |
| ENSMUSG00000029754 | Dlx6 | Enriched in Het dLGN | distal-less homeobox 6 [Source:MGI Symbol;Acc:MGI:101927] | 218.7 | -1.884 | 0.01892 |
| ENSMUSG00000032346 | Ooep | Enriched in Het dLGN | oocyte expressed protein [Source:MGI Symbol;Acc:MGI:1915218] | 122.5 | -1.833 | 0.001642 |
| ENSMUSG00000042379 | Esm1 | Enriched in Het dLGN | endothelial cell-specific molecule 1 [Source:MGI Symbol;Acc:MGI:1918940] | 27.46 | -1.815 | 0.03317 |
| ENSMUSG00000020415 | Pttg1 | Enriched in Het dLGN | pituitary tumor-transforming gene 1 [Source:MGI Symbol;Acc:MGI:1353578] | 492.2 | -1.783 | 1.17E-10 |
| ENSMUSG00000032291 | Crabp1 | Enriched in Het dLGN | cellular retinoic acid binding protein I [Source:MGI Symbol;Acc:MGI:88490] | 103.7 | -1.755 | 0.01016 |
| ENSMUSG00000024553 | Galr1 | Enriched in Het dLGN | galanin receptor 1 [Source:MGI Symbol;Acc:MGI:1096364] | 29.72 | -1.669 | 0.01519 |
| ENSMUSG00000010476 | Ebf3 | Enriched in Het dLGN | early B cell factor 3 [Source:MGI Symbol;Acc:MGI:894289] | 46.9 | -1.661 | 0.04444 |
| ENSMUSG00000084989 | Crocc2 | Enriched in Het dLGN | ciliary rootlet coiled-coil, rootletin family member 2 [Source:MGI Symbol;Acc:MGI:3045962] | 42.84 | -1.606 | 0.03411 |
| ENSMUSG00000004366 | Sst | Enriched in Het dLGN | somatostatin [Source:MGI Symbol;Acc:MGI:98326] | 2053 | -1.563 | 0.0165 |
| ENSMUSG00000023391 | Dlx2 | Enriched in Het dLGN | distal-less homeobox 2 [Source:MGI Symbol;Acc:MGI:94902] | 78.78 | -1.521 | 0.02103 |
| ENSMUSG00000034459 | Ifit1 | Enriched in Het dLGN | interferon-induced protein with tetratricopeptide repeats 1 [Source:MGI Symbol;Acc:MGI:99450] | 32.72 | -1.516 | 0.01227 |
| ENSMUSG00000041046 | Ramp3 | Enriched in Het dLGN | receptor (calcitonin) activity modifying protein 3 [Source:MGI Symbol;Acc:MGI:1860292] | 1157 | -1.438 | 0.04666 |
| ENSMUSG00000020890 | Gucy2e | Enriched in Het dLGN | guanylate cyclase 2e [Source:MGI Symbol;Acc:MGI:105123] | 154 | -1.371 | 0.01309 |
| ENSMUSG00000046861 | Hectd3 | Enriched in Het dLGN | HECT domain E3 ubiquitin protein ligase 3 [Source:MGI Symbol;Acc:MGI:1923858] | 1039 | -1.282 | 0.003999 |
| ENSMUSG00000026247 | Ecel1 | Enriched in Het dLGN | endothelin converting enzyme-like 1 [Source:MGI Symbol;Acc:MGI:1343461] | 618.5 | -1.252 | 0.00035 |
| ENSMUSG00000032514 | Ttc21a | Enriched in Het dLGN | tetratricopeptide repeat domain 21A [Source:MGI Symbol;Acc:MGI:1921302] | 72.95 | -1.199 | 0.01892 |
| ENSMUSG00000062372 | Otof | Enriched in Het dLGN | otoferlin [Source:MGI Symbol;Acc:MGI:1891247] | 401.8 | -1.186 | 0.002189 |
| ENSMUSG00000026896 | Ifih1 | Enriched in Het dLGN | interferon induced with helicase C domain 1 [Source:MGI Symbol;Acc:MGI:1918836] | 62.95 | -1.168 | 0.0004839 |
| ENSMUSG00000069874 | Irgm2 | Enriched in Het dLGN | immunity-related GTPase family M member 2 [Source:MGI Symbol;Acc:MGI:1926262] | 72.85 | -1.161 | 0.01991 |
| ENSMUSG00000042514 | Klhl14 | Enriched in Het dLGN | kelch-like 14 [Source:MGI Symbol;Acc:MGI:1921249] | 179.9 | -1.156 | 0.02446 |
| ENSMUSG00000032221 | Mns1 | Enriched in Het dLGN | meiosis-specific nuclear structural protein 1 [Source:MGI Symbol;Acc:MGI:107933] | 208.3 | -1.105 | 0.000002882 |
| ENSMUSG00000064354 | mt-Co2 | Enriched in Het dLGN | mitochondrially encoded cytochrome c oxidase II [Source:MGI Symbol;Acc:MGI:102503] | 177.7 | -1.074 | 0.03418 |
| ENSMUSG00000029561 | Oasl2 | Enriched in Het dLGN | 2'-5' oligoadenylate synthetase-like 2 [Source:MGI Symbol;Acc:MGI:1344390] | 93.26 | -1.069 | 0.01124 |
| ENSMUSG00000028883 | Sema3a | Enriched in Het dLGN | sema domain, immunoglobulin domain (Ig), short basic domain, secreted, (semaphorin) 3A [Source:MGI Symbol;Acc:MGI:10755 | 140.8 | -1.061 | 0.01048 |
| ENSMUSG00000040152 | Thbs1 | Enriched in Het dLGN | thrombospondin 1 [Source:MGI Symbol;Acc:MGI:98737] | 216.9 | -1.036 | 0.01214 |
| ENSMUSG00000034684 | Sema3f | Enriched in Het dLGN | sema domain, immunoglobulin domain (Ig), short basic domain, secreted, (semaphorin) 3F [Source:MGI Symbol;Acc:MGI:10963 | 541.6 | -1.01 | 0.007169 |
| ENSMUSG00000040565 | Btaf1 | Enriched in KO dLGN | B-TFIID TATA-box binding protein associated factor 1 [Source:MGI Symbol;Acc:MGI:2147538] | 1188 | 1.077 | 0.01416 |
| ENSMUSG00000044017 | Adgrd1 | Enriched in KO dLGN | adhesion G protein-coupled receptor D1 [Source:MGI Symbol;Acc:MGI:3041203] | 36.41 | 1.294 | 0.04073 |
| ENSMUSG00000096351 | Samd11 | Enriched in KO dLGN | sterile alpha motif domain containing 11 [Source:MGI Symbol;Acc:MGI:2446220] | 502.3 | 1.306 | 0.03228 |
| ENSMUSG00000043091 | Tuba1c | Enriched in KO dLGN | tubulin, alpha 1C [Source:MGI Symbol;Acc:MGI:1095409] | 1206 | 1.893 | 0.02361 |
| ENSMUSG00000102439 | Flg | Enriched in KO dLGN | filaggrin [Source:MGI Symbol;Acc:MGI:95553] | 27.09 | 2.807 | 0.001642 |
| ENSMUSG00000061808 | Ttr | Enriched in KO dLGN | transthyretin [Source:MGI Symbol;Acc:MGI:98865] | 106700 | 3.704 | 0.02195 |
| ENSMUSG00000072476 | Gm9008 | Enriched in KO dLGN | predicted pseudogene 9008 [Source:MGI Symbol;Acc:MGI:3644000] | 110.4 | 7.086 | 0.005016 |
| ENSMUSG00000061780 | Cfd | Enriched in KO dLGN | complement factor D (adipsin) [Source:MGI Symbol;Acc:MGI:87931] | 38.12 | 10.09 | 8.368E-40 |

**Contrast numerator: WT_dLGN. Contrast denominator: Het_dLGN.**

| ensembl_gene_id | external_gene_name | Significance | description | baseMean | logFC | adj.P.Val |
|---|---|---|---|---|---|---|
| ENSMUSG00000026697 | Myoc | Enriched in Het dLGN | myocilin [Source:MGI Symbol;Acc:MGI:1202864] | 1257 | -2.175 | 0.01475 |
| ENSMUSG00000075217 | Fads2b | Enriched in Het dLGN | fatty acid desaturase 2B [Source:MGI Symbol;Acc:MGI:2687041] | 36.49 | -1.777 | 0.01581 |
| ENSMUSG00000030711 | Sult1a1 | Enriched in Het dLGN | sulfotransferase family 1A, phenol-preferring, member 1 [Source:MGI Symbol;Acc:MGI:102896] | 136.1 | -1.596 | 0.002537 |
| ENSMUSG00000031351 | Zfp185 | Enriched in Het dLGN | zinc finger protein 185 [Source:MGI Symbol;Acc:MGI:108095] | 72.07 | -1.565 | 0.002629 |
| ENSMUSG00000004885 | Crabp2 | Enriched in Het dLGN | cellular retinoic acid binding protein II [Source:MGI Symbol;Acc:MGI:88491] | 297.3 | -1.466 | 0.004985 |
| ENSMUSG00000020264 | Slc36a2 | Enriched in Het dLGN | solute carrier family 36 (proton/amino acid symporter), member 2 [Source:MGI Symbol;Acc:MGI:1891430] | 39.45 | -1.384 | 0.04202 |
| ENSMUSG00000046794 | Ppp1r3b | Enriched in Het dLGN | protein phosphatase 1, regulatory subunit 3B [Source:MGI Symbol;Acc:MGI:2177268] | 170.4 | -1.374 | 0.01705 |
| ENSMUSG00000048572 | Tmem252 | Enriched in Het dLGN | transmembrane protein 252 [Source:MGI Symbol;Acc:MGI:3583948] | 130.6 | -1.31 | 0.01789 |
| ENSMUSG00000027611 | Procr | Enriched in Het dLGN | protein C receptor, endothelial [Source:MGI Symbol;Acc:MGI:104596] | 62.55 | -1.283 | 0.006457 |
| ENSMUSG00000079396 | Gm3411 | Enriched in Het dLGN | predicted gene 3411 [Source:MGI Symbol;Acc:MGI:3781589] | 44.47 | -1.215 | 0.03195 |
| ENSMUSG00000072572 | Slc39a2 | Enriched in Het dLGN | solute carrier family 39 (zinc transporter), member 2 [Source:MGI Symbol;Acc:MGI:2684326] | 391.8 | -1.13 | 0.002259 |
| ENSMUSG00000045930 | Clec14a | Enriched in Het dLGN | C-type lectin domain family 14, member a [Source:MGI Symbol;Acc:MGI:1914114] | 339.3 | -1.057 | 0.002629 |
| ENSMUSG00000070867 | Trabd2b | Enriched in Het dLGN | TraB domain containing 2B [Source:MGI Symbol;Acc:MGI:3650152] | 98.97 | -1.057 | 0.0422 |
| ENSMUSG00000043631 | Ecm2 | Enriched in Het dLGN | extracellular matrix protein 2, female organ and adipocyte specific [Source:MGI Symbol;Acc:MGI:3039578] | 224.7 | -1.026 | 0.007824 |
| ENSMUSG00000073643 | Wdfy1 | Enriched in Het dLGN | WD repeat and FYVE domain containing 1 [Source:MGI Symbol;Acc:MGI:1916618] | 1394 | -1.022 | 0.001173 |
| ENSMUSG00000078963 | Hsbp1l1 | Enriched in WT dLGN | heat shock factor binding protein 1-like 1 [Source:MGI Symbol;Acc:MGI:1913505] | 136.9 | 1.046 | 0.01078 |
| ENSMUSG00000029343 | Crybb1 | Enriched in WT dLGN | crystallin, beta B1 [Source:MGI Symbol;Acc:MGI:104992] | 382.9 | 1.087 | 0.02971 |
| ENSMUSG00000035403 | Crb2 | Enriched in WT dLGN | crumbs family member 2 [Source:MGI Symbol;Acc:MGI:2679260] | 303 | 1.091 | 0.03734 |
| ENSMUSG00000030402 | Ppm1n | Enriched in WT dLGN | protein phosphatase, Mg2+/Mn2+ dependent, 1N (putative) [Source:MGI Symbol;Acc:MGI:2142330] | 112.8 | 1.118 | 0.04459 |
| ENSMUSG00000035594 | Chrna5 | Enriched in WT dLGN | cholinergic receptor, nicotinic, alpha polypeptide 5 [Source:MGI Symbol;Acc:MGI:87889] | 89.56 | 1.127 | 0.04599 |
| ENSMUSG00000033722 | BC034090 | Enriched in WT dLGN | cDNA sequence BC034090 [Source:MGI Symbol;Acc:MGI:2672904] | 679.9 | 1.16 | 0.04599 |
| ENSMUSG00000004655 | Aqp1 | Enriched in WT dLGN | aquaporin 1 [Source:MGI Symbol;Acc:MGI:103201] | 760 | 1.163 | 0.001173 |
| ENSMUSG00000050786 | Ccdc126 | Enriched in WT dLGN | coiled-coil domain containing 126 [Source:MGI Symbol;Acc:MGI:1889376] | 362.7 | 1.186 | 0.01299 |
| ENSMUSG00000035504 | Reep6 | Enriched in WT dLGN | receptor accessory protein 6 [Source:MGI Symbol;Acc:MGI:1917585] | 630.1 | 1.19 | 0.03803 |
| ENSMUSG00000014198 | Zfp385c | Enriched in WT dLGN | zinc finger protein 385C [Source:MGI Symbol;Acc:MGI:3608347] | 77.61 | 1.206 | 0.0344 |
| ENSMUSG00000010021 | Kif19a | Enriched in WT dLGN | kinesin family member 19A [Source:MGI Symbol;Acc:MGI:2447024] | 575.9 | 1.289 | 0.03885 |
| ENSMUSG00000028906 | Epb41 | Enriched in WT dLGN | erythrocyte membrane protein band 4.1 [Source:MGI Symbol;Acc:MGI:95401] | 2662 | 1.292 | 0.04599 |
| ENSMUSG00000091636 | Akain1 | Enriched in WT dLGN | A kinase (PRKA) anchor inhibitor 1 [Source:MGI Symbol;Acc:MGI:2444600] | 44.47 | 1.292 | 0.04824 |
| ENSMUSG00000021903 | Galnt15 | Enriched in WT dLGN | polypeptide N-acetylgalactosaminyltransferase 15 [Source:MGI Symbol;Acc:MGI:1926004] | 59.51 | 1.324 | 0.03691 |
| ENSMUSG00000020212 | Mdm1 | Enriched in WT dLGN | transformed mouse 3T3 cell double minute 1 [Source:MGI Symbol;Acc:MGI:96951] | 635.6 | 1.331 | 0.03191 |
| ENSMUSG00000048617 | Rtbdn | Enriched in WT dLGN | retbindin [Source:MGI Symbol;Acc:MGI:2443686] | 224.4 | 1.335 | 0.02332 |
| ENSMUSG00000021009 | Ptpn21 | Enriched in WT dLGN | protein tyrosine phosphatase, non-receptor type 21 [Source:MGI Symbol;Acc:MGI:1344406] | 363.1 | 1.353 | 0.001744 |
| ENSMUSG00000021384 | Susd3 | Enriched in WT dLGN | sushi domain containing 3 [Source:MGI Symbol;Acc:MGI:1913579] | 91.32 | 1.394 | 0.04508 |
| ENSMUSG00000046110 | Serinc4 | Enriched in WT dLGN | serine incorporator 4 [Source:MGI Symbol;Acc:MGI:2441842] | 77.44 | 1.432 | 0.01143 |
| ENSMUSG00000022860 | Chodl | Enriched in WT dLGN | chondrolectin [Source:MGI Symbol;Acc:MGI:2179069] | 54.84 | 1.447 | 0.006696 |
| ENSMUSG00000070337 | Gpr179 | Enriched in WT dLGN | G protein-coupled receptor 179 [Source:MGI Symbol;Acc:MGI:2443409] | 191.6 | 1.461 | 0.01737 |
| ENSMUSG00000041460 | Cacna2d4 | Enriched in WT dLGN | calcium channel, voltage-dependent, alpha 2/delta subunit 4 [Source:MGI Symbol;Acc:MGI:2442632] | 764.1 | 1.48 | 0.0212 |
| ENSMUSG00000030230 | Plcz1 | Enriched in WT dLGN | phospholipase C, zeta 1 [Source:MGI Symbol;Acc:MGI:2150308] | 49.01 | 1.496 | 0.04429 |
| ENSMUSG00000100486 | Gm4131 | Enriched in WT dLGN | predicted gene 4131 [Source:MGI Symbol;Acc:MGI:3782307] | 601.5 | 1.502 | 0.009821 |
| ENSMUSG00000038963 | Slco4a1 | Enriched in WT dLGN | solute carrier organic anion transporter family, member 4a1 [Source:MGI Symbol;Acc:MGI:1351866] | 158.8 | 1.54 | 0.001876 |
| ENSMUSG00000025161 | Slc16a3 | Enriched in WT dLGN | solute carrier family 16 (monocarboxylic acid transporters), member 3 [Source:MGI Symbol;Acc:MGI:1933438] | 145.2 | 1.57 | 0.01473 |
| ENSMUSG00000056888 | Glipr1 | Enriched in WT dLGN | GLI pathogenesis-related 1 (glioma) [Source:MGI Symbol;Acc:MGI:1920940] | 49.31 | 1.586 | 0.007303 |
| ENSMUSG00000042258 | Isl1 | Enriched in WT dLGN | ISL1 transcription factor, LIM/homeodomain [Source:MGI Symbol;Acc:MGI:101791] | 509.4 | 1.609 | 0.005472 |
| ENSMUSG00000054580 | Pla2r1 | Enriched in WT dLGN | phospholipase A2 receptor 1 [Source:MGI Symbol;Acc:MGI:102468] | 74.41 | 1.642 | 0.02904 |
| ENSMUSG00000048038 | Ccdc187 | Enriched in WT dLGN | coiled-coil domain containing 187 [Source:MGI Symbol;Acc:MGI:3045295] | 109.5 | 1.678 | 0.01473 |
| ENSMUSG00000000093 | Tbx2 | Enriched in WT dLGN | T-box 2 [Source:MGI Symbol;Acc:MGI:98494] | 343.9 | 1.683 | 0.02501 |
| ENSMUSG00000045534 | Kcna5 | Enriched in WT dLGN | potassium voltage-gated channel, shaker-related subfamily, member 5 [Source:MGI Symbol;Acc:MGI:96662] | 103.5 | 1.707 | 0.01525 |
| ENSMUSG00000018862 | Otop3 | Enriched in WT dLGN | otopetrin 3 [Source:MGI Symbol;Acc:MGI:1916852] | 45.5 | 1.73 | 0.02005 |
| ENSMUSG00000046593 | Tmem215 | Enriched in WT dLGN | transmembrane protein 215 [Source:MGI Symbol;Acc:MGI:2444167] | 205.2 | 1.774 | 0.0344 |
| ENSMUSG00000028977 | Casz1 | Enriched in WT dLGN | castor zinc finger 1 [Source:MGI Symbol;Acc:MGI:1196251] | 549.7 | 1.807 | 0.01834 |
| ENSMUSG00000026830 | Ermn | Enriched in WT dLGN | ermin, ERM-like protein [Source:MGI Symbol;Acc:MGI:1925017] | 43.78 | 1.809 | 0.03767 |
| ENSMUSG00000029086 | Prom1 | Enriched in WT dLGN | prominin 1 [Source:MGI Symbol;Acc:MGI:1100886] | 1240 | 1.816 | 0.03381 |
| ENSMUSG00000025064 | Col17a1 | Enriched in WT dLGN | collagen, type XVII, alpha 1 [Source:MGI Symbol;Acc:MGI:88450] | 64.36 | 1.867 | 0.01373 |
| ENSMUSG00000028736 | Pax7 | Enriched in WT dLGN | paired box 7 [Source:MGI Symbol;Acc:MGI:97491] | 45.56 | 1.875 | 0.03081 |
| ENSMUSG00000020890 | Gucy2e | Enriched in WT dLGN | guanylate cyclase 2e [Source:MGI Symbol;Acc:MGI:105123] | 320.2 | 1.966 | 0.04538 |
| ENSMUSG00000010476 | Ebf3 | Enriched in WT dLGN | early B cell factor 3 [Source:MGI Symbol;Acc:MGI:894289] | 117.7 | 1.968 | 0.009347 |
| ENSMUSG00000043850 | Clrn1 | Enriched in WT dLGN | clarin 1 [Source:MGI Symbol;Acc:MGI:2388124] | 47.71 | 1.971 | 0.01329 |
| ENSMUSG00000062859 | Tcp11 | Enriched in WT dLGN | t-complex protein 11 [Source:MGI Symbol;Acc:MGI:98544] | 42.53 | 1.973 | 0.01338 |
| ENSMUSG00000063681 | Crb1 | Enriched in WT dLGN | crumbs family member 1, photoreceptor morphogenesis associated [Source:MGI Symbol;Acc:MGI:2136343] | 392.5 | 2.026 | 0.02006 |
| ENSMUSG00000021337 | Scgn | Enriched in WT dLGN | secretagogin, EF-hand calcium binding protein [Source:MGI Symbol;Acc:MGI:2384873] | 85.16 | 2.068 | 0.01501 |
| ENSMUSG00000027270 | Lamp5 | Enriched in WT dLGN | lysosomal-associated membrane protein family, member 5 [Source:MGI Symbol;Acc:MGI:1923411] | 77.62 | 2.101 | 0.01571 |
| ENSMUSG00000075256 | Cerkl | Enriched in WT dLGN | ceramide kinase-like [Source:MGI Symbol;Acc:MGI:3037816] | 109.5 | 2.111 | 0.005508 |
| ENSMUSG00000039714 | Cplx3 | Enriched in WT dLGN | complexin 3 [Source:MGI Symbol;Acc:MGI:2384571] | 475.5 | 2.178 | 0.01737 |
| ENSMUSG00000040258 | Nxph4 | Enriched in WT dLGN | neurexophilin 4 [Source:MGI Symbol;Acc:MGI:1336197] | 152.4 | 2.189 | 0.01796 |
| ENSMUSG00000066975 | Cryba4 | Enriched in WT dLGN | crystallin, beta A4 [Source:MGI Symbol;Acc:MGI:102716] | 178.6 | 2.207 | 0.01885 |
| ENSMUSG00000058626 | Capn11 | Enriched in WT dLGN | calpain 11 [Source:MGI Symbol;Acc:MGI:1352490] | 21.77 | 2.25 | 0.01779 |
| ENSMUSG00000021803 | Cdhr1 | Enriched in WT dLGN | cadherin-related family member 1 [Source:MGI Symbol;Acc:MGI:2157782] | 1773 | 2.288 | 0.006542 |
| ENSMUSG00000040714 | Klc3 | Enriched in WT dLGN | kinesin light chain 3 [Source:MGI Symbol;Acc:MGI:1277971] | 241.3 | 2.29 | 0.03883 |
| ENSMUSG00000038115 | Ano2 | Enriched in WT dLGN | anoctamin 2 [Source:MGI Symbol;Acc:MGI:2387214] | 230.7 | 2.415 | 0.0218 |
| ENSMUSG00000074991 | Gabrr3 | Enriched in WT dLGN | gamma-aminobutyric acid (GABA) receptor, rho 3 [Source:MGI Symbol;Acc:MGI:3588203] | 58.13 | 2.461 | 0.005913 |
| ENSMUSG00000021123 | Rdh12 | Enriched in WT dLGN | retinol dehydrogenase 12 [Source:MGI Symbol;Acc:MGI:1925224] | 217.6 | 2.556 | 0.01211 |
| ENSMUSG00000038151 | Prdm1 | Enriched in WT dLGN | PR domain containing 1, with ZNF domain [Source:MGI Symbol;Acc:MGI:99655] | 226 | 2.601 | 0.00426 |
| ENSMUSG00000096351 | Samd11 | Enriched in WT dLGN | sterile alpha motif domain containing 11 [Source:MGI Symbol;Acc:MGI:2446220] | 834.8 | 2.618 | 0.0009793 |
| ENSMUSG00000020782 | Llgl2 | Enriched in WT dLGN | LLGL2 scribble cell polarity complex component [Source:MGI Symbol;Acc:MGI:1918843] | 188.5 | 2.648 | 0.02001 |
| ENSMUSG00000021363 | Mak | Enriched in WT dLGN | male germ cell-associated kinase [Source:MGI Symbol;Acc:MGI:96913] | 254.5 | 2.662 | 0.007378 |
| ENSMUSG00000038811 | Gngt2 | Enriched in WT dLGN | guanine nucleotide binding protein (G protein), gamma transducing activity polypeptide 2 [Source:MGI Symbol;Acc:MGI:89358 | 227.6 | 2.664 | 0.02825 |
| ENSMUSG00000079550 | Mpp4 | Enriched in WT dLGN | membrane protein, palmitoylated 4 (MAGUK p55 subfamily member 4) [Source:MGI Symbol;Acc:MGI:2386681] | 296.3 | 2.76 | 0.006504 |
| ENSMUSG00000028943 | Espn | Enriched in WT dLGN | espin [Source:MGI Symbol;Acc:MGI:1861630] | 64.68 | 2.802 | 0.004985 |
| ENSMUSG00000064330 | Pde6h | Enriched in WT dLGN | phosphodiesterase 6H, cGMP-specific, cone, gamma [Source:MGI Symbol;Acc:MGI:1925850] | 78.93 | 2.891 | 0.01737 |
| ENSMUSG00000056947 | Mab21l1 | Enriched in WT dLGN | mab-21-like 1 [Source:MGI Symbol;Acc:MGI:1333773] | 285.9 | 2.996 | 0.008368 |
| ENSMUSG00000030905 | Crym | Enriched in WT dLGN | crystallin, mu [Source:MGI Symbol;Acc:MGI:102675] | 649.3 | 3.056 | 0.01588 |
| ENSMUSG00000030206 | Gsg1 | Enriched in WT dLGN | germ cell associated 1 [Source:MGI Symbol;Acc:MGI:1194499] | 31.97 | 3.068 | 0.03198 |
| ENSMUSG00000021359 | Tfap2a | Enriched in WT dLGN | transcription factor AP-2, alpha [Source:MGI Symbol;Acc:MGI:104671] | 104 | 3.076 | 0.008368 |
| ENSMUSG00000071648 | Rom1 | Enriched in WT dLGN | rod outer segment membrane protein 1 [Source:MGI Symbol;Acc:MGI:97998] | 1310 | 3.174 | 0.007318 |
| ENSMUSG00000034452 | Slc24a1 | Enriched in WT dLGN | solute carrier family 24 (sodium/potassium/calcium exchanger), member 1 [Source:MGI Symbol;Acc:MGI:2384871] | 124.7 | 3.187 | 0.006346 |
| ENSMUSG00000020080 | Hkdc1 | Enriched in WT dLGN | hexokinase domain containing 1 [Source:MGI Symbol;Acc:MGI:2384910] | 92.62 | 3.232 | 0.04916 |
| ENSMUSG00000026609 | Ush2a | Enriched in WT dLGN | usherin [Source:MGI Symbol;Acc:MGI:1341292] | 180.8 | 3.306 | 0.004985 |
| ENSMUSG00000044429 | Cryga | Enriched in WT dLGN | crystallin, gamma A [Source:MGI Symbol;Acc:MGI:88521] | 37.52 | 3.308 | 0.002896 |
| ENSMUSG00000068154 | Insm1 | Enriched in WT dLGN | insulinoma-associated 1 [Source:MGI Symbol;Acc:MGI:1859980] | 509.4 | 3.387 | 0.001173 |
| ENSMUSG00000031142 | Cacna1f | Enriched in WT dLGN | calcium channel, voltage-dependent, alpha 1F subunit [Source:MGI Symbol;Acc:MGI:1859639] | 80.7 | 3.396 | 0.0001159 |
| ENSMUSG00000067220 | Cnga1 | Enriched in WT dLGN | cyclic nucleotide gated channel alpha 1 [Source:MGI Symbol;Acc:MGI:88436] | 375.3 | 3.453 | 0.006745 |
| ENSMUSG00000029352 | Crybb3 | Enriched in WT dLGN | crystallin, beta B3 [Source:MGI Symbol;Acc:MGI:102717] | 300.7 | 3.501 | 0.001173 |
| ENSMUSG00000022237 | Ankrd33b | Enriched in WT dLGN | ankyrin repeat domain 33B [Source:MGI Symbol;Acc:MGI:1917904] | 537.3 | 3.506 | 0.006242 |
| ENSMUSG00000057132 | Rpgrip1 | Enriched in WT dLGN | retinitis pigmentosa GTPase regulator interacting protein 1 [Source:MGI Symbol;Acc:MGI:1932134] | 807.3 | 3.556 | 0.0009445 |
| ENSMUSG00000037446 | Tulp1 | Enriched in WT dLGN | tubby like protein 1 [Source:MGI Symbol;Acc:MGI:109571] | 1171 | 3.614 | 0.002347 |
| ENSMUSG00000097050 | Gm9918 | Enriched in WT dLGN | predicted gene 9918 [Source:MGI Symbol;Acc:MGI:3646315] | 72.57 | 3.748 | 0.008368 |
| ENSMUSG00000029410 | Ppef2 | Enriched in WT dLGN | protein phosphatase, EF hand calcium-binding domain 2 [Source:MGI Symbol;Acc:MGI:1342304] | 245.8 | 3.774 | 0.005609 |
| ENSMUSG00000040632 | Nrl | Enriched in WT dLGN | neural retina leucine zipper gene [Source:MGI Symbol;Acc:MGI:102567] | 1510 | 3.827 | 0.002914 |
| ENSMUSG00000056494 | Cngb3 | Enriched in WT dLGN | cyclic nucleotide gated channel beta 3 [Source:MGI Symbol;Acc:MGI:1353562] | 36.69 | 3.929 | 0.04832 |
| ENSMUSG00000024992 | Pde6c | Enriched in WT dLGN | phosphodiesterase 6C, cGMP specific, cone, alpha prime [Source:MGI Symbol;Acc:MGI:105956] | 75.25 | 3.935 | 0.001092 |
| ENSMUSG00000020907 | Rcvrn | Enriched in WT dLGN | recoverin [Source:MGI Symbol;Acc:MGI:97883] | 470.4 | 3.964 | 0.004985 |
| ENSMUSG00000031789 | Cngb1 | Enriched in WT dLGN | cyclic nucleotide gated channel beta 1 [Source:MGI Symbol;Acc:MGI:2664102] | 674.9 | 4.076 | 0.002581 |
| ENSMUSG00000023439 | Gnb3 | Enriched in WT dLGN | guanine nucleotide binding protein (G protein), beta 3 [Source:MGI Symbol;Acc:MGI:95785] | 519.4 | 4.125 | 0.003401 |
| ENSMUSG00000045493 | Bhlhe23 | Enriched in WT dLGN | basic helix-loop-helix family, member e23 [Source:MGI Symbol;Acc:MGI:2153710] | 65.86 | 4.252 | 0.009449 |
| ENSMUSG00000093865 | Lrit3 | Enriched in WT dLGN | leucine-rich repeat, immunoglobulin-like and transmembrane domains 3 [Source:MGI Symbol;Acc:MGI:2685267] | 30.16 | 4.265 | 0.001173 |
| ENSMUSG00000032343 | Impg1 | Enriched in WT dLGN | interphotoreceptor matrix proteoglycan 1 [Source:MGI Symbol;Acc:MGI:1926876] | 167.2 | 4.358 | 0.001092 |
| ENSMUSG00000037161 | Mgarp | Enriched in WT dLGN | mitochondria localized glutamic acid rich protein [Source:MGI Symbol;Acc:MGI:1914999] | 458.7 | 4.361 | 0.006504 |
| ENSMUSG00000056043 | Rgs9bp | Enriched in WT dLGN | regulator of G-protein signalling 9 binding protein [Source:MGI Symbol;Acc:MGI:2384418] | 211.6 | 4.376 | 0.001359 |
| ENSMUSG00000034829 | Nxnl1 | Enriched in WT dLGN | nucleoredoxin-like 1 [Source:MGI Symbol;Acc:MGI:1924446] | 201.4 | 4.415 | 0.009366 |
| ENSMUSG00000092349 | Smim40 | Enriched in WT dLGN | small integral membrane protein 40 [Source:MGI Symbol;Acc:MGI:3054967] | 129.7 | 4.435 | 0.001574 |
| ENSMUSG00000056055 | Sag | Enriched in WT dLGN | S-antigen, retina and pineal gland (arrestin) [Source:MGI Symbol;Acc:MGI:98227] | 2556 | 4.448 | 0.005873 |
| ENSMUSG00000035270 | Impg2 | Enriched in WT dLGN | interphotoreceptor matrix proteoglycan 2 [Source:MGI Symbol;Acc:MGI:3044955] | 690.4 | 4.465 | 0.001776 |
| ENSMUSG00000031293 | Rs1 | Enriched in WT dLGN | retinoschisis (X-linked, juvenile) 1 (human) [Source:MGI Symbol;Acc:MGI:1336189] | 736.2 | 4.493 | 0.003401 |
| ENSMUSG00000046049 | Rp1l1 | Enriched in WT dLGN | retinitis pigmentosa 1 homolog like 1 [Source:MGI Symbol;Acc:MGI:2384303] | 539.3 | 4.508 | 0.008667 |
| ENSMUSG00000025386 | Pde6g | Enriched in WT dLGN | phosphodiesterase 6G, cGMP-specific, rod, gamma [Source:MGI Symbol;Acc:MGI:97526] | 759.4 | 4.522 | 0.008666 |
| ENSMUSG00000047298 | Kcnv2 | Enriched in WT dLGN | potassium channel, subfamily V, member 2 [Source:MGI Symbol;Acc:MGI:2670981] | 331.8 | 4.559 | 0.02336 |
| ENSMUSG00000103119 | Gm37583 | Enriched in WT dLGN | predicted gene, 37583 [Source:MGI Symbol;Acc:MGI:5610811] | 105.6 | 4.571 | 0.001173 |
| ENSMUSG00000030324 | Rho | Enriched in WT dLGN | rhodopsin [Source:MGI Symbol;Acc:MGI:97914] | 4213 | 4.573 | 0.01199 |
| ENSMUSG00000058831 | Opn1sw | Enriched in WT dLGN | opsin 1 (cone pigments), short-wave-sensitive (color blindness, tritan) [Source:MGI Symbol;Acc:MGI:99438] | 155.8 | 4.583 | 0.009835 |
| ENSMUSG00000042240 | Crybb2 | Enriched in WT dLGN | crystallin, beta B2 [Source:MGI Symbol;Acc:MGI:88519] | 255.7 | 4.61 | 0.0006541 |
| ENSMUSG00000025927 | Tfap2b | Enriched in WT dLGN | transcription factor AP-2 beta [Source:MGI Symbol;Acc:MGI:104672] | 370.9 | 4.617 | 0.006504 |
| ENSMUSG00000110344 | Gm45716 | Enriched in WT dLGN | predicted gene 45716 [Source:MGI Symbol;Acc:MGI:5804831] | 43.95 | 4.807 | 0.00002877 |
| ENSMUSG00000040554 | Aipl1 | Enriched in WT dLGN | aryl hydrocarbon receptor-interacting protein-like 1 [Source:MGI Symbol;Acc:MGI:2148800] | 316.7 | 4.818 | 0.001544 |
| ENSMUSG00000036480 | Prss56 | Enriched in WT dLGN | protease, serine 56 [Source:MGI Symbol;Acc:MGI:1916703] | 140.8 | 4.866 | 0.005021 |
| ENSMUSG00000005649 | Cabp5 | Enriched in WT dLGN | calcium binding protein 5 [Source:MGI Symbol;Acc:MGI:1352746] | 61.32 | 4.897 | 0.001464 |

| | | | | | | |
|---|---|---|---|---|---|---|
| ENSMUSG00000029663 | Gngt1 | Enriched in WT dLGN | guanine nucleotide binding protein (G protein), gamma transducing activity polypeptide 1 [Source:MGI Symbol;Acc:MGI:10916 | 688.6 | 4.906 | 0.01779 |
| ENSMUSG00000074365 | Crxos | Enriched in WT dLGN | cone-rod homeobox, opposite strand [Source:MGI Symbol;Acc:MGI:2451355] | 48.47 | 4.941 | 0.01215 |
| ENSMUSG00000047034 | Ankrd33 | Enriched in WT dLGN | ankyrin repeat domain 33 [Source:MGI Symbol;Acc:MGI:2443398] | 128.9 | 4.954 | 0.02542 |
| ENSMUSG00000031450 | Grk1 | Enriched in WT dLGN | G protein-coupled receptor kinase 1 [Source:MGI Symbol;Acc:MGI:1345146] | 407.8 | 4.961 | 0.008286 |
| ENSMUSG00000006007 | Pdc | Enriched in WT dLGN | phosducin [Source:MGI Symbol;Acc:MGI:98090] | 2322 | 4.996 | 0.006685 |
| ENSMUSG00000048015 | Neurod4 | Enriched in WT dLGN | neurogenic differentiation 4 [Source:MGI Symbol;Acc:MGI:108055] | 512.1 | 5.043 | 0.002707 |
| ENSMUSG00000033501 | Crygs | Enriched in WT dLGN | crystallin, gamma S [Source:MGI Symbol;Acc:MGI:1298216] | 142.2 | 5.056 | 0.0009793 |
| ENSMUSG00000021239 | Vsx2 | Enriched in WT dLGN | visual system homeobox 2 [Source:MGI Symbol;Acc:MGI:88401] | 917.1 | 5.058 | 0.001861 |
| ENSMUSG00000051860 | Samd7 | Enriched in WT dLGN | sterile alpha motif domain containing 7 [Source:MGI Symbol;Acc:MGI:1923203] | 304.5 | 5.073 | 0.001773 |
| ENSMUSG00000086322 | E130218I03Rik | Enriched in WT dLGN | RIKEN cDNA E130218I03 gene [Source:MGI Symbol;Acc:MGI:3528958] | 327.2 | 5.094 | 0.0001776 |
| ENSMUSG00000043418 | Lrit2 | Enriched in WT dLGN | leucine-rich repeat, immunoglobulin-like and transmembrane domains 2 [Source:MGI Symbol;Acc:MGI:2444885] | 118.4 | 5.111 | 0.0009791 |
| ENSMUSG00000085139 | A730046J19Rik | Enriched in WT dLGN | RIKEN cDNA A730046J19 gene [Source:MGI Symbol;Acc:MGI:2442684] | 62.16 | 5.118 | 0.006346 |
| ENSMUSG00000064356 | mt-Atp8 | Enriched in WT dLGN | mitochondrially encoded ATP synthase 8 [Source:MGI Symbol;Acc:MGI:99926] | 32.09 | 5.163 | 0.005873 |
| ENSMUSG00000041578 | Crx | Enriched in WT dLGN | cone-rod homeobox [Source:MGI Symbol;Acc:MGI:1194883] | 1591 | 5.185 | 0.01081 |
| ENSMUSG00000070683 | Lactbl1 | Enriched in WT dLGN | lactamase, beta-like 1 [Source:MGI Symbol;Acc:MGI:2448566] | 62.79 | 5.202 | 0.001406 |
| ENSMUSG00000041534 | Rbp3 | Enriched in WT dLGN | retinol binding protein 3, interstitial [Source:MGI Symbol;Acc:MGI:97878] | 2863 | 5.22 | 0.001942 |
| ENSMUSG00000026468 | Lhx4 | Enriched in WT dLGN | LIM homeobox protein 4 [Source:MGI Symbol;Acc:MGI:101776] | 133.7 | 5.282 | 0.001092 |
| ENSMUSG00000023978 | Prph2 | Enriched in WT dLGN | peripherin 2 [Source:MGI Symbol;Acc:MGI:102791] | 1888 | 5.302 | 0.001861 |
| ENSMUSG00000024518 | Rax | Enriched in WT dLGN | retina and anterior neural fold homeobox [Source:MGI Symbol;Acc:MGI:109632] | 219.8 | 5.324 | 0.006346 |
| ENSMUSG00000041044 | Lrit1 | Enriched in WT dLGN | leucine-rich repeat, immunoglobulin-like and transmembrane domains 1 [Source:MGI Symbol;Acc:MGI:2385320] | 351.3 | 5.325 | 0.006542 |
| ENSMUSG00000000617 | Grm6 | Enriched in WT dLGN | glutamate receptor, metabotropic 6 [Source:MGI Symbol;Acc:MGI:1351343] | 184.6 | 5.34 | 0.001304 |
| ENSMUSG00000008932 | Slc1a7 | Enriched in WT dLGN | solute carrier family 1 (glutamate transporter), member 7 [Source:MGI Symbol;Acc:MGI:2444087] | 187 | 5.342 | 0.0006541 |
| ENSMUSG00000024519 | Cplx4 | Enriched in WT dLGN | complexin 4 [Source:MGI Symbol;Acc:MGI:2685803] | 134.4 | 5.391 | 0.003603 |
| ENSMUSG00000067438 | Hmx1 | Enriched in WT dLGN | H6 homeobox 1 [Source:MGI Symbol;Acc:MGI:107178] | 52.16 | 5.465 | 0.003568 |
| ENSMUSG00000025900 | Rp1 | Enriched in WT dLGN | retinitis pigmentosa 1 (human) [Source:MGI Symbol;Acc:MGI:1341105] | 1607 | 5.476 | 0.0007934 |
| ENSMUSG00000029491 | Pde6b | Enriched in WT dLGN | phosphodiesterase 6B, cGMP, rod receptor, beta polypeptide [Source:MGI Symbol;Acc:MGI:97525] | 1069 | 5.573 | 0.003524 |
| ENSMUSG00000027530 | Fabp12 | Enriched in WT dLGN | fatty acid binding protein 12 [Source:MGI Symbol;Acc:MGI:1922747] | 73.46 | 5.647 | 0.005725 |
| ENSMUSG00000053773 | Rdh8 | Enriched in WT dLGN | retinol dehydrogenase 8 [Source:MGI Symbol;Acc:MGI:2685028] | 53.24 | 5.875 | 0.02682 |
| ENSMUSG00000040478 | Prdm13 | Enriched in WT dLGN | PR domain containing 13 [Source:MGI Symbol;Acc:MGI:2448528] | 65.96 | 6.082 | 0.004551 |
| ENSMUSG00000044375 | Pcare | Enriched in WT dLGN | photoreceptor cilium actin regulator [Source:MGI Symbol;Acc:MGI:2385061] | 194 | 6.238 | 0.001381 |
| ENSMUSG00000032292 | Nr2e3 | Enriched in WT dLGN | nuclear receptor subfamily 2, group E, member 3 [Source:MGI Symbol;Acc:MGI:1346317] | 1659 | 6.332 | 0.001173 |
| ENSMUSG00000049908 | Gja8 | Enriched in WT dLGN | gap junction protein, alpha 8 [Source:MGI Symbol;Acc:MGI:99953] | 37.76 | 6.529 | 0.009666 |
| ENSMUSG00000025389 | Mip | Enriched in WT dLGN | major intrinsic protein of lens fiber [Source:MGI Symbol;Acc:MGI:96990] | 71.98 | 6.649 | 0.001359 |
| ENSMUSG00000006546 | Cryba2 | Enriched in WT dLGN | crystallin, beta A2 [Source:MGI Symbol;Acc:MGI:104336] | 138.5 | 7.62 | 0.0006541 |
| ENSMUSG00000025952 | Crygc | Enriched in WT dLGN | crystallin, gamma C [Source:MGI Symbol;Acc:MGI:88523] | 87.43 | 7.64 | 0.001637 |
| ENSMUSG00000024041 | Cryaa | Enriched in WT dLGN | crystallin, alpha A [Source:MGI Symbol;Acc:MGI:88515] | 1048 | 8.072 | 0.0001159 |
| ENSMUSG00000070870 | Cryge | Enriched in WT dLGN | crystallin, gamma E [Source:MGI Symbol;Acc:MGI:88525] | 106.4 | 8.132 | 0.0009445 |
| ENSMUSG00000025945 | Crygf | Enriched in WT dLGN | crystallin, gamma F [Source:MGI Symbol;Acc:MGI:88526] | 38.99 | 8.15 | 0.0001776 |
| ENSMUSG00000073658 | Crygb | Enriched in WT dLGN | crystallin, gamma B [Source:MGI Symbol;Acc:MGI:88522] | 153.5 | 8.169 | 0.0001159 |
| ENSMUSG00000067299 | Crygd | Enriched in WT dLGN | crystallin, gamma D [Source:MGI Symbol;Acc:MGI:88524] | 144.9 | 8.563 | 0.0006541 |
| ENSMUSG00000000724 | Cryba1 | Enriched in WT dLGN | crystallin, beta A1 [Source:MGI Symbol;Acc:MGI:88518] | 473.5 | 9.01 | 0.0001746 |

**Contrast numerator: KO_SCN. Contrast denominator: Het_SCN.**

| ensembl_gene_id | external_gene_name | Significance | description | baseMean | logFC | adj.P.Val |
|---|---|---|---|---|---|---|
| ENSMUSG00000041578 | Crx | Enriched in Het SCN | cone-rod homeobox [Source:MGI Symbol;Acc:MGI:1194883] | 174.2 | -6.188 | 0.00877 |
| ENSMUSG00000021848 | Otx2 | Enriched in Het SCN | orthodenticle homeobox 2 [Source:MGI Symbol;Acc:MGI:97451] | 109.6 | -5.258 | 0.01218 |
| ENSMUSG00000025386 | Pde6g | Enriched in Het SCN | phosphodiesterase 6G, cGMP-specific, rod, gamma [Source:MGI Symbol;Acc:MGI:97526] | 98.91 | -4.61 | 0.01546 |
| ENSMUSG00000037161 | Mgarp | Enriched in Het SCN | mitochondria localized glutamic acid rich protein [Source:MGI Symbol;Acc:MGI:1914999] | 51.11 | -4.597 | 0.02594 |
| ENSMUSG00000034384 | Barhl2 | Enriched in Het SCN | BarH like homeobox 2 [Source:MGI Symbol;Acc:MGI:1859314] | 64.55 | -4.571 | 0.004537 |
| ENSMUSG00000006007 | Pdc | Enriched in Het SCN | phosducin [Source:MGI Symbol;Acc:MGI:98090] | 250.2 | -4.475 | 0.04073 |
| ENSMUSG00000060461 | Dppa5a | Enriched in Het SCN | developmental pluripotency associated 5A [Source:MGI Symbol;Acc:MGI:101800] | 58.21 | -4.413 | 0.000001905 |
| ENSMUSG00000032292 | Nr2e3 | Enriched in Het SCN | nuclear receptor subfamily 2, group E, member 3 [Source:MGI Symbol;Acc:MGI:1346317] | 228.3 | -4.025 | 0.04811 |
| ENSMUSG00000032446 | Eomes | Enriched in Het SCN | eomesodermin [Source:MGI Symbol;Acc:MGI:1201683] | 160.9 | -3.724 | 0.000305 |
| ENSMUSG00000026890 | Lhx6 | Enriched in Het SCN | LIM homeobox protein 6 [Source:MGI Symbol;Acc:MGI:1306803] | 590.5 | -3.544 | 0.003639 |
| ENSMUSG00000032269 | Htr3a | Enriched in Het SCN | 5-hydroxytryptamine (serotonin) receptor 3A [Source:MGI Symbol;Acc:MGI:96282] | 50.15 | -3.365 | 0.00008932 |
| ENSMUSG00000043969 | Emx2 | Enriched in Het SCN | empty spiracles homeobox 2 [Source:MGI Symbol;Acc:MGI:95388] | 53.84 | -3.266 | 0.003391 |
| ENSMUSG00000071230 | Npw | Enriched in Het SCN | neuropeptide W [Source:MGI Symbol;Acc:MGI:2685781] | 38.09 | -3.114 | 0.003001 |
| ENSMUSG00000049796 | Crh | Enriched in Het SCN | corticotropin releasing hormone [Source:MGI Symbol;Acc:MGI:88496] | 143.4 | -3.029 | 3.735E-07 |
| ENSMUSG00000091519 | Skor2 | Enriched in Het SCN | SKI family transcriptional corepressor 2 [Source:MGI Symbol;Acc:MGI:3645984] | 63.1 | -2.939 | 0.0008393 |
| ENSMUSG00000054146 | Krt15 | Enriched in Het SCN | keratin 15 [Source:MGI Symbol;Acc:MGI:96689] | 39.18 | -2.904 | 0.001604 |
| ENSMUSG00000051354 | Samd3 | Enriched in Het SCN | sterile alpha motif domain containing 3 [Source:MGI Symbol;Acc:MGI:2685469] | 51.06 | -2.626 | 0.001604 |
| ENSMUSG00000032346 | Ooep | Enriched in Het SCN | oocyte expressed protein [Source:MGI Symbol;Acc:MGI:1915218] | 104.6 | -2.62 | 0.00001531 |
| ENSMUSG00000024518 | Rax | Enriched in Het SCN | retina and anterior neural fold homeobox [Source:MGI Symbol;Acc:MGI:109632] | 58.51 | -2.546 | 0.01136 |
| ENSMUSG00000035033 | Tbr1 | Enriched in Het SCN | T-box brain transcription factor 1 [Source:MGI Symbol;Acc:MGI:107404] | 133.4 | -2.388 | 0.03194 |
| ENSMUSG00000040148 | Hmx3 | Enriched in Het SCN | H6 homeobox 3 [Source:MGI Symbol;Acc:MGI:107160] | 196.4 | -2.208 | 0.0002422 |
| ENSMUSG00000050100 | Hmx2 | Enriched in Het SCN | H6 homeobox 2 [Source:MGI Symbol;Acc:MGI:107159] | 176.8 | -2.179 | 0.000006336 |
| ENSMUSG00000096225 | Lhx8 | Enriched in Het SCN | LIM homeobox protein 8 [Source:MGI Symbol;Acc:MGI:1096343] | 273.9 | -2.163 | 0.03233 |
| ENSMUSG00000020950 | Foxg1 | Enriched in Het SCN | forkhead box G1 [Source:MGI Symbol;Acc:MGI:1347464] | 152.1 | -2.116 | 0.004537 |
| ENSMUSG00000024650 | Slc22a6 | Enriched in Het SCN | solute carrier family 22 (organic anion transporter), member 6 [Source:MGI Symbol;Acc:MGI:892001] | 190.8 | -1.892 | 0.000888 |
| ENSMUSG00000021950 | Anxa8 | Enriched in Het SCN | annexin A8 [Source:MGI Symbol;Acc:MGI:1201374] | 66.97 | -1.882 | 0.009012 |
| ENSMUSG00000030270 | Cpne9 | Enriched in Het SCN | copine family member IX [Source:MGI Symbol;Acc:MGI:2443052] | 89.72 | -1.856 | 0.008277 |
| ENSMUSG00000020415 | Pttg1 | Enriched in Het SCN | pituitary tumor-transforming gene 1 [Source:MGI Symbol;Acc:MGI:1353578] | 488.8 | -1.792 | 6.991E-08 |
| ENSMUSG00000032105 | Pdzd3 | Enriched in Het SCN | PDZ domain containing 3 [Source:MGI Symbol;Acc:MGI:2429554] | 73.28 | -1.604 | 0.02144 |
| ENSMUSG00000040258 | Nxph4 | Enriched in Het SCN | neurexophilin 4 [Source:MGI Symbol;Acc:MGI:1336197] | 420.8 | -1.586 | 0.04073 |
| ENSMUSG00000044229 | Nxpe4 | Enriched in Het SCN | neurexophilin and PC-esterase domain family, member 4 [Source:MGI Symbol;Acc:MGI:1924792] | 931.1 | -1.559 | 4.943E-08 |
| ENSMUSG00000030074 | Gxylt2 | Enriched in Het SCN | glucoside xylosyltransferase 2 [Source:MGI Symbol;Acc:MGI:2682940] | 92.12 | -1.532 | 0.02344 |
| ENSMUSG00000029373 | Pf4 | Enriched in Het SCN | platelet factor 4 [Source:MGI Symbol;Acc:MGI:1888711] | 34.49 | -1.528 | 0.03222 |
| ENSMUSG00000066438 | Plekhd1 | Enriched in Het SCN | pleckstrin homology domain containing, family D (with coiled-coil domains) member 1 [Source:MGI Symbol;Acc:MGI:3036228] | 118 | -1.509 | 0.0173 |
| ENSMUSG00000039109 | F13a1 | Enriched in Het SCN | coagulation factor XIII, A1 subunit [Source:MGI Symbol;Acc:MGI:1921395] | 96.92 | -1.503 | 0.02269 |
| ENSMUSG00000020053 | Igf1 | Enriched in Het SCN | insulin-like growth factor 1 [Source:MGI Symbol;Acc:MGI:96432] | 219.8 | -1.489 | 0.000005299 |
| ENSMUSG00000029843 | Slc13a4 | Enriched in Het SCN | solute carrier family 13 (sodium/sulfate symporters), member 4 [Source:MGI Symbol;Acc:MGI:2442367] | 473.7 | -1.446 | 0.03305 |
| ENSMUSG00000046861 | Hectd3 | Enriched in Het SCN | HECT domain E3 ubiquitin protein ligase 3 [Source:MGI Symbol;Acc:MGI:1923858] | 1309 | -1.394 | 0.0006749 |
| ENSMUSG00000030108 | Slc6a13 | Enriched in Het SCN | solute carrier family 6 (neurotransmitter transporter, GABA), member 13 [Source:MGI Symbol;Acc:MGI:95629] | 300.4 | -1.344 | 0.0005386 |
| ENSMUSG00000104282 | Gm37460 | Enriched in Het SCN | predicted gene, 37460 [Source:MGI Symbol;Acc:MGI:5610688] | 62.82 | -1.327 | 0.03515 |
| ENSMUSG00000026051 | Ecrg4 | Enriched in Het SCN | ECRG4 augurin precursor [Source:MGI Symbol;Acc:MGI:1926146] | 179.1 | -1.297 | 0.03133 |
| ENSMUSG00000004347 | Pde1c | Enriched in Het SCN | phosphodiesterase 1C [Source:MGI Symbol;Acc:MGI:108413] | 615.2 | -1.269 | 0.0001551 |
| ENSMUSG00000078639 | Gm12695 | Enriched in Het SCN | predicted gene 12695 [Source:MGI Symbol;Acc:MGI:3650206] | 48.91 | -1.254 | 0.007815 |
| ENSMUSG00000037206 | Islr | Enriched in Het SCN | immunoglobulin superfamily containing leucine-rich repeat [Source:MGI Symbol;Acc:MGI:1349645] | 506.8 | -1.235 | 0.0002949 |
| ENSMUSG00000075707 | Dio3 | Enriched in Het SCN | deiodinase, iodothyronine type III [Source:MGI Symbol;Acc:MGI:1306782] | 178.8 | -1.231 | 0.02774 |
| ENSMUSG00000052504 | Epha3 | Enriched in Het SCN | Eph receptor A3 [Source:MGI Symbol;Acc:MGI:99612] | 159.3 | -1.23 | 0.001046 |
| ENSMUSG00000071648 | Rom1 | Enriched in Het SCN | rod outer segment membrane protein 1 [Source:MGI Symbol;Acc:MGI:97998] | 260.9 | -1.221 | 0.04073 |
| ENSMUSG00000001506 | Col1a1 | Enriched in Het SCN | collagen, type I, alpha 1 [Source:MGI Symbol;Acc:MGI:88467] | 1820 | -1.211 | 0.002187 |
| ENSMUSG00000042498 | Radx | Enriched in Het SCN | RPA1 related single stranded DNA binding protein, X-linked [Source:MGI Symbol;Acc:MGI:2147848] | 164.8 | -1.175 | 0.02801 |
| ENSMUSG00000019230 | Lhx9 | Enriched in Het SCN | LIM homeobox protein 9 [Source:MGI Symbol;Acc:MGI:1316721] | 181.9 | -1.108 | 0.0006749 |
| ENSMUSG00000035279 | Ssc5d | Enriched in Het SCN | scavenger receptor cysteine rich family, 5 domains [Source:MGI Symbol;Acc:MGI:3606211] | 132.8 | -1.068 | 0.009592 |
| ENSMUSG00000004885 | Crabp2 | Enriched in Het SCN | cellular retinoic acid binding protein II [Source:MGI Symbol;Acc:MGI:88491] | 130 | -1.057 | 0.005939 |
| ENSMUSG00000046167 | Gldn | Enriched in Het SCN | gliomedin [Source:MGI Symbol;Acc:MGI:2388361] | 103.7 | -1.047 | 0.009869 |
| ENSMUSG00000032332 | Col12a1 | Enriched in Het SCN | collagen, type XII, alpha 1 [Source:MGI Symbol;Acc:MGI:88448] | 564.2 | -1.036 | 2.927E-12 |
| ENSMUSG00000031303 | Map3k15 | Enriched in Het SCN | mitogen-activated protein kinase kinase kinase 15 [Source:MGI Symbol;Acc:MGI:2448588] | 172.1 | -1.034 | 0.0007223 |
| ENSMUSG00000051236 | Msrb3 | Enriched in KO SCN | methionine sulfoxide reductase B3 [Source:MGI Symbol;Acc:MGI:2443538] | 284.8 | 1.051 | 0.002823 |
| ENSMUSG00000032135 | Mcam | Enriched in KO SCN | melanoma cell adhesion molecule [Source:MGI Symbol;Acc:MGI:1933966] | 652.5 | 1.105 | 0.0001462 |
| ENSMUSG00000022836 | Mylk | Enriched in KO SCN | myosin, light polypeptide kinase [Source:MGI Symbol;Acc:MGI:894806] | 480.5 | 1.11 | 0.0004494 |
| ENSMUSG00000067818 | Myl9 | Enriched in KO SCN | myosin, light polypeptide 9, regulatory [Source:MGI Symbol;Acc:MGI:2138915] | 1067 | 1.112 | 0.02872 |
| ENSMUSG00000046280 | She | Enriched in KO SCN | src homology 2 domain-containing transforming protein E [Source:MGI Symbol;Acc:MGI:1099462] | 173.6 | 1.167 | 0.0005076 |
| ENSMUSG00000020154 | Ptprb | Enriched in KO SCN | protein tyrosine phosphatase, receptor type, B [Source:MGI Symbol;Acc:MGI:97809] | 1226 | 1.215 | 0.01867 |
| ENSMUSG00000022371 | Col14a1 | Enriched in KO SCN | collagen, type XIV, alpha 1 [Source:MGI Symbol;Acc:MGI:1341272] | 56.5 | 1.28 | 0.01546 |
| ENSMUSG00000045903 | Npas4 | Enriched in KO SCN | neuronal PAS domain protein 4 [Source:MGI Symbol;Acc:MGI:2664186] | 356.2 | 1.286 | 4.943E-08 |
| ENSMUSG00000004328 | Hif3a | Enriched in KO SCN | hypoxia inducible factor 3, alpha subunit [Source:MGI Symbol;Acc:MGI:1859778] | 172.9 | 1.289 | 0.0267 |
| ENSMUSG00000024049 | Myom1 | Enriched in KO SCN | myomesin 1 [Source:MGI Symbol;Acc:MGI:1341430] | 96.55 | 1.334 | 0.002136 |
| ENSMUSG00000037166 | Ppp1r14a | Enriched in KO SCN | protein phosphatase 1, regulatory inhibitor subunit 14A [Source:MGI Symbol;Acc:MGI:1931139] | 48.01 | 1.374 | 0.0013 |
| ENSMUSG00000001930 | Vwf | Enriched in KO SCN | Von Willebrand factor [Source:MGI Symbol;Acc:MGI:98941] | 904 | 1.488 | 0.02074 |
| ENSMUSG00000020788 | Atp2a3 | Enriched in KO SCN | ATPase, Ca++ transporting, ubiquitous [Source:MGI Symbol;Acc:MGI:1194503] | 316.3 | 1.559 | 0.0007678 |
| ENSMUSG00000068196 | Col8a1 | Enriched in KO SCN | collagen, type VIII, alpha 1 [Source:MGI Symbol;Acc:MGI:88463] | 461.7 | 1.562 | 0.0001109 |
| ENSMUSG00000048489 | Depp1 | Enriched in KO SCN | DEPP1 autophagy regulator [Source:MGI Symbol;Acc:MGI:1918730] | 57.45 | 1.588 | 0.0003643 |
| ENSMUSG00000041445 | Mmrn2 | Enriched in KO SCN | multimerin 2 [Source:MGI Symbol;Acc:MGI:2385618] | 192.3 | 1.631 | 0.000001675 |
| ENSMUSG00000015467 | Egfl8 | Enriched in KO SCN | EGF-like domain 8 [Source:MGI Symbol;Acc:MGI:1932094] | 142.2 | 1.738 | 0.000008988 |
| ENSMUSG00000055044 | Pdlim1 | Enriched in KO SCN | PDZ and LIM domain 1 (elfin) [Source:MGI Symbol;Acc:MGI:1860611] | 82.6 | 1.748 | 0.00161 |
| ENSMUSG00000021904 | Sema3g | Enriched in KO SCN | sema domain, immunoglobulin domain (Ig), short basic domain, secreted, (semaphorin) 3G [Source:MGI Symbol;Acc:MGI:304124 | 264.1 | 1.796 | 0.00005502 |
| ENSMUSG00000031377 | Bmx | Enriched in KO SCN | BMX non-receptor tyrosine kinase [Source:MGI Symbol;Acc:MGI:1101778] | 81.29 | 1.858 | 0.006665 |
| ENSMUSG00000028776 | Tinagl1 | Enriched in KO SCN | tubulointerstitial nephritis antigen-like 1 [Source:MGI Symbol;Acc:MGI:2137617] | 228 | 2.17 | 0.000003181 |
| ENSMUSG00000057123 | Gja5 | Enriched in KO SCN | gap junction protein, alpha 5 [Source:MGI Symbol;Acc:MGI:95716] | 245.6 | 2.23 | 0.000001561 |
| ENSMUSG00000020787 | P2rx1 | Enriched in KO SCN | purinergic receptor P2X, ligand-gated ion channel, 1 [Source:MGI Symbol;Acc:MGI:1098235] | 30.44 | 2.261 | 0.01786 |
| ENSMUSG00000030048 | Gkn3 | Enriched in KO SCN | gastrokine 3 [Source:MGI Symbol;Acc:MGI:1916138] | 73.89 | 2.42 | 1.563E-07 |
| ENSMUSG00000032085 | Tagln | Enriched in KO SCN | transgelin [Source:MGI Symbol;Acc:MGI:106012] | 483.2 | 2.734 | 0.006641 |
| ENSMUSG00000021186 | Fbln5 | Enriched in KO SCN | fibulin 5 [Source:MGI Symbol;Acc:MGI:1346091] | 999.8 | 2.747 | 0.0008535 |
| ENSMUSG00000019326 | Aoc3 | Enriched in KO SCN | amine oxidase, copper containing 3 [Source:MGI Symbol;Acc:MGI:1306797] | 110.9 | 2.871 | 0.0001363 |
| ENSMUSG00000029675 | Eln | Enriched in KO SCN | elastin [Source:MGI Symbol;Acc:MGI:95317] | 11630 | 2.882 | 0.00006029 |
| ENSMUSG00000025610 | Map3k7cl | Enriched in KO SCN | Map3k7 C-terminal like [Source:MGI Symbol;Acc:MGI:2446584] | 74.1 | 2.911 | 0.0005205 |
| ENSMUSG00000027656 | Ccn5 | Enriched in KO SCN | cellular communication network factor 5 [Source:MGI Symbol;Acc:MGI:1328326] | 25.21 | 3.162 | 0.0007089 |
| ENSMUSG00000001349 | Cnn1 | Enriched in KO SCN | calponin 1 [Source:MGI Symbol;Acc:MGI:104979] | 58.85 | 3.463 | 0.00136 |
| ENSMUSG00000018830 | Myh11 | Enriched in KO SCN | myosin, heavy polypeptide 11, smooth muscle [Source:MGI Symbol;Acc:MGI:102643] | 356.5 | 3.726 | 0.000001561 |
| ENSMUSG00000035783 | Acta2 | Enriched in KO SCN | actin, alpha 2, smooth muscle, aorta [Source:MGI Symbol;Acc:MGI:87909] | 1013 | 3.921 | 0.00005502 |
| ENSMUSG00000059430 | Actg2 | Enriched in KO SCN | actin, gamma 2, smooth muscle, enteric [Source:MGI Symbol;Acc:MGI:104589] | 63.21 | 4.279 | 5.168E-07 |
| ENSMUSG00000021678 | F2rl1 | Enriched in KO SCN | coagulation factor II (thrombin) receptor-like 1 [Source:MGI Symbol;Acc:MGI:101910] | 20.96 | 4.749 | 0.00002567 |
| ENSMUSG00000072476 | Gm9008 | Enriched in KO SCN | predicted pseudogene 9008 [Source:MGI Symbol;Acc:MGI:3644000] | 92.94 | 5.519 | 0.001173 |

**Contrast numerator: WT_SCN. Contrast denominator: Het_SCN.**

| ensembl_gene_id | external_gene_name | Significance | description | baseMean | logFC | adj.P.Val |
|---|---|---|---|---|---|---|
| ENSMUSG00000105243 | Gm43444 | Enriched in Het SCN | predicted gene 43444 [Source:MGI Symbol;Acc:MGI:5663581] | 59.97 | -9 | 3.251E-14 |
| ENSMUSG00000040264 | Gbp2b | Enriched in Het SCN | guanylate binding protein 2b [Source:MGI Symbol;Acc:MGI:95666] | 60.56 | -8 | 7.138E-11 |
| ENSMUSG00000108900 | Ccdc194 | Enriched in Het SCN | coiled-coil domain containing 194 [Source:MGI Symbol;Acc:MGI:3588239] | 27 | -4.566 | 0.000008395 |
| ENSMUSG00000071230 | Npw | Enriched in Het SCN | neuropeptide W [Source:MGI Symbol;Acc:MGI:2685781] | 36.76 | -2.98 | 0.003493 |
| ENSMUSG00000027199 | Gatm | Enriched in Het SCN | glycine amidinotransferase (L-arginine:glycine amidinotransferase) [Source:MGI Symbol;Acc:MGI:1914342] | 67.56 | -2.698 | 0.02605 |
| ENSMUSG00000072572 | Slc39a2 | Enriched in Het SCN | solute carrier family 39 (zinc transporter), member 2 [Source:MGI Symbol;Acc:MGI:2684326] | 446.3 | -2.698 | 1.422E-27 |
| ENSMUSG00000040752 | Myh6 | Enriched in Het SCN | myosin, heavy polypeptide 6, cardiac muscle, alpha [Source:MGI Symbol;Acc:MGI:97255] | 71.63 | -1.947 | 4.711E-08 |
| ENSMUSG00000030787 | Lyve1 | Enriched in Het SCN | lymphatic vessel endothelial hyaluronan receptor 1 [Source:MGI Symbol;Acc:MGI:2136348] | 28.36 | -1.86 | 0.008633 |
| ENSMUSG00000030703 | Gdpd3 | Enriched in Het SCN | glycerophosphodiester phosphodiesterase domain containing 3 [Source:MGI Symbol;Acc:MGI:1915866] | 78.27 | -1.7 | 0.02841 |
| ENSMUSG00000027209 | Fam227b | Enriched in Het SCN | family with sequence similarity 227, member B [Source:MGI Symbol;Acc:MGI:1923073] | 58.97 | -1.629 | 0.001283 |
| ENSMUSG00000093985 | Gm10406 | Enriched in Het SCN | predicted gene 10406 [Source:MGI Symbol;Acc:MGI:3711272] | 82.84 | -1.603 | 0.00002278 |
| ENSMUSG00000035296 | Sgcg | Enriched in Het SCN | sarcoglycan, gamma (dystrophin-associated glycoprotein) [Source:MGI Symbol;Acc:MGI:1346524] | 153.4 | -1.593 | 0.000002008 |
| ENSMUSG00000028037 | Ifi44 | Enriched in Het SCN | interferon-induced protein 44 [Source:MGI Symbol;Acc:MGI:2443016] | 59.75 | -1.469 | 0.02841 |
| ENSMUSG00000073643 | Wdfy1 | Enriched in Het SCN | WD repeat and FYVE domain containing 1 [Source:MGI Symbol;Acc:MGI:1916618] | 1911 | -1.453 | 2.883E-07 |
| ENSMUSG00000032532 | Cck | Enriched in Het SCN | cholecystokinin [Source:MGI Symbol;Acc:MGI:88297] | 572.1 | -1.41 | 0.03349 |
| ENSMUSG00000034579 | Pla2g3 | Enriched in Het SCN | phospholipase A2, group III [Source:MGI Symbol;Acc:MGI:2444945] | 978.7 | -1.351 | 0.005655 |
| ENSMUSG00000050377 | Il31ra | Enriched in Het SCN | interleukin 31 receptor A [Source:MGI Symbol;Acc:MGI:2180511] | 3262 | -1.325 | 0.000001884 |
| ENSMUSG00000038457 | Tmem255b | Enriched in Het SCN | transmembrane protein 255B [Source:MGI Symbol;Acc:MGI:2685533] | 200.5 | -1.134 | 0.00001175 |
| ENSMUSG00000114378 | Gm49355 | Enriched in Het SCN | predicted gene, 49355 [Source:MGI Symbol;Acc:MGI:6121560] | 88.31 | -1.117 | 0.0008644 |
| ENSMUSG00000043336 | Filip1l | Enriched in Het SCN | filamin A interacting protein 1-like [Source:MGI Symbol;Acc:MGI:1925999] | 1255 | -1.083 | 0.01586 |
| ENSMUSG00000022748 | Cmss1 | Enriched in Het SCN | cms small ribosomal subunit 1 [Source:MGI Symbol;Acc:MGI:1913747] | 33320 | -1.057 | 0.00143 |
| ENSMUSG00000054966 | Lmntd1 | Enriched in Het SCN | lamin tail domain containing 1 [Source:MGI Symbol;Acc:MGI:1921321] | 226.8 | -1.006 | 0.005507 |
| ENSMUSG00000050919 | Zfp366 | Enriched in WT SCN | zinc finger protein 366 [Source:MGI Symbol;Acc:MGI:2178429] | 141.8 | 1.037 | 0.001283 |
| ENSMUSG00000041479 | Syt15 | Enriched in WT SCN | synaptotagmin XV [Source:MGI Symbol;Acc:MGI:2442166] | 56.07 | 1.095 | 0.003346 |
| ENSMUSG00000073158 | 9030624G23Rik | Enriched in WT SCN | RIKEN cDNA 9030624G23 gene [Source:MGI Symbol;Acc:MGI:1914058] | 82.65 | 1.109 | 0.000007068 |
| ENSMUSG00000047591 | Mafa | Enriched in WT SCN | v-maf musculoaponeurotic fibrosarcoma oncogene family, protein A (avian) [Source:MGI Symbol;Acc:MGI:26733( | 77.89 | 1.354 | 0.03902 |
| ENSMUSG00000021880 | Rnase6 | Enriched in WT SCN | ribonuclease, RNase A family, 6 [Source:MGI Symbol;Acc:MGI:1925666] | 59.48 | 1.485 | 0.04315 |
| ENSMUSG00000095681 | Gm8281 | Enriched in WT SCN | predicted gene, 8281 [Source:MGI Symbol;Acc:MGI:3647811] | 99.93 | 1.524 | 1.449E-11 |
| ENSMUSG00000002324 | Rec8 | Enriched in WT SCN | REC8 meiotic recombination protein [Source:MGI Symbol;Acc:MGI:1929645] | 521.6 | 1.79 | 0.03726 |
| ENSMUSG00000091519 | Skor2 | Enriched in WT SCN | SKI family transcriptional corepressor 2 [Source:MGI Symbol;Acc:MGI:3645984] | 111.3 | 2.021 | 0.001409 |
| ENSMUSG00000112449 | Srp54b | Enriched in WT SCN | signal recognition particle 54B [Source:MGI Symbol;Acc:MGI:3714357] | 36.45 | 2.404 | 0.0191 |
| ENSMUSG00000074252 | Gm10654 | Enriched in WT SCN | predicted gene 10654 [Source:MGI Symbol;Acc:MGI:3643366] | 74.35 | 2.527 | 0.0002376 |
| ENSMUSG00000058626 | Capn11 | Enriched in WT SCN | calpain 11 [Source:MGI Symbol;Acc:MGI:1352490] | 51.73 | 3.151 | 8.132E-12 |

**Contrast numerator: WT_SCN. Contrast denominator: KO_SCN.**

| ensembl_gene_id | external_gene_name | Significance | description | baseMean | logFC | adj.P.Val |
|---|---|---|---|---|---|---|
| ENSMUSG00000105243 | Gm43444 | Enriched in KO SCN | predicted gene 43444 [Source:MGI Symbol;Acc:MGI:5663581] | 104 | -10.02 | 2.472E-18 |
| ENSMUSG00000079012 | Serpina3m | Enriched in KO SCN | serine (or cysteine) peptidase inhibitor, clade A, member 3M [Source:MGI Symbol;Acc:MGI:98378] | 49.99 | -8.32 | 1.463E-11 |
| ENSMUSG00000072476 | Gm9008 | Enriched in KO SCN | predicted pseudogene 9008 [Source:MGI Symbol;Acc:MGI:3644000] | 125.5 | -7.88 | 1.174E-21 |
| ENSMUSG00000061780 | Cfd | Enriched in KO SCN | complement factor D (adipsin) [Source:MGI Symbol;Acc:MGI:87931] | 63.99 | -6.364 | 0.000004397 |
| ENSMUSG00000027513 | Pck1 | Enriched in KO SCN | phosphoenolpyruvate carboxykinase 1, cytosolic [Source:MGI Symbol;Acc:MGI:97501] | 57.72 | -6.01 | 0.02616 |
| ENSMUSG00000022057 | Adamdec1 | Enriched in KO SCN | ADAM-like, decysin 1 [Source:MGI Symbol;Acc:MGI:1917650] | 37.58 | -4.03 | 0.00004709 |
| ENSMUSG00000062515 | Fabp4 | Enriched in KO SCN | fatty acid binding protein 4, adipocyte [Source:MGI Symbol;Acc:MGI:88038] | 191 | -3.751 | 0.03895 |
| ENSMUSG00000072572 | Slc39a2 | Enriched in KO SCN | solute carrier family 39 (zinc transporter), member 2 [Source:MGI Symbol;Acc:MGI:2684326] | 538.3 | -3.529 | 5.118E-53 |
| ENSMUSG00000059430 | Actg2 | Enriched in KO SCN | actin, gamma 2, smooth muscle, enteric [Source:MGI Symbol;Acc:MGI:104589] | 106.7 | -3.377 | 3.684E-07 |
| ENSMUSG00000040752 | Myh6 | Enriched in KO SCN | myosin, heavy polypeptide 6, cardiac muscle, alpha [Source:MGI Symbol;Acc:MGI:97255] | 104.3 | -2.799 | 2.182E-13 |
| ENSMUSG00000030278 | Cidec | Enriched in KO SCN | cell death-inducing DFFA-like effector c [Source:MGI Symbol;Acc:MGI:95585] | 42.51 | -2.786 | 0.001936 |
| ENSMUSG00000018830 | Myh11 | Enriched in KO SCN | myosin, heavy polypeptide 11, smooth muscle [Source:MGI Symbol;Acc:MGI:102643] | 601.6 | -2.785 | 0.0004079 |
| ENSMUSG00000001349 | Cnn1 | Enriched in KO SCN | calponin 1 [Source:MGI Symbol;Acc:MGI:104979] | 97.66 | -2.718 | 0.001207 |
| ENSMUSG00000027656 | Ccn5 | Enriched in KO SCN | cellular communication network factor 5 [Source:MGI Symbol;Acc:MGI:1328326] | 42.22 | -2.69 | 0.008836 |
| ENSMUSG00000035783 | Acta2 | Enriched in KO SCN | actin, alpha 2, smooth muscle, aorta [Source:MGI Symbol;Acc:MGI:87909] | 1711 | -2.687 | 0.005279 |
| ENSMUSG00000032517 | Mobp | Enriched in KO SCN | myelin-associated oligodendrocytic basic protein [Source:MGI Symbol;Acc:MGI:108511] | 3016 | -2.598 | 0.00003269 |
| ENSMUSG00000076439 | Mog | Enriched in KO SCN | myelin oligodendrocyte glycoprotein [Source:MGI Symbol;Acc:MGI:97435] | 158.7 | -2.565 | 3.297E-09 |
| ENSMUSG00000032085 | Tagln | Enriched in KO SCN | transgelin [Source:MGI Symbol;Acc:MGI:106012] | 761.8 | -2.327 | 0.001413 |
| ENSMUSG00000029675 | Eln | Enriched in KO SCN | elastin [Source:MGI Symbol;Acc:MGI:95317] | 18970 | -2.322 | 0.001558 |
| ENSMUSG00000019326 | Aoc3 | Enriched in KO SCN | amine oxidase, copper containing 3 [Source:MGI Symbol;Acc:MGI:1306797] | 175.2 | -2.314 | 0.01959 |
| ENSMUSG00000027559 | Car3 | Enriched in KO SCN | carbonic anhydrase 3 [Source:MGI Symbol;Acc:MGI:88270] | 195.5 | -2.297 | 4.416E-08 |
| ENSMUSG00000025610 | Map3k7cl | Enriched in KO SCN | Map3k7 C-terminal like [Source:MGI Symbol;Acc:MGI:2446584] | 120.5 | -2.241 | 0.003825 |
| ENSMUSG00000021186 | Fbln5 | Enriched in KO SCN | fibulin 5 [Source:MGI Symbol;Acc:MGI:1346091] | 1586 | -2.144 | 0.002835 |
| ENSMUSG00000035296 | Sgcg | Enriched in KO SCN | sarcoglycan, gamma (dystrophin-associated glycoprotein) [Source:MGI Symbol;Acc:MGI:1346524] | 188 | -2.119 | 1.308E-12 |
| ENSMUSG00000035896 | Rnase1 | Enriched in KO SCN | ribonuclease, RNase A family, 1 (pancreatic) [Source:MGI Symbol;Acc:MGI:97919] | 59.34 | -2.034 | 0.0003202 |
| ENSMUSG00000020411 | Nipal4 | Enriched in KO SCN | NIPA-like domain containing 4 [Source:MGI Symbol;Acc:MGI:2444671] | 64.94 | -1.952 | 0.002987 |
| ENSMUSG00000030048 | Gkn3 | Enriched in KO SCN | gastrokine 3 [Source:MGI Symbol;Acc:MGI:1916138] | 110.9 | -1.927 | 0.000003817 |
| ENSMUSG00000038457 | Tmem255b | Enriched in KO SCN | transmembrane protein 255B [Source:MGI Symbol;Acc:MGI:2685533] | 320.6 | -1.918 | 1.209E-20 |
| ENSMUSG00000057123 | Gja5 | Enriched in KO SCN | gap junction protein, alpha 5 [Source:MGI Symbol;Acc:MGI:95716] | 378.1 | -1.912 | 0.000001841 |
| ENSMUSG00000040483 | Xaf1 | Enriched in KO SCN | XIAP associated factor 1 [Source:MGI Symbol;Acc:MGI:3772572] | 139.9 | -1.866 | 1.967E-07 |
| ENSMUSG00000028776 | Tinagl1 | Enriched in KO SCN | tubulointerstitial nephritis antigen-like 1 [Source:MGI Symbol;Acc:MGI:2137617] | 347.2 | -1.838 | 0.00002255 |
| ENSMUSG00000050854 | Tmem125 | Enriched in KO SCN | transmembrane protein 125 [Source:MGI Symbol;Acc:MGI:1923409] | 63.86 | -1.793 | 0.01066 |
| ENSMUSG00000030703 | Gdpd3 | Enriched in KO SCN | glycerophosphodiester phosphodiesterase domain containing 3 [Source:MGI Symbol;Acc:MGI:1915866] | 76.86 | -1.779 | 0.0008913 |
| ENSMUSG00000034579 | Pla2g3 | Enriched in KO SCN | phospholipase A2, group III [Source:MGI Symbol;Acc:MGI:2444945] | 1696 | -1.756 | 0.002109 |
| ENSMUSG00000037625 | Cldn11 | Enriched in KO SCN | claudin 11 [Source:MGI Symbol;Acc:MGI:106925] | 1805 | -1.727 | 0.0007629 |
| ENSMUSG00000073643 | Wdfy1 | Enriched in KO SCN | WD repeat and FYVE domain containing 1 [Source:MGI Symbol;Acc:MGI:1916618] | 2291 | -1.709 | 1.563E-33 |
| ENSMUSG00000022074 | Tnfrsf10b | Enriched in KO SCN | tumor necrosis factor receptor superfamily, member 10b [Source:MGI Symbol;Acc:MGI:1341090] | 61.6 | -1.691 | 0.00008403 |
| ENSMUSG00000041607 | Mbp | Enriched in KO SCN | myelin basic protein [Source:MGI Symbol;Acc:MGI:96925] | 30260 | -1.685 | 0.00007435 |
| ENSMUSG00000043336 | Filip1l | Enriched in KO SCN | filamin A interacting protein 1-like [Source:MGI Symbol;Acc:MGI:1925999] | 968.3 | -1.672 | 8.171E-09 |
| ENSMUSG00000037166 | Ppp1r14a | Enriched in KO SCN | protein phosphatase 1, regulatory inhibitor subunit 14A [Source:MGI Symbol;Acc:MGI:1931139] | 58.89 | -1.633 | 0.0009841 |
| ENSMUSG00000050377 | Il31ra | Enriched in KO SCN | interleukin 31 receptor A [Source:MGI Symbol;Acc:MGI:2180511] | 2301 | -1.627 | 1.036E-14 |
| ENSMUSG00000073680 | Tmem88b | Enriched in KO SCN | transmembrane protein 88B [Source:MGI Symbol;Acc:MGI:2444329] | 513 | -1.602 | 2.838E-10 |
| ENSMUSG00000093985 | Gm10406 | Enriched in KO SCN | predicted gene 10406 [Source:MGI Symbol;Acc:MGI:3711272] | 74.9 | -1.602 | 0.001271 |
| ENSMUSG00000023019 | Gpd1 | Enriched in KO SCN | glycerol-3-phosphate dehydrogenase 1 (soluble) [Source:MGI Symbol;Acc:MGI:95679] | 238.4 | -1.57 | 4.515E-08 |
| ENSMUSG00000114378 | Gm49355 | Enriched in KO SCN | predicted gene, 49355 [Source:MGI Symbol;Acc:MGI:6121560] | 105.4 | -1.566 | 0.000003104 |
| ENSMUSG00000036634 | Mag | Enriched in KO SCN | myelin-associated glycoprotein [Source:MGI Symbol;Acc:MGI:96912] | 2003 | -1.542 | 0.001917 |
| ENSMUSG00000054966 | Lmntd1 | Enriched in KO SCN | lamin tail domain containing 1 [Source:MGI Symbol;Acc:MGI:1921321] | 245 | -1.542 | 9.82E-14 |
| ENSMUSG00000025219 | Fgf8 | Enriched in KO SCN | fibroblast growth factor 8 [Source:MGI Symbol;Acc:MGI:99604] | 54.86 | -1.528 | 0.04116 |
| ENSMUSG00000043333 | Rhbdl2 | Enriched in KO SCN | rhomboid like 2 [Source:MGI Symbol;Acc:MGI:3608413] | 49.82 | -1.527 | 0.004983 |
| ENSMUSG00000050423 | Ppp1r3g | Enriched in KO SCN | protein phosphatase 1, regulatory subunit 3G [Source:MGI Symbol;Acc:MGI:1923737] | 69.13 | -1.519 | 0.01259 |
| ENSMUSG00000026697 | Myoc | Enriched in KO SCN | myocilin [Source:MGI Symbol;Acc:MGI:1202864] | 427.5 | -1.472 | 0.00001872 |
| ENSMUSG00000068196 | Col8a1 | Enriched in KO SCN | collagen, type VIII, alpha 1 [Source:MGI Symbol;Acc:MGI:88463] | 665.2 | -1.472 | 1.954E-11 |
| ENSMUSG00000034177 | Rnf43 | Enriched in KO SCN | ring finger protein 43 [Source:MGI Symbol;Acc:MGI:2442609] | 60.6 | -1.442 | 0.003796 |
| ENSMUSG00000004328 | Hif3a | Enriched in KO SCN | hypoxia inducible factor 3, alpha subunit [Source:MGI Symbol;Acc:MGI:1859778] | 231.5 | -1.412 | 0.000005637 |
| ENSMUSG00000031425 | Plp1 | Enriched in KO SCN | proteolipid protein (myelin) 1 [Source:MGI Symbol;Acc:MGI:97623] | 16320 | -1.376 | 0.002649 |
| ENSMUSG00000031377 | Bmx | Enriched in KO SCN | BMX non-receptor tyrosine kinase [Source:MGI Symbol;Acc:MGI:1101778] | 118.5 | -1.373 | 0.007712 |
| ENSMUSG00000018566 | Slc2a4 | Enriched in KO SCN | solute carrier family 2 (facilitated glucose transporter), member 4 [Source:MGI Symbol;Acc:MGI:95758] | 131.9 | -1.357 | 0.0003041 |
| ENSMUSG00000027375 | Mal | Enriched in KO SCN | myelin and lymphocyte protein, T cell differentiation protein [Source:MGI Symbol;Acc:MGI:892970] | 485.9 | -1.351 | 1.409E-07 |
| ENSMUSG00000026208 | Des | Enriched in KO SCN | desmin [Source:MGI Symbol;Acc:MGI:94885] | 113 | -1.35 | 0.01358 |
| ENSMUSG00000045903 | Npas4 | Enriched in KO SCN | neuronal PAS domain protein 4 [Source:MGI Symbol;Acc:MGI:2664186] | 439.5 | -1.323 | 9.491E-10 |
| ENSMUSG00000006403 | Adamts4 | Enriched in KO SCN | a disintegrin-like and metallopeptidase (reprolysin type) with thrombospondin type 1 motif, 4 [Source:MGI Symbol;Acc:MGI:133994 | 532.4 | -1.299 | 0.0002997 |
| ENSMUSG00000024049 | Myom1 | Enriched in KO SCN | myomesin 1 [Source:MGI Symbol;Acc:MGI:1341430] | 126.1 | -1.298 | 0.001245 |
| ENSMUSG00000026830 | Ermn | Enriched in KO SCN | ermin, ERM-like protein [Source:MGI Symbol;Acc:MGI:1925017] | 82.5 | -1.295 | 0.003292 |
| ENSMUSG00000073406 | H2-Bl | Enriched in KO SCN | histocompatibility 2, blastocyst [Source:MGI Symbol;Acc:MGI:892004] | 179.1 | -1.221 | 0.01873 |
| ENSMUSG00000038173 | Enpp6 | Enriched in KO SCN | ectonucleotide pyrophosphatase/phosphodiesterase 6 [Source:MGI Symbol;Acc:MGI:2445171] | 1524 | -1.219 | 0.00001872 |
| ENSMUSG00000032420 | Nt5e | Enriched in KO SCN | 5' nucleotidase, ecto [Source:MGI Symbol;Acc:MGI:99782] | 150 | -1.206 | 0.01824 |
| ENSMUSG00000015467 | Egfl8 | Enriched in KO SCN | EGF-like domain 8 [Source:MGI Symbol;Acc:MGI:1932094] | 206.7 | -1.203 | 0.001901 |
| ENSMUSG00000021091 | Serpina3n | Enriched in KO SCN | serine (or cysteine) peptidase inhibitor, clade A, member 3N [Source:MGI Symbol;Acc:MGI:105045] | 2536 | -1.177 | 2.035E-11 |
| ENSMUSG00000033579 | Fa2h | Enriched in KO SCN | fatty acid 2-hydroxylase [Source:MGI Symbol;Acc:MGI:2443327] | 576.3 | -1.15 | 0.0004079 |
| ENSMUSG00000026475 | Rgs16 | Enriched in KO SCN | regulator of G-protein signaling 16 [Source:MGI Symbol;Acc:MGI:108407] | 3923 | -1.149 | 0.0411 |
| ENSMUSG00000021453 | Gadd45g | Enriched in KO SCN | growth arrest and DNA-damage-inducible 45 gamma [Source:MGI Symbol;Acc:MGI:1346325] | 1188 | -1.143 | 0.00001301 |
| ENSMUSG00000034000 | Neu4 | Enriched in KO SCN | sialidase 4 [Source:MGI Symbol;Acc:MGI:2661364] | 530.5 | -1.13 | 1.09E-10 |
| ENSMUSG00000036098 | Myrf | Enriched in KO SCN | myelin regulatory factor [Source:MGI Symbol;Acc:MGI:2684944] | 1079 | -1.128 | 0.000008361 |
| ENSMUSG00000007594 | Hapln4 | Enriched in KO SCN | hyaluronan and proteoglycan link protein 4 [Source:MGI Symbol;Acc:MGI:2679531] | 659.3 | -1.123 | 0.002625 |
| ENSMUSG00000067818 | Myl9 | Enriched in KO SCN | myosin, light polypeptide 9, regulatory [Source:MGI Symbol;Acc:MGI:2138915] | 1338 | -1.11 | 0.02861 |
| ENSMUSG00000022748 | Cmss1 | Enriched in KO SCN | cms small ribosomal subunit 1 [Source:MGI Symbol;Acc:MGI:1913747] | 23820 | -1.097 | 1.513E-08 |
| ENSMUSG00000006154 | Eps8l1 | Enriched in KO SCN | EPS8-like 1 [Source:MGI Symbol;Acc:MGI:1914675] | 385.1 | -1.067 | 0.004422 |
| ENSMUSG00000032854 | Ugt8a | Enriched in KO SCN | UDP galactosyltransferase 8A [Source:MGI Symbol;Acc:MGI:109522] | 1497 | -1.057 | 0.00005151 |
| ENSMUSG00000074170 | Plekhf1 | Enriched in KO SCN | pleckstrin homology domain containing, family F (with FYVE domain) member 1 [Source:MGI Symbol;Acc:MGI:1919537] | 112.8 | -1.054 | 0.01876 |
| ENSMUSG00000035274 | Tpbg | Enriched in KO SCN | trophoblast glycoprotein [Source:MGI Symbol;Acc:MGI:1341264] | 1115 | -1.047 | 3.492E-12 |
| ENSMUSG00000031636 | Pdlim3 | Enriched in KO SCN | PDZ and LIM domain 3 [Source:MGI Symbol;Acc:MGI:1859274] | 190 | -1.021 | 0.001178 |
| ENSMUSG00000048540 | Nhlh2 | Enriched in WT SCN | nescient helix loop helix 2 [Source:MGI Symbol;Acc:MGI:97324] | 351.1 | 1.002 | 0.00394 |
| ENSMUSG00000037206 | Islr | Enriched in WT SCN | immunoglobulin superfamily containing leucine-rich repeat [Source:MGI Symbol;Acc:MGI:1349645] | 378.2 | 1.017 | 0.003423 |
| ENSMUSG00000040258 | Nxph4 | Enriched in WT SCN | neurexophilin 4 [Source:MGI Symbol;Acc:MGI:1336197] | 276.3 | 1.02 | 0.03286 |
| ENSMUSG00000022325 | Pop1 | Enriched in WT SCN | processing of precursor 1, ribonuclease P/MRP family, (S. cerevisiae) [Source:MGI Symbol;Acc:MGI:1914974] | 260.1 | 1.043 | 0.000006939 |
| ENSMUSG00000045996 | Polr2k | Enriched in WT SCN | polymerase (RNA) II (DNA directed) polypeptide K [Source:MGI Symbol;Acc:MGI:102725] | 401.4 | 1.057 | 0.00006091 |
| ENSMUSG00000056380 | Gpr50 | Enriched in WT SCN | G-protein-coupled receptor 50 [Source:MGI Symbol;Acc:MGI:1333877] | 533.3 | 1.058 | 0.004093 |
| ENSMUSG00000033792 | Atp7a | Enriched in WT SCN | ATPase, Cu++ transporting, alpha polypeptide [Source:MGI Symbol;Acc:MGI:99400] | 225.9 | 1.061 | 0.005924 |
| ENSMUSG00000047259 | Mc4r | Enriched in WT SCN | melanocortin 4 receptor [Source:MGI Symbol;Acc:MGI:99457] | 116.6 | 1.076 | 0.0423 |
| ENSMUSG00000091345 | Col6a5 | Enriched in WT SCN | collagen, type VI, alpha 5 [Source:MGI Symbol;Acc:MGI:3648134] | 290.4 | 1.08 | 0.00002641 |
| ENSMUSG00000027004 | Frzb | Enriched in WT SCN | frizzled-related protein [Source:MGI Symbol;Acc:MGI:892032] | 185.8 | 1.083 | 0.04116 |
| ENSMUSG00000038115 | Ano2 | Enriched in WT SCN | anoctamin 2 [Source:MGI Symbol;Acc:MGI:2387214] | 58.92 | 1.102 | 0.04205 |
| ENSMUSG00000026774 | Potegl | Enriched in WT SCN | POTE ankyrin domain family, member G like [Source:MGI Symbol;Acc:MGI:1918231] | 80.66 | 1.114 | 0.01077 |
| ENSMUSG00000018927 | Ccl6 | Enriched in WT SCN | chemokine (C-C motif) ligand 6 [Source:MGI Symbol;Acc:MGI:98263] | 54.45 | 1.116 | 0.03801 |
| ENSMUSG00000035279 | Ssc5d | Enriched in WT SCN | scavenger receptor cysteine rich family, 5 domains [Source:MGI Symbol;Acc:MGI:3606211] | 136.1 | 1.118 | 0.001508 |
| ENSMUSG00000057777 | Mab21l2 | Enriched in WT SCN | mab-21-like 2 [Source:MGI Symbol;Acc:MGI:1346022] | 158.3 | 1.123 | 0.008415 |
| ENSMUSG00000063681 | Crb1 | Enriched in WT SCN | crumbs family member 1, photoreceptor morphogenesis associated [Source:MGI Symbol;Acc:MGI:2136343] | 714.3 | 1.127 | 4.305E-12 |
| ENSMUSG00000024907 | Gal | Enriched in WT SCN | galanin and GMAP prepropeptide [Source:MGI Symbol;Acc:MGI:95637] | 832.6 | 1.152 | 0.0009358 |
| ENSMUSG00000026826 | Nr4a2 | Enriched in WT SCN | nuclear receptor subfamily 4, group A, member 2 [Source:MGI Symbol;Acc:MGI:1352456] | 257.3 | 1.159 | 0.00001089 |
| ENSMUSG00000108049 | Gm44168 | Enriched in WT SCN | predicted gene, 44168 [Source:MGI Symbol;Acc:MGI:5690560] | 79.91 | 1.165 | 0.006222 |
| ENSMUSG00000042436 | Mfap4 | Enriched in WT SCN | microfibrillar-associated protein 4 [Source:MGI Symbol;Acc:MGI:1342276] | 207.8 | 1.184 | 0.04426 |
| ENSMUSG00000034652 | Cd300a | Enriched in WT SCN | CD300A molecule [Source:MGI Symbol;Acc:MGI:2443411] | 138.3 | 1.192 | 0.001365 |
| ENSMUSG00000079941 | Cox5b-ps | Enriched in WT SCN | cytochrome c oxidase subunit 5B, pseudogene [Source:MGI Symbol;Acc:MGI:3649411] | 491.4 | 1.198 | 6.263E-09 |
| ENSMUSG00000031112 | Stk26 | Enriched in WT SCN | serine/threonine kinase 26 [Source:MGI Symbol;Acc:MGI:1917665] | 103 | 1.211 | 0.01165 |
| ENSMUSG00000024043 | Arhgap28 | Enriched in WT SCN | Rho GTPase activating protein 28 [Source:MGI Symbol;Acc:MGI:2147003] | 92.58 | 1.241 | 0.009889 |
| ENSMUSG00000032332 | Col12a1 | Enriched in WT SCN | collagen, type XII, alpha 1 [Source:MGI Symbol;Acc:MGI:88448] | 598.9 | 1.241 | 9.395E-14 |
| ENSMUSG00000015812 | Gnrh1 | Enriched in WT SCN | gonadotropin releasing hormone 1 [Source:MGI Symbol;Acc:MGI:95789] | 121.2 | 1.258 | 0.02971 |
| ENSMUSG00000052504 | Epha3 | Enriched in WT SCN | Eph receptor A3 [Source:MGI Symbol;Acc:MGI:99612] | 134.6 | 1.289 | 0.0007833 |
| ENSMUSG00000021996 | Esd | Enriched in WT SCN | esterase D/formylglutathione hydrolase [Source:MGI Symbol;Acc:MGI:95421] | 1798 | 1.298 | 1.472E-34 |
| ENSMUSG00000026051 | Ecrg4 | Enriched in WT SCN | ECRG4 augurin precursor [Source:MGI Symbol;Acc:MGI:1926146] | 152.6 | 1.298 | 0.002467 |
| ENSMUSG00000031303 | Map3k15 | Enriched in WT SCN | mitogen-activated protein kinase kinase kinase 15 [Source:MGI Symbol;Acc:MGI:2448588] | 188.4 | 1.31 | 0.00001374 |
| ENSMUSG00000029595 | Lhx5 | Enriched in WT SCN | LIM homeobox protein 5 [Source:MGI Symbol;Acc:MGI:107792] | 498.8 | 1.316 | 0.000007681 |
| ENSMUSG00000053219 | Raet1e | Enriched in WT SCN | retinoic acid early transcript 1E [Source:MGI Symbol;Acc:MGI:2675273] | 393.7 | 1.326 | 1.228E-12 |
| ENSMUSG00000021798 | Ldb3 | Enriched in WT SCN | LIM domain binding 3 [Source:MGI Symbol;Acc:MGI:1344412] | 158.8 | 1.332 | 0.00006127 |
| ENSMUSG00000060459 | Kng2 | Enriched in WT SCN | kininogen 2 [Source:MGI Symbol;Acc:MGI:3027157] | 115.9 | 1.366 | 0.00001872 |
| ENSMUSG00000046861 | Hectd3 | Enriched in WT SCN | HECT domain E3 ubiquitin protein ligase 3 [Source:MGI Symbol;Acc:MGI:1923858] | 1208 | 1.382 | 0.000003907 |
| ENSMUSG00000020268 | Lyrm7 | Enriched in WT SCN | LYR motif containing 7 [Source:MGI Symbol;Acc:MGI:1922780] | 247.7 | 1.384 | 1.685E-12 |
| ENSMUSG00000075707 | Dio3 | Enriched in WT SCN | deiodinase, iodothyronine type III [Source:MGI Symbol;Acc:MGI:1306782] | 156.3 | 1.386 | 0.0001171 |
| ENSMUSG00000072494 | Ppp1r3e | Enriched in WT SCN | protein phosphatase 1, regulatory subunit 3E [Source:MGI Symbol;Acc:MGI:2145790] | 1821 | 1.41 | 5.214E-37 |
| ENSMUSG00000036760 | Kcnk9 | Enriched in WT SCN | potassium channel, subfamily K, member 9 [Source:MGI Symbol;Acc:MGI:3521816] | 3435 | 1.419 | 0.001277 |
| ENSMUSG00000019768 | Esr1 | Enriched in WT SCN | estrogen receptor 1 (alpha) [Source:MGI Symbol;Acc:MGI:1352467] | 656.1 | 1.424 | 0.02002 |
| ENSMUSG00000032572 | Col6a4 | Enriched in WT SCN | collagen, type VI, alpha 4 [Source:MGI Symbol;Acc:MGI:1915803] | 73.76 | 1.474 | 0.001656 |
| ENSMUSG00000069874 | Irgm2 | Enriched in WT SCN | immunity-related GTPase family M member 2 [Source:MGI Symbol;Acc:MGI:1926262] | 109.3 | 1.5 | 0.000003631 |
| ENSMUSG00000004347 | Pde1c | Enriched in WT SCN | phosphodiesterase 1C [Source:MGI Symbol;Acc:MGI:108413] | 643 | 1.557 | 6.372E-09 |
| ENSMUSG00000001506 | Col1a1 | Enriched in WT SCN | collagen, type I, alpha 1 [Source:MGI Symbol;Acc:MGI:88467] | 1557 | 1.572 | 1.454E-10 |

| | | | | | | |
|---|---|---|---|---|---|---|
| ENSMUSG00000030270 | Cpne9 | Enriched in WT SCN | copine family member IX [Source:MGI Symbol;Acc:MGI:2443052] | 73.85 | 1.63 | 0.0004531 |
| ENSMUSG00000019230 | Lhx9 | Enriched in WT SCN | LIM homeobox protein 9 [Source:MGI Symbol;Acc:MGI:1316721] | 221.8 | 1.638 | 0.000008953 |
| ENSMUSG00000048562 | Sp8 | Enriched in WT SCN | trans-acting transcription factor 8 [Source:MGI Symbol;Acc:MGI:2443471] | 190.6 | 1.638 | 0.0004765 |
| ENSMUSG00000041479 | Syt15 | Enriched in WT SCN | synaptotagmin XV [Source:MGI Symbol;Acc:MGI:2442166] | 109.1 | 1.643 | 0.02911 |
| ENSMUSG00000046167 | Gldn | Enriched in WT SCN | gliomedin [Source:MGI Symbol;Acc:MGI:2388361] | 120.6 | 1.643 | 0.005747 |
| ENSMUSG00000096039 | D830030K20Rik | Enriched in WT SCN | RIKEN cDNA D830030K20 gene [Source:MGI Symbol;Acc:MGI:2443830] | 55.57 | 1.691 | 0.0003272 |
| ENSMUSG00000058626 | Capn11 | Enriched in WT SCN | calpain 11 [Source:MGI Symbol;Acc:MGI:1352490] | 104.7 | 1.709 | 0.01667 |
| ENSMUSG00000073733 | Cplane2 | Enriched in WT SCN | ciliogenesis and planar polarity effector 2 [Source:MGI Symbol;Acc:MGI:1923416] | 116.8 | 1.717 | 3.56E-09 |
| ENSMUSG00000091754 | Gm3636 | Enriched in WT SCN | predicted gene 3636 [Source:MGI Symbol;Acc:MGI:3781812] | 55.75 | 1.756 | 0.00003131 |
| ENSMUSG00000044229 | Nxpe4 | Enriched in WT SCN | neurexophilin and PC-esterase domain family, member 4 [Source:MGI Symbol;Acc:MGI:1924792] | 908 | 1.777 | 2.421E-49 |
| ENSMUSG00000100486 | Gm4131 | Enriched in WT SCN | predicted gene 4131 [Source:MGI Symbol;Acc:MGI:3782307] | 623.8 | 1.821 | 9.255E-28 |
| ENSMUSG00000062713 | Sim2 | Enriched in WT SCN | single-minded family bHLH transcription factor 2 [Source:MGI Symbol;Acc:MGI:98307] | 63.54 | 1.866 | 0.003387 |
| ENSMUSG00000056055 | Sag | Enriched in WT SCN | S-antigen, retina and pineal gland (arrestin) [Source:MGI Symbol;Acc:MGI:98227] | 119.7 | 1.871 | 0.01336 |
| ENSMUSG00000042498 | Radx | Enriched in WT SCN | RPA1 related single stranded DNA binding protein, X-linked [Source:MGI Symbol;Acc:MGI:2147848] | 216.8 | 1.892 | 1.135E-09 |
| ENSMUSG00000096225 | Lhx8 | Enriched in WT SCN | LIM homeobox protein 8 [Source:MGI Symbol;Acc:MGI:1096343] | 138.6 | 1.956 | 0.0006418 |
| ENSMUSG00000047591 | Mafa | Enriched in WT SCN | v-maf musculoaponeurotic fibrosarcoma oncogene family, protein A (avian) [Source:MGI Symbol;Acc:MGI:2673307] | 82.09 | 1.977 | 0.003292 |
| ENSMUSG00000020415 | Pttg1 | Enriched in WT SCN | pituitary tumor-transforming gene 1 [Source:MGI Symbol;Acc:MGI:1353578] | 453.6 | 2.001 | 2.77E-23 |
| ENSMUSG00000032105 | Pdzd3 | Enriched in WT SCN | PDZ domain containing 3 [Source:MGI Symbol;Acc:MGI:2429554] | 72.53 | 2.032 | 0.00002844 |
| ENSMUSG00000067220 | Cnga1 | Enriched in WT SCN | cyclic nucleotide gated channel alpha 1 [Source:MGI Symbol;Acc:MGI:88436] | 69.21 | 2.077 | 6.307E-07 |
| ENSMUSG00000033227 | Wnt6 | Enriched in WT SCN | wingless-type MMTV integration site family, member 6 [Source:MGI Symbol;Acc:MGI:98960] | 71.41 | 2.09 | 0.02401 |
| ENSMUSG00000095681 | Gm8281 | Enriched in WT SCN | predicted gene, 8281 [Source:MGI Symbol;Acc:MGI:3647811] | 129.9 | 2.14 | 0.00000785 |
| ENSMUSG00000052234 | Epx | Enriched in WT SCN | eosinophil peroxidase [Source:MGI Symbol;Acc:MGI:107569] | 82.23 | 2.162 | 0.01592 |
| ENSMUSG00000044177 | Wfikkn2 | Enriched in WT SCN | WAP, follistatin/kazal, immunoglobulin, kunitz and netrin domain containing 2 [Source:MGI Symbol;Acc:MGI:2669209] | 101.9 | 2.259 | 0.00001862 |
| ENSMUSG00000030324 | Rho | Enriched in WT SCN | rhodopsin [Source:MGI Symbol;Acc:MGI:97914] | 122.3 | 2.424 | 0.0004531 |
| ENSMUSG00000037446 | Tulp1 | Enriched in WT SCN | tubby like protein 1 [Source:MGI Symbol;Acc:MGI:109571] | 69.21 | 2.439 | 0.001365 |
| ENSMUSG00000059343 | Aldoart1 | Enriched in WT SCN | aldolase 1 A, retrogene 1 [Source:MGI Symbol;Acc:MGI:2447811] | 59.22 | 2.59 | 8.599E-09 |
| ENSMUSG00000036412 | Arsi | Enriched in WT SCN | arylsulfatase i [Source:MGI Symbol;Acc:MGI:2670959] | 69.64 | 2.6 | 0.00002255 |
| ENSMUSG00000021950 | Anxa8 | Enriched in WT SCN | annexin A8 [Source:MGI Symbol;Acc:MGI:1201374] | 88.49 | 2.701 | 1.2E-10 |
| ENSMUSG00000032346 | Ooep | Enriched in WT SCN | oocyte expressed protein [Source:MGI Symbol;Acc:MGI:1915218] | 91.4 | 2.784 | 2.098E-10 |
| ENSMUSG00000040148 | Hmx3 | Enriched in WT SCN | H6 homeobox 3 [Source:MGI Symbol;Acc:MGI:107160] | 232.5 | 2.823 | 9.338E-10 |
| ENSMUSG00000019890 | Nts | Enriched in WT SCN | neurotensin [Source:MGI Symbol;Acc:MGI:1328351] | 1769 | 2.83 | 0.00001486 |
| ENSMUSG00000050100 | Hmx2 | Enriched in WT SCN | H6 homeobox 2 [Source:MGI Symbol;Acc:MGI:107159] | 203.6 | 2.83 | 2.838E-10 |
| ENSMUSG00000035033 | Tbr1 | Enriched in WT SCN | T-box brain transcription factor 1 [Source:MGI Symbol;Acc:MGI:107404] | 66.65 | 2.838 | 0.0001096 |
| ENSMUSG00000049796 | Crh | Enriched in WT SCN | corticotropin releasing hormone [Source:MGI Symbol;Acc:MGI:88496] | 120.9 | 3.016 | 1.6E-12 |
| ENSMUSG00000020950 | Foxg1 | Enriched in WT SCN | forkhead box G1 [Source:MGI Symbol;Acc:MGI:1347464] | 214.1 | 3.212 | 9.704E-12 |
| ENSMUSG00000024650 | Slc22a6 | Enriched in WT SCN | solute carrier family 22 (organic anion transporter), member 6 [Source:MGI Symbol;Acc:MGI:892001] | 135.3 | 3.643 | 0.04091 |
| ENSMUSG00000015090 | Ptgds | Enriched in WT SCN | prostaglandin D2 synthase (brain) [Source:MGI Symbol;Acc:MGI:99261] | 13560 | 3.991 | 0.01351 |
| ENSMUSG00000026890 | Lhx6 | Enriched in WT SCN | LIM homeobox protein 6 [Source:MGI Symbol;Acc:MGI:1306803] | 305.8 | 4.076 | 2.32E-21 |
| ENSMUSG00000023978 | Prph2 | Enriched in WT SCN | peripherin 2 [Source:MGI Symbol;Acc:MGI:102791] | 71.04 | 4.561 | 0.01705 |
| ENSMUSG00000091519 | Skor2 | Enriched in WT SCN | SKI family transcriptional corepressor 2 [Source:MGI Symbol;Acc:MGI:3645984] | 103.9 | 4.857 | 3.629E-15 |
| ENSMUSG00000032292 | Nr2e3 | Enriched in WT SCN | nuclear receptor subfamily 2, group E, member 3 [Source:MGI Symbol;Acc:MGI:1346317] | 77.5 | 4.88 | 0.004093 |
| ENSMUSG00000060461 | Dppa5a | Enriched in WT SCN | developmental pluripotency associated 5A [Source:MGI Symbol;Acc:MGI:101800] | 40.14 | 4.905 | 6.677E-08 |
| ENSMUSG00000006007 | Pdc | Enriched in WT SCN | phosducin [Source:MGI Symbol;Acc:MGI:98090] | 75.02 | 4.928 | 0.003387 |
| ENSMUSG00000021848 | Otx2 | Enriched in WT SCN | orthodenticle homeobox 2 [Source:MGI Symbol;Acc:MGI:97451] | 46.83 | 4.992 | 1.466E-10 |
| ENSMUSG00000032446 | Eomes | Enriched in WT SCN | eomesodermin [Source:MGI Symbol;Acc:MGI:1201683] | 248.4 | 5.409 | 3.545E-12 |
| ENSMUSG00000061808 | Ttr | Enriched in WT SCN | transthyretin [Source:MGI Symbol;Acc:MGI:98865] | 373.7 | 5.448 | 0.00003419 |
| ENSMUSG00000029491 | Pde6b | Enriched in WT SCN | phosphodiesterase 6B, cGMP, rod receptor, beta polypeptide [Source:MGI Symbol;Acc:MGI:97525] | 45.6 | 5.506 | 9.482E-07 |
| ENSMUSG00000041578 | Crx | Enriched in WT SCN | cone-rod homeobox [Source:MGI Symbol;Acc:MGI:1194883] | 60.51 | 6.364 | 2.247E-08 |
| ENSMUSG00000024041 | Cryaa | Enriched in WT SCN | crystallin, alpha A [Source:MGI Symbol;Acc:MGI:88515] | 117.9 | 6.54 | 0.000598 |
| ENSMUSG00000021880 | Rnase6 | Enriched in WT SCN | ribonuclease, RNase A family, 6 [Source:MGI Symbol;Acc:MGI:1925666] | 60.46 | 6.827 | 1.208E-11 |
| ENSMUSG00000000724 | Cryba1 | Enriched in WT SCN | crystallin, beta A1 [Source:MGI Symbol;Acc:MGI:88518] | 52.79 | 7.435 | 0.002438 |

**Contrast numerator: WT_retina. Contrast denominator: Het_retina.**

| ensembl_gene_id | external_gene_name | Significance | description | baseMean | logFC | adj.P.Val |
|---|---|---|---|---|---|---|
| ENSMUSG00000111176 | Gm47233 | Enriched in Het retina | predicted gene, 47233 [Source:MGI Symbol;Acc:MGI:6096050] | 84.91 | -8.904 | 0.000038 |
| ENSMUSG00000028635 | Edn2 | Enriched in Het retina | endothelin 2 [Source:MGI Symbol;Acc:MGI:95284] | 60.04 | -6.789 | 0.03417 |
| ENSMUSG00000093805 | Gal3st2b | Enriched in Het retina | galactose-3-O-sulfotransferase 2B [Source:MGI Symbol;Acc:MGI:3711964] | 42.46 | -6.757 | 4.868E-09 |
| ENSMUSG00000035296 | Sgcg | Enriched in Het retina | sarcoglycan, gamma (dystrophin-associated glycoprotein) [Source:MGI Symbol;Acc:MGI:1346524] | 147.1 | -5.554 | 1.537E-35 |
| ENSMUSG00000072572 | Slc39a2 | Enriched in Het retina | solute carrier family 39 (zinc transporter), member 2 [Source:MGI Symbol;Acc:MGI:2684326] | 537.7 | -4.492 | 6.787E-74 |
| ENSMUSG00000041782 | Lad1 | Enriched in Het retina | ladinin [Source:MGI Symbol;Acc:MGI:109343] | 80.64 | -3.473 | 0.01992 |
| ENSMUSG00000037225 | Fgf2 | Enriched in Het retina | fibroblast growth factor 2 [Source:MGI Symbol;Acc:MGI:95516] | 818.4 | -2.648 | 0.02915 |
| ENSMUSG00000090799 | Klhl33 | Enriched in Het retina | kelch-like 33 [Source:MGI Symbol;Acc:MGI:3644593] | 1060 | -2.421 | 1.716E-54 |
| ENSMUSG00000062515 | Fabp4 | Enriched in Het retina | fatty acid binding protein 4, adipocyte [Source:MGI Symbol;Acc:MGI:88038] | 66.59 | -2.27 | 0.0002753 |
| ENSMUSG00000029304 | Spp1 | Enriched in Het retina | secreted phosphoprotein 1 [Source:MGI Symbol;Acc:MGI:98389] | 1333 | -1.874 | 0.000009676 |
| ENSMUSG00000034837 | Gnat1 | Enriched in Het retina | guanine nucleotide binding protein, alpha transducing 1 [Source:MGI Symbol;Acc:MGI:95778] | 10670 | -1.86 | 1.158E-17 |
| ENSMUSG00000025496 | Drd4 | Enriched in Het retina | dopamine receptor D4 [Source:MGI Symbol;Acc:MGI:94926] | 333 | -1.833 | 0.0009242 |
| ENSMUSG00000071076 | Jund | Enriched in Het retina | jun D proto-oncogene [Source:MGI Symbol;Acc:MGI:96648] | 61.49 | -1.833 | 0.003988 |
| ENSMUSG00000045246 | Kcng4 | Enriched in Het retina | potassium voltage-gated channel, subfamily G, member 4 [Source:MGI Symbol;Acc:MGI:1913983] | 347.2 | -1.786 | 0.000605 |
| ENSMUSG00000020581 | Agr2 | Enriched in Het retina | anterior gradient 2 [Source:MGI Symbol;Acc:MGI:1344405] | 65.57 | -1.765 | 0.000003669 |
| ENSMUSG00000030703 | Gdpd3 | Enriched in Het retina | glycerophosphodiester phosphodiesterase domain containing 3 [Source:MGI Symbol;Acc:MGI:1915866] | 127.1 | -1.733 | 0.003099 |
| ENSMUSG00000108398 | Gm30191 | Enriched in Het retina | predicted gene, 30191 [Source:MGI Symbol;Acc:MGI:5589350] | 282.1 | -1.629 | 2.388E-14 |
| ENSMUSG00000026173 | Plcd4 | Enriched in Het retina | phospholipase C, delta 4 [Source:MGI Symbol;Acc:MGI:107469] | 295.3 | -1.619 | 7.851E-08 |
| ENSMUSG00000007655 | Cav1 | Enriched in Het retina | caveolin 1, caveolae protein [Source:MGI Symbol;Acc:MGI:102709] | 948.7 | -1.608 | 3.644E-11 |
| ENSMUSG00000031448 | Adprhl1 | Enriched in Het retina | ADP-ribosylhydrolase like 1 [Source:MGI Symbol;Acc:MGI:2442168] | 51.69 | -1.599 | 0.000006127 |
| ENSMUSG00000073643 | Wdfy1 | Enriched in Het retina | WD repeat and FYVE domain containing 1 [Source:MGI Symbol;Acc:MGI:1916618] | 2068 | -1.591 | 4.634E-08 |
| ENSMUSG00000041828 | Abca8a | Enriched in Het retina | ATP-binding cassette, sub-family A (ABC1), member 8a [Source:MGI Symbol;Acc:MGI:2386846] | 1511 | -1.47 | 0.00004533 |
| ENSMUSG00000031394 | Opn1mw | Enriched in Het retina | opsin 1 (cone pigments), medium-wave-sensitive (color blindness, deutan) [Source:MGI Symbol;Acc:MGI:1097692] | 260.2 | -1.446 | 0.0005535 |
| ENSMUSG00000024411 | Aqp4 | Enriched in Het retina | aquaporin 4 [Source:MGI Symbol;Acc:MGI:107387] | 230 | -1.411 | 0.000001503 |
| ENSMUSG00000030787 | Lyve1 | Enriched in Het retina | lymphatic vessel endothelial hyaluronan receptor 1 [Source:MGI Symbol;Acc:MGI:2136348] | 94.8 | -1.404 | 0.0000221 |
| ENSMUSG00000022026 | Olfm4 | Enriched in Het retina | olfactomedin 4 [Source:MGI Symbol;Acc:MGI:2685142] | 55.58 | -1.371 | 0.00002496 |
| ENSMUSG00000034452 | Slc24a1 | Enriched in Het retina | solute carrier family 24 (sodium/potassium/calcium exchanger), member 1 [Source:MGI Symbol;Acc:MGI:2384871] | 4554 | -1.368 | 0.000000379 |
| ENSMUSG00000019772 | Vip | Enriched in Het retina | vasoactive intestinal polypeptide [Source:MGI Symbol;Acc:MGI:98933] | 152.5 | -1.359 | 8.153E-07 |
| ENSMUSG00000007682 | Dio2 | Enriched in Het retina | deiodinase, iodothyronine, type II [Source:MGI Symbol;Acc:MGI:1338833] | 481.2 | -1.339 | 7.83E-14 |
| ENSMUSG00000023484 | Prph | Enriched in Het retina | peripherin [Source:MGI Symbol;Acc:MGI:97774] | 98.02 | -1.306 | 0.04264 |
| ENSMUSG00000042073 | Abhd14b | Enriched in Het retina | abhydrolase domain containing 14b [Source:MGI Symbol;Acc:MGI:1923741] | 1662 | -1.266 | 0.001968 |
| ENSMUSG00000097789 | Gm2115 | Enriched in Het retina | predicted gene 2115 [Source:MGI Symbol;Acc:MGI:3780284] | 104.3 | -1.244 | 0.003557 |
| ENSMUSG00000048108 | Tmem72 | Enriched in Het retina | transmembrane protein 72 [Source:MGI Symbol;Acc:MGI:2442707] | 78.01 | -1.221 | 0.04065 |
| ENSMUSG00000040693 | Slco4c1 | Enriched in Het retina | solute carrier organic anion transporter family, member 4C1 [Source:MGI Symbol;Acc:MGI:2442784] | 65.33 | -1.217 | 0.001833 |
| ENSMUSG00000025739 | Gng13 | Enriched in Het retina | guanine nucleotide binding protein (G protein), gamma 13 [Source:MGI Symbol;Acc:MGI:1925616] | 474.3 | -1.193 | 0.000006127 |
| ENSMUSG00000114378 | Gm49355 | Enriched in Het retina | predicted gene, 49355 [Source:MGI Symbol;Acc:MGI:6121560] | 76.55 | -1.19 | 0.0009242 |
| ENSMUSG00000085007 | Smim43 | Enriched in Het retina | small integral membrane protein 43 [Source:MGI Symbol;Acc:MGI:3650339] | 61.35 | -1.187 | 0.01156 |
| ENSMUSG00000033207 | Mamdc2 | Enriched in Het retina | MAM domain containing 2 [Source:MGI Symbol;Acc:MGI:1918988] | 120.8 | -1.186 | 0.002242 |
| ENSMUSG00000009376 | Met | Enriched in Het retina | met proto-oncogene [Source:MGI Symbol;Acc:MGI:96969] | 235.5 | -1.148 | 0.008127 |
| ENSMUSG00000050211 | Pla2g4e | Enriched in Het retina | phospholipase A2, group IVE [Source:MGI Symbol;Acc:MGI:1919144] | 232 | -1.127 | 0.00004987 |
| ENSMUSG00000031373 | Car5b | Enriched in Het retina | carbonic anhydrase 5b, mitochondrial [Source:MGI Symbol;Acc:MGI:1926249] | 95.43 | -1.123 | 0.003512 |
| ENSMUSG00000027004 | Frzb | Enriched in Het retina | frizzled-related protein [Source:MGI Symbol;Acc:MGI:892032] | 638.8 | -1.112 | 9.051E-10 |
| ENSMUSG00000050663 | Trhde | Enriched in Het retina | TRH-degrading enzyme [Source:MGI Symbol;Acc:MGI:2384311] | 597 | -1.111 | 5.068E-13 |
| ENSMUSG00000040690 | Col16a1 | Enriched in Het retina | collagen, type XVI, alpha 1 [Source:MGI Symbol;Acc:MGI:1095396] | 233.1 | -1.101 | 0.0001034 |
| ENSMUSG00000028174 | Rpe65 | Enriched in Het retina | retinal pigment epithelium 65 [Source:MGI Symbol;Acc:MGI:98001] | 174.1 | -1.067 | 0.02789 |
| ENSMUSG00000023267 | Gabrr2 | Enriched in Het retina | gamma-aminobutyric acid (GABA) C receptor, subunit rho 2 [Source:MGI Symbol;Acc:MGI:95626] | 149.5 | -1.062 | 0.01607 |
| ENSMUSG00000004630 | Pcp2 | Enriched in Het retina | Purkinje cell protein 2 (L7) [Source:MGI Symbol;Acc:MGI:97508] | 2586 | -1.061 | 7.253E-15 |
| ENSMUSG00000038094 | Atp13a4 | Enriched in Het retina | ATPase type 13A4 [Source:MGI Symbol;Acc:MGI:1924456] | 55.52 | -1.022 | 0.01445 |
| ENSMUSG00000019122 | Ccl9 | Enriched in Het retina | chemokine (C-C motif) ligand 9 [Source:MGI Symbol;Acc:MGI:104533] | 73.74 | -1.017 | 0.0229 |
| ENSMUSG00000024331 | Dsc2 | Enriched in Het retina | desmocollin 2 [Source:MGI Symbol;Acc:MGI:103221] | 91.49 | -1.01 | 0.03583 |
| ENSMUSG00000019935 | Slc17a8 | Enriched in Het retina | solute carrier family 17 (sodium-dependent inorganic phosphate cotransporter), member 8 [Source:MGI Symbol;Acc:MGI:30396 | 212.7 | -1.005 | 0.0005773 |
| ENSMUSG00000031004 | Mki67 | Enriched in WT retina | antigen identified by monoclonal antibody Ki 67 [Source:MGI Symbol;Acc:MGI:106035] | 424.4 | 1.034 | 0.0009881 |
| ENSMUSG00000028661 | Epha8 | Enriched in WT retina | Eph receptor A8 [Source:MGI Symbol;Acc:MGI:109378] | 1885 | 1.037 | 0.000004608 |
| ENSMUSG00000073125 | Xlr3b | Enriched in WT retina | X-linked lymphocyte-regulated 3B [Source:MGI Symbol;Acc:MGI:109505] | 176.7 | 1.037 | 0.02232 |
| ENSMUSG00000027326 | Knl1 | Enriched in WT retina | kinetochore scaffold 1 [Source:MGI Symbol;Acc:MGI:1923714] | 41.46 | 1.07 | 0.04907 |
| ENSMUSG00000026879 | Gsn | Enriched in WT retina | gelsolin [Source:MGI Symbol;Acc:MGI:95851] | 371.7 | 1.074 | 0.007761 |
| ENSMUSG00000073733 | Cplane2 | Enriched in WT retina | ciliogenesis and planar polarity effector 2 [Source:MGI Symbol;Acc:MGI:1923416] | 71.65 | 1.108 | 0.0002585 |
| ENSMUSG00000033952 | Aspm | Enriched in WT retina | abnormal spindle microtubule assembly [Source:MGI Symbol;Acc:MGI:1334448] | 48.03 | 1.126 | 0.0148 |
| ENSMUSG00000071497 | Nutf2-ps1 | Enriched in WT retina | nuclear transport factor 2, pseudogene 1 [Source:MGI Symbol;Acc:MGI:108008] | 134.3 | 1.183 | 0.01844 |
| ENSMUSG00000045440 | Insm2 | Enriched in WT retina | insulinoma-associated 2 [Source:MGI Symbol;Acc:MGI:1930787] | 572.1 | 1.19 | 0.000002754 |
| ENSMUSG00000044447 | Dock5 | Enriched in WT retina | dedicator of cytokinesis 5 [Source:MGI Symbol;Acc:MGI:2652871] | 151.7 | 1.193 | 0.008556 |
| ENSMUSG00000031073 | Fgf15 | Enriched in WT retina | fibroblast growth factor 15 [Source:MGI Symbol;Acc:MGI:1096383] | 75.49 | 1.201 | 0.002386 |
| ENSMUSG00000035783 | Acta2 | Enriched in WT retina | actin, alpha 2, smooth muscle, aorta [Source:MGI Symbol;Acc:MGI:87909] | 117.7 | 1.225 | 0.001488 |
| ENSMUSG00000031995 | St14 | Enriched in WT retina | suppression of tumorigenicity 14 (colon carcinoma) [Source:MGI Symbol;Acc:MGI:1338881] | 249.4 | 1.259 | 0.0001694 |
| ENSMUSG00000041301 | Cftr | Enriched in WT retina | cystic fibrosis transmembrane conductance regulator [Source:MGI Symbol;Acc:MGI:88388] | 76.3 | 1.28 | 0.002821 |
| ENSMUSG00000013415 | Igf2bp1 | Enriched in WT retina | insulin-like growth factor 2 mRNA binding protein 1 [Source:MGI Symbol;Acc:MGI:1890357] | 49.77 | 1.304 | 0.0001131 |
| ENSMUSG00000021799 | Opn4 | Enriched in WT retina | opsin 4 (melanopsin) [Source:MGI Symbol;Acc:MGI:1353425] | 134.3 | 1.35 | 0.00001748 |
| ENSMUSG00000044468 | Tent5c | Enriched in WT retina | terminal nucleotidyltransferase 5C [Source:MGI Symbol;Acc:MGI:1921895] | 261.8 | 1.454 | 0.03357 |
| ENSMUSG00000054252 | Fgfr3 | Enriched in WT retina | fibroblast growth factor receptor 3 [Source:MGI Symbol;Acc:MGI:95524] | 136 | 1.576 | 0.0006824 |
| ENSMUSG00000053198 | Prx | Enriched in WT retina | periaxin [Source:MGI Symbol;Acc:MGI:108176] | 99.8 | 1.585 | 0.009017 |
| ENSMUSG00000091956 | C2cd4b | Enriched in WT retina | C2 calcium-dependent domain containing 4B [Source:MGI Symbol;Acc:MGI:1922947] | 135.9 | 1.688 | 8.233E-11 |
| ENSMUSG00000028776 | Tinagl1 | Enriched in WT retina | tubulointerstitial nephritis antigen-like 1 [Source:MGI Symbol;Acc:MGI:2137617] | 54.86 | 1.759 | 0.02579 |
| ENSMUSG00000033249 | Hsf4 | Enriched in WT retina | heat shock transcription factor 4 [Source:MGI Symbol;Acc:MGI:1347058] | 76.54 | 1.925 | 0.003402 |
| ENSMUSG00000041731 | Pgm5 | Enriched in WT retina | phosphoglucomutase 5 [Source:MGI Symbol;Acc:MGI:1925668] | 50.91 | 2.315 | 0.002721 |
| ENSMUSG00000096014 | Sox1 | Enriched in WT retina | SRY (sex determining region Y)-box 1 [Source:MGI Symbol;Acc:MGI:98357] | 49.13 | 2.681 | 0.001958 |
| ENSMUSG00000025229 | Pitx3 | Enriched in WT retina | paired-like homeodomain transcription factor 3 [Source:MGI Symbol;Acc:MGI:1100498] | 27.93 | 2.684 | 0.04369 |
| ENSMUSG00000039457 | Ppl | Enriched in WT retina | periplakin [Source:MGI Symbol;Acc:MGI:1194898] | 53.94 | 3.019 | 0.01445 |
| ENSMUSG00000033501 | Crygs | Enriched in WT retina | crystallin, gamma S [Source:MGI Symbol;Acc:MGI:1298216] | 6851 | 3.028 | 0.03713 |
| ENSMUSG00000038135 | Crygn | Enriched in WT retina | crystallin, gamma N [Source:MGI Symbol;Acc:MGI:2449167] | 858.9 | 3.033 | 0.03001 |
| ENSMUSG00000042240 | Crybb2 | Enriched in WT retina | crystallin, beta B2 [Source:MGI Symbol;Acc:MGI:88519] | 11780 | 3.055 | 0.04045 |
| ENSMUSG00000024041 | Cryaa | Enriched in WT retina | crystallin, alpha A [Source:MGI Symbol;Acc:MGI:88515] | 47370 | 3.15 | 0.02247 |
| ENSMUSG00000029343 | Crybb1 | Enriched in WT retina | crystallin, beta B1 [Source:MGI Symbol;Acc:MGI:104992] | 8124 | 3.244 | 0.009093 |
| ENSMUSG00000038840 | Birc7 | Enriched in WT retina | baculoviral IAP repeat-containing 7 (livin) [Source:MGI Symbol;Acc:MGI:2676458] | 26.6 | 3.255 | 0.001815 |
| ENSMUSG00000059900 | Tmem40 | Enriched in WT retina | transmembrane protein 40 [Source:MGI Symbol;Acc:MGI:2137870] | 110.3 | 3.271 | 0.011 |
| ENSMUSG00000066975 | Cryba4 | Enriched in WT retina | crystallin, beta A4 [Source:MGI Symbol;Acc:MGI:102716] | 6251 | 3.29 | 0.01056 |
| ENSMUSG00000029352 | Crybb3 | Enriched in WT retina | crystallin, beta B3 [Source:MGI Symbol;Acc:MGI:102717] | 13510 | 3.313 | 0.007036 |
| ENSMUSG00000058626 | Capn11 | Enriched in WT retina | calpain 11 [Source:MGI Symbol;Acc:MGI:1352490] | 44.11 | 3.349 | 7.723E-09 |
| ENSMUSG00000006546 | Cryba2 | Enriched in WT retina | crystallin, beta A2 [Source:MGI Symbol;Acc:MGI:104336] | 6329 | 3.423 | 0.0123 |
| ENSMUSG00000025389 | Mip | Enriched in WT retina | major intrinsic protein of lens fiber [Source:MGI Symbol;Acc:MGI:96990] | 3122 | 3.471 | 0.03147 |
| ENSMUSG00000000724 | Cryba1 | Enriched in WT retina | crystallin, beta A1 [Source:MGI Symbol;Acc:MGI:88518] | 22200 | 3.488 | 0.006842 |
| ENSMUSG00000025952 | Crygc | Enriched in WT retina | crystallin, gamma C [Source:MGI Symbol;Acc:MGI:88523] | 4377 | 3.552 | 0.001954 |
| ENSMUSG00000032556 | Bfsp2 | Enriched in WT retina | beaded filament structural protein 2, phakinin [Source:MGI Symbol;Acc:MGI:1333828] | 1058 | 3.56 | 0.0001309 |
| ENSMUSG00000027420 | Bfsp1 | Enriched in WT retina | beaded filament structural protein 1, in lens-CP94 [Source:MGI Symbol;Acc:MGI:101770] | 1424 | 3.65 | 0.00001818 |
| ENSMUSG00000073658 | Crygb | Enriched in WT retina | crystallin, gamma B [Source:MGI Symbol;Acc:MGI:88522] | 7986 | 3.663 | 0.001011 |
| ENSMUSG00000067299 | Crygd | Enriched in WT retina | crystallin, gamma D [Source:MGI Symbol;Acc:MGI:88524] | 7701 | 3.7 | 0.0008175 |
| ENSMUSG00000070870 | Cryge | Enriched in WT retina | crystallin, gamma E [Source:MGI Symbol;Acc:MGI:88525] | 5510 | 3.751 | 0.0006279 |
| ENSMUSG00000048582 | Gja3 | Enriched in WT retina | gap junction protein, alpha 3 [Source:MGI Symbol;Acc:MGI:95714] | 260.7 | 3.805 | 0.04154 |
| ENSMUSG00000044429 | Cryga | Enriched in WT retina | crystallin, gamma A [Source:MGI Symbol;Acc:MGI:88521] | 1736 | 3.871 | 0.00009709 |
| ENSMUSG00000032401 | Lctl | Enriched in WT retina | lactase-like [Source:MGI Symbol;Acc:MGI:2183549] | 115.4 | 4.004 | 0.000452 |
| ENSMUSG00000025945 | Crygf | Enriched in WT retina | crystallin, gamma F [Source:MGI Symbol;Acc:MGI:88526] | 2381 | 4.382 | 0.00001908 |
| ENSMUSG00000026253 | Chrng | Enriched in WT retina | cholinergic receptor, nicotinic, gamma polypeptide [Source:MGI Symbol;Acc:MGI:87895] | 24.56 | 4.89 | 0.03061 |

**Contrast numerator: KO_retina. Contrast denominator: Het_retina.**

| ensembl_gene_id | external_gene_name | Significance | description | baseMean | logFC | adj.P.Val |
|---|---|---|---|---|---|---|
| ENSMUSG00000093805 | Gal3st2b | Enriched in Het retina | galactose-3-O-sulfotransferase 2B [Source:MGI Symbol;Acc:MGI:3711964] | 45.05 | -7.862 | 2.239E-08 |
| ENSMUSG00000021799 | Opn4 | Enriched in Het retina | opsin 4 (melanopsin) [Source:MGI Symbol;Acc:MGI:1353425] | 66.87 | -5.452 | 2.654E-15 |
| ENSMUSG00000041479 | Syt15 | Enriched in Het retina | synaptotagmin XV [Source:MGI Symbol;Acc:MGI:2442166] | 97.75 | -3.667 | 2.079E-16 |
| ENSMUSG00000020415 | Pttg1 | Enriched in Het retina | pituitary tumor-transforming gene 1 [Source:MGI Symbol;Acc:MGI:1353578] | 360.6 | -2.216 | 0.002411 |
| ENSMUSG00000032221 | Mns1 | Enriched in Het retina | meiosis-specific nuclear structural protein 1 [Source:MGI Symbol;Acc:MGI:107933] | 531 | -1.78 | 0.0001942 |
| ENSMUSG00000026173 | Plcd4 | Enriched in Het retina | phospholipase C, delta 4 [Source:MGI Symbol;Acc:MGI:107469] | 310 | -1.694 | 8.022E-07 |
| ENSMUSG00000020268 | Lyrm7 | Enriched in Het retina | LYR motif containing 7 [Source:MGI Symbol;Acc:MGI:1922780] | 299.9 | -1.083 | 1.915E-09 |
| ENSMUSG00000001025 | S100a6 | Enriched in KO retina | S100 calcium binding protein A6 (calcyclin) [Source:MGI Symbol;Acc:MGI:1339467] | 509.7 | 1.029 | 0.002243 |
| ENSMUSG00000040489 | Sox30 | Enriched in KO retina | SRY (sex determining region Y)-box 30 [Source:MGI Symbol;Acc:MGI:1341157] | 238.8 | 1.655 | 0.0007092 |
| ENSMUSG00000053441 | Adamts19 | Enriched in KO retina | a disintegrin-like and metallopeptidase (reprolysin type) with thrombospondin type 1 motif, 19 [Source:MGI Symbol;Acc:MGI:244287 | 213.9 | 1.933 | 0.000002329 |
| ENSMUSG00000107256 | Gm13821 | Enriched in KO retina | predicted gene 13821 [Source:MGI Symbol;Acc:MGI:3652003] | 44.2 | 2.093 | 0.04726 |
| ENSMUSG00000036480 | Prss56 | Enriched in KO retina | protease, serine 56 [Source:MGI Symbol;Acc:MGI:1916703] | 3197 | 2.703 | 0.01787 |
| ENSMUSG00000112449 | Srp54b | Enriched in KO retina | signal recognition particle 54B [Source:MGI Symbol;Acc:MGI:3714357] | 44.06 | 4.595 | 0.03169 |

**Contrast numerator: WT_retina. Contrast denominator: KO_retina.**

| ensembl_gene_id | external_gene_name | Significance | description | baseMean | logFC | adj.P.Val |
|---|---|---|---|---|---|---|
| ENSMUSG00000111176 | Gm47233 | Enriched in KO retina | predicted gene, 47233 [Source:MGI Symbol;Acc:MGI:6096050] | 76.38 | -9.703 | 0.000004132 |
| ENSMUSG00000072476 | Gm9008 | Enriched in KO retina | predicted pseudogene 9008 [Source:MGI Symbol;Acc:MGI:3644000] | 159.4 | -7.225 | 1.927E-26 |
| ENSMUSG00000035296 | Sgcg | Enriched in KO retina | sarcoglycan, gamma (dystrophin-associated glycoprotein) [Source:MGI Symbol;Acc:MGI:1346524] | 161 | -5.857 | 1.791E-37 |
| ENSMUSG00000072572 | Slc39a2 | Enriched in KO retina | solute carrier family 39 (zinc transporter), member 2 [Source:MGI Symbol;Acc:MGI:2684326] | 499.7 | -4.881 | 2.661E-147 |
| ENSMUSG00000112449 | Srp54b | Enriched in KO retina | signal recognition particle 54B [Source:MGI Symbol;Acc:MGI:3714357] | 71.25 | -4.613 | 0.01189 |
| ENSMUSG00000090799 | Klhl33 | Enriched in KO retina | kelch-like 33 [Source:MGI Symbol;Acc:MGI:3644593] | 1318 | -3.054 | 1.031E-130 |
| ENSMUSG00000037225 | Fgf2 | Enriched in KO retina | fibroblast growth factor 2 [Source:MGI Symbol;Acc:MGI:95516] | 795.2 | -2.894 | 0.00009183 |
| ENSMUSG00000062515 | Fabp4 | Enriched in KO retina | fatty acid binding protein 4, adipocyte [Source:MGI Symbol;Acc:MGI:88038] | 90.43 | -2.859 | 1.748E-09 |
| ENSMUSG00000040489 | Sox30 | Enriched in KO retina | SRY (sex determining region Y)-box 30 [Source:MGI Symbol;Acc:MGI:1341157] | 295.3 | -2.593 | 4.734E-30 |
| ENSMUSG00000029304 | Spp1 | Enriched in KO retina | secreted phosphoprotein 1 [Source:MGI Symbol;Acc:MGI:98389] | 1439 | -2.461 | 0.005293 |
| ENSMUSG00000021091 | Serpina3n | Enriched in KO retina | serine (or cysteine) peptidase inhibitor, clade A, member 3N [Source:MGI Symbol;Acc:MGI:105045] | 1098 | -2.385 | 0.003163 |
| ENSMUSG00000023484 | Prph | Enriched in KO retina | peripherin [Source:MGI Symbol;Acc:MGI:97774] | 121.1 | -1.973 | 0.000004974 |
| ENSMUSG00000073643 | Wdfy1 | Enriched in KO retina | WD repeat and FYVE domain containing 1 [Source:MGI Symbol;Acc:MGI:1916618] | 2372 | -1.905 | 2.47E-14 |
| ENSMUSG00000030703 | Gdpd3 | Enriched in KO retina | glycerophosphodiester phosphodiesterase domain containing 3 [Source:MGI Symbol;Acc:MGI:191586 | 102.8 | -1.772 | 0.0004243 |
| ENSMUSG00000030787 | Lyve1 | Enriched in KO retina | lymphatic vessel endothelial hyaluronan receptor 1 [Source:MGI Symbol;Acc:MGI:2136348] | 97.33 | -1.645 | 0.000001519 |
| ENSMUSG00000025044 | Msr1 | Enriched in KO retina | macrophage scavenger receptor 1 [Source:MGI Symbol;Acc:MGI:98257] | 62.67 | -1.633 | 0.006607 |
| ENSMUSG00000001506 | Col1a1 | Enriched in KO retina | collagen, type I, alpha 1 [Source:MGI Symbol;Acc:MGI:88467] | 111.9 | -1.554 | 0.0005169 |
| ENSMUSG00000037071 | Scd1 | Enriched in KO retina | stearoyl-Coenzyme A desaturase 1 [Source:MGI Symbol;Acc:MGI:98239] | 425.2 | -1.43 | 1.446E-07 |
| ENSMUSG00000033491 | Prss35 | Enriched in KO retina | protease, serine 35 [Source:MGI Symbol;Acc:MGI:2444800] | 100.6 | -1.388 | 0.0003968 |
| ENSMUSG00000007682 | Dio2 | Enriched in KO retina | deiodinase, iodothyronine, type II [Source:MGI Symbol;Acc:MGI:1338833] | 461.3 | -1.376 | 7.163E-19 |
| ENSMUSG00000025496 | Drd4 | Enriched in KO retina | dopamine receptor D4 [Source:MGI Symbol;Acc:MGI:94926] | 225.8 | -1.311 | 0.000003831 |
| ENSMUSG00000020932 | Gfap | Enriched in KO retina | glial fibrillary acidic protein [Source:MGI Symbol;Acc:MGI:95697] | 3878 | -1.299 | 3.733E-29 |
| ENSMUSG00000036242 | Armh4 | Enriched in KO retina | armadillo-like helical domain containing 4 [Source:MGI Symbol;Acc:MGI:1914669] | 1191 | -1.292 | 7.046E-36 |
| ENSMUSG00000068699 | Flnc | Enriched in KO retina | filamin C, gamma [Source:MGI Symbol;Acc:MGI:95557] | 63.5 | -1.277 | 0.01796 |
| ENSMUSG00000043461 | Sptssb | Enriched in KO retina | serine palmitoyltransferase, small subunit B [Source:MGI Symbol;Acc:MGI:1913433] | 69.86 | -1.253 | 0.004362 |
| ENSMUSG00000071637 | Cebpd | Enriched in KO retina | CCAAT/enhancer binding protein (C/EBP), delta [Source:MGI Symbol;Acc:MGI:103573] | 469.1 | -1.223 | 0.04256 |
| ENSMUSG00000050663 | Trhde | Enriched in KO retina | TRH-degrading enzyme [Source:MGI Symbol;Acc:MGI:2384311] | 647.2 | -1.216 | 2.841E-16 |
| ENSMUSG00000061462 | Obscn | Enriched in KO retina | obscurin, cytoskeletal calmodulin and titin-interacting RhoGEF [Source:MGI Symbol;Acc:MGI:2681862] | 95.91 | -1.215 | 0.004153 |
| ENSMUSG00000038457 | Tmem255b | Enriched in KO retina | transmembrane protein 255B [Source:MGI Symbol;Acc:MGI:2685533] | 122.6 | -1.201 | 0.0006391 |
| ENSMUSG00000108398 | Gm30191 | Enriched in KO retina | predicted gene, 30191 [Source:MGI Symbol;Acc:MGI:5589350] | 168.7 | -1.159 | 0.001412 |
| ENSMUSG00000037705 | Tecta | Enriched in KO retina | tectorin alpha [Source:MGI Symbol;Acc:MGI:109575] | 67.14 | -1.155 | 0.02551 |
| ENSMUSG00000058183 | Mmel1 | Enriched in KO retina | membrane metallo-endopeptidase-like 1 [Source:MGI Symbol;Acc:MGI:1351603] | 152.5 | -1.149 | 0.001481 |
| ENSMUSG00000003469 | Phyhip | Enriched in KO retina | phytanoyl-CoA hydroxylase interacting protein [Source:MGI Symbol;Acc:MGI:1860417] | 194 | -1.143 | 0.000001264 |
| ENSMUSG00000042010 | Acacb | Enriched in KO retina | acetyl-Coenzyme A carboxylase beta [Source:MGI Symbol;Acc:MGI:2140940] | 191.5 | -1.11 | 0.00001216 |
| ENSMUSG00000022074 | Tnfrsf10b | Enriched in KO retina | tumor necrosis factor receptor superfamily, member 10b [Source:MGI Symbol;Acc:MGI:1341090] | 86.04 | -1.099 | 0.01254 |
| ENSMUSG00000033318 | Gstt2 | Enriched in KO retina | glutathione S-transferase, theta 2 [Source:MGI Symbol;Acc:MGI:106188] | 122.8 | -1.075 | 0.0008262 |
| ENSMUSG00000004630 | Pcp2 | Enriched in KO retina | Purkinje cell protein 2 (L7) [Source:MGI Symbol;Acc:MGI:97508] | 2493 | -1.069 | 9.631E-23 |
| ENSMUSG00000025739 | Gng13 | Enriched in KO retina | guanine nucleotide binding protein (G protein), gamma 13 [Source:MGI Symbol;Acc:MGI:1925616] | 414.2 | -1.056 | 1.505E-08 |
| ENSMUSG00000022431 | Ribc2 | Enriched in KO retina | RIB43A domain with coiled-coils 2 [Source:MGI Symbol;Acc:MGI:1914997] | 72.25 | -1.035 | 0.0202 |
| ENSMUSG00000033949 | Trim36 | Enriched in KO retina | tripartite motif-containing 36 [Source:MGI Symbol;Acc:MGI:106264] | 2485 | -1.035 | 8.328E-30 |
| ENSMUSG00000034837 | Gnat1 | Enriched in KO retina | guanine nucleotide binding protein, alpha transducing 1 [Source:MGI Symbol;Acc:MGI:95778] | 6752 | -1.035 | 0.0001575 |
| ENSMUSG00000022324 | Matn2 | Enriched in KO retina | matrilin 2 [Source:MGI Symbol;Acc:MGI:109613] | 150.5 | -1.032 | 0.01308 |
| ENSMUSG00000030790 | Adm | Enriched in KO retina | adrenomedullin [Source:MGI Symbol;Acc:MGI:108058] | 277.9 | -1.03 | 0.0002529 |
| ENSMUSG00000036585 | Fgf1 | Enriched in KO retina | fibroblast growth factor 1 [Source:MGI Symbol;Acc:MGI:95515] | 417.5 | -1.026 | 0.00009183 |
| ENSMUSG00000096255 | Dynlt1b | Enriched in KO retina | dynein light chain Tctex-type 1B [Source:MGI Symbol;Acc:MGI:98643] | 179.8 | -1.019 | 0.004232 |
| ENSMUSG00000012819 | Cdh23 | Enriched in KO retina | cadherin 23 (otocadherin) [Source:MGI Symbol;Acc:MGI:1890219] | 142.1 | -1.017 | 0.0008221 |
| ENSMUSG00000114378 | Gm49355 | Enriched in KO retina | predicted gene, 49355 [Source:MGI Symbol;Acc:MGI:6121560] | 64.18 | -1.017 | 0.04875 |
| ENSMUSG00000034361 | Cpne2 | Enriched in KO retina | copine II [Source:MGI Symbol;Acc:MGI:2387578] | 125 | -1.012 | 0.006116 |
| ENSMUSG00000021699 | Pde4d | Enriched in KO retina | phosphodiesterase 4D, cAMP specific [Source:MGI Symbol;Acc:MGI:99555] | 2444 | -1.008 | 0.000769 |
| ENSMUSG00000032807 | Alox12b | Enriched in KO retina | arachidonate 12-lipoxygenase, 12R type [Source:MGI Symbol;Acc:MGI:1274782] | 67.94 | -1.007 | 0.01948 |
| ENSMUSG00000038473 | Nos1ap | Enriched in KO retina | nitric oxide synthase 1 (neuronal) adaptor protein [Source:MGI Symbol;Acc:MGI:1917979] | 843.7 | -1.006 | 1.283E-08 |
| ENSMUSG00000043541 | Casc1 | Enriched in KO retina | cancer susceptibility candidate 1 [Source:MGI Symbol;Acc:MGI:2444480] | 136.7 | -1.006 | 0.001648 |
| ENSMUSG00000026605 | Cenpf | Enriched in WT retina | centromere protein F [Source:MGI Symbol;Acc:MGI:1313302] | 146.5 | 1.006 | 0.005706 |
| ENSMUSG00000039543 | Cfap70 | Enriched in WT retina | cilia and flagella associated protein 70 [Source:MGI Symbol;Acc:MGI:1923920] | 115 | 1.012 | 0.003768 |
| ENSMUSG00000046861 | Hectd3 | Enriched in WT retina | HECT domain E3 ubiquitin protein ligase 3 [Source:MGI Symbol;Acc:MGI:1923858] | 1410 | 1.024 | 0.003511 |
| ENSMUSG00000020649 | Rrm2 | Enriched in WT retina | ribonucleotide reductase M2 [Source:MGI Symbol;Acc:MGI:98181] | 186.3 | 1.027 | 0.0009788 |
| ENSMUSG00000068762 | Gstm6 | Enriched in WT retina | glutathione S-transferase, mu 6 [Source:MGI Symbol;Acc:MGI:1309467] | 116.4 | 1.035 | 0.004656 |
| ENSMUSG00000049357 | Brd8dc | Enriched in WT retina | BRD8 domain containing [Source:MGI Symbol;Acc:MGI:3045347] | 81.87 | 1.056 | 0.007817 |
| ENSMUSG00000020799 | Tekt1 | Enriched in WT retina | tektin 1 [Source:MGI Symbol;Acc:MGI:1333819] | 93.29 | 1.066 | 0.03053 |
| ENSMUSG00000020000 | Moxd1 | Enriched in WT retina | monooxygenase, DBH-like 1 [Source:MGI Symbol;Acc:MGI:1921582] | 747.2 | 1.095 | 2.15E-12 |
| ENSMUSG00000027875 | Hmgcs2 | Enriched in WT retina | 3-hydroxy-3-methylglutaryl-Coenzyme A synthase 2 [Source:MGI Symbol;Acc:MGI:101939] | 112.6 | 1.111 | 0.002023 |
| ENSMUSG00000072774 | Zfp951 | Enriched in WT retina | zinc finger protein 951 [Source:MGI Symbol;Acc:MGI:2441896] | 161.5 | 1.122 | 0.002095 |
| ENSMUSG00000045440 | Insm2 | Enriched in WT retina | insulinoma-associated 2 [Source:MGI Symbol;Acc:MGI:1930787] | 788.1 | 1.135 | 0.004994 |
| ENSMUSG00000073733 | Cplane2 | Enriched in WT retina | ciliogenesis and planar polarity effector 2 [Source:MGI Symbol;Acc:MGI:1923416] | 93.14 | 1.17 | 0.002068 |
| ENSMUSG00000018733 | Pex12 | Enriched in WT retina | peroxisomal biogenesis factor 12 [Source:MGI Symbol;Acc:MGI:2144177] | 131.5 | 1.193 | 0.007983 |
| ENSMUSG00000053219 | Raet1e | Enriched in WT retina | retinoic acid early transcript 1E [Source:MGI Symbol;Acc:MGI:2675273] | 446.2 | 1.225 | 1.793E-16 |
| ENSMUSG00000020481 | Ankrd36 | Enriched in WT retina | ankyrin repeat domain 36 [Source:MGI Symbol;Acc:MGI:1923639] | 131.5 | 1.228 | 0.0001432 |
| ENSMUSG00000031073 | Fgf15 | Enriched in WT retina | fibroblast growth factor 15 [Source:MGI Symbol;Acc:MGI:1096383] | 104.6 | 1.242 | 0.0006665 |
| ENSMUSG00000013415 | Igf2bp1 | Enriched in WT retina | insulin-like growth factor 2 mRNA binding protein 1 [Source:MGI Symbol;Acc:MGI:1890357] | 71.86 | 1.312 | 0.007371 |
| ENSMUSG00000063796 | Slc22a8 | Enriched in WT retina | solute carrier family 22 (organic anion transporter), member 8 [Source:MGI Symbol;Acc:MGI:1336187 | 60.99 | 1.328 | 0.01338 |
| ENSMUSG00000079941 | Cox5b-ps | Enriched in WT retina | cytochrome c oxidase subunit 5B, pseudogene [Source:MGI Symbol;Acc:MGI:3649411] | 555.2 | 1.338 | 5.786E-19 |
| ENSMUSG00000064354 | mt-Co2 | Enriched in WT retina | mitochondrially encoded cytochrome c oxidase II [Source:MGI Symbol;Acc:MGI:102503] | 75.49 | 1.38 | 0.00252 |
| ENSMUSG00000003484 | Cyp4f18 | Enriched in WT retina | cytochrome P450, family 4, subfamily f, polypeptide 18 [Source:MGI Symbol;Acc:MGI:1919304] | 52.89 | 1.393 | 0.03924 |
| ENSMUSG00000026774 | Potegl | Enriched in WT retina | POTE ankyrin domain family, member G like [Source:MGI Symbol;Acc:MGI:1918231] | 59.31 | 1.406 | 0.00865 |
| ENSMUSG00000072494 | Ppp1r3e | Enriched in WT retina | protein phosphatase 1, regulatory subunit 3E [Source:MGI Symbol;Acc:MGI:2145790] | 891 | 1.428 | 6.994E-43 |
| ENSMUSG00000031995 | St14 | Enriched in WT retina | suppression of tumorigenicity 14 (colon carcinoma) [Source:MGI Symbol;Acc:MGI:1338881] | 354.7 | 1.434 | 5.879E-07 |
| ENSMUSG00000020268 | Lyrm7 | Enriched in WT retina | LYR motif containing 7 [Source:MGI Symbol;Acc:MGI:1922780] | 321.5 | 1.438 | 1.265E-16 |
| ENSMUSG00000032572 | Col6a4 | Enriched in WT retina | collagen, type VI, alpha 4 [Source:MGI Symbol;Acc:MGI:1915803] | 60.52 | 1.452 | 0.002726 |
| ENSMUSG00000054404 | Slfn5 | Enriched in WT retina | schlafen 5 [Source:MGI Symbol;Acc:MGI:1329004] | 55.73 | 1.551 | 0.01341 |
| ENSMUSG00000059343 | Aldoart1 | Enriched in WT retina | aldolase 1 A, retrogene 1 [Source:MGI Symbol;Acc:MGI:2447811] | 157.3 | 1.661 | 8.795E-09 |
| ENSMUSG00000022584 | Ly6c2 | Enriched in WT retina | lymphocyte antigen 6 complex, locus C2 [Source:MGI Symbol;Acc:MGI:3712069] | 40.57 | 1.673 | 0.04933 |
| ENSMUSG00000035295 | Wdr38 | Enriched in WT retina | WD repeat domain 38 [Source:MGI Symbol;Acc:MGI:1923896] | 43.25 | 1.751 | 0.01813 |
| ENSMUSG00000100486 | Gm4131 | Enriched in WT retina | predicted gene 4131 [Source:MGI Symbol;Acc:MGI:3782307] | 283.8 | 1.788 | 6.95E-18 |
| ENSMUSG00000091956 | C2cd4b | Enriched in WT retina | C2 calcium-dependent domain containing 4B [Source:MGI Symbol;Acc:MGI:1922947] | 191.6 | 1.79 | 3.351E-10 |
| ENSMUSG00000028571 | Cyp2j13 | Enriched in WT retina | cytochrome P450, family 2, subfamily j, polypeptide 13 [Source:MGI Symbol;Acc:MGI:2385197] | 255.5 | 1.804 | 8.738E-21 |
| ENSMUSG00000038086 | Hspb2 | Enriched in WT retina | heat shock protein 2 [Source:MGI Symbol;Acc:MGI:1916503] | 36.39 | 1.916 | 0.04733 |
| ENSMUSG00000064358 | mt-Co3 | Enriched in WT retina | mitochondrially encoded cytochrome c oxidase III [Source:MGI Symbol;Acc:MGI:102502] | 43.46 | 2.101 | 0.0002142 |
| ENSMUSG00000054252 | Fgfr3 | Enriched in WT retina | fibroblast growth factor receptor 3 [Source:MGI Symbol;Acc:MGI:95524] | 182.7 | 2.11 | 4.602E-09 |
| ENSMUSG00000041731 | Pgm5 | Enriched in WT retina | phosphoglucomutase 5 [Source:MGI Symbol;Acc:MGI:1925668] | 82.91 | 2.243 | 0.0001749 |
| ENSMUSG00000109446 | Gm9195 | Enriched in WT retina | predicted gene 9195 [Source:MGI Symbol;Acc:MGI:3779838] | 49.8 | 2.568 | 0.00001202 |
| ENSMUSG00000034248 | Slc25a37 | Enriched in WT retina | solute carrier family 25, member 37 [Source:MGI Symbol;Acc:MGI:1914962] | 1002 | 2.606 | 2.044E-125 |
| ENSMUSG00000096014 | Sox1 | Enriched in WT retina | SRY (sex determining region Y)-box 1 [Source:MGI Symbol;Acc:MGI:98357] | 76.85 | 2.698 | 0.000006151 |
| ENSMUSG00000032221 | Mns1 | Enriched in WT retina | meiosis-specific nuclear structural protein 1 [Source:MGI Symbol;Acc:MGI:107933] | 676.8 | 2.861 | 1.187E-07 |
| ENSMUSG00000029343 | Crybb1 | Enriched in WT retina | crystallin, beta B1 [Source:MGI Symbol;Acc:MGI:104992] | 14370 | 2.985 | 0.03 |
| ENSMUSG00000039630 | Hnrnpu | Enriched in WT retina | heterogeneous nuclear ribonucleoprotein U [Source:MGI Symbol;Acc:MGI:1858195] | 678.4 | 3.006 | 0.008583 |
| ENSMUSG00000030093 | Wnt7a | Enriched in WT retina | wingless-type MMTV integration site family, member 7A [Source:MGI Symbol;Acc:MGI:98961] | 41.52 | 3.029 | 0.0005169 |
| ENSMUSG00000022382 | Wnt7b | Enriched in WT retina | wingless-type MMTV integration site family, member 7B [Source:MGI Symbol;Acc:MGI:98962] | 235.3 | 3.108 | 0.005453 |
| ENSMUSG00000041479 | Syt15 | Enriched in WT retina | synaptotagmin XV [Source:MGI Symbol;Acc:MGI:2442166] | 51.71 | 3.108 | 1.067E-08 |
| ENSMUSG00000020415 | Pttg1 | Enriched in WT retina | pituitary tumor-transforming gene 1 [Source:MGI Symbol;Acc:MGI:1353578] | 367.1 | 3.136 | 3.368E-65 |
| ENSMUSG00000022383 | Ppara | Enriched in WT retina | peroxisome proliferator activated receptor alpha [Source:MGI Symbol;Acc:MGI:104740] | 178.1 | 3.178 | 0.01044 |
| ENSMUSG00000029352 | Crybb3 | Enriched in WT retina | crystallin, beta B3 [Source:MGI Symbol;Acc:MGI:102717] | 23310 | 3.309 | 0.01162 |
| ENSMUSG00000059900 | Tmem40 | Enriched in WT retina | transmembrane protein 40 [Source:MGI Symbol;Acc:MGI:2137870] | 185.1 | 3.346 | 0.01838 |
| ENSMUSG00000021390 | Ogn | Enriched in WT retina | osteoglycin [Source:MGI Symbol;Acc:MGI:109278] | 132.5 | 3.351 | 0.04074 |
| ENSMUSG00000032401 | Lctl | Enriched in WT retina | lactase-like [Source:MGI Symbol;Acc:MGI:2183549] | 213.4 | 3.359 | 0.01465 |
| ENSMUSG00000024041 | Cryaa | Enriched in WT retina | crystallin, alpha A [Source:MGI Symbol;Acc:MGI:88515] | 80160 | 3.497 | 0.01015 |
| ENSMUSG00000066975 | Cryba4 | Enriched in WT retina | crystallin, beta A4 [Source:MGI Symbol;Acc:MGI:102716] | 10720 | 3.623 | 0.006056 |
| ENSMUSG00000000724 | Cryba1 | Enriched in WT retina | crystallin, beta A1 [Source:MGI Symbol;Acc:MGI:88518] | 38740 | 3.633 | 0.009305 |
| ENSMUSG00000033501 | Crygs | Enriched in WT retina | crystallin, gamma S [Source:MGI Symbol;Acc:MGI:1298216] | 11410 | 3.84 | 0.00551 |
| ENSMUSG00000006546 | Cryba2 | Enriched in WT retina | crystallin, beta A2 [Source:MGI Symbol;Acc:MGI:104336] | 10800 | 3.976 | 0.001512 |
| ENSMUSG00000038135 | Crygn | Enriched in WT retina | crystallin, gamma N [Source:MGI Symbol;Acc:MGI:2449167] | 1424 | 4.013 | 0.002726 |
| ENSMUSG00000032556 | Bfsp2 | Enriched in WT retina | beaded filament structural protein 2, phakinin [Source:MGI Symbol;Acc:MGI:1333828] | 1745 | 4.043 | 0.004505 |
| ENSMUSG00000042240 | Crybb2 | Enriched in WT retina | crystallin, beta B2 [Source:MGI Symbol;Acc:MGI:88519] | 19310 | 4.052 | 0.003561 |
| ENSMUSG00000048582 | Gja3 | Enriched in WT retina | gap junction protein, alpha 3 [Source:MGI Symbol;Acc:MGI:95714] | 424.6 | 4.676 | 0.03558 |
| ENSMUSG00000025389 | Mip | Enriched in WT retina | major intrinsic protein of lens fiber [Source:MGI Symbol;Acc:MGI:96990] | 5109 | 5.435 | 0.0001796 |
| ENSMUSG00000025229 | Pitx3 | Enriched in WT retina | paired-like homeodomain transcription factor 3 [Source:MGI Symbol;Acc:MGI:1100498] | 41.62 | 5.973 | 4.739E-09 |
| ENSMUSG00000049908 | Gja8 | Enriched in WT retina | gap junction protein, alpha 8 [Source:MGI Symbol;Acc:MGI:99953] | 2039 | 6.476 | 0.00003837 |
| ENSMUSG00000021799 | Opn4 | Enriched in WT retina | opsin 4 (melanopsin) [Source:MGI Symbol;Acc:MGI:1353425] | 133.7 | 6.621 | 8.71E-32 |
| ENSMUSG00000026253 | Chrng | Enriched in WT retina | cholinergic receptor, nicotinic, gamma polypeptide [Source:MGI Symbol;Acc:MGI:87895] | 42.78 | 6.977 | 0.008776 |
