## Supplementary material for "Melanopsin regulates axonal translation underlying retinohypothalamic circuit assembly": Table_S2_TRAP_seq_tissue_contrasts_significant_genes

**Contrast numerator: P8_Het_dLGN. Contrast denominator: P8_Het_retina.**

| ensembl_gene_id | description | mgi_symbol | Significance | deseq_logfc | deseq_adjp | deseq_basemean |
|---|---|---|---|---|---|---|
| ENSMUSG00000021685 | orthopedia homeobox [Source:MGI Symbol;Acc:MGI:99835] | Otp | Enriched in dLGN | 19.26 | 0.000000031 | 81.46 |
| ENSMUSG00000032532 | cholecystokinin [Source:MGI Symbol;Acc:MGI:88297] | Cck | Enriched in dLGN | 6.994 | 4.895E-41 | 445.5 |
| ENSMUSG00000027360 | histidine decarboxylase [Source:MGI Symbol;Acc:MGI:96062] | Hdc | Enriched in dLGN | 6.234 | 9.892E-31 | 44.93 |
| ENSMUSG00000078816 | protein kinase C, gamma [Source:MGI Symbol;Acc:MGI:97597] | Prkcg | Enriched in dLGN | 5.684 | 9.174E-51 | 571.8 |
| ENSMUSG00000004892 | brevican [Source:MGI Symbol;Acc:MGI:1096385] | Bcan | Enriched in dLGN | 4.526 | 3.089E-57 | 546.7 |
| ENSMUSG00000062372 | otoferlin [Source:MGI Symbol;Acc:MGI:1891247] | Otof | Enriched in dLGN | 4.384 | 3.198E-20 | 68.05 |
| ENSMUSG00000004366 | somatostatin [Source:MGI Symbol;Acc:MGI:98326] | Sst | Enriched in dLGN | 4.27 | 4.309E-15 | 62.89 |
| ENSMUSG00000010825 | glutamate receptor, ionotropic, delta 2 (Grid2) interacting protein 1 [Source:MGI Symbol;Acc:MGI:2176213] | Grid2ip | Enriched in dLGN | 4.249 | 3.094E-39 | 357.2 |
| ENSMUSG00000068696 | G-protein coupled receptor 88 [Source:MGI Symbol;Acc:MGI:1927653] | Gpr88 | Enriched in dLGN | 4.075 | 1.058E-32 | 66.64 |
| ENSMUSG00000073406 | histocompatibility 2, blastocyst [Source:MGI Symbol;Acc:MGI:892004] | H2-Bl | Enriched in dLGN | 4.01 | 3.066E-07 | 18.45 |
| ENSMUSG00000064179 | troponin T1, skeletal, slow [Source:MGI Symbol;Acc:MGI:1333868] | Tnnt1 | Enriched in dLGN | 3.955 | 9.887E-19 | 308.7 |
| ENSMUSG00000037428 | VGF nerve growth factor inducible [Source:MGI Symbol;Acc:MGI:1343180] | Vgf | Enriched in dLGN | 3.95 | 9.626E-43 | 1249 |
| ENSMUSG00000040181 | flavin containing monooxygenase 1 [Source:MGI Symbol;Acc:MGI:1310002] | Fmo1 | Enriched in dLGN | 3.914 | 3.671E-17 | 18.97 |
| ENSMUSG00000020838 | solute carrier family 6 (neurotransmitter transporter, serotonin), member 4 [Source:MGI Symbol;Acc:MGI:96285] | Slc6a4 | Enriched in dLGN | 3.735 | 4.291E-15 | 23.26 |
| ENSMUSG00000032503 | cyclic AMP-regulated phosphoprotein, 21 [Source:MGI Symbol;Acc:MGI:107562] | Arpp21 | Enriched in dLGN | 3.564 | 6.45E-20 | 594.6 |
| ENSMUSG00000110086 | small integral membrane protein 32 [Source:MGI Symbol;Acc:MGI:5791459] | Smim32 | Enriched in dLGN | 3.525 | 1.236E-17 | 24.37 |
| ENSMUSG00000036856 | wingless-type MMTV integration site family, member 4 [Source:MGI Symbol;Acc:MGI:98957] | Wnt4 | Enriched in dLGN | 3.441 | 1.384E-27 | 80.14 |
| ENSMUSG00000026278 | BCL2-related ovarian killer [Source:MGI Symbol;Acc:MGI:1858494] | Bok | Enriched in dLGN | 3.1 | 2.389E-21 | 219.1 |
| ENSMUSG00000056306 | serine rich and transmembrane domain containing 1 [Source:MGI Symbol;Acc:MGI:3607715] | Sertm1 | Enriched in dLGN | 3.038 | 1.296E-21 | 70.23 |
| ENSMUSG00000068373 | RIKEN cDNA D430041D05 gene [Source:MGI Symbol;Acc:MGI:2181743] | D430041D05Rik | Enriched in dLGN | 2.972 | 3.354E-56 | 780.2 |
| ENSMUSG00000024617 | calcium/calmodulin-dependent protein kinase II alpha [Source:MGI Symbol;Acc:MGI:88256] | Camk2a | Enriched in dLGN | 2.794 | 2.231E-20 | 1425 |
| ENSMUSG00000022112 | glypican 5 [Source:MGI Symbol;Acc:MGI:1194894] | Gpc5 | Enriched in dLGN | 2.687 | 3.572E-08 | 390.3 |
| ENSMUSG00000000794 | potassium intermediate/small conductance calcium-activated channel, subfamily N, member 3 [Source:MGI Symbol;Acc:MGI:2153183] | Kcnn3 | Enriched in dLGN | 2.685 | 2.114E-19 | 249.5 |
| ENSMUSG00000053004 | histamine receptor H1 [Source:MGI Symbol;Acc:MGI:107619] | Hrh1 | Enriched in dLGN | 2.664 | 0.00000378 | 32.83 |
| ENSMUSG00000045201 | leucine rich repeat containing 3B [Source:MGI Symbol;Acc:MGI:2384996] | Lrrc3b | Enriched in dLGN | 2.659 | 7.158E-08 | 34.56 |
| ENSMUSG00000073294 | EZH inhibitory protein [Source:MGI Symbol;Acc:MGI:2147968] | Ezhip | Enriched in dLGN | 2.653 | 0.02067 | 8.404 |
| ENSMUSG00000074923 | p21 (RAC1) activated kinase 6 [Source:MGI Symbol;Acc:MGI:2679420] | Pak6 | Enriched in dLGN | 2.646 | 3.479E-25 | 281.4 |
| ENSMUSG00000022696 | SID1 transmembrane family, member 1 [Source:MGI Symbol;Acc:MGI:2443155] | Sidt1 | Enriched in dLGN | 2.637 | 1.441E-14 | 43.24 |
| ENSMUSG00000039546 | adherens junction associated protein 1 [Source:MGI Symbol;Acc:MGI:2685419] | Ajap1 | Enriched in dLGN | 2.619 | 1.301E-23 | 293.4 |
| ENSMUSG00000063646 | janus kinase and microtubule interacting protein 1 [Source:MGI Symbol;Acc:MGI:1923321] | Jakmip1 | Enriched in dLGN | 2.595 | 2.567E-08 | 47 |
| ENSMUSG00000029516 | citron [Source:MGI Symbol;Acc:MGI:105313] | Cit | Enriched in dLGN | 2.591 | 2.781E-28 | 2284 |
| ENSMUSG00000029471 | calcium/calmodulin-dependent protein kinase kinase 2, beta [Source:MGI Symbol;Acc:MGI:2444812] | Camkk2 | Enriched in dLGN | 2.578 | 1.567E-32 | 438.3 |
| ENSMUSG00000042817 | FMS-like tyrosine kinase 3 [Source:MGI Symbol;Acc:MGI:95559] | Flt3 | Enriched in dLGN | 2.569 | 1.106E-08 | 42.78 |
| ENSMUSG00000028758 | kinesin family member 17 [Source:MGI Symbol;Acc:MGI:1098229] | Kif17 | Enriched in dLGN | 2.536 | 5.512E-23 | 102.3 |
| ENSMUSG00000031137 | fibroblast growth factor 13 [Source:MGI Symbol;Acc:MGI:109178] | Fgf13 | Enriched in dLGN | 2.5 | 1.538E-16 | 419.6 |
| ENSMUSG00000046182 | GSG1-like [Source:MGI Symbol;Acc:MGI:2685483] | Gsg1l | Enriched in dLGN | 2.491 | 6.094E-26 | 471.7 |
| ENSMUSG00000020374 | RasGEF domain family, member 1C [Source:MGI Symbol;Acc:MGI:1921813] | Rasgef1c | Enriched in dLGN | 2.455 | 1.374E-15 | 39.33 |
| ENSMUSG00000010086 | ring finger protein 112 [Source:MGI Symbol;Acc:MGI:106611] | Rnf112 | Enriched in dLGN | 2.45 | 2.893E-14 | 340.8 |
| ENSMUSG00000067028 | contactin associated protein-like 5B [Source:MGI Symbol;Acc:MGI:3664583] | Cntnap5b | Enriched in dLGN | 2.443 | 7.959E-09 | 198.7 |
| ENSMUSG00000079388 | RIKEN cDNA 2610042L04 gene [Source:MGI Symbol;Acc:MGI:1914305] | 2610042L04Rik | Enriched in dLGN | 2.43 | 0.003146 | 4.418 |
| ENSMUSG00000059203 | interleukin 1 receptor accessory protein-like 2 [Source:MGI Symbol;Acc:MGI:1913106] | Il1rapl2 | Enriched in dLGN | 2.392 | 4.156E-07 | 143.9 |
| ENSMUSG00000035355 | potassium voltage-gated channel, subfamily H (eag-related), member 4 [Source:MGI Symbol;Acc:MGI:2156184] | Kcnh4 | Enriched in dLGN | 2.371 | 0.01167 | 7.674 |
| ENSMUSG00000050069 | gremlin 2, DAN family BMP antagonist [Source:MGI Symbol;Acc:MGI:1344367] | Grem2 | Enriched in dLGN | 2.361 | 1.6E-14 | 70.97 |
| ENSMUSG00000008496 | POU domain, class 2, transcription factor 2 [Source:MGI Symbol;Acc:MGI:101897] | Pou2f2 | Enriched in dLGN | 2.35 | 6.248E-13 | 295.3 |
| ENSMUSG00000052026 | solute carrier family 6 (neurotransmitter transporter, L-proline), member 7 [Source:MGI Symbol;Acc:MGI:2147363] | Slc6a7 | Enriched in dLGN | 2.305 | 9.559E-23 | 217.9 |
| ENSMUSG00000055409 | NEL-like 1 [Source:MGI Symbol;Acc:MGI:2443902] | Nell1 | Enriched in dLGN | 2.299 | 5.01E-08 | 543.3 |
| ENSMUSG00000046480 | sodium channel, type IV, beta [Source:MGI Symbol;Acc:MGI:2687406] | Scn4b | Enriched in dLGN | 2.294 | 0.00002944 | 167.6 |
| ENSMUSG00000039278 | proprotein convertase subtilisin/kexin type 1 inhibitor [Source:MGI Symbol;Acc:MGI:1353431] | Pcsk1n | Enriched in dLGN | 2.292 | 8.423E-13 | 2810 |
| ENSMUSG00000018849 | WW, C2 and coiled-coil domain containing 1 [Source:MGI Symbol;Acc:MGI:2388637] | Wwc1 | Enriched in dLGN | 2.285 | 2.364E-32 | 625.4 |
| ENSMUSG00000026564 | dual specificity phosphatase 27 (putative) [Source:MGI Symbol;Acc:MGI:2685055] | Dusp27 | Enriched in dLGN | 2.271 | 5.339E-07 | 48.02 |
| ENSMUSG00000004187 | kinesin family member C2 [Source:MGI Symbol;Acc:MGI:109187] | Kifc2 | Enriched in dLGN | 2.269 | 4.897E-35 | 769 |
| ENSMUSG00000070644 | ethanolamine kinase 2 [Source:MGI Symbol;Acc:MGI:2443760] | Etnk2 | Enriched in dLGN | 2.236 | 2.003E-11 | 35.13 |
| ENSMUSG00000033854 | potassium channel, subfamily K, member 10 [Source:MGI Symbol;Acc:MGI:1919508] | Kcnk10 | Enriched in dLGN | 2.223 | 6.793E-19 | 191.6 |
| ENSMUSG00000021596 | multiple C2 domains, transmembrane 1 [Source:MGI Symbol;Acc:MGI:1926021] | Mctp1 | Enriched in dLGN | 2.196 | 5.468E-10 | 101.7 |
| ENSMUSG00000025854 | FAM20C, golgi associated secretory pathway kinase [Source:MGI Symbol;Acc:MGI:2136853] | Fam20c | Enriched in dLGN | 2.149 | 5.879E-23 | 465.1 |
| ENSMUSG00000039004 | bone morphogenetic protein 6 [Source:MGI Symbol;Acc:MGI:88182] | Bmp6 | Enriched in dLGN | 2.118 | 1.733E-07 | 58.77 |
| ENSMUSG00000001227 | sema domain, transmembrane domain (TM), and cytoplasmic domain, (semaphorin) 6B [Source:MGI Symbol;Acc:MGI:1202889] | Sema6b | Enriched in dLGN | 2.059 | 1.246E-23 | 529.4 |
| ENSMUSG00000034282 | envoplakin [Source:MGI Symbol;Acc:MGI:107507] | Evpl | Enriched in dLGN | 2.041 | 4.822E-11 | 114.4 |
| ENSMUSG00000033022 | cysteine dioxygenase 1, cytosolic [Source:MGI Symbol;Acc:MGI:105925] | Cdo1 | Enriched in dLGN | 2.029 | 8.341E-09 | 55.15 |
| ENSMUSG00000027200 | sema domain, transmembrane domain (TM), and cytoplasmic domain, (semaphorin) 6D [Source:MGI Symbol;Acc:MGI:2387661] | Sema6d | Enriched in dLGN | 2.018 | 2.888E-11 | 731.9 |
| ENSMUSG00000037703 | leucine zipper, putative tumor suppressor family member 3 [Source:MGI Symbol;Acc:MGI:2656976] | Lzts3 | Enriched in dLGN | 1.992 | 3.042E-16 | 572.4 |
| ENSMUSG00000036377 | capping protein inhibiting regulator of actin [Source:MGI Symbol;Acc:MGI:2444817] | Cracd | Enriched in dLGN | 1.992 | 1.078E-14 | 1779 |
| ENSMUSG00000026888 | growth factor receptor bound protein 14 [Source:MGI Symbol;Acc:MGI:1355324] | Grb14 | Enriched in dLGN | 1.982 | 4.115E-08 | 102.4 |
| ENSMUSG00000066129 | kinase non-catalytic C-lobe domain (KIND) containing 1 [Source:MGI Symbol;Acc:MGI:1923734] | Kndc1 | Enriched in dLGN | 1.978 | 2.811E-15 | 714.5 |
| ENSMUSG00000045731 | prepronociceptin [Source:MGI Symbol;Acc:MGI:105308] | Pnoc | Enriched in dLGN | 1.964 | 4.784E-08 | 50.56 |
| ENSMUSG00000028909 | protein tyrosine phosphatase, receptor type, U [Source:MGI Symbol;Acc:MGI:1321151] | Ptpru | Enriched in dLGN | 1.947 | 1.365E-18 | 212.6 |
| ENSMUSG00000043165 | loricrin [Source:MGI Symbol;Acc:MGI:96816] | Lor | Enriched in dLGN | 1.915 | 0.003828 | 17.17 |
| ENSMUSG00000038860 | GTPase activating RANGAP domain-like 3 [Source:MGI Symbol;Acc:MGI:2139309] | Garnl3 | Enriched in dLGN | 1.892 | 3.83E-10 | 238.5 |
| ENSMUSG00000038453 | SRC kinase signaling inhibitor 1 [Source:MGI Symbol;Acc:MGI:1933179] | Srcin1 | Enriched in dLGN | 1.885 | 3.722E-24 | 3243 |
| ENSMUSG00000028007 | sorting nexin 7 [Source:MGI Symbol;Acc:MGI:1923811] | Snx7 | Enriched in dLGN | 1.884 | 1.327E-10 | 54.52 |
| ENSMUSG00000022208 | junctophilin 4 [Source:MGI Symbol;Acc:MGI:2443113] | Jph4 | Enriched in dLGN | 1.883 | 2.994E-13 | 961.5 |
| ENSMUSG00000034353 | receptor (calcitonin) activity modifying protein 1 [Source:MGI Symbol;Acc:MGI:1858418] | Ramp1 | Enriched in dLGN | 1.859 | 0.000006162 | 54.71 |
| ENSMUSG00000047415 | G protein-coupled receptor 68 [Source:MGI Symbol;Acc:MGI:2441763] | Gpr68 | Enriched in dLGN | 1.853 | 7.102E-09 | 34.85 |
| ENSMUSG00000027394 | tubulin tyrosine ligase [Source:MGI Symbol;Acc:MGI:1916987] | Ttl | Enriched in dLGN | 1.851 | 3.372E-12 | 343 |
| ENSMUSG00000029757 | dynein cytoplasmic 1 intermediate chain 1 [Source:MGI Symbol;Acc:MGI:107743] | Dync1i1 | Enriched in dLGN | 1.843 | 0.000001046 | 339.1 |
| ENSMUSG00000000489 | platelet derived growth factor, B polypeptide [Source:MGI Symbol;Acc:MGI:97528] | Pdgfb | Enriched in dLGN | 1.84 | 6.234E-10 | 70.92 |
| ENSMUSG00000039579 | glutamate receptor ionotropic, NMDA3A [Source:MGI Symbol;Acc:MGI:1933206] | Grin3a | Enriched in dLGN | 1.838 | 1.72E-10 | 314.9 |
| ENSMUSG00000019590 | cytochrome b-561 [Source:MGI Symbol;Acc:MGI:103253] | Cyb561 | Enriched in dLGN | 1.813 | 2.923E-08 | 123.2 |
| ENSMUSG00000028876 | Eph receptor A10 [Source:MGI Symbol;Acc:MGI:3586824] | Epha10 | Enriched in dLGN | 1.812 | 8.539E-13 | 202.5 |
| ENSMUSG00000028944 | protein kinase, AMP-activated, gamma 2 non-catalytic subunit [Source:MGI Symbol;Acc:MGI:1336153] | Prkag2 | Enriched in dLGN | 1.801 | 6.308E-11 | 280.9 |
| ENSMUSG00000040794 | C1q and tumor necrosis factor related protein 4 [Source:MGI Symbol;Acc:MGI:1914695] | C1qtnf4 | Enriched in dLGN | 1.795 | 1.517E-12 | 207.8 |
| ENSMUSG00000026347 | transmembrane protein 163 [Source:MGI Symbol;Acc:MGI:1919410] | Tmem163 | Enriched in dLGN | 1.781 | 3.895E-10 | 376 |
| ENSMUSG00000034226 | ras homolog family member V [Source:MGI Symbol;Acc:MGI:2444227] | Rhov | Enriched in dLGN | 1.778 | 3.871E-07 | 54.65 |
| ENSMUSG00000020990 | cyclin-dependent kinase-like 1 (CDC2-related kinase) [Source:MGI Symbol;Acc:MGI:1918341] | Cdkl1 | Enriched in dLGN | 1.778 | 6.088E-07 | 53.67 |
| ENSMUSG00000034664 | integrin alpha 2b [Source:MGI Symbol;Acc:MGI:96601] | Itga2b | Enriched in dLGN | 1.774 | 0.0004057 | 10.87 |
| ENSMUSG00000020732 | RAB37, member RAS oncogene family [Source:MGI Symbol;Acc:MGI:1929945] | Rab37 | Enriched in dLGN | 1.774 | 0.003275 | 50.93 |
| ENSMUSG00000046834 | keratin 1 [Source:MGI Symbol;Acc:MGI:96698] | Krt1 | Enriched in dLGN | 1.76 | 0.00003105 | 16.62 |
| ENSMUSG00000056367 | ARP3 actin-related protein 3B [Source:MGI Symbol;Acc:MGI:2661120] | Actr3b | Enriched in dLGN | 1.758 | 1.006E-17 | 140.3 |
| ENSMUSG00000030685 | potassium channel tetramerisation domain containing 13 [Source:MGI Symbol;Acc:MGI:1923739] | Kctd13 | Enriched in dLGN | 1.752 | 1.286E-11 | 195.9 |
| ENSMUSG00000032492 | parathyroid hormone 1 receptor [Source:MGI Symbol;Acc:MGI:97801] | Pth1r | Enriched in dLGN | 1.743 | 8.197E-08 | 46.94 |
| ENSMUSG00000047746 | F-box protein 40 [Source:MGI Symbol;Acc:MGI:2443753] | Fbxo40 | Enriched in dLGN | 1.743 | 0.003844 | 15.08 |
| ENSMUSG00000042429 | adenosine A1 receptor [Source:MGI Symbol;Acc:MGI:99401] | Adora1 | Enriched in dLGN | 1.741 | 9.8E-09 | 800.2 |
| ENSMUSG00000038244 | microtubule associated monooxygenase, calponin and LIM domain containing 2 [Source:MGI Symbol;Acc:MGI:2444947] | Mical2 | Enriched in dLGN | 1.709 | 1.207E-07 | 255.1 |
| ENSMUSG00000007207 | syntaxin 1A (brain) [Source:MGI Symbol;Acc:MGI:109355] | Stx1a | Enriched in dLGN | 1.705 | 1.747E-09 | 278.7 |
| ENSMUSG00000054013 | transmembrane protein 179 [Source:MGI Symbol;Acc:MGI:2144891] | Tmem179 | Enriched in dLGN | 1.705 | 6.431E-09 | 349.1 |
| ENSMUSG00000022537 | transmembrane protein 44 [Source:MGI Symbol;Acc:MGI:1924489] | Tmem44 | Enriched in dLGN | 1.702 | 4.229E-14 | 164.5 |
| ENSMUSG00000035226 | regulating synaptic membrane exocytosis 4 [Source:MGI Symbol;Acc:MGI:2674366] | Rims4 | Enriched in dLGN | 1.692 | 1.024E-21 | 674.1 |
| ENSMUSG00000037362 | cellular communication network factor 3 [Source:MGI Symbol;Acc:MGI:109185] | Ccn3 | Enriched in dLGN | 1.686 | 0.02191 | 15.41 |
| ENSMUSG00000032890 | regulating synaptic membrane exocytosis 3 [Source:MGI Symbol;Acc:MGI:2443331] | Rims3 | Enriched in dLGN | 1.677 | 6.916E-10 | 1313 |
| ENSMUSG00000027797 | doublecortin-like kinase 1 [Source:MGI Symbol;Acc:MGI:1330861] | Dclk1 | Enriched in dLGN | 1.674 | 2.208E-14 | 1464 |
| ENSMUSG00000024525 | inositol monophosphatase 2 [Source:MGI Symbol;Acc:MGI:2149728] | Impa2 | Enriched in dLGN | 1.669 | 0.000001799 | 23.08 |
| ENSMUSG00000028833 | neurochondrin [Source:MGI Symbol;Acc:MGI:1347351] | Ncdn | Enriched in dLGN | 1.663 | 1.072E-07 | 1887 |
| ENSMUSG00000060275 | neuregulin 2 [Source:MGI Symbol;Acc:MGI:1098246] | Nrg2 | Enriched in dLGN | 1.659 | 0.00009441 | 123.8 |
| ENSMUSG00000042388 | DLG associated protein 3 [Source:MGI Symbol;Acc:MGI:3039563] | Dlgap3 | Enriched in dLGN | 1.654 | 1.409E-12 | 1641 |
| ENSMUSG00000063296 | transmembrane protein 117 [Source:MGI Symbol;Acc:MGI:2444580] | Tmem117 | Enriched in dLGN | 1.654 | 0.00009173 | 103 |
| ENSMUSG00000056413 | ArfGAP with dual PH domains 1 [Source:MGI Symbol;Acc:MGI:2442201] | Adap1 | Enriched in dLGN | 1.653 | 2.476E-20 | 552.2 |
| ENSMUSG00000048385 | scratch family zinc finger 1 [Source:MGI Symbol;Acc:MGI:2176606] | Scrt1 | Enriched in dLGN | 1.653 | 3.423E-09 | 1666 |
| ENSMUSG00000021259 | cytochrome P450, family 46, subfamily a, polypeptide 1 [Source:MGI Symbol;Acc:MGI:1341877] | Cyp46a1 | Enriched in dLGN | 1.641 | 1.109E-12 | 248.2 |
| ENSMUSG00000026380 | transcription factor CP2-like 1 [Source:MGI Symbol;Acc:MGI:2444691] | Tfcp2l1 | Enriched in dLGN | 1.636 | 0.003687 | 32.75 |
| ENSMUSG00000035576 | L3MBTL1 histone methyl-lysine binding protein [Source:MGI Symbol;Acc:MGI:2676663] | L3mbtl1 | Enriched in dLGN | 1.622 | 0.00000439 | 85.42 |
| ENSMUSG00000043424 | eukaryotic translation initiation factor 3, subunit J2 [Source:MGI Symbol;Acc:MGI:3704486] | Eif3j2 | Enriched in dLGN | 1.622 | 0.00003421 | 59.28 |
| ENSMUSG00000038623 | transmembrane 6 superfamily member 1 [Source:MGI Symbol;Acc:MGI:1933209] | Tm6sf1 | Enriched in dLGN | 1.621 | 0.00005544 | 52.29 |
| ENSMUSG00000031963 | BMP-binding endothelial regulator [Source:MGI Symbol;Acc:MGI:1920480] | Bmper | Enriched in dLGN | 1.586 | 0.00004451 | 44.47 |
| ENSMUSG00000027525 | phosphatase and actin regulator 3 [Source:MGI Symbol;Acc:MGI:1921439] | Phactr3 | Enriched in dLGN | 1.585 | 2.06E-17 | 523.5 |
| ENSMUSG00000049252 | low density lipoprotein-related protein 1B [Source:MGI Symbol;Acc:MGI:2151136] | Lrp1b | Enriched in dLGN | 1.574 | 0.000013 | 1165 |
| ENSMUSG00000067242 | leucine-rich repeat LGI family, member 1 [Source:MGI Symbol;Acc:MGI:1861691] | Lgi1 | Enriched in dLGN | 1.567 | 0.000003213 | 72.29 |
| ENSMUSG00000056222 | sparc/osteonectin, cwcv and kazal-like domains proteoglycan 1 [Source:MGI Symbol;Acc:MGI:105371] | Spock1 | Enriched in dLGN | 1.567 | 0.000003374 | 1204 |
| ENSMUSG00000041020 | MAP7 domain containing 2 [Source:MGI Symbol;Acc:MGI:1917474] | Map7d2 | Enriched in dLGN | 1.563 | 3.035E-09 | 736.5 |
| ENSMUSG00000030270 | copine family member IX [Source:MGI Symbol;Acc:MGI:2443052] | Cpne9 | Enriched in dLGN | 1.549 | 0.001168 | 175.2 |
| ENSMUSG00000003872 | lin-7 homolog B, crumbs cell polarity complex component [Source:MGI Symbol;Acc:MGI:1330858] | Lin7b | Enriched in dLGN | 1.544 | 0.00002176 | 29.61 |
| ENSMUSG00000042766 | tripartite motif-containing 46 [Source:MGI Symbol;Acc:MGI:2673000] | Trim46 | Enriched in dLGN | 1.543 | 4.789E-15 | 200.1 |
| ENSMUSG00000022044 | stathmin-like 4 [Source:MGI Symbol;Acc:MGI:1931224] | Stmn4 | Enriched in dLGN | 1.54 | 0.000008537 | 638.3 |
| ENSMUSG00000027584 | opioid receptor-like 1 [Source:MGI Symbol;Acc:MGI:97440] | Oprl1 | Enriched in dLGN | 1.538 | 1.526E-11 | 154 |
| ENSMUSG00000004263 | atrophin 1 [Source:MGI Symbol;Acc:MGI:104725] | Atn1 | Enriched in dLGN | 1.535 | 0.00579 | 25.12 |
| ENSMUSG00000028865 | CD164 sialomucin-like 2 [Source:MGI Symbol;Acc:MGI:1916905] | Cd164l2 | Enriched in dLGN | 1.528 | 0.04088 | 5.202 |
| ENSMUSG00000005045 | chromodomain helicase DNA binding protein 5 [Source:MGI Symbol;Acc:MGI:3036258] | Chd5 | Enriched in dLGN | 1.526 | 6.704E-11 | 1862 |

| | | | | | | |
|---|---|---|---|---|---|---|
| ENSMUSG00000005357 | solute carrier family 1 (high affinity aspartate/glutamate transporter), member 6 [Source:MGI Symbol;Acc:MGI:1096331] | Slc1a6 | Enriched in dLGN | 1.525 | 0.000007296 | 52.8 |
| ENSMUSG00000024907 | galanin and GMAP prepropeptide [Source:MGI Symbol;Acc:MGI:95637] | Gal | Enriched in dLGN | 1.525 | 0.002499 | 30 |
| ENSMUSG00000061911 | myelin transcription factor 1-like [Source:MGI Symbol;Acc:MGI:1100511] | Myt1l | Enriched in dLGN | 1.52 | 1.074E-10 | 1383 |
| ENSMUSG00000039419 | contactin associated protein-like 2 [Source:MGI Symbol;Acc:MGI:1914047] | Cntnap2 | Enriched in dLGN | 1.504 | 0.001635 | 1307 |
| ENSMUSG00000029875 | coiled-coil domain containing 184 [Source:MGI Symbol;Acc:MGI:2146066] | Ccdc184 | Enriched in dLGN | 1.499 | 0.000001139 | 201 |
| ENSMUSG00000039976 | TBC1 domain family, member 16 [Source:MGI Symbol;Acc:MGI:2652878] | Tbc1d16 | Enriched in dLGN | 1.493 | 3.763E-08 | 1281 |
| ENSMUSG00000042807 | HECT, C2 and WW domain containing E3 ubiquitin protein ligase 2 [Source:MGI Symbol;Acc:MGI:2685817] | Hecw2 | Enriched in dLGN | 1.493 | 0.000008802 | 320.7 |
| ENSMUSG00000044813 | src homology 2 domain-containing transforming protein B [Source:MGI Symbol;Acc:MGI:98294] | Shb | Enriched in dLGN | 1.485 | 8.272E-07 | 96.59 |
| ENSMUSG00000022840 | adenylate cyclase 5 [Source:MGI Symbol;Acc:MGI:99673] | Adcy5 | Enriched in dLGN | 1.482 | 5.179E-08 | 2027 |
| ENSMUSG00000006218 | family with sequence similarity 131, member C [Source:MGI Symbol;Acc:MGI:2685539] | Fam131c | Enriched in dLGN | 1.479 | 0.0001021 | 42.58 |
| ENSMUSG00000031910 | hyaluronan synthase 3 [Source:MGI Symbol;Acc:MGI:109599] | Has3 | Enriched in dLGN | 1.474 | 0.002412 | 10.81 |
| ENSMUSG00000105867 | predicted gene 42517 [Source:MGI Symbol;Acc:MGI:5662654] | Gm42517 | Enriched in dLGN | 1.47 | 0.000003202 | 89.61 |
| ENSMUSG00000025576 | RNA binding protein, fox-1 homolog (C. elegans) 3 [Source:MGI Symbol;Acc:MGI:106368] | Rbfox3 | Enriched in dLGN | 1.464 | 3.173E-09 | 388.1 |
| ENSMUSG00000005716 | parvalbumin [Source:MGI Symbol;Acc:MGI:97821] | Pvalb | Enriched in dLGN | 1.44 | 0.04758 | 26.7 |
| ENSMUSG00000062184 | heparan sulfate 6-O-sulfotransferase 2 [Source:MGI Symbol;Acc:MGI:1354959] | Hs6st2 | Enriched in dLGN | 1.434 | 0.00001537 | 235.4 |
| ENSMUSG00000029330 | CDP-diacylglycerol synthase 1 [Source:MGI Symbol;Acc:MGI:1921846] | Cds1 | Enriched in dLGN | 1.424 | 2.172E-09 | 337.7 |
| ENSMUSG00000041115 | IQ motif and Sec7 domain 2 [Source:MGI Symbol;Acc:MGI:3528396] | Iqsec2 | Enriched in dLGN | 1.42 | 1.892E-15 | 368.7 |
| ENSMUSG00000030089 | solute carrier family 41, member 3 [Source:MGI Symbol;Acc:MGI:1918949] | Slc41a3 | Enriched in dLGN | 1.419 | 0.0001597 | 52.15 |
| ENSMUSG00000019194 | sodium channel, voltage-gated, type I, beta [Source:MGI Symbol;Acc:MGI:98247] | Scn1b | Enriched in dLGN | 1.411 | 0.00001219 | 470 |
| ENSMUSG00000046793 | G protein-coupled receptor 61 [Source:MGI Symbol;Acc:MGI:2441719] | Gpr61 | Enriched in dLGN | 1.392 | 0.00004573 | 53.44 |
| ENSMUSG00000031760 | metallothionein 3 [Source:MGI Symbol;Acc:MGI:97173] | Mt3 | Enriched in dLGN | 1.389 | 0.0003196 | 131.4 |
| ENSMUSG00000036564 | N-myc downstream regulated gene 4 [Source:MGI Symbol;Acc:MGI:2384590] | Ndrg4 | Enriched in dLGN | 1.387 | 2.654E-07 | 2611 |
| ENSMUSG00000051146 | calcium/calmodulin-dependent protein kinase II inhibitor 2 [Source:MGI Symbol;Acc:MGI:1920297] | Camk2n2 | Enriched in dLGN | 1.383 | 6.085E-10 | 513.1 |
| ENSMUSG00000035653 | leucine rich repeat and fibronectin type III domain containing 5 [Source:MGI Symbol;Acc:MGI:2144814] | Lrfn5 | Enriched in dLGN | 1.381 | 0.0002812 | 564.4 |
| ENSMUSG00000003273 | carbonic anhydrase 11 [Source:MGI Symbol;Acc:MGI:1336193] | Car11 | Enriched in dLGN | 1.38 | 1.017E-07 | 105.3 |
| ENSMUSG00000028069 | G patch domain containing 4 [Source:MGI Symbol;Acc:MGI:1913864] | Gpatch4 | Enriched in dLGN | 1.378 | 9.96E-10 | 93.76 |
| ENSMUSG00000028339 | collagen, type XV, alpha 1 [Source:MGI Symbol;Acc:MGI:88449] | Col15a1 | Enriched in dLGN | 1.372 | 0.002307 | 38.43 |
| ENSMUSG00000037348 | progestin and adipoQ receptor family member VII [Source:MGI Symbol;Acc:MGI:1919154] | Paqr7 | Enriched in dLGN | 1.367 | 0.000001479 | 163 |
| ENSMUSG00000024524 | guanine nucleotide binding protein, alpha stimulating, olfactory type [Source:MGI Symbol;Acc:MGI:95774] | Gnal | Enriched in dLGN | 1.363 | 3.631E-10 | 618.2 |
| ENSMUSG00000015354 | procollagen C-endopeptidase enhancer 2 [Source:MGI Symbol;Acc:MGI:1923727] | Pcolce2 | Enriched in dLGN | 1.362 | 0.01342 | 23.56 |
| ENSMUSG00000021991 | calcium channel, voltage-dependent, alpha2/delta subunit 3 [Source:MGI Symbol;Acc:MGI:1338890] | Cacna2d3 | Enriched in dLGN | 1.358 | 0.002644 | 366.8 |
| ENSMUSG00000042532 | golgi autoantigen, golgin subfamily a, 7B [Source:MGI Symbol;Acc:MGI:1918396] | Golga7b | Enriched in dLGN | 1.355 | 1.409E-10 | 283.6 |
| ENSMUSG00000053137 | mitogen-activated protein kinase 11 [Source:MGI Symbol;Acc:MGI:1338024] | Mapk11 | Enriched in dLGN | 1.355 | 0.00007913 | 63.14 |
| ENSMUSG00000019990 | phosphodiesterase 7B [Source:MGI Symbol;Acc:MGI:1352752] | Pde7b | Enriched in dLGN | 1.353 | 0.0002698 | 194.2 |
| ENSMUSG00000031104 | RAB33A, member RAS oncogene family [Source:MGI Symbol;Acc:MGI:109493] | Rab33a | Enriched in dLGN | 1.35 | 0.0001692 | 43.76 |
| ENSMUSG00000043811 | reticulon 4 receptor [Source:MGI Symbol;Acc:MGI:2136886] | Rtn4r | Enriched in dLGN | 1.346 | 0.0001772 | 85.48 |
| ENSMUSG00000022577 | lymphocyte antigen 6 complex, locus H [Source:MGI Symbol;Acc:MGI:1346030] | Ly6h | Enriched in dLGN | 1.343 | 0.0001249 | 411.3 |
| ENSMUSG00000027298 | TYRO3 protein tyrosine kinase 3 [Source:MGI Symbol;Acc:MGI:104294] | Tyro3 | Enriched in dLGN | 1.337 | 7.674E-08 | 178.1 |
| ENSMUSG00000020331 | hyperpolarization-activated, cyclic nucleotide-gated K+ 2 [Source:MGI Symbol;Acc:MGI:1298210] | Hcn2 | Enriched in dLGN | 1.322 | 3.699E-09 | 598.2 |
| ENSMUSG00000043051 | disrupted in schizophrenia 1 [Source:MGI Symbol;Acc:MGI:2447658] | Disc1 | Enriched in dLGN | 1.32 | 0.002314 | 34.75 |
| ENSMUSG00000047261 | growth associated protein 43 [Source:MGI Symbol;Acc:MGI:95639] | Gap43 | Enriched in dLGN | 1.319 | 0.000003236 | 876.7 |
| ENSMUSG00000024064 | polypeptide N-acetylgalactosaminyltransferase 14 [Source:MGI Symbol;Acc:MGI:1918935] | Galnt14 | Enriched in dLGN | 1.318 | 0.0003803 | 190 |
| ENSMUSG00000063694 | cytochrome c, somatic [Source:MGI Symbol;Acc:MGI:88578] | Cycs | Enriched in dLGN | 1.314 | 0.00008352 | 116.9 |
| ENSMUSG00000108591 | promoter of Mat2a antisense radiation induced circulating long non-coding RNA [Source:MGI Symbol;Acc:MGI:1925358] | Particl | Enriched in dLGN | 1.31 | 0.00003845 | 62.29 |
| ENSMUSG00000020658 | EFR3 homolog B [Source:MGI Symbol;Acc:MGI:2444851] | Efr3b | Enriched in dLGN | 1.304 | 1.051E-10 | 856.7 |
| ENSMUSG00000042751 | nicotinamide nucleotide adenylyltransferase 2 [Source:MGI Symbol;Acc:MGI:2444155] | Nmnat2 | Enriched in dLGN | 1.304 | 1.716E-07 | 932 |
| ENSMUSG00000019785 | clavesin 2 [Source:MGI Symbol;Acc:MGI:2443223] | Clvs2 | Enriched in dLGN | 1.291 | 0.0000248 | 86.26 |
| ENSMUSG00000050963 | K+ voltage-gated channel, subfamily S, 2 [Source:MGI Symbol;Acc:MGI:1197011] | Kcns2 | Enriched in dLGN | 1.289 | 0.000001487 | 98.76 |
| ENSMUSG00000019966 | kit ligand [Source:MGI Symbol;Acc:MGI:96974] | Kitl | Enriched in dLGN | 1.288 | 0.0007649 | 453.1 |
| ENSMUSG00000035064 | eukaryotic elongation factor-2 kinase [Source:MGI Symbol;Acc:MGI:1195261] | Eef2k | Enriched in dLGN | 1.285 | 7.784E-09 | 118.4 |
| ENSMUSG00000031111 | immunoglobulin superfamily, member 1 [Source:MGI Symbol;Acc:MGI:2147913] | Igsf1 | Enriched in dLGN | 1.274 | 0.007382 | 83.92 |
| ENSMUSG00000025743 | syndecan 3 [Source:MGI Symbol;Acc:MGI:1349163] | Sdc3 | Enriched in dLGN | 1.272 | 0.000002419 | 1949 |
| ENSMUSG00000061702 | transmembrane protein 91 [Source:MGI Symbol;Acc:MGI:2443589] | Tmem91 | Enriched in dLGN | 1.27 | 0.006045 | 41.88 |
| ENSMUSG00000087408 | ceramide synthase 1 [Source:MGI Symbol;Acc:MGI:2136690] | Cers1 | Enriched in dLGN | 1.267 | 3.272E-10 | 94.13 |
| ENSMUSG00000058153 | seizure related 6 homolog like [Source:MGI Symbol;Acc:MGI:1935121] | Sez6l | Enriched in dLGN | 1.259 | 8.579E-07 | 1179 |
| ENSMUSG00000061518 | cytochrome c oxidase subunit 5B [Source:MGI Symbol;Acc:MGI:88475] | Cox5b | Enriched in dLGN | 1.252 | 0.0000272 | 455.8 |
| ENSMUSG00000059518 | zinc finger, HIT domain containing 1 [Source:MGI Symbol;Acc:MGI:1917353] | Znhit1 | Enriched in dLGN | 1.251 | 7.461E-07 | 135.3 |
| ENSMUSG00000053297 | expressed sequence AI854703 [Source:MGI Symbol;Acc:MGI:2141510] | AI854703 | Enriched in dLGN | 1.248 | 0.000002343 | 90.96 |
| ENSMUSG00000028949 | SWI/SNF related, matrix associated, actin dependent regulator of chromatin, subfamily d, member 3 [Source:MGI Symbol;Acc:MGI:19142 | Smarcd3 | Enriched in dLGN | 1.244 | 9.146E-14 | 234.8 |
| ENSMUSG00000044024 | RELT-like 2 [Source:MGI Symbol;Acc:MGI:1918044] | Rell2 | Enriched in dLGN | 1.239 | 5.543E-09 | 285.3 |
| ENSMUSG00000010608 | RNA binding motif protein 25 [Source:MGI Symbol;Acc:MGI:1914289] | Rbm25 | Enriched in dLGN | 1.235 | 0.00008042 | 1039 |
| ENSMUSG00000031906 | sphingomyelin phosphodiesterase 3, neutral [Source:MGI Symbol;Acc:MGI:1927578] | Smpd3 | Enriched in dLGN | 1.234 | 0.00001758 | 615.6 |
| ENSMUSG00000002771 | glutamate receptor, ionotropic, NMDA2D (epsilon 4) [Source:MGI Symbol;Acc:MGI:95823] | Grin2d | Enriched in dLGN | 1.233 | 2.994E-07 | 466.7 |
| ENSMUSG00000079523 | thymosin, beta 10 [Source:MGI Symbol;Acc:MGI:109146] | Tmsb10 | Enriched in dLGN | 1.233 | 0.0008871 | 881.5 |
| ENSMUSG00000025375 | apoptosis-associated tyrosine kinase [Source:MGI Symbol;Acc:MGI:1197518] | Aatk | Enriched in dLGN | 1.229 | 2.086E-07 | 1628 |
| ENSMUSG00000020219 | translocase of inner mitochondrial membrane 13 [Source:MGI Symbol;Acc:MGI:1353432] | Timm13 | Enriched in dLGN | 1.228 | 5.559E-09 | 98.93 |
| ENSMUSG00000020866 | calcium channel, voltage-dependent, T type, alpha 1G subunit [Source:MGI Symbol;Acc:MGI:1201678] | Cacna1g | Enriched in dLGN | 1.221 | 1.317E-08 | 1369 |
| ENSMUSG00000037341 | solute carrier family 9 (sodium/hydrogen exchanger), member 7 [Source:MGI Symbol;Acc:MGI:2444530] | Slc9a7 | Enriched in dLGN | 1.214 | 0.0001056 | 237.7 |
| ENSMUSG00000028532 | cache domain containing 1 [Source:MGI Symbol;Acc:MGI:2444177] | Cachd1 | Enriched in dLGN | 1.208 | 0.000129 | 291.5 |
| ENSMUSG00000024038 | NADH:ubiquinone oxidoreductase core subunit V3 [Source:MGI Symbol;Acc:MGI:1890894] | Ndufv3 | Enriched in dLGN | 1.207 | 0.000001361 | 374.2 |
| ENSMUSG00000036815 | dipeptidylpeptidase 10 [Source:MGI Symbol;Acc:MGI:2442409] | Dpp10 | Enriched in dLGN | 1.205 | 0.004358 | 576.4 |
| ENSMUSG00000028610 | DMRT-like family B with proline-rich C-terminal, 1 [Source:MGI Symbol;Acc:MGI:1927125] | Dmrtb1 | Enriched in dLGN | 1.204 | 0.006926 | 17.02 |
| ENSMUSG00000045045 | leucine rich repeat and fibronectin type III domain containing 4 [Source:MGI Symbol;Acc:MGI:2385612] | Lrfn4 | Enriched in dLGN | 1.203 | 0.000001413 | 212.7 |
| ENSMUSG00000023067 | cyclin-dependent kinase inhibitor 1A (P21) [Source:MGI Symbol;Acc:MGI:104556] | Cdkn1a | Enriched in dLGN | 1.201 | 0.001541 | 78.16 |
| ENSMUSG00000034265 | zinc finger, DHHC domain containing 14 [Source:MGI Symbol;Acc:MGI:2653229] | Zdhhc14 | Enriched in dLGN | 1.191 | 0.002536 | 188.1 |
| ENSMUSG00000061451 | transmembrane protein 151A [Source:MGI Symbol;Acc:MGI:2147713] | Tmem151a | Enriched in dLGN | 1.187 | 0.000002077 | 350.6 |
| ENSMUSG00000066705 | FXYD domain-containing ion transport regulator 6 [Source:MGI Symbol;Acc:MGI:1890226] | Fxyd6 | Enriched in dLGN | 1.182 | 0.01203 | 380.3 |
| ENSMUSG00000023017 | acid-sensing (proton-gated) ion channel 1 [Source:MGI Symbol;Acc:MGI:1194915] | Asic1 | Enriched in dLGN | 1.179 | 0.0000014 | 275.4 |
| ENSMUSG00000000627 | sema domain, immunoglobulin domain (Ig), TM domain, and short cytoplasmic domain [Source:MGI Symbol;Acc:MGI:1340055] | Sema4f | Enriched in dLGN | 1.179 | 0.000009759 | 125.4 |
| ENSMUSG00000032946 | RAS, guanyl releasing protein 2 [Source:MGI Symbol;Acc:MGI:1333849] | Rasgrp2 | Enriched in dLGN | 1.174 | 0.0003598 | 149.8 |
| ENSMUSG00000045348 | neuronal tyrosine-phosphorylated phosphoinositide 3-kinase adaptor 1 [Source:MGI Symbol;Acc:MGI:2443880] | Nyap1 | Enriched in dLGN | 1.169 | 3.578E-08 | 501.1 |
| ENSMUSG00000032024 | CXADR-like membrane protein [Source:MGI Symbol;Acc:MGI:1918816] | Clmp | Enriched in dLGN | 1.163 | 0.00001318 | 151.4 |
| ENSMUSG00000028149 | RAP1, GTP-GDP dissociation stimulator 1 [Source:MGI Symbol;Acc:MGI:2385189] | Rap1gds1 | Enriched in dLGN | 1.149 | 1.416E-07 | 847.3 |
| ENSMUSG00000075702 | selenoprotein M [Source:MGI Symbol;Acc:MGI:2149786] | Selenom | Enriched in dLGN | 1.147 | 0.000076 | 136.2 |
| ENSMUSG00000023827 | 1-acylglycerol-3-phosphate O-acyltransferase 4 (lysophosphatidic acid acyltransferase, delta) [Source:MGI Symbol;Acc:MGI:1915512] | Agpat4 | Enriched in dLGN | 1.145 | 0.000002943 | 114.1 |
| ENSMUSG00000018012 | Rac family small GTPase 3 [Source:MGI Symbol;Acc:MGI:2180784] | Rac3 | Enriched in dLGN | 1.143 | 0.002202 | 122.2 |
| ENSMUSG00000034171 | fatty acid amide hydrolase [Source:MGI Symbol;Acc:MGI:109609] | Faah | Enriched in dLGN | 1.139 | 6.161E-09 | 134 |
| ENSMUSG00000004113 | calcium channel, voltage-dependent, N type, alpha 1B subunit [Source:MGI Symbol;Acc:MGI:88296] | Cacna1b | Enriched in dLGN | 1.136 | 0.000001136 | 1028 |
| ENSMUSG00000035051 | DEAH (Asp-Glu-Ala-Asp/His) box polypeptide 57 [Source:MGI Symbol;Acc:MGI:2147067] | Dhx57 | Enriched in dLGN | 1.134 | 0.000001627 | 248.4 |
| ENSMUSG00000036948 | trafficking protein particle complex 14 [Source:MGI Symbol;Acc:MGI:2385896] | Trappc14 | Enriched in dLGN | 1.127 | 2.192E-08 | 146.1 |
| ENSMUSG00000038296 | polypeptide N-acetylgalactosaminyltransferase 18 [Source:MGI Symbol;Acc:MGI:2446239] | Galnt18 | Enriched in dLGN | 1.123 | 0.009904 | 139.9 |
| ENSMUSG00000033597 | CASK interacting protein 1 [Source:MGI Symbol;Acc:MGI:2442952] | Caskin1 | Enriched in dLGN | 1.121 | 1.023E-08 | 1073 |
| ENSMUSG00000036615 | regulatory factor X-associated protein [Source:MGI Symbol;Acc:MGI:2180854] | Rfxap | Enriched in dLGN | 1.121 | 9.643E-08 | 118.3 |
| ENSMUSG00000046378 | aspartate beta-hydroxylase domain containing 1 [Source:MGI Symbol;Acc:MGI:2685014] | Asphd1 | Enriched in dLGN | 1.117 | 0.002494 | 56.65 |
| ENSMUSG00000047428 | delta like non-canonical Notch ligand 2 [Source:MGI Symbol;Acc:MGI:2146838] | Dlk2 | Enriched in dLGN | 1.115 | 0.0005903 | 45.68 |
| ENSMUSG00000041263 | RUN and SH3 domain containing 1 [Source:MGI Symbol;Acc:MGI:1919546] | Rusc1 | Enriched in dLGN | 1.109 | 0.0001972 | 448.9 |
| ENSMUSG00000015599 | tau tubulin kinase 1 [Source:MGI Symbol;Acc:MGI:2147036] | Ttbk1 | Enriched in dLGN | 1.108 | 4.012E-07 | 1894 |
| ENSMUSG00000055805 | formin-like 1 [Source:MGI Symbol;Acc:MGI:1888994] | Fmnl1 | Enriched in dLGN | 1.106 | 1.898E-07 | 238.1 |
| ENSMUSG00000091337 | EP300 interacting inhibitor of differentiation 1 [Source:MGI Symbol;Acc:MGI:1889651] | Eid1 | Enriched in dLGN | 1.101 | 0.00002179 | 1279 |
| ENSMUSG00000018411 | microtubule-associated protein tau [Source:MGI Symbol;Acc:MGI:97180] | Mapt | Enriched in dLGN | 1.097 | 0.0001531 | 1946 |
| ENSMUSG00000063446 | phospholipid phosphatase related 1 [Source:MGI Symbol;Acc:MGI:2445015] | Plppr1 | Enriched in dLGN | 1.096 | 0.04133 | 101.6 |
| ENSMUSG00000028849 | MAP7 domain containing 1 [Source:MGI Symbol;Acc:MGI:2384297] | Map7d1 | Enriched in dLGN | 1.086 | 6.467E-09 | 1844 |
| ENSMUSG00000027203 | deoxyuridine triphosphatase [Source:MGI Symbol;Acc:MGI:1346051] | Dut | Enriched in dLGN | 1.076 | 0.00000343 | 92.41 |
| ENSMUSG00000075318 | sodium channel, voltage-gated, type II, alpha [Source:MGI Symbol;Acc:MGI:98248] | Scn2a | Enriched in dLGN | 1.074 | 0.000001413 | 1334 |
| ENSMUSG00000018405 | mitochondrial rRNA methyltransferase 1 [Source:MGI Symbol;Acc:MGI:2443470] | Mrm1 | Enriched in dLGN | 1.073 | 0.003089 | 29.07 |
| ENSMUSG00000035640 | calcium channel, voltage-dependent, beta subunit associated regulatory protein [Source:MGI Symbol;Acc:MGI:1354170] | Cbarp | Enriched in dLGN | 1.068 | 0.00004508 | 1169 |
| ENSMUSG00000005986 | ankyrin repeat domain 13 family, member D [Source:MGI Symbol;Acc:MGI:1915673] | Ankrd13d | Enriched in dLGN | 1.067 | 0.000005415 | 148.5 |
| ENSMUSG00000029161 | cell growth regulator with EF hand domain 1 [Source:MGI Symbol;Acc:MGI:1915817] | Cgref1 | Enriched in dLGN | 1.066 | 0.002162 | 115.3 |
| ENSMUSG00000034271 | Jun dimerization protein 2 [Source:MGI Symbol;Acc:MGI:1932093] | Jdp2 | Enriched in dLGN | 1.065 | 8.859E-08 | 161.1 |
| ENSMUSG00000004933 | megakaryocyte-associated tyrosine kinase [Source:MGI Symbol;Acc:MGI:99259] | Matk | Enriched in dLGN | 1.06 | 0.00008251 | 175.5 |
| ENSMUSG00000031561 | teneurin transmembrane protein 3 [Source:MGI Symbol;Acc:MGI:1345183] | Tenm3 | Enriched in dLGN | 1.057 | 0.000004933 | 2385 |
| ENSMUSG00000031841 | cadherin 13 [Source:MGI Symbol;Acc:MGI:99551] | Cdh13 | Enriched in dLGN | 1.054 | 0.0127 | 448.3 |
| ENSMUSG00000029120 | protein phosphatase 2, regulatory subunit B, gamma [Source:MGI Symbol;Acc:MGI:2442660] | Ppp2r2c | Enriched in dLGN | 1.049 | 6.865E-08 | 1258 |
| ENSMUSG00000033316 | polypeptide N-acetylgalactosaminyltransferase 9 [Source:MGI Symbol;Acc:MGI:2677965] | Galnt9 | Enriched in dLGN | 1.048 | 0.00006358 | 204.6 |
| ENSMUSG00000018474 | chromodomain helicase DNA binding protein 3 [Source:MGI Symbol;Acc:MGI:1344395] | Chd3 | Enriched in dLGN | 1.045 | 7.97E-08 | 5030 |
| ENSMUSG00000009394 | synapsin II [Source:MGI Symbol;Acc:MGI:103020] | Syn2 | Enriched in dLGN | 1.042 | 0.0007051 | 611.4 |
| ENSMUSG00000021448 | src homology 2 domain-containing transforming protein C3 [Source:MGI Symbol;Acc:MGI:106179] | Shc3 | Enriched in dLGN | 1.041 | 0.000007498 | 933.3 |
| ENSMUSG00000056124 | UDP-Gal:betaGlcNAc beta 1,4-galactosyltransferase, polypeptide 6 [Source:MGI Symbol;Acc:MGI:1928380] | B4galt6 | Enriched in dLGN | 1.037 | 3.835E-09 | 417.7 |
| ENSMUSG00000058740 | potassium channel, subfamily T, member 1 [Source:MGI Symbol;Acc:MGI:1924627] | Kcnt1 | Enriched in dLGN | 1.037 | 0.00337 | 138.9 |
| ENSMUSG00000041035 | deleted in primary ciliary dyskinesia [Source:MGI Symbol;Acc:MGI:1924407] | Dpcd | Enriched in dLGN | 1.036 | 0.0001344 | 135.1 |
| ENSMUSG00000037990 | SH3 domain containing ring finger 3 [Source:MGI Symbol;Acc:MGI:2444637] | Sh3rf3 | Enriched in dLGN | 1.035 | 0.0001406 | 202.2 |
| ENSMUSG00000044117 | bMERB domain containing 1 [Source:MGI Symbol;Acc:MGI:1914504] | Bmerb1 | Enriched in dLGN | 1.027 | 0.000005587 | 457.3 |
| ENSMUSG00000027220 | synaptotagmin XIII [Source:MGI Symbol;Acc:MGI:1933945] | Syt13 | Enriched in dLGN | 1.025 | 2.029E-07 | 763.5 |
| ENSMUSG00000030303 | fatty acyl CoA reductase 2 [Source:MGI Symbol;Acc:MGI:2687035] | Far2 | Enriched in dLGN | 1.025 | 0.04036 | 96.51 |
| ENSMUSG00000020327 | fibroblast growth factor 22 [Source:MGI Symbol;Acc:MGI:1914362] | Fgf22 | Enriched in dLGN | 1.025 | 0.04874 | 11.32 |
| ENSMUSG00000096847 | transmembrane protein 151B [Source:MGI Symbol;Acc:MGI:2685169] | Tmem151b | Enriched in dLGN | 1.023 | 0.0001425 | 1096 |
| ENSMUSG00000034656 | calcium channel, voltage-dependent, P/Q type, alpha 1A subunit [Source:MGI Symbol;Acc:MGI:109482] | Cacna1a | Enriched in dLGN | 1.022 | 2.039E-07 | 1321 |
| ENSMUSG00000021270 | heat shock protein 90, alpha (cytosolic), class A member 1 [Source:MGI Symbol;Acc:MGI:96250] | Hsp90aa1 | Enriched in dLGN | 1.019 | 0.004114 | 3186 |

| | | | | | | |
|---|---|---|---|---|---|---|
| ENSMUSG00000027134 | lysophosphatidylcholine acyltransferase 4 [Source:MGI Symbol;Acc:MGI:2138993] | Lpcat4 | Enriched in dLGN | 1.016 | 0.000008401 | 130.5 |
| ENSMUSG00000020083 | family with sequence similarity 241, member B [Source:MGI Symbol;Acc:MGI:1917144] | Fam241b | Enriched in dLGN | 1.014 | 0.0002627 | 42.28 |
| ENSMUSG00000045763 | brain abundant, membrane attached signal protein 1 [Source:MGI Symbol;Acc:MGI:1917600] | Basp1 | Enriched in dLGN | 1.011 | 0.00009227 | 3099 |
| ENSMUSG00000027669 | guanine nucleotide binding protein (G protein), beta 4 [Source:MGI Symbol;Acc:MGI:104581] | Gnb4 | Enriched in dLGN | 1.007 | 0.00001195 | 103.7 |
| ENSMUSG00000075590 | nuclear receptor binding protein 2 [Source:MGI Symbol;Acc:MGI:2385017] | Nrbp2 | Enriched in dLGN | 1.005 | 0.00003249 | 333.6 |
| ENSMUSG00000041073 | NAC alpha domain containing [Source:MGI Symbol;Acc:MGI:3603030] | Nacad | Enriched in retina | -1.007 | 0.00002032 | 343.1 |
| ENSMUSG00000002055 | sperm associated antigen 5 [Source:MGI Symbol;Acc:MGI:1927470] | Spag5 | Enriched in retina | -1.015 | 0.01233 | 34.31 |
| ENSMUSG00000010936 | Vac14 homolog (S. cerevisiae) [Source:MGI Symbol;Acc:MGI:2157980] | Vac14 | Enriched in retina | -1.017 | 0.000007118 | 108.7 |
| ENSMUSG00000003746 | mannosidase 1, alpha [Source:MGI Symbol;Acc:MGI:104677] | Man1a | Enriched in retina | -1.023 | 0.007325 | 79.36 |
| ENSMUSG00000025889 | synuclein, alpha [Source:MGI Symbol;Acc:MGI:1277151] | Snca | Enriched in retina | -1.032 | 0.01184 | 118.2 |
| ENSMUSG00000024446 | ribonuclease P 21 subunit [Source:MGI Symbol;Acc:MGI:1914926] | Rpp21 | Enriched in retina | -1.05 | 0.03694 | 21.09 |
| ENSMUSG00000016253 | negative elongation factor complex member C/D, Th1l [Source:MGI Symbol;Acc:MGI:1926424] | Nelfcd | Enriched in retina | -1.075 | 7.939E-07 | 75.48 |
| ENSMUSG00000010066 | calcium channel, voltage-dependent, alpha 2/delta subunit 2 [Source:MGI Symbol;Acc:MGI:1929813] | Cacna2d2 | Enriched in retina | -1.102 | 0.00002907 | 439.4 |
| ENSMUSG00000021728 | embigin [Source:MGI Symbol;Acc:MGI:95321] | Emb | Enriched in retina | -1.108 | 0.006045 | 88.85 |
| ENSMUSG00000049303 | synaptotagmin XII [Source:MGI Symbol;Acc:MGI:2159601] | Syt12 | Enriched in retina | -1.12 | 0.003407 | 74.94 |
| ENSMUSG00000036062 | PHD finger protein 24 [Source:MGI Symbol;Acc:MGI:2140712] | Phf24 | Enriched in retina | -1.146 | 0.000001321 | 370 |
| ENSMUSG00000046321 | heparan sulfate (glucosamine) 3-O-sulfotransferase 2 [Source:MGI Symbol;Acc:MGI:1333802] | Hs3st2 | Enriched in retina | -1.173 | 0.003626 | 41.56 |
| ENSMUSG00000046546 | family with sequence similarity 43, member A [Source:MGI Symbol;Acc:MGI:2676309] | Fam43a | Enriched in retina | -1.177 | 0.003484 | 83.9 |
| ENSMUSG00000050822 | solute carrier family 29 (nucleoside transporters), member 4 [Source:MGI Symbol;Acc:MGI:2385330] | Slc29a4 | Enriched in retina | -1.242 | 0.0005858 | 126.5 |
| ENSMUSG00000023262 | aminoacylase 1 [Source:MGI Symbol;Acc:MGI:87913] | Acy1 | Enriched in retina | -1.311 | 0.01056 | 15.88 |
| ENSMUSG00000041959 | S100 calcium binding protein A10 (calpactin) [Source:MGI Symbol;Acc:MGI:1339468] | S100a10 | Enriched in retina | -1.373 | 0.0268 | 41.51 |
| ENSMUSG00000042115 | kelch domain containing 8A [Source:MGI Symbol;Acc:MGI:2442630] | Klhdc8a | Enriched in retina | -1.397 | 2.497E-07 | 181.2 |
| ENSMUSG00000021070 | bradykinin receptor, beta 2 [Source:MGI Symbol;Acc:MGI:102845] | Bdkrb2 | Enriched in retina | -1.417 | 0.0458 | 6.429 |
| ENSMUSG00000027971 | N-deacetylase/N-sulfotransferase (heparin glucosaminyl) 4 [Source:MGI Symbol;Acc:MGI:1932545] | Ndst4 | Enriched in retina | -1.426 | 0.008732 | 48.89 |
| ENSMUSG00000026463 | ATPase, Ca++ transporting, plasma membrane 4 [Source:MGI Symbol;Acc:MGI:88111] | Atp2b4 | Enriched in retina | -1.46 | 0.00005017 | 561.4 |
| ENSMUSG00000025876 | unc-5 netrin receptor A [Source:MGI Symbol;Acc:MGI:894682] | Unc5a | Enriched in retina | -1.478 | 0.000007498 | 196.1 |
| ENSMUSG00000017897 | EYA transcriptional coactivator and phosphatase 2 [Source:MGI Symbol;Acc:MGI:109341] | Eya2 | Enriched in retina | -1.494 | 0.0002415 | 21.41 |
| ENSMUSG00000044469 | tumor necrosis factor, alpha-induced protein 8-like 1 [Source:MGI Symbol;Acc:MGI:1913693] | Tnfaip8l1 | Enriched in retina | -1.496 | 0.003194 | 9.788 |
| ENSMUSG00000039461 | T cell leukemia translocation altered gene [Source:MGI Symbol;Acc:MGI:1918829] | Tcta | Enriched in retina | -1.519 | 6.112E-08 | 50.88 |
| ENSMUSG00000041577 | proline arginine-rich end leucine-rich repeat [Source:MGI Symbol;Acc:MGI:2151110] | Prelp | Enriched in retina | -1.605 | 0.005827 | 118.3 |
| ENSMUSG00000001794 | calpain, small subunit 1 [Source:MGI Symbol;Acc:MGI:88266] | Capns1 | Enriched in retina | -1.637 | 0.000001097 | 150.2 |
| ENSMUSG00000022861 | diacylglycerol kinase, gamma [Source:MGI Symbol;Acc:MGI:105060] | Dgkg | Enriched in retina | -1.681 | 0.0001605 | 135 |
| ENSMUSG00000020734 | glutamate receptor, ionotropic, NMDA2C (epsilon 3) [Source:MGI Symbol;Acc:MGI:95822] | Grin2c | Enriched in retina | -1.707 | 0.0005617 | 127.2 |
| ENSMUSG00000035407 | KN motif and ankyrin repeat domains 4 [Source:MGI Symbol;Acc:MGI:3043381] | Kank4 | Enriched in retina | -1.789 | 0.001214 | 22.26 |
| ENSMUSG00000006307 | lysine (K)-specific methyltransferase 2B [Source:MGI Symbol;Acc:MGI:109565] | Kmt2b | Enriched in retina | -1.792 | 2.854E-10 | 848.2 |
| ENSMUSG00000006764 | tryptophan hydroxylase 2 [Source:MGI Symbol;Acc:MGI:2651811] | Tph2 | Enriched in retina | -1.794 | 0.03437 | 3.884 |
| ENSMUSG00000033082 | C-type lectin domain family 1, member a [Source:MGI Symbol;Acc:MGI:2444151] | Clec1a | Enriched in retina | -1.832 | 0.008523 | 6.168 |
| ENSMUSG00000001119 | collagen, type VI, alpha 1 [Source:MGI Symbol;Acc:MGI:88459] | Col6a1 | Enriched in retina | -1.912 | 0.0001462 | 82.8 |
| ENSMUSG00000047963 | starch binding domain 1 [Source:MGI Symbol;Acc:MGI:1261768] | Stbd1 | Enriched in retina | -1.926 | 0.00211 | 7.985 |
| ENSMUSG00000039954 | serine/threonine kinase 32A [Source:MGI Symbol;Acc:MGI:2442403] | Stk32a | Enriched in retina | -1.94 | 0.000006111 | 77.08 |
| ENSMUSG00000002633 | sonic hedgehog [Source:MGI Symbol;Acc:MGI:98297] | Shh | Enriched in retina | -1.959 | 0.000005466 | 32.28 |
| ENSMUSG00000030528 | Bloom syndrome, RecQ like helicase [Source:MGI Symbol;Acc:MGI:1328362] | Blm | Enriched in retina | -2.009 | 3.084E-07 | 50.26 |
| ENSMUSG00000027314 | delta like canonical Notch ligand 4 [Source:MGI Symbol;Acc:MGI:1859388] | Dll4 | Enriched in retina | -2.047 | 0.0008383 | 11.29 |
| ENSMUSG00000050296 | ATP-binding cassette, sub-family A (ABC1), member 12 [Source:MGI Symbol;Acc:MGI:2676312] | Abca12 | Enriched in retina | -2.117 | 0.02527 | 4.035 |
| ENSMUSG00000001504 | Iroquois homeobox 2 [Source:MGI Symbol;Acc:MGI:1197526] | Irx2 | Enriched in retina | -2.131 | 0.000009021 | 24.13 |
| ENSMUSG00000024287 | THO complex 1 [Source:MGI Symbol;Acc:MGI:1919668] | Thoc1 | Enriched in retina | -2.202 | 4.296E-08 | 109.9 |
| ENSMUSG00000009734 | POU domain, class 6, transcription factor 2 [Source:MGI Symbol;Acc:MGI:2443631] | Pou6f2 | Enriched in retina | -2.219 | 1.348E-07 | 127.9 |
| ENSMUSG00000060969 | Iroquois homeobox 1 [Source:MGI Symbol;Acc:MGI:1197515] | Irx1 | Enriched in retina | -2.272 | 0.0001385 | 29.96 |
| ENSMUSG00000058925 | coiled-coil domain containing 192 [Source:MGI Symbol;Acc:MGI:1922694] | Ccdc192 | Enriched in retina | -2.274 | 0.0009751 | 7.152 |
| ENSMUSG00000020806 | rhomboid 5 homolog 2 [Source:MGI Symbol;Acc:MGI:2442473] | Rhbdf2 | Enriched in retina | -2.291 | 0.000004648 | 43.1 |
| ENSMUSG00000030110 | ret proto-oncogene [Source:MGI Symbol;Acc:MGI:97902] | Ret | Enriched in retina | -2.355 | 8.797E-13 | 188.3 |
| ENSMUSG00000031734 | Iroquois related homeobox 3 [Source:MGI Symbol;Acc:MGI:1197522] | Irx3 | Enriched in retina | -2.356 | 0.000003691 | 16.38 |
| ENSMUSG00000026778 | protein kinase C, theta [Source:MGI Symbol;Acc:MGI:97601] | Prkcq | Enriched in retina | -2.389 | 4.541E-09 | 111.9 |
| ENSMUSG00000032584 | macrophage stimulating 1 receptor (c-met-related tyrosine kinase) [Source:MGI Symbol;Acc:MGI:99614] | Mst1r | Enriched in retina | -2.462 | 0.0142 | 3.48 |
| ENSMUSG00000061048 | cadherin 3 [Source:MGI Symbol;Acc:MGI:88356] | Cdh3 | Enriched in retina | -2.514 | 0.0003301 | 8.894 |
| ENSMUSG00000041592 | sidekick cell adhesion molecule 2 [Source:MGI Symbol;Acc:MGI:2443847] | Sdk2 | Enriched in retina | -2.565 | 8.776E-21 | 398.3 |
| ENSMUSG00000046719 | neurexophilin 3 [Source:MGI Symbol;Acc:MGI:1336188] | Nxph3 | Enriched in retina | -2.624 | 3.423E-09 | 27.7 |
| ENSMUSG00000078958 | ATPase, H+ transporting, lysosomal accessory protein 1-like [Source:MGI Symbol;Acc:MGI:3648665] | Atp6ap1l | Enriched in retina | -2.633 | 0.00729 | 5.851 |
| ENSMUSG00000038530 | regulator of G-protein signaling 4 [Source:MGI Symbol;Acc:MGI:108409] | Rgs4 | Enriched in retina | -2.679 | 3.43E-10 | 578.5 |
| ENSMUSG00000068220 | lectin, galactose binding, soluble 1 [Source:MGI Symbol;Acc:MGI:96777] | Lgals1 | Enriched in retina | -2.707 | 0.000006043 | 30.47 |
| ENSMUSG00000030830 | integrin alpha L [Source:MGI Symbol;Acc:MGI:96606] | Itgal | Enriched in retina | -2.716 | 0.0004073 | 6.323 |
| ENSMUSG00000025401 | myosin IA [Source:MGI Symbol;Acc:MGI:107732] | Myo1a | Enriched in retina | -2.867 | 0.01534 | 4.176 |
| ENSMUSG00000062168 | protein phosphatase with EF hand calcium-binding domain 1 [Source:MGI Symbol;Acc:MGI:1097157] | Ppef1 | Enriched in retina | -2.88 | 0.001591 | 6.202 |
| ENSMUSG00000012889 | podocan-like 1 [Source:MGI Symbol;Acc:MGI:2685352] | Podnl1 | Enriched in retina | -3.024 | 0.006953 | 3.541 |
| ENSMUSG00000096740 | LBH domain containing 1 [Source:MGI Symbol;Acc:MGI:5516029] | Lbhd1 | Enriched in retina | -3.163 | 0.002118 | 104.6 |
| ENSMUSG00000002980 | basal cell adhesion molecule [Source:MGI Symbol;Acc:MGI:1929940] | Bcam | Enriched in retina | -3.245 | 2.062E-12 | 83.63 |
| ENSMUSG00000019971 | centrosomal protein 290 [Source:MGI Symbol;Acc:MGI:2384917] | Cep290 | Enriched in retina | -3.279 | 4.669E-18 | 749.1 |
| ENSMUSG00000025432 | advillin [Source:MGI Symbol;Acc:MGI:1333798] | Avil | Enriched in retina | -3.385 | 0.001129 | 4.817 |
| ENSMUSG00000031737 | Iroquois homeobox 5 [Source:MGI Symbol;Acc:MGI:1859086] | Irx5 | Enriched in retina | -3.496 | 1.441E-14 | 31.09 |
| ENSMUSG00000056296 | synaptoporin [Source:MGI Symbol;Acc:MGI:1919253] | Synpr | Enriched in retina | -3.623 | 1.026E-18 | 203.6 |
| ENSMUSG00000037962 | refilin A [Source:MGI Symbol;Acc:MGI:1920371] | Rflna | Enriched in retina | -3.785 | 0.00454 | 3.272 |
| ENSMUSG00000023484 | peripherin [Source:MGI Symbol;Acc:MGI:97774] | Prph | Enriched in retina | -4.204 | 0.00000878 | 64.1 |
| ENSMUSG00000035296 | sarcoglycan, gamma (dystrophin-associated glycoprotein) [Source:MGI Symbol;Acc:MGI:1346524] | Sgcg | Enriched in retina | -4.319 | 0.004637 | 9.668 |
| ENSMUSG00000048070 | phosphoinositide-interacting regulator of transient receptor potential channels [Source:MGI Symbol;Acc:MGI:2443635] | Pirt | Enriched in retina | -4.669 | 1.113E-07 | 10.02 |
| ENSMUSG00000034115 | sodium channel, voltage-gated, type XI, alpha [Source:MGI Symbol;Acc:MGI:1345149] | Scn11a | Enriched in retina | -4.69 | 0.00006547 | 5.567 |
| ENSMUSG00000002100 | myosin binding protein C, cardiac [Source:MGI Symbol;Acc:MGI:102844] | Mybpc3 | Enriched in retina | -5.311 | 0.0001172 | 4.298 |
| ENSMUSG00000004098 | collagen, type V, alpha 3 [Source:MGI Symbol;Acc:MGI:1858212] | Col5a3 | Enriched in retina | -5.33 | 4.628E-23 | 67.35 |
| ENSMUSG00000032387 | RNA binding protein with multiple splicing 2 [Source:MGI Symbol;Acc:MGI:1919223] | Rbpms2 | Enriched in retina | -5.67 | 2.729E-22 | 47.72 |
| ENSMUSG00000069372 | cortexin 3 [Source:MGI Symbol;Acc:MGI:3642816] | Ctxn3 | Enriched in retina | -5.678 | 3.611E-10 | 38.01 |
| ENSMUSG00000031586 | RNA binding protein gene with multiple splicing [Source:MGI Symbol;Acc:MGI:1334446] | Rbpms | Enriched in retina | -5.805 | 2.938E-31 | 81.55 |
| ENSMUSG00000051279 | growth differentiation factor 6 [Source:MGI Symbol;Acc:MGI:95689] | Gdf6 | Enriched in retina | -6.941 | 0.000000192 | 11.98 |
| ENSMUSG00000031738 | Iroquois homeobox 6 [Source:MGI Symbol;Acc:MGI:1927642] | Irx6 | Enriched in retina | -7.761 | 3.052E-16 | 50.97 |
| ENSMUSG00000031688 | POU domain, class 4, transcription factor 2 [Source:MGI Symbol;Acc:MGI:102524] | Pou4f2 | Enriched in retina | -8.308 | 6.697E-13 | 45.2 |
| ENSMUSG00000032446 | eomesodermin [Source:MGI Symbol;Acc:MGI:1201683] | Eomes | Enriched in retina | -8.787 | 1.445E-13 | 60.2 |
| ENSMUSG00000021799 | opsin 4 (melanopsin) [Source:MGI Symbol;Acc:MGI:1353425] | Opn4 | Enriched in retina | -9.109 | 3.172E-11 | 25.82 |
| ENSMUSG00000031965 | T-box 20 [Source:MGI Symbol;Acc:MGI:1888496] | Tbx20 | Enriched in retina | -10.6 | 8.339E-15 | 45.45 |

**Contrast numerator: P8_KO_dLGN. Contrast denominator: P8_KO_retina.**

| ensembl_gene_id | description | mgi_symbol | Significance | deseq_logfc | deseq_adjp | deseq_basemean |
|---|---|---|---|---|---|---|
| ENSMUSG00000032532 | cholecystokinin [Source:MGI Symbol;Acc:MGI:88297] | Cck | Enriched in dLGN | 8.027 | 1.242E-45 | 445.5 |
| ENSMUSG00000029155 | spermatogenesis associated 18 [Source:MGI Symbol;Acc:MGI:1920722] | Spata18 | Enriched in dLGN | 7.746 | 9.994E-07 | 10.42 |
| ENSMUSG00000078816 | protein kinase C, gamma [Source:MGI Symbol;Acc:MGI:97597] | Prkcg | Enriched in dLGN | 5.938 | 1.229E-47 | 571.8 |
| ENSMUSG00000027360 | histidine decarboxylase [Source:MGI Symbol;Acc:MGI:96062] | Hdc | Enriched in dLGN | 5.843 | 1.216E-27 | 44.93 |
| ENSMUSG00000024798 | 5-hydroxytryptamine (serotonin) receptor 7 [Source:MGI Symbol;Acc:MGI:99841] | Htr7 | Enriched in dLGN | 5.67 | 1.165E-46 | 109.3 |
| ENSMUSG00000041144 | dynein, axonemal, heavy chain 7B [Source:MGI Symbol;Acc:MGI:2684953] | Dnah7b | Enriched in dLGN | 5.037 | 0.000002143 | 449.4 |
| ENSMUSG00000004892 | brevican [Source:MGI Symbol;Acc:MGI:1096385] | Bcan | Enriched in dLGN | 4.982 | 1.142E-62 | 546.7 |
| ENSMUSG00000048616 | noggin [Source:MGI Symbol;Acc:MGI:104327] | Nog | Enriched in dLGN | 4.662 | 2.383E-33 | 53.91 |
| ENSMUSG00000105594 | predicted gene 43398 [Source:MGI Symbol;Acc:MGI:5663535] | Gm43398 | Enriched in dLGN | 4.549 | 7.215E-25 | 117 |
| ENSMUSG00000036634 | myelin-associated glycoprotein [Source:MGI Symbol;Acc:MGI:96912] | Mag | Enriched in dLGN | 4.539 | 6.904E-12 | 357.6 |
| ENSMUSG00000010825 | glutamate receptor, ionotropic, delta 2 (Grid2) interacting protein 1 [Source:MGI Symbol;Acc:MGI:2176213] | Grid2ip | Enriched in dLGN | 4.519 | 5.917E-40 | 357.2 |
| ENSMUSG00000020838 | solute carrier family 6 (neurotransmitter transporter, serotonin), member 4 [Source:MGI Symbol;Acc:MGI:96285] | Slc6a4 | Enriched in dLGN | 4.509 | 4.448E-16 | 23.26 |
| ENSMUSG00000051864 | TBC1 domain family, member 22a [Source:MGI Symbol;Acc:MGI:1289265] | Tbc1d22a | Enriched in dLGN | 4.473 | 2.387E-09 | 351.4 |
| ENSMUSG00000073406 | histocompatibility 2, blastocyst [Source:MGI Symbol;Acc:MGI:892004] | H2-Bl | Enriched in dLGN | 4.435 | 1.143E-09 | 18.45 |
| ENSMUSG00000068696 | G-protein coupled receptor 88 [Source:MGI Symbol;Acc:MGI:1927653] | Gpr88 | Enriched in dLGN | 4.369 | 8.445E-32 | 66.64 |
| ENSMUSG00000019767 | coiled-coil domain containing 170 [Source:MGI Symbol;Acc:MGI:2685067] | Ccdc170 | Enriched in dLGN | 4.145 | 0.001739 | 16.02 |
| ENSMUSG00000038242 | aldehyde oxidase 4 [Source:MGI Symbol;Acc:MGI:1919122] | Aox4 | Enriched in dLGN | 4.128 | 1.075E-08 | 8.603 |
| ENSMUSG00000037428 | VGF nerve growth factor inducible [Source:MGI Symbol;Acc:MGI:1343180] | Vgf | Enriched in dLGN | 4.093 | 1.32E-42 | 1249 |
| ENSMUSG00000110086 | small integral membrane protein 32 [Source:MGI Symbol;Acc:MGI:5791459] | Smim32 | Enriched in dLGN | 4.043 | 1.081E-23 | 24.37 |
| ENSMUSG00000026278 | BCL2-related ovarian killer [Source:MGI Symbol;Acc:MGI:1858494] | Bok | Enriched in dLGN | 4.039 | 6.348E-33 | 219.1 |
| ENSMUSG00000041688 | angiomotin [Source:MGI Symbol;Acc:MGI:108440] | Amot | Enriched in dLGN | 3.987 | 3.672E-16 | 327.4 |
| ENSMUSG00000052372 | interleukin 1 receptor accessory protein-like 1 [Source:MGI Symbol;Acc:MGI:2687319] | Il1rapl1 | Enriched in dLGN | 3.964 | 3.262E-10 | 1897 |
| ENSMUSG00000033350 | carbohydrate sulfotransferase 2 [Source:MGI Symbol;Acc:MGI:1891160] | Chst2 | Enriched in dLGN | 3.914 | 3.412E-34 | 1902 |
| ENSMUSG00000036856 | wingless-type MMTV integration site family, member 4 [Source:MGI Symbol;Acc:MGI:98957] | Wnt4 | Enriched in dLGN | 3.84 | 2.037E-29 | 80.14 |
| ENSMUSG00000032503 | cyclic AMP-regulated phosphoprotein, 21 [Source:MGI Symbol;Acc:MGI:107562] | Arpp21 | Enriched in dLGN | 3.754 | 3.147E-20 | 594.6 |
| ENSMUSG00000062372 | otoferlin [Source:MGI Symbol;Acc:MGI:1891247] | Otof | Enriched in dLGN | 3.649 | 1.497E-11 | 68.05 |
| ENSMUSG00000108601 | predicted gene 44645 [Source:MGI Symbol;Acc:MGI:5753221] | Gm44645 | Enriched in dLGN | 3.593 | 4.108E-19 | 130.8 |
| ENSMUSG00000030340 | sodium channel, nonvoltage-gated 1 alpha [Source:MGI Symbol;Acc:MGI:101782] | Scnn1a | Enriched in dLGN | 3.58 | 3.781E-18 | 44.62 |
| ENSMUSG00000028744 | solute carrier family 66 member 1 [Source:MGI Symbol;Acc:MGI:2384837] | Slc66a1 | Enriched in dLGN | 3.461 | 3.598E-08 | 195.7 |
| ENSMUSG00000027344 | fibrous sheath-interacting protein 1 [Source:MGI Symbol;Acc:MGI:1918563] | Fsip1 | Enriched in dLGN | 3.409 | 1.705E-10 | 17.62 |
| ENSMUSG00000056306 | serine rich and transmembrane domain containing 1 [Source:MGI Symbol;Acc:MGI:3607715] | Sertm1 | Enriched in dLGN | 3.398 | 7.212E-23 | 70.23 |
| ENSMUSG00000063239 | glutamate receptor, metabotropic 4 [Source:MGI Symbol;Acc:MGI:1351341] | Grm4 | Enriched in dLGN | 3.291 | 6.825E-47 | 424.8 |
| ENSMUSG00000024937 | EH domain binding protein 1-like 1 [Source:MGI Symbol;Acc:MGI:3612340] | Ehbp1l1 | Enriched in dLGN | 3.291 | 1.087E-25 | 192.9 |
| ENSMUSG00000040181 | flavin containing monooxygenase 1 [Source:MGI Symbol;Acc:MGI:1310002] | Fmo1 | Enriched in dLGN | 3.254 | 8.774E-09 | 18.97 |
| ENSMUSG00000032965 | intraflagellar transport 57 [Source:MGI Symbol;Acc:MGI:1921166] | Ift57 | Enriched in dLGN | 3.245 | 2.242E-49 | 254.5 |
| ENSMUSG00000012017 | scavenger receptor class F, member 2 [Source:MGI Symbol;Acc:MGI:1858430] | Scarf2 | Enriched in dLGN | 3.211 | 9.805E-21 | 58.27 |
| ENSMUSG00000000560 | gamma-aminobutyric acid (GABA) A receptor, subunit alpha 2 [Source:MGI Symbol;Acc:MGI:95614] | Gabra2 | Enriched in dLGN | 3.182 | 1.025E-14 | 394.2 |
| ENSMUSG00000096257 | coiled-coil glutamate-rich protein 2 [Source:MGI Symbol;Acc:MGI:3645242] | Ccer2 | Enriched in dLGN | 3.15 | 1.13E-08 | 12.8 |
| ENSMUSG00000046182 | GSG1-like [Source:MGI Symbol;Acc:MGI:2685483] | Gsg1l | Enriched in dLGN | 3.143 | 5.972E-38 | 471.7 |
| ENSMUSG00000027716 | transient receptor potential cation channel, subfamily C, member 3 [Source:MGI Symbol;Acc:MGI:109526] | Trpc3 | Enriched in dLGN | 3.14 | 8.838E-28 | 212.4 |
| ENSMUSG00000039115 | integrin alpha 9 [Source:MGI Symbol;Acc:MGI:104756] | Itga9 | Enriched in dLGN | 3.125 | 0.001 | 1811 |
| ENSMUSG00000027104 | activating transcription factor 2 [Source:MGI Symbol;Acc:MGI:109349] | Atf2 | Enriched in dLGN | 3.098 | 7.35E-12 | 703.5 |
| ENSMUSG00000020902 | netrin 1 [Source:MGI Symbol;Acc:MGI:105088] | Ntn1 | Enriched in dLGN | 3.083 | 1.712E-11 | 66.38 |
| ENSMUSG00000060882 | potassium voltage-gated channel, Shal-related family, member 2 [Source:MGI Symbol;Acc:MGI:102663] | Kcnd2 | Enriched in dLGN | 3.06 | 3.611E-12 | 981.4 |
| ENSMUSG00000065968 | interferon induced transmembrane protein 7 [Source:MGI Symbol;Acc:MGI:1921732] | Ifitm7 | Enriched in dLGN | 3.026 | 6.887E-07 | 15.26 |
| ENSMUSG00000117286 | predicted gene 1043 [Source:MGI Symbol;Acc:MGI:2685889] | Gm1043 | Enriched in dLGN | 3.023 | 2.894E-21 | 237.8 |
| ENSMUSG00000070644 | ethanolamine kinase 2 [Source:MGI Symbol;Acc:MGI:2443760] | Etnk2 | Enriched in dLGN | 2.974 | 4.543E-17 | 35.13 |
| ENSMUSG00000079184 | M-phase phosphoprotein 8 [Source:MGI Symbol;Acc:MGI:1922589] | Mphosph8 | Enriched in dLGN | 2.952 | 3.749E-12 | 685.9 |
| ENSMUSG00000039546 | adherens junction associated protein 1 [Source:MGI Symbol;Acc:MGI:2685419] | Ajap1 | Enriched in dLGN | 2.942 | 1.131E-27 | 293.4 |
| ENSMUSG00000074652 | myosin, heavy chain 7B, cardiac muscle, beta [Source:MGI Symbol;Acc:MGI:3710243] | Myh7b | Enriched in dLGN | 2.924 | 6.097E-16 | 65.13 |
| ENSMUSG00000068373 | RIKEN cDNA D430041D05 gene [Source:MGI Symbol;Acc:MGI:2181743] | D430041D05Rik | Enriched in dLGN | 2.905 | 6.329E-49 | 780.2 |
| ENSMUSG00000028758 | kinesin family member 17 [Source:MGI Symbol;Acc:MGI:1098229] | Kif17 | Enriched in dLGN | 2.888 | 5.767E-27 | 102.3 |
| ENSMUSG00000036377 | capping protein inhibiting regulator of actin [Source:MGI Symbol;Acc:MGI:2444817] | Cracd | Enriched in dLGN | 2.887 | 5.593E-28 | 1779 |
| ENSMUSG00000042817 | FMS-like tyrosine kinase 3 [Source:MGI Symbol;Acc:MGI:95559] | Flt3 | Enriched in dLGN | 2.814 | 1.545E-08 | 42.78 |
| ENSMUSG00000029471 | calcium/calmodulin-dependent protein kinase kinase 2, beta [Source:MGI Symbol;Acc:MGI:2444812] | Camkk2 | Enriched in dLGN | 2.794 | 5.285E-35 | 438.3 |
| ENSMUSG00000052026 | solute carrier family 6 (neurotransmitter transporter, L-proline), member 7 [Source:MGI Symbol;Acc:MGI:2147363] | Slc6a7 | Enriched in dLGN | 2.791 | 3.297E-29 | 217.9 |
| ENSMUSG00000091002 | transcription elongation regulator 1-like [Source:MGI Symbol;Acc:MGI:1917821] | Tcerg1l | Enriched in dLGN | 2.766 | 1.361E-14 | 243.7 |
| ENSMUSG00000045731 | prepronociceptin [Source:MGI Symbol;Acc:MGI:105308] | Pnoc | Enriched in dLGN | 2.766 | 2.934E-13 | 50.56 |
| ENSMUSG00000022112 | glypican 5 [Source:MGI Symbol;Acc:MGI:1194894] | Gpc5 | Enriched in dLGN | 2.747 | 8.28E-08 | 390.3 |
| ENSMUSG00000007594 | hyaluronan and proteoglycan link protein 4 [Source:MGI Symbol;Acc:MGI:2679531] | Hapln4 | Enriched in dLGN | 2.719 | 0.000002423 | 156.8 |
| ENSMUSG00000055078 | gamma-aminobutyric acid (GABA) A receptor, subunit alpha 5 [Source:MGI Symbol;Acc:MGI:95617] | Gabra5 | Enriched in dLGN | 2.711 | 2.327E-09 | 60.37 |
| ENSMUSG00000029516 | citron [Source:MGI Symbol;Acc:MGI:105313] | Cit | Enriched in dLGN | 2.695 | 3.311E-28 | 2284 |
| ENSMUSG00000090053 | paralemmin A kinase anchor protein [Source:MGI Symbol;Acc:MGI:5141924] | Pakap | Enriched in dLGN | 2.69 | 0.0006887 | 58.1 |
| ENSMUSG00000040794 | C1q and tumor necrosis factor related protein 4 [Source:MGI Symbol;Acc:MGI:1914695] | C1qtnf4 | Enriched in dLGN | 2.648 | 2.476E-24 | 207.8 |
| ENSMUSG00000024109 | neurexin I [Source:MGI Symbol;Acc:MGI:1096391] | Nrxn1 | Enriched in dLGN | 2.634 | 2.087E-16 | 3720 |
| ENSMUSG00000031910 | hyaluronan synthase 3 [Source:MGI Symbol;Acc:MGI:109599] | Has3 | Enriched in dLGN | 2.634 | 9.284E-07 | 10.81 |
| ENSMUSG00000034282 | envoplakin [Source:MGI Symbol;Acc:MGI:107507] | Evpl | Enriched in dLGN | 2.629 | 5.838E-17 | 114.4 |
| ENSMUSG00000033022 | cysteine dioxygenase 1, cytosolic [Source:MGI Symbol;Acc:MGI:105925] | Cdo1 | Enriched in dLGN | 2.623 | 4.606E-12 | 55.15 |
| ENSMUSG00000063646 | janus kinase and microtubule interacting protein 1 [Source:MGI Symbol;Acc:MGI:1923321] | Jakmip1 | Enriched in dLGN | 2.621 | 0.000000189 | 47 |
| ENSMUSG00000025854 | FAM20C, golgi associated secretory pathway kinase [Source:MGI Symbol;Acc:MGI:2136853] | Fam20c | Enriched in dLGN | 2.596 | 2.604E-30 | 465.1 |
| ENSMUSG00000046480 | sodium channel, type IV, beta [Source:MGI Symbol;Acc:MGI:2687406] | Scn4b | Enriched in dLGN | 2.595 | 0.000002978 | 167.6 |
| ENSMUSG00000030592 | ryanodine receptor 1, skeletal muscle [Source:MGI Symbol;Acc:MGI:99659] | Ryr1 | Enriched in dLGN | 2.57 | 0.000001835 | 72.59 |
| ENSMUSG00000025856 | platelet derived growth factor, alpha [Source:MGI Symbol;Acc:MGI:97527] | Pdgfa | Enriched in dLGN | 2.552 | 4.533E-26 | 235.2 |
| ENSMUSG00000024617 | calcium/calmodulin-dependent protein kinase II alpha [Source:MGI Symbol;Acc:MGI:88256] | Camk2a | Enriched in dLGN | 2.546 | 1.345E-15 | 1425 |
| ENSMUSG00000066129 | kinase non-catalytic C-lobe domain (KIND) containing 1 [Source:MGI Symbol;Acc:MGI:1923734] | Kndc1 | Enriched in dLGN | 2.543 | 1.069E-22 | 714.5 |
| ENSMUSG00000022408 | family with sequence similarity 83, member F [Source:MGI Symbol;Acc:MGI:2146227] | Fam83f | Enriched in dLGN | 2.536 | 5.519E-07 | 19.97 |
| ENSMUSG00000044933 | somatostatin receptor 3 [Source:MGI Symbol;Acc:MGI:98329] | Sstr3 | Enriched in dLGN | 2.536 | 0.000005522 | 21.77 |
| ENSMUSG00000056258 | potassium voltage-gated channel, subfamily Q, member 3 [Source:MGI Symbol;Acc:MGI:1336181] | Kcnq3 | Enriched in dLGN | 2.529 | 9.562E-15 | 1087 |
| ENSMUSG00000116652 | RIKEN cDNA B830017H08 gene [Source:MGI Symbol;Acc:MGI:3045365] | B830017H08Rik | Enriched in dLGN | 2.524 | 0.000001213 | 15.95 |
| ENSMUSG00000020107 | anaphase promoting complex subunit 16 [Source:MGI Symbol;Acc:MGI:1289325] | Anapc16 | Enriched in dLGN | 2.509 | 5.204E-12 | 134.3 |
| ENSMUSG00000025422 | ArfGAP with GTPase domain, ankyrin repeat and PH domain 2 [Source:MGI Symbol;Acc:MGI:3580016] | Agap2 | Enriched in dLGN | 2.497 | 1.924E-35 | 2518 |
| ENSMUSG00000049112 | oxytocin receptor [Source:MGI Symbol;Acc:MGI:109147] | Oxtr | Enriched in dLGN | 2.493 | 2.883E-09 | 27.31 |
| ENSMUSG00000037362 | cellular communication network factor 3 [Source:MGI Symbol;Acc:MGI:109185] | Ccn3 | Enriched in dLGN | 2.46 | 0.001503 | 15.41 |
| ENSMUSG00000038623 | transmembrane 6 superfamily member 1 [Source:MGI Symbol;Acc:MGI:1933209] | Tm6sf1 | Enriched in dLGN | 2.456 | 1.789E-09 | 52.29 |
| ENSMUSG00000039004 | bone morphogenetic protein 6 [Source:MGI Symbol;Acc:MGI:88182] | Bmp6 | Enriched in dLGN | 2.451 | 7.19E-09 | 58.77 |
| ENSMUSG00000040938 | solute carrier family 16 (monocarboxylic acid transporters), member 11 [Source:MGI Symbol;Acc:MGI:2663709] | Slc16a11 | Enriched in dLGN | 2.436 | 0.0005862 | 9.503 |
| ENSMUSG00000000794 | potassium intermediate/small conductance calcium-activated channel, subfamily N, member 3 [Source:MGI Symbol;Acc:MGI:2153183] | Kcnn3 | Enriched in dLGN | 2.435 | 6.198E-14 | 249.5 |
| ENSMUSG00000041605 | innate immunity activator [Source:MGI Symbol;Acc:MGI:1921579] | Inava | Enriched in dLGN | 2.415 | 0.0000374 | 19.29 |
| ENSMUSG00000052861 | dynein, axonemal, heavy chain 6 [Source:MGI Symbol;Acc:MGI:107744] | Dnah6 | Enriched in dLGN | 2.415 | 0.00008669 | 52.06 |
| ENSMUSG00000033854 | potassium channel, subfamily K, member 10 [Source:MGI Symbol;Acc:MGI:1919508] | Kcnk10 | Enriched in dLGN | 2.396 | 4.532E-19 | 191.6 |
| ENSMUSG00000039278 | proprotein convertase subtilisin/kexin type 1 inhibitor [Source:MGI Symbol;Acc:MGI:1353431] | Pcsk1n | Enriched in dLGN | 2.388 | 6.022E-13 | 2810 |
| ENSMUSG00000018849 | WW, C2 and coiled-coil domain containing 1 [Source:MGI Symbol;Acc:MGI:2388637] | Wwc1 | Enriched in dLGN | 2.363 | 2.424E-32 | 625.4 |
| ENSMUSG00000064202 | spermatogenesis associated 6 like [Source:MGI Symbol;Acc:MGI:1918036] | Spata6l | Enriched in dLGN | 2.357 | 4.075E-08 | 24.41 |
| ENSMUSG00000021270 | heat shock protein 90, alpha (cytosolic), class A member 1 [Source:MGI Symbol;Acc:MGI:96250] | Hsp90aa1 | Enriched in dLGN | 2.354 | 5.5E-12 | 3186 |
| ENSMUSG00000055409 | NEL-like 1 [Source:MGI Symbol;Acc:MGI:2443902] | Nell1 | Enriched in dLGN | 2.353 | 9.378E-08 | 543.3 |
| ENSMUSG00000022454 | NEL-like 2 [Source:MGI Symbol;Acc:MGI:1858510] | Nell2 | Enriched in dLGN | 2.349 | 2.27E-17 | 1350 |
| ENSMUSG00000074923 | p21 (RAC1) activated kinase 6 [Source:MGI Symbol;Acc:MGI:2679420] | Pak6 | Enriched in dLGN | 2.344 | 3.607E-18 | 281.4 |
| ENSMUSG00000037341 | solute carrier family 9 (sodium/hydrogen exchanger), member 7 [Source:MGI Symbol;Acc:MGI:2444530] | Slc9a7 | Enriched in dLGN | 2.34 | 4.864E-14 | 237.7 |
| ENSMUSG00000056305 | ubiquitin specific peptidase 39 [Source:MGI Symbol;Acc:MGI:107622] | Usp39 | Enriched in dLGN | 2.336 | 5.629E-17 | 113.7 |
| ENSMUSG00000004187 | kinesin family member C2 [Source:MGI Symbol;Acc:MGI:109187] | Kifc2 | Enriched in dLGN | 2.316 | 8.507E-34 | 769 |
| ENSMUSG00000043165 | loricrin [Source:MGI Symbol;Acc:MGI:96816] | Lor | Enriched in dLGN | 2.309 | 0.0003631 | 17.17 |
| ENSMUSG00000029875 | coiled-coil domain containing 184 [Source:MGI Symbol;Acc:MGI:2146066] | Ccdc184 | Enriched in dLGN | 2.297 | 3.052E-13 | 201 |
| ENSMUSG00000010086 | ring finger protein 112 [Source:MGI Symbol;Acc:MGI:106611] | Rnf112 | Enriched in dLGN | 2.292 | 1.219E-11 | 340.8 |
| ENSMUSG00000030685 | potassium channel tetramerisation domain containing 13 [Source:MGI Symbol;Acc:MGI:1923739] | Kctd13 | Enriched in dLGN | 2.283 | 7.493E-18 | 195.9 |
| ENSMUSG00000032492 | parathyroid hormone 1 receptor [Source:MGI Symbol;Acc:MGI:97801] | Pth1r | Enriched in dLGN | 2.278 | 1.497E-11 | 46.94 |
| ENSMUSG00000051335 | glucose-fructose oxidoreductase domain containing 1 [Source:MGI Symbol;Acc:MGI:2145304] | Gfod1 | Enriched in dLGN | 2.26 | 1.4E-31 | 253.7 |
| ENSMUSG00000030849 | fibroblast growth factor receptor 2 [Source:MGI Symbol;Acc:MGI:95523] | Fgfr2 | Enriched in dLGN | 2.26 | 0.00009995 | 937.9 |
| ENSMUSG00000019158 | transmembrane protein 160 [Source:MGI Symbol;Acc:MGI:1916344] | Tmem160 | Enriched in dLGN | 2.25 | 7.416E-07 | 90.04 |
| ENSMUSG00000027394 | tubulin tyrosine ligase [Source:MGI Symbol;Acc:MGI:1916987] | Ttl | Enriched in dLGN | 2.232 | 5.197E-16 | 343 |
| ENSMUSG00000048142 | N-acetyltransferase 8-like [Source:MGI Symbol;Acc:MGI:2447776] | Nat8l | Enriched in dLGN | 2.217 | 5.396E-14 | 599.1 |
| ENSMUSG00000028007 | sorting nexin 7 [Source:MGI Symbol;Acc:MGI:1923811] | Snx7 | Enriched in dLGN | 2.216 | 6.378E-12 | 54.52 |
| ENSMUSG00000005357 | solute carrier family 1 (high affinity aspartate/glutamate transporter), member 6 [Source:MGI Symbol;Acc:MGI:1096331] | Slc1a6 | Enriched in dLGN | 2.214 | 4.042E-10 | 52.8 |
| ENSMUSG00000037703 | leucine zipper, putative tumor suppressor family member 3 [Source:MGI Symbol;Acc:MGI:2656976] | Lzts3 | Enriched in dLGN | 2.212 | 1.243E-18 | 572.4 |
| ENSMUSG00000078235 | family with sequence similarity 43, member B [Source:MGI Symbol;Acc:MGI:3651622] | Fam43b | Enriched in dLGN | 2.197 | 6.859E-10 | 60.59 |
| ENSMUSG00000043857 | mannoside acetylglucosaminyltransferase 5, isoenzyme B [Source:MGI Symbol;Acc:MGI:3606200] | Mgat5b | Enriched in dLGN | 2.191 | 6.643E-21 | 479 |
| ENSMUSG00000051331 | calcium channel, voltage-dependent, L type, alpha 1C subunit [Source:MGI Symbol;Acc:MGI:103013] | Cacna1c | Enriched in dLGN | 2.179 | 3.087E-10 | 1223 |
| ENSMUSG00000001227 | sema domain, transmembrane domain (TM), and cytoplasmic domain, (semaphorin) 6B [Source:MGI Symbol;Acc:MGI:1202889] | Sema6b | Enriched in dLGN | 2.172 | 1.649E-24 | 529.4 |
| ENSMUSG00000025468 | calcyon neuron-specific vesicular protein [Source:MGI Symbol;Acc:MGI:1915816] | Caly | Enriched in dLGN | 2.17 | 2.702E-10 | 238.8 |
| ENSMUSG00000105867 | predicted gene 42517 [Source:MGI Symbol;Acc:MGI:5662654] | Gm42517 | Enriched in dLGN | 2.168 | 5.204E-12 | 89.61 |
| ENSMUSG00000021260 | hedgehog interacting protein-like 1 [Source:MGI Symbol;Acc:MGI:1919265] | Hhipl1 | Enriched in dLGN | 2.166 | 3.172E-13 | 71.28 |
| ENSMUSG00000070509 | repulsive guidance molecule family member A [Source:MGI Symbol;Acc:MGI:2679262] | Rgma | Enriched in dLGN | 2.15 | 1.217E-20 | 395.9 |
| ENSMUSG00000046058 | EP300 interacting inhibitor of differentiation 2 [Source:MGI Symbol;Acc:MGI:2681174] | Eid2 | Enriched in dLGN | 2.138 | 5.108E-09 | 137.6 |
| ENSMUSG00000026888 | growth factor receptor bound protein 14 [Source:MGI Symbol;Acc:MGI:1355324] | Grb14 | Enriched in dLGN | 2.135 | 3.277E-08 | 102.4 |
| ENSMUSG00000026347 | transmembrane protein 163 [Source:MGI Symbol;Acc:MGI:1919410] | Tmem163 | Enriched in dLGN | 2.131 | 5.702E-13 | 376 |
| ENSMUSG00000025762 | La ribonucleoprotein 1B [Source:MGI Symbol;Acc:MGI:1914604] | Larp1b | Enriched in dLGN | 2.13 | 6.816E-13 | 100 |
| ENSMUSG00000102189 | predicted gene, 37194 [Source:MGI Symbol;Acc:MGI:5610422] | Gm37194 | Enriched in dLGN | 2.122 | 0.0007265 | 32.7 |

| | | | | | | |
|---|---|---|---|---|---|---|
| ENSMUSG00000046442 | protein phosphatase 1E (PP2C domain containing) [Source:MGI Symbol;Acc:MGI:2444096] | Ppm1e | Enriched in dLGN | 2.116 | 2.204E-17 | 844.2 |
| ENSMUSG00000038860 | GTPase activating RANGAP domain-like 3 [Source:MGI Symbol;Acc:MGI:2139309] | Garnl3 | Enriched in dLGN | 2.11 | 3.08E-11 | 238.5 |
| ENSMUSG00000050069 | gremlin 2, DAN family BMP antagonist [Source:MGI Symbol;Acc:MGI:1344367] | Grem2 | Enriched in dLGN | 2.105 | 1.732E-09 | 70.97 |
| ENSMUSG00000059518 | zinc finger, HIT domain containing 1 [Source:MGI Symbol;Acc:MGI:1917353] | Znhit1 | Enriched in dLGN | 2.1 | 1.457E-16 | 135.3 |
| ENSMUSG00000019590 | cytochrome b-561 [Source:MGI Symbol;Acc:MGI:103253] | Cyb561 | Enriched in dLGN | 2.095 | 7.013E-10 | 123.2 |
| ENSMUSG00000020354 | sarcoglycan, delta (dystrophin-associated glycoprotein) [Source:MGI Symbol;Acc:MGI:1346525] | Sgcd | Enriched in dLGN | 2.095 | 0.00005773 | 245.7 |
| ENSMUSG00000028876 | Eph receptor A10 [Source:MGI Symbol;Acc:MGI:3586824] | Epha10 | Enriched in dLGN | 2.089 | 1.254E-15 | 202.5 |
| ENSMUSG00000030098 | glutamate receptor interacting protein 2 [Source:MGI Symbol;Acc:MGI:2681173] | Grip2 | Enriched in dLGN | 2.087 | 3.742E-22 | 119.7 |
| ENSMUSG00000021792 | peroxiredoxin like 2A [Source:MGI Symbol;Acc:MGI:1917814] | Prxl2a | Enriched in dLGN | 2.084 | 5.016E-25 | 278.9 |
| ENSMUSG00000025318 | junctophilin 3 [Source:MGI Symbol;Acc:MGI:1891497] | Jph3 | Enriched in dLGN | 2.082 | 1.744E-20 | 1064 |
| ENSMUSG00000007207 | syntaxin 1A (brain) [Source:MGI Symbol;Acc:MGI:109355] | Stx1a | Enriched in dLGN | 2.081 | 1.56E-12 | 278.7 |
| ENSMUSG00000056413 | ArfGAP with dual PH domains 1 [Source:MGI Symbol;Acc:MGI:2442201] | Adap1 | Enriched in dLGN | 2.076 | 3.773E-29 | 552.2 |
| ENSMUSG00000078532 | Na+/K+ transporting ATPase interacting 1 [Source:MGI Symbol;Acc:MGI:1914399] | Nkain1 | Enriched in dLGN | 2.076 | 3.728E-13 | 207.8 |
| ENSMUSG00000059203 | interleukin 1 receptor accessory protein-like 2 [Source:MGI Symbol;Acc:MGI:1913106] | Il1rapl2 | Enriched in dLGN | 2.069 | 0.0000569 | 143.9 |
| ENSMUSG00000030270 | copine family member IX [Source:MGI Symbol;Acc:MGI:2443052] | Cpne9 | Enriched in dLGN | 2.063 | 0.00002723 | 175.2 |
| ENSMUSG00000062044 | lemur tyrosine kinase 3 [Source:MGI Symbol;Acc:MGI:3039582] | Lmtk3 | Enriched in dLGN | 2.058 | 1.578E-14 | 2447 |
| ENSMUSG00000026525 | opsin 3 [Source:MGI Symbol;Acc:MGI:1338022] | Opn3 | Enriched in dLGN | 2.053 | 0.0008889 | 17.62 |
| ENSMUSG00000031603 | fibroblast growth factor 20 [Source:MGI Symbol;Acc:MGI:1891346] | Fgf20 | Enriched in dLGN | 2.048 | 0.0009637 | 11.6 |
| ENSMUSG00000034353 | receptor (calcitonin) activity modifying protein 1 [Source:MGI Symbol;Acc:MGI:1858418] | Ramp1 | Enriched in dLGN | 2.043 | 0.00003262 | 54.71 |
| ENSMUSG00000054027 | 5'-nucleotidase domain containing 3 [Source:MGI Symbol;Acc:MGI:3513266] | Nt5dc3 | Enriched in dLGN | 2.042 | 8.286E-12 | 395 |
| ENSMUSG00000034226 | ras homolog family member V [Source:MGI Symbol;Acc:MGI:2444227] | Rhov | Enriched in dLGN | 2.04 | 2.934E-08 | 54.65 |
| ENSMUSG00000055692 | transmembrane protein 191 [Source:MGI Symbol;Acc:MGI:107238] | Tmem191 | Enriched in dLGN | 2.037 | 9.366E-09 | 172.4 |
| ENSMUSG00000046834 | keratin 1 [Source:MGI Symbol;Acc:MGI:96698] | Krt1 | Enriched in dLGN | 2.029 | 0.00001128 | 16.62 |
| ENSMUSG00000028833 | neurochondrin [Source:MGI Symbol;Acc:MGI:1347351] | Ncdn | Enriched in dLGN | 2.026 | 2.78E-10 | 1887 |
| ENSMUSG00000028793 | ring finger protein 19B [Source:MGI Symbol;Acc:MGI:1922484] | Rnf19b | Enriched in dLGN | 2.022 | 5.796E-13 | 239.7 |
| ENSMUSG00000034336 | internexin neuronal intermediate filament protein, alpha [Source:MGI Symbol;Acc:MGI:96568] | Ina | Enriched in dLGN | 2.004 | 1.663E-10 | 1938 |
| ENSMUSG00000075702 | selenoprotein M [Source:MGI Symbol;Acc:MGI:2149786] | Selenom | Enriched in dLGN | 2.001 | 5.196E-12 | 136.2 |
| ENSMUSG00000038260 | transient receptor potential cation channel, subfamily M, member 4 [Source:MGI Symbol;Acc:MGI:1915917] | Trpm4 | Enriched in dLGN | 1.999 | 1.603E-07 | 48.54 |
| ENSMUSG00000021062 | RAB15, member RAS oncogene family [Source:MGI Symbol;Acc:MGI:1916865] | Rab15 | Enriched in dLGN | 1.996 | 1.771E-15 | 344 |
| ENSMUSG00000095098 | coiled-coil domain containing 85B [Source:MGI Symbol;Acc:MGI:2147607] | Ccdc85b | Enriched in dLGN | 1.996 | 1.186E-12 | 384.5 |
| ENSMUSG00000054013 | transmembrane protein 179 [Source:MGI Symbol;Acc:MGI:2144891] | Tmem179 | Enriched in dLGN | 1.995 | 2.937E-11 | 349.1 |
| ENSMUSG00000030706 | mitochondrial ribosomal protein L48 [Source:MGI Symbol;Acc:MGI:1289321] | Mrpl48 | Enriched in dLGN | 1.99 | 4.739E-08 | 153.1 |
| ENSMUSG00000036882 | Rho GTPase activating protein 33 [Source:MGI Symbol;Acc:MGI:2673998] | Arhgap33 | Enriched in dLGN | 1.985 | 5.262E-15 | 502.5 |
| ENSMUSG00000053004 | histamine receptor H1 [Source:MGI Symbol;Acc:MGI:107619] | Hrh1 | Enriched in dLGN | 1.981 | 0.01333 | 32.83 |
| ENSMUSG00000003872 | lin-7 homolog B, crumbs cell polarity complex component [Source:MGI Symbol;Acc:MGI:1330858] | Lin7b | Enriched in dLGN | 1.974 | 2.948E-07 | 29.61 |
| ENSMUSG00000079396 | predicted gene 3411 [Source:MGI Symbol;Acc:MGI:3781589] | Gm3411 | Enriched in dLGN | 1.972 | 0.01034 | 6.451 |
| ENSMUSG00000033467 | cytokine receptor-like factor 2 [Source:MGI Symbol;Acc:MGI:1889506] | Crlf2 | Enriched in dLGN | 1.971 | 1.543E-10 | 87.19 |
| ENSMUSG00000056310 | tRNA-yW synthesizing protein 1 homolog (S. cerevisiae) [Source:MGI Symbol;Acc:MGI:2141161] | Tyw1 | Enriched in dLGN | 1.969 | 1.28E-10 | 150.1 |
| ENSMUSG00000047415 | G protein-coupled receptor 68 [Source:MGI Symbol;Acc:MGI:2441763] | Gpr68 | Enriched in dLGN | 1.964 | 7.234E-08 | 34.85 |
| ENSMUSG00000022623 | SH3 and multiple ankyrin repeat domains 3 [Source:MGI Symbol;Acc:MGI:1930016] | Shank3 | Enriched in dLGN | 1.959 | 7.506E-10 | 4014 |
| ENSMUSG00000030265 | Kirsten rat sarcoma viral oncogene homolog [Source:MGI Symbol;Acc:MGI:96680] | Kras | Enriched in dLGN | 1.958 | 1.254E-15 | 342.3 |
| ENSMUSG00000030089 | solute carrier family 41, member 3 [Source:MGI Symbol;Acc:MGI:1918949] | Slc41a3 | Enriched in dLGN | 1.958 | 3.292E-07 | 52.15 |
| ENSMUSG00000021596 | multiple C2 domains, transmembrane 1 [Source:MGI Symbol;Acc:MGI:1926021] | Mctp1 | Enriched in dLGN | 1.953 | 5.496E-07 | 101.7 |
| ENSMUSG00000029346 | SRR1 domain containing [Source:MGI Symbol;Acc:MGI:1917368] | Srrd | Enriched in dLGN | 1.942 | 1.26E-09 | 45.97 |
| ENSMUSG00000027584 | opioid receptor-like 1 [Source:MGI Symbol;Acc:MGI:97440] | Oprl1 | Enriched in dLGN | 1.936 | 1.94E-15 | 154 |
| ENSMUSG00000042429 | adenosine A1 receptor [Source:MGI Symbol;Acc:MGI:99401] | Adora1 | Enriched in dLGN | 1.935 | 7.524E-10 | 800.2 |
| ENSMUSG00000021259 | cytochrome P450, family 46, subfamily a, polypeptide 1 [Source:MGI Symbol;Acc:MGI:1341877] | Cyp46a1 | Enriched in dLGN | 1.929 | 1.293E-15 | 248.2 |
| ENSMUSG00000048385 | scratch family zinc finger 1 [Source:MGI Symbol;Acc:MGI:2176606] | Scrt1 | Enriched in dLGN | 1.929 | 1.896E-11 | 1666 |
| ENSMUSG00000020736 | 5',3'-nucleotidase, cytosolic [Source:MGI Symbol;Acc:MGI:1354954] | Nt5c | Enriched in dLGN | 1.928 | 1.898E-12 | 100.9 |
| ENSMUSG00000021892 | SH3-domain binding protein 5 (BTK-associated) [Source:MGI Symbol;Acc:MGI:1344391] | Sh3bp5 | Enriched in dLGN | 1.926 | 1.168E-09 | 251.2 |
| ENSMUSG00000049422 | coiled-coil-helix-coiled-coil-helix domain containing 10 [Source:MGI Symbol;Acc:MGI:2143558] | Chchd10 | Enriched in dLGN | 1.921 | 1.43E-08 | 224.5 |
| ENSMUSG00000030889 | von Willebrand factor A domain containing 3A [Source:MGI Symbol;Acc:MGI:3041229] | Vwa3a | Enriched in dLGN | 1.918 | 0.01738 | 26.65 |
| ENSMUSG00000029467 | ATPase, Ca++ transporting, cardiac muscle, slow twitch 2 [Source:MGI Symbol;Acc:MGI:88110] | Atp2a2 | Enriched in dLGN | 1.915 | 1.616E-17 | 2995 |
| ENSMUSG00000035226 | regulating synaptic membrane exocytosis 4 [Source:MGI Symbol;Acc:MGI:2674366] | Rims4 | Enriched in dLGN | 1.912 | 1.489E-25 | 674.1 |
| ENSMUSG00000048537 | pleckstrin homology like domain, family B, member 1 [Source:MGI Symbol;Acc:MGI:2143230] | Phldb1 | Enriched in dLGN | 1.911 | 1.936E-15 | 593 |
| ENSMUSG00000029330 | CDP-diacylglycerol synthase 1 [Source:MGI Symbol;Acc:MGI:1921846] | Cds1 | Enriched in dLGN | 1.909 | 4.201E-15 | 337.7 |
| ENSMUSG00000034472 | RASD family, member 2 [Source:MGI Symbol;Acc:MGI:1922391] | Rasd2 | Enriched in dLGN | 1.909 | 1.694E-09 | 116.4 |
| ENSMUSG00000020431 | adenylate cyclase 1 [Source:MGI Symbol;Acc:MGI:99677] | Adcy1 | Enriched in dLGN | 1.909 | 7.022E-09 | 5232 |
| ENSMUSG00000000489 | platelet derived growth factor, B polypeptide [Source:MGI Symbol;Acc:MGI:97528] | Pdgfb | Enriched in dLGN | 1.907 | 1.729E-09 | 70.92 |
| ENSMUSG00000057897 | calcium/calmodulin-dependent protein kinase II, beta [Source:MGI Symbol;Acc:MGI:88257] | Camk2b | Enriched in dLGN | 1.906 | 5.445E-22 | 2031 |
| ENSMUSG00000041263 | RUN and SH3 domain containing 1 [Source:MGI Symbol;Acc:MGI:1919546] | Rusc1 | Enriched in dLGN | 1.894 | 1.941E-10 | 448.9 |
| ENSMUSG00000039579 | glutamate receptor ionotropic, NMDA3A [Source:MGI Symbol;Acc:MGI:1933206] | Grin3a | Enriched in dLGN | 1.893 | 5.292E-10 | 314.9 |
| ENSMUSG00000038244 | microtubule associated monooxygenase, calponin and LIM domain containing 2 [Source:MGI Symbol;Acc:MGI:2444947] | Mical2 | Enriched in dLGN | 1.891 | 2.163E-08 | 255.1 |
| ENSMUSG00000029419 | apical junction component 1 [Source:MGI Symbol;Acc:MGI:2685842] | Ajm1 | Enriched in dLGN | 1.887 | 5.447E-11 | 326.7 |
| ENSMUSG00000090958 | leucine rich repeat containing 32 [Source:MGI Symbol;Acc:MGI:93882] | Lrrc32 | Enriched in dLGN | 1.884 | 0.00002547 | 22.04 |
| ENSMUSG00000020993 | trafficking protein particle complex 6B [Source:MGI Symbol;Acc:MGI:1925482] | Trappc6b | Enriched in dLGN | 1.867 | 3.401E-08 | 162.5 |
| ENSMUSG00000052253 | zinc finger protein 622 [Source:MGI Symbol;Acc:MGI:1289282] | Zfp622 | Enriched in dLGN | 1.866 | 7.251E-14 | 157.1 |
| ENSMUSG00000009292 | transient receptor potential cation channel, subfamily M, member 2 [Source:MGI Symbol;Acc:MGI:1351901] | Trpm2 | Enriched in dLGN | 1.865 | 3.772E-11 | 209.9 |
| ENSMUSG00000062184 | heparan sulfate 6-O-sulfotransferase 2 [Source:MGI Symbol;Acc:MGI:1354959] | Hs6st2 | Enriched in dLGN | 1.86 | 5.736E-08 | 235.4 |
| ENSMUSG00000022208 | junctophilin 4 [Source:MGI Symbol;Acc:MGI:2443113] | Jph4 | Enriched in dLGN | 1.856 | 4.465E-12 | 961.5 |
| ENSMUSG00000048281 | deleted in lymphocytic leukemia, 7 [Source:MGI Symbol;Acc:MGI:2447771] | Dleu7 | Enriched in dLGN | 1.856 | 0.001867 | 12.86 |
| ENSMUSG00000028909 | protein tyrosine phosphatase, receptor type, U [Source:MGI Symbol;Acc:MGI:1321151] | Ptpru | Enriched in dLGN | 1.844 | 6.728E-15 | 212.6 |
| ENSMUSG00000050821 | family with sequence similarity 131, member A [Source:MGI Symbol;Acc:MGI:1925658] | Fam131a | Enriched in dLGN | 1.835 | 1.086E-10 | 179.2 |
| ENSMUSG00000022892 | amyloid beta (A4) precursor protein [Source:MGI Symbol;Acc:MGI:88059] | App | Enriched in dLGN | 1.834 | 1.241E-08 | 4592 |
| ENSMUSG00000022840 | adenylate cyclase 5 [Source:MGI Symbol;Acc:MGI:99673] | Adcy5 | Enriched in dLGN | 1.832 | 4.36E-11 | 2027 |
| ENSMUSG00000031026 | tripartite motif-containing 66 [Source:MGI Symbol;Acc:MGI:2152406] | Trim66 | Enriched in dLGN | 1.824 | 1.11E-08 | 221.5 |
| ENSMUSG00000028681 | patched 2 [Source:MGI Symbol;Acc:MGI:1095405] | Ptch2 | Enriched in dLGN | 1.82 | 0.02181 | 9.607 |
| ENSMUSG00000026156 | beta-1,3-glucuronyltransferase 2 (glucuronosyltransferase S) [Source:MGI Symbol;Acc:MGI:2389490] | B3gat2 | Enriched in dLGN | 1.817 | 1.564E-15 | 140.5 |
| ENSMUSG00000046793 | G protein-coupled receptor 61 [Source:MGI Symbol;Acc:MGI:2441719] | Gpr61 | Enriched in dLGN | 1.816 | 7.163E-07 | 53.44 |
| ENSMUSG00000023067 | cyclin-dependent kinase inhibitor 1A (P21) [Source:MGI Symbol;Acc:MGI:104556] | Cdkn1a | Enriched in dLGN | 1.813 | 0.000002226 | 78.16 |
| ENSMUSG00000024907 | galanin and GMAP prepropeptide [Source:MGI Symbol;Acc:MGI:95637] | Gal | Enriched in dLGN | 1.813 | 0.0009828 | 30 |
| ENSMUSG00000038295 | autophagy related 9B [Source:MGI Symbol;Acc:MGI:2685420] | Atg9b | Enriched in dLGN | 1.813 | 0.002133 | 13.88 |
| ENSMUSG00000032946 | RAS, guanyl releasing protein 2 [Source:MGI Symbol;Acc:MGI:1333849] | Rasgrp2 | Enriched in dLGN | 1.81 | 6.221E-08 | 149.8 |
| ENSMUSG00000092167 | predicted gene 3696 [Source:MGI Symbol;Acc:MGI:3781872] | Gm3696 | Enriched in dLGN | 1.806 | 0.03914 | 6.523 |
| ENSMUSG00000058153 | seizure related 6 homolog like [Source:MGI Symbol;Acc:MGI:1935121] | Sez6l | Enriched in dLGN | 1.801 | 4.667E-12 | 1179 |
| ENSMUSG00000037348 | progestin and adipoQ receptor family member VII [Source:MGI Symbol;Acc:MGI:1919154] | Paqr7 | Enriched in dLGN | 1.801 | 1.159E-09 | 163 |
| ENSMUSG00000008475 | actin related protein 2/3 complex, subunit 5 [Source:MGI Symbol;Acc:MGI:1915021] | Arpc5 | Enriched in dLGN | 1.8 | 3.573E-14 | 264.7 |
| ENSMUSG00000030811 | F-box and leucine-rich repeat protein 19 [Source:MGI Symbol;Acc:MGI:3039600] | Fbxl19 | Enriched in dLGN | 1.795 | 1.555E-07 | 714.5 |
| ENSMUSG00000056367 | ARP3 actin-related protein 3B [Source:MGI Symbol;Acc:MGI:2661120] | Actr3b | Enriched in dLGN | 1.789 | 6.129E-15 | 140.3 |
| ENSMUSG00000042766 | tripartite motif-containing 46 [Source:MGI Symbol;Acc:MGI:2673000] | Trim46 | Enriched in dLGN | 1.786 | 4.98E-18 | 200.1 |
| ENSMUSG00000042249 | G protein-coupled receptor kinase 3 [Source:MGI Symbol;Acc:MGI:87941] | Grk3 | Enriched in dLGN | 1.78 | 1.493E-14 | 419.2 |
| ENSMUSG00000020990 | cyclin-dependent kinase-like 1 (CDC2-related kinase) [Source:MGI Symbol;Acc:MGI:1918341] | Cdkl1 | Enriched in dLGN | 1.78 | 0.00001374 | 53.67 |
| ENSMUSG00000038738 | SH3 and multiple ankyrin repeat domains 1 [Source:MGI Symbol;Acc:MGI:3613677] | Shank1 | Enriched in dLGN | 1.777 | 4.571E-11 | 6658 |
| ENSMUSG00000048483 | zinc finger, DHHC-type containing 22 [Source:MGI Symbol;Acc:MGI:2685108] | Zdhhc22 | Enriched in dLGN | 1.777 | 5.623E-10 | 612.1 |
| ENSMUSG00000024873 | cornichon family AMPA receptor auxiliary protein 2 [Source:MGI Symbol;Acc:MGI:1277225] | Cnih2 | Enriched in dLGN | 1.773 | 2.479E-17 | 220.1 |
| ENSMUSG00000041037 | immunity-related GTPase family, Q [Source:MGI Symbol;Acc:MGI:2667176] | Irgq | Enriched in dLGN | 1.771 | 2.688E-13 | 618.2 |
| ENSMUSG00000019785 | clavesin 2 [Source:MGI Symbol;Acc:MGI:2443223] | Clvs2 | Enriched in dLGN | 1.769 | 1.374E-07 | 86.26 |
| ENSMUSG00000019966 | kit ligand [Source:MGI Symbol;Acc:MGI:96974] | Kitl | Enriched in dLGN | 1.767 | 0.000005658 | 453.1 |
| ENSMUSG00000048807 | solute carrier family 35, member E4 [Source:MGI Symbol;Acc:MGI:2144150] | Slc35e4 | Enriched in dLGN | 1.766 | 6.262E-14 | 70.33 |
| ENSMUSG00000032878 | coiled-coil domain containing 85A [Source:MGI Symbol;Acc:MGI:2445069] | Ccdc85a | Enriched in dLGN | 1.766 | 6.338E-08 | 409.4 |
| ENSMUSG00000022696 | SID1 transmembrane family, member 1 [Source:MGI Symbol;Acc:MGI:2443155] | Sidt1 | Enriched in dLGN | 1.754 | 0.00007531 | 43.24 |
| ENSMUSG00000027200 | sema domain, transmembrane domain (TM), and cytoplasmic domain, (semaphorin) 6D [Source:MGI Symbol;Acc:MGI:2387661] | Sema6d | Enriched in dLGN | 1.753 | 5.31E-08 | 731.9 |
| ENSMUSG00000002771 | glutamate receptor, ionotropic, NMDA2D (epsilon 4) [Source:MGI Symbol;Acc:MGI:95823] | Grin2d | Enriched in dLGN | 1.752 | 8.044E-13 | 466.7 |
| ENSMUSG00000022054 | neurofilament, medium polypeptide [Source:MGI Symbol;Acc:MGI:97314] | Nefm | Enriched in dLGN | 1.744 | 0.0005862 | 5610 |
| ENSMUSG00000023868 | phosphodiesterase 10A [Source:MGI Symbol;Acc:MGI:1345143] | Pde10a | Enriched in dLGN | 1.742 | 3.402E-07 | 520.6 |
| ENSMUSG00000046287 | paraneoplastic antigen MA3 [Source:MGI Symbol;Acc:MGI:2180565] | Pnma3 | Enriched in dLGN | 1.737 | 5.606E-07 | 110.9 |
| ENSMUSG00000020953 | cochlin [Source:MGI Symbol;Acc:MGI:1278313] | Coch | Enriched in dLGN | 1.737 | 0.001249 | 80.94 |
| ENSMUSG00000031520 | vascular endothelial growth factor C [Source:MGI Symbol;Acc:MGI:109124] | Vegfc | Enriched in dLGN | 1.735 | 0.0009829 | 30.52 |
| ENSMUSG00000027581 | stathmin-like 3 [Source:MGI Symbol;Acc:MGI:1277137] | Stmn3 | Enriched in dLGN | 1.734 | 6.537E-07 | 1780 |
| ENSMUSG00000051146 | calcium/calmodulin-dependent protein kinase II inhibitor 2 [Source:MGI Symbol;Acc:MGI:1920297] | Camk2n2 | Enriched in dLGN | 1.732 | 4.178E-14 | 513.1 |
| ENSMUSG00000028944 | protein kinase, AMP-activated, gamma 2 non-catalytic subunit [Source:MGI Symbol;Acc:MGI:1336153] | Prkag2 | Enriched in dLGN | 1.73 | 3.278E-09 | 280.9 |
| ENSMUSG00000046791 | ribosomal oxygenase 1 [Source:MGI Symbol;Acc:MGI:1919202] | Riox1 | Enriched in dLGN | 1.73 | 4.387E-09 | 41.2 |
| ENSMUSG00000059974 | neurotrimin [Source:MGI Symbol;Acc:MGI:2446259] | Ntm | Enriched in dLGN | 1.729 | 3.644E-08 | 2316 |
| ENSMUSG00000070858 | predicted gene 1673 [Source:MGI Symbol;Acc:MGI:2686519] | Gm1673 | Enriched in dLGN | 1.729 | 0.000557 | 55.45 |
| ENSMUSG00000091337 | EP300 interacting inhibitor of differentiation 1 [Source:MGI Symbol;Acc:MGI:1889651] | Eid1 | Enriched in dLGN | 1.723 | 3.709E-11 | 1279 |
| ENSMUSG00000039166 | A kinase (PRKA) anchor protein 7 [Source:MGI Symbol;Acc:MGI:1859150] | Akap7 | Enriched in dLGN | 1.721 | 0.000008327 | 348.8 |
| ENSMUSG00000005045 | chromodomain helicase DNA binding protein 5 [Source:MGI Symbol;Acc:MGI:3036258] | Chd5 | Enriched in dLGN | 1.716 | 1.2E-12 | 1862 |
| ENSMUSG00000032890 | regulating synaptic membrane exocytosis 3 [Source:MGI Symbol;Acc:MGI:2443331] | Rims3 | Enriched in dLGN | 1.713 | 1.316E-09 | 1313 |
| ENSMUSG00000024524 | guanine nucleotide binding protein, alpha stimulating, olfactory type [Source:MGI Symbol;Acc:MGI:95774] | Gnal | Enriched in dLGN | 1.708 | 2.72E-14 | 618.2 |
| ENSMUSG00000043811 | reticulon 4 receptor [Source:MGI Symbol;Acc:MGI:2136886] | Rtn4r | Enriched in dLGN | 1.707 | 0.000003453 | 85.48 |
| ENSMUSG00000067028 | contactin associated protein-like 5B [Source:MGI Symbol;Acc:MGI:3664583] | Cntnap5b | Enriched in dLGN | 1.707 | 0.0003109 | 198.7 |
| ENSMUSG00000051359 | neurocalcin delta [Source:MGI Symbol;Acc:MGI:1196326] | Ncald | Enriched in dLGN | 1.705 | 2.443E-15 | 710.7 |
| ENSMUSG00000050541 | adrenergic receptor, alpha 1b [Source:MGI Symbol;Acc:MGI:104774] | Adra1b | Enriched in dLGN | 1.704 | 0.0001243 | 73.77 |
| ENSMUSG00000030411 | NOVA alternative splicing regulator 2 [Source:MGI Symbol;Acc:MGI:104296] | Nova2 | Enriched in dLGN | 1.703 | 1.813E-15 | 858.7 |
| ENSMUSG00000018965 | tyrosine 3-monooxygenase/tryptophan 5-monooxygenase activation protein, eta polypeptide [Source:MGI Symbol;Acc:MGI:109194] | Ywhah | Enriched in dLGN | 1.701 | 0.000002427 | 3239 |
| ENSMUSG00000014786 | solute carrier family 9 (sodium/hydrogen exchanger), member 5 [Source:MGI Symbol;Acc:MGI:2685542] | Slc9a5 | Enriched in dLGN | 1.7 | 0.00004176 | 110.7 |
| ENSMUSG00000038020 | Rap guanine nucleotide exchange factor (GEF)-like 1 [Source:MGI Symbol;Acc:MGI:3611446] | Rapgefl1 | Enriched in dLGN | 1.697 | 1.11E-15 | 205.6 |
| ENSMUSG00000047746 | F-box protein 40 [Source:MGI Symbol;Acc:MGI:2443753] | Fbxo40 | Enriched in dLGN | 1.696 | 0.01748 | 15.08 |

| | | | | | | |
|---|---|---|---|---|---|---|
| ENSMUSG00000004642 | stem-loop binding protein [Source:MGI Symbol;Acc:MGI:108402] | Slbp | Enriched in dLGN | 1.695 | 2.816E-10 | 85.78 |
| ENSMUSG00000039976 | TBC1 domain family, member 16 [Source:MGI Symbol;Acc:MGI:2652878] | Tbc1d16 | Enriched in dLGN | 1.693 | 1.4E-09 | 1281 |
| ENSMUSG00000022577 | lymphocyte antigen 6 complex, locus H [Source:MGI Symbol;Acc:MGI:1346030] | Ly6h | Enriched in dLGN | 1.686 | 0.00000235 | 411.3 |
| ENSMUSG00000061518 | cytochrome c oxidase subunit 5B [Source:MGI Symbol;Acc:MGI:88475] | Cox5b | Enriched in dLGN | 1.685 | 3.118E-08 | 455.8 |
| ENSMUSG00000030704 | RAB6A, member RAS oncogene family [Source:MGI Symbol;Acc:MGI:894313] | Rab6a | Enriched in dLGN | 1.682 | 0.00000133 | 1003 |
| ENSMUSG00000108591 | promoter of Mat2a antisense radiation induced circulating long non-coding RNA [Source:MGI Symbol;Acc:MGI:1925358] | Particl | Enriched in dLGN | 1.678 | 2.223E-07 | 62.29 |
| ENSMUSG00000031353 | retinoblastoma binding protein 7, chromatin remodeling factor [Source:MGI Symbol;Acc:MGI:1194910] | Rbbp7 | Enriched in dLGN | 1.675 | 3.132E-16 | 405.8 |
| ENSMUSG00000019464 | prostaglandin E receptor 1 (subtype EP1) [Source:MGI Symbol;Acc:MGI:97793] | Ptger1 | Enriched in dLGN | 1.675 | 0.002592 | 8.868 |
| ENSMUSG00000038011 | dynein, axonemal, heavy chain 10 [Source:MGI Symbol;Acc:MGI:1860299] | Dnah10 | Enriched in dLGN | 1.675 | 0.002714 | 44.22 |
| ENSMUSG00000004263 | atrophin 1 [Source:MGI Symbol;Acc:MGI:104725] | Atn1 | Enriched in dLGN | 1.672 | 0.001814 | 25.12 |
| ENSMUSG00000020982 | nuclear export mediator factor [Source:MGI Symbol;Acc:MGI:1918305] | Nemf | Enriched in dLGN | 1.664 | 0.00003582 | 270.7 |
| ENSMUSG00000035228 | coiled-coil domain containing 106 [Source:MGI Symbol;Acc:MGI:2385900] | Ccdc106 | Enriched in dLGN | 1.663 | 4.807E-10 | 90.05 |
| ENSMUSG00000087408 | ceramide synthase 1 [Source:MGI Symbol;Acc:MGI:2136690] | Cers1 | Enriched in dLGN | 1.658 | 7.486E-15 | 94.13 |
| ENSMUSG00000031104 | RAB33A, member RAS oncogene family [Source:MGI Symbol;Acc:MGI:109493] | Rab33a | Enriched in dLGN | 1.657 | 0.0000143 | 43.76 |
| ENSMUSG00000036564 | N-myc downstream regulated gene 4 [Source:MGI Symbol;Acc:MGI:2384590] | Ndrg4 | Enriched in dLGN | 1.655 | 2.276E-09 | 2611 |
| ENSMUSG00000027500 | stathmin-like 2 [Source:MGI Symbol;Acc:MGI:98241] | Stmn2 | Enriched in dLGN | 1.654 | 0.00000763 | 1265 |
| ENSMUSG00000031351 | zinc finger protein 185 [Source:MGI Symbol;Acc:MGI:108095] | Zfp185 | Enriched in dLGN | 1.645 | 0.0005422 | 546.2 |
| ENSMUSG00000036615 | regulatory factor X-associated protein [Source:MGI Symbol;Acc:MGI:2180854] | Rfxap | Enriched in dLGN | 1.639 | 3.251E-14 | 118.3 |
| ENSMUSG00000045201 | leucine rich repeat containing 3B [Source:MGI Symbol;Acc:MGI:2384996] | Lrrc3b | Enriched in dLGN | 1.637 | 0.005932 | 34.56 |
| ENSMUSG00000027298 | TYRO3 protein tyrosine kinase 3 [Source:MGI Symbol;Acc:MGI:104294] | Tyro3 | Enriched in dLGN | 1.632 | 1.5E-10 | 178.1 |
| ENSMUSG00000028039 | ephrin A3 [Source:MGI Symbol;Acc:MGI:106644] | Efna3 | Enriched in dLGN | 1.628 | 1.44E-11 | 71.32 |
| ENSMUSG00000025743 | syndecan 3 [Source:MGI Symbol;Acc:MGI:1349163] | Sdc3 | Enriched in dLGN | 1.627 | 3.488E-09 | 1949 |
| ENSMUSG00000053297 | expressed sequence AI854703 [Source:MGI Symbol;Acc:MGI:2141510] | AI854703 | Enriched in dLGN | 1.624 | 3.706E-09 | 90.96 |
| ENSMUSG00000029364 | WD repeat and SOCS box-containing 2 [Source:MGI Symbol;Acc:MGI:2144041] | Wsb2 | Enriched in dLGN | 1.618 | 8.145E-09 | 865.5 |
| ENSMUSG00000021278 | amnionless [Source:MGI Symbol;Acc:MGI:1934943] | Amn | Enriched in dLGN | 1.618 | 0.001777 | 16.05 |
| ENSMUSG00000038552 | fibronectin type III domain containing 4 [Source:MGI Symbol;Acc:MGI:1917195] | Fndc4 | Enriched in dLGN | 1.613 | 4.279E-16 | 238.1 |
| ENSMUSG00000067629 | synaptic Ras GTPase activating protein 1 homolog (rat) [Source:MGI Symbol;Acc:MGI:3039785] | Syngap1 | Enriched in dLGN | 1.611 | 1.285E-24 | 1366 |
| ENSMUSG00000015354 | procollagen C-endopeptidase enhancer 2 [Source:MGI Symbol;Acc:MGI:1923727] | Pcolce2 | Enriched in dLGN | 1.611 | 0.00533 | 23.56 |
| ENSMUSG00000003863 | protein tyrosine phosphatase, receptor type, f polypeptide (PTPRF), interacting protein (liprin), alpha 3 [Source:MGI Symbol;Acc:MGI:1924037] | Ppfia3 | Enriched in dLGN | 1.607 | 9.472E-18 | 864.8 |
| ENSMUSG00000022940 | phosphatidylinositol glycan anchor biosynthesis, class P [Source:MGI Symbol;Acc:MGI:1860433] | Pigp | Enriched in dLGN | 1.607 | 0.0008085 | 39.52 |
| ENSMUSG00000061911 | myelin transcription factor 1-like [Source:MGI Symbol;Acc:MGI:1100511] | Myt1l | Enriched in dLGN | 1.606 | 5.597E-11 | 1383 |
| ENSMUSG00000045083 | leucine rich repeat and Ig domain containing 2 [Source:MGI Symbol;Acc:MGI:2442298] | Lingo2 | Enriched in dLGN | 1.606 | 0.00181 | 769.1 |
| ENSMUSG00000028063 | lamin A [Source:MGI Symbol;Acc:MGI:96794] | Lmna | Enriched in dLGN | 1.605 | 3.644E-08 | 135.1 |
| ENSMUSG00000042724 | mitogen-activated protein kinase kinase kinase 9 [Source:MGI Symbol;Acc:MGI:2449952] | Map3k9 | Enriched in dLGN | 1.603 | 1.158E-16 | 571.6 |
| ENSMUSG00000040105 | phospholipid phosphatase 6 [Source:MGI Symbol;Acc:MGI:1921661] | Plpp6 | Enriched in dLGN | 1.603 | 1.262E-08 | 114.9 |
| ENSMUSG00000063802 | HSPA (heat shock 70kDa) binding protein, cytoplasmic cochaperone 1 [Source:MGI Symbol;Acc:MGI:1913495] | Hspbp1 | Enriched in dLGN | 1.595 | 1.116E-12 | 128.5 |
| ENSMUSG00000047261 | growth associated protein 43 [Source:MGI Symbol;Acc:MGI:95639] | Gap43 | Enriched in dLGN | 1.594 | 4.442E-08 | 876.7 |
| ENSMUSG00000027221 | carbohydrate sulfotransferase 1 [Source:MGI Symbol;Acc:MGI:1924219] | Chst1 | Enriched in dLGN | 1.591 | 1.533E-09 | 247 |
| ENSMUSG00000097185 | predicted gene, 26596 [Source:MGI Symbol;Acc:MGI:5477090] | Gm26596 | Enriched in dLGN | 1.59 | 0.001148 | 16.73 |
| ENSMUSG00000038094 | ATPase type 13A4 [Source:MGI Symbol;Acc:MGI:1924456] | Atp13a4 | Enriched in dLGN | 1.589 | 0.02059 | 39.55 |
| ENSMUSG00000026687 | aldehyde dehydrogenase 9, subfamily A1 [Source:MGI Symbol;Acc:MGI:1861622] | Aldh9a1 | Enriched in dLGN | 1.587 | 3.578E-08 | 102.9 |
| ENSMUSG00000034685 | family with sequence similarity 171, member A2 [Source:MGI Symbol;Acc:MGI:2448496] | Fam171a2 | Enriched in dLGN | 1.585 | 3.118E-07 | 403.3 |
| ENSMUSG00000038587 | A kinase (PRKA) anchor protein (gravin) 12 [Source:MGI Symbol;Acc:MGI:1932576] | Akap12 | Enriched in dLGN | 1.581 | 0.00006469 | 1056 |
| ENSMUSG00000045763 | brain abundant, membrane attached signal protein 1 [Source:MGI Symbol;Acc:MGI:1917600] | Basp1 | Enriched in dLGN | 1.578 | 1.048E-09 | 3099 |
| ENSMUSG00000057614 | guanine nucleotide binding protein (G protein), alpha inhibiting 1 [Source:MGI Symbol;Acc:MGI:95771] | Gnai1 | Enriched in dLGN | 1.577 | 7.016E-07 | 448.7 |
| ENSMUSG00000028945 | Ras homolog enriched in brain [Source:MGI Symbol;Acc:MGI:97912] | Rheb | Enriched in dLGN | 1.575 | 1.725E-15 | 175.6 |
| ENSMUSG00000038453 | SRC kinase signaling inhibitor 1 [Source:MGI Symbol;Acc:MGI:1933179] | Srcin1 | Enriched in dLGN | 1.572 | 8.424E-16 | 3243 |
| ENSMUSG00000056222 | sparc/osteonectin, cwcv and kazal-like domains proteoglycan 1 [Source:MGI Symbol;Acc:MGI:105371] | Spock1 | Enriched in dLGN | 1.57 | 0.000008473 | 1204 |
| ENSMUSG00000027220 | synaptotagmin XIII [Source:MGI Symbol;Acc:MGI:1933945] | Syt13 | Enriched in dLGN | 1.568 | 4.028E-15 | 763.5 |
| ENSMUSG00000103009 | predicted gene, 56350 [Source:MGI Symbol;Acc:MGI:6849158] | Gm56350 | Enriched in dLGN | 1.568 | 6.156E-07 | 44.44 |
| ENSMUSG00000056856 | janus kinase and microtubule interacting protein 3 [Source:MGI Symbol;Acc:MGI:1921254] | Jakmip3 | Enriched in dLGN | 1.567 | 1.85E-09 | 256.3 |
| ENSMUSG00000003273 | carbonic anhydrase 11 [Source:MGI Symbol;Acc:MGI:1336193] | Car11 | Enriched in dLGN | 1.564 | 6.384E-09 | 105.3 |
| ENSMUSG00000036966 | SPRY domain containing 3 [Source:MGI Symbol;Acc:MGI:2446175] | Spryd3 | Enriched in dLGN | 1.562 | 8.742E-15 | 481.5 |
| ENSMUSG00000045045 | leucine rich repeat and fibronectin type III domain containing 4 [Source:MGI Symbol;Acc:MGI:2385612] | Lrfn4 | Enriched in dLGN | 1.56 | 7.526E-10 | 212.7 |
| ENSMUSG00000035828 | proviral integration site 3 [Source:MGI Symbol;Acc:MGI:1355297] | Pim3 | Enriched in dLGN | 1.56 | 1.457E-08 | 118.5 |
| ENSMUSG00000038777 | sema domain, transmembrane domain (TM), and cytoplasmic domain, (semaphorin) 6C [Source:MGI Symbol;Acc:MGI:1338032] | Sema6c | Enriched in dLGN | 1.556 | 0.00009809 | 156.7 |
| ENSMUSG00000002006 | PDZ domain containing 4 [Source:MGI Symbol;Acc:MGI:2443483] | Pdzd4 | Enriched in dLGN | 1.555 | 1.382E-10 | 936.4 |
| ENSMUSG00000072663 | sperm flagellar 2 [Source:MGI Symbol;Acc:MGI:2443727] | Spef2 | Enriched in dLGN | 1.555 | 0.04439 | 23.31 |
| ENSMUSG00000021377 | DEK proto-oncogene (DNA binding) [Source:MGI Symbol;Acc:MGI:1926209] | Dek | Enriched in dLGN | 1.554 | 4.51E-12 | 361.9 |
| ENSMUSG00000035064 | eukaryotic elongation factor-2 kinase [Source:MGI Symbol;Acc:MGI:1195261] | Eef2k | Enriched in dLGN | 1.55 | 6.211E-11 | 118.4 |
| ENSMUSG00000020327 | fibroblast growth factor 22 [Source:MGI Symbol;Acc:MGI:1914362] | Fgf22 | Enriched in dLGN | 1.55 | 0.005751 | 11.32 |
| ENSMUSG00000044024 | RELT-like 2 [Source:MGI Symbol;Acc:MGI:1918044] | Rell2 | Enriched in dLGN | 1.549 | 1.349E-12 | 285.3 |
| ENSMUSG00000031561 | teneurin transmembrane protein 3 [Source:MGI Symbol;Acc:MGI:1345183] | Tenm3 | Enriched in dLGN | 1.549 | 3.376E-11 | 2385 |
| ENSMUSG00000061702 | transmembrane protein 91 [Source:MGI Symbol;Acc:MGI:2443589] | Tmem91 | Enriched in dLGN | 1.547 | 0.002383 | 41.88 |
| ENSMUSG00000021102 | glutaredoxin 5 [Source:MGI Symbol;Acc:MGI:1920296] | Glrx5 | Enriched in dLGN | 1.545 | 9.227E-07 | 170.8 |
| ENSMUSG00000008496 | POU domain, class 2, transcription factor 2 [Source:MGI Symbol;Acc:MGI:101897] | Pou2f2 | Enriched in dLGN | 1.539 | 0.00001295 | 295.3 |
| ENSMUSG00000044576 | GRB2 associated regulator of MAPK1 subtype 2 [Source:MGI Symbol;Acc:MGI:2685290] | Garem2 | Enriched in dLGN | 1.537 | 0.000005956 | 62.67 |
| ENSMUSG00000006218 | family with sequence similarity 131, member C [Source:MGI Symbol;Acc:MGI:2685539] | Fam131c | Enriched in dLGN | 1.533 | 0.000187 | 42.58 |
| ENSMUSG00000036087 | SLAIN motif family, member 2 [Source:MGI Symbol;Acc:MGI:1923241] | Slain2 | Enriched in dLGN | 1.532 | 1.825E-09 | 209.6 |
| ENSMUSG00000018509 | centromere protein V [Source:MGI Symbol;Acc:MGI:1920389] | Cenpv | Enriched in dLGN | 1.523 | 0.0002442 | 40.11 |
| ENSMUSG00000031760 | metallothionein 3 [Source:MGI Symbol;Acc:MGI:97173] | Mt3 | Enriched in dLGN | 1.522 | 0.00021 | 131.4 |
| ENSMUSG00000025790 | solute carrier organic anion transporter family, member 3a1 [Source:MGI Symbol;Acc:MGI:1351867] | Slco3a1 | Enriched in dLGN | 1.519 | 0.0000681 | 185.2 |
| ENSMUSG00000054428 | ATPase inhibitory factor 1 [Source:MGI Symbol;Acc:MGI:1196457] | Atpif1 | Enriched in dLGN | 1.514 | 0.000000774 | 829.4 |
| ENSMUSG00000020658 | EFR3 homolog B [Source:MGI Symbol;Acc:MGI:2444851] | Efr3b | Enriched in dLGN | 1.511 | 5.05E-13 | 856.7 |
| ENSMUSG00000029126 | neuron specific gene family member 1 [Source:MGI Symbol;Acc:MGI:109149] | Nsg1 | Enriched in dLGN | 1.511 | 1.086E-07 | 719.7 |
| ENSMUSG00000039057 | myosin XVI [Source:MGI Symbol;Acc:MGI:2685951] | Myo16 | Enriched in dLGN | 1.511 | 0.000004251 | 431.2 |
| ENSMUSG00000042388 | DLG associated protein 3 [Source:MGI Symbol;Acc:MGI:3039563] | Dlgap3 | Enriched in dLGN | 1.51 | 6.57E-10 | 1641 |
| ENSMUSG00000053192 | myeloid/lymphoid or mixed-lineage leukemia; translocated to, 11 [Source:MGI Symbol;Acc:MGI:1929671] | Mllt11 | Enriched in dLGN | 1.506 | 7.372E-09 | 499.1 |
| ENSMUSG00000037541 | SH3 and multiple ankyrin repeat domains 2 [Source:MGI Symbol;Acc:MGI:2671987] | Shank2 | Enriched in dLGN | 1.504 | 2.064E-24 | 1896 |
| ENSMUSG00000035576 | L3MBTL1 histone methyl-lysine binding protein [Source:MGI Symbol;Acc:MGI:2676663] | L3mbtl1 | Enriched in dLGN | 1.504 | 0.0001141 | 85.42 |
| ENSMUSG00000003469 | phytanoyl-CoA hydroxylase interacting protein [Source:MGI Symbol;Acc:MGI:1860417] | Phyhip | Enriched in dLGN | 1.504 | 0.0005422 | 112 |
| ENSMUSG00000028677 | ring finger protein 220 [Source:MGI Symbol;Acc:MGI:1913993] | Rnf220 | Enriched in dLGN | 1.5 | 0.000004865 | 731.2 |
| ENSMUSG00000029475 | lysine (K)-specific demethylase 2B [Source:MGI Symbol;Acc:MGI:1354737] | Kdm2b | Enriched in dLGN | 1.498 | 3.385E-11 | 344.7 |
| ENSMUSG00000026972 | arrestin domain containing 1 [Source:MGI Symbol;Acc:MGI:2446136] | Arrdc1 | Enriched in dLGN | 1.497 | 0.0008824 | 25.53 |
| ENSMUSG00000024429 | guanine nucleotide binding protein-like 1 [Source:MGI Symbol;Acc:MGI:95764] | Gnl1 | Enriched in dLGN | 1.492 | 6.08E-12 | 740.8 |
| ENSMUSG00000075467 | DNL-type zinc finger [Source:MGI Symbol;Acc:MGI:106559] | Dnlz | Enriched in dLGN | 1.477 | 4.152E-09 | 120.6 |
| ENSMUSG00000026833 | olfactomedin 1 [Source:MGI Symbol;Acc:MGI:1860437] | Olfm1 | Enriched in dLGN | 1.475 | 3.733E-08 | 1863 |
| ENSMUSG00000026991 | plakophilin 4 [Source:MGI Symbol;Acc:MGI:109281] | Pkp4 | Enriched in dLGN | 1.474 | 7.612E-14 | 894.5 |
| ENSMUSG00000049252 | low density lipoprotein-related protein 1B [Source:MGI Symbol;Acc:MGI:2151136] | Lrp1b | Enriched in dLGN | 1.472 | 0.0001121 | 1165 |
| ENSMUSG00000073433 | Rho GDP dissociation inhibitor (GDI) gamma [Source:MGI Symbol;Acc:MGI:108430] | Arhgdig | Enriched in dLGN | 1.471 | 0.000002487 | 185.1 |
| ENSMUSG00000028478 | clathrin, light polypeptide (Lca) [Source:MGI Symbol;Acc:MGI:894297] | Clta | Enriched in dLGN | 1.466 | 7.603E-16 | 866.9 |
| ENSMUSG00000028849 | MAP7 domain containing 1 [Source:MGI Symbol;Acc:MGI:2384297] | Map7d1 | Enriched in dLGN | 1.465 | 1.389E-14 | 1844 |
| ENSMUSG00000040350 | tripartite motif-containing 7 [Source:MGI Symbol;Acc:MGI:2137353] | Trim7 | Enriched in dLGN | 1.461 | 0.01356 | 12.75 |
| ENSMUSG00000034664 | integrin alpha 2b [Source:MGI Symbol;Acc:MGI:96601] | Itga2b | Enriched in dLGN | 1.461 | 0.01407 | 10.87 |
| ENSMUSG00000084845 | transmembrane protein 240 [Source:MGI Symbol;Acc:MGI:3648074] | Tmem240 | Enriched in dLGN | 1.46 | 0.00001217 | 76.95 |
| ENSMUSG00000054000 | tumor suppressor candidate 1 [Source:MGI Symbol;Acc:MGI:2684283] | Tusc1 | Enriched in dLGN | 1.458 | 0.006078 | 40.94 |
| ENSMUSG00000019194 | sodium channel, voltage-gated, type I, beta [Source:MGI Symbol;Acc:MGI:98247] | Scn1b | Enriched in dLGN | 1.456 | 0.00001505 | 470 |
| ENSMUSG00000020481 | ankyrin repeat domain 36 [Source:MGI Symbol;Acc:MGI:1923639] | Ankrd36 | Enriched in dLGN | 1.447 | 0.001574 | 15.79 |
| ENSMUSG00000074793 | heat shock protein 12B [Source:MGI Symbol;Acc:MGI:1919880] | Hspa12b | Enriched in dLGN | 1.446 | 0.0002716 | 37.37 |
| ENSMUSG00000000627 | sema domain, immunoglobulin domain (Ig), TM domain, and short cytoplasmic domain [Source:MGI Symbol;Acc:MGI:1340055] | Sema4f | Enriched in dLGN | 1.439 | 1.345E-07 | 125.4 |
| ENSMUSG00000056076 | eukaryotic translation initiation factor 3, subunit B [Source:MGI Symbol;Acc:MGI:106478] | Eif3b | Enriched in dLGN | 1.437 | 3.147E-14 | 530.2 |
| ENSMUSG00000027189 | tripartite motif-containing 44 [Source:MGI Symbol;Acc:MGI:1931835] | Trim44 | Enriched in dLGN | 1.437 | 2.263E-07 | 2112 |
| ENSMUSG00000034145 | transmembrane protein 63c [Source:MGI Symbol;Acc:MGI:2444386] | Tmem63c | Enriched in dLGN | 1.436 | 0.000000835 | 106.4 |
| ENSMUSG00000006711 | RIKEN cDNA D130043K22 gene [Source:MGI Symbol;Acc:MGI:3036268] | D130043K22Rik | Enriched in dLGN | 1.434 | 0.0002914 | 133.1 |
| ENSMUSG00000054793 | cell adhesion molecule 4 [Source:MGI Symbol;Acc:MGI:2449088] | Cadm4 | Enriched in dLGN | 1.431 | 7.67E-11 | 1063 |
| ENSMUSG00000025384 | Fanconi anemia core complex associated protein 100 [Source:MGI Symbol;Acc:MGI:1919135] | Faap100 | Enriched in dLGN | 1.429 | 0.000001511 | 187.1 |
| ENSMUSG00000034171 | fatty acid amide hydrolase [Source:MGI Symbol;Acc:MGI:109609] | Faah | Enriched in dLGN | 1.425 | 3.329E-11 | 134 |
| ENSMUSG00000018411 | microtubule-associated protein tau [Source:MGI Symbol;Acc:MGI:97180] | Mapt | Enriched in dLGN | 1.422 | 0.000001377 | 1946 |
| ENSMUSG00000020882 | calcium channel, voltage-dependent, beta 1 subunit [Source:MGI Symbol;Acc:MGI:102522] | Cacnb1 | Enriched in dLGN | 1.421 | 4.703E-15 | 246.5 |
| ENSMUSG00000025375 | apoptosis-associated tyrosine kinase [Source:MGI Symbol;Acc:MGI:1197518] | Aatk | Enriched in dLGN | 1.418 | 5.437E-09 | 1628 |
| ENSMUSG00000079003 | sterile alpha motif domain containing 1 [Source:MGI Symbol;Acc:MGI:2142433] | Samd1 | Enriched in dLGN | 1.414 | 1.617E-07 | 225.7 |
| ENSMUSG00000013150 | glucose-fructose oxidoreductase domain containing 2 [Source:MGI Symbol;Acc:MGI:1917825] | Gfod2 | Enriched in dLGN | 1.411 | 1.867E-13 | 156.4 |
| ENSMUSG00000007944 | tetratricopeptide repeat domain 9B [Source:MGI Symbol;Acc:MGI:1920282] | Ttc9b | Enriched in dLGN | 1.411 | 0.000001173 | 147 |
| ENSMUSG00000044813 | src homology 2 domain-containing transforming protein B [Source:MGI Symbol;Acc:MGI:98294] | Shb | Enriched in dLGN | 1.411 | 0.000009609 | 96.59 |
| ENSMUSG00000003604 | apoptosis, caspase activation inhibitor [Source:MGI Symbol;Acc:MGI:1921518] | Aven | Enriched in dLGN | 1.411 | 0.00003542 | 49.12 |
| ENSMUSG00000031805 | Janus kinase 3 [Source:MGI Symbol;Acc:MGI:99928] | Jak3 | Enriched in dLGN | 1.411 | 0.0003412 | 28.96 |
| ENSMUSG00000028610 | DMRT-like family B with proline-rich C-terminal, 1 [Source:MGI Symbol;Acc:MGI:1927125] | Dmrtb1 | Enriched in dLGN | 1.411 | 0.00655 | 17.02 |
| ENSMUSG00000021156 | zinc finger, MYND domain containing 11 [Source:MGI Symbol;Acc:MGI:1913755] | Zmynd11 | Enriched in dLGN | 1.41 | 0.00001596 | 980.2 |
| ENSMUSG00000028949 | SWI/SNF related, matrix associated, actin dependent regulator of chromatin, subfamily d, member 3 [Source:MGI Symbol;Acc:MGI:1914243] | Smarcd3 | Enriched in dLGN | 1.403 | 1.397E-14 | 234.8 |
| ENSMUSG00000026443 | leucine rich repeat protein 2, neuronal [Source:MGI Symbol;Acc:MGI:106037] | Lrrn2 | Enriched in dLGN | 1.403 | 3.19E-08 | 489.2 |
| ENSMUSG00000024935 | solute carrier family 1 (neuronal/epithelial high affinity glutamate transporter, system Xag), member 1 [Source:MGI Symbol;Acc:MGI:105083] | Slc1a1 | Enriched in dLGN | 1.401 | 1.854E-08 | 323.7 |
| ENSMUSG00000026558 | uridine-cytidine kinase 2 [Source:MGI Symbol;Acc:MGI:1931744] | Uck2 | Enriched in dLGN | 1.399 | 5.187E-08 | 349 |
| ENSMUSG00000031700 | glutamic pyruvate transaminase (alanine aminotransferase) 2 [Source:MGI Symbol;Acc:MGI:1915391] | Gpt2 | Enriched in dLGN | 1.398 | 2.833E-08 | 138.5 |
| ENSMUSG00000023827 | 1-acylglycerol-3-phosphate O-acyltransferase 4 (lysophosphatidic acid acyltransferase, delta) [Source:MGI Symbol;Acc:MGI:1915512] | Agpat4 | Enriched in dLGN | 1.398 | 1.392E-07 | 114.1 |
| ENSMUSG00000047013 | F-box protein 41 [Source:MGI Symbol;Acc:MGI:1261912] | Fbxo41 | Enriched in dLGN | 1.397 | 0.000002394 | 562.6 |
| ENSMUSG00000071073 | leucine rich repeat containing 73 [Source:MGI Symbol;Acc:MGI:2684934] | Lrrc73 | Enriched in dLGN | 1.396 | 8.586E-09 | 70.3 |
| ENSMUSG00000030515 | threonyl-tRNA synthetase-like 2 [Source:MGI Symbol;Acc:MGI:2444486] | Tarsl2 | Enriched in dLGN | 1.393 | 1.954E-09 | 105.5 |
| ENSMUSG00000020374 | RasGEF domain family, member 1C [Source:MGI Symbol;Acc:MGI:1921813] | Rasgef1c | Enriched in dLGN | 1.392 | 0.0001963 | 39.33 |

| | | | | | | |
|---|---|---|---|---|---|---|
| ENSMUSG00000033316 | polypeptide N-acetylgalactosaminyltransferase 9 [Source:MGI Symbol;Acc:MGI:2677965] | Galnt9 | Enriched in dLGN | 1.391 | 1.731E-07 | 204.6 |
| ENSMUSG00000074182 | zinc finger, HIT type 6 [Source:MGI Symbol;Acc:MGI:1916996] | Znhit6 | Enriched in dLGN | 1.39 | 0.000001521 | 173.8 |
| ENSMUSG00000057716 | transmembrane protein 178B [Source:MGI Symbol;Acc:MGI:3647581] | Tmem178b | Enriched in dLGN | 1.387 | 8.044E-13 | 1516 |
| ENSMUSG00000053746 | peptidyl-tRNA hydrolase 1 homolog [Source:MGI Symbol;Acc:MGI:1913779] | Ptrh1 | Enriched in dLGN | 1.386 | 0.02413 | 9.59 |
| ENSMUSG00000031422 | mortality factor 4 like 2 [Source:MGI Symbol;Acc:MGI:1927167] | Morf4l2 | Enriched in dLGN | 1.379 | 2.046E-14 | 607.4 |
| ENSMUSG00000034156 | TSPO associated protein 1 [Source:MGI Symbol;Acc:MGI:2450877] | Tspoap1 | Enriched in dLGN | 1.378 | 4.492E-07 | 1203 |
| ENSMUSG00000041115 | IQ motif and Sec7 domain 2 [Source:MGI Symbol;Acc:MGI:3528396] | Iqsec2 | Enriched in dLGN | 1.373 | 3.58E-13 | 368.7 |
| ENSMUSG00000032615 | 5',3'-nucleotidase, mitochondrial [Source:MGI Symbol;Acc:MGI:1917127] | Nt5m | Enriched in dLGN | 1.371 | 0.000003174 | 139.9 |
| ENSMUSG00000034220 | glypican 1 [Source:MGI Symbol;Acc:MGI:1194891] | Gpc1 | Enriched in dLGN | 1.367 | 4.233E-10 | 632.1 |
| ENSMUSG00000079523 | thymosin, beta 10 [Source:MGI Symbol;Acc:MGI:109146] | Tmsb10 | Enriched in dLGN | 1.365 | 0.0003513 | 881.5 |
| ENSMUSG00000038267 | solute carrier family 22, member 23 [Source:MGI Symbol;Acc:MGI:1920352] | Slc22a23 | Enriched in dLGN | 1.364 | 1.354E-08 | 369.7 |
| ENSMUSG00000052584 | stress-associated endoplasmic reticulum protein family member 2 [Source:MGI Symbol;Acc:MGI:1919911] | Serp2 | Enriched in dLGN | 1.361 | 0.000001616 | 134.2 |
| ENSMUSG00000024038 | NADH:ubiquinone oxidoreductase core subunit V3 [Source:MGI Symbol;Acc:MGI:1890894] | Ndufv3 | Enriched in dLGN | 1.359 | 1.345E-07 | 374.2 |
| ENSMUSG00000024064 | polypeptide N-acetylgalactosaminyltransferase 14 [Source:MGI Symbol;Acc:MGI:1918935] | Galnt14 | Enriched in dLGN | 1.358 | 0.0005182 | 190 |
| ENSMUSG00000035969 | RUN and SH3 domain containing 2 [Source:MGI Symbol;Acc:MGI:2140371] | Rusc2 | Enriched in dLGN | 1.357 | 3.498E-09 | 986.7 |
| ENSMUSG00000042532 | golgi autoantigen, golgin subfamily a, 7B [Source:MGI Symbol;Acc:MGI:1918396] | Golga7b | Enriched in dLGN | 1.356 | 1.147E-09 | 283.6 |
| ENSMUSG00000011589 | fibronectin type 3 and SPRY domain-containing protein [Source:MGI Symbol;Acc:MGI:1934858] | Fsd1 | Enriched in dLGN | 1.353 | 0.00021 | 96.92 |
| ENSMUSG00000063873 | solute carrier family 24 (sodium/potassium/calcium exchanger), member 3 [Source:MGI Symbol;Acc:MGI:2137513] | Slc24a3 | Enriched in dLGN | 1.352 | 0.000002281 | 1240 |
| ENSMUSG00000046378 | aspartate beta-hydroxylase domain containing 1 [Source:MGI Symbol;Acc:MGI:2685014] | Asphd1 | Enriched in dLGN | 1.349 | 0.0003573 | 56.65 |
| ENSMUSG00000033788 | dysferlin [Source:MGI Symbol;Acc:MGI:1349385] | Dysf | Enriched in dLGN | 1.347 | 0.0008525 | 67.23 |
| ENSMUSG00000091811 | InaF motif containing 1 [Source:MGI Symbol;Acc:MGI:1913550] | Inafm1 | Enriched in dLGN | 1.345 | 0.0000657 | 82.23 |
| ENSMUSG00000005986 | ankyrin repeat domain 13 family, member D [Source:MGI Symbol;Acc:MGI:1915673] | Ankrd13d | Enriched in dLGN | 1.342 | 4.614E-08 | 148.5 |
| ENSMUSG00000004031 | bone morphogenic protein/retinoic acid inducible neural-specific 2 [Source:MGI Symbol;Acc:MGI:2443333] | Brinp2 | Enriched in dLGN | 1.342 | 0.000005818 | 323.9 |
| ENSMUSG00000048720 | TBC1D12: TBC1 domain family, member 12 [Source:MGI Symbol;Acc:MGI:2384803] | Tbc1d12 | Enriched in dLGN | 1.34 | 3.152E-07 | 117.8 |
| ENSMUSG00000020216 | junctional sarcoplasmic reticulum protein 1 [Source:MGI Symbol;Acc:MGI:1916700] | Jsrp1 | Enriched in dLGN | 1.335 | 0.03926 | 6.626 |
| ENSMUSG00000030376 | solute carrier family 8 (sodium/calcium exchanger), member 2 [Source:MGI Symbol;Acc:MGI:107996] | Slc8a2 | Enriched in dLGN | 1.333 | 4.397E-09 | 330.3 |
| ENSMUSG00000019861 | golgi associated PDZ and coiled-coil motif containing [Source:MGI Symbol;Acc:MGI:2149946] | Gopc | Enriched in dLGN | 1.327 | 7.634E-16 | 304.7 |
| ENSMUSG00000117406 | netrin 3 [Source:MGI Symbol;Acc:MGI:1341188] | Ntn3 | Enriched in dLGN | 1.327 | 0.0005977 | 52.98 |
| ENSMUSG00000033595 | leucine-rich repeat LGI family, member 3 [Source:MGI Symbol;Acc:MGI:2182619] | Lgi3 | Enriched in dLGN | 1.327 | 0.001544 | 185.5 |
| ENSMUSG00000032435 | dynein cytoplasmic 1 light intermediate chain 1 [Source:MGI Symbol;Acc:MGI:2135610] | Dync1li1 | Enriched in dLGN | 1.325 | 6.208E-21 | 559.8 |
| ENSMUSG00000029833 | tripartite motif-containing 24 [Source:MGI Symbol;Acc:MGI:109275] | Trim24 | Enriched in dLGN | 1.316 | 0.000005824 | 198.7 |
| ENSMUSG00000075318 | sodium channel, voltage-gated, type II, alpha [Source:MGI Symbol;Acc:MGI:98248] | Scn2a | Enriched in dLGN | 1.313 | 1.065E-08 | 1334 |
| ENSMUSG00000038607 | guanine nucleotide binding protein (G protein), gamma 10 [Source:MGI Symbol;Acc:MGI:1336169] | Gng10 | Enriched in dLGN | 1.313 | 0.0003336 | 28.82 |
| ENSMUSG00000018405 | mitochondrial rRNA methyltransferase 1 [Source:MGI Symbol;Acc:MGI:2443470] | Mrm1 | Enriched in dLGN | 1.311 | 0.0005427 | 29.07 |
| ENSMUSG00000004113 | calcium channel, voltage-dependent, N type, alpha 1B subunit [Source:MGI Symbol;Acc:MGI:88296] | Cacna1b | Enriched in dLGN | 1.31 | 5.034E-08 | 1028 |
| ENSMUSG00000025487 | proteasome (prosome, macropain) 26S subunit, non-ATPase, 13 [Source:MGI Symbol;Acc:MGI:1345192] | Psmd13 | Enriched in dLGN | 1.308 | 3.407E-09 | 179.8 |
| ENSMUSG00000020866 | calcium channel, voltage-dependent, T type, alpha 1G subunit [Source:MGI Symbol;Acc:MGI:1201678] | Cacna1g | Enriched in dLGN | 1.308 | 4.067E-09 | 1369 |
| ENSMUSG00000027525 | phosphatase and actin regulator 3 [Source:MGI Symbol;Acc:MGI:1921439] | Phactr3 | Enriched in dLGN | 1.307 | 6.112E-11 | 523.5 |
| ENSMUSG00000054459 | visinin-like 1 [Source:MGI Symbol;Acc:MGI:1349453] | Vsnl1 | Enriched in dLGN | 1.307 | 0.01119 | 746.5 |
| ENSMUSG00000028532 | cache domain containing 1 [Source:MGI Symbol;Acc:MGI:2444177] | Cachd1 | Enriched in dLGN | 1.305 | 0.00007674 | 291.5 |
| ENSMUSG00000040502 | membrane associated ring-CH-type finger 9 [Source:MGI Symbol;Acc:MGI:2446144] | Marchf9 | Enriched in dLGN | 1.304 | 0.000001644 | 235.6 |
| ENSMUSG00000026207 | SPEG complex locus [Source:MGI Symbol;Acc:MGI:109282] | Speg | Enriched in dLGN | 1.303 | 9.366E-09 | 395.1 |
| ENSMUSG00000025940 | transmembrane protein 70 [Source:MGI Symbol;Acc:MGI:1915068] | Tmem70 | Enriched in dLGN | 1.303 | 0.0000358 | 50.14 |
| ENSMUSG00000017831 | RAB5A, member RAS oncogene family [Source:MGI Symbol;Acc:MGI:105926] | Rab5a | Enriched in dLGN | 1.3 | 0.000004483 | 208.5 |
| ENSMUSG00000042712 | transcription elongation factor A like 9 [Source:MGI Symbol;Acc:MGI:109567] | Tceal9 | Enriched in dLGN | 1.299 | 0.000004121 | 169.6 |
| ENSMUSG00000090071 | cyclin-dependent kinase 5, regulatory subunit 2 (p39) [Source:MGI Symbol;Acc:MGI:1330828] | Cdk5r2 | Enriched in dLGN | 1.296 | 0.000001589 | 1118 |
| ENSMUSG00000026959 | glutamate receptor, ionotropic, NMDA1 (zeta 1) [Source:MGI Symbol;Acc:MGI:95819] | Grin1 | Enriched in dLGN | 1.296 | 0.0002137 | 942.6 |
| ENSMUSG00000042751 | nicotinamide nucleotide adenylyltransferase 2 [Source:MGI Symbol;Acc:MGI:2444155] | Nmnat2 | Enriched in dLGN | 1.288 | 9.899E-07 | 932 |
| ENSMUSG00000058740 | potassium channel, subfamily T, member 1 [Source:MGI Symbol;Acc:MGI:1924627] | Kcnt1 | Enriched in dLGN | 1.288 | 0.0005104 | 138.9 |
| ENSMUSG00000009394 | synapsin II [Source:MGI Symbol;Acc:MGI:103020] | Syn2 | Enriched in dLGN | 1.286 | 0.00005617 | 611.4 |
| ENSMUSG00000052539 | membrane associated guanylate kinase, WW and PDZ domain containing 3 [Source:MGI Symbol;Acc:MGI:1923484] | Magi3 | Enriched in dLGN | 1.284 | 4.672E-11 | 426.4 |
| ENSMUSG00000029814 | insulin-like growth factor 2 mRNA binding protein 3 [Source:MGI Symbol;Acc:MGI:1890359] | Igf2bp3 | Enriched in dLGN | 1.284 | 0.01456 | 57.12 |
| ENSMUSG00000033597 | CASK interacting protein 1 [Source:MGI Symbol;Acc:MGI:2442952] | Caskin1 | Enriched in dLGN | 1.281 | 2.247E-10 | 1073 |
| ENSMUSG00000023017 | acid-sensing (proton-gated) ion channel 1 [Source:MGI Symbol;Acc:MGI:1194915] | Asic1 | Enriched in dLGN | 1.281 | 6.464E-07 | 275.4 |
| ENSMUSG00000042804 | G protein-coupled receptor 153 [Source:MGI Symbol;Acc:MGI:1916157] | Gpr153 | Enriched in dLGN | 1.28 | 0.0000148 | 1042 |
| ENSMUSG00000045009 | proline-rich transmembrane protein 3 [Source:MGI Symbol;Acc:MGI:2444810] | Prrt3 | Enriched in dLGN | 1.279 | 0.00007855 | 111.1 |
| ENSMUSG00000066705 | FXYD domain-containing ion transport regulator 6 [Source:MGI Symbol;Acc:MGI:1890226] | Fxyd6 | Enriched in dLGN | 1.279 | 0.008957 | 380.3 |
| ENSMUSG00000034271 | Jun dimerization protein 2 [Source:MGI Symbol;Acc:MGI:1932093] | Jdp2 | Enriched in dLGN | 1.278 | 1.313E-09 | 161.1 |
| ENSMUSG00000092216 | predicted gene, 19345 [Source:MGI Symbol;Acc:MGI:5011530] | Gm19345 | Enriched in dLGN | 1.277 | 0.001287 | 23.03 |
| ENSMUSG00000034839 | La ribonucleoprotein 6, translational regulator [Source:MGI Symbol;Acc:MGI:1914807] | Larp6 | Enriched in dLGN | 1.275 | 0.000000117 | 87.58 |
| ENSMUSG00000018507 | transient receptor potential cation channel, subfamily V, member 2 [Source:MGI Symbol;Acc:MGI:1341836] | Trpv2 | Enriched in dLGN | 1.275 | 0.0001497 | 57.51 |
| ENSMUSG00000026203 | DnaJ heat shock protein family (Hsp40) member B2 [Source:MGI Symbol;Acc:MGI:1928739] | Dnajb2 | Enriched in dLGN | 1.274 | 4.948E-09 | 239.7 |
| ENSMUSG00000028680 | polo like kinase 3 [Source:MGI Symbol;Acc:MGI:109604] | Plk3 | Enriched in dLGN | 1.274 | 0.0003991 | 41.32 |
| ENSMUSG00000050164 | melanin-concentrating hormone receptor 1 [Source:MGI Symbol;Acc:MGI:2180756] | Mchr1 | Enriched in dLGN | 1.274 | 0.03772 | 17.75 |
| ENSMUSG00000027223 | mitogen-activated protein kinase 8 interacting protein 1 [Source:MGI Symbol;Acc:MGI:1309464] | Mapk8ip1 | Enriched in dLGN | 1.272 | 5.31E-08 | 1583 |
| ENSMUSG00000058297 | sparc/osteonectin, cwcv and kazal-like domains proteoglycan 2 [Source:MGI Symbol;Acc:MGI:1891351] | Spock2 | Enriched in dLGN | 1.27 | 0.0006763 | 7309 |
| ENSMUSG00000026499 | acyl-Coenzyme A binding domain containing 3 [Source:MGI Symbol;Acc:MGI:2181074] | Acbd3 | Enriched in dLGN | 1.267 | 0.000003543 | 390.9 |
| ENSMUSG00000002319 | importin 4 [Source:MGI Symbol;Acc:MGI:1923001] | Ipo4 | Enriched in dLGN | 1.266 | 2.044E-08 | 114.9 |
| ENSMUSG00000024758 | reticulon 3 [Source:MGI Symbol;Acc:MGI:1339970] | Rtn3 | Enriched in dLGN | 1.263 | 1.339E-10 | 1535 |
| ENSMUSG00000027203 | deoxyuridine triphosphatase [Source:MGI Symbol;Acc:MGI:1346051] | Dut | Enriched in dLGN | 1.262 | 0.000001125 | 92.41 |
| ENSMUSG00000032024 | CXADR-like membrane protein [Source:MGI Symbol;Acc:MGI:1918816] | Clmp | Enriched in dLGN | 1.262 | 0.00001111 | 151.4 |
| ENSMUSG00000062661 | neuronal calcium sensor 1 [Source:MGI Symbol;Acc:MGI:109166] | Ncs1 | Enriched in dLGN | 1.261 | 6.605E-08 | 1107 |
| ENSMUSG00000020029 | nudix (nucleoside diphosphate linked moiety X)-type motif 4 [Source:MGI Symbol;Acc:MGI:1918457] | Nudt4 | Enriched in dLGN | 1.259 | 0.000008653 | 752.5 |
| ENSMUSG00000033768 | neurexin II [Source:MGI Symbol;Acc:MGI:1096362] | Nrxn2 | Enriched in dLGN | 1.258 | 0.000000006 | 5038 |
| ENSMUSG00000006576 | solute carrier family 4 (anion exchanger), member 3 [Source:MGI Symbol;Acc:MGI:109350] | Slc4a3 | Enriched in dLGN | 1.256 | 4.366E-09 | 825.7 |
| ENSMUSG00000078591 | heparan sulfate (glucosamine) 3-O-sulfotransferase 4 [Source:MGI Symbol;Acc:MGI:1333792] | Hs3st4 | Enriched in dLGN | 1.255 | 0.02244 | 324.6 |
| ENSMUSG00000026885 | tubulin tyrosine ligase-like family, member 11 [Source:MGI Symbol;Acc:MGI:1921660] | Ttll11 | Enriched in dLGN | 1.254 | 0.00005813 | 160.9 |
| ENSMUSG00000047428 | delta like non-canonical Notch ligand 2 [Source:MGI Symbol;Acc:MGI:2146838] | Dlk2 | Enriched in dLGN | 1.252 | 0.0002445 | 45.68 |
| undefined | undefined | undefined | Enriched in dLGN | 1.251 | 0.000266 | 439.9 |
| ENSMUSG00000021114 | ATPase, H+ transporting, lysosomal V1 subunit D [Source:MGI Symbol;Acc:MGI:1921084] | Atp6v1d | Enriched in dLGN | 1.25 | 0.00003891 | 284.8 |
| ENSMUSG00000024462 | gamma-aminobutyric acid (GABA) B receptor, 1 [Source:MGI Symbol;Acc:MGI:1860139] | Gabbr1 | Enriched in dLGN | 1.241 | 2.165E-11 | 2409 |
| ENSMUSG00000047155 | cytochrome P450, family 4, subfamily x, polypeptide 1 [Source:MGI Symbol;Acc:MGI:1932403] | Cyp4x1 | Enriched in dLGN | 1.241 | 0.001238 | 55.68 |
| ENSMUSG00000025869 | NOP16 nucleolar protein [Source:MGI Symbol;Acc:MGI:107862] | Nop16 | Enriched in dLGN | 1.238 | 0.00006962 | 53.74 |
| ENSMUSG00000028337 | coronin, actin binding protein 2A [Source:MGI Symbol;Acc:MGI:1345966] | Coro2a | Enriched in dLGN | 1.237 | 0.00003767 | 177.4 |
| ENSMUSG00000028785 | hippocalcin [Source:MGI Symbol;Acc:MGI:1336200] | Hpca | Enriched in dLGN | 1.236 | 0.000001693 | 294.3 |
| ENSMUSG00000018012 | Rac family small GTPase 3 [Source:MGI Symbol;Acc:MGI:2180784] | Rac3 | Enriched in dLGN | 1.236 | 0.00155 | 122.2 |
| ENSMUSG00000033615 | complexin 1 [Source:MGI Symbol;Acc:MGI:104727] | Cplx1 | Enriched in dLGN | 1.236 | 0.002218 | 2304 |
| ENSMUSG00000018865 | sulfotransferase family 4A, member 1 [Source:MGI Symbol;Acc:MGI:1888971] | Sult4a1 | Enriched in dLGN | 1.235 | 2.803E-07 | 795.2 |
| ENSMUSG00000026858 | mitoguardin 2 [Source:MGI Symbol;Acc:MGI:1922035] | Miga2 | Enriched in dLGN | 1.234 | 1.204E-09 | 319.8 |
| ENSMUSG00000039156 | stromal interaction molecule 2 [Source:MGI Symbol;Acc:MGI:2151156] | Stim2 | Enriched in dLGN | 1.233 | 4.62E-10 | 187.8 |
| ENSMUSG00000022438 | parvin, beta [Source:MGI Symbol;Acc:MGI:2153063] | Parvb | Enriched in dLGN | 1.232 | 0.0000573 | 253.3 |
| ENSMUSG00000055629 | beta-1,4-N-acetyl-galactosaminyl transferase 4 [Source:MGI Symbol;Acc:MGI:2652891] | B4galnt4 | Enriched in dLGN | 1.229 | 3.344E-07 | 491.4 |
| ENSMUSG00000020544 | cytochrome c oxidase assembly protein 11, copper chaperone [Source:MGI Symbol;Acc:MGI:1917052] | Cox11 | Enriched in dLGN | 1.228 | 0.0001878 | 38.98 |
| ENSMUSG00000016252 | ATP synthase, H+ transporting, mitochondrial F1 complex, epsilon subunit [Source:MGI Symbol;Acc:MGI:1855697] | Atp5e | Enriched in dLGN | 1.222 | 0.002074 | 153.2 |
| ENSMUSG00000061451 | transmembrane protein 151A [Source:MGI Symbol;Acc:MGI:2147713] | Tmem151a | Enriched in dLGN | 1.221 | 0.000003553 | 350.6 |
| ENSMUSG00000054079 | UTP18 small subunit processome component [Source:MGI Symbol;Acc:MGI:1923402] | Utp18 | Enriched in dLGN | 1.22 | 1.493E-07 | 90.78 |
| ENSMUSG00000020083 | family with sequence similarity 241, member B [Source:MGI Symbol;Acc:MGI:1917144] | Fam241b | Enriched in dLGN | 1.22 | 0.0001268 | 42.28 |
| ENSMUSG00000022012 | ecto-NOX disulfide-thiol exchanger 1 [Source:MGI Symbol;Acc:MGI:2444896] | Enox1 | Enriched in dLGN | 1.218 | 0.01815 | 297.2 |
| ENSMUSG00000020331 | hyperpolarization-activated, cyclic nucleotide-gated K+ 2 [Source:MGI Symbol;Acc:MGI:1298210] | Hcn2 | Enriched in dLGN | 1.216 | 2.667E-07 | 598.2 |
| ENSMUSG00000029060 | mindbomb E3 ubiquitin protein ligase 2 [Source:MGI Symbol;Acc:MGI:2679684] | Mib2 | Enriched in dLGN | 1.216 | 9.745E-07 | 340.9 |
| ENSMUSG00000029716 | transferrin receptor 2 [Source:MGI Symbol;Acc:MGI:1354956] | Tfr2 | Enriched in dLGN | 1.216 | 0.008284 | 15.12 |
| ENSMUSG00000045174 | APC membrane recruitment 3 [Source:MGI Symbol;Acc:MGI:3026939] | Amer3 | Enriched in dLGN | 1.215 | 0.0001207 | 95.69 |
| ENSMUSG00000022353 | MTSS I-BAR domain containing 1 [Source:MGI Symbol;Acc:MGI:2384818] | Mtss1 | Enriched in dLGN | 1.214 | 1.898E-12 | 629.5 |
| ENSMUSG00000021400 | Werner helicase interacting protein 1 [Source:MGI Symbol;Acc:MGI:1926153] | Wrnip1 | Enriched in dLGN | 1.212 | 3.756E-08 | 192.4 |
| ENSMUSG00000020496 | ring finger protein 187 [Source:MGI Symbol;Acc:MGI:1914224] | Rnf187 | Enriched in dLGN | 1.203 | 2.326E-07 | 1303 |
| ENSMUSG00000030282 | cytidine monophospho-N-acetylneuraminic acid synthetase [Source:MGI Symbol;Acc:MGI:1337124] | Cmas | Enriched in dLGN | 1.197 | 1.045E-12 | 276.8 |
| ENSMUSG00000039033 | taspase, threonine aspartase 1 [Source:MGI Symbol;Acc:MGI:1923062] | Tasp1 | Enriched in dLGN | 1.196 | 0.03359 | 247 |
| ENSMUSG00000033545 | zinc and ring finger 1 [Source:MGI Symbol;Acc:MGI:2177308] | Znrf1 | Enriched in dLGN | 1.191 | 0.000007464 | 710.6 |
| ENSMUSG00000051518 | ribosomal protein S19 binding protein 1 [Source:MGI Symbol;Acc:MGI:1913788] | Rps19bp1 | Enriched in dLGN | 1.187 | 0.0001403 | 50.07 |
| ENSMUSG00000024597 | solute carrier family 12, member 2 [Source:MGI Symbol;Acc:MGI:101924] | Slc12a2 | Enriched in dLGN | 1.187 | 0.0003406 | 331.5 |
| ENSMUSG00000035051 | DEAH (Asp-Glu-Ala-Asp/His) box polypeptide 57 [Source:MGI Symbol;Acc:MGI:2147067] | Dhx57 | Enriched in dLGN | 1.186 | 0.000002565 | 248.4 |
| ENSMUSG00000047547 | clathrin, light polypeptide (Lcb) [Source:MGI Symbol;Acc:MGI:1921575] | Cltb | Enriched in dLGN | 1.184 | 1.591E-13 | 896 |
| ENSMUSG00000025576 | RNA binding protein, fox-1 homolog (C. elegans) 3 [Source:MGI Symbol;Acc:MGI:106368] | Rbfox3 | Enriched in dLGN | 1.184 | 0.000008435 | 388.1 |
| ENSMUSG00000041375 | coiled-coil domain containing 9 [Source:MGI Symbol;Acc:MGI:1921443] | Ccdc9 | Enriched in dLGN | 1.174 | 1.918E-07 | 178.7 |
| ENSMUSG00000050967 | cellular repressor of E1A-stimulated genes 2 [Source:MGI Symbol;Acc:MGI:1928333] | Creg2 | Enriched in dLGN | 1.17 | 0.0003536 | 188.8 |
| ENSMUSG00000035640 | calcium channel, voltage-dependent, beta subunit associated regulatory protein [Source:MGI Symbol;Acc:MGI:1354170] | Cbarp | Enriched in dLGN | 1.169 | 0.0000152 | 1169 |
| ENSMUSG00000005575 | ubiquitin-conjugating enzyme E2M [Source:MGI Symbol;Acc:MGI:108278] | Ube2m | Enriched in dLGN | 1.168 | 7.696E-09 | 351 |
| ENSMUSG00000045733 | shadow of prion protein [Source:MGI Symbol;Acc:MGI:3582583] | Sprn | Enriched in dLGN | 1.166 | 0.000001412 | 239 |
| ENSMUSG00000032249 | acidic (leucine-rich) nuclear phosphoprotein 32 family, member A [Source:MGI Symbol;Acc:MGI:108447] | Anp32a | Enriched in dLGN | 1.165 | 2.129E-10 | 803.6 |
| ENSMUSG00000020868 | xylosyltransferase II [Source:MGI Symbol;Acc:MGI:2444797] | Xylt2 | Enriched in dLGN | 1.163 | 6.76E-08 | 293.3 |
| ENSMUSG00000022848 | solute carrier family 49 member 4 [Source:MGI Symbol;Acc:MGI:2387188] | Slc49a4 | Enriched in dLGN | 1.161 | 0.00005153 | 225.2 |
| ENSMUSG00000019471 | cell division cycle 37 [Source:MGI Symbol;Acc:MGI:109531] | Cdc37 | Enriched in dLGN | 1.158 | 0.000002128 | 613 |
| ENSMUSG00000040390 | mitogen-activated protein kinase kinase kinase 10 [Source:MGI Symbol;Acc:MGI:1346879] | Map3k10 | Enriched in dLGN | 1.156 | 0.00003034 | 1531 |
| ENSMUSG00000063200 | nucleolar protein 7 [Source:MGI Symbol;Acc:MGI:1917328] | Nol7 | Enriched in dLGN | 1.156 | 0.00004521 | 166.5 |
| ENSMUSG00000035653 | leucine rich repeat and fibronectin type III domain containing 5 [Source:MGI Symbol;Acc:MGI:2144814] | Lrfn5 | Enriched in dLGN | 1.154 | 0.004434 | 564.4 |
| ENSMUSG00000002997 | protein kinase, cAMP dependent regulatory, type II beta [Source:MGI Symbol;Acc:MGI:97760] | Prkar2b | Enriched in dLGN | 1.145 | 0.00009738 | 198.8 |
| ENSMUSG00000030854 | protein tyrosine phosphatase, non-receptor type 5 [Source:MGI Symbol;Acc:MGI:97807] | Ptpn5 | Enriched in dLGN | 1.143 | 4.617E-07 | 407 |
| ENSMUSG00000032557 | ubiquitin-like modifier activating enzyme 5 [Source:MGI Symbol;Acc:MGI:1913913] | Uba5 | Enriched in dLGN | 1.142 | 1.942E-09 | 128.6 |

| | | | | | | |
|---|---|---|---|---|---|---|
| ENSMUSG00000024858 | G protein-coupled receptor kinase 2 [Source:MGI Symbol;Acc:MGI:87940] | Grk2 | Enriched in dLGN | 1.138 | 2.738E-08 | 566.1 |
| ENSMUSG00000033020 | polymerase (RNA) II (DNA directed) polypeptide F [Source:MGI Symbol;Acc:MGI:1349393] | Polr2f | Enriched in dLGN | 1.135 | 0.00001057 | 105.7 |
| ENSMUSG00000045348 | neuronal tyrosine-phosphorylated phosphoinositide 3-kinase adaptor 1 [Source:MGI Symbol;Acc:MGI:2443880] | Nyap1 | Enriched in dLGN | 1.124 | 4.169E-07 | 501.1 |
| ENSMUSG00000043719 | collagen, type VI, alpha 6 [Source:MGI Symbol;Acc:MGI:2444259] | Col6a6 | Enriched in dLGN | 1.124 | 0.01477 | 894 |
| ENSMUSG00000034993 | vesicle amine transport 1 [Source:MGI Symbol;Acc:MGI:1349450] | Vat1 | Enriched in dLGN | 1.122 | 0.0005656 | 429.8 |
| ENSMUSG00000040640 | ELKS/RAB6-interacting/CAST family member 2 [Source:MGI Symbol;Acc:MGI:1098749] | Erc2 | Enriched in dLGN | 1.122 | 0.002365 | 1202 |
| ENSMUSG00000020116 | partner of NOB1 homolog [Source:MGI Symbol;Acc:MGI:1913499] | Pno1 | Enriched in dLGN | 1.119 | 0.001776 | 43.22 |
| ENSMUSG00000055717 | SLAIN motif family, member 1 [Source:MGI Symbol;Acc:MGI:2145578] | Slain1 | Enriched in dLGN | 1.118 | 0.00004592 | 164.4 |
| ENSMUSG00000064254 | ethylmalonic encephalopathy 1 [Source:MGI Symbol;Acc:MGI:1913321] | Ethe1 | Enriched in dLGN | 1.116 | 0.01 | 23.39 |
| ENSMUSG00000031906 | sphingomyelin phosphodiesterase 3, neutral [Source:MGI Symbol;Acc:MGI:1927578] | Smpd3 | Enriched in dLGN | 1.115 | 0.0002379 | 615.6 |
| ENSMUSG00000024095 | heterogeneous nuclear ribonucleoprotein L-like [Source:MGI Symbol;Acc:MGI:1919942] | Hnrnpll | Enriched in dLGN | 1.114 | 0.00003724 | 186.5 |
| ENSMUSG00000028541 | UDP-Gal:betaGlcNAc beta 1,4- galactosyltransferase, polypeptide 2 [Source:MGI Symbol;Acc:MGI:1858493] | B4galt2 | Enriched in dLGN | 1.114 | 0.00005975 | 235.2 |
| ENSMUSG00000018451 | RIKEN cDNA 6330403K07 gene [Source:MGI Symbol;Acc:MGI:1918001] | 6330403K07Rik | Enriched in dLGN | 1.111 | 0.01506 | 664.5 |
| ENSMUSG00000078794 | dishevelled-binding antagonist of beta-catenin 3 [Source:MGI Symbol;Acc:MGI:3654828] | Dact3 | Enriched in dLGN | 1.108 | 0.0002108 | 843 |
| ENSMUSG00000013033 | adhesion G protein-coupled receptor L1 [Source:MGI Symbol;Acc:MGI:1929461] | Adgrl1 | Enriched in dLGN | 1.104 | 1.489E-09 | 3350 |
| ENSMUSG00000063146 | CAP-GLY domain containing linker protein 2 [Source:MGI Symbol;Acc:MGI:1313136] | Clip2 | Enriched in dLGN | 1.104 | 0.00002384 | 788.5 |
| ENSMUSG00000052934 | F-box protein 31 [Source:MGI Symbol;Acc:MGI:1354708] | Fbxo31 | Enriched in dLGN | 1.101 | 9.874E-11 | 417.4 |
| ENSMUSG00000035278 | pleckstrin homology domain containing, family J member 1 [Source:MGI Symbol;Acc:MGI:1925920] | Plekhj1 | Enriched in dLGN | 1.1 | 0.0003182 | 59.38 |
| ENSMUSG00000073557 | protein phosphatase 1, regulatory subunit 12B [Source:MGI Symbol;Acc:MGI:1916417] | Ppp1r12b | Enriched in dLGN | 1.099 | 4.524E-09 | 691.5 |
| ENSMUSG00000036948 | trafficking protein particle complex 14 [Source:MGI Symbol;Acc:MGI:2385896] | Trappc14 | Enriched in dLGN | 1.098 | 3.472E-07 | 146.1 |
| ENSMUSG00000029560 | sorting nexin 8 [Source:MGI Symbol;Acc:MGI:2443816] | Snx8 | Enriched in dLGN | 1.094 | 1.195E-07 | 134.3 |
| ENSMUSG00000022055 | neurofilament, light polypeptide [Source:MGI Symbol;Acc:MGI:97313] | Nefl | Enriched in dLGN | 1.092 | 0.003967 | 7224 |
| ENSMUSG00000031897 | proteasome (prosome, macropain) subunit, beta type 10 [Source:MGI Symbol;Acc:MGI:1096380] | Psmb10 | Enriched in dLGN | 1.088 | 0.001442 | 45.08 |
| ENSMUSG00000032997 | chondroitin polymerizing factor [Source:MGI Symbol;Acc:MGI:106576] | Chpf | Enriched in dLGN | 1.087 | 0.00001274 | 256.5 |
| ENSMUSG00000032463 | Fas apoptotic inhibitory molecule [Source:MGI Symbol;Acc:MGI:1344387] | Faim | Enriched in dLGN | 1.087 | 0.0000221 | 101.9 |
| ENSMUSG00000032366 | tropomyosin 1, alpha [Source:MGI Symbol;Acc:MGI:98809] | Tpm1 | Enriched in dLGN | 1.084 | 1.44E-11 | 654.7 |
| ENSMUSG00000019990 | phosphodiesterase 7B [Source:MGI Symbol;Acc:MGI:1352752] | Pde7b | Enriched in dLGN | 1.082 | 0.008122 | 194.2 |
| ENSMUSG00000028149 | RAP1, GTP-GDP dissociation stimulator 1 [Source:MGI Symbol;Acc:MGI:2385189] | Rap1gds1 | Enriched in dLGN | 1.08 | 0.000003342 | 847.3 |
| ENSMUSG00000025103 | BTB (POZ) domain containing 1 [Source:MGI Symbol;Acc:MGI:1933765] | Btbd1 | Enriched in dLGN | 1.078 | 0.00007814 | 230.6 |
| ENSMUSG00000075590 | nuclear receptor binding protein 2 [Source:MGI Symbol;Acc:MGI:2385017] | Nrbp2 | Enriched in dLGN | 1.076 | 0.00002207 | 333.6 |
| ENSMUSG00000043398 | G protein-coupled receptor 135 [Source:MGI Symbol;Acc:MGI:2676315] | Gpr135 | Enriched in dLGN | 1.076 | 0.01521 | 34.44 |
| ENSMUSG00000030284 | cysteine-rich with EGF-like domains 1 [Source:MGI Symbol;Acc:MGI:2152539] | Creld1 | Enriched in dLGN | 1.071 | 0.000004851 | 242.9 |
| ENSMUSG00000035109 | SHC (Src homology 2 domain containing) family, member 4 [Source:MGI Symbol;Acc:MGI:2655364] | Shc4 | Enriched in dLGN | 1.07 | 0.001244 | 41.77 |
| ENSMUSG00000057858 | family with sequence similarity 204, member A [Source:MGI Symbol;Acc:MGI:1289174] | Fam204a | Enriched in dLGN | 1.065 | 0.00008742 | 72.57 |
| ENSMUSG00000023353 | ArfGAP with GTPase domain, ankyrin repeat and PH domain 3 [Source:MGI Symbol;Acc:MGI:2183446] | Agap3 | Enriched in dLGN | 1.06 | 7.795E-09 | 835.7 |
| ENSMUSG00000079020 | solute carrier family 45, member 4 [Source:MGI Symbol;Acc:MGI:2146236] | Slc45a4 | Enriched in dLGN | 1.06 | 5.249E-08 | 386.8 |
| ENSMUSG00000079598 | C-type lectin domain family 2, member L [Source:MGI Symbol;Acc:MGI:2141402] | Clec2l | Enriched in dLGN | 1.058 | 0.001834 | 252.6 |
| ENSMUSG00000064247 | phosphatidylinositol-specific phospholipase C, X domain containing 1 [Source:MGI Symbol;Acc:MGI:2685422] | Plcxd1 | Enriched in dLGN | 1.055 | 0.008039 | 33.08 |
| ENSMUSG00000027602 | microtubule-associated protein 1 light chain 3 alpha [Source:MGI Symbol;Acc:MGI:1915661] | Map1lc3a | Enriched in dLGN | 1.048 | 0.000129 | 233.2 |
| ENSMUSG00000053137 | mitogen-activated protein kinase 11 [Source:MGI Symbol;Acc:MGI:1338024] | Mapk11 | Enriched in dLGN | 1.048 | 0.005922 | 63.14 |
| ENSMUSG00000021549 | RAS p21 protein activator 1 [Source:MGI Symbol;Acc:MGI:97860] | Rasa1 | Enriched in dLGN | 1.043 | 0.00003736 | 293.7 |
| ENSMUSG00000061859 | PATJ, crumbs cell polarity complex component [Source:MGI Symbol;Acc:MGI:1277960] | Patj | Enriched in dLGN | 1.043 | 0.02039 | 609.6 |
| ENSMUSG00000039372 | membrane associated ring-CH-type finger 4 [Source:MGI Symbol;Acc:MGI:2683550] | Marchf4 | Enriched in dLGN | 1.042 | 0.00001101 | 252.8 |
| ENSMUSG00000038077 | potassium voltage-gated channel, shaker-related, subfamily, member 6 [Source:MGI Symbol;Acc:MGI:96663] | Kcna6 | Enriched in dLGN | 1.038 | 4.212E-07 | 491.8 |
| ENSMUSG00000020635 | FK506 binding protein 1b [Source:MGI Symbol;Acc:MGI:1336205] | Fkbp1b | Enriched in dLGN | 1.037 | 0.01534 | 66.86 |
| ENSMUSG00000023106 | density-regulated protein [Source:MGI Symbol;Acc:MGI:1915434] | Denr | Enriched in dLGN | 1.035 | 0.000005672 | 114.8 |
| ENSMUSG00000027669 | guanine nucleotide binding protein (G protein), beta 4 [Source:MGI Symbol;Acc:MGI:104581] | Gnb4 | Enriched in dLGN | 1.028 | 0.00008659 | 103.7 |
| ENSMUSG00000015092 | endothelial differentiation-related factor 1 [Source:MGI Symbol;Acc:MGI:1891227] | Edf1 | Enriched in dLGN | 1.026 | 0.00005937 | 190.3 |
| ENSMUSG00000048332 | lipoma HMGIC fusion partner [Source:MGI Symbol;Acc:MGI:1920048] | Lhfp | Enriched in dLGN | 1.026 | 0.02386 | 294.9 |
| ENSMUSG00000034194 | R3H domain and coiled-coil containing 1 [Source:MGI Symbol;Acc:MGI:1919093] | R3hcc1 | Enriched in dLGN | 1.025 | 0.0001448 | 178.2 |
| ENSMUSG00000050751 | piggyBac transposable element derived 5 [Source:MGI Symbol;Acc:MGI:2429955] | Pgbd5 | Enriched in dLGN | 1.024 | 0.0000041 | 318 |
| ENSMUSG00000052033 | prefoldin 4 [Source:MGI Symbol;Acc:MGI:1923512] | Pfdn4 | Enriched in dLGN | 1.024 | 0.02488 | 41.11 |
| ENSMUSG00000026188 | transmembrane protein 169 [Source:MGI Symbol;Acc:MGI:2442781] | Tmem169 | Enriched in dLGN | 1.023 | 0.00002421 | 139.3 |
| ENSMUSG00000031963 | BMP-binding endothelial regulator [Source:MGI Symbol;Acc:MGI:1920480] | Bmper | Enriched in dLGN | 1.023 | 0.03578 | 44.47 |
| ENSMUSG00000048661 | LEM domain containing 3 [Source:MGI Symbol;Acc:MGI:3580376] | Lemd3 | Enriched in dLGN | 1.022 | 0.000004924 | 190.2 |
| ENSMUSG00000025437 | ubiquitin specific peptidase 33 [Source:MGI Symbol;Acc:MGI:2159711] | Usp33 | Enriched in dLGN | 1.02 | 0.0002761 | 653.5 |
| ENSMUSG00000033585 | necdin, MAGE family member [Source:MGI Symbol;Acc:MGI:97290] | Ndn | Enriched in dLGN | 1.019 | 0.01025 | 549 |
| ENSMUSG00000021476 | hyaluronic acid binding protein 4 [Source:MGI Symbol;Acc:MGI:1891713] | Habp4 | Enriched in dLGN | 1.018 | 1.405E-07 | 661.9 |
| ENSMUSG00000018474 | chromodomain helicase DNA binding protein 3 [Source:MGI Symbol;Acc:MGI:1344395] | Chd3 | Enriched in dLGN | 1.018 | 5.137E-07 | 5030 |
| ENSMUSG00000024120 | leucine-rich PPR-motif containing [Source:MGI Symbol;Acc:MGI:1919666] | Lrpprc | Enriched in dLGN | 1.017 | 0.000001144 | 294.4 |
| ENSMUSG00000084786 | ubiquitin-like 5 [Source:MGI Symbol;Acc:MGI:1913427] | Ubl5 | Enriched in dLGN | 1.017 | 0.00023 | 89.51 |
| ENSMUSG00000066607 | inhibitory synaptic factor 1 [Source:MGI Symbol;Acc:MGI:2442108] | Insyn1 | Enriched in dLGN | 1.016 | 0.00009771 | 582.5 |
| ENSMUSG00000044117 | bMERB domain containing 1 [Source:MGI Symbol;Acc:MGI:1914504] | Bmerb1 | Enriched in dLGN | 1.013 | 0.00002533 | 457.3 |
| ENSMUSG00000057649 | bromodomain containing 9 [Source:MGI Symbol;Acc:MGI:2145317] | Brd9 | Enriched in dLGN | 1.011 | 0.0024 | 227.3 |
| ENSMUSG00000027408 | carboxypeptidase X 1 (M14 family) [Source:MGI Symbol;Acc:MGI:1934569] | Cpxm1 | Enriched in dLGN | 1.011 | 0.003716 | 23.78 |
| ENSMUSG00000041837 | programmed cell death 7 [Source:MGI Symbol;Acc:MGI:1859170] | Pdcd7 | Enriched in dLGN | 1.008 | 0.00005058 | 140.5 |
| ENSMUSG00000001089 | leucine zipper protein 1 [Source:MGI Symbol;Acc:MGI:107629] | Luzp1 | Enriched in dLGN | 1.007 | 0.001619 | 485.1 |
| ENSMUSG00000031790 | matrix metallopeptidase 15 [Source:MGI Symbol;Acc:MGI:109320] | Mmp15 | Enriched in dLGN | 1.006 | 0.0005836 | 357.5 |
| ENSMUSG00000029120 | protein phosphatase 2, regulatory subunit B, gamma [Source:MGI Symbol;Acc:MGI:2442660] | Ppp2r2c | Enriched in dLGN | 1.003 | 8.682E-07 | 1258 |
| ENSMUSG00000040616 | transmembrane protein 51 [Source:MGI Symbol;Acc:MGI:2384874] | Tmem51 | Enriched in dLGN | 1.003 | 0.007948 | 24.76 |
| ENSMUSG00000028416 | BCL2-associated athanogene 1 [Source:MGI Symbol;Acc:MGI:108047] | Bag1 | Enriched in dLGN | 1.002 | 0.0001667 | 349.9 |
| ENSMUSG00000031357 | synapse associated protein 1 [Source:MGI Symbol;Acc:MGI:1914293] | Syap1 | Enriched in dLGN | 1.002 | 0.002304 | 90.22 |
| ENSMUSG00000021294 | kinesin family member 26A [Source:MGI Symbol;Acc:MGI:2447072] | Kif26a | Enriched in dLGN | 1.001 | 0.00008866 | 99.43 |
| ENSMUSG00000030844 | regulator of G-protein signalling 10 [Source:MGI Symbol;Acc:MGI:1915115] | Rgs10 | Enriched in dLGN | 1.001 | 0.04637 | 70.22 |
| ENSMUSG00000026883 | disabled 2 interacting protein [Source:MGI Symbol;Acc:MGI:1916851] | Dab2ip | Enriched in retina | -1.001 | 6.894E-09 | 820.5 |
| ENSMUSG00000020814 | matrix-remodelling associated 7 [Source:MGI Symbol;Acc:MGI:1914872] | Mxra7 | Enriched in retina | -1.012 | 0.001808 | 522.1 |
| ENSMUSG00000042570 | MIER family member 2 [Source:MGI Symbol;Acc:MGI:1917677] | Mier2 | Enriched in retina | -1.017 | 0.0002694 | 131.4 |
| ENSMUSG00000010936 | Vac14 homolog (S. cerevisiae) [Source:MGI Symbol;Acc:MGI:2157980] | Vac14 | Enriched in retina | -1.024 | 0.0001442 | 108.7 |
| ENSMUSG00000052373 | membrane protein, palmitoylated 3 (MAGUK p55 subfamily member 3) [Source:MGI Symbol;Acc:MGI:1328354] | Mpp3 | Enriched in retina | -1.026 | 0.0005101 | 85.12 |
| ENSMUSG00000031442 | mcf.2 transforming sequence-like [Source:MGI Symbol;Acc:MGI:103263] | Mcf2l | Enriched in retina | -1.047 | 1.121E-07 | 987.2 |
| ENSMUSG00000007036 | abhydrolase domain containing 16A [Source:MGI Symbol;Acc:MGI:99476] | Abhd16a | Enriched in retina | -1.05 | 0.00003262 | 134 |
| ENSMUSG00000030621 | malic enzyme 3, NADP(+)-dependent, mitochondrial [Source:MGI Symbol;Acc:MGI:1916679] | Me3 | Enriched in retina | -1.059 | 0.009135 | 69.42 |
| ENSMUSG00000034163 | zinc finger, C3H1-type containing [Source:MGI Symbol;Acc:MGI:2446143] | Zfc3h1 | Enriched in retina | -1.06 | 0.001973 | 428.2 |
| ENSMUSG00000022377 | ArfGAP with SH3 domain, ankyrin repeat and PH domain1 [Source:MGI Symbol;Acc:MGI:1342335] | Asap1 | Enriched in retina | -1.064 | 0.0001892 | 410.6 |
| ENSMUSG00000030314 | autophagy related 7 [Source:MGI Symbol;Acc:MGI:1921494] | Atg7 | Enriched in retina | -1.072 | 0.00008425 | 93.47 |
| ENSMUSG00000052752 | TNF receptor-associated factor 7 [Source:MGI Symbol;Acc:MGI:3042141] | Traf7 | Enriched in retina | -1.074 | 0.0002088 | 103.5 |
| ENSMUSG00000036980 | TATA-box binding protein associated factor 6 [Source:MGI Symbol;Acc:MGI:109129] | Taf6 | Enriched in retina | -1.084 | 0.00001319 | 74.4 |
| ENSMUSG00000036062 | PHD finger protein 24 [Source:MGI Symbol;Acc:MGI:2140712] | Phf24 | Enriched in retina | -1.086 | 0.00002704 | 370 |
| ENSMUSG00000039461 | T cell leukemia translocation altered gene [Source:MGI Symbol;Acc:MGI:1918829] | Tcta | Enriched in retina | -1.095 | 0.001376 | 50.88 |
| ENSMUSG00000042115 | kelch domain containing 8A [Source:MGI Symbol;Acc:MGI:2442630] | Klhdc8a | Enriched in retina | -1.107 | 0.0001411 | 181.2 |
| ENSMUSG00000025155 | dihydrouridine synthase 1-like (S. cerevisiae) [Source:MGI Symbol;Acc:MGI:1915980] | Dus1l | Enriched in retina | -1.13 | 0.001868 | 60.74 |
| ENSMUSG00000035382 | proprotein convertase subtilisin/kexin type 7 [Source:MGI Symbol;Acc:MGI:107421] | Pcsk7 | Enriched in retina | -1.146 | 0.00009171 | 88.69 |
| ENSMUSG00000028854 | solute carrier family 9 (sodium/hydrogen exchanger), member 1 [Source:MGI Symbol;Acc:MGI:102462] | Slc9a1 | Enriched in retina | -1.149 | 1.419E-07 | 175.5 |
| ENSMUSG00000022475 | histone deacetylase 7 [Source:MGI Symbol;Acc:MGI:1891835] | Hdac7 | Enriched in retina | -1.178 | 0.002471 | 108.1 |
| ENSMUSG00000042581 | thrombospondin, type I, domain containing 7B [Source:MGI Symbol;Acc:MGI:2443925] | Thsd7b | Enriched in retina | -1.193 | 0.009887 | 235.9 |
| ENSMUSG00000033717 | adrenergic receptor, alpha 2a [Source:MGI Symbol;Acc:MGI:87934] | Adra2a | Enriched in retina | -1.198 | 0.0005735 | 85.56 |
| ENSMUSG00000040565 | B-TFIID TATA-box binding protein associated factor 1 [Source:MGI Symbol;Acc:MGI:2147538] | Btaf1 | Enriched in retina | -1.198 | 0.0003097 | 279.3 |
| ENSMUSG00000021375 | kinesin family member 13A [Source:MGI Symbol;Acc:MGI:1098264] | Kif13a | Enriched in retina | -1.203 | 5.972E-12 | 781.7 |
| ENSMUSG00000000326 | catechol-O-methyltransferase [Source:MGI Symbol;Acc:MGI:88470] | Comt | Enriched in retina | -1.207 | 0.00004123 | 103.3 |
| ENSMUSG00000022885 | beta galactoside alpha 2,6 sialyltransferase 1 [Source:MGI Symbol;Acc:MGI:108470] | St6gal1 | Enriched in retina | -1.224 | 0.001301 | 86.45 |
| ENSMUSG00000012296 | tight junction associated protein 1 [Source:MGI Symbol;Acc:MGI:1921344] | Tjap1 | Enriched in retina | -1.23 | 0.005604 | 89.08 |
| ENSMUSG00000028104 | polymerase (RNA) III (DNA directed) polypeptide G like [Source:MGI Symbol;Acc:MGI:1917120] | Polr3gl | Enriched in retina | -1.237 | 0.00319 | 85.99 |
| ENSMUSG00000039183 | nucleotide binding protein 2 [Source:MGI Symbol;Acc:MGI:1347072] | Nubp2 | Enriched in retina | -1.239 | 0.003833 | 28.88 |
| ENSMUSG00000090799 | kelch-like 33 [Source:MGI Symbol;Acc:MGI:3644593] | Klhl33 | Enriched in retina | -1.248 | 0.02744 | 42.5 |
| ENSMUSG00000044505 | leucine rich repeat and Ig domain containing 4 [Source:MGI Symbol;Acc:MGI:2444651] | Lingo4 | Enriched in retina | -1.27 | 0.02661 | 15.71 |
| ENSMUSG00000026494 | kinesin family member 26B [Source:MGI Symbol;Acc:MGI:2447076] | Kif26b | Enriched in retina | -1.28 | 0.0001601 | 391.1 |
| ENSMUSG00000024187 | family with sequence similarity 234, member A [Source:MGI Symbol;Acc:MGI:2146854] | Fam234a | Enriched in retina | -1.289 | 0.000002921 | 87.13 |
| ENSMUSG00000026411 | transmembrane protein 9 [Source:MGI Symbol;Acc:MGI:1913491] | Tmem9 | Enriched in retina | -1.301 | 0.00000201 | 120.8 |
| ENSMUSG00000035392 | DENN/MADD domain containing 1A [Source:MGI Symbol;Acc:MGI:2442794] | Dennd1a | Enriched in retina | -1.315 | 0.0005341 | 395.9 |
| ENSMUSG00000050822 | solute carrier family 29 (nucleoside transporters), member 4 [Source:MGI Symbol;Acc:MGI:2385330] | Slc29a4 | Enriched in retina | -1.329 | 0.000596 | 126.5 |
| ENSMUSG00000039683 | sidekick cell adhesion molecule 1 [Source:MGI Symbol;Acc:MGI:2444413] | Sdk1 | Enriched in retina | -1.345 | 0.003782 | 374.1 |
| ENSMUSG00000010066 | calcium channel, voltage-dependent, alpha 2/delta subunit 2 [Source:MGI Symbol;Acc:MGI:1929813] | Cacna2d2 | Enriched in retina | -1.371 | 0.000001161 | 439.4 |
| ENSMUSG00000058248 | potassium voltage-gated channel, subfamily H (eag-related), member 1 [Source:MGI Symbol;Acc:MGI:1341721] | Kcnh1 | Enriched in retina | -1.382 | 0.001419 | 103 |
| ENSMUSG00000037148 | Rho GTPase activating protein 10 [Source:MGI Symbol;Acc:MGI:1925764] | Arhgap10 | Enriched in retina | -1.413 | 0.0004145 | 198.1 |
| ENSMUSG00000046727 | cysteine-rich transmembrane module containing 1 [Source:MGI Symbol;Acc:MGI:1913310] | Cystm1 | Enriched in retina | -1.43 | 0.03827 | 23.26 |
| ENSMUSG00000072476 | predicted pseudogene 9008 [Source:MGI Symbol;Acc:MGI:3644000] | Gm9008 | Enriched in retina | -1.48 | 0.02748 | 8.19 |
| ENSMUSG00000020701 | transmembrane protein 132E [Source:MGI Symbol;Acc:MGI:2685490] | Tmem132e | Enriched in retina | -1.486 | 0.000004695 | 113.1 |
| ENSMUSG00000044469 | tumor necrosis factor, alpha-induced protein 8-like 1 [Source:MGI Symbol;Acc:MGI:1913693] | Tnfaip8l1 | Enriched in retina | -1.494 | 0.02343 | 9.788 |
| ENSMUSG00000032468 | armadillo repeat containing 8 [Source:MGI Symbol;Acc:MGI:1921375] | Armc8 | Enriched in retina | -1.497 | 0.000000429 | 227.6 |
| ENSMUSG00000026463 | ATPase, Ca++ transporting, plasma membrane 4 [Source:MGI Symbol;Acc:MGI:88111] | Atp2b4 | Enriched in retina | -1.503 | 0.00008047 | 561.4 |
| ENSMUSG00000046321 | heparan sulfate (glucosamine) 3-O-sulfotransferase 2 [Source:MGI Symbol;Acc:MGI:1333802] | Hs3st2 | Enriched in retina | -1.509 | 0.001438 | 41.56 |
| ENSMUSG00000022894 | a disintegrin-like and metallopeptidase (reprolysin type) with thrombospondin type 1 motif, 5 (aggrecanase-2) [Source:MGI Symbol;Acc:MGI:13463 | Adamts5 | Enriched in retina | -1.522 | 0.0002554 | 56.57 |
| ENSMUSG00000031684 | solute carrier family 10 (sodium/bile acid cotransporter family), member 7 [Source:MGI Symbol;Acc:MGI:1924025] | Slc10a7 | Enriched in retina | -1.535 | 0.001321 | 73.07 |
| ENSMUSG00000022999 | limb region 1 like [Source:MGI Symbol;Acc:MGI:1289247] | Lmbr1l | Enriched in retina | -1.543 | 0.002695 | 52.69 |
| ENSMUSG00000026778 | protein kinase C, theta [Source:MGI Symbol;Acc:MGI:97601] | Prkcq | Enriched in retina | -1.564 | 0.0006667 | 111.9 |
| ENSMUSG00000003746 | mannosidase 1, alpha [Source:MGI Symbol;Acc:MGI:104677] | Man1a | Enriched in retina | -1.598 | 0.0003663 | 79.36 |
| ENSMUSG00000020734 | glutamate receptor, ionotropic, NMDA2C (epsilon 3) [Source:MGI Symbol;Acc:MGI:95822] | Grin2c | Enriched in retina | -1.608 | 0.003354 | 127.2 |
| ENSMUSG00000036585 | fibroblast growth factor 1 [Source:MGI Symbol;Acc:MGI:95515] | Fgf1 | Enriched in retina | -1.612 | 0.002386 | 148.8 |

| | | | | | | |
|---|---|---|---|---|---|---|
| ENSMUSG00000061048 | cadherin 3 [Source:MGI Symbol;Acc:MGI:88356] | Cdh3 | Enriched in retina | -1.639 | 0.04236 | 8.894 |
| ENSMUSG00000025876 | unc-5 netrin receptor A [Source:MGI Symbol;Acc:MGI:894682] | Unc5a | Enriched in retina | -1.639 | 0.000003855 | 196.1 |
| ENSMUSG00000046997 | splA/ryanodine receptor domain and SOCS box containing 4 [Source:MGI Symbol;Acc:MGI:2183445] | Spsb4 | Enriched in retina | -1.646 | 0.0003065 | 36.7 |
| ENSMUSG00000071379 | hippocalcin-like 1 [Source:MGI Symbol;Acc:MGI:1855689] | Hpcal1 | Enriched in retina | -1.665 | 0.0003916 | 98.63 |
| ENSMUSG00000035407 | KN motif and ankyrin repeat domains 4 [Source:MGI Symbol;Acc:MGI:3043381] | Kank4 | Enriched in retina | -1.666 | 0.01429 | 22.26 |
| ENSMUSG00000001794 | calpain, small subunit 1 [Source:MGI Symbol;Acc:MGI:88266] | Capns1 | Enriched in retina | -1.674 | 0.00001213 | 150.2 |
| ENSMUSG00000041439 | major facilitator superfamily domain containing 6 [Source:MGI Symbol;Acc:MGI:1922925] | Mfsd6 | Enriched in retina | -1.675 | 0.000002301 | 415.4 |
| ENSMUSG00000030110 | ret proto-oncogene [Source:MGI Symbol;Acc:MGI:97902] | Ret | Enriched in retina | -1.712 | 0.000001489 | 188.3 |
| ENSMUSG00000029005 | dorsal inhibitory axon guidance protein [Source:MGI Symbol;Acc:MGI:1917683] | Draxin | Enriched in retina | -1.725 | 0.00006427 | 35.71 |
| ENSMUSG00000075334 | reprimo, TP53 dependent G2 arrest mediator candidate [Source:MGI Symbol;Acc:MGI:1915124] | Rprm | Enriched in retina | -1.737 | 0.0002965 | 67.3 |
| ENSMUSG00000028796 | polyhomeotic 2 [Source:MGI Symbol;Acc:MGI:1860454] | Phc2 | Enriched in retina | -1.772 | 5.196E-12 | 280 |
| ENSMUSG00000023495 | poly(rC) binding protein 4 [Source:MGI Symbol;Acc:MGI:1890471] | Pcbp4 | Enriched in retina | -1.778 | 4.949E-20 | 550 |
| ENSMUSG00000037992 | retinoic acid receptor, alpha [Source:MGI Symbol;Acc:MGI:97856] | Rara | Enriched in retina | -1.787 | 4.809E-13 | 152.2 |
| ENSMUSG00000043391 | RIKEN cDNA 2510009E07 gene [Source:MGI Symbol;Acc:MGI:1919440] | 2510009E07Rik | Enriched in retina | -1.792 | 6.248E-22 | 739.1 |
| ENSMUSG00000033209 | tetratricopeptide repeat domain 28 [Source:MGI Symbol;Acc:MGI:2140873] | Ttc28 | Enriched in retina | -1.799 | 2.089E-08 | 597.6 |
| ENSMUSG00000028080 | LPS-responsive beige-like anchor [Source:MGI Symbol;Acc:MGI:1933162] | Lrba | Enriched in retina | -1.806 | 4.349E-08 | 681.5 |
| ENSMUSG00000068758 | interleukin 3 receptor, alpha chain [Source:MGI Symbol;Acc:MGI:96553] | Il3ra | Enriched in retina | -1.863 | 0.03973 | 7.8 |
| ENSMUSG00000001504 | Iroquois homeobox 2 [Source:MGI Symbol;Acc:MGI:1197526] | Irx2 | Enriched in retina | -1.867 | 0.00112 | 24.13 |
| ENSMUSG00000049625 | TRAF-interacting protein with forkhead-associated domain, family member B [Source:MGI Symbol;Acc:MGI:2385852] | Tifab | Enriched in retina | -1.878 | 0.03319 | 7.994 |
| ENSMUSG00000042129 | Ras association (RalGDS/AF-6) domain family member 4 [Source:MGI Symbol;Acc:MGI:2386853] | Rassf4 | Enriched in retina | -2.098 | 3.791E-07 | 89.04 |
| ENSMUSG00000002058 | unc-119 lipid binding chaperone [Source:MGI Symbol;Acc:MGI:1328357] | Unc119 | Enriched in retina | -2.116 | 3.504E-16 | 481.3 |
| ENSMUSG00000038530 | regulator of G-protein signaling 4 [Source:MGI Symbol;Acc:MGI:108409] | Rgs4 | Enriched in retina | -2.214 | 0.000001322 | 578.5 |
| ENSMUSG00000020810 | cytoglobin [Source:MGI Symbol;Acc:MGI:2149481] | Cygb | Enriched in retina | -2.257 | 1.026E-09 | 213.7 |
| ENSMUSG00000070687 | 5-hydroxytryptamine (serotonin) receptor 1D [Source:MGI Symbol;Acc:MGI:96276] | Htr1d | Enriched in retina | -2.307 | 0.0003023 | 19.28 |
| ENSMUSG00000039137 | whirlin [Source:MGI Symbol;Acc:MGI:2682003] | Whrn | Enriched in retina | -2.367 | 4.201E-15 | 351.2 |
| ENSMUSG00000073991 | cyclic nucleotide binding domain containing 1 [Source:MGI Symbol;Acc:MGI:3650508] | Cnbd1 | Enriched in retina | -2.496 | 0.006339 | 9.142 |
| ENSMUSG00000066189 | calcium channel, voltage-dependent, gamma subunit 3 [Source:MGI Symbol;Acc:MGI:1859165] | Cacng3 | Enriched in retina | -2.5 | 2.295E-13 | 109.7 |
| ENSMUSG00000002980 | basal cell adhesion molecule [Source:MGI Symbol;Acc:MGI:1929940] | Bcam | Enriched in retina | -2.612 | 2.241E-07 | 83.63 |
| ENSMUSG00000031734 | Iroquois related homeobox 3 [Source:MGI Symbol;Acc:MGI:1197522] | Irx3 | Enriched in retina | -2.728 | 0.00002174 | 16.38 |
| ENSMUSG00000039954 | serine/threonine kinase 32A [Source:MGI Symbol;Acc:MGI:2442403] | Stk32a | Enriched in retina | -2.858 | 4.523E-07 | 77.08 |
| ENSMUSG00000041592 | sidekick cell adhesion molecule 2 [Source:MGI Symbol;Acc:MGI:2443847] | Sdk2 | Enriched in retina | -2.867 | 1.343E-21 | 398.3 |
| ENSMUSG00000030830 | integrin alpha L [Source:MGI Symbol;Acc:MGI:96606] | Itgal | Enriched in retina | -2.87 | 0.0025 | 6.323 |
| ENSMUSG00000025432 | advillin [Source:MGI Symbol;Acc:MGI:1333798] | Avil | Enriched in retina | -2.985 | 0.01408 | 4.817 |
| ENSMUSG00000042073 | abhydrolase domain containing 14b [Source:MGI Symbol;Acc:MGI:1923741] | Abhd14b | Enriched in retina | -3.015 | 0.0002786 | 38.48 |
| ENSMUSG00000006764 | tryptophan hydroxylase 2 [Source:MGI Symbol;Acc:MGI:2651811] | Tph2 | Enriched in retina | -3.033 | 0.04814 | 3.884 |
| ENSMUSG00000046719 | neurexophilin 3 [Source:MGI Symbol;Acc:MGI:1336188] | Nxph3 | Enriched in retina | -3.192 | 2.067E-07 | 27.7 |
| ENSMUSG00000090523 | glycophorin C [Source:MGI Symbol;Acc:MGI:1098566] | Gypc | Enriched in retina | -3.207 | 0.01759 | 5.436 |
| ENSMUSG00000037705 | tectorin alpha [Source:MGI Symbol;Acc:MGI:109575] | Tecta | Enriched in retina | -3.335 | 0.003064 | 4.094 |
| ENSMUSG00000034402 | potassium voltage-gated channel, subfamily H (eag-related), member 5 [Source:MGI Symbol;Acc:MGI:3584508] | Kcnh5 | Enriched in retina | -3.346 | 4.656E-18 | 240.3 |
| ENSMUSG00000023484 | peripherin [Source:MGI Symbol;Acc:MGI:97774] | Prph | Enriched in retina | -3.427 | 0.0006701 | 64.1 |
| ENSMUSG00000016624 | PHD finger protein 21B [Source:MGI Symbol;Acc:MGI:2443812] | Phf21b | Enriched in retina | -3.714 | 1.873E-16 | 142.2 |
| ENSMUSG00000031727 | polyamine modulated factor 1 binding protein 1 [Source:MGI Symbol;Acc:MGI:1930136] | Pmfbp1 | Enriched in retina | -3.765 | 0.0008465 | 36.63 |
| ENSMUSG00000032128 | roundabout guidance receptor 3 [Source:MGI Symbol;Acc:MGI:1343102] | Robo3 | Enriched in retina | -3.968 | 0.0001787 | 9.898 |
| ENSMUSG00000056296 | synaptoporin [Source:MGI Symbol;Acc:MGI:1919253] | Synpr | Enriched in retina | -3.98 | 4.401E-15 | 203.6 |
| ENSMUSG00000031737 | Iroquois homeobox 5 [Source:MGI Symbol;Acc:MGI:1859086] | Irx5 | Enriched in retina | -4.074 | 6.756E-12 | 31.09 |
| ENSMUSG00000031586 | RNA binding protein gene with multiple splicing [Source:MGI Symbol;Acc:MGI:1334446] | Rbpms | Enriched in retina | -4.225 | 3.356E-17 | 81.55 |
| ENSMUSG00000096351 | sterile alpha motif domain containing 11 [Source:MGI Symbol;Acc:MGI:2446220] | Samd11 | Enriched in retina | -4.277 | 3.077E-16 | 401.5 |
| ENSMUSG00000004098 | collagen, type V, alpha 3 [Source:MGI Symbol;Acc:MGI:1858212] | Col5a3 | Enriched in retina | -4.454 | 5.68E-14 | 67.35 |
| ENSMUSG00000035296 | sarcoglycan, gamma (dystrophin-associated glycoprotein) [Source:MGI Symbol;Acc:MGI:1346524] | Sgcg | Enriched in retina | -4.481 | 0.003788 | 9.668 |
| ENSMUSG00000058925 | coiled-coil domain containing 192 [Source:MGI Symbol;Acc:MGI:1922694] | Ccdc192 | Enriched in retina | -4.628 | 0.0003573 | 7.152 |
| ENSMUSG00000069372 | cortexin 3 [Source:MGI Symbol;Acc:MGI:3642816] | Ctxn3 | Enriched in retina | -5.192 | 0.00001172 | 38.01 |
| ENSMUSG00000048070 | phosphoinositide-interacting regulator of transient receptor potential channels [Source:MGI Symbol;Acc:MGI:2443635] | Pirt | Enriched in retina | -5.393 | 0.0001966 | 10.02 |
| ENSMUSG00000034115 | sodium channel, voltage-gated, type XI, alpha [Source:MGI Symbol;Acc:MGI:1345149] | Scn11a | Enriched in retina | -5.431 | 0.0001152 | 5.567 |
| ENSMUSG00000032387 | RNA binding protein with multiple splicing 2 [Source:MGI Symbol;Acc:MGI:1919223] | Rbpms2 | Enriched in retina | -5.932 | 2.154E-10 | 47.72 |
| ENSMUSG00000034777 | ventral anterior homeobox 2 [Source:MGI Symbol;Acc:MGI:1346018] | Vax2 | Enriched in retina | -6.094 | 2.78E-10 | 45.1 |
| ENSMUSG00000051279 | growth differentiation factor 6 [Source:MGI Symbol;Acc:MGI:95689] | Gdf6 | Enriched in retina | -6.132 | 0.000006718 | 11.98 |
| ENSMUSG00000002100 | myosin binding protein C, cardiac [Source:MGI Symbol;Acc:MGI:102844] | Mybpc3 | Enriched in retina | -6.616 | 0.00003386 | 4.298 |
| ENSMUSG00000032446 | eomesodermin [Source:MGI Symbol;Acc:MGI:1201683] | Eomes | Enriched in retina | -7.632 | 3.707E-08 | 60.2 |
| ENSMUSG00000067438 | H6 homeobox 1 [Source:MGI Symbol;Acc:MGI:107178] | Hmx1 | Enriched in retina | -7.844 | 4.764E-09 | 37.32 |
| ENSMUSG00000031965 | T-box 20 [Source:MGI Symbol;Acc:MGI:1888496] | Tbx20 | Enriched in retina | -8.136 | 1.187E-08 | 45.45 |
| ENSMUSG00000031688 | POU domain, class 4, transcription factor 2 [Source:MGI Symbol;Acc:MGI:102524] | Pou4f2 | Enriched in retina | -8.959 | 6.793E-11 | 45.2 |
| ENSMUSG00000031738 | Iroquois homeobox 6 [Source:MGI Symbol;Acc:MGI:1927642] | Irx6 | Enriched in retina | -9.404 | 1.444E-12 | 50.97 |

**Contrast numerator: P8_Het_dLGN. Contrast denominator: P8_Het_SCN.**

| ensembl_gene_id | description | mgi_symbol | Significance | deseq_logfc | deseq_adjp | deseq_basemean |
|---|---|---|---|---|---|---|
| ENSMUSG00000018486 | wingless-type MMTV integration site family, member 9B [Source:MGI Symbol;Acc:MGI:1197020] | Wnt9b | Enriched in dLGN | 3.403 | 2.555E-09 | 165.2 |
| ENSMUSG00000060257 | scratch family zinc finger 2 [Source:MGI Symbol;Acc:MGI:2139287] | Scrt2 | Enriched in dLGN | 3.12 | 1.055E-23 | 141.5 |
| ENSMUSG00000064179 | troponin T1, skeletal, slow [Source:MGI Symbol;Acc:MGI:1333868] | Tnnt1 | Enriched in dLGN | 2.914 | 4.201E-09 | 308.7 |
| ENSMUSG00000070570 | solute carrier family 17 (sodium-dependent inorganic phosphate cotransporter), member 7 [Source:MGI Symbol;Acc:MGI:1920211] | Slc17a7 | Enriched in dLGN | 2.722 | 0.000000015 | 1629 |
| ENSMUSG00000048385 | scratch family zinc finger 1 [Source:MGI Symbol;Acc:MGI:2176606] | Scrt1 | Enriched in dLGN | 2.514 | 7.381E-19 | 1666 |
| ENSMUSG00000068696 | G-protein coupled receptor 88 [Source:MGI Symbol;Acc:MGI:1927653] | Gpr88 | Enriched in dLGN | 2.106 | 6.521E-09 | 66.64 |
| ENSMUSG00000039114 | neuritin 1 [Source:MGI Symbol;Acc:MGI:1915654] | Nrn1 | Enriched in dLGN | 1.97 | 0.000001702 | 742.9 |
| ENSMUSG00000020431 | adenylate cyclase 1 [Source:MGI Symbol;Acc:MGI:99677] | Adcy1 | Enriched in dLGN | 1.879 | 1.124E-08 | 5232 |
| ENSMUSG00000041046 | receptor (calcitonin) activity modifying protein 3 [Source:MGI Symbol;Acc:MGI:1860292] | Ramp3 | Enriched in dLGN | 1.76 | 0.00002663 | 121.1 |
| ENSMUSG00000037428 | VGF nerve growth factor inducible [Source:MGI Symbol;Acc:MGI:1343180] | Vgf | Enriched in dLGN | 1.672 | 8.865E-08 | 1249 |
| ENSMUSG00000020806 | rhomboid 5 homolog 2 [Source:MGI Symbol;Acc:MGI:2442473] | Rhbdf2 | Enriched in dLGN | 1.656 | 0.03065 | 43.1 |
| ENSMUSG00000021848 | orthodenticle homeobox 2 [Source:MGI Symbol;Acc:MGI:97451] | Otx2 | Enriched in dLGN | 1.649 | 0.02658 | 248.7 |
| ENSMUSG00000021290 | ATP synthase membrane subunit 6.8PL [Source:MGI Symbol;Acc:MGI:1917507] | Atp5mpl | Enriched in dLGN | 1.637 | 0.0012 | 181.4 |
| ENSMUSG00000029632 | Ndufa4, mitochondrial complex associated [Source:MGI Symbol;Acc:MGI:107686] | Ndufa4 | Enriched in dLGN | 1.633 | 0.00004108 | 183.2 |
| ENSMUSG00000040794 | C1q and tumor necrosis factor related protein 4 [Source:MGI Symbol;Acc:MGI:1914695] | C1qtnf4 | Enriched in dLGN | 1.408 | 4.654E-07 | 207.8 |
| ENSMUSG00000032249 | acidic (leucine-rich) nuclear phosphoprotein 32 family, member A [Source:MGI Symbol;Acc:MGI:108447] | Anp32a | Enriched in dLGN | 1.394 | 3.302E-14 | 803.6 |
| ENSMUSG00000075324 | fidgetin [Source:MGI Symbol;Acc:MGI:1890647] | Fign | Enriched in dLGN | 1.392 | 0.002417 | 251.6 |
| ENSMUSG00000021660 | basic transcription factor 3 [Source:MGI Symbol;Acc:MGI:1202875] | Btf3 | Enriched in dLGN | 1.324 | 0.0000103 | 71.1 |
| ENSMUSG00000016427 | NADH:ubiquinone oxidoreductase subunit A1 [Source:MGI Symbol;Acc:MGI:1929511] | Ndufa1 | Enriched in dLGN | 1.287 | 0.000111 | 84.15 |
| ENSMUSG00000020949 | FK506 binding protein 3 [Source:MGI Symbol;Acc:MGI:1353460] | Fkbp3 | Enriched in dLGN | 1.281 | 0.002016 | 160.5 |
| ENSMUSG00000000088 | cytochrome c oxidase subunit 5A [Source:MGI Symbol;Acc:MGI:88474] | Cox5a | Enriched in dLGN | 1.221 | 0.001749 | 112.1 |
| ENSMUSG00000032435 | dynein cytoplasmic 1 light intermediate chain 1 [Source:MGI Symbol;Acc:MGI:2135610] | Dync1li1 | Enriched in dLGN | 1.205 | 4.742E-17 | 559.8 |
| ENSMUSG00000063253 | short coiled-coil protein [Source:MGI Symbol;Acc:MGI:1927654] | Scoc | Enriched in dLGN | 1.197 | 0.000007368 | 113.5 |
| ENSMUSG00000022892 | amyloid beta (A4) precursor protein [Source:MGI Symbol;Acc:MGI:88059] | App | Enriched in dLGN | 1.195 | 0.0005464 | 4592 |
| ENSMUSG00000031104 | RAB33A, member RAS oncogene family [Source:MGI Symbol;Acc:MGI:109493] | Rab33a | Enriched in dLGN | 1.194 | 0.007422 | 43.76 |
| ENSMUSG00000066150 | solute carrier family 31, member 1 [Source:MGI Symbol;Acc:MGI:1333843] | Slc31a1 | Enriched in dLGN | 1.183 | 0.01796 | 94.7 |
| ENSMUSG00000019943 | ATPase, Ca++ transporting, plasma membrane 1 [Source:MGI Symbol;Acc:MGI:104653] | Atp2b1 | Enriched in dLGN | 1.182 | 3.124E-09 | 2706 |
| ENSMUSG00000022180 | solute carrier family 7 (cationic amino acid transporter, y+ system), member 8 [Source:MGI Symbol;Acc:MGI:1355323] | Slc7a8 | Enriched in dLGN | 1.171 | 0.0001829 | 249.8 |
| ENSMUSG00000024038 | NADH:ubiquinone oxidoreductase core subunit V3 [Source:MGI Symbol;Acc:MGI:1890894] | Ndufv3 | Enriched in dLGN | 1.16 | 0.00002085 | 374.2 |
| ENSMUSG00000027203 | deoxyuridine triphosphatase [Source:MGI Symbol;Acc:MGI:1346051] | Dut | Enriched in dLGN | 1.137 | 0.00002453 | 92.41 |
| ENSMUSG00000028832 | stathmin 1 [Source:MGI Symbol;Acc:MGI:96739] | Stmn1 | Enriched in dLGN | 1.107 | 0.006983 | 644 |
| ENSMUSG00000048483 | zinc finger, DHHC-type containing 22 [Source:MGI Symbol;Acc:MGI:2685108] | Zdhhc22 | Enriched in dLGN | 1.103 | 0.0003555 | 612.1 |
| ENSMUSG00000036751 | cytochrome c oxidase, subunit 6B1 [Source:MGI Symbol;Acc:MGI:107460] | Cox6b1 | Enriched in dLGN | 1.103 | 0.000761 | 143.6 |
| ENSMUSG00000062760 | shisa like 1 [Source:MGI Symbol;Acc:MGI:1919551] | Shisal1 | Enriched in dLGN | 1.086 | 7.441E-07 | 955.5 |
| ENSMUSG00000061518 | cytochrome c oxidase subunit 5B [Source:MGI Symbol;Acc:MGI:88475] | Cox5b | Enriched in dLGN | 1.063 | 0.001326 | 455.8 |
| ENSMUSG00000030127 | COP9 signalosome subunit 7A [Source:MGI Symbol;Acc:MGI:1349400] | Cops7a | Enriched in dLGN | 1.019 | 0.0004838 | 167 |
| ENSMUSG00000029066 | mitochondrial ribosomal protein L20 [Source:MGI Symbol;Acc:MGI:2137221] | Mrpl20 | Enriched in dLGN | 1.002 | 0.01227 | 48.19 |
| ENSMUSG00000076432 | tyrosine 3-monooxygenase/tryptophan 5-monooxygenase activation protein theta [Source:MGI Symbol;Acc:MGI:891963] | Ywhaq | Enriched in dLGN | 1.001 | 0.008313 | 236.1 |
| ENSMUSG00000023169 | solute carrier family 38, member 1 [Source:MGI Symbol;Acc:MGI:2145895] | Slc38a1 | Enriched in SCN | -1.009 | 0.0001784 | 1408 |
| ENSMUSG00000029992 | glutamine fructose-6-phosphate transaminase 1 [Source:MGI Symbol;Acc:MGI:95698] | Gfpt1 | Enriched in SCN | -1.012 | 0.00002619 | 537.8 |
| ENSMUSG00000021687 | secretory carrier membrane protein 1 [Source:MGI Symbol;Acc:MGI:1349480] | Scamp1 | Enriched in SCN | -1.025 | 0.0004763 | 570.2 |
| ENSMUSG00000024921 | SWI/SNF related, matrix associated, actin dependent regulator of chromatin, subfamily a, member 2 [Source:MGI Symbol;Acc:MGI:996 | Smarca2 | Enriched in SCN | -1.029 | 0.000003716 | 1435 |
| ENSMUSG00000042873 | lipoma HMGIC fusion partner-like protein 4 [Source:MGI Symbol;Acc:MGI:3057108] | Lhfpl4 | Enriched in SCN | -1.049 | 0.000276 | 1362 |
| ENSMUSG00000007670 | KH-type splicing regulatory protein [Source:MGI Symbol;Acc:MGI:1336214] | Khsrp | Enriched in SCN | -1.051 | 0.00001994 | 695.6 |
| ENSMUSG00000024579 | prenylcysteine oxidase 1 like [Source:MGI Symbol;Acc:MGI:3606062] | Pcyox1l | Enriched in SCN | -1.053 | 0.004103 | 127.8 |
| ENSMUSG00000027983 | cytochrome P450, family 2, subfamily u, polypeptide 1 [Source:MGI Symbol;Acc:MGI:1918769] | Cyp2u1 | Enriched in SCN | -1.056 | 0.0005259 | 119.4 |
| ENSMUSG00000108358 | predicted gene 44509 [Source:MGI Symbol;Acc:MGI:5753085] | Gm44509 | Enriched in SCN | -1.062 | 0.04022 | 119 |
| ENSMUSG00000027879 | SEC22 homolog B, vesicle trafficking protein [Source:MGI Symbol;Acc:MGI:1338759] | Sec22b | Enriched in SCN | -1.07 | 0.01019 | 253.4 |
| ENSMUSG00000052557 | giant axonal neuropathy [Source:MGI Symbol;Acc:MGI:1890619] | Gan | Enriched in SCN | -1.076 | 0.0002929 | 388.9 |
| ENSMUSG00000034940 | synergin, gamma [Source:MGI Symbol;Acc:MGI:1354742] | Synrg | Enriched in SCN | -1.079 | 4.235E-10 | 418.7 |
| ENSMUSG00000075703 | selenoprotein I [Source:MGI Symbol;Acc:MGI:107898] | Selenoi | Enriched in SCN | -1.096 | 0.0001392 | 269.3 |
| ENSMUSG00000060935 | transmembrane protein 263 [Source:MGI Symbol;Acc:MGI:2143652] | Tmem263 | Enriched in SCN | -1.11 | 0.0004029 | 126.6 |
| ENSMUSG00000022521 | CREB binding protein [Source:MGI Symbol;Acc:MGI:1098280] | Crebbp | Enriched in SCN | -1.11 | 0.00001383 | 2003 |
| ENSMUSG00000032425 | zinc finger protein 949 [Source:MGI Symbol;Acc:MGI:1918890] | Zfp949 | Enriched in SCN | -1.111 | 0.00876 | 260.3 |
| ENSMUSG00000024400 | WD repeat domain 33 [Source:MGI Symbol;Acc:MGI:1921570] | Wdr33 | Enriched in SCN | -1.135 | 0.00001822 | 635.9 |
| ENSMUSG00000021770 | sterile alpha motif domain containing 8 [Source:MGI Symbol;Acc:MGI:1914880] | Samd8 | Enriched in SCN | -1.148 | 0.0004542 | 676.8 |
| ENSMUSG00000069662 | myristoylated alanine rich protein kinase C substrate [Source:MGI Symbol;Acc:MGI:96907] | Marcks | Enriched in SCN | -1.163 | 0.01281 | 2471 |
| ENSMUSG00000032905 | autophagy related 12 [Source:MGI Symbol;Acc:MGI:1914776] | Atg12 | Enriched in SCN | -1.166 | 0.00005589 | 183.2 |
| ENSMUSG00000020922 | LSM12 homolog [Source:MGI Symbol;Acc:MGI:1919592] | Lsm12 | Enriched in SCN | -1.225 | 0.00412 | 397.5 |
| ENSMUSG00000049728 | zinc finger protein 668 [Source:MGI Symbol;Acc:MGI:2442943] | Zfp668 | Enriched in SCN | -1.237 | 0.0001969 | 123.8 |
| ENSMUSG00000041040 | family with sequence similarity 117, member B [Source:MGI Symbol;Acc:MGI:1920000] | Fam117b | Enriched in SCN | -1.265 | 0.000004017 | 418.6 |
| ENSMUSG00000034377 | tubby like protein 4 [Source:MGI Symbol;Acc:MGI:1916092] | Tulp4 | Enriched in SCN | -1.322 | 0.0133 | 2896 |
| ENSMUSG00000031226 | polysaccharide biosynthesis domain containing 1 [Source:MGI Symbol;Acc:MGI:1914933] | Pbdc1 | Enriched in SCN | -1.33 | 0.002338 | 236 |
| ENSMUSG00000039542 | neural cell adhesion molecule 1 [Source:MGI Symbol;Acc:MGI:97281] | Ncam1 | Enriched in SCN | -1.358 | 0.0000923 | 5804 |
| ENSMUSG00000024261 | synaptotagmin IV [Source:MGI Symbol;Acc:MGI:101759] | Syt4 | Enriched in SCN | -1.375 | 0.00009877 | 328.7 |
| ENSMUSG00000039770 | yippee like 5 [Source:MGI Symbol;Acc:MGI:1916937] | Ypel5 | Enriched in SCN | -1.378 | 0.0004002 | 207.1 |
| ENSMUSG00000022048 | dihydropyrimidinase-like 2 [Source:MGI Symbol;Acc:MGI:1349763] | Dpysl2 | Enriched in SCN | -1.423 | 0.0006753 | 6238 |
| ENSMUSG00000017929 | UDP-Gal:betaGlcNAc beta 1,4-galactosyltransferase, polypeptide 5 [Source:MGI Symbol;Acc:MGI:1927169] | B4galt5 | Enriched in SCN | -1.433 | 0.000002492 | 566.4 |
| ENSMUSG00000040928 | S100P binding protein [Source:MGI Symbol;Acc:MGI:1921898] | S100pbp | Enriched in SCN | -1.437 | 8.611E-07 | 257.9 |
| ENSMUSG00000028106 | regulation of nuclear pre-mRNA domain containing 2 [Source:MGI Symbol;Acc:MGI:1922387] | Rprd2 | Enriched in SCN | -1.451 | 1.06E-08 | 443.4 |
| ENSMUSG00000028381 | UDP-glucose ceramide glucosyltransferase [Source:MGI Symbol;Acc:MGI:1332243] | Ugcg | Enriched in SCN | -1.475 | 0.0003471 | 361.5 |
| ENSMUSG00000010554 | methyltransferase like 16 [Source:MGI Symbol;Acc:MGI:1914743] | Mettl16 | Enriched in SCN | -1.506 | 0.000001918 | 322.2 |
| ENSMUSG00000042105 | inositol polyphosphate-5-phosphatase F [Source:MGI Symbol;Acc:MGI:2141867] | Inpp5f | Enriched in SCN | -1.559 | 1.414E-07 | 994.9 |
| ENSMUSG00000031302 | neuroligin 3 [Source:MGI Symbol;Acc:MGI:2444609] | Nlgn3 | Enriched in SCN | -1.566 | 0.00009745 | 1941 |
| ENSMUSG00000026020 | NOP58 ribonucleoprotein [Source:MGI Symbol;Acc:MGI:1933184] | Nop58 | Enriched in SCN | -1.579 | 2.424E-08 | 363.9 |
| ENSMUSG00000044667 | phospholipid phosphatase related 4 [Source:MGI Symbol;Acc:MGI:106530] | Plppr4 | Enriched in SCN | -1.632 | 7.347E-08 | 669.3 |
| ENSMUSG00000021139 | predicted gene 20498 [Source:MGI Symbol;Acc:MGI:5141963] | Gm20498 | Enriched in SCN | -2.055 | 0.00004168 | 272.3 |
| ENSMUSG00000032076 | cell adhesion molecule 1 [Source:MGI Symbol;Acc:MGI:1889272] | Cadm1 | Enriched in SCN | -2.056 | 5.221E-14 | 2724 |
| ENSMUSG00000095139 | POU domain, class 3, transcription factor 2 [Source:MGI Symbol;Acc:MGI:101895] | Pou3f2 | Enriched in SCN | -2.316 | 2.243E-10 | 124.7 |
| ENSMUSG00000059854 | HYDIN, axonemal central pair apparatus protein [Source:MGI Symbol;Acc:MGI:2389007] | Hydin | Enriched in SCN | -3.594 | 3.872E-09 | 115.1 |
| ENSMUSG00000021685 | orthopedia homeobox [Source:MGI Symbol;Acc:MGI:99835] | Otp | Enriched in SCN | -11.09 | 0.002454 | 81.46 |

**Contrast numerator: P8_KO_dLGN. Contrast denominator: P8_KO_SCN.**

| ensembl_gene_id | description | mgi_symbol | Significance | deseq_logfc | deseq_adjp | deseq_basemean |
|---|---|---|---|---|---|---|
| ENSMUSG00000041144 | dynein, axonemal, heavy chain 7B [Source:MGI Symbol;Acc:MGI:2684953] | Dnah7b | Enriched in dLGN | 3.916 | 0.0006168 | 449.4 |
| ENSMUSG00000027716 | transient receptor potential cation channel, subfamily C, member 3 [Source:MGI Symbol;Acc:MGI:109526] | Trpc3 | Enriched in dLGN | 3.764 | 7.04E-30 | 212.4 |
| ENSMUSG00000065968 | interferon induced transmembrane protein 7 [Source:MGI Symbol;Acc:MGI:1921732] | Ifitm7 | Enriched in dLGN | 3.688 | 0.00003836 | 15.26 |
| ENSMUSG00000020000 | monooxygenase, DBH-like 1 [Source:MGI Symbol;Acc:MGI:1921582] | Moxd1 | Enriched in dLGN | 3.353 | 0.0001891 | 20.89 |
| ENSMUSG00000068696 | G-protein coupled receptor 88 [Source:MGI Symbol;Acc:MGI:1927653] | Gpr88 | Enriched in dLGN | 3.097 | 2.756E-11 | 66.64 |
| ENSMUSG00000002908 | potassium intermediate/small conductance calcium-activated channel, subfamily N, member 1 [Source:MGI Symbol;Acc:MGI:1933993] | Kcnn1 | Enriched in dLGN | 3.003 | 7.806E-19 | 72.47 |
| ENSMUSG00000073406 | histocompatibility 2, blastocyst [Source:MGI Symbol;Acc:MGI:892004] | H2-Bl | Enriched in dLGN | 2.895 | 0.0003504 | 18.45 |
| ENSMUSG00000028744 | solute carrier family 66 member 1 [Source:MGI Symbol;Acc:MGI:2384837] | Slc66a1 | Enriched in dLGN | 2.893 | 0.00001646 | 195.7 |
| ENSMUSG00000032965 | intraflagellar transport 57 [Source:MGI Symbol;Acc:MGI:1921166] | Ift57 | Enriched in dLGN | 2.848 | 2.918E-31 | 254.5 |
| ENSMUSG00000051864 | TBC1 domain family, member 22a [Source:MGI Symbol;Acc:MGI:1289265] | Tbc1d22a | Enriched in dLGN | 2.354 | 0.005099 | 351.4 |
| ENSMUSG00000014313 | cytochrome c oxidase subunit 6C [Source:MGI Symbol;Acc:MGI:104614] | Cox6c | Enriched in dLGN | 2.259 | 4.972E-08 | 345.4 |
| ENSMUSG00000062070 | phosphoglycerate kinase 1 [Source:MGI Symbol;Acc:MGI:97555] | Pgk1 | Enriched in dLGN | 2.242 | 1.136E-09 | 300 |
| ENSMUSG00000040794 | C1q and tumor necrosis factor related protein 4 [Source:MGI Symbol;Acc:MGI:1914695] | C1qtnf4 | Enriched in dLGN | 2.201 | 7.907E-15 | 207.8 |
| ENSMUSG00000028461 | coiled-coil domain containing 107 [Source:MGI Symbol;Acc:MGI:1913423] | Ccdc107 | Enriched in dLGN | 2.19 | 0.04779 | 309.5 |
| ENSMUSG00000064288 | H4 clustered histone 12 [Source:MGI Symbol;Acc:MGI:2448439] | H4c12 | Enriched in dLGN | 2.126 | 0.01837 | 10.87 |
| ENSMUSG00000005357 | solute carrier family 1 (high affinity aspartate/glutamate transporter), member 6 [Source:MGI Symbol;Acc:MGI:1096331] | Slc1a6 | Enriched in dLGN | 2.06 | 7.113E-07 | 52.8 |
| ENSMUSG00000039166 | A kinase (PRKA) anchor protein 7 [Source:MGI Symbol;Acc:MGI:1859150] | Akap7 | Enriched in dLGN | 2.045 | 3.866E-07 | 348.8 |
| ENSMUSG00000096257 | coiled-coil glutamate-rich protein 2 [Source:MGI Symbol;Acc:MGI:3645242] | Ccer2 | Enriched in dLGN | 2.034 | 0.00378 | 12.8 |
| ENSMUSG00000091405 | H4 clustered histone 14 [Source:MGI Symbol;Acc:MGI:2140113] | H4c14 | Enriched in dLGN | 2.028 | 0.0002026 | 36.46 |
| ENSMUSG00000011752 | phosphoglycerate mutase 1 [Source:MGI Symbol;Acc:MGI:97552] | Pgam1 | Enriched in dLGN | 1.951 | 3.131E-07 | 217.3 |
| ENSMUSG00000032249 | acidic (leucine-rich) nuclear phosphoprotein 32 family, member A [Source:MGI Symbol;Acc:MGI:108447] | Anp32a | Enriched in dLGN | 1.947 | 1.308E-24 | 803.6 |
| ENSMUSG00000016252 | ATP synthase, H+ transporting, mitochondrial F1 complex, epsilon subunit [Source:MGI Symbol;Acc:MGI:1855697] | Atp5e | Enriched in dLGN | 1.925 | 0.000004388 | 153.2 |
| ENSMUSG00000024935 | solute carrier family 1 (neuronal/epithelial high affinity glutamate transporter, system Xag), member 1 [Source:MGI Symbol;Acc:MGI:1050 | Slc1a1 | Enriched in dLGN | 1.909 | 1.348E-12 | 323.7 |
| ENSMUSG00000022892 | amyloid beta (A4) precursor protein [Source:MGI Symbol;Acc:MGI:88059] | App | Enriched in dLGN | 1.895 | 1.014E-08 | 4592 |
| ENSMUSG00000046442 | protein phosphatase 1E (PP2C domain containing) [Source:MGI Symbol;Acc:MGI:2444096] | Ppm1e | Enriched in dLGN | 1.891 | 4.275E-13 | 844.2 |
| ENSMUSG00000020814 | matrix-remodelling associated 7 [Source:MGI Symbol;Acc:MGI:1914872] | Mxra7 | Enriched in dLGN | 1.875 | 1.661E-08 | 522.1 |
| ENSMUSG00000030685 | potassium channel tetramerisation domain containing 13 [Source:MGI Symbol;Acc:MGI:1923739] | Kctd13 | Enriched in dLGN | 1.864 | 8.717E-11 | 195.9 |
| ENSMUSG00000084786 | ubiquitin-like 5 [Source:MGI Symbol;Acc:MGI:1913427] | Ubl5 | Enriched in dLGN | 1.834 | 3.74E-08 | 89.51 |
| ENSMUSG00000032435 | dynein cytoplasmic 1 light intermediate chain 1 [Source:MGI Symbol;Acc:MGI:2135610] | Dync1li1 | Enriched in dLGN | 1.812 | 5.998E-32 | 559.8 |
| ENSMUSG00000020029 | nudix (nucleoside diphosphate linked moiety X)-type motif 4 [Source:MGI Symbol;Acc:MGI:1918457] | Nudt4 | Enriched in dLGN | 1.757 | 1.243E-09 | 752.5 |
| ENSMUSG00000021270 | heat shock protein 90, alpha (cytosolic), class A member 1 [Source:MGI Symbol;Acc:MGI:96250] | Hsp90aa1 | Enriched in dLGN | 1.714 | 0.000002431 | 3186 |
| ENSMUSG00000053192 | myeloid/lymphoid or mixed-lineage leukemia; translocated to, 11 [Source:MGI Symbol;Acc:MGI:1929671] | Mllt11 | Enriched in dLGN | 1.704 | 5.008E-10 | 499.1 |
| ENSMUSG00000040105 | phospholipid phosphatase 6 [Source:MGI Symbol;Acc:MGI:1921661] | Plpp6 | Enriched in dLGN | 1.7 | 1.081E-07 | 114.9 |
| ENSMUSG00000054843 | attractin like 1 [Source:MGI Symbol;Acc:MGI:2147749] | Atrnl1 | Enriched in dLGN | 1.676 | 0.000001039 | 629.5 |
| ENSMUSG00000027104 | activating transcription factor 2 [Source:MGI Symbol;Acc:MGI:109349] | Atf2 | Enriched in dLGN | 1.662 | 0.0009202 | 703.5 |
| ENSMUSG00000050248 | EvC ciliary complex subunit 2 [Source:MGI Symbol;Acc:MGI:1915775] | Evc2 | Enriched in dLGN | 1.66 | 2.039E-08 | 56.13 |
| ENSMUSG00000056305 | ubiquitin specific peptidase 39 [Source:MGI Symbol;Acc:MGI:107622] | Usp39 | Enriched in dLGN | 1.644 | 1.275E-07 | 113.7 |
| ENSMUSG00000061518 | cytochrome c oxidase subunit 5B [Source:MGI Symbol;Acc:MGI:88475] | Cox5b | Enriched in dLGN | 1.628 | 4.574E-07 | 455.8 |
| ENSMUSG00000049422 | coiled-coil-helix-coiled-coil-helix domain containing 10 [Source:MGI Symbol;Acc:MGI:2143558] | Chchd10 | Enriched in dLGN | 1.625 | 0.00001205 | 224.5 |
| ENSMUSG00000028793 | ring finger protein 19B [Source:MGI Symbol;Acc:MGI:1922484] | Rnf19b | Enriched in dLGN | 1.619 | 8.836E-08 | 239.7 |
| ENSMUSG00000079184 | M-phase phosphoprotein 8 [Source:MGI Symbol;Acc:MGI:1922589] | Mphosph8 | Enriched in dLGN | 1.616 | 0.0005872 | 685.9 |
| ENSMUSG00000020116 | partner of NOB1 homolog [Source:MGI Symbol;Acc:MGI:1913499] | Pno1 | Enriched in dLGN | 1.608 | 0.0003587 | 43.22 |
| ENSMUSG00000025869 | NOP16 nucleolar protein [Source:MGI Symbol;Acc:MGI:107862] | Nop16 | Enriched in dLGN | 1.563 | 0.00006046 | 53.74 |
| ENSMUSG00000020107 | anaphase promoting complex subunit 16 [Source:MGI Symbol;Acc:MGI:1289325] | Anapc16 | Enriched in dLGN | 1.559 | 0.000105 | 134.3 |
| ENSMUSG00000024907 | galanin and GMAP prepropeptide [Source:MGI Symbol;Acc:MGI:95637] | Gal | Enriched in dLGN | 1.542 | 0.02195 | 30 |
| ENSMUSG00000021892 | SH3-domain binding protein 5 (BTK-associated) [Source:MGI Symbol;Acc:MGI:1344391] | Sh3bp5 | Enriched in dLGN | 1.535 | 0.000006535 | 251.2 |
| ENSMUSG00000046711 | high mobility group AT-hook 1 [Source:MGI Symbol;Acc:MGI:96160] | Hmga1 | Enriched in dLGN | 1.504 | 0.00007675 | 154.5 |
| ENSMUSG00000039221 | ribosomal protein L22 like 1 [Source:MGI Symbol;Acc:MGI:1915278] | Rpl22l1 | Enriched in dLGN | 1.492 | 0.009757 | 25.35 |
| ENSMUSG00000030704 | RAB6A, member RAS oncogene family [Source:MGI Symbol;Acc:MGI:894313] | Rab6a | Enriched in dLGN | 1.485 | 0.00005974 | 1003 |
| ENSMUSG00000020458 | reticulon 4 [Source:MGI Symbol;Acc:MGI:1915835] | Rtn4 | Enriched in dLGN | 1.484 | 0.00002851 | 1440 |
| ENSMUSG00000020358 | heterogeneous nuclear ribonucleoprotein A/B [Source:MGI Symbol;Acc:MGI:1330294] | Hnrnpab | Enriched in dLGN | 1.483 | 1.404E-08 | 542 |
| ENSMUSG00000075702 | selenoprotein M [Source:MGI Symbol;Acc:MGI:2149786] | Selenom | Enriched in dLGN | 1.479 | 0.000007293 | 136.2 |
| ENSMUSG00000048616 | noggin [Source:MGI Symbol;Acc:MGI:104327] | Nog | Enriched in dLGN | 1.438 | 0.0004898 | 53.91 |
| ENSMUSG00000027341 | transmembrane protein 230 [Source:MGI Symbol;Acc:MGI:1917862] | Tmem230 | Enriched in dLGN | 1.426 | 0.008374 | 38.18 |
| ENSMUSG00000031950 | gamma-aminobutyric acid (GABA) A receptor-associated protein-like 2 [Source:MGI Symbol;Acc:MGI:1890602] | Gabarapl2 | Enriched in dLGN | 1.418 | 0.00001248 | 147.2 |
| ENSMUSG00000047617 | non-homologous end joining factor [Source:MGI Symbol;Acc:MGI:2442831] | Paxx | Enriched in dLGN | 1.408 | 0.0003417 | 66.93 |
| ENSMUSG00000032366 | tropomyosin 1, alpha [Source:MGI Symbol;Acc:MGI:98809] | Tpm1 | Enriched in dLGN | 1.404 | 1.835E-16 | 654.7 |
| ENSMUSG00000030706 | mitochondrial ribosomal protein L48 [Source:MGI Symbol;Acc:MGI:1289321] | Mrpl48 | Enriched in dLGN | 1.403 | 0.0005795 | 153.1 |
| ENSMUSG00000046058 | EP300 interacting inhibitor of differentiation 2 [Source:MGI Symbol;Acc:MGI:2681174] | Eid2 | Enriched in dLGN | 1.381 | 0.0007357 | 137.6 |
| ENSMUSG00000028447 | dynactin 3 [Source:MGI Symbol;Acc:MGI:1859251] | Dctn3 | Enriched in dLGN | 1.376 | 0.000002153 | 123.3 |
| ENSMUSG00000024038 | NADH:ubiquinone oxidoreductase core subunit V3 [Source:MGI Symbol;Acc:MGI:1890894] | Ndufv3 | Enriched in dLGN | 1.374 | 0.000000575 | 374.2 |
| ENSMUSG00000001089 | leucine zipper protein 1 [Source:MGI Symbol;Acc:MGI:107629] | Luzp1 | Enriched in dLGN | 1.373 | 0.00003682 | 485.1 |
| ENSMUSG00000028563 | TM2 domain containing 1 [Source:MGI Symbol;Acc:MGI:2137022] | Tm2d1 | Enriched in dLGN | 1.371 | 0.002135 | 48.93 |
| ENSMUSG00000045763 | brain abundant, membrane attached signal protein 1 [Source:MGI Symbol;Acc:MGI:1917600] | Basp1 | Enriched in dLGN | 1.342 | 7.462E-07 | 3099 |
| ENSMUSG00000020736 | 5',3'-nucleotidase, cytosolic [Source:MGI Symbol;Acc:MGI:1354954] | Nt5c | Enriched in dLGN | 1.342 | 0.00002255 | 100.9 |
| ENSMUSG00000030876 | methyltransferase like 9 [Source:MGI Symbol;Acc:MGI:1914862] | Mettl9 | Enriched in dLGN | 1.323 | 0.001891 | 224.2 |
| ENSMUSG00000059518 | zinc finger, HIT domain containing 1 [Source:MGI Symbol;Acc:MGI:1917353] | Znhit1 | Enriched in dLGN | 1.322 | 0.000004187 | 135.3 |
| ENSMUSG00000028648 | NADH:ubiquinone oxidoreductase core subunit S5 [Source:MGI Symbol;Acc:MGI:1890889] | Ndufs5 | Enriched in dLGN | 1.316 | 0.006342 | 101.9 |
| ENSMUSG00000027602 | microtubule-associated protein 1 light chain 3 alpha [Source:MGI Symbol;Acc:MGI:1915661] | Map1lc3a | Enriched in dLGN | 1.314 | 0.000007024 | 233.2 |
| ENSMUSG00000020481 | ankyrin repeat domain 36 [Source:MGI Symbol;Acc:MGI:1923639] | Ankrd36 | Enriched in dLGN | 1.314 | 0.03536 | 15.79 |
| ENSMUSG00000021792 | peroxiredoxin like 2A [Source:MGI Symbol;Acc:MGI:1917814] | Prxl2a | Enriched in dLGN | 1.309 | 6.685E-09 | 278.9 |
| ENSMUSG00000051518 | ribosomal protein S19 binding protein 1 [Source:MGI Symbol;Acc:MGI:1913788] | Rps19bp1 | Enriched in dLGN | 1.309 | 0.0005481 | 50.07 |
| ENSMUSG00000026238 | prothymosin alpha [Source:MGI Symbol;Acc:MGI:97803] | Ptma | Enriched in dLGN | 1.306 | 4.359E-08 | 652.5 |
| ENSMUSG00000019861 | golgi associated PDZ and coiled-coil motif containing [Source:MGI Symbol;Acc:MGI:2149946] | Gopc | Enriched in dLGN | 1.303 | 1.845E-12 | 304.7 |
| ENSMUSG00000028478 | clathrin, light polypeptide (Lca) [Source:MGI Symbol;Acc:MGI:894297] | Clta | Enriched in dLGN | 1.301 | 1.026E-11 | 866.9 |
| ENSMUSG00000037148 | Rho GTPase activating protein 10 [Source:MGI Symbol;Acc:MGI:1925764] | Arhgap10 | Enriched in dLGN | 1.239 | 0.01309 | 198.1 |
| ENSMUSG00000051817 | SRY (sex determining region Y)-box 12 [Source:MGI Symbol;Acc:MGI:98360] | Sox12 | Enriched in dLGN | 1.237 | 0.01883 | 619.9 |
| ENSMUSG00000078235 | family with sequence similarity 43, member B [Source:MGI Symbol;Acc:MGI:3651622] | Fam43b | Enriched in dLGN | 1.233 | 0.00331 | 60.59 |
| ENSMUSG00000036377 | capping protein inhibiting regulator of actin [Source:MGI Symbol;Acc:MGI:2444817] | Cracd | Enriched in dLGN | 1.231 | 0.00002161 | 1779 |
| ENSMUSG00000028416 | BCL2-associated athanogene 1 [Source:MGI Symbol;Acc:MGI:108047] | Bag1 | Enriched in dLGN | 1.227 | 0.00001617 | 349.9 |
| ENSMUSG00000078249 | high mobility group AT-hook 1B [Source:MGI Symbol;Acc:MGI:96161] | Hmga1b | Enriched in dLGN | 1.225 | 0.001655 | 54.25 |
| ENSMUSG00000019471 | cell division cycle 37 [Source:MGI Symbol;Acc:MGI:109531] | Cdc37 | Enriched in dLGN | 1.222 | 0.00000217 | 613 |
| ENSMUSG00000025451 | polyadenylate binding protein-interacting protein 1 [Source:MGI Symbol;Acc:MGI:2384993] | Paip1 | Enriched in dLGN | 1.218 | 1.034E-07 | 231.7 |
| ENSMUSG00000004642 | stem-loop binding protein [Source:MGI Symbol;Acc:MGI:108402] | Slbp | Enriched in dLGN | 1.213 | 0.0001579 | 85.78 |
| ENSMUSG00000020108 | DNA-damage-inducible transcript 4 [Source:MGI Symbol;Acc:MGI:1921997] | Ddit4 | Enriched in dLGN | 1.213 | 0.0005159 | 181.8 |
| ENSMUSG00000025103 | BTB (POZ) domain containing 1 [Source:MGI Symbol;Acc:MGI:1933765] | Btbd1 | Enriched in dLGN | 1.212 | 0.00004051 | 230.6 |
| ENSMUSG00000034557 | zinc finger, FYVE domain containing 9 [Source:MGI Symbol;Acc:MGI:2652838] | Zfyve9 | Enriched in dLGN | 1.21 | 0.004441 | 559.4 |
| ENSMUSG00000031357 | synapse associated protein 1 [Source:MGI Symbol;Acc:MGI:1914293] | Syap1 | Enriched in dLGN | 1.207 | 0.001405 | 90.22 |
| ENSMUSG00000054428 | ATPase inhibitory factor 1 [Source:MGI Symbol;Acc:MGI:1196457] | Atpif1 | Enriched in dLGN | 1.205 | 0.0002719 | 829.4 |
| ENSMUSG00000019877 | serine incorporator 1 [Source:MGI Symbol;Acc:MGI:1926228] | Serinc1 | Enriched in dLGN | 1.203 | 0.00348 | 662.3 |
| ENSMUSG00000027502 | replication termination factor 2 [Source:MGI Symbol;Acc:MGI:1913654] | Rtf2 | Enriched in dLGN | 1.188 | 0.00002178 | 124.2 |
| ENSMUSG00000041688 | angiomotin [Source:MGI Symbol;Acc:MGI:108440] | Amot | Enriched in dLGN | 1.182 | 0.0429 | 327.4 |
| ENSMUSG00000027248 | protein disulfide isomerase associated 3 [Source:MGI Symbol;Acc:MGI:95834] | Pdia3 | Enriched in dLGN | 1.176 | 0.0009234 | 401 |
| ENSMUSG00000036087 | SLAIN motif family, member 2 [Source:MGI Symbol;Acc:MGI:1923241] | Slain2 | Enriched in dLGN | 1.165 | 0.00003705 | 209.6 |
| ENSMUSG00000046791 | ribosomal oxygenase 1 [Source:MGI Symbol;Acc:MGI:1919202] | Riox1 | Enriched in dLGN | 1.163 | 0.001274 | 41.2 |
| ENSMUSG00000023106 | density-regulated protein [Source:MGI Symbol;Acc:MGI:1915434] | Denr | Enriched in dLGN | 1.16 | 0.00003026 | 114.8 |
| ENSMUSG00000029634 | ring finger protein (C3H2C3 type) 6 [Source:MGI Symbol;Acc:MGI:1921382] | Rnf6 | Enriched in dLGN | 1.155 | 0.00001023 | 280.9 |
| ENSMUSG00000029467 | ATPase, Ca++ transporting, cardiac muscle, slow twitch 2 [Source:MGI Symbol;Acc:MGI:88110] | Atp2a2 | Enriched in dLGN | 1.15 | 0.000002131 | 2995 |
| ENSMUSG00000026188 | transmembrane protein 169 [Source:MGI Symbol;Acc:MGI:2442781] | Tmem169 | Enriched in dLGN | 1.145 | 0.00004284 | 139.3 |
| ENSMUSG00000002058 | unc-119 lipid binding chaperone [Source:MGI Symbol;Acc:MGI:1328357] | Unc119 | Enriched in dLGN | 1.145 | 0.0004015 | 481.3 |
| ENSMUSG00000025208 | mitochondrial ribosomal protein L43 [Source:MGI Symbol;Acc:MGI:2137229] | Mrpl43 | Enriched in dLGN | 1.129 | 0.001683 | 70.84 |
| ENSMUSG00000030515 | threonyl-tRNA synthetase-like 2 [Source:MGI Symbol;Acc:MGI:2444486] | Tarsl2 | Enriched in dLGN | 1.122 | 0.00005495 | 105.5 |
| ENSMUSG00000018965 | tyrosine 3-monooxygenase/tryptophan 5-monooxygenase activation protein, eta polypeptide [Source:MGI Symbol;Acc:MGI:109194] | Ywhah | Enriched in dLGN | 1.119 | 0.004931 | 3239 |
| ENSMUSG00000028945 | Ras homolog enriched in brain [Source:MGI Symbol;Acc:MGI:97912] | Rheb | Enriched in dLGN | 1.112 | 0.000001563 | 175.6 |
| ENSMUSG00000032393 | dipeptidylpeptidase 8 [Source:MGI Symbol;Acc:MGI:1921638] | Dpp8 | Enriched in dLGN | 1.11 | 0.00001561 | 799.2 |
| ENSMUSG00000091337 | EP300 interacting inhibitor of differentiation 1 [Source:MGI Symbol;Acc:MGI:1889651] | Eid1 | Enriched in dLGN | 1.106 | 0.00009393 | 1279 |
| ENSMUSG00000020349 | protein phosphatase 2 (formerly 2A), catalytic subunit, alpha isoform [Source:MGI Symbol;Acc:MGI:1321159] | Ppp2ca | Enriched in dLGN | 1.105 | 0.0001025 | 828.3 |
| ENSMUSG00000028466 | cAMP responsive element binding protein 3 [Source:MGI Symbol;Acc:MGI:99946] | Creb3 | Enriched in dLGN | 1.104 | 0.0004654 | 89.41 |
| ENSMUSG00000038525 | armadillo repeat containing 10 [Source:MGI Symbol;Acc:MGI:1914461] | Armc10 | Enriched in dLGN | 1.101 | 0.008693 | 41.51 |
| ENSMUSG00000021114 | ATPase, H+ transporting, lysosomal V1 subunit D [Source:MGI Symbol;Acc:MGI:1921084] | Atp6v1d | Enriched in dLGN | 1.094 | 0.001277 | 284.8 |
| ENSMUSG00000027203 | deoxyuridine triphosphatase [Source:MGI Symbol;Acc:MGI:1346051] | Dut | Enriched in dLGN | 1.087 | 0.00058 | 92.41 |
| ENSMUSG00000003604 | apoptosis, caspase activation inhibitor [Source:MGI Symbol;Acc:MGI:1921518] | Aven | Enriched in dLGN | 1.079 | 0.01091 | 49.12 |
| ENSMUSG00000052253 | zinc finger protein 622 [Source:MGI Symbol;Acc:MGI:1289282] | Zfp622 | Enriched in dLGN | 1.074 | 0.0001689 | 157.1 |
| ENSMUSG00000025762 | La ribonucleoprotein 1B [Source:MGI Symbol;Acc:MGI:1914604] | Larp1b | Enriched in dLGN | 1.07 | 0.002471 | 100 |
| ENSMUSG00000021102 | glutaredoxin 5 [Source:MGI Symbol;Acc:MGI:1920296] | Glrx5 | Enriched in dLGN | 1.067 | 0.00279 | 170.8 |
| ENSMUSG00000001016 | interleukin enhancer binding factor 2 [Source:MGI Symbol;Acc:MGI:1915031] | Ilf2 | Enriched in dLGN | 1.063 | 0.0003062 | 131.5 |
| ENSMUSG00000009630 | protein phosphatase 2 (formerly 2A), catalytic subunit, beta isoform [Source:MGI Symbol;Acc:MGI:1321161] | Ppp2cb | Enriched in dLGN | 1.06 | 0.000002093 | 209.1 |
| ENSMUSG00000026687 | aldehyde dehydrogenase 9, subfamily A1 [Source:MGI Symbol;Acc:MGI:1861622] | Aldh9a1 | Enriched in dLGN | 1.058 | 0.001505 | 102.9 |
| ENSMUSG00000040462 | amplified in osteosarcoma [Source:MGI Symbol;Acc:MGI:1924301] | Os9 | Enriched in dLGN | 1.056 | 0.00005771 | 698.2 |
| ENSMUSG00000008475 | actin related protein 2/3 complex, subunit 5 [Source:MGI Symbol;Acc:MGI:1915021] | Arpc5 | Enriched in dLGN | 1.047 | 0.00008198 | 264.7 |
| ENSMUSG00000027091 | zinc finger CCCH-type containing 15 [Source:MGI Symbol;Acc:MGI:1919747] | Zc3h15 | Enriched in dLGN | 1.042 | 0.0001152 | 331.8 |
| ENSMUSG00000058569 | transmembrane p24 trafficking protein 9 [Source:MGI Symbol;Acc:MGI:1914761] | Tmed9 | Enriched in dLGN | 1.041 | 0.0004241 | 135.9 |
| ENSMUSG00000026499 | acyl-Coenzyme A binding domain containing 3 [Source:MGI Symbol;Acc:MGI:2181074] | Acbd3 | Enriched in dLGN | 1.036 | 0.0005412 | 390.9 |
| ENSMUSG00000037563 | ribosomal protein S16 [Source:MGI Symbol;Acc:MGI:98118] | Rps16 | Enriched in dLGN | 1.021 | 0.01425 | 85.5 |
| ENSMUSG00000047547 | clathrin, light polypeptide (Lcb) [Source:MGI Symbol;Acc:MGI:1921575] | Cltb | Enriched in dLGN | 1.02 | 3.307E-09 | 896 |
| ENSMUSG00000015092 | endothelial differentiation-related factor 1 [Source:MGI Symbol;Acc:MGI:1891227] | Edf1 | Enriched in dLGN | 1.016 | 0.0003985 | 190.3 |

| | | | | | | |
|---|---|---|---|---|---|---|
| ENSMUSG00000036615 | regulatory factor X-associated protein [Source:MGI Symbol;Acc:MGI:2180854] | Rfxap | Enriched in dLGN | 1.012 | 0.00007202 | 118.3 |
| ENSMUSG00000035228 | coiled-coil domain containing 106 [Source:MGI Symbol;Acc:MGI:2385900] | Ccdc106 | Enriched in dLGN | 1.004 | 0.001419 | 90.05 |
| ENSMUSG00000020544 | cytochrome c oxidase assembly protein 11, copper chaperone [Source:MGI Symbol;Acc:MGI:1917052] | Cox11 | Enriched in dLGN | 1.001 | 0.01911 | 38.98 |
| ENSMUSG00000048154 | lysine (K)-specific methyltransferase 2D [Source:MGI Symbol;Acc:MGI:2682319] | Kmt2d | Enriched in SCN | -1.005 | 0.01665 | 4240 |
| ENSMUSG00000020455 | tripartite motif-containing 11 [Source:MGI Symbol;Acc:MGI:2137355] | Trim11 | Enriched in SCN | -1.067 | 0.02299 | 100.1 |
| ENSMUSG00000025607 | coatomer protein complex, subunit gamma 2 [Source:MGI Symbol;Acc:MGI:1858683] | Copg2 | Enriched in SCN | -1.101 | 0.02756 | 319.5 |
| ENSMUSG00000024002 | bromodomain containing 4 [Source:MGI Symbol;Acc:MGI:1888520] | Brd4 | Enriched in SCN | -1.13 | 0.002092 | 1667 |
| ENSMUSG00000046546 | family with sequence similarity 43, member A [Source:MGI Symbol;Acc:MGI:2676309] | Fam43a | Enriched in SCN | -1.283 | 0.004464 | 83.9 |
| ENSMUSG00000046637 | tetratricopeptide repeat domain 34 [Source:MGI Symbol;Acc:MGI:2445205] | Ttc34 | Enriched in SCN | -1.373 | 0.02534 | 13.93 |
| ENSMUSG00000027014 | CWC22 spliceosome-associated protein [Source:MGI Symbol;Acc:MGI:2136773] | Cwc22 | Enriched in SCN | -1.402 | 0.009197 | 206.1 |
| ENSMUSG00000010342 | testis expressed gene 14 [Source:MGI Symbol;Acc:MGI:1933227] | Tex14 | Enriched in SCN | -1.508 | 0.03383 | 109.9 |
| ENSMUSG00000000560 | gamma-aminobutyric acid (GABA) A receptor, subunit alpha 2 [Source:MGI Symbol;Acc:MGI:95614] | Gabra2 | Enriched in SCN | -1.529 | 0.0007366 | 394.2 |
| ENSMUSG00000042606 | HIRA interacting protein 3 [Source:MGI Symbol;Acc:MGI:2142364] | Hirip3 | Enriched in SCN | -1.573 | 0.000209 | 132.7 |
| ENSMUSG00000032425 | zinc finger protein 949 [Source:MGI Symbol;Acc:MGI:1918890] | Zfp949 | Enriched in SCN | -1.623 | 0.00007451 | 260.3 |
| ENSMUSG00000029245 | Eph receptor A5 [Source:MGI Symbol;Acc:MGI:99654] | Epha5 | Enriched in SCN | -1.9 | 0.000003691 | 636.4 |
| ENSMUSG00000096141 | dynein, axonemal, heavy chain 7A [Source:MGI Symbol;Acc:MGI:2685838] | Dnah7a | Enriched in SCN | -2.042 | 0.009713 | 25.06 |
| ENSMUSG00000052428 | transmembrane and coiled-coil domains 1 [Source:MGI Symbol;Acc:MGI:1921173] | Tmco1 | Enriched in SCN | -2.288 | 0.0005163 | 95.8 |
| ENSMUSG00000072663 | sperm flagellar 2 [Source:MGI Symbol;Acc:MGI:2443727] | Spef2 | Enriched in SCN | -2.305 | 0.001315 | 23.31 |
| ENSMUSG00000019767 | coiled-coil domain containing 170 [Source:MGI Symbol;Acc:MGI:2685067] | Ccdc170 | Enriched in SCN | -2.609 | 0.008825 | 16.02 |
| ENSMUSG00000029442 | WD repeat domain 66 [Source:MGI Symbol;Acc:MGI:1918495] | Wdr66 | Enriched in SCN | -2.735 | 7.323E-08 | 54.47 |
| ENSMUSG00000029601 | IQ motif containing D [Source:MGI Symbol;Acc:MGI:1922982] | Iqcd | Enriched in SCN | -3.025 | 0.003281 | 36.37 |
| ENSMUSG00000050677 | coiled-coil domain containing 96 [Source:MGI Symbol;Acc:MGI:1913967] | Ccdc96 | Enriched in SCN | -3.241 | 0.000007024 | 66.05 |
| ENSMUSG00000030276 | tubulin tyrosine ligase-like family, member 3 [Source:MGI Symbol;Acc:MGI:2141418] | Ttll3 | Enriched in SCN | -3.339 | 2.763E-07 | 73.6 |

**Contrast numerator: P8_Het_SCN. Contrast denominator: P8_Het_retina.**

| ensembl_gene_id | description | mgi_symbol | Significance | deseq_logfc | deseq_adjp | deseq_basemean |
|---|---|---|---|---|---|---|
| ENSMUSG00000004892 | brevican [Source:MGI Symbol;Acc:MGI:1096385] | Bcan | Enriched in SCN | 4.853 | 6.417E-70 | 546.7 |
| ENSMUSG00000059854 | HYDIN, axonemal central pair apparatus protein [Source:MGI Symbol;Acc:MGI:2389007] | Hydin | Enriched in SCN | 4.608 | 1.007E-16 | 115.1 |
| ENSMUSG00000062372 | otoferlin [Source:MGI Symbol;Acc:MGI:1891247] | Otof | Enriched in SCN | 4.044 | 3.431E-18 | 68.05 |
| ENSMUSG00000027360 | histidine decarboxylase [Source:MGI Symbol;Acc:MGI:96062] | Hdc | Enriched in SCN | 4.005 | 1.741E-12 | 44.93 |
| ENSMUSG00000021139 | predicted gene 20498 [Source:MGI Symbol;Acc:MGI:5141963] | Gm20498 | Enriched in SCN | 3.961 | 1.116E-19 | 272.3 |
| ENSMUSG00000095139 | POU domain, class 3, transcription factor 2 [Source:MGI Symbol;Acc:MGI:101895] | Pou3f2 | Enriched in SCN | 3.914 | 6.506E-33 | 124.7 |
| ENSMUSG00000053004 | histamine receptor H1 [Source:MGI Symbol;Acc:MGI:107619] | Hrh1 | Enriched in SCN | 3.91 | 1.578E-14 | 32.83 |
| ENSMUSG00000048721 | fibronectin type III domain containing 9 [Source:MGI Symbol;Acc:MGI:2443410] | Fndc9 | Enriched in SCN | 3.408 | 0.00006945 | 15.76 |
| ENSMUSG00000078816 | protein kinase C, gamma [Source:MGI Symbol;Acc:MGI:97597] | Prkcg | Enriched in SCN | 3.399 | 1.006E-18 | 571.8 |
| ENSMUSG00000040181 | flavin containing monooxygenase 1 [Source:MGI Symbol;Acc:MGI:1310002] | Fmo1 | Enriched in SCN | 3.287 | 7.714E-13 | 18.97 |
| ENSMUSG00000073294 | EZH inhibitory protein [Source:MGI Symbol;Acc:MGI:2147968] | Ezhip | Enriched in SCN | 3.218 | 0.00209 | 8.404 |
| ENSMUSG00000031111 | immunoglobulin superfamily, member 1 [Source:MGI Symbol;Acc:MGI:2147913] | Igsf1 | Enriched in SCN | 3.133 | 1.435E-15 | 83.92 |
| ENSMUSG00000000794 | potassium intermediate/small conductance calcium-activated channel, subfamily N, member 3 [Source:MGI Symbol;Acc:MGI:2153183] | Kcnn3 | Enriched in SCN | 3.035 | 1.817E-26 | 249.5 |
| ENSMUSG00000038112 | expressed sequence AW551984 [Source:MGI Symbol;Acc:MGI:2143322] | AW551984 | Enriched in SCN | 2.945 | 0.00006894 | 188.6 |
| ENSMUSG00000024617 | calcium/calmodulin-dependent protein kinase II alpha [Source:MGI Symbol;Acc:MGI:88256] | Camk2a | Enriched in SCN | 2.939 | 8.853E-24 | 1425 |
| ENSMUSG00000022696 | SID1 transmembrane family, member 1 [Source:MGI Symbol;Acc:MGI:2443155] | Sidt1 | Enriched in SCN | 2.917 | 3.48E-20 | 43.24 |
| ENSMUSG00000010086 | ring finger protein 112 [Source:MGI Symbol;Acc:MGI:106611] | Rnf112 | Enriched in SCN | 2.869 | 1.597E-20 | 340.8 |
| ENSMUSG00000066705 | FXYD domain-containing ion transport regulator 6 [Source:MGI Symbol;Acc:MGI:1890226] | Fxyd6 | Enriched in SCN | 2.836 | 6.737E-12 | 380.3 |
| ENSMUSG00000044667 | phospholipid phosphatase related 4 [Source:MGI Symbol;Acc:MGI:106530] | Plppr4 | Enriched in SCN | 2.809 | 5.549E-25 | 669.3 |
| ENSMUSG00000042425 | FERM and PDZ domain containing 3 [Source:MGI Symbol;Acc:MGI:3646547] | Frmpd3 | Enriched in SCN | 2.76 | 5.524E-10 | 191.4 |
| ENSMUSG00000108358 | predicted gene 44509 [Source:MGI Symbol;Acc:MGI:5753085] | Gm44509 | Enriched in SCN | 2.758 | 6.893E-12 | 119 |
| ENSMUSG00000036856 | wingless-type MMTV integration site family, member 4 [Source:MGI Symbol;Acc:MGI:98957] | Wnt4 | Enriched in SCN | 2.745 | 1.125E-18 | 80.14 |
| ENSMUSG00000021750 | family with sequence similarity 107, member A [Source:MGI Symbol;Acc:MGI:3041256] | Fam107a | Enriched in SCN | 2.717 | 0.000103 | 164.6 |
| ENSMUSG00000045201 | leucine rich repeat containing 3B [Source:MGI Symbol;Acc:MGI:2384996] | Lrrc3b | Enriched in SCN | 2.681 | 1.122E-08 | 34.56 |
| ENSMUSG00000022208 | junctophilin 4 [Source:MGI Symbol;Acc:MGI:2443113] | Jph4 | Enriched in SCN | 2.662 | 1.863E-27 | 961.5 |
| ENSMUSG00000074923 | p21 (RAC1) activated kinase 6 [Source:MGI Symbol;Acc:MGI:2679420] | Pak6 | Enriched in SCN | 2.629 | 1.327E-26 | 281.4 |
| ENSMUSG00000038453 | SRC kinase signaling inhibitor 1 [Source:MGI Symbol;Acc:MGI:1933179] | Srcin1 | Enriched in SCN | 2.628 | 2.644E-49 | 3243 |
| ENSMUSG00000063646 | janus kinase and microtubule interacting protein 1 [Source:MGI Symbol;Acc:MGI:1923321] | Jakmip1 | Enriched in SCN | 2.602 | 6.773E-09 | 47 |
| ENSMUSG00000018486 | wingless-type MMTV integration site family, member 9B [Source:MGI Symbol;Acc:MGI:1197020] | Wnt9b | Enriched in SCN | 2.56 | 0.00002208 | 165.2 |
| ENSMUSG00000042105 | inositol polyphosphate-5-phosphatase F [Source:MGI Symbol;Acc:MGI:2141867] | Inpp5f | Enriched in SCN | 2.551 | 1.058E-21 | 994.9 |
| ENSMUSG00000039278 | proprotein convertase subtilisin/kexin type 1 inhibitor [Source:MGI Symbol;Acc:MGI:1353431] | Pcsk1n | Enriched in SCN | 2.512 | 3.916E-16 | 2810 |
| ENSMUSG00000054027 | 5'-nucleotidase domain containing 3 [Source:MGI Symbol;Acc:MGI:3513266] | Nt5dc3 | Enriched in SCN | 2.5 | 7.501E-20 | 395 |
| ENSMUSG00000005045 | chromodomain helicase DNA binding protein 5 [Source:MGI Symbol;Acc:MGI:3036258] | Chd5 | Enriched in SCN | 2.496 | 1.055E-29 | 1862 |
| ENSMUSG00000022112 | glypican 5 [Source:MGI Symbol;Acc:MGI:1194894] | Gpc5 | Enriched in SCN | 2.495 | 1.266E-07 | 390.3 |
| ENSMUSG00000054013 | transmembrane protein 179 [Source:MGI Symbol;Acc:MGI:2144891] | Tmem179 | Enriched in SCN | 2.49 | 1.366E-19 | 349.1 |
| ENSMUSG00000068373 | RIKEN cDNA D430041D05 gene [Source:MGI Symbol;Acc:MGI:2181743] | D430041D05Rik | Enriched in SCN | 2.489 | 7.771E-42 | 780.2 |
| ENSMUSG00000034282 | envoplakin [Source:MGI Symbol;Acc:MGI:107507] | Evpl | Enriched in SCN | 2.483 | 2.499E-17 | 114.4 |
| ENSMUSG00000073406 | histocompatibility 2, blastocyst [Source:MGI Symbol;Acc:MGI:892004] | H2-Bl | Enriched in SCN | 2.482 | 0.003037 | 18.45 |
| ENSMUSG00000061702 | transmembrane protein 91 [Source:MGI Symbol;Acc:MGI:2443589] | Tmem91 | Enriched in SCN | 2.474 | 1.832E-10 | 41.88 |
| ENSMUSG00000110086 | small integral membrane protein 32 [Source:MGI Symbol;Acc:MGI:5791459] | Smim32 | Enriched in SCN | 2.42 | 3.043E-08 | 24.37 |
| ENSMUSG00000070644 | ethanolamine kinase 2 [Source:MGI Symbol;Acc:MGI:2443760] | Etnk2 | Enriched in SCN | 2.412 | 7.519E-15 | 35.13 |
| ENSMUSG00000028758 | kinesin family member 17 [Source:MGI Symbol;Acc:MGI:1098229] | Kif17 | Enriched in SCN | 2.406 | 9.073E-22 | 102.3 |
| ENSMUSG00000055430 | nucleosome assembly protein 1-like 5 [Source:MGI Symbol;Acc:MGI:1923555] | Nap1l5 | Enriched in SCN | 2.395 | 1.059E-13 | 1237 |
| ENSMUSG00000032532 | cholecystokinin [Source:MGI Symbol;Acc:MGI:88297] | Cck | Enriched in SCN | 2.371 | 0.00002074 | 445.5 |
| ENSMUSG00000035355 | potassium voltage-gated channel, subfamily H (eag-related), member 4 [Source:MGI Symbol;Acc:MGI:2156184] | Kcnh4 | Enriched in SCN | 2.369 | 0.006538 | 7.674 |
| ENSMUSG00000029516 | citron [Source:MGI Symbol;Acc:MGI:105313] | Cit | Enriched in SCN | 2.361 | 6.647E-25 | 2284 |
| ENSMUSG00000035431 | somatostatin receptor 1 [Source:MGI Symbol;Acc:MGI:98327] | Sstr1 | Enriched in SCN | 2.353 | 0.000003501 | 18.08 |
| ENSMUSG00000067028 | contactin associated protein-like 5B [Source:MGI Symbol;Acc:MGI:3664583] | Cntnap5b | Enriched in SCN | 2.343 | 9.872E-09 | 198.7 |
| ENSMUSG00000070304 | sodium channel, voltage-gated, type II, beta [Source:MGI Symbol;Acc:MGI:106921] | Scn2b | Enriched in SCN | 2.341 | 5.642E-08 | 2099 |
| ENSMUSG00000052026 | solute carrier family 6 (neurotransmitter transporter, L-proline), member 7 [Source:MGI Symbol;Acc:MGI:2147363] | Slc6a7 | Enriched in SCN | 2.34 | 6.772E-25 | 217.9 |
| ENSMUSG00000020090 | neuropeptide FF receptor 1 [Source:MGI Symbol;Acc:MGI:2685082] | Npffr1 | Enriched in SCN | 2.336 | 0.000006938 | 15.66 |
| ENSMUSG00000019590 | cytochrome b-561 [Source:MGI Symbol;Acc:MGI:103253] | Cyb561 | Enriched in SCN | 2.321 | 2.989E-14 | 123.2 |
| ENSMUSG00000031226 | polysaccharide biosynthesis domain containing 1 [Source:MGI Symbol;Acc:MGI:1914933] | Pbdc1 | Enriched in SCN | 2.278 | 4.93E-10 | 236 |
| ENSMUSG00000037428 | VGF nerve growth factor inducible [Source:MGI Symbol;Acc:MGI:1343180] | Vgf | Enriched in SCN | 2.278 | 2.463E-15 | 1249 |
| ENSMUSG00000035576 | L3MBTL1 histone methyl-lysine binding protein [Source:MGI Symbol;Acc:MGI:2676663] | L3mbtl1 | Enriched in SCN | 2.245 | 4.385E-12 | 85.42 |
| ENSMUSG00000008496 | POU domain, class 2, transcription factor 2 [Source:MGI Symbol;Acc:MGI:101897] | Pou2f2 | Enriched in SCN | 2.228 | 2.053E-12 | 295.3 |
| undefined | undefined | undefined | Enriched in SCN | 2.216 | 2.094E-13 | 1196 |
| ENSMUSG00000029602 | RAS protein activator like 1 (GAP1 like) [Source:MGI Symbol;Acc:MGI:1330842] | Rasal1 | Enriched in SCN | 2.206 | 2.413E-10 | 51.4 |
| ENSMUSG00000020374 | RasGEF domain family, member 1C [Source:MGI Symbol;Acc:MGI:1921813] | Rasgef1c | Enriched in SCN | 2.202 | 5.226E-14 | 39.33 |
| ENSMUSG00000032890 | regulating synaptic membrane exocytosis 3 [Source:MGI Symbol;Acc:MGI:2443331] | Rims3 | Enriched in SCN | 2.176 | 3.588E-17 | 1313 |
| ENSMUSG00000021534 | RIKEN cDNA 1700001L19 gene [Source:MGI Symbol;Acc:MGI:1916565] | 1700001L19Rik | Enriched in SCN | 2.168 | 0.000001116 | 10.84 |
| ENSMUSG00000075703 | selenoprotein I [Source:MGI Symbol;Acc:MGI:107898] | Selenoi | Enriched in SCN | 2.164 | 3.969E-19 | 269.3 |
| ENSMUSG00000032017 | glutamate receptor, ionotropic, kainate 4 [Source:MGI Symbol;Acc:MGI:95817] | Grik4 | Enriched in SCN | 2.151 | 2.947E-10 | 185.2 |
| ENSMUSG00000075324 | fidgetin [Source:MGI Symbol;Acc:MGI:1890647] | Fign | Enriched in SCN | 2.15 | 5.043E-08 | 251.6 |
| ENSMUSG00000022048 | dihydropyrimidinase-like 2 [Source:MGI Symbol;Acc:MGI:1349763] | Dpysl2 | Enriched in SCN | 2.13 | 4.08E-09 | 6238 |
| ENSMUSG00000026020 | NOP58 ribonucleoprotein [Source:MGI Symbol;Acc:MGI:1933184] | Nop58 | Enriched in SCN | 2.122 | 6.31E-17 | 363.9 |
| ENSMUSG00000037703 | leucine zipper, putative tumor suppressor family member 3 [Source:MGI Symbol;Acc:MGI:2656976] | Lzts3 | Enriched in SCN | 2.122 | 2.17E-19 | 572.4 |
| ENSMUSG00000050069 | gremlin 2, DAN family BMP antagonist [Source:MGI Symbol;Acc:MGI:1344367] | Grem2 | Enriched in SCN | 2.117 | 7.473E-13 | 70.97 |
| ENSMUSG00000021448 | src homology 2 domain-containing transforming protein C3 [Source:MGI Symbol;Acc:MGI:106179] | Shc3 | Enriched in SCN | 2.117 | 1.325E-23 | 933.3 |
| ENSMUSG00000031302 | neuroligin 3 [Source:MGI Symbol;Acc:MGI:2444609] | Nlgn3 | Enriched in SCN | 2.108 | 2.795E-09 | 1941 |
| ENSMUSG00000018849 | WW, C2 and coiled-coil domain containing 1 [Source:MGI Symbol;Acc:MGI:2388637] | Wwc1 | Enriched in SCN | 2.108 | 5.06E-29 | 625.4 |
| ENSMUSG00000020922 | LSM12 homolog [Source:MGI Symbol;Acc:MGI:1919592] | Lsm12 | Enriched in SCN | 2.099 | 4.055E-09 | 397.5 |
| ENSMUSG00000004263 | atrophin 1 [Source:MGI Symbol;Acc:MGI:104725] | Atn1 | Enriched in SCN | 2.094 | 0.00004853 | 25.12 |
| ENSMUSG00000004187 | kinesin family member C2 [Source:MGI Symbol;Acc:MGI:109187] | Kifc2 | Enriched in SCN | 2.085 | 3.12E-31 | 769 |
| ENSMUSG00000034226 | ras homolog family member V [Source:MGI Symbol;Acc:MGI:2444227] | Rhov | Enriched in SCN | 2.068 | 2.684E-10 | 54.65 |
| ENSMUSG00000102692 | dachsous cadherin related 2 [Source:MGI Symbol;Acc:MGI:2685263] | Dchs2 | Enriched in SCN | 2.066 | 0.00008642 | 117.9 |
| ENSMUSG00000042873 | lipoma HMGIC fusion partner-like protein 4 [Source:MGI Symbol;Acc:MGI:3057108] | Lhfpl4 | Enriched in SCN | 2.063 | 5.987E-17 | 1362 |
| ENSMUSG00000021596 | multiple C2 domains, transmembrane 1 [Source:MGI Symbol;Acc:MGI:1926021] | Mctp1 | Enriched in SCN | 2.035 | 2.677E-09 | 101.7 |
| ENSMUSG00000034037 | FYVE, RhoGEF and PH domain containing 5 [Source:MGI Symbol;Acc:MGI:2443369] | Fgd5 | Enriched in SCN | 2.034 | 4.17E-09 | 83.07 |
| ENSMUSG00000051853 | ADP-ribosylation factor 3 [Source:MGI Symbol;Acc:MGI:99432] | Arf3 | Enriched in SCN | 2.033 | 1.792E-12 | 2113 |
| ENSMUSG00000037362 | cellular communication network factor 3 [Source:MGI Symbol;Acc:MGI:109185] | Ccn3 | Enriched in SCN | 2.019 | 0.001547 | 15.41 |
| ENSMUSG00000042216 | small G protein signaling modulator 1 [Source:MGI Symbol;Acc:MGI:107320] | Sgsm1 | Enriched in SCN | 2.009 | 2.494E-18 | 412.2 |
| ENSMUSG00000040752 | myosin, heavy polypeptide 6, cardiac muscle, alpha [Source:MGI Symbol;Acc:MGI:97255] | Myh6 | Enriched in SCN | 2 | 0.001846 | 9.823 |
| ENSMUSG00000046491 | C1q and tumor necrosis factor related protein 2 [Source:MGI Symbol;Acc:MGI:1916433] | C1qtnf2 | Enriched in SCN | 1.998 | 0.01251 | 4.439 |
| ENSMUSG00000010554 | methyltransferase like 16 [Source:MGI Symbol;Acc:MGI:1914743] | Mettl16 | Enriched in SCN | 1.984 | 1.111E-12 | 322.2 |
| ENSMUSG00000017929 | UDP-Gal:betaGlcNAc beta 1,4-galactosyltransferase, polypeptide 5 [Source:MGI Symbol;Acc:MGI:1927169] | B4galt5 | Enriched in SCN | 1.927 | 1.002E-12 | 566.4 |
| ENSMUSG00000079388 | RIKEN cDNA 2610042L04 gene [Source:MGI Symbol;Acc:MGI:1914305] | 2610042L04Rik | Enriched in SCN | 1.922 | 0.01575 | 4.418 |
| ENSMUSG00000009575 | chromobox 5 [Source:MGI Symbol;Acc:MGI:109372] | Cbx5 | Enriched in SCN | 1.922 | 1.873E-10 | 2146 |
| ENSMUSG00000058793 | CDP-diacylglycerol synthase (phosphatidate cytidylyltransferase) 2 [Source:MGI Symbol;Acc:MGI:1332236] | Cds2 | Enriched in SCN | 1.904 | 5.023E-11 | 2925 |
| ENSMUSG00000021852 | solute carrier family 35, member F4 [Source:MGI Symbol;Acc:MGI:1922538] | Slc35f4 | Enriched in SCN | 1.899 | 0.00006144 | 56.44 |
| ENSMUSG00000027584 | opioid receptor-like 1 [Source:MGI Symbol;Acc:MGI:97440] | Oprl1 | Enriched in SCN | 1.882 | 4.814E-19 | 154 |
| ENSMUSG00000037994 | solute carrier family 9, subfamily B (NHA2, cation proton antiporter 2), member 2 [Source:MGI Symbol;Acc:MGI:2140077] | Slc9b2 | Enriched in SCN | 1.878 | 0.002071 | 11.02 |
| ENSMUSG00000009394 | synapsin II [Source:MGI Symbol;Acc:MGI:103020] | Syn2 | Enriched in SCN | 1.865 | 2.405E-11 | 611.4 |
| ENSMUSG00000058420 | synaptotagmin XVII [Source:MGI Symbol;Acc:MGI:104966] | Syt17 | Enriched in SCN | 1.851 | 2.562E-08 | 49.03 |
| ENSMUSG00000034664 | integrin alpha 2b [Source:MGI Symbol;Acc:MGI:96601] | Itga2b | Enriched in SCN | 1.832 | 0.00008288 | 10.87 |
| ENSMUSG00000105867 | predicted gene 42517 [Source:MGI Symbol;Acc:MGI:5662654] | Gm42517 | Enriched in SCN | 1.826 | 5.376E-10 | 89.61 |
| ENSMUSG00000069662 | myristoylated alanine rich protein kinase C substrate [Source:MGI Symbol;Acc:MGI:96907] | Marcks | Enriched in SCN | 1.817 | 0.000003077 | 2471 |
| ENSMUSG00000055471 | anaplastic lymphoma kinase [Source:MGI Symbol;Acc:MGI:103305] | Alk | Enriched in SCN | 1.815 | 0.00002063 | 211 |
| ENSMUSG00000053297 | expressed sequence AI854703 [Source:MGI Symbol;Acc:MGI:2141510] | AI854703 | Enriched in SCN | 1.812 | 1.809E-14 | 90.96 |
| ENSMUSG00000043165 | loricrin [Source:MGI Symbol;Acc:MGI:96816] | Lor | Enriched in SCN | 1.797 | 0.00516 | 17.17 |
| ENSMUSG00000006711 | RIKEN cDNA D130043K22 gene [Source:MGI Symbol;Acc:MGI:3036268] | D130043K22Rik | Enriched in SCN | 1.797 | 6.086E-08 | 133.1 |
| ENSMUSG00000039579 | glutamate receptor ionotropic, NMDA3A [Source:MGI Symbol;Acc:MGI:1933206] | Grin3a | Enriched in SCN | 1.792 | 9.642E-11 | 314.9 |
| ENSMUSG00000063296 | transmembrane protein 117 [Source:MGI Symbol;Acc:MGI:2444580] | Tmem117 | Enriched in SCN | 1.789 | 0.000007537 | 103 |
| ENSMUSG00000052557 | giant axonal neuropathy [Source:MGI Symbol;Acc:MGI:1890619] | Gan | Enriched in SCN | 1.789 | 1.636E-12 | 388.9 |
| ENSMUSG00000022577 | lymphocyte antigen 6 complex, locus H [Source:MGI Symbol;Acc:MGI:1346030] | Ly6h | Enriched in SCN | 1.768 | 6.275E-08 | 411.3 |
| ENSMUSG00000035640 | calcium channel, voltage-dependent, beta subunit associated regulatory protein [Source:MGI Symbol;Acc:MGI:1354170] | Cbarp | Enriched in SCN | 1.758 | 3.213E-13 | 1169 |
| ENSMUSG00000022421 | neuronal pentraxin receptor [Source:MGI Symbol;Acc:MGI:1920590] | Nptxr | Enriched in SCN | 1.752 | 4.544E-08 | 1078 |
| ENSMUSG00000021662 | Rho guanine nucleotide exchange factor (GEF) 28 [Source:MGI Symbol;Acc:MGI:1346016] | Arhgef28 | Enriched in SCN | 1.744 | 2.929E-15 | 213.6 |
| ENSMUSG00000049252 | low density lipoprotein-related protein 1B [Source:MGI Symbol;Acc:MGI:2151136] | Lrp1b | Enriched in SCN | 1.742 | 3.959E-07 | 1165 |
| ENSMUSG00000032076 | cell adhesion molecule 1 [Source:MGI Symbol;Acc:MGI:1889272] | Cadm1 | Enriched in SCN | 1.74 | 2.395E-11 | 2724 |
| ENSMUSG00000041115 | IQ motif and Sec7 domain 2 [Source:MGI Symbol;Acc:MGI:3528396] | Iqsec2 | Enriched in SCN | 1.739 | 7.973E-25 | 368.7 |
| ENSMUSG00000029580 | actin, beta [Source:MGI Symbol;Acc:MGI:87904] | Actb | Enriched in SCN | 1.733 | 0.0001087 | 6299 |
| ENSMUSG00000028341 | nuclear receptor subfamily 4, group A, member 3 [Source:MGI Symbol;Acc:MGI:1352457] | Nr4a3 | Enriched in SCN | 1.732 | 0.000005137 | 47.01 |
| ENSMUSG00000031963 | BMP-binding endothelial regulator [Source:MGI Symbol;Acc:MGI:1920480] | Bmper | Enriched in SCN | 1.716 | 0.000001836 | 44.47 |
| ENSMUSG00000036760 | potassium channel, subfamily K, member 9 [Source:MGI Symbol;Acc:MGI:3521816] | Kcnk9 | Enriched in SCN | 1.713 | 8.609E-08 | 406.3 |
| ENSMUSG00000066129 | kinase non-catalytic C-lobe domain (KIND) containing 1 [Source:MGI Symbol;Acc:MGI:1923734] | Kndc1 | Enriched in SCN | 1.713 | 2.605E-12 | 714.5 |
| ENSMUSG00000047415 | G protein-coupled receptor 68 [Source:MGI Symbol;Acc:MGI:2441763] | Gpr68 | Enriched in SCN | 1.707 | 1.357E-08 | 34.85 |
| ENSMUSG00000043388 | transmembrane protein 130 [Source:MGI Symbol;Acc:MGI:3607706] | Tmem130 | Enriched in SCN | 1.7 | 3.191E-07 | 1062 |
| ENSMUSG00000033022 | cysteine dioxygenase 1, cytosolic [Source:MGI Symbol;Acc:MGI:105925] | Cdo1 | Enriched in SCN | 1.696 | 6.686E-07 | 55.15 |
| ENSMUSG00000059203 | interleukin 1 receptor accessory protein-like 2 [Source:MGI Symbol;Acc:MGI:1913106] | Il1rapl2 | Enriched in SCN | 1.695 | 0.0002948 | 143.9 |
| ENSMUSG00000051391 | tyrosine 3-monooxygenase/tryptophan 5-monooxygenase activation protein, gamma polypeptide [Source:MGI Symbol;Acc:MGI:108109] | Ywhag | Enriched in SCN | 1.685 | 4.965E-09 | 5433 |
| ENSMUSG00000023017 | acid-sensing (proton-gated) ion channel 1 [Source:MGI Symbol;Acc:MGI:1194915] | Asic1 | Enriched in SCN | 1.672 | 1.528E-13 | 275.4 |
| ENSMUSG00000028381 | UDP-glucose ceramide glucosyltransferase [Source:MGI Symbol;Acc:MGI:1332243] | Ugcg | Enriched in SCN | 1.662 | 0.000003796 | 361.5 |
| ENSMUSG00000037348 | progestin and adipoQ receptor family member VII [Source:MGI Symbol;Acc:MGI:1919154] | Paqr7 | Enriched in SCN | 1.643 | 6.657E-10 | 163 |
| ENSMUSG00000001227 | sema domain, transmembrane domain (TM), and cytoplasmic domain, (semaphorin) 6B [Source:MGI Symbol;Acc:MGI:1202889] | Sema6b | Enriched in SCN | 1.641 | 5.324E-16 | 529.4 |
| ENSMUSG00000022840 | adenylate cyclase 5 [Source:MGI Symbol;Acc:MGI:99673] | Adcy5 | Enriched in SCN | 1.639 | 3.251E-10 | 2027 |

| | | | | | | |
|---|---|---|---|---|---|---|
| ENSMUSG00000042846 | leucine rich repeat transmembrane neuronal 3 [Source:MGI Symbol;Acc:MGI:2389177] | Lrrtm3 | Enriched in SCN | 1.635 | 0.00002781 | 135.3 |
| ENSMUSG00000032425 | zinc finger protein 949 [Source:MGI Symbol;Acc:MGI:1918890] | Zfp949 | Enriched in SCN | 1.623 | 0.000004001 | 260.3 |
| ENSMUSG00000030854 | protein tyrosine phosphatase, non-receptor type 5 [Source:MGI Symbol;Acc:MGI:97807] | Ptpn5 | Enriched in SCN | 1.613 | 6.137E-16 | 407 |
| ENSMUSG00000051495 | interferon regulatory factor 2 binding protein 2 [Source:MGI Symbol;Acc:MGI:2443921] | Irf2bp2 | Enriched in SCN | 1.607 | 5.202E-07 | 827.5 |
| ENSMUSG00000026494 | kinesin family member 26B [Source:MGI Symbol;Acc:MGI:2447076] | Kif26b | Enriched in SCN | 1.606 | 1.928E-08 | 391.1 |
| ENSMUSG00000072572 | solute carrier family 39 (zinc transporter), member 2 [Source:MGI Symbol;Acc:MGI:2684326] | Slc39a2 | Enriched in SCN | 1.604 | 0.04171 | 19.9 |
| ENSMUSG00000066607 | inhibitory synaptic factor 1 [Source:MGI Symbol;Acc:MGI:2442108] | Insyn1 | Enriched in SCN | 1.596 | 1.107E-11 | 582.5 |
| ENSMUSG00000067242 | leucine-rich repeat LGI family, member 1 [Source:MGI Symbol;Acc:MGI:1861691] | Lgi1 | Enriched in SCN | 1.589 | 3.965E-07 | 72.29 |
| ENSMUSG00000028064 | sema domain, immunoglobulin domain (Ig), transmembrane domain (TM) and short cytoplasmic domain, (semaphorin) 4A [Source:MGI Symbol;Acc:MGI:1075 | Sema4a | Enriched in SCN | 1.583 | 3.453E-10 | 82.17 |
| ENSMUSG00000028681 | patched 2 [Source:MGI Symbol;Acc:MGI:1095405] | Ptch2 | Enriched in SCN | 1.576 | 0.01414 | 9.607 |
| ENSMUSG00000037905 | Bri3 binding protein [Source:MGI Symbol;Acc:MGI:1924059] | Bri3bp | Enriched in SCN | 1.573 | 1.729E-08 | 774.6 |
| ENSMUSG00000037503 | family with sequence similarity 168, member B [Source:MGI Symbol;Acc:MGI:2448487] | Fam168b | Enriched in SCN | 1.553 | 2.275E-09 | 1846 |
| ENSMUSG00000027797 | doublecortin-like kinase 1 [Source:MGI Symbol;Acc:MGI:1330861] | Dclk1 | Enriched in SCN | 1.552 | 3.201E-13 | 1464 |
| ENSMUSG00000027983 | cytochrome P450, family 2, subfamily u, polypeptide 1 [Source:MGI Symbol;Acc:MGI:1918769] | Cyp2u1 | Enriched in SCN | 1.549 | 3.687E-10 | 119.4 |
| ENSMUSG00000058297 | sparc/osteonectin, cwcv and kazal-like domains proteoglycan 2 [Source:MGI Symbol;Acc:MGI:1891351] | Spock2 | Enriched in SCN | 1.547 | 0.000004888 | 7309 |
| ENSMUSG00000031760 | metallothionein 3 [Source:MGI Symbol;Acc:MGI:97173] | Mt3 | Enriched in SCN | 1.54 | 0.00002412 | 131.4 |
| ENSMUSG00000027624 | erythrocyte membrane protein band 4.1 like 1 [Source:MGI Symbol;Acc:MGI:103010] | Epb41l1 | Enriched in SCN | 1.537 | 1.085E-16 | 1373 |
| ENSMUSG00000034993 | vesicle amine transport 1 [Source:MGI Symbol;Acc:MGI:1349450] | Vat1 | Enriched in SCN | 1.519 | 1.254E-07 | 429.8 |
| ENSMUSG00000024002 | bromodomain containing 4 [Source:MGI Symbol;Acc:MGI:1888520] | Brd4 | Enriched in SCN | 1.514 | 0.000001104 | 1667 |
| ENSMUSG00000061911 | myelin transcription factor 1-like [Source:MGI Symbol;Acc:MGI:1100511] | Myt1l | Enriched in SCN | 1.51 | 3.148E-11 | 1383 |
| ENSMUSG00000040928 | S100P binding protein [Source:MGI Symbol;Acc:MGI:1921898] | S100pbp | Enriched in SCN | 1.506 | 5.055E-09 | 257.9 |
| ENSMUSG00000045348 | neuronal tyrosine-phosphorylated phosphoinositide 3-kinase adaptor 1 [Source:MGI Symbol;Acc:MGI:2443880] | Nyap1 | Enriched in SCN | 1.504 | 6.243E-14 | 501.1 |
| ENSMUSG00000042751 | nicotinamide nucleotide adenylyltransferase 2 [Source:MGI Symbol;Acc:MGI:2444155] | Nmnat2 | Enriched in SCN | 1.503 | 2.464E-10 | 932 |
| ENSMUSG00000033854 | potassium channel, subfamily K, member 10 [Source:MGI Symbol;Acc:MGI:1919508] | Kcnk10 | Enriched in SCN | 1.499 | 1.736E-09 | 191.6 |
| ENSMUSG00000042766 | tripartite motif-containing 46 [Source:MGI Symbol;Acc:MGI:2673000] | Trim46 | Enriched in SCN | 1.488 | 5.246E-15 | 200.1 |
| ENSMUSG00000019785 | clavesin 2 [Source:MGI Symbol;Acc:MGI:2443223] | Clvs2 | Enriched in SCN | 1.486 | 1.575E-07 | 86.26 |
| ENSMUSG00000024579 | prenylcysteine oxidase 1 like [Source:MGI Symbol;Acc:MGI:3606062] | Pcyox1l | Enriched in SCN | 1.485 | 7.202E-07 | 127.8 |
| ENSMUSG00000025134 | Aly/REF export factor [Source:MGI Symbol;Acc:MGI:1341044] | Alyref | Enriched in SCN | 1.479 | 0.000001347 | 194.3 |
| ENSMUSG00000040102 | kelch-like 42 [Source:MGI Symbol;Acc:MGI:2444786] | Klhl42 | Enriched in SCN | 1.475 | 9.478E-07 | 203.4 |
| ENSMUSG00000046793 | G protein-coupled receptor 61 [Source:MGI Symbol;Acc:MGI:2441719] | Gpr61 | Enriched in SCN | 1.474 | 0.000003924 | 53.44 |
| ENSMUSG00000049728 | zinc finger protein 668 [Source:MGI Symbol;Acc:MGI:2442943] | Zfp668 | Enriched in SCN | 1.467 | 1.361E-07 | 123.8 |
| ENSMUSG00000058740 | potassium channel, subfamily T, member 1 [Source:MGI Symbol;Acc:MGI:1924627] | Kcnt1 | Enriched in SCN | 1.466 | 0.000004975 | 138.9 |
| ENSMUSG00000032905 | autophagy related 12 [Source:MGI Symbol;Acc:MGI:1914776] | Atg12 | Enriched in SCN | 1.466 | 2.121E-09 | 183.2 |
| ENSMUSG00000060268 | armadillo-like helical domain containing 1 [Source:MGI Symbol;Acc:MGI:2686507] | Armh1 | Enriched in SCN | 1.464 | 0.006453 | 6.738 |
| ENSMUSG00000061601 | piccolo (presynaptic cytomatrix protein) [Source:MGI Symbol;Acc:MGI:1349390] | Pclo | Enriched in SCN | 1.463 | 0.00002035 | 4892 |
| ENSMUSG00000005506 | CUGBP, Elav-like family member 1 [Source:MGI Symbol;Acc:MGI:1342295] | Celf1 | Enriched in SCN | 1.463 | 9.733E-08 | 951.6 |
| ENSMUSG00000020083 | family with sequence similarity 241, member B [Source:MGI Symbol;Acc:MGI:1917144] | Fam241b | Enriched in SCN | 1.462 | 1.155E-09 | 42.28 |
| ENSMUSG00000039546 | adherens junction associated protein 1 [Source:MGI Symbol;Acc:MGI:2685419] | Ajap1 | Enriched in SCN | 1.46 | 3.967E-08 | 293.4 |
| ENSMUSG00000000627 | sema domain, immunoglobulin domain (Ig), TM domain, and short cytoplasmic domain [Source:MGI Symbol;Acc:MGI:1340055] | Sema4f | Enriched in SCN | 1.46 | 3.577E-09 | 125.4 |
| ENSMUSG00000028909 | protein tyrosine phosphatase, receptor type, U [Source:MGI Symbol;Acc:MGI:1321151] | Ptpru | Enriched in SCN | 1.447 | 5.08E-11 | 212.6 |
| ENSMUSG00000025375 | apoptosis-associated tyrosine kinase [Source:MGI Symbol;Acc:MGI:1197518] | Aatk | Enriched in SCN | 1.438 | 1.908E-10 | 1628 |
| ENSMUSG00000006218 | family with sequence similarity 131, member C [Source:MGI Symbol;Acc:MGI:2685539] | Fam131c | Enriched in SCN | 1.434 | 0.00006218 | 42.58 |
| ENSMUSG00000022415 | synaptogyrin 1 [Source:MGI Symbol;Acc:MGI:1328323] | Syngr1 | Enriched in SCN | 1.433 | 4.215E-11 | 1440 |
| ENSMUSG00000021395 | spindlin 1 [Source:MGI Symbol;Acc:MGI:109242] | Spin1 | Enriched in SCN | 1.432 | 0.000002859 | 935.2 |
| ENSMUSG00000047428 | delta like non-canonical Notch ligand 2 [Source:MGI Symbol;Acc:MGI:2146838] | Dlk2 | Enriched in SCN | 1.427 | 0.000001293 | 45.68 |
| ENSMUSG00000038065 | maturin, neural progenitor differentiation regulator homolog (Xenopus) [Source:MGI Symbol;Acc:MGI:1915485] | Mturn | Enriched in SCN | 1.427 | 1.809E-08 | 967.7 |
| ENSMUSG00000039542 | neural cell adhesion molecule 1 [Source:MGI Symbol;Acc:MGI:97281] | Ncam1 | Enriched in SCN | 1.417 | 0.000005705 | 5804 |
| ENSMUSG00000032503 | cyclic AMP-regulated phosphoprotein, 21 [Source:MGI Symbol;Acc:MGI:107562] | Arpp21 | Enriched in SCN | 1.415 | 0.0004758 | 594.6 |
| ENSMUSG00000022812 | glycogen synthase kinase 3 beta [Source:MGI Symbol;Acc:MGI:1861437] | Gsk3b | Enriched in SCN | 1.414 | 0.0000339 | 2445 |
| ENSMUSG00000048148 | NACHT and WD repeat domain containing 1 [Source:MGI Symbol;Acc:MGI:2442268] | Nwd1 | Enriched in SCN | 1.411 | 1.094E-09 | 275.4 |
| ENSMUSG00000087408 | ceramide synthase 1 [Source:MGI Symbol;Acc:MGI:2136690] | Cers1 | Enriched in SCN | 1.411 | 7.258E-14 | 94.13 |
| ENSMUSG00000004933 | megakaryocyte-associated tyrosine kinase [Source:MGI Symbol;Acc:MGI:99259] | Matk | Enriched in SCN | 1.408 | 1.425E-08 | 175.5 |
| ENSMUSG00000046378 | aspartate beta-hydroxylase domain containing 1 [Source:MGI Symbol;Acc:MGI:2685014] | Asphd1 | Enriched in SCN | 1.403 | 0.00004801 | 56.65 |
| ENSMUSG00000056124 | UDP-Gal:betaGlcNAc beta 1,4-galactosyltransferase, polypeptide 6 [Source:MGI Symbol;Acc:MGI:1928380] | B4galt6 | Enriched in SCN | 1.4 | 1.246E-17 | 417.7 |
| ENSMUSG00000035064 | eukaryotic elongation factor-2 kinase [Source:MGI Symbol;Acc:MGI:1195261] | Eef2k | Enriched in SCN | 1.392 | 3.553E-11 | 118.4 |
| ENSMUSG00000042532 | golgi autoantigen, golgin subfamily a, 7B [Source:MGI Symbol;Acc:MGI:1918396] | Golga7b | Enriched in SCN | 1.391 | 7.209E-12 | 283.6 |
| ENSMUSG00000027200 | sema domain, transmembrane domain (TM), and cytoplasmic domain, (semaphorin) 6D [Source:MGI Symbol;Acc:MGI:2387661] | Sema6d | Enriched in SCN | 1.387 | 0.000004031 | 731.9 |
| ENSMUSG00000000489 | platelet derived growth factor, B polypeptide [Source:MGI Symbol;Acc:MGI:97528] | Pdgfb | Enriched in SCN | 1.383 | 0.000002771 | 70.92 |
| ENSMUSG00000043051 | disrupted in schizophrenia 1 [Source:MGI Symbol;Acc:MGI:2447658] | Disc1 | Enriched in SCN | 1.38 | 0.0006038 | 34.75 |
| ENSMUSG00000042246 | transmembrane channel-like gene family 7 [Source:MGI Symbol;Acc:MGI:2443317] | Tmc7 | Enriched in SCN | 1.374 | 0.00001301 | 45.35 |
| ENSMUSG00000026434 | nuclear casein kinase and cyclin-dependent kinase substrate 1 [Source:MGI Symbol;Acc:MGI:1934811] | Nucks1 | Enriched in SCN | 1.372 | 0.0009723 | 1384 |
| ENSMUSG00000024400 | WD repeat domain 33 [Source:MGI Symbol;Acc:MGI:1921570] | Wdr33 | Enriched in SCN | 1.372 | 4.82E-09 | 635.9 |
| ENSMUSG00000048486 | fat storage-inducing transmembrane protein 2 [Source:MGI Symbol;Acc:MGI:2444508] | Fitm2 | Enriched in SCN | 1.371 | 3.191E-07 | 173 |
| ENSMUSG00000041037 | immunity-related GTPase family, Q [Source:MGI Symbol;Acc:MGI:2667176] | Irgq | Enriched in SCN | 1.371 | 2.085E-09 | 618.2 |
| ENSMUSG00000022698 | N(alpha)-acetyltransferase 50, NatE catalytic subunit [Source:MGI Symbol;Acc:MGI:1919367] | Naa50 | Enriched in SCN | 1.37 | 0.000003192 | 245.8 |
| ENSMUSG00000051790 | neuroligin 2 [Source:MGI Symbol;Acc:MGI:2681835] | Nlgn2 | Enriched in SCN | 1.368 | 2.223E-12 | 1807 |
| ENSMUSG00000028782 | adhesion G protein-coupled receptor B2 [Source:MGI Symbol;Acc:MGI:2451244] | Adgrb2 | Enriched in SCN | 1.366 | 4.66E-10 | 1121 |
| ENSMUSG00000025743 | syndecan 3 [Source:MGI Symbol;Acc:MGI:1349163] | Sdc3 | Enriched in SCN | 1.362 | 1.262E-07 | 1949 |
| ENSMUSG00000018547 | phosphatidylinositol-5-phosphate 4-kinase, type II, beta [Source:MGI Symbol;Acc:MGI:1934234] | Pip4k2b | Enriched in SCN | 1.362 | 6.03E-09 | 806.6 |
| ENSMUSG00000026458 | protein tyrosine phosphatase, receptor type, f polypeptide (PTPRF), interacting protein (liprin), alpha 4 [Source:MGI Symbol;Acc:MGI:1915757] | Ppfia4 | Enriched in SCN | 1.36 | 3.99E-18 | 888.5 |
| ENSMUSG00000031841 | cadherin 13 [Source:MGI Symbol;Acc:MGI:99551] | Cdh13 | Enriched in SCN | 1.357 | 0.0005119 | 448.3 |
| ENSMUSG00000027525 | phosphatase and actin regulator 3 [Source:MGI Symbol;Acc:MGI:1921439] | Phactr3 | Enriched in SCN | 1.356 | 1.159E-13 | 523.5 |
| ENSMUSG00000060275 | neuregulin 2 [Source:MGI Symbol;Acc:MGI:1098246] | Nrg2 | Enriched in SCN | 1.353 | 0.001142 | 123.8 |
| ENSMUSG00000062373 | transmembrane protein 65 [Source:MGI Symbol;Acc:MGI:1922118] | Tmem65 | Enriched in SCN | 1.343 | 6.148E-11 | 323.4 |
| ENSMUSG00000044912 | synaptotagmin XVI [Source:MGI Symbol;Acc:MGI:2673872] | Syt16 | Enriched in SCN | 1.341 | 0.0000013 | 222.1 |
| ENSMUSG00000096847 | transmembrane protein 151B [Source:MGI Symbol;Acc:MGI:2685169] | Tmem151b | Enriched in SCN | 1.337 | 1.114E-07 | 1096 |
| ENSMUSG00000038860 | GTPase activating RANGAP domain-like 3 [Source:MGI Symbol;Acc:MGI:2139309] | Garnl3 | Enriched in SCN | 1.336 | 0.000008047 | 238.5 |
| ENSMUSG00000020661 | DNA methyltransferase 3A [Source:MGI Symbol;Acc:MGI:1261827] | Dnmt3a | Enriched in SCN | 1.323 | 3.27E-10 | 3814 |
| ENSMUSG00000039770 | yippee like 5 [Source:MGI Symbol;Acc:MGI:1916937] | Ypel5 | Enriched in SCN | 1.311 | 0.00008325 | 207.1 |
| ENSMUSG00000039159 | ubiquitin-conjugating enzyme E2H [Source:MGI Symbol;Acc:MGI:104632] | Ube2h | Enriched in SCN | 1.299 | 2.387E-10 | 558.1 |
| ENSMUSG00000027394 | tubulin tyrosine ligase [Source:MGI Symbol;Acc:MGI:1916987] | Ttl | Enriched in SCN | 1.298 | 0.00000088 | 343 |
| ENSMUSG00000023827 | 1-acylglycerol-3-phosphate O-acyltransferase 4 (lysophosphatidic acid acyltransferase, delta) [Source:MGI Symbol;Acc:MGI:1915512] | Agpat4 | Enriched in SCN | 1.298 | 1.285E-08 | 114.1 |
| ENSMUSG00000022340 | syntabulin (syntaxin-interacting) [Source:MGI Symbol;Acc:MGI:2442392] | Sybu | Enriched in SCN | 1.293 | 1.856E-08 | 257.9 |
| ENSMUSG00000028949 | SWI/SNF related, matrix associated, actin dependent regulator of chromatin, subfamily d, member 3 [Source:MGI Symbol;Acc:MGI:1914243] | Smarcd3 | Enriched in SCN | 1.293 | 2.914E-16 | 234.8 |
| ENSMUSG00000028876 | Eph receptor A10 [Source:MGI Symbol;Acc:MGI:3586824] | Epha10 | Enriched in SCN | 1.292 | 3.058E-07 | 202.5 |
| ENSMUSG00000051146 | calcium/calmodulin-dependent protein kinase II inhibitor 2 [Source:MGI Symbol;Acc:MGI:1920297] | Camk2n2 | Enriched in SCN | 1.292 | 2.542E-09 | 513.1 |
| ENSMUSG00000047261 | growth associated protein 43 [Source:MGI Symbol;Acc:MGI:95639] | Gap43 | Enriched in SCN | 1.273 | 0.000003113 | 876.7 |
| ENSMUSG00000038884 | shiftless antiviral inhibitor of ribosomal frameshifting [Source:MGI Symbol;Acc:MGI:2441788] | Shfl | Enriched in SCN | 1.267 | 0.000004214 | 133.8 |
| ENSMUSG00000002771 | glutamate receptor, ionotropic, NMDA2D (epsilon 4) [Source:MGI Symbol;Acc:MGI:95823] | Grin2d | Enriched in SCN | 1.262 | 4.285E-08 | 466.7 |
| ENSMUSG00000025576 | RNA binding protein, fox-1 homolog (C. elegans) 3 [Source:MGI Symbol;Acc:MGI:106368] | Rbfox3 | Enriched in SCN | 1.258 | 1.753E-07 | 388.1 |
| ENSMUSG00000036815 | dipeptidylpeptidase 10 [Source:MGI Symbol;Acc:MGI:2442409] | Dpp10 | Enriched in SCN | 1.255 | 0.001706 | 576.4 |
| ENSMUSG00000036377 | capping protein inhibiting regulator of actin [Source:MGI Symbol;Acc:MGI:2444817] | Cracd | Enriched in SCN | 1.238 | 0.000001541 | 1779 |
| ENSMUSG00000022044 | stathmin-like 4 [Source:MGI Symbol;Acc:MGI:1931224] | Stmn4 | Enriched in SCN | 1.229 | 0.0002845 | 638.3 |
| ENSMUSG00000004558 | N-myc downstream regulated gene 2 [Source:MGI Symbol;Acc:MGI:1352498] | Ndrg2 | Enriched in SCN | 1.228 | 0.00002502 | 450.7 |
| ENSMUSG00000038077 | potassium voltage-gated channel, shaker-related, subfamily, member 6 [Source:MGI Symbol;Acc:MGI:96663] | Kcna6 | Enriched in SCN | 1.227 | 2.691E-11 | 491.8 |
| ENSMUSG00000092216 | predicted gene, 19345 [Source:MGI Symbol;Acc:MGI:5011530] | Gm19345 | Enriched in SCN | 1.225 | 0.0004727 | 23.03 |
| ENSMUSG00000058153 | seizure related 6 homolog like [Source:MGI Symbol;Acc:MGI:1935121] | Sez6l | Enriched in SCN | 1.223 | 7.009E-07 | 1179 |
| ENSMUSG00000063511 | small nuclear ribonucleoprotein 70 (U1) [Source:MGI Symbol;Acc:MGI:98341] | Snrnp70 | Enriched in SCN | 1.221 | 0.004293 | 1544 |
| ENSMUSG00000030411 | NOVA alternative splicing regulator 2 [Source:MGI Symbol;Acc:MGI:104296] | Nova2 | Enriched in SCN | 1.22 | 2.034E-09 | 858.7 |
| ENSMUSG00000043424 | eukaryotic translation initiation factor 3, subunit J2 [Source:MGI Symbol;Acc:MGI:3704486] | Eif3j2 | Enriched in SCN | 1.219 | 0.00179 | 59.28 |
| ENSMUSG00000071359 | TATA box binding protein-like 1 [Source:MGI Symbol;Acc:MGI:1339946] | Tbpl1 | Enriched in SCN | 1.218 | 0.00008251 | 195 |
| ENSMUSG00000055409 | NEL-like 1 [Source:MGI Symbol;Acc:MGI:2443902] | Nell1 | Enriched in SCN | 1.215 | 0.004615 | 543.3 |
| ENSMUSG00000021770 | sterile alpha motif domain containing 8 [Source:MGI Symbol;Acc:MGI:1914880] | Samd8 | Enriched in SCN | 1.215 | 0.00002404 | 676.8 |
| ENSMUSG00000068099 | small integral membrane protein 45 [Source:MGI Symbol;Acc:MGI:1923755] | Smim45 | Enriched in SCN | 1.206 | 0.00002657 | 83.12 |
| ENSMUSG00000033597 | CASK interacting protein 1 [Source:MGI Symbol;Acc:MGI:2442952] | Caskin1 | Enriched in SCN | 1.204 | 1.46E-10 | 1073 |
| ENSMUSG00000046182 | GSG1-like [Source:MGI Symbol;Acc:MGI:2685483] | Gsg1l | Enriched in SCN | 1.201 | 6.527E-07 | 471.7 |
| ENSMUSG00000003273 | carbonic anhydrase 11 [Source:MGI Symbol;Acc:MGI:1336193] | Car11 | Enriched in SCN | 1.194 | 0.000002785 | 105.3 |
| ENSMUSG00000040811 | echinoderm microtubule associated protein like 2 [Source:MGI Symbol;Acc:MGI:1919455] | Eml2 | Enriched in SCN | 1.192 | 1.963E-07 | 740.8 |
| ENSMUSG00000047155 | cytochrome P450, family 4, subfamily x, polypeptide 1 [Source:MGI Symbol;Acc:MGI:1932403] | Cyp4x1 | Enriched in SCN | 1.191 | 0.0002087 | 55.68 |
| ENSMUSG00000027162 | lin-7 homolog C, crumbs cell polarity complex component [Source:MGI Symbol;Acc:MGI:1330839] | Lin7c | Enriched in SCN | 1.189 | 0.00005609 | 403.1 |
| ENSMUSG00000018411 | microtubule-associated protein tau [Source:MGI Symbol;Acc:MGI:97180] | Mapt | Enriched in SCN | 1.187 | 0.00001656 | 1946 |
| ENSMUSG00000035051 | DEAH (Asp-Glu-Ala-Asp/His) box polypeptide 57 [Source:MGI Symbol;Acc:MGI:2147067] | Dhx57 | Enriched in SCN | 1.186 | 1.312E-07 | 248.4 |
| ENSMUSG00000024921 | SWI/SNF related, matrix associated, actin dependent regulator of chromatin, subfamily a, member 2 [Source:MGI Symbol;Acc:MGI:99603] | Smarca2 | Enriched in SCN | 1.183 | 3.012E-09 | 1435 |
| ENSMUSG00000034353 | receptor (calcitonin) activity modifying protein 1 [Source:MGI Symbol;Acc:MGI:1858418] | Ramp1 | Enriched in SCN | 1.18 | 0.003877 | 54.71 |
| ENSMUSG00000029471 | calcium/calmodulin-dependent protein kinase kinase 2, beta [Source:MGI Symbol;Acc:MGI:2444812] | Camkk2 | Enriched in SCN | 1.178 | 1.516E-07 | 438.3 |
| ENSMUSG00000040722 | secretory carrier membrane protein 5 [Source:MGI Symbol;Acc:MGI:1928948] | Scamp5 | Enriched in SCN | 1.172 | 3.125E-09 | 968.8 |
| ENSMUSG00000029757 | dynein cytoplasmic 1 intermediate chain 1 [Source:MGI Symbol;Acc:MGI:107743] | Dync1i1 | Enriched in SCN | 1.169 | 0.002017 | 339.1 |
| ENSMUSG00000010825 | glutamate receptor, ionotropic, delta 2 (Grid2) interacting protein 1 [Source:MGI Symbol;Acc:MGI:2176213] | Grid2ip | Enriched in SCN | 1.169 | 0.001696 | 357.2 |
| ENSMUSG00000040761 | spen family transcription repressor [Source:MGI Symbol;Acc:MGI:1891706] | Spen | Enriched in SCN | 1.169 | 0.0001662 | 2216 |
| ENSMUSG00000041040 | family with sequence similarity 117, member B [Source:MGI Symbol;Acc:MGI:1920000] | Fam117b | Enriched in SCN | 1.169 | 0.000002013 | 418.6 |
| ENSMUSG00000026972 | arrestin domain containing 1 [Source:MGI Symbol;Acc:MGI:2446136] | Arrdc1 | Enriched in SCN | 1.166 | 0.003418 | 25.53 |
| ENSMUSG00000020440 | ADP-ribosylation factor 5 [Source:MGI Symbol;Acc:MGI:99434] | Arf5 | Enriched in SCN | 1.16 | 0.00002327 | 360.3 |
| ENSMUSG00000033020 | polymerase (RNA) II (DNA directed) polypeptide F [Source:MGI Symbol;Acc:MGI:1349393] | Polr2f | Enriched in SCN | 1.154 | 0.000001254 | 105.7 |
| ENSMUSG00000026278 | BCL2-related ovarian killer [Source:MGI Symbol;Acc:MGI:1858494] | Bok | Enriched in SCN | 1.147 | 0.0013 | 219.1 |
| ENSMUSG00000030806 | syntaxin 1B [Source:MGI Symbol;Acc:MGI:1930705] | Stx1b | Enriched in SCN | 1.147 | 0.00008247 | 1893 |
| ENSMUSG00000020658 | EFR3 homolog B [Source:MGI Symbol;Acc:MGI:2444851] | Efr3b | Enriched in SCN | 1.146 | 5.362E-09 | 856.7 |
| ENSMUSG00000062929 | cofilin 2, muscle [Source:MGI Symbol;Acc:MGI:101763] | Cfl2 | Enriched in SCN | 1.143 | 0.0006238 | 177.2 |
| ENSMUSG00000005986 | ankyrin repeat domain 13 family, member D [Source:MGI Symbol;Acc:MGI:1915673] | Ankrd13d | Enriched in SCN | 1.143 | 2.495E-07 | 148.5 |

| | | | | | | |
|---|---|---|---|---|---|---|
| ENSMUSG00000074698 | casein kinase 2, alpha 1 polypeptide [Source:MGI Symbol;Acc:MGI:88543] | Csnk2a1 | Enriched in SCN | 1.142 | 0.00004853 | 310.6 |
| ENSMUSG00000021294 | kinesin family member 26A [Source:MGI Symbol;Acc:MGI:2447072] | Kif26a | Enriched in SCN | 1.14 | 2.375E-07 | 99.43 |
| ENSMUSG00000015354 | procollagen C-endopeptidase enhancer 2 [Source:MGI Symbol;Acc:MGI:1923727] | Pcolce2 | Enriched in SCN | 1.132 | 0.03332 | 23.56 |
| ENSMUSG00000000568 | heterogeneous nuclear ribonucleoprotein D [Source:MGI Symbol;Acc:MGI:101947] | Hnrnpd | Enriched in SCN | 1.131 | 4.854E-07 | 722.6 |
| ENSMUSG00000039047 | phosphatidylinositol glycan anchor biosynthesis, class K [Source:MGI Symbol;Acc:MGI:1913863] | Pigk | Enriched in SCN | 1.125 | 9.658E-08 | 388.2 |
| ENSMUSG00000029578 | WD repeat domain, phosphoinositide interacting 2 [Source:MGI Symbol;Acc:MGI:1923831] | Wipi2 | Enriched in SCN | 1.12 | 1.293E-08 | 376.9 |
| ENSMUSG00000035226 | regulating synaptic membrane exocytosis 4 [Source:MGI Symbol;Acc:MGI:2674366] | Rims4 | Enriched in SCN | 1.119 | 2.25E-10 | 674.1 |
| ENSMUSG00000042202 | solute carrier family 35, member E2 [Source:MGI Symbol;Acc:MGI:2444240] | Slc35e2 | Enriched in SCN | 1.118 | 0.000002007 | 518.7 |
| ENSMUSG00000021974 | fibroblast growth factor 9 [Source:MGI Symbol;Acc:MGI:104723] | Fgf9 | Enriched in SCN | 1.116 | 0.005211 | 263.9 |
| ENSMUSG00000029095 | actin-binding LIM protein 2 [Source:MGI Symbol;Acc:MGI:2385758] | Ablim2 | Enriched in SCN | 1.116 | 0.00001168 | 268.3 |
| ENSMUSG00000030067 | forkhead box P1 [Source:MGI Symbol;Acc:MGI:1914004] | Foxp1 | Enriched in SCN | 1.114 | 0.0002717 | 337.2 |
| ENSMUSG00000048895 | cyclin-dependent kinase 5, regulatory subunit 1 (p35) [Source:MGI Symbol;Acc:MGI:101764] | Cdk5r1 | Enriched in SCN | 1.113 | 0.0002174 | 1376 |
| ENSMUSG00000056429 | trans-golgi network protein [Source:MGI Symbol;Acc:MGI:105080] | Tgoln1 | Enriched in SCN | 1.107 | 0.0004038 | 521.3 |
| ENSMUSG00000056367 | ARP3 actin-related protein 3B [Source:MGI Symbol;Acc:MGI:2661120] | Actr3b | Enriched in SCN | 1.104 | 1.233E-07 | 140.3 |
| ENSMUSG00000041020 | MAP7 domain containing 2 [Source:MGI Symbol;Acc:MGI:1917474] | Map7d2 | Enriched in SCN | 1.103 | 0.00002393 | 736.5 |
| ENSMUSG00000020297 | neuron specific gene family member 2 [Source:MGI Symbol;Acc:MGI:1202070] | Nsg2 | Enriched in SCN | 1.095 | 2.271E-09 | 922.9 |
| ENSMUSG00000073755 | RIKEN cDNA 5730409E04Rik gene [Source:MGI Symbol;Acc:MGI:3609248] | 5730409E04Rik | Enriched in SCN | 1.089 | 0.00001107 | 353.7 |
| ENSMUSG00000025854 | FAM20C, golgi associated secretory pathway kinase [Source:MGI Symbol;Acc:MGI:2136853] | Fam20c | Enriched in SCN | 1.088 | 0.000001026 | 465.1 |
| ENSMUSG00000095098 | coiled-coil domain containing 85B [Source:MGI Symbol;Acc:MGI:2147607] | Ccdc85b | Enriched in SCN | 1.087 | 0.00007907 | 384.5 |
| ENSMUSG00000055302 | Morf4 family associated protein 1 [Source:MGI Symbol;Acc:MGI:1914818] | Mrfap1 | Enriched in SCN | 1.085 | 0.0002434 | 1483 |
| ENSMUSG00000029348 | aspartate beta-hydroxylase domain containing 2 [Source:MGI Symbol;Acc:MGI:1920148] | Asphd2 | Enriched in SCN | 1.084 | 0.00008858 | 128.4 |
| ENSMUSG00000068923 | synaptotagmin XI [Source:MGI Symbol;Acc:MGI:1859547] | Syt11 | Enriched in SCN | 1.079 | 0.000004555 | 1678 |
| ENSMUSG00000073433 | Rho GDP dissociation inhibitor (GDI) gamma [Source:MGI Symbol;Acc:MGI:108430] | Arhgdig | Enriched in SCN | 1.077 | 0.000248 | 185.1 |
| ENSMUSG00000020654 | adenylate cyclase 3 [Source:MGI Symbol;Acc:MGI:99675] | Adcy3 | Enriched in SCN | 1.077 | 1.106E-08 | 175 |
| ENSMUSG00000071072 | prostaglandin E synthase 3 [Source:MGI Symbol;Acc:MGI:1929282] | Ptges3 | Enriched in SCN | 1.075 | 0.0001135 | 305.5 |
| ENSMUSG00000041313 | solute carrier family 7 (cationic amino acid transporter, y+ system), member 1 [Source:MGI Symbol;Acc:MGI:88117] | Slc7a1 | Enriched in SCN | 1.067 | 0.00285 | 345.4 |
| ENSMUSG00000039419 | contactin associated protein-like 2 [Source:MGI Symbol;Acc:MGI:1914047] | Cntnap2 | Enriched in SCN | 1.056 | 0.02456 | 1307 |
| ENSMUSG00000019370 | calmodulin 3 [Source:MGI Symbol;Acc:MGI:103249] | Calm3 | Enriched in SCN | 1.056 | 0.00002558 | 6755 |
| ENSMUSG00000020331 | hyperpolarization-activated, cyclic nucleotide-gated K+ 2 [Source:MGI Symbol;Acc:MGI:1298210] | Hcn2 | Enriched in SCN | 1.054 | 0.000001692 | 598.2 |
| ENSMUSG00000074657 | kinesin family member 5A [Source:MGI Symbol;Acc:MGI:109564] | Kif5a | Enriched in SCN | 1.05 | 0.0007454 | 6146 |
| ENSMUSG00000034342 | Casitas B-lineage lymphoma [Source:MGI Symbol;Acc:MGI:88279] | Cbl | Enriched in SCN | 1.05 | 9.821E-07 | 484.4 |
| ENSMUSG00000027669 | guanine nucleotide binding protein (G protein), beta 4 [Source:MGI Symbol;Acc:MGI:104581] | Gnb4 | Enriched in SCN | 1.05 | 7.818E-07 | 103.7 |
| ENSMUSG00000041654 | solute carrier family 39 (metal ion transporter), member 11 [Source:MGI Symbol;Acc:MGI:1917056] | Slc39a11 | Enriched in SCN | 1.049 | 0.0008746 | 62.34 |
| ENSMUSG00000050967 | cellular repressor of E1A-stimulated genes 2 [Source:MGI Symbol;Acc:MGI:1928333] | Creg2 | Enriched in SCN | 1.049 | 0.0002356 | 188.8 |
| ENSMUSG00000064179 | troponin T1, skeletal, slow [Source:MGI Symbol;Acc:MGI:1333868] | Tnnt1 | Enriched in SCN | 1.041 | 0.04726 | 308.7 |
| ENSMUSG00000000532 | activin A receptor, type 1B [Source:MGI Symbol;Acc:MGI:1338944] | Acvr1b | Enriched in SCN | 1.036 | 8.1E-11 | 463 |
| ENSMUSG00000026915 | spermatid perinuclear RNA binding protein [Source:MGI Symbol;Acc:MGI:104626] | Strbp | Enriched in SCN | 1.034 | 0.0002854 | 2337 |
| ENSMUSG00000030019 | F-box and leucine-rich repeat protein 14 [Source:MGI Symbol;Acc:MGI:2141676] | Fbxl14 | Enriched in SCN | 1.031 | 0.003046 | 241.9 |
| ENSMUSG00000031137 | fibroblast growth factor 13 [Source:MGI Symbol;Acc:MGI:109178] | Fgf13 | Enriched in SCN | 1.029 | 0.001065 | 419.6 |
| ENSMUSG00000056222 | sparc/osteonectin, cwcv and kazal-like domains proteoglycan 1 [Source:MGI Symbol;Acc:MGI:105371] | Spock1 | Enriched in SCN | 1.027 | 0.002244 | 1204 |
| ENSMUSG00000061536 | SEC22 homolog C, vesicle trafficking protein [Source:MGI Symbol;Acc:MGI:2447871] | Sec22c | Enriched in SCN | 1.027 | 0.0006565 | 263.2 |
| ENSMUSG00000035770 | dynein, cytoplasmic 1 light intermediate chain 2 [Source:MGI Symbol;Acc:MGI:107738] | Dync1li2 | Enriched in SCN | 1.019 | 0.0002039 | 1131 |
| ENSMUSG00000075318 | sodium channel, voltage-gated, type II, alpha [Source:MGI Symbol;Acc:MGI:98248] | Scn2a | Enriched in SCN | 1.005 | 0.00000296 | 1334 |
| ENSMUSG00000014592 | calmodulin binding transcription activator 1 [Source:MGI Symbol;Acc:MGI:2140230] | Camta1 | Enriched in SCN | 1.003 | 8.136E-10 | 1945 |
| ENSMUSG00000039195 | bublin coiled coil protein [Source:MGI Symbol;Acc:MGI:1920987] | Bbln | Enriched in retina | -1.001 | 0.002197 | 50.47 |
| ENSMUSG00000033436 | armadillo repeat containing, X-linked 2 [Source:MGI Symbol;Acc:MGI:1914666] | Armcx2 | Enriched in retina | -1.003 | 0.000005693 | 262.7 |
| ENSMUSG00000048332 | lipoma HMGIC fusion partner [Source:MGI Symbol;Acc:MGI:1920048] | Lhfp | Enriched in retina | -1.011 | 0.01613 | 294.9 |
| ENSMUSG00000034891 | synuclein, beta [Source:MGI Symbol;Acc:MGI:1889011] | Sncb | Enriched in retina | -1.018 | 0.001316 | 939.2 |
| ENSMUSG00000028630 | dual-specificity tyrosine-(Y)-phosphorylation regulated kinase 2 [Source:MGI Symbol;Acc:MGI:1330301] | Dyrk2 | Enriched in retina | -1.025 | 0.00004804 | 685.8 |
| ENSMUSG00000023262 | aminoacylase 1 [Source:MGI Symbol;Acc:MGI:87913] | Acy1 | Enriched in retina | -1.028 | 0.03196 | 15.88 |
| ENSMUSG00000018634 | corticotropin releasing hormone receptor 1 [Source:MGI Symbol;Acc:MGI:88498] | Crhr1 | Enriched in retina | -1.032 | 0.005733 | 47.56 |
| ENSMUSG00000027314 | delta like canonical Notch ligand 4 [Source:MGI Symbol;Acc:MGI:1859388] | Dll4 | Enriched in retina | -1.054 | 0.03967 | 11.29 |
| ENSMUSG00000039461 | T cell leukemia translocation altered gene [Source:MGI Symbol;Acc:MGI:1918829] | Tcta | Enriched in retina | -1.077 | 0.000005104 | 50.88 |
| ENSMUSG00000022885 | beta galactoside alpha 2,6 sialyltransferase 1 [Source:MGI Symbol;Acc:MGI:108470] | St6gal1 | Enriched in retina | -1.093 | 0.000604 | 86.45 |
| ENSMUSG00000030782 | transforming growth factor beta 1 induced transcript 1 [Source:MGI Symbol;Acc:MGI:102784] | Tgfb1i1 | Enriched in retina | -1.101 | 0.0005399 | 24.56 |
| ENSMUSG00000004952 | RAS p21 protein activator 4 [Source:MGI Symbol;Acc:MGI:1858600] | Rasa4 | Enriched in retina | -1.116 | 0.002279 | 77.45 |
| ENSMUSG00000028631 | potassium voltage-gated channel, subfamily Q, member 4 [Source:MGI Symbol;Acc:MGI:1926803] | Kcnq4 | Enriched in retina | -1.134 | 0.001101 | 45.84 |
| ENSMUSG00000021219 | regulator of G-protein signaling 6 [Source:MGI Symbol;Acc:MGI:1354730] | Rgs6 | Enriched in retina | -1.163 | 0.008101 | 225 |
| ENSMUSG00000028937 | acyl-CoA thioesterase 7 [Source:MGI Symbol;Acc:MGI:1917275] | Acot7 | Enriched in retina | -1.174 | 0.01432 | 554 |
| ENSMUSG00000002055 | sperm associated antigen 5 [Source:MGI Symbol;Acc:MGI:1927470] | Spag5 | Enriched in retina | -1.18 | 0.001478 | 34.31 |
| ENSMUSG00000052560 | copine VIII [Source:MGI Symbol;Acc:MGI:1914121] | Cpne8 | Enriched in retina | -1.189 | 0.0003105 | 103.9 |
| ENSMUSG00000033174 | monoglyceride lipase [Source:MGI Symbol;Acc:MGI:1346042] | Mgll | Enriched in retina | -1.191 | 0.000002542 | 422.8 |
| ENSMUSG00000024446 | ribonuclease P 21 subunit [Source:MGI Symbol;Acc:MGI:1914926] | Rpp21 | Enriched in retina | -1.224 | 0.006181 | 21.09 |
| ENSMUSG00000052387 | transient receptor potential cation channel, subfamily M, member 3 [Source:MGI Symbol;Acc:MGI:2443101] | Trpm3 | Enriched in retina | -1.228 | 0.0003495 | 2184 |
| ENSMUSG00000041592 | sidekick cell adhesion molecule 2 [Source:MGI Symbol;Acc:MGI:2443847] | Sdk2 | Enriched in retina | -1.234 | 0.000004332 | 398.3 |
| ENSMUSG00000033615 | complexin 1 [Source:MGI Symbol;Acc:MGI:104727] | Cplx1 | Enriched in retina | -1.241 | 0.0008611 | 2304 |
| ENSMUSG00000041046 | receptor (calcitonin) activity modifying protein 3 [Source:MGI Symbol;Acc:MGI:1860292] | Ramp3 | Enriched in retina | -1.274 | 0.0006774 | 121.1 |
| ENSMUSG00000032011 | thymus cell antigen 1, theta [Source:MGI Symbol;Acc:MGI:98747] | Thy1 | Enriched in retina | -1.307 | 0.0001922 | 952.8 |
| ENSMUSG00000039662 | isoprenylcysteine carboxyl methyltransferase [Source:MGI Symbol;Acc:MGI:1888594] | Icmt | Enriched in retina | -1.325 | 9.641E-07 | 734.9 |
| ENSMUSG00000000326 | catechol-O-methyltransferase [Source:MGI Symbol;Acc:MGI:88470] | Comt | Enriched in retina | -1.353 | 2.898E-09 | 103.3 |
| ENSMUSG00000060240 | cell cycle exit and neuronal differentiation 1 [Source:MGI Symbol;Acc:MGI:1929898] | Cend1 | Enriched in retina | -1.367 | 0.001598 | 473.2 |
| ENSMUSG00000035407 | KN motif and ankyrin repeat domains 4 [Source:MGI Symbol;Acc:MGI:3043381] | Kank4 | Enriched in retina | -1.373 | 0.004353 | 22.26 |
| ENSMUSG00000003657 | calbindin 2 [Source:MGI Symbol;Acc:MGI:101914] | Calb2 | Enriched in retina | -1.373 | 0.0002748 | 847.8 |
| ENSMUSG00000020836 | coronin 6 [Source:MGI Symbol;Acc:MGI:2183448] | Coro6 | Enriched in retina | -1.393 | 0.04848 | 39.45 |
| ENSMUSG00000048899 | ribosomal modification protein rimK-like family member A [Source:MGI Symbol;Acc:MGI:3040686] | Rimkla | Enriched in retina | -1.41 | 2.903E-09 | 161 |
| ENSMUSG00000050822 | solute carrier family 29 (nucleoside transporters), member 4 [Source:MGI Symbol;Acc:MGI:2385330] | Slc29a4 | Enriched in retina | -1.431 | 0.00002388 | 126.5 |
| ENSMUSG00000011256 | a disintegrin and metallopeptidase domain 19 (meltrin beta) [Source:MGI Symbol;Acc:MGI:105377] | Adam19 | Enriched in retina | -1.449 | 6.136E-07 | 194.7 |
| ENSMUSG00000020838 | solute carrier family 6 (neurotransmitter transporter, serotonin), member 4 [Source:MGI Symbol;Acc:MGI:96285] | Slc6a4 | Enriched in retina | -1.467 | 0.02143 | 23.26 |
| ENSMUSG00000021728 | embigin [Source:MGI Symbol;Acc:MGI:95321] | Emb | Enriched in retina | -1.475 | 0.00006913 | 88.85 |
| ENSMUSG00000022055 | neurofilament, light polypeptide [Source:MGI Symbol;Acc:MGI:97313] | Nefl | Enriched in retina | -1.515 | 0.000007701 | 7224 |
| ENSMUSG00000040373 | calcium channel, voltage-dependent, gamma subunit 5 [Source:MGI Symbol;Acc:MGI:2157946] | Cacng5 | Enriched in retina | -1.573 | 2.225E-09 | 203.8 |
| ENSMUSG00000024942 | calpain 1 [Source:MGI Symbol;Acc:MGI:88263] | Capn1 | Enriched in retina | -1.576 | 0.0004866 | 59.73 |
| ENSMUSG00000022610 | mitogen-activated protein kinase 12 [Source:MGI Symbol;Acc:MGI:1353438] | Mapk12 | Enriched in retina | -1.581 | 0.0157 | 13.25 |
| ENSMUSG00000020396 | neurofilament, heavy polypeptide [Source:MGI Symbol;Acc:MGI:97309] | Nefh | Enriched in retina | -1.584 | 0.0001674 | 1828 |
| ENSMUSG00000020635 | FK506 binding protein 1b [Source:MGI Symbol;Acc:MGI:1336205] | Fkbp1b | Enriched in retina | -1.586 | 0.00004598 | 66.86 |
| ENSMUSG00000017167 | contactin associated protein-like 1 [Source:MGI Symbol;Acc:MGI:1858201] | Cntnap1 | Enriched in retina | -1.622 | 0.000004686 | 622.8 |
| ENSMUSG00000042115 | kelch domain containing 8A [Source:MGI Symbol;Acc:MGI:2442630] | Klhdc8a | Enriched in retina | -1.647 | 2.403E-10 | 181.2 |
| ENSMUSG00000005716 | parvalbumin [Source:MGI Symbol;Acc:MGI:97821] | Pvalb | Enriched in retina | -1.661 | 0.0499 | 26.7 |
| ENSMUSG00000006307 | lysine (K)-specific methyltransferase 2B [Source:MGI Symbol;Acc:MGI:109565] | Kmt2b | Enriched in retina | -1.716 | 4.151E-10 | 848.2 |
| ENSMUSG00000020953 | cochlin [Source:MGI Symbol;Acc:MGI:1278313] | Coch | Enriched in retina | -1.832 | 0.0003475 | 80.94 |
| ENSMUSG00000001794 | calpain, small subunit 1 [Source:MGI Symbol;Acc:MGI:88266] | Capns1 | Enriched in retina | -1.84 | 6.964E-09 | 150.2 |
| ENSMUSG00000030528 | Bloom syndrome, RecQ like helicase [Source:MGI Symbol;Acc:MGI:1328362] | Blm | Enriched in retina | -1.897 | 8.986E-08 | 50.26 |
| ENSMUSG00000024287 | THO complex 1 [Source:MGI Symbol;Acc:MGI:1919668] | Thoc1 | Enriched in retina | -1.957 | 1.575E-07 | 109.9 |
| ENSMUSG00000001333 | syncoilin [Source:MGI Symbol;Acc:MGI:1916078] | Sync | Enriched in retina | -1.962 | 0.000016 | 17.38 |
| ENSMUSG00000019966 | kit ligand [Source:MGI Symbol;Acc:MGI:96974] | Kitl | Enriched in retina | -1.985 | 4.677E-08 | 453.1 |
| ENSMUSG00000027071 | purinergic receptor P2X, ligand-gated ion channel, 3 [Source:MGI Symbol;Acc:MGI:1097160] | P2rx3 | Enriched in retina | -2.023 | 0.002565 | 12.15 |
| ENSMUSG00000017897 | EYA transcriptional coactivator and phosphatase 2 [Source:MGI Symbol;Acc:MGI:109341] | Eya2 | Enriched in retina | -2.063 | 5.863E-07 | 21.41 |
| ENSMUSG00000018340 | annexin A6 [Source:MGI Symbol;Acc:MGI:88255] | Anxa6 | Enriched in retina | -2.111 | 0.00002916 | 254.2 |
| ENSMUSG00000062168 | protein phosphatase with EF hand calcium-binding domain 1 [Source:MGI Symbol;Acc:MGI:1097157] | Ppef1 | Enriched in retina | -2.121 | 0.003149 | 6.202 |
| ENSMUSG00000032584 | macrophage stimulating 1 receptor (c-met-related tyrosine kinase) [Source:MGI Symbol;Acc:MGI:99614] | Mst1r | Enriched in retina | -2.124 | 0.008988 | 3.48 |
| ENSMUSG00000060257 | scratch family zinc finger 2 [Source:MGI Symbol;Acc:MGI:2139287] | Scrt2 | Enriched in retina | -2.124 | 8.781E-13 | 141.5 |
| ENSMUSG00000068699 | filamin C, gamma [Source:MGI Symbol;Acc:MGI:95557] | Flnc | Enriched in retina | -2.183 | 0.00004299 | 29.35 |
| ENSMUSG00000030110 | ret proto-oncogene [Source:MGI Symbol;Acc:MGI:97902] | Ret | Enriched in retina | -2.23 | 1.328E-12 | 188.3 |
| ENSMUSG00000024697 | guanine nucleotide binding protein, alpha 14 [Source:MGI Symbol;Acc:MGI:95769] | Gna14 | Enriched in retina | -2.24 | 0.02062 | 3.559 |
| ENSMUSG00000047746 | F-box protein 40 [Source:MGI Symbol;Acc:MGI:2443753] | Fbxo40 | Enriched in retina | -2.241 | 0.02456 | 15.08 |
| ENSMUSG00000020732 | RAB37, member RAS oncogene family [Source:MGI Symbol;Acc:MGI:1929945] | Rab37 | Enriched in retina | -2.267 | 0.02206 | 50.93 |
| ENSMUSG00000009734 | POU domain, class 6, transcription factor 2 [Source:MGI Symbol;Acc:MGI:2443631] | Pou6f2 | Enriched in retina | -2.279 | 1.026E-08 | 127.9 |
| ENSMUSG00000002980 | basal cell adhesion molecule [Source:MGI Symbol;Acc:MGI:1929940] | Bcam | Enriched in retina | -2.284 | 1.151E-07 | 83.63 |
| ENSMUSG00000040703 | cytochrome P450, family 2, subfamily s, polypeptide 1 [Source:MGI Symbol;Acc:MGI:1921384] | Cyp2s1 | Enriched in retina | -2.297 | 0.00000445 | 19.07 |
| ENSMUSG00000025930 | musculin [Source:MGI Symbol;Acc:MGI:1333884] | Msc | Enriched in retina | -2.308 | 0.007779 | 4.237 |
| ENSMUSG00000060969 | Iroquois homeobox 1 [Source:MGI Symbol;Acc:MGI:1197515] | Irx1 | Enriched in retina | -2.333 | 0.00003635 | 29.96 |
| ENSMUSG00000021948 | protein kinase C, delta [Source:MGI Symbol;Acc:MGI:97598] | Prkcd | Enriched in retina | -2.478 | 0.00002387 | 727.9 |
| ENSMUSG00000056296 | synaptoporin [Source:MGI Symbol;Acc:MGI:1919253] | Synpr | Enriched in retina | -2.489 | 5.865E-11 | 203.6 |
| ENSMUSG00000030270 | copine family member IX [Source:MGI Symbol;Acc:MGI:2443052] | Cpne9 | Enriched in retina | -2.503 | 0.000001048 | 175.2 |
| ENSMUSG00000038530 | regulator of G-protein signaling 4 [Source:MGI Symbol;Acc:MGI:108409] | Rgs4 | Enriched in retina | -2.52 | 9.523E-10 | 578.5 |
| ENSMUSG00000050296 | ATP-binding cassette, sub-family A (ABC1), member 12 [Source:MGI Symbol;Acc:MGI:2676312] | Abca12 | Enriched in retina | -2.531 | 0.002487 | 4.035 |
| ENSMUSG00000047963 | starch binding domain 1 [Source:MGI Symbol;Acc:MGI:1261768] | Stbd1 | Enriched in retina | -2.558 | 0.000009137 | 7.985 |
| ENSMUSG00000022054 | neurofilament, medium polypeptide [Source:MGI Symbol;Acc:MGI:97314] | Nefm | Enriched in retina | -2.752 | 1.195E-09 | 5610 |
| ENSMUSG00000020216 | junctional sarcoplasmic reticulum protein 1 [Source:MGI Symbol;Acc:MGI:1916700] | Jsrp1 | Enriched in retina | -2.82 | 0.006368 | 6.626 |
| ENSMUSG00000006764 | tryptophan hydroxylase 2 [Source:MGI Symbol;Acc:MGI:2651811] | Tph2 | Enriched in retina | -2.879 | 0.001735 | 3.884 |
| ENSMUSG00000019971 | centrosomal protein 290 [Source:MGI Symbol;Acc:MGI:2384917] | Cep290 | Enriched in retina | -2.917 | 1.315E-15 | 749.1 |
| ENSMUSG00000025401 | myosin IA [Source:MGI Symbol;Acc:MGI:107732] | Myo1a | Enriched in retina | -2.939 | 0.004578 | 4.176 |
| ENSMUSG00000058925 | coiled-coil domain containing 192 [Source:MGI Symbol;Acc:MGI:1922694] | Ccdc192 | Enriched in retina | -3.003 | 0.00006452 | 7.152 |
| ENSMUSG00000025366 | extended synaptotagmin-like protein 1 [Source:MGI Symbol;Acc:MGI:1344426] | Esyt1 | Enriched in retina | -3.142 | 2.285E-13 | 159.4 |
| ENSMUSG00000001504 | Iroquois homeobox 2 [Source:MGI Symbol;Acc:MGI:1197526] | Irx2 | Enriched in retina | -3.178 | 1.225E-10 | 24.13 |
| ENSMUSG00000031734 | Iroquois related homeobox 3 [Source:MGI Symbol;Acc:MGI:1197522] | Irx3 | Enriched in retina | -3.262 | 2.754E-09 | 16.38 |
| ENSMUSG00000033082 | C-type lectin domain family 1, member a [Source:MGI Symbol;Acc:MGI:2444151] | Clec1a | Enriched in retina | -3.268 | 0.00002062 | 6.168 |
| ENSMUSG00000026778 | protein kinase C, theta [Source:MGI Symbol;Acc:MGI:97601] | Prkcq | Enriched in retina | -3.268 | 8.89E-17 | 111.9 |

| | | | | | | |
|---|---|---|---|---|---|---|
| ENSMUSG00000090223 | Purkinje cell protein 4 [Source:MGI Symbol;Acc:MGI:97509] | Pcp4 | Enriched in retina | -3.323 | 0.000000198 | 93.25 |
| ENSMUSG00000041959 | S100 calcium binding protein A10 (calpactin) [Source:MGI Symbol;Acc:MGI:1339468] | S100a10 | Enriched in retina | -3.417 | 1.571E-09 | 41.51 |
| ENSMUSG00000034115 | sodium channel, voltage-gated, type XI, alpha [Source:MGI Symbol;Acc:MGI:1345149] | Scn11a | Enriched in retina | -3.421 | 0.00000453 | 5.567 |
| ENSMUSG00000004098 | collagen, type V, alpha 3 [Source:MGI Symbol;Acc:MGI:1858212] | Col5a3 | Enriched in retina | -3.457 | 1.057E-13 | 67.35 |
| ENSMUSG00000043155 | 4-hydroxyphenylpyruvate dioxygenase-like [Source:MGI Symbol;Acc:MGI:2444646] | Hpdl | Enriched in retina | -3.613 | 0.01299 | 2.85 |
| ENSMUSG00000070570 | solute carrier family 17 (sodium-dependent inorganic phosphate cotransporter), member 7 [Source:MGI Symbol;Acc:MGI:1920211] | Slc17a7 | Enriched in retina | -3.681 | 4.701E-17 | 1629 |
| ENSMUSG00000001119 | collagen, type VI, alpha 1 [Source:MGI Symbol;Acc:MGI:88459] | Col6a1 | Enriched in retina | -3.991 | 2.252E-15 | 82.8 |
| ENSMUSG00000078958 | ATPase, H+ transporting, lysosomal accessory protein 1-like [Source:MGI Symbol;Acc:MGI:3648665] | Atp6ap1l | Enriched in retina | -4.05 | 0.0003255 | 5.851 |
| ENSMUSG00000024871 | double C2, gamma [Source:MGI Symbol;Acc:MGI:1926250] | Doc2g | Enriched in retina | -4.163 | 0.00293 | 6.218 |
| ENSMUSG00000061048 | cadherin 3 [Source:MGI Symbol;Acc:MGI:88356] | Cdh3 | Enriched in retina | -4.284 | 0.000002678 | 8.894 |
| ENSMUSG00000025432 | advillin [Source:MGI Symbol;Acc:MGI:1333798] | Avil | Enriched in retina | -4.348 | 0.0006103 | 4.817 |
| ENSMUSG00000051279 | growth differentiation factor 6 [Source:MGI Symbol;Acc:MGI:95689] | Gdf6 | Enriched in retina | -4.735 | 1.171E-11 | 11.98 |
| ENSMUSG00000031737 | Iroquois homeobox 5 [Source:MGI Symbol;Acc:MGI:1859086] | Irx5 | Enriched in retina | -4.831 | 4.061E-19 | 31.09 |
| ENSMUSG00000032387 | RNA binding protein with multiple splicing 2 [Source:MGI Symbol;Acc:MGI:1919223] | Rbpms2 | Enriched in retina | -4.968 | 1.07E-31 | 47.72 |
| ENSMUSG00000030830 | integrin alpha L [Source:MGI Symbol;Acc:MGI:96606] | Itgal | Enriched in retina | -5.032 | 0.00001931 | 6.323 |
| ENSMUSG00000023484 | peripherin [Source:MGI Symbol;Acc:MGI:97774] | Prph | Enriched in retina | -5.251 | 3.55E-08 | 64.1 |
| ENSMUSG00000031586 | RNA binding protein gene with multiple splicing [Source:MGI Symbol;Acc:MGI:1334446] | Rbpms | Enriched in retina | -5.357 | 1.198E-33 | 81.55 |
| ENSMUSG00000032446 | eomesodermin [Source:MGI Symbol;Acc:MGI:1201683] | Eomes | Enriched in retina | -5.419 | 2.752E-12 | 60.2 |
| ENSMUSG00000021848 | orthodenticle homeobox 2 [Source:MGI Symbol;Acc:MGI:97451] | Otx2 | Enriched in retina | -5.765 | 1.699E-25 | 248.7 |
| ENSMUSG00000048070 | phosphoinositide-interacting regulator of transient receptor potential channels [Source:MGI Symbol;Acc:MGI:2443635] | Pirt | Enriched in retina | -5.767 | 8.609E-09 | 10.02 |
| ENSMUSG00000068220 | lectin, galactose binding, soluble 1 [Source:MGI Symbol;Acc:MGI:96777] | Lgals1 | Enriched in retina | -5.79 | 7.981E-15 | 30.47 |
| ENSMUSG00000002100 | myosin binding protein C, cardiac [Source:MGI Symbol;Acc:MGI:102844] | Mybpc3 | Enriched in retina | -6.939 | 0.000001709 | 4.298 |
| ENSMUSG00000069372 | cortexin 3 [Source:MGI Symbol;Acc:MGI:3642816] | Ctxn3 | Enriched in retina | -8.613 | 5.655E-10 | 38.01 |
| ENSMUSG00000031738 | Iroquois homeobox 6 [Source:MGI Symbol;Acc:MGI:1927642] | Irx6 | Enriched in retina | -9.283 | 5.807E-13 | 50.97 |
| ENSMUSG00000031965 | T-box 20 [Source:MGI Symbol;Acc:MGI:1888496] | Tbx20 | Enriched in retina | -9.741 | 4.228E-13 | 45.45 |
| ENSMUSG00000031688 | POU domain, class 4, transcription factor 2 [Source:MGI Symbol;Acc:MGI:102524] | Pou4f2 | Enriched in retina | -10.2 | 6.078E-15 | 45.2 |
| ENSMUSG00000021799 | opsin 4 (melanopsin) [Source:MGI Symbol;Acc:MGI:1353425] | Opn4 | Enriched in retina | -10.26 | 1.748E-14 | 25.82 |

**Contrast numerator: P8_KO_SCN. Contrast denominator: P8_KO_retina.**

| ensembl_gene_id | description | mgi_symbol | Significance | deseq_logfc | deseq_adjp | deseq_basemean |
|---|---|---|---|---|---|---|
| ENSMUSG00000004892 | brevican [Source:MGI Symbol;Acc:MGI:1096385] | Bcan | Enriched in SCN | 5.457 | 1.476E-85 | 546.7 |
| ENSMUSG00000024798 | 5-hydroxytryptamine (serotonin) receptor 7 [Source:MGI Symbol;Acc:MGI:99841] | Htr7 | Enriched in SCN | 5.054 | 2.973E-40 | 109.3 |
| ENSMUSG00000038011 | dynein, axonemal, heavy chain 10 [Source:MGI Symbol;Acc:MGI:1860299] | Dnah10 | Enriched in SCN | 4.861 | 1.701E-33 | 44.22 |
| ENSMUSG00000062372 | otoferlin [Source:MGI Symbol;Acc:MGI:1891247] | Otof | Enriched in SCN | 4.838 | 1.629E-23 | 68.05 |
| ENSMUSG00000052861 | dynein, axonemal, heavy chain 6 [Source:MGI Symbol;Acc:MGI:107744] | Dnah6 | Enriched in SCN | 4.798 | 1.407E-21 | 52.06 |
| ENSMUSG00000053004 | histamine receptor H1 [Source:MGI Symbol;Acc:MGI:107619] | Hrh1 | Enriched in SCN | 4.717 | 2.455E-17 | 32.83 |
| ENSMUSG00000000560 | gamma-aminobutyric acid (GABA) A receptor, subunit alpha 2 [Source:MGI Symbol;Acc:MGI:95614] | Gabra2 | Enriched in SCN | 4.711 | 2.289E-36 | 394.2 |
| ENSMUSG00000090053 | paralemmin A kinase anchor protein [Source:MGI Symbol;Acc:MGI:5141924] | Pakap | Enriched in SCN | 4.604 | 2.269E-11 | 58.1 |
| ENSMUSG00000042385 | granzyme K [Source:MGI Symbol;Acc:MGI:1298232] | Gzmk | Enriched in SCN | 4.516 | 0.0002294 | 4.859 |
| ENSMUSG00000078816 | protein kinase C, gamma [Source:MGI Symbol;Acc:MGI:97597] | Prkcg | Enriched in SCN | 4.043 | 5.853E-24 | 571.8 |
| ENSMUSG00000048721 | fibronectin type III domain containing 9 [Source:MGI Symbol;Acc:MGI:2443410] | Fndc9 | Enriched in SCN | 3.974 | 0.000009289 | 15.76 |
| ENSMUSG00000036634 | myelin-associated glycoprotein [Source:MGI Symbol;Acc:MGI:96912] | Mag | Enriched in SCN | 3.82 | 2.261E-09 | 357.6 |
| ENSMUSG00000020354 | sarcoglycan, delta (dystrophin-associated glycoprotein) [Source:MGI Symbol;Acc:MGI:1346525] | Sgcd | Enriched in SCN | 3.709 | 3.318E-16 | 245.7 |
| ENSMUSG00000020902 | netrin 1 [Source:MGI Symbol;Acc:MGI:105088] | Ntn1 | Enriched in SCN | 3.68 | 3.603E-19 | 66.38 |
| ENSMUSG00000027360 | histidine decarboxylase [Source:MGI Symbol;Acc:MGI:96062] | Hdc | Enriched in SCN | 3.654 | 1.103E-10 | 44.93 |
| ENSMUSG00000022112 | glypican 5 [Source:MGI Symbol;Acc:MGI:1194894] | Gpc5 | Enriched in SCN | 3.566 | 1.91E-14 | 390.3 |
| ENSMUSG00000037362 | cellular communication network factor 3 [Source:MGI Symbol;Acc:MGI:109185] | Ccn3 | Enriched in SCN | 3.485 | 1.007E-08 | 15.41 |
| ENSMUSG00000041605 | innate immunity activator [Source:MGI Symbol;Acc:MGI:1921579] | Inava | Enriched in SCN | 3.468 | 5.931E-13 | 19.29 |
| ENSMUSG00000050122 | von Willebrand factor A domain containing 3B [Source:MGI Symbol;Acc:MGI:1918103] | Vwa3b | Enriched in SCN | 3.439 | 0.000001306 | 6.63 |
| ENSMUSG00000021499 | cation channel, sperm associated 3 [Source:MGI Symbol;Acc:MGI:1924106] | Catsper3 | Enriched in SCN | 3.36 | 0.00006046 | 4.868 |
| ENSMUSG00000040181 | flavin containing monooxygenase 1 [Source:MGI Symbol;Acc:MGI:1310002] | Fmo1 | Enriched in SCN | 3.337 | 2.777E-11 | 18.97 |
| ENSMUSG00000030592 | ryanodine receptor 1, skeletal muscle [Source:MGI Symbol;Acc:MGI:99659] | Ryr1 | Enriched in SCN | 3.335 | 3.596E-12 | 72.59 |
| ENSMUSG00000038094 | ATPase type 13A4 [Source:MGI Symbol;Acc:MGI:1924456] | Atp13a4 | Enriched in SCN | 3.312 | 4.858E-10 | 39.55 |
| ENSMUSG00000060882 | potassium voltage-gated channel, Shal-related family, member 2 [Source:MGI Symbol;Acc:MGI:102663] | Kcnd2 | Enriched in SCN | 3.297 | 5.665E-16 | 981.4 |
| ENSMUSG00000000794 | potassium intermediate/small conductance calcium-activated channel, subfamily N, member 3 [Source:MGI Symbol;Acc:MGI:2153183] | Kcnn3 | Enriched in SCN | 3.288 | 7.163E-30 | 249.5 |
| ENSMUSG00000044933 | somatostatin receptor 3 [Source:MGI Symbol;Acc:MGI:98329] | Sstr3 | Enriched in SCN | 3.286 | 1.209E-12 | 21.77 |
| ENSMUSG00000048616 | noggin [Source:MGI Symbol;Acc:MGI:104327] | Nog | Enriched in SCN | 3.224 | 7.544E-17 | 53.91 |
| ENSMUSG00000067028 | contactin associated protein-like 5B [Source:MGI Symbol;Acc:MGI:3664583] | Cntnap5b | Enriched in SCN | 3.214 | 2.048E-15 | 198.7 |
| ENSMUSG00000010086 | ring finger protein 112 [Source:MGI Symbol;Acc:MGI:106611] | Rnf112 | Enriched in SCN | 3.204 | 1.142E-25 | 340.8 |
| ENSMUSG00000022696 | SID1 transmembrane family, member 1 [Source:MGI Symbol;Acc:MGI:2443155] | Sidt1 | Enriched in SCN | 3.151 | 1.221E-21 | 43.24 |
| ENSMUSG00000045083 | leucine rich repeat and Ig domain containing 2 [Source:MGI Symbol;Acc:MGI:2442298] | Lingo2 | Enriched in SCN | 3.1 | 4.717E-12 | 769.1 |
| ENSMUSG00000024617 | calcium/calmodulin-dependent protein kinase II alpha [Source:MGI Symbol;Acc:MGI:88256] | Camk2a | Enriched in SCN | 3.03 | 5.88E-25 | 1425 |
| ENSMUSG00000117286 | predicted gene 1043 [Source:MGI Symbol;Acc:MGI:2685889] | Gm1043 | Enriched in SCN | 3.021 | 3.172E-24 | 237.8 |
| ENSMUSG00000035576 | L3MBTL1 histone methyl-lysine binding protein [Source:MGI Symbol;Acc:MGI:2676663] | L3mbtl1 | Enriched in SCN | 3.008 | 6.669E-21 | 85.42 |
| ENSMUSG00000061702 | transmembrane protein 91 [Source:MGI Symbol;Acc:MGI:2443589] | Tmem91 | Enriched in SCN | 2.996 | 4.289E-14 | 41.88 |
| ENSMUSG00000066705 | FXYD domain-containing ion transport regulator 6 [Source:MGI Symbol;Acc:MGI:1890226] | Fxyd6 | Enriched in SCN | 2.983 | 4.432E-13 | 380.3 |
| ENSMUSG00000030385 | RIKEN cDNA 2900092C05 gene [Source:MGI Symbol;Acc:MGI:1920340] | 2900092C05Rik | Enriched in SCN | 2.958 | 0.0004118 | 3.887 |
| ENSMUSG00000108601 | predicted gene 44645 [Source:MGI Symbol;Acc:MGI:5753221] | Gm44645 | Enriched in SCN | 2.956 | 5.494E-15 | 130.8 |
| ENSMUSG00000052026 | solute carrier family 6 (neurotransmitter transporter, L-proline), member 7 [Source:MGI Symbol;Acc:MGI:2147363] | Slc6a7 | Enriched in SCN | 2.953 | 6.858E-38 | 217.9 |
| ENSMUSG00000063646 | janus kinase and microtubule interacting protein 1 [Source:MGI Symbol;Acc:MGI:1923321] | Jakmip1 | Enriched in SCN | 2.916 | 2.303E-10 | 47 |
| ENSMUSG00000010342 | testis expressed gene 14 [Source:MGI Symbol;Acc:MGI:1933227] | Tex14 | Enriched in SCN | 2.908 | 3.682E-07 | 109.9 |
| ENSMUSG00000021596 | multiple C2 domains, transmembrane 1 [Source:MGI Symbol;Acc:MGI:1926021] | Mctp1 | Enriched in SCN | 2.851 | 4.838E-17 | 101.7 |
| ENSMUSG00000052372 | interleukin 1 receptor accessory protein-like 1 [Source:MGI Symbol;Acc:MGI:2687319] | Il1rapl1 | Enriched in SCN | 2.808 | 0.000002893 | 1897 |
| ENSMUSG00000036856 | wingless-type MMTV integration site family, member 4 [Source:MGI Symbol;Acc:MGI:98957] | Wnt4 | Enriched in SCN | 2.749 | 1.131E-16 | 80.14 |
| ENSMUSG00000031026 | tripartite motif-containing 66 [Source:MGI Symbol;Acc:MGI:2152406] | Trim66 | Enriched in SCN | 2.747 | 6.037E-23 | 221.5 |
| ENSMUSG00000030889 | von Willebrand factor A domain containing 3A [Source:MGI Symbol;Acc:MGI:3041229] | Vwa3a | Enriched in SCN | 2.737 | 0.00009168 | 26.65 |
| ENSMUSG00000055078 | gamma-aminobutyric acid (GABA) A receptor, subunit alpha 5 [Source:MGI Symbol;Acc:MGI:95617] | Gabra5 | Enriched in SCN | 2.7 | 2.951E-11 | 60.37 |
| ENSMUSG00000045201 | leucine rich repeat containing 3B [Source:MGI Symbol;Acc:MGI:2384996] | Lrrc3b | Enriched in SCN | 2.672 | 2.827E-08 | 34.56 |
| ENSMUSG00000004263 | atrophin 1 [Source:MGI Symbol;Acc:MGI:104725] | Atn1 | Enriched in SCN | 2.668 | 5.011E-09 | 25.12 |
| ENSMUSG00000025607 | coatomer protein complex, subunit gamma 2 [Source:MGI Symbol;Acc:MGI:1858683] | Copg2 | Enriched in SCN | 2.651 | 1.922E-11 | 319.5 |
| ENSMUSG00000040938 | solute carrier family 16 (monocarboxylic acid transporters), member 11 [Source:MGI Symbol;Acc:MGI:2663709] | Slc16a11 | Enriched in SCN | 2.644 | 0.00001942 | 9.503 |
| ENSMUSG00000070644 | ethanolamine kinase 2 [Source:MGI Symbol;Acc:MGI:2443760] | Etnk2 | Enriched in SCN | 2.642 | 8.085E-16 | 35.13 |
| ENSMUSG00000003469 | phytanoyl-CoA hydroxylase interacting protein [Source:MGI Symbol;Acc:MGI:1860417] | Phyhip | Enriched in SCN | 2.628 | 1.889E-13 | 112 |
| ENSMUSG00000058740 | potassium channel, subfamily T, member 1 [Source:MGI Symbol;Acc:MGI:1924627] | Kcnt1 | Enriched in SCN | 2.605 | 4.053E-17 | 138.9 |
| ENSMUSG00000059203 | interleukin 1 receptor accessory protein-like 2 [Source:MGI Symbol;Acc:MGI:1913106] | Il1rapl2 | Enriched in SCN | 2.574 | 2.228E-08 | 143.9 |
| ENSMUSG00000018451 | RIKEN cDNA 6330403K07 gene [Source:MGI Symbol;Acc:MGI:1918001] | 6330403K07Rik | Enriched in SCN | 2.519 | 7.5E-11 | 664.5 |
| ENSMUSG00000007594 | hyaluronan and proteoglycan link protein 4 [Source:MGI Symbol;Acc:MGI:2679531] | Hapln4 | Enriched in SCN | 2.509 | 0.000002564 | 156.8 |
| ENSMUSG00000055471 | anaplastic lymphoma kinase [Source:MGI Symbol;Acc:MGI:103305] | Alk | Enriched in SCN | 2.5 | 2.613E-11 | 211 |
| ENSMUSG00000061462 | obscurin, cytoskeletal calmodulin and titin-interacting RhoGEF [Source:MGI Symbol;Acc:MGI:2681862] | Obscn | Enriched in SCN | 2.499 | 4.373E-07 | 66.02 |
| ENSMUSG00000036198 | Rho GTPase activating protein 36 [Source:MGI Symbol;Acc:MGI:1922654] | Arhgap36 | Enriched in SCN | 2.493 | 0.001315 | 77.62 |
| ENSMUSG00000028681 | patched 2 [Source:MGI Symbol;Acc:MGI:1095405] | Ptch2 | Enriched in SCN | 2.483 | 0.00008513 | 9.607 |
| ENSMUSG00000046287 | paraneoplastic antigen MA3 [Source:MGI Symbol;Acc:MGI:2180565] | Pnma3 | Enriched in SCN | 2.477 | 1.317E-16 | 110.9 |
| ENSMUSG00000005045 | chromodomain helicase DNA binding protein 5 [Source:MGI Symbol;Acc:MGI:3036258] | Chd5 | Enriched in SCN | 2.451 | 1.045E-28 | 1862 |
| ENSMUSG00000019590 | cytochrome b-561 [Source:MGI Symbol;Acc:MGI:103253] | Cyb561 | Enriched in SCN | 2.433 | 1.628E-15 | 123.2 |
| ENSMUSG00000051331 | calcium channel, voltage-dependent, L type, alpha 1C subunit [Source:MGI Symbol;Acc:MGI:103013] | Cacna1c | Enriched in SCN | 2.433 | 2.398E-14 | 1223 |
| ENSMUSG00000034156 | TSPO associated protein 1 [Source:MGI Symbol;Acc:MGI:2450877] | Tspoap1 | Enriched in SCN | 2.432 | 2.046E-23 | 1203 |
| ENSMUSG00000028758 | kinesin family member 17 [Source:MGI Symbol;Acc:MGI:1098229] | Kif17 | Enriched in SCN | 2.424 | 8.379E-21 | 102.3 |
| ENSMUSG00000039278 | proprotein convertase subtilisin/kexin type 1 inhibitor [Source:MGI Symbol;Acc:MGI:1353431] | Pcsk1n | Enriched in SCN | 2.424 | 3.873E-15 | 2810 |
| ENSMUSG00000042425 | FERM and PDZ domain containing 3 [Source:MGI Symbol;Acc:MGI:3646547] | Frmpd3 | Enriched in SCN | 2.416 | 7.979E-08 | 191.4 |
| ENSMUSG00000052428 | transmembrane and coiled-coil domains 1 [Source:MGI Symbol;Acc:MGI:1921173] | Tmco1 | Enriched in SCN | 2.4 | 0.00001105 | 95.8 |
| ENSMUSG00000024937 | EH domain binding protein 1-like 1 [Source:MGI Symbol;Acc:MGI:3612340] | Ehbp1l1 | Enriched in SCN | 2.398 | 7.253E-16 | 192.9 |
| ENSMUSG00000032297 | CUGBP, Elav-like family member 6 [Source:MGI Symbol;Acc:MGI:1923433] | Celf6 | Enriched in SCN | 2.389 | 3.021E-13 | 185.4 |
| ENSMUSG00000022208 | junctophilin 4 [Source:MGI Symbol;Acc:MGI:2443113] | Jph4 | Enriched in SCN | 2.384 | 2.439E-22 | 961.5 |
| ENSMUSG00000019785 | clavesin 2 [Source:MGI Symbol;Acc:MGI:2443223] | Clvs2 | Enriched in SCN | 2.373 | 1.868E-16 | 86.26 |
| ENSMUSG00000054013 | transmembrane protein 179 [Source:MGI Symbol;Acc:MGI:2144891] | Tmem179 | Enriched in SCN | 2.37 | 5.064E-18 | 349.1 |
| ENSMUSG00000024109 | neurexin I [Source:MGI Symbol;Acc:MGI:1096391] | Nrxn1 | Enriched in SCN | 2.353 | 4.433E-15 | 3720 |
| ENSMUSG00000096141 | dynein, axonemal, heavy chain 7A [Source:MGI Symbol;Acc:MGI:2685838] | Dnah7a | Enriched in SCN | 2.353 | 0.000001065 | 25.06 |
| ENSMUSG00000049252 | low density lipoprotein-related protein 1B [Source:MGI Symbol;Acc:MGI:2151136] | Lrp1b | Enriched in SCN | 2.328 | 6.21E-12 | 1165 |
| ENSMUSG00000055692 | transmembrane protein 191 [Source:MGI Symbol;Acc:MGI:107238] | Tmem191 | Enriched in SCN | 2.319 | 7.263E-13 | 172.4 |
| ENSMUSG00000038242 | aldehyde oxidase 4 [Source:MGI Symbol;Acc:MGI:1919122] | Aox4 | Enriched in SCN | 2.318 | 0.005136 | 8.603 |
| ENSMUSG00000021541 | transient receptor potential cation channel, subfamily C, member 7 [Source:MGI Symbol;Acc:MGI:1349470] | Trpc7 | Enriched in SCN | 2.314 | 9.847E-09 | 62.32 |
| ENSMUSG00000043419 | ring finger protein 227 [Source:MGI Symbol;Acc:MGI:1915359] | Rnf227 | Enriched in SCN | 2.31 | 1.131E-16 | 254.4 |
| ENSMUSG00000044533 | ribosomal protein S2 [Source:MGI Symbol;Acc:MGI:105110] | Rps2 | Enriched in SCN | 2.305 | 9.058E-08 | 542 |
| ENSMUSG00000074652 | myosin, heavy chain 7B, cardiac muscle, beta [Source:MGI Symbol;Acc:MGI:3710243] | Myh7b | Enriched in SCN | 2.302 | 4.609E-11 | 65.13 |
| ENSMUSG00000062044 | lemur tyrosine kinase 3 [Source:MGI Symbol;Acc:MGI:3039582] | Lmtk3 | Enriched in SCN | 2.301 | 1.814E-20 | 2447 |
| ENSMUSG00000046491 | C1q and tumor necrosis factor related protein 2 [Source:MGI Symbol;Acc:MGI:1916433] | C1qtnf2 | Enriched in SCN | 2.286 | 0.005936 | 4.439 |
| ENSMUSG00000050821 | family with sequence similarity 131, member A [Source:MGI Symbol;Acc:MGI:1925658] | Fam131a | Enriched in SCN | 2.277 | 6.535E-19 | 179.2 |
| ENSMUSG00000031841 | cadherin 13 [Source:MGI Symbol;Acc:MGI:99551] | Cdh13 | Enriched in SCN | 2.265 | 1.913E-09 | 448.3 |
| ENSMUSG00000020142 | solute carrier family 1 (glutamate/neutral amino acid transporter), member 4 [Source:MGI Symbol;Acc:MGI:2135601] | Slc1a4 | Enriched in SCN | 2.239 | 1.32E-19 | 230.7 |
| ENSMUSG00000048537 | pleckstrin homology like domain, family B, member 1 [Source:MGI Symbol;Acc:MGI:2143230] | Phldb1 | Enriched in SCN | 2.236 | 5.535E-24 | 593 |
| ENSMUSG00000034226 | ras homolog family member V [Source:MGI Symbol;Acc:MGI:2444227] | Rhov | Enriched in SCN | 2.22 | 1.668E-11 | 54.65 |
| ENSMUSG00000029602 | RAS protein activator like 1 (GAP1 like) [Source:MGI Symbol;Acc:MGI:1330842] | Rasal1 | Enriched in SCN | 2.218 | 2.205E-10 | 51.4 |
| ENSMUSG00000102692 | dachsous cadherin related 2 [Source:MGI Symbol;Acc:MGI:2685263] | Dchs2 | Enriched in SCN | 2.217 | 0.00002415 | 117.9 |
| ENSMUSG00000058019 | carboxylesterase 5A [Source:MGI Symbol;Acc:MGI:1915185] | Ces5a | Enriched in SCN | 2.213 | 0.02068 | 3.783 |
| ENSMUSG00000034282 | envoplakin [Source:MGI Symbol;Acc:MGI:107507] | Evpl | Enriched in SCN | 2.201 | 1.025E-13 | 114.4 |
| ENSMUSG00000021852 | solute carrier family 35, member F4 [Source:MGI Symbol;Acc:MGI:1922538] | Slc35f4 | Enriched in SCN | 2.2 | 0.000003626 | 56.44 |
| ENSMUSG00000042216 | small G protein signaling modulator 1 [Source:MGI Symbol;Acc:MGI:107320] | Sgsm1 | Enriched in SCN | 2.19 | 2.005E-21 | 412.2 |
| ENSMUSG00000006711 | RIKEN cDNA D130043K22 gene [Source:MGI Symbol;Acc:MGI:3036268] | D130043K22Rik | Enriched in SCN | 2.187 | 8.909E-11 | 133.1 |
| ENSMUSG00000091002 | transcription elongation regulator 1-like [Source:MGI Symbol;Acc:MGI:1917821] | Tcerg1l | Enriched in SCN | 2.185 | 1.253E-10 | 243.7 |
| ENSMUSG00000053395 | calcium channel, voltage-dependent, gamma subunit 8 [Source:MGI Symbol;Acc:MGI:1932376] | Cacng8 | Enriched in SCN | 2.177 | 3.812E-12 | 79.33 |
| ENSMUSG00000034664 | integrin alpha 2b [Source:MGI Symbol;Acc:MGI:96601] | Itga2b | Enriched in SCN | 2.171 | 0.000002287 | 10.87 |
| ENSMUSG00000105867 | predicted gene 42517 [Source:MGI Symbol;Acc:MGI:5662654] | Gm42517 | Enriched in SCN | 2.161 | 4.972E-14 | 89.61 |
| ENSMUSG00000025422 | ArfGAP with GTPase domain, ankyrin repeat and PH domain 2 [Source:MGI Symbol;Acc:MGI:3580016] | Agap2 | Enriched in SCN | 2.16 | 2.63E-30 | 2518 |
| ENSMUSG00000040752 | myosin, heavy polypeptide 6, cardiac muscle, alpha [Source:MGI Symbol;Acc:MGI:97255] | Myh6 | Enriched in SCN | 2.158 | 0.0007065 | 9.823 |
| ENSMUSG00000038884 | shiftless antiviral inhibitor of ribosomal frameshifting [Source:MGI Symbol;Acc:MGI:2441788] | Shfl | Enriched in SCN | 2.154 | 2.741E-16 | 133.8 |
| ENSMUSG00000037428 | VGF nerve growth factor inducible [Source:MGI Symbol;Acc:MGI:1343180] | Vgf | Enriched in SCN | 2.152 | 8.901E-14 | 1249 |
| ENSMUSG00000029516 | citron [Source:MGI Symbol;Acc:MGI:105313] | Cit | Enriched in SCN | 2.143 | 1.674E-20 | 2284 |
| ENSMUSG00000094370 | predicted gene 3373 [Source:MGI Symbol;Acc:MGI:3781551] | Gm3373 | Enriched in SCN | 2.129 | 0.004375 | 3.809 |
| ENSMUSG00000069911 | inhibitory synaptic factor family member 2B [Source:MGI Symbol;Acc:MGI:3643491] | Insyn2b | Enriched in SCN | 2.128 | 3.304E-07 | 58.95 |
| ENSMUSG00000056856 | janus kinase and microtubule interacting protein 3 [Source:MGI Symbol;Acc:MGI:1921254] | Jakmip3 | Enriched in SCN | 2.115 | 9.611E-20 | 256.3 |
| ENSMUSG00000050541 | adrenergic receptor, alpha 1b [Source:MGI Symbol;Acc:MGI:104774] | Adra1b | Enriched in SCN | 2.106 | 7.901E-08 | 73.77 |
| ENSMUSG00000042846 | leucine rich repeat transmembrane neuronal 3 [Source:MGI Symbol;Acc:MGI:2389177] | Lrrtm3 | Enriched in SCN | 2.103 | 7.828E-08 | 135.3 |
| ENSMUSG00000009394 | synapsin II [Source:MGI Symbol;Acc:MGI:103020] | Syn2 | Enriched in SCN | 2.094 | 7.358E-14 | 611.4 |
| ENSMUSG00000020704 | acid-sensing (proton-gated) ion channel 2 [Source:MGI Symbol;Acc:MGI:1100867] | Asic2 | Enriched in SCN | 2.093 | 3.049E-07 | 975.3 |
| ENSMUSG00000066129 | kinase non-catalytic C-lobe domain (KIND) containing 1 [Source:MGI Symbol;Acc:MGI:1923734] | Kndc1 | Enriched in SCN | 2.077 | 2.408E-17 | 714.5 |
| ENSMUSG00000036923 | storkhead box 1 [Source:MGI Symbol;Acc:MGI:2684909] | Stox1 | Enriched in SCN | 2.074 | 9.516E-09 | 18.87 |
| ENSMUSG00000025468 | calcyon neuron-specific vesicular protein [Source:MGI Symbol;Acc:MGI:1915816] | Caly | Enriched in SCN | 2.062 | 8.716E-11 | 238.8 |
| ENSMUSG00000050677 | coiled-coil domain containing 96 [Source:MGI Symbol;Acc:MGI:1913967] | Ccdc96 | Enriched in SCN | 2.046 | 0.0003479 | 66.05 |
| ENSMUSG00000057897 | calcium/calmodulin-dependent protein kinase II, beta [Source:MGI Symbol;Acc:MGI:88257] | Camk2b | Enriched in SCN | 2.04 | 1.189E-28 | 2031 |
| ENSMUSG00000008496 | POU domain, class 2, transcription factor 2 [Source:MGI Symbol;Acc:MGI:101897] | Pou2f2 | Enriched in SCN | 2.038 | 1.223E-10 | 295.3 |
| ENSMUSG00000018849 | WW, C2 and coiled-coil domain containing 1 [Source:MGI Symbol;Acc:MGI:2388637] | Wwc1 | Enriched in SCN | 2.032 | 3.045E-27 | 625.4 |
| ENSMUSG00000031860 | pre B cell leukemia homeobox 4 [Source:MGI Symbol;Acc:MGI:1931321] | Pbx4 | Enriched in SCN | 2.022 | 3.544E-11 | 37.95 |
| ENSMUSG00000036815 | dipeptidylpeptidase 10 [Source:MGI Symbol;Acc:MGI:2442409] | Dpp10 | Enriched in SCN | 2.016 | 1.828E-07 | 576.4 |
| ENSMUSG00000117694 | small nucleolar RNA host gene 4 [Source:MGI Symbol;Acc:MGI:4937091] | Snhg4 | Enriched in SCN | 2.016 | 0.000000808 | 124.9 |
| ENSMUSG00000031760 | metallothionein 3 [Source:MGI Symbol;Acc:MGI:97173] | Mt3 | Enriched in SCN | 2.014 | 2.686E-08 | 131.4 |
| ENSMUSG00000044576 | GRB2 associated regulator of MAPK1 subtype 2 [Source:MGI Symbol;Acc:MGI:2685290] | Garem2 | Enriched in SCN | 2.008 | 2.917E-12 | 62.67 |

| | | | | | | |
|---|---|---|---|---|---|---|
| ENSMUSG00000068373 | RIKEN cDNA D430041D05 gene [Source:MGI Symbol;Acc:MGI:2181743] | D430041D05Rik | Enriched in SCN | 2.006 | 1.755E-26 | 780.2 |
| ENSMUSG00000026173 | phospholipase C, delta 4 [Source:MGI Symbol;Acc:MGI:107469] | Plcd4 | Enriched in SCN | 2.006 | 0.000008762 | 71.97 |
| ENSMUSG00000043165 | loricrin [Source:MGI Symbol;Acc:MGI:96816] | Lor | Enriched in SCN | 1.997 | 0.0008478 | 17.17 |
| ENSMUSG00000038860 | GTPase activating RANGAP domain-like 3 [Source:MGI Symbol;Acc:MGI:2139309] | Garnl3 | Enriched in SCN | 1.99 | 1.361E-11 | 238.5 |
| ENSMUSG00000033022 | cysteine dioxygenase 1, cytosolic [Source:MGI Symbol;Acc:MGI:105925] | Cdo1 | Enriched in SCN | 1.981 | 3.415E-08 | 55.15 |
| ENSMUSG00000025856 | platelet derived growth factor, alpha [Source:MGI Symbol;Acc:MGI:97527] | Pdgfa | Enriched in SCN | 1.98 | 5.267E-18 | 235.2 |
| ENSMUSG00000047155 | cytochrome P450, family 4, subfamily x, polypeptide 1 [Source:MGI Symbol;Acc:MGI:1932403] | Cyp4x1 | Enriched in SCN | 1.979 | 5.508E-10 | 55.68 |
| ENSMUSG00000116652 | RIKEN cDNA B830017H08 gene [Source:MGI Symbol;Acc:MGI:3045365] | B830017H08Rik | Enriched in SCN | 1.97 | 0.00008546 | 15.95 |
| ENSMUSG00000034353 | receptor (calcitonin) activity modifying protein 1 [Source:MGI Symbol;Acc:MGI:1858418] | Ramp1 | Enriched in SCN | 1.969 | 0.000005526 | 54.71 |
| ENSMUSG00000074923 | p21 (RAC1) activated kinase 6 [Source:MGI Symbol;Acc:MGI:2679420] | Pak6 | Enriched in SCN | 1.965 | 4.684E-15 | 281.4 |
| ENSMUSG00000063296 | transmembrane protein 117 [Source:MGI Symbol;Acc:MGI:2444580] | Tmem117 | Enriched in SCN | 1.964 | 8.536E-07 | 103 |
| ENSMUSG00000031727 | polyamine modulated factor 1 binding protein 1 [Source:MGI Symbol;Acc:MGI:1930136] | Pmfbp1 | Enriched in SCN | 1.959 | 0.0005304 | 36.63 |
| ENSMUSG00000061578 | kinase suppressor of ras 2 [Source:MGI Symbol;Acc:MGI:3610315] | Ksr2 | Enriched in SCN | 1.954 | 7.933E-19 | 833.8 |
| ENSMUSG00000049112 | oxytocin receptor [Source:MGI Symbol;Acc:MGI:109147] | Oxtr | Enriched in SCN | 1.951 | 4.416E-07 | 27.31 |
| ENSMUSG00000020524 | glutamate receptor, ionotropic, AMPA1 (alpha 1) [Source:MGI Symbol;Acc:MGI:95808] | Gria1 | Enriched in SCN | 1.948 | 2.567E-18 | 870.1 |
| ENSMUSG00000030844 | regulator of G-protein signalling 10 [Source:MGI Symbol;Acc:MGI:1915115] | Rgs10 | Enriched in SCN | 1.94 | 0.000001487 | 70.22 |
| ENSMUSG00000022577 | lymphocyte antigen 6 complex, locus H [Source:MGI Symbol;Acc:MGI:1346030] | Ly6h | Enriched in SCN | 1.937 | 2.49E-09 | 411.3 |
| ENSMUSG00000031659 | adenylate cyclase 7 [Source:MGI Symbol;Acc:MGI:102891] | Adcy7 | Enriched in SCN | 1.925 | 9.197E-13 | 65.54 |
| ENSMUSG00000063446 | phospholipid phosphatase related 1 [Source:MGI Symbol;Acc:MGI:2445015] | Plppr1 | Enriched in SCN | 1.919 | 0.00007558 | 101.6 |
| ENSMUSG00000037703 | leucine zipper, putative tumor suppressor family member 3 [Source:MGI Symbol;Acc:MGI:2656976] | Lzts3 | Enriched in SCN | 1.915 | 4.671E-16 | 572.4 |
| ENSMUSG00000004187 | kinesin family member C2 [Source:MGI Symbol;Acc:MGI:109187] | Kifc2 | Enriched in SCN | 1.905 | 5.126E-26 | 769 |
| ENSMUSG00000046378 | aspartate beta-hydroxylase domain containing 1 [Source:MGI Symbol;Acc:MGI:2685014] | Asphd1 | Enriched in SCN | 1.897 | 8.901E-09 | 56.65 |
| ENSMUSG00000021448 | src homology 2 domain-containing transforming protein C3 [Source:MGI Symbol;Acc:MGI:106179] | Shc3 | Enriched in SCN | 1.889 | 6.222E-19 | 933.3 |
| ENSMUSG00000046793 | G protein-coupled receptor 61 [Source:MGI Symbol;Acc:MGI:2441719] | Gpr61 | Enriched in SCN | 1.886 | 9.24E-09 | 53.44 |
| ENSMUSG00000029095 | actin-binding LIM protein 2 [Source:MGI Symbol;Acc:MGI:2385758] | Ablim2 | Enriched in SCN | 1.881 | 2.562E-14 | 268.3 |
| ENSMUSG00000043388 | transmembrane protein 130 [Source:MGI Symbol;Acc:MGI:3607706] | Tmem130 | Enriched in SCN | 1.87 | 1.682E-08 | 1062 |
| ENSMUSG00000074673 | tubulin tyrosine ligase-like family, member 9 [Source:MGI Symbol;Acc:MGI:1913589] | Ttll9 | Enriched in SCN | 1.869 | 0.002407 | 6.47 |
| ENSMUSG00000028865 | CD164 sialomucin-like 2 [Source:MGI Symbol;Acc:MGI:1916905] | Cd164l2 | Enriched in SCN | 1.865 | 0.01769 | 5.202 |
| ENSMUSG00000032502 | src homology three (SH3) and cysteine rich domain [Source:MGI Symbol;Acc:MGI:1201400] | Stac | Enriched in SCN | 1.857 | 0.001484 | 39.11 |
| ENSMUSG00000060275 | neuregulin 2 [Source:MGI Symbol;Acc:MGI:1098246] | Nrg2 | Enriched in SCN | 1.854 | 0.000004537 | 123.8 |
| ENSMUSG00000054863 | TAFA chemokine like family member 5 [Source:MGI Symbol;Acc:MGI:2146182] | Tafa5 | Enriched in SCN | 1.85 | 3.608E-11 | 499.7 |
| ENSMUSG00000018339 | glutathione peroxidase 3 [Source:MGI Symbol;Acc:MGI:105102] | Gpx3 | Enriched in SCN | 1.847 | 0.003412 | 82.52 |
| ENSMUSG00000057716 | transmembrane protein 178B [Source:MGI Symbol;Acc:MGI:3647581] | Tmem178b | Enriched in SCN | 1.846 | 1.625E-25 | 1516 |
| ENSMUSG00000056724 | neurobeachin-like 2 [Source:MGI Symbol;Acc:MGI:2448554] | Nbeal2 | Enriched in SCN | 1.842 | 4.125E-10 | 63.07 |
| ENSMUSG00000020374 | RasGEF domain family, member 1C [Source:MGI Symbol;Acc:MGI:1921813] | Rasgef1c | Enriched in SCN | 1.833 | 1.047E-09 | 39.33 |
| ENSMUSG00000038453 | SRC kinase signaling inhibitor 1 [Source:MGI Symbol;Acc:MGI:1933179] | Srcin1 | Enriched in SCN | 1.827 | 4.479E-24 | 3243 |
| ENSMUSG00000027584 | opioid receptor-like 1 [Source:MGI Symbol;Acc:MGI:97440] | Oprl1 | Enriched in SCN | 1.82 | 1.454E-16 | 154 |
| ENSMUSG00000025790 | solute carrier organic anion transporter family, member 3a1 [Source:MGI Symbol;Acc:MGI:1351867] | Slco3a1 | Enriched in SCN | 1.818 | 1.191E-07 | 185.2 |
| ENSMUSG00000020090 | neuropeptide FF receptor 1 [Source:MGI Symbol;Acc:MGI:2685082] | Npffr1 | Enriched in SCN | 1.818 | 0.001261 | 15.66 |
| ENSMUSG00000033061 | regulated endocrine-specific protein 18 [Source:MGI Symbol;Acc:MGI:1098222] | Resp18 | Enriched in SCN | 1.809 | 0.001663 | 74.08 |
| ENSMUSG00000102189 | predicted gene, 37194 [Source:MGI Symbol;Acc:MGI:5610422] | Gm37194 | Enriched in SCN | 1.802 | 0.002178 | 32.7 |
| ENSMUSG00000032503 | cyclic AMP-regulated phosphoprotein, 21 [Source:MGI Symbol;Acc:MGI:107562] | Arpp21 | Enriched in SCN | 1.792 | 0.000007943 | 594.6 |
| ENSMUSG00000028341 | nuclear receptor subfamily 4, group A, member 3 [Source:MGI Symbol;Acc:MGI:1352457] | Nr4a3 | Enriched in SCN | 1.79 | 0.00002025 | 47.01 |
| ENSMUSG00000029245 | Eph receptor A5 [Source:MGI Symbol;Acc:MGI:99654] | Epha5 | Enriched in SCN | 1.785 | 8.607E-07 | 636.4 |
| ENSMUSG00000052477 | RIKEN cDNA C130026I21 gene [Source:MGI Symbol;Acc:MGI:3612702] | C130026I21Rik | Enriched in SCN | 1.785 | 0.01154 | 6.977 |
| ENSMUSG00000047428 | delta like non-canonical Notch ligand 2 [Source:MGI Symbol;Acc:MGI:2146838] | Dlk2 | Enriched in SCN | 1.782 | 4.858E-10 | 45.68 |
| ENSMUSG00000059146 | neurotrophic tyrosine kinase, receptor, type 3 [Source:MGI Symbol;Acc:MGI:97385] | Ntrk3 | Enriched in SCN | 1.781 | 5.473E-07 | 795.7 |
| ENSMUSG00000029419 | apical junction component 1 [Source:MGI Symbol;Acc:MGI:2685842] | Ajm1 | Enriched in SCN | 1.776 | 2.932E-11 | 326.7 |
| ENSMUSG00000031700 | glutamic pyruvate transaminase (alanine aminotransferase) 2 [Source:MGI Symbol;Acc:MGI:1915391] | Gpt2 | Enriched in SCN | 1.772 | 3.644E-16 | 138.5 |
| ENSMUSG00000031963 | BMP-binding endothelial regulator [Source:MGI Symbol;Acc:MGI:1920480] | Bmper | Enriched in SCN | 1.765 | 0.000003921 | 44.47 |
| ENSMUSG00000032017 | glutamate receptor, ionotropic, kainate 4 [Source:MGI Symbol;Acc:MGI:95817] | Grik4 | Enriched in SCN | 1.758 | 2.762E-07 | 185.2 |
| ENSMUSG00000037348 | progestin and adipoQ receptor family member VII [Source:MGI Symbol;Acc:MGI:1919154] | Paqr7 | Enriched in SCN | 1.747 | 1.275E-10 | 163 |
| ENSMUSG00000030098 | glutamate receptor interacting protein 2 [Source:MGI Symbol;Acc:MGI:2681173] | Grip2 | Enriched in SCN | 1.746 | 1.011E-17 | 119.7 |
| ENSMUSG00000058153 | seizure related 6 homolog like [Source:MGI Symbol;Acc:MGI:1935121] | Sez6l | Enriched in SCN | 1.737 | 7.428E-13 | 1179 |
| ENSMUSG00000034145 | transmembrane protein 63c [Source:MGI Symbol;Acc:MGI:2444386] | Tmem63c | Enriched in SCN | 1.73 | 2.252E-11 | 106.4 |
| ENSMUSG00000041078 | glutamate receptor, ionotropic, delta 1 [Source:MGI Symbol;Acc:MGI:95812] | Grid1 | Enriched in SCN | 1.725 | 0.00000228 | 524.9 |
| ENSMUSG00000021278 | amnionless [Source:MGI Symbol;Acc:MGI:1934943] | Amn | Enriched in SCN | 1.72 | 0.00004966 | 16.05 |
| ENSMUSG00000032036 | kirre like nephrin family adhesion molecule 3 [Source:MGI Symbol;Acc:MGI:1914953] | Kirrel3 | Enriched in SCN | 1.709 | 0.001217 | 576.5 |
| ENSMUSG00000030276 | tubulin tyrosine ligase-like family, member 3 [Source:MGI Symbol;Acc:MGI:2141418] | Ttll3 | Enriched in SCN | 1.708 | 0.000809 | 73.6 |
| ENSMUSG00000042249 | G protein-coupled receptor kinase 3 [Source:MGI Symbol;Acc:MGI:87941] | Grk3 | Enriched in SCN | 1.695 | 3.321E-15 | 419.2 |
| ENSMUSG00000027200 | sema domain, transmembrane domain (TM), and cytoplasmic domain, (semaphorin) 6D [Source:MGI Symbol;Acc:MGI:2387661] | Sema6d | Enriched in SCN | 1.69 | 1.548E-08 | 731.9 |
| ENSMUSG00000058420 | synaptotagmin XVII [Source:MGI Symbol;Acc:MGI:104966] | Syt17 | Enriched in SCN | 1.685 | 0.000001794 | 49.03 |
| undefined | undefined | undefined | Enriched in SCN | 1.685 | 0.000002601 | 71.4 |
| ENSMUSG00000022012 | ecto-NOX disulfide-thiol exchanger 1 [Source:MGI Symbol;Acc:MGI:2444896] | Enox1 | Enriched in SCN | 1.68 | 0.0002199 | 297.2 |
| ENSMUSG00000028909 | protein tyrosine phosphatase, receptor type, U [Source:MGI Symbol;Acc:MGI:1321151] | Ptpru | Enriched in SCN | 1.678 | 2.825E-14 | 212.6 |
| ENSMUSG00000004933 | megakaryocyte-associated tyrosine kinase [Source:MGI Symbol;Acc:MGI:99259] | Matk | Enriched in SCN | 1.675 | 9.851E-12 | 175.5 |
| ENSMUSG00000066191 | ankyrin repeat and sterile alpha motif domain containing 6 [Source:MGI Symbol;Acc:MGI:1922941] | Anks6 | Enriched in SCN | 1.673 | 1.415E-07 | 55.31 |
| ENSMUSG00000034993 | vesicle amine transport 1 [Source:MGI Symbol;Acc:MGI:1349450] | Vat1 | Enriched in SCN | 1.67 | 6.435E-09 | 429.8 |
| ENSMUSG00000042873 | lipoma HMGIC fusion partner-like protein 4 [Source:MGI Symbol;Acc:MGI:3057108] | Lhfpl4 | Enriched in SCN | 1.666 | 2.414E-11 | 1362 |
| ENSMUSG00000036377 | capping protein inhibiting regulator of actin [Source:MGI Symbol;Acc:MGI:2444817] | Cracd | Enriched in SCN | 1.656 | 5.826E-11 | 1779 |
| ENSMUSG00000029442 | WD repeat domain 66 [Source:MGI Symbol;Acc:MGI:1918495] | Wdr66 | Enriched in SCN | 1.649 | 0.00001014 | 54.47 |
| ENSMUSG00000074785 | plexin C1 [Source:MGI Symbol;Acc:MGI:1890127] | Plxnc1 | Enriched in SCN | 1.646 | 1.791E-12 | 283.7 |
| ENSMUSG00000055409 | NEL-like 1 [Source:MGI Symbol;Acc:MGI:2443902] | Nell1 | Enriched in SCN | 1.642 | 0.00009837 | 543.3 |
| ENSMUSG00000028197 | collagen, type XXIV, alpha 1 [Source:MGI Symbol;Acc:MGI:1918605] | Col24a1 | Enriched in SCN | 1.64 | 0.01927 | 14.18 |
| ENSMUSG00000032890 | regulating synaptic membrane exocytosis 3 [Source:MGI Symbol;Acc:MGI:2443331] | Rims3 | Enriched in SCN | 1.636 | 4.74E-10 | 1313 |
| ENSMUSG00000025576 | RNA binding protein, fox-1 homolog (C. elegans) 3 [Source:MGI Symbol;Acc:MGI:106368] | Rbfox3 | Enriched in SCN | 1.635 | 5.919E-12 | 388.1 |
| ENSMUSG00000039546 | adherens junction associated protein 1 [Source:MGI Symbol;Acc:MGI:2685419] | Ajap1 | Enriched in SCN | 1.635 | 4.747E-10 | 293.4 |
| ENSMUSG00000023868 | phosphodiesterase 10A [Source:MGI Symbol;Acc:MGI:1345143] | Pde10a | Enriched in SCN | 1.617 | 0.00000039 | 520.6 |
| ENSMUSG00000035064 | eukaryotic elongation factor-2 kinase [Source:MGI Symbol;Acc:MGI:1195261] | Eef2k | Enriched in SCN | 1.611 | 4.833E-14 | 118.4 |
| ENSMUSG00000022840 | adenylate cyclase 5 [Source:MGI Symbol;Acc:MGI:99673] | Adcy5 | Enriched in SCN | 1.61 | 6.435E-10 | 2027 |
| ENSMUSG00000050069 | gremlin 2, DAN family BMP antagonist [Source:MGI Symbol;Acc:MGI:1344367] | Grem2 | Enriched in SCN | 1.609 | 6.495E-07 | 70.97 |
| ENSMUSG00000051335 | glucose-fructose oxidoreductase domain containing 1 [Source:MGI Symbol;Acc:MGI:2145304] | Gfod1 | Enriched in SCN | 1.604 | 8.919E-18 | 253.7 |
| ENSMUSG00000027221 | carbohydrate sulfotransferase 1 [Source:MGI Symbol;Acc:MGI:1924219] | Chst1 | Enriched in SCN | 1.601 | 2.503E-11 | 247 |
| ENSMUSG00000042766 | tripartite motif-containing 46 [Source:MGI Symbol;Acc:MGI:2673000] | Trim46 | Enriched in SCN | 1.597 | 8.186E-17 | 200.1 |
| ENSMUSG00000003872 | lin-7 homolog B, crumbs cell polarity complex component [Source:MGI Symbol;Acc:MGI:1330858] | Lin7b | Enriched in SCN | 1.594 | 0.00000934 | 29.61 |
| ENSMUSG00000018509 | centromere protein V [Source:MGI Symbol;Acc:MGI:1920389] | Cenpv | Enriched in SCN | 1.592 | 0.00002505 | 40.11 |
| ENSMUSG00000020990 | cyclin-dependent kinase-like 1 (CDC2-related kinase) [Source:MGI Symbol;Acc:MGI:1918341] | Cdkl1 | Enriched in SCN | 1.585 | 0.00002492 | 53.67 |
| ENSMUSG00000055540 | Eph receptor A6 [Source:MGI Symbol;Acc:MGI:108034] | Epha6 | Enriched in SCN | 1.581 | 0.0001269 | 266.7 |
| ENSMUSG00000053166 | cadherin 22 [Source:MGI Symbol;Acc:MGI:1341843] | Cdh22 | Enriched in SCN | 1.58 | 0.000008257 | 180 |
| ENSMUSG00000025318 | junctophilin 3 [Source:MGI Symbol;Acc:MGI:1891497] | Jph3 | Enriched in SCN | 1.571 | 1.513E-13 | 1064 |
| ENSMUSG00000054027 | 5'-nucleotidase domain containing 3 [Source:MGI Symbol;Acc:MGI:3513266] | Nt5dc3 | Enriched in SCN | 1.569 | 3.102E-08 | 395 |
| ENSMUSG00000061911 | myelin transcription factor 1-like [Source:MGI Symbol;Acc:MGI:1100511] | Myt1l | Enriched in SCN | 1.567 | 5.771E-12 | 1383 |
| ENSMUSG00000040606 | kazrin, periplakin interacting protein [Source:MGI Symbol;Acc:MGI:1918779] | Kazn | Enriched in SCN | 1.566 | 2.237E-08 | 697.4 |
| ENSMUSG00000034730 | adhesion G protein-coupled receptor B1 [Source:MGI Symbol;Acc:MGI:1933736] | Adgrb1 | Enriched in SCN | 1.565 | 2.039E-20 | 1290 |
| ENSMUSG00000078591 | heparan sulfate (glucosamine) 3-O-sulfotransferase 4 [Source:MGI Symbol;Acc:MGI:1333792] | Hs3st4 | Enriched in SCN | 1.564 | 0.001388 | 324.6 |
| ENSMUSG00000033467 | cytokine receptor-like factor 2 [Source:MGI Symbol;Acc:MGI:1889506] | Crlf2 | Enriched in SCN | 1.563 | 8.301E-08 | 87.19 |
| ENSMUSG00000047415 | G protein-coupled receptor 68 [Source:MGI Symbol;Acc:MGI:2441763] | Gpr68 | Enriched in SCN | 1.563 | 0.00000242 | 34.85 |
| ENSMUSG00000034037 | FYVE, RhoGEF and PH domain containing 5 [Source:MGI Symbol;Acc:MGI:2443369] | Fgd5 | Enriched in SCN | 1.561 | 0.00001613 | 83.07 |
| ENSMUSG00000087408 | ceramide synthase 1 [Source:MGI Symbol;Acc:MGI:2136690] | Cers1 | Enriched in SCN | 1.559 | 2.608E-15 | 94.13 |
| ENSMUSG00000039579 | glutamate receptor ionotropic, NMDA3A [Source:MGI Symbol;Acc:MGI:1933206] | Grin3a | Enriched in SCN | 1.556 | 3.929E-08 | 314.9 |
| ENSMUSG00000022861 | diacylglycerol kinase, gamma [Source:MGI Symbol;Acc:MGI:105060] | Dgkg | Enriched in SCN | 1.556 | 0.0001543 | 135 |
| ENSMUSG00000031351 | zinc finger protein 185 [Source:MGI Symbol;Acc:MGI:108095] | Zfp185 | Enriched in SCN | 1.549 | 0.0004513 | 546.2 |
| ENSMUSG00000020175 | RAB36, member RAS oncogene family [Source:MGI Symbol;Acc:MGI:1924127] | Rab36 | Enriched in SCN | 1.548 | 4.554E-07 | 78.31 |
| ENSMUSG00000048142 | N-acetyltransferase 8-like [Source:MGI Symbol;Acc:MGI:2447776] | Nat8l | Enriched in SCN | 1.54 | 4.901E-08 | 599.1 |
| ENSMUSG00000030854 | protein tyrosine phosphatase, non-receptor type 5 [Source:MGI Symbol;Acc:MGI:97807] | Ptpn5 | Enriched in SCN | 1.537 | 2.87E-14 | 407 |
| ENSMUSG00000029420 | RIMS binding protein 2 [Source:MGI Symbol;Acc:MGI:2443235] | Rimbp2 | Enriched in SCN | 1.536 | 1.667E-17 | 390.3 |
| ENSMUSG00000029875 | coiled-coil domain containing 184 [Source:MGI Symbol;Acc:MGI:2146066] | Ccdc184 | Enriched in SCN | 1.534 | 4.009E-07 | 201 |
| ENSMUSG00000030811 | F-box and leucine-rich repeat protein 19 [Source:MGI Symbol;Acc:MGI:3039600] | Fbxl19 | Enriched in SCN | 1.532 | 0.000001809 | 714.5 |
| ENSMUSG00000110086 | small integral membrane protein 32 [Source:MGI Symbol;Acc:MGI:5791459] | Smim32 | Enriched in SCN | 1.531 | 0.002433 | 24.37 |
| ENSMUSG00000043719 | collagen, type VI, alpha 6 [Source:MGI Symbol;Acc:MGI:2444259] | Col6a6 | Enriched in SCN | 1.525 | 0.0002043 | 894 |
| ENSMUSG00000027624 | erythrocyte membrane protein band 4.1 like 1 [Source:MGI Symbol;Acc:MGI:103010] | Epb41l1 | Enriched in SCN | 1.522 | 2.391E-16 | 1373 |
| ENSMUSG00000023827 | 1-acylglycerol-3-phosphate O-acyltransferase 4 (lysophosphatidic acid acyltransferase, delta) [Source:MGI Symbol;Acc:MGI:1915512] | Agpat4 | Enriched in SCN | 1.505 | 2.189E-10 | 114.1 |
| ENSMUSG00000023017 | acid-sensing (proton-gated) ion channel 1 [Source:MGI Symbol;Acc:MGI:1194915] | Asic1 | Enriched in SCN | 1.499 | 8.709E-11 | 275.4 |
| ENSMUSG00000058793 | CDP-diacylglycerol synthase (phosphatidate cytidylyltransferase) 2 [Source:MGI Symbol;Acc:MGI:1332236] | Cds2 | Enriched in SCN | 1.49 | 4.197E-07 | 2925 |
| ENSMUSG00000070858 | predicted gene 1673 [Source:MGI Symbol;Acc:MGI:2686519] | Gm1673 | Enriched in SCN | 1.489 | 0.001161 | 55.45 |
| ENSMUSG00000026972 | arrestin domain containing 1 [Source:MGI Symbol;Acc:MGI:2446136] | Arrdc1 | Enriched in SCN | 1.481 | 0.0002268 | 25.53 |
| ENSMUSG00000108154 | predicted gene, 44033 [Source:MGI Symbol;Acc:MGI:5690425] | Gm44033 | Enriched in SCN | 1.479 | 0.006246 | 7.498 |
| ENSMUSG00000038295 | autophagy related 9B [Source:MGI Symbol;Acc:MGI:2685420] | Atg9b | Enriched in SCN | 1.478 | 0.004927 | 13.88 |
| ENSMUSG00000022623 | SH3 and multiple ankyrin repeat domains 3 [Source:MGI Symbol;Acc:MGI:1930016] | Shank3 | Enriched in SCN | 1.472 | 0.000001075 | 4014 |
| ENSMUSG00000059326 | colony stimulating factor 2 receptor, alpha, low-affinity (granulocyte-macrophage) [Source:MGI Symbol;Acc:MGI:1339754] | Csf2ra | Enriched in SCN | 1.471 | 5.488E-07 | 47.6 |
| ENSMUSG00000040952 | ribosomal protein S19 [Source:MGI Symbol;Acc:MGI:1333780] | Rps19 | Enriched in SCN | 1.461 | 0.00002673 | 155.7 |
| ENSMUSG00000005986 | ankyrin repeat domain 13 family, member D [Source:MGI Symbol;Acc:MGI:1915673] | Ankrd13d | Enriched in SCN | 1.456 | 5.715E-11 | 148.5 |
| ENSMUSG00000033510 | OTU domain containing 7A [Source:MGI Symbol;Acc:MGI:2158505] | Otud7a | Enriched in SCN | 1.453 | 0.000001185 | 416.2 |
| ENSMUSG00000073565 | proline rich 16 [Source:MGI Symbol;Acc:MGI:1918623] | Prr16 | Enriched in SCN | 1.453 | 0.0003239 | 79.02 |
| ENSMUSG00000075590 | nuclear receptor binding protein 2 [Source:MGI Symbol;Acc:MGI:2385017] | Nrbp2 | Enriched in SCN | 1.446 | 1.331E-10 | 333.6 |
| ENSMUSG00000032434 | CKLF-like MARVEL transmembrane domain containing 6 [Source:MGI Symbol;Acc:MGI:2447165] | Cmtm6 | Enriched in SCN | 1.446 | 0.00003418 | 131.1 |
| ENSMUSG00000026443 | leucine rich repeat protein 2, neuronal [Source:MGI Symbol;Acc:MGI:106037] | Lrrn2 | Enriched in SCN | 1.439 | 7.601E-10 | 489.2 |
| ENSMUSG00000018921 | proline, glutamic acid and leucine rich protein 1 [Source:MGI Symbol;Acc:MGI:1922523] | Pelp1 | Enriched in SCN | 1.437 | 4.304E-14 | 296.1 |

| | | | | | | |
|---|---|---|---|---|---|---|
| ENSMUSG00000049119 | family with sequence similarity 110, member B [Source:MGI Symbol;Acc:MGI:1916593] | Fam110b | Enriched in SCN | 1.434 | 0.00001121 | 150.5 |
| ENSMUSG00000117406 | netrin 3 [Source:MGI Symbol;Acc:MGI:1341188] | Ntn3 | Enriched in SCN | 1.434 | 0.00002821 | 52.98 |
| ENSMUSG00000052584 | stress-associated endoplasmic reticulum protein family member 2 [Source:MGI Symbol;Acc:MGI:1919911] | Serp2 | Enriched in SCN | 1.427 | 2.01E-08 | 134.2 |
| ENSMUSG00000040867 | brain-enriched guanylate kinase-associated [Source:MGI Symbol;Acc:MGI:3044626] | Begain | Enriched in SCN | 1.426 | 4.113E-07 | 445 |
| ENSMUSG00000036106 | proline rich 5 (renal) [Source:MGI Symbol;Acc:MGI:1924714] | Prr5 | Enriched in SCN | 1.422 | 3.387E-07 | 50.83 |
| ENSMUSG00000024248 | cytochrome c oxidase subunit 7A2 like [Source:MGI Symbol;Acc:MGI:106015] | Cox7a2l | Enriched in SCN | 1.421 | 0.000001809 | 252.8 |
| ENSMUSG00000043843 | transmembrane protein 145 [Source:MGI Symbol;Acc:MGI:3607779] | Tmem145 | Enriched in SCN | 1.42 | 1.918E-07 | 182.6 |
| ENSMUSG00000068099 | small integral membrane protein 45 [Source:MGI Symbol;Acc:MGI:1923755] | Smim45 | Enriched in SCN | 1.419 | 5.338E-08 | 83.12 |
| ENSMUSG00000036246 | Gem-interacting protein [Source:MGI Symbol;Acc:MGI:1926066] | Gmip | Enriched in SCN | 1.414 | 0.00001026 | 61.04 |
| ENSMUSG00000000627 | sema domain, immunoglobulin domain (Ig), TM domain, and short cytoplasmic domain [Source:MGI Symbol;Acc:MGI:1340055] | Sema4f | Enriched in SCN | 1.412 | 1.41E-08 | 125.4 |
| ENSMUSG00000004558 | N-myc downstream regulated gene 2 [Source:MGI Symbol;Acc:MGI:1352498] | Ndrg2 | Enriched in SCN | 1.404 | 1.031E-07 | 450.7 |
| ENSMUSG00000002771 | glutamate receptor, ionotropic, NMDA2D (epsilon 4) [Source:MGI Symbol;Acc:MGI:95823] | Grin2d | Enriched in SCN | 1.402 | 1.079E-09 | 466.7 |
| ENSMUSG00000021662 | Rho guanine nucleotide exchange factor (GEF) 28 [Source:MGI Symbol;Acc:MGI:1346016] | Arhgef28 | Enriched in SCN | 1.399 | 1.032E-09 | 213.6 |
| ENSMUSG00000019990 | phosphodiesterase 7B [Source:MGI Symbol;Acc:MGI:1352752] | Pde7b | Enriched in SCN | 1.394 | 0.0001156 | 194.2 |
| ENSMUSG00000020863 | LUC7-like 3 (S. cerevisiae) [Source:MGI Symbol;Acc:MGI:1914934] | Luc7l3 | Enriched in SCN | 1.391 | 0.00005552 | 973.3 |
| ENSMUSG00000051146 | calcium/calmodulin-dependent protein kinase II inhibitor 2 [Source:MGI Symbol;Acc:MGI:1920297] | Camk2n2 | Enriched in SCN | 1.385 | 1.531E-10 | 513.1 |
| ENSMUSG00000041115 | IQ motif and Sec7 domain 2 [Source:MGI Symbol;Acc:MGI:3528396] | Iqsec2 | Enriched in SCN | 1.384 | 1.411E-15 | 368.7 |
| ENSMUSG00000037503 | family with sequence similarity 168, member B [Source:MGI Symbol;Acc:MGI:2448487] | Fam168b | Enriched in SCN | 1.381 | 0.000000132 | 1846 |
| ENSMUSG00000026083 | eukaryotic translation initiation factor 5B [Source:MGI Symbol;Acc:MGI:2441772] | Eif5b | Enriched in SCN | 1.381 | 0.000001546 | 566 |
| ENSMUSG00000025375 | apoptosis-associated tyrosine kinase [Source:MGI Symbol;Acc:MGI:1197518] | Aatk | Enriched in SCN | 1.372 | 1.307E-09 | 1628 |
| ENSMUSG00000063239 | glutamate receptor, metabotropic 4 [Source:MGI Symbol;Acc:MGI:1351341] | Grm4 | Enriched in SCN | 1.368 | 4.157E-09 | 424.8 |
| ENSMUSG00000028680 | polo like kinase 3 [Source:MGI Symbol;Acc:MGI:109604] | Plk3 | Enriched in SCN | 1.367 | 0.00001188 | 41.32 |
| ENSMUSG00000028876 | Eph receptor A10 [Source:MGI Symbol;Acc:MGI:3586824] | Epha10 | Enriched in SCN | 1.357 | 6.357E-08 | 202.5 |
| ENSMUSG00000034245 | histone deacetylase 11 [Source:MGI Symbol;Acc:MGI:2385252] | Hdac11 | Enriched in SCN | 1.353 | 1.183E-09 | 513.4 |
| ENSMUSG00000001520 | nuclear receptor interacting protein 2 [Source:MGI Symbol;Acc:MGI:1891884] | Nrip2 | Enriched in SCN | 1.351 | 0.002909 | 17.02 |
| ENSMUSG00000042606 | HIRA interacting protein 3 [Source:MGI Symbol;Acc:MGI:2142364] | Hirip3 | Enriched in SCN | 1.35 | 0.0001323 | 132.7 |
| ENSMUSG00000029346 | SRR1 domain containing [Source:MGI Symbol;Acc:MGI:1917368] | Srrd | Enriched in SCN | 1.348 | 0.000008063 | 45.97 |
| ENSMUSG00000097185 | predicted gene, 26596 [Source:MGI Symbol;Acc:MGI:5477090] | Gm26596 | Enriched in SCN | 1.348 | 0.002422 | 16.73 |
| undefined | undefined | undefined | Enriched in SCN | 1.347 | 0.00001789 | 439.9 |
| ENSMUSG00000018507 | transient receptor potential cation channel, subfamily V, member 2 [Source:MGI Symbol;Acc:MGI:1341836] | Trpv2 | Enriched in SCN | 1.343 | 0.000006668 | 57.51 |
| ENSMUSG00000041119 | phosphodiesterase 9A [Source:MGI Symbol;Acc:MGI:1277179] | Pde9a | Enriched in SCN | 1.34 | 0.00001952 | 114 |
| ENSMUSG00000020083 | family with sequence similarity 241, member B [Source:MGI Symbol;Acc:MGI:1917144] | Fam241b | Enriched in SCN | 1.333 | 7.603E-07 | 42.28 |
| ENSMUSG00000053297 | expressed sequence AI854703 [Source:MGI Symbol;Acc:MGI:2141510] | AI854703 | Enriched in SCN | 1.326 | 2.306E-07 | 90.96 |
| ENSMUSG00000053580 | tetratricopeptide repeat, ankyrin repeat and coiled-coil containing 2 [Source:MGI Symbol;Acc:MGI:2444121] | Tanc2 | Enriched in SCN | 1.321 | 1.898E-15 | 1227 |
| ENSMUSG00000028949 | SWI/SNF related, matrix associated, actin dependent regulator of chromatin, subfamily d, member 3 [Source:MGI Symbol;Acc:MGI:1914243] | Smarcd3 | Enriched in SCN | 1.319 | 1.908E-15 | 234.8 |
| ENSMUSG00000004113 | calcium channel, voltage-dependent, N type, alpha 1B subunit [Source:MGI Symbol;Acc:MGI:88296] | Cacna1b | Enriched in SCN | 1.319 | 3.013E-09 | 1028 |
| ENSMUSG00000049804 | armadillo repeat containing, X-linked 4 [Source:MGI Symbol;Acc:MGI:2147887] | Armcx4 | Enriched in SCN | 1.318 | 0.000001147 | 1119 |
| ENSMUSG00000037509 | Rho guanine nucleotide exchange factor (GEF) 4 [Source:MGI Symbol;Acc:MGI:2442507] | Arhgef4 | Enriched in SCN | 1.317 | 2.608E-15 | 371.4 |
| ENSMUSG00000031910 | hyaluronan synthase 3 [Source:MGI Symbol;Acc:MGI:109599] | Has3 | Enriched in SCN | 1.314 | 0.01893 | 10.81 |
| ENSMUSG00000033350 | carbohydrate sulfotransferase 2 [Source:MGI Symbol;Acc:MGI:1891160] | Chst2 | Enriched in SCN | 1.313 | 0.00004161 | 1902 |
| ENSMUSG00000073557 | protein phosphatase 1, regulatory subunit 12B [Source:MGI Symbol;Acc:MGI:1916417] | Ppp1r12b | Enriched in SCN | 1.312 | 1.183E-14 | 691.5 |
| ENSMUSG00000038244 | microtubule associated monooxygenase, calponin and LIM domain containing 2 [Source:MGI Symbol;Acc:MGI:2444947] | Mical2 | Enriched in SCN | 1.311 | 0.00005331 | 255.1 |
| ENSMUSG00000038738 | SH3 and multiple ankyrin repeat domains 1 [Source:MGI Symbol;Acc:MGI:3613677] | Shank1 | Enriched in SCN | 1.31 | 0.000000318 | 6658 |
| ENSMUSG00000062184 | heparan sulfate 6-O-sulfotransferase 2 [Source:MGI Symbol;Acc:MGI:1354959] | Hs6st2 | Enriched in SCN | 1.31 | 0.00006206 | 235.4 |
| ENSMUSG00000031603 | fibroblast growth factor 20 [Source:MGI Symbol;Acc:MGI:1891346] | Fgf20 | Enriched in SCN | 1.309 | 0.02798 | 11.6 |
| ENSMUSG00000032878 | coiled-coil domain containing 85A [Source:MGI Symbol;Acc:MGI:2445069] | Ccdc85a | Enriched in SCN | 1.306 | 0.00002477 | 409.4 |
| ENSMUSG00000074182 | zinc finger, HIT type 6 [Source:MGI Symbol;Acc:MGI:1916996] | Znhit6 | Enriched in SCN | 1.305 | 0.00000104 | 173.8 |
| ENSMUSG00000028373 | astrotactin 2 [Source:MGI Symbol;Acc:MGI:1889277] | Astn2 | Enriched in SCN | 1.302 | 0.001445 | 621.4 |
| ENSMUSG00000059361 | neurensin 2 [Source:MGI Symbol;Acc:MGI:2684969] | Nrsn2 | Enriched in SCN | 1.301 | 0.001013 | 106.4 |
| ENSMUSG00000036264 | follistatin-like 4 [Source:MGI Symbol;Acc:MGI:2443199] | Fstl4 | Enriched in SCN | 1.3 | 0.002128 | 119.1 |
| ENSMUSG00000044646 | zinc finger and BTB domain containing 7C [Source:MGI Symbol;Acc:MGI:2443302] | Zbtb7c | Enriched in SCN | 1.298 | 0.004576 | 113.9 |
| ENSMUSG00000092216 | predicted gene, 19345 [Source:MGI Symbol;Acc:MGI:5011530] | Gm19345 | Enriched in SCN | 1.297 | 0.00023 | 23.03 |
| ENSMUSG00000056367 | ARP3 actin-related protein 3B [Source:MGI Symbol;Acc:MGI:2661120] | Actr3b | Enriched in SCN | 1.296 | 4.995E-09 | 140.3 |
| ENSMUSG00000054708 | ankyrin repeat domain 24 [Source:MGI Symbol;Acc:MGI:1890394] | Ankrd24 | Enriched in SCN | 1.292 | 0.000006668 | 154.8 |
| ENSMUSG00000006218 | family with sequence similarity 131, member C [Source:MGI Symbol;Acc:MGI:2685539] | Fam131c | Enriched in SCN | 1.289 | 0.000535 | 42.58 |
| ENSMUSG00000024299 | a disintegrin-like and metallopeptidase (reprolysin type) with thrombospondin type 1 motif, 10 [Source:MGI Symbol;Acc:MGI:2449112] | Adamts10 | Enriched in SCN | 1.289 | 0.002314 | 70.69 |
| ENSMUSG00000070003 | single stranded DNA binding protein 4 [Source:MGI Symbol;Acc:MGI:1924150] | Ssbp4 | Enriched in SCN | 1.284 | 1.271E-08 | 196.3 |
| ENSMUSG00000038777 | sema domain, transmembrane domain (TM), and cytoplasmic domain, (semaphorin) 6C [Source:MGI Symbol;Acc:MGI:1338032] | Sema6c | Enriched in SCN | 1.283 | 0.0005952 | 156.7 |
| ENSMUSG00000022211 | capping protein regulator and myosin 1 linker 3 [Source:MGI Symbol;Acc:MGI:2448573] | Carmil3 | Enriched in SCN | 1.278 | 0.000005221 | 317.9 |
| ENSMUSG00000026259 | neuronal guanine nucleotide exchange factor [Source:MGI Symbol;Acc:MGI:1858414] | Ngef | Enriched in SCN | 1.275 | 0.0001958 | 72.19 |
| ENSMUSG00000020654 | adenylate cyclase 3 [Source:MGI Symbol;Acc:MGI:99675] | Adcy3 | Enriched in SCN | 1.272 | 2.249E-11 | 175 |
| ENSMUSG00000029060 | mindbomb E3 ubiquitin protein ligase 2 [Source:MGI Symbol;Acc:MGI:2679684] | Mib2 | Enriched in SCN | 1.271 | 2.394E-08 | 340.9 |
| ENSMUSG00000051359 | neurocalcin delta [Source:MGI Symbol;Acc:MGI:1196326] | Ncald | Enriched in SCN | 1.27 | 4.104E-10 | 710.7 |
| ENSMUSG00000046613 | von Willebrand factor A domain containing 5B2 [Source:MGI Symbol;Acc:MGI:2681859] | Vwa5b2 | Enriched in SCN | 1.268 | 0.00002626 | 119.4 |
| ENSMUSG00000058297 | sparc/osteonectin, cwcv and kazal-like domains proteoglycan 2 [Source:MGI Symbol;Acc:MGI:1891351] | Spock2 | Enriched in SCN | 1.268 | 0.0002304 | 7309 |
| ENSMUSG00000024873 | cornichon family AMPA receptor auxiliary protein 2 [Source:MGI Symbol;Acc:MGI:1277225] | Cnih2 | Enriched in SCN | 1.259 | 1.907E-10 | 220.1 |
| ENSMUSG00000028337 | coronin, actin binding protein 2A [Source:MGI Symbol;Acc:MGI:1345966] | Coro2a | Enriched in SCN | 1.258 | 0.000003559 | 177.4 |
| ENSMUSG00000035640 | calcium channel, voltage-dependent, beta subunit associated regulatory protein [Source:MGI Symbol;Acc:MGI:1354170] | Cbarp | Enriched in SCN | 1.256 | 3.833E-07 | 1169 |
| ENSMUSG00000061028 | CLK4-associating serine/arginine rich protein [Source:MGI Symbol;Acc:MGI:1855695] | Clasrp | Enriched in SCN | 1.252 | 0.003265 | 182.6 |
| ENSMUSG00000045174 | APC membrane recruitment 3 [Source:MGI Symbol;Acc:MGI:3026939] | Amer3 | Enriched in SCN | 1.248 | 0.00000774 | 95.69 |
| ENSMUSG00000033854 | potassium channel, subfamily K, member 10 [Source:MGI Symbol;Acc:MGI:1919508] | Kcnk10 | Enriched in SCN | 1.244 | 0.000002793 | 191.6 |
| ENSMUSG00000001227 | sema domain, transmembrane domain (TM), and cytoplasmic domain, (semaphorin) 6B [Source:MGI Symbol;Acc:MGI:1202889] | Sema6b | Enriched in SCN | 1.239 | 2.163E-09 | 529.4 |
| ENSMUSG00000042817 | FMS-like tyrosine kinase 3 [Source:MGI Symbol;Acc:MGI:95559] | Flt3 | Enriched in SCN | 1.234 | 0.02486 | 42.78 |
| ENSMUSG00000053746 | peptidyl-tRNA hydrolase 1 homolog [Source:MGI Symbol;Acc:MGI:1913779] | Ptrh1 | Enriched in SCN | 1.227 | 0.02795 | 9.59 |
| ENSMUSG00000003273 | carbonic anhydrase 11 [Source:MGI Symbol;Acc:MGI:1336193] | Car11 | Enriched in SCN | 1.226 | 0.000001903 | 105.3 |
| ENSMUSG00000026156 | beta-1,3-glucuronyltransferase 2 (glucuronosyltransferase S) [Source:MGI Symbol;Acc:MGI:2389490] | B3gat2 | Enriched in SCN | 1.216 | 1.676E-08 | 140.5 |
| ENSMUSG00000021991 | calcium channel, voltage-dependent, alpha2/delta subunit 3 [Source:MGI Symbol;Acc:MGI:1338890] | Cacna2d3 | Enriched in SCN | 1.216 | 0.005453 | 366.8 |
| ENSMUSG00000033768 | neurexin II [Source:MGI Symbol;Acc:MGI:1096362] | Nrxn2 | Enriched in SCN | 1.213 | 1.673E-09 | 5038 |
| ENSMUSG00000047013 | F-box protein 41 [Source:MGI Symbol;Acc:MGI:1261912] | Fbxo41 | Enriched in SCN | 1.205 | 0.00001298 | 562.6 |
| ENSMUSG00000045348 | neuronal tyrosine-phosphorylated phosphoinositide 3-kinase adaptor 1 [Source:MGI Symbol;Acc:MGI:2443880] | Nyap1 | Enriched in SCN | 1.2 | 3.925E-09 | 501.1 |
| ENSMUSG00000042751 | nicotinamide nucleotide adenylyltransferase 2 [Source:MGI Symbol;Acc:MGI:2444155] | Nmnat2 | Enriched in SCN | 1.198 | 9.616E-07 | 932 |
| ENSMUSG00000034570 | inositol polyphosphate 5-phosphatase J [Source:MGI Symbol;Acc:MGI:2158663] | Inpp5j | Enriched in SCN | 1.198 | 0.001156 | 112.1 |
| ENSMUSG00000041556 | F-box protein 2 [Source:MGI Symbol;Acc:MGI:2446216] | Fbxo2 | Enriched in SCN | 1.193 | 0.005863 | 92.09 |
| ENSMUSG00000023911 | FLYWCH family member 2 [Source:MGI Symbol;Acc:MGI:1924167] | Flywch2 | Enriched in SCN | 1.192 | 0.0001365 | 50.53 |
| ENSMUSG00000032507 | F-box and leucine-rich repeat protein 2 [Source:MGI Symbol;Acc:MGI:1919429] | Fbxl2 | Enriched in SCN | 1.191 | 0.0001577 | 73.88 |
| ENSMUSG00000037784 | DAZ interacting protein 1-like [Source:MGI Symbol;Acc:MGI:1919757] | Dzip1l | Enriched in SCN | 1.188 | 1.182E-07 | 90.29 |
| ENSMUSG00000047261 | growth associated protein 43 [Source:MGI Symbol;Acc:MGI:95639] | Gap43 | Enriched in SCN | 1.187 | 0.00001683 | 876.7 |
| ENSMUSG00000007207 | syntaxin 1A (brain) [Source:MGI Symbol;Acc:MGI:109355] | Stx1a | Enriched in SCN | 1.183 | 0.00003415 | 278.7 |
| ENSMUSG00000002799 | jagged 2 [Source:MGI Symbol;Acc:MGI:1098270] | Jag2 | Enriched in SCN | 1.182 | 2.971E-10 | 295.8 |
| ENSMUSG00000040412 | endosome-lysosome associated apoptosis and autophagy regulator 1 [Source:MGI Symbol;Acc:MGI:1923930] | Elapor1 | Enriched in SCN | 1.179 | 0.00009798 | 93.25 |
| ENSMUSG00000038623 | transmembrane 6 superfamily member 1 [Source:MGI Symbol;Acc:MGI:1933209] | Tm6sf1 | Enriched in SCN | 1.178 | 0.00589 | 52.29 |
| ENSMUSG00000095098 | coiled-coil domain containing 85B [Source:MGI Symbol;Acc:MGI:2147607] | Ccdc85b | Enriched in SCN | 1.176 | 0.00001651 | 384.5 |
| ENSMUSG00000027394 | tubulin tyrosine ligase [Source:MGI Symbol;Acc:MGI:1916987] | Ttl | Enriched in SCN | 1.163 | 0.00001596 | 343 |
| ENSMUSG00000044912 | synaptotagmin XVI [Source:MGI Symbol;Acc:MGI:2673872] | Syt16 | Enriched in SCN | 1.162 | 0.00004106 | 222.1 |
| ENSMUSG00000041037 | immunity-related GTPase family, Q [Source:MGI Symbol;Acc:MGI:2667176] | Irgq | Enriched in SCN | 1.159 | 6.081E-07 | 618.2 |
| ENSMUSG00000026494 | kinesin family member 26B [Source:MGI Symbol;Acc:MGI:2447076] | Kif26b | Enriched in SCN | 1.152 | 0.00009441 | 391.1 |
| ENSMUSG00000028782 | adhesion G protein-coupled receptor B2 [Source:MGI Symbol;Acc:MGI:2451244] | Adgrb2 | Enriched in SCN | 1.15 | 2.121E-07 | 1121 |
| ENSMUSG00000032537 | Eph receptor B1 [Source:MGI Symbol;Acc:MGI:1096337] | Ephb1 | Enriched in SCN | 1.149 | 0.00007045 | 295.7 |
| ENSMUSG00000041654 | solute carrier family 39 (metal ion transporter), member 11 [Source:MGI Symbol;Acc:MGI:1917056] | Slc39a11 | Enriched in SCN | 1.148 | 0.000352 | 62.34 |
| ENSMUSG00000038077 | potassium voltage-gated channel, shaker-related, subfamily, member 6 [Source:MGI Symbol;Acc:MGI:96663] | Kcna6 | Enriched in SCN | 1.147 | 8.036E-10 | 491.8 |
| ENSMUSG00000033597 | CASK interacting protein 1 [Source:MGI Symbol;Acc:MGI:2442952] | Caskin1 | Enriched in SCN | 1.147 | 1.159E-09 | 1073 |
| ENSMUSG00000011589 | fibronectin type 3 and SPRY domain-containing protein [Source:MGI Symbol;Acc:MGI:1934858] | Fsd1 | Enriched in SCN | 1.147 | 0.0007456 | 96.92 |
| ENSMUSG00000029916 | acylglycerol kinase [Source:MGI Symbol;Acc:MGI:1917173] | Agk | Enriched in SCN | 1.144 | 0.0007274 | 70.67 |
| ENSMUSG00000034686 | proline rich 7 (synaptic) [Source:MGI Symbol;Acc:MGI:3487246] | Prr7 | Enriched in SCN | 1.143 | 7.955E-07 | 94.05 |
| ENSMUSG00000038042 | protein tyrosine phosphatase domain containing 1 [Source:MGI Symbol;Acc:MGI:2145430] | Ptpdc1 | Enriched in SCN | 1.14 | 0.00000134 | 145.1 |
| ENSMUSG00000031376 | ATPase, Ca++ transporting, plasma membrane 3 [Source:MGI Symbol;Acc:MGI:1347353] | Atp2b3 | Enriched in SCN | 1.138 | 0.0005279 | 308 |
| ENSMUSG00000022199 | solute carrier family 22 (organic cation transporter), member 17 [Source:MGI Symbol;Acc:MGI:1926225] | Slc22a17 | Enriched in SCN | 1.136 | 0.000001809 | 805.9 |
| ENSMUSG00000053137 | mitogen-activated protein kinase 11 [Source:MGI Symbol;Acc:MGI:1338024] | Mapk11 | Enriched in SCN | 1.133 | 0.0007584 | 63.14 |
| ENSMUSG00000049044 | Rap guanine nucleotide exchange factor (GEF) 4 [Source:MGI Symbol;Acc:MGI:1917723] | Rapgef4 | Enriched in SCN | 1.126 | 0.00001725 | 306 |
| ENSMUSG00000026278 | BCL2-related ovarian killer [Source:MGI Symbol;Acc:MGI:1858494] | Bok | Enriched in SCN | 1.125 | 0.0025 | 219.1 |
| ENSMUSG00000061046 | hydroxyacylglutathione hydrolase-like [Source:MGI Symbol;Acc:MGI:1919877] | Haghl | Enriched in SCN | 1.123 | 0.00004307 | 162.5 |
| ENSMUSG00000046727 | cysteine-rich transmembrane module containing 1 [Source:MGI Symbol;Acc:MGI:1913310] | Cystm1 | Enriched in SCN | 1.123 | 0.003605 | 23.26 |
| ENSMUSG00000029669 | tetraspanin 12 [Source:MGI Symbol;Acc:MGI:1889818] | Tspan12 | Enriched in SCN | 1.12 | 0.000004905 | 93.15 |
| ENSMUSG00000051177 | phospholipase C, beta 1 [Source:MGI Symbol;Acc:MGI:97613] | Plcb1 | Enriched in SCN | 1.12 | 0.007025 | 366.6 |
| ENSMUSG00000038020 | Rap guanine nucleotide exchange factor (GEF)-like 1 [Source:MGI Symbol;Acc:MGI:3611446] | Rapgefl1 | Enriched in SCN | 1.119 | 3.783E-08 | 205.6 |
| ENSMUSG00000064254 | ethylmalonic encephalopathy 1 [Source:MGI Symbol;Acc:MGI:1913321] | Ethe1 | Enriched in SCN | 1.117 | 0.001697 | 23.39 |
| ENSMUSG00000054477 | potassium intermediate/small conductance calcium-activated channel, subfamily N, member 2 [Source:MGI Symbol;Acc:MGI:2153182] | Kcnn2 | Enriched in SCN | 1.114 | 0.002553 | 198.9 |
| ENSMUSG00000056258 | potassium voltage-gated channel, subfamily Q, member 3 [Source:MGI Symbol;Acc:MGI:1336181] | Kcnq3 | Enriched in SCN | 1.111 | 0.0006292 | 1087 |
| ENSMUSG00000045318 | adrenergic receptor, alpha 2c [Source:MGI Symbol;Acc:MGI:87936] | Adra2c | Enriched in SCN | 1.108 | 0.01552 | 33.8 |
| ENSMUSG00000026458 | protein tyrosine phosphatase, receptor type, f polypeptide (PTPRF), interacting protein (liprin), alpha 4 [Source:MGI Symbol;Acc:MGI:1915757] | Ppfia4 | Enriched in SCN | 1.105 | 4.007E-12 | 888.5 |
| ENSMUSG00000003528 | solute carrier family 25 (mitochondrial carrier, citrate transporter), member 1 [Source:MGI Symbol;Acc:MGI:1345283] | Slc25a1 | Enriched in SCN | 1.104 | 0.0001529 | 63.13 |
| ENSMUSG00000035051 | DEAH (Asp-Glu-Ala-Asp/His) box polypeptide 57 [Source:MGI Symbol;Acc:MGI:2147067] | Dhx57 | Enriched in SCN | 1.098 | 0.000002162 | 248.4 |
| ENSMUSG00000032425 | zinc finger protein 949 [Source:MGI Symbol;Acc:MGI:1918890] | Zfp949 | Enriched in SCN | 1.097 | 0.00258 | 260.3 |
| ENSMUSG00000026959 | glutamate receptor, ionotropic, NMDA1 (zeta 1) [Source:MGI Symbol;Acc:MGI:95819] | Grin1 | Enriched in SCN | 1.095 | 0.0008141 | 942.6 |
| ENSMUSG00000040350 | tripartite motif-containing 7 [Source:MGI Symbol;Acc:MGI:2137353] | Trim7 | Enriched in SCN | 1.093 | 0.04716 | 12.75 |
| ENSMUSG00000043460 | leucine rich repeat and fibronectin type III, extracellular 2 [Source:MGI Symbol;Acc:MGI:3608416] | Elfn2 | Enriched in SCN | 1.081 | 0.00005554 | 990.9 |
| ENSMUSG00000025743 | syndecan 3 [Source:MGI Symbol;Acc:MGI:1349163] | Sdc3 | Enriched in SCN | 1.076 | 0.00004318 | 1949 |
| ENSMUSG00000022421 | neuronal pentraxin receptor [Source:MGI Symbol;Acc:MGI:1920590] | Nptxr | Enriched in SCN | 1.074 | 0.001311 | 1078 |

| | | | | | | |
|---|---|---|---|---|---|---|
| ENSMUSG00000027525 | phosphatase and actin regulator 3 [Source:MGI Symbol;Acc:MGI:1921439] | Phactr3 | Enriched in SCN | 1.072 | 1.165E-08 | 523.5 |
| ENSMUSG00000032193 | low density lipoprotein receptor [Source:MGI Symbol;Acc:MGI:96765] | Ldlr | Enriched in SCN | 1.07 | 0.0002615 | 63.28 |
| ENSMUSG00000039787 | cerebral endothelial cell adhesion molecule [Source:MGI Symbol;Acc:MGI:2139134] | Cercam | Enriched in SCN | 1.067 | 0.0005773 | 23.25 |
| ENSMUSG00000042532 | golgi autoantigen, golgin subfamily a, 7B [Source:MGI Symbol;Acc:MGI:1918396] | Golga7b | Enriched in SCN | 1.053 | 5.289E-07 | 283.6 |
| ENSMUSG00000028541 | UDP-Gal:betaGlcNAc beta 1,4- galactosyltransferase, polypeptide 2 [Source:MGI Symbol;Acc:MGI:1858493] | B4galt2 | Enriched in SCN | 1.052 | 0.00003595 | 235.2 |
| ENSMUSG00000002279 | lipase maturation factor 1 [Source:MGI Symbol;Acc:MGI:1923733] | Lmf1 | Enriched in SCN | 1.049 | 0.002172 | 68.12 |
| ENSMUSG00000039477 | trinucleotide repeat containing 18 [Source:MGI Symbol;Acc:MGI:3648294] | Tnrc18 | Enriched in SCN | 1.048 | 2.933E-08 | 2428 |
| ENSMUSG00000053519 | Kv channel-interacting protein 1 [Source:MGI Symbol;Acc:MGI:1917607] | Kcnip1 | Enriched in SCN | 1.047 | 0.02903 | 534.9 |
| ENSMUSG00000046709 | mitogen-activated protein kinase 10 [Source:MGI Symbol;Acc:MGI:1346863] | Mapk10 | Enriched in SCN | 1.045 | 0.00004897 | 662.3 |
| ENSMUSG00000040640 | ELKS/RAB6-interacting/CAST family member 2 [Source:MGI Symbol;Acc:MGI:1098749] | Erc2 | Enriched in SCN | 1.045 | 0.002181 | 1202 |
| ENSMUSG00000028944 | protein kinase, AMP-activated, gamma 2 non-catalytic subunit [Source:MGI Symbol;Acc:MGI:1336153] | Prkag2 | Enriched in SCN | 1.042 | 0.0002547 | 280.9 |
| ENSMUSG00000057649 | bromodomain containing 9 [Source:MGI Symbol;Acc:MGI:2145317] | Brd9 | Enriched in SCN | 1.042 | 0.0005791 | 227.3 |
| ENSMUSG00000070509 | repulsive guidance molecule family member A [Source:MGI Symbol;Acc:MGI:2679262] | Rgma | Enriched in SCN | 1.037 | 0.000005103 | 395.9 |
| ENSMUSG00000033545 | zinc and ring finger 1 [Source:MGI Symbol;Acc:MGI:2177308] | Znrf1 | Enriched in SCN | 1.037 | 0.00002925 | 710.6 |
| ENSMUSG00000038267 | solute carrier family 22, member 23 [Source:MGI Symbol;Acc:MGI:1920352] | Slc22a23 | Enriched in SCN | 1.031 | 0.000005465 | 369.7 |
| ENSMUSG00000025348 | integrin alpha 7 [Source:MGI Symbol;Acc:MGI:102700] | Itga7 | Enriched in SCN | 1.031 | 0.007645 | 44.33 |
| ENSMUSG00000056310 | tRNA-yW synthesizing protein 1 homolog (S. cerevisiae) [Source:MGI Symbol;Acc:MGI:2141161] | Tyw1 | Enriched in SCN | 1.029 | 0.0006541 | 150.1 |
| ENSMUSG00000041837 | programmed cell death 7 [Source:MGI Symbol;Acc:MGI:1859170] | Pdcd7 | Enriched in SCN | 1.023 | 0.000005766 | 140.5 |
| ENSMUSG00000055629 | beta-1,4-N-acetyl-galactosaminyl transferase 4 [Source:MGI Symbol;Acc:MGI:2652891] | B4galnt4 | Enriched in SCN | 1.023 | 0.000006232 | 491.4 |
| ENSMUSG00000040390 | mitogen-activated protein kinase kinase kinase 10 [Source:MGI Symbol;Acc:MGI:1346879] | Map3k10 | Enriched in SCN | 1.023 | 0.00007652 | 1531 |
| ENSMUSG00000051323 | protocadherin 19 [Source:MGI Symbol;Acc:MGI:2685563] | Pcdh19 | Enriched in SCN | 1.02 | 0.002109 | 636.5 |
| ENSMUSG00000040035 | dispatched RND transporter family member 2 [Source:MGI Symbol;Acc:MGI:2388733] | Disp2 | Enriched in SCN | 1.015 | 0.0004813 | 1280 |
| ENSMUSG00000040133 | G protein-coupled receptor 176 [Source:MGI Symbol;Acc:MGI:2685858] | Gpr176 | Enriched in SCN | 1.012 | 0.0003498 | 77.99 |
| ENSMUSG00000013150 | glucose-fructose oxidoreductase domain containing 2 [Source:MGI Symbol;Acc:MGI:1917825] | Gfod2 | Enriched in SCN | 1.011 | 3.268E-08 | 156.4 |
| ENSMUSG00000036760 | potassium channel, subfamily K, member 9 [Source:MGI Symbol;Acc:MGI:3521816] | Kcnk9 | Enriched in SCN | 1.009 | 0.003146 | 406.3 |
| ENSMUSG00000054793 | cell adhesion molecule 4 [Source:MGI Symbol;Acc:MGI:2449088] | Cadm4 | Enriched in SCN | 1.008 | 0.000001549 | 1063 |
| ENSMUSG00000003863 | protein tyrosine phosphatase, receptor type, f polypeptide (PTPRF), interacting protein (liprin), alpha 3 [Source:MGI Symbol;Acc:MGI:1924037] | Ppfia3 | Enriched in SCN | 1 | 2.745E-08 | 864.8 |
| ENSMUSG00000060240 | cell cycle exit and neuronal differentiation 1 [Source:MGI Symbol;Acc:MGI:1929898] | Cend1 | Enriched in retina | -1.014 | 0.02568 | 473.2 |
| ENSMUSG00000068699 | filamin C, gamma [Source:MGI Symbol;Acc:MGI:95557] | Flnc | Enriched in retina | -1.015 | 0.07348 | 29.35 |
| ENSMUSG00000033209 | tetratricopeptide repeat domain 28 [Source:MGI Symbol;Acc:MGI:2140873] | Ttc28 | Enriched in retina | -1.022 | 0.0008545 | 597.6 |
| ENSMUSG00000029608 | rabphilin 3A [Source:MGI Symbol;Acc:MGI:102788] | Rph3a | Enriched in retina | -1.023 | 0.0009133 | 1196 |
| ENSMUSG00000041592 | sidekick cell adhesion molecule 2 [Source:MGI Symbol;Acc:MGI:2443847] | Sdk2 | Enriched in retina | -1.024 | 0.0002087 | 398.3 |
| ENSMUSG00000092032 | vomeronasal 2, receptor 59 [Source:MGI Symbol;Acc:MGI:3643747] | Vmn2r59 | Enriched in retina | -1.025 | 0.1989 | 5.159 |
| ENSMUSG00000026321 | tumor necrosis factor receptor superfamily, member 11a, NFKB activator [Source:MGI Symbol;Acc:MGI:1314891] | Tnfrsf11a | Enriched in retina | -1.027 | 0.01643 | 53.11 |
| ENSMUSG00000027296 | inositol 1,4,5-trisphosphate 3-kinase A [Source:MGI Symbol;Acc:MGI:1333822] | Itpka | Enriched in retina | -1.036 | 0.1549 | 9.715 |
| ENSMUSG00000048899 | ribosomal modification protein rimK-like family member A [Source:MGI Symbol;Acc:MGI:3040686] | Rimkla | Enriched in retina | -1.051 | 0.00002886 | 161 |
| ENSMUSG00000064220 | H2A clustered histone 18 [Source:MGI Symbol;Acc:MGI:96097] | H2ac18 | Enriched in retina | -1.051 | 0.1599 | 6.736 |
| ENSMUSG00000052752 | TNF receptor-associated factor 7 [Source:MGI Symbol;Acc:MGI:3042141] | Traf7 | Enriched in retina | -1.055 | 0.00002505 | 103.5 |
| ENSMUSG00000032468 | armadillo repeat containing 8 [Source:MGI Symbol;Acc:MGI:1921375] | Armc8 | Enriched in retina | -1.056 | 0.00007181 | 227.6 |
| ENSMUSG00000034994 | eukaryotic translation elongation factor 2 [Source:MGI Symbol;Acc:MGI:95288] | Eef2 | Enriched in retina | -1.059 | 2.703E-08 | 4640 |
| ENSMUSG00000037151 | leucine rich repeat containing 20 [Source:MGI Symbol;Acc:MGI:2387182] | Lrrc20 | Enriched in retina | -1.065 | 0.001515 | 64.28 |
| ENSMUSG00000028631 | potassium voltage-gated channel, subfamily Q, member 4 [Source:MGI Symbol;Acc:MGI:1926803] | Kcnq4 | Enriched in retina | -1.069 | 0.001685 | 45.84 |
| ENSMUSG00000021219 | regulator of G-protein signaling 6 [Source:MGI Symbol;Acc:MGI:1354730] | Rgs6 | Enriched in retina | -1.069 | 0.01874 | 225 |
| ENSMUSG00000062300 | nectin cell adhesion molecule 2 [Source:MGI Symbol;Acc:MGI:97822] | Nectin2 | Enriched in retina | -1.074 | 0.0009139 | 66.46 |
| ENSMUSG00000029503 | purinergic receptor P2X, ligand-gated ion channel, 2 [Source:MGI Symbol;Acc:MGI:2665170] | P2rx2 | Enriched in retina | -1.075 | 0.2311 | 6.203 |
| ENSMUSG00000031065 | cyclin-dependent kinase 16 [Source:MGI Symbol;Acc:MGI:97516] | Cdk16 | Enriched in retina | -1.086 | 4.061E-09 | 785.5 |
| ENSMUSG00000036980 | TATA-box binding protein associated factor 6 [Source:MGI Symbol;Acc:MGI:109129] | Taf6 | Enriched in retina | -1.117 | 4.086E-08 | 74.4 |
| ENSMUSG00000044505 | leucine rich repeat and Ig domain containing 4 [Source:MGI Symbol;Acc:MGI:2444651] | Lingo4 | Enriched in retina | -1.121 | 0.01646 | 15.71 |
| ENSMUSG00000027577 | cholinergic receptor, nicotinic, alpha polypeptide 4 [Source:MGI Symbol;Acc:MGI:87888] | Chrna4 | Enriched in retina | -1.127 | 0.0002793 | 546.3 |
| ENSMUSG00000046841 | cytoskeleton-associated protein 4 [Source:MGI Symbol;Acc:MGI:2444926] | Ckap4 | Enriched in retina | -1.144 | 6.329E-08 | 323.5 |
| ENSMUSG00000040703 | cytochrome P450, family 2, subfamily s, polypeptide 1 [Source:MGI Symbol;Acc:MGI:1921384] | Cyp2s1 | Enriched in retina | -1.148 | 0.03657 | 19.07 |
| ENSMUSG00000018634 | corticotropin releasing hormone receptor 1 [Source:MGI Symbol;Acc:MGI:88498] | Crhr1 | Enriched in retina | -1.159 | 0.003993 | 47.56 |
| ENSMUSG00000030352 | tetraspanin 9 [Source:MGI Symbol;Acc:MGI:1924558] | Tspan9 | Enriched in retina | -1.168 | 1.408E-08 | 538.1 |
| ENSMUSG00000039137 | whirlin [Source:MGI Symbol;Acc:MGI:2682003] | Whrn | Enriched in retina | -1.186 | 0.00001414 | 351.2 |
| ENSMUSG00000022885 | beta galactoside alpha 2,6 sialyltransferase 1 [Source:MGI Symbol;Acc:MGI:108470] | St6gal1 | Enriched in retina | -1.187 | 0.0003944 | 86.45 |
| ENSMUSG00000052387 | transient receptor potential cation channel, subfamily M, member 3 [Source:MGI Symbol;Acc:MGI:2443101] | Trpm3 | Enriched in retina | -1.189 | 0.0005788 | 2184 |
| ENSMUSG00000052302 | TBC1 domain family, member 30 [Source:MGI Symbol;Acc:MGI:1921944] | Tbc1d30 | Enriched in retina | -1.215 | 2.427E-09 | 314.2 |
| ENSMUSG00000032011 | thymus cell antigen 1, theta [Source:MGI Symbol;Acc:MGI:98747] | Thy1 | Enriched in retina | -1.216 | 0.0006922 | 952.8 |
| ENSMUSG00000056596 | TMF1-regulated nuclear protein 1 [Source:MGI Symbol;Acc:MGI:1916789] | Trnp1 | Enriched in retina | -1.219 | 0.00005053 | 640.5 |
| ENSMUSG00000039461 | T cell leukemia translocation altered gene [Source:MGI Symbol;Acc:MGI:1918829] | Tcta | Enriched in retina | -1.229 | 0.00001803 | 50.88 |
| ENSMUSG00000036913 | tripartite motif-containing 67 [Source:MGI Symbol;Acc:MGI:3045323] | Trim67 | Enriched in retina | -1.229 | 0.001478 | 572.3 |
| ENSMUSG00000027071 | purinergic receptor P2X, ligand-gated ion channel, 3 [Source:MGI Symbol;Acc:MGI:1097160] | P2rx3 | Enriched in retina | -1.25 | 0.1189 | 12.15 |
| ENSMUSG00000072476 | predicted pseudogene 9008 [Source:MGI Symbol;Acc:MGI:3644000] | Gm9008 | Enriched in retina | -1.255 | 0.03042 | 8.19 |
| ENSMUSG00000030782 | transforming growth factor beta 1 induced transcript 1 [Source:MGI Symbol;Acc:MGI:102784] | Tgfb1i1 | Enriched in retina | -1.261 | 0.001802 | 24.56 |
| ENSMUSG00000020635 | FK506 binding protein 1b [Source:MGI Symbol;Acc:MGI:1336205] | Fkbp1b | Enriched in retina | -1.267 | 0.003481 | 66.86 |
| ENSMUSG00000039252 | leucine-rich repeat LGI family, member 2 [Source:MGI Symbol;Acc:MGI:2180196] | Lgi2 | Enriched in retina | -1.272 | 0.0007875 | 332.3 |
| ENSMUSG00000051502 | UFM1-specific peptidase 1 [Source:MGI Symbol;Acc:MGI:1917490] | Ufsp1 | Enriched in retina | -1.277 | 0.00009835 | 52.27 |
| ENSMUSG00000078249 | high mobility group AT-hook 1B [Source:MGI Symbol;Acc:MGI:96161] | Hmga1b | Enriched in retina | -1.284 | 0.00003091 | 54.25 |
| ENSMUSG00000026525 | opsin 3 [Source:MGI Symbol;Acc:MGI:1338022] | Opn3 | Enriched in retina | -1.289 | 0.2618 | 17.62 |
| ENSMUSG00000000326 | catechol-O-methyltransferase [Source:MGI Symbol;Acc:MGI:88470] | Comt | Enriched in retina | -1.291 | 0.000000476 | 103.3 |
| ENSMUSG00000006764 | tryptophan hydroxylase 2 [Source:MGI Symbol;Acc:MGI:2651811] | Tph2 | Enriched in retina | -1.291 | 0.1559 | 3.884 |
| ENSMUSG00000033436 | armadillo repeat containing, X-linked 2 [Source:MGI Symbol;Acc:MGI:1914666] | Armcx2 | Enriched in retina | -1.305 | 1.366E-08 | 262.7 |
| ENSMUSG00000028926 | cyclin-dependent kinase 14 [Source:MGI Symbol;Acc:MGI:894318] | Cdk14 | Enriched in retina | -1.311 | 8.243E-07 | 747.6 |
| ENSMUSG00000022999 | limb region 1 like [Source:MGI Symbol;Acc:MGI:1289247] | Lmbr1l | Enriched in retina | -1.331 | 0.004371 | 52.69 |
| ENSMUSG00000049625 | TRAF-interacting protein with forkhead-associated domain, family member B [Source:MGI Symbol;Acc:MGI:2385852] | Tifab | Enriched in retina | -1.338 | 0.04811 | 7.994 |
| ENSMUSG00000025155 | dihydrouridine synthase 1-like (S. cerevisiae) [Source:MGI Symbol;Acc:MGI:1915980] | Dus1l | Enriched in retina | -1.343 | 0.00004899 | 60.74 |
| ENSMUSG00000048385 | scratch family zinc finger 1 [Source:MGI Symbol;Acc:MGI:2176606] | Scrt1 | Enriched in retina | -1.369 | 0.000001083 | 1666 |
| ENSMUSG00000094500 | small integral membrane protein 18 [Source:MGI Symbol;Acc:MGI:1919882] | Smim18 | Enriched in retina | -1.369 | 0.1389 | 4.253 |
| ENSMUSG00000041046 | receptor (calcitonin) activity modifying protein 3 [Source:MGI Symbol;Acc:MGI:1860292] | Ramp3 | Enriched in retina | -1.375 | 0.0009669 | 121.1 |
| ENSMUSG00000019796 | low density lipoprotein receptor-related protein 11 [Source:MGI Symbol;Acc:MGI:2442989] | Lrp11 | Enriched in retina | -1.378 | 9.7E-09 | 271 |
| ENSMUSG00000030598 | F-box protein 17 [Source:MGI Symbol;Acc:MGI:1354707] | Fbxo17 | Enriched in retina | -1.391 | 0.0191 | 10.55 |
| ENSMUSG00000029005 | dorsal inhibitory axon guidance protein [Source:MGI Symbol;Acc:MGI:1917683] | Draxin | Enriched in retina | -1.429 | 0.00003726 | 35.71 |
| ENSMUSG00000001794 | calpain, small subunit 1 [Source:MGI Symbol;Acc:MGI:88266] | Capns1 | Enriched in retina | -1.448 | 0.00001905 | 150.2 |
| ENSMUSG00000042129 | Ras association (RalGDS/AF-6) domain family member 4 [Source:MGI Symbol;Acc:MGI:2386853] | Rassf4 | Enriched in retina | -1.477 | 0.00001243 | 89.04 |
| ENSMUSG00000040373 | calcium channel, voltage-dependent, gamma subunit 5 [Source:MGI Symbol;Acc:MGI:2157946] | Cacng5 | Enriched in retina | -1.501 | 9.058E-08 | 203.8 |
| ENSMUSG00000022055 | neurofilament, light polypeptide [Source:MGI Symbol;Acc:MGI:97313] | Nefl | Enriched in retina | -1.514 | 0.00000808 | 7224 |
| ENSMUSG00000021256 | vasohibin 1 [Source:MGI Symbol;Acc:MGI:2442543] | Vash1 | Enriched in retina | -1.54 | 2.902E-13 | 304.6 |
| ENSMUSG00000004098 | collagen, type V, alpha 3 [Source:MGI Symbol;Acc:MGI:1858212] | Col5a3 | Enriched in retina | -1.551 | 0.001172 | 67.35 |
| ENSMUSG00000071234 | synapse differentiation inducing 1 like [Source:MGI Symbol;Acc:MGI:2685107] | Syndig1l | Enriched in retina | -1.554 | 0.000002951 | 79.49 |
| ENSMUSG00000058925 | coiled-coil domain containing 192 [Source:MGI Symbol;Acc:MGI:1922694] | Ccdc192 | Enriched in retina | -1.559 | 0.02875 | 7.152 |
| ENSMUSG00000041439 | major facilitator superfamily domain containing 6 [Source:MGI Symbol;Acc:MGI:1922925] | Mfsd6 | Enriched in retina | -1.582 | 9.587E-07 | 415.4 |
| ENSMUSG00000066189 | calcium channel, voltage-dependent, gamma subunit 3 [Source:MGI Symbol;Acc:MGI:1859165] | Cacng3 | Enriched in retina | -1.585 | 1.136E-09 | 109.7 |
| ENSMUSG00000033174 | monoglyceride lipase [Source:MGI Symbol;Acc:MGI:1346042] | Mgll | Enriched in retina | -1.592 | 7.881E-10 | 422.8 |
| ENSMUSG00000022894 | a disintegrin-like and metallopeptidase (reprolysin type) with thrombospondin type 1 motif, 5 (aggrecanase-2) [Source:MGI Symbol;Acc:MGI:13463 | Adamts5 | Enriched in retina | -1.601 | 0.000003615 | 56.57 |
| ENSMUSG00000035407 | KN motif and ankyrin repeat domains 4 [Source:MGI Symbol;Acc:MGI:3043381] | Kank4 | Enriched in retina | -1.601 | 0.004158 | 22.26 |
| ENSMUSG00000046668 | CXXC finger 5 [Source:MGI Symbol;Acc:MGI:1914643] | Cxxc5 | Enriched in retina | -1.606 | 1.588E-20 | 291.6 |
| ENSMUSG00000056665 | thioesterase superfamily member 6 [Source:MGI Symbol;Acc:MGI:1925301] | Them6 | Enriched in retina | -1.614 | 0.0001733 | 58.01 |
| ENSMUSG00000033615 | complexin 1 [Source:MGI Symbol;Acc:MGI:104727] | Cplx1 | Enriched in retina | -1.632 | 0.00001026 | 2304 |
| ENSMUSG00000034883 | leucine rich repeat protein 1 [Source:MGI Symbol;Acc:MGI:1916956] | Lrr1 | Enriched in retina | -1.637 | 0.2834 | 14.15 |
| ENSMUSG00000073991 | cyclic nucleotide binding domain containing 1 [Source:MGI Symbol;Acc:MGI:3650508] | Cnbd1 | Enriched in retina | -1.674 | 0.01549 | 9.142 |
| ENSMUSG00000039183 | nucleotide binding protein 2 [Source:MGI Symbol;Acc:MGI:1347072] | Nubp2 | Enriched in retina | -1.68 | 0.00001341 | 28.88 |
| ENSMUSG00000024871 | double C2, gamma [Source:MGI Symbol;Acc:MGI:1926250] | Doc2g | Enriched in retina | -1.684 | 0.02772 | 6.218 |
| ENSMUSG00000021948 | protein kinase C, delta [Source:MGI Symbol;Acc:MGI:97598] | Prkcd | Enriched in retina | -1.731 | 0.002339 | 727.9 |
| ENSMUSG00000034402 | potassium voltage-gated channel, subfamily H (eag-related), member 5 [Source:MGI Symbol;Acc:MGI:3584508] | Kcnh5 | Enriched in retina | -1.743 | 5.236E-08 | 240.3 |
| ENSMUSG00000044469 | tumor necrosis factor, alpha-induced protein 8-like 1 [Source:MGI Symbol;Acc:MGI:1913693] | Tnfaip8l1 | Enriched in retina | -1.744 | 0.002415 | 9.788 |
| ENSMUSG00000030499 | potassium channel tetramerisation domain containing 15 [Source:MGI Symbol;Acc:MGI:2385276] | Kctd15 | Enriched in retina | -1.76 | 5.847E-10 | 127.8 |
| ENSMUSG00000050822 | solute carrier family 29 (nucleoside transporters), member 4 [Source:MGI Symbol;Acc:MGI:2385330] | Slc29a4 | Enriched in retina | -1.823 | 1.588E-07 | 126.5 |
| ENSMUSG00000039662 | isoprenylcysteine carboxyl methyltransferase [Source:MGI Symbol;Acc:MGI:1888594] | Icmt | Enriched in retina | -1.827 | 9.146E-12 | 734.9 |
| ENSMUSG00000001333 | syncoilin [Source:MGI Symbol;Acc:MGI:1916078] | Sync | Enriched in retina | -1.844 | 0.0004025 | 17.38 |
| ENSMUSG00000020810 | cytoglobin [Source:MGI Symbol;Acc:MGI:2149481] | Cygb | Enriched in retina | -1.864 | 1.338E-08 | 213.7 |
| ENSMUSG00000042115 | kelch domain containing 8A [Source:MGI Symbol;Acc:MGI:2442630] | Klhdc8a | Enriched in retina | -1.89 | 1.25E-12 | 181.2 |
| ENSMUSG00000035296 | sarcoglycan, gamma (dystrophin-associated glycoprotein) [Source:MGI Symbol;Acc:MGI:1346524] | Sgcg | Enriched in retina | -1.89 | 0.07971 | 9.668 |
| ENSMUSG00000056296 | synaptoporin [Source:MGI Symbol;Acc:MGI:1919253] | Synpr | Enriched in retina | -1.945 | 0.000001031 | 203.6 |
| ENSMUSG00000070687 | 5-hydroxytryptamine (serotonin) receptor 1D [Source:MGI Symbol;Acc:MGI:96276] | Htr1d | Enriched in retina | -1.947 | 0.00009045 | 19.28 |
| ENSMUSG00000042073 | abhydrolase domain containing 14b [Source:MGI Symbol;Acc:MGI:1923741] | Abhd14b | Enriched in retina | -1.964 | 0.001772 | 38.48 |
| ENSMUSG00000025141 | myeloid-associated differentiation marker-like 2 [Source:MGI Symbol;Acc:MGI:1915765] | Myadml2 | Enriched in retina | -1.967 | 0.01765 | 14.75 |
| ENSMUSG00000023495 | poly(rC) binding protein 4 [Source:MGI Symbol;Acc:MGI:1890471] | Pcbp4 | Enriched in retina | -1.972 | 4.332E-28 | 550 |
| ENSMUSG00000025432 | advillin [Source:MGI Symbol;Acc:MGI:1333798] | Avil | Enriched in retina | -1.992 | 0.03232 | 4.817 |
| ENSMUSG00000002980 | basal cell adhesion molecule [Source:MGI Symbol;Acc:MGI:1929940] | Bcam | Enriched in retina | -1.994 | 0.000008407 | 83.63 |
| ENSMUSG00000001504 | Iroquois homeobox 2 [Source:MGI Symbol;Acc:MGI:1197526] | Irx2 | Enriched in retina | -2.008 | 0.00006085 | 24.13 |
| ENSMUSG00000011256 | a disintegrin and metallopeptidase domain 19 (meltrin beta) [Source:MGI Symbol;Acc:MGI:105377] | Adam19 | Enriched in retina | -2.049 | 1.859E-11 | 194.7 |
| ENSMUSG00000016356 | collagen, type XX, alpha 1 [Source:MGI Symbol;Acc:MGI:1920618] | Col20a1 | Enriched in retina | -2.055 | 0.00402 | 25.93 |
| ENSMUSG00000002908 | potassium intermediate/small conductance calcium-activated channel, subfamily N, member 1 [Source:MGI Symbol;Acc:MGI:1933993] | Kcnn1 | Enriched in retina | -2.124 | 6.101E-12 | 72.47 |
| ENSMUSG00000090523 | glycophorin C [Source:MGI Symbol;Acc:MGI:1098566] | Gypc | Enriched in retina | -2.151 | 0.01885 | 5.436 |
| ENSMUSG00000024942 | calpain 1 [Source:MGI Symbol;Acc:MGI:88263] | Capn1 | Enriched in retina | -2.166 | 0.0001201 | 59.73 |
| ENSMUSG00000078958 | ATPase, H+ transporting, lysosomal accessory protein 1-like [Source:MGI Symbol;Acc:MGI:3648665] | Atp6ap1l | Enriched in retina | -2.292 | 0.03117 | 5.851 |
| ENSMUSG00000037992 | retinoic acid receptor, alpha [Source:MGI Symbol;Acc:MGI:97856] | Rara | Enriched in retina | -2.297 | 2.68E-23 | 152.2 |
| ENSMUSG00000030830 | integrin alpha L [Source:MGI Symbol;Acc:MGI:96606] | Itgal | Enriched in retina | -2.332 | 0.0009505 | 6.323 |

| | | | | | | |
|---|---|---|---|---|---|---|
| ENSMUSG00000030110 | ret proto-oncogene [Source:MGI Symbol;Acc:MGI:97902] | Ret | Enriched in retina | -2.429 | 1.204E-13 | 188.3 |
| ENSMUSG00000038530 | regulator of G-protein signaling 4 [Source:MGI Symbol;Acc:MGI:108409] | Rgs4 | Enriched in retina | -2.429 | 6.841E-09 | 578.5 |
| ENSMUSG00000016624 | PHD finger protein 21B [Source:MGI Symbol;Acc:MGI:2443812] | Phf21b | Enriched in retina | -2.462 | 7.107E-11 | 142.2 |
| ENSMUSG00000043391 | RIKEN cDNA 2510009E07 gene [Source:MGI Symbol;Acc:MGI:1919440] | 2510009E07Rik | Enriched in retina | -2.505 | 1.129E-47 | 739.1 |
| ENSMUSG00000026778 | protein kinase C, theta [Source:MGI Symbol;Acc:MGI:97601] | Prkcq | Enriched in retina | -2.593 | 9.768E-10 | 111.9 |
| ENSMUSG00000022054 | neurofilament, medium polypeptide [Source:MGI Symbol;Acc:MGI:97314] | Nefm | Enriched in retina | -2.597 | 1.189E-08 | 5610 |
| ENSMUSG00000019966 | kit ligand [Source:MGI Symbol;Acc:MGI:96974] | Kitl | Enriched in retina | -2.622 | 2.042E-11 | 453.1 |
| ENSMUSG00000031734 | Iroquois related homeobox 3 [Source:MGI Symbol;Acc:MGI:1197522] | Irx3 | Enriched in retina | -2.684 | 0.000004025 | 16.38 |
| ENSMUSG00000047963 | starch binding domain 1 [Source:MGI Symbol;Acc:MGI:1261768] | Stbd1 | Enriched in retina | -2.706 | 0.0002608 | 7.985 |
| ENSMUSG00000037705 | tectorin alpha [Source:MGI Symbol;Acc:MGI:109575] | Tecta | Enriched in retina | -2.778 | 0.003182 | 4.094 |
| ENSMUSG00000090223 | Purkinje cell protein 4 [Source:MGI Symbol;Acc:MGI:97509] | Pcp4 | Enriched in retina | -2.965 | 0.00004593 | 93.25 |
| ENSMUSG00000025366 | extended synaptotagmin-like protein 1 [Source:MGI Symbol;Acc:MGI:1344426] | Esyt1 | Enriched in retina | -3.141 | 8.22E-11 | 159.4 |
| ENSMUSG00000030270 | copine family member IX [Source:MGI Symbol;Acc:MGI:2443052] | Cpne9 | Enriched in retina | -3.236 | 0.000002371 | 175.2 |
| ENSMUSG00000034115 | sodium channel, voltage-gated, type XI, alpha [Source:MGI Symbol;Acc:MGI:1345149] | Scn11a | Enriched in retina | -3.336 | 0.001039 | 5.567 |
| ENSMUSG00000032128 | roundabout guidance receptor 3 [Source:MGI Symbol;Acc:MGI:1343102] | Robo3 | Enriched in retina | -3.43 | 0.0000639 | 9.898 |
| ENSMUSG00000020216 | junctional sarcoplasmic reticulum protein 1 [Source:MGI Symbol;Acc:MGI:1916700] | Jsrp1 | Enriched in retina | -3.707 | 0.008563 | 6.626 |
| ENSMUSG00000031737 | Iroquois homeobox 5 [Source:MGI Symbol;Acc:MGI:1859086] | Irx5 | Enriched in retina | -3.711 | 2.723E-14 | 31.09 |
| ENSMUSG00000020838 | solute carrier family 6 (neurotransmitter transporter, serotonin), member 4 [Source:MGI Symbol;Acc:MGI:96285] | Slc6a4 | Enriched in retina | -3.895 | 0.00465 | 23.26 |
| ENSMUSG00000031586 | RNA binding protein gene with multiple splicing [Source:MGI Symbol;Acc:MGI:1334446] | Rbpms | Enriched in retina | -3.906 | 2.9E-19 | 81.55 |
| ENSMUSG00000047746 | F-box protein 40 [Source:MGI Symbol;Acc:MGI:2443753] | Fbxo40 | Enriched in retina | -3.98 | 0.005825 | 15.08 |
| ENSMUSG00000032387 | RNA binding protein with multiple splicing 2 [Source:MGI Symbol;Acc:MGI:1919223] | Rbpms2 | Enriched in retina | -4.546 | 9.526E-21 | 47.72 |
| ENSMUSG00000025389 | major intrinsic protein of lens fiber [Source:MGI Symbol;Acc:MGI:96990] | Mip | Enriched in retina | -4.746 | 0.03401 | 36.23 |
| ENSMUSG00000068220 | lectin, galactose binding, soluble 1 [Source:MGI Symbol;Acc:MGI:96777] | Lgals1 | Enriched in retina | -4.892 | 1.624E-07 | 30.47 |
| ENSMUSG00000023484 | peripherin [Source:MGI Symbol;Acc:MGI:97774] | Prph | Enriched in retina | -5.01 | 2.293E-07 | 64.1 |
| ENSMUSG00000032446 | eomesodermin [Source:MGI Symbol;Acc:MGI:1201683] | Eomes | Enriched in retina | -5.064 | 2.581E-09 | 60.2 |
| ENSMUSG00000061048 | cadherin 3 [Source:MGI Symbol;Acc:MGI:88356] | Cdh3 | Enriched in retina | -5.318 | 0.0001371 | 8.894 |
| ENSMUSG00000048070 | phosphoinositide-interacting regulator of transient receptor potential channels [Source:MGI Symbol;Acc:MGI:2443635] | Pirt | Enriched in retina | -5.327 | 0.00001131 | 10.02 |
| ENSMUSG00000069372 | cortexin 3 [Source:MGI Symbol;Acc:MGI:3642816] | Ctxn3 | Enriched in retina | -5.642 | 2.964E-07 | 38.01 |
| ENSMUSG00000002100 | myosin binding protein C, cardiac [Source:MGI Symbol;Acc:MGI:102844] | Mybpc3 | Enriched in retina | -6.304 | 0.00001472 | 4.298 |
| ENSMUSG00000048108 | transmembrane protein 72 [Source:MGI Symbol;Acc:MGI:2442707] | Tmem72 | Enriched in retina | -6.96 | 0.00002895 | 209.8 |
| ENSMUSG00000067438 | H6 homeobox 1 [Source:MGI Symbol;Acc:MGI:107178] | Hmx1 | Enriched in retina | -7.122 | 2.903E-10 | 37.32 |
| ENSMUSG00000051279 | growth differentiation factor 6 [Source:MGI Symbol;Acc:MGI:95689] | Gdf6 | Enriched in retina | -7.502 | 7.933E-09 | 11.98 |
| ENSMUSG00000034777 | ventral anterior homeobox 2 [Source:MGI Symbol;Acc:MGI:1346018] | Vax2 | Enriched in retina | -7.876 | 8.107E-10 | 45.1 |
| ENSMUSG00000031965 | T-box 20 [Source:MGI Symbol;Acc:MGI:1888496] | Tbx20 | Enriched in retina | -8.258 | 1.079E-09 | 45.45 |
| ENSMUSG00000031688 | POU domain, class 4, transcription factor 2 [Source:MGI Symbol;Acc:MGI:102524] | Pou4f2 | Enriched in retina | -9.365 | 9.404E-13 | 45.2 |
| ENSMUSG00000031738 | Iroquois homeobox 6 [Source:MGI Symbol;Acc:MGI:1927642] | Irx6 | Enriched in retina | -9.591 | 1.014E-13 | 50.97 |

**Contrast numerator: P15_Het_dLGN. Contrast denominator: P15_Het_retina.**

| ensembl_gene_id | description | mgi_symbol | Significance | deseq_logfc | deseq_adjp | deseq_basemean |
|---|---|---|---|---|---|---|
| ENSMUSG00000078816 | protein kinase C, gamma [Source:MGI Symbol;Acc:MGI:97597] | Prkcg | Enriched in dLGN | 8.153 | 9.37E-107 | 571.8 |
| ENSMUSG00000030785 | cytochrome c oxidase subunit 6A2 [Source:MGI Symbol;Acc:MGI:104649] | Cox6a2 | Enriched in dLGN | 6.954 | 1.569E-15 | 11.74 |
| ENSMUSG00000031994 | a disintegrin-like and metallopeptidase (reprolysin type) with thrombospondin type 1 motif, 8 [Source:MGI Symbol;Acc:MGI:1353468] | Adamts8 | Enriched in dLGN | 6.94 | 1.814E-30 | 52.93 |
| ENSMUSG00000032532 | cholecystokinin [Source:MGI Symbol;Acc:MGI:88297] | Cck | Enriched in dLGN | 6.905 | 1.035E-46 | 445.5 |
| ENSMUSG00000021948 | protein kinase C, delta [Source:MGI Symbol;Acc:MGI:97598] | Prkcd | Enriched in dLGN | 6.493 | 2.771E-48 | 727.9 |
| ENSMUSG00000055027 | SET and MYND domain containing 1 [Source:MGI Symbol;Acc:MGI:104790] | Smyd1 | Enriched in dLGN | 6.239 | 4.184E-36 | 65.99 |
| ENSMUSG00000022468 | endonuclease, polyU-specific [Source:MGI Symbol;Acc:MGI:97746] | Endou | Enriched in dLGN | 5.984 | 8.974E-26 | 34.93 |
| ENSMUSG00000029211 | gamma-aminobutyric acid (GABA) A receptor, subunit alpha 4 [Source:MGI Symbol;Acc:MGI:95616] | Gabra4 | Enriched in dLGN | 5.728 | 1.961E-66 | 154.6 |
| ENSMUSG00000000296 | tumor protein D52-like 1 [Source:MGI Symbol;Acc:MGI:1298386] | Tpd52l1 | Enriched in dLGN | 4.913 | 1.001E-45 | 61.04 |
| ENSMUSG00000050541 | adrenergic receptor, alpha 1b [Source:MGI Symbol;Acc:MGI:104774] | Adra1b | Enriched in dLGN | 4.889 | 2.678E-42 | 73.77 |
| ENSMUSG00000064179 | troponin T1, skeletal, slow [Source:MGI Symbol;Acc:MGI:1333868] | Tnnt1 | Enriched in dLGN | 4.844 | 5.971E-36 | 308.7 |
| ENSMUSG00000094132 | predicted gene 3264 [Source:MGI Symbol;Acc:MGI:3781442] | Gm3264 | Enriched in dLGN | 4.839 | 0.0008726 | 2.94 |
| ENSMUSG00000032776 | multiple C2 domains, transmembrane 2 [Source:MGI Symbol;Acc:MGI:2685335] | Mctp2 | Enriched in dLGN | 4.821 | 1.574E-21 | 28.35 |
| ENSMUSG00000026525 | opsin 3 [Source:MGI Symbol;Acc:MGI:1338022] | Opn3 | Enriched in dLGN | 4.78 | 1.941E-24 | 17.62 |
| ENSMUSG00000020733 | solute carrier family 9 (sodium/hydrogen exchanger), member 3 regulator 1 [Source:MGI Symbol;Acc:MGI:1349482] | Slc9a3r1 | Enriched in dLGN | 4.53 | 1.686E-88 | 214.9 |
| ENSMUSG00000054196 | collagen triple helix repeat containing 1 [Source:MGI Symbol;Acc:MGI:1915838] | Cthrc1 | Enriched in dLGN | 4.516 | 1.419E-30 | 38 |
| ENSMUSG00000030340 | sodium channel, nonvoltage-gated 1 alpha [Source:MGI Symbol;Acc:MGI:101782] | Scnn1a | Enriched in dLGN | 4.482 | 2.947E-32 | 44.62 |
| ENSMUSG00000027360 | histidine decarboxylase [Source:MGI Symbol;Acc:MGI:96062] | Hdc | Enriched in dLGN | 4.435 | 2.63E-20 | 44.93 |
| ENSMUSG00000060780 | leucine rich repeat transmembrane neuronal 1 [Source:MGI Symbol;Acc:MGI:2389173] | Lrrtm1 | Enriched in dLGN | 4.313 | 3.033E-94 | 250 |
| ENSMUSG00000028082 | SH3 domain protein D19 [Source:MGI Symbol;Acc:MGI:1350923] | Sh3d19 | Enriched in dLGN | 4.302 | 7.096E-42 | 657.1 |
| ENSMUSG00000030854 | protein tyrosine phosphatase, non-receptor type 5 [Source:MGI Symbol;Acc:MGI:97807] | Ptpn5 | Enriched in dLGN | 4.251 | 7.903E-96 | 407 |
| ENSMUSG00000048616 | noggin [Source:MGI Symbol;Acc:MGI:104327] | Nog | Enriched in dLGN | 4.238 | 3.729E-30 | 53.91 |
| ENSMUSG00000017417 | plexin domain containing 1 [Source:MGI Symbol;Acc:MGI:1919574] | Plxdc1 | Enriched in dLGN | 4.234 | 1.496E-45 | 77.27 |
| ENSMUSG00000037428 | VGF nerve growth factor inducible [Source:MGI Symbol;Acc:MGI:1343180] | Vgf | Enriched in dLGN | 4.209 | 6.12E-56 | 1249 |
| ENSMUSG00000027347 | RAS guanyl releasing protein 1 [Source:MGI Symbol;Acc:MGI:1314635] | Rasgrp1 | Enriched in dLGN | 4.208 | 2.759E-32 | 353.4 |
| ENSMUSG00000056258 | potassium voltage-gated channel, subfamily Q, member 3 [Source:MGI Symbol;Acc:MGI:1336181] | Kcnq3 | Enriched in dLGN | 4.204 | 5.249E-50 | 1087 |
| ENSMUSG00000032355 | muscular LMNA-interacting protein [Source:MGI Symbol;Acc:MGI:1916892] | Mlip | Enriched in dLGN | 4.169 | 3.84E-15 | 22.7 |
| ENSMUSG00000038128 | calcium/calmodulin-dependent protein kinase IV [Source:MGI Symbol;Acc:MGI:88258] | Camk4 | Enriched in dLGN | 4.163 | 1.402E-75 | 937 |
| ENSMUSG00000036972 | zinc finger protein of the cerebellum 4 [Source:MGI Symbol;Acc:MGI:107201] | Zic4 | Enriched in dLGN | 4.145 | 1.111E-30 | 142.7 |
| ENSMUSG00000035506 | solute carrier family 12 (potassium/chloride transporters), member 8 [Source:MGI Symbol;Acc:MGI:2443672] | Slc12a8 | Enriched in dLGN | 4.08 | 2.238E-12 | 17.24 |
| ENSMUSG00000040624 | pleckstrin homology domain containing, family G (with RhoGef domain) member 1 [Source:MGI Symbol;Acc:MGI:2676551] | Plekhg1 | Enriched in dLGN | 3.985 | 8.967E-58 | 465 |
| ENSMUSG00000096257 | coiled-coil glutamate-rich protein 2 [Source:MGI Symbol;Acc:MGI:3645242] | Ccer2 | Enriched in dLGN | 3.975 | 2.093E-14 | 12.8 |
| ENSMUSG00000004892 | brevican [Source:MGI Symbol;Acc:MGI:1096385] | Bcan | Enriched in dLGN | 3.967 | 3.507E-51 | 546.7 |
| ENSMUSG00000024727 | transient receptor potential cation channel, subfamily M, member 6 [Source:MGI Symbol;Acc:MGI:2675603] | Trpm6 | Enriched in dLGN | 3.928 | 5.756E-10 | 19.86 |
| ENSMUSG00000068696 | G-protein coupled receptor 88 [Source:MGI Symbol;Acc:MGI:1927653] | Gpr88 | Enriched in dLGN | 3.908 | 2.645E-44 | 66.64 |
| ENSMUSG00000045731 | prepronociceptin [Source:MGI Symbol;Acc:MGI:105308] | Pnoc | Enriched in dLGN | 3.899 | 6.719E-27 | 50.56 |
| ENSMUSG00000021209 | protein phosphatase 4, regulatory subunit 4 [Source:MGI Symbol;Acc:MGI:1921771] | Ppp4r4 | Enriched in dLGN | 3.889 | 1.34E-52 | 139.4 |
| ENSMUSG00000003665 | hyaluronan synthase 1 [Source:MGI Symbol;Acc:MGI:106590] | Has1 | Enriched in dLGN | 3.83 | 0.001831 | 2.393 |
| ENSMUSG00000042353 | Fras1 related extracellular matrix protein 3 [Source:MGI Symbol;Acc:MGI:2685641] | Frem3 | Enriched in dLGN | 3.825 | 1.018E-24 | 57.69 |
| ENSMUSG00000066129 | kinase non-catalytic C-lobe domain (KIND) containing 1 [Source:MGI Symbol;Acc:MGI:1923734] | Kndc1 | Enriched in dLGN | 3.817 | 7.12E-65 | 714.5 |
| ENSMUSG00000029054 | gamma-aminobutyric acid (GABA) A receptor, subunit delta [Source:MGI Symbol;Acc:MGI:95622] | Gabrd | Enriched in dLGN | 3.791 | 1.29E-20 | 52.02 |
| ENSMUSG00000042514 | kelch-like 14 [Source:MGI Symbol;Acc:MGI:1921249] | Klhl14 | Enriched in dLGN | 3.648 | 4.709E-07 | 9.277 |
| ENSMUSG00000040723 | RCSD domain containing 1 [Source:MGI Symbol;Acc:MGI:2676394] | Rcsd1 | Enriched in dLGN | 3.619 | 7.351E-17 | 27.5 |
| ENSMUSG00000067860 | zinc finger protein of the cerebellum 3 [Source:MGI Symbol;Acc:MGI:106676] | Zic3 | Enriched in dLGN | 3.588 | 1.294E-20 | 26.92 |
| ENSMUSG00000051022 | heparan sulfate (glucosamine) 3-O-sulfotransferase 1 [Source:MGI Symbol;Acc:MGI:1201606] | Hs3st1 | Enriched in dLGN | 3.574 | 3.454E-46 | 132 |
| ENSMUSG00000027489 | N-terminal EF-hand calcium binding protein 3 [Source:MGI Symbol;Acc:MGI:1861721] | Necab3 | Enriched in dLGN | 3.565 | 7.003E-29 | 36.72 |
| ENSMUSG00000020059 | synaptonemal complex protein 3 [Source:MGI Symbol;Acc:MGI:109542] | Sycp3 | Enriched in dLGN | 3.494 | 0.001209 | 2.11 |
| ENSMUSG00000068373 | RIKEN cDNA D430041D05 gene [Source:MGI Symbol;Acc:MGI:2181743] | D430041D05Rik | Enriched in dLGN | 3.457 | 1.891E-86 | 780.2 |
| ENSMUSG00000030075 | contactin 3 [Source:MGI Symbol;Acc:MGI:99534] | Cntn3 | Enriched in dLGN | 3.45 | 1.802E-23 | 116.4 |
| ENSMUSG00000021259 | cytochrome P450, family 46, subfamily a, polypeptide 1 [Source:MGI Symbol;Acc:MGI:1341877] | Cyp46a1 | Enriched in dLGN | 3.421 | 1.132E-63 | 248.2 |
| ENSMUSG00000001763 | tetraspanin 33 [Source:MGI Symbol;Acc:MGI:1919012] | Tspan33 | Enriched in dLGN | 3.42 | 5.688E-37 | 62.53 |
| ENSMUSG00000049176 | FERM and PDZ domain containing 4 [Source:MGI Symbol;Acc:MGI:3042378] | Frmpd4 | Enriched in dLGN | 3.403 | 3.747E-26 | 381.5 |
| ENSMUSG00000004849 | adaptor protein complex AP-1, sigma 1 [Source:MGI Symbol;Acc:MGI:1098244] | Ap1s1 | Enriched in dLGN | 3.392 | 1.213E-44 | 288.6 |
| ENSMUSG00000052026 | solute carrier family 6 (neurotransmitter transporter, L-proline), member 7 [Source:MGI Symbol;Acc:MGI:2147363] | Slc6a7 | Enriched in dLGN | 3.364 | 1.796E-49 | 217.9 |
| ENSMUSG00000034324 | transmembrane protein 132C [Source:MGI Symbol;Acc:MGI:2443061] | Tmem132c | Enriched in dLGN | 3.338 | 9.46E-41 | 360 |
| ENSMUSG00000046480 | sodium channel, type IV, beta [Source:MGI Symbol;Acc:MGI:2687406] | Scn4b | Enriched in dLGN | 3.302 | 1.433E-13 | 167.6 |
| ENSMUSG00000033044 | dehydrogenase/reductase (SDR family) member 7C [Source:MGI Symbol;Acc:MGI:1915710] | Dhrs7c | Enriched in dLGN | 3.244 | 0.0007891 | 6.079 |
| ENSMUSG00000020838 | solute carrier family 6 (neurotransmitter transporter, serotonin), member 4 [Source:MGI Symbol;Acc:MGI:96285] | Slc6a4 | Enriched in dLGN | 3.234 | 0.0006455 | 23.26 |
| ENSMUSG00000039783 | kynurenine 3-monooxygenase (kynurenine 3-hydroxylase) [Source:MGI Symbol;Acc:MGI:2138151] | Kmo | Enriched in dLGN | 3.231 | 6.148E-08 | 4.13 |
| ENSMUSG00000028833 | neurochondrin [Source:MGI Symbol;Acc:MGI:1347351] | Ncdn | Enriched in dLGN | 3.206 | 1.047E-31 | 1887 |
| ENSMUSG00000020732 | RAB37, member RAS oncogene family [Source:MGI Symbol;Acc:MGI:1929945] | Rab37 | Enriched in dLGN | 3.185 | 6.937E-12 | 50.93 |
| ENSMUSG00000027965 | olfactomedin 3 [Source:MGI Symbol;Acc:MGI:2387329] | Olfm3 | Enriched in dLGN | 3.183 | 8.697E-22 | 175.9 |
| ENSMUSG00000019828 | glutamate receptor, metabotropic 1 [Source:MGI Symbol;Acc:MGI:1351338] | Grm1 | Enriched in dLGN | 3.171 | 1.426E-25 | 572.7 |
| ENSMUSG00000018821 | arginine vasopressin-induced 1 [Source:MGI Symbol;Acc:MGI:1916784] | Avpi1 | Enriched in dLGN | 3.159 | 4.07E-23 | 34.89 |
| ENSMUSG00000019762 | iodotyrosine deiodinase [Source:MGI Symbol;Acc:MGI:1917587] | Iyd | Enriched in dLGN | 3.154 | 0.00001218 | 6.399 |
| ENSMUSG00000022454 | NEL-like 2 [Source:MGI Symbol;Acc:MGI:1858510] | Nell2 | Enriched in dLGN | 3.135 | 9.444E-39 | 1350 |
| ENSMUSG00000022494 | shisa family member 9 [Source:MGI Symbol;Acc:MGI:1919805] | Shisa9 | Enriched in dLGN | 3.13 | 6.824E-21 | 606.7 |
| ENSMUSG00000040794 | C1q and tumor necrosis factor related protein 4 [Source:MGI Symbol;Acc:MGI:1914695] | C1qtnf4 | Enriched in dLGN | 3.128 | 6.739E-41 | 207.8 |
| ENSMUSG00000034825 | nuclear receptor interacting protein 3 [Source:MGI Symbol;Acc:MGI:1925843] | Nrip3 | Enriched in dLGN | 3.116 | 8.488E-23 | 285.5 |
| ENSMUSG00000032503 | cyclic AMP-regulated phosphoprotein, 21 [Source:MGI Symbol;Acc:MGI:107562] | Arpp21 | Enriched in dLGN | 3.11 | 4.445E-18 | 594.6 |
| ENSMUSG00000055409 | NEL-like 1 [Source:MGI Symbol;Acc:MGI:2443902] | Nell1 | Enriched in dLGN | 3.11 | 1.225E-16 | 543.3 |
| ENSMUSG00000050315 | synaptopodin 2 [Source:MGI Symbol;Acc:MGI:2153070] | Synpo2 | Enriched in dLGN | 3.093 | 4.067E-27 | 581.4 |
| ENSMUSG00000020644 | inhibitor of DNA binding 2 [Source:MGI Symbol;Acc:MGI:96397] | Id2 | Enriched in dLGN | 3.093 | 1.365E-15 | 36.74 |
| ENSMUSG00000026688 | microsomal glutathione S-transferase 3 [Source:MGI Symbol;Acc:MGI:1913697] | Mgst3 | Enriched in dLGN | 3.025 | 3.547E-11 | 88.46 |
| ENSMUSG00000026235 | Eph receptor A4 [Source:MGI Symbol;Acc:MGI:98277] | Epha4 | Enriched in dLGN | 2.979 | 1.905E-32 | 500.9 |
| ENSMUSG00000063434 | sortilin-related VPS10 domain containing receptor 3 [Source:MGI Symbol;Acc:MGI:1913923] | Sorcs3 | Enriched in dLGN | 2.979 | 8.332E-19 | 332.9 |
| ENSMUSG00000024529 | lysyl oxidase [Source:MGI Symbol;Acc:MGI:96817] | Lox | Enriched in dLGN | 2.977 | 9.79E-08 | 9.017 |
| ENSMUSG00000079396 | predicted gene 3411 [Source:MGI Symbol;Acc:MGI:3781589] | Gm3411 | Enriched in dLGN | 2.964 | 0.0001005 | 6.451 |
| ENSMUSG00000050830 | von Willebrand factor C domain containing 2 [Source:MGI Symbol;Acc:MGI:2442987] | Vwc2 | Enriched in dLGN | 2.939 | 3.547E-22 | 127 |
| ENSMUSG00000040713 | cellular repressor of E1A-stimulated genes 1 [Source:MGI Symbol;Acc:MGI:1344382] | Creg1 | Enriched in dLGN | 2.892 | 3.11E-24 | 76.45 |
| ENSMUSG00000024109 | neurexin I [Source:MGI Symbol;Acc:MGI:1096391] | Nrxn1 | Enriched in dLGN | 2.888 | 4.796E-25 | 3720 |
| ENSMUSG00000056306 | serine rich and transmembrane domain containing 1 [Source:MGI Symbol;Acc:MGI:3607715] | Sertm1 | Enriched in dLGN | 2.872 | 3.267E-21 | 70.23 |
| ENSMUSG00000040612 | immunoglobulin-like domain containing receptor 2 [Source:MGI Symbol;Acc:MGI:1196370] | Ildr2 | Enriched in dLGN | 2.827 | 1.745E-24 | 1307 |
| ENSMUSG00000059857 | netrin G1 [Source:MGI Symbol;Acc:MGI:1934028] | Ntng1 | Enriched in dLGN | 2.813 | 1.512E-13 | 904 |
| ENSMUSG00000004951 | heat shock protein 1 [Source:MGI Symbol;Acc:MGI:96240] | Hspb1 | Enriched in dLGN | 2.806 | 0.000001949 | 22.09 |
| ENSMUSG00000073406 | histocompatibility 2, blastocyst [Source:MGI Symbol;Acc:MGI:892004] | H2-Bl | Enriched in dLGN | 2.806 | 0.00009447 | 18.45 |
| ENSMUSG00000021373 | CAP, adenylate cyclase-associated protein, 2 (yeast) [Source:MGI Symbol;Acc:MGI:1914502] | Cap2 | Enriched in dLGN | 2.804 | 1.059E-28 | 147 |
| ENSMUSG00000058927 | predicted gene 10053 [Source:MGI Symbol;Acc:MGI:3704493] | Gm10053 | Enriched in dLGN | 2.761 | 2.793E-09 | 16.38 |
| ENSMUSG00000026824 | potassium inwardly-rectifying channel, subfamily J, member 3 [Source:MGI Symbol;Acc:MGI:104742] | Kcnj3 | Enriched in dLGN | 2.752 | 4.349E-21 | 206.8 |
| ENSMUSG00000030730 | ATPase, Ca++ transporting, cardiac muscle, fast twitch 1 [Source:MGI Symbol;Acc:MGI:105058] | Atp2a1 | Enriched in dLGN | 2.723 | 0.00003638 | 39.3 |
| ENSMUSG00000033685 | uncoupling protein 2 (mitochondrial, proton carrier) [Source:MGI Symbol;Acc:MGI:109354] | Ucp2 | Enriched in dLGN | 2.711 | 6.716E-16 | 149.6 |
| ENSMUSG00000061859 | PATJ, crumbs cell polarity complex component [Source:MGI Symbol;Acc:MGI:1277960] | Patj | Enriched in dLGN | 2.706 | 1.162E-14 | 609.6 |
| ENSMUSG00000046182 | GSG1-like [Source:MGI Symbol;Acc:MGI:2685483] | Gsg1l | Enriched in dLGN | 2.685 | 2.452E-34 | 471.7 |
| ENSMUSG00000027200 | sema domain, transmembrane domain (TM), and cytoplasmic domain, (semaphorin) 6D [Source:MGI Symbol;Acc:MGI:2387661] | Sema6d | Enriched in dLGN | 2.68 | 1.556E-22 | 731.9 |
| ENSMUSG00000110711 | predicted gene 45760 [Source:MGI Symbol;Acc:MGI:5804875] | Gm45760 | Enriched in dLGN | 2.67 | 0.0006678 | 2.941 |
| ENSMUSG00000046834 | keratin 1 [Source:MGI Symbol;Acc:MGI:96698] | Krt1 | Enriched in dLGN | 2.664 | 4.01E-12 | 16.62 |
| ENSMUSG00000058589 | ankyrin repeat and sterile alpha motif domain containing 1B [Source:MGI Symbol;Acc:MGI:1924781] | Anks1b | Enriched in dLGN | 2.658 | 1.485E-17 | 1182 |
| ENSMUSG00000029153 | OCIA domain containing 2 [Source:MGI Symbol;Acc:MGI:1916377] | Ociad2 | Enriched in dLGN | 2.655 | 1.849E-21 | 70.58 |
| ENSMUSG00000033316 | polypeptide N-acetylgalactosaminyltransferase 9 [Source:MGI Symbol;Acc:MGI:2677965] | Galnt9 | Enriched in dLGN | 2.649 | 5.278E-32 | 204.6 |
| ENSMUSG00000060402 | carbohydrate sulfotransferase 8 [Source:MGI Symbol;Acc:MGI:1916197] | Chst8 | Enriched in dLGN | 2.639 | 3.574E-17 | 67.64 |
| ENSMUSG00000033615 | complexin 1 [Source:MGI Symbol;Acc:MGI:104727] | Cplx1 | Enriched in dLGN | 2.62 | 1.277E-15 | 2304 |
| ENSMUSG00000005357 | solute carrier family 1 (high affinity aspartate/glutamate transporter), member 6 [Source:MGI Symbol;Acc:MGI:1096331] | Slc1a6 | Enriched in dLGN | 2.619 | 6.514E-16 | 52.8 |
| ENSMUSG00000035849 | keratin 222 [Source:MGI Symbol;Acc:MGI:2442728] | Krt222 | Enriched in dLGN | 2.61 | 4.5E-20 | 127 |
| ENSMUSG00000056413 | ArfGAP with dual PH domains 1 [Source:MGI Symbol;Acc:MGI:2442201] | Adap1 | Enriched in dLGN | 2.605 | 1.332E-56 | 552.2 |
| ENSMUSG00000029657 | heat shock 105kDa/110kDa protein 1 [Source:MGI Symbol;Acc:MGI:105053] | Hsph1 | Enriched in dLGN | 2.604 | 2.027E-23 | 555.2 |
| ENSMUSG00000065968 | interferon induced transmembrane protein 7 [Source:MGI Symbol;Acc:MGI:1921732] | Ifitm7 | Enriched in dLGN | 2.603 | 0.000000247 | 15.26 |
| ENSMUSG00000020589 | CYFIP related Rac1 interactor A [Source:MGI Symbol;Acc:MGI:1261783] | Cyria | Enriched in dLGN | 2.599 | 2.664E-30 | 283.7 |
| ENSMUSG00000023905 | tumor necrosis factor receptor superfamily, member 12a [Source:MGI Symbol;Acc:MGI:1351484] | Tnfrsf12a | Enriched in dLGN | 2.599 | 9.074E-08 | 7.873 |
| ENSMUSG00000074736 | synapse differentiation inducing 1 [Source:MGI Symbol;Acc:MGI:3702158] | Syndig1 | Enriched in dLGN | 2.596 | 1.432E-12 | 50.55 |
| ENSMUSG00000037568 | vasohibin 2 [Source:MGI Symbol;Acc:MGI:2444826] | Vash2 | Enriched in dLGN | 2.595 | 2.267E-23 | 93.06 |
| ENSMUSG00000024524 | guanine nucleotide binding protein, alpha stimulating, olfactory type [Source:MGI Symbol;Acc:MGI:95774] | Gnal | Enriched in dLGN | 2.594 | 4.023E-41 | 618.2 |
| ENSMUSG00000039115 | integrin alpha 9 [Source:MGI Symbol;Acc:MGI:104756] | Itga9 | Enriched in dLGN | 2.567 | 0.001604 | 1811 |
| ENSMUSG00000038244 | microtubule associated monooxygenase, calponin and LIM domain containing 2 [Source:MGI Symbol;Acc:MGI:2444947] | Mical2 | Enriched in dLGN | 2.56 | 1.98E-19 | 255.1 |
| ENSMUSG00000004187 | kinesin family member C2 [Source:MGI Symbol;Acc:MGI:109187] | Kifc2 | Enriched in dLGN | 2.538 | 2.341E-52 | 769 |
| ENSMUSG00000028434 | erythrocyte membrane protein band 4.1 like 4b [Source:MGI Symbol;Acc:MGI:1859149] | Epb41l4b | Enriched in dLGN | 2.529 | 1.274E-29 | 333.4 |
| ENSMUSG00000027076 | translocase of inner mitochondrial membrane 10 [Source:MGI Symbol;Acc:MGI:1353429] | Timm10 | Enriched in dLGN | 2.524 | 1.944E-13 | 29.72 |
| ENSMUSG00000028024 | glutamyl aminopeptidase [Source:MGI Symbol;Acc:MGI:106645] | Enpep | Enriched in dLGN | 2.502 | 8.913E-07 | 10.7 |
| ENSMUSG00000045733 | shadow of prion protein [Source:MGI Symbol;Acc:MGI:3582583] | Sprn | Enriched in dLGN | 2.497 | 5.424E-37 | 239 |
| ENSMUSG00000042567 | NIMA (never in mitosis gene a)- related kinase 10 [Source:MGI Symbol;Acc:MGI:2685128] | Nek10 | Enriched in dLGN | 2.495 | 0.001273 | 47 |
| ENSMUSG00000007594 | hyaluronan and proteoglycan link protein 4 [Source:MGI Symbol;Acc:MGI:2679531] | Hapln4 | Enriched in dLGN | 2.492 | 4.428E-08 | 156.8 |
| ENSMUSG00000028680 | polo like kinase 3 [Source:MGI Symbol;Acc:MGI:109604] | Plk3 | Enriched in dLGN | 2.474 | 7.31E-18 | 41.32 |
| ENSMUSG00000031760 | metallothionein 3 [Source:MGI Symbol;Acc:MGI:97173] | Mt3 | Enriched in dLGN | 2.473 | 4.666E-15 | 131.4 |
| ENSMUSG00000026888 | growth factor receptor bound protein 14 [Source:MGI Symbol;Acc:MGI:1355324] | Grb14 | Enriched in dLGN | 2.468 | 9.888E-14 | 102.4 |
| ENSMUSG00000030089 | solute carrier family 41, member 3 [Source:MGI Symbol;Acc:MGI:1918949] | Slc41a3 | Enriched in dLGN | 2.461 | 2.989E-14 | 52.15 |
| ENSMUSG00000026616 | complement receptor 2 [Source:MGI Symbol;Acc:MGI:88489] | Cr2 | Enriched in dLGN | 2.447 | 0.006835 | 3.229 |
| ENSMUSG00000007653 | gamma-aminobutyric acid (GABA) A receptor, subunit beta 2 [Source:MGI Symbol;Acc:MGI:95620] | Gabrb2 | Enriched in dLGN | 2.439 | 3.56E-13 | 322.6 |

| | | | | | | |
|---|---|---|---|---|---|---|
| ENSMUSG00000056222 | sparc/osteonectin, cwcv and kazal-like domains proteoglycan 1 [Source:MGI Symbol;Acc:MGI:105371] | Spock1 | Enriched in dLGN | 2.418 | 2.712E-16 | 1204 |
| ENSMUSG00000034226 | ras homolog family member V [Source:MGI Symbol;Acc:MGI:2444227] | Rhov | Enriched in dLGN | 2.407 | 4.564E-12 | 54.65 |
| ENSMUSG00000025468 | calcyon neuron-specific vesicular protein [Source:MGI Symbol;Acc:MGI:1915816] | Caly | Enriched in dLGN | 2.405 | 1.478E-16 | 238.8 |
| ENSMUSG00000028718 | Scl/Tal1 interrupting locus [Source:MGI Symbol;Acc:MGI:107477] | Stil | Enriched in dLGN | 2.405 | 0.0002671 | 5.074 |
| ENSMUSG00000032965 | intraflagellar transport 57 [Source:MGI Symbol;Acc:MGI:1921166] | Ift57 | Enriched in dLGN | 2.401 | 2.363E-35 | 254.5 |
| ENSMUSG00000070803 | Cbp/p300-interacting transactivator, with Glu/Asp-rich carboxy-terminal domain, 4 [Source:MGI Symbol;Acc:MGI:1861694] | Cited4 | Enriched in dLGN | 2.374 | 0.00002553 | 9.191 |
| ENSMUSG00000041112 | engulfment and cell motility 1 [Source:MGI Symbol;Acc:MGI:2153044] | Elmo1 | Enriched in dLGN | 2.369 | 1.147E-15 | 598.8 |
| ENSMUSG00000062937 | methylthioadenosine phosphorylase [Source:MGI Symbol;Acc:MGI:1914152] | Mtap | Enriched in dLGN | 2.358 | 4.94E-10 | 29.38 |
| ENSMUSG00000030889 | von Willebrand factor A domain containing 3A [Source:MGI Symbol;Acc:MGI:3041229] | Vwa3a | Enriched in dLGN | 2.343 | 0.0005699 | 26.65 |
| ENSMUSG00000041670 | regulating synaptic membrane exocytosis 1 [Source:MGI Symbol;Acc:MGI:2152971] | Rims1 | Enriched in dLGN | 2.336 | 3.789E-23 | 1881 |
| ENSMUSG00000019943 | ATPase, Ca++ transporting, plasma membrane 1 [Source:MGI Symbol;Acc:MGI:104653] | Atp2b1 | Enriched in dLGN | 2.316 | 7.338E-43 | 2706 |
| ENSMUSG00000031137 | fibroblast growth factor 13 [Source:MGI Symbol;Acc:MGI:109178] | Fgf13 | Enriched in dLGN | 2.315 | 1.118E-16 | 419.6 |
| ENSMUSG00000041020 | MAP7 domain containing 2 [Source:MGI Symbol;Acc:MGI:1917474] | Map7d2 | Enriched in dLGN | 2.314 | 3.036E-23 | 736.5 |
| ENSMUSG00000031778 | chemokine (C-X3-C motif) ligand 1 [Source:MGI Symbol;Acc:MGI:1097153] | Cx3cl1 | Enriched in dLGN | 2.314 | 4.394E-14 | 522 |
| ENSMUSG00000028367 | thioredoxin 1 [Source:MGI Symbol;Acc:MGI:98874] | Txn1 | Enriched in dLGN | 2.313 | 2.958E-16 | 92.9 |
| ENSMUSG00000001227 | sema domain, transmembrane domain (TM), and cytoplasmic domain, (semaphorin) 6B [Source:MGI Symbol;Acc:MGI:1202889] | Sema6b | Enriched in dLGN | 2.3 | 4.206E-35 | 529.4 |
| ENSMUSG00000036564 | N-myc downstream regulated gene 4 [Source:MGI Symbol;Acc:MGI:2384590] | Ndrg4 | Enriched in dLGN | 2.298 | 2.589E-22 | 2611 |
| ENSMUSG00000021591 | glutaredoxin [Source:MGI Symbol;Acc:MGI:2135625] | Glrx | Enriched in dLGN | 2.292 | 1.195E-18 | 37.82 |
| ENSMUSG00000027716 | transient receptor potential cation channel, subfamily C, member 3 [Source:MGI Symbol;Acc:MGI:109526] | Trpc3 | Enriched in dLGN | 2.278 | 4.853E-19 | 212.4 |
| ENSMUSG00000031438 | ring finger protein 128 [Source:MGI Symbol;Acc:MGI:1914139] | Rnf128 | Enriched in dLGN | 2.276 | 0.000001428 | 18.97 |
| ENSMUSG00000029467 | ATPase, Ca++ transporting, cardiac muscle, slow twitch 2 [Source:MGI Symbol;Acc:MGI:88110] | Atp2a2 | Enriched in dLGN | 2.271 | 2.813E-31 | 2995 |
| ENSMUSG00000041263 | RUN and SH3 domain containing 1 [Source:MGI Symbol;Acc:MGI:1919546] | Rusc1 | Enriched in dLGN | 2.271 | 8.513E-19 | 448.9 |
| ENSMUSG00000034570 | inositol polyphosphate 5-phosphatase J [Source:MGI Symbol;Acc:MGI:2158663] | Inpp5j | Enriched in dLGN | 2.271 | 3.162E-14 | 112.1 |
| ENSMUSG00000020990 | cyclin-dependent kinase-like 1 (CDC2-related kinase) [Source:MGI Symbol;Acc:MGI:1918341] | Cdkl1 | Enriched in dLGN | 2.266 | 1.293E-12 | 53.67 |
| ENSMUSG00000024190 | dual specificity phosphatase 1 [Source:MGI Symbol;Acc:MGI:105120] | Dusp1 | Enriched in dLGN | 2.265 | 2.25E-26 | 44.73 |
| ENSMUSG00000044576 | GRB2 associated regulator of MAPK1 subtype 2 [Source:MGI Symbol;Acc:MGI:2685290] | Garem2 | Enriched in dLGN | 2.256 | 2.832E-12 | 62.67 |
| ENSMUSG00000044835 | ankyrin repeat domain 45 [Source:MGI Symbol;Acc:MGI:1921094] | Ankrd45 | Enriched in dLGN | 2.253 | 7.939E-17 | 83.71 |
| ENSMUSG00000052372 | interleukin 1 receptor accessory protein-like 1 [Source:MGI Symbol;Acc:MGI:2687319] | Il1rapl1 | Enriched in dLGN | 2.246 | 0.00006604 | 1897 |
| ENSMUSG00000027577 | cholinergic receptor, nicotinic, alpha polypeptide 4 [Source:MGI Symbol;Acc:MGI:87888] | Chrna4 | Enriched in dLGN | 2.244 | 4.769E-17 | 546.3 |
| ENSMUSG00000030350 | protein arginine N-methyltransferase 8 [Source:MGI Symbol;Acc:MGI:3043083] | Prmt8 | Enriched in dLGN | 2.241 | 1.141E-24 | 303.1 |
| ENSMUSG00000034353 | receptor (calcitonin) activity modifying protein 1 [Source:MGI Symbol;Acc:MGI:1858418] | Ramp1 | Enriched in dLGN | 2.241 | 1.942E-09 | 54.71 |
| ENSMUSG00000022376 | adenylate cyclase 8 [Source:MGI Symbol;Acc:MGI:1341110] | Adcy8 | Enriched in dLGN | 2.24 | 8.77E-14 | 297.1 |
| ENSMUSG00000020331 | hyperpolarization-activated, cyclic nucleotide-gated K+ 2 [Source:MGI Symbol;Acc:MGI:1298210] | Hcn2 | Enriched in dLGN | 2.239 | 2.229E-30 | 598.2 |
| ENSMUSG00000004031 | bone morphogenic protein/retinoic acid inducible neural-specific 2 [Source:MGI Symbol;Acc:MGI:2443333] | Brinp2 | Enriched in dLGN | 2.238 | 1.184E-19 | 323.9 |
| ENSMUSG00000062184 | heparan sulfate 6-O-sulfotransferase 2 [Source:MGI Symbol;Acc:MGI:1354959] | Hs6st2 | Enriched in dLGN | 2.236 | 2.025E-14 | 235.4 |
| ENSMUSG00000019966 | kit ligand [Source:MGI Symbol;Acc:MGI:96974] | Kitl | Enriched in dLGN | 2.231 | 1.39E-11 | 453.1 |
| ENSMUSG00000040653 | protein phosphatase 1, regulatory inhibitor subunit 14C [Source:MGI Symbol;Acc:MGI:1923392] | Ppp1r14c | Enriched in dLGN | 2.208 | 2.389E-19 | 150.8 |
| ENSMUSG00000024907 | galanin and GMAP prepropeptide [Source:MGI Symbol;Acc:MGI:95637] | Gal | Enriched in dLGN | 2.205 | 0.000002477 | 30 |
| ENSMUSG00000041216 | clavesin 1 [Source:MGI Symbol;Acc:MGI:1921688] | Clvs1 | Enriched in dLGN | 2.204 | 1.751E-13 | 163.7 |
| ENSMUSG00000035735 | diacylglycerol lipase, alpha [Source:MGI Symbol;Acc:MGI:2677061] | Dagla | Enriched in dLGN | 2.201 | 9.424E-32 | 435.4 |
| ENSMUSG00000071369 | mitogen-activated protein kinase kinase kinase 5 [Source:MGI Symbol;Acc:MGI:1346876] | Map3k5 | Enriched in dLGN | 2.201 | 8.577E-22 | 236 |
| ENSMUSG00000029632 | Ndufa4, mitochondrial complex associated [Source:MGI Symbol;Acc:MGI:107686] | Ndufa4 | Enriched in dLGN | 2.193 | 6.104E-13 | 183.2 |
| ENSMUSG00000045009 | proline-rich transmembrane protein 3 [Source:MGI Symbol;Acc:MGI:2444810] | Prrt3 | Enriched in dLGN | 2.192 | 1.468E-17 | 111.1 |
| ENSMUSG00000022112 | glypican 5 [Source:MGI Symbol;Acc:MGI:1194894] | Gpc5 | Enriched in dLGN | 2.191 | 0.000001098 | 390.3 |
| ENSMUSG00000050138 | potassium channel, subfamily K, member 12 [Source:MGI Symbol;Acc:MGI:2684043] | Kcnk12 | Enriched in dLGN | 2.185 | 1.615E-10 | 38.23 |
| ENSMUSG00000043811 | reticulon 4 receptor [Source:MGI Symbol;Acc:MGI:2136886] | Rtn4r | Enriched in dLGN | 2.171 | 6.926E-12 | 85.48 |
| ENSMUSG00000031970 | dysbindin (dystrobrevin binding protein 1) domain containing 1 [Source:MGI Symbol;Acc:MGI:1919435] | Dbndd1 | Enriched in dLGN | 2.167 | 5.671E-21 | 82.55 |
| ENSMUSG00000019194 | sodium channel, voltage-gated, type I, beta [Source:MGI Symbol;Acc:MGI:98247] | Scn1b | Enriched in dLGN | 2.163 | 5.937E-15 | 470 |
| ENSMUSG00000035000 | dipeptidylpeptidase 4 [Source:MGI Symbol;Acc:MGI:94919] | Dpp4 | Enriched in dLGN | 2.145 | 0.00009184 | 9.243 |
| ENSMUSG00000036578 | FXYD domain-containing ion transport regulator 7 [Source:MGI Symbol;Acc:MGI:1889006] | Fxyd7 | Enriched in dLGN | 2.135 | 3.36E-10 | 140.4 |
| ENSMUSG00000061451 | transmembrane protein 151A [Source:MGI Symbol;Acc:MGI:2147713] | Tmem151a | Enriched in dLGN | 2.126 | 1.361E-22 | 350.6 |
| ENSMUSG00000028610 | DMRT-like family B with proline-rich C-terminal, 1 [Source:MGI Symbol;Acc:MGI:1927125] | Dmrtb1 | Enriched in dLGN | 2.121 | 0.000001975 | 17.02 |
| ENSMUSG00000026643 | N-myristoyltransferase 2 [Source:MGI Symbol;Acc:MGI:1202298] | Nmt2 | Enriched in dLGN | 2.117 | 8.259E-37 | 203.8 |
| ENSMUSG00000040430 | phosphatidylinositol transfer protein, cytoplasmic 1 [Source:MGI Symbol;Acc:MGI:1919045] | Pitpnc1 | Enriched in dLGN | 2.115 | 1.153E-27 | 764.9 |
| ENSMUSG00000021947 | crystallin, lambda 1 [Source:MGI Symbol;Acc:MGI:1915881] | Cryl1 | Enriched in dLGN | 2.105 | 2.936E-10 | 28.77 |
| ENSMUSG00000044017 | adhesion G protein-coupled receptor D1 [Source:MGI Symbol;Acc:MGI:3041203] | Adgrd1 | Enriched in dLGN | 2.104 | 0.001235 | 5.834 |
| ENSMUSG00000028149 | RAP1, GTP-GDP dissociation stimulator 1 [Source:MGI Symbol;Acc:MGI:2385189] | Rap1gds1 | Enriched in dLGN | 2.102 | 2.562E-28 | 847.3 |
| ENSMUSG00000034612 | carbohydrate sulfotransferase 11 [Source:MGI Symbol;Acc:MGI:1927166] | Chst11 | Enriched in dLGN | 2.096 | 2.127E-17 | 236.6 |
| ENSMUSG00000006057 | ATP synthase, H+ transporting, mitochondrial F0 complex, subunit C1 (subunit 9) [Source:MGI Symbol;Acc:MGI:107653] | Atp5g1 | Enriched in dLGN | 2.096 | 5.922E-12 | 76.53 |
| ENSMUSG00000032966 | FK506 binding protein 1a [Source:MGI Symbol;Acc:MGI:95541] | Fkbp1a | Enriched in dLGN | 2.095 | 9.382E-23 | 430.5 |
| ENSMUSG00000026959 | glutamate receptor, ionotropic, NMDA1 (zeta 1) [Source:MGI Symbol;Acc:MGI:95819] | Grin1 | Enriched in dLGN | 2.085 | 5.045E-13 | 942.6 |
| ENSMUSG00000028773 | fatty acid binding protein 3, muscle and heart [Source:MGI Symbol;Acc:MGI:95476] | Fabp3 | Enriched in dLGN | 2.08 | 0.00001646 | 75.72 |
| ENSMUSG00000036330 | solute carrier family 18 (vesicular monoamine), member 1 [Source:MGI Symbol;Acc:MGI:106684] | Slc18a1 | Enriched in dLGN | 2.078 | 0.001218 | 5.605 |
| ENSMUSG00000034839 | La ribonucleoprotein 6, translational regulator [Source:MGI Symbol;Acc:MGI:1914807] | Larp6 | Enriched in dLGN | 2.063 | 4.656E-24 | 87.58 |
| ENSMUSG00000025428 | ATP synthase, H+ transporting, mitochondrial F1 complex, alpha subunit 1 [Source:MGI Symbol;Acc:MGI:88115] | Atp5a1 | Enriched in dLGN | 2.062 | 3.448E-13 | 1689 |
| ENSMUSG00000000901 | matrix metallopeptidase 11 [Source:MGI Symbol;Acc:MGI:97008] | Mmp11 | Enriched in dLGN | 2.035 | 0.00002266 | 8.781 |
| ENSMUSG00000044024 | RELT-like 2 [Source:MGI Symbol;Acc:MGI:1918044] | Rell2 | Enriched in dLGN | 2.03 | 5.701E-28 | 285.3 |
| ENSMUSG00000029189 | sel-1 suppressor of lin-12-like 3 (C. elegans) [Source:MGI Symbol;Acc:MGI:1916941] | Sel1l3 | Enriched in dLGN | 2.025 | 3.647E-15 | 154 |
| ENSMUSG00000042807 | HECT, C2 and WW domain containing E3 ubiquitin protein ligase 2 [Source:MGI Symbol;Acc:MGI:2685817] | Hecw2 | Enriched in dLGN | 2.023 | 1.085E-11 | 320.7 |
| ENSMUSG00000037606 | oxysterol binding protein-like 5 [Source:MGI Symbol;Acc:MGI:1930265] | Osbpl5 | Enriched in dLGN | 2.013 | 2.893E-35 | 213.9 |
| ENSMUSG00000038552 | fibronectin type III domain containing 4 [Source:MGI Symbol;Acc:MGI:1917195] | Fndc4 | Enriched in dLGN | 2.01 | 7.667E-32 | 238.1 |
| ENSMUSG00000027134 | lysophosphatidylcholine acyltransferase 4 [Source:MGI Symbol;Acc:MGI:2138993] | Lpcat4 | Enriched in dLGN | 2.006 | 6.427E-27 | 130.5 |
| ENSMUSG00000039809 | gamma-aminobutyric acid (GABA) B receptor, 2 [Source:MGI Symbol;Acc:MGI:2386030] | Gabbr2 | Enriched in dLGN | 2 | 2.048E-25 | 1664 |
| ENSMUSG00000049612 | oligodendrocyte myelin glycoprotein [Source:MGI Symbol;Acc:MGI:106586] | Omg | Enriched in dLGN | 2 | 1.93E-09 | 36.01 |
| ENSMUSG00000075702 | selenoprotein M [Source:MGI Symbol;Acc:MGI:2149786] | Selenom | Enriched in dLGN | 1.999 | 5.454E-16 | 136.2 |
| ENSMUSG00000029769 | coiled-coil domain containing 136 [Source:MGI Symbol;Acc:MGI:1918128] | Ccdc136 | Enriched in dLGN | 1.997 | 1.863E-23 | 1034 |
| ENSMUSG00000007944 | tetratricopeptide repeat domain 9B [Source:MGI Symbol;Acc:MGI:1920282] | Ttc9b | Enriched in dLGN | 1.996 | 3.867E-15 | 147 |
| ENSMUSG00000031772 | contactin associated protein-like 4 [Source:MGI Symbol;Acc:MGI:2183572] | Cntnap4 | Enriched in dLGN | 1.996 | 2.035E-09 | 164.6 |
| ENSMUSG00000033595 | leucine-rich repeat LGI family, member 3 [Source:MGI Symbol;Acc:MGI:2182619] | Lgi3 | Enriched in dLGN | 1.994 | 1.257E-09 | 185.5 |
| ENSMUSG00000024953 | peroxiredoxin 5 [Source:MGI Symbol;Acc:MGI:1859821] | Prdx5 | Enriched in dLGN | 1.989 | 4.476E-19 | 344.2 |
| ENSMUSG00000016252 | ATP synthase, H+ transporting, mitochondrial F1 complex, epsilon subunit [Source:MGI Symbol;Acc:MGI:1855697] | Atp5e | Enriched in dLGN | 1.988 | 1.008E-09 | 153.2 |
| ENSMUSG00000026904 | solute carrier family 4, sodium bicarbonate cotransporter-like, member 10 [Source:MGI Symbol;Acc:MGI:2150150] | Slc4a10 | Enriched in dLGN | 1.984 | 1.631E-15 | 601.1 |
| ENSMUSG00000030718 | protein phosphatase methylesterase 1 [Source:MGI Symbol;Acc:MGI:1919840] | Ppme1 | Enriched in dLGN | 1.979 | 7.751E-15 | 576.9 |
| ENSMUSG00000040811 | echinoderm microtubule associated protein like 2 [Source:MGI Symbol;Acc:MGI:1919455] | Eml2 | Enriched in dLGN | 1.978 | 2.487E-21 | 740.8 |
| ENSMUSG00000113149 | predicted gene, 49383 [Source:MGI Symbol;Acc:MGI:6121605] | Gm49383 | Enriched in dLGN | 1.978 | 0.002167 | 6.916 |
| ENSMUSG00000020163 | ubiquinol-cytochrome c reductase, complex III subunit XI [Source:MGI Symbol;Acc:MGI:1913844] | Uqcr11 | Enriched in dLGN | 1.976 | 2.208E-09 | 153.3 |
| ENSMUSG00000027894 | solute carrier family 6 (neurotransmitter transporter), member 17 [Source:MGI Symbol;Acc:MGI:2442535] | Slc6a17 | Enriched in dLGN | 1.97 | 2.287E-18 | 1006 |
| ENSMUSG00000034220 | glypican 1 [Source:MGI Symbol;Acc:MGI:1194891] | Gpc1 | Enriched in dLGN | 1.963 | 1.636E-25 | 632.1 |
| ENSMUSG00000047746 | F-box protein 40 [Source:MGI Symbol;Acc:MGI:2443753] | Fbxo40 | Enriched in dLGN | 1.958 | 0.0001325 | 15.08 |
| ENSMUSG00000028710 | ATP synthase mitochondrial F1 complex assembly factor 1 [Source:MGI Symbol;Acc:MGI:2180560] | Atpaf1 | Enriched in dLGN | 1.955 | 2.605E-19 | 89.62 |
| ENSMUSG00000002068 | cyclin E1 [Source:MGI Symbol;Acc:MGI:88316] | Ccne1 | Enriched in dLGN | 1.947 | 6.098E-09 | 29.03 |
| ENSMUSG00000004558 | N-myc downstream regulated gene 2 [Source:MGI Symbol;Acc:MGI:1352498] | Ndrg2 | Enriched in dLGN | 1.946 | 2.755E-16 | 450.7 |
| ENSMUSG00000021520 | ubiquinol-cytochrome c reductase binding protein [Source:MGI Symbol;Acc:MGI:1914780] | Uqcrb | Enriched in dLGN | 1.943 | 0.000000305 | 49.09 |
| ENSMUSG00000039546 | adherens junction associated protein 1 [Source:MGI Symbol;Acc:MGI:2685419] | Ajap1 | Enriched in dLGN | 1.931 | 5.645E-15 | 293.4 |
| ENSMUSG00000008153 | calsyntenin 3 [Source:MGI Symbol;Acc:MGI:2178323] | Clstn3 | Enriched in dLGN | 1.931 | 1.12E-12 | 1064 |
| ENSMUSG00000027792 | butyrylcholinesterase [Source:MGI Symbol;Acc:MGI:894278] | Bche | Enriched in dLGN | 1.926 | 0.00000244 | 17.98 |
| ENSMUSG00000018965 | tyrosine 3-monooxygenase/tryptophan 5-monooxygenase activation protein, eta polypeptide [Source:MGI Symbol;Acc:MGI:109194] | Ywhah | Enriched in dLGN | 1.923 | 4.601E-10 | 3239 |
| ENSMUSG00000027698 | neutral cholesterol ester hydrolase 1 [Source:MGI Symbol;Acc:MGI:2443191] | Nceh1 | Enriched in dLGN | 1.919 | 6.474E-12 | 190.1 |
| ENSMUSG00000042298 | tetratricopeptide repeat domain 19 [Source:MGI Symbol;Acc:MGI:1920045] | Ttc19 | Enriched in dLGN | 1.917 | 4.712E-16 | 462.6 |
| ENSMUSG00000025094 | solute carrier family 18 (vesicular monoamine), member 2 [Source:MGI Symbol;Acc:MGI:106677] | Slc18a2 | Enriched in dLGN | 1.917 | 0.00009947 | 54.24 |
| ENSMUSG00000029765 | plexin A4 [Source:MGI Symbol;Acc:MGI:2179061] | Plxna4 | Enriched in dLGN | 1.915 | 7.997E-17 | 780.1 |
| ENSMUSG00000024935 | solute carrier family 1 (neuronal/epithelial high affinity glutamate transporter, system Xag), member 1 [Source:MGI Symbol;Acc:MGI:1050 | Slc1a1 | Enriched in dLGN | 1.914 | 3.309E-20 | 323.7 |
| ENSMUSG00000049422 | coiled-coil-helix-coiled-coil-helix domain containing 10 [Source:MGI Symbol;Acc:MGI:2143558] | Chchd10 | Enriched in dLGN | 1.91 | 4.345E-11 | 224.5 |
| ENSMUSG00000026321 | tumor necrosis factor receptor superfamily, member 11a, NFKB activator [Source:MGI Symbol;Acc:MGI:1314891] | Tnfrsf11a | Enriched in dLGN | 1.895 | 5.849E-09 | 53.11 |
| ENSMUSG00000038264 | sema domain, immunoglobulin domain (Ig), and GPI membrane anchor, (semaphorin) 7A [Source:MGI Symbol;Acc:MGI:1306826] | Sema7a | Enriched in dLGN | 1.894 | 6.689E-14 | 229.8 |
| ENSMUSG00000022602 | activity regulated cytoskeletal-associated protein [Source:MGI Symbol;Acc:MGI:88067] | Arc | Enriched in dLGN | 1.891 | 0.000000399 | 44.21 |
| ENSMUSG00000027556 | carbonic anhydrase 1 [Source:MGI Symbol;Acc:MGI:88268] | Car1 | Enriched in dLGN | 1.885 | 0.00005975 | 12.27 |
| ENSMUSG00000000884 | guanine nucleotide binding protein (G protein), beta polypeptide 1-like [Source:MGI Symbol;Acc:MGI:1338057] | Gnb1l | Enriched in dLGN | 1.885 | 0.006886 | 294.8 |
| ENSMUSG00000111375 | BTB (POZ) domain containing 8 [Source:MGI Symbol;Acc:MGI:3646208] | Btbd8 | Enriched in dLGN | 1.878 | 1.755E-11 | 197 |
| ENSMUSG00000020321 | malate dehydrogenase 1, NAD (soluble) [Source:MGI Symbol;Acc:MGI:97051] | Mdh1 | Enriched in dLGN | 1.876 | 0.000001979 | 617.5 |
| ENSMUSG00000079941 | cytochrome c oxidase subunit 5B, pseudogene [Source:MGI Symbol;Acc:MGI:3649411] | Cox5b-ps | Enriched in dLGN | 1.866 | 4.831E-07 | 35.35 |
| ENSMUSG00000028444 | ciliary neurotrophic factor receptor [Source:MGI Symbol;Acc:MGI:99605] | Cntfr | Enriched in dLGN | 1.863 | 5.614E-16 | 219.4 |
| ENSMUSG00000015656 | heat shock protein 8 [Source:MGI Symbol;Acc:MGI:105384] | Hspa8 | Enriched in dLGN | 1.862 | 0.000009677 | 1562 |
| ENSMUSG00000051146 | calcium/calmodulin-dependent protein kinase II inhibitor 2 [Source:MGI Symbol;Acc:MGI:1920297] | Camk2n2 | Enriched in dLGN | 1.859 | 1.333E-20 | 513.1 |
| ENSMUSG00000022366 | solute carrier family 22 (organic cation transporter), member 22 [Source:MGI Symbol;Acc:MGI:2446114] | Slc22a22 | Enriched in dLGN | 1.858 | 0.004299 | 26.67 |
| ENSMUSG00000064202 | spermatogenesis associated 6 like [Source:MGI Symbol;Acc:MGI:1918036] | Spata6l | Enriched in dLGN | 1.852 | 3.705E-07 | 24.41 |
| ENSMUSG00000022587 | lymphocyte antigen 6 complex, locus E [Source:MGI Symbol;Acc:MGI:106651] | Ly6e | Enriched in dLGN | 1.85 | 1.25E-10 | 95.38 |
| ENSMUSG00000033847 | phospholipase A2, group IVC (cytosolic, calcium-independent) [Source:MGI Symbol;Acc:MGI:1196403] | Pla2g4c | Enriched in dLGN | 1.846 | 0.03952 | 7.739 |
| ENSMUSG00000051359 | neurocalcin delta [Source:MGI Symbol;Acc:MGI:1196326] | Ncald | Enriched in dLGN | 1.837 | 7.056E-23 | 710.7 |
| ENSMUSG00000044252 | oxysterol binding protein-like 1A [Source:MGI Symbol;Acc:MGI:1927551] | Osbpl1a | Enriched in dLGN | 1.837 | 3.734E-17 | 434.2 |
| ENSMUSG00000009894 | synaptosomal-associated protein, 47 [Source:MGI Symbol;Acc:MGI:1915076] | Snap47 | Enriched in dLGN | 1.835 | 1.542E-10 | 533.6 |
| ENSMUSG00000006711 | RIKEN cDNA D130043K22 gene [Source:MGI Symbol;Acc:MGI:3036268] | D130043K22Rik | Enriched in dLGN | 1.832 | 6.834E-09 | 133.1 |
| ENSMUSG00000054459 | visinin-like 1 [Source:MGI Symbol;Acc:MGI:1349453] | Vsnl1 | Enriched in dLGN | 1.832 | 0.00001163 | 746.5 |
| ENSMUSG00000009394 | synapsin II [Source:MGI Symbol;Acc:MGI:103020] | Syn2 | Enriched in dLGN | 1.831 | 3.978E-12 | 611.4 |
| ENSMUSG00000027104 | activating transcription factor 2 [Source:MGI Symbol;Acc:MGI:109349] | Atf2 | Enriched in dLGN | 1.831 | 0.000005923 | 703.5 |
| ENSMUSG00000028251 | thiosulfate sulfurtransferase (rhodanese)-like domain containing 3 [Source:MGI Symbol;Acc:MGI:1924282] | Tstd3 | Enriched in dLGN | 1.83 | 1.293E-07 | 35.11 |
| ENSMUSG00000030500 | solute carrier family 17 (sodium-dependent inorganic phosphate cotransporter), member 6 [Source:MGI Symbol;Acc:MGI:2156052] | Slc17a6 | Enriched in dLGN | 1.827 | 0.000005218 | 605.1 |
| ENSMUSG00000015217 | high mobility group box 3 [Source:MGI Symbol;Acc:MGI:1098219] | Hmgb3 | Enriched in dLGN | 1.823 | 3.254E-11 | 141 |

| | | | | | | |
|---|---|---|---|---|---|---|
| ENSMUSG00000019990 | phosphodiesterase 7B [Source:MGI Symbol;Acc:MGI:1352752] | Pde7b | Enriched in dLGN | 1.822 | 1.779E-08 | 194.2 |
| ENSMUSG00000061518 | cytochrome c oxidase subunit 5B [Source:MGI Symbol;Acc:MGI:88475] | Cox5b | Enriched in dLGN | 1.818 | 3.27E-12 | 455.8 |
| ENSMUSG00000041697 | cytochrome c oxidase subunit 6A1 [Source:MGI Symbol;Acc:MGI:103099] | Cox6a1 | Enriched in dLGN | 1.818 | 2.467E-10 | 168 |
| ENSMUSG00000028351 | bone morphogenic protein/retinoic acid inducible neural specific 1 [Source:MGI Symbol;Acc:MGI:1928478] | Brinp1 | Enriched in dLGN | 1.815 | 1.508E-09 | 586.1 |
| ENSMUSG00000028648 | NADH:ubiquinone oxidoreductase core subunit S5 [Source:MGI Symbol;Acc:MGI:1890889] | Ndufs5 | Enriched in dLGN | 1.81 | 8.028E-09 | 101.9 |
| ENSMUSG00000028949 | SWI/SNF related, matrix associated, actin dependent regulator of chromatin, subfamily d, member 3 [Source:MGI Symbol;Acc:MGI:191424 | Smarcd3 | Enriched in dLGN | 1.803 | 5.528E-30 | 234.8 |
| ENSMUSG00000049556 | leucine rich repeat and Ig domain containing 1 [Source:MGI Symbol;Acc:MGI:1915522] | Lingo1 | Enriched in dLGN | 1.796 | 1.482E-19 | 811.2 |
| ENSMUSG00000040118 | calcium channel, voltage-dependent, alpha2/delta subunit 1 [Source:MGI Symbol;Acc:MGI:88295] | Cacna2d1 | Enriched in dLGN | 1.795 | 6.248E-11 | 541.2 |
| ENSMUSG00000025790 | solute carrier organic anion transporter family, member 3a1 [Source:MGI Symbol;Acc:MGI:1351867] | Slco3a1 | Enriched in dLGN | 1.792 | 1.583E-08 | 185.2 |
| ENSMUSG00000056201 | cofilin 1, non-muscle [Source:MGI Symbol;Acc:MGI:101757] | Cfl1 | Enriched in dLGN | 1.79 | 3.617E-10 | 762.6 |
| ENSMUSG00000053192 | myeloid/lymphoid or mixed-lineage leukemia; translocated to, 11 [Source:MGI Symbol;Acc:MGI:1929671] | Mllt11 | Enriched in dLGN | 1.788 | 8.972E-16 | 499.1 |
| ENSMUSG00000036934 | RIKEN cDNA 4921524J17 gene [Source:MGI Symbol;Acc:MGI:1913964] | 4921524J17Rik | Enriched in dLGN | 1.786 | 2.697E-08 | 42.54 |
| ENSMUSG00000091722 | siah E3 ubiquitin protein ligase family member 3 [Source:MGI Symbol;Acc:MGI:2685758] | Siah3 | Enriched in dLGN | 1.785 | 1.926E-11 | 62.95 |
| ENSMUSG00000028334 | N-acetylneuraminic acid synthase (sialic acid synthase) [Source:MGI Symbol;Acc:MGI:2149820] | Nans | Enriched in dLGN | 1.783 | 4.029E-09 | 106.3 |
| ENSMUSG00000063802 | HSPA (heat shock 70kDa) binding protein, cytoplasmic cochaperone 1 [Source:MGI Symbol;Acc:MGI:1913495] | Hspbp1 | Enriched in dLGN | 1.778 | 1.986E-20 | 128.5 |
| ENSMUSG00000026526 | fumarate hydratase 1 [Source:MGI Symbol;Acc:MGI:95530] | Fh1 | Enriched in dLGN | 1.776 | 7.003E-08 | 124 |
| ENSMUSG00000030609 | apoptosis enhancing nuclease [Source:MGI Symbol;Acc:MGI:1915298] | Aen | Enriched in dLGN | 1.772 | 3.455E-21 | 72.83 |
| ENSMUSG00000036751 | cytochrome c oxidase, subunit 6B1 [Source:MGI Symbol;Acc:MGI:107460] | Cox6b1 | Enriched in dLGN | 1.77 | 1.327E-12 | 143.6 |
| ENSMUSG00000038421 | Fc receptor-like A [Source:MGI Symbol;Acc:MGI:2138647] | Fcrla | Enriched in dLGN | 1.77 | 0.004335 | 28.16 |
| ENSMUSG00000032564 | copine IV [Source:MGI Symbol;Acc:MGI:1921270] | Cpne4 | Enriched in dLGN | 1.765 | 0.000001545 | 126.3 |
| ENSMUSG00000027581 | stathmin-like 3 [Source:MGI Symbol;Acc:MGI:1277137] | Stmn3 | Enriched in dLGN | 1.747 | 6.07E-09 | 1780 |
| ENSMUSG00000019179 | malate dehydrogenase 2, NAD (mitochondrial) [Source:MGI Symbol;Acc:MGI:97050] | Mdh2 | Enriched in dLGN | 1.743 | 4.47E-08 | 626.3 |
| ENSMUSG00000058400 | pyroglutamylated RFamide peptide receptor [Source:MGI Symbol;Acc:MGI:2677633] | Qrfpr | Enriched in dLGN | 1.743 | 0.0001073 | 24.15 |
| ENSMUSG00000071866 | peptidylprolyl isomerase A [Source:MGI Symbol;Acc:MGI:97749] | Ppia | Enriched in dLGN | 1.743 | 0.0005841 | 211.2 |
| ENSMUSG00000073433 | Rho GDP dissociation inhibitor (GDI) gamma [Source:MGI Symbol;Acc:MGI:108430] | Arhgdig | Enriched in dLGN | 1.738 | 2.299E-10 | 185.1 |
| ENSMUSG00000026347 | transmembrane protein 163 [Source:MGI Symbol;Acc:MGI:1919410] | Tmem163 | Enriched in dLGN | 1.735 | 2.35E-11 | 376 |
| ENSMUSG00000063931 | peptidase D [Source:MGI Symbol;Acc:MGI:97542] | Pepd | Enriched in dLGN | 1.735 | 9.56E-08 | 102.6 |
| ENSMUSG00000020736 | 5',3'-nucleotidase, cytosolic [Source:MGI Symbol;Acc:MGI:1354954] | Nt5c | Enriched in dLGN | 1.73 | 9.491E-13 | 100.9 |
| ENSMUSG00000029185 | family with sequence similarity 114, member A1 [Source:MGI Symbol;Acc:MGI:1915553] | Fam114a1 | Enriched in dLGN | 1.724 | 8.776E-07 | 29.67 |
| ENSMUSG00000001380 | histidyl-tRNA synthetase [Source:MGI Symbol;Acc:MGI:108087] | Hars | Enriched in dLGN | 1.713 | 6.58E-19 | 256 |
| ENSMUSG00000053024 | contactin 2 [Source:MGI Symbol;Acc:MGI:104518] | Cntn2 | Enriched in dLGN | 1.709 | 2.541E-11 | 303.9 |
| ENSMUSG00000035885 | cytochrome c oxidase subunit 8A [Source:MGI Symbol;Acc:MGI:105959] | Cox8a | Enriched in dLGN | 1.707 | 4.629E-13 | 292.6 |
| ENSMUSG00000047547 | clathrin, light polypeptide (Lcb) [Source:MGI Symbol;Acc:MGI:1921575] | Cltb | Enriched in dLGN | 1.705 | 1.667E-35 | 896 |
| ENSMUSG00000032946 | RAS, guanyl releasing protein 2 [Source:MGI Symbol;Acc:MGI:1333849] | Rasgrp2 | Enriched in dLGN | 1.693 | 1.452E-08 | 149.8 |
| ENSMUSG00000059361 | neurensin 2 [Source:MGI Symbol;Acc:MGI:2684969] | Nrsn2 | Enriched in dLGN | 1.692 | 0.000001407 | 106.4 |
| ENSMUSG00000036158 | prickle planar cell polarity protein 1 [Source:MGI Symbol;Acc:MGI:1916034] | Prickle1 | Enriched in dLGN | 1.691 | 4.499E-13 | 385.1 |
| ENSMUSG00000061576 | dipeptidylpeptidase 6 [Source:MGI Symbol;Acc:MGI:94921] | Dpp6 | Enriched in dLGN | 1.691 | 2.307E-07 | 1354 |
| ENSMUSG00000042182 | BEN domain containing 6 [Source:MGI Symbol;Acc:MGI:2444572] | Bend6 | Enriched in dLGN | 1.689 | 4.842E-15 | 256 |
| ENSMUSG00000032366 | tropomyosin 1, alpha [Source:MGI Symbol;Acc:MGI:98809] | Tpm1 | Enriched in dLGN | 1.684 | 2.575E-35 | 654.7 |
| ENSMUSG00000053329 | glutamine amidotransferase like class 1 domain containing 3A [Source:MGI Symbol;Acc:MGI:1351861] | Gatd3a | Enriched in dLGN | 1.678 | 6.918E-17 | 113.5 |
| ENSMUSG00000024083 | praja ring finger ubiquitin ligase 2 [Source:MGI Symbol;Acc:MGI:2159342] | Pja2 | Enriched in dLGN | 1.675 | 8.584E-11 | 1111 |
| ENSMUSG00000029120 | protein phosphatase 2, regulatory subunit B, gamma [Source:MGI Symbol;Acc:MGI:2442660] | Ppp2r2c | Enriched in dLGN | 1.674 | 1.508E-22 | 1258 |
| ENSMUSG00000052981 | ubiquitin-conjugating enzyme E2Q family-like 1 [Source:MGI Symbol;Acc:MGI:1924230] | Ube2ql1 | Enriched in dLGN | 1.669 | 6.26E-15 | 310.3 |
| ENSMUSG00000001986 | glutamate receptor, ionotropic, AMPA3 (alpha 3) [Source:MGI Symbol;Acc:MGI:95810] | Gria3 | Enriched in dLGN | 1.668 | 1.388E-11 | 283.8 |
| ENSMUSG00000002771 | glutamate receptor, ionotropic, NMDA2D (epsilon 4) [Source:MGI Symbol;Acc:MGI:95823] | Grin2d | Enriched in dLGN | 1.665 | 1.762E-14 | 466.7 |
| ENSMUSG00000063694 | cytochrome c, somatic [Source:MGI Symbol;Acc:MGI:88578] | Cycs | Enriched in dLGN | 1.665 | 1.274E-08 | 116.9 |
| ENSMUSG00000027500 | stathmin-like 2 [Source:MGI Symbol;Acc:MGI:98241] | Stmn2 | Enriched in dLGN | 1.658 | 1.768E-07 | 1265 |
| ENSMUSG00000029465 | actin related protein 2/3 complex, subunit 3 [Source:MGI Symbol;Acc:MGI:1928375] | Arpc3 | Enriched in dLGN | 1.658 | 2.073E-07 | 143.2 |
| ENSMUSG00000029641 | RAS-like, family 11, member A [Source:MGI Symbol;Acc:MGI:1916145] | Rasl11a | Enriched in dLGN | 1.655 | 0.00007873 | 21.43 |
| ENSMUSG00000019831 | WASP family, member 1 [Source:MGI Symbol;Acc:MGI:1890563] | Wasf1 | Enriched in dLGN | 1.647 | 4.293E-24 | 290.1 |
| ENSMUSG00000025762 | La ribonucleoprotein 1B [Source:MGI Symbol;Acc:MGI:1914604] | Larp1b | Enriched in dLGN | 1.647 | 1.615E-10 | 100 |
| ENSMUSG00000020116 | partner of NOB1 homolog [Source:MGI Symbol;Acc:MGI:1913499] | Pno1 | Enriched in dLGN | 1.646 | 3.781E-09 | 43.22 |
| ENSMUSG00000046442 | protein phosphatase 1E (PP2C domain containing) [Source:MGI Symbol;Acc:MGI:2444096] | Ppm1e | Enriched in dLGN | 1.644 | 5.869E-14 | 844.2 |
| ENSMUSG00000034488 | EGF-like repeats and discoidin I-like domains 3 [Source:MGI Symbol;Acc:MGI:1329025] | Edil3 | Enriched in dLGN | 1.641 | 0.000001149 | 564.8 |
| ENSMUSG00000024359 | heat shock protein 9 [Source:MGI Symbol;Acc:MGI:96245] | Hspa9 | Enriched in dLGN | 1.64 | 9.066E-09 | 574.8 |
| ENSMUSG00000089717 | olfactory receptor family 13 subfamily F member 5 [Source:MGI Symbol;Acc:MGI:3030109] | Or13f5 | Enriched in dLGN | 1.639 | 0.02318 | 4.484 |
| ENSMUSG00000026496 | poly (ADP-ribose) polymerase family, member 1 [Source:MGI Symbol;Acc:MGI:1340806] | Parp1 | Enriched in dLGN | 1.629 | 9.287E-20 | 378.8 |
| ENSMUSG00000028161 | protein phosphatase 3, catalytic subunit, alpha isoform [Source:MGI Symbol;Acc:MGI:107164] | Ppp3ca | Enriched in dLGN | 1.623 | 3.414E-21 | 1200 |
| ENSMUSG00000032959 | phosphatidylethanolamine binding protein 1 [Source:MGI Symbol;Acc:MGI:1344408] | Pebp1 | Enriched in dLGN | 1.623 | 2.757E-07 | 164.1 |
| ENSMUSG00000031603 | fibroblast growth factor 20 [Source:MGI Symbol;Acc:MGI:1891346] | Fgf20 | Enriched in dLGN | 1.621 | 0.004419 | 11.6 |
| ENSMUSG00000044117 | bMERB domain containing 1 [Source:MGI Symbol;Acc:MGI:1914504] | Bmerb1 | Enriched in dLGN | 1.618 | 2.7E-16 | 457.3 |
| ENSMUSG00000034171 | fatty acid amide hydrolase [Source:MGI Symbol;Acc:MGI:109609] | Faah | Enriched in dLGN | 1.617 | 4.413E-21 | 134 |
| ENSMUSG00000050856 | ATP synthase, H+ transporting, mitochondrial F1F0 complex, subunit E [Source:MGI Symbol;Acc:MGI:106636] | Atp5k | Enriched in dLGN | 1.617 | 0.000006295 | 126.9 |
| ENSMUSG00000025968 | NADH:ubiquinone oxidoreductase core subunit S1 [Source:MGI Symbol;Acc:MGI:2443241] | Ndufs1 | Enriched in dLGN | 1.614 | 1.051E-11 | 337.9 |
| ENSMUSG00000035960 | apurinic/apyrimidinic endonuclease 1 [Source:MGI Symbol;Acc:MGI:88042] | Apex1 | Enriched in dLGN | 1.613 | 0.00004016 | 35.5 |
| ENSMUSG00000031231 | cytochrome c oxidase subunit 7B [Source:MGI Symbol;Acc:MGI:1913392] | Cox7b | Enriched in dLGN | 1.605 | 0.0006304 | 113.9 |
| ENSMUSG00000000088 | cytochrome c oxidase subunit 5A [Source:MGI Symbol;Acc:MGI:88474] | Cox5a | Enriched in dLGN | 1.602 | 5.215E-08 | 112.1 |
| ENSMUSG00000028410 | DnaJ heat shock protein family (Hsp40) member A1 [Source:MGI Symbol;Acc:MGI:1270129] | Dnaja1 | Enriched in dLGN | 1.596 | 0.00002769 | 251.8 |
| ENSMUSG00000010914 | pyruvate dehydrogenase complex, component X [Source:MGI Symbol;Acc:MGI:1351627] | Pdhx | Enriched in dLGN | 1.588 | 8.507E-14 | 152.7 |
| ENSMUSG00000029033 | ArfGAP with coiled-coil, ankyrin repeat and PH domains 3 [Source:MGI Symbol;Acc:MGI:2153589] | Acap3 | Enriched in dLGN | 1.585 | 1.541E-25 | 352.1 |
| ENSMUSG00000033061 | regulated endocrine-specific protein 18 [Source:MGI Symbol;Acc:MGI:1098222] | Resp18 | Enriched in dLGN | 1.583 | 0.002201 | 74.08 |
| ENSMUSG00000033389 | Rho GTPase activating protein 44 [Source:MGI Symbol;Acc:MGI:2144423] | Arhgap44 | Enriched in dLGN | 1.582 | 1.627E-17 | 522.2 |
| ENSMUSG00000079184 | M-phase phosphoprotein 8 [Source:MGI Symbol;Acc:MGI:1922589] | Mphosph8 | Enriched in dLGN | 1.581 | 0.00003348 | 685.9 |
| ENSMUSG00000025940 | transmembrane protein 70 [Source:MGI Symbol;Acc:MGI:1915068] | Tmem70 | Enriched in dLGN | 1.579 | 2.162E-11 | 50.14 |
| ENSMUSG00000031391 | L1 cell adhesion molecule [Source:MGI Symbol;Acc:MGI:96721] | L1cam | Enriched in dLGN | 1.579 | 0.00001756 | 803.2 |
| ENSMUSG00000044005 | glutaminase 2 (liver, mitochondrial) [Source:MGI Symbol;Acc:MGI:2143539] | Gls2 | Enriched in dLGN | 1.578 | 0.000009756 | 17.76 |
| ENSMUSG00000043556 | F-box and leucine-rich repeat protein 7 [Source:MGI Symbol;Acc:MGI:3052506] | Fbxl7 | Enriched in dLGN | 1.574 | 0.001429 | 47.6 |
| ENSMUSG00000067242 | leucine-rich repeat LGI family, member 1 [Source:MGI Symbol;Acc:MGI:1861691] | Lgi1 | Enriched in dLGN | 1.569 | 1.209E-07 | 72.29 |
| ENSMUSG00000042817 | FMS-like tyrosine kinase 3 [Source:MGI Symbol;Acc:MGI:95559] | Flt3 | Enriched in dLGN | 1.563 | 0.000134 | 42.78 |
| ENSMUSG00000067455 | H4 clustered histone 11 [Source:MGI Symbol;Acc:MGI:2448436] | H4c11 | Enriched in dLGN | 1.562 | 0.002706 | 5.483 |
| ENSMUSG00000034930 | rhotekin [Source:MGI Symbol;Acc:MGI:107371] | Rtkn | Enriched in dLGN | 1.56 | 8.977E-10 | 71.08 |
| ENSMUSG00000026858 | mitoguardin 2 [Source:MGI Symbol;Acc:MGI:1922035] | Miga2 | Enriched in dLGN | 1.558 | 7.706E-20 | 319.8 |
| ENSMUSG00000082229 | nucleosome assembly protein 1-like 2 [Source:MGI Symbol;Acc:MGI:106654] | Nap1l2 | Enriched in dLGN | 1.557 | 0.000000708 | 277.1 |
| ENSMUSG00000029840 | myotrophin [Source:MGI Symbol;Acc:MGI:99445] | Mtpn | Enriched in dLGN | 1.552 | 1.002E-13 | 335.2 |
| ENSMUSG00000073838 | Tu translation elongation factor, mitochondrial [Source:MGI Symbol;Acc:MGI:1923686] | Tufm | Enriched in dLGN | 1.552 | 1.372E-07 | 100.2 |
| ENSMUSG00000032011 | thymus cell antigen 1, theta [Source:MGI Symbol;Acc:MGI:98747] | Thy1 | Enriched in dLGN | 1.546 | 0.000001101 | 952.8 |
| ENSMUSG00000062825 | actin, gamma, cytoplasmic 1 [Source:MGI Symbol;Acc:MGI:87906] | Actg1 | Enriched in dLGN | 1.543 | 0.00009369 | 407 |
| ENSMUSG00000039480 | 5'-nucleotidase domain containing 1 [Source:MGI Symbol;Acc:MGI:2442446] | Nt5dc1 | Enriched in dLGN | 1.542 | 0.0001665 | 84.48 |
| ENSMUSG00000022270 | reticulophagy regulator 1 [Source:MGI Symbol;Acc:MGI:1913520] | Retreg1 | Enriched in dLGN | 1.541 | 7.349E-09 | 162.5 |
| ENSMUSG00000038462 | ubiquinol-cytochrome c reductase, Rieske iron-sulfur polypeptide 1 [Source:MGI Symbol;Acc:MGI:1913944] | Uqcrfs1 | Enriched in dLGN | 1.536 | 1.638E-10 | 172.4 |
| ENSMUSG00000025980 | heat shock protein 1 (chaperonin) [Source:MGI Symbol;Acc:MGI:96242] | Hspd1 | Enriched in dLGN | 1.535 | 1.134E-10 | 264 |
| ENSMUSG00000036850 | mitochondrial ribosomal protein L41 [Source:MGI Symbol;Acc:MGI:1333816] | Mrpl41 | Enriched in dLGN | 1.535 | 2.053E-10 | 53.68 |
| ENSMUSG00000021114 | ATPase, H+ transporting, lysosomal V1 subunit D [Source:MGI Symbol;Acc:MGI:1921084] | Atp6v1d | Enriched in dLGN | 1.529 | 9.017E-10 | 284.8 |
| ENSMUSG00000035964 | transmembrane protein 59-like [Source:MGI Symbol;Acc:MGI:1915187] | Tmem59l | Enriched in dLGN | 1.527 | 5.473E-08 | 166 |
| ENSMUSG00000063882 | ubiquinol-cytochrome c reductase hinge protein [Source:MGI Symbol;Acc:MGI:1913826] | Uqcrh | Enriched in dLGN | 1.526 | 1.962E-12 | 256.5 |
| ENSMUSG00000007021 | synaptogyrin 3 [Source:MGI Symbol;Acc:MGI:1341881] | Syngr3 | Enriched in dLGN | 1.525 | 3.277E-10 | 261.6 |
| ENSMUSG00000035049 | ribosomal RNA processing 12 homolog [Source:MGI Symbol;Acc:MGI:2147437] | Rrp12 | Enriched in dLGN | 1.523 | 1.525E-12 | 124.1 |
| ENSMUSG00000022337 | ER membrane protein complex subunit 2 [Source:MGI Symbol;Acc:MGI:1913986] | Emc2 | Enriched in dLGN | 1.523 | 6.544E-11 | 104.6 |
| ENSMUSG00000034566 | ATP synthase, H+ transporting, mitochondrial F0 complex, subunit D [Source:MGI Symbol;Acc:MGI:1918929] | Atp5h | Enriched in dLGN | 1.521 | 1.21E-08 | 216.1 |
| ENSMUSG00000024758 | reticulon 3 [Source:MGI Symbol;Acc:MGI:1339970] | Rtn3 | Enriched in dLGN | 1.519 | 2.24E-19 | 1535 |
| ENSMUSG00000005510 | NADH:ubiquinone oxidoreductase core subunit S3 [Source:MGI Symbol;Acc:MGI:1915599] | Ndufs3 | Enriched in dLGN | 1.519 | 5.001E-12 | 101.1 |
| ENSMUSG00000007564 | protein phosphatase 2, regulatory subunit A, alpha [Source:MGI Symbol;Acc:MGI:1926334] | Ppp2r1a | Enriched in dLGN | 1.512 | 2.102E-07 | 634.9 |
| ENSMUSG00000022205 | SUB1 homolog, transcriptional regulator [Source:MGI Symbol;Acc:MGI:104811] | Sub1 | Enriched in dLGN | 1.511 | 2.295E-18 | 399.4 |
| ENSMUSG00000033352 | mitogen-activated protein kinase kinase 4 [Source:MGI Symbol;Acc:MGI:1346869] | Map2k4 | Enriched in dLGN | 1.502 | 1.869E-15 | 341.7 |
| ENSMUSG00000029126 | neuron specific gene family member 1 [Source:MGI Symbol;Acc:MGI:109149] | Nsg1 | Enriched in dLGN | 1.502 | 8.903E-10 | 719.7 |
| ENSMUSG00000023089 | NADH:ubiquinone oxidoreductase subunit A5 [Source:MGI Symbol;Acc:MGI:1915452] | Ndufa5 | Enriched in dLGN | 1.501 | 1.184E-08 | 166.1 |
| ENSMUSG00000036438 | calmodulin 2 [Source:MGI Symbol;Acc:MGI:103250] | Calm2 | Enriched in dLGN | 1.499 | 0.0001059 | 1256 |
| ENSMUSG00000039989 | chromobox 4 [Source:MGI Symbol;Acc:MGI:1195985] | Cbx4 | Enriched in dLGN | 1.493 | 4.698E-23 | 266.9 |
| ENSMUSG00000030337 | vesicle-associated membrane protein 1 [Source:MGI Symbol;Acc:MGI:1313276] | Vamp1 | Enriched in dLGN | 1.493 | 4.124E-10 | 282.4 |
| ENSMUSG00000031841 | cadherin 13 [Source:MGI Symbol;Acc:MGI:99551] | Cdh13 | Enriched in dLGN | 1.491 | 0.00003803 | 448.3 |
| ENSMUSG00000025092 | heat shock protein 12A [Source:MGI Symbol;Acc:MGI:1920692] | Hspa12a | Enriched in dLGN | 1.49 | 5.894E-12 | 913 |
| ENSMUSG00000029649 | proteasome maturation protein [Source:MGI Symbol;Acc:MGI:1913787] | Pomp | Enriched in dLGN | 1.489 | 4.628E-09 | 109 |
| ENSMUSG00000027108 | Obg-like ATPase 1 [Source:MGI Symbol;Acc:MGI:1914309] | Ola1 | Enriched in dLGN | 1.487 | 0.02071 | 454.9 |
| ENSMUSG00000117853 | vomeronasal 1 receptor, 88 [Source:MGI Symbol;Acc:MGI:3852416] | Vmn1r88 | Enriched in dLGN | 1.486 | 0.04341 | 3.474 |
| ENSMUSG00000029659 | mesenteric estrogen dependent adipogenesis [Source:MGI Symbol;Acc:MGI:1917967] | Medag | Enriched in dLGN | 1.484 | 0.0001069 | 25.4 |
| ENSMUSG00000029223 | ubiquitin carboxy-terminal hydrolase L1 [Source:MGI Symbol;Acc:MGI:103149] | Uchl1 | Enriched in dLGN | 1.483 | 0.000003407 | 1008 |
| ENSMUSG00000025189 | cyclin M1 [Source:MGI Symbol;Acc:MGI:1891366] | Cnnm1 | Enriched in dLGN | 1.477 | 4.281E-07 | 258.6 |
| ENSMUSG00000039838 | solute carrier family 45, member 1 [Source:MGI Symbol;Acc:MGI:2653235] | Slc45a1 | Enriched in dLGN | 1.475 | 2.957E-20 | 187.4 |
| ENSMUSG00000005683 | citrate synthase [Source:MGI Symbol;Acc:MGI:88529] | Cs | Enriched in dLGN | 1.475 | 1.801E-10 | 414 |
| ENSMUSG00000075318 | sodium channel, voltage-gated, type II, alpha [Source:MGI Symbol;Acc:MGI:98248] | Scn2a | Enriched in dLGN | 1.468 | 9.425E-14 | 1334 |
| ENSMUSG00000063873 | solute carrier family 24 (sodium/potassium/calcium exchanger), member 3 [Source:MGI Symbol;Acc:MGI:2137513] | Slc24a3 | Enriched in dLGN | 1.467 | 2.034E-09 | 1240 |
| ENSMUSG00000046058 | EP300 interacting inhibitor of differentiation 2 [Source:MGI Symbol;Acc:MGI:2681174] | Eid2 | Enriched in dLGN | 1.467 | 0.000007251 | 137.6 |
| ENSMUSG00000056091 | ST3 beta-galactoside alpha-2,3-sialyltransferase 5 [Source:MGI Symbol;Acc:MGI:1339963] | St3gal5 | Enriched in dLGN | 1.466 | 3.178E-15 | 176.3 |
| ENSMUSG00000032657 | endosomal transmembrane epsin interactor 3 [Source:MGI Symbol;Acc:MGI:1915771] | Entrep3 | Enriched in dLGN | 1.466 | 1.81E-08 | 83.78 |
| ENSMUSG00000020949 | FK506 binding protein 3 [Source:MGI Symbol;Acc:MGI:1353460] | Fkbp3 | Enriched in dLGN | 1.459 | 0.000005486 | 160.5 |
| ENSMUSG00000014313 | cytochrome c oxidase subunit 6C [Source:MGI Symbol;Acc:MGI:104614] | Cox6c | Enriched in dLGN | 1.459 | 0.000006076 | 345.4 |
| ENSMUSG00000054428 | ATPase inhibitory factor 1 [Source:MGI Symbol;Acc:MGI:1196457] | Atpif1 | Enriched in dLGN | 1.458 | 3.311E-08 | 829.4 |
| ENSMUSG00000036815 | dipeptidylpeptidase 10 [Source:MGI Symbol;Acc:MGI:2442409] | Dpp10 | Enriched in dLGN | 1.455 | 0.00006936 | 576.4 |

| | | | | | | |
|---|---|---|---|---|---|---|
| ENSMUSG00000042682 | selenoprotein K [Source:MGI Symbol;Acc:MGI:1931466] | Selenok | Enriched in dLGN | 1.454 | 0.00001069 | 254.3 |
| ENSMUSG00000061080 | limbic system-associated membrane protein [Source:MGI Symbol;Acc:MGI:1261760] | Lsamp | Enriched in dLGN | 1.451 | 0.00001014 | 3253 |
| ENSMUSG00000052698 | talin 2 [Source:MGI Symbol;Acc:MGI:1917799] | Tln2 | Enriched in dLGN | 1.444 | 8.403E-23 | 1565 |
| ENSMUSG00000025142 | alveolar soft part sarcoma chromosome region, candidate 1 (human) [Source:MGI Symbol;Acc:MGI:1916188] | Aspscr1 | Enriched in dLGN | 1.444 | 5.835E-12 | 101.1 |
| ENSMUSG00000027220 | synaptotagmin XIII [Source:MGI Symbol;Acc:MGI:1933945] | Syt13 | Enriched in dLGN | 1.443 | 1.503E-16 | 763.5 |
| ENSMUSG00000027203 | deoxyuridine triphosphatase [Source:MGI Symbol;Acc:MGI:1346051] | Dut | Enriched in dLGN | 1.442 | 2.704E-12 | 92.41 |
| ENSMUSG00000026750 | proteasome (prosome, macropain) subunit, beta type 7 [Source:MGI Symbol;Acc:MGI:107637] | Psmb7 | Enriched in dLGN | 1.441 | 7.697E-09 | 169.1 |
| ENSMUSG00000024777 | protein phosphatase 2, regulatory subunit B', beta [Source:MGI Symbol;Acc:MGI:2388480] | Ppp2r5b | Enriched in dLGN | 1.44 | 5.18E-19 | 428 |
| ENSMUSG00000031561 | teneurin transmembrane protein 3 [Source:MGI Symbol;Acc:MGI:1345183] | Tenm3 | Enriched in dLGN | 1.436 | 2.111E-12 | 2385 |
| ENSMUSG00000017781 | phosphatidylinositol transfer protein, alpha [Source:MGI Symbol;Acc:MGI:99887] | Pitpna | Enriched in dLGN | 1.434 | 3.406E-16 | 792.9 |
| ENSMUSG00000006932 | catenin (cadherin associated protein), beta 1 [Source:MGI Symbol;Acc:MGI:88276] | Ctnnb1 | Enriched in dLGN | 1.434 | 2.034E-08 | 1065 |
| ENSMUSG00000049971 | glycosyltransferase 1 domain containing 1 [Source:MGI Symbol;Acc:MGI:2442755] | Glt1d1 | Enriched in dLGN | 1.429 | 4.336E-10 | 53.34 |
| ENSMUSG00000030417 | programmed cell death 5 [Source:MGI Symbol;Acc:MGI:1913538] | Pdcd5 | Enriched in dLGN | 1.429 | 3.209E-08 | 70.53 |
| ENSMUSG00000029875 | coiled-coil domain containing 184 [Source:MGI Symbol;Acc:MGI:2146066] | Ccdc184 | Enriched in dLGN | 1.428 | 4.104E-07 | 201 |
| ENSMUSG00000002379 | NADH:ubiquinone oxidoreductase subunit A11 [Source:MGI Symbol;Acc:MGI:1917125] | Ndufa11 | Enriched in dLGN | 1.428 | 0.001137 | 75.68 |
| ENSMUSG00000018446 | complement component 1, q subcomponent binding protein [Source:MGI Symbol;Acc:MGI:1194505] | C1qbp | Enriched in dLGN | 1.427 | 2.145E-07 | 91.52 |
| ENSMUSG00000019951 | UHRF1 (ICBP90) binding protein 1-like [Source:MGI Symbol;Acc:MGI:2442888] | Uhrf1bp1l | Enriched in dLGN | 1.426 | 2.724E-24 | 276.6 |
| ENSMUSG00000059409 | protein phosphatase 2, regulatory subunit B', delta [Source:MGI Symbol;Acc:MGI:2388481] | Ppp2r5d | Enriched in dLGN | 1.424 | 1.957E-14 | 362.3 |
| ENSMUSG00000040896 | potassium voltage-gated channel, Shal-related family, member 3 [Source:MGI Symbol;Acc:MGI:1928743] | Kcnd3 | Enriched in dLGN | 1.422 | 2.124E-08 | 579.5 |
| ENSMUSG00000040373 | calcium channel, voltage-dependent, gamma subunit 5 [Source:MGI Symbol;Acc:MGI:2157946] | Cacng5 | Enriched in dLGN | 1.421 | 1.702E-09 | 203.8 |
| ENSMUSG00000100486 | predicted gene 4131 [Source:MGI Symbol;Acc:MGI:3782307] | Gm4131 | Enriched in dLGN | 1.419 | 6.653E-10 | 68.64 |
| ENSMUSG00000038555 | receptor accessory protein 2 [Source:MGI Symbol;Acc:MGI:2385070] | Reep2 | Enriched in dLGN | 1.418 | 9.671E-13 | 302.7 |
| ENSMUSG00000003072 | ATP synthase, H+ transporting, mitochondrial F1 complex, delta subunit [Source:MGI Symbol;Acc:MGI:1913293] | Atp5d | Enriched in dLGN | 1.415 | 5.726E-10 | 259.8 |
| ENSMUSG00000024127 | prolyl endopeptidase-like [Source:MGI Symbol;Acc:MGI:2441932] | Prepl | Enriched in dLGN | 1.413 | 8.785E-07 | 815.8 |
| ENSMUSG00000019505 | ubiquitin B [Source:MGI Symbol;Acc:MGI:98888] | Ubb | Enriched in dLGN | 1.411 | 0.0003141 | 652 |
| ENSMUSG00000024064 | polypeptide N-acetylgalactosaminyltransferase 14 [Source:MGI Symbol;Acc:MGI:1918935] | Galnt14 | Enriched in dLGN | 1.409 | 0.00001849 | 190 |
| ENSMUSG00000031701 | DnaJ heat shock protein family (Hsp40) member A2 [Source:MGI Symbol;Acc:MGI:1931882] | Dnaja2 | Enriched in dLGN | 1.408 | 2.871E-08 | 401 |
| ENSMUSG00000027827 | potassium voltage-gated channel, shaker-related subfamily, beta member 1 [Source:MGI Symbol;Acc:MGI:109155] | Kcnab1 | Enriched in dLGN | 1.407 | 0.00006488 | 138 |
| ENSMUSG00000038690 | ATP synthase, H+ transporting, mitochondrial F0 complex, subunit F2 [Source:MGI Symbol;Acc:MGI:1927558] | Atp5j2 | Enriched in dLGN | 1.406 | 7.338E-09 | 138.3 |
| ENSMUSG00000029752 | asparagine synthetase [Source:MGI Symbol;Acc:MGI:1350929] | Asns | Enriched in dLGN | 1.406 | 0.0001324 | 106.2 |
| ENSMUSG00000049493 | plastin 1 (I-isoform) [Source:MGI Symbol;Acc:MGI:104809] | Pls1 | Enriched in dLGN | 1.404 | 0.004094 | 15.52 |
| ENSMUSG00000004642 | stem-loop binding protein [Source:MGI Symbol;Acc:MGI:108402] | Slbp | Enriched in dLGN | 1.402 | 1.176E-09 | 85.78 |
| ENSMUSG00000027131 | ER membrane protein complex subunit 4 [Source:MGI Symbol;Acc:MGI:1915282] | Emc4 | Enriched in dLGN | 1.401 | 0.000001195 | 81.85 |
| ENSMUSG00000047879 | ubiquitin specific peptidase 14 [Source:MGI Symbol;Acc:MGI:1928898] | Usp14 | Enriched in dLGN | 1.4 | 5.225E-10 | 157.3 |
| ENSMUSG00000022890 | ATP synthase, H+ transporting, mitochondrial F0 complex, subunit F [Source:MGI Symbol;Acc:MGI:107777] | Atp5j | Enriched in dLGN | 1.4 | 0.000000698 | 249.2 |
| ENSMUSG00000062380 | tubulin, beta 3 class III [Source:MGI Symbol;Acc:MGI:107813] | Tubb3 | Enriched in dLGN | 1.399 | 0.005535 | 1342 |
| ENSMUSG00000045624 | ESF1 nucleolar pre-rRNA processing protein homolog [Source:MGI Symbol;Acc:MGI:1913830] | Esf1 | Enriched in dLGN | 1.397 | 1.881E-08 | 125.5 |
| ENSMUSG00000001507 | integrin alpha 3 [Source:MGI Symbol;Acc:MGI:96602] | Itga3 | Enriched in dLGN | 1.392 | 9.676E-11 | 148.8 |
| ENSMUSG00000025089 | glial cell line derived neurotrophic factor family receptor alpha 1 [Source:MGI Symbol;Acc:MGI:1100842] | Gfra1 | Enriched in dLGN | 1.388 | 1.691E-11 | 184.8 |
| ENSMUSG00000021737 | proteasome (prosome, macropain) 26S subunit, non-ATPase, 6 [Source:MGI Symbol;Acc:MGI:1913663] | Psmd6 | Enriched in dLGN | 1.388 | 4.453E-08 | 147.1 |
| ENSMUSG00000079523 | thymosin, beta 10 [Source:MGI Symbol;Acc:MGI:109146] | Tmsb10 | Enriched in dLGN | 1.382 | 0.00002169 | 881.5 |
| ENSMUSG00000073676 | heat shock protein 1 (chaperonin 10) [Source:MGI Symbol;Acc:MGI:104680] | Hspe1 | Enriched in dLGN | 1.382 | 0.00002807 | 62.31 |
| ENSMUSG00000026260 | NADH:ubiquinone oxidoreductase subunit A10 [Source:MGI Symbol;Acc:MGI:1914523] | Ndufa10 | Enriched in dLGN | 1.38 | 0.000006407 | 263.7 |
| ENSMUSG00000033900 | microtubule-associated protein 9 [Source:MGI Symbol;Acc:MGI:2442208] | Map9 | Enriched in dLGN | 1.378 | 8.019E-13 | 524.3 |
| ENSMUSG00000031007 | ATPase, H+ transporting, lysosomal accessory protein 2 [Source:MGI Symbol;Acc:MGI:1917745] | Atp6ap2 | Enriched in dLGN | 1.374 | 0.000003085 | 242.6 |
| ENSMUSG00000031176 | dynein light chain Tctex-type 3 [Source:MGI Symbol;Acc:MGI:1914367] | Dynlt3 | Enriched in dLGN | 1.373 | 0.00001057 | 112.8 |
| ENSMUSG00000079598 | C-type lectin domain family 2, member L [Source:MGI Symbol;Acc:MGI:2141402] | Clec2l | Enriched in dLGN | 1.371 | 9.034E-07 | 252.6 |
| ENSMUSG00000030849 | fibroblast growth factor receptor 2 [Source:MGI Symbol;Acc:MGI:95523] | Fgfr2 | Enriched in dLGN | 1.371 | 0.007178 | 937.9 |
| ENSMUSG00000032435 | dynein cytoplasmic 1 light intermediate chain 1 [Source:MGI Symbol;Acc:MGI:2135610] | Dync1li1 | Enriched in dLGN | 1.37 | 5.349E-29 | 559.8 |
| ENSMUSG00000047221 | family with sequence similarity 185, member A [Source:MGI Symbol;Acc:MGI:2140983] | Fam185a | Enriched in dLGN | 1.37 | 0.000002534 | 76.93 |
| ENSMUSG00000047126 | clathrin, heavy polypeptide (Hc) [Source:MGI Symbol;Acc:MGI:2388633] | Cltc | Enriched in dLGN | 1.367 | 3.668E-07 | 1529 |
| ENSMUSG00000026430 | Ras association (RalGDS/AF-6) domain family member 5 [Source:MGI Symbol;Acc:MGI:1926375] | Rassf5 | Enriched in dLGN | 1.366 | 2.225E-09 | 93.47 |
| ENSMUSG00000035653 | leucine rich repeat and fibronectin type III domain containing 5 [Source:MGI Symbol;Acc:MGI:2144814] | Lrfn5 | Enriched in dLGN | 1.365 | 0.00005541 | 564.4 |
| ENSMUSG00000030647 | NADH:ubiquinone oxidoreductase subunit C2 [Source:MGI Symbol;Acc:MGI:1344370] | Ndufc2 | Enriched in dLGN | 1.364 | 0.0001216 | 77.57 |
| ENSMUSG00000032279 | isocitrate dehydrogenase 3 (NAD+) alpha [Source:MGI Symbol;Acc:MGI:1915084] | Idh3a | Enriched in dLGN | 1.362 | 0.0008029 | 302 |
| ENSMUSG00000026585 | kinesin-associated protein 3 [Source:MGI Symbol;Acc:MGI:107566] | Kifap3 | Enriched in dLGN | 1.359 | 9.477E-08 | 663.9 |
| ENSMUSG00000022892 | amyloid beta (A4) precursor protein [Source:MGI Symbol;Acc:MGI:88059] | App | Enriched in dLGN | 1.357 | 0.000001597 | 4592 |
| ENSMUSG00000023861 | mitochondrial pyruvate carrier 1 [Source:MGI Symbol;Acc:MGI:1915240] | Mpc1 | Enriched in dLGN | 1.353 | 0.000007329 | 73.22 |
| ENSMUSG00000072964 | basic helix-loop-helix domain containing, class B9 [Source:MGI Symbol;Acc:MGI:1917487] | Bhlhb9 | Enriched in dLGN | 1.351 | 0.000003581 | 158.1 |
| ENSMUSG00000042216 | small G protein signaling modulator 1 [Source:MGI Symbol;Acc:MGI:107320] | Sgsm1 | Enriched in dLGN | 1.349 | 2.438E-09 | 412.2 |
| ENSMUSG00000020029 | nudix (nucleoside diphosphate linked moiety X)-type motif 4 [Source:MGI Symbol;Acc:MGI:1918457] | Nudt4 | Enriched in dLGN | 1.346 | 2.343E-08 | 752.5 |
| ENSMUSG00000059734 | NADH:ubiquinone oxidoreductase core subunit S8 [Source:MGI Symbol;Acc:MGI:2385079] | Ndufs8 | Enriched in dLGN | 1.344 | 0.00002622 | 73.63 |
| ENSMUSG00000071014 | NADH:ubiquinone oxidoreductase subunit B6 [Source:MGI Symbol;Acc:MGI:2684983] | Ndufb6 | Enriched in dLGN | 1.342 | 0.00001153 | 50.89 |
| ENSMUSG00000024862 | kinesin light chain 2 [Source:MGI Symbol;Acc:MGI:107953] | Klc2 | Enriched in dLGN | 1.341 | 5.195E-13 | 682.5 |
| ENSMUSG00000060212 | pecanex homolog 2 [Source:MGI Symbol;Acc:MGI:2445010] | Pcnx2 | Enriched in dLGN | 1.341 | 2.025E-09 | 282.9 |
| ENSMUSG00000015961 | adenylosuccinate synthetase, non muscle [Source:MGI Symbol;Acc:MGI:87948] | Adss | Enriched in dLGN | 1.339 | 5.268E-10 | 159.9 |
| ENSMUSG00000029860 | zyxin [Source:MGI Symbol;Acc:MGI:103072] | Zyx | Enriched in dLGN | 1.339 | 3.579E-09 | 118.6 |
| ENSMUSG00000025651 | ubiquinol-cytochrome c reductase core protein 1 [Source:MGI Symbol;Acc:MGI:107876] | Uqcrc1 | Enriched in dLGN | 1.339 | 0.00001971 | 409.8 |
| ENSMUSG00000043091 | tubulin, alpha 1C [Source:MGI Symbol;Acc:MGI:1095409] | Tuba1c | Enriched in dLGN | 1.338 | 0.02535 | 41.77 |
| ENSMUSG00000006731 | beta-1,4-N-acetyl-galactosaminyl transferase 1 [Source:MGI Symbol;Acc:MGI:1342057] | B4galnt1 | Enriched in dLGN | 1.335 | 3.016E-10 | 230.2 |
| ENSMUSG00000038489 | polymerase (RNA) II (DNA directed) polypeptide L [Source:MGI Symbol;Acc:MGI:1913741] | Polr2l | Enriched in dLGN | 1.332 | 0.000001395 | 28.51 |
| ENSMUSG00000027603 | gamma-glutamyltransferase 7 [Source:MGI Symbol;Acc:MGI:1913385] | Ggt7 | Enriched in dLGN | 1.33 | 1.693E-08 | 194.7 |
| ENSMUSG00000049775 | thymosin, beta 4, X chromosome [Source:MGI Symbol;Acc:MGI:99510] | Tmsb4x | Enriched in dLGN | 1.329 | 4.807E-11 | 317.2 |
| ENSMUSG00000000563 | ATP synthase peripheral stalk-membrane subunit b [Source:MGI Symbol;Acc:MGI:1100495] | Atp5pb | Enriched in dLGN | 1.329 | 0.000000142 | 236 |
| ENSMUSG00000062995 | islet cell autoantigen 1 [Source:MGI Symbol;Acc:MGI:96391] | Ica1 | Enriched in dLGN | 1.329 | 0.002921 | 217.1 |
| ENSMUSG00000026860 | SH3-domain GRB2-like endophilin B2 [Source:MGI Symbol;Acc:MGI:2385131] | Sh3glb2 | Enriched in dLGN | 1.327 | 7.258E-10 | 236.6 |
| ENSMUSG00000052738 | succinate-CoA ligase, GDP-forming, alpha subunit [Source:MGI Symbol;Acc:MGI:1927234] | Suclg1 | Enriched in dLGN | 1.326 | 0.00002124 | 138.6 |
| ENSMUSG00000062542 | synaptotagmin IX [Source:MGI Symbol;Acc:MGI:1926373] | Syt9 | Enriched in dLGN | 1.325 | 7.262E-12 | 283.3 |
| ENSMUSG00000019849 | prolyl endopeptidase [Source:MGI Symbol;Acc:MGI:1270863] | Prep | Enriched in dLGN | 1.324 | 1.762E-07 | 207.5 |
| ENSMUSG00000001089 | leucine zipper protein 1 [Source:MGI Symbol;Acc:MGI:107629] | Luzp1 | Enriched in dLGN | 1.324 | 4.658E-07 | 485.1 |
| ENSMUSG00000021250 | FBJ osteosarcoma oncogene [Source:MGI Symbol;Acc:MGI:95574] | Fos | Enriched in dLGN | 1.319 | 0.00001421 | 57.19 |
| ENSMUSG00000005161 | peroxiredoxin 2 [Source:MGI Symbol;Acc:MGI:109486] | Prdx2 | Enriched in dLGN | 1.318 | 0.00001099 | 321.2 |
| ENSMUSG00000000168 | dihydrolipoamide S-acetyltransferase (E2 component of pyruvate dehydrogenase complex) [Source:MGI Symbol;Acc:MGI:2385311] | Dlat | Enriched in dLGN | 1.317 | 1.502E-10 | 196.1 |
| ENSMUSG00000060279 | adaptor-related protein complex 2, alpha 1 subunit [Source:MGI Symbol;Acc:MGI:101921] | Ap2a1 | Enriched in dLGN | 1.317 | 2.453E-10 | 671.9 |
| ENSMUSG00000042750 | brain expressed X-linked 2 [Source:MGI Symbol;Acc:MGI:1338017] | Bex2 | Enriched in dLGN | 1.316 | 5.733E-08 | 798 |
| ENSMUSG00000027533 | fatty acid binding protein 5, epidermal [Source:MGI Symbol;Acc:MGI:101790] | Fabp5 | Enriched in dLGN | 1.316 | 0.01836 | 258.1 |
| ENSMUSG00000015476 | proline-rich transmembrane protein 1 [Source:MGI Symbol;Acc:MGI:1932118] | Prrt1 | Enriched in dLGN | 1.313 | 0.000007046 | 115.5 |
| ENSMUSG00000016349 | eukaryotic translation elongation factor 1 alpha 2 [Source:MGI Symbol;Acc:MGI:1096317] | Eef1a2 | Enriched in dLGN | 1.312 | 0.000009096 | 1496 |
| ENSMUSG00000028790 | KH domain containing, RNA binding, signal transduction associated 1 [Source:MGI Symbol;Acc:MGI:893579] | Khdrbs1 | Enriched in dLGN | 1.31 | 4.729E-11 | 341 |
| ENSMUSG00000026568 | mitochondrial pyruvate carrier 2 [Source:MGI Symbol;Acc:MGI:1917706] | Mpc2 | Enriched in dLGN | 1.31 | 4.524E-07 | 84.48 |
| ENSMUSG00000044894 | ubiquinol-cytochrome c reductase, complex III subunit VII [Source:MGI Symbol;Acc:MGI:107807] | Uqcrq | Enriched in dLGN | 1.308 | 0.000001026 | 236.5 |
| ENSMUSG00000019210 | ATPase, H+ transporting, lysosomal V1 subunit E1 [Source:MGI Symbol;Acc:MGI:894326] | Atp6v1e1 | Enriched in dLGN | 1.306 | 0.00002408 | 270.4 |
| ENSMUSG00000071528 | ATP synthase membrane subunit DAPIT [Source:MGI Symbol;Acc:MGI:1891435] | Atp5md | Enriched in dLGN | 1.306 | 0.0004203 | 71.32 |
| ENSMUSG00000019861 | golgi associated PDZ and coiled-coil motif containing [Source:MGI Symbol;Acc:MGI:2149946] | Gopc | Enriched in dLGN | 1.304 | 2.072E-20 | 304.7 |
| ENSMUSG00000055022 | contactin 1 [Source:MGI Symbol;Acc:MGI:105980] | Cntn1 | Enriched in dLGN | 1.301 | 4.369E-08 | 1753 |
| ENSMUSG00000026576 | ATPase, Na+/K+ transporting, beta 1 polypeptide [Source:MGI Symbol;Acc:MGI:88108] | Atp1b1 | Enriched in dLGN | 1.299 | 0.00007133 | 3270 |
| ENSMUSG00000078812 | eukaryotic translation initiation factor 5A [Source:MGI Symbol;Acc:MGI:106248] | Eif5a | Enriched in dLGN | 1.297 | 5.177E-09 | 424.1 |
| ENSMUSG00000025545 | citrate lyase beta like [Source:MGI Symbol;Acc:MGI:1916884] | Clybl | Enriched in dLGN | 1.297 | 0.0002367 | 35.95 |
| ENSMUSG00000038335 | TSR1 20S rRNA accumulation [Source:MGI Symbol;Acc:MGI:2144566] | Tsr1 | Enriched in dLGN | 1.292 | 9.21E-17 | 103.4 |
| ENSMUSG00000020042 | BTB (POZ) domain containing 11 [Source:MGI Symbol;Acc:MGI:1921257] | Btbd11 | Enriched in dLGN | 1.292 | 0.000001013 | 318.5 |
| ENSMUSG00000045348 | neuronal tyrosine-phosphorylated phosphoinositide 3-kinase adaptor 1 [Source:MGI Symbol;Acc:MGI:2443880] | Nyap1 | Enriched in dLGN | 1.291 | 3.614E-11 | 501.1 |
| ENSMUSG00000028020 | glycine receptor, beta subunit [Source:MGI Symbol;Acc:MGI:95751] | Glrb | Enriched in dLGN | 1.291 | 4.703E-10 | 323.7 |
| ENSMUSG00000003380 | Rab acceptor 1 (prenylated) [Source:MGI Symbol;Acc:MGI:1201692] | Rabac1 | Enriched in dLGN | 1.289 | 1.239E-10 | 110.7 |
| ENSMUSG00000040455 | ubiquitin specific petidase 45 [Source:MGI Symbol;Acc:MGI:101850] | Usp45 | Enriched in dLGN | 1.289 | 0.000000964 | 101.7 |
| ENSMUSG00000039629 | striatin interacting protein 2 [Source:MGI Symbol;Acc:MGI:2444363] | Strip2 | Enriched in dLGN | 1.289 | 0.00007177 | 138 |
| ENSMUSG00000044628 | ring finger protein 208 [Source:MGI Symbol;Acc:MGI:1916096] | Rnf208 | Enriched in dLGN | 1.288 | 2.358E-09 | 128.2 |
| ENSMUSG00000033793 | ATPase, H+ transporting, lysosomal V1 subunit H [Source:MGI Symbol;Acc:MGI:1914864] | Atp6v1h | Enriched in dLGN | 1.286 | 0.000006472 | 178.8 |
| ENSMUSG00000028212 | cyclin E2 [Source:MGI Symbol;Acc:MGI:1329034] | Ccne2 | Enriched in dLGN | 1.284 | 0.004031 | 12.53 |
| ENSMUSG00000031432 | phosphoribosyl pyrophosphate synthetase 1 [Source:MGI Symbol;Acc:MGI:97775] | Prps1 | Enriched in dLGN | 1.283 | 6.055E-07 | 65.65 |
| ENSMUSG00000034880 | mitochondrial ribosomal protein L34 [Source:MGI Symbol;Acc:MGI:2137227] | Mrpl34 | Enriched in dLGN | 1.283 | 0.000002696 | 34.19 |
| ENSMUSG00000003273 | carbonic anhydrase 11 [Source:MGI Symbol;Acc:MGI:1336193] | Car11 | Enriched in dLGN | 1.282 | 5.018E-08 | 105.3 |
| ENSMUSG00000061603 | A kinase (PRKA) anchor protein 6 [Source:MGI Symbol;Acc:MGI:3050566] | Akap6 | Enriched in dLGN | 1.28 | 1.983E-10 | 2172 |
| ENSMUSG00000018865 | sulfotransferase family 4A, member 1 [Source:MGI Symbol;Acc:MGI:1888971] | Sult4a1 | Enriched in dLGN | 1.275 | 5.227E-10 | 795.2 |
| ENSMUSG00000026377 | nucleolar protein interacting with the FHA domain of MKI67 [Source:MGI Symbol;Acc:MGI:1915199] | Nifk | Enriched in dLGN | 1.274 | 2.419E-07 | 61.02 |
| ENSMUSG00000064329 | sodium channel, voltage-gated, type I, alpha [Source:MGI Symbol;Acc:MGI:98246] | Scn1a | Enriched in dLGN | 1.274 | 0.00001516 | 703.3 |
| ENSMUSG00000025393 | ATP synthase, H+ transporting mitochondrial F1 complex, beta subunit [Source:MGI Symbol;Acc:MGI:107801] | Atp5b | Enriched in dLGN | 1.274 | 0.00003526 | 1784 |
| ENSMUSG00000042109 | cold shock domain containing C2, RNA binding [Source:MGI Symbol;Acc:MGI:2146027] | Csdc2 | Enriched in dLGN | 1.273 | 9.909E-12 | 601.3 |
| ENSMUSG00000061032 | ribosomal RNA processing 1 [Source:MGI Symbol;Acc:MGI:1203500] | Rrp1 | Enriched in dLGN | 1.272 | 5.225E-10 | 886 |
| ENSMUSG00000016526 | dual-specificity tyrosine-(Y)-phosphorylation regulated kinase 3 [Source:MGI Symbol;Acc:MGI:1330300] | Dyrk3 | Enriched in dLGN | 1.272 | 0.0001113 | 17.86 |
| ENSMUSG00000063406 | transmembrane p24 trafficking protein 5 [Source:MGI Symbol;Acc:MGI:1921586] | Tmed5 | Enriched in dLGN | 1.269 | 0.00002707 | 35.86 |
| ENSMUSG00000034343 | ubiquitin-conjugating enzyme E2F (putative) [Source:MGI Symbol;Acc:MGI:1915171] | Ube2f | Enriched in dLGN | 1.268 | 2.583E-10 | 95.96 |
| ENSMUSG00000031672 | glutamatic-oxaloacetic transaminase 2, mitochondrial [Source:MGI Symbol;Acc:MGI:95792] | Got2 | Enriched in dLGN | 1.266 | 0.00002756 | 253.7 |
| ENSMUSG00000029777 | glycyl-tRNA synthetase [Source:MGI Symbol;Acc:MGI:2449057] | Gars | Enriched in dLGN | 1.265 | 3.765E-10 | 453.4 |
| ENSMUSG00000025487 | proteasome (prosome, macropain) 26S subunit, non-ATPase, 13 [Source:MGI Symbol;Acc:MGI:1345192] | Psmd13 | Enriched in dLGN | 1.26 | 1.625E-11 | 179.8 |
| ENSMUSG00000020349 | protein phosphatase 2 (formerly 2A), catalytic subunit, alpha isoform [Source:MGI Symbol;Acc:MGI:1321159] | Ppp2ca | Enriched in dLGN | 1.259 | 2.779E-08 | 828.3 |
| ENSMUSG00000056004 | endosome-lysosome associated apoptosis and autophagy regulator family member 2 [Source:MGI Symbol;Acc:MGI:2443264] | Elapor2 | Enriched in dLGN | 1.258 | 0.000002224 | 229.7 |
| ENSMUSG00000021748 | pyruvate dehydrogenase (lipoamide) beta [Source:MGI Symbol;Acc:MGI:1915513] | Pdhb | Enriched in dLGN | 1.255 | 9.81E-08 | 160.8 |
| ENSMUSG00000033998 | potassium channel, subfamily K, member 1 [Source:MGI Symbol;Acc:MGI:109322] | Kcnk1 | Enriched in dLGN | 1.255 | 0.000004612 | 107.5 |

| | | | | | | |
|---|---|---|---|---|---|---|
| ENSMUSG00000026895 | NADH:ubiquinone oxidoreductase subunit A8 [Source:MGI Symbol;Acc:MGI:1915625] | Ndufa8 | Enriched in dLGN | 1.252 | 2.045E-12 | 174.6 |
| ENSMUSG00000033294 | NOC4 like [Source:MGI Symbol;Acc:MGI:2140843] | Noc4l | Enriched in dLGN | 1.251 | 0.00005427 | 33.71 |
| ENSMUSG00000017167 | contactin associated protein-like 1 [Source:MGI Symbol;Acc:MGI:1858201] | Cntnap1 | Enriched in dLGN | 1.25 | 0.0001246 | 622.8 |
| ENSMUSG00000027254 | microtubule-associated protein 1 A [Source:MGI Symbol;Acc:MGI:1306776] | Map1a | Enriched in dLGN | 1.246 | 0.0001713 | 6967 |
| ENSMUSG00000040136 | ATP-binding cassette, sub-family C (CFTR/MRP), member 8 [Source:MGI Symbol;Acc:MGI:1352629] | Abcc8 | Enriched in dLGN | 1.241 | 0.0001419 | 86.74 |
| ENSMUSG00000023033 | sodium channel, voltage-gated, type VIII, alpha [Source:MGI Symbol;Acc:MGI:103169] | Scn8a | Enriched in dLGN | 1.239 | 1.772E-11 | 1192 |
| ENSMUSG00000028447 | dynactin 3 [Source:MGI Symbol;Acc:MGI:1859251] | Dctn3 | Enriched in dLGN | 1.239 | 2.007E-10 | 123.3 |
| ENSMUSG00000027206 | COP9 signalosome subunit 2 [Source:MGI Symbol;Acc:MGI:1330276] | Cops2 | Enriched in dLGN | 1.236 | 1.099E-09 | 139.2 |
| ENSMUSG00000047649 | CD3E antigen, epsilon polypeptide associated protein [Source:MGI Symbol;Acc:MGI:1917583] | Cd3eap | Enriched in dLGN | 1.229 | 0.00003899 | 71.75 |
| ENSMUSG00000028528 | DnaJ heat shock protein family (Hsp40) member C6 [Source:MGI Symbol;Acc:MGI:1919935] | Dnajc6 | Enriched in dLGN | 1.228 | 1.705E-11 | 781.5 |
| ENSMUSG00000023904 | host cell factor C1 regulator 1 (XPO1-dependent) [Source:MGI Symbol;Acc:MGI:2663619] | Hcfc1r1 | Enriched in dLGN | 1.228 | 0.00009337 | 96.23 |
| ENSMUSG00000028811 | tyrosyl-tRNA synthetase [Source:MGI Symbol;Acc:MGI:2147627] | Yars | Enriched in dLGN | 1.225 | 3.691E-25 | 315 |
| ENSMUSG00000024527 | AFG3-like AAA ATPase 2 [Source:MGI Symbol;Acc:MGI:1916847] | Afg3l2 | Enriched in dLGN | 1.225 | 2.765E-15 | 309.3 |
| ENSMUSG00000031790 | matrix metallopeptidase 15 [Source:MGI Symbol;Acc:MGI:109320] | Mmp15 | Enriched in dLGN | 1.225 | 6.757E-07 | 357.5 |
| ENSMUSG00000024130 | ATP-binding cassette, sub-family A (ABC1), member 3 [Source:MGI Symbol;Acc:MGI:1351617] | Abca3 | Enriched in dLGN | 1.224 | 5.937E-15 | 605.6 |
| ENSMUSG00000027189 | tripartite motif-containing 44 [Source:MGI Symbol;Acc:MGI:1931835] | Trim44 | Enriched in dLGN | 1.223 | 4.255E-07 | 2112 |
| ENSMUSG00000041453 | ribosomal protein L21 [Source:MGI Symbol;Acc:MGI:1278340] | Rpl21 | Enriched in dLGN | 1.219 | 0.00005325 | 129.9 |
| ENSMUSG00000039943 | phospholipase C, beta 4 [Source:MGI Symbol;Acc:MGI:107464] | Plcb4 | Enriched in dLGN | 1.212 | 0.000002974 | 926.4 |
| ENSMUSG00000028937 | acyl-CoA thioesterase 7 [Source:MGI Symbol;Acc:MGI:1917275] | Acot7 | Enriched in dLGN | 1.211 | 0.005383 | 554 |
| ENSMUSG00000021361 | transmembrane protein 14C [Source:MGI Symbol;Acc:MGI:1913404] | Tmem14c | Enriched in dLGN | 1.205 | 1.874E-07 | 47.27 |
| ENSMUSG00000034187 | N-ethylmaleimide sensitive fusion protein [Source:MGI Symbol;Acc:MGI:104560] | Nsf | Enriched in dLGN | 1.205 | 5.763E-07 | 1114 |
| ENSMUSG00000028998 | translocase of outer mitochondrial membrane 7 [Source:MGI Symbol;Acc:MGI:1913419] | Tomm7 | Enriched in dLGN | 1.205 | 0.00004893 | 121.4 |
| ENSMUSG00000068099 | small integral membrane protein 45 [Source:MGI Symbol;Acc:MGI:1923755] | Smim45 | Enriched in dLGN | 1.204 | 1.246E-07 | 83.12 |
| ENSMUSG00000018340 | annexin A6 [Source:MGI Symbol;Acc:MGI:88255] | Anxa6 | Enriched in dLGN | 1.204 | 0.004481 | 254.2 |
| ENSMUSG00000018770 | ATP synthase, H+ transporting, mitochondrial F0 complex, subunit C3 (subunit 9) [Source:MGI Symbol;Acc:MGI:2442035] | Atp5g3 | Enriched in dLGN | 1.203 | 2.872E-07 | 265.3 |
| ENSMUSG00000062785 | potassium voltage gated channel, Shaw-related subfamily, member 3 [Source:MGI Symbol;Acc:MGI:96669] | Kcnc3 | Enriched in dLGN | 1.201 | 1.407E-09 | 437.6 |
| ENSMUSG00000052852 | receptor accessory protein 1 [Source:MGI Symbol;Acc:MGI:1098827] | Reep1 | Enriched in dLGN | 1.2 | 6.573E-09 | 542 |
| ENSMUSG00000031422 | mortality factor 4 like 2 [Source:MGI Symbol;Acc:MGI:1927167] | Morf4l2 | Enriched in dLGN | 1.198 | 2.591E-14 | 607.4 |
| ENSMUSG00000036510 | cadherin 8 [Source:MGI Symbol;Acc:MGI:107434] | Cdh8 | Enriched in dLGN | 1.197 | 0.000786 | 272 |
| ENSMUSG00000049097 | ankyrin repeat domain 34A [Source:MGI Symbol;Acc:MGI:3617846] | Ankrd34a | Enriched in dLGN | 1.195 | 0.0000851 | 84.03 |
| ENSMUSG00000022024 | SGT1, suppressor of G2 allele of SKP1 (S. cerevisiae) [Source:MGI Symbol;Acc:MGI:1915205] | Sugt1 | Enriched in dLGN | 1.194 | 2.617E-10 | 199.6 |
| ENSMUSG00000079426 | actin related protein 2/3 complex, subunit 4 [Source:MGI Symbol;Acc:MGI:1915339] | Arpc4 | Enriched in dLGN | 1.194 | 1.989E-07 | 89.11 |
| ENSMUSG00000021290 | ATP synthase membrane subunit 6.8PL [Source:MGI Symbol;Acc:MGI:1917507] | Atp5mpl | Enriched in dLGN | 1.194 | 0.003409 | 181.4 |
| ENSMUSG00000060166 | zinc finger, DHHC domain containing 8 [Source:MGI Symbol;Acc:MGI:1338012] | Zdhhc8 | Enriched in dLGN | 1.192 | 4.565E-12 | 169.3 |
| ENSMUSG00000034285 | nipsnap homolog 1 [Source:MGI Symbol;Acc:MGI:1278344] | Nipsnap1 | Enriched in dLGN | 1.191 | 1.759E-09 | 81.27 |
| ENSMUSG00000067925 | retrotransposon Gag like 8A [Source:MGI Symbol;Acc:MGI:1913408] | Rtl8a | Enriched in dLGN | 1.186 | 0.00009131 | 28.45 |
| ENSMUSG00000028719 | cytidine monophosphate (UMP-CMP) kinase 1 [Source:MGI Symbol;Acc:MGI:1913838] | Cmpk1 | Enriched in dLGN | 1.185 | 0.00000044 | 226.9 |
| ENSMUSG00000032330 | cytochrome c oxidase subunit 7A2 [Source:MGI Symbol;Acc:MGI:1316715] | Cox7a2 | Enriched in dLGN | 1.184 | 0.000003404 | 76.71 |
| ENSMUSG00000032908 | sphingosine-1-phosphate phosphatase 2 [Source:MGI Symbol;Acc:MGI:3589109] | Sgpp2 | Enriched in dLGN | 1.184 | 0.0006383 | 115.4 |
| ENSMUSG00000029108 | protocadherin 7 [Source:MGI Symbol;Acc:MGI:1860487] | Pcdh7 | Enriched in dLGN | 1.183 | 0.00001455 | 1224 |
| ENSMUSG00000063480 | SNU13 homolog, small nuclear ribonucleoprotein (U4/U6.U5) [Source:MGI Symbol;Acc:MGI:893586] | Snu13 | Enriched in dLGN | 1.183 | 0.0005013 | 38.94 |
| ENSMUSG00000027635 | DSN1 homolog, MIS12 kinetochore complex component [Source:MGI Symbol;Acc:MGI:1914184] | Dsn1 | Enriched in dLGN | 1.182 | 0.03297 | 50.45 |
| ENSMUSG00000014294 | NADH:ubiquinone oxidoreductase subunit A2 [Source:MGI Symbol;Acc:MGI:1343103] | Ndufa2 | Enriched in dLGN | 1.18 | 1.323E-07 | 115.8 |
| ENSMUSG00000025268 | MAGE family member D2 [Source:MGI Symbol;Acc:MGI:1933391] | Maged2 | Enriched in dLGN | 1.177 | 0.001301 | 189.8 |
| ENSMUSG00000045613 | cholinergic receptor, muscarinic 2, cardiac [Source:MGI Symbol;Acc:MGI:88397] | Chrm2 | Enriched in dLGN | 1.176 | 0.0008709 | 71.41 |
| ENSMUSG00000020664 | dihydrolipoamide dehydrogenase [Source:MGI Symbol;Acc:MGI:107450] | Dld | Enriched in dLGN | 1.175 | 0.00002616 | 258.4 |
| ENSMUSG00000021116 | eukaryotic translation initiation factor 2, subunit 1 alpha [Source:MGI Symbol;Acc:MGI:95299] | Eif2s1 | Enriched in dLGN | 1.175 | 0.000111 | 89.93 |
| ENSMUSG00000028555 | tetratricopeptide repeat domain 39A [Source:MGI Symbol;Acc:MGI:2444350] | Ttc39a | Enriched in dLGN | 1.171 | 0.0004524 | 25.31 |
| ENSMUSG00000023106 | density-regulated protein [Source:MGI Symbol;Acc:MGI:1915434] | Denr | Enriched in dLGN | 1.17 | 8.131E-11 | 114.8 |
| ENSMUSG00000009013 | dynein light chain LC8-type 1 [Source:MGI Symbol;Acc:MGI:1861457] | Dynll1 | Enriched in dLGN | 1.169 | 1.184E-08 | 315.9 |
| ENSMUSG00000030515 | threonyl-tRNA synthetase-like 2 [Source:MGI Symbol;Acc:MGI:2444486] | Tarsl2 | Enriched in dLGN | 1.168 | 7.667E-09 | 105.5 |
| ENSMUSG00000028373 | astrotactin 2 [Source:MGI Symbol;Acc:MGI:1889277] | Astn2 | Enriched in dLGN | 1.166 | 0.001991 | 621.4 |
| ENSMUSG00000040990 | SH3-domain kinase binding protein 1 [Source:MGI Symbol;Acc:MGI:1889583] | Sh3kbp1 | Enriched in dLGN | 1.165 | 1.152E-08 | 327.5 |
| ENSMUSG00000036186 | divergent protein kinase domain 1B [Source:MGI Symbol;Acc:MGI:1927576] | Dipk1b | Enriched in dLGN | 1.162 | 0.000339 | 28.18 |
| ENSMUSG00000059278 | N(alpha)-acetyltransferase 38, NatC auxiliary subunit [Source:MGI Symbol;Acc:MGI:1925554] | Naa38 | Enriched in dLGN | 1.162 | 0.0008934 | 31.83 |
| ENSMUSG00000061046 | hydroxyacylglutathione hydrolase-like [Source:MGI Symbol;Acc:MGI:1919877] | Haghl | Enriched in dLGN | 1.158 | 0.000004371 | 162.5 |
| ENSMUSG00000030683 | seizure related 6 homolog like 2 [Source:MGI Symbol;Acc:MGI:2385295] | Sez6l2 | Enriched in dLGN | 1.157 | 9.965E-07 | 601.7 |
| ENSMUSG00000025855 | protein kinase, cAMP dependent regulatory, type I beta [Source:MGI Symbol;Acc:MGI:97759] | Prkar1b | Enriched in dLGN | 1.156 | 2.075E-08 | 871 |
| ENSMUSG00000025781 | ATP synthase, H+ transporting, mitochondrial F1 complex, gamma polypeptide 1 [Source:MGI Symbol;Acc:MGI:1261437] | Atp5c1 | Enriched in dLGN | 1.156 | 6.039E-08 | 290 |
| ENSMUSG00000005580 | adenylate cyclase 9 [Source:MGI Symbol;Acc:MGI:108450] | Adcy9 | Enriched in dLGN | 1.156 | 2.718E-07 | 235.6 |
| ENSMUSG00000025410 | dynactin 2 [Source:MGI Symbol;Acc:MGI:107733] | Dctn2 | Enriched in dLGN | 1.155 | 2.414E-11 | 382.2 |
| ENSMUSG00000026204 | protein tyrosine phosphatase, receptor type, N [Source:MGI Symbol;Acc:MGI:102765] | Ptprn | Enriched in dLGN | 1.154 | 0.000113 | 802.8 |
| ENSMUSG00000035674 | NADH:ubiquinone oxidoreductase subunit A3 [Source:MGI Symbol;Acc:MGI:1913341] | Ndufa3 | Enriched in dLGN | 1.152 | 0.0005974 | 56.67 |
| ENSMUSG00000014077 | calcineurin-like EF hand protein 1 [Source:MGI Symbol;Acc:MGI:1927185] | Chp1 | Enriched in dLGN | 1.149 | 3.952E-07 | 152.7 |
| ENSMUSG00000031971 | centriole, cilia and spindle associated protein [Source:MGI Symbol;Acc:MGI:1920670] | Ccsap | Enriched in dLGN | 1.147 | 4.952E-10 | 95.08 |
| ENSMUSG00000033419 | synaptosomal-associated protein 91 [Source:MGI Symbol;Acc:MGI:109132] | Snap91 | Enriched in dLGN | 1.146 | 1.2E-10 | 939.7 |
| ENSMUSG00000026032 | NADH:ubiquinone oxidoreductase subunit B3 [Source:MGI Symbol;Acc:MGI:1913745] | Ndufb3 | Enriched in dLGN | 1.145 | 0.000009051 | 49.01 |
| ENSMUSG00000020869 | leucine rich repeat containing 59 [Source:MGI Symbol;Acc:MGI:2138133] | Lrrc59 | Enriched in dLGN | 1.144 | 9.885E-09 | 321.9 |
| ENSMUSG00000074656 | eukaryotic translation initiation factor 2, subunit 2 (beta) [Source:MGI Symbol;Acc:MGI:1914454] | Eif2s2 | Enriched in dLGN | 1.142 | 3.404E-07 | 150.9 |
| ENSMUSG00000027430 | D-tyrosyl-tRNA deacylase 1 [Source:MGI Symbol;Acc:MGI:1913294] | Dtd1 | Enriched in dLGN | 1.142 | 0.00008714 | 130.9 |
| ENSMUSG00000041762 | G protein-coupled receptor 155 [Source:MGI Symbol;Acc:MGI:1915776] | Gpr155 | Enriched in dLGN | 1.14 | 0.000002444 | 55.46 |
| ENSMUSG00000002808 | ependymin related protein 1 (zebrafish) [Source:MGI Symbol;Acc:MGI:2145369] | Epdr1 | Enriched in dLGN | 1.138 | 0.0000573 | 73.68 |
| ENSMUSG00000027236 | eukaryotic translation initiation factor 3, subunit J1 [Source:MGI Symbol;Acc:MGI:1925905] | Eif3j1 | Enriched in dLGN | 1.136 | 0.00001885 | 85.54 |
| ENSMUSG00000091264 | small integral membrane protein 13 [Source:MGI Symbol;Acc:MGI:2652854] | Smim13 | Enriched in dLGN | 1.134 | 0.000009194 | 346 |
| ENSMUSG00000036309 | S-phase kinase-associated protein 1 [Source:MGI Symbol;Acc:MGI:103575] | Skp1 | Enriched in dLGN | 1.134 | 0.00003074 | 534.7 |
| ENSMUSG00000033918 | presenilin associated, rhomboid-like [Source:MGI Symbol;Acc:MGI:1277152] | Parl | Enriched in dLGN | 1.132 | 7.966E-08 | 65.89 |
| ENSMUSG00000028488 | SH3-domain GRB2-like 2 [Source:MGI Symbol;Acc:MGI:700009] | Sh3gl2 | Enriched in dLGN | 1.131 | 0.0001245 | 586.9 |
| ENSMUSG00000022982 | superoxide dismutase 1, soluble [Source:MGI Symbol;Acc:MGI:98351] | Sod1 | Enriched in dLGN | 1.13 | 0.000000918 | 224.4 |
| ENSMUSG00000024425 | Nedd4 family interacting protein 1 [Source:MGI Symbol;Acc:MGI:1929601] | Ndfip1 | Enriched in dLGN | 1.13 | 0.000003121 | 747.3 |
| ENSMUSG00000036989 | tripartite motif-containing 3 [Source:MGI Symbol;Acc:MGI:1860040] | Trim3 | Enriched in dLGN | 1.129 | 0.000006596 | 276 |
| ENSMUSG00000016346 | potassium voltage-gated channel, subfamily Q, member 2 [Source:MGI Symbol;Acc:MGI:1309503] | Kcnq2 | Enriched in dLGN | 1.128 | 4.935E-08 | 1096 |
| ENSMUSG00000021606 | NADH:ubiquinone oxidoreductase core subunit S6 [Source:MGI Symbol;Acc:MGI:107932] | Ndufs6 | Enriched in dLGN | 1.126 | 7.315E-08 | 70.27 |
| ENSMUSG00000002032 | transmembrane protein 25 [Source:MGI Symbol;Acc:MGI:1918937] | Tmem25 | Enriched in dLGN | 1.126 | 0.00001461 | 70.33 |
| ENSMUSG00000021037 | AHA1, activator of heat shock protein ATPase 1 [Source:MGI Symbol;Acc:MGI:2387603] | Ahsa1 | Enriched in dLGN | 1.123 | 6.704E-11 | 199 |
| ENSMUSG00000038604 | RHO family interacting cell polarization regulator 1 [Source:MGI Symbol;Acc:MGI:1922937] | Ripor1 | Enriched in dLGN | 1.123 | 3.282E-09 | 200.2 |
| ENSMUSG00000021156 | zinc finger, MYND domain containing 11 [Source:MGI Symbol;Acc:MGI:1913755] | Zmynd11 | Enriched in dLGN | 1.122 | 0.00007331 | 980.2 |
| ENSMUSG00000019877 | serine incorporator 1 [Source:MGI Symbol;Acc:MGI:1926228] | Serinc1 | Enriched in dLGN | 1.122 | 0.000411 | 662.3 |
| ENSMUSG00000021771 | voltage-dependent anion channel 2 [Source:MGI Symbol;Acc:MGI:106915] | Vdac2 | Enriched in dLGN | 1.12 | 0.001268 | 243.1 |
| ENSMUSG00000033307 | macrophage migration inhibitory factor (glycosylation-inhibiting factor) [Source:MGI Symbol;Acc:MGI:96982] | Mif | Enriched in dLGN | 1.12 | 0.006788 | 90.39 |
| ENSMUSG00000022013 | DnaJ heat shock protein family (Hsp40) member C15 [Source:MGI Symbol;Acc:MGI:1913398] | Dnajc15 | Enriched in dLGN | 1.116 | 0.00002365 | 113.6 |
| ENSMUSG00000028062 | late endosomal/lysosomal adaptor, MAPK and MTOR activator 2 [Source:MGI Symbol;Acc:MGI:1932697] | Lamtor2 | Enriched in dLGN | 1.116 | 0.00005335 | 47.8 |
| ENSMUSG00000001472 | transcription factor 25 (basic helix-loop-helix) [Source:MGI Symbol;Acc:MGI:1914105] | Tcf25 | Enriched in dLGN | 1.115 | 1.941E-16 | 1701 |
| ENSMUSG00000031963 | BMP-binding endothelial regulator [Source:MGI Symbol;Acc:MGI:1920480] | Bmper | Enriched in dLGN | 1.115 | 0.005405 | 44.47 |
| ENSMUSG00000016319 | solute carrier family 25 (mitochondrial carrier, adenine nucleotide translocator), member 5 [Source:MGI Symbol;Acc:MGI:1353496] | Slc25a5 | Enriched in dLGN | 1.114 | 0.002724 | 198.8 |
| ENSMUSG00000000776 | polymerase (RNA) III (DNA directed) polypeptide D [Source:MGI Symbol;Acc:MGI:1914315] | Polr3d | Enriched in dLGN | 1.111 | 8.1E-09 | 108.5 |
| ENSMUSG00000024091 | vesicle-associated membrane protein, associated protein A [Source:MGI Symbol;Acc:MGI:1353561] | Vapa | Enriched in dLGN | 1.111 | 1.398E-08 | 341.6 |
| ENSMUSG00000039431 | myotubularin related protein 7 [Source:MGI Symbol;Acc:MGI:1891693] | Mtmr7 | Enriched in dLGN | 1.111 | 5.204E-07 | 199.1 |
| ENSMUSG00000025964 | a disintegrin and metallopeptidase domain 23 [Source:MGI Symbol;Acc:MGI:1345162] | Adam23 | Enriched in dLGN | 1.109 | 0.00004402 | 982.6 |
| ENSMUSG00000028532 | cache domain containing 1 [Source:MGI Symbol;Acc:MGI:2444177] | Cachd1 | Enriched in dLGN | 1.109 | 0.00009184 | 291.5 |
| ENSMUSG00000006717 | acyl-CoA thioesterase 13 [Source:MGI Symbol;Acc:MGI:1914084] | Acot13 | Enriched in dLGN | 1.109 | 0.0008203 | 30.71 |
| ENSMUSG00000021024 | proteasome subunit alpha 6 [Source:MGI Symbol;Acc:MGI:1347006] | Psma6 | Enriched in dLGN | 1.106 | 0.01049 | 71.31 |
| ENSMUSG00000039983 | coiled-coil domain containing 32 [Source:MGI Symbol;Acc:MGI:2685477] | Ccdc32 | Enriched in dLGN | 1.101 | 0.00001204 | 49.02 |
| ENSMUSG00000032306 | mannose phosphate isomerase [Source:MGI Symbol;Acc:MGI:97075] | Mpi | Enriched in dLGN | 1.1 | 1.36E-08 | 76.39 |
| ENSMUSG00000028416 | BCL2-associated athanogene 1 [Source:MGI Symbol;Acc:MGI:108047] | Bag1 | Enriched in dLGN | 1.1 | 7.803E-07 | 349.9 |
| ENSMUSG00000027305 | NADH:ubiquinone oxidoreductase complex assembly factor 1 [Source:MGI Symbol;Acc:MGI:1916952] | Ndufaf1 | Enriched in dLGN | 1.099 | 0.000001528 | 32.64 |
| ENSMUSG00000020849 | tyrosine 3-monooxygenase/tryptophan 5-monooxygenase activation protein, epsilon polypeptide [Source:MGI Symbol;Acc:MGI:894689] | Ywhae | Enriched in dLGN | 1.097 | 6.012E-07 | 2150 |
| ENSMUSG00000024099 | NADH:ubiquinone oxidoreductase core subunit V2 [Source:MGI Symbol;Acc:MGI:1920150] | Ndufv2 | Enriched in dLGN | 1.096 | 5.468E-08 | 217.5 |
| ENSMUSG00000024403 | ATPase, H+ transporting, lysosomal V1 subunit G2 [Source:MGI Symbol;Acc:MGI:1913487] | Atp6v1g2 | Enriched in dLGN | 1.094 | 0.000002225 | 338.6 |
| ENSMUSG00000087687 | PET100 homolog [Source:MGI Symbol;Acc:MGI:3615306] | Pet100 | Enriched in dLGN | 1.094 | 0.00000881 | 59.25 |
| ENSMUSG00000018848 | arginyl-tRNA synthetase [Source:MGI Symbol;Acc:MGI:1914297] | Rars | Enriched in dLGN | 1.093 | 1.196E-11 | 102.9 |
| ENSMUSG00000022241 | threonyl-tRNA synthetase [Source:MGI Symbol;Acc:MGI:106314] | Tars | Enriched in dLGN | 1.093 | 0.000000399 | 160.9 |
| ENSMUSG00000062352 | integrin beta 1 binding protein 1 [Source:MGI Symbol;Acc:MGI:1306802] | Itgb1bp1 | Enriched in dLGN | 1.093 | 0.00002403 | 59.33 |
| ENSMUSG00000009863 | succinate dehydrogenase complex, subunit B, iron sulfur (Ip) [Source:MGI Symbol;Acc:MGI:1914930] | Sdhb | Enriched in dLGN | 1.092 | 0.00000694 | 134.3 |
| ENSMUSG00000005873 | receptor accessory protein 5 [Source:MGI Symbol;Acc:MGI:1270152] | Reep5 | Enriched in dLGN | 1.092 | 0.00002868 | 705.3 |
| ENSMUSG00000001383 | zinc finger, matrin type 2 [Source:MGI Symbol;Acc:MGI:1913742] | Zmat2 | Enriched in dLGN | 1.091 | 2.549E-08 | 273.6 |
| ENSMUSG00000048332 | lipoma HMGIC fusion partner [Source:MGI Symbol;Acc:MGI:1920048] | Lhfp | Enriched in dLGN | 1.091 | 0.003493 | 294.9 |
| ENSMUSG00000017631 | active BCR-related gene [Source:MGI Symbol;Acc:MGI:107771] | Abr | Enriched in dLGN | 1.089 | 9.091E-10 | 1239 |
| ENSMUSG00000071662 | polymerase (RNA) II (DNA directed) polypeptide G [Source:MGI Symbol;Acc:MGI:1914960] | Polr2g | Enriched in dLGN | 1.085 | 0.000009443 | 40.09 |
| ENSMUSG00000024121 | ATPase, H+ transporting, lysosomal V0 subunit C [Source:MGI Symbol;Acc:MGI:88116] | Atp6v0c | Enriched in dLGN | 1.085 | 0.0001329 | 333.4 |
| ENSMUSG00000022965 | interferon gamma receptor 2 [Source:MGI Symbol;Acc:MGI:107654] | Ifngr2 | Enriched in dLGN | 1.084 | 0.0000147 | 119.8 |
| ENSMUSG00000027602 | microtubule-associated protein 1 light chain 3 alpha [Source:MGI Symbol;Acc:MGI:1915661] | Map1lc3a | Enriched in dLGN | 1.083 | 0.00003049 | 233.2 |
| ENSMUSG00000013921 | CAP-GLY domain containing linker protein 3 [Source:MGI Symbol;Acc:MGI:1923936] | Clip3 | Enriched in dLGN | 1.08 | 0.0005198 | 1062 |
| ENSMUSG00000051223 | basic leucine zipper and W2 domains 1 [Source:MGI Symbol;Acc:MGI:1914132] | Bzw1 | Enriched in dLGN | 1.078 | 0.00004023 | 393.6 |
| ENSMUSG00000042766 | tripartite motif-containing 46 [Source:MGI Symbol;Acc:MGI:2673000] | Trim46 | Enriched in dLGN | 1.076 | 3.357E-08 | 200.1 |
| ENSMUSG00000027406 | isocitrate dehydrogenase 3 (NAD+) beta [Source:MGI Symbol;Acc:MGI:2158650] | Idh3b | Enriched in dLGN | 1.076 | 0.00002989 | 236.2 |
| ENSMUSG00000049760 | mitochondrial contact site and cristae organizing system subunit 13 [Source:MGI Symbol;Acc:MGI:2442174] | Micos13 | Enriched in dLGN | 1.075 | 0.000007842 | 44.94 |
| ENSMUSG00000002102 | proteasome (prosome, macropain) 26S subunit, ATPase 3 [Source:MGI Symbol;Acc:MGI:1098754] | Psmc3 | Enriched in dLGN | 1.075 | 0.00001057 | 304.4 |
| ENSMUSG00000062006 | ribosomal protein L34 [Source:MGI Symbol;Acc:MGI:1915686] | Rpl34 | Enriched in dLGN | 1.075 | 0.002204 | 135.1 |

| | | | | | | |
|---|---|---|---|---|---|---|
| ENSMUSG00000038762 | ATP-binding cassette, sub-family F (GCN20), member 1 [Source:MGI Symbol;Acc:MGI:1351658] | Abcf1 | Enriched in dLGN | 1.074 | 1.92E-09 | 576.3 |
| ENSMUSG00000030946 | phospholysine phosphohistidine inorganic pyrophosphate phosphatase [Source:MGI Symbol;Acc:MGI:1923679] | Lhpp | Enriched in dLGN | 1.073 | 0.0001148 | 34.61 |
| ENSMUSG00000024942 | calpain 1 [Source:MGI Symbol;Acc:MGI:88263] | Capn1 | Enriched in dLGN | 1.073 | 0.005126 | 59.73 |
| ENSMUSG00000036398 | protein phosphatase 1, regulatory inhibitor subunit 11 [Source:MGI Symbol;Acc:MGI:1923747] | Ppp1r11 | Enriched in dLGN | 1.07 | 0.000002564 | 110.1 |
| ENSMUSG00000024044 | erythrocyte membrane protein band 4.1 like 3 [Source:MGI Symbol;Acc:MGI:103008] | Epb41l3 | Enriched in dLGN | 1.07 | 0.00005961 | 1430 |
| ENSMUSG00000028692 | aldo-keto reductase family 1, member A1 (aldehyde reductase) [Source:MGI Symbol;Acc:MGI:1929955] | Akr1a1 | Enriched in dLGN | 1.069 | 4.027E-09 | 579.7 |
| ENSMUSG00000016541 | ataxin 10 [Source:MGI Symbol;Acc:MGI:1859293] | Atxn10 | Enriched in dLGN | 1.068 | 0.0007674 | 415.1 |
| ENSMUSG00000014769 | proteasome (prosome, macropain) subunit, beta type 1 [Source:MGI Symbol;Acc:MGI:104884] | Psmb1 | Enriched in dLGN | 1.066 | 0.000008684 | 166 |
| ENSMUSG00000026527 | regulator of G protein signaling 7 [Source:MGI Symbol;Acc:MGI:1346089] | Rgs7 | Enriched in dLGN | 1.066 | 0.007355 | 427.5 |
| ENSMUSG00000024581 | N-ethylmaleimide sensitive fusion protein attachment protein gamma [Source:MGI Symbol;Acc:MGI:104561] | Napg | Enriched in dLGN | 1.065 | 1.289E-10 | 325.8 |
| ENSMUSG00000019785 | clavesin 2 [Source:MGI Symbol;Acc:MGI:2443223] | Clvs2 | Enriched in dLGN | 1.063 | 0.0002591 | 86.26 |
| ENSMUSG00000060739 | NSA2 ribosome biogenesis homolog [Source:MGI Symbol;Acc:MGI:1913883] | Nsa2 | Enriched in dLGN | 1.058 | 0.0001044 | 65.27 |
| ENSMUSG00000045287 | reticulon 4 receptor-like 1 [Source:MGI Symbol;Acc:MGI:2661375] | Rtn4rl1 | Enriched in dLGN | 1.057 | 0.000005726 | 303.8 |
| ENSMUSG00000035864 | synaptotagmin I [Source:MGI Symbol;Acc:MGI:99667] | Syt1 | Enriched in dLGN | 1.057 | 0.00001189 | 3835 |
| ENSMUSG00000030092 | contactin 6 [Source:MGI Symbol;Acc:MGI:1858223] | Cntn6 | Enriched in dLGN | 1.057 | 0.002073 | 202.1 |
| ENSMUSG00000069237 | family with sequence similarity 8, member A1 [Source:MGI Symbol;Acc:MGI:2145496] | Fam8a1 | Enriched in dLGN | 1.056 | 0.000007351 | 177.6 |
| ENSMUSG00000030738 | eukaryotic translation initiation factor 3, subunit C [Source:MGI Symbol;Acc:MGI:1926966] | Eif3c | Enriched in dLGN | 1.055 | 0.00003076 | 682.4 |
| ENSMUSG00000033585 | necdin, MAGE family member [Source:MGI Symbol;Acc:MGI:97290] | Ndn | Enriched in dLGN | 1.055 | 0.001431 | 549 |
| ENSMUSG00000036067 | solute carrier family 2 (facilitated glucose transporter), member 6 [Source:MGI Symbol;Acc:MGI:2443286] | Slc2a6 | Enriched in dLGN | 1.054 | 0.001108 | 33.47 |
| ENSMUSG00000020267 | histidine triad nucleotide binding protein 1 [Source:MGI Symbol;Acc:MGI:1321133] | Hint1 | Enriched in dLGN | 1.053 | 0.00002073 | 255.7 |
| ENSMUSG00000020153 | NADH:ubiquinone oxidoreductase core subunit S7 [Source:MGI Symbol;Acc:MGI:1922656] | Ndufs7 | Enriched in dLGN | 1.052 | 0.000006201 | 124 |
| ENSMUSG00000024583 | thioredoxin-like 1 [Source:MGI Symbol;Acc:MGI:1860078] | Txnl1 | Enriched in dLGN | 1.05 | 8.917E-08 | 112.7 |
| ENSMUSG00000020340 | cytoplasmic FMR1 interacting protein 2 [Source:MGI Symbol;Acc:MGI:1924134] | Cyfip2 | Enriched in dLGN | 1.049 | 6.477E-07 | 1444 |
| ENSMUSG00000039601 | regulator of calcineurin 2 [Source:MGI Symbol;Acc:MGI:1858219] | Rcan2 | Enriched in dLGN | 1.043 | 0.00005694 | 311.5 |
| ENSMUSG00000060240 | cell cycle exit and neuronal differentiation 1 [Source:MGI Symbol;Acc:MGI:1929898] | Cend1 | Enriched in dLGN | 1.043 | 0.009235 | 473.2 |
| ENSMUSG00000033460 | armadillo repeat containing, X-linked 1 [Source:MGI Symbol;Acc:MGI:1925498] | Armcx1 | Enriched in dLGN | 1.042 | 0.000000633 | 200.9 |
| ENSMUSG00000078713 | translocase of outer mitochondrial membrane 5 [Source:MGI Symbol;Acc:MGI:1915762] | Tomm5 | Enriched in dLGN | 1.042 | 0.000699 | 36.78 |
| ENSMUSG00000079480 | peptidyl-prolyl cis/trans isomerase, NIMA-interacting, 4 (parvulin) [Source:MGI Symbol;Acc:MGI:1916963] | Pin4 | Enriched in dLGN | 1.042 | 0.003947 | 13.6 |
| ENSMUSG00000028426 | RAD23 homolog B, nucleotide excision repair protein [Source:MGI Symbol;Acc:MGI:105128] | Rad23b | Enriched in dLGN | 1.04 | 3.036E-10 | 394.2 |
| ENSMUSG00000026833 | olfactomedin 1 [Source:MGI Symbol;Acc:MGI:1860437] | Olfm1 | Enriched in dLGN | 1.039 | 0.00001077 | 1863 |
| ENSMUSG00000068882 | small RNA binding exonuclease protection factor La [Source:MGI Symbol;Acc:MGI:98423] | Ssb | Enriched in dLGN | 1.039 | 0.00002213 | 214.7 |
| ENSMUSG00000021816 | protein phosphatase 3, catalytic subunit, beta isoform [Source:MGI Symbol;Acc:MGI:107163] | Ppp3cb | Enriched in dLGN | 1.038 | 3.814E-12 | 533.5 |
| ENSMUSG00000024208 | ubiquinol-cytochrome c reductase complex assembly factor 2 [Source:MGI Symbol;Acc:MGI:1914517] | Uqcc2 | Enriched in dLGN | 1.038 | 0.00023 | 101.2 |
| ENSMUSG00000017778 | cytochrome c oxidase subunit 7C [Source:MGI Symbol;Acc:MGI:103226] | Cox7c | Enriched in dLGN | 1.038 | 0.0005213 | 35.38 |
| ENSMUSG00000035835 | phospholipid phosphatase related 3 [Source:MGI Symbol;Acc:MGI:2388640] | Plppr3 | Enriched in dLGN | 1.036 | 0.00003967 | 329.9 |
| ENSMUSG00000057181 | RIKEN cDNA 5730455P16 gene [Source:MGI Symbol;Acc:MGI:1917841] | 5730455P16Rik | Enriched in dLGN | 1.033 | 4.261E-07 | 171.8 |
| ENSMUSG00000044442 | N-6 adenine-specific DNA methyltransferase 1 (putative) [Source:MGI Symbol;Acc:MGI:1915018] | N6amt1 | Enriched in dLGN | 1.031 | 0.00002223 | 39.22 |
| ENSMUSG00000029036 | ATPase family, AAA domain containing 3A [Source:MGI Symbol;Acc:MGI:1919214] | Atad3a | Enriched in dLGN | 1.03 | 4.059E-07 | 104.5 |
| ENSMUSG00000026188 | transmembrane protein 169 [Source:MGI Symbol;Acc:MGI:2442781] | Tmem169 | Enriched in dLGN | 1.026 | 2.339E-07 | 139.3 |
| ENSMUSG00000029430 | RAN, member RAS oncogene family [Source:MGI Symbol;Acc:MGI:1333112] | Ran | Enriched in dLGN | 1.026 | 0.0001621 | 230.2 |
| ENSMUSG00000052926 | ribonuclease H2, large subunit [Source:MGI Symbol;Acc:MGI:1916974] | Rnaseh2a | Enriched in dLGN | 1.025 | 0.00001428 | 40.8 |
| ENSMUSG00000020297 | neuron specific gene family member 2 [Source:MGI Symbol;Acc:MGI:1202070] | Nsg2 | Enriched in dLGN | 1.023 | 3.938E-09 | 922.9 |
| ENSMUSG00000028478 | clathrin, light polypeptide (Lca) [Source:MGI Symbol;Acc:MGI:894297] | Clta | Enriched in dLGN | 1.022 | 2.25E-10 | 866.9 |
| ENSMUSG00000027184 | cell cycle associated protein 1 [Source:MGI Symbol;Acc:MGI:1858234] | Caprin1 | Enriched in dLGN | 1.021 | 4.443E-07 | 681.9 |
| ENSMUSG00000031950 | gamma-aminobutyric acid (GABA) A receptor-associated protein-like 2 [Source:MGI Symbol;Acc:MGI:1890602] | Gabarapl2 | Enriched in dLGN | 1.021 | 0.000008425 | 147.2 |
| ENSMUSG00000016427 | NADH:ubiquinone oxidoreductase subunit A1 [Source:MGI Symbol;Acc:MGI:1929511] | Ndufa1 | Enriched in dLGN | 1.02 | 0.00001773 | 84.15 |
| ENSMUSG00000113902 | NADH:ubiquinone oxidoreductase subunit B1 [Source:MGI Symbol;Acc:MGI:3780865] | Ndufb1 | Enriched in dLGN | 1.02 | 0.003156 | 43.14 |
| ENSMUSG00000025260 | hydroxysteroid (17-beta) dehydrogenase 10 [Source:MGI Symbol;Acc:MGI:1333871] | Hsd17b10 | Enriched in dLGN | 1.02 | 0.009836 | 20.9 |
| ENSMUSG00000039347 | ATPase, H+ transporting, lysosomal V0 subunit E2 [Source:MGI Symbol;Acc:MGI:1923502] | Atp6v0e2 | Enriched in dLGN | 1.019 | 0.000000253 | 554.6 |
| ENSMUSG00000030301 | coiled-coil domain containing 91 [Source:MGI Symbol;Acc:MGI:1914265] | Ccdc91 | Enriched in dLGN | 1.018 | 0.000001902 | 147.9 |
| ENSMUSG00000028470 | histidine triad nucleotide binding protein 2 [Source:MGI Symbol;Acc:MGI:1916167] | Hint2 | Enriched in dLGN | 1.014 | 0.002237 | 22.68 |
| ENSMUSG00000059343 | aldolase 1 A, retrogene 1 [Source:MGI Symbol;Acc:MGI:2447811] | Aldoart1 | Enriched in dLGN | 1.012 | 0.009246 | 19.45 |
| ENSMUSG00000026103 | glutaminase [Source:MGI Symbol;Acc:MGI:95752] | Gls | Enriched in dLGN | 1.011 | 8.243E-07 | 697 |
| ENSMUSG00000041997 | tousled-like kinase 1 [Source:MGI Symbol;Acc:MGI:2441683] | Tlk1 | Enriched in dLGN | 1.011 | 0.000001689 | 318.7 |
| ENSMUSG00000023932 | cell division cycle 5-like (S. pombe) [Source:MGI Symbol;Acc:MGI:1918952] | Cdc5l | Enriched in dLGN | 1.011 | 0.000001801 | 151.3 |
| ENSMUSG00000026748 | plexin domain containing 2 [Source:MGI Symbol;Acc:MGI:1914698] | Plxdc2 | Enriched in dLGN | 1.009 | 0.001602 | 262.9 |
| ENSMUSG00000027712 | annexin A5 [Source:MGI Symbol;Acc:MGI:106008] | Anxa5 | Enriched in dLGN | 1.009 | 0.01373 | 102.3 |
| ENSMUSG00000025287 | acyl-CoA thioesterase 9 [Source:MGI Symbol;Acc:MGI:1928939] | Acot9 | Enriched in dLGN | 1.007 | 0.02462 | 23.21 |
| ENSMUSG00000020312 | SHC (Src homology 2 domain containing) transforming protein 2 [Source:MGI Symbol;Acc:MGI:106180] | Shc2 | Enriched in dLGN | 1.003 | 1.653E-07 | 242.8 |
| ENSMUSG00000000171 | succinate dehydrogenase complex, subunit D, integral membrane protein [Source:MGI Symbol;Acc:MGI:1914175] | Sdhd | Enriched in dLGN | 1.002 | 6.69E-08 | 85.24 |
| ENSMUSG00000015092 | endothelial differentiation-related factor 1 [Source:MGI Symbol;Acc:MGI:1891227] | Edf1 | Enriched in dLGN | 1.002 | 0.000002891 | 190.3 |
| ENSMUSG00000004561 | methyltransferase like 17 [Source:MGI Symbol;Acc:MGI:1098577] | Mettl17 | Enriched in retina | -1.005 | 0.03401 | 47.96 |
| ENSMUSG00000027341 | transmembrane protein 230 [Source:MGI Symbol;Acc:MGI:1917862] | Tmem230 | Enriched in retina | -1.01 | 0.0004803 | 38.18 |
| ENSMUSG00000028013 | pyrophosphatase (inorganic) 2 [Source:MGI Symbol;Acc:MGI:1922026] | Ppa2 | Enriched in retina | -1.016 | 0.00001256 | 141.1 |
| ENSMUSG00000029422 | arginine/serine-rich coiled-coil 2 [Source:MGI Symbol;Acc:MGI:1913489] | Rsrc2 | Enriched in retina | -1.023 | 0.0003313 | 465.1 |
| ENSMUSG00000032384 | casein kinase 1, gamma 1 [Source:MGI Symbol;Acc:MGI:2660884] | Csnk1g1 | Enriched in retina | -1.023 | 0.000002004 | 220.4 |
| ENSMUSG00000045095 | membrane associated guanylate kinase, WW and PDZ domain containing 1 [Source:MGI Symbol;Acc:MGI:1203522] | Magi1 | Enriched in retina | -1.032 | 0.00001315 | 743.1 |
| ENSMUSG00000031399 | FAM3 metabolism regulating signaling molecule A [Source:MGI Symbol;Acc:MGI:1913544] | Fam3a | Enriched in retina | -1.036 | 0.001349 | 44.59 |
| ENSMUSG00000032558 | nephronophthisis 3 (adolescent) [Source:MGI Symbol;Acc:MGI:1921275] | Nphp3 | Enriched in retina | -1.037 | 0.03282 | 34.65 |
| ENSMUSG00000022668 | GTP-binding protein 8 (putative) [Source:MGI Symbol;Acc:MGI:1913317] | Gtpbp8 | Enriched in retina | -1.039 | 0.004974 | 56.85 |
| ENSMUSG00000028576 | intraflagellar transport 74 [Source:MGI Symbol;Acc:MGI:1914944] | Ift74 | Enriched in retina | -1.041 | 0.0003379 | 79.32 |
| ENSMUSG00000015980 | leucine rich repeat containing 27 [Source:MGI Symbol;Acc:MGI:1923862] | Lrrc27 | Enriched in retina | -1.045 | 0.004273 | 44.92 |
| ENSMUSG00000039756 | deoxynucleotidyltransferase, terminal, interacting protein 2 [Source:MGI Symbol;Acc:MGI:1923173] | Dnttip2 | Enriched in retina | -1.046 | 0.0006241 | 112.1 |
| ENSMUSG00000024054 | SMC hinge domain containing 1 [Source:MGI Symbol;Acc:MGI:1921605] | Smchd1 | Enriched in retina | -1.047 | 0.000001801 | 205.8 |
| ENSMUSG00000056832 | tetratricopeptide repeat domain 26 [Source:MGI Symbol;Acc:MGI:2444853] | Ttc26 | Enriched in retina | -1.048 | 0.009274 | 27.07 |
| ENSMUSG00000036572 | UPF3 regulator of nonsense transcripts homolog B (yeast) [Source:MGI Symbol;Acc:MGI:1915384] | Upf3b | Enriched in retina | -1.049 | 0.001044 | 271.8 |
| ENSMUSG00000033799 | transcription activation suppressor family member 2 [Source:MGI Symbol;Acc:MGI:2145274] | Tasor2 | Enriched in retina | -1.051 | 0.000009131 | 156.7 |
| ENSMUSG00000003541 | immediate early response 3 [Source:MGI Symbol;Acc:MGI:104814] | Ier3 | Enriched in retina | -1.053 | 0.003185 | 21.94 |
| ENSMUSG00000034707 | glucosamine (N-acetyl)-6-sulfatase [Source:MGI Symbol;Acc:MGI:1922862] | Gns | Enriched in retina | -1.058 | 3.584E-09 | 343.2 |
| ENSMUSG00000036023 | poly (ADP-ribose) polymerase family, member 2 [Source:MGI Symbol;Acc:MGI:1341112] | Parp2 | Enriched in retina | -1.06 | 0.006061 | 46.73 |
| ENSMUSG00000035407 | KN motif and ankyrin repeat domains 4 [Source:MGI Symbol;Acc:MGI:3043381] | Kank4 | Enriched in retina | -1.061 | 0.01487 | 22.26 |
| ENSMUSG00000030592 | ryanodine receptor 1, skeletal muscle [Source:MGI Symbol;Acc:MGI:99659] | Ryr1 | Enriched in retina | -1.066 | 0.04343 | 72.59 |
| ENSMUSG00000025035 | ADP-ribosylation factor-like 3 [Source:MGI Symbol;Acc:MGI:1929699] | Arl3 | Enriched in retina | -1.086 | 1.401E-07 | 316.8 |
| ENSMUSG00000068250 | antagonist of mitotic exit network 1 [Source:MGI Symbol;Acc:MGI:2442933] | Amn1 | Enriched in retina | -1.089 | 0.02829 | 40.08 |
| ENSMUSG00000024891 | solute carrier family 29 (nucleoside transporters), member 2 [Source:MGI Symbol;Acc:MGI:1345278] | Slc29a2 | Enriched in retina | -1.14 | 0.00008799 | 40.3 |
| ENSMUSG00000045968 | transmembrane epididymal family member 2 [Source:MGI Symbol;Acc:MGI:1923273] | Teddm2 | Enriched in retina | -1.161 | 0.006267 | 15.55 |
| ENSMUSG00000031631 | cilia and flagella associated protein 97 [Source:MGI Symbol;Acc:MGI:1914006] | Cfap97 | Enriched in retina | -1.17 | 0.0000261 | 94.5 |
| ENSMUSG00000049878 | rearranged L-myc fusion sequence [Source:MGI Symbol;Acc:MGI:1924705] | Rlf | Enriched in retina | -1.189 | 3.209E-08 | 212.5 |
| ENSMUSG00000049946 | cDNA sequence BC030500 [Source:MGI Symbol;Acc:MGI:2447770] | BC030500 | Enriched in retina | -1.2 | 0.001639 | 32.35 |
| ENSMUSG00000033543 | general transcription factor II A, 2 [Source:MGI Symbol;Acc:MGI:1933289] | Gtf2a2 | Enriched in retina | -1.214 | 0.001327 | 44.7 |
| ENSMUSG00000036676 | transmembrane and tetratricopeptide repeat containing 3 [Source:MGI Symbol;Acc:MGI:3036255] | Tmtc3 | Enriched in retina | -1.224 | 3.651E-07 | 82.34 |
| ENSMUSG00000034274 | THO complex 5 [Source:MGI Symbol;Acc:MGI:1351333] | Thoc5 | Enriched in retina | -1.225 | 1.174E-07 | 125.8 |
| ENSMUSG00000066568 | LSM14A mRNA processing body assembly factor [Source:MGI Symbol;Acc:MGI:1914320] | Lsm14a | Enriched in retina | -1.244 | 1.601E-18 | 280.1 |
| ENSMUSG00000036879 | phosphorylase kinase beta [Source:MGI Symbol;Acc:MGI:97578] | Phkb | Enriched in retina | -1.291 | 0.00001725 | 174.5 |
| ENSMUSG00000028051 | hyperpolarization-activated, cyclic nucleotide-gated K+ 3 [Source:MGI Symbol;Acc:MGI:1298211] | Hcn3 | Enriched in retina | -1.294 | 0.000003652 | 158.3 |
| ENSMUSG00000073792 | asparagine-linked glycosylation 6 (alpha-1,3,-glucosyltransferase) [Source:MGI Symbol;Acc:MGI:2444031] | Alg6 | Enriched in retina | -1.296 | 0.001078 | 28.53 |
| ENSMUSG00000005886 | nuclear receptor coactivator 2 [Source:MGI Symbol;Acc:MGI:1276533] | Ncoa2 | Enriched in retina | -1.301 | 9.98E-16 | 793.7 |
| ENSMUSG00000040565 | B-TFIID TATA-box binding protein associated factor 1 [Source:MGI Symbol;Acc:MGI:2147538] | Btaf1 | Enriched in retina | -1.332 | 8.858E-07 | 279.3 |
| ENSMUSG00000031536 | polymerase (DNA directed), beta [Source:MGI Symbol;Acc:MGI:97740] | Polb | Enriched in retina | -1.411 | 0.0001427 | 159.7 |
| ENSMUSG00000031872 | brain expressed, associated with Nedd4, 1 [Source:MGI Symbol;Acc:MGI:1929597] | Bean1 | Enriched in retina | -1.452 | 0.0001145 | 45.05 |
| ENSMUSG00000054469 | lysocardiolipin acyltransferase 1 [Source:MGI Symbol;Acc:MGI:2684937] | Lclat1 | Enriched in retina | -1.455 | 2.435E-07 | 128 |
| ENSMUSG00000020908 | myosin, heavy polypeptide 3, skeletal muscle, embryonic [Source:MGI Symbol;Acc:MGI:1339709] | Myh3 | Enriched in retina | -1.459 | 0.04595 | 16.98 |
| ENSMUSG00000042508 | cyclin D binding myb like transcription factor 1 [Source:MGI Symbol;Acc:MGI:1344415] | Dmtf1 | Enriched in retina | -1.466 | 0.00001981 | 197.7 |
| ENSMUSG00000027014 | CWC22 spliceosome-associated protein [Source:MGI Symbol;Acc:MGI:2136773] | Cwc22 | Enriched in retina | -1.471 | 0.000487 | 206.1 |
| ENSMUSG00000034075 | zinc finger, DHHC domain containing 5 [Source:MGI Symbol;Acc:MGI:1923573] | Zdhhc5 | Enriched in retina | -1.471 | 4.275E-17 | 421.5 |
| ENSMUSG00000037801 | IQ motif containing H [Source:MGI Symbol;Acc:MGI:1925500] | Iqch | Enriched in retina | -1.482 | 0.02841 | 9.49 |
| ENSMUSG00000053914 | lysine (K)-specific demethylase 4D [Source:MGI Symbol;Acc:MGI:3606484] | Kdm4d | Enriched in retina | -1.489 | 0.01198 | 20.63 |
| ENSMUSG00000070806 | zinc finger, MYND domain containing 12 [Source:MGI Symbol;Acc:MGI:2140259] | Zmynd12 | Enriched in retina | -1.546 | 0.01628 | 9.762 |
| ENSMUSG00000027674 | peroxisomal biogenesis factor 5-like [Source:MGI Symbol;Acc:MGI:1916672] | Pex5l | Enriched in retina | -1.552 | 0.00003998 | 789.1 |
| ENSMUSG00000030110 | ret proto-oncogene [Source:MGI Symbol;Acc:MGI:97902] | Ret | Enriched in retina | -1.59 | 9.734E-08 | 188.3 |
| ENSMUSG00000056851 | poly(rC) binding protein 2 [Source:MGI Symbol;Acc:MGI:108202] | Pcbp2 | Enriched in retina | -1.611 | 2.627E-17 | 828.1 |
| ENSMUSG00000042557 | transcriptional regulator, SIN3A (yeast) [Source:MGI Symbol;Acc:MGI:107157] | Sin3a | Enriched in retina | -1.63 | 6.246E-24 | 223.5 |
| ENSMUSG00000014846 | tubulin polymerization-promoting protein family member 3 [Source:MGI Symbol;Acc:MGI:1915221] | Tppp3 | Enriched in retina | -1.658 | 1.814E-07 | 139.8 |
| ENSMUSG00000013846 | ST3 beta-galactoside alpha-2,3-sialyltransferase 1 [Source:MGI Symbol;Acc:MGI:98304] | St3gal1 | Enriched in retina | -1.659 | 4.704E-14 | 259.6 |
| ENSMUSG00000026739 | Bmi1 polycomb ring finger oncogene [Source:MGI Symbol;Acc:MGI:88174] | Bmi1 | Enriched in retina | -1.712 | 2.277E-15 | 140.4 |
| ENSMUSG00000041548 | heat shock protein 8 [Source:MGI Symbol;Acc:MGI:2135756] | Hspb8 | Enriched in retina | -1.724 | 0.0003731 | 19.43 |
| ENSMUSG00000049511 | 5-hydroxytryptamine (serotonin) receptor 1B [Source:MGI Symbol;Acc:MGI:96274] | Htr1b | Enriched in retina | -1.748 | 7.804E-07 | 142.5 |
| ENSMUSG00000038079 | transmembrane protein 237 [Source:MGI Symbol;Acc:MGI:2138365] | Tmem237 | Enriched in retina | -1.757 | 1.63E-20 | 98.82 |
| ENSMUSG00000024666 | transmembrane protein 138 [Source:MGI Symbol;Acc:MGI:1920232] | Tmem138 | Enriched in retina | -1.776 | 0.00004607 | 25.53 |
| ENSMUSG00000032599 | inositol hexaphosphate kinase 2 [Source:MGI Symbol;Acc:MGI:1923750] | Ip6k2 | Enriched in retina | -1.783 | 1.509E-10 | 280.5 |
| ENSMUSG00000034402 | potassium voltage-gated channel, subfamily H (eag-related), member 5 [Source:MGI Symbol;Acc:MGI:3584508] | Kcnh5 | Enriched in retina | -1.837 | 3.272E-10 | 240.3 |
| ENSMUSG00000035517 | tudor domain containing 7 [Source:MGI Symbol;Acc:MGI:2140279] | Tdrd7 | Enriched in retina | -1.914 | 3.274E-29 | 468.1 |
| ENSMUSG00000075334 | reprimo, TP53 dependent G2 arrest mediator candidate [Source:MGI Symbol;Acc:MGI:1915124] | Rprm | Enriched in retina | -1.917 | 0.000008653 | 67.3 |
| ENSMUSG00000052301 | double C2, alpha [Source:MGI Symbol;Acc:MGI:109446] | Doc2a | Enriched in retina | -1.918 | 0.0006384 | 56.86 |
| ENSMUSG00000056260 | ligand dependent nuclear receptor interacting factor 1 [Source:MGI Symbol;Acc:MGI:2445214] | Lrif1 | Enriched in retina | -1.973 | 0.00001062 | 40.61 |
| ENSMUSG00000068615 | gap junction protein, delta 2 [Source:MGI Symbol;Acc:MGI:1334209] | Gjd2 | Enriched in retina | -1.988 | 1.04E-22 | 192.5 |
| ENSMUSG00000047141 | zinc finger protein 654 [Source:MGI Symbol;Acc:MGI:1919270] | Zfp654 | Enriched in retina | -1.999 | 5.108E-14 | 224.3 |
| ENSMUSG00000024856 | CDK2-associated protein 2 [Source:MGI Symbol;Acc:MGI:1098779] | Cdk2ap2 | Enriched in retina | -2.043 | 2.261E-09 | 59.04 |

| | | | | | | |
|---|---|---|---|---|---|---|
| ENSMUSG00000061462 | obscurin, cytoskeletal calmodulin and titin-interacting RhoGEF [Source:MGI Symbol;Acc:MGI:2681862] | Obscn | Enriched in retina | -2.05 | 0.00004463 | 66.02 |
| ENSMUSG00000036585 | fibroblast growth factor 1 [Source:MGI Symbol;Acc:MGI:95515] | Fgf1 | Enriched in retina | -2.061 | 9.721E-08 | 148.8 |
| ENSMUSG00000030189 | Y box protein 3 [Source:MGI Symbol;Acc:MGI:2137670] | Ybx3 | Enriched in retina | -2.065 | 7.667E-13 | 527 |
| ENSMUSG00000072572 | solute carrier family 39 (zinc transporter), member 2 [Source:MGI Symbol;Acc:MGI:2684326] | Slc39a2 | Enriched in retina | -2.072 | 0.004813 | 19.9 |
| ENSMUSG00000030703 | glycerophosphodiester phosphodiesterase domain containing 3 [Source:MGI Symbol;Acc:MGI:1915866] | Gdpd3 | Enriched in retina | -2.102 | 0.005735 | 6.428 |
| ENSMUSG00000036995 | ArfGAP with SH3 domain, ankyrin repeat and PH domain 3 [Source:MGI Symbol;Acc:MGI:2684986] | Asap3 | Enriched in retina | -2.202 | 7.994E-10 | 47.68 |
| ENSMUSG00000040703 | cytochrome P450, family 2, subfamily s, polypeptide 1 [Source:MGI Symbol;Acc:MGI:1921384] | Cyp2s1 | Enriched in retina | -2.204 | 6.176E-07 | 19.07 |
| ENSMUSG00000027971 | N-deacetylase/N-sulfotransferase (heparin glucosaminyl) 4 [Source:MGI Symbol;Acc:MGI:1932545] | Ndst4 | Enriched in retina | -2.244 | 0.00001257 | 48.89 |
| ENSMUSG00000032192 | guanine nucleotide binding protein (G protein), beta 5 [Source:MGI Symbol;Acc:MGI:101848] | Gnb5 | Enriched in retina | -2.247 | 2.933E-25 | 441.8 |
| ENSMUSG00000060969 | Iroquois homeobox 1 [Source:MGI Symbol;Acc:MGI:1197515] | Irx1 | Enriched in retina | -2.302 | 0.00004547 | 29.96 |
| ENSMUSG00000035620 | RIC8 guanine nucleotide exchange factor B [Source:MGI Symbol;Acc:MGI:2682307] | Ric8b | Enriched in retina | -2.303 | 5.787E-20 | 318 |
| ENSMUSG00000022257 | lysosomal-associated protein transmembrane 4B [Source:MGI Symbol;Acc:MGI:1890494] | Laptm4b | Enriched in retina | -2.317 | 5.115E-29 | 165.8 |
| ENSMUSG00000022722 | ADP-ribosylation factor-like 6 [Source:MGI Symbol;Acc:MGI:1927136] | Arl6 | Enriched in retina | -2.345 | 2.506E-13 | 147.9 |
| ENSMUSG00000067367 | Ly1 antibody reactive clone [Source:MGI Symbol;Acc:MGI:107470] | Lyar | Enriched in retina | -2.347 | 1.81E-21 | 128.3 |
| ENSMUSG00000041147 | breast cancer 2, early onset [Source:MGI Symbol;Acc:MGI:109337] | Brca2 | Enriched in retina | -2.37 | 1.722E-11 | 53.68 |
| ENSMUSG00000000552 | zinc finger protein 385A [Source:MGI Symbol;Acc:MGI:1352495] | Zfp385a | Enriched in retina | -2.385 | 3.543E-37 | 618.4 |
| ENSMUSG00000028476 | reversion-inducing-cysteine-rich protein with kazal motifs [Source:MGI Symbol;Acc:MGI:1855698] | Reck | Enriched in retina | -2.396 | 5.488E-08 | 71.55 |
| ENSMUSG00000040489 | SRY (sex determining region Y)-box 30 [Source:MGI Symbol;Acc:MGI:1341157] | Sox30 | Enriched in retina | -2.43 | 0.002142 | 11.69 |
| ENSMUSG00000107402 | RIKEN cDNA 4732416N19 gene [Source:MGI Symbol;Acc:MGI:2444634] | 4732416N19Rik | Enriched in retina | -2.498 | 0.008044 | 6.196 |
| ENSMUSG00000031734 | Iroquois related homeobox 3 [Source:MGI Symbol;Acc:MGI:1197522] | Irx3 | Enriched in retina | -2.545 | 2.654E-07 | 16.38 |
| ENSMUSG00000020014 | cilia and flagella associated protein 54 [Source:MGI Symbol;Acc:MGI:1922208] | Cfap54 | Enriched in retina | -2.548 | 1.184E-07 | 91.99 |
| ENSMUSG00000038457 | transmembrane protein 255B [Source:MGI Symbol;Acc:MGI:2685533] | Tmem255b | Enriched in retina | -2.575 | 1.123E-16 | 59.61 |
| ENSMUSG00000062300 | nectin cell adhesion molecule 2 [Source:MGI Symbol;Acc:MGI:97822] | Nectin2 | Enriched in retina | -2.579 | 1.081E-14 | 66.46 |
| ENSMUSG00000020806 | rhomboid 5 homolog 2 [Source:MGI Symbol;Acc:MGI:2442473] | Rhbdf2 | Enriched in retina | -2.621 | 2.829E-08 | 43.1 |
| ENSMUSG00000026463 | ATPase, Ca++ transporting, plasma membrane 4 [Source:MGI Symbol;Acc:MGI:88111] | Atp2b4 | Enriched in retina | -2.634 | 2.342E-16 | 561.4 |
| ENSMUSG00000078963 | heat shock factor binding protein 1-like 1 [Source:MGI Symbol;Acc:MGI:1913505] | Hsbp1l1 | Enriched in retina | -2.642 | 1.281E-09 | 35.03 |
| ENSMUSG00000001504 | Iroquois homeobox 2 [Source:MGI Symbol;Acc:MGI:1197526] | Irx2 | Enriched in retina | -2.643 | 2.123E-08 | 24.13 |
| ENSMUSG00000040666 | SH3-binding domain glutamic acid-rich protein [Source:MGI Symbol;Acc:MGI:1354740] | Sh3bgr | Enriched in retina | -2.672 | 0.000008683 | 12.69 |
| ENSMUSG00000025432 | advillin [Source:MGI Symbol;Acc:MGI:1333798] | Avil | Enriched in retina | -2.702 | 0.002164 | 4.817 |
| ENSMUSG00000001120 | poly(rC) binding protein 3 [Source:MGI Symbol;Acc:MGI:1890470] | Pcbp3 | Enriched in retina | -2.721 | 1.56E-44 | 628.3 |
| ENSMUSG00000034981 | prostate androgen-regulated mucin-like protein 1 [Source:MGI Symbol;Acc:MGI:2443349] | Parm1 | Enriched in retina | -2.793 | 5.96E-10 | 144.1 |
| ENSMUSG00000025876 | unc-5 netrin receptor A [Source:MGI Symbol;Acc:MGI:894682] | Unc5a | Enriched in retina | -2.807 | 9.087E-20 | 196.1 |
| ENSMUSG00000068220 | lectin, galactose binding, soluble 1 [Source:MGI Symbol;Acc:MGI:96777] | Lgals1 | Enriched in retina | -2.824 | 9.065E-07 | 30.47 |
| ENSMUSG00000021541 | transient receptor potential cation channel, subfamily C, member 7 [Source:MGI Symbol;Acc:MGI:1349470] | Trpc7 | Enriched in retina | -2.907 | 0.000001129 | 62.32 |
| ENSMUSG00000050822 | solute carrier family 29 (nucleoside transporters), member 4 [Source:MGI Symbol;Acc:MGI:2385330] | Slc29a4 | Enriched in retina | -2.913 | 3.304E-18 | 126.5 |
| ENSMUSG00000026778 | protein kinase C, theta [Source:MGI Symbol;Acc:MGI:97601] | Prkcq | Enriched in retina | -2.928 | 1.806E-14 | 111.9 |
| ENSMUSG00000058966 | TLC domain containing 3B [Source:MGI Symbol;Acc:MGI:1916202] | Tlcd3b | Enriched in retina | -3.053 | 2.492E-39 | 987.5 |
| ENSMUSG00000029276 | glomulin, FKBP associated protein [Source:MGI Symbol;Acc:MGI:2141180] | Glmn | Enriched in retina | -3.081 | 3.563E-13 | 122.5 |
| ENSMUSG00000039954 | serine/threonine kinase 32A [Source:MGI Symbol;Acc:MGI:2442403] | Stk32a | Enriched in retina | -3.376 | 8.556E-15 | 77.08 |
| ENSMUSG00000047986 | paralemmin 3 [Source:MGI Symbol;Acc:MGI:1921587] | Palm3 | Enriched in retina | -3.479 | 1.371E-08 | 25.27 |
| ENSMUSG00000039158 | AT-hook transcription factor [Source:MGI Symbol;Acc:MGI:2140340] | Akna | Enriched in retina | -3.517 | 2.328E-29 | 134.6 |
| ENSMUSG00000002980 | basal cell adhesion molecule [Source:MGI Symbol;Acc:MGI:1929940] | Bcam | Enriched in retina | -3.575 | 2.396E-16 | 83.63 |
| ENSMUSG00000091402 | retinal degeneration 3-like [Source:MGI Symbol;Acc:MGI:2675860] | Rd3l | Enriched in retina | -3.636 | 2.25E-14 | 41.89 |
| ENSMUSG00000020810 | cytoglobin [Source:MGI Symbol;Acc:MGI:2149481] | Cygb | Enriched in retina | -3.636 | 1.973E-30 | 213.7 |
| ENSMUSG00000096696 | zinc finger protein 960 [Source:MGI Symbol;Acc:MGI:3052731] | Zfp960 | Enriched in retina | -3.664 | 0.000112 | 4 |
| ENSMUSG00000001333 | syncoilin [Source:MGI Symbol;Acc:MGI:1916078] | Sync | Enriched in retina | -3.963 | 1.203E-07 | 17.38 |
| ENSMUSG00000046719 | neurexophilin 3 [Source:MGI Symbol;Acc:MGI:1336188] | Nxph3 | Enriched in retina | -4.045 | 3.257E-15 | 27.7 |
| ENSMUSG00000096351 | sterile alpha motif domain containing 11 [Source:MGI Symbol;Acc:MGI:2446220] | Samd11 | Enriched in retina | -4.655 | 2.246E-25 | 401.5 |
| ENSMUSG00000058925 | coiled-coil domain containing 192 [Source:MGI Symbol;Acc:MGI:1922694] | Ccdc192 | Enriched in retina | -4.677 | 0.000005125 | 7.152 |
| ENSMUSG00000079357 | predicted gene 11100 [Source:MGI Symbol;Acc:MGI:3779338] | Gm11100 | Enriched in retina | -4.691 | 0.000004362 | 4.575 |
| ENSMUSG00000031737 | Iroquois homeobox 5 [Source:MGI Symbol;Acc:MGI:1859086] | Irx5 | Enriched in retina | -4.722 | 2.763E-15 | 31.09 |
| ENSMUSG00000031586 | RNA binding protein gene with multiple splicing [Source:MGI Symbol;Acc:MGI:1334446] | Rbpms | Enriched in retina | -4.877 | 6.181E-28 | 81.55 |
| ENSMUSG00000032387 | RNA binding protein with multiple splicing 2 [Source:MGI Symbol;Acc:MGI:1919223] | Rbpms2 | Enriched in retina | -5.34 | 3.186E-24 | 47.72 |
| ENSMUSG00000033196 | myosin, heavy polypeptide 2, skeletal muscle, adult [Source:MGI Symbol;Acc:MGI:1339710] | Myh2 | Enriched in retina | -5.354 | 0.00008309 | 4.066 |
| ENSMUSG00000051279 | growth differentiation factor 6 [Source:MGI Symbol;Acc:MGI:95689] | Gdf6 | Enriched in retina | -5.385 | 2.183E-09 | 11.98 |
| ENSMUSG00000029304 | secreted phosphoprotein 1 [Source:MGI Symbol;Acc:MGI:98389] | Spp1 | Enriched in retina | -5.581 | 5.086E-09 | 33.23 |
| ENSMUSG00000042073 | abhydrolase domain containing 14b [Source:MGI Symbol;Acc:MGI:1923741] | Abhd14b | Enriched in retina | -5.784 | 8.892E-16 | 38.48 |
| ENSMUSG00000023484 | peripherin [Source:MGI Symbol;Acc:MGI:97774] | Prph | Enriched in retina | -5.956 | 4.707E-13 | 64.1 |
| ENSMUSG00000002100 | myosin binding protein C, cardiac [Source:MGI Symbol;Acc:MGI:102844] | Mybpc3 | Enriched in retina | -6.511 | 0.000001051 | 4.298 |
| ENSMUSG00000035296 | sarcoglycan, gamma (dystrophin-associated glycoprotein) [Source:MGI Symbol;Acc:MGI:1346524] | Sgcg | Enriched in retina | -6.983 | 5.532E-09 | 9.668 |
| ENSMUSG00000069372 | cortexin 3 [Source:MGI Symbol;Acc:MGI:3642816] | Ctxn3 | Enriched in retina | -7.203 | 2.521E-13 | 38.01 |
| ENSMUSG00000021799 | opsin 4 (melanopsin) [Source:MGI Symbol;Acc:MGI:1353425] | Opn4 | Enriched in retina | -7.262 | 6.38E-13 | 25.82 |
| ENSMUSG00000031738 | Iroquois homeobox 6 [Source:MGI Symbol;Acc:MGI:1927642] | Irx6 | Enriched in retina | -8.251 | 2.534E-16 | 50.97 |
| ENSMUSG00000031965 | T-box 20 [Source:MGI Symbol;Acc:MGI:1888496] | Tbx20 | Enriched in retina | -8.71 | 2.505E-14 | 45.45 |
| ENSMUSG00000032446 | eomesodermin [Source:MGI Symbol;Acc:MGI:1201683] | Eomes | Enriched in retina | -8.796 | 1.775E-12 | 60.2 |
| ENSMUSG00000031688 | POU domain, class 4, transcription factor 2 [Source:MGI Symbol;Acc:MGI:102524] | Pou4f2 | Enriched in retina | -9.506 | 4.292E-17 | 45.2 |

**Contrast numerator: P15_KO_dLGN. Contrast denominator: P15_KO_retina.**

| ensembl_gene_id | description | mgi_symbol | Significance | deseq_logfc | deseq_adjp | deseq_basemean |
|---|---|---|---|---|---|---|
| ENSMUSG00000030785 | cytochrome c oxidase subunit 6A2 [Source:MGI Symbol;Acc:MGI:104649] | Cox6a2 | Enriched in dLGN | 8.517 | 1.958E-11 | 11.74 |
| ENSMUSG00000078816 | protein kinase C, gamma [Source:MGI Symbol;Acc:MGI:97597] | Prkcg | Enriched in dLGN | 8.457 | 9.316E-88 | 571.8 |
| ENSMUSG00000023828 | solute carrier family 22 (organic cation transporter), member 3 [Source:MGI Symbol;Acc:MGI:1333817] | Slc22a3 | Enriched in dLGN | 7.697 | 5.088E-17 | 27.45 |
| ENSMUSG00000032532 | cholecystokinin [Source:MGI Symbol;Acc:MGI:88297] | Cck | Enriched in dLGN | 7.084 | 6.012E-39 | 445.5 |
| ENSMUSG00000021948 | protein kinase C, delta [Source:MGI Symbol;Acc:MGI:97598] | Prkcd | Enriched in dLGN | 7.058 | 2.616E-44 | 727.9 |
| ENSMUSG00000022468 | endonuclease, polyU-specific [Source:MGI Symbol;Acc:MGI:97746] | Endou | Enriched in dLGN | 6.922 | 4.783E-24 | 34.93 |
| ENSMUSG00000031994 | a disintegrin-like and metallopeptidase (reprolysin type) with thrombospondin type 1 motif, 8 [Source:MGI Symbol;Acc:MGI:1353468] | Adamts8 | Enriched in dLGN | 6.782 | 2.85E-25 | 52.93 |
| ENSMUSG00000039783 | kynurenine 3-monooxygenase (kynurenine 3-hydroxylase) [Source:MGI Symbol;Acc:MGI:2138151] | Kmo | Enriched in dLGN | 6.521 | 5.042E-10 | 4.13 |
| ENSMUSG00000055027 | SET and MYND domain containing 1 [Source:MGI Symbol;Acc:MGI:104790] | Smyd1 | Enriched in dLGN | 6.078 | 4.578E-29 | 65.99 |
| ENSMUSG00000062713 | single-minded family bHLH transcription factor 2 [Source:MGI Symbol;Acc:MGI:98307] | Sim2 | Enriched in dLGN | 5.962 | 0.00004244 | 5.359 |
| ENSMUSG00000026525 | opsin 3 [Source:MGI Symbol;Acc:MGI:1338022] | Opn3 | Enriched in dLGN | 5.916 | 9.675E-25 | 17.62 |
| ENSMUSG00000029211 | gamma-aminobutyric acid (GABA) A receptor, subunit alpha 4 [Source:MGI Symbol;Acc:MGI:95616] | Gabra4 | Enriched in dLGN | 5.823 | 5.414E-56 | 154.6 |
| ENSMUSG00000039103 | nexilin [Source:MGI Symbol;Acc:MGI:1916060] | Nexn | Enriched in dLGN | 5.51 | 4.708E-25 | 125.8 |
| ENSMUSG00000027347 | RAS guanyl releasing protein 1 [Source:MGI Symbol;Acc:MGI:1314635] | Rasgrp1 | Enriched in dLGN | 5.409 | 1.108E-40 | 353.4 |
| ENSMUSG00000032776 | multiple C2 domains, transmembrane 2 [Source:MGI Symbol;Acc:MGI:2685335] | Mctp2 | Enriched in dLGN | 5.072 | 1.636E-19 | 28.35 |
| ENSMUSG00000025081 | tudor domain containing 1 [Source:MGI Symbol;Acc:MGI:1933218] | Tdrd1 | Enriched in dLGN | 4.798 | 0.000009359 | 5.686 |
| ENSMUSG00000026278 | BCL2-related ovarian killer [Source:MGI Symbol;Acc:MGI:1858494] | Bok | Enriched in dLGN | 4.774 | 7.232E-42 | 219.1 |
| ENSMUSG00000030340 | sodium channel, nonvoltage-gated 1 alpha [Source:MGI Symbol;Acc:MGI:101782] | Scnn1a | Enriched in dLGN | 4.73 | 2.33E-26 | 44.62 |
| ENSMUSG00000028176 | leucine rich repeat containing 7 [Source:MGI Symbol;Acc:MGI:2676665] | Lrrc7 | Enriched in dLGN | 4.726 | 5.802E-38 | 677.2 |
| ENSMUSG00000049225 | pyruvate dehydrogenase phosphatase catalytic subunit 1 [Source:MGI Symbol;Acc:MGI:2685870] | Pdp1 | Enriched in dLGN | 4.613 | 4.225E-84 | 541.9 |
| ENSMUSG00000028082 | SH3 domain protein D19 [Source:MGI Symbol;Acc:MGI:1350923] | Sh3d19 | Enriched in dLGN | 4.586 | 2.66E-37 | 657.1 |
| ENSMUSG00000040723 | RCSD domain containing 1 [Source:MGI Symbol;Acc:MGI:2676394] | Rcsd1 | Enriched in dLGN | 4.557 | 1.883E-19 | 27.5 |
| ENSMUSG00000050541 | adrenergic receptor, alpha 1b [Source:MGI Symbol;Acc:MGI:104774] | Adra1b | Enriched in dLGN | 4.556 | 4.563E-31 | 73.77 |
| ENSMUSG00000020733 | solute carrier family 9 (sodium/hydrogen exchanger), member 3 regulator 1 [Source:MGI Symbol;Acc:MGI:1349482] | Slc9a3r1 | Enriched in dLGN | 4.545 | 1.538E-70 | 214.9 |
| ENSMUSG00000000296 | tumor protein D52-like 1 [Source:MGI Symbol;Acc:MGI:1298386] | Tpd52l1 | Enriched in dLGN | 4.543 | 1.479E-34 | 61.04 |
| ENSMUSG00000064179 | troponin T1, skeletal, slow [Source:MGI Symbol;Acc:MGI:1333868] | Tnnt1 | Enriched in dLGN | 4.524 | 5.296E-25 | 308.7 |
| ENSMUSG00000056258 | potassium voltage-gated channel, subfamily Q, member 3 [Source:MGI Symbol;Acc:MGI:1336181] | Kcnq3 | Enriched in dLGN | 4.457 | 5.785E-44 | 1087 |
| ENSMUSG00000022014 | epithelial stromal interaction 1 (breast) [Source:MGI Symbol;Acc:MGI:1915168] | Epsti1 | Enriched in dLGN | 4.367 | 1.484E-08 | 6.466 |
| ENSMUSG00000020732 | RAB37, member RAS oncogene family [Source:MGI Symbol;Acc:MGI:1929945] | Rab37 | Enriched in dLGN | 4.322 | 3.172E-16 | 50.93 |
| ENSMUSG00000029370 | Ras association (RalGDS/AF-6) domain family member 6 [Source:MGI Symbol;Acc:MGI:1920496] | Rassf6 | Enriched in dLGN | 4.284 | 1.417E-07 | 5.633 |
| ENSMUSG00000068696 | G-protein coupled receptor 88 [Source:MGI Symbol;Acc:MGI:1927653] | Gpr88 | Enriched in dLGN | 4.229 | 1.175E-40 | 66.64 |
| ENSMUSG00000033044 | dehydrogenase/reductase (SDR family) member 7C [Source:MGI Symbol;Acc:MGI:1915710] | Dhrs7c | Enriched in dLGN | 4.22 | 0.001365 | 6.079 |
| ENSMUSG00000054196 | collagen triple helix repeat containing 1 [Source:MGI Symbol;Acc:MGI:1915838] | Cthrc1 | Enriched in dLGN | 4.219 | 1.678E-23 | 38 |
| ENSMUSG00000045731 | prepronociceptin [Source:MGI Symbol;Acc:MGI:105308] | Pnoc | Enriched in dLGN | 4.218 | 9.414E-24 | 50.56 |
| ENSMUSG00000020734 | glutamate receptor, ionotropic, NMDA2C (epsilon 3) [Source:MGI Symbol;Acc:MGI:95822] | Grin2c | Enriched in dLGN | 4.193 | 5.611E-24 | 127.2 |
| ENSMUSG00000046480 | sodium channel, type IV, beta [Source:MGI Symbol;Acc:MGI:2687406] | Scn4b | Enriched in dLGN | 4.189 | 1.244E-16 | 167.6 |
| ENSMUSG00000035506 | solute carrier family 12 (potassium/chloride transporters), member 8 [Source:MGI Symbol;Acc:MGI:2443672] | Slc12a8 | Enriched in dLGN | 4.182 | 1.504E-10 | 17.24 |
| ENSMUSG00000066129 | kinase non-catalytic C-lobe domain (KIND) containing 1 [Source:MGI Symbol;Acc:MGI:1923734] | Kndc1 | Enriched in dLGN | 4.141 | 1.906E-59 | 714.5 |
| ENSMUSG00000029054 | gamma-aminobutyric acid (GABA) A receptor, subunit delta [Source:MGI Symbol;Acc:MGI:95622] | Gabrd | Enriched in dLGN | 4.07 | 1.68E-18 | 52.02 |
| ENSMUSG00000036972 | zinc finger protein of the cerebellum 4 [Source:MGI Symbol;Acc:MGI:107201] | Zic4 | Enriched in dLGN | 4.029 | 5.035E-24 | 142.7 |
| ENSMUSG00000004892 | brevican [Source:MGI Symbol;Acc:MGI:1096385] | Bcan | Enriched in dLGN | 4.017 | 1.416E-40 | 546.7 |
| ENSMUSG00000022900 | immunoglobulin-like domain containing receptor 1 [Source:MGI Symbol;Acc:MGI:2146574] | Ildr1 | Enriched in dLGN | 3.986 | 0.00004004 | 4.707 |
| ENSMUSG00000032503 | cyclic AMP-regulated phosphoprotein, 21 [Source:MGI Symbol;Acc:MGI:107562] | Arpp21 | Enriched in dLGN | 3.968 | 1.419E-22 | 594.6 |
| ENSMUSG00000048616 | noggin [Source:MGI Symbol;Acc:MGI:104327] | Nog | Enriched in dLGN | 3.92 | 2.987E-21 | 53.91 |
| ENSMUSG00000026688 | microsomal glutathione S-transferase 3 [Source:MGI Symbol;Acc:MGI:1913697] | Mgst3 | Enriched in dLGN | 3.92 | 2.543E-14 | 88.46 |
| ENSMUSG00000027360 | histidine decarboxylase [Source:MGI Symbol;Acc:MGI:96062] | Hdc | Enriched in dLGN | 3.896 | 1.334E-14 | 44.93 |
| ENSMUSG00000038128 | calcium/calmodulin-dependent protein kinase IV [Source:MGI Symbol;Acc:MGI:88258] | Camk4 | Enriched in dLGN | 3.878 | 5.075E-52 | 937 |
| ENSMUSG00000027489 | N-terminal EF-hand calcium binding protein 3 [Source:MGI Symbol;Acc:MGI:1861721] | Necab3 | Enriched in dLGN | 3.853 | 1.186E-26 | 36.72 |
| ENSMUSG00000040624 | pleckstrin homology domain containing, family G (with RhoGef domain) member 1 [Source:MGI Symbol;Acc:MGI:2676551] | Plekhg1 | Enriched in dLGN | 3.843 | 9.784E-43 | 465 |
| ENSMUSG00000034825 | nuclear receptor interacting protein 3 [Source:MGI Symbol;Acc:MGI:1925843] | Nrip3 | Enriched in dLGN | 3.793 | 2.947E-26 | 285.5 |
| ENSMUSG00000068373 | RIKEN cDNA D430041D05 gene [Source:MGI Symbol;Acc:MGI:2181743] | D430041D05Rik | Enriched in dLGN | 3.765 | 2.739E-80 | 780.2 |
| ENSMUSG00000019966 | kit ligand [Source:MGI Symbol;Acc:MGI:96974] | Kitl | Enriched in dLGN | 3.75 | 2.832E-24 | 453.1 |
| ENSMUSG00000024727 | transient receptor potential cation channel, subfamily M, member 6 [Source:MGI Symbol;Acc:MGI:2675603] | Trpm6 | Enriched in dLGN | 3.69 | 1.047E-07 | 19.86 |
| ENSMUSG00000021209 | protein phosphatase 4, regulatory subunit 4 [Source:MGI Symbol;Acc:MGI:1921771] | Ppp4r4 | Enriched in dLGN | 3.673 | 4.17E-39 | 139.4 |
| ENSMUSG00000040710 | ST8 alpha-N-acetyl-neuraminide alpha-2,8-sialyltransferase 4 [Source:MGI Symbol;Acc:MGI:106018] | St8sia4 | Enriched in dLGN | 3.665 | 7.896E-27 | 69.4 |
| ENSMUSG00000060882 | potassium voltage-gated channel, Shal-related family, member 2 [Source:MGI Symbol;Acc:MGI:102663] | Kcnd2 | Enriched in dLGN | 3.657 | 2.33E-17 | 981.4 |
| ENSMUSG00000049176 | FERM and PDZ domain containing 4 [Source:MGI Symbol;Acc:MGI:3042378] | Frmpd4 | Enriched in dLGN | 3.654 | 6.859E-24 | 381.5 |
| ENSMUSG00000042353 | Fras1 related extracellular matrix protein 3 [Source:MGI Symbol;Acc:MGI:2685641] | Frem3 | Enriched in dLGN | 3.638 | 4.167E-19 | 57.69 |
| ENSMUSG00000021259 | cytochrome P450, family 46, subfamily a, polypeptide 1 [Source:MGI Symbol;Acc:MGI:1341877] | Cyp46a1 | Enriched in dLGN | 3.6 | 3.425E-55 | 248.2 |
| ENSMUSG00000067860 | zinc finger protein of the cerebellum 3 [Source:MGI Symbol;Acc:MGI:106676] | Zic3 | Enriched in dLGN | 3.587 | 6.265E-18 | 26.92 |
| ENSMUSG00000018821 | arginine vasopressin-induced 1 [Source:MGI Symbol;Acc:MGI:1916784] | Avpi1 | Enriched in dLGN | 3.526 | 1.833E-22 | 34.89 |
| ENSMUSG00000030303 | fatty acyl CoA reductase 2 [Source:MGI Symbol;Acc:MGI:2687035] | Far2 | Enriched in dLGN | 3.469 | 9.711E-14 | 96.51 |
| ENSMUSG00000046834 | keratin 1 [Source:MGI Symbol;Acc:MGI:96698] | Krt1 | Enriched in dLGN | 3.461 | 2.142E-13 | 16.62 |
| ENSMUSG00000022454 | NEL-like 2 [Source:MGI Symbol;Acc:MGI:1858510] | Nell2 | Enriched in dLGN | 3.452 | 1.005E-36 | 1350 |
| ENSMUSG00000033685 | uncoupling protein 2 (mitochondrial, proton carrier) [Source:MGI Symbol;Acc:MGI:109354] | Ucp2 | Enriched in dLGN | 3.448 | 6.653E-20 | 149.6 |
| ENSMUSG00000040612 | immunoglobulin-like domain containing receptor 2 [Source:MGI Symbol;Acc:MGI:1196370] | Ildr2 | Enriched in dLGN | 3.438 | 4.762E-28 | 1307 |
| ENSMUSG00000073406 | histocompatibility 2, blastocyst [Source:MGI Symbol;Acc:MGI:892004] | H2-Bl | Enriched in dLGN | 3.431 | 0.00003652 | 18.45 |
| ENSMUSG00000096257 | coiled-coil glutamate-rich protein 2 [Source:MGI Symbol;Acc:MGI:3645242] | Ccer2 | Enriched in dLGN | 3.424 | 4.744E-10 | 12.8 |
| ENSMUSG00000037996 | solute carrier family 24 (sodium/potassium/calcium exchanger), member 2 [Source:MGI Symbol;Acc:MGI:1923626] | Slc24a2 | Enriched in dLGN | 3.401 | 1.726E-19 | 1329 |
| ENSMUSG00000033615 | complexin 1 [Source:MGI Symbol;Acc:MGI:104727] | Cplx1 | Enriched in dLGN | 3.337 | 1.705E-19 | 2304 |
| ENSMUSG00000030889 | von Willebrand factor A domain containing 3A [Source:MGI Symbol;Acc:MGI:3041229] | Vwa3a | Enriched in dLGN | 3.305 | 0.00004345 | 26.65 |
| ENSMUSG00000027965 | olfactomedin 3 [Source:MGI Symbol;Acc:MGI:2387329] | Olfm3 | Enriched in dLGN | 3.297 | 8.769E-19 | 175.9 |
| ENSMUSG00000026235 | Eph receptor A4 [Source:MGI Symbol;Acc:MGI:98277] | Epha4 | Enriched in dLGN | 3.283 | 6.609E-31 | 500.9 |
| ENSMUSG00000026888 | growth factor receptor bound protein 14 [Source:MGI Symbol;Acc:MGI:1355324] | Grb14 | Enriched in dLGN | 3.277 | 1.952E-18 | 102.4 |
| ENSMUSG00000051022 | heparan sulfate (glucosamine) 3-O-sulfotransferase 1 [Source:MGI Symbol;Acc:MGI:1201606] | Hs3st1 | Enriched in dLGN | 3.269 | 2.483E-31 | 132 |
| ENSMUSG00000021719 | regulator of G-protein signalling 7 binding protein [Source:MGI Symbol;Acc:MGI:106334] | Rgs7bp | Enriched in dLGN | 3.23 | 4.039E-29 | 598.8 |
| ENSMUSG00000031760 | metallothionein 3 [Source:MGI Symbol;Acc:MGI:97173] | Mt3 | Enriched in dLGN | 3.222 | 3.753E-19 | 131.4 |
| ENSMUSG00000030075 | contactin 3 [Source:MGI Symbol;Acc:MGI:99534] | Cntn3 | Enriched in dLGN | 3.212 | 5.861E-17 | 116.4 |
| ENSMUSG00000026301 | IQ motif containing with AAA domain [Source:MGI Symbol;Acc:MGI:1922168] | Iqca | Enriched in dLGN | 3.212 | 0.002434 | 31.31 |
| ENSMUSG00000020589 | CYFIP related Rac1 interactor A [Source:MGI Symbol;Acc:MGI:1261783] | Cyria | Enriched in dLGN | 3.183 | 1.333E-35 | 283.7 |
| ENSMUSG00000003665 | hyaluronan synthase 1 [Source:MGI Symbol;Acc:MGI:106590] | Has1 | Enriched in dLGN | 3.174 | 0.005065 | 2.393 |
| ENSMUSG00000024524 | guanine nucleotide binding protein, alpha stimulating, olfactory type [Source:MGI Symbol;Acc:MGI:95774] | Gnal | Enriched in dLGN | 3.147 | 3.988E-47 | 618.2 |
| ENSMUSG00000055409 | NEL-like 1 [Source:MGI Symbol;Acc:MGI:2443902] | Nell1 | Enriched in dLGN | 3.131 | 1.842E-13 | 543.3 |
| ENSMUSG00000040164 | K+ voltage-gated channel, subfamily S, 1 [Source:MGI Symbol;Acc:MGI:1197019] | Kcns1 | Enriched in dLGN | 3.128 | 3.706E-09 | 9.63 |
| ENSMUSG00000060402 | carbohydrate sulfotransferase 8 [Source:MGI Symbol;Acc:MGI:1916197] | Chst8 | Enriched in dLGN | 3.121 | 3.85E-18 | 67.64 |
| ENSMUSG00000040794 | C1q and tumor necrosis factor related protein 4 [Source:MGI Symbol;Acc:MGI:1914695] | C1qtnf4 | Enriched in dLGN | 3.11 | 2.148E-32 | 207.8 |
| ENSMUSG00000065968 | interferon induced transmembrane protein 7 [Source:MGI Symbol;Acc:MGI:1921732] | Ifitm7 | Enriched in dLGN | 3.103 | 1.509E-08 | 15.26 |
| ENSMUSG00000025475 | adhesion G protein-coupled receptor A1 [Source:MGI Symbol;Acc:MGI:1277167] | Adgra1 | Enriched in dLGN | 3.093 | 5.226E-40 | 360.1 |
| ENSMUSG00000001763 | tetraspanin 33 [Source:MGI Symbol;Acc:MGI:1919012] | Tspan33 | Enriched in dLGN | 3.093 | 8.703E-26 | 62.53 |
| ENSMUSG00000029657 | heat shock 105kDa/110kDa protein 1 [Source:MGI Symbol;Acc:MGI:105053] | Hsph1 | Enriched in dLGN | 3.091 | 1.133E-25 | 555.2 |
| ENSMUSG00000074736 | synapse differentiation inducing 1 [Source:MGI Symbol;Acc:MGI:3702158] | Syndig1 | Enriched in dLGN | 3.06 | 1.032E-13 | 50.55 |
| ENSMUSG00000024529 | lysyl oxidase [Source:MGI Symbol;Acc:MGI:96817] | Lox | Enriched in dLGN | 3.052 | 0.000000556 | 9.017 |
| ENSMUSG00000005357 | solute carrier family 1 (high affinity aspartate/glutamate transporter), member 6 [Source:MGI Symbol;Acc:MGI:1096331] | Slc1a6 | Enriched in dLGN | 3.042 | 2.589E-16 | 52.8 |
| ENSMUSG00000007594 | hyaluronan and proteoglycan link protein 4 [Source:MGI Symbol;Acc:MGI:2679531] | Hapln4 | Enriched in dLGN | 3.018 | 4.321E-09 | 156.8 |
| ENSMUSG00000027200 | sema domain, transmembrane domain (TM), and cytoplasmic domain, (semaphorin) 6D [Source:MGI Symbol;Acc:MGI:2387661] | Sema6d | Enriched in dLGN | 3.004 | 3.51E-22 | 731.9 |
| ENSMUSG00000029153 | OCIA domain containing 2 [Source:MGI Symbol;Acc:MGI:1916377] | Ociad2 | Enriched in dLGN | 3.004 | 3.543E-21 | 70.58 |
| ENSMUSG00000020990 | cyclin-dependent kinase-like 1 (CDC2-related kinase) [Source:MGI Symbol;Acc:MGI:1918341] | Cdkl1 | Enriched in dLGN | 2.989 | 1.014E-16 | 53.67 |
| ENSMUSG00000019828 | glutamate receptor, metabotropic 1 [Source:MGI Symbol;Acc:MGI:1351338] | Grm1 | Enriched in dLGN | 2.979 | 5.126E-18 | 572.7 |
| ENSMUSG00000036578 | FXYD domain-containing ion transport regulator 7 [Source:MGI Symbol;Acc:MGI:1889006] | Fxyd7 | Enriched in dLGN | 2.964 | 9.66E-15 | 140.4 |
| ENSMUSG00000040713 | cellular repressor of E1A-stimulated genes 1 [Source:MGI Symbol;Acc:MGI:1344382] | Creg1 | Enriched in dLGN | 2.962 | 9.938E-21 | 76.45 |
| ENSMUSG00000023905 | tumor necrosis factor receptor superfamily, member 12a [Source:MGI Symbol;Acc:MGI:1351484] | Tnfrsf12a | Enriched in dLGN | 2.956 | 8.784E-08 | 7.873 |
| ENSMUSG00000036745 | tubulin tyrosine ligase-like family, member 7 [Source:MGI Symbol;Acc:MGI:1918142] | Ttll7 | Enriched in dLGN | 2.954 | 7.552E-22 | 648.1 |
| ENSMUSG00000058927 | predicted gene 10053 [Source:MGI Symbol;Acc:MGI:3704493] | Gm10053 | Enriched in dLGN | 2.953 | 3.161E-08 | 16.38 |
| ENSMUSG00000031438 | ring finger protein 128 [Source:MGI Symbol;Acc:MGI:1914139] | Rnf128 | Enriched in dLGN | 2.949 | 2.747E-08 | 18.97 |
| ENSMUSG00000061859 | PATJ, crumbs cell polarity complex component [Source:MGI Symbol;Acc:MGI:1277960] | Patj | Enriched in dLGN | 2.946 | 1.159E-13 | 609.6 |
| ENSMUSG00000034570 | inositol polyphosphate 5-phosphatase J [Source:MGI Symbol;Acc:MGI:2158663] | Inpp5j | Enriched in dLGN | 2.939 | 2.932E-18 | 112.1 |
| ENSMUSG00000038244 | microtubule associated monooxygenase, calponin and LIM domain containing 2 [Source:MGI Symbol;Acc:MGI:2444947] | Mical2 | Enriched in dLGN | 2.919 | 8.118E-20 | 255.1 |
| ENSMUSG00000049612 | oligodendrocyte myelin glycoprotein [Source:MGI Symbol;Acc:MGI:106586] | Omg | Enriched in dLGN | 2.918 | 2.501E-15 | 36.01 |
| ENSMUSG00000019762 | iodotyrosine deiodinase [Source:MGI Symbol;Acc:MGI:1917587] | Iyd | Enriched in dLGN | 2.917 | 0.0002831 | 6.399 |
| ENSMUSG00000056413 | ArfGAP with dual PH domains 1 [Source:MGI Symbol;Acc:MGI:2442201] | Adap1 | Enriched in dLGN | 2.914 | 5.076E-55 | 552.2 |
| ENSMUSG00000028718 | Scl/Tal1 interrupting locus [Source:MGI Symbol;Acc:MGI:107477] | Stil | Enriched in dLGN | 2.912 | 0.00006938 | 5.074 |
| ENSMUSG00000027577 | cholinergic receptor, nicotinic, alpha polypeptide 4 [Source:MGI Symbol;Acc:MGI:87888] | Chrna4 | Enriched in dLGN | 2.904 | 5.469E-22 | 546.3 |
| ENSMUSG00000026384 | protein tyrosine phosphatase, non-receptor type 4 [Source:MGI Symbol;Acc:MGI:1099792] | Ptpn4 | Enriched in dLGN | 2.895 | 1.964E-23 | 1013 |
| ENSMUSG00000020059 | synaptonemal complex protein 3 [Source:MGI Symbol;Acc:MGI:109542] | Sycp3 | Enriched in dLGN | 2.892 | 0.004046 | 2.11 |
| ENSMUSG00000041020 | MAP7 domain containing 2 [Source:MGI Symbol;Acc:MGI:1917474] | Map7d2 | Enriched in dLGN | 2.889 | 5.657E-28 | 736.5 |
| ENSMUSG00000062184 | heparan sulfate 6-O-sulfotransferase 2 [Source:MGI Symbol;Acc:MGI:1354959] | Hs6st2 | Enriched in dLGN | 2.888 | 1.733E-18 | 235.4 |
| ENSMUSG00000028464 | tropomyosin 2, beta [Source:MGI Symbol;Acc:MGI:98810] | Tpm2 | Enriched in dLGN | 2.876 | 8.18E-09 | 25.35 |
| ENSMUSG00000033061 | regulated endocrine-specific protein 18 [Source:MGI Symbol;Acc:MGI:1098222] | Resp18 | Enriched in dLGN | 2.851 | 5.654E-07 | 74.08 |
| ENSMUSG00000021591 | glutaredoxin [Source:MGI Symbol;Acc:MGI:2135625] | Glrx | Enriched in dLGN | 2.849 | 4.564E-23 | 37.82 |
| ENSMUSG00000044835 | ankyrin repeat domain 45 [Source:MGI Symbol;Acc:MGI:1921094] | Ankrd45 | Enriched in dLGN | 2.846 | 8.184E-21 | 83.71 |
| ENSMUSG00000034353 | receptor (calcitonin) activity modifying protein 1 [Source:MGI Symbol;Acc:MGI:1858418] | Ramp1 | Enriched in dLGN | 2.846 | 1.459E-11 | 54.71 |
| ENSMUSG00000045733 | shadow of prion protein [Source:MGI Symbol;Acc:MGI:3582583] | Sprn | Enriched in dLGN | 2.837 | 2.121E-37 | 239 |
| ENSMUSG00000071866 | peptidylprolyl isomerase A [Source:MGI Symbol;Acc:MGI:97749] | Ppia | Enriched in dLGN | 2.814 | 5.067E-07 | 211.2 |
| ENSMUSG00000062937 | methylthioadenosine phosphorylase [Source:MGI Symbol;Acc:MGI:1914152] | Mtap | Enriched in dLGN | 2.813 | 2.917E-11 | 29.38 |
| ENSMUSG00000035849 | keratin 222 [Source:MGI Symbol;Acc:MGI:2442728] | Krt222 | Enriched in dLGN | 2.807 | 1.921E-18 | 127 |
| ENSMUSG00000037492 | zinc finger, matrin type 4 [Source:MGI Symbol;Acc:MGI:2443497] | Zmat4 | Enriched in dLGN | 2.807 | 9.317E-16 | 529.7 |

| | | | | | | |
|---|---|---|---|---|---|---|
| ENSMUSG00000030500 | solute carrier family 17 (sodium-dependent inorganic phosphate cotransporter), member 6 [Source:MGI Symbol;Acc:MGI:2156052] | Slc17a6 | Enriched in dLGN | 2.806 | 3.253E-10 | 605.1 |
| ENSMUSG00000026824 | potassium inwardly-rectifying channel, subfamily J, member 3 [Source:MGI Symbol;Acc:MGI:104742] | Kcnj3 | Enriched in dLGN | 2.801 | 1.573E-17 | 206.8 |
| ENSMUSG00000110711 | predicted gene 45760 [Source:MGI Symbol;Acc:MGI:5804875] | Gm45760 | Enriched in dLGN | 2.798 | 0.001839 | 2.941 |
| ENSMUSG00000115293 | Eef1akmt4-endothelin converting enzyme 2 readthrough [Source:MGI Symbol;Acc:MGI:6121518] | Eef1ece2 | Enriched in dLGN | 2.795 | 1.852E-09 | 44.17 |
| ENSMUSG00000027076 | translocase of inner mitochondrial membrane 10 [Source:MGI Symbol;Acc:MGI:1353429] | Timm10 | Enriched in dLGN | 2.79 | 2.798E-13 | 29.72 |
| ENSMUSG00000024109 | neurexin I [Source:MGI Symbol;Acc:MGI:1096391] | Nrxn1 | Enriched in dLGN | 2.786 | 1.85E-18 | 3720 |
| ENSMUSG00000053930 | shisa family member 6 [Source:MGI Symbol;Acc:MGI:2685725] | Shisa6 | Enriched in dLGN | 2.764 | 6.426E-16 | 186.6 |
| ENSMUSG00000028773 | fatty acid binding protein 3, muscle and heart [Source:MGI Symbol;Acc:MGI:95476] | Fabp3 | Enriched in dLGN | 2.762 | 3.012E-07 | 75.72 |
| ENSMUSG00000033316 | polypeptide N-acetylgalactosaminyltransferase 9 [Source:MGI Symbol;Acc:MGI:2677965] | Galnt9 | Enriched in dLGN | 2.761 | 1.406E-27 | 204.6 |
| ENSMUSG00000061451 | transmembrane protein 151A [Source:MGI Symbol;Acc:MGI:2147713] | Tmem151a | Enriched in dLGN | 2.758 | 3.209E-29 | 350.6 |
| ENSMUSG00000044017 | adhesion G protein-coupled receptor D1 [Source:MGI Symbol;Acc:MGI:3041203] | Adgrd1 | Enriched in dLGN | 2.736 | 0.000355 | 5.834 |
| ENSMUSG00000019194 | sodium channel, voltage-gated, type I, beta [Source:MGI Symbol;Acc:MGI:98247] | Scn1b | Enriched in dLGN | 2.734 | 2.437E-18 | 470 |
| ENSMUSG00000028364 | tenascin C [Source:MGI Symbol;Acc:MGI:101922] | Tnc | Enriched in dLGN | 2.729 | 1.052E-09 | 268.5 |
| ENSMUSG00000041263 | RUN and SH3 domain containing 1 [Source:MGI Symbol;Acc:MGI:1919546] | Rusc1 | Enriched in dLGN | 2.726 | 6.188E-21 | 448.9 |
| ENSMUSG00000056222 | sparc/osteonectin, cwcv and kazal-like domains proteoglycan 1 [Source:MGI Symbol;Acc:MGI:105371] | Spock1 | Enriched in dLGN | 2.724 | 3.712E-16 | 1204 |
| ENSMUSG00000025468 | calcyon neuron-specific vesicular protein [Source:MGI Symbol;Acc:MGI:1915816] | Caly | Enriched in dLGN | 2.721 | 1.337E-16 | 238.8 |
| ENSMUSG00000015656 | heat shock protein 8 [Source:MGI Symbol;Acc:MGI:105384] | Hspa8 | Enriched in dLGN | 2.721 | 6.739E-09 | 1562 |
| ENSMUSG00000028367 | thioredoxin 1 [Source:MGI Symbol;Acc:MGI:98874] | Txn1 | Enriched in dLGN | 2.718 | 1.816E-17 | 92.9 |
| ENSMUSG00000046182 | GSG1-like [Source:MGI Symbol;Acc:MGI:2685483] | Gsg1l | Enriched in dLGN | 2.71 | 1.092E-27 | 471.7 |
| ENSMUSG00000004031 | bone morphogenic protein/retinoic acid inducible neural-specific 2 [Source:MGI Symbol;Acc:MGI:2443333] | Brinp2 | Enriched in dLGN | 2.685 | 3.793E-22 | 323.9 |
| ENSMUSG00000025428 | ATP synthase, H+ transporting, mitochondrial F1 complex, alpha subunit 1 [Source:MGI Symbol;Acc:MGI:88115] | Atp5a1 | Enriched in dLGN | 2.678 | 4.736E-17 | 1689 |
| ENSMUSG00000050315 | synaptopodin 2 [Source:MGI Symbol;Acc:MGI:2153070] | Synpo2 | Enriched in dLGN | 2.675 | 2.325E-16 | 581.4 |
| ENSMUSG00000043811 | reticulon 4 receptor [Source:MGI Symbol;Acc:MGI:2136886] | Rtn4r | Enriched in dLGN | 2.67 | 7.957E-14 | 85.48 |
| ENSMUSG00000007653 | gamma-aminobutyric acid (GABA) A receptor, subunit beta 2 [Source:MGI Symbol;Acc:MGI:95620] | Gabrb2 | Enriched in dLGN | 2.67 | 1.596E-12 | 322.6 |
| ENSMUSG00000022494 | shisa family member 9 [Source:MGI Symbol;Acc:MGI:1919805] | Shisa9 | Enriched in dLGN | 2.664 | 2.028E-12 | 606.7 |
| ENSMUSG00000075702 | selenoprotein M [Source:MGI Symbol;Acc:MGI:2149786] | Selenom | Enriched in dLGN | 2.654 | 7.453E-22 | 136.2 |
| ENSMUSG00000041112 | engulfment and cell motility 1 [Source:MGI Symbol;Acc:MGI:2153044] | Elmo1 | Enriched in dLGN | 2.644 | 2.473E-15 | 598.8 |
| ENSMUSG00000027894 | solute carrier family 6 (neurotransmitter transporter), member 17 [Source:MGI Symbol;Acc:MGI:2442535] | Slc6a17 | Enriched in dLGN | 2.636 | 2.956E-25 | 1006 |
| ENSMUSG00000021373 | CAP, adenylate cyclase-associated protein, 2 (yeast) [Source:MGI Symbol;Acc:MGI:1914502] | Cap2 | Enriched in dLGN | 2.623 | 1.924E-20 | 147 |
| ENSMUSG00000033595 | leucine-rich repeat LGI family, member 3 [Source:MGI Symbol;Acc:MGI:2182619] | Lgi3 | Enriched in dLGN | 2.613 | 1.478E-12 | 185.5 |
| ENSMUSG00000028149 | RAP1, GTP-GDP dissociation stimulator 1 [Source:MGI Symbol;Acc:MGI:2385189] | Rap1gds1 | Enriched in dLGN | 2.61 | 5.378E-34 | 847.3 |
| ENSMUSG00000020321 | malate dehydrogenase 1, NAD (soluble) [Source:MGI Symbol;Acc:MGI:97051] | Mdh1 | Enriched in dLGN | 2.599 | 3.932E-09 | 617.5 |
| ENSMUSG00000022587 | lymphocyte antigen 6 complex, locus E [Source:MGI Symbol;Acc:MGI:106651] | Ly6e | Enriched in dLGN | 2.584 | 1.13E-15 | 95.38 |
| ENSMUSG00000067242 | leucine-rich repeat LGI family, member 1 [Source:MGI Symbol;Acc:MGI:1861691] | Lgi1 | Enriched in dLGN | 2.582 | 4.132E-15 | 72.29 |
| ENSMUSG00000028351 | bone morphogenic protein/retinoic acid inducible neural specific 1 [Source:MGI Symbol;Acc:MGI:1928478] | Brinp1 | Enriched in dLGN | 2.58 | 1.694E-14 | 586.1 |
| ENSMUSG00000050967 | cellular repressor of E1A-stimulated genes 2 [Source:MGI Symbol;Acc:MGI:1928333] | Creg2 | Enriched in dLGN | 2.577 | 1.452E-18 | 188.8 |
| ENSMUSG00000037568 | vasohibin 2 [Source:MGI Symbol;Acc:MGI:2444826] | Vash2 | Enriched in dLGN | 2.576 | 6.125E-19 | 93.06 |
| ENSMUSG00000006057 | ATP synthase, H+ transporting, mitochondrial F0 complex, subunit C1 (subunit 9) [Source:MGI Symbol;Acc:MGI:107653] | Atp5g1 | Enriched in dLGN | 2.576 | 4.989E-14 | 76.53 |
| ENSMUSG00000059857 | netrin G1 [Source:MGI Symbol;Acc:MGI:1934028] | Ntng1 | Enriched in dLGN | 2.568 | 3.026E-09 | 904 |
| ENSMUSG00000056201 | cofilin 1, non-muscle [Source:MGI Symbol;Acc:MGI:101757] | Cfl1 | Enriched in dLGN | 2.563 | 1.064E-15 | 762.6 |
| ENSMUSG00000028410 | DnaJ heat shock protein family (Hsp40) member A1 [Source:MGI Symbol;Acc:MGI:1270129] | Dnaja1 | Enriched in dLGN | 2.559 | 1.176E-09 | 251.8 |
| ENSMUSG00000027712 | annexin A5 [Source:MGI Symbol;Acc:MGI:106008] | Anxa5 | Enriched in dLGN | 2.551 | 8.029E-09 | 102.3 |
| ENSMUSG00000027792 | butyrylcholinesterase [Source:MGI Symbol;Acc:MGI:894278] | Bche | Enriched in dLGN | 2.545 | 1.103E-08 | 17.98 |
| ENSMUSG00000024907 | galanin and GMAP prepropeptide [Source:MGI Symbol;Acc:MGI:95637] | Gal | Enriched in dLGN | 2.544 | 0.000001732 | 30 |
| ENSMUSG00000009894 | synaptosomal-associated protein, 47 [Source:MGI Symbol;Acc:MGI:1915076] | Snap47 | Enriched in dLGN | 2.543 | 2.476E-15 | 533.6 |
| ENSMUSG00000032966 | FK506 binding protein 1a [Source:MGI Symbol;Acc:MGI:95541] | Fkbp1a | Enriched in dLGN | 2.542 | 4.81E-26 | 430.5 |
| ENSMUSG00000028434 | erythrocyte membrane protein band 4.1 like 4b [Source:MGI Symbol;Acc:MGI:1859149] | Epb41l4b | Enriched in dLGN | 2.541 | 6.859E-24 | 333.4 |
| ENSMUSG00000031970 | dysbindin (dystrobrevin binding protein 1) domain containing 1 [Source:MGI Symbol;Acc:MGI:1919435] | Dbndd1 | Enriched in dLGN | 2.531 | 3.865E-22 | 82.55 |
| ENSMUSG00000043629 | RIKEN cDNA 1700019D03 gene [Source:MGI Symbol;Acc:MGI:1914330] | 1700019D03Rik | Enriched in dLGN | 2.529 | 2.94E-11 | 31.11 |
| ENSMUSG00000034226 | ras homolog family member V [Source:MGI Symbol;Acc:MGI:2444227] | Rhov | Enriched in dLGN | 2.523 | 1.946E-10 | 54.65 |
| ENSMUSG00000018965 | tyrosine 3-monooxygenase/tryptophan 5-monooxygenase activation protein, eta polypeptide [Source:MGI Symbol;Acc:MGI:109194] | Ywhah | Enriched in dLGN | 2.515 | 4.224E-13 | 3239 |
| ENSMUSG00000039115 | integrin alpha 9 [Source:MGI Symbol;Acc:MGI:104756] | Itga9 | Enriched in dLGN | 2.511 | 0.00641 | 1811 |
| ENSMUSG00000006711 | RIKEN cDNA D130043K22 gene [Source:MGI Symbol;Acc:MGI:3036268] | D130043K22Rik | Enriched in dLGN | 2.508 | 1.095E-12 | 133.1 |
| ENSMUSG00000035000 | dipeptidylpeptidase 4 [Source:MGI Symbol;Acc:MGI:94919] | Dpp4 | Enriched in dLGN | 2.488 | 0.00001678 | 9.243 |
| ENSMUSG00000039809 | gamma-aminobutyric acid (GABA) B receptor, 2 [Source:MGI Symbol;Acc:MGI:2386030] | Gabbr2 | Enriched in dLGN | 2.487 | 1.8E-30 | 1664 |
| ENSMUSG00000028710 | ATP synthase mitochondrial F1 complex assembly factor 1 [Source:MGI Symbol;Acc:MGI:2180560] | Atpaf1 | Enriched in dLGN | 2.471 | 3.078E-24 | 89.62 |
| ENSMUSG00000030718 | protein phosphatase methylesterase 1 [Source:MGI Symbol;Acc:MGI:1919840] | Ppme1 | Enriched in dLGN | 2.467 | 8.437E-18 | 576.9 |
| ENSMUSG00000050830 | von Willebrand factor C domain containing 2 [Source:MGI Symbol;Acc:MGI:2442987] | Vwc2 | Enriched in dLGN | 2.464 | 5.044E-13 | 127 |
| ENSMUSG00000029632 | Ndufa4, mitochondrial complex associated [Source:MGI Symbol;Acc:MGI:107686] | Ndufa4 | Enriched in dLGN | 2.462 | 7.865E-13 | 183.2 |
| ENSMUSG00000020331 | hyperpolarization-activated, cyclic nucleotide-gated K+ 2 [Source:MGI Symbol;Acc:MGI:1298210] | Hcn2 | Enriched in dLGN | 2.457 | 1.445E-28 | 598.2 |
| ENSMUSG00000063434 | sortilin-related VPS10 domain containing receptor 3 [Source:MGI Symbol;Acc:MGI:1913923] | Sorcs3 | Enriched in dLGN | 2.452 | 1.366E-10 | 332.9 |
| ENSMUSG00000041216 | clavesin 1 [Source:MGI Symbol;Acc:MGI:1921688] | Clvs1 | Enriched in dLGN | 2.447 | 3.222E-13 | 163.7 |
| ENSMUSG00000022269 | membrane associated ring-CH-type finger 11 [Source:MGI Symbol;Acc:MGI:3608327] | Marchf11 | Enriched in dLGN | 2.444 | 0.000001709 | 40.63 |
| ENSMUSG00000024953 | peroxiredoxin 5 [Source:MGI Symbol;Acc:MGI:1859821] | Prdx5 | Enriched in dLGN | 2.439 | 3.358E-22 | 344.2 |
| ENSMUSG00000002797 | gamma-glutamyl cyclotransferase [Source:MGI Symbol;Acc:MGI:95700] | Ggct | Enriched in dLGN | 2.439 | 6.685E-14 | 25.71 |
| ENSMUSG00000053192 | myeloid/lymphoid or mixed-lineage leukemia; translocated to, 11 [Source:MGI Symbol;Acc:MGI:1929671] | Mllt11 | Enriched in dLGN | 2.427 | 2.454E-22 | 499.1 |
| ENSMUSG00000031137 | fibroblast growth factor 13 [Source:MGI Symbol;Acc:MGI:109178] | Fgf13 | Enriched in dLGN | 2.427 | 1.51E-14 | 419.6 |
| ENSMUSG00000029223 | ubiquitin carboxy-terminal hydrolase L1 [Source:MGI Symbol;Acc:MGI:103149] | Uchl1 | Enriched in dLGN | 2.427 | 8.425E-12 | 1008 |
| ENSMUSG00000027698 | neutral cholesterol ester hydrolase 1 [Source:MGI Symbol;Acc:MGI:2443191] | Nceh1 | Enriched in dLGN | 2.425 | 8.893E-15 | 190.1 |
| ENSMUSG00000032564 | copine IV [Source:MGI Symbol;Acc:MGI:1921270] | Cpne4 | Enriched in dLGN | 2.421 | 3.259E-09 | 126.3 |
| ENSMUSG00000071862 | leucine rich repeat transmembrane neuronal 2 [Source:MGI Symbol;Acc:MGI:2389174] | Lrrtm2 | Enriched in dLGN | 2.419 | 2.026E-16 | 78.72 |
| ENSMUSG00000079941 | cytochrome c oxidase subunit 5B, pseudogene [Source:MGI Symbol;Acc:MGI:3649411] | Cox5b-ps | Enriched in dLGN | 2.413 | 9.817E-09 | 35.35 |
| ENSMUSG00000032011 | thymus cell antigen 1, theta [Source:MGI Symbol;Acc:MGI:98747] | Thy1 | Enriched in dLGN | 2.408 | 9.7E-12 | 952.8 |
| ENSMUSG00000029189 | sel-1 suppressor of lin-12-like 3 (C. elegans) [Source:MGI Symbol;Acc:MGI:1916941] | Sel1l3 | Enriched in dLGN | 2.406 | 6.668E-17 | 154 |
| ENSMUSG00000026643 | N-myristoyltransferase 2 [Source:MGI Symbol;Acc:MGI:1202298] | Nmt2 | Enriched in dLGN | 2.393 | 4.223E-38 | 203.8 |
| ENSMUSG00000059361 | neurensin 2 [Source:MGI Symbol;Acc:MGI:2684969] | Nrsn2 | Enriched in dLGN | 2.39 | 9.821E-10 | 106.4 |
| ENSMUSG00000027556 | carbonic anhydrase 1 [Source:MGI Symbol;Acc:MGI:88268] | Car1 | Enriched in dLGN | 2.389 | 0.000003884 | 12.27 |
| ENSMUSG00000028251 | thiosulfate sulfurtransferase (rhodanese)-like domain containing 3 [Source:MGI Symbol;Acc:MGI:1924282] | Tstd3 | Enriched in dLGN | 2.388 | 4.693E-10 | 35.11 |
| ENSMUSG00000031409 | transcription elongation factor A (SII)-like 6 [Source:MGI Symbol;Acc:MGI:1923939] | Tceal6 | Enriched in dLGN | 2.387 | 1.005E-14 | 116.9 |
| ENSMUSG00000020736 | 5',3'-nucleotidase, cytosolic [Source:MGI Symbol;Acc:MGI:1354954] | Nt5c | Enriched in dLGN | 2.385 | 1.497E-18 | 100.9 |
| ENSMUSG00000028024 | glutamyl aminopeptidase [Source:MGI Symbol;Acc:MGI:106645] | Enpep | Enriched in dLGN | 2.368 | 0.00002901 | 10.7 |
| ENSMUSG00000031231 | cytochrome c oxidase subunit 7B [Source:MGI Symbol;Acc:MGI:1913392] | Cox7b | Enriched in dLGN | 2.367 | 0.000005215 | 113.9 |
| ENSMUSG00000024935 | solute carrier family 1 (neuronal/epithelial high affinity glutamate transporter, system Xag), member 1 [Source:MGI Symbol;Acc:MGI:105083] | Slc1a1 | Enriched in dLGN | 2.356 | 5.983E-24 | 323.7 |
| ENSMUSG00000071369 | mitogen-activated protein kinase kinase kinase 5 [Source:MGI Symbol;Acc:MGI:1346876] | Map3k5 | Enriched in dLGN | 2.356 | 6.418E-20 | 236 |
| ENSMUSG00000042807 | HECT, C2 and WW domain containing E3 ubiquitin protein ligase 2 [Source:MGI Symbol;Acc:MGI:2685817] | Hecw2 | Enriched in dLGN | 2.356 | 1.86E-12 | 320.7 |
| ENSMUSG00000021520 | ubiquinol-cytochrome c reductase binding protein [Source:MGI Symbol;Acc:MGI:1914780] | Uqcrb | Enriched in dLGN | 2.355 | 2.466E-08 | 49.09 |
| ENSMUSG00000030350 | protein arginine N-methyltransferase 8 [Source:MGI Symbol;Acc:MGI:3043083] | Prmt8 | Enriched in dLGN | 2.349 | 1.318E-21 | 303.1 |
| ENSMUSG00000026959 | glutamate receptor, ionotropic, NMDA1 (zeta 1) [Source:MGI Symbol;Acc:MGI:95819] | Grin1 | Enriched in dLGN | 2.346 | 6.378E-13 | 942.6 |
| ENSMUSG00000064202 | spermatogenesis associated 6 like [Source:MGI Symbol;Acc:MGI:1918036] | Spata6l | Enriched in dLGN | 2.346 | 2.535E-09 | 24.41 |
| ENSMUSG00000045009 | proline-rich transmembrane protein 3 [Source:MGI Symbol;Acc:MGI:2444810] | Prrt3 | Enriched in dLGN | 2.339 | 6.661E-16 | 111.1 |
| ENSMUSG00000028744 | solute carrier family 66 member 1 [Source:MGI Symbol;Acc:MGI:2384837] | Slc66a1 | Enriched in dLGN | 2.339 | 0.0001916 | 195.7 |
| ENSMUSG00000019943 | ATPase, Ca++ transporting, plasma membrane 1 [Source:MGI Symbol;Acc:MGI:104653] | Atp2b1 | Enriched in dLGN | 2.329 | 4.616E-34 | 2706 |
| ENSMUSG00000073838 | Tu translation elongation factor, mitochondrial [Source:MGI Symbol;Acc:MGI:1923686] | Tufm | Enriched in dLGN | 2.324 | 1.365E-12 | 100.2 |
| ENSMUSG00000070803 | Cbp/p300-interacting transactivator, with Glu/Asp-rich carboxy-terminal domain, 4 [Source:MGI Symbol;Acc:MGI:1861694] | Cited4 | Enriched in dLGN | 2.31 | 0.0001406 | 9.191 |
| ENSMUSG00000029659 | mesenteric estrogen dependent adipogenesis [Source:MGI Symbol;Acc:MGI:1917967] | Medag | Enriched in dLGN | 2.306 | 4.41E-08 | 25.4 |
| ENSMUSG00000079593 | RIKEN cDNA 4933416I08 gene [Source:MGI Symbol;Acc:MGI:1918409] | 4933416I08Rik | Enriched in dLGN | 2.304 | 0.00227 | 3.881 |
| ENSMUSG00000028873 | cell division cycle associated 8 [Source:MGI Symbol;Acc:MGI:1196274] | Cdca8 | Enriched in dLGN | 2.295 | 0.001227 | 5.365 |
| ENSMUSG00000007021 | synaptogyrin 3 [Source:MGI Symbol;Acc:MGI:1341881] | Syngr3 | Enriched in dLGN | 2.282 | 3.29E-17 | 261.6 |
| ENSMUSG00000031391 | L1 cell adhesion molecule [Source:MGI Symbol;Acc:MGI:96721] | L1cam | Enriched in dLGN | 2.282 | 2.539E-08 | 803.2 |
| ENSMUSG00000027581 | stathmin-like 3 [Source:MGI Symbol;Acc:MGI:1277137] | Stmn3 | Enriched in dLGN | 2.281 | 1.395E-11 | 1780 |
| ENSMUSG00000027716 | transient receptor potential cation channel, subfamily C, member 3 [Source:MGI Symbol;Acc:MGI:109526] | Trpc3 | Enriched in dLGN | 2.276 | 1.887E-15 | 212.4 |
| ENSMUSG00000007944 | tetratricopeptide repeat domain 9B [Source:MGI Symbol;Acc:MGI:1920282] | Ttc9b | Enriched in dLGN | 2.26 | 4.446E-15 | 147 |
| ENSMUSG00000037703 | leucine zipper, putative tumor suppressor family member 3 [Source:MGI Symbol;Acc:MGI:2656976] | Lzts3 | Enriched in dLGN | 2.257 | 3.533E-19 | 572.4 |
| ENSMUSG00000029465 | actin related protein 2/3 complex, subunit 3 [Source:MGI Symbol;Acc:MGI:1928375] | Arpc3 | Enriched in dLGN | 2.257 | 2.452E-10 | 143.2 |
| ENSMUSG00000038421 | Fc receptor-like A [Source:MGI Symbol;Acc:MGI:2138647] | Fcrla | Enriched in dLGN | 2.245 | 0.001029 | 28.16 |
| ENSMUSG00000026347 | transmembrane protein 163 [Source:MGI Symbol;Acc:MGI:1919410] | Tmem163 | Enriched in dLGN | 2.243 | 1.786E-14 | 376 |
| ENSMUSG00000001227 | sema domain, transmembrane domain (TM), and cytoplasmic domain, (semaphorin) 6B [Source:MGI Symbol;Acc:MGI:1202889] | Sema6b | Enriched in dLGN | 2.241 | 2.469E-26 | 529.4 |
| ENSMUSG00000007564 | protein phosphatase 2, regulatory subunit A, alpha [Source:MGI Symbol;Acc:MGI:1926334] | Ppp2r1a | Enriched in dLGN | 2.24 | 5.476E-12 | 634.9 |
| ENSMUSG00000017412 | calcium channel, voltage-dependent, beta 4 subunit [Source:MGI Symbol;Acc:MGI:103301] | Cacnb4 | Enriched in dLGN | 2.238 | 4.876E-10 | 413.3 |
| ENSMUSG00000027134 | lysophosphatidylcholine acyltransferase 4 [Source:MGI Symbol;Acc:MGI:2138993] | Lpcat4 | Enriched in dLGN | 2.233 | 3.953E-26 | 130.5 |
| ENSMUSG00000036751 | cytochrome c oxidase, subunit 6B1 [Source:MGI Symbol;Acc:MGI:107460] | Cox6b1 | Enriched in dLGN | 2.231 | 1.733E-15 | 143.6 |
| ENSMUSG00000049422 | coiled-coil-helix-coiled-coil-helix domain containing 10 [Source:MGI Symbol;Acc:MGI:2143558] | Chchd10 | Enriched in dLGN | 2.225 | 9.381E-12 | 224.5 |
| ENSMUSG00000004187 | kinesin family member C2 [Source:MGI Symbol;Acc:MGI:109187] | Kifc2 | Enriched in dLGN | 2.224 | 4.763E-32 | 769 |
| ENSMUSG00000024127 | prolyl endopeptidase-like [Source:MGI Symbol;Acc:MGI:2441932] | Prepl | Enriched in dLGN | 2.22 | 3.845E-12 | 815.8 |
| ENSMUSG00000044024 | RELT-like 2 [Source:MGI Symbol;Acc:MGI:1918044] | Rell2 | Enriched in dLGN | 2.215 | 5.382E-26 | 285.3 |
| ENSMUSG00000025980 | heat shock protein 1 (chaperonin) [Source:MGI Symbol;Acc:MGI:96242] | Hspd1 | Enriched in dLGN | 2.215 | 7.845E-17 | 264 |
| ENSMUSG00000001380 | histidyl-tRNA synthetase [Source:MGI Symbol;Acc:MGI:108087] | Hars | Enriched in dLGN | 2.206 | 2.465E-24 | 256 |
| ENSMUSG00000032279 | isocitrate dehydrogenase 3 (NAD+) alpha [Source:MGI Symbol;Acc:MGI:1915084] | Idh3a | Enriched in dLGN | 2.205 | 9.076E-07 | 302 |
| ENSMUSG00000000088 | cytochrome c oxidase subunit 5A [Source:MGI Symbol;Acc:MGI:88474] | Cox5a | Enriched in dLGN | 2.201 | 2.652E-11 | 112.1 |
| ENSMUSG00000024083 | praja ring finger ubiquitin ligase 2 [Source:MGI Symbol;Acc:MGI:2159342] | Pja2 | Enriched in dLGN | 2.196 | 3.536E-14 | 1111 |
| ENSMUSG00000063694 | cytochrome c, somatic [Source:MGI Symbol;Acc:MGI:88578] | Cycs | Enriched in dLGN | 2.196 | 2.504E-11 | 116.9 |
| ENSMUSG00000028648 | NADH:ubiquinone oxidoreductase core subunit S5 [Source:MGI Symbol;Acc:MGI:1890889] | Ndufs5 | Enriched in dLGN | 2.193 | 4.33E-10 | 101.9 |
| ENSMUSG00000002068 | cyclin E1 [Source:MGI Symbol;Acc:MGI:88316] | Ccne1 | Enriched in dLGN | 2.192 | 6.577E-09 | 29.03 |
| ENSMUSG00000029752 | asparagine synthetase [Source:MGI Symbol;Acc:MGI:1350929] | Asns | Enriched in dLGN | 2.192 | 6.593E-08 | 106.2 |
| ENSMUSG00000034220 | glypican 1 [Source:MGI Symbol;Acc:MGI:1194891] | Gpc1 | Enriched in dLGN | 2.187 | 7.337E-25 | 632.1 |
| ENSMUSG00000036186 | divergent protein kinase domain 1B [Source:MGI Symbol;Acc:MGI:1927576] | Dipk1b | Enriched in dLGN | 2.185 | 4.261E-10 | 28.18 |
| ENSMUSG00000038264 | sema domain, immunoglobulin domain (Ig), and GPI membrane anchor, (semaphorin) 7A [Source:MGI Symbol;Acc:MGI:1306826] | Sema7a | Enriched in dLGN | 2.176 | 1.896E-14 | 229.8 |
| ENSMUSG00000058589 | ankyrin repeat and sterile alpha motif domain containing 1B [Source:MGI Symbol;Acc:MGI:1924781] | Anks1b | Enriched in dLGN | 2.173 | 0.000000001 | 1182 |

| | | | | | | |
|---|---|---|---|---|---|---|
| ENSMUSG00000016252 | ATP synthase, H+ transporting, mitochondrial F1 complex, epsilon subunit [Source:MGI Symbol;Acc:MGI:1855697] | Atp5e | Enriched in dLGN | 2.171 | 3.825E-09 | 153.2 |
| ENSMUSG00000082229 | nucleosome assembly protein 1-like 2 [Source:MGI Symbol;Acc:MGI:106654] | Nap1l2 | Enriched in dLGN | 2.17 | 6.424E-10 | 277.1 |
| ENSMUSG00000023904 | host cell factor C1 regulator 1 (XPO1-dependent) [Source:MGI Symbol;Acc:MGI:2663619] | Hcfc1r1 | Enriched in dLGN | 2.162 | 3.275E-10 | 96.23 |
| ENSMUSG00000008153 | calsyntenin 3 [Source:MGI Symbol;Acc:MGI:2178323] | Clstn3 | Enriched in dLGN | 2.157 | 2.013E-12 | 1064 |
| ENSMUSG00000019179 | malate dehydrogenase 2, NAD (mitochondrial) [Source:MGI Symbol;Acc:MGI:97050] | Mdh2 | Enriched in dLGN | 2.154 | 1.802E-09 | 626.3 |
| ENSMUSG00000062825 | actin, gamma, cytoplasmic 1 [Source:MGI Symbol;Acc:MGI:87906] | Actg1 | Enriched in dLGN | 2.153 | 9.849E-07 | 407 |
| ENSMUSG00000036438 | calmodulin 2 [Source:MGI Symbol;Acc:MGI:103250] | Calm2 | Enriched in dLGN | 2.15 | 5.934E-07 | 1256 |
| ENSMUSG00000043091 | tubulin, alpha 1C [Source:MGI Symbol;Acc:MGI:1095409] | Tuba1c | Enriched in dLGN | 2.148 | 0.000948 | 41.77 |
| ENSMUSG00000040759 | CKLF-like MARVEL transmembrane domain containing 5 [Source:MGI Symbol;Acc:MGI:2447164] | Cmtm5 | Enriched in dLGN | 2.146 | 0.00001496 | 15.12 |
| ENSMUSG00000044252 | oxysterol binding protein-like 1A [Source:MGI Symbol;Acc:MGI:1927551] | Osbpl1a | Enriched in dLGN | 2.145 | 2.879E-18 | 434.2 |
| ENSMUSG00000026585 | kinesin-associated protein 3 [Source:MGI Symbol;Acc:MGI:107566] | Kifap3 | Enriched in dLGN | 2.14 | 4.711E-14 | 663.9 |
| ENSMUSG00000006932 | catenin (cadherin associated protein), beta 1 [Source:MGI Symbol;Acc:MGI:88276] | Ctnnb1 | Enriched in dLGN | 2.139 | 6.472E-14 | 1065 |
| ENSMUSG00000044117 | bMERB domain containing 1 [Source:MGI Symbol;Acc:MGI:1914504] | Bmerb1 | Enriched in dLGN | 2.134 | 6.143E-22 | 457.3 |
| ENSMUSG00000041670 | regulating synaptic membrane exocytosis 1 [Source:MGI Symbol;Acc:MGI:2152971] | Rims1 | Enriched in dLGN | 2.131 | 2.012E-15 | 1881 |
| ENSMUSG00000028212 | cyclin E2 [Source:MGI Symbol;Acc:MGI:1329034] | Ccne2 | Enriched in dLGN | 2.13 | 0.000009959 | 12.53 |
| ENSMUSG00000040811 | echinoderm microtubule associated protein like 2 [Source:MGI Symbol;Acc:MGI:1919455] | Eml2 | Enriched in dLGN | 2.124 | 3.634E-19 | 740.8 |
| ENSMUSG00000026526 | fumarate hydratase 1 [Source:MGI Symbol;Acc:MGI:95530] | Fh1 | Enriched in dLGN | 2.123 | 9.504E-09 | 124 |
| ENSMUSG00000029120 | protein phosphatase 2, regulatory subunit B, gamma [Source:MGI Symbol;Acc:MGI:2442660] | Ppp2r2c | Enriched in dLGN | 2.121 | 5.607E-28 | 1258 |
| ENSMUSG00000055373 | fucosyltransferase 9 [Source:MGI Symbol;Acc:MGI:1330859] | Fut9 | Enriched in dLGN | 2.105 | 2.725E-09 | 265.5 |
| ENSMUSG00000019210 | ATPase, H+ transporting, lysosomal V1 subunit E1 [Source:MGI Symbol;Acc:MGI:894326] | Atp6v1e1 | Enriched in dLGN | 2.097 | 9.782E-10 | 270.4 |
| ENSMUSG00000061518 | cytochrome c oxidase subunit 5B [Source:MGI Symbol;Acc:MGI:88475] | Cox5b | Enriched in dLGN | 2.093 | 1.235E-12 | 455.8 |
| ENSMUSG00000032965 | intraflagellar transport 57 [Source:MGI Symbol;Acc:MGI:1921166] | Ift57 | Enriched in dLGN | 2.084 | 7.634E-22 | 254.5 |
| ENSMUSG00000042182 | BEN domain containing 6 [Source:MGI Symbol;Acc:MGI:2444572] | Bend6 | Enriched in dLGN | 2.08 | 9.133E-18 | 256 |
| ENSMUSG00000034566 | ATP synthase, H+ transporting, mitochondrial F0 complex, subunit D [Source:MGI Symbol;Acc:MGI:1918929] | Atp5h | Enriched in dLGN | 2.077 | 3.022E-12 | 216.1 |
| ENSMUSG00000022620 | arylsulfatase A [Source:MGI Symbol;Acc:MGI:88077] | Arsa | Enriched in dLGN | 2.075 | 6.135E-12 | 62.33 |
| ENSMUSG00000031007 | ATPase, H+ transporting, lysosomal accessory protein 2 [Source:MGI Symbol;Acc:MGI:1917745] | Atp6ap2 | Enriched in dLGN | 2.073 | 2.437E-10 | 242.6 |
| ENSMUSG00000038552 | fibronectin type III domain containing 4 [Source:MGI Symbol;Acc:MGI:1917195] | Fndc4 | Enriched in dLGN | 2.069 | 2.965E-27 | 238.1 |
| ENSMUSG00000033352 | mitogen-activated protein kinase kinase 4 [Source:MGI Symbol;Acc:MGI:1346869] | Map2k4 | Enriched in dLGN | 2.066 | 6.552E-23 | 341.7 |
| ENSMUSG00000026904 | solute carrier family 4, sodium bicarbonate cotransporter-like, member 10 [Source:MGI Symbol;Acc:MGI:2150150] | Slc4a10 | Enriched in dLGN | 2.062 | 2.548E-13 | 601.1 |
| ENSMUSG00000051359 | neurocalcin delta [Source:MGI Symbol;Acc:MGI:1196326] | Ncald | Enriched in dLGN | 2.056 | 1.206E-22 | 710.7 |
| ENSMUSG00000035964 | transmembrane protein 59-like [Source:MGI Symbol;Acc:MGI:1915187] | Tmem59l | Enriched in dLGN | 2.052 | 6.968E-11 | 166 |
| ENSMUSG00000005161 | peroxiredoxin 2 [Source:MGI Symbol;Acc:MGI:109486] | Prdx2 | Enriched in dLGN | 2.044 | 8.115E-10 | 321.2 |
| ENSMUSG00000032959 | phosphatidylethanolamine binding protein 1 [Source:MGI Symbol;Acc:MGI:1344408] | Pebp1 | Enriched in dLGN | 2.042 | 8.424E-09 | 164.1 |
| ENSMUSG00000041697 | cytochrome c oxidase subunit 6A1 [Source:MGI Symbol;Acc:MGI:103099] | Cox6a1 | Enriched in dLGN | 2.036 | 3.463E-10 | 168 |
| ENSMUSG00000042388 | DLG associated protein 3 [Source:MGI Symbol;Acc:MGI:3039563] | Dlgap3 | Enriched in dLGN | 2.034 | 1.56E-17 | 1641 |
| ENSMUSG00000022376 | adenylate cyclase 8 [Source:MGI Symbol;Acc:MGI:1341110] | Adcy8 | Enriched in dLGN | 2.022 | 2.75E-09 | 297.1 |
| ENSMUSG00000037606 | oxysterol binding protein-like 5 [Source:MGI Symbol;Acc:MGI:1930265] | Osbpl5 | Enriched in dLGN | 2.016 | 2.246E-28 | 213.9 |
| ENSMUSG00000005683 | citrate synthase [Source:MGI Symbol;Acc:MGI:88529] | Cs | Enriched in dLGN | 2.016 | 6.5E-15 | 414 |
| ENSMUSG00000022321 | cadherin 10 [Source:MGI Symbol;Acc:MGI:107436] | Cdh10 | Enriched in dLGN | 2.014 | 2.435E-08 | 144.1 |
| ENSMUSG00000024359 | heat shock protein 9 [Source:MGI Symbol;Acc:MGI:96245] | Hspa9 | Enriched in dLGN | 2.011 | 3.548E-10 | 574.8 |
| ENSMUSG00000019877 | serine incorporator 1 [Source:MGI Symbol;Acc:MGI:1926228] | Serinc1 | Enriched in dLGN | 2.001 | 1.053E-08 | 662.3 |
| ENSMUSG00000075318 | sodium channel, voltage-gated, type II, alpha [Source:MGI Symbol;Acc:MGI:98248] | Scn2a | Enriched in dLGN | 2 | 1.575E-19 | 1334 |
| ENSMUSG00000063802 | HSPA (heat shock 70kDa) binding protein, cytoplasmic cochaperone 1 [Source:MGI Symbol;Acc:MGI:1913495] | Hspbp1 | Enriched in dLGN | 1.994 | 1.713E-20 | 128.5 |
| ENSMUSG00000030337 | vesicle-associated membrane protein 1 [Source:MGI Symbol;Acc:MGI:1313276] | Vamp1 | Enriched in dLGN | 1.994 | 1.196E-13 | 282.4 |
| ENSMUSG00000019852 | ARFGEF family member 3 [Source:MGI Symbol;Acc:MGI:106387] | Arfgef3 | Enriched in dLGN | 1.985 | 8.538E-22 | 824.9 |
| ENSMUSG00000031149 | PRA1 domain family 2 [Source:MGI Symbol;Acc:MGI:1859607] | Praf2 | Enriched in dLGN | 1.982 | 0.0003905 | 22.15 |
| ENSMUSG00000019505 | ubiquitin B [Source:MGI Symbol;Acc:MGI:98888] | Ubb | Enriched in dLGN | 1.975 | 0.000005901 | 652 |
| ENSMUSG00000032366 | tropomyosin 1, alpha [Source:MGI Symbol;Acc:MGI:98809] | Tpm1 | Enriched in dLGN | 1.972 | 5.327E-38 | 654.7 |
| ENSMUSG00000026260 | NADH:ubiquinone oxidoreductase subunit A10 [Source:MGI Symbol;Acc:MGI:1914523] | Ndufa10 | Enriched in dLGN | 1.968 | 6.961E-09 | 263.7 |
| ENSMUSG00000014313 | cytochrome c oxidase subunit 6C [Source:MGI Symbol;Acc:MGI:104614] | Cox6c | Enriched in dLGN | 1.968 | 4.159E-08 | 345.4 |
| ENSMUSG00000031176 | dynein light chain Tctex-type 3 [Source:MGI Symbol;Acc:MGI:1914367] | Dynlt3 | Enriched in dLGN | 1.967 | 9.452E-09 | 112.8 |
| ENSMUSG00000049556 | leucine rich repeat and Ig domain containing 1 [Source:MGI Symbol;Acc:MGI:1915522] | Lingo1 | Enriched in dLGN | 1.966 | 2.38E-18 | 811.2 |
| ENSMUSG00000111375 | BTB (POZ) domain containing 8 [Source:MGI Symbol;Acc:MGI:3646208] | Btbd8 | Enriched in dLGN | 1.965 | 3.884E-10 | 197 |
| ENSMUSG00000038462 | ubiquinol-cytochrome c reductase, Rieske iron-sulfur polypeptide 1 [Source:MGI Symbol;Acc:MGI:1913944] | Uqcrfs1 | Enriched in dLGN | 1.964 | 3.119E-13 | 172.4 |
| ENSMUSG00000021114 | ATPase, H+ transporting, lysosomal V1 subunit D [Source:MGI Symbol;Acc:MGI:1921084] | Atp6v1d | Enriched in dLGN | 1.964 | 2.122E-12 | 284.8 |
| ENSMUSG00000020949 | FK506 binding protein 3 [Source:MGI Symbol;Acc:MGI:1353460] | Fkbp3 | Enriched in dLGN | 1.963 | 3.828E-08 | 160.5 |
| ENSMUSG00000024873 | cornichon family AMPA receptor auxiliary protein 2 [Source:MGI Symbol;Acc:MGI:1277225] | Cnih2 | Enriched in dLGN | 1.961 | 2.801E-21 | 220.1 |
| ENSMUSG00000036815 | dipeptidylpeptidase 10 [Source:MGI Symbol;Acc:MGI:2442409] | Dpp10 | Enriched in dLGN | 1.958 | 0.000001547 | 576.4 |
| ENSMUSG00000073433 | Rho GDP dissociation inhibitor (GDI) gamma [Source:MGI Symbol;Acc:MGI:108430] | Arhgdig | Enriched in dLGN | 1.957 | 2.946E-10 | 185.1 |
| ENSMUSG00000024587 | asparaginyl-tRNA synthetase [Source:MGI Symbol;Acc:MGI:1917473] | Nars | Enriched in dLGN | 1.956 | 5.214E-13 | 419 |
| ENSMUSG00000032946 | RAS, guanyl releasing protein 2 [Source:MGI Symbol;Acc:MGI:1333849] | Rasgrp2 | Enriched in dLGN | 1.952 | 7.057E-09 | 149.8 |
| ENSMUSG00000025092 | heat shock protein 12A [Source:MGI Symbol;Acc:MGI:1920692] | Hspa12a | Enriched in dLGN | 1.949 | 1.083E-15 | 913 |
| ENSMUSG00000063931 | peptidase D [Source:MGI Symbol;Acc:MGI:97542] | Pepd | Enriched in dLGN | 1.949 | 8.26E-08 | 102.6 |
| ENSMUSG00000062526 | metallophosphoesterase 1 [Source:MGI Symbol;Acc:MGI:2661311] | Mppe1 | Enriched in dLGN | 1.947 | 8.498E-10 | 32.36 |
| ENSMUSG00000029840 | myotrophin [Source:MGI Symbol;Acc:MGI:99445] | Mtpn | Enriched in dLGN | 1.945 | 7.174E-17 | 335.2 |
| ENSMUSG00000025968 | NADH:ubiquinone oxidoreductase core subunit S1 [Source:MGI Symbol;Acc:MGI:2443241] | Ndufs1 | Enriched in dLGN | 1.938 | 3.663E-13 | 337.9 |
| ENSMUSG00000031772 | contactin associated protein-like 4 [Source:MGI Symbol;Acc:MGI:2183572] | Cntnap4 | Enriched in dLGN | 1.926 | 2.111E-07 | 164.6 |
| ENSMUSG00000040373 | calcium channel, voltage-dependent, gamma subunit 5 [Source:MGI Symbol;Acc:MGI:2157946] | Cacng5 | Enriched in dLGN | 1.923 | 2.719E-13 | 203.8 |
| ENSMUSG00000022856 | transmembrane protein 41a [Source:MGI Symbol;Acc:MGI:1913914] | Tmem41a | Enriched in dLGN | 1.923 | 1.036E-12 | 44.73 |
| ENSMUSG00000000168 | dihydrolipoamide S-acetyltransferase (E2 component of pyruvate dehydrogenase complex) [Source:MGI Symbol;Acc:MGI:2385311] | Dlat | Enriched in dLGN | 1.918 | 6.382E-17 | 196.1 |
| ENSMUSG00000036067 | solute carrier family 2 (facilitated glucose transporter), member 6 [Source:MGI Symbol;Acc:MGI:2443286] | Slc2a6 | Enriched in dLGN | 1.916 | 7.203E-08 | 33.47 |
| ENSMUSG00000027131 | ER membrane protein complex subunit 4 [Source:MGI Symbol;Acc:MGI:1915282] | Emc4 | Enriched in dLGN | 1.913 | 2.35E-09 | 81.85 |
| ENSMUSG00000027108 | Obg-like ATPase 1 [Source:MGI Symbol;Acc:MGI:1914309] | Ola1 | Enriched in dLGN | 1.909 | 0.00745 | 454.9 |
| ENSMUSG00000010914 | pyruvate dehydrogenase complex, component X [Source:MGI Symbol;Acc:MGI:1351627] | Pdhx | Enriched in dLGN | 1.907 | 6.444E-16 | 152.7 |
| ENSMUSG00000026750 | proteasome (prosome, macropain) subunit, beta type 7 [Source:MGI Symbol;Acc:MGI:107637] | Psmb7 | Enriched in dLGN | 1.903 | 8.869E-12 | 169.1 |
| ENSMUSG00000053024 | contactin 2 [Source:MGI Symbol;Acc:MGI:104518] | Cntn2 | Enriched in dLGN | 1.903 | 4.737E-11 | 303.9 |
| ENSMUSG00000049775 | thymosin, beta 4, X chromosome [Source:MGI Symbol;Acc:MGI:99510] | Tmsb4x | Enriched in dLGN | 1.899 | 3.197E-17 | 317.2 |
| ENSMUSG00000039546 | adherens junction associated protein 1 [Source:MGI Symbol;Acc:MGI:2685419] | Ajap1 | Enriched in dLGN | 1.898 | 1.195E-11 | 293.4 |
| ENSMUSG00000047126 | clathrin, heavy polypeptide (Hc) [Source:MGI Symbol;Acc:MGI:2388633] | Cltc | Enriched in dLGN | 1.894 | 3.025E-10 | 1529 |
| ENSMUSG00000049971 | glycosyltransferase 1 domain containing 1 [Source:MGI Symbol;Acc:MGI:2442755] | Glt1d1 | Enriched in dLGN | 1.891 | 2.644E-14 | 53.34 |
| ENSMUSG00000000563 | ATP synthase peripheral stalk-membrane subunit b [Source:MGI Symbol;Acc:MGI:1100495] | Atp5pb | Enriched in dLGN | 1.891 | 1.616E-11 | 236 |
| ENSMUSG00000022110 | succinate-Coenzyme A ligase, ADP-forming, beta subunit [Source:MGI Symbol;Acc:MGI:1306775] | Sucla2 | Enriched in dLGN | 1.89 | 0.0000001 | 256.6 |
| ENSMUSG00000030647 | NADH:ubiquinone oxidoreductase subunit C2 [Source:MGI Symbol;Acc:MGI:1344370] | Ndufc2 | Enriched in dLGN | 1.89 | 0.000001348 | 77.57 |
| ENSMUSG00000025287 | acyl-CoA thioesterase 9 [Source:MGI Symbol;Acc:MGI:1928939] | Acot9 | Enriched in dLGN | 1.887 | 0.00006174 | 23.21 |
| ENSMUSG00000024777 | protein phosphatase 2, regulatory subunit B', beta [Source:MGI Symbol;Acc:MGI:2388480] | Ppp2r5b | Enriched in dLGN | 1.883 | 2.884E-25 | 428 |
| ENSMUSG00000034488 | EGF-like repeats and discoidin I-like domains 3 [Source:MGI Symbol;Acc:MGI:1329025] | Edil3 | Enriched in dLGN | 1.882 | 6.937E-07 | 564.8 |
| ENSMUSG00000016349 | eukaryotic translation elongation factor 1 alpha 2 [Source:MGI Symbol;Acc:MGI:1096317] | Eef1a2 | Enriched in dLGN | 1.88 | 1.196E-08 | 1496 |
| ENSMUSG00000036850 | mitochondrial ribosomal protein L41 [Source:MGI Symbol;Acc:MGI:1333816] | Mrpl41 | Enriched in dLGN | 1.872 | 1.053E-12 | 53.68 |
| ENSMUSG00000054428 | ATPase inhibitory factor 1 [Source:MGI Symbol;Acc:MGI:1196457] | Atpif1 | Enriched in dLGN | 1.872 | 2.584E-10 | 829.4 |
| ENSMUSG00000015961 | adenylosuccinate synthetase, non muscle [Source:MGI Symbol;Acc:MGI:87948] | Adss | Enriched in dLGN | 1.869 | 5.936E-15 | 159.9 |
| ENSMUSG00000029875 | coiled-coil domain containing 184 [Source:MGI Symbol;Acc:MGI:2146066] | Ccdc184 | Enriched in dLGN | 1.869 | 3.305E-09 | 201 |
| ENSMUSG00000047879 | ubiquitin specific peptidase 14 [Source:MGI Symbol;Acc:MGI:1928898] | Usp14 | Enriched in dLGN | 1.866 | 9.509E-14 | 157.3 |
| ENSMUSG00000002808 | ependymin related protein 1 (zebrafish) [Source:MGI Symbol;Acc:MGI:2145369] | Epdr1 | Enriched in dLGN | 1.865 | 2.178E-09 | 73.68 |
| ENSMUSG00000031701 | DnaJ heat shock protein family (Hsp40) member A2 [Source:MGI Symbol;Acc:MGI:1931882] | Dnaja2 | Enriched in dLGN | 1.864 | 5.341E-11 | 401 |
| ENSMUSG00000025189 | cyclin M1 [Source:MGI Symbol;Acc:MGI:1891366] | Cnnm1 | Enriched in dLGN | 1.864 | 1.245E-08 | 258.6 |
| ENSMUSG00000001986 | glutamate receptor, ionotropic, AMPA3 (alpha 3) [Source:MGI Symbol;Acc:MGI:95810] | Gria3 | Enriched in dLGN | 1.862 | 1.642E-11 | 283.8 |
| ENSMUSG00000024758 | reticulon 3 [Source:MGI Symbol;Acc:MGI:1339970] | Rtn3 | Enriched in dLGN | 1.857 | 1.962E-22 | 1535 |
| ENSMUSG00000040430 | phosphatidylinositol transfer protein, cytoplasmic 1 [Source:MGI Symbol;Acc:MGI:1919045] | Pitpnc1 | Enriched in dLGN | 1.847 | 5.148E-17 | 764.9 |
| ENSMUSG00000056091 | ST3 beta-galactoside alpha-2,3-sialyltransferase 5 [Source:MGI Symbol;Acc:MGI:1339963] | St3gal5 | Enriched in dLGN | 1.843 | 3.536E-19 | 176.3 |
| ENSMUSG00000025790 | solute carrier organic anion transporter family, member 3a1 [Source:MGI Symbol;Acc:MGI:1351867] | Slco3a1 | Enriched in dLGN | 1.84 | 2.241E-07 | 185.2 |
| ENSMUSG00000032908 | sphingosine-1-phosphate phosphatase 2 [Source:MGI Symbol;Acc:MGI:3589109] | Sgpp2 | Enriched in dLGN | 1.837 | 0.000001579 | 115.4 |
| ENSMUSG00000033389 | Rho GTPase activating protein 44 [Source:MGI Symbol;Acc:MGI:2144423] | Arhgap44 | Enriched in dLGN | 1.836 | 2E-18 | 522.2 |
| ENSMUSG00000029769 | coiled-coil domain containing 136 [Source:MGI Symbol;Acc:MGI:1918128] | Ccdc136 | Enriched in dLGN | 1.823 | 1.132E-15 | 1034 |
| ENSMUSG00000005873 | receptor accessory protein 5 [Source:MGI Symbol;Acc:MGI:1270152] | Reep5 | Enriched in dLGN | 1.823 | 2.81E-10 | 705.3 |
| ENSMUSG00000028444 | ciliary neurotrophic factor receptor [Source:MGI Symbol;Acc:MGI:99605] | Cntfr | Enriched in dLGN | 1.822 | 2.243E-12 | 219.4 |
| ENSMUSG00000053297 | expressed sequence AI854703 [Source:MGI Symbol;Acc:MGI:2141510] | AI854703 | Enriched in dLGN | 1.82 | 2.35E-12 | 90.96 |
| ENSMUSG00000036309 | S-phase kinase-associated protein 1 [Source:MGI Symbol;Acc:MGI:103575] | Skp1 | Enriched in dLGN | 1.82 | 1.644E-09 | 534.7 |
| ENSMUSG00000028078 | doublecortin-like kinase 2 [Source:MGI Symbol;Acc:MGI:1918012] | Dclk2 | Enriched in dLGN | 1.818 | 1.087E-11 | 340.9 |
| ENSMUSG00000051146 | calcium/calmodulin-dependent protein kinase II inhibitor 2 [Source:MGI Symbol;Acc:MGI:1920297] | Camk2n2 | Enriched in dLGN | 1.816 | 1.197E-15 | 513.1 |
| ENSMUSG00000059734 | NADH:ubiquinone oxidoreductase core subunit S8 [Source:MGI Symbol;Acc:MGI:2385079] | Ndufs8 | Enriched in dLGN | 1.807 | 3.906E-07 | 73.63 |
| ENSMUSG00000029765 | plexin A4 [Source:MGI Symbol;Acc:MGI:2179061] | Plxna4 | Enriched in dLGN | 1.803 | 4.615E-12 | 780.1 |
| ENSMUSG00000051951 | X-linked Kx blood group related 4 [Source:MGI Symbol;Acc:MGI:3528744] | Xkr4 | Enriched in dLGN | 1.8 | 0.000003026 | 863.8 |
| ENSMUSG00000064329 | sodium channel, voltage-gated, type I, alpha [Source:MGI Symbol;Acc:MGI:98246] | Scn1a | Enriched in dLGN | 1.799 | 4.167E-08 | 703.3 |
| ENSMUSG00000040118 | calcium channel, voltage-dependent, alpha2/delta subunit 1 [Source:MGI Symbol;Acc:MGI:88295] | Cacna2d1 | Enriched in dLGN | 1.797 | 6.773E-09 | 541.2 |
| ENSMUSG00000028020 | glycine receptor, beta subunit [Source:MGI Symbol;Acc:MGI:95751] | Glrb | Enriched in dLGN | 1.794 | 7.762E-15 | 323.7 |
| ENSMUSG00000025268 | MAGE family member D2 [Source:MGI Symbol;Acc:MGI:1933391] | Maged2 | Enriched in dLGN | 1.794 | 0.000009608 | 189.8 |
| ENSMUSG00000017781 | phosphatidylinositol transfer protein, alpha [Source:MGI Symbol;Acc:MGI:99887] | Pitpna | Enriched in dLGN | 1.79 | 1.56E-19 | 792.9 |
| ENSMUSG00000031672 | glutamatic-oxaloacetic transaminase 2, mitochondrial [Source:MGI Symbol;Acc:MGI:95792] | Got2 | Enriched in dLGN | 1.783 | 1.104E-07 | 253.7 |
| ENSMUSG00000033307 | macrophage migration inhibitory factor (glycosylation-inhibiting factor) [Source:MGI Symbol;Acc:MGI:96982] | Mif | Enriched in dLGN | 1.778 | 0.00008419 | 90.39 |
| ENSMUSG00000022892 | amyloid beta (A4) precursor protein [Source:MGI Symbol;Acc:MGI:88059] | App | Enriched in dLGN | 1.777 | 2.019E-08 | 4592 |
| ENSMUSG00000020116 | partner of NOB1 homolog [Source:MGI Symbol;Acc:MGI:1913499] | Pno1 | Enriched in dLGN | 1.775 | 1.097E-08 | 43.22 |
| ENSMUSG00000013921 | CAP-GLY domain containing linker protein 3 [Source:MGI Symbol;Acc:MGI:1923936] | Clip3 | Enriched in dLGN | 1.774 | 2.485E-07 | 1062 |
| ENSMUSG00000027104 | activating transcription factor 2 [Source:MGI Symbol;Acc:MGI:109349] | Atf2 | Enriched in dLGN | 1.772 | 0.0001099 | 703.5 |
| ENSMUSG00000025393 | ATP synthase, H+ transporting mitochondrial F1 complex, beta subunit [Source:MGI Symbol;Acc:MGI:107801] | Atp5b | Enriched in dLGN | 1.77 | 2.581E-07 | 1784 |
| ENSMUSG00000018865 | sulfotransferase family 4A, member 1 [Source:MGI Symbol;Acc:MGI:1888971] | Sult4a1 | Enriched in dLGN | 1.767 | 1.74E-14 | 795.2 |
| ENSMUSG00000002771 | glutamate receptor, ionotropic, NMDA2D (epsilon 4) [Source:MGI Symbol;Acc:MGI:95823] | Grin2d | Enriched in dLGN | 1.766 | 7.096E-13 | 466.7 |
| ENSMUSG00000015217 | high mobility group box 3 [Source:MGI Symbol;Acc:MGI:1098219] | Hmgb3 | Enriched in dLGN | 1.764 | 1.153E-08 | 141 |
| ENSMUSG00000034187 | N-ethylmaleimide sensitive fusion protein [Source:MGI Symbol;Acc:MGI:104560] | Nsf | Enriched in dLGN | 1.763 | 5.518E-11 | 1114 |

| | | | | | | |
|---|---|---|---|---|---|---|
| ENSMUSG00000026568 | mitochondrial pyruvate carrier 2 [Source:MGI Symbol;Acc:MGI:1917706] | Mpc2 | Enriched in dLGN | 1.763 | 7.541E-10 | 84.48 |
| ENSMUSG00000073676 | heat shock protein 1 (chaperonin 10) [Source:MGI Symbol;Acc:MGI:104680] | Hspe1 | Enriched in dLGN | 1.762 | 0.000001066 | 62.31 |
| ENSMUSG00000025375 | apoptosis-associated tyrosine kinase [Source:MGI Symbol;Acc:MGI:1197518] | Aatk | Enriched in dLGN | 1.755 | 1.358E-13 | 1628 |
| ENSMUSG00000032657 | endosomal transmembrane epsin interactor 3 [Source:MGI Symbol;Acc:MGI:1915771] | Entrep3 | Enriched in dLGN | 1.753 | 1.875E-09 | 83.78 |
| ENSMUSG00000061576 | dipeptidylpeptidase 6 [Source:MGI Symbol;Acc:MGI:94921] | Dpp6 | Enriched in dLGN | 1.751 | 0.000002109 | 1354 |
| ENSMUSG00000002032 | transmembrane protein 25 [Source:MGI Symbol;Acc:MGI:1918937] | Tmem25 | Enriched in dLGN | 1.746 | 1.292E-09 | 70.33 |
| ENSMUSG00000028843 | SH3 domain binding glutamic acid-rich protein-like 3 [Source:MGI Symbol;Acc:MGI:1920973] | Sh3bgrl3 | Enriched in dLGN | 1.744 | 0.0000712 | 64.15 |
| ENSMUSG00000079426 | actin related protein 2/3 complex, subunit 4 [Source:MGI Symbol;Acc:MGI:1915339] | Arpc4 | Enriched in dLGN | 1.743 | 5.867E-12 | 89.11 |
| ENSMUSG00000035885 | cytochrome c oxidase subunit 8A [Source:MGI Symbol;Acc:MGI:105959] | Cox8a | Enriched in dLGN | 1.743 | 6.331E-11 | 292.6 |
| ENSMUSG00000022205 | SUB1 homolog, transcriptional regulator [Source:MGI Symbol;Acc:MGI:104811] | Sub1 | Enriched in dLGN | 1.742 | 2.284E-19 | 399.4 |
| ENSMUSG00000023089 | NADH:ubiquinone oxidoreductase subunit A5 [Source:MGI Symbol;Acc:MGI:1915452] | Ndufa5 | Enriched in dLGN | 1.74 | 3.797E-09 | 166.1 |
| ENSMUSG00000060212 | pecanex homolog 2 [Source:MGI Symbol;Acc:MGI:2445010] | Pcnx2 | Enriched in dLGN | 1.739 | 3.94E-12 | 282.9 |
| ENSMUSG00000091722 | siah E3 ubiquitin protein ligase family member 3 [Source:MGI Symbol;Acc:MGI:2685758] | Siah3 | Enriched in dLGN | 1.739 | 3.751E-09 | 62.95 |
| ENSMUSG00000047547 | clathrin, light polypeptide (Lcb) [Source:MGI Symbol;Acc:MGI:1921575] | Cltb | Enriched in dLGN | 1.738 | 5.554E-29 | 896 |
| ENSMUSG00000003072 | ATP synthase, H+ transporting, mitochondrial F1 complex, delta subunit [Source:MGI Symbol;Acc:MGI:1913293] | Atp5d | Enriched in dLGN | 1.738 | 1.195E-11 | 259.8 |
| ENSMUSG00000019951 | UHRF1 (ICBP90) binding protein 1-like [Source:MGI Symbol;Acc:MGI:2442888] | Uhrf1bp1l | Enriched in dLGN | 1.737 | 1.136E-28 | 276.6 |
| ENSMUSG00000059409 | protein phosphatase 2, regulatory subunit B', delta [Source:MGI Symbol;Acc:MGI:2388481] | Ppp2r5d | Enriched in dLGN | 1.735 | 1.068E-16 | 362.3 |
| ENSMUSG00000002379 | NADH:ubiquinone oxidoreductase subunit A11 [Source:MGI Symbol;Acc:MGI:1917125] | Ndufa11 | Enriched in dLGN | 1.732 | 0.0003978 | 75.68 |
| ENSMUSG00000063253 | short coiled-coil protein [Source:MGI Symbol;Acc:MGI:1927654] | Scoc | Enriched in dLGN | 1.731 | 9.456E-14 | 113.5 |
| ENSMUSG00000063406 | transmembrane p24 trafficking protein 5 [Source:MGI Symbol;Acc:MGI:1921586] | Tmed5 | Enriched in dLGN | 1.731 | 7.643E-08 | 35.86 |
| ENSMUSG00000076432 | tyrosine 3-monooxygenase/tryptophan 5-monooxygenase activation protein theta [Source:MGI Symbol;Acc:MGI:891963] | Ywhaq | Enriched in dLGN | 1.731 | 1.233E-07 | 236.1 |
| ENSMUSG00000035960 | apurinic/apyrimidinic endonuclease 1 [Source:MGI Symbol;Acc:MGI:88042] | Apex1 | Enriched in dLGN | 1.724 | 0.00007828 | 35.5 |
| ENSMUSG00000000884 | guanine nucleotide binding protein (G protein), beta polypeptide 1-like [Source:MGI Symbol;Acc:MGI:1338057] | Gnb1l | Enriched in dLGN | 1.719 | 0.03056 | 294.8 |
| ENSMUSG00000041115 | IQ motif and Sec7 domain 2 [Source:MGI Symbol;Acc:MGI:3528396] | Iqsec2 | Enriched in dLGN | 1.714 | 5.86E-21 | 368.7 |
| ENSMUSG00000028370 | pregnancy-associated plasma protein A [Source:MGI Symbol;Acc:MGI:97479] | Pappa | Enriched in dLGN | 1.713 | 0.00001651 | 69.09 |
| ENSMUSG00000027220 | synaptotagmin XIII [Source:MGI Symbol;Acc:MGI:1933945] | Syt13 | Enriched in dLGN | 1.71 | 4.919E-18 | 763.5 |
| ENSMUSG00000048100 | TATA-box binding protein associated factor 13 [Source:MGI Symbol;Acc:MGI:1913500] | Taf13 | Enriched in dLGN | 1.709 | 0.000003477 | 38.39 |
| ENSMUSG00000004902 | solute carrier family 25 (mitochondrial carrier), member 18 [Source:MGI Symbol;Acc:MGI:1919053] | Slc25a18 | Enriched in dLGN | 1.705 | 0.0001171 | 68.62 |
| ENSMUSG00000028033 | potassium voltage-gated channel, subfamily Q, member 5 [Source:MGI Symbol;Acc:MGI:1924937] | Kcnq5 | Enriched in dLGN | 1.704 | 0.00007438 | 180.8 |
| ENSMUSG00000039838 | solute carrier family 45, member 1 [Source:MGI Symbol;Acc:MGI:2653235] | Slc45a1 | Enriched in dLGN | 1.7 | 3.647E-21 | 187.4 |
| ENSMUSG00000038555 | receptor accessory protein 2 [Source:MGI Symbol;Acc:MGI:2385070] | Reep2 | Enriched in dLGN | 1.7 | 2.618E-14 | 302.7 |
| ENSMUSG00000063882 | ubiquinol-cytochrome c reductase hinge protein [Source:MGI Symbol;Acc:MGI:1913826] | Uqcrh | Enriched in dLGN | 1.698 | 3.483E-12 | 256.5 |
| ENSMUSG00000045624 | ESF1 nucleolar pre-rRNA processing protein homolog [Source:MGI Symbol;Acc:MGI:1913830] | Esf1 | Enriched in dLGN | 1.697 | 6.954E-10 | 125.5 |
| ENSMUSG00000016319 | solute carrier family 25 (mitochondrial carrier, adenine nucleotide translocator), member 5 [Source:MGI Symbol;Acc:MGI:1353496] | Slc25a5 | Enriched in dLGN | 1.688 | 0.00003895 | 198.8 |
| ENSMUSG00000025940 | transmembrane protein 70 [Source:MGI Symbol;Acc:MGI:1915068] | Tmem70 | Enriched in dLGN | 1.686 | 6.214E-11 | 50.14 |
| ENSMUSG00000033793 | ATPase, H+ transporting, lysosomal V1 subunit H [Source:MGI Symbol;Acc:MGI:1914864] | Atp6v1h | Enriched in dLGN | 1.684 | 0.000000109 | 178.8 |
| ENSMUSG00000019158 | transmembrane protein 160 [Source:MGI Symbol;Acc:MGI:1916344] | Tmem160 | Enriched in dLGN | 1.684 | 0.0002414 | 90.04 |
| ENSMUSG00000060279 | adaptor-related protein complex 2, alpha 1 subunit [Source:MGI Symbol;Acc:MGI:101921] | Ap2a1 | Enriched in dLGN | 1.682 | 6.311E-13 | 671.9 |
| ENSMUSG00000021737 | proteasome (prosome, macropain) 26S subunit, non-ATPase, 6 [Source:MGI Symbol;Acc:MGI:1913663] | Psmd6 | Enriched in dLGN | 1.681 | 2.434E-09 | 147.1 |
| ENSMUSG00000038174 | hyccin PI4KA lipid kinase complex subunit 2 [Source:MGI Symbol;Acc:MGI:1098784] | Hycc2 | Enriched in dLGN | 1.681 | 0.00002079 | 246 |
| ENSMUSG00000001089 | leucine zipper protein 1 [Source:MGI Symbol;Acc:MGI:107629] | Luzp1 | Enriched in dLGN | 1.679 | 1.143E-08 | 485.1 |
| ENSMUSG00000004789 | dihydrolipoamide S-succinyltransferase (E2 component of 2-oxo-glutarate complex) [Source:MGI Symbol;Acc:MGI:1926170] | Dlst | Enriched in dLGN | 1.678 | 7.701E-21 | 409.2 |
| ENSMUSG00000039431 | myotubularin related protein 7 [Source:MGI Symbol;Acc:MGI:1891693] | Mtmr7 | Enriched in dLGN | 1.673 | 6.813E-12 | 199.1 |
| ENSMUSG00000052738 | succinate-CoA ligase, GDP-forming, alpha subunit [Source:MGI Symbol;Acc:MGI:1927234] | Suclg1 | Enriched in dLGN | 1.673 | 0.000001432 | 138.6 |
| ENSMUSG00000006731 | beta-1,4-N-acetyl-galactosaminyl transferase 1 [Source:MGI Symbol;Acc:MGI:1342057] | B4galnt1 | Enriched in dLGN | 1.67 | 2.159E-12 | 230.2 |
| ENSMUSG00000003380 | Rab acceptor 1 (prenylated) [Source:MGI Symbol;Acc:MGI:1201692] | Rabac1 | Enriched in dLGN | 1.669 | 1.244E-13 | 110.7 |
| ENSMUSG00000033208 | S100 protein, beta polypeptide, neural [Source:MGI Symbol;Acc:MGI:98217] | S100b | Enriched in dLGN | 1.668 | 0.0001845 | 86.39 |
| ENSMUSG00000026430 | Ras association (RalGDS/AF-6) domain family member 5 [Source:MGI Symbol;Acc:MGI:1926375] | Rassf5 | Enriched in dLGN | 1.667 | 3.408E-11 | 93.47 |
| ENSMUSG00000029777 | glycyl-tRNA synthetase [Source:MGI Symbol;Acc:MGI:2449057] | Gars | Enriched in dLGN | 1.666 | 1.712E-13 | 453.4 |
| ENSMUSG00000051223 | basic leucine zipper and W2 domains 1 [Source:MGI Symbol;Acc:MGI:1914132] | Bzw1 | Enriched in dLGN | 1.666 | 9.504E-09 | 393.6 |
| ENSMUSG00000031841 | cadherin 13 [Source:MGI Symbol;Acc:MGI:99551] | Cdh13 | Enriched in dLGN | 1.666 | 0.0000413 | 448.3 |
| ENSMUSG00000024862 | kinesin light chain 2 [Source:MGI Symbol;Acc:MGI:107953] | Klc2 | Enriched in dLGN | 1.664 | 1.948E-15 | 682.5 |
| ENSMUSG00000021024 | proteasome subunit alpha 6 [Source:MGI Symbol;Acc:MGI:1347006] | Psma6 | Enriched in dLGN | 1.663 | 0.0004272 | 71.31 |
| ENSMUSG00000035049 | ribosomal RNA processing 12 homolog [Source:MGI Symbol;Acc:MGI:2147437] | Rrp12 | Enriched in dLGN | 1.658 | 9.352E-12 | 124.1 |
| ENSMUSG00000019990 | phosphodiesterase 7B [Source:MGI Symbol;Acc:MGI:1352752] | Pde7b | Enriched in dLGN | 1.658 | 0.000005635 | 194.2 |
| ENSMUSG00000018326 | tyrosine 3-monooxygenase/tryptophan 5-monooxygenase activation protein, beta polypeptide [Source:MGI Symbol;Acc:MGI:1891917] | Ywhab | Enriched in dLGN | 1.657 | 1.335E-09 | 949.4 |
| ENSMUSG00000091264 | small integral membrane protein 13 [Source:MGI Symbol;Acc:MGI:2652854] | Smim13 | Enriched in dLGN | 1.656 | 4.884E-09 | 346 |
| ENSMUSG00000026858 | mitoguardin 2 [Source:MGI Symbol;Acc:MGI:1922035] | Miga2 | Enriched in dLGN | 1.652 | 1.143E-17 | 319.8 |
| ENSMUSG00000020340 | cytoplasmic FMR1 interacting protein 2 [Source:MGI Symbol;Acc:MGI:1924134] | Cyfip2 | Enriched in dLGN | 1.652 | 1.823E-12 | 1444 |
| ENSMUSG00000024190 | dual specificity phosphatase 1 [Source:MGI Symbol;Acc:MGI:105120] | Dusp1 | Enriched in dLGN | 1.65 | 8.334E-12 | 44.73 |
| ENSMUSG00000000901 | matrix metallopeptidase 11 [Source:MGI Symbol;Acc:MGI:97008] | Mmp11 | Enriched in dLGN | 1.649 | 0.001256 | 8.781 |
| ENSMUSG00000024966 | stress-induced phosphoprotein 1 [Source:MGI Symbol;Acc:MGI:109130] | Stip1 | Enriched in dLGN | 1.644 | 0.000001774 | 255 |
| ENSMUSG00000039601 | regulator of calcineurin 2 [Source:MGI Symbol;Acc:MGI:1858219] | Rcan2 | Enriched in dLGN | 1.641 | 1.009E-08 | 311.5 |
| ENSMUSG00000062070 | phosphoglycerate kinase 1 [Source:MGI Symbol;Acc:MGI:97555] | Pgk1 | Enriched in dLGN | 1.639 | 1.561E-07 | 300 |
| ENSMUSG00000022024 | SGT1, suppressor of G2 allele of SKP1 (S. cerevisiae) [Source:MGI Symbol;Acc:MGI:1915205] | Sugt1 | Enriched in dLGN | 1.635 | 4.039E-15 | 199.6 |
| ENSMUSG00000016541 | ataxin 10 [Source:MGI Symbol;Acc:MGI:1859293] | Atxn10 | Enriched in dLGN | 1.634 | 0.000003277 | 415.1 |
| ENSMUSG00000020349 | protein phosphatase 2 (formerly 2A), catalytic subunit, alpha isoform [Source:MGI Symbol;Acc:MGI:1321159] | Ppp2ca | Enriched in dLGN | 1.631 | 1.39E-10 | 828.3 |
| ENSMUSG00000030751 | proteasome subunit alpha 1 [Source:MGI Symbol;Acc:MGI:1347005] | Psma1 | Enriched in dLGN | 1.631 | 0.00001083 | 72.84 |
| ENSMUSG00000057766 | ankyrin repeat domain 29 [Source:MGI Symbol;Acc:MGI:2687055] | Ankrd29 | Enriched in dLGN | 1.627 | 6.515E-07 | 58.3 |
| ENSMUSG00000032330 | cytochrome c oxidase subunit 7A2 [Source:MGI Symbol;Acc:MGI:1316715] | Cox7a2 | Enriched in dLGN | 1.626 | 7.431E-09 | 76.71 |
| ENSMUSG00000020089 | pyrophosphatase (inorganic) 1 [Source:MGI Symbol;Acc:MGI:97831] | Ppa1 | Enriched in dLGN | 1.624 | 0.0001375 | 125.6 |
| ENSMUSG00000039480 | 5'-nucleotidase domain containing 1 [Source:MGI Symbol;Acc:MGI:2442446] | Nt5dc1 | Enriched in dLGN | 1.624 | 0.0003935 | 84.48 |
| ENSMUSG00000055022 | contactin 1 [Source:MGI Symbol;Acc:MGI:105980] | Cntn1 | Enriched in dLGN | 1.62 | 1.251E-09 | 1753 |
| ENSMUSG00000020953 | cochlin [Source:MGI Symbol;Acc:MGI:1278313] | Coch | Enriched in dLGN | 1.618 | 0.001227 | 80.94 |
| ENSMUSG00000019849 | prolyl endopeptidase [Source:MGI Symbol;Acc:MGI:1270863] | Prep | Enriched in dLGN | 1.614 | 1.261E-08 | 207.5 |
| ENSMUSG00000034171 | fatty acid amide hydrolase [Source:MGI Symbol;Acc:MGI:109609] | Faah | Enriched in dLGN | 1.613 | 1.611E-17 | 134 |
| ENSMUSG00000030683 | seizure related 6 homolog like 2 [Source:MGI Symbol;Acc:MGI:2385295] | Sez6l2 | Enriched in dLGN | 1.61 | 1.036E-09 | 601.7 |
| ENSMUSG00000020267 | histidine triad nucleotide binding protein 1 [Source:MGI Symbol;Acc:MGI:1321133] | Hint1 | Enriched in dLGN | 1.608 | 4.858E-09 | 255.7 |
| ENSMUSG00000036934 | RIKEN cDNA 4921524J17 gene [Source:MGI Symbol;Acc:MGI:1913964] | 4921524J17Rik | Enriched in dLGN | 1.606 | 0.000007057 | 42.54 |
| ENSMUSG00000019872 | sphingomyelin phosphodiesterase, acid-like 3A [Source:MGI Symbol;Acc:MGI:1931437] | Smpdl3a | Enriched in dLGN | 1.606 | 0.004014 | 14.04 |
| ENSMUSG00000053329 | glutamine amidotransferase like class 1 domain containing 3A [Source:MGI Symbol;Acc:MGI:1351861] | Gatd3a | Enriched in dLGN | 1.605 | 6.847E-13 | 113.5 |
| ENSMUSG00000031633 | solute carrier family 25 (mitochondrial carrier, adenine nucleotide translocator), member 4 [Source:MGI Symbol;Acc:MGI:1353495] | Slc25a4 | Enriched in dLGN | 1.591 | 3.154E-07 | 1489 |
| ENSMUSG00000027206 | COP9 signalosome subunit 2 [Source:MGI Symbol;Acc:MGI:1330276] | Cops2 | Enriched in dLGN | 1.585 | 1.173E-12 | 139.2 |
| ENSMUSG00000006717 | acyl-CoA thioesterase 13 [Source:MGI Symbol;Acc:MGI:1914084] | Acot13 | Enriched in dLGN | 1.585 | 0.00001081 | 30.71 |
| ENSMUSG00000028692 | aldo-keto reductase family 1, member A1 (aldehyde reductase) [Source:MGI Symbol;Acc:MGI:1929955] | Akr1a1 | Enriched in dLGN | 1.584 | 4.476E-15 | 579.7 |
| ENSMUSG00000024425 | Nedd4 family interacting protein 1 [Source:MGI Symbol;Acc:MGI:1929601] | Ndfip1 | Enriched in dLGN | 1.581 | 4.988E-09 | 747.3 |
| ENSMUSG00000070287 | solute carrier family 35, member G2 [Source:MGI Symbol;Acc:MGI:2685365] | Slc35g2 | Enriched in dLGN | 1.581 | 0.000001638 | 38.17 |
| ENSMUSG00000025487 | proteasome (prosome, macropain) 26S subunit, non-ATPase, 13 [Source:MGI Symbol;Acc:MGI:1345192] | Psmd13 | Enriched in dLGN | 1.577 | 4.75E-14 | 179.8 |
| ENSMUSG00000021947 | crystallin, lambda 1 [Source:MGI Symbol;Acc:MGI:1915881] | Cryl1 | Enriched in dLGN | 1.576 | 0.000009406 | 28.77 |
| ENSMUSG00000033900 | microtubule-associated protein 9 [Source:MGI Symbol;Acc:MGI:2442208] | Map9 | Enriched in dLGN | 1.575 | 2.862E-13 | 524.3 |
| ENSMUSG00000033998 | potassium channel, subfamily K, member 1 [Source:MGI Symbol;Acc:MGI:109322] | Kcnk1 | Enriched in dLGN | 1.575 | 2.223E-07 | 107.5 |
| ENSMUSG00000005312 | ubiquilin 1 [Source:MGI Symbol;Acc:MGI:1860276] | Ubqln1 | Enriched in dLGN | 1.574 | 4.753E-16 | 342.6 |
| ENSMUSG00000002332 | dehydrogenase/reductase (SDR family) member 1 [Source:MGI Symbol;Acc:MGI:1196314] | Dhrs1 | Enriched in dLGN | 1.572 | 0.0001627 | 49.92 |
| ENSMUSG00000020664 | dihydrolipoamide dehydrogenase [Source:MGI Symbol;Acc:MGI:107450] | Dld | Enriched in dLGN | 1.571 | 4.348E-07 | 258.4 |
| ENSMUSG00000015932 | destrin [Source:MGI Symbol;Acc:MGI:1929270] | Dstn | Enriched in dLGN | 1.571 | 0.000001719 | 300.9 |
| ENSMUSG00000063904 | dipeptidylpeptidase 3 [Source:MGI Symbol;Acc:MGI:1922471] | Dpp3 | Enriched in dLGN | 1.568 | 5.936E-12 | 154.1 |
| ENSMUSG00000030103 | basic helix-loop-helix family, member e40 [Source:MGI Symbol;Acc:MGI:1097714] | Bhlhe40 | Enriched in dLGN | 1.567 | 0.000001047 | 95.28 |
| ENSMUSG00000061603 | A kinase (PRKA) anchor protein 6 [Source:MGI Symbol;Acc:MGI:3050566] | Akap6 | Enriched in dLGN | 1.565 | 4.727E-12 | 2172 |
| ENSMUSG00000027457 | syntaphilin [Source:MGI Symbol;Acc:MGI:2139270] | Snph | Enriched in dLGN | 1.564 | 2.084E-13 | 637.1 |
| ENSMUSG00000030557 | myocyte enhancer factor 2A [Source:MGI Symbol;Acc:MGI:99532] | Mef2a | Enriched in dLGN | 1.564 | 9.93E-11 | 532.8 |
| ENSMUSG00000021748 | pyruvate dehydrogenase (lipoamide) beta [Source:MGI Symbol;Acc:MGI:1915513] | Pdhb | Enriched in dLGN | 1.562 | 2.581E-09 | 160.8 |
| ENSMUSG00000021290 | ATP synthase membrane subunit 6.8PL [Source:MGI Symbol;Acc:MGI:1917507] | Atp5mpl | Enriched in dLGN | 1.561 | 0.000535 | 181.4 |
| ENSMUSG00000049493 | plastin 1 (I-isoform) [Source:MGI Symbol;Acc:MGI:104809] | Pls1 | Enriched in dLGN | 1.558 | 0.002471 | 15.52 |
| ENSMUSG00000042298 | tetratricopeptide repeat domain 19 [Source:MGI Symbol;Acc:MGI:1920045] | Ttc19 | Enriched in dLGN | 1.556 | 7.193E-09 | 462.6 |
| ENSMUSG00000006732 | methyltransferase like 1 [Source:MGI Symbol;Acc:MGI:1339986] | Mettl1 | Enriched in dLGN | 1.556 | 0.000004142 | 23.99 |
| ENSMUSG00000025477 | inositol polyphosphate-5-phosphatase A [Source:MGI Symbol;Acc:MGI:2686961] | Inpp5a | Enriched in dLGN | 1.555 | 2.966E-09 | 299.2 |
| ENSMUSG00000025781 | ATP synthase, H+ transporting, mitochondrial F1 complex, gamma polypeptide 1 [Source:MGI Symbol;Acc:MGI:1261437] | Atp5c1 | Enriched in dLGN | 1.554 | 5.901E-11 | 290 |
| ENSMUSG00000057614 | guanine nucleotide binding protein (G protein), alpha inhibiting 1 [Source:MGI Symbol;Acc:MGI:95771] | Gnai1 | Enriched in dLGN | 1.553 | 6.629E-07 | 448.7 |
| ENSMUSG00000029860 | zyxin [Source:MGI Symbol;Acc:MGI:103072] | Zyx | Enriched in dLGN | 1.549 | 9.671E-10 | 118.6 |
| ENSMUSG00000027406 | isocitrate dehydrogenase 3 (NAD+) beta [Source:MGI Symbol;Acc:MGI:2158650] | Idh3b | Enriched in dLGN | 1.549 | 6.138E-08 | 236.2 |
| ENSMUSG00000039989 | chromobox 4 [Source:MGI Symbol;Acc:MGI:1195985] | Cbx4 | Enriched in dLGN | 1.544 | 9.094E-20 | 266.9 |
| ENSMUSG00000055839 | elongin B [Source:MGI Symbol;Acc:MGI:1914923] | Elob | Enriched in dLGN | 1.544 | 5.485E-08 | 99.23 |
| ENSMUSG00000027602 | microtubule-associated protein 1 light chain 3 alpha [Source:MGI Symbol;Acc:MGI:1915661] | Map1lc3a | Enriched in dLGN | 1.543 | 2.135E-09 | 233.2 |
| ENSMUSG00000027184 | cell cycle associated protein 1 [Source:MGI Symbol;Acc:MGI:1858234] | Caprin1 | Enriched in dLGN | 1.541 | 6.93E-12 | 681.9 |
| ENSMUSG00000025762 | La ribonucleoprotein 1B [Source:MGI Symbol;Acc:MGI:1914604] | Larp1b | Enriched in dLGN | 1.541 | 8.888E-08 | 100 |
| ENSMUSG00000029066 | mitochondrial ribosomal protein L20 [Source:MGI Symbol;Acc:MGI:2137221] | Mrpl20 | Enriched in dLGN | 1.541 | 1.306E-07 | 48.19 |
| ENSMUSG00000021643 | small EDRK-rich factor 1 [Source:MGI Symbol;Acc:MGI:1337114] | Serf1 | Enriched in dLGN | 1.541 | 0.000006279 | 40.29 |
| ENSMUSG00000072964 | basic helix-loop-helix domain containing, class B9 [Source:MGI Symbol;Acc:MGI:1917487] | Bhlhb9 | Enriched in dLGN | 1.54 | 0.000001972 | 158.1 |
| ENSMUSG00000029998 | prenylcysteine oxidase 1 [Source:MGI Symbol;Acc:MGI:1914131] | Pcyox1 | Enriched in dLGN | 1.537 | 1.884E-07 | 268.2 |
| ENSMUSG00000060739 | NSA2 ribosome biogenesis homolog [Source:MGI Symbol;Acc:MGI:1913883] | Nsa2 | Enriched in dLGN | 1.537 | 2.111E-07 | 65.27 |
| ENSMUSG00000025410 | dynactin 2 [Source:MGI Symbol;Acc:MGI:107733] | Dctn2 | Enriched in dLGN | 1.534 | 1.938E-15 | 382.2 |
| ENSMUSG00000029247 | phosphoribosylaminoimidazole carboxylase, phosphoribosylaminoribosylaminoimidazole, succinocarboxamide synthetase [Source:MGI Symbol;Acc:MGI:1914304] | Paics | Enriched in dLGN | 1.533 | 0.00002094 | 114.1 |
| ENSMUSG00000020029 | nudix (nucleoside diphosphate linked moiety X)-type motif 4 [Source:MGI Symbol;Acc:MGI:1918457] | Nudt4 | Enriched in dLGN | 1.531 | 1.66E-08 | 752.5 |
| ENSMUSG00000062995 | islet cell autoantigen 1 [Source:MGI Symbol;Acc:MGI:96391] | Ica1 | Enriched in dLGN | 1.531 | 0.002258 | 217.1 |
| ENSMUSG00000027603 | gamma-glutamyltransferase 7 [Source:MGI Symbol;Acc:MGI:1913385] | Ggt7 | Enriched in dLGN | 1.53 | 7.486E-09 | 194.7 |
| ENSMUSG00000018589 | glycine receptor, alpha 2 subunit [Source:MGI Symbol;Acc:MGI:95748] | Glra2 | Enriched in dLGN | 1.527 | 0.0001778 | 84.2 |
| ENSMUSG00000056004 | endosome-lysosome associated apoptosis and autophagy regulator family member 2 [Source:MGI Symbol;Acc:MGI:2443264] | Elapor2 | Enriched in dLGN | 1.526 | 2.452E-07 | 229.7 |
| ENSMUSG00000034557 | zinc finger, FYVE domain containing 9 [Source:MGI Symbol;Acc:MGI:2652838] | Zfyve9 | Enriched in dLGN | 1.526 | 0.00003197 | 559.4 |

| | | | | | | |
|---|---|---|---|---|---|---|
| ENSMUSG00000042682 | selenoprotein K [Source:MGI Symbol;Acc:MGI:1931466] | Selenok | Enriched in dLGN | 1.522 | 0.00004212 | 254.3 |
| ENSMUSG00000033460 | armadillo repeat containing, X-linked 1 [Source:MGI Symbol;Acc:MGI:1925498] | Armcx1 | Enriched in dLGN | 1.521 | 7.69E-11 | 200.9 |
| ENSMUSG00000096847 | transmembrane protein 151B [Source:MGI Symbol;Acc:MGI:2685169] | Tmem151b | Enriched in dLGN | 1.52 | 8.995E-09 | 1096 |
| ENSMUSG00000026833 | olfactomedin 1 [Source:MGI Symbol;Acc:MGI:1860437] | Olfm1 | Enriched in dLGN | 1.519 | 7.631E-09 | 1863 |
| ENSMUSG00000016346 | potassium voltage-gated channel, subfamily Q, member 2 [Source:MGI Symbol;Acc:MGI:1309503] | Kcnq2 | Enriched in dLGN | 1.515 | 6.092E-11 | 1096 |
| ENSMUSG00000020869 | leucine rich repeat containing 59 [Source:MGI Symbol;Acc:MGI:2138133] | Lrrc59 | Enriched in dLGN | 1.514 | 1.165E-11 | 321.9 |
| ENSMUSG00000055805 | formin-like 1 [Source:MGI Symbol;Acc:MGI:1888994] | Fmnl1 | Enriched in dLGN | 1.511 | 1.86E-12 | 238.1 |
| ENSMUSG00000044221 | G-rich RNA sequence binding factor 1 [Source:MGI Symbol;Acc:MGI:106479] | Grsf1 | Enriched in dLGN | 1.51 | 8.628E-12 | 242.3 |
| ENSMUSG00000028528 | DnaJ heat shock protein family (Hsp40) member C6 [Source:MGI Symbol;Acc:MGI:1919935] | Dnajc6 | Enriched in dLGN | 1.508 | 1.765E-13 | 781.5 |
| ENSMUSG00000020635 | FK506 binding protein 1b [Source:MGI Symbol;Acc:MGI:1336205] | Fkbp1b | Enriched in dLGN | 1.507 | 0.0001196 | 66.86 |
| ENSMUSG00000028998 | translocase of outer mitochondrial membrane 7 [Source:MGI Symbol;Acc:MGI:1913419] | Tomm7 | Enriched in dLGN | 1.504 | 0.000004934 | 121.4 |
| ENSMUSG00000079598 | C-type lectin domain family 2, member L [Source:MGI Symbol;Acc:MGI:2141402] | Clec2l | Enriched in dLGN | 1.501 | 0.000001869 | 252.6 |
| ENSMUSG00000033294 | NOC4 like [Source:MGI Symbol;Acc:MGI:2140843] | Noc4l | Enriched in dLGN | 1.497 | 0.000008701 | 33.71 |
| ENSMUSG00000028041 | a disintegrin and metallopeptidase domain 15 (metargidin) [Source:MGI Symbol;Acc:MGI:1333882] | Adam15 | Enriched in dLGN | 1.496 | 5.522E-13 | 177.5 |
| ENSMUSG00000025613 | chaperonin containing Tcp1, subunit 8 (theta) [Source:MGI Symbol;Acc:MGI:107183] | Cct8 | Enriched in dLGN | 1.496 | 0.000001269 | 259.2 |
| ENSMUSG00000021037 | AHA1, activator of heat shock protein ATPase 1 [Source:MGI Symbol;Acc:MGI:2387603] | Ahsa1 | Enriched in dLGN | 1.492 | 6.248E-15 | 199 |
| ENSMUSG00000049760 | mitochondrial contact site and cristae organizing system subunit 13 [Source:MGI Symbol;Acc:MGI:2442174] | Micos13 | Enriched in dLGN | 1.492 | 1.001E-08 | 44.94 |
| ENSMUSG00000051851 | retrotransposon Gag like 8C [Source:MGI Symbol;Acc:MGI:1920115] | Rtl8c | Enriched in dLGN | 1.491 | 0.00008434 | 53.92 |
| ENSMUSG00000020496 | ring finger protein 187 [Source:MGI Symbol;Acc:MGI:1914224] | Rnf187 | Enriched in dLGN | 1.489 | 4.616E-11 | 1303 |
| ENSMUSG00000040455 | ubiquitin specific peptidase 45 [Source:MGI Symbol;Acc:MGI:101850] | Usp45 | Enriched in dLGN | 1.489 | 2.694E-07 | 101.7 |
| ENSMUSG00000026204 | protein tyrosine phosphatase, receptor type, N [Source:MGI Symbol;Acc:MGI:102765] | Ptprn | Enriched in dLGN | 1.489 | 0.000008465 | 802.8 |
| ENSMUSG00000023033 | sodium channel, voltage-gated, type VIII, alpha [Source:MGI Symbol;Acc:MGI:103169] | Scn8a | Enriched in dLGN | 1.487 | 7.655E-13 | 1192 |
| ENSMUSG00000028811 | tyrosyl-tRNA synthetase [Source:MGI Symbol;Acc:MGI:2147627] | Yars | Enriched in dLGN | 1.486 | 2.913E-29 | 315 |
| ENSMUSG00000028790 | KH domain containing, RNA binding, signal transduction associated 1 [Source:MGI Symbol;Acc:MGI:893579] | Khdrbs1 | Enriched in dLGN | 1.486 | 2.579E-11 | 341 |
| ENSMUSG00000001056 | NHP2 ribonucleoprotein [Source:MGI Symbol;Acc:MGI:1098547] | Nhp2 | Enriched in dLGN | 1.484 | 9.282E-07 | 27.46 |
| ENSMUSG00000017167 | contactin associated protein-like 1 [Source:MGI Symbol;Acc:MGI:1858201] | Cntnap1 | Enriched in dLGN | 1.483 | 0.00004982 | 622.8 |
| ENSMUSG00000022765 | synaptosomal-associated protein 29 [Source:MGI Symbol;Acc:MGI:1914724] | Snap29 | Enriched in dLGN | 1.482 | 4.962E-09 | 76.62 |
| ENSMUSG00000022337 | ER membrane protein complex subunit 2 [Source:MGI Symbol;Acc:MGI:1913986] | Emc2 | Enriched in dLGN | 1.482 | 1.082E-08 | 104.6 |
| ENSMUSG00000063873 | solute carrier family 24 (sodium/potassium/calcium exchanger), member 3 [Source:MGI Symbol;Acc:MGI:2137513] | Slc24a3 | Enriched in dLGN | 1.479 | 9.357E-08 | 1240 |
| ENSMUSG00000068749 | proteasome subunit alpha 5 [Source:MGI Symbol;Acc:MGI:1347009] | Psma5 | Enriched in dLGN | 1.479 | 0.00006705 | 53.28 |
| ENSMUSG00000014769 | proteasome (prosome, macropain) subunit, beta type 1 [Source:MGI Symbol;Acc:MGI:104884] | Psmb1 | Enriched in dLGN | 1.473 | 3.039E-08 | 166 |
| ENSMUSG00000041453 | ribosomal protein L21 [Source:MGI Symbol;Acc:MGI:1278340] | Rpl21 | Enriched in dLGN | 1.472 | 0.00001053 | 129.9 |
| ENSMUSG00000027282 | mitochondrial carrier 2 [Source:MGI Symbol;Acc:MGI:1929260] | Mtch2 | Enriched in dLGN | 1.471 | 8.581E-10 | 103.4 |
| ENSMUSG00000044628 | ring finger protein 208 [Source:MGI Symbol;Acc:MGI:1916096] | Rnf208 | Enriched in dLGN | 1.471 | 1.548E-09 | 128.2 |
| ENSMUSG00000039347 | ATPase, H+ transporting, lysosomal V0 subunit E2 [Source:MGI Symbol;Acc:MGI:1923502] | Atp6v0e2 | Enriched in dLGN | 1.47 | 2.45E-11 | 554.6 |
| ENSMUSG00000018882 | mitochondrial ribosomal protein L45 [Source:MGI Symbol;Acc:MGI:1914286] | Mrpl45 | Enriched in dLGN | 1.466 | 4.438E-09 | 64.61 |
| ENSMUSG00000022890 | ATP synthase, H+ transporting, mitochondrial F0 complex, subunit F [Source:MGI Symbol;Acc:MGI:107777] | Atp5j | Enriched in dLGN | 1.466 | 0.000003919 | 249.2 |
| ENSMUSG00000025777 | ganglioside-induced differentiation-associated-protein 1 [Source:MGI Symbol;Acc:MGI:1338002] | Gdap1 | Enriched in dLGN | 1.465 | 7.469E-11 | 211.7 |
| ENSMUSG00000003812 | deoxyribonuclease II alpha [Source:MGI Symbol;Acc:MGI:1329019] | Dnase2a | Enriched in dLGN | 1.464 | 0.00005262 | 35.72 |
| ENSMUSG00000034173 | coiled-coil-helix-coiled-coil-helix domain containing 2-like [Source:MGI Symbol;Acc:MGI:1919220] | Chchd2l | Enriched in dLGN | 1.464 | 0.003842 | 15.6 |
| ENSMUSG00000029403 | cyclin-dependent kinase-like 2 (CDC2-related kinase) [Source:MGI Symbol;Acc:MGI:1858227] | Cdkl2 | Enriched in dLGN | 1.461 | 8.615E-12 | 131.1 |
| ENSMUSG00000034343 | ubiquitin-conjugating enzyme E2F (putative) [Source:MGI Symbol;Acc:MGI:1915171] | Ube2f | Enriched in dLGN | 1.461 | 3.863E-11 | 95.96 |
| ENSMUSG00000024064 | polypeptide N-acetylgalactosaminyltransferase 14 [Source:MGI Symbol;Acc:MGI:1918935] | Galnt14 | Enriched in dLGN | 1.458 | 0.00007962 | 190 |
| ENSMUSG00000009013 | dynein light chain LC8-type 1 [Source:MGI Symbol;Acc:MGI:1861457] | Dynll1 | Enriched in dLGN | 1.457 | 1.769E-10 | 315.9 |
| ENSMUSG00000029649 | proteasome maturation protein [Source:MGI Symbol;Acc:MGI:1913787] | Pomp | Enriched in dLGN | 1.455 | 3.088E-07 | 109 |
| ENSMUSG00000045967 | G protein-coupled receptor 158 [Source:MGI Symbol;Acc:MGI:2441697] | Gpr158 | Enriched in dLGN | 1.454 | 0.000004949 | 404.3 |
| ENSMUSG00000032435 | dynein cytoplasmic 1 light intermediate chain 1 [Source:MGI Symbol;Acc:MGI:2135610] | Dync1li1 | Enriched in dLGN | 1.453 | 3.406E-26 | 559.8 |
| ENSMUSG00000046442 | protein phosphatase 1E (PP2C domain containing) [Source:MGI Symbol;Acc:MGI:2444096] | Ppm1e | Enriched in dLGN | 1.452 | 5.668E-09 | 844.2 |
| ENSMUSG00000031561 | teneurin transmembrane protein 3 [Source:MGI Symbol;Acc:MGI:1345183] | Tenm3 | Enriched in dLGN | 1.45 | 3.874E-10 | 2385 |
| ENSMUSG00000024099 | NADH:ubiquinone oxidoreductase core subunit V2 [Source:MGI Symbol;Acc:MGI:1920150] | Ndufv2 | Enriched in dLGN | 1.446 | 9.623E-11 | 217.5 |
| ENSMUSG00000041762 | G protein-coupled receptor 155 [Source:MGI Symbol;Acc:MGI:1915776] | Gpr155 | Enriched in dLGN | 1.446 | 3.798E-08 | 55.46 |
| ENSMUSG00000024130 | ATP-binding cassette, sub-family A (ABC1), member 3 [Source:MGI Symbol;Acc:MGI:1351617] | Abca3 | Enriched in dLGN | 1.445 | 2.525E-16 | 605.6 |
| ENSMUSG00000020849 | tyrosine 3-monooxygenase/tryptophan 5-monooxygenase activation protein, epsilon polypeptide [Source:MGI Symbol;Acc:MGI:894689] | Ywhae | Enriched in dLGN | 1.445 | 4.303E-09 | 2150 |
| ENSMUSG00000021917 | signal peptidase complex subunit 1 homolog (S. cerevisiae) [Source:MGI Symbol;Acc:MGI:1916269] | Spcs1 | Enriched in dLGN | 1.44 | 0.00002233 | 46.82 |
| ENSMUSG00000038335 | TSR1 20S rRNA accumulation [Source:MGI Symbol;Acc:MGI:2144566] | Tsr1 | Enriched in dLGN | 1.435 | 1.132E-17 | 103.4 |
| ENSMUSG00000036510 | cadherin 8 [Source:MGI Symbol;Acc:MGI:107434] | Cdh8 | Enriched in dLGN | 1.433 | 0.0002918 | 272 |
| ENSMUSG00000038845 | prohibitin [Source:MGI Symbol;Acc:MGI:97572] | Phb | Enriched in dLGN | 1.432 | 4.689E-08 | 71.59 |
| ENSMUSG00000035776 | CD99 antigen-like 2 [Source:MGI Symbol;Acc:MGI:2177151] | Cd99l2 | Enriched in dLGN | 1.428 | 2.231E-11 | 280.3 |
| ENSMUSG00000062190 | LanC (bacterial lantibiotic synthetase component C)-like 2 [Source:MGI Symbol;Acc:MGI:1919085] | Lancl2 | Enriched in dLGN | 1.428 | 8.306E-10 | 173 |
| ENSMUSG00000020629 | acireductone dioxygenase 1 [Source:MGI Symbol;Acc:MGI:2144929] | Adi1 | Enriched in dLGN | 1.428 | 0.000000312 | 28.33 |
| ENSMUSG00000014077 | calcineurin-like EF hand protein 1 [Source:MGI Symbol;Acc:MGI:1927185] | Chp1 | Enriched in dLGN | 1.426 | 1.421E-08 | 152.7 |
| ENSMUSG00000039943 | phospholipase C, beta 4 [Source:MGI Symbol;Acc:MGI:107464] | Plcb4 | Enriched in dLGN | 1.426 | 9.557E-07 | 926.4 |
| ENSMUSG00000026496 | poly (ADP-ribose) polymerase family, member 1 [Source:MGI Symbol;Acc:MGI:1340806] | Parp1 | Enriched in dLGN | 1.415 | 2.878E-12 | 378.8 |
| ENSMUSG00000038690 | ATP synthase, H+ transporting, mitochondrial F0 complex, subunit F2 [Source:MGI Symbol;Acc:MGI:1927558] | Atp5j2 | Enriched in dLGN | 1.414 | 2.635E-07 | 138.3 |
| ENSMUSG00000029185 | family with sequence similarity 114, member A1 [Source:MGI Symbol;Acc:MGI:1915553] | Fam114a1 | Enriched in dLGN | 1.412 | 0.0003386 | 29.67 |
| ENSMUSG00000028373 | astrotactin 2 [Source:MGI Symbol;Acc:MGI:1889277] | Astn2 | Enriched in dLGN | 1.41 | 0.000817 | 621.4 |
| ENSMUSG00000029001 | F-box protein 44 [Source:MGI Symbol;Acc:MGI:1354744] | Fbxo44 | Enriched in dLGN | 1.409 | 1.928E-09 | 104.3 |
| ENSMUSG00000034880 | mitochondrial ribosomal protein L34 [Source:MGI Symbol;Acc:MGI:2137227] | Mrpl34 | Enriched in dLGN | 1.409 | 0.000003522 | 34.19 |
| ENSMUSG00000032462 | phosphatidylinositol-4,5-bisphosphate 3-kinase catalytic subunit beta [Source:MGI Symbol;Acc:MGI:1922019] | Pik3cb | Enriched in dLGN | 1.404 | 1.721E-11 | 99.32 |
| ENSMUSG00000025240 | SAC1 suppressor of actin mutations 1-like (yeast) [Source:MGI Symbol;Acc:MGI:1933169] | Sacm1l | Enriched in dLGN | 1.402 | 0.00001075 | 163.5 |
| ENSMUSG00000031068 | glutaredoxin 3 [Source:MGI Symbol;Acc:MGI:1353653] | Glrx3 | Enriched in dLGN | 1.401 | 0.000001442 | 113.9 |
| ENSMUSG00000024403 | ATPase, H+ transporting, lysosomal V1 subunit G2 [Source:MGI Symbol;Acc:MGI:1913487] | Atp6v1g2 | Enriched in dLGN | 1.399 | 6.574E-08 | 338.6 |
| ENSMUSG00000058355 | ATP-binding cassette, sub-family E (OABP), member 1 [Source:MGI Symbol;Acc:MGI:1195458] | Abce1 | Enriched in dLGN | 1.397 | 0.00001175 | 169.9 |
| ENSMUSG00000045948 | mitochondrial ribosomal protein S12 [Source:MGI Symbol;Acc:MGI:1346333] | Mrps12 | Enriched in dLGN | 1.396 | 0.00002254 | 42.54 |
| ENSMUSG00000039983 | coiled-coil domain containing 32 [Source:MGI Symbol;Acc:MGI:2685477] | Ccdc32 | Enriched in dLGN | 1.393 | 4.584E-07 | 49.02 |
| ENSMUSG00000021250 | FBJ osteosarcoma oncogene [Source:MGI Symbol;Acc:MGI:95574] | Fos | Enriched in dLGN | 1.393 | 0.00004187 | 57.19 |
| ENSMUSG00000022270 | reticulophagy regulator 1 [Source:MGI Symbol;Acc:MGI:1913520] | Retreg1 | Enriched in dLGN | 1.39 | 0.000003371 | 162.5 |
| ENSMUSG00000027002 | NCK-associated protein 1 [Source:MGI Symbol;Acc:MGI:1355333] | Nckap1 | Enriched in dLGN | 1.389 | 1.307E-09 | 682.6 |
| ENSMUSG00000071014 | NADH:ubiquinone oxidoreductase subunit B6 [Source:MGI Symbol;Acc:MGI:2684983] | Ndufb6 | Enriched in dLGN | 1.389 | 0.000047 | 50.89 |
| ENSMUSG00000063063 | catenin (cadherin associated protein), alpha 2 [Source:MGI Symbol;Acc:MGI:88275] | Ctnna2 | Enriched in dLGN | 1.388 | 0.002574 | 1024 |
| ENSMUSG00000028719 | cytidine monophosphate (UMP-CMP) kinase 1 [Source:MGI Symbol;Acc:MGI:1913838] | Cmpk1 | Enriched in dLGN | 1.387 | 1.187E-07 | 226.9 |
| ENSMUSG00000020326 | cyclin G1 [Source:MGI Symbol;Acc:MGI:102890] | Ccng1 | Enriched in dLGN | 1.386 | 0.000001836 | 214.5 |
| ENSMUSG00000031422 | mortality factor 4 like 2 [Source:MGI Symbol;Acc:MGI:1927167] | Morf4l2 | Enriched in dLGN | 1.385 | 4.928E-15 | 607.4 |
| ENSMUSG00000040035 | dispatched RND transporter family member 2 [Source:MGI Symbol;Acc:MGI:2388733] | Disp2 | Enriched in dLGN | 1.384 | 0.000003275 | 1280 |
| ENSMUSG00000027254 | microtubule-associated protein 1 A [Source:MGI Symbol;Acc:MGI:1306776] | Map1a | Enriched in dLGN | 1.384 | 0.0002074 | 6967 |
| ENSMUSG00000041763 | tripeptidyl peptidase II [Source:MGI Symbol;Acc:MGI:102724] | Tpp2 | Enriched in dLGN | 1.383 | 2.441E-08 | 165.9 |
| ENSMUSG00000029430 | RAN, member RAS oncogene family [Source:MGI Symbol;Acc:MGI:1333112] | Ran | Enriched in dLGN | 1.383 | 0.000004423 | 230.2 |
| ENSMUSG00000023106 | density-regulated protein [Source:MGI Symbol;Acc:MGI:1915434] | Denr | Enriched in dLGN | 1.381 | 2.243E-12 | 114.8 |
| ENSMUSG00000018770 | ATP synthase, H+ transporting, mitochondrial F0 complex, subunit C3 (subunit 9) [Source:MGI Symbol;Acc:MGI:2442035] | Atp5g3 | Enriched in dLGN | 1.38 | 1.686E-07 | 265.3 |
| ENSMUSG00000022099 | dematin actin binding protein [Source:MGI Symbol;Acc:MGI:99670] | Dmtn | Enriched in dLGN | 1.378 | 1.319E-10 | 452.3 |
| ENSMUSG00000021771 | voltage-dependent anion channel 2 [Source:MGI Symbol;Acc:MGI:106915] | Vdac2 | Enriched in dLGN | 1.377 | 0.0003705 | 243.1 |
| ENSMUSG00000025208 | mitochondrial ribosomal protein L43 [Source:MGI Symbol;Acc:MGI:2137229] | Mrpl43 | Enriched in dLGN | 1.374 | 7.886E-08 | 70.84 |
| ENSMUSG00000020309 | ChaC, cation transport regulator 2 [Source:MGI Symbol;Acc:MGI:1915294] | Chac2 | Enriched in dLGN | 1.374 | 0.0007185 | 18.28 |
| ENSMUSG00000031950 | gamma-aminobutyric acid (GABA) A receptor-associated protein-like 2 [Source:MGI Symbol;Acc:MGI:1890602] | Gabarapl2 | Enriched in dLGN | 1.373 | 6.201E-08 | 147.2 |
| ENSMUSG00000032101 | DEAD box helicase 25 [Source:MGI Symbol;Acc:MGI:1353582] | Ddx25 | Enriched in dLGN | 1.373 | 0.000002206 | 127.8 |
| ENSMUSG00000022295 | ATPase, H+ transporting, lysosomal V1 subunit C1 [Source:MGI Symbol;Acc:MGI:1913585] | Atp6v1c1 | Enriched in dLGN | 1.37 | 2.821E-08 | 229 |
| ENSMUSG00000030061 | ubiquitin-like modifier activating enzyme 3 [Source:MGI Symbol;Acc:MGI:1341217] | Uba3 | Enriched in dLGN | 1.368 | 0.000002671 | 78.39 |
| ENSMUSG00000002102 | proteasome (prosome, macropain) 26S subunit, ATPase 3 [Source:MGI Symbol;Acc:MGI:1098754] | Psmc3 | Enriched in dLGN | 1.366 | 5.388E-07 | 304.4 |
| ENSMUSG00000003949 | hepatic leukemia factor [Source:MGI Symbol;Acc:MGI:96108] | Hlf | Enriched in dLGN | 1.361 | 0.0000364 | 354.9 |
| ENSMUSG00000004285 | ATPase, H+ transporting, lysosomal V1 subunit F [Source:MGI Symbol;Acc:MGI:1913394] | Atp6v1f | Enriched in dLGN | 1.36 | 0.0008956 | 78.37 |
| ENSMUSG00000038486 | synaptic vesicle glycoprotein 2 a [Source:MGI Symbol;Acc:MGI:1927139] | Sv2a | Enriched in dLGN | 1.359 | 1.148E-08 | 1691 |
| ENSMUSG00000068099 | small integral membrane protein 45 [Source:MGI Symbol;Acc:MGI:1923755] | Smim45 | Enriched in dLGN | 1.356 | 1.008E-07 | 83.12 |
| ENSMUSG00000026377 | nucleolar protein interacting with the FHA domain of MKI67 [Source:MGI Symbol;Acc:MGI:1915199] | Nifk | Enriched in dLGN | 1.356 | 8.463E-07 | 61.02 |
| ENSMUSG00000028426 | RAD23 homolog B, nucleotide excision repair protein [Source:MGI Symbol;Acc:MGI:105128] | Rad23b | Enriched in dLGN | 1.354 | 1.765E-13 | 394.2 |
| ENSMUSG00000071528 | ATP synthase membrane subunit DAPIT [Source:MGI Symbol;Acc:MGI:1891435] | Atp5md | Enriched in dLGN | 1.354 | 0.001065 | 71.32 |
| ENSMUSG00000027827 | potassium voltage-gated channel, shaker-related subfamily, beta member 1 [Source:MGI Symbol;Acc:MGI:109155] | Kcnab1 | Enriched in dLGN | 1.353 | 0.000602 | 138 |
| ENSMUSG00000025066 | SWI5 dependent recombination repair 1 [Source:MGI Symbol;Acc:MGI:1915038] | Sfr1 | Enriched in dLGN | 1.352 | 6.563E-11 | 217.3 |
| ENSMUSG00000025899 | alkB homolog 8, tRNA methyltransferase [Source:MGI Symbol;Acc:MGI:1914917] | Alkbh8 | Enriched in dLGN | 1.35 | 2.75E-11 | 153.7 |
| ENSMUSG00000026860 | SH3-domain GRB2-like endophilin B2 [Source:MGI Symbol;Acc:MGI:2385131] | Sh3glb2 | Enriched in dLGN | 1.349 | 2.378E-08 | 236.6 |
| ENSMUSG00000036667 | TRPM8 channel-associated factor 1 [Source:MGI Symbol;Acc:MGI:1914665] | Tcaf1 | Enriched in dLGN | 1.349 | 0.000028 | 849.1 |
| ENSMUSG00000022965 | interferon gamma receptor 2 [Source:MGI Symbol;Acc:MGI:107654] | Ifngr2 | Enriched in dLGN | 1.348 | 0.000001061 | 119.8 |
| ENSMUSG00000001891 | UDP-glucose pyrophosphorylase 2 [Source:MGI Symbol;Acc:MGI:2183447] | Ugp2 | Enriched in dLGN | 1.348 | 0.00001606 | 62.12 |
| ENSMUSG00000028416 | BCL2-associated athanogene 1 [Source:MGI Symbol;Acc:MGI:108047] | Bag1 | Enriched in dLGN | 1.347 | 6.01E-08 | 349.9 |
| ENSMUSG00000030614 | transmembrane protein 126B [Source:MGI Symbol;Acc:MGI:1915722] | Tmem126b | Enriched in dLGN | 1.346 | 0.000006896 | 86.83 |
| ENSMUSG00000017778 | cytochrome c oxidase subunit 7C [Source:MGI Symbol;Acc:MGI:103226] | Cox7c | Enriched in dLGN | 1.345 | 0.00002639 | 35.38 |
| ENSMUSG00000024044 | erythrocyte membrane protein band 4.1 like 3 [Source:MGI Symbol;Acc:MGI:103008] | Epb41l3 | Enriched in dLGN | 1.343 | 0.000006797 | 1430 |
| ENSMUSG00000113902 | NADH:ubiquinone oxidoreductase subunit B1 [Source:MGI Symbol;Acc:MGI:3780865] | Ndufb1 | Enriched in dLGN | 1.342 | 0.0003754 | 43.14 |
| ENSMUSG00000059343 | aldolase 1 A, retrogene 1 [Source:MGI Symbol;Acc:MGI:2447811] | Aldoart1 | Enriched in dLGN | 1.342 | 0.002149 | 19.45 |
| ENSMUSG00000045427 | heterogeneous nuclear ribonucleoprotein H2 [Source:MGI Symbol;Acc:MGI:1201779] | Hnrnph2 | Enriched in dLGN | 1.341 | 0.00000364 | 111 |
| ENSMUSG00000036298 | solute carrier family 2 (facilitated glucose transporter), member 13 [Source:MGI Symbol;Acc:MGI:2146030] | Slc2a13 | Enriched in dLGN | 1.341 | 0.00009627 | 258.3 |
| ENSMUSG00000038530 | regulator of G-protein signaling 4 [Source:MGI Symbol;Acc:MGI:108409] | Rgs4 | Enriched in dLGN | 1.341 | 0.002678 | 578.5 |
| ENSMUSG00000018848 | arginyl-tRNA synthetase [Source:MGI Symbol;Acc:MGI:1914297] | Rars | Enriched in dLGN | 1.339 | 3.514E-14 | 102.9 |
| ENSMUSG00000028161 | protein phosphatase 3, catalytic subunit, alpha isoform [Source:MGI Symbol;Acc:MGI:107164] | Ppp3ca | Enriched in dLGN | 1.339 | 8.048E-12 | 1200 |
| ENSMUSG00000020102 | solute carrier family 16 (monocarboxylic acid transporters), member 7 [Source:MGI Symbol;Acc:MGI:1330284] | Slc16a7 | Enriched in dLGN | 1.337 | 0.0002486 | 145.4 |
| ENSMUSG00000036398 | protein phosphatase 1, regulatory inhibitor subunit 11 [Source:MGI Symbol;Acc:MGI:1923747] | Ppp1r11 | Enriched in dLGN | 1.336 | 1.269E-07 | 110.1 |
| ENSMUSG00000044894 | ubiquinol-cytochrome c reductase, complex III subunit VII [Source:MGI Symbol;Acc:MGI:107807] | Uqcrq | Enriched in dLGN | 1.334 | 0.00001005 | 236.5 |
| ENSMUSG00000056486 | chimerin 1 [Source:MGI Symbol;Acc:MGI:1915674] | Chn1 | Enriched in dLGN | 1.33 | 1.067E-14 | 504.4 |
| ENSMUSG00000030612 | mitochondrial ribosomal protein L46 [Source:MGI Symbol;Acc:MGI:1914558] | Mrpl46 | Enriched in dLGN | 1.329 | 0.00001145 | 33.73 |

| | | | | | | |
|---|---|---|---|---|---|---|
| ENSMUSG00000071658 | guanine nucleotide binding protein (G protein), gamma 3 [Source:MGI Symbol;Acc:MGI:102704] | Gng3 | Enriched in dLGN | 1.326 | 0.000004555 | 413.1 |
| ENSMUSG00000030738 | eukaryotic translation initiation factor 3, subunit C [Source:MGI Symbol;Acc:MGI:1926966] | Eif3c | Enriched in dLGN | 1.324 | 0.000002978 | 682.4 |
| ENSMUSG00000033419 | synaptosomal-associated protein 91 [Source:MGI Symbol;Acc:MGI:109132] | Snap91 | Enriched in dLGN | 1.321 | 4.283E-11 | 939.7 |
| ENSMUSG00000079317 | trafficking protein particle complex 2 [Source:MGI Symbol;Acc:MGI:1913476] | Trappc2 | Enriched in dLGN | 1.321 | 0.0001987 | 15.83 |
| ENSMUSG00000008892 | voltage-dependent anion channel 3 [Source:MGI Symbol;Acc:MGI:106922] | Vdac3 | Enriched in dLGN | 1.321 | 0.0002857 | 103.5 |
| ENSMUSG00000014294 | NADH:ubiquinone oxidoreductase subunit A2 [Source:MGI Symbol;Acc:MGI:1343103] | Ndufa2 | Enriched in dLGN | 1.319 | 1.258E-07 | 115.8 |
| ENSMUSG00000040896 | potassium voltage-gated channel, Shal-related family, member 3 [Source:MGI Symbol;Acc:MGI:1928743] | Kcnd3 | Enriched in dLGN | 1.319 | 0.000004398 | 579.5 |
| ENSMUSG00000024581 | N-ethylmaleimide sensitive fusion protein attachment protein gamma [Source:MGI Symbol;Acc:MGI:104561] | Napg | Enriched in dLGN | 1.316 | 7.406E-13 | 325.8 |
| ENSMUSG00000041997 | tousled-like kinase 1 [Source:MGI Symbol;Acc:MGI:2441683] | Tlk1 | Enriched in dLGN | 1.316 | 1.699E-08 | 318.7 |
| ENSMUSG00000001942 | sialic acid acetylesterase [Source:MGI Symbol;Acc:MGI:104803] | Siae | Enriched in dLGN | 1.316 | 0.0008233 | 30.58 |
| ENSMUSG00000024208 | ubiquinol-cytochrome c reductase complex assembly factor 2 [Source:MGI Symbol;Acc:MGI:1914517] | Uqcc2 | Enriched in dLGN | 1.314 | 0.00002618 | 101.2 |
| ENSMUSG00000019861 | golgi associated PDZ and coiled-coil motif containing [Source:MGI Symbol;Acc:MGI:2149946] | Gopc | Enriched in dLGN | 1.313 | 6.888E-17 | 304.7 |
| ENSMUSG00000026490 | CDC42 binding protein kinase alpha [Source:MGI Symbol;Acc:MGI:2441841] | Cdc42bpa | Enriched in dLGN | 1.311 | 2.819E-11 | 1511 |
| ENSMUSG00000048967 | YjeF N-terminal domain containing 3 [Source:MGI Symbol;Acc:MGI:2681845] | Yjefn3 | Enriched in dLGN | 1.311 | 0.001125 | 14.2 |
| ENSMUSG00000025979 | MOB family member 4, phocein [Source:MGI Symbol;Acc:MGI:104899] | Mob4 | Enriched in dLGN | 1.31 | 0.00002406 | 70.17 |
| ENSMUSG00000020458 | reticulon 4 [Source:MGI Symbol;Acc:MGI:1915835] | Rtn4 | Enriched in dLGN | 1.31 | 0.00006913 | 1440 |
| ENSMUSG00000040048 | NADH:ubiquinone oxidoreductase subunit B10 [Source:MGI Symbol;Acc:MGI:1915592] | Ndufb10 | Enriched in dLGN | 1.309 | 4.442E-07 | 201.2 |
| ENSMUSG00000030127 | COP9 signalosome subunit 7A [Source:MGI Symbol;Acc:MGI:1349400] | Cops7a | Enriched in dLGN | 1.308 | 2.888E-08 | 167 |
| ENSMUSG00000036158 | prickle planar cell polarity protein 1 [Source:MGI Symbol;Acc:MGI:1916034] | Prickle1 | Enriched in dLGN | 1.308 | 9.017E-07 | 385.1 |
| ENSMUSG00000049303 | synaptotagmin XII [Source:MGI Symbol;Acc:MGI:2159601] | Syt12 | Enriched in dLGN | 1.308 | 0.0001105 | 74.94 |
| ENSMUSG00000034088 | high density lipoprotein (HDL) binding protein [Source:MGI Symbol;Acc:MGI:99256] | Hdlbp | Enriched in dLGN | 1.303 | 4.812E-10 | 727.2 |
| ENSMUSG00000015981 | serine/threonine kinase 32C [Source:MGI Symbol;Acc:MGI:2385336] | Stk32c | Enriched in dLGN | 1.303 | 3.12E-09 | 234.7 |
| ENSMUSG00000027223 | mitogen-activated protein kinase 8 interacting protein 1 [Source:MGI Symbol;Acc:MGI:1309464] | Mapk8ip1 | Enriched in dLGN | 1.302 | 1.533E-08 | 1583 |
| ENSMUSG00000058388 | putative homeodomain transcription factor 1 [Source:MGI Symbol;Acc:MGI:1332671] | Phtf1 | Enriched in dLGN | 1.3 | 3.904E-10 | 177.4 |
| ENSMUSG00000004798 | unc-51 like kinase 2 [Source:MGI Symbol;Acc:MGI:1352758] | Ulk2 | Enriched in dLGN | 1.299 | 2.617E-10 | 388.4 |
| ENSMUSG00000074656 | eukaryotic translation initiation factor 2, subunit 2 (beta) [Source:MGI Symbol;Acc:MGI:1914454] | Eif2s2 | Enriched in dLGN | 1.296 | 1.713E-07 | 150.9 |
| ENSMUSG00000022982 | superoxide dismutase 1, soluble [Source:MGI Symbol;Acc:MGI:98351] | Sod1 | Enriched in dLGN | 1.295 | 5.495E-07 | 224.4 |
| ENSMUSG00000054162 | sparc/osteonectin, cwcv and kazal-like domains proteoglycan 3 [Source:MGI Symbol;Acc:MGI:1920152] | Spock3 | Enriched in dLGN | 1.295 | 0.0002849 | 404.4 |
| ENSMUSG00000115987 | vacuolar protein sorting 28 [Source:MGI Symbol;Acc:MGI:1914164] | Vps28 | Enriched in dLGN | 1.294 | 0.0000132 | 105.7 |
| ENSMUSG00000021792 | peroxiredoxin like 2A [Source:MGI Symbol;Acc:MGI:1917814] | Prxl2a | Enriched in dLGN | 1.293 | 5.817E-11 | 278.9 |
| ENSMUSG00000020577 | tetraspanin 13 [Source:MGI Symbol;Acc:MGI:1913359] | Tspan13 | Enriched in dLGN | 1.293 | 2.148E-08 | 273.9 |
| ENSMUSG00000027236 | eukaryotic translation initiation factor 3, subunit J1 [Source:MGI Symbol;Acc:MGI:1925905] | Eif3j1 | Enriched in dLGN | 1.293 | 0.00001163 | 85.54 |
| ENSMUSG00000006315 | transmembrane protein 147 [Source:MGI Symbol;Acc:MGI:1915011] | Tmem147 | Enriched in dLGN | 1.291 | 0.00001174 | 46.25 |
| ENSMUSG00000022094 | solute carrier family 39 (zinc transporter), member 14 [Source:MGI Symbol;Acc:MGI:2384851] | Slc39a14 | Enriched in dLGN | 1.29 | 8.077E-09 | 73.63 |
| ENSMUSG00000010803 | gamma-aminobutyric acid (GABA) A receptor, subunit alpha 1 [Source:MGI Symbol;Acc:MGI:95613] | Gabra1 | Enriched in dLGN | 1.29 | 0.0006931 | 231 |
| ENSMUSG00000039275 | forkhead box K2 [Source:MGI Symbol;Acc:MGI:1916087] | Foxk2 | Enriched in dLGN | 1.288 | 5.168E-20 | 465.7 |
| ENSMUSG00000043461 | serine palmitoyltransferase, small subunit B [Source:MGI Symbol;Acc:MGI:1913433] | Sptssb | Enriched in dLGN | 1.286 | 0.02454 | 9.484 |
| ENSMUSG00000063480 | SNU13 homolog, small nuclear ribonucleoprotein (U4/U6.U5) [Source:MGI Symbol;Acc:MGI:893586] | Snu13 | Enriched in dLGN | 1.285 | 0.0005154 | 38.94 |
| ENSMUSG00000021816 | protein phosphatase 3, catalytic subunit, beta isoform [Source:MGI Symbol;Acc:MGI:107163] | Ppp3cb | Enriched in dLGN | 1.284 | 1.309E-14 | 533.5 |
| ENSMUSG00000009535 | RNA (guanine-7-) methyltransferase [Source:MGI Symbol;Acc:MGI:1915147] | Rnmt | Enriched in dLGN | 1.28 | 1.878E-10 | 138.4 |
| ENSMUSG00000027523 | GNAS (guanine nucleotide binding protein, alpha stimulating) complex locus [Source:MGI Symbol;Acc:MGI:95777] | Gnas | Enriched in dLGN | 1.28 | 1.104E-08 | 3503 |
| ENSMUSG00000027133 | NOP10 ribonucleoprotein [Source:MGI Symbol;Acc:MGI:1913431] | Nop10 | Enriched in dLGN | 1.279 | 0.0006074 | 25.83 |
| ENSMUSG00000048332 | lipoma HMGIC fusion partner [Source:MGI Symbol;Acc:MGI:1920048] | Lhfp | Enriched in dLGN | 1.279 | 0.002204 | 294.9 |
| ENSMUSG00000044442 | N-6 adenine-specific DNA methyltransferase 1 (putative) [Source:MGI Symbol;Acc:MGI:1915018] | N6amt1 | Enriched in dLGN | 1.276 | 6.061E-07 | 39.22 |
| ENSMUSG00000001383 | zinc finger, matrin type 2 [Source:MGI Symbol;Acc:MGI:1913742] | Zmat2 | Enriched in dLGN | 1.275 | 5.827E-09 | 273.6 |
| ENSMUSG00000017221 | proteasome (prosome, macropain) 26S subunit, non-ATPase, 3 [Source:MGI Symbol;Acc:MGI:98858] | Psmd3 | Enriched in dLGN | 1.273 | 2.56E-08 | 193.7 |
| ENSMUSG00000028249 | syndecan binding protein [Source:MGI Symbol;Acc:MGI:1337026] | Sdcbp | Enriched in dLGN | 1.273 | 0.00004085 | 76 |
| ENSMUSG00000029447 | chaperonin containing Tcp1, subunit 6a (zeta) [Source:MGI Symbol;Acc:MGI:107943] | Cct6a | Enriched in dLGN | 1.267 | 6.724E-07 | 356.4 |
| ENSMUSG00000039629 | striatin interacting protein 2 [Source:MGI Symbol;Acc:MGI:2444363] | Strip2 | Enriched in dLGN | 1.267 | 0.0005295 | 138 |
| ENSMUSG00000021116 | eukaryotic translation initiation factor 2, subunit 1 alpha [Source:MGI Symbol;Acc:MGI:95299] | Eif2s1 | Enriched in dLGN | 1.266 | 0.0001805 | 89.93 |
| ENSMUSG00000030731 | synaptotagmin III [Source:MGI Symbol;Acc:MGI:99665] | Syt3 | Enriched in dLGN | 1.265 | 2.091E-08 | 150.8 |
| ENSMUSG00000004929 | thimet oligopeptidase 1 [Source:MGI Symbol;Acc:MGI:1354165] | Thop1 | Enriched in dLGN | 1.265 | 3.796E-07 | 79.81 |
| ENSMUSG00000030327 | NECAP endocytosis associated 1 [Source:MGI Symbol;Acc:MGI:1914852] | Necap1 | Enriched in dLGN | 1.265 | 0.0001391 | 167.4 |
| ENSMUSG00000026188 | transmembrane protein 169 [Source:MGI Symbol;Acc:MGI:2442781] | Tmem169 | Enriched in dLGN | 1.264 | 7.25E-09 | 139.3 |
| ENSMUSG00000027340 | solute carrier family 23 (nucleobase transporters), member 2 [Source:MGI Symbol;Acc:MGI:1859682] | Slc23a2 | Enriched in dLGN | 1.263 | 5.848E-08 | 495.3 |
| ENSMUSG00000028318 | polymerase (RNA) I polypeptide E [Source:MGI Symbol;Acc:MGI:1929022] | Polr1e | Enriched in dLGN | 1.262 | 0.000006242 | 28.6 |
| ENSMUSG00000058400 | pyroglutamylated RFamide peptide receptor [Source:MGI Symbol;Acc:MGI:2677633] | Qrfpr | Enriched in dLGN | 1.262 | 0.0095 | 24.15 |
| ENSMUSG00000028062 | late endosomal/lysosomal adaptor, MAPK and MTOR activator 2 [Source:MGI Symbol;Acc:MGI:1932697] | Lamtor2 | Enriched in dLGN | 1.261 | 0.00002394 | 47.8 |
| ENSMUSG00000021282 | eukaryotic translation initiation factor 5 [Source:MGI Symbol;Acc:MGI:95309] | Eif5 | Enriched in dLGN | 1.26 | 7.319E-07 | 396.4 |
| ENSMUSG00000029610 | aminoacyl tRNA synthetase complex-interacting multifunctional protein 2 [Source:MGI Symbol;Acc:MGI:2385237] | Aimp2 | Enriched in dLGN | 1.26 | 0.000349 | 26.12 |
| ENSMUSG00000034024 | chaperonin containing Tcp1, subunit 2 (beta) [Source:MGI Symbol;Acc:MGI:107186] | Cct2 | Enriched in dLGN | 1.259 | 4.736E-09 | 313.7 |
| ENSMUSG00000020598 | neuronal cell adhesion molecule [Source:MGI Symbol;Acc:MGI:104750] | Nrcam | Enriched in dLGN | 1.258 | 4.103E-07 | 667.9 |
| ENSMUSG00000037152 | NADH:ubiquinone oxidoreductase subunit C1 [Source:MGI Symbol;Acc:MGI:1913627] | Ndufc1 | Enriched in dLGN | 1.258 | 0.00101 | 49.47 |
| ENSMUSG00000045348 | neuronal tyrosine-phosphorylated phosphoinositide 3-kinase adaptor 1 [Source:MGI Symbol;Acc:MGI:2443880] | Nyap1 | Enriched in dLGN | 1.257 | 1.432E-08 | 501.1 |
| ENSMUSG00000024955 | estrogen related receptor, alpha [Source:MGI Symbol;Acc:MGI:1346831] | Esrra | Enriched in dLGN | 1.257 | 1.187E-07 | 88.56 |
| ENSMUSG00000027248 | protein disulfide isomerase associated 3 [Source:MGI Symbol;Acc:MGI:95834] | Pdia3 | Enriched in dLGN | 1.257 | 0.00004244 | 401 |
| ENSMUSG00000025868 | HIG1 domain family, member 2A [Source:MGI Symbol;Acc:MGI:1914294] | Higd2a | Enriched in dLGN | 1.256 | 0.0002991 | 47.28 |
| ENSMUSG00000024527 | AFG3-like AAA ATPase 2 [Source:MGI Symbol;Acc:MGI:1916847] | Afg3l2 | Enriched in dLGN | 1.255 | 5.448E-13 | 309.3 |
| ENSMUSG00000022789 | dynamin 1-like [Source:MGI Symbol;Acc:MGI:1921256] | Dnm1l | Enriched in dLGN | 1.254 | 0.000003362 | 380.5 |
| ENSMUSG00000054034 | transcription elongation factor A (SII)-like 5 [Source:MGI Symbol;Acc:MGI:3036236] | Tceal5 | Enriched in dLGN | 1.253 | 0.004214 | 58.79 |
| ENSMUSG00000030417 | programmed cell death 5 [Source:MGI Symbol;Acc:MGI:1913538] | Pdcd5 | Enriched in dLGN | 1.251 | 0.00001586 | 70.53 |
| ENSMUSG00000022231 | sema domain, seven thrombospondin repeats (type 1 and type 1-like), transmembrane domain (TM) and short cytoplasmic domain, (semaphorin) 5A [Source:MGI Symbol;Acc:MGI:10 | Sema5a | Enriched in dLGN | 1.247 | 0.00003216 | 795.5 |
| ENSMUSG00000045613 | cholinergic receptor, muscarinic 2, cardiac [Source:MGI Symbol;Acc:MGI:88397] | Chrm2 | Enriched in dLGN | 1.247 | 0.001214 | 71.41 |
| ENSMUSG00000029036 | ATPase family, AAA domain containing 3A [Source:MGI Symbol;Acc:MGI:1919214] | Atad3a | Enriched in dLGN | 1.246 | 2.094E-08 | 104.5 |
| ENSMUSG00000030301 | coiled-coil domain containing 91 [Source:MGI Symbol;Acc:MGI:1914265] | Ccdc91 | Enriched in dLGN | 1.246 | 1.336E-07 | 147.9 |
| ENSMUSG00000048385 | scratch family zinc finger 1 [Source:MGI Symbol;Acc:MGI:2176606] | Scrt1 | Enriched in dLGN | 1.246 | 0.0000171 | 1666 |
| ENSMUSG00000025917 | COP9 signalosome subunit 5 [Source:MGI Symbol;Acc:MGI:1349415] | Cops5 | Enriched in dLGN | 1.246 | 0.000911 | 96.98 |
| ENSMUSG00000005580 | adenylate cyclase 9 [Source:MGI Symbol;Acc:MGI:108450] | Adcy9 | Enriched in dLGN | 1.245 | 7.319E-07 | 235.6 |
| ENSMUSG00000028782 | adhesion G protein-coupled receptor B2 [Source:MGI Symbol;Acc:MGI:2451244] | Adgrb2 | Enriched in dLGN | 1.244 | 1.163E-07 | 1121 |
| ENSMUSG00000020720 | proteasome (prosome, macropain) 26S subunit, non-ATPase, 12 [Source:MGI Symbol;Acc:MGI:1914247] | Psmd12 | Enriched in dLGN | 1.244 | 0.00002927 | 128.2 |
| ENSMUSG00000079184 | M-phase phosphoprotein 8 [Source:MGI Symbol;Acc:MGI:1922589] | Mphosph8 | Enriched in dLGN | 1.244 | 0.004311 | 685.9 |
| ENSMUSG00000020153 | NADH:ubiquinone oxidoreductase core subunit S7 [Source:MGI Symbol;Acc:MGI:1922656] | Ndufs7 | Enriched in dLGN | 1.242 | 0.00001431 | 124 |
| ENSMUSG00000060450 | ring finger protein 14 [Source:MGI Symbol;Acc:MGI:1929668] | Rnf14 | Enriched in dLGN | 1.24 | 2.381E-10 | 686.9 |
| ENSMUSG00000074797 | inosine triphosphatase (nucleoside triphosphate pyrophosphatase) [Source:MGI Symbol;Acc:MGI:96622] | Itpa | Enriched in dLGN | 1.239 | 0.00000204 | 70.22 |
| ENSMUSG00000028656 | CAP, adenylate cyclase-associated protein 1 (yeast) [Source:MGI Symbol;Acc:MGI:88262] | Cap1 | Enriched in dLGN | 1.239 | 0.0001886 | 243.2 |
| ENSMUSG00000038084 | OPA1, mitochondrial dynamin like GTPase [Source:MGI Symbol;Acc:MGI:1921393] | Opa1 | Enriched in dLGN | 1.237 | 0.00002608 | 499.1 |
| ENSMUSG00000028447 | dynactin 3 [Source:MGI Symbol;Acc:MGI:1859251] | Dctn3 | Enriched in dLGN | 1.235 | 1.512E-08 | 123.3 |
| ENSMUSG00000031202 | RAB39B, member RAS oncogene family [Source:MGI Symbol;Acc:MGI:1915040] | Rab39b | Enriched in dLGN | 1.235 | 0.0000766 | 57.43 |
| ENSMUSG00000039385 | cadherin 6 [Source:MGI Symbol;Acc:MGI:107435] | Cdh6 | Enriched in dLGN | 1.235 | 0.0006756 | 237 |
| ENSMUSG00000013736 | tRNA nucleotidyl transferase, CCA-adding, 1 [Source:MGI Symbol;Acc:MGI:1917297] | Trnt1 | Enriched in dLGN | 1.234 | 4.742E-07 | 70.42 |
| ENSMUSG00000020312 | SHC (Src homology 2 domain containing) transforming protein 2 [Source:MGI Symbol;Acc:MGI:106180] | Shc2 | Enriched in dLGN | 1.233 | 1.053E-08 | 242.8 |
| ENSMUSG00000023328 | acetylcholinesterase [Source:MGI Symbol;Acc:MGI:87876] | Ache | Enriched in dLGN | 1.232 | 0.0001549 | 333.5 |
| ENSMUSG00000028555 | tetratricopeptide repeat domain 39A [Source:MGI Symbol;Acc:MGI:2444350] | Ttc39a | Enriched in dLGN | 1.232 | 0.0008747 | 25.31 |
| ENSMUSG00000021548 | cyclin H [Source:MGI Symbol;Acc:MGI:1913921] | Ccnh | Enriched in dLGN | 1.231 | 0.0001253 | 69.07 |
| ENSMUSG00000026664 | phytanoyl-CoA hydroxylase [Source:MGI Symbol;Acc:MGI:891978] | Phyh | Enriched in dLGN | 1.23 | 0.000002246 | 93.1 |
| ENSMUSG00000068739 | seryl-aminoacyl-tRNA synthetase [Source:MGI Symbol;Acc:MGI:102809] | Sars | Enriched in dLGN | 1.229 | 2.226E-11 | 493.2 |
| ENSMUSG00000024583 | thioredoxin-like 1 [Source:MGI Symbol;Acc:MGI:1860078] | Txnl1 | Enriched in dLGN | 1.228 | 1.873E-08 | 112.7 |
| ENSMUSG00000042097 | zinc finger protein 239 [Source:MGI Symbol;Acc:MGI:1306812] | Zfp239 | Enriched in dLGN | 1.228 | 0.00002467 | 26.99 |
| ENSMUSG00000039768 | DnaJ heat shock protein family (Hsp40) member C11 [Source:MGI Symbol;Acc:MGI:2443386] | Dnajc11 | Enriched in dLGN | 1.226 | 2.51E-10 | 251.9 |
| ENSMUSG00000020818 | major facilitator superfamily domain containing 11 [Source:MGI Symbol;Acc:MGI:1917150] | Mfsd11 | Enriched in dLGN | 1.226 | 2.725E-07 | 44.36 |
| ENSMUSG00000021606 | NADH:ubiquinone oxidoreductase core subunit S6 [Source:MGI Symbol;Acc:MGI:107932] | Ndufs6 | Enriched in dLGN | 1.224 | 1.068E-07 | 70.27 |
| ENSMUSG00000027091 | zinc finger CCCH-type containing 15 [Source:MGI Symbol;Acc:MGI:1919747] | Zc3h15 | Enriched in dLGN | 1.223 | 0.000000141 | 331.8 |
| ENSMUSG00000084786 | ubiquitin-like 5 [Source:MGI Symbol;Acc:MGI:1913427] | Ubl5 | Enriched in dLGN | 1.223 | 0.000000242 | 89.51 |
| ENSMUSG00000026000 | LanC (bacterial lantibiotic synthetase component C)-like 1 [Source:MGI Symbol;Acc:MGI:1336997] | Lancl1 | Enriched in dLGN | 1.222 | 1.488E-09 | 119.7 |
| ENSMUSG00000021963 | Sin3-associated polypeptide 18 [Source:MGI Symbol;Acc:MGI:1277978] | Sap18 | Enriched in dLGN | 1.222 | 4.192E-08 | 76.48 |
| ENSMUSG00000040990 | SH3-domain kinase binding protein 1 [Source:MGI Symbol;Acc:MGI:1889583] | Sh3kbp1 | Enriched in dLGN | 1.22 | 8.187E-08 | 327.5 |
| ENSMUSG00000025439 | chloride channel, nucleotide-sensitive, 1A [Source:MGI Symbol;Acc:MGI:109638] | Clns1a | Enriched in dLGN | 1.22 | 5.529E-07 | 102.4 |
| ENSMUSG00000025366 | extended synaptotagmin-like protein 1 [Source:MGI Symbol;Acc:MGI:1344426] | Esyt1 | Enriched in dLGN | 1.219 | 0.004787 | 159.4 |
| ENSMUSG00000030591 | proteasome (prosome, macropain) 26S subunit, non-ATPase, 8 [Source:MGI Symbol;Acc:MGI:1888669] | Psmd8 | Enriched in dLGN | 1.217 | 5.279E-08 | 174.8 |
| ENSMUSG00000021218 | guanosine diphosphate (GDP) dissociation inhibitor 2 [Source:MGI Symbol;Acc:MGI:99845] | Gdi2 | Enriched in dLGN | 1.214 | 0.000001646 | 530.5 |
| ENSMUSG00000019494 | COP9 signalosome subunit 6 [Source:MGI Symbol;Acc:MGI:1349439] | Cops6 | Enriched in dLGN | 1.214 | 0.0005223 | 134 |
| ENSMUSG00000009630 | protein phosphatase 2 (formerly 2A), catalytic subunit, beta isoform [Source:MGI Symbol;Acc:MGI:1321161] | Ppp2cb | Enriched in dLGN | 1.212 | 3.408E-11 | 209.1 |
| ENSMUSG00000027502 | replication termination factor 2 [Source:MGI Symbol;Acc:MGI:1913654] | Rtf2 | Enriched in dLGN | 1.212 | 3.281E-10 | 124.2 |
| ENSMUSG00000028470 | histidine triad nucleotide binding protein 2 [Source:MGI Symbol;Acc:MGI:1916167] | Hint2 | Enriched in dLGN | 1.212 | 0.000856 | 22.68 |
| ENSMUSG00000062006 | ribosomal protein L34 [Source:MGI Symbol;Acc:MGI:1915686] | Rpl34 | Enriched in dLGN | 1.212 | 0.001916 | 135.1 |
| ENSMUSG00000041360 | pumilio RNA-binding family member 3 [Source:MGI Symbol;Acc:MGI:106253] | Pum3 | Enriched in dLGN | 1.211 | 5.657E-07 | 123.2 |
| ENSMUSG00000039100 | membrane associated ring-CH-type finger 6 [Source:MGI Symbol;Acc:MGI:2442773] | Marchf6 | Enriched in dLGN | 1.209 | 4.288E-12 | 751.7 |
| ENSMUSG00000006651 | amyloid beta (A4) precursor-like protein 1 [Source:MGI Symbol;Acc:MGI:88046] | Aplp1 | Enriched in dLGN | 1.209 | 0.000001073 | 2009 |
| ENSMUSG00000062270 | mortality factor 4 like 1 [Source:MGI Symbol;Acc:MGI:1096551] | Morf4l1 | Enriched in dLGN | 1.209 | 0.000008754 | 259.3 |
| ENSMUSG00000041645 | DEAD box helicase 24 [Source:MGI Symbol;Acc:MGI:1351337] | Ddx24 | Enriched in dLGN | 1.204 | 7.998E-09 | 586.1 |
| ENSMUSG00000029108 | protocadherin 7 [Source:MGI Symbol;Acc:MGI:1860487] | Pcdh7 | Enriched in dLGN | 1.204 | 0.00009132 | 1224 |
| ENSMUSG00000075700 | selenoprotein T [Source:MGI Symbol;Acc:MGI:1916477] | Selenot | Enriched in dLGN | 1.203 | 0.00000118 | 383.6 |
| ENSMUSG00000030007 | chaperonin containing Tcp1, subunit 7 (eta) [Source:MGI Symbol;Acc:MGI:107184] | Cct7 | Enriched in dLGN | 1.203 | 0.000003863 | 351.8 |
| ENSMUSG00000011752 | phosphoglycerate mutase 1 [Source:MGI Symbol;Acc:MGI:97552] | Pgam1 | Enriched in dLGN | 1.2 | 0.0002969 | 217.3 |
| ENSMUSG00000020817 | rabaptin, RAB GTPase binding effector protein 1 [Source:MGI Symbol;Acc:MGI:1860236] | Rabep1 | Enriched in dLGN | 1.195 | 0.00004066 | 777.4 |
| ENSMUSG00000042712 | transcription elongation factor A like 9 [Source:MGI Symbol;Acc:MGI:109567] | Tceal9 | Enriched in dLGN | 1.194 | 0.000006939 | 169.6 |
| ENSMUSG00000090841 | myosin, light polypeptide 6, alkali, smooth muscle and non-muscle [Source:MGI Symbol;Acc:MGI:109318] | Myl6 | Enriched in dLGN | 1.194 | 0.00003422 | 274.2 |
| ENSMUSG00000024387 | casein kinase 2, beta polypeptide [Source:MGI Symbol;Acc:MGI:88548] | Csnk2b | Enriched in dLGN | 1.19 | 0.002342 | 97.86 |

| | | | | | | |
|---|---|---|---|---|---|---|
| ENSMUSG00000029388 | eukaryotic translation initiation factor 2B, subunit 1 (alpha) [Source:MGI Symbol;Acc:MGI:2384802] | Eif2b1 | Enriched in dLGN | 1.189 | 0.00002967 | 40.06 |
| ENSMUSG00000062542 | synaptotagmin IX [Source:MGI Symbol;Acc:MGI:1926373] | Syt9 | Enriched in dLGN | 1.188 | 4.604E-08 | 283.3 |
| ENSMUSG00000042660 | WD repeat domain 55 [Source:MGI Symbol;Acc:MGI:1915186] | Wdr55 | Enriched in dLGN | 1.186 | 0.00003106 | 26.69 |
| ENSMUSG00000068039 | t-complex protein 1 [Source:MGI Symbol;Acc:MGI:98535] | Tcp1 | Enriched in dLGN | 1.185 | 0.00004479 | 455.1 |
| ENSMUSG00000061474 | mitochondrial ribosomal protein S36 [Source:MGI Symbol;Acc:MGI:1913378] | Mrps36 | Enriched in dLGN | 1.185 | 0.009834 | 23.77 |
| ENSMUSG00000021178 | protease (prosome, macropain) 26S subunit, ATPase 1 [Source:MGI Symbol;Acc:MGI:106054] | Psmc1 | Enriched in dLGN | 1.184 | 0.00005836 | 275.2 |
| ENSMUSG00000031533 | mitochondrial ribosomal protein S31 [Source:MGI Symbol;Acc:MGI:1913153] | Mrps31 | Enriched in dLGN | 1.183 | 0.00004687 | 35.82 |
| ENSMUSG00000061080 | limbic system-associated membrane protein [Source:MGI Symbol;Acc:MGI:1261760] | Lsamp | Enriched in dLGN | 1.183 | 0.001641 | 3253 |
| ENSMUSG00000030879 | mitochondrial ribosomal protein L17 [Source:MGI Symbol;Acc:MGI:1351608] | Mrpl17 | Enriched in dLGN | 1.18 | 4.249E-07 | 76.99 |
| ENSMUSG00000054843 | attractin like 1 [Source:MGI Symbol;Acc:MGI:2147749] | Atrnl1 | Enriched in dLGN | 1.18 | 0.000209 | 629.5 |
| ENSMUSG00000047649 | CD3E antigen, epsilon polypeptide associated protein [Source:MGI Symbol;Acc:MGI:1917583] | Cd3eap | Enriched in dLGN | 1.18 | 0.0003803 | 71.75 |
| ENSMUSG00000008763 | mannosidase, alpha, class 1A, member 2 [Source:MGI Symbol;Acc:MGI:104676] | Man1a2 | Enriched in dLGN | 1.177 | 1.809E-07 | 612.4 |
| ENSMUSG00000061313 | DDHD domain containing 2 [Source:MGI Symbol;Acc:MGI:1919358] | Ddhd2 | Enriched in dLGN | 1.177 | 7.635E-07 | 151.6 |
| ENSMUSG00000030105 | ADP-ribosylation factor-like 8B [Source:MGI Symbol;Acc:MGI:1914416] | Arl8b | Enriched in dLGN | 1.176 | 2.145E-11 | 268.8 |
| ENSMUSG00000004207 | prosaposin [Source:MGI Symbol;Acc:MGI:97783] | Psap | Enriched in dLGN | 1.176 | 0.000196 | 1320 |
| ENSMUSG00000050730 | Rho GTPase activating protein 42 [Source:MGI Symbol;Acc:MGI:1918794] | Arhgap42 | Enriched in dLGN | 1.176 | 0.0008179 | 71.18 |
| ENSMUSG00000032046 | abhydrolase domain containing 12 [Source:MGI Symbol;Acc:MGI:1923442] | Abhd12 | Enriched in dLGN | 1.174 | 1.216E-09 | 313.9 |
| ENSMUSG00000021752 | potassium channel tetramerisation domain containing 6 [Source:MGI Symbol;Acc:MGI:1918643] | Kctd6 | Enriched in dLGN | 1.174 | 0.00003109 | 62.84 |
| ENSMUSG00000025869 | NOP16 nucleolar protein [Source:MGI Symbol;Acc:MGI:107862] | Nop16 | Enriched in dLGN | 1.169 | 0.00002447 | 53.74 |
| ENSMUSG00000021194 | chromogranin A [Source:MGI Symbol;Acc:MGI:88394] | Chga | Enriched in dLGN | 1.169 | 0.001826 | 812.4 |
| ENSMUSG00000022201 | zinc finger RNA binding protein [Source:MGI Symbol;Acc:MGI:1341890] | Zfr | Enriched in dLGN | 1.165 | 4.831E-08 | 1105 |
| ENSMUSG00000027531 | inositol (myo)-1(or 4)-monophosphatase 1 [Source:MGI Symbol;Acc:MGI:1933158] | Impa1 | Enriched in dLGN | 1.162 | 9.531E-09 | 75.71 |
| ENSMUSG00000030946 | phospholysine phosphohistidine inorganic pyrophosphate phosphatase [Source:MGI Symbol;Acc:MGI:1923679] | Lhpp | Enriched in dLGN | 1.16 | 0.00007768 | 34.61 |
| ENSMUSG00000033845 | mitochondrial ribosomal protein L15 [Source:MGI Symbol;Acc:MGI:1351639] | Mrpl15 | Enriched in dLGN | 1.159 | 2.567E-07 | 86.2 |
| ENSMUSG00000003161 | sorcin [Source:MGI Symbol;Acc:MGI:98419] | Sri | Enriched in dLGN | 1.157 | 0.00000765 | 62.39 |
| ENSMUSG00000037149 | DEAD box helicase 1 [Source:MGI Symbol;Acc:MGI:2144727] | Ddx1 | Enriched in dLGN | 1.154 | 3.987E-07 | 410.1 |
| ENSMUSG00000035790 | centrosomal protein 19 [Source:MGI Symbol;Acc:MGI:1914244] | Cep19 | Enriched in dLGN | 1.153 | 1.956E-11 | 80.5 |
| ENSMUSG00000027203 | deoxyuridine triphosphatase [Source:MGI Symbol;Acc:MGI:1346051] | Dut | Enriched in dLGN | 1.15 | 5.379E-07 | 92.41 |
| ENSMUSG00000009863 | succinate dehydrogenase complex, subunit B, iron sulfur (Ip) [Source:MGI Symbol;Acc:MGI:1914930] | Sdhb | Enriched in dLGN | 1.15 | 0.00002244 | 134.3 |
| ENSMUSG00000031303 | mitogen-activated protein kinase kinase kinase 15 [Source:MGI Symbol;Acc:MGI:2448588] | Map3k15 | Enriched in dLGN | 1.149 | 0.02011 | 11.32 |
| ENSMUSG00000028577 | phospholipase A2, activating protein [Source:MGI Symbol;Acc:MGI:104810] | Plaa | Enriched in dLGN | 1.148 | 2.424E-08 | 186.2 |
| ENSMUSG00000015668 | PDZ domain containing 11 [Source:MGI Symbol;Acc:MGI:1919871] | Pdzd11 | Enriched in dLGN | 1.148 | 0.0003923 | 27.95 |
| ENSMUSG00000029613 | eukaryotic translation initiation factor 2 alpha kinase 1 [Source:MGI Symbol;Acc:MGI:1353448] | Eif2ak1 | Enriched in dLGN | 1.147 | 1.835E-08 | 145.1 |
| ENSMUSG00000020114 | cullin associated and neddylation disassociated 1 [Source:MGI Symbol;Acc:MGI:1261820] | Cand1 | Enriched in dLGN | 1.146 | 1.126E-08 | 459.1 |
| ENSMUSG00000019923 | ZW10 interactor [Source:MGI Symbol;Acc:MGI:1289227] | Zwint | Enriched in dLGN | 1.146 | 0.000008974 | 1513 |
| ENSMUSG00000005732 | RAN binding protein 1 [Source:MGI Symbol;Acc:MGI:96269] | Ranbp1 | Enriched in dLGN | 1.146 | 0.00004345 | 62.92 |
| ENSMUSG00000024194 | cutA divalent cation tolerance homolog [Source:MGI Symbol;Acc:MGI:1914925] | Cuta | Enriched in dLGN | 1.143 | 0.000003266 | 66.24 |
| ENSMUSG00000022956 | ATP synthase, H+ transporting, mitochondrial F1 complex, O subunit [Source:MGI Symbol;Acc:MGI:106341] | Atp5o | Enriched in dLGN | 1.142 | 0.0002604 | 36.29 |
| ENSMUSG00000030691 | FCH and double SH3 domains 2 [Source:MGI Symbol;Acc:MGI:2448475] | Fchsd2 | Enriched in dLGN | 1.141 | 4.654E-08 | 298.3 |
| ENSMUSG00000033436 | armadillo repeat containing, X-linked 2 [Source:MGI Symbol;Acc:MGI:1914666] | Armcx2 | Enriched in dLGN | 1.141 | 6.156E-07 | 262.7 |
| ENSMUSG00000010376 | neural precursor cell expressed, developmentally down-regulated gene 8 [Source:MGI Symbol;Acc:MGI:97301] | Nedd8 | Enriched in dLGN | 1.141 | 0.000072 | 191.2 |
| ENSMUSG00000071662 | polymerase (RNA) II (DNA directed) polypeptide G [Source:MGI Symbol;Acc:MGI:1914960] | Polr2g | Enriched in dLGN | 1.137 | 0.00001798 | 40.09 |
| ENSMUSG00000028138 | alcohol dehydrogenase 5 (class III), chi polypeptide [Source:MGI Symbol;Acc:MGI:87929] | Adh5 | Enriched in dLGN | 1.135 | 0.0001014 | 68.21 |
| ENSMUSG00000054942 | mitoguardin 1 [Source:MGI Symbol;Acc:MGI:1924567] | Miga1 | Enriched in dLGN | 1.134 | 7.934E-12 | 313 |
| ENSMUSG00000079562 | macrophage erythroblast attacher [Source:MGI Symbol;Acc:MGI:1891748] | Maea | Enriched in dLGN | 1.134 | 7.276E-11 | 195.6 |
| ENSMUSG00000055943 | ER membrane protein complex subunit 7 [Source:MGI Symbol;Acc:MGI:1920274] | Emc7 | Enriched in dLGN | 1.133 | 0.00004822 | 203.9 |
| ENSMUSG00000030452 | non imprinted in Prader-Willi/Angelman syndrome 2 homolog (human) [Source:MGI Symbol;Acc:MGI:1913918] | Nipa2 | Enriched in dLGN | 1.133 | 0.002424 | 70.36 |
| ENSMUSG00000002010 | isocitrate dehydrogenase 3 (NAD+), gamma [Source:MGI Symbol;Acc:MGI:1099463] | Idh3g | Enriched in dLGN | 1.132 | 0.009643 | 76.06 |
| ENSMUSG00000027346 | glycerophosphocholine phosphodiesterase 1 [Source:MGI Symbol;Acc:MGI:104898] | Gpcpd1 | Enriched in dLGN | 1.131 | 0.00001193 | 161 |
| ENSMUSG00000026245 | phenylalanyl-tRNA synthetase, beta subunit [Source:MGI Symbol;Acc:MGI:1346035] | Farsb | Enriched in dLGN | 1.128 | 0.000001132 | 92.51 |
| ENSMUSG00000040907 | ATPase, Na+/K+ transporting, alpha 3 polypeptide [Source:MGI Symbol;Acc:MGI:88107] | Atp1a3 | Enriched in dLGN | 1.127 | 0.000007964 | 7574 |
| ENSMUSG00000023932 | cell division cycle 5-like (S. pombe) [Source:MGI Symbol;Acc:MGI:1918952] | Cdc5l | Enriched in dLGN | 1.125 | 0.000001478 | 151.3 |
| ENSMUSG00000004642 | stem-loop binding protein [Source:MGI Symbol;Acc:MGI:108402] | Slbp | Enriched in dLGN | 1.124 | 0.00001644 | 85.78 |
| ENSMUSG00000005510 | NADH:ubiquinone oxidoreductase core subunit S3 [Source:MGI Symbol;Acc:MGI:1915599] | Ndufs3 | Enriched in dLGN | 1.123 | 0.000005899 | 101.1 |
| ENSMUSG00000038717 | ATP synthase, H+ transporting, mitochondrial F0 complex, subunit G [Source:MGI Symbol;Acc:MGI:1351597] | Atp5l | Enriched in dLGN | 1.123 | 0.001033 | 38.82 |
| ENSMUSG00000091955 | predicted pseudogene 9844 [Source:MGI Symbol;Acc:MGI:3704288] | Gm9844 | Enriched in dLGN | 1.123 | 0.002225 | 233.8 |
| ENSMUSG00000034285 | nipsnap homolog 1 [Source:MGI Symbol;Acc:MGI:1278344] | Nipsnap1 | Enriched in dLGN | 1.122 | 1.631E-07 | 81.27 |
| ENSMUSG00000024120 | leucine-rich PPR-motif containing [Source:MGI Symbol;Acc:MGI:1919666] | Lrpprc | Enriched in dLGN | 1.118 | 5.888E-09 | 294.4 |
| ENSMUSG00000036615 | regulatory factor X-associated protein [Source:MGI Symbol;Acc:MGI:2180854] | Rfxap | Enriched in dLGN | 1.117 | 1.586E-07 | 118.3 |
| ENSMUSG00000022536 | glyoxylate reductase 1 homolog (Arabidopsis) [Source:MGI Symbol;Acc:MGI:1921272] | Glyr1 | Enriched in dLGN | 1.116 | 1.294E-11 | 384.5 |
| ENSMUSG00000041236 | VPS41 HOPS complex subunit [Source:MGI Symbol;Acc:MGI:1929215] | Vps41 | Enriched in dLGN | 1.116 | 3.443E-09 | 365.7 |
| ENSMUSG00000038762 | ATP-binding cassette, sub-family F (GCN20), member 1 [Source:MGI Symbol;Acc:MGI:1351658] | Abcf1 | Enriched in dLGN | 1.115 | 2.963E-08 | 576.3 |
| ENSMUSG00000029559 | RIKEN cDNA 2210016L21 gene [Source:MGI Symbol;Acc:MGI:1919607] | 2210016L21Rik | Enriched in dLGN | 1.115 | 6.551E-07 | 211.9 |
| ENSMUSG00000016308 | ubiquitin-conjugating enzyme E2A [Source:MGI Symbol;Acc:MGI:102959] | Ube2a | Enriched in dLGN | 1.114 | 0.00000358 | 44.52 |
| ENSMUSG00000011256 | a disintegrin and metallopeptidase domain 19 (meltrin beta) [Source:MGI Symbol;Acc:MGI:105377] | Adam19 | Enriched in dLGN | 1.114 | 0.0002808 | 194.7 |
| ENSMUSG00000033124 | autophagy related 9A [Source:MGI Symbol;Acc:MGI:2138446] | Atg9a | Enriched in dLGN | 1.113 | 2.486E-13 | 274.4 |
| ENSMUSG00000036257 | patatin-like phospholipase domain containing 8 [Source:MGI Symbol;Acc:MGI:1914702] | Pnpla8 | Enriched in dLGN | 1.113 | 0.000003296 | 213.7 |
| ENSMUSG00000030335 | mitochondrial ribosomal protein L51 [Source:MGI Symbol;Acc:MGI:1913743] | Mrpl51 | Enriched in dLGN | 1.112 | 0.00002756 | 68.55 |
| ENSMUSG00000070858 | predicted gene 1673 [Source:MGI Symbol;Acc:MGI:2686519] | Gm1673 | Enriched in dLGN | 1.11 | 0.02824 | 55.45 |
| ENSMUSG00000000339 | RNA 3'-terminal phosphate cyclase [Source:MGI Symbol;Acc:MGI:1913618] | Rtca | Enriched in dLGN | 1.109 | 0.000001016 | 93.2 |
| ENSMUSG00000021660 | basic transcription factor 3 [Source:MGI Symbol;Acc:MGI:1202875] | Btf3 | Enriched in dLGN | 1.108 | 0.000007483 | 71.1 |
| ENSMUSG00000019087 | ATPase, H+ transporting, lysosomal accessory protein 1 [Source:MGI Symbol;Acc:MGI:109629] | Atp6ap1 | Enriched in dLGN | 1.106 | 0.00008136 | 276.8 |
| ENSMUSG00000056553 | protein tyrosine phosphatase, receptor type, N polypeptide 2 [Source:MGI Symbol;Acc:MGI:107418] | Ptprn2 | Enriched in dLGN | 1.1 | 0.0003283 | 1089 |
| ENSMUSG00000040136 | ATP-binding cassette, sub-family C (CFTR/MRP), member 8 [Source:MGI Symbol;Acc:MGI:1352629] | Abcc8 | Enriched in dLGN | 1.098 | 0.002652 | 86.74 |
| ENSMUSG00000033917 | glycerophosphodiester phosphodiesterase 1 [Source:MGI Symbol;Acc:MGI:1891827] | Gde1 | Enriched in dLGN | 1.097 | 3.088E-07 | 234.5 |
| ENSMUSG00000000171 | succinate dehydrogenase complex, subunit D, integral membrane protein [Source:MGI Symbol;Acc:MGI:1914175] | Sdhd | Enriched in dLGN | 1.096 | 7.044E-08 | 85.24 |
| ENSMUSG00000062997 | ribosomal protein L35 [Source:MGI Symbol;Acc:MGI:1913739] | Rpl35 | Enriched in dLGN | 1.096 | 0.0004997 | 91.03 |
| ENSMUSG00000016833 | mitochondrial ribosomal protein S18C [Source:MGI Symbol;Acc:MGI:1915985] | Mrps18c | Enriched in dLGN | 1.096 | 0.02532 | 8.671 |
| ENSMUSG00000032324 | tetraspanin 3 [Source:MGI Symbol;Acc:MGI:1928098] | Tspan3 | Enriched in dLGN | 1.095 | 0.00002959 | 417.4 |
| ENSMUSG00000031839 | heat shock factor binding protein 1 [Source:MGI Symbol;Acc:MGI:1915446] | Hsbp1 | Enriched in dLGN | 1.094 | 0.000006754 | 392.5 |
| ENSMUSG00000030663 | RIKEN cDNA 1110004F10 gene [Source:MGI Symbol;Acc:MGI:1929274] | 1110004F10Rik | Enriched in dLGN | 1.091 | 5.373E-07 | 195 |
| ENSMUSG00000030284 | cysteine-rich with EGF-like domains 1 [Source:MGI Symbol;Acc:MGI:2152539] | Creld1 | Enriched in dLGN | 1.091 | 0.000001001 | 242.9 |
| ENSMUSG00000031347 | centrin 2 [Source:MGI Symbol;Acc:MGI:1347085] | Cetn2 | Enriched in dLGN | 1.091 | 0.00001121 | 57.78 |
| ENSMUSG00000062352 | integrin beta 1 binding protein 1 [Source:MGI Symbol;Acc:MGI:1306802] | Itgb1bp1 | Enriched in dLGN | 1.087 | 0.00001356 | 59.33 |
| ENSMUSG00000028419 | charged multivesicular body protein 5 [Source:MGI Symbol;Acc:MGI:1924209] | Chmp5 | Enriched in dLGN | 1.086 | 0.00006915 | 102.8 |
| ENSMUSG00000023460 | RAB12, member RAS oncogene family [Source:MGI Symbol;Acc:MGI:894284] | Rab12 | Enriched in dLGN | 1.085 | 3.429E-10 | 311.7 |
| ENSMUSG00000015094 | neural proliferation, differentiation and control 1 [Source:MGI Symbol;Acc:MGI:1099802] | Npdc1 | Enriched in dLGN | 1.085 | 5.184E-08 | 346.6 |
| ENSMUSG00000032555 | topoisomerase (DNA) II binding protein 1 [Source:MGI Symbol;Acc:MGI:1920018] | Topbp1 | Enriched in dLGN | 1.083 | 1.719E-07 | 78.01 |
| ENSMUSG00000021285 | protein phosphatase 1, regulatory subunit 13B [Source:MGI Symbol;Acc:MGI:1336199] | Ppp1r13b | Enriched in dLGN | 1.08 | 8.59E-08 | 497.8 |
| ENSMUSG00000078249 | high mobility group AT-hook 1B [Source:MGI Symbol;Acc:MGI:96161] | Hmga1b | Enriched in dLGN | 1.08 | 0.00001495 | 54.25 |
| ENSMUSG00000027804 | peptidylprolyl isomerase D (cyclophilin D) [Source:MGI Symbol;Acc:MGI:1914988] | Ppid | Enriched in dLGN | 1.079 | 0.00009143 | 95.78 |
| ENSMUSG00000068882 | small RNA binding exonuclease protection factor La [Source:MGI Symbol;Acc:MGI:98423] | Ssb | Enriched in dLGN | 1.078 | 0.00008256 | 214.7 |
| ENSMUSG00000035864 | synaptotagmin I [Source:MGI Symbol;Acc:MGI:99667] | Syt1 | Enriched in dLGN | 1.076 | 0.00008003 | 3835 |
| ENSMUSG00000028478 | clathrin, light polypeptide (Lca) [Source:MGI Symbol;Acc:MGI:894297] | Clta | Enriched in dLGN | 1.074 | 3.463E-09 | 866.9 |
| ENSMUSG00000004933 | megakaryocyte-associated tyrosine kinase [Source:MGI Symbol;Acc:MGI:99259] | Matk | Enriched in dLGN | 1.071 | 0.00008953 | 175.5 |
| ENSMUSG00000026887 | mitochondrial ribosome recycling factor [Source:MGI Symbol;Acc:MGI:1915121] | Mrrf | Enriched in dLGN | 1.07 | 0.00003525 | 35.97 |
| ENSMUSG00000044627 | SWI5 recombination repair homolog (yeast) [Source:MGI Symbol;Acc:MGI:1920181] | Swi5 | Enriched in dLGN | 1.068 | 6.787E-08 | 130.5 |
| ENSMUSG00000062785 | potassium voltage gated channel, Shaw-related subfamily, member 3 [Source:MGI Symbol;Acc:MGI:96669] | Kcnc3 | Enriched in dLGN | 1.068 | 0.000002088 | 437.6 |
| ENSMUSG00000021877 | ADP-ribosylation factor 4 [Source:MGI Symbol;Acc:MGI:99433] | Arf4 | Enriched in dLGN | 1.067 | 0.000001093 | 151 |
| ENSMUSG00000031059 | NADH:ubiquinone oxidoreductase subunit B11 [Source:MGI Symbol;Acc:MGI:1349919] | Ndufb11 | Enriched in dLGN | 1.067 | 0.00001132 | 155.4 |
| ENSMUSG00000029641 | RAS-like, family 11, member A [Source:MGI Symbol;Acc:MGI:1916145] | Rasl11a | Enriched in dLGN | 1.067 | 0.02457 | 21.43 |
| ENSMUSG00000021518 | phosphatidylserine synthase 1 [Source:MGI Symbol;Acc:MGI:1276575] | Ptdss1 | Enriched in dLGN | 1.066 | 2.423E-09 | 227.3 |
| ENSMUSG00000031948 | lysyl-tRNA synthetase [Source:MGI Symbol;Acc:MGI:1934754] | Kars | Enriched in dLGN | 1.065 | 0.00006439 | 171 |
| ENSMUSG00000018470 | potassium voltage-gated channel, shaker-related subfamily, beta member 3 [Source:MGI Symbol;Acc:MGI:1336208] | Kcnab3 | Enriched in dLGN | 1.064 | 0.001668 | 53.86 |
| ENSMUSG00000001376 | VPS50 EARP/GARPII complex subunit [Source:MGI Symbol;Acc:MGI:1920538] | Vps50 | Enriched in dLGN | 1.06 | 0.00008493 | 171.6 |
| ENSMUSG00000026254 | eukaryotic translation initiation factor 4E member 2 [Source:MGI Symbol;Acc:MGI:1914440] | Eif4e2 | Enriched in dLGN | 1.059 | 9.434E-15 | 227.8 |
| ENSMUSG00000031774 | proteasome activator subunit 3 interacting protein 1 [Source:MGI Symbol;Acc:MGI:1919637] | Psme3ip1 | Enriched in dLGN | 1.059 | 9.592E-10 | 123.3 |
| ENSMUSG00000022884 | eukaryotic translation initiation factor 4A2 [Source:MGI Symbol;Acc:MGI:106906] | Eif4a2 | Enriched in dLGN | 1.059 | 6.914E-07 | 619.6 |
| ENSMUSG00000032112 | trafficking protein particle complex 4 [Source:MGI Symbol;Acc:MGI:1926211] | Trappc4 | Enriched in dLGN | 1.059 | 0.0001124 | 41.09 |
| ENSMUSG00000000787 | DEAD box helicase 3, X-linked [Source:MGI Symbol;Acc:MGI:103064] | Ddx3x | Enriched in dLGN | 1.058 | 0.00008574 | 602.8 |
| ENSMUSG00000022370 | mitochondrial ribosomal protein L13 [Source:MGI Symbol;Acc:MGI:2137218] | Mrpl13 | Enriched in dLGN | 1.058 | 0.0002836 | 50.25 |
| ENSMUSG00000038722 | BUD31 homolog [Source:MGI Symbol;Acc:MGI:2141291] | Bud31 | Enriched in dLGN | 1.055 | 9.907E-07 | 48.97 |
| ENSMUSG00000027305 | NADH:ubiquinone oxidoreductase complex assembly factor 1 [Source:MGI Symbol;Acc:MGI:1916952] | Ndufaf1 | Enriched in dLGN | 1.051 | 0.00001857 | 32.64 |
| ENSMUSG00000060904 | ADP-ribosylation factor-like 1 [Source:MGI Symbol;Acc:MGI:99436] | Arl1 | Enriched in dLGN | 1.051 | 0.0002673 | 131.5 |
| ENSMUSG00000024887 | N-acylsphingosine amidohydrolase 2 [Source:MGI Symbol;Acc:MGI:1859310] | Asah2 | Enriched in dLGN | 1.051 | 0.01311 | 39.79 |
| ENSMUSG00000023110 | protein arginine N-methyltransferase 5 [Source:MGI Symbol;Acc:MGI:1351645] | Prmt5 | Enriched in dLGN | 1.05 | 0.0000173 | 156.1 |
| ENSMUSG00000056596 | TMF1-regulated nuclear protein 1 [Source:MGI Symbol;Acc:MGI:1916789] | Trnp1 | Enriched in dLGN | 1.05 | 0.0006046 | 640.5 |
| ENSMUSG00000040536 | N-terminal EF-hand calcium binding protein 1 [Source:MGI Symbol;Acc:MGI:1916602] | Necab1 | Enriched in dLGN | 1.05 | 0.002576 | 133 |
| ENSMUSG00000014606 | solute carrier family 25 (mitochondrial carrier oxoglutarate carrier), member 11 [Source:MGI Symbol;Acc:MGI:1915113] | Slc25a11 | Enriched in dLGN | 1.049 | 4.83E-08 | 129.2 |
| ENSMUSG00000003199 | MPN domain containing [Source:MGI Symbol;Acc:MGI:1915297] | Mpnd | Enriched in dLGN | 1.049 | 0.000000649 | 241.7 |
| ENSMUSG00000033278 | protein tyrosine phosphatase, receptor type, M [Source:MGI Symbol;Acc:MGI:102694] | Ptprm | Enriched in dLGN | 1.049 | 0.002328 | 370.1 |
| ENSMUSG00000015092 | endothelial differentiation-related factor 1 [Source:MGI Symbol;Acc:MGI:1891227] | Edf1 | Enriched in dLGN | 1.048 | 0.00001306 | 190.3 |
| ENSMUSG00000021486 | PRELI domain containing 1 [Source:MGI Symbol;Acc:MGI:1913744] | Prelid1 | Enriched in dLGN | 1.045 | 0.00001239 | 247.5 |
| ENSMUSG00000031617 | transmembrane protein 184C [Source:MGI Symbol;Acc:MGI:2384562] | Tmem184c | Enriched in dLGN | 1.045 | 0.0001244 | 115.9 |
| ENSMUSG00000020526 | zinc finger, HIT type 3 [Source:MGI Symbol;Acc:MGI:3051596] | Znhit3 | Enriched in dLGN | 1.045 | 0.0003544 | 29.29 |
| ENSMUSG00000001569 | nucleolar protein with MIF4G domain 1 [Source:MGI Symbol;Acc:MGI:1861749] | Nom1 | Enriched in dLGN | 1.043 | 8.665E-09 | 142.3 |
| ENSMUSG00000027673 | NADH:ubiquinone oxidoreductase subunit B5 [Source:MGI Symbol;Acc:MGI:1913296] | Ndufb5 | Enriched in dLGN | 1.042 | 0.000001351 | 138.6 |
| ENSMUSG00000004070 | heme oxygenase 2 [Source:MGI Symbol;Acc:MGI:109373] | Hmox2 | Enriched in dLGN | 1.04 | 0.0004513 | 96.26 |

| | | | | | | |
|---|---|---|---|---|---|---|
| ENSMUSG00000032940 | RNA binding motif protein 11 [Source:MGI Symbol;Acc:MGI:2447622] | Rbm11 | Enriched in dLGN | 1.04 | 0.03748 | 9.326 |
| ENSMUSG00000021193 | pitrilysin metallepetidase 1 [Source:MGI Symbol;Acc:MGI:1916867] | Pitrm1 | Enriched in dLGN | 1.039 | 0.00002496 | 115.9 |
| ENSMUSG00000032096 | archain 1 [Source:MGI Symbol;Acc:MGI:2387591] | Arcn1 | Enriched in dLGN | 1.039 | 0.0009175 | 316.7 |
| ENSMUSG00000051671 | cytochrome c oxidase assembly factor 6 [Source:MGI Symbol;Acc:MGI:1915142] | Coa6 | Enriched in dLGN | 1.039 | 0.01272 | 13.32 |
| ENSMUSG00000026527 | regulator of G protein signaling 7 [Source:MGI Symbol;Acc:MGI:1346089] | Rgs7 | Enriched in dLGN | 1.038 | 0.02087 | 427.5 |
| ENSMUSG00000032336 | neuroplastin [Source:MGI Symbol;Acc:MGI:108077] | Nptn | Enriched in dLGN | 1.035 | 7.897E-07 | 662.1 |
| ENSMUSG00000032078 | ZPR1 zinc finger [Source:MGI Symbol;Acc:MGI:1330262] | Zpr1 | Enriched in dLGN | 1.035 | 0.000008332 | 88.62 |
| ENSMUSG00000029014 | DnaJ heat shock protein family (Hsp40) member C2 [Source:MGI Symbol;Acc:MGI:99470] | Dnajc2 | Enriched in dLGN | 1.035 | 0.00002935 | 134.3 |
| ENSMUSG00000028488 | SH3-domain GRB2-like 2 [Source:MGI Symbol;Acc:MGI:700009] | Sh3gl2 | Enriched in dLGN | 1.035 | 0.001966 | 586.9 |
| ENSMUSG00000022792 | tyrosyl-tRNA synthetase 2 (mitochondrial) [Source:MGI Symbol;Acc:MGI:1917370] | Yars2 | Enriched in dLGN | 1.034 | 0.001552 | 29.46 |
| ENSMUSG00000025236 | ADP-dependent glucokinase [Source:MGI Symbol;Acc:MGI:1919391] | Adpgk | Enriched in dLGN | 1.033 | 0.00008564 | 56.67 |
| ENSMUSG00000030264 | THUMP domain containing 3 [Source:MGI Symbol;Acc:MGI:1277973] | Thumpd3 | Enriched in dLGN | 1.033 | 0.003164 | 45.23 |
| ENSMUSG00000017801 | MAX-like protein X [Source:MGI Symbol;Acc:MGI:108398] | Mlx | Enriched in dLGN | 1.03 | 0.00582 | 24.55 |
| ENSMUSG00000042590 | importin 11 [Source:MGI Symbol;Acc:MGI:2442377] | Ipo11 | Enriched in dLGN | 1.028 | 0.00002891 | 139.3 |
| ENSMUSG00000022964 | transmembrane protein 50B [Source:MGI Symbol;Acc:MGI:1925225] | Tmem50b | Enriched in dLGN | 1.028 | 0.00003369 | 152.6 |
| ENSMUSG00000026914 | proteasome (prosome, macropain) 26S subunit, non-ATPase, 14 [Source:MGI Symbol;Acc:MGI:1913284] | Psmd14 | Enriched in dLGN | 1.027 | 0.00006276 | 139.6 |
| ENSMUSG00000090553 | small nuclear ribonucleoprotein E [Source:MGI Symbol;Acc:MGI:98346] | Snrpe | Enriched in dLGN | 1.027 | 0.0003217 | 27.79 |
| ENSMUSG00000020149 | RAB1A, member RAS oncogene family [Source:MGI Symbol;Acc:MGI:97842] | Rab1a | Enriched in dLGN | 1.026 | 0.0000173 | 300.1 |
| ENSMUSG00000003923 | transcription factor A, mitochondrial [Source:MGI Symbol;Acc:MGI:107810] | Tfam | Enriched in dLGN | 1.025 | 0.00002214 | 69.43 |
| ENSMUSG00000054452 | TLE family member 5, transcriptional modulator [Source:MGI Symbol;Acc:MGI:95806] | Tle5 | Enriched in dLGN | 1.024 | 1.434E-09 | 670.5 |
| ENSMUSG00000030057 | cellular nucleic acid binding protein [Source:MGI Symbol;Acc:MGI:88431] | Cnbp | Enriched in dLGN | 1.024 | 0.00004826 | 267.3 |
| ENSMUSG00000022706 | mitochondrial ribosomal protein L40 [Source:MGI Symbol;Acc:MGI:1332635] | Mrpl40 | Enriched in dLGN | 1.023 | 0.00003907 | 48.44 |
| ENSMUSG00000021987 | myotubularin related protein 6 [Source:MGI Symbol;Acc:MGI:2145637] | Mtmr6 | Enriched in dLGN | 1.022 | 5.747E-09 | 223.6 |
| ENSMUSG00000040022 | RAB11 family interacting protein 2 (class I) [Source:MGI Symbol;Acc:MGI:1922248] | Rab11fip2 | Enriched in dLGN | 1.022 | 3.663E-07 | 182.3 |
| ENSMUSG00000059278 | N(alpha)-acetyltransferase 38, NatC auxiliary subunit [Source:MGI Symbol;Acc:MGI:1925554] | Naa38 | Enriched in dLGN | 1.022 | 0.009374 | 31.83 |
| ENSMUSG00000057666 | glyceraldehyde-3-phosphate dehydrogenase [Source:MGI Symbol;Acc:MGI:95640] | Gapdh | Enriched in dLGN | 1.021 | 0.04502 | 93.82 |
| ENSMUSG00000040620 | DEAH (Asp-Glu-Ala-His) box polypeptide 33 [Source:MGI Symbol;Acc:MGI:2445102] | Dhx33 | Enriched in dLGN | 1.02 | 0.00002254 | 72.8 |
| ENSMUSG00000026825 | dynamin 1 [Source:MGI Symbol;Acc:MGI:107384] | Dnm1 | Enriched in dLGN | 1.02 | 0.0001203 | 2096 |
| ENSMUSG00000033597 | CASK interacting protein 1 [Source:MGI Symbol;Acc:MGI:2442952] | Caskin1 | Enriched in dLGN | 1.019 | 4.657E-07 | 1073 |
| ENSMUSG00000045287 | reticulon 4 receptor-like 1 [Source:MGI Symbol;Acc:MGI:2661375] | Rtn4rl1 | Enriched in dLGN | 1.019 | 0.00009879 | 303.8 |
| ENSMUSG00000038279 | NOP2 nucleolar protein [Source:MGI Symbol;Acc:MGI:107891] | Nop2 | Enriched in dLGN | 1.018 | 0.00002292 | 78.7 |
| ENSMUSG00000039000 | ubiquitin protein ligase E3C [Source:MGI Symbol;Acc:MGI:2140998] | Ube3c | Enriched in dLGN | 1.016 | 9.613E-10 | 441.8 |
| ENSMUSG00000022962 | phosphoribosylglycinamide formyltransferase [Source:MGI Symbol;Acc:MGI:95654] | Gart | Enriched in dLGN | 1.016 | 0.000007709 | 92.71 |
| ENSMUSG00000028964 | Parkinson disease (autosomal recessive, early onset) 7 [Source:MGI Symbol;Acc:MGI:2135637] | Park7 | Enriched in dLGN | 1.016 | 0.00001038 | 178.9 |
| ENSMUSG00000039197 | adenosine kinase [Source:MGI Symbol;Acc:MGI:87930] | Adk | Enriched in dLGN | 1.016 | 0.01785 | 98.44 |
| ENSMUSG00000028945 | Ras homolog enriched in brain [Source:MGI Symbol;Acc:MGI:97912] | Rheb | Enriched in dLGN | 1.015 | 1.163E-07 | 175.6 |
| ENSMUSG00000020261 | solute carrier family 36 (proton/amino acid symporter), member 1 [Source:MGI Symbol;Acc:MGI:2445299] | Slc36a1 | Enriched in dLGN | 1.015 | 0.00005332 | 223.9 |
| ENSMUSG00000034891 | synuclein, beta [Source:MGI Symbol;Acc:MGI:1889011] | Sncb | Enriched in dLGN | 1.015 | 0.001886 | 939.2 |
| ENSMUSG00000078941 | adenylate kinase 6 [Source:MGI Symbol;Acc:MGI:5510732] | Ak6 | Enriched in dLGN | 1.015 | 0.005816 | 18.09 |
| ENSMUSG00000000776 | polymerase (RNA) III (DNA directed) polypeptide D [Source:MGI Symbol;Acc:MGI:1914315] | Polr3d | Enriched in dLGN | 1.014 | 0.000001426 | 108.5 |
| ENSMUSG00000033628 | phosphatidylinositol 3-kinase catalytic subunit type 3 [Source:MGI Symbol;Acc:MGI:2445019] | Pik3c3 | Enriched in dLGN | 1.014 | 0.0001844 | 127.1 |
| ENSMUSG00000030432 | ribosomal protein L28 [Source:MGI Symbol;Acc:MGI:101839] | Rpl28 | Enriched in dLGN | 1.013 | 0.001056 | 78.46 |
| ENSMUSG00000026032 | NADH:ubiquinone oxidoreductase subunit B3 [Source:MGI Symbol;Acc:MGI:1913745] | Ndufb3 | Enriched in dLGN | 1.009 | 0.0004125 | 49.01 |
| ENSMUSG00000031918 | myotubularin related protein 2 [Source:MGI Symbol;Acc:MGI:1924366] | Mtmr2 | Enriched in dLGN | 1.008 | 0.0001254 | 157.4 |
| ENSMUSG00000039747 | ORAI calcium release-activated calcium modulator 2 [Source:MGI Symbol;Acc:MGI:2443195] | Orai2 | Enriched in dLGN | 1.007 | 0.00002327 | 138.6 |
| ENSMUSG00000048076 | ADP-ribosylation factor 1 [Source:MGI Symbol;Acc:MGI:99431] | Arf1 | Enriched in dLGN | 1.007 | 0.0004902 | 460.6 |
| ENSMUSG00000030079 | RuvB-like protein 1 [Source:MGI Symbol;Acc:MGI:1928760] | Ruvbl1 | Enriched in dLGN | 1.007 | 0.0009855 | 70.63 |
| ENSMUSG00000030092 | contactin 6 [Source:MGI Symbol;Acc:MGI:1858223] | Cntn6 | Enriched in dLGN | 1.007 | 0.008994 | 202.1 |
| ENSMUSG00000015759 | cornichon family AMPA receptor auxiliary protein 1 [Source:MGI Symbol;Acc:MGI:1277202] | Cnih1 | Enriched in dLGN | 1.005 | 0.0002458 | 68.86 |
| ENSMUSG00000001366 | f-box protein 9 [Source:MGI Symbol;Acc:MGI:1918788] | Fbxo9 | Enriched in dLGN | 1.004 | 1.402E-12 | 320.2 |
| ENSMUSG00000053838 | NudC domain containing 3 [Source:MGI Symbol;Acc:MGI:2144158] | Nudcd3 | Enriched in dLGN | 1.001 | 2.707E-10 | 600.9 |
| ENSMUSG00000039278 | proprotein convertase subtilisin/kexin type 1 inhibitor [Source:MGI Symbol;Acc:MGI:1353431] | Pcsk1n | Enriched in dLGN | 1 | 0.003411 | 2810 |
| ENSMUSG00000050663 | TRH-degrading enzyme [Source:MGI Symbol;Acc:MGI:2384311] | Trhde | Enriched in retina | -1 | 0.005032 | 309.3 |
| ENSMUSG00000036676 | transmembrane and tetratricopeptide repeat containing 3 [Source:MGI Symbol;Acc:MGI:3036255] | Tmtc3 | Enriched in retina | -1.001 | 0.00006816 | 82.34 |
| ENSMUSG00000039756 | deoxynucleotidyltransferase, terminal, interacting protein 2 [Source:MGI Symbol;Acc:MGI:1923173] | Dnttip2 | Enriched in retina | -1.003 | 0.002663 | 112.1 |
| ENSMUSG00000020694 | tousled-like kinase 2 (Arabidopsis) [Source:MGI Symbol;Acc:MGI:1346023] | Tlk2 | Enriched in retina | -1.012 | 0.000009107 | 331.4 |
| ENSMUSG00000040269 | mitochondrial ribosomal protein S28 [Source:MGI Symbol;Acc:MGI:1913480] | Mrps28 | Enriched in retina | -1.016 | 0.03677 | 40.7 |
| ENSMUSG00000037787 | cytochrome c oxidase assembly factor 8 [Source:MGI Symbol;Acc:MGI:1915270] | Coa8 | Enriched in retina | -1.018 | 0.0004781 | 74.03 |
| ENSMUSG00000031872 | brain expressed, associated with Nedd4, 1 [Source:MGI Symbol;Acc:MGI:1929597] | Bean1 | Enriched in retina | -1.032 | 0.009647 | 45.05 |
| ENSMUSG00000066568 | LSM14A mRNA processing body assembly factor [Source:MGI Symbol;Acc:MGI:1914320] | Lsm14a | Enriched in retina | -1.035 | 1.507E-11 | 280.1 |
| ENSMUSG00000025035 | ADP-ribosylation factor-like 3 [Source:MGI Symbol;Acc:MGI:1929699] | Arl3 | Enriched in retina | -1.054 | 0.000004429 | 316.8 |
| ENSMUSG00000055436 | serine and arginine-rich splicing factor 11 [Source:MGI Symbol;Acc:MGI:1916457] | Srsf11 | Enriched in retina | -1.055 | 0.001558 | 305.3 |
| ENSMUSG00000025049 | TATA-box binding protein associated factor 5 [Source:MGI Symbol;Acc:MGI:2442144] | Taf5 | Enriched in retina | -1.061 | 0.0001051 | 41.18 |
| ENSMUSG00000039910 | Cbp/p300-interacting transactivator, with Glu/Asp-rich carboxy-terminal domain, 2 [Source:MGI Symbol;Acc:MGI:1306784] | Cited2 | Enriched in retina | -1.062 | 0.00003639 | 198.6 |
| ENSMUSG00000028292 | arginyl-tRNA synthetase 2, mitochondrial [Source:MGI Symbol;Acc:MGI:1923596] | Rars2 | Enriched in retina | -1.069 | 0.01041 | 49.79 |
| ENSMUSG00000008496 | POU domain, class 2, transcription factor 2 [Source:MGI Symbol;Acc:MGI:101897] | Pou2f2 | Enriched in retina | -1.07 | 0.003508 | 295.3 |
| ENSMUSG00000034848 | tetratricopeptide repeat domain 21B [Source:MGI Symbol;Acc:MGI:1920918] | Ttc21b | Enriched in retina | -1.072 | 0.01066 | 76.94 |
| ENSMUSG00000028911 | serine and arginine-rich splicing factor 4 [Source:MGI Symbol;Acc:MGI:1890577] | Srsf4 | Enriched in retina | -1.072 | 5.364E-12 | 388.9 |
| ENSMUSG00000024067 | dpy-30, histone methyltransferase complex regulatory subunit [Source:MGI Symbol;Acc:MGI:1913560] | Dpy30 | Enriched in retina | -1.078 | 0.0001753 | 73.16 |
| ENSMUSG00000020166 | CCR4-NOT transcription complex, subunit 2 [Source:MGI Symbol;Acc:MGI:1919318] | Cnot2 | Enriched in retina | -1.079 | 0.0000057 | 172.3 |
| ENSMUSG00000023939 | mitochondrial ribosomal protein L14 [Source:MGI Symbol;Acc:MGI:1333864] | Mrpl14 | Enriched in retina | -1.084 | 0.005026 | 19.2 |
| ENSMUSG00000021552 | G kinase anchoring protein 1 [Source:MGI Symbol;Acc:MGI:1891694] | Gkap1 | Enriched in retina | -1.088 | 0.000082 | 95.16 |
| ENSMUSG00000020009 | interferon gamma receptor 1 [Source:MGI Symbol;Acc:MGI:107655] | Ifngr1 | Enriched in retina | -1.091 | 0.00001251 | 51.07 |
| ENSMUSG00000022788 | FYVE, RhoGEF and PH domain containing 4 [Source:MGI Symbol;Acc:MGI:2183747] | Fgd4 | Enriched in retina | -1.103 | 0.00000082 | 345 |
| ENSMUSG00000010110 | syntaxin 5A [Source:MGI Symbol;Acc:MGI:1928483] | Stx5a | Enriched in retina | -1.104 | 0.001148 | 74.48 |
| ENSMUSG00000035933 | component of oligomeric golgi complex 5 [Source:MGI Symbol;Acc:MGI:2145130] | Cog5 | Enriched in retina | -1.104 | 0.000584 | 161 |
| ENSMUSG00000060427 | zinc finger protein 868 [Source:MGI Symbol;Acc:MGI:2142546] | Zfp868 | Enriched in retina | -1.107 | 0.003244 | 33.49 |
| ENSMUSG00000005823 | G protein-coupled receptor 108 [Source:MGI Symbol;Acc:MGI:1925558] | Gpr108 | Enriched in retina | -1.113 | 0.00007506 | 51.17 |
| ENSMUSG00000045410 | aldo-keto reductase family 1, member E1 [Source:MGI Symbol;Acc:MGI:1914758] | Akr1e1 | Enriched in retina | -1.113 | 0.00001367 | 70.94 |
| ENSMUSG00000042535 | GTP binding protein 1 [Source:MGI Symbol;Acc:MGI:109443] | Gtpbp1 | Enriched in retina | -1.115 | 9.074E-11 | 209.5 |
| ENSMUSG00000034126 | protein-O-mannosyltransferase 2 [Source:MGI Symbol;Acc:MGI:2444430] | Pomt2 | Enriched in retina | -1.119 | 1.263E-07 | 138.2 |
| ENSMUSG00000020140 | leucine rich repeat containing G protein coupled receptor 5 [Source:MGI Symbol;Acc:MGI:1341817] | Lgr5 | Enriched in retina | -1.123 | 0.003122 | 46.54 |
| ENSMUSG00000034402 | potassium voltage-gated channel, subfamily H (eag-related), member 5 [Source:MGI Symbol;Acc:MGI:3584508] | Kcnh5 | Enriched in retina | -1.125 | 0.0006916 | 240.3 |
| ENSMUSG00000046311 | zinc finger protein 62 [Source:MGI Symbol;Acc:MGI:99662] | Zfp62 | Enriched in retina | -1.131 | 0.00009193 | 116.3 |
| ENSMUSG00000066149 | cell division cycle 26 [Source:MGI Symbol;Acc:MGI:1913690] | Cdc26 | Enriched in retina | -1.132 | 0.00003033 | 49.15 |
| ENSMUSG00000040234 | transmembrane 7 superfamily member 3 [Source:MGI Symbol;Acc:MGI:1914873] | Tm7sf3 | Enriched in retina | -1.14 | 0.000001296 | 102.7 |
| ENSMUSG00000060743 | H3.3 histone A [Source:MGI Symbol;Acc:MGI:1097686] | H3f3a | Enriched in retina | -1.144 | 0.0006007 | 111.3 |
| ENSMUSG00000028576 | intraflagellar transport 74 [Source:MGI Symbol;Acc:MGI:1914944] | Ift74 | Enriched in retina | -1.146 | 0.0001603 | 79.32 |
| ENSMUSG00000037646 | vacuolar protein sorting 13B [Source:MGI Symbol;Acc:MGI:1916380] | Vps13b | Enriched in retina | -1.155 | 0.000001597 | 601.4 |
| ENSMUSG00000005886 | nuclear receptor coactivator 2 [Source:MGI Symbol;Acc:MGI:1276533] | Ncoa2 | Enriched in retina | -1.155 | 2.149E-10 | 793.7 |
| ENSMUSG00000018428 | A kinase (PRKA) anchor protein 1 [Source:MGI Symbol;Acc:MGI:104729] | Akap1 | Enriched in retina | -1.157 | 0.000009518 | 231.2 |
| ENSMUSG00000028013 | pyrophosphatase (inorganic) 2 [Source:MGI Symbol;Acc:MGI:1922026] | Ppa2 | Enriched in retina | -1.169 | 0.000003186 | 141.1 |
| ENSMUSG00000026239 | phosphodiesterase 6D, cGMP-specific, rod, delta [Source:MGI Symbol;Acc:MGI:1270843] | Pde6d | Enriched in retina | -1.17 | 0.00001806 | 100.7 |
| ENSMUSG00000049511 | 5-hydroxytryptamine (serotonin) receptor 1B [Source:MGI Symbol;Acc:MGI:96274] | Htr1b | Enriched in retina | -1.183 | 0.003311 | 142.5 |
| ENSMUSG00000092607 | sodium channel modifier 1 [Source:MGI Symbol;Acc:MGI:1341284] | Scnm1 | Enriched in retina | -1.196 | 0.002479 | 37.88 |
| ENSMUSG00000015980 | leucine rich repeat containing 27 [Source:MGI Symbol;Acc:MGI:1923862] | Lrrc27 | Enriched in retina | -1.206 | 0.001944 | 44.92 |
| ENSMUSG00000034075 | zinc finger, DHHC domain containing 5 [Source:MGI Symbol;Acc:MGI:1923573] | Zdhhc5 | Enriched in retina | -1.208 | 5.877E-10 | 421.5 |
| ENSMUSG00000023892 | zinc finger protein 51 [Source:MGI Symbol;Acc:MGI:99198] | Zfp51 | Enriched in retina | -1.212 | 0.01007 | 15.06 |
| ENSMUSG00000024483 | ankyrin repeat and KH domain containing 1 [Source:MGI Symbol;Acc:MGI:1921733] | Ankhd1 | Enriched in retina | -1.212 | 1.531E-08 | 373.2 |
| ENSMUSG00000029422 | arginine/serine-rich coiled-coil 2 [Source:MGI Symbol;Acc:MGI:1913489] | Rsrc2 | Enriched in retina | -1.219 | 0.0001134 | 465.1 |
| ENSMUSG00000049878 | rearranged L-myc fusion sequence [Source:MGI Symbol;Acc:MGI:1924705] | Rlf | Enriched in retina | -1.22 | 2.237E-07 | 212.5 |
| ENSMUSG00000026739 | Bmi1 polycomb ring finger oncogene [Source:MGI Symbol;Acc:MGI:88174] | Bmi1 | Enriched in retina | -1.22 | 1.138E-07 | 140.4 |
| ENSMUSG00000032384 | casein kinase 1, gamma 1 [Source:MGI Symbol;Acc:MGI:2660884] | Csnk1g1 | Enriched in retina | -1.227 | 1.638E-07 | 220.4 |
| ENSMUSG00000030619 | embryonic ectoderm development [Source:MGI Symbol;Acc:MGI:95286] | Eed | Enriched in retina | -1.233 | 0.00005282 | 53.97 |
| ENSMUSG00000067995 | general transcription factor IIF, polypeptide 2 [Source:MGI Symbol;Acc:MGI:1915955] | Gtf2f2 | Enriched in retina | -1.234 | 0.00005185 | 71.17 |
| ENSMUSG00000028051 | hyperpolarization-activated, cyclic nucleotide-gated K+ 3 [Source:MGI Symbol;Acc:MGI:1298211] | Hcn3 | Enriched in retina | -1.245 | 0.00005888 | 158.3 |
| ENSMUSG00000024347 | pleckstrin and Sec7 domain containing 2 [Source:MGI Symbol;Acc:MGI:1921252] | Psd2 | Enriched in retina | -1.272 | 0.00184 | 258.6 |
| ENSMUSG00000000028 | cell division cycle 45 [Source:MGI Symbol;Acc:MGI:1338073] | Cdc45 | Enriched in retina | -1.273 | 0.02986 | 7.114 |
| ENSMUSG00000035314 | glycerophosphodiester phosphodiesterase domain containing 5 [Source:MGI Symbol;Acc:MGI:2686926] | Gdpd5 | Enriched in retina | -1.289 | 0.001131 | 328.8 |
| ENSMUSG00000020547 | basic leucine zipper and W2 domains 2 [Source:MGI Symbol;Acc:MGI:1914162] | Bzw2 | Enriched in retina | -1.297 | 0.00005578 | 169.6 |
| ENSMUSG00000035845 | asparagine-linked glycosylation 12 (alpha-1,6-mannosyltransferase) [Source:MGI Symbol;Acc:MGI:2385025] | Alg12 | Enriched in retina | -1.305 | 0.0009781 | 28.22 |
| ENSMUSG00000039913 | p21 (RAC1) activated kinase 5 [Source:MGI Symbol;Acc:MGI:1920334] | Pak5 | Enriched in retina | -1.319 | 0.002696 | 253.3 |
| ENSMUSG00000048772 | transmembrane protein 53 [Source:MGI Symbol;Acc:MGI:1916027] | Tmem53 | Enriched in retina | -1.343 | 0.04558 | 7.714 |
| ENSMUSG00000019726 | lysosomal trafficking regulator [Source:MGI Symbol;Acc:MGI:107448] | Lyst | Enriched in retina | -1.352 | 4.746E-11 | 465.9 |
| ENSMUSG00000056851 | poly(rC) binding protein 2 [Source:MGI Symbol;Acc:MGI:108202] | Pcbp2 | Enriched in retina | -1.358 | 2.477E-10 | 828.1 |
| ENSMUSG00000026778 | protein kinase C, theta [Source:MGI Symbol;Acc:MGI:97601] | Prkcq | Enriched in retina | -1.367 | 0.00168 | 111.9 |
| ENSMUSG00000029012 | origin recognition complex, subunit 5 [Source:MGI Symbol;Acc:MGI:1347044] | Orc5 | Enriched in retina | -1.372 | 0.00559 | 47.78 |
| ENSMUSG00000036023 | poly (ADP-ribose) polymerase family, member 2 [Source:MGI Symbol;Acc:MGI:1341112] | Parp2 | Enriched in retina | -1.379 | 0.0007928 | 46.73 |
| ENSMUSG00000000276 | diacylglycerol kinase, epsilon [Source:MGI Symbol;Acc:MGI:1889276] | Dgke | Enriched in retina | -1.387 | 2.593E-07 | 375.4 |
| ENSMUSG00000045968 | transmembrane epididymal family member 2 [Source:MGI Symbol;Acc:MGI:1923273] | Teddm2 | Enriched in retina | -1.401 | 0.001952 | 15.55 |
| ENSMUSG00000070806 | zinc finger, MYND domain containing 12 [Source:MGI Symbol;Acc:MGI:2140259] | Zmynd12 | Enriched in retina | -1.408 | 0.02329 | 9.762 |
| ENSMUSG00000014075 | dynein light chain Tctex-type 2B [Source:MGI Symbol;Acc:MGI:1913311] | Dynlt2b | Enriched in retina | -1.414 | 0.01304 | 16.09 |
| ENSMUSG00000040936 | unc-51-like kinase 4 [Source:MGI Symbol;Acc:MGI:1921622] | Ulk4 | Enriched in retina | -1.428 | 0.004342 | 132.6 |
| ENSMUSG00000028080 | LPS-responsive beige-like anchor [Source:MGI Symbol;Acc:MGI:1933162] | Lrba | Enriched in retina | -1.433 | 0.000005319 | 681.5 |
| ENSMUSG00000027259 | adenosine deaminase-like [Source:MGI Symbol;Acc:MGI:1923144] | Adal | Enriched in retina | -1.434 | 0.000001134 | 54.84 |
| ENSMUSG00000020069 | heterogeneous nuclear ribonucleoprotein H3 [Source:MGI Symbol;Acc:MGI:1926462] | Hnrnph3 | Enriched in retina | -1.436 | 0.000001194 | 152.6 |
| ENSMUSG00000025217 | beta-transducin repeat containing protein [Source:MGI Symbol;Acc:MGI:1338871] | Btrc | Enriched in retina | -1.443 | 0.000004574 | 344.2 |
| ENSMUSG00000037788 | vesicular, overexpressed in cancer, prosurvival protein 1 [Source:MGI Symbol;Acc:MGI:2141658] | Vopp1 | Enriched in retina | -1.46 | 5.603E-11 | 391.5 |

| | | | | | | |
|---|---|---|---|---|---|---|
| ENSMUSG00000064115 | cell adhesion molecule 2 [Source:MGI Symbol;Acc:MGI:2442722] | Cadm2 | Enriched in retina | -1.462 | 0.0003121 | 1919 |
| ENSMUSG00000056832 | tetratricopeptide repeat domain 26 [Source:MGI Symbol;Acc:MGI:2444853] | Ttc26 | Enriched in retina | -1.469 | 0.0002875 | 27.07 |
| ENSMUSG00000031399 | FAM3 metabolism regulating signaling molecule A [Source:MGI Symbol;Acc:MGI:1913544] | Fam3a | Enriched in retina | -1.469 | 0.00001919 | 44.59 |
| ENSMUSG00000040591 | centriolar satellite-associated tubulin polyglutamylase complex regulator 1 [Source:MGI Symbol;Acc:MGI:1915079] | Cstpp1 | Enriched in retina | -1.479 | 0.000001919 | 214.8 |
| ENSMUSG00000020067 | myopalladin [Source:MGI Symbol;Acc:MGI:1916052] | Mypn | Enriched in retina | -1.481 | 0.004254 | 14.16 |
| ENSMUSG00000034610 | terminal uridylyl transferase 4 [Source:MGI Symbol;Acc:MGI:2445126] | Tut4 | Enriched in retina | -1.485 | 1.042E-07 | 281.6 |
| ENSMUSG00000046380 | jerky [Source:MGI Symbol;Acc:MGI:106214] | Jrk | Enriched in retina | -1.5 | 0.000001229 | 43.1 |
| ENSMUSG00000021519 | mitochondrial transcription termination factor 3 [Source:MGI Symbol;Acc:MGI:1913660] | Mterf3 | Enriched in retina | -1.525 | 0.001382 | 36.34 |
| ENSMUSG00000034349 | structural maintenance of chromosomes 4 [Source:MGI Symbol;Acc:MGI:1917349] | Smc4 | Enriched in retina | -1.526 | 0.00002657 | 62.95 |
| ENSMUSG00000024891 | solute carrier family 29 (nucleoside transporters), member 2 [Source:MGI Symbol;Acc:MGI:1345278] | Slc29a2 | Enriched in retina | -1.565 | 4.588E-07 | 40.3 |
| ENSMUSG00000038122 | TBC1 domain family, member 32 [Source:MGI Symbol;Acc:MGI:2442827] | Tbc1d32 | Enriched in retina | -1.57 | 2.665E-07 | 100.5 |
| ENSMUSG00000106918 | mitochondrial ribosomal protein L33 [Source:MGI Symbol;Acc:MGI:2137225] | Mrpl33 | Enriched in retina | -1.594 | 0.0000277 | 55.34 |
| ENSMUSG00000019897 | coiled-coil domain containing 59 [Source:MGI Symbol;Acc:MGI:1289302] | Ccdc59 | Enriched in retina | -1.603 | 0.000001019 | 84.14 |
| ENSMUSG00000002679 | mediator complex subunit 6 [Source:MGI Symbol;Acc:MGI:1917042] | Med6 | Enriched in retina | -1.605 | 0.000367 | 28.53 |
| ENSMUSG00000024500 | protein phosphatase 2, regulatory subunit B, beta [Source:MGI Symbol;Acc:MGI:1920180] | Ppp2r2b | Enriched in retina | -1.626 | 0.00001014 | 744.1 |
| ENSMUSG00000031922 | centrosomal protein 57 [Source:MGI Symbol;Acc:MGI:1915551] | Cep57 | Enriched in retina | -1.627 | 0.000001367 | 52.14 |
| ENSMUSG00000039168 | death-associated protein [Source:MGI Symbol;Acc:MGI:1918190] | Dap | Enriched in retina | -1.643 | 0.0006245 | 35.43 |
| ENSMUSG00000041992 | Rap guanine nucleotide exchange factor (GEF) 5 [Source:MGI Symbol;Acc:MGI:2444365] | Rapgef5 | Enriched in retina | -1.66 | 4.888E-13 | 377.8 |
| ENSMUSG00000034295 | formin homology 2 domain containing 3 [Source:MGI Symbol;Acc:MGI:1925847] | Fhod3 | Enriched in retina | -1.661 | 1.774E-08 | 470.1 |
| ENSMUSG00000038079 | transmembrane protein 237 [Source:MGI Symbol;Acc:MGI:2138365] | Tmem237 | Enriched in retina | -1.679 | 9.406E-18 | 98.82 |
| ENSMUSG00000042557 | transcriptional regulator, SIN3A (yeast) [Source:MGI Symbol;Acc:MGI:107157] | Sin3a | Enriched in retina | -1.681 | 3.253E-22 | 223.5 |
| ENSMUSG00000039753 | F-box and leucine-rich repeat protein 5 [Source:MGI Symbol;Acc:MGI:2152883] | Fbxl5 | Enriched in retina | -1.688 | 8.758E-13 | 292.1 |
| ENSMUSG00000068250 | antagonist of mitotic exit network 1 [Source:MGI Symbol;Acc:MGI:2442933] | Amn1 | Enriched in retina | -1.691 | 0.0008354 | 40.08 |
| ENSMUSG00000033849 | UDP-Gal:betaGlcNAc beta 1,3-galactosyltransferase, polypeptide 2 [Source:MGI Symbol;Acc:MGI:1349461] | B3galt2 | Enriched in retina | -1.693 | 1.473E-08 | 115.1 |
| ENSMUSG00000028197 | collagen, type XXIV, alpha 1 [Source:MGI Symbol;Acc:MGI:1918605] | Col24a1 | Enriched in retina | -1.697 | 0.01774 | 14.18 |
| ENSMUSG00000031937 | V-set and transmembrane domain containing 5 [Source:MGI Symbol;Acc:MGI:1916387] | Vstm5 | Enriched in retina | -1.698 | 0.001213 | 21.85 |
| ENSMUSG00000046985 | transmembrane anterior posterior transformation 1 [Source:MGI Symbol;Acc:MGI:2683537] | Tapt1 | Enriched in retina | -1.704 | 2.842E-09 | 186.8 |
| ENSMUSG00000039860 | serine/arginine repetitive matrix 3 [Source:MGI Symbol;Acc:MGI:1920309] | Srrm3 | Enriched in retina | -1.706 | 1.624E-11 | 438.4 |
| ENSMUSG00000040703 | cytochrome P450, family 2, subfamily s, polypeptide 1 [Source:MGI Symbol;Acc:MGI:1921384] | Cyp2s1 | Enriched in retina | -1.714 | 0.0003166 | 19.07 |
| ENSMUSG00000042508 | cyclin D binding myb like transcription factor 1 [Source:MGI Symbol;Acc:MGI:1344415] | Dmtf1 | Enriched in retina | -1.722 | 0.000005491 | 197.7 |
| ENSMUSG00000054469 | lysocardiolipin acyltransferase 1 [Source:MGI Symbol;Acc:MGI:2684937] | Lclat1 | Enriched in retina | -1.731 | 1.098E-08 | 128 |
| ENSMUSG00000044229 | neurexophilin and PC-esterase domain family, member 4 [Source:MGI Symbol;Acc:MGI:1924792] | Nxpe4 | Enriched in retina | -1.739 | 1.571E-08 | 105.5 |
| ENSMUSG00000030779 | retinoblastoma binding protein 6, ubiquitin ligase [Source:MGI Symbol;Acc:MGI:894835] | Rbbp6 | Enriched in retina | -1.739 | 1.934E-10 | 688.1 |
| ENSMUSG00000027674 | peroxisomal biogenesis factor 5-like [Source:MGI Symbol;Acc:MGI:1916672] | Pex5l | Enriched in retina | -1.74 | 0.00004187 | 789.1 |
| ENSMUSG00000032409 | ataxia telangiectasia and Rad3 related [Source:MGI Symbol;Acc:MGI:108028] | Atr | Enriched in retina | -1.747 | 1.893E-07 | 87.66 |
| ENSMUSG00000060969 | Iroquois homeobox 1 [Source:MGI Symbol;Acc:MGI:1197515] | Irx1 | Enriched in retina | -1.758 | 0.007152 | 29.96 |
| ENSMUSG00000022471 | X-ray repair complementing defective repair in Chinese hamster cells 6 [Source:MGI Symbol;Acc:MGI:95606] | Xrcc6 | Enriched in retina | -1.778 | 0.000006122 | 97.01 |
| ENSMUSG00000031536 | polymerase (DNA directed), beta [Source:MGI Symbol;Acc:MGI:97740] | Polb | Enriched in retina | -1.78 | 0.0000116 | 159.7 |
| ENSMUSG00000034163 | zinc finger, C3H1-type containing [Source:MGI Symbol;Acc:MGI:2446143] | Zfc3h1 | Enriched in retina | -1.787 | 8.191E-09 | 428.2 |
| ENSMUSG00000045095 | membrane associated guanylate kinase, WW and PDZ domain containing 1 [Source:MGI Symbol;Acc:MGI:1203522] | Magi1 | Enriched in retina | -1.826 | 1.706E-12 | 743.1 |
| ENSMUSG00000055116 | aryl hydrocarbon receptor nuclear translocator-like [Source:MGI Symbol;Acc:MGI:1096381] | Arntl | Enriched in retina | -1.834 | 0.00000031 | 222.7 |
| ENSMUSG00000024666 | transmembrane protein 138 [Source:MGI Symbol;Acc:MGI:1920232] | Tmem138 | Enriched in retina | -1.84 | 0.00003097 | 25.53 |
| ENSMUSG00000029817 | transformer 2 alpha [Source:MGI Symbol;Acc:MGI:1933972] | Tra2a | Enriched in retina | -1.859 | 0.00003255 | 258.1 |
| ENSMUSG00000037818 | abhydrolase domain containing 18 [Source:MGI Symbol;Acc:MGI:1915468] | Abhd18 | Enriched in retina | -1.876 | 0.00003138 | 66.82 |
| ENSMUSG00000013846 | ST3 beta-galactoside alpha-2,3-sialyltransferase 1 [Source:MGI Symbol;Acc:MGI:98304] | St3gal1 | Enriched in retina | -1.886 | 9.47E-15 | 259.6 |
| ENSMUSG00000036181 | H1.2 linker histone, cluster member [Source:MGI Symbol;Acc:MGI:1931526] | H1f2 | Enriched in retina | -1.899 | 4.85E-09 | 187.7 |
| ENSMUSG00000035517 | tudor domain containing 7 [Source:MGI Symbol;Acc:MGI:2140279] | Tdrd7 | Enriched in retina | -1.933 | 2.345E-24 | 468.1 |
| ENSMUSG00000057914 | calcium channel, voltage-dependent, beta 2 subunit [Source:MGI Symbol;Acc:MGI:894644] | Cacnb2 | Enriched in retina | -1.935 | 3.018E-08 | 527.9 |
| ENSMUSG00000023495 | poly(rC) binding protein 4 [Source:MGI Symbol;Acc:MGI:1890471] | Pcbp4 | Enriched in retina | -1.953 | 1.125E-25 | 550 |
| ENSMUSG00000016356 | collagen, type XX, alpha 1 [Source:MGI Symbol;Acc:MGI:1920618] | Col20a1 | Enriched in retina | -1.96 | 0.00008254 | 25.93 |
| ENSMUSG00000054728 | phosphatase and actin regulator 1 [Source:MGI Symbol;Acc:MGI:2659021] | Phactr1 | Enriched in retina | -1.962 | 4.027E-12 | 986.4 |
| ENSMUSG00000036572 | UPF3 regulator of nonsense transcripts homolog B (yeast) [Source:MGI Symbol;Acc:MGI:1915384] | Upf3b | Enriched in retina | -1.967 | 1.65E-08 | 271.8 |
| ENSMUSG00000023235 | chemokine (C-C motif) ligand 25 [Source:MGI Symbol;Acc:MGI:1099448] | Ccl25 | Enriched in retina | -1.973 | 0.00001824 | 57.94 |
| ENSMUSG00000043243 | niban apoptosis regulator 3 [Source:MGI Symbol;Acc:MGI:3686743] | Niban3 | Enriched in retina | -1.978 | 0.01419 | 8.227 |
| ENSMUSG00000032558 | nephronophthisis 3 (adolescent) [Source:MGI Symbol;Acc:MGI:1921275] | Nphp3 | Enriched in retina | -1.983 | 0.0001608 | 34.65 |
| ENSMUSG00000035620 | RIC8 guanine nucleotide exchange factor B [Source:MGI Symbol;Acc:MGI:2682307] | Ric8b | Enriched in retina | -1.984 | 5.661E-13 | 318 |
| ENSMUSG00000021916 | glycosyltransferase 8 domain containing 1 [Source:MGI Symbol;Acc:MGI:1923735] | Glt8d1 | Enriched in retina | -1.991 | 6.936E-07 | 54.33 |
| ENSMUSG00000060568 | family with sequence similarity 78, member B [Source:MGI Symbol;Acc:MGI:2443050] | Fam78b | Enriched in retina | -2.004 | 7.708E-10 | 234.4 |
| ENSMUSG00000024424 | tetratricopeptide repeat domain 39C [Source:MGI Symbol;Acc:MGI:1919997] | Ttc39c | Enriched in retina | -2.009 | 3.482E-09 | 109.8 |
| ENSMUSG00000036822 | topoisomerase I binding, arginine/serine-rich [Source:MGI Symbol;Acc:MGI:2146189] | Topors | Enriched in retina | -2.01 | 6.79E-20 | 180.5 |
| ENSMUSG00000037801 | IQ motif containing H [Source:MGI Symbol;Acc:MGI:1925500] | Iqch | Enriched in retina | -2.013 | 0.002242 | 9.49 |
| ENSMUSG00000042396 | RNA binding motif protein 7 [Source:MGI Symbol;Acc:MGI:1914260] | Rbm7 | Enriched in retina | -2.016 | 2.01E-13 | 65.46 |
| ENSMUSG00000068615 | gap junction protein, delta 2 [Source:MGI Symbol;Acc:MGI:1334209] | Gjd2 | Enriched in retina | -2.033 | 5.637E-20 | 192.5 |
| ENSMUSG00000040490 | leucine rich repeat and fibronectin type III domain containing 2 [Source:MGI Symbol;Acc:MGI:1917780] | Lrfn2 | Enriched in retina | -2.038 | 0.000005539 | 166.4 |
| ENSMUSG00000073792 | asparagine-linked glycosylation 6 (alpha-1,3,-glucosyltransferase) [Source:MGI Symbol;Acc:MGI:2444031] | Alg6 | Enriched in retina | -2.046 | 4.557E-07 | 28.53 |
| ENSMUSG00000032599 | inositol hexaphosphate kinase 2 [Source:MGI Symbol;Acc:MGI:1923750] | Ip6k2 | Enriched in retina | -2.05 | 4.715E-11 | 280.5 |
| ENSMUSG00000028572 | hook microtubule tethering protein 1 [Source:MGI Symbol;Acc:MGI:1925213] | Hook1 | Enriched in retina | -2.051 | 9.006E-13 | 348.9 |
| ENSMUSG00000047141 | zinc finger protein 654 [Source:MGI Symbol;Acc:MGI:1919270] | Zfp654 | Enriched in retina | -2.055 | 3.075E-12 | 224.3 |
| ENSMUSG00000019865 | neuromedin B receptor [Source:MGI Symbol;Acc:MGI:1100525] | Nmbr | Enriched in retina | -2.069 | 0.0006074 | 8.779 |
| ENSMUSG00000050822 | solute carrier family 29 (nucleoside transporters), member 4 [Source:MGI Symbol;Acc:MGI:2385330] | Slc29a4 | Enriched in retina | -2.101 | 1.066E-08 | 126.5 |
| ENSMUSG00000027895 | potassium voltage gated channel, Shaw-related subfamily, member 4 [Source:MGI Symbol;Acc:MGI:96670] | Kcnc4 | Enriched in retina | -2.11 | 3.04E-09 | 208.4 |
| ENSMUSG00000022257 | lysosomal-associated protein transmembrane 4B [Source:MGI Symbol;Acc:MGI:1890494] | Laptm4b | Enriched in retina | -2.119 | 8.904E-22 | 165.8 |
| ENSMUSG00000031636 | PDZ and LIM domain 3 [Source:MGI Symbol;Acc:MGI:1859274] | Pdlim3 | Enriched in retina | -2.128 | 0.00002626 | 14.05 |
| ENSMUSG00000036995 | ArfGAP with SH3 domain, ankyrin repeat and PH domain 3 [Source:MGI Symbol;Acc:MGI:2684986] | Asap3 | Enriched in retina | -2.164 | 7.781E-09 | 47.68 |
| ENSMUSG00000001333 | syncoilin [Source:MGI Symbol;Acc:MGI:1916078] | Sync | Enriched in retina | -2.169 | 0.0006408 | 17.38 |
| ENSMUSG00000001504 | Iroquois homeobox 2 [Source:MGI Symbol;Acc:MGI:1197526] | Irx2 | Enriched in retina | -2.172 | 0.00004242 | 24.13 |
| ENSMUSG00000021125 | arginase type II [Source:MGI Symbol;Acc:MGI:1330806] | Arg2 | Enriched in retina | -2.174 | 0.00004189 | 11.96 |
| ENSMUSG00000096696 | zinc finger protein 960 [Source:MGI Symbol;Acc:MGI:3052731] | Zfp960 | Enriched in retina | -2.179 | 0.003586 | 4 |
| ENSMUSG00000040621 | gem nuclear organelle associated protein 8 [Source:MGI Symbol;Acc:MGI:2384300] | Gemin8 | Enriched in retina | -2.18 | 0.0006955 | 20.06 |
| ENSMUSG00000056260 | ligand dependent nuclear receptor interacting factor 1 [Source:MGI Symbol;Acc:MGI:2445214] | Lrif1 | Enriched in retina | -2.191 | 0.00000388 | 40.61 |
| ENSMUSG00000000552 | zinc finger protein 385A [Source:MGI Symbol;Acc:MGI:1352495] | Zfp385a | Enriched in retina | -2.211 | 3.746E-26 | 618.4 |
| ENSMUSG00000078963 | heat shock factor binding protein 1-like 1 [Source:MGI Symbol;Acc:MGI:1913505] | Hsbp1l1 | Enriched in retina | -2.232 | 9.434E-07 | 35.03 |
| ENSMUSG00000021619 | autophagy related 10 [Source:MGI Symbol;Acc:MGI:1914045] | Atg10 | Enriched in retina | -2.252 | 0.00008475 | 39.83 |
| ENSMUSG00000043668 | TOX high mobility group box family member 3 [Source:MGI Symbol;Acc:MGI:3039593] | Tox3 | Enriched in retina | -2.269 | 6.961E-09 | 105.2 |
| ENSMUSG00000030189 | Y box protein 3 [Source:MGI Symbol;Acc:MGI:2137670] | Ybx3 | Enriched in retina | -2.29 | 1.295E-12 | 527 |
| ENSMUSG00000038457 | transmembrane protein 255B [Source:MGI Symbol;Acc:MGI:2685533] | Tmem255b | Enriched in retina | -2.302 | 2.321E-11 | 59.61 |
| ENSMUSG00000026646 | suppressor of variegation 3-9 2 [Source:MGI Symbol;Acc:MGI:1890396] | Suv39h2 | Enriched in retina | -2.34 | 6.97E-10 | 51.55 |
| ENSMUSG00000030592 | ryanodine receptor 1, skeletal muscle [Source:MGI Symbol;Acc:MGI:99659] | Ryr1 | Enriched in retina | -2.352 | 0.00004471 | 72.59 |
| ENSMUSG00000031257 | NADPH oxidase 1 [Source:MGI Symbol;Acc:MGI:2450016] | Nox1 | Enriched in retina | -2.375 | 0.000209 | 7.043 |
| ENSMUSG00000020994 | pinin [Source:MGI Symbol;Acc:MGI:1100514] | Pnn | Enriched in retina | -2.394 | 2.243E-12 | 615.1 |
| ENSMUSG00000032192 | guanine nucleotide binding protein (G protein), beta 5 [Source:MGI Symbol;Acc:MGI:101848] | Gnb5 | Enriched in retina | -2.41 | 1.943E-23 | 441.8 |
| ENSMUSG00000029190 | DNA segment, Chr 5, ERATO Doi 579, expressed [Source:MGI Symbol;Acc:MGI:1261849] | D5Ertd579e | Enriched in retina | -2.414 | 0.000006739 | 1250 |
| ENSMUSG00000027016 | zinc finger protein 385B [Source:MGI Symbol;Acc:MGI:2444734] | Zfp385b | Enriched in retina | -2.433 | 4.297E-08 | 428.2 |
| ENSMUSG00000022722 | ADP-ribosylation factor-like 6 [Source:MGI Symbol;Acc:MGI:1927136] | Arl6 | Enriched in retina | -2.441 | 1.072E-12 | 147.9 |
| ENSMUSG00000024856 | CDK2-associated protein 2 [Source:MGI Symbol;Acc:MGI:1098779] | Cdk2ap2 | Enriched in retina | -2.466 | 2.706E-11 | 59.04 |
| ENSMUSG00000042369 | RNA binding motif protein 45 [Source:MGI Symbol;Acc:MGI:2387367] | Rbm45 | Enriched in retina | -2.467 | 3.812E-07 | 110.5 |
| ENSMUSG00000025876 | unc-5 netrin receptor A [Source:MGI Symbol;Acc:MGI:894682] | Unc5a | Enriched in retina | -2.508 | 2.035E-13 | 196.1 |
| ENSMUSG00000042050 | dynein 2 intermediate chain 1 [Source:MGI Symbol;Acc:MGI:2445085] | Dync2i1 | Enriched in retina | -2.511 | 9.414E-16 | 228.8 |
| ENSMUSG00000022803 | popeye domain containing 2 [Source:MGI Symbol;Acc:MGI:1930150] | Popdc2 | Enriched in retina | -2.519 | 0.02208 | 4.598 |
| ENSMUSG00000047495 | DLG associated protein 2 [Source:MGI Symbol;Acc:MGI:2443181] | Dlgap2 | Enriched in retina | -2.552 | 2.225E-07 | 201 |
| ENSMUSG00000053062 | junction adhesion molecule 2 [Source:MGI Symbol;Acc:MGI:1933820] | Jam2 | Enriched in retina | -2.572 | 4.424E-16 | 304.1 |
| ENSMUSG00000027014 | CWC22 spliceosome-associated protein [Source:MGI Symbol;Acc:MGI:2136773] | Cwc22 | Enriched in retina | -2.665 | 7.246E-09 | 206.1 |
| ENSMUSG00000055150 | zinc finger protein 78 [Source:MGI Symbol;Acc:MGI:107783] | Zfp78 | Enriched in retina | -2.672 | 2.038E-08 | 33.89 |
| ENSMUSG00000067367 | Ly1 antibody reactive clone [Source:MGI Symbol;Acc:MGI:107470] | Lyar | Enriched in retina | -2.695 | 8.551E-24 | 128.3 |
| ENSMUSG00000027601 | mitochondrial fission regulator 1 [Source:MGI Symbol;Acc:MGI:1914722] | Mtfr1 | Enriched in retina | -2.696 | 4.24E-11 | 98.99 |
| ENSMUSG00000020435 | oxysterol binding protein 2 [Source:MGI Symbol;Acc:MGI:1921559] | Osbp2 | Enriched in retina | -2.71 | 2.454E-22 | 628.7 |
| ENSMUSG00000031734 | Iroquois related homeobox 3 [Source:MGI Symbol;Acc:MGI:1197522] | Irx3 | Enriched in retina | -2.712 | 0.000003113 | 16.38 |
| ENSMUSG00000028188 | spermatogenesis associated 1 [Source:MGI Symbol;Acc:MGI:1918201] | Spata1 | Enriched in retina | -2.716 | 0.001575 | 64.99 |
| ENSMUSG00000022092 | protein phosphatase 3, catalytic subunit, gamma isoform [Source:MGI Symbol;Acc:MGI:107162] | Ppp3cc | Enriched in retina | -2.742 | 3.828E-17 | 226.2 |
| ENSMUSG00000004561 | methyltransferase like 17 [Source:MGI Symbol;Acc:MGI:1098577] | Mettl17 | Enriched in retina | -2.747 | 4.985E-08 | 47.96 |
| ENSMUSG00000072572 | solute carrier family 39 (zinc transporter), member 2 [Source:MGI Symbol;Acc:MGI:2684326] | Slc39a2 | Enriched in retina | -2.784 | 0.0002679 | 19.9 |
| ENSMUSG00000002980 | basal cell adhesion molecule [Source:MGI Symbol;Acc:MGI:1929940] | Bcam | Enriched in retina | -2.804 | 5.428E-09 | 83.63 |
| ENSMUSG00000001120 | poly(rC) binding protein 3 [Source:MGI Symbol;Acc:MGI:1890470] | Pcbp3 | Enriched in retina | -2.816 | 6.761E-39 | 628.3 |
| ENSMUSG00000091402 | retinal degeneration 3-like [Source:MGI Symbol;Acc:MGI:2675860] | Rd3l | Enriched in retina | -2.827 | 2.102E-09 | 41.89 |
| ENSMUSG00000032743 | katanin interacting protein [Source:MGI Symbol;Acc:MGI:2442760] | Katnip | Enriched in retina | -2.857 | 2.031E-25 | 322.2 |
| ENSMUSG00000037325 | Bardet-Biedl syndrome 7 (human) [Source:MGI Symbol;Acc:MGI:1918742] | Bbs7 | Enriched in retina | -2.876 | 7.656E-15 | 164.6 |
| ENSMUSG00000092210 | RIKEN cDNA A930009A15 gene [Source:MGI Symbol;Acc:MGI:1925048] | A930009A15Rik | Enriched in retina | -2.88 | 0.0002004 | 13.52 |
| ENSMUSG00000037973 | ITPR interacting domain containing 1 [Source:MGI Symbol;Acc:MGI:2685304] | Itprid1 | Enriched in retina | -2.885 | 0.04157 | 2.748 |
| ENSMUSG00000025269 | apurinic/apyrimidinic endonuclease 2 [Source:MGI Symbol;Acc:MGI:1924872] | Apex2 | Enriched in retina | -2.885 | 2.696E-08 | 107 |
| ENSMUSG00000030703 | glycerophosphodiester phosphodiesterase domain containing 3 [Source:MGI Symbol;Acc:MGI:1915866] | Gdpd3 | Enriched in retina | -2.89 | 0.0005375 | 6.428 |
| ENSMUSG00000021928 | emopamil binding protein-like [Source:MGI Symbol;Acc:MGI:1915427] | Ebpl | Enriched in retina | -2.907 | 2.75E-18 | 170.4 |
| ENSMUSG00000096883 | shisa family member 8 [Source:MGI Symbol;Acc:MGI:2146080] | Shisa8 | Enriched in retina | -3.079 | 2.001E-19 | 27.13 |
| ENSMUSG00000040489 | SRY (sex determining region Y)-box 30 [Source:MGI Symbol;Acc:MGI:1341157] | Sox30 | Enriched in retina | -3.096 | 0.0001817 | 11.69 |
| ENSMUSG00000058966 | TLC domain containing 3B [Source:MGI Symbol;Acc:MGI:1916202] | Tlcd3b | Enriched in retina | -3.101 | 1.928E-32 | 987.5 |
| ENSMUSG00000061462 | obscurin, cytoskeletal calmodulin and titin-interacting RhoGEF [Source:MGI Symbol;Acc:MGI:2681862] | Obscn | Enriched in retina | -3.15 | 2.624E-08 | 66.02 |
| ENSMUSG00000079418 | autophagy related 4A, cysteine peptidase [Source:MGI Symbol;Acc:MGI:2147903] | Atg4a | Enriched in retina | -3.156 | 0.00216 | 66.59 |
| ENSMUSG00000021013 | tetratricopeptide repeat domain 8 [Source:MGI Symbol;Acc:MGI:1923510] | Ttc8 | Enriched in retina | -3.176 | 5.081E-15 | 161.9 |
| ENSMUSG00000029304 | secreted phosphoprotein 1 [Source:MGI Symbol;Acc:MGI:98389] | Spp1 | Enriched in retina | -3.213 | 0.001663 | 33.23 |

| | | | | | | |
|---|---|---|---|---|---|---|
| ENSMUSG00000038208 | post-GPI attachment to proteins 3 [Source:MGI Symbol;Acc:MGI:2444461] | Pgap3 | Enriched in retina | -3.22 | 1.382E-13 | 48.45 |
| ENSMUSG00000050786 | coiled-coil domain containing 126 [Source:MGI Symbol;Acc:MGI:1889376] | Ccdc126 | Enriched in retina | -3.23 | 2.772E-28 | 193.5 |
| ENSMUSG00000051736 | family with sequence similarity 229, member B [Source:MGI Symbol;Acc:MGI:1913587] | Fam229b | Enriched in retina | -3.258 | 9.977E-07 | 21.72 |
| ENSMUSG00000060579 | fragile histidine triad gene [Source:MGI Symbol;Acc:MGI:1277947] | Fhit | Enriched in retina | -3.323 | 5.573E-11 | 135.3 |
| ENSMUSG00000038119 | cell adhesion molecule-related/down-regulated by oncogenes [Source:MGI Symbol;Acc:MGI:1926387] | Cdon | Enriched in retina | -3.334 | 3.715E-21 | 299.2 |
| ENSMUSG00000038515 | GH regulated TBC protein 1 [Source:MGI Symbol;Acc:MGI:1914040] | Grtp1 | Enriched in retina | -3.361 | 6.439E-18 | 106.5 |
| ENSMUSG00000021733 | solute carrier family 4, sodium bicarbonate cotransporter, member 7 [Source:MGI Symbol;Acc:MGI:2443878] | Slc4a7 | Enriched in retina | -3.363 | 3.184E-31 | 436.9 |
| ENSMUSG00000028248 | PNN interacting serine/arginine-rich [Source:MGI Symbol;Acc:MGI:1913875] | Pnisr | Enriched in retina | -3.376 | 6.514E-14 | 892.2 |
| ENSMUSG00000029276 | glomulin, FKBP associated protein [Source:MGI Symbol;Acc:MGI:2141180] | Glmn | Enriched in retina | -3.414 | 2.015E-13 | 122.5 |
| ENSMUSG00000028397 | lysine (K)-specific demethylase 4C [Source:MGI Symbol;Acc:MGI:1924054] | Kdm4c | Enriched in retina | -3.444 | 2.87E-26 | 443.3 |
| ENSMUSG00000028730 | cilia and flagella associated protein 57 [Source:MGI Symbol;Acc:MGI:2686209] | Cfap57 | Enriched in retina | -3.452 | 0.000004377 | 18.37 |
| ENSMUSG00000102349 | predicted gene, 37376 [Source:MGI Symbol;Acc:MGI:5610604] | Gm37376 | Enriched in retina | -3.483 | 0.0004328 | 32.91 |
| ENSMUSG00000031174 | retinitis pigmentosa GTPase regulator [Source:MGI Symbol;Acc:MGI:1344037] | Rpgr | Enriched in retina | -3.56 | 2.075E-11 | 194.1 |
| ENSMUSG00000039375 | WD repeat domain 17 [Source:MGI Symbol;Acc:MGI:1924662] | Wdr17 | Enriched in retina | -3.646 | 6.562E-25 | 263.9 |
| ENSMUSG00000041460 | calcium channel, voltage-dependent, alpha 2/delta subunit 4 [Source:MGI Symbol;Acc:MGI:2442632] | Cacna2d4 | Enriched in retina | -3.656 | 4.34E-26 | 641.5 |
| ENSMUSG00000080316 | sperm acrosome associated 6 [Source:MGI Symbol;Acc:MGI:1922452] | Spaca6 | Enriched in retina | -3.669 | 8.123E-11 | 101 |
| ENSMUSG00000044362 | coiled-coil domain containing 89 [Source:MGI Symbol;Acc:MGI:1917304] | Ccdc89 | Enriched in retina | -3.67 | 3.356E-09 | 17.87 |
| ENSMUSG00000002100 | myosin binding protein C, cardiac [Source:MGI Symbol;Acc:MGI:102844] | Mybpc3 | Enriched in retina | -3.686 | 0.001589 | 4.298 |
| ENSMUSG00000045103 | dystrophin, muscular dystrophy [Source:MGI Symbol;Acc:MGI:94909] | Dmd | Enriched in retina | -3.713 | 1.904E-16 | 1673 |
| ENSMUSG00000023484 | peripherin [Source:MGI Symbol;Acc:MGI:97774] | Prph | Enriched in retina | -3.744 | 0.00003602 | 64.1 |
| ENSMUSG00000093806 | acetylserotonin O-methyltransferase [Source:MGI Symbol;Acc:MGI:96090] | Asmt | Enriched in retina | -3.792 | 0.01378 | 3.293 |
| ENSMUSG00000045319 | proline and serine rich 2 [Source:MGI Symbol;Acc:MGI:2442238] | Proser2 | Enriched in retina | -3.809 | 1.107E-21 | 42.91 |
| ENSMUSG00000043639 | RNA binding motif protein 20 [Source:MGI Symbol;Acc:MGI:1920963] | Rbm20 | Enriched in retina | -3.816 | 7.148E-18 | 125.3 |
| ENSMUSG00000047632 | fibroblast growth factor binding protein 3 [Source:MGI Symbol;Acc:MGI:1919764] | Fgfbp3 | Enriched in retina | -3.844 | 4.823E-09 | 20.4 |
| ENSMUSG00000047986 | paralemmin 3 [Source:MGI Symbol;Acc:MGI:1921587] | Palm3 | Enriched in retina | -3.862 | 3.879E-07 | 25.27 |
| ENSMUSG00000037148 | Rho GTPase activating protein 10 [Source:MGI Symbol;Acc:MGI:1925764] | Arhgap10 | Enriched in retina | -3.894 | 2.965E-27 | 198.1 |
| ENSMUSG00000005233 | SPC25, NDC80 kinetochore complex component, homolog (S. cerevisiae) [Source:MGI Symbol;Acc:MGI:1913692] | Spc25 | Enriched in retina | -3.895 | 1.087E-11 | 60.05 |
| ENSMUSG00000046667 | RNA binding motif protein 12 B1 [Source:MGI Symbol;Acc:MGI:1919647] | Rbm12b1 | Enriched in retina | -3.906 | 4.154E-12 | 35.33 |
| ENSMUSG00000043727 | RIKEN cDNA F830045P16 gene [Source:MGI Symbol;Acc:MGI:3045317] | F830045P16Rik | Enriched in retina | -3.921 | 0.00001106 | 4.358 |
| ENSMUSG00000043541 | cancer susceptibility candidate 1 [Source:MGI Symbol;Acc:MGI:2444480] | Casc1 | Enriched in retina | -3.944 | 3.59E-08 | 13.93 |
| ENSMUSG00000034452 | solute carrier family 24 (sodium/potassium/calcium exchanger), member 1 [Source:MGI Symbol;Acc:MGI:2384871] | Slc24a1 | Enriched in retina | -3.998 | 0.00262 | 1018 |
| ENSMUSG00000096351 | sterile alpha motif domain containing 11 [Source:MGI Symbol;Acc:MGI:2446220] | Samd11 | Enriched in retina | -4.056 | 2.243E-16 | 401.5 |
| ENSMUSG00000037143 | cilia and flagella associated protein 61 [Source:MGI Symbol;Acc:MGI:1926024] | Cfap61 | Enriched in retina | -4.07 | 6.754E-22 | 108.6 |
| ENSMUSG00000027401 | transglutaminase 3, E polypeptide [Source:MGI Symbol;Acc:MGI:98732] | Tgm3 | Enriched in retina | -4.117 | 0.0003698 | 3.6 |
| ENSMUSG00000045776 | leucine-rich repeats and transmembrane domains 1 [Source:MGI Symbol;Acc:MGI:2442106] | Lrtm1 | Enriched in retina | -4.167 | 0.0004418 | 127.3 |
| ENSMUSG00000048337 | neuropeptide Y receptor Y4 [Source:MGI Symbol;Acc:MGI:105374] | Npy4r | Enriched in retina | -4.174 | 0.0356 | 1.332 |
| ENSMUSG00000042961 | EGF-like, fibronectin type III and laminin G domains [Source:MGI Symbol;Acc:MGI:2146149] | Egflam | Enriched in retina | -4.192 | 4.532E-48 | 370.2 |
| ENSMUSG00000063681 | crumbs family member 1, photoreceptor morphogenesis associated [Source:MGI Symbol;Acc:MGI:2136343] | Crb1 | Enriched in retina | -4.274 | 6.005E-13 | 382.8 |
| ENSMUSG00000051444 | Bardet-Biedl syndrome 12 (human) [Source:MGI Symbol;Acc:MGI:2686651] | Bbs12 | Enriched in retina | -4.347 | 1.427E-10 | 89.99 |
| ENSMUSG00000031789 | cyclic nucleotide gated channel beta 1 [Source:MGI Symbol;Acc:MGI:2664102] | Cngb1 | Enriched in retina | -4.362 | 0.000008313 | 733.4 |
| ENSMUSG00000045275 | Leber congenital amaurosis 5-like [Source:MGI Symbol;Acc:MGI:3041157] | Lca5l | Enriched in retina | -4.401 | 1.03E-13 | 37.83 |
| ENSMUSG00000034837 | G protein subunit alpha transducin 1 [Source:MGI Symbol;Acc:MGI:95778] | Gnat1 | Enriched in retina | -4.497 | 0.001828 | 1840 |
| ENSMUSG00000069372 | cortexin 3 [Source:MGI Symbol;Acc:MGI:3642816] | Ctxn3 | Enriched in retina | -4.501 | 2.329E-08 | 38.01 |
| ENSMUSG00000075256 | ceramide kinase-like [Source:MGI Symbol;Acc:MGI:3037816] | Cerkl | Enriched in retina | -4.578 | 7.563E-16 | 144.8 |
| ENSMUSG00000036602 | ALX homeobox 1 [Source:MGI Symbol;Acc:MGI:104621] | Alx1 | Enriched in retina | -4.608 | 0.0181 | 1.229 |
| ENSMUSG00000045493 | basic helix-loop-helix family, member e23 [Source:MGI Symbol;Acc:MGI:2153710] | Bhlhe23 | Enriched in retina | -4.675 | 2.216E-15 | 30.34 |
| ENSMUSG00000041193 | phospholipase A2, group V [Source:MGI Symbol;Acc:MGI:101899] | Pla2g5 | Enriched in retina | -4.687 | 2.01E-18 | 65.41 |
| ENSMUSG00000031738 | Iroquois homeobox 6 [Source:MGI Symbol;Acc:MGI:1927642] | Irx6 | Enriched in retina | -4.695 | 5.257E-17 | 50.97 |
| ENSMUSG00000041488 | syntaxin 3 [Source:MGI Symbol;Acc:MGI:103077] | Stx3 | Enriched in retina | -4.725 | 2.527E-34 | 591.3 |
| ENSMUSG00000032420 | 5' nucleotidase, ecto [Source:MGI Symbol;Acc:MGI:99782] | Nt5e | Enriched in retina | -4.731 | 1.059E-21 | 156.5 |
| ENSMUSG00000024227 | PDZ and pleckstrin homology domains 1 [Source:MGI Symbol;Acc:MGI:1916489] | Pdzph1 | Enriched in retina | -4.736 | 4.449E-08 | 255.2 |
| ENSMUSG00000031293 | retinoschisis (X-linked, juvenile) 1 (human) [Source:MGI Symbol;Acc:MGI:1336189] | Rs1 | Enriched in retina | -4.75 | 0.0001675 | 1028 |
| ENSMUSG00000072295 | C2 calcium dependent domain containing 6 [Source:MGI Symbol;Acc:MGI:1920713] | C2cd6 | Enriched in retina | -4.769 | 9.569E-07 | 7.861 |
| ENSMUSG00000035270 | interphotoreceptor matrix proteoglycan 2 [Source:MGI Symbol;Acc:MGI:3044955] | Impg2 | Enriched in retina | -4.84 | 4.271E-09 | 754.3 |
| ENSMUSG00000025916 | protein phosphatase 1, regulatory subunit 42 [Source:MGI Symbol;Acc:MGI:1921138] | Ppp1r42 | Enriched in retina | -4.84 | 8.335E-11 | 17.65 |
| ENSMUSG00000006007 | phosducin [Source:MGI Symbol;Acc:MGI:98090] | Pdc | Enriched in retina | -4.857 | 0.0003524 | 804 |
| ENSMUSG00000042282 | guanylate cyclase 2f [Source:MGI Symbol;Acc:MGI:105119] | Gucy2f | Enriched in retina | -4.873 | 1.352E-12 | 221.8 |
| ENSMUSG00000030324 | rhodopsin [Source:MGI Symbol;Acc:MGI:97914] | Rho | Enriched in retina | -4.875 | 0.000143 | 5636 |
| ENSMUSG00000041578 | cone-rod homeobox [Source:MGI Symbol;Acc:MGI:1194883] | Crx | Enriched in retina | -4.914 | 0.00004482 | 944.1 |
| ENSMUSG00000020890 | guanylate cyclase 2e [Source:MGI Symbol;Acc:MGI:105123] | Gucy2e | Enriched in retina | -4.918 | 2.112E-09 | 507.5 |
| ENSMUSG00000086962 | predicted gene 12248 [Source:MGI Symbol;Acc:MGI:3651124] | Gm12248 | Enriched in retina | -4.946 | 0.0001583 | 8.028 |
| ENSMUSG00000056055 | S-antigen, retina and pineal gland (arrestin) [Source:MGI Symbol;Acc:MGI:98227] | Sag | Enriched in retina | -4.959 | 0.000009172 | 2015 |
| ENSMUSG00000042073 | abhydrolase domain containing 14b [Source:MGI Symbol;Acc:MGI:1923741] | Abhd14b | Enriched in retina | -4.974 | 1.13E-15 | 38.48 |
| ENSMUSG00000108841 | FERM and PDZ domain containing 2 [Source:MGI Symbol;Acc:MGI:2685472] | Frmpd2 | Enriched in retina | -4.976 | 1.943E-23 | 339.6 |
| ENSMUSG00000004630 | Purkinje cell protein 2 (L7) [Source:MGI Symbol;Acc:MGI:97508] | Pcp2 | Enriched in retina | -4.993 | 6.649E-08 | 42.06 |
| ENSMUSG00000041044 | leucine-rich repeat, immunoglobulin-like and transmembrane domains 1 [Source:MGI Symbol;Acc:MGI:2385320] | Lrit1 | Enriched in retina | -5.03 | 0.00003685 | 386.5 |
| ENSMUSG00000021123 | retinol dehydrogenase 12 [Source:MGI Symbol;Acc:MGI:1925224] | Rdh12 | Enriched in retina | -5.039 | 4.269E-09 | 285 |
| ENSMUSG00000023978 | peripherin 2 [Source:MGI Symbol;Acc:MGI:102791] | Prph2 | Enriched in retina | -5.049 | 0.0001312 | 1752 |
| ENSMUSG00000048015 | neurogenic differentiation 4 [Source:MGI Symbol;Acc:MGI:108055] | Neurod4 | Enriched in retina | -5.053 | 1.033E-09 | 168.9 |
| ENSMUSG00000047298 | potassium channel, subfamily V, member 2 [Source:MGI Symbol;Acc:MGI:2670981] | Kcnv2 | Enriched in retina | -5.079 | 0.000005373 | 537.7 |
| ENSMUSG00000058743 | potassium inwardly-rectifying channel, subfamily J, member 14 [Source:MGI Symbol;Acc:MGI:2384820] | Kcnj14 | Enriched in retina | -5.087 | 4.322E-18 | 91.1 |
| ENSMUSG00000032292 | nuclear receptor subfamily 2, group E, member 3 [Source:MGI Symbol;Acc:MGI:1346317] | Nr2e3 | Enriched in retina | -5.099 | 0.00002774 | 617.6 |
| ENSMUSG00000029663 | guanine nucleotide binding protein (G protein), gamma transducing activity polypeptide 1 [Source:MGI Symbol;Acc:MGI:109165] | Gngt1 | Enriched in retina | -5.1 | 0.0001027 | 593.7 |
| ENSMUSG00000096740 | LBH domain containing 1 [Source:MGI Symbol;Acc:MGI:5516029] | Lbhd1 | Enriched in retina | -5.122 | 2.613E-07 | 104.6 |
| ENSMUSG00000023979 | guanylate cyclase activator 1B [Source:MGI Symbol;Acc:MGI:1194489] | Guca1b | Enriched in retina | -5.132 | 2.911E-07 | 220.2 |
| ENSMUSG00000024575 | phosphodiesterase 6A, cGMP-specific, rod, alpha [Source:MGI Symbol;Acc:MGI:97524] | Pde6a | Enriched in retina | -5.134 | 0.00006046 | 1301 |
| ENSMUSG00000031450 | G protein-coupled receptor kinase 1 [Source:MGI Symbol;Acc:MGI:1345146] | Grk1 | Enriched in retina | -5.135 | 0.0000637 | 1629 |
| ENSMUSG00000046049 | retinitis pigmentosa 1 homolog like 1 [Source:MGI Symbol;Acc:MGI:2384303] | Rp1l1 | Enriched in retina | -5.153 | 0.00003455 | 1013 |
| ENSMUSG00000025900 | retinitis pigmentosa 1 (human) [Source:MGI Symbol;Acc:MGI:1341105] | Rp1 | Enriched in retina | -5.155 | 0.000003459 | 3382 |
| ENSMUSG00000021384 | sushi domain containing 3 [Source:MGI Symbol;Acc:MGI:1913579] | Susd3 | Enriched in retina | -5.16 | 1.184E-17 | 69.76 |
| ENSMUSG00000041534 | retinol binding protein 3, interstitial [Source:MGI Symbol;Acc:MGI:97878] | Rbp3 | Enriched in retina | -5.184 | 0.00005268 | 3079 |
| ENSMUSG00000027530 | fatty acid binding protein 12 [Source:MGI Symbol;Acc:MGI:1922747] | Fabp12 | Enriched in retina | -5.19 | 8.322E-07 | 148.6 |
| ENSMUSG00000024519 | complexin 4 [Source:MGI Symbol;Acc:MGI:2685803] | Cplx4 | Enriched in retina | -5.231 | 0.00003325 | 325.1 |
| ENSMUSG00000054580 | phospholipase A2 receptor 1 [Source:MGI Symbol;Acc:MGI:102468] | Pla2r1 | Enriched in retina | -5.234 | 1.639E-10 | 229.2 |
| ENSMUSG00000049353 | retinal degeneration 3 [Source:MGI Symbol;Acc:MGI:1921273] | Rd3 | Enriched in retina | -5.235 | 1.238E-13 | 236.2 |
| ENSMUSG00000038115 | anoctamin 2 [Source:MGI Symbol;Acc:MGI:2387214] | Ano2 | Enriched in retina | -5.243 | 1.952E-19 | 281.6 |
| ENSMUSG00000069170 | adhesion G protein-coupled receptor V1 [Source:MGI Symbol;Acc:MGI:1274784] | Adgrv1 | Enriched in retina | -5.253 | 2.888E-24 | 786.2 |
| ENSMUSG00000034829 | nucleoredoxin-like 1 [Source:MGI Symbol;Acc:MGI:1924446] | Nxnl1 | Enriched in retina | -5.257 | 1.034E-07 | 200.2 |
| ENSMUSG00000005649 | calcium binding protein 5 [Source:MGI Symbol;Acc:MGI:1352746] | Cabp5 | Enriched in retina | -5.31 | 4.259E-17 | 50.22 |
| ENSMUSG00000097050 | predicted gene 9918 [Source:MGI Symbol;Acc:MGI:3646315] | Gm9918 | Enriched in retina | -5.324 | 3.181E-08 | 57.93 |
| ENSMUSG00000102900 | predicted gene, 37811 [Source:MGI Symbol;Acc:MGI:5611039] | Gm37811 | Enriched in retina | -5.326 | 7.508E-11 | 16.7 |
| ENSMUSG00000030523 | transient receptor potential cation channel, subfamily M, member 1 [Source:MGI Symbol;Acc:MGI:1330305] | Trpm1 | Enriched in retina | -5.345 | 0.000001944 | 420.9 |
| ENSMUSG00000036395 | galactosidase, beta 1-like 2 [Source:MGI Symbol;Acc:MGI:2388283] | Glb1l2 | Enriched in retina | -5.364 | 9.924E-18 | 81.64 |
| ENSMUSG00000075410 | photoreceptor disc component [Source:MGI Symbol;Acc:MGI:3649529] | Prcd | Enriched in retina | -5.371 | 1.217E-29 | 168.2 |
| ENSMUSG00000106887 | RIKEN cDNA A930005G22 gene [Source:MGI Symbol;Acc:MGI:3528178] | A930005G22Rik | Enriched in retina | -5.386 | 3.138E-07 | 34.64 |
| ENSMUSG00000056947 | mab-21-like 1 [Source:MGI Symbol;Acc:MGI:1333773] | Mab21l1 | Enriched in retina | -5.431 | 3.337E-08 | 403.6 |
| ENSMUSG00000053773 | retinol dehydrogenase 8 [Source:MGI Symbol;Acc:MGI:2685028] | Rdh8 | Enriched in retina | -5.434 | 2.452E-11 | 87.45 |
| ENSMUSG00000075330 | RIKEN cDNA A930003A15 gene [Source:MGI Symbol;Acc:MGI:1915412] | A930003A15Rik | Enriched in retina | -5.434 | 5.336E-16 | 40.93 |
| ENSMUSG00000029491 | phosphodiesterase 6B, cGMP, rod receptor, beta polypeptide [Source:MGI Symbol;Acc:MGI:97525] | Pde6b | Enriched in retina | -5.443 | 0.000007062 | 1205 |
| ENSMUSG00000029601 | IQ motif containing D [Source:MGI Symbol;Acc:MGI:1922982] | Iqcd | Enriched in retina | -5.447 | 3.036E-13 | 36.37 |
| ENSMUSG00000021803 | cadherin-related family member 1 [Source:MGI Symbol;Acc:MGI:2157782] | Cdhr1 | Enriched in retina | -5.471 | 1.932E-14 | 1092 |
| ENSMUSG00000032343 | interphotoreceptor matrix proteoglycan 1 [Source:MGI Symbol;Acc:MGI:1926876] | Impg1 | Enriched in retina | -5.475 | 3.048E-10 | 255.4 |
| ENSMUSG00000020907 | recoverin [Source:MGI Symbol;Acc:MGI:97883] | Rcvrn | Enriched in retina | -5.511 | 4.568E-09 | 323.5 |
| ENSMUSG00000057132 | retinitis pigmentosa GTPase regulator interacting protein 1 [Source:MGI Symbol;Acc:MGI:1932134] | Rpgrip1 | Enriched in retina | -5.527 | 3.463E-10 | 1656 |
| ENSMUSG00000067220 | cyclic nucleotide gated channel alpha 1 [Source:MGI Symbol;Acc:MGI:88436] | Cnga1 | Enriched in retina | -5.527 | 1.068E-12 | 478.2 |
| ENSMUSG00000070683 | lactamase, beta-like 1 [Source:MGI Symbol;Acc:MGI:2448566] | Lactbl1 | Enriched in retina | -5.534 | 0.000000914 | 222.9 |
| ENSMUSG00000043418 | leucine-rich repeat, immunoglobulin-like and transmembrane domains 2 [Source:MGI Symbol;Acc:MGI:2444885] | Lrit2 | Enriched in retina | -5.551 | 3.277E-07 | 108.6 |
| ENSMUSG00000035504 | receptor accessory protein 6 [Source:MGI Symbol;Acc:MGI:1917585] | Reep6 | Enriched in retina | -5.559 | 2.688E-17 | 1054 |
| ENSMUSG00000031142 | calcium channel, voltage-dependent, alpha 1F subunit [Source:MGI Symbol;Acc:MGI:1859639] | Cacna1f | Enriched in retina | -5.59 | 1.199E-07 | 357 |
| ENSMUSG00000021363 | male germ cell-associated kinase [Source:MGI Symbol;Acc:MGI:96913] | Mak | Enriched in retina | -5.591 | 2.051E-12 | 259.3 |
| ENSMUSG00000093865 | leucine-rich repeat, immunoglobulin-like and transmembrane domains 3 [Source:MGI Symbol;Acc:MGI:2685267] | Lrit3 | Enriched in retina | -5.621 | 0.000006097 | 120.3 |
| ENSMUSG00000051860 | sterile alpha motif domain containing 7 [Source:MGI Symbol;Acc:MGI:1923203] | Samd7 | Enriched in retina | -5.625 | 2.549E-10 | 192 |
| ENSMUSG00000026609 | usherin [Source:MGI Symbol;Acc:MGI:1341292] | Ush2a | Enriched in retina | -5.644 | 8.979E-12 | 1475 |
| ENSMUSG00000056043 | regulator of G-protein signalling 9 binding protein [Source:MGI Symbol;Acc:MGI:2384418] | Rgs9bp | Enriched in retina | -5.653 | 2.772E-09 | 666 |
| ENSMUSG00000047034 | ankyrin repeat domain 33 [Source:MGI Symbol;Acc:MGI:2443398] | Ankrd33 | Enriched in retina | -5.675 | 4.651E-10 | 105.8 |
| ENSMUSG00000049811 | family with sequence similarity 161, member A [Source:MGI Symbol;Acc:MGI:1921123] | Fam161a | Enriched in retina | -5.688 | 4.654E-22 | 823.6 |
| ENSMUSG00000086322 | RIKEN cDNA E130218I03 gene [Source:MGI Symbol;Acc:MGI:3528958] | E130218I03Rik | Enriched in retina | -5.709 | 3.175E-26 | 276.3 |
| ENSMUSG00000000617 | glutamate receptor, metabotropic 6 [Source:MGI Symbol;Acc:MGI:1351343] | Grm6 | Enriched in retina | -5.755 | 1.641E-08 | 178.8 |
| ENSMUSG00000073551 | serine peptidase inhibitor, Kazal type 13 [Source:MGI Symbol;Acc:MGI:3642511] | Spink13 | Enriched in retina | -5.777 | 1.81E-10 | 7.681 |
| ENSMUSG00000037446 | tubby like protein 1 [Source:MGI Symbol;Acc:MGI:109571] | Tulp1 | Enriched in retina | -5.779 | 5.362E-11 | 663.6 |
| ENSMUSG00000037161 | mitochondria localized glutamic acid rich protein [Source:MGI Symbol;Acc:MGI:1914999] | Mgarp | Enriched in retina | -5.786 | 7.193E-09 | 530.5 |
| ENSMUSG00000071648 | rod outer segment membrane protein 1 [Source:MGI Symbol;Acc:MGI:97998] | Rom1 | Enriched in retina | -5.787 | 2.18E-22 | 555.4 |
| ENSMUSG00000036480 | protease, serine 56 [Source:MGI Symbol;Acc:MGI:1916703] | Prss56 | Enriched in retina | -5.805 | 9.61E-08 | 103.2 |
| ENSMUSG00000110344 | small integral membrane protein 36 [Source:MGI Symbol;Acc:MGI:5804831] | Smim36 | Enriched in retina | -5.828 | 4.993E-10 | 107.5 |
| ENSMUSG00000040554 | aryl hydrocarbon receptor-interacting protein-like 1 [Source:MGI Symbol;Acc:MGI:2148800] | Aipl1 | Enriched in retina | -5.85 | 0.000001001 | 368.2 |
| ENSMUSG00000029410 | protein phosphatase, EF hand calcium-binding domain 2 [Source:MGI Symbol;Acc:MGI:1342304] | Ppef2 | Enriched in retina | -5.928 | 5.698E-22 | 246.6 |
| ENSMUSG00000024992 | phosphodiesterase 6C, cGMP specific, cone, alpha prime [Source:MGI Symbol;Acc:MGI:105956] | Pde6c | Enriched in retina | -5.93 | 3.407E-22 | 100.5 |
| ENSMUSG00000021396 | nucleoredoxin-like 2 [Source:MGI Symbol;Acc:MGI:1922374] | Nxnl2 | Enriched in retina | -5.931 | 1.568E-13 | 84.76 |

| | | | | | | |
|---|---|---|---|---|---|---|
| ENSMUSG00000035296 | sarcoglycan, gamma (dystrophin-associated glycoprotein) [Source:MGI Symbol;Acc:MGI:1346524] | Sgcg | Enriched in retina | -5.938 | 1.195E-11 | 9.668 |
| ENSMUSG00000040632 | neural retina leucine zipper gene [Source:MGI Symbol;Acc:MGI:102567] | Nrl | Enriched in retina | -5.948 | 1.222E-10 | 505.3 |
| ENSMUSG00000044375 | photoreceptor cilium actin regulator [Source:MGI Symbol;Acc:MGI:2385061] | Pcare | Enriched in retina | -6.038 | 1.139E-07 | 962.8 |
| ENSMUSG00000111755 | predicted gene, 47160 [Source:MGI Symbol;Acc:MGI:6095933] | Gm47160 | Enriched in retina | -6.071 | 0.000005881 | 97.35 |
| ENSMUSG00000021359 | transcription factor AP-2, alpha [Source:MGI Symbol;Acc:MGI:104671] | Tfap2a | Enriched in retina | -6.1 | 2.977E-14 | 139.8 |
| ENSMUSG00000108398 | predicted gene, 30191 [Source:MGI Symbol;Acc:MGI:5589350] | Gm30191 | Enriched in retina | -6.118 | 0.000003627 | 4.829 |
| ENSMUSG00000008932 | solute carrier family 1 (glutamate transporter), member 7 [Source:MGI Symbol;Acc:MGI:2444087] | Slc1a7 | Enriched in retina | -6.128 | 7.032E-13 | 185.9 |
| ENSMUSG00000024842 | calcium binding protein 4 [Source:MGI Symbol;Acc:MGI:1920910] | Cabp4 | Enriched in retina | -6.151 | 3.407E-13 | 95.11 |
| ENSMUSG00000026468 | LIM homeobox protein 4 [Source:MGI Symbol;Acc:MGI:101776] | Lhx4 | Enriched in retina | -6.261 | 1.038E-23 | 73.9 |
| ENSMUSG00000079550 | membrane protein, palmitoylated 4 (MAGUK p55 subfamily member 4) [Source:MGI Symbol;Acc:MGI:2386681] | Mpp4 | Enriched in retina | -6.333 | 4.888E-13 | 514.2 |
| ENSMUSG00000048617 | retbindin [Source:MGI Symbol;Acc:MGI:2443686] | Rtbdn | Enriched in retina | -6.384 | 3.543E-28 | 183 |
| ENSMUSG00000051985 | immunoglobulin-like and fibronectin type III domain containing 1 [Source:MGI Symbol;Acc:MGI:3045352] | Igfn1 | Enriched in retina | -6.439 | 2.944E-14 | 21.94 |
| ENSMUSG00000024518 | retina and anterior neural fold homeobox [Source:MGI Symbol;Acc:MGI:109632] | Rax | Enriched in retina | -6.468 | 8.227E-23 | 128.9 |
| ENSMUSG00000114882 | predicted gene, 48581 [Source:MGI Symbol;Acc:MGI:6098148] | Gm48581 | Enriched in retina | -6.507 | 4.821E-09 | 15.29 |
| ENSMUSG00000031688 | POU domain, class 4, transcription factor 2 [Source:MGI Symbol;Acc:MGI:102524] | Pou4f2 | Enriched in retina | -6.662 | 9.752E-14 | 45.2 |
| ENSMUSG00000025515 | mucin 2 [Source:MGI Symbol;Acc:MGI:1339364] | Muc2 | Enriched in retina | -6.743 | 9.963E-14 | 78.5 |
| ENSMUSG00000074037 | melanocortin 1 receptor [Source:MGI Symbol;Acc:MGI:99456] | Mc1r | Enriched in retina | -6.783 | 1.443E-10 | 11.83 |
| ENSMUSG00000049908 | gap junction protein, alpha 8 [Source:MGI Symbol;Acc:MGI:99953] | Gja8 | Enriched in retina | -6.835 | 0.00004436 | 178.2 |
| ENSMUSG00000060890 | arrestin 3, retinal [Source:MGI Symbol;Acc:MGI:2159617] | Arr3 | Enriched in retina | -7.014 | 1.58E-12 | 38.76 |
| ENSMUSG00000042240 | crystallin, beta B2 [Source:MGI Symbol;Acc:MGI:88519] | Crybb2 | Enriched in retina | -7.538 | 1.18E-08 | 184.8 |
| ENSMUSG00000025389 | major intrinsic protein of lens fiber [Source:MGI Symbol;Acc:MGI:96990] | Mip | Enriched in retina | -9.371 | 0.00002441 | 36.23 |
| ENSMUSG00000070870 | crystallin, gamma E [Source:MGI Symbol;Acc:MGI:88525] | Cryge | Enriched in retina | -9.74 | 0.000004142 | 45.26 |

**Contrast numerator: P15_Het_dLGN. Contrast denominator: P15_Het_SCN.**

| ensembl_gene_id | description | mgi_symbol | Significance | deseq_logfc | deseq_adjp | deseq_basemean |
|---|---|---|---|---|---|---|
| ENSMUSG00000029054 | gamma-aminobutyric acid (GABA) A receptor, subunit delta [Source:MGI Symbol;Acc:MGI:95622] | Gabrd | Enriched in dLGN | 6.433 | 9.133E-20 | 52.02 |
| ENSMUSG00000022468 | endonuclease, polyU-specific [Source:MGI Symbol;Acc:MGI:97746] | Endou | Enriched in dLGN | 6.137 | 6.294E-15 | 34.93 |
| ENSMUSG00000032532 | cholecystokinin [Source:MGI Symbol;Acc:MGI:88297] | Cck | Enriched in dLGN | 5.839 | 1.059E-29 | 445.5 |
| ENSMUSG00000021948 | protein kinase C, delta [Source:MGI Symbol;Acc:MGI:97598] | Prkcd | Enriched in dLGN | 5.615 | 3.634E-32 | 727.9 |
| ENSMUSG00000064179 | troponin T1, skeletal, slow [Source:MGI Symbol;Acc:MGI:1333868] | Tnnt1 | Enriched in dLGN | 5.614 | 1.635E-37 | 308.7 |
| ENSMUSG00000039783 | kynurenine 3-monooxygenase (kynurenine 3-hydroxylase) [Source:MGI Symbol;Acc:MGI:2138151] | Kmo | Enriched in dLGN | 5.551 | 0.00001343 | 4.13 |
| ENSMUSG00000110711 | predicted gene 45760 [Source:MGI Symbol;Acc:MGI:5804875] | Gm45760 | Enriched in dLGN | 5.021 | 0.0003321 | 2.941 |
| ENSMUSG00000033044 | dehydrogenase/reductase (SDR family) member 7C [Source:MGI Symbol;Acc:MGI:1915710] | Dhrs7c | Enriched in dLGN | 4.826 | 0.001052 | 6.079 |
| ENSMUSG00000030785 | cytochrome c oxidase subunit 6A2 [Source:MGI Symbol;Acc:MGI:104649] | Cox6a2 | Enriched in dLGN | 4.558 | 8.91E-08 | 11.74 |
| ENSMUSG00000050315 | synaptopodin 2 [Source:MGI Symbol;Acc:MGI:2153070] | Synpo2 | Enriched in dLGN | 4.488 | 5.088E-46 | 581.4 |
| ENSMUSG00000004951 | heat shock protein 1 [Source:MGI Symbol;Acc:MGI:96240] | Hspb1 | Enriched in dLGN | 4.432 | 0.000001437 | 22.09 |
| ENSMUSG00000025172 | ankyrin repeat domain 2 (stretch responsive muscle) [Source:MGI Symbol;Acc:MGI:1861447] | Ankrd2 | Enriched in dLGN | 4.362 | 0.001506 | 4.542 |
| ENSMUSG00000026688 | microsomal glutathione S-transferase 3 [Source:MGI Symbol;Acc:MGI:1913697] | Mgst3 | Enriched in dLGN | 4.29 | 5.377E-16 | 88.46 |
| ENSMUSG00000028024 | glutamyl aminopeptidase [Source:MGI Symbol;Acc:MGI:106645] | Enpep | Enriched in dLGN | 4.274 | 2.801E-07 | 10.7 |
| ENSMUSG00000091402 | retinal degeneration 3-like [Source:MGI Symbol;Acc:MGI:2675860] | Rd3l | Enriched in dLGN | 4.242 | 0.001311 | 41.89 |
| ENSMUSG00000018470 | potassium voltage-gated channel, shaker-related subfamily, beta member 3 [Source:MGI Symbol;Acc:MGI:1336208] | Kcnab3 | Enriched in dLGN | 4.175 | 1.125E-19 | 53.86 |
| ENSMUSG00000032776 | multiple C2 domains, transmembrane 2 [Source:MGI Symbol;Acc:MGI:2685335] | Mctp2 | Enriched in dLGN | 4.164 | 2.378E-11 | 28.35 |
| ENSMUSG00000020953 | cochlin [Source:MGI Symbol;Acc:MGI:1278313] | Coch | Enriched in dLGN | 4.058 | 3.513E-14 | 80.94 |
| ENSMUSG00000026525 | opsin 3 [Source:MGI Symbol;Acc:MGI:1338022] | Opn3 | Enriched in dLGN | 4.045 | 2.338E-12 | 17.62 |
| ENSMUSG00000031994 | a disintegrin-like and metallopeptidase (reprolysin type) with thrombospondin type 1 motif, 8 [Source:MGI Symbol;Acc:MGI:1353468] | Adamts8 | Enriched in dLGN | 4.021 | 5.113E-15 | 52.93 |
| ENSMUSG00000025366 | extended synaptotagmin-like protein 1 [Source:MGI Symbol;Acc:MGI:1344426] | Esyt1 | Enriched in dLGN | 3.99 | 2.098E-19 | 159.4 |
| ENSMUSG00000078816 | protein kinase C, gamma [Source:MGI Symbol;Acc:MGI:97597] | Prkcg | Enriched in dLGN | 3.87 | 1.017E-28 | 571.8 |
| ENSMUSG00000055027 | SET and MYND domain containing 1 [Source:MGI Symbol;Acc:MGI:104790] | Smyd1 | Enriched in dLGN | 3.748 | 5.134E-14 | 65.99 |
| ENSMUSG00000033615 | complexin 1 [Source:MGI Symbol;Acc:MGI:104727] | Cplx1 | Enriched in dLGN | 3.744 | 7.879E-28 | 2304 |
| ENSMUSG00000045287 | reticulon 4 receptor-like 1 [Source:MGI Symbol;Acc:MGI:2661375] | Rtn4rl1 | Enriched in dLGN | 3.704 | 2.854E-40 | 303.8 |
| ENSMUSG00000003541 | immediate early response 3 [Source:MGI Symbol;Acc:MGI:104814] | Ier3 | Enriched in dLGN | 3.688 | 0.000004546 | 21.94 |
| ENSMUSG00000068696 | G-protein coupled receptor 88 [Source:MGI Symbol;Acc:MGI:1927653] | Gpr88 | Enriched in dLGN | 3.661 | 3.859E-27 | 66.64 |
| ENSMUSG00000029641 | RAS-like, family 11, member A [Source:MGI Symbol;Acc:MGI:1916145] | Rasl11a | Enriched in dLGN | 3.601 | 3.655E-09 | 21.43 |
| ENSMUSG00000038264 | sema domain, immunoglobulin domain (Ig), and GPI membrane anchor, (semaphorin) 7A [Source:MGI Symbol;Acc:MGI:1306826] | Sema7a | Enriched in dLGN | 3.574 | 1.906E-33 | 229.8 |
| ENSMUSG00000027347 | RAS guanyl releasing protein 1 [Source:MGI Symbol;Acc:MGI:1314635] | Rasgrp1 | Enriched in dLGN | 3.552 | 1.825E-20 | 353.4 |
| ENSMUSG00000029211 | gamma-aminobutyric acid (GABA) A receptor, subunit alpha 4 [Source:MGI Symbol;Acc:MGI:95616] | Gabra4 | Enriched in dLGN | 3.52 | 1.276E-23 | 154.6 |
| ENSMUSG00000035506 | solute carrier family 12 (potassium/chloride transporters), member 8 [Source:MGI Symbol;Acc:MGI:2443672] | Slc12a8 | Enriched in dLGN | 3.502 | 9.542E-07 | 17.24 |
| ENSMUSG00000030340 | sodium channel, nonvoltage-gated 1 alpha [Source:MGI Symbol;Acc:MGI:101782] | Scnn1a | Enriched in dLGN | 3.462 | 4.951E-15 | 44.62 |
| ENSMUSG00000054196 | collagen triple helix repeat containing 1 [Source:MGI Symbol;Acc:MGI:1915838] | Cthrc1 | Enriched in dLGN | 3.456 | 5.008E-14 | 38 |
| ENSMUSG00000031231 | cytochrome c oxidase subunit 7B [Source:MGI Symbol;Acc:MGI:1913392] | Cox7b | Enriched in dLGN | 3.438 | 2.963E-11 | 113.9 |
| ENSMUSG00000040624 | pleckstrin homology domain containing, family G (with RhoGef domain) member 1 [Source:MGI Symbol;Acc:MGI:2676551] | Plekhg1 | Enriched in dLGN | 3.361 | 1.657E-35 | 465 |
| ENSMUSG00000059857 | netrin G1 [Source:MGI Symbol;Acc:MGI:1934028] | Ntng1 | Enriched in dLGN | 3.301 | 4.529E-16 | 904 |
| ENSMUSG00000023905 | tumor necrosis factor receptor superfamily, member 12a [Source:MGI Symbol;Acc:MGI:1351484] | Tnfrsf12a | Enriched in dLGN | 3.292 | 0.0000051 | 7.873 |
| ENSMUSG00000089717 | olfactory receptor family 13 subfamily F member 5 [Source:MGI Symbol;Acc:MGI:3030109] | Or13f5 | Enriched in dLGN | 3.244 | 0.002305 | 4.484 |
| ENSMUSG00000029632 | Ndufa4, mitochondrial complex associated [Source:MGI Symbol;Acc:MGI:107686] | Ndufa4 | Enriched in dLGN | 3.24 | 1.391E-21 | 183.2 |
| ENSMUSG00000026235 | Eph receptor A4 [Source:MGI Symbol;Acc:MGI:98277] | Epha4 | Enriched in dLGN | 3.23 | 4.833E-32 | 500.9 |
| ENSMUSG00000029659 | mesenteric estrogen dependent adipogenesis [Source:MGI Symbol;Acc:MGI:1917967] | Medag | Enriched in dLGN | 3.222 | 1.308E-10 | 25.4 |
| ENSMUSG00000039629 | striatin interacting protein 2 [Source:MGI Symbol;Acc:MGI:2444363] | Strip2 | Enriched in dLGN | 3.212 | 3.219E-18 | 138 |
| ENSMUSG00000027716 | transient receptor potential cation channel, subfamily C, member 3 [Source:MGI Symbol;Acc:MGI:109526] | Trpc3 | Enriched in dLGN | 3.141 | 1.659E-25 | 212.4 |
| ENSMUSG00000032578 | cytokine inducible SH2-containing protein [Source:MGI Symbol;Acc:MGI:103159] | Cish | Enriched in dLGN | 3.123 | 0.000003443 | 23.3 |
| ENSMUSG00000071866 | peptidylprolyl isomerase A [Source:MGI Symbol;Acc:MGI:97749] | Ppia | Enriched in dLGN | 3.118 | 5.851E-09 | 211.2 |
| ENSMUSG00000045625 | phosphatidylinositol glycan anchor biosynthesis, class Z [Source:MGI Symbol;Acc:MGI:2443822] | Pigz | Enriched in dLGN | 3.106 | 4.699E-07 | 18.26 |
| ENSMUSG00000016319 | solute carrier family 25 (mitochondrial carrier, adenine nucleotide translocator), member 5 [Source:MGI Symbol;Acc:MGI:1353496] | Slc25a5 | Enriched in dLGN | 3.103 | 8.123E-15 | 198.8 |
| ENSMUSG00000032908 | sphingosine-1-phosphate phosphatase 2 [Source:MGI Symbol;Acc:MGI:3589109] | Sgpp2 | Enriched in dLGN | 3.079 | 1.374E-15 | 115.4 |
| ENSMUSG00000014313 | cytochrome c oxidase subunit 6C [Source:MGI Symbol;Acc:MGI:104614] | Cox6c | Enriched in dLGN | 3.034 | 1.783E-18 | 345.4 |
| ENSMUSG00000027556 | carbonic anhydrase 1 [Source:MGI Symbol;Acc:MGI:88268] | Car1 | Enriched in dLGN | 3.025 | 0.000001711 | 12.27 |
| ENSMUSG00000062070 | phosphoglycerate kinase 1 [Source:MGI Symbol;Acc:MGI:97555] | Pgk1 | Enriched in dLGN | 3.018 | 2.65E-23 | 300 |
| ENSMUSG00000020733 | solute carrier family 9 (sodium/hydrogen exchanger), member 3 regulator 1 [Source:MGI Symbol;Acc:MGI:1349482] | Slc9a3r1 | Enriched in dLGN | 3.017 | 6.916E-36 | 214.9 |
| ENSMUSG00000028082 | SH3 domain protein D19 [Source:MGI Symbol;Acc:MGI:1350923] | Sh3d19 | Enriched in dLGN | 3.014 | 8.979E-19 | 657.1 |
| ENSMUSG00000065968 | interferon induced transmembrane protein 7 [Source:MGI Symbol;Acc:MGI:1921732] | Ifitm7 | Enriched in dLGN | 3.014 | 0.000001375 | 15.26 |
| ENSMUSG00000039115 | integrin alpha 9 [Source:MGI Symbol;Acc:MGI:104756] | Itga9 | Enriched in dLGN | 3.003 | 0.0004832 | 1811 |
| ENSMUSG00000042353 | Fras1 related extracellular matrix protein 3 [Source:MGI Symbol;Acc:MGI:2685641] | Frem3 | Enriched in dLGN | 2.958 | 2.735E-11 | 57.69 |
| ENSMUSG00000020059 | synaptonemal complex protein 3 [Source:MGI Symbol;Acc:MGI:109542] | Sycp3 | Enriched in dLGN | 2.926 | 0.0441 | 2.11 |
| ENSMUSG00000030500 | solute carrier family 17 (sodium-dependent inorganic phosphate cotransporter), member 6 [Source:MGI Symbol;Acc:MGI:2156052] | Slc17a6 | Enriched in dLGN | 2.911 | 4.334E-12 | 605.1 |
| ENSMUSG00000022820 | NADH:ubiquinone oxidoreductase subunit B4 [Source:MGI Symbol;Acc:MGI:1915444] | Ndufb4 | Enriched in dLGN | 2.907 | 0.00009944 | 32.87 |
| ENSMUSG00000027489 | N-terminal EF-hand calcium binding protein 3 [Source:MGI Symbol;Acc:MGI:1861721] | Necab3 | Enriched in dLGN | 2.899 | 3.294E-14 | 36.72 |
| ENSMUSG00000028718 | Scl/Tal1 interrupting locus [Source:MGI Symbol;Acc:MGI:107477] | Stil | Enriched in dLGN | 2.884 | 0.008243 | 5.074 |
| ENSMUSG00000040713 | cellular repressor of E1A-stimulated genes 1 [Source:MGI Symbol;Acc:MGI:1344382] | Creg1 | Enriched in dLGN | 2.874 | 7.782E-18 | 76.45 |
| ENSMUSG00000058927 | predicted gene 10053 [Source:MGI Symbol;Acc:MGI:3704493] | Gm10053 | Enriched in dLGN | 2.833 | 0.000001398 | 16.38 |
| ENSMUSG00000029153 | OCIA domain containing 2 [Source:MGI Symbol;Acc:MGI:1916377] | Ociad2 | Enriched in dLGN | 2.824 | 2.671E-17 | 70.58 |
| ENSMUSG00000032892 | RAN guanine nucleotide release factor [Source:MGI Symbol;Acc:MGI:1889073] | Rangrf | Enriched in dLGN | 2.812 | 0.004007 | 6.22 |
| ENSMUSG00000024064 | polypeptide N-acetylgalactosaminyltransferase 14 [Source:MGI Symbol;Acc:MGI:1918935] | Galnt14 | Enriched in dLGN | 2.803 | 3.832E-14 | 190 |
| ENSMUSG00000028773 | fatty acid binding protein 3, muscle and heart [Source:MGI Symbol;Acc:MGI:95476] | Fabp3 | Enriched in dLGN | 2.793 | 2.097E-07 | 75.72 |
| ENSMUSG00000019762 | iodotyrosine deiodinase [Source:MGI Symbol;Acc:MGI:1917587] | Iyd | Enriched in dLGN | 2.792 | 0.001573 | 6.399 |
| ENSMUSG00000038128 | calcium/calmodulin-dependent protein kinase IV [Source:MGI Symbol;Acc:MGI:88258] | Camk4 | Enriched in dLGN | 2.773 | 2.398E-30 | 937 |
| ENSMUSG00000027131 | ER membrane protein complex subunit 4 [Source:MGI Symbol;Acc:MGI:1915282] | Emc4 | Enriched in dLGN | 2.769 | 1.849E-15 | 81.85 |
| ENSMUSG00000020838 | solute carrier family 6 (neurotransmitter transporter, serotonin), member 4 [Source:MGI Symbol;Acc:MGI:96285] | Slc6a4 | Enriched in dLGN | 2.726 | 0.03214 | 23.26 |
| ENSMUSG00000053025 | synaptic vesicle glycoprotein 2 b [Source:MGI Symbol;Acc:MGI:1927338] | Sv2b | Enriched in dLGN | 2.711 | 8.892E-32 | 914.9 |
| ENSMUSG00000029769 | coiled-coil domain containing 136 [Source:MGI Symbol;Acc:MGI:1918128] | Ccdc136 | Enriched in dLGN | 2.689 | 2.569E-36 | 1034 |
| ENSMUSG00000021259 | cytochrome P450, family 46, subfamily a, polypeptide 1 [Source:MGI Symbol;Acc:MGI:1341877] | Cyp46a1 | Enriched in dLGN | 2.668 | 1.635E-32 | 248.2 |
| ENSMUSG00000027533 | fatty acid binding protein 5, epidermal [Source:MGI Symbol;Acc:MGI:101790] | Fabp5 | Enriched in dLGN | 2.668 | 0.000004529 | 258.1 |
| ENSMUSG00000028149 | RAP1, GTP-GDP dissociation stimulator 1 [Source:MGI Symbol;Acc:MGI:2385189] | Rap1gds1 | Enriched in dLGN | 2.661 | 1.655E-38 | 847.3 |
| ENSMUSG00000091387 | glucosaminyl (N-acetyl) transferase 4, core 2 (beta-1,6-N-acetylglucosaminyltransferase) [Source:MGI Symbol;Acc:MGI:2684919] | Gcnt4 | Enriched in dLGN | 2.645 | 0.00001023 | 28.71 |
| ENSMUSG00000033307 | macrophage migration inhibitory factor (glycosylation-inhibiting factor) [Source:MGI Symbol;Acc:MGI:96982] | Mif | Enriched in dLGN | 2.639 | 8.674E-09 | 90.39 |
| ENSMUSG00000032011 | thymus cell antigen 1, theta [Source:MGI Symbol;Acc:MGI:98747] | Thy1 | Enriched in dLGN | 2.609 | 2.928E-15 | 952.8 |
| ENSMUSG00000036578 | FXYD domain-containing ion transport regulator 7 [Source:MGI Symbol;Acc:MGI:1889006] | Fxyd7 | Enriched in dLGN | 2.607 | 3.341E-12 | 140.4 |
| ENSMUSG00000028833 | neurochondrin [Source:MGI Symbol;Acc:MGI:1347351] | Ncdn | Enriched in dLGN | 2.6 | 3.86E-19 | 1887 |
| ENSMUSG00000021520 | ubiquinol-cytochrome c reductase binding protein [Source:MGI Symbol;Acc:MGI:1914780] | Uqcrb | Enriched in dLGN | 2.595 | 5.312E-09 | 49.09 |
| ENSMUSG00000040723 | RCSD domain containing 1 [Source:MGI Symbol;Acc:MGI:2676394] | Rcsd1 | Enriched in dLGN | 2.589 | 1.772E-07 | 27.5 |
| ENSMUSG00000019943 | ATPase, Ca++ transporting, plasma membrane 1 [Source:MGI Symbol;Acc:MGI:104653] | Atp2b1 | Enriched in dLGN | 2.544 | 5.088E-46 | 2706 |
| ENSMUSG00000061859 | PATJ, crumbs cell polarity complex component [Source:MGI Symbol;Acc:MGI:1277960] | Patj | Enriched in dLGN | 2.543 | 1.113E-11 | 609.6 |
| ENSMUSG00000070803 | Cbp/p300-interacting transactivator, with Glu/Asp-rich carboxy-terminal domain, 4 [Source:MGI Symbol;Acc:MGI:1861694] | Cited4 | Enriched in dLGN | 2.543 | 0.0009066 | 9.191 |
| ENSMUSG00000020116 | partner of NOB1 homolog [Source:MGI Symbol;Acc:MGI:1913499] | Pno1 | Enriched in dLGN | 2.519 | 1.007E-11 | 43.22 |
| ENSMUSG00000000088 | cytochrome c oxidase subunit 5A [Source:MGI Symbol;Acc:MGI:88474] | Cox5a | Enriched in dLGN | 2.513 | 4.515E-14 | 112.1 |
| ENSMUSG00000021290 | ATP synthase membrane subunit 6.8PL [Source:MGI Symbol;Acc:MGI:1917507] | Atp5mpl | Enriched in dLGN | 2.511 | 9.02E-09 | 181.4 |
| ENSMUSG00000018821 | arginine vasopressin-induced 1 [Source:MGI Symbol;Acc:MGI:1916784] | Avpi1 | Enriched in dLGN | 2.51 | 1.25E-11 | 34.89 |
| ENSMUSG00000028251 | thiosulfate sulfurtransferase (rhodanese)-like domain containing 3 [Source:MGI Symbol;Acc:MGI:1924282] | Tstd3 | Enriched in dLGN | 2.503 | 6.306E-09 | 35.11 |
| ENSMUSG00000036330 | solute carrier family 18 (vesicular monoamine), member 1 [Source:MGI Symbol;Acc:MGI:106684] | Slc18a1 | Enriched in dLGN | 2.502 | 0.004339 | 5.605 |
| ENSMUSG00000038717 | ATP synthase, H+ transporting, mitochondrial F0 complex, subunit G [Source:MGI Symbol;Acc:MGI:1351597] | Atp5l | Enriched in dLGN | 2.486 | 3.24E-09 | 38.82 |
| ENSMUSG00000020321 | malate dehydrogenase 1, NAD (soluble) [Source:MGI Symbol;Acc:MGI:97051] | Mdh1 | Enriched in dLGN | 2.483 | 2.271E-09 | 617.5 |
| ENSMUSG00000022180 | solute carrier family 7 (cationic amino acid transporter, y+ system), member 8 [Source:MGI Symbol;Acc:MGI:1355323] | Slc7a8 | Enriched in dLGN | 2.47 | 2.323E-19 | 249.8 |
| ENSMUSG00000060780 | leucine rich repeat transmembrane neuronal 1 [Source:MGI Symbol;Acc:MGI:2389173] | Lrrtm1 | Enriched in dLGN | 2.469 | 3.815E-29 | 250 |
| ENSMUSG00000028488 | SH3-domain GRB2-like 2 [Source:MGI Symbol;Acc:MGI:700009] | Sh3gl2 | Enriched in dLGN | 2.463 | 1.308E-15 | 586.9 |
| ENSMUSG00000051022 | heparan sulfate (glucosamine) 3-O-sulfotransferase 1 [Source:MGI Symbol;Acc:MGI:1201606] | Hs3st1 | Enriched in dLGN | 2.461 | 1.113E-18 | 132 |
| ENSMUSG00000006057 | ATP synthase, H+ transporting, mitochondrial F0 complex, subunit C1 (subunit 9) [Source:MGI Symbol;Acc:MGI:107653] | Atp5g1 | Enriched in dLGN | 2.457 | 2.644E-12 | 76.53 |
| ENSMUSG00000017417 | plexin domain containing 1 [Source:MGI Symbol;Acc:MGI:1919574] | Plxdc1 | Enriched in dLGN | 2.454 | 2.655E-14 | 77.27 |
| ENSMUSG00000078566 | BCL2/adenovirus E1B interacting protein 3 [Source:MGI Symbol;Acc:MGI:109326] | Bnip3 | Enriched in dLGN | 2.427 | 3.875E-09 | 111.7 |
| ENSMUSG00000028648 | NADH:ubiquinone oxidoreductase core subunit S5 [Source:MGI Symbol;Acc:MGI:1890889] | Ndufs5 | Enriched in dLGN | 2.415 | 8.615E-12 | 101.9 |
| ENSMUSG00000014243 | zinc finger SWIM-type containing 7 [Source:MGI Symbol;Acc:MGI:1916997] | Zswim7 | Enriched in dLGN | 2.415 | 0.000001731 | 26.22 |
| ENSMUSG00000025092 | heat shock protein 12A [Source:MGI Symbol;Acc:MGI:1920692] | Hspa12a | Enriched in dLGN | 2.407 | 6.8E-26 | 913 |
| ENSMUSG00000021591 | glutaredoxin [Source:MGI Symbol;Acc:MGI:2135625] | Glrx | Enriched in dLGN | 2.407 | 2.634E-13 | 37.82 |
| ENSMUSG00000028334 | N-acetylneuraminic acid synthase (sialic acid synthase) [Source:MGI Symbol;Acc:MGI:2149820] | Nans | Enriched in dLGN | 2.4 | 5.49E-12 | 106.3 |
| ENSMUSG00000022454 | NEL-like 2 [Source:MGI Symbol;Acc:MGI:1858510] | Nell2 | Enriched in dLGN | 2.395 | 1.34E-20 | 1350 |
| ENSMUSG00000024190 | dual specificity phosphatase 1 [Source:MGI Symbol;Acc:MGI:105120] | Dusp1 | Enriched in dLGN | 2.39 | 2.186E-18 | 44.73 |
| ENSMUSG00000027076 | translocase of inner mitochondrial membrane 10 [Source:MGI Symbol;Acc:MGI:1353429] | Timm10 | Enriched in dLGN | 2.379 | 1.149E-08 | 29.72 |
| ENSMUSG00000038421 | Fc receptor-like A [Source:MGI Symbol;Acc:MGI:2138647] | Fcrla | Enriched in dLGN | 2.367 | 0.0008755 | 28.16 |
| ENSMUSG00000096257 | coiled-coil glutamate-rich protein 2 [Source:MGI Symbol;Acc:MGI:3645242] | Ccer2 | Enriched in dLGN | 2.354 | 0.00004456 | 12.8 |
| ENSMUSG00000062937 | methylthioadenosine phosphorylase [Source:MGI Symbol;Acc:MGI:1914152] | Mtap | Enriched in dLGN | 2.345 | 5.204E-07 | 29.38 |
| ENSMUSG00000004849 | adaptor protein complex AP-1, sigma 1 [Source:MGI Symbol;Acc:MGI:1098244] | Ap1s1 | Enriched in dLGN | 2.335 | 6.605E-19 | 288.6 |
| ENSMUSG00000041697 | cytochrome c oxidase subunit 6A1 [Source:MGI Symbol;Acc:MGI:103099] | Cox6a1 | Enriched in dLGN | 2.329 | 2.046E-13 | 168 |
| ENSMUSG00000036158 | prickle planar cell polarity protein 1 [Source:MGI Symbol;Acc:MGI:1916034] | Prickle1 | Enriched in dLGN | 2.326 | 7.075E-20 | 385.1 |
| ENSMUSG00000073838 | Tu translation elongation factor, mitochondrial [Source:MGI Symbol;Acc:MGI:1923686] | Tufm | Enriched in dLGN | 2.317 | 1.285E-11 | 100.2 |
| ENSMUSG00000016252 | ATP synthase, H+ transporting, mitochondrial F1 complex, epsilon subunit [Source:MGI Symbol;Acc:MGI:1855697] | Atp5e | Enriched in dLGN | 2.314 | 9.269E-11 | 153.2 |
| ENSMUSG00000059040 | enolase 1B, retrotransposed [Source:MGI Symbol;Acc:MGI:3648653] | Eno1b | Enriched in dLGN | 2.308 | 0.000005365 | 124.1 |
| ENSMUSG00000033316 | polypeptide N-acetylgalactosaminyltransferase 9 [Source:MGI Symbol;Acc:MGI:2677965] | Galnt9 | Enriched in dLGN | 2.306 | 5.507E-20 | 204.6 |
| ENSMUSG00000029189 | sel-1 suppressor of lin-12-like 3 (C. elegans) [Source:MGI Symbol;Acc:MGI:1916941] | Sel1l3 | Enriched in dLGN | 2.305 | 4.077E-15 | 154 |
| ENSMUSG00000025393 | ATP synthase, H+ transporting mitochondrial F1 complex, beta subunit [Source:MGI Symbol;Acc:MGI:107801] | Atp5b | Enriched in dLGN | 2.298 | 5.542E-13 | 1784 |
| ENSMUSG00000033998 | potassium channel, subfamily K, member 1 [Source:MGI Symbol;Acc:MGI:109322] | Kcnk1 | Enriched in dLGN | 2.283 | 2.596E-13 | 107.5 |
| ENSMUSG00000030089 | solute carrier family 41, member 3 [Source:MGI Symbol;Acc:MGI:1918949] | Slc41a3 | Enriched in dLGN | 2.277 | 2.358E-09 | 52.15 |
| ENSMUSG00000054843 | attractin like 1 [Source:MGI Symbol;Acc:MGI:2147749] | Atrnl1 | Enriched in dLGN | 2.274 | 2.168E-14 | 629.5 |
| ENSMUSG00000044252 | oxysterol binding protein-like 1A [Source:MGI Symbol;Acc:MGI:1927551] | Osbpl1a | Enriched in dLGN | 2.261 | 1.527E-21 | 434.2 |

| | | | | | | |
|---|---|---|---|---|---|---|
| ENSMUSG00000059734 | NADH:ubiquinone oxidoreductase core subunit S8 [Source:MGI Symbol;Acc:MGI:2385079] | Ndufs8 | Enriched in dLGN | 2.257 | 9.501E-10 | 73.63 |
| ENSMUSG00000028367 | thioredoxin 1 [Source:MGI Symbol;Acc:MGI:98874] | Txn1 | Enriched in dLGN | 2.253 | 4.51E-12 | 92.9 |
| ENSMUSG00000113149 | predicted gene, 49383 [Source:MGI Symbol;Acc:MGI:6121605] | Gm49383 | Enriched in dLGN | 2.252 | 0.007884 | 6.916 |
| ENSMUSG00000028710 | ATP synthase mitochondrial F1 complex assembly factor 1 [Source:MGI Symbol;Acc:MGI:2180560] | Atpaf1 | Enriched in dLGN | 2.25 | 3.931E-18 | 89.62 |
| ENSMUSG00000020431 | adenylate cyclase 1 [Source:MGI Symbol;Acc:MGI:99677] | Adcy1 | Enriched in dLGN | 2.244 | 8.482E-14 | 5232 |
| ENSMUSG00000034324 | transmembrane protein 132C [Source:MGI Symbol;Acc:MGI:2443061] | Tmem132c | Enriched in dLGN | 2.241 | 1.137E-16 | 360 |
| ENSMUSG00000029066 | mitochondrial ribosomal protein L20 [Source:MGI Symbol;Acc:MGI:2137221] | Mrpl20 | Enriched in dLGN | 2.24 | 5.323E-10 | 48.19 |
| ENSMUSG00000024935 | solute carrier family 1 (neuronal/epithelial high affinity glutamate transporter, system Xag), member 1 [Source:MGI Symbol;Acc:MGI:105083] | Slc1a1 | Enriched in dLGN | 2.23 | 2.472E-22 | 323.7 |
| ENSMUSG00000034825 | nuclear receptor interacting protein 3 [Source:MGI Symbol;Acc:MGI:1925843] | Nrip3 | Enriched in dLGN | 2.225 | 1.016E-10 | 285.5 |
| ENSMUSG00000063524 | enolase 1, alpha non-neuron [Source:MGI Symbol;Acc:MGI:95393] | Eno1 | Enriched in dLGN | 2.225 | 6.135E-10 | 583.3 |
| ENSMUSG00000026888 | growth factor receptor bound protein 14 [Source:MGI Symbol;Acc:MGI:1355324] | Grb14 | Enriched in dLGN | 2.224 | 5.157E-09 | 102.4 |
| ENSMUSG00000049422 | coiled-coil-helix-coiled-coil-helix domain containing 10 [Source:MGI Symbol;Acc:MGI:2143558] | Chchd10 | Enriched in dLGN | 2.223 | 1.638E-12 | 224.5 |
| ENSMUSG00000062785 | potassium voltage gated channel, Shaw-related subfamily, member 3 [Source:MGI Symbol;Acc:MGI:96669] | Kcnc3 | Enriched in dLGN | 2.222 | 3.47E-25 | 437.6 |
| ENSMUSG00000111375 | BTB (POZ) domain containing 8 [Source:MGI Symbol;Acc:MGI:3646208] | Btbd8 | Enriched in dLGN | 2.212 | 8.216E-13 | 197 |
| ENSMUSG00000045009 | proline-rich transmembrane protein 3 [Source:MGI Symbol;Acc:MGI:2444810] | Prrt3 | Enriched in dLGN | 2.208 | 2.769E-14 | 111.1 |
| ENSMUSG00000002032 | transmembrane protein 25 [Source:MGI Symbol;Acc:MGI:1918937] | Tmem25 | Enriched in dLGN | 2.205 | 1.03E-12 | 70.33 |
| ENSMUSG00000020644 | inhibitor of DNA binding 2 [Source:MGI Symbol;Acc:MGI:96397] | Id2 | Enriched in dLGN | 2.205 | 0.000001007 | 36.74 |
| ENSMUSG00000027495 | family with sequence similarity 210, member B [Source:MGI Symbol;Acc:MGI:1914267] | Fam210b | Enriched in dLGN | 2.201 | 5.085E-09 | 55.78 |
| ENSMUSG00000079480 | peptidyl-prolyl cis/trans isomerase, NIMA-interacting, 4 (parvulin) [Source:MGI Symbol;Acc:MGI:1916963] | Pin4 | Enriched in dLGN | 2.201 | 0.000021 | 13.6 |
| ENSMUSG00000026260 | NADH:ubiquinone oxidoreductase subunit A10 [Source:MGI Symbol;Acc:MGI:1914523] | Ndufa10 | Enriched in dLGN | 2.197 | 2.598E-11 | 263.7 |
| ENSMUSG00000011752 | phosphoglycerate mutase 1 [Source:MGI Symbol;Acc:MGI:97552] | Pgam1 | Enriched in dLGN | 2.193 | 7.976E-12 | 217.3 |
| ENSMUSG00000064329 | sodium channel, voltage-gated, type I, alpha [Source:MGI Symbol;Acc:MGI:98246] | Scn1a | Enriched in dLGN | 2.185 | 1.179E-12 | 703.3 |
| ENSMUSG00000025545 | citrate lyase beta like [Source:MGI Symbol;Acc:MGI:1916884] | Clybl | Enriched in dLGN | 2.185 | 0.000001883 | 35.95 |
| ENSMUSG00000035849 | keratin 222 [Source:MGI Symbol;Acc:MGI:2442728] | Krt222 | Enriched in dLGN | 2.18 | 5.897E-12 | 127 |
| ENSMUSG00000027712 | annexin A5 [Source:MGI Symbol;Acc:MGI:106008] | Anxa5 | Enriched in dLGN | 2.179 | 0.00000058 | 102.3 |
| ENSMUSG00000037606 | oxysterol binding protein-like 5 [Source:MGI Symbol;Acc:MGI:1930265] | Osbpl5 | Enriched in dLGN | 2.178 | 4.002E-30 | 213.9 |
| ENSMUSG00000027133 | NOP10 ribonucleoprotein [Source:MGI Symbol;Acc:MGI:1913431] | Nop10 | Enriched in dLGN | 2.174 | 8.063E-07 | 25.83 |
| ENSMUSG00000037568 | vasohibin 2 [Source:MGI Symbol;Acc:MGI:2444826] | Vash2 | Enriched in dLGN | 2.172 | 1.685E-12 | 93.06 |
| ENSMUSG00000032279 | isocitrate dehydrogenase 3 (NAD+) alpha [Source:MGI Symbol;Acc:MGI:1915084] | Idh3a | Enriched in dLGN | 2.165 | 4.267E-07 | 302 |
| ENSMUSG00000021373 | CAP, adenylate cyclase-associated protein, 2 (yeast) [Source:MGI Symbol;Acc:MGI:1914502] | Cap2 | Enriched in dLGN | 2.163 | 1.703E-14 | 147 |
| ENSMUSG00000032564 | copine IV [Source:MGI Symbol;Acc:MGI:1921270] | Cpne4 | Enriched in dLGN | 2.163 | 1.325E-07 | 126.3 |
| ENSMUSG00000024727 | transient receptor potential cation channel, subfamily M, member 6 [Source:MGI Symbol;Acc:MGI:2675603] | Trpm6 | Enriched in dLGN | 2.162 | 0.002257 | 19.86 |
| ENSMUSG00000020163 | ubiquinol-cytochrome c reductase, complex III subunit XI [Source:MGI Symbol;Acc:MGI:1913844] | Uqcr11 | Enriched in dLGN | 2.161 | 2.296E-09 | 153.3 |
| ENSMUSG00000026568 | mitochondrial pyruvate carrier 2 [Source:MGI Symbol;Acc:MGI:1917706] | Mpc2 | Enriched in dLGN | 2.152 | 1.868E-12 | 84.48 |
| ENSMUSG00000055782 | ATP-binding cassette, sub-family D (ALD), member 2 [Source:MGI Symbol;Acc:MGI:1349467] | Abcd2 | Enriched in dLGN | 2.138 | 9.125E-12 | 85.55 |
| ENSMUSG00000029752 | asparagine synthetase [Source:MGI Symbol;Acc:MGI:1350929] | Asns | Enriched in dLGN | 2.137 | 1.602E-07 | 106.2 |
| ENSMUSG00000023861 | mitochondrial pyruvate carrier 1 [Source:MGI Symbol;Acc:MGI:1915240] | Mpc1 | Enriched in dLGN | 2.133 | 2.063E-09 | 73.22 |
| ENSMUSG00000025428 | ATP synthase, H+ transporting, mitochondrial F1 complex, alpha subunit 1 [Source:MGI Symbol;Acc:MGI:88115] | Atp5a1 | Enriched in dLGN | 2.129 | 1.369E-12 | 1689 |
| ENSMUSG00000005161 | peroxiredoxin 2 [Source:MGI Symbol;Acc:MGI:109486] | Prdx2 | Enriched in dLGN | 2.124 | 3.628E-11 | 321.2 |
| ENSMUSG00000024841 | eukaryotic translation initiation factor 1A domain containing [Source:MGI Symbol;Acc:MGI:1917110] | Eif1ad | Enriched in dLGN | 2.119 | 9.348E-12 | 57.89 |
| ENSMUSG00000041112 | engulfment and cell motility 1 [Source:MGI Symbol;Acc:MGI:2153044] | Elmo1 | Enriched in dLGN | 2.118 | 2.614E-11 | 598.8 |
| ENSMUSG00000019505 | ubiquitin B [Source:MGI Symbol;Acc:MGI:98888] | Ubb | Enriched in dLGN | 2.117 | 2.346E-07 | 652 |
| ENSMUSG00000038552 | fibronectin type III domain containing 4 [Source:MGI Symbol;Acc:MGI:1917195] | Fndc4 | Enriched in dLGN | 2.102 | 2.816E-26 | 238.1 |
| ENSMUSG00000040430 | phosphatidylinositol transfer protein, cytoplasmic 1 [Source:MGI Symbol;Acc:MGI:1919045] | Pitpnc1 | Enriched in dLGN | 2.101 | 1.221E-23 | 764.9 |
| ENSMUSG00000105827 | H2B clustered histone 18 [Source:MGI Symbol;Acc:MGI:2448413] | H2bc18 | Enriched in dLGN | 2.098 | 0.00001344 | 42.81 |
| ENSMUSG00000019210 | ATPase, H+ transporting, lysosomal V1 subunit E1 [Source:MGI Symbol;Acc:MGI:894326] | Atp6v1e1 | Enriched in dLGN | 2.075 | 3.269E-10 | 270.4 |
| ENSMUSG00000023904 | host cell factor C1 regulator 1 (XPO1-dependent) [Source:MGI Symbol;Acc:MGI:2663619] | Hcfc1r1 | Enriched in dLGN | 2.075 | 3.975E-09 | 96.23 |
| ENSMUSG00000018362 | karyopherin (importin) alpha 2 [Source:MGI Symbol;Acc:MGI:103561] | Kpna2 | Enriched in dLGN | 2.074 | 0.00002755 | 67.24 |
| ENSMUSG00000030350 | protein arginine N-methyltransferase 8 [Source:MGI Symbol;Acc:MGI:3043083] | Prmt8 | Enriched in dLGN | 2.072 | 1.104E-17 | 303.1 |
| ENSMUSG00000017167 | contactin associated protein-like 1 [Source:MGI Symbol;Acc:MGI:1858201] | Cntnap1 | Enriched in dLGN | 2.072 | 1.075E-09 | 622.8 |
| ENSMUSG00000030695 | aldolase A, fructose-bisphosphate [Source:MGI Symbol;Acc:MGI:87994] | Aldoa | Enriched in dLGN | 2.069 | 3.168E-12 | 1713 |
| ENSMUSG00000020949 | FK506 binding protein 3 [Source:MGI Symbol;Acc:MGI:1353460] | Fkbp3 | Enriched in dLGN | 2.065 | 3.655E-09 | 160.5 |
| ENSMUSG00000039943 | phospholipase C, beta 4 [Source:MGI Symbol;Acc:MGI:107464] | Plcb4 | Enriched in dLGN | 2.063 | 3.173E-14 | 926.4 |
| ENSMUSG00000045733 | shadow of prion protein [Source:MGI Symbol;Acc:MGI:3582583] | Sprn | Enriched in dLGN | 2.056 | 2.634E-21 | 239 |
| ENSMUSG00000028271 | general transcription factor IIB [Source:MGI Symbol;Acc:MGI:2385191] | Gtf2b | Enriched in dLGN | 2.056 | 3.711E-07 | 36.1 |
| ENSMUSG00000113902 | NADH:ubiquinone oxidoreductase subunit B1 [Source:MGI Symbol;Acc:MGI:3780865] | Ndufb1 | Enriched in dLGN | 2.053 | 0.000000567 | 43.14 |
| ENSMUSG00000019179 | malate dehydrogenase 2, NAD (mitochondrial) [Source:MGI Symbol;Acc:MGI:97050] | Mdh2 | Enriched in dLGN | 2.052 | 1.286E-09 | 626.3 |
| ENSMUSG00000078963 | heat shock factor binding protein 1-like 1 [Source:MGI Symbol;Acc:MGI:1913505] | Hsbp1l1 | Enriched in dLGN | 2.052 | 0.005006 | 35.03 |
| ENSMUSG00000062542 | synaptotagmin IX [Source:MGI Symbol;Acc:MGI:1926373] | Syt9 | Enriched in dLGN | 2.048 | 4.744E-21 | 283.3 |
| ENSMUSG00000034566 | ATP synthase, H+ transporting, mitochondrial F0 complex, subunit D [Source:MGI Symbol;Acc:MGI:1918929] | Atp5h | Enriched in dLGN | 2.047 | 2.513E-12 | 216.1 |
| ENSMUSG00000034880 | mitochondrial ribosomal protein L34 [Source:MGI Symbol;Acc:MGI:2137227] | Mrpl34 | Enriched in dLGN | 2.033 | 1.059E-08 | 34.19 |
| ENSMUSG00000001089 | leucine zipper protein 1 [Source:MGI Symbol;Acc:MGI:107629] | Luzp1 | Enriched in dLGN | 2.032 | 3.412E-13 | 485.1 |
| ENSMUSG00000021209 | protein phosphatase 4, regulatory subunit 4 [Source:MGI Symbol;Acc:MGI:1921771] | Ppp4r4 | Enriched in dLGN | 2.03 | 1.278E-13 | 139.4 |
| ENSMUSG00000002379 | NADH:ubiquinone oxidoreductase subunit A11 [Source:MGI Symbol;Acc:MGI:1917125] | Ndufa11 | Enriched in dLGN | 2.03 | 0.00002931 | 75.68 |
| ENSMUSG00000030718 | protein phosphatase methylesterase 1 [Source:MGI Symbol;Acc:MGI:1919840] | Ppme1 | Enriched in dLGN | 2.028 | 1.119E-13 | 576.9 |
| ENSMUSG00000035864 | synaptotagmin I [Source:MGI Symbol;Acc:MGI:99667] | Syt1 | Enriched in dLGN | 2.025 | 3.618E-16 | 3835 |
| ENSMUSG00000034520 | gap junction protein, gamma 1 [Source:MGI Symbol;Acc:MGI:95718] | Gjc1 | Enriched in dLGN | 2.022 | 6.838E-08 | 53.26 |
| ENSMUSG00000020140 | leucine rich repeat containing G protein coupled receptor 5 [Source:MGI Symbol;Acc:MGI:1341817] | Lgr5 | Enriched in dLGN | 2.021 | 0.00004265 | 46.54 |
| ENSMUSG00000040794 | C1q and tumor necrosis factor related protein 4 [Source:MGI Symbol;Acc:MGI:1914695] | C1qtnf4 | Enriched in dLGN | 2.02 | 2.73E-15 | 207.8 |
| ENSMUSG00000046834 | keratin 1 [Source:MGI Symbol;Acc:MGI:96698] | Krt1 | Enriched in dLGN | 2.016 | 0.00002228 | 16.62 |
| ENSMUSG00000030647 | NADH:ubiquinone oxidoreductase subunit C2 [Source:MGI Symbol;Acc:MGI:1344370] | Ndufc2 | Enriched in dLGN | 2.014 | 0.000000452 | 77.57 |
| ENSMUSG00000059409 | protein phosphatase 2, regulatory subunit B', delta [Source:MGI Symbol;Acc:MGI:2388481] | Ppp2r5d | Enriched in dLGN | 2.013 | 1.187E-22 | 362.3 |
| ENSMUSG00000024953 | peroxiredoxin 5 [Source:MGI Symbol;Acc:MGI:1859821] | Prdx5 | Enriched in dLGN | 2.012 | 9.439E-17 | 344.2 |
| ENSMUSG00000027108 | Obg-like ATPase 1 [Source:MGI Symbol;Acc:MGI:1914309] | Ola1 | Enriched in dLGN | 2.006 | 0.003439 | 454.9 |
| ENSMUSG00000034891 | synuclein, beta [Source:MGI Symbol;Acc:MGI:1889011] | Sncb | Enriched in dLGN | 2.003 | 2.118E-11 | 939.2 |
| ENSMUSG00000032965 | intraflagellar transport 57 [Source:MGI Symbol;Acc:MGI:1921166] | Ift57 | Enriched in dLGN | 2.002 | 3.015E-20 | 254.5 |
| ENSMUSG00000030869 | NADH:ubiquinone oxidoreductase subunit AB1 [Source:MGI Symbol;Acc:MGI:1917566] | Ndufab1 | Enriched in dLGN | 1.998 | 3.797E-09 | 76.72 |
| ENSMUSG00000027965 | olfactomedin 3 [Source:MGI Symbol;Acc:MGI:2387329] | Olfm3 | Enriched in dLGN | 1.997 | 5.014E-08 | 175.9 |
| ENSMUSG00000071014 | NADH:ubiquinone oxidoreductase subunit B6 [Source:MGI Symbol;Acc:MGI:2684983] | Ndufb6 | Enriched in dLGN | 1.995 | 4.082E-08 | 50.89 |
| ENSMUSG00000040373 | calcium channel, voltage-dependent, gamma subunit 5 [Source:MGI Symbol;Acc:MGI:2157946] | Cacng5 | Enriched in dLGN | 1.982 | 3.513E-14 | 203.8 |
| ENSMUSG00000023089 | NADH:ubiquinone oxidoreductase subunit A5 [Source:MGI Symbol;Acc:MGI:1915452] | Ndufa5 | Enriched in dLGN | 1.982 | 1.33E-11 | 166.1 |
| ENSMUSG00000027674 | peroxisomal biogenesis factor 5-like [Source:MGI Symbol;Acc:MGI:1916672] | Pex5l | Enriched in dLGN | 1.978 | 0.000001071 | 789.1 |
| ENSMUSG00000001056 | NHP2 ribonucleoprotein [Source:MGI Symbol;Acc:MGI:1098547] | Nhp2 | Enriched in dLGN | 1.974 | 7.535E-07 | 27.46 |
| ENSMUSG00000059343 | aldolase 1 A, retrogene 1 [Source:MGI Symbol;Acc:MGI:2447811] | Aldoart1 | Enriched in dLGN | 1.973 | 0.00006135 | 19.45 |
| ENSMUSG00000008822 | acylphosphatase 1, erythrocyte (common) type [Source:MGI Symbol;Acc:MGI:1913454] | Acyp1 | Enriched in dLGN | 1.972 | 0.0003199 | 16.49 |
| ENSMUSG00000021253 | transforming growth factor, beta 3 [Source:MGI Symbol;Acc:MGI:98727] | Tgfb3 | Enriched in dLGN | 1.971 | 1.901E-11 | 74.63 |
| ENSMUSG00000073676 | heat shock protein 1 (chaperonin 10) [Source:MGI Symbol;Acc:MGI:104680] | Hspe1 | Enriched in dLGN | 1.97 | 3.083E-07 | 62.31 |
| ENSMUSG00000020635 | FK506 binding protein 1b [Source:MGI Symbol;Acc:MGI:1336205] | Fkbp1b | Enriched in dLGN | 1.97 | 0.000001402 | 66.86 |
| ENSMUSG00000030189 | Y box protein 3 [Source:MGI Symbol;Acc:MGI:2137670] | Ybx3 | Enriched in dLGN | 1.965 | 3.04E-09 | 527 |
| ENSMUSG00000015656 | heat shock protein 8 [Source:MGI Symbol;Acc:MGI:105384] | Hspa8 | Enriched in dLGN | 1.964 | 0.00001103 | 1562 |
| ENSMUSG00000040536 | N-terminal EF-hand calcium binding protein 1 [Source:MGI Symbol;Acc:MGI:1916602] | Necab1 | Enriched in dLGN | 1.958 | 2.922E-08 | 133 |
| ENSMUSG00000022956 | ATP synthase, H+ transporting, mitochondrial F1 complex, O subunit [Source:MGI Symbol;Acc:MGI:106341] | Atp5o | Enriched in dLGN | 1.958 | 1.075E-07 | 36.29 |
| ENSMUSG00000057666 | glyceraldehyde-3-phosphate dehydrogenase [Source:MGI Symbol;Acc:MGI:95640] | Gapdh | Enriched in dLGN | 1.947 | 0.0001003 | 93.82 |
| ENSMUSG00000010914 | pyruvate dehydrogenase complex, component X [Source:MGI Symbol;Acc:MGI:1351627] | Pdhx | Enriched in dLGN | 1.941 | 4.012E-15 | 152.7 |
| ENSMUSG00000042298 | tetratricopeptide repeat domain 19 [Source:MGI Symbol;Acc:MGI:1920045] | Ttc19 | Enriched in dLGN | 1.941 | 2.655E-14 | 462.6 |
| ENSMUSG00000031672 | glutamatic-oxaloacetic transaminase 2, mitochondrial [Source:MGI Symbol;Acc:MGI:95792] | Got2 | Enriched in dLGN | 1.934 | 2.475E-09 | 253.7 |
| ENSMUSG00000019877 | serine incorporator 1 [Source:MGI Symbol;Acc:MGI:1926228] | Serinc1 | Enriched in dLGN | 1.932 | 5.183E-09 | 662.3 |
| ENSMUSG00000060402 | carbohydrate sulfotransferase 8 [Source:MGI Symbol;Acc:MGI:1916197] | Chst8 | Enriched in dLGN | 1.927 | 3.846E-08 | 67.64 |
| ENSMUSG00000000901 | matrix metallopeptidase 11 [Source:MGI Symbol;Acc:MGI:97008] | Mmp11 | Enriched in dLGN | 1.927 | 0.00216 | 8.781 |
| ENSMUSG00000027350 | chromogranin B [Source:MGI Symbol;Acc:MGI:88395] | Chgb | Enriched in dLGN | 1.924 | 2.574E-13 | 1564 |
| ENSMUSG00000043461 | serine palmitoyltransferase, small subunit B [Source:MGI Symbol;Acc:MGI:1913433] | Sptssb | Enriched in dLGN | 1.917 | 0.004811 | 9.484 |
| ENSMUSG00000000563 | ATP synthase peripheral stalk-membrane subunit b [Source:MGI Symbol;Acc:MGI:1100495] | Atp5pb | Enriched in dLGN | 1.911 | 5.558E-12 | 236 |
| ENSMUSG00000032503 | cyclic AMP-regulated phosphoprotein, 21 [Source:MGI Symbol;Acc:MGI:107562] | Arpp21 | Enriched in dLGN | 1.907 | 9.986E-07 | 594.6 |
| ENSMUSG00000048495 | tRNA-yW synthesizing protein 5 [Source:MGI Symbol;Acc:MGI:1915986] | Tyw5 | Enriched in dLGN | 1.903 | 0.0002271 | 22.16 |
| ENSMUSG00000057766 | ankyrin repeat domain 29 [Source:MGI Symbol;Acc:MGI:2687055] | Ankrd29 | Enriched in dLGN | 1.902 | 8.577E-08 | 58.3 |
| ENSMUSG00000028289 | Eph receptor A7 [Source:MGI Symbol;Acc:MGI:95276] | Epha7 | Enriched in dLGN | 1.899 | 1.439E-09 | 197.6 |
| ENSMUSG00000040612 | immunoglobulin-like domain containing receptor 2 [Source:MGI Symbol;Acc:MGI:1196370] | Ildr2 | Enriched in dLGN | 1.896 | 1.94E-10 | 1307 |
| ENSMUSG00000040111 | GRAM domain containing 1B [Source:MGI Symbol;Acc:MGI:1925037] | Gramd1b | Enriched in dLGN | 1.895 | 5.192E-18 | 1123 |
| ENSMUSG00000022890 | ATP synthase, H+ transporting, mitochondrial F0 complex, subunit F [Source:MGI Symbol;Acc:MGI:107777] | Atp5j | Enriched in dLGN | 1.895 | 5.327E-10 | 249.2 |
| ENSMUSG00000031391 | L1 cell adhesion molecule [Source:MGI Symbol;Acc:MGI:96721] | L1cam | Enriched in dLGN | 1.895 | 0.000001091 | 803.2 |
| ENSMUSG00000045731 | prepronociceptin [Source:MGI Symbol;Acc:MGI:105308] | Pnoc | Enriched in dLGN | 1.894 | 0.000001144 | 50.56 |
| ENSMUSG00000027603 | gamma-glutamyltransferase 7 [Source:MGI Symbol;Acc:MGI:1913385] | Ggt7 | Enriched in dLGN | 1.893 | 4.409E-13 | 194.7 |
| ENSMUSG00000033918 | presenilin associated, rhomboid-like [Source:MGI Symbol;Acc:MGI:1277152] | Parl | Enriched in dLGN | 1.879 | 7.013E-12 | 65.89 |
| ENSMUSG00000036751 | cytochrome c oxidase, subunit 6B1 [Source:MGI Symbol;Acc:MGI:107460] | Cox6b1 | Enriched in dLGN | 1.876 | 2.077E-11 | 143.6 |
| ENSMUSG00000038462 | ubiquinol-cytochrome c reductase, Rieske iron-sulfur polypeptide 1 [Source:MGI Symbol;Acc:MGI:1913944] | Uqcrfs1 | Enriched in dLGN | 1.872 | 2.829E-12 | 172.4 |
| ENSMUSG00000044894 | ubiquinol-cytochrome c reductase, complex III subunit VII [Source:MGI Symbol;Acc:MGI:107807] | Uqcrq | Enriched in dLGN | 1.87 | 1.103E-10 | 236.5 |
| ENSMUSG00000027360 | histidine decarboxylase [Source:MGI Symbol;Acc:MGI:96062] | Hdc | Enriched in dLGN | 1.867 | 0.00004351 | 44.93 |
| ENSMUSG00000056258 | potassium voltage-gated channel, subfamily Q, member 3 [Source:MGI Symbol;Acc:MGI:1336181] | Kcnq3 | Enriched in dLGN | 1.864 | 1.046E-09 | 1087 |
| ENSMUSG00000025968 | NADH:ubiquinone oxidoreductase core subunit S1 [Source:MGI Symbol;Acc:MGI:2443241] | Ndufs1 | Enriched in dLGN | 1.86 | 6.629E-13 | 337.9 |
| ENSMUSG00000015932 | destrin [Source:MGI Symbol;Acc:MGI:1929270] | Dstn | Enriched in dLGN | 1.859 | 5.812E-09 | 300.9 |
| ENSMUSG00000034640 | TCDD-inducible poly(ADP-ribose) polymerase [Source:MGI Symbol;Acc:MGI:2159210] | Tiparp | Enriched in dLGN | 1.858 | 4.127E-08 | 38.05 |
| ENSMUSG00000032435 | dynein cytoplasmic 1 light intermediate chain 1 [Source:MGI Symbol;Acc:MGI:2135610] | Dync1li1 | Enriched in dLGN | 1.852 | 2.854E-40 | 559.8 |
| ENSMUSG00000037152 | NADH:ubiquinone oxidoreductase subunit C1 [Source:MGI Symbol;Acc:MGI:1913627] | Ndufc1 | Enriched in dLGN | 1.851 | 0.000005422 | 49.47 |
| ENSMUSG00000027406 | isocitrate dehydrogenase 3 (NAD+) beta [Source:MGI Symbol;Acc:MGI:2158650] | Idh3b | Enriched in dLGN | 1.85 | 3.25E-11 | 236.2 |
| ENSMUSG00000050541 | adrenergic receptor, alpha 1b [Source:MGI Symbol;Acc:MGI:104774] | Adra1b | Enriched in dLGN | 1.848 | 1.376E-07 | 73.77 |
| ENSMUSG00000032303 | cholinergic receptor, nicotinic, alpha polypeptide 3 [Source:MGI Symbol;Acc:MGI:87887] | Chrna3 | Enriched in dLGN | 1.847 | 0.0003687 | 38.91 |
| ENSMUSG00000025260 | hydroxysteroid (17-beta) dehydrogenase 10 [Source:MGI Symbol;Acc:MGI:1333871] | Hsd17b10 | Enriched in dLGN | 1.845 | 0.0003039 | 20.9 |
| ENSMUSG00000002068 | cyclin E1 [Source:MGI Symbol;Acc:MGI:88316] | Ccne1 | Enriched in dLGN | 1.84 | 0.000004707 | 29.03 |

| | | | | | | |
|---|---|---|---|---|---|---|
| ENSMUSG00000023456 | triosephosphate isomerase 1 [Source:MGI Symbol;Acc:MGI:98797] | Tpi1 | Enriched in dLGN | 1.838 | 4.839E-09 | 483.5 |
| ENSMUSG00000048332 | lipoma HMGIC fusion partner [Source:MGI Symbol;Acc:MGI:1920048] | Lhfp | Enriched in dLGN | 1.835 | 0.00000363 | 294.9 |
| ENSMUSG00000061482 | H4 clustered histone 4 [Source:MGI Symbol;Acc:MGI:2448423] | H4c4 | Enriched in dLGN | 1.831 | 0.00002967 | 85.46 |
| ENSMUSG00000079426 | actin related protein 2/3 complex, subunit 4 [Source:MGI Symbol;Acc:MGI:1915339] | Arpc4 | Enriched in dLGN | 1.83 | 4.098E-11 | 89.11 |
| ENSMUSG00000031007 | ATPase, H+ transporting, lysosomal accessory protein 2 [Source:MGI Symbol;Acc:MGI:1917745] | Atp6ap2 | Enriched in dLGN | 1.83 | 6.907E-09 | 242.6 |
| ENSMUSG00000032959 | phosphatidylethanolamine binding protein 1 [Source:MGI Symbol;Acc:MGI:1344408] | Pebp1 | Enriched in dLGN | 1.83 | 1.129E-07 | 164.1 |
| ENSMUSG00000041763 | tripeptidyl peptidase II [Source:MGI Symbol;Acc:MGI:102724] | Tpp2 | Enriched in dLGN | 1.829 | 1.457E-12 | 165.9 |
| ENSMUSG00000061518 | cytochrome c oxidase subunit 5B [Source:MGI Symbol;Acc:MGI:88475] | Cox5b | Enriched in dLGN | 1.829 | 7.341E-11 | 455.8 |
| ENSMUSG00000018965 | tyrosine 3-monooxygenase/tryptophan 5-monooxygenase activation protein, eta polypeptide [Source:MGI Symbol;Acc:MGI:109194] | Ywhah | Enriched in dLGN | 1.827 | 2.404E-08 | 3239 |
| ENSMUSG00000030612 | mitochondrial ribosomal protein L46 [Source:MGI Symbol;Acc:MGI:1914558] | Mrpl46 | Enriched in dLGN | 1.826 | 5.057E-07 | 33.73 |
| ENSMUSG00000036438 | calmodulin 2 [Source:MGI Symbol;Acc:MGI:103250] | Calm2 | Enriched in dLGN | 1.824 | 0.000007696 | 1256 |
| ENSMUSG00000020309 | ChaC, cation transport regulator 2 [Source:MGI Symbol;Acc:MGI:1915294] | Chac2 | Enriched in dLGN | 1.824 | 0.0002612 | 18.28 |
| ENSMUSG00000026526 | fumarate hydratase 1 [Source:MGI Symbol;Acc:MGI:95530] | Fh1 | Enriched in dLGN | 1.823 | 6.026E-07 | 124 |
| ENSMUSG00000024524 | guanine nucleotide binding protein, alpha stimulating, olfactory type [Source:MGI Symbol;Acc:MGI:95774] | Gnal | Enriched in dLGN | 1.82 | 3.016E-18 | 618.2 |
| ENSMUSG00000033917 | glycerophosphodiester phosphodiesterase 1 [Source:MGI Symbol;Acc:MGI:1891827] | Gde1 | Enriched in dLGN | 1.819 | 2.1E-17 | 234.5 |
| ENSMUSG00000060429 | syntrophin, basic 1 [Source:MGI Symbol;Acc:MGI:101781] | Sntb1 | Enriched in dLGN | 1.819 | 4.544E-10 | 115.5 |
| ENSMUSG00000046442 | protein phosphatase 1E (PP2C domain containing) [Source:MGI Symbol;Acc:MGI:2444096] | Ppm1e | Enriched in dLGN | 1.816 | 9.197E-15 | 844.2 |
| ENSMUSG00000024387 | casein kinase 2, beta polypeptide [Source:MGI Symbol;Acc:MGI:88548] | Csnk2b | Enriched in dLGN | 1.815 | 0.000004377 | 97.86 |
| ENSMUSG00000026496 | poly (ADP-ribose) polymerase family, member 1 [Source:MGI Symbol;Acc:MGI:1340806] | Parp1 | Enriched in dLGN | 1.81 | 9.447E-20 | 378.8 |
| ENSMUSG00000029918 | mitochondrial ribosomal protein S33 [Source:MGI Symbol;Acc:MGI:1338046] | Mrps33 | Enriched in dLGN | 1.81 | 0.00006322 | 26.36 |
| ENSMUSG00000056201 | cofilin 1, non-muscle [Source:MGI Symbol;Acc:MGI:101757] | Cfl1 | Enriched in dLGN | 1.795 | 3.833E-09 | 762.6 |
| ENSMUSG00000029649 | proteasome maturation protein [Source:MGI Symbol;Acc:MGI:1913787] | Pomp | Enriched in dLGN | 1.793 | 6.85E-10 | 109 |
| ENSMUSG00000051671 | cytochrome c oxidase assembly factor 6 [Source:MGI Symbol;Acc:MGI:1915142] | Coa6 | Enriched in dLGN | 1.791 | 0.0006758 | 13.32 |
| ENSMUSG00000049612 | oligodendrocyte myelin glycoprotein [Source:MGI Symbol;Acc:MGI:106586] | Omg | Enriched in dLGN | 1.786 | 0.000004732 | 36.01 |
| ENSMUSG00000052302 | TBC1 domain family, member 30 [Source:MGI Symbol;Acc:MGI:1921944] | Tbc1d30 | Enriched in dLGN | 1.784 | 2.417E-19 | 314.2 |
| ENSMUSG00000034488 | EGF-like repeats and discoidin I-like domains 3 [Source:MGI Symbol;Acc:MGI:1329025] | Edil3 | Enriched in dLGN | 1.784 | 7.039E-07 | 564.8 |
| ENSMUSG00000079317 | trafficking protein particle complex 2 [Source:MGI Symbol;Acc:MGI:1913476] | Trappc2 | Enriched in dLGN | 1.782 | 0.0001916 | 15.83 |
| ENSMUSG00000040907 | ATPase, Na+/K+ transporting, alpha 3 polypeptide [Source:MGI Symbol;Acc:MGI:88107] | Atp1a3 | Enriched in dLGN | 1.777 | 1.375E-14 | 7574 |
| ENSMUSG00000045427 | heterogeneous nuclear ribonucleoprotein H2 [Source:MGI Symbol;Acc:MGI:1201779] | Hnrnph2 | Enriched in dLGN | 1.777 | 2.046E-09 | 111 |
| ENSMUSG00000022889 | mitochondrial ribosomal protein L39 [Source:MGI Symbol;Acc:MGI:1351620] | Mrpl39 | Enriched in dLGN | 1.775 | 2.422E-08 | 63.69 |
| ENSMUSG00000070953 | Rab9 effector protein with kelch motifs [Source:MGI Symbol;Acc:MGI:2139530] | Rabepk | Enriched in dLGN | 1.775 | 0.000006618 | 41.33 |
| ENSMUSG00000055839 | elongin B [Source:MGI Symbol;Acc:MGI:1914923] | Elob | Enriched in dLGN | 1.774 | 4.393E-09 | 99.23 |
| ENSMUSG00000016427 | NADH:ubiquinone oxidoreductase subunit A1 [Source:MGI Symbol;Acc:MGI:1929511] | Ndufa1 | Enriched in dLGN | 1.772 | 2.369E-10 | 84.15 |
| ENSMUSG00000025781 | ATP synthase, H+ transporting, mitochondrial F1 complex, gamma polypeptide 1 [Source:MGI Symbol;Acc:MGI:1261437] | Atp5c1 | Enriched in dLGN | 1.77 | 4.21E-14 | 290 |
| ENSMUSG00000036850 | mitochondrial ribosomal protein L41 [Source:MGI Symbol;Acc:MGI:1333816] | Mrpl41 | Enriched in dLGN | 1.77 | 2.037E-09 | 53.68 |
| ENSMUSG00000020664 | dihydrolipoamide dehydrogenase [Source:MGI Symbol;Acc:MGI:107450] | Dld | Enriched in dLGN | 1.77 | 4.576E-09 | 258.4 |
| ENSMUSG00000084786 | ubiquitin-like 5 [Source:MGI Symbol;Acc:MGI:1913427] | Ubl5 | Enriched in dLGN | 1.766 | 1.657E-11 | 89.51 |
| ENSMUSG00000030591 | proteasome (prosome, macropain) 26S subunit, non-ATPase, 8 [Source:MGI Symbol;Acc:MGI:1888669] | Psmd8 | Enriched in dLGN | 1.757 | 3.901E-14 | 174.8 |
| ENSMUSG00000026032 | NADH:ubiquinone oxidoreductase subunit B3 [Source:MGI Symbol;Acc:MGI:1913745] | Ndufb3 | Enriched in dLGN | 1.757 | 2.564E-08 | 49.01 |
| ENSMUSG00000036934 | RIKEN cDNA 4921524J17 gene [Source:MGI Symbol;Acc:MGI:1913964] | 4921524J17Rik | Enriched in dLGN | 1.753 | 0.000004474 | 42.54 |
| ENSMUSG00000003949 | hepatic leukemia factor [Source:MGI Symbol;Acc:MGI:96108] | Hlf | Enriched in dLGN | 1.751 | 1.894E-08 | 354.9 |
| ENSMUSG00000025651 | ubiquinol-cytochrome c reductase core protein 1 [Source:MGI Symbol;Acc:MGI:107876] | Uqcrc1 | Enriched in dLGN | 1.748 | 1.599E-07 | 409.8 |
| ENSMUSG00000020115 | TANK-binding kinase 1 [Source:MGI Symbol;Acc:MGI:1929658] | Tbk1 | Enriched in dLGN | 1.744 | 3.817E-09 | 121.8 |
| ENSMUSG00000038244 | microtubule associated monooxygenase, calponin and LIM domain containing 2 [Source:MGI Symbol;Acc:MGI:2444947] | Mical2 | Enriched in dLGN | 1.741 | 2.082E-08 | 255.1 |
| ENSMUSG00000030423 | processing of precursor 4, ribonuclease P/MRP family, (S. cerevisiae) [Source:MGI Symbol;Acc:MGI:1913411] | Pop4 | Enriched in dLGN | 1.74 | 0.00003739 | 40.41 |
| ENSMUSG00000032551 | RIKEN cDNA 1110059G10 gene [Source:MGI Symbol;Acc:MGI:1913452] | 1110059G10Rik | Enriched in dLGN | 1.738 | 0.00527 | 11.31 |
| ENSMUSG00000029467 | ATPase, Ca++ transporting, cardiac muscle, slow twitch 2 [Source:MGI Symbol;Acc:MGI:88110] | Atp2a2 | Enriched in dLGN | 1.737 | 6.994E-17 | 2995 |
| ENSMUSG00000022965 | interferon gamma receptor 2 [Source:MGI Symbol;Acc:MGI:107654] | Ifngr2 | Enriched in dLGN | 1.731 | 1.549E-09 | 119.8 |
| ENSMUSG00000021771 | voltage-dependent anion channel 2 [Source:MGI Symbol;Acc:MGI:106915] | Vdac2 | Enriched in dLGN | 1.727 | 0.000003265 | 243.1 |
| ENSMUSG00000036932 | apoptosis-inducing factor, mitochondrion-associated 1 [Source:MGI Symbol;Acc:MGI:1349419] | Aifm1 | Enriched in dLGN | 1.725 | 8.547E-07 | 96.3 |
| ENSMUSG00000024644 | CNDP dipeptidase 2 (metallopeptidase M20 family) [Source:MGI Symbol;Acc:MGI:1913304] | Cndp2 | Enriched in dLGN | 1.724 | 0.000000129 | 120.2 |
| ENSMUSG00000021748 | pyruvate dehydrogenase (lipoamide) beta [Source:MGI Symbol;Acc:MGI:1915513] | Pdhb | Enriched in dLGN | 1.717 | 7.375E-11 | 160.8 |
| ENSMUSG00000003072 | ATP synthase, H+ transporting, mitochondrial F1 complex, delta subunit [Source:MGI Symbol;Acc:MGI:1913293] | Atp5d | Enriched in dLGN | 1.716 | 5.622E-12 | 259.8 |
| ENSMUSG00000001763 | tetraspanin 33 [Source:MGI Symbol;Acc:MGI:1919012] | Tspan33 | Enriched in dLGN | 1.716 | 3.659E-09 | 62.53 |
| ENSMUSG00000021576 | programmed cell death 6 [Source:MGI Symbol;Acc:MGI:109283] | Pdcd6 | Enriched in dLGN | 1.716 | 0.00001856 | 37.47 |
| ENSMUSG00000047126 | clathrin, heavy polypeptide (Hc) [Source:MGI Symbol;Acc:MGI:2388633] | Cltc | Enriched in dLGN | 1.713 | 1.439E-09 | 1529 |
| ENSMUSG00000006717 | acyl-CoA thioesterase 13 [Source:MGI Symbol;Acc:MGI:1914084] | Acot13 | Enriched in dLGN | 1.704 | 0.00002799 | 30.71 |
| ENSMUSG00000007564 | protein phosphatase 2, regulatory subunit A, alpha [Source:MGI Symbol;Acc:MGI:1926334] | Ppp2r1a | Enriched in dLGN | 1.703 | 3.769E-08 | 634.9 |
| ENSMUSG00000031778 | chemokine (C-X3-C motif) ligand 1 [Source:MGI Symbol;Acc:MGI:1097153] | Cx3cl1 | Enriched in dLGN | 1.703 | 2.642E-07 | 522 |
| ENSMUSG00000024044 | erythrocyte membrane protein band 4.1 like 3 [Source:MGI Symbol;Acc:MGI:103008] | Epb41l3 | Enriched in dLGN | 1.701 | 8.807E-10 | 1430 |
| ENSMUSG00000031505 | NAD(P)HX dehydratase [Source:MGI Symbol;Acc:MGI:1913353] | Naxd | Enriched in dLGN | 1.701 | 0.0003919 | 43.91 |
| ENSMUSG00000030061 | ubiquitin-like modifier activating enzyme 3 [Source:MGI Symbol;Acc:MGI:1341217] | Uba3 | Enriched in dLGN | 1.692 | 1.088E-07 | 78.39 |
| ENSMUSG00000019814 | LTV1 ribosome biogenesis factor [Source:MGI Symbol;Acc:MGI:2447810] | Ltv1 | Enriched in dLGN | 1.691 | 0.00005151 | 46.86 |
| ENSMUSG00000062825 | actin, gamma, cytoplasmic 1 [Source:MGI Symbol;Acc:MGI:87906] | Actg1 | Enriched in dLGN | 1.691 | 0.00006637 | 407 |
| ENSMUSG00000055409 | NEL-like 1 [Source:MGI Symbol;Acc:MGI:2443902] | Nell1 | Enriched in dLGN | 1.69 | 0.00004102 | 543.3 |
| ENSMUSG00000031176 | dynein light chain Tctex-type 3 [Source:MGI Symbol;Acc:MGI:1914367] | Dynlt3 | Enriched in dLGN | 1.684 | 0.000001132 | 112.8 |
| ENSMUSG00000017778 | cytochrome c oxidase subunit 7C [Source:MGI Symbol;Acc:MGI:103226] | Cox7c | Enriched in dLGN | 1.681 | 0.000005669 | 35.38 |
| ENSMUSG00000059278 | N(alpha)-acetyltransferase 38, NatC auxiliary subunit [Source:MGI Symbol;Acc:MGI:1925554] | Naa38 | Enriched in dLGN | 1.68 | 0.0000703 | 31.83 |
| ENSMUSG00000043091 | tubulin, alpha 1C [Source:MGI Symbol;Acc:MGI:1095409] | Tuba1c | Enriched in dLGN | 1.68 | 0.01178 | 41.77 |
| ENSMUSG00000031633 | solute carrier family 25 (mitochondrial carrier, adenine nucleotide translocator), member 4 [Source:MGI Symbol;Acc:MGI:1353495] | Slc25a4 | Enriched in dLGN | 1.678 | 8.299E-09 | 1489 |
| ENSMUSG00000027134 | lysophosphatidylcholine acyltransferase 4 [Source:MGI Symbol;Acc:MGI:2138993] | Lpcat4 | Enriched in dLGN | 1.677 | 3.487E-15 | 130.5 |
| ENSMUSG00000025917 | COP9 signalosome subunit 5 [Source:MGI Symbol;Acc:MGI:1349415] | Cops5 | Enriched in dLGN | 1.677 | 0.00001072 | 96.98 |
| ENSMUSG00000078941 | adenylate kinase 6 [Source:MGI Symbol;Acc:MGI:5510732] | Ak6 | Enriched in dLGN | 1.676 | 0.0002437 | 18.09 |
| ENSMUSG00000026904 | solute carrier family 4, sodium bicarbonate cotransporter-like, member 10 [Source:MGI Symbol;Acc:MGI:2150150] | Slc4a10 | Enriched in dLGN | 1.674 | 5.003E-10 | 601.1 |
| ENSMUSG00000056091 | ST3 beta-galactoside alpha-2,3-sialyltransferase 5 [Source:MGI Symbol;Acc:MGI:1339963] | St3gal5 | Enriched in dLGN | 1.671 | 7.049E-15 | 176.3 |
| ENSMUSG00000020153 | NADH:ubiquinone oxidoreductase core subunit S7 [Source:MGI Symbol;Acc:MGI:1922656] | Ndufs7 | Enriched in dLGN | 1.671 | 2.871E-10 | 124 |
| ENSMUSG00000032355 | muscular LMNA-interacting protein [Source:MGI Symbol;Acc:MGI:1916892] | Mlip | Enriched in dLGN | 1.671 | 0.004052 | 22.7 |
| ENSMUSG00000024414 | mitochondrial ribosomal protein L27 [Source:MGI Symbol;Acc:MGI:2137224] | Mrpl27 | Enriched in dLGN | 1.67 | 0.00006714 | 26.69 |
| ENSMUSG00000030127 | COP9 signalosome subunit 7A [Source:MGI Symbol;Acc:MGI:1349400] | Cops7a | Enriched in dLGN | 1.669 | 3.05E-12 | 167 |
| ENSMUSG00000027346 | glycerophosphocholine phosphodiesterase 1 [Source:MGI Symbol;Acc:MGI:104898] | Gpcpd1 | Enriched in dLGN | 1.667 | 4.973E-10 | 161 |
| ENSMUSG00000028444 | ciliary neurotrophic factor receptor [Source:MGI Symbol;Acc:MGI:99605] | Cntfr | Enriched in dLGN | 1.662 | 1.06E-10 | 219.4 |
| ENSMUSG00000000326 | catechol-O-methyltransferase [Source:MGI Symbol;Acc:MGI:88470] | Comt | Enriched in dLGN | 1.661 | 2.972E-11 | 103.3 |
| ENSMUSG00000061474 | mitochondrial ribosomal protein S36 [Source:MGI Symbol;Acc:MGI:1913378] | Mrps36 | Enriched in dLGN | 1.658 | 0.002143 | 23.77 |
| ENSMUSG00000038722 | BUD31 homolog [Source:MGI Symbol;Acc:MGI:2141291] | Bud31 | Enriched in dLGN | 1.656 | 5.636E-09 | 48.97 |
| ENSMUSG00000020225 | transmembrane BAX inhibitor motif containing 4 [Source:MGI Symbol;Acc:MGI:1915462] | Tmbim4 | Enriched in dLGN | 1.654 | 0.0001757 | 25.14 |
| ENSMUSG00000031971 | centriole, cilia and spindle associated protein [Source:MGI Symbol;Acc:MGI:1920670] | Ccsap | Enriched in dLGN | 1.652 | 2.339E-13 | 95.08 |
| ENSMUSG00000027698 | neutral cholesterol ester hydrolase 1 [Source:MGI Symbol;Acc:MGI:2443191] | Nceh1 | Enriched in dLGN | 1.652 | 6.978E-08 | 190.1 |
| ENSMUSG00000027273 | synaptosomal-associated protein 25 [Source:MGI Symbol;Acc:MGI:98331] | Snap25 | Enriched in dLGN | 1.651 | 2.43E-09 | 1966 |
| ENSMUSG00000052456 | guided entry of tail-anchored proteins factor 3, ATPase [Source:MGI Symbol;Acc:MGI:1928379] | Get3 | Enriched in dLGN | 1.65 | 1.113E-08 | 120.5 |
| ENSMUSG00000022587 | lymphocyte antigen 6 complex, locus E [Source:MGI Symbol;Acc:MGI:106651] | Ly6e | Enriched in dLGN | 1.65 | 2.892E-07 | 95.38 |
| ENSMUSG00000021832 | proteasome (prosome, macropain) 26S subunit, ATPase, 6 [Source:MGI Symbol;Acc:MGI:1914339] | Psmc6 | Enriched in dLGN | 1.649 | 8.96E-08 | 131.6 |
| ENSMUSG00000035674 | NADH:ubiquinone oxidoreductase subunit A3 [Source:MGI Symbol;Acc:MGI:1913341] | Ndufa3 | Enriched in dLGN | 1.649 | 0.00002139 | 56.67 |
| ENSMUSG00000075702 | selenoprotein M [Source:MGI Symbol;Acc:MGI:2149786] | Selenom | Enriched in dLGN | 1.646 | 2.22E-09 | 136.2 |
| ENSMUSG00000002102 | proteasome (prosome, macropain) 26S subunit, ATPase 3 [Source:MGI Symbol;Acc:MGI:1098754] | Psmc3 | Enriched in dLGN | 1.642 | 4.703E-10 | 304.4 |
| ENSMUSG00000035443 | thymocyte nuclear protein 1 [Source:MGI Symbol;Acc:MGI:1925112] | Thyn1 | Enriched in dLGN | 1.642 | 0.00005339 | 32.79 |
| ENSMUSG00000029860 | zyxin [Source:MGI Symbol;Acc:MGI:103072] | Zyx | Enriched in dLGN | 1.641 | 3.977E-10 | 118.6 |
| ENSMUSG00000022241 | threonyl-tRNA synthetase [Source:MGI Symbol;Acc:MGI:106314] | Tars | Enriched in dLGN | 1.64 | 3.872E-11 | 160.9 |
| ENSMUSG00000029185 | family with sequence similarity 114, member A1 [Source:MGI Symbol;Acc:MGI:1915553] | Fam114a1 | Enriched in dLGN | 1.64 | 0.0001054 | 29.67 |
| ENSMUSG00000032459 | mitochondrial ribosomal protein S22 [Source:MGI Symbol;Acc:MGI:1928137] | Mrps22 | Enriched in dLGN | 1.636 | 0.002222 | 35.07 |
| ENSMUSG00000032336 | neuroplastin [Source:MGI Symbol;Acc:MGI:108077] | Nptn | Enriched in dLGN | 1.635 | 1.469E-16 | 662.1 |
| ENSMUSG00000032330 | cytochrome c oxidase subunit 7A2 [Source:MGI Symbol;Acc:MGI:1316715] | Cox7a2 | Enriched in dLGN | 1.634 | 3.432E-08 | 76.71 |
| ENSMUSG00000034187 | N-ethylmaleimide sensitive fusion protein [Source:MGI Symbol;Acc:MGI:104560] | Nsf | Enriched in dLGN | 1.628 | 1.345E-10 | 1114 |
| ENSMUSG00000031996 | amyloid beta (A4) precursor-like protein 2 [Source:MGI Symbol;Acc:MGI:88047] | Aplp2 | Enriched in dLGN | 1.624 | 5.958E-09 | 2704 |
| ENSMUSG00000031617 | transmembrane protein 184C [Source:MGI Symbol;Acc:MGI:2384562] | Tmem184c | Enriched in dLGN | 1.621 | 1.315E-08 | 115.9 |
| ENSMUSG00000026154 | succinate dehydrogenase complex assembly factor 4 [Source:MGI Symbol;Acc:MGI:1915252] | Sdhaf4 | Enriched in dLGN | 1.62 | 0.001548 | 17.29 |
| ENSMUSG00000023175 | basigin [Source:MGI Symbol;Acc:MGI:88208] | Bsg | Enriched in dLGN | 1.619 | 1.957E-11 | 872.5 |
| ENSMUSG00000039385 | cadherin 6 [Source:MGI Symbol;Acc:MGI:107435] | Cdh6 | Enriched in dLGN | 1.616 | 0.000004866 | 237 |
| ENSMUSG00000020029 | nudix (nucleoside diphosphate linked moiety X)-type motif 4 [Source:MGI Symbol;Acc:MGI:1918457] | Nudt4 | Enriched in dLGN | 1.614 | 2.981E-10 | 752.5 |
| ENSMUSG00000028447 | dynactin 3 [Source:MGI Symbol;Acc:MGI:1859251] | Dctn3 | Enriched in dLGN | 1.613 | 9.497E-13 | 123.3 |
| ENSMUSG00000005510 | NADH:ubiquinone oxidoreductase core subunit S3 [Source:MGI Symbol;Acc:MGI:1915599] | Ndufs3 | Enriched in dLGN | 1.609 | 3.211E-10 | 101.1 |
| ENSMUSG00000020806 | rhomboid 5 homolog 2 [Source:MGI Symbol;Acc:MGI:2442473] | Rhbdf2 | Enriched in dLGN | 1.609 | 0.03469 | 43.1 |
| ENSMUSG00000044835 | ankyrin repeat domain 45 [Source:MGI Symbol;Acc:MGI:1921094] | Ankrd45 | Enriched in dLGN | 1.607 | 2.384E-07 | 83.71 |
| ENSMUSG00000049097 | ankyrin repeat domain 34A [Source:MGI Symbol;Acc:MGI:3617846] | Ankrd34a | Enriched in dLGN | 1.605 | 0.000002317 | 84.03 |
| ENSMUSG00000027827 | potassium voltage-gated channel, shaker-related subfamily, beta member 1 [Source:MGI Symbol;Acc:MGI:109155] | Kcnab1 | Enriched in dLGN | 1.604 | 0.00003821 | 138 |
| ENSMUSG00000021368 | TBC1 domain family, member 7 [Source:MGI Symbol;Acc:MGI:1914296] | Tbc1d7 | Enriched in dLGN | 1.603 | 0.000001813 | 35.71 |
| ENSMUSG00000024172 | beta galactoside alpha 2,6 sialyltransferase 2 [Source:MGI Symbol;Acc:MGI:2445190] | St6gal2 | Enriched in dLGN | 1.602 | 7.406E-07 | 150.3 |
| ENSMUSG00000032306 | mannose phosphate isomerase [Source:MGI Symbol;Acc:MGI:97075] | Mpi | Enriched in dLGN | 1.601 | 3.724E-11 | 76.39 |
| ENSMUSG00000021114 | ATPase, H+ transporting, lysosomal V1 subunit D [Source:MGI Symbol;Acc:MGI:1921084] | Atp6v1d | Enriched in dLGN | 1.601 | 3.504E-09 | 284.8 |
| ENSMUSG00000040620 | DEAH (Asp-Glu-Ala-His) box polypeptide 33 [Source:MGI Symbol;Acc:MGI:2445102] | Dhx33 | Enriched in dLGN | 1.6 | 1.04E-08 | 72.8 |
| ENSMUSG00000031647 | microfibrillar-associated protein 3-like [Source:MGI Symbol;Acc:MGI:1918556] | Mfap3l | Enriched in dLGN | 1.598 | 1.654E-08 | 250.9 |
| ENSMUSG00000000296 | tumor protein D52-like 1 [Source:MGI Symbol;Acc:MGI:1298386] | Tpd52l1 | Enriched in dLGN | 1.598 | 0.000001302 | 61.04 |
| ENSMUSG00000063931 | peptidase D [Source:MGI Symbol;Acc:MGI:97542] | Pepd | Enriched in dLGN | 1.597 | 0.00001217 | 102.6 |
| ENSMUSG00000022376 | adenylate cyclase 8 [Source:MGI Symbol;Acc:MGI:1341110] | Adcy8 | Enriched in dLGN | 1.596 | 0.000001247 | 297.1 |
| ENSMUSG00000031634 | UFM1-specific peptidase 2 [Source:MGI Symbol;Acc:MGI:1913679] | Ufsp2 | Enriched in dLGN | 1.594 | 0.00002709 | 89.61 |
| ENSMUSG00000002728 | N(alpha)-acetyltransferase 20, NatB catalytic subunit [Source:MGI Symbol;Acc:MGI:1915127] | Naa20 | Enriched in dLGN | 1.593 | 0.00001278 | 46.04 |
| ENSMUSG00000029777 | glycyl-tRNA synthetase [Source:MGI Symbol;Acc:MGI:2449057] | Gars | Enriched in dLGN | 1.592 | 3.546E-13 | 453.4 |
| ENSMUSG00000044117 | bMERB domain containing 1 [Source:MGI Symbol;Acc:MGI:1914504] | Bmerb1 | Enriched in dLGN | 1.587 | 1.636E-13 | 457.3 |
| ENSMUSG00000024425 | Nedd4 family interacting protein 1 [Source:MGI Symbol;Acc:MGI:1929601] | Ndfip1 | Enriched in dLGN | 1.587 | 5.595E-10 | 747.3 |

| | | | | | | |
|---|---|---|---|---|---|---|
| ENSMUSG00000028410 | DnaJ heat shock protein family (Hsp40) member A1 [Source:MGI Symbol;Acc:MGI:1270129] | Dnaja1 | Enriched in dLGN | 1.587 | 0.0001244 | 251.8 |
| ENSMUSG00000048616 | noggin [Source:MGI Symbol;Acc:MGI:104327] | Nog | Enriched in dLGN | 1.584 | 0.000008592 | 53.91 |
| ENSMUSG00000021024 | proteasome subunit alpha 6 [Source:MGI Symbol;Acc:MGI:1347006] | Psma6 | Enriched in dLGN | 1.58 | 0.001064 | 71.31 |
| ENSMUSG00000016349 | eukaryotic translation elongation factor 1 alpha 2 [Source:MGI Symbol;Acc:MGI:1096317] | Eef1a2 | Enriched in dLGN | 1.579 | 3.914E-07 | 1496 |
| ENSMUSG00000031137 | fibroblast growth factor 13 [Source:MGI Symbol;Acc:MGI:109178] | Fgf13 | Enriched in dLGN | 1.575 | 2.597E-07 | 419.6 |
| ENSMUSG00000022892 | amyloid beta (A4) precursor protein [Source:MGI Symbol;Acc:MGI:88059] | App | Enriched in dLGN | 1.573 | 1.222E-07 | 4592 |
| ENSMUSG00000024099 | NADH:ubiquinone oxidoreductase core subunit V2 [Source:MGI Symbol;Acc:MGI:1920150] | Ndufv2 | Enriched in dLGN | 1.568 | 3.452E-12 | 217.5 |
| ENSMUSG00000040136 | ATP-binding cassette, sub-family C (CFTR/MRP), member 8 [Source:MGI Symbol;Acc:MGI:1352629] | Abcc8 | Enriched in dLGN | 1.568 | 0.00002649 | 86.74 |
| ENSMUSG00000079508 | apolipoprotein O [Source:MGI Symbol;Acc:MGI:1915566] | Apoo | Enriched in dLGN | 1.568 | 0.009451 | 29.79 |
| ENSMUSG00000031701 | DnaJ heat shock protein family (Hsp40) member A2 [Source:MGI Symbol;Acc:MGI:1931882] | Dnaja2 | Enriched in dLGN | 1.567 | 9.403E-09 | 401 |
| ENSMUSG00000000168 | dihydrolipoamide S-acetyltransferase (E2 component of pyruvate dehydrogenase complex) [Source:MGI Symbol;Acc:MGI:2385311] | Dlat | Enriched in dLGN | 1.562 | 1.013E-11 | 196.1 |
| ENSMUSG00000050830 | von Willebrand factor C domain containing 2 [Source:MGI Symbol;Acc:MGI:2442987] | Vwc2 | Enriched in dLGN | 1.562 | 0.000003246 | 127 |
| ENSMUSG00000025940 | transmembrane protein 70 [Source:MGI Symbol;Acc:MGI:1915068] | Tmem70 | Enriched in dLGN | 1.561 | 8.165E-08 | 50.14 |
| ENSMUSG00000001380 | histidyl-tRNA synthetase [Source:MGI Symbol;Acc:MGI:108087] | Hars | Enriched in dLGN | 1.56 | 3.641E-13 | 256 |
| ENSMUSG00000062352 | integrin beta 1 binding protein 1 [Source:MGI Symbol;Acc:MGI:1306802] | Itgb1bp1 | Enriched in dLGN | 1.56 | 4.033E-08 | 59.33 |
| ENSMUSG00000052698 | talin 2 [Source:MGI Symbol;Acc:MGI:1917799] | Tln2 | Enriched in dLGN | 1.558 | 3.273E-23 | 1565 |
| ENSMUSG00000038690 | ATP synthase, H+ transporting, mitochondrial F0 complex, subunit F2 [Source:MGI Symbol;Acc:MGI:1927558] | Atp5j2 | Enriched in dLGN | 1.558 | 1.112E-08 | 138.3 |
| ENSMUSG00000090841 | myosin, light polypeptide 6, alkali, smooth muscle and non-muscle [Source:MGI Symbol;Acc:MGI:109318] | Myl6 | Enriched in dLGN | 1.558 | 2.064E-08 | 274.2 |
| ENSMUSG00000002718 | chromosome segregation 1-like (S. cerevisiae) [Source:MGI Symbol;Acc:MGI:1339951] | Cse1l | Enriched in dLGN | 1.558 | 4.014E-07 | 100.5 |
| ENSMUSG00000029247 | phosphoribosylaminoimidazole carboxylase, phosphoribosylaminoribosylaminoimidazole, succinocarboxamide synthetase [Source:MGI Symbol;Acc:MGI:1914304] | Paics | Enriched in dLGN | 1.558 | 0.00001591 | 114.1 |
| ENSMUSG00000031059 | NADH:ubiquinone oxidoreductase subunit B11 [Source:MGI Symbol;Acc:MGI:1349919] | Ndufb11 | Enriched in dLGN | 1.556 | 1.991E-10 | 155.4 |
| ENSMUSG00000020589 | CYFIP related Rac1 interactor A [Source:MGI Symbol;Acc:MGI:1261783] | Cyria | Enriched in dLGN | 1.555 | 5.567E-10 | 283.7 |
| ENSMUSG00000023572 | cyclin D-type binding-protein 1 [Source:MGI Symbol;Acc:MGI:109595] | Ccndbp1 | Enriched in dLGN | 1.549 | 1.073E-07 | 105.3 |
| ENSMUSG00000062981 | mitochondrial ribosomal protein L42 [Source:MGI Symbol;Acc:MGI:1333774] | Mrpl42 | Enriched in dLGN | 1.549 | 0.0004232 | 43.37 |
| ENSMUSG00000032366 | tropomyosin 1, alpha [Source:MGI Symbol;Acc:MGI:98809] | Tpm1 | Enriched in dLGN | 1.544 | 2.723E-25 | 654.7 |
| ENSMUSG00000063694 | cytochrome c, somatic [Source:MGI Symbol;Acc:MGI:88578] | Cycs | Enriched in dLGN | 1.544 | 0.000002185 | 116.9 |
| ENSMUSG00000070319 | eukaryotic translation initiation factor 3, subunit G [Source:MGI Symbol;Acc:MGI:1858258] | Eif3g | Enriched in dLGN | 1.544 | 0.000004697 | 174.2 |
| ENSMUSG00000019951 | UHRF1 (ICBP90) binding protein 1-like [Source:MGI Symbol;Acc:MGI:2442888] | Uhrf1bp1l | Enriched in dLGN | 1.542 | 1.556E-21 | 276.6 |
| ENSMUSG00000004285 | ATPase, H+ transporting, lysosomal V1 subunit F [Source:MGI Symbol;Acc:MGI:1913394] | Atp6v1f | Enriched in dLGN | 1.539 | 0.0002033 | 78.37 |
| ENSMUSG00000031668 | eukaryotic translation initiation factor 2 alpha kinase 3 [Source:MGI Symbol;Acc:MGI:1341830] | Eif2ak3 | Enriched in dLGN | 1.537 | 3.333E-11 | 198.6 |
| ENSMUSG00000019087 | ATPase, H+ transporting, lysosomal accessory protein 1 [Source:MGI Symbol;Acc:MGI:109629] | Atp6ap1 | Enriched in dLGN | 1.533 | 1.244E-08 | 276.8 |
| ENSMUSG00000030291 | mediator complex subunit 21 [Source:MGI Symbol;Acc:MGI:1347064] | Med21 | Enriched in dLGN | 1.533 | 0.002474 | 15.67 |
| ENSMUSG00000022337 | ER membrane protein complex subunit 2 [Source:MGI Symbol;Acc:MGI:1913986] | Emc2 | Enriched in dLGN | 1.529 | 1.96E-08 | 104.6 |
| ENSMUSG00000038555 | receptor accessory protein 2 [Source:MGI Symbol;Acc:MGI:2385070] | Reep2 | Enriched in dLGN | 1.524 | 2.893E-12 | 302.7 |
| ENSMUSG00000075706 | glutathione peroxidase 4 [Source:MGI Symbol;Acc:MGI:104767] | Gpx4 | Enriched in dLGN | 1.524 | 1.738E-08 | 170.9 |
| ENSMUSG00000032249 | acidic (leucine-rich) nuclear phosphoprotein 32 family, member A [Source:MGI Symbol;Acc:MGI:108447] | Anp32a | Enriched in dLGN | 1.522 | 1.317E-18 | 803.6 |
| ENSMUSG00000035885 | cytochrome c oxidase subunit 8A [Source:MGI Symbol;Acc:MGI:105959] | Cox8a | Enriched in dLGN | 1.522 | 3.506E-09 | 292.6 |
| ENSMUSG00000024981 | acyl-CoA synthetase long-chain family member 5 [Source:MGI Symbol;Acc:MGI:1919129] | Acsl5 | Enriched in dLGN | 1.521 | 7.668E-07 | 65.86 |
| ENSMUSG00000015668 | PDZ domain containing 11 [Source:MGI Symbol;Acc:MGI:1919871] | Pdzd11 | Enriched in dLGN | 1.521 | 0.00007283 | 27.95 |
| ENSMUSG00000026527 | regulator of G protein signaling 7 [Source:MGI Symbol;Acc:MGI:1346089] | Rgs7 | Enriched in dLGN | 1.517 | 0.0003259 | 427.5 |
| ENSMUSG00000035372 | RIKEN cDNA 1810055G02 gene [Source:MGI Symbol;Acc:MGI:1919306] | 1810055G02Rik | Enriched in dLGN | 1.516 | 0.0002269 | 34.57 |
| ENSMUSG00000020481 | ankyrin repeat domain 36 [Source:MGI Symbol;Acc:MGI:1923639] | Ankrd36 | Enriched in dLGN | 1.516 | 0.004144 | 15.79 |
| ENSMUSG00000005683 | citrate synthase [Source:MGI Symbol;Acc:MGI:88529] | Cs | Enriched in dLGN | 1.509 | 1.703E-09 | 414 |
| ENSMUSG00000027203 | deoxyuridine triphosphatase [Source:MGI Symbol;Acc:MGI:1346051] | Dut | Enriched in dLGN | 1.508 | 6.312E-10 | 92.41 |
| ENSMUSG00000063531 | sema domain, immunoglobulin domain (Ig), short basic domain, secreted, (semaphorin) 3E [Source:MGI Symbol;Acc:MGI:1340034] | Sema3e | Enriched in dLGN | 1.507 | 0.001202 | 112.6 |
| ENSMUSG00000039809 | gamma-aminobutyric acid (GABA) B receptor, 2 [Source:MGI Symbol;Acc:MGI:2386030] | Gabbr2 | Enriched in dLGN | 1.506 | 2.634E-13 | 1664 |
| ENSMUSG00000037361 | splicing factor 3B, subunit 6 [Source:MGI Symbol;Acc:MGI:1913305] | Sf3b6 | Enriched in dLGN | 1.504 | 0.0008757 | 35.23 |
| ENSMUSG00000010095 | solute carrier family 3 (activators of dibasic and neutral amino acid transport), member 2 [Source:MGI Symbol;Acc:MGI:96955] | Slc3a2 | Enriched in dLGN | 1.502 | 2.18E-11 | 248.5 |
| ENSMUSG00000021194 | chromogranin A [Source:MGI Symbol;Acc:MGI:88394] | Chga | Enriched in dLGN | 1.501 | 0.00001835 | 812.4 |
| ENSMUSG00000060279 | adaptor-related protein complex 2, alpha 1 subunit [Source:MGI Symbol;Acc:MGI:101921] | Ap2a1 | Enriched in dLGN | 1.499 | 1.688E-11 | 671.9 |
| ENSMUSG00000044221 | G-rich RNA sequence binding factor 1 [Source:MGI Symbol;Acc:MGI:106479] | Grsf1 | Enriched in dLGN | 1.496 | 1.199E-11 | 242.3 |
| ENSMUSG00000083282 | cathepsin F [Source:MGI Symbol;Acc:MGI:1861434] | Ctsf | Enriched in dLGN | 1.496 | 3.871E-09 | 193.6 |
| ENSMUSG00000031918 | myotubularin related protein 2 [Source:MGI Symbol;Acc:MGI:1924366] | Mtmr2 | Enriched in dLGN | 1.496 | 4.108E-08 | 157.4 |
| ENSMUSG00000033419 | synaptosomal-associated protein 91 [Source:MGI Symbol;Acc:MGI:109132] | Snap91 | Enriched in dLGN | 1.494 | 3.842E-15 | 939.7 |
| ENSMUSG00000013921 | CAP-GLY domain containing linker protein 3 [Source:MGI Symbol;Acc:MGI:1923936] | Clip3 | Enriched in dLGN | 1.492 | 0.000004831 | 1062 |
| ENSMUSG00000028837 | proteasome (prosome, macropain) subunit, beta type 2 [Source:MGI Symbol;Acc:MGI:1347045] | Psmb2 | Enriched in dLGN | 1.492 | 0.0001542 | 65.51 |
| ENSMUSG00000021606 | NADH:ubiquinone oxidoreductase core subunit S6 [Source:MGI Symbol;Acc:MGI:107932] | Ndufs6 | Enriched in dLGN | 1.49 | 6.306E-09 | 70.27 |
| ENSMUSG00000030652 | demethyl-Q 7 [Source:MGI Symbol;Acc:MGI:107207] | Coq7 | Enriched in dLGN | 1.489 | 0.000003688 | 54.52 |
| ENSMUSG00000063882 | ubiquinol-cytochrome c reductase hinge protein [Source:MGI Symbol;Acc:MGI:1913826] | Uqcrh | Enriched in dLGN | 1.488 | 3.956E-10 | 256.5 |
| ENSMUSG00000091722 | siah E3 ubiquitin protein ligase family member 3 [Source:MGI Symbol;Acc:MGI:2685758] | Siah3 | Enriched in dLGN | 1.488 | 0.000001451 | 62.95 |
| ENSMUSG00000049775 | thymosin, beta 4, X chromosome [Source:MGI Symbol;Acc:MGI:99510] | Tmsb4x | Enriched in dLGN | 1.485 | 2.386E-11 | 317.2 |
| ENSMUSG00000025980 | heat shock protein 1 (chaperonin) [Source:MGI Symbol;Acc:MGI:96242] | Hspd1 | Enriched in dLGN | 1.485 | 1.324E-08 | 264 |
| ENSMUSG00000036989 | tripartite motif-containing 3 [Source:MGI Symbol;Acc:MGI:1860040] | Trim3 | Enriched in dLGN | 1.485 | 4.73E-08 | 276 |
| ENSMUSG00000022658 | transgelin 3 [Source:MGI Symbol;Acc:MGI:1926784] | Tagln3 | Enriched in dLGN | 1.485 | 0.00001233 | 267.2 |
| ENSMUSG00000031774 | proteasome activator subunit 3 interacting protein 1 [Source:MGI Symbol;Acc:MGI:1919637] | Psme3ip1 | Enriched in dLGN | 1.484 | 6.554E-14 | 123.3 |
| ENSMUSG00000026103 | glutaminase [Source:MGI Symbol;Acc:MGI:95752] | Gls | Enriched in dLGN | 1.483 | 1.113E-11 | 697 |
| ENSMUSG00000021361 | transmembrane protein 14C [Source:MGI Symbol;Acc:MGI:1913404] | Tmem14c | Enriched in dLGN | 1.483 | 1.979E-07 | 47.27 |
| ENSMUSG00000019689 | formation of mitochondrial complex V assembly factor 1 [Source:MGI Symbol;Acc:MGI:1913367] | Fmc1 | Enriched in dLGN | 1.479 | 0.0006553 | 21.78 |
| ENSMUSG00000020514 | mitochondrial ribosomal protein L22 [Source:MGI Symbol;Acc:MGI:1333794] | Mrpl22 | Enriched in dLGN | 1.478 | 0.000878 | 23.4 |
| ENSMUSG00000027248 | protein disulfide isomerase associated 3 [Source:MGI Symbol;Acc:MGI:95834] | Pdia3 | Enriched in dLGN | 1.476 | 5.046E-07 | 401 |
| ENSMUSG00000026585 | kinesin-associated protein 3 [Source:MGI Symbol;Acc:MGI:107566] | Kifap3 | Enriched in dLGN | 1.475 | 5.923E-08 | 663.9 |
| ENSMUSG00000035000 | dipeptidylpeptidase 4 [Source:MGI Symbol;Acc:MGI:94919] | Dpp4 | Enriched in dLGN | 1.475 | 0.02962 | 9.243 |
| ENSMUSG00000032966 | FK506 binding protein 1a [Source:MGI Symbol;Acc:MGI:95541] | Fkbp1a | Enriched in dLGN | 1.473 | 2.256E-10 | 430.5 |
| ENSMUSG00000045624 | ESF1 nucleolar pre-rRNA processing protein homolog [Source:MGI Symbol;Acc:MGI:1913830] | Esf1 | Enriched in dLGN | 1.471 | 1.849E-07 | 125.5 |
| ENSMUSG00000020089 | pyrophosphatase (inorganic) 1 [Source:MGI Symbol;Acc:MGI:97831] | Ppa1 | Enriched in dLGN | 1.469 | 0.0005296 | 125.6 |
| ENSMUSG00000036955 | kinesin family binding protein [Source:MGI Symbol;Acc:MGI:1919570] | Kifbp | Enriched in dLGN | 1.467 | 1.5E-10 | 286.8 |
| ENSMUSG00000035198 | tubulin, gamma 1 [Source:MGI Symbol;Acc:MGI:101834] | Tubg1 | Enriched in dLGN | 1.466 | 0.0003968 | 105.4 |
| ENSMUSG00000034171 | fatty acid amide hydrolase [Source:MGI Symbol;Acc:MGI:109609] | Faah | Enriched in dLGN | 1.464 | 2.369E-13 | 134 |
| ENSMUSG00000036748 | CUE domain containing 2 [Source:MGI Symbol;Acc:MGI:1914366] | Cuedc2 | Enriched in dLGN | 1.464 | 3.345E-08 | 145.7 |
| ENSMUSG00000019373 | COP9 signalosome subunit 3 [Source:MGI Symbol;Acc:MGI:1349409] | Cops3 | Enriched in dLGN | 1.462 | 0.00003381 | 92.31 |
| ENSMUSG00000022564 | glutamate receptor, ionotropic, N-methyl D-aspartate-associated protein 1 (glutamate binding) [Source:MGI Symbol;Acc:MGI:1913418] | Grina | Enriched in dLGN | 1.46 | 1.671E-09 | 437.4 |
| ENSMUSG00000024120 | leucine-rich PPR-motif containing [Source:MGI Symbol;Acc:MGI:1919666] | Lrpprc | Enriched in dLGN | 1.459 | 5.887E-14 | 294.4 |
| ENSMUSG00000024777 | protein phosphatase 2, regulatory subunit B', beta [Source:MGI Symbol;Acc:MGI:2388480] | Ppp2r5b | Enriched in dLGN | 1.458 | 2.88E-16 | 428 |
| ENSMUSG00000000056 | nuclear prelamin A recognition factor [Source:MGI Symbol;Acc:MGI:1914858] | Narf | Enriched in dLGN | 1.458 | 7.049E-15 | 273.8 |
| ENSMUSG00000032267 | ubiquitin specific peptidase 28 [Source:MGI Symbol;Acc:MGI:2442293] | Usp28 | Enriched in dLGN | 1.457 | 0.000006275 | 93.89 |
| ENSMUSG00000033294 | NOC4 like [Source:MGI Symbol;Acc:MGI:2140843] | Noc4l | Enriched in dLGN | 1.457 | 0.0001494 | 33.71 |
| ENSMUSG00000023033 | sodium channel, voltage-gated, type VIII, alpha [Source:MGI Symbol;Acc:MGI:103169] | Scn8a | Enriched in dLGN | 1.455 | 1.021E-13 | 1192 |
| ENSMUSG00000026824 | potassium inwardly-rectifying channel, subfamily J, member 3 [Source:MGI Symbol;Acc:MGI:104742] | Kcnj3 | Enriched in dLGN | 1.454 | 0.00000627 | 206.8 |
| ENSMUSG00000029388 | eukaryotic translation initiation factor 2B, subunit 1 (alpha) [Source:MGI Symbol;Acc:MGI:2384802] | Eif2b1 | Enriched in dLGN | 1.453 | 0.00002521 | 40.06 |
| ENSMUSG00000031603 | fibroblast growth factor 20 [Source:MGI Symbol;Acc:MGI:1891346] | Fgf20 | Enriched in dLGN | 1.45 | 0.04455 | 11.6 |
| ENSMUSG00000016308 | ubiquitin-conjugating enzyme E2A [Source:MGI Symbol;Acc:MGI:102959] | Ube2a | Enriched in dLGN | 1.448 | 8.857E-07 | 44.52 |
| ENSMUSG00000031533 | mitochondrial ribosomal protein S31 [Source:MGI Symbol;Acc:MGI:1913153] | Mrps31 | Enriched in dLGN | 1.445 | 0.00007604 | 35.82 |
| ENSMUSG00000022706 | mitochondrial ribosomal protein L40 [Source:MGI Symbol;Acc:MGI:1332635] | Mrpl40 | Enriched in dLGN | 1.444 | 0.000000631 | 48.44 |
| ENSMUSG00000025410 | dynactin 2 [Source:MGI Symbol;Acc:MGI:107733] | Dctn2 | Enriched in dLGN | 1.443 | 3.27E-14 | 382.2 |
| ENSMUSG00000027170 | eukaryotic translation initiation factor 3, subunit M [Source:MGI Symbol;Acc:MGI:1351744] | Eif3m | Enriched in dLGN | 1.44 | 4.553E-09 | 71.19 |
| ENSMUSG00000029657 | heat shock 105kDa/110kDa protein 1 [Source:MGI Symbol;Acc:MGI:105053] | Hsph1 | Enriched in dLGN | 1.44 | 4.318E-07 | 555.2 |
| ENSMUSG00000046798 | claudin 12 [Source:MGI Symbol;Acc:MGI:1929288] | Cldn12 | Enriched in dLGN | 1.439 | 2.062E-10 | 124.6 |
| ENSMUSG00000003380 | Rab acceptor 1 (prenylated) [Source:MGI Symbol;Acc:MGI:1201692] | Rabac1 | Enriched in dLGN | 1.439 | 4.884E-10 | 110.7 |
| ENSMUSG00000020458 | reticulon 4 [Source:MGI Symbol;Acc:MGI:1915835] | Rtn4 | Enriched in dLGN | 1.439 | 0.000003323 | 1440 |
| ENSMUSG00000067455 | H4 clustered histone 11 [Source:MGI Symbol;Acc:MGI:2448436] | H4c11 | Enriched in dLGN | 1.438 | 0.04612 | 5.483 |
| ENSMUSG00000024887 | N-acylsphingosine amidohydrolase 2 [Source:MGI Symbol;Acc:MGI:1859310] | Asah2 | Enriched in dLGN | 1.436 | 0.001425 | 39.79 |
| ENSMUSG00000029610 | aminoacyl tRNA synthetase complex-interacting multifunctional protein 2 [Source:MGI Symbol;Acc:MGI:2385237] | Aimp2 | Enriched in dLGN | 1.435 | 0.001078 | 26.12 |
| ENSMUSG00000026833 | olfactomedin 1 [Source:MGI Symbol;Acc:MGI:1860437] | Olfm1 | Enriched in dLGN | 1.432 | 7.401E-09 | 1863 |
| ENSMUSG00000021548 | cyclin H [Source:MGI Symbol;Acc:MGI:1913921] | Ccnh | Enriched in dLGN | 1.429 | 0.00004828 | 69.07 |
| ENSMUSG00000040048 | NADH:ubiquinone oxidoreductase subunit B10 [Source:MGI Symbol;Acc:MGI:1915592] | Ndufb10 | Enriched in dLGN | 1.428 | 2.863E-08 | 201.2 |
| ENSMUSG00000022450 | NADH:ubiquinone oxidoreductase subunit A6 [Source:MGI Symbol;Acc:MGI:1914380] | Ndufa6 | Enriched in dLGN | 1.427 | 1.72E-08 | 126.1 |
| ENSMUSG00000053329 | glutamine amidotransferase like class 1 domain containing 3A [Source:MGI Symbol;Acc:MGI:1351861] | Gatd3a | Enriched in dLGN | 1.426 | 8.78E-10 | 113.5 |
| ENSMUSG00000033389 | Rho GTPase activating protein 44 [Source:MGI Symbol;Acc:MGI:2144423] | Arhgap44 | Enriched in dLGN | 1.425 | 1.571E-12 | 522.2 |
| ENSMUSG00000026959 | glutamate receptor, ionotropic, NMDA1 (zeta 1) [Source:MGI Symbol;Acc:MGI:95819] | Grin1 | Enriched in dLGN | 1.425 | 0.000005101 | 942.6 |
| ENSMUSG00000026456 | cytochrome b5 reductase 1 [Source:MGI Symbol;Acc:MGI:1919267] | Cyb5r1 | Enriched in dLGN | 1.424 | 0.01025 | 33.56 |
| ENSMUSG00000037072 | selenoprotein F [Source:MGI Symbol;Acc:MGI:1927947] | Selenof | Enriched in dLGN | 1.415 | 1.802E-09 | 179.7 |
| ENSMUSG00000026750 | proteasome (prosome, macropain) subunit, beta type 7 [Source:MGI Symbol;Acc:MGI:107637] | Psmb7 | Enriched in dLGN | 1.415 | 0.000000324 | 169.1 |
| ENSMUSG00000025240 | SAC1 suppressor of actin mutations 1-like (yeast) [Source:MGI Symbol;Acc:MGI:1933169] | Sacm1l | Enriched in dLGN | 1.413 | 0.00001058 | 163.5 |
| ENSMUSG00000018446 | complement component 1, q subcomponent binding protein [Source:MGI Symbol;Acc:MGI:1194505] | C1qbp | Enriched in dLGN | 1.41 | 0.00000713 | 91.52 |
| ENSMUSG00000027293 | EH-domain containing 4 [Source:MGI Symbol;Acc:MGI:1919619] | Ehd4 | Enriched in dLGN | 1.41 | 0.00001224 | 74.05 |
| ENSMUSG00000079941 | cytochrome c oxidase subunit 5B, pseudogene [Source:MGI Symbol;Acc:MGI:3649411] | Cox5b-ps | Enriched in dLGN | 1.402 | 0.001336 | 35.35 |
| ENSMUSG00000034088 | high density lipoprotein (HDL) binding protein [Source:MGI Symbol;Acc:MGI:99256] | Hdlbp | Enriched in dLGN | 1.401 | 2.867E-12 | 727.2 |
| ENSMUSG00000013160 | ATPase, H+ transporting, lysosomal V0 subunit D1 [Source:MGI Symbol;Acc:MGI:1201778] | Atp6v0d1 | Enriched in dLGN | 1.399 | 4.924E-08 | 268.5 |
| ENSMUSG00000021037 | AHA1, activator of heat shock protein ATPase 1 [Source:MGI Symbol;Acc:MGI:2387603] | Ahsa1 | Enriched in dLGN | 1.398 | 1.162E-12 | 199 |
| ENSMUSG00000076432 | tyrosine 3-monooxygenase/tryptophan 5-monooxygenase activation protein theta [Source:MGI Symbol;Acc:MGI:891963] | Ywhaq | Enriched in dLGN | 1.397 | 0.00001589 | 236.1 |
| ENSMUSG00000019494 | COP9 signalosome subunit 6 [Source:MGI Symbol;Acc:MGI:1349439] | Cops6 | Enriched in dLGN | 1.396 | 0.0000491 | 134 |
| ENSMUSG00000021314 | amphiphysin [Source:MGI Symbol;Acc:MGI:103574] | Amph | Enriched in dLGN | 1.394 | 2.857E-16 | 841.6 |
| ENSMUSG00000020869 | leucine rich repeat containing 59 [Source:MGI Symbol;Acc:MGI:2138133] | Lrrc59 | Enriched in dLGN | 1.394 | 2.022E-10 | 321.9 |
| ENSMUSG00000026492 | transcription factor B2, mitochondrial [Source:MGI Symbol;Acc:MGI:107937] | Tfb2m | Enriched in dLGN | 1.393 | 0.0004515 | 22.6 |
| ENSMUSG00000025630 | hypoxanthine guanine phosphoribosyl transferase [Source:MGI Symbol;Acc:MGI:96217] | Hprt | Enriched in dLGN | 1.391 | 2.876E-09 | 174.6 |
| ENSMUSG00000020333 | acyl-CoA synthetase long-chain family member 6 [Source:MGI Symbol;Acc:MGI:894291] | Acsl6 | Enriched in dLGN | 1.39 | 7.232E-10 | 594.1 |
| ENSMUSG00000046182 | GSG1-like [Source:MGI Symbol;Acc:MGI:2685483] | Gsg1l | Enriched in dLGN | 1.39 | 6.032E-09 | 471.7 |
| ENSMUSG00000025757 | heat shock protein 4 like [Source:MGI Symbol;Acc:MGI:107422] | Hspa4l | Enriched in dLGN | 1.389 | 2.427E-07 | 545.2 |

| | | | | | | |
|---|---|---|---|---|---|---|
| ENSMUSG00000019861 | golgi associated PDZ and coiled-coil motif containing [Source:MGI Symbol;Acc:MGI:2149946] | Gopc | Enriched in dLGN | 1.388 | 6.807E-18 | 304.7 |
| ENSMUSG00000006024 | N-ethylmaleimide sensitive fusion protein attachment protein alpha [Source:MGI Symbol;Acc:MGI:104563] | Napa | Enriched in dLGN | 1.387 | 1.445E-11 | 230.6 |
| ENSMUSG00000022962 | phosphoribosylglycinamide formyltransferase [Source:MGI Symbol;Acc:MGI:95654] | Gart | Enriched in dLGN | 1.387 | 4.259E-08 | 92.71 |
| ENSMUSG00000052738 | succinate-CoA ligase, GDP-forming, alpha subunit [Source:MGI Symbol;Acc:MGI:1927234] | Suclg1 | Enriched in dLGN | 1.386 | 0.0000608 | 138.6 |
| ENSMUSG00000019849 | prolyl endopeptidase [Source:MGI Symbol;Acc:MGI:1270863] | Prep | Enriched in dLGN | 1.383 | 7.853E-07 | 207.5 |
| ENSMUSG00000021660 | basic transcription factor 3 [Source:MGI Symbol;Acc:MGI:1202875] | Btf3 | Enriched in dLGN | 1.38 | 0.000001073 | 71.1 |
| ENSMUSG00000041020 | MAP7 domain containing 2 [Source:MGI Symbol;Acc:MGI:1917474] | Map7d2 | Enriched in dLGN | 1.378 | 4.954E-08 | 736.5 |
| ENSMUSG00000021764 | NADH:ubiquinone oxidoreductase core subunit S4 [Source:MGI Symbol;Acc:MGI:1343135] | Ndufs4 | Enriched in dLGN | 1.378 | 0.000001443 | 153.5 |
| ENSMUSG00000025189 | cyclin M1 [Source:MGI Symbol;Acc:MGI:1891366] | Cnnm1 | Enriched in dLGN | 1.378 | 0.00001315 | 258.6 |
| ENSMUSG00000026825 | dynamin 1 [Source:MGI Symbol;Acc:MGI:107384] | Dnm1 | Enriched in dLGN | 1.376 | 2.119E-08 | 2096 |
| ENSMUSG00000021737 | proteasome (prosome, macropain) 26S subunit, non-ATPase, 6 [Source:MGI Symbol;Acc:MGI:1913663] | Psmd6 | Enriched in dLGN | 1.376 | 0.00000123 | 147.1 |
| ENSMUSG00000028470 | histidine triad nucleotide binding protein 2 [Source:MGI Symbol;Acc:MGI:1916167] | Hint2 | Enriched in dLGN | 1.375 | 0.001021 | 22.68 |
| ENSMUSG00000000339 | RNA 3'-terminal phosphate cyclase [Source:MGI Symbol;Acc:MGI:1913618] | Rtca | Enriched in dLGN | 1.374 | 1.569E-08 | 93.2 |
| ENSMUSG00000042682 | selenoprotein K [Source:MGI Symbol;Acc:MGI:1931466] | Selenok | Enriched in dLGN | 1.374 | 0.000128 | 254.3 |
| ENSMUSG00000024158 | hydroxyacyl glutathione hydrolase [Source:MGI Symbol;Acc:MGI:95745] | Hagh | Enriched in dLGN | 1.373 | 0.0000225 | 60.65 |
| ENSMUSG00000027963 | exostosin-like glycosyltransferase 2 [Source:MGI Symbol;Acc:MGI:1889574] | Extl2 | Enriched in dLGN | 1.372 | 6.441E-08 | 151.6 |
| ENSMUSG00000050856 | ATP synthase, H+ transporting, mitochondrial F1F0 complex, subunit E [Source:MGI Symbol;Acc:MGI:106636] | Atp5k | Enriched in dLGN | 1.372 | 0.0005749 | 126.9 |
| ENSMUSG00000028555 | tetratricopeptide repeat domain 39A [Source:MGI Symbol;Acc:MGI:2444350] | Ttc39a | Enriched in dLGN | 1.37 | 0.001041 | 25.31 |
| ENSMUSG00000025035 | ADP-ribosylation factor-like 3 [Source:MGI Symbol;Acc:MGI:1929699] | Arl3 | Enriched in dLGN | 1.366 | 9.247E-09 | 316.8 |
| ENSMUSG00000042660 | WD repeat domain 55 [Source:MGI Symbol;Acc:MGI:1915186] | Wdr55 | Enriched in dLGN | 1.364 | 0.0002742 | 26.69 |
| ENSMUSG00000026664 | phytanoyl-CoA hydroxylase [Source:MGI Symbol;Acc:MGI:891978] | Phyh | Enriched in dLGN | 1.363 | 7.705E-07 | 93.1 |
| ENSMUSG00000026709 | aspartyl-tRNA synthetase 2 (mitochondrial) [Source:MGI Symbol;Acc:MGI:2442510] | Dars2 | Enriched in dLGN | 1.363 | 0.00007126 | 36.15 |
| ENSMUSG00000031309 | ribosomal protein S6 kinase polypeptide 3 [Source:MGI Symbol;Acc:MGI:104557] | Rps6ka3 | Enriched in dLGN | 1.361 | 0.000006848 | 168.6 |
| ENSMUSG00000071528 | ATP synthase membrane subunit DAPIT [Source:MGI Symbol;Acc:MGI:1891435] | Atp5md | Enriched in dLGN | 1.361 | 0.00112 | 71.32 |
| ENSMUSG00000032398 | small nuclear RNA activating complex, polypeptide 5 [Source:MGI Symbol;Acc:MGI:1914282] | Snapc5 | Enriched in dLGN | 1.361 | 0.01208 | 21.39 |
| ENSMUSG00000040389 | WD repeat domain 47 [Source:MGI Symbol;Acc:MGI:2139593] | Wdr47 | Enriched in dLGN | 1.359 | 9.6E-10 | 344.7 |
| ENSMUSG00000031820 | BRISC and BRCA1 A complex member 1 [Source:MGI Symbol;Acc:MGI:1915501] | Babam1 | Enriched in dLGN | 1.359 | 3.05E-09 | 108.4 |
| ENSMUSG00000060904 | ADP-ribosylation factor-like 1 [Source:MGI Symbol;Acc:MGI:99436] | Arl1 | Enriched in dLGN | 1.358 | 0.000004148 | 131.5 |
| ENSMUSG00000028351 | bone morphogenic protein/retinoic acid inducible neural specific 1 [Source:MGI Symbol;Acc:MGI:1928478] | Brinp1 | Enriched in dLGN | 1.358 | 0.00003187 | 586.1 |
| ENSMUSG00000026188 | transmembrane protein 169 [Source:MGI Symbol;Acc:MGI:2442781] | Tmem169 | Enriched in dLGN | 1.354 | 3.264E-09 | 139.3 |
| ENSMUSG00000005357 | solute carrier family 1 (high affinity aspartate/glutamate transporter), member 6 [Source:MGI Symbol;Acc:MGI:1096331] | Slc1a6 | Enriched in dLGN | 1.353 | 0.0002689 | 52.8 |
| ENSMUSG00000031950 | gamma-aminobutyric acid (GABA) A receptor-associated protein-like 2 [Source:MGI Symbol;Acc:MGI:1890602] | Gabarapl2 | Enriched in dLGN | 1.351 | 0.000000121 | 147.2 |
| ENSMUSG00000028062 | late endosomal/lysosomal adaptor, MAPK and MTOR activator 2 [Source:MGI Symbol;Acc:MGI:1932697] | Lamtor2 | Enriched in dLGN | 1.351 | 0.00005131 | 47.8 |
| ENSMUSG00000022231 | sema domain, seven thrombospondin repeats (type 1 and type 1-like), transmembrane domain (TM) and short cytoplasmic domain, (semaphorin) 5A [Source:MGI Symbol;Acc:MGI:10 | Sema5a | Enriched in dLGN | 1.35 | 0.000001963 | 795.5 |
| ENSMUSG00000028318 | polymerase (RNA) I polypeptide E [Source:MGI Symbol;Acc:MGI:1929022] | Polr1e | Enriched in dLGN | 1.35 | 0.0001626 | 28.6 |
| ENSMUSG00000049556 | leucine rich repeat and Ig domain containing 1 [Source:MGI Symbol;Acc:MGI:1915522] | Lingo1 | Enriched in dLGN | 1.348 | 3.33E-10 | 811.2 |
| ENSMUSG00000041444 | Rho GTPase activating protein 32 [Source:MGI Symbol;Acc:MGI:2450166] | Arhgap32 | Enriched in dLGN | 1.348 | 1.07E-09 | 970.5 |
| ENSMUSG00000026766 | methylmalonic aciduria (cobalamin deficiency) cblD type, with homocystinuria [Source:MGI Symbol;Acc:MGI:1923786] | Mmadhc | Enriched in dLGN | 1.348 | 0.00001181 | 58.08 |
| ENSMUSG00000057388 | mitochondrial ribosomal protein L18 [Source:MGI Symbol;Acc:MGI:1914931] | Mrpl18 | Enriched in dLGN | 1.345 | 0.000004857 | 67.46 |
| ENSMUSG00000058355 | ATP-binding cassette, sub-family E (OABP), member 1 [Source:MGI Symbol;Acc:MGI:1195458] | Abce1 | Enriched in dLGN | 1.344 | 0.00003185 | 169.9 |
| ENSMUSG00000058267 | mitochondrial ribosomal protein S14 [Source:MGI Symbol;Acc:MGI:1928141] | Mrps14 | Enriched in dLGN | 1.344 | 0.00007261 | 45.2 |
| ENSMUSG00000059991 | neuronal pentraxin 2 [Source:MGI Symbol;Acc:MGI:1858209] | Nptx2 | Enriched in dLGN | 1.343 | 0.00005709 | 106.4 |
| ENSMUSG00000032883 | acyl-CoA synthetase long-chain family member 3 [Source:MGI Symbol;Acc:MGI:1921455] | Acsl3 | Enriched in dLGN | 1.338 | 7.628E-07 | 522.5 |
| ENSMUSG00000047547 | clathrin, light polypeptide (Lcb) [Source:MGI Symbol;Acc:MGI:1921575] | Cltb | Enriched in dLGN | 1.337 | 2.15E-19 | 896 |
| ENSMUSG00000031167 | RNA binding motif (RNP1, RRM) protein 3 [Source:MGI Symbol;Acc:MGI:1099460] | Rbm3 | Enriched in dLGN | 1.337 | 0.0002321 | 106.4 |
| ENSMUSG00000039983 | coiled-coil domain containing 32 [Source:MGI Symbol;Acc:MGI:2685477] | Ccdc32 | Enriched in dLGN | 1.336 | 0.000009857 | 49.02 |
| ENSMUSG00000026087 | mitochondrial ribosomal protein L30 [Source:MGI Symbol;Acc:MGI:1333820] | Mrpl30 | Enriched in dLGN | 1.336 | 0.00001395 | 45.34 |
| ENSMUSG00000019828 | glutamate receptor, metabotropic 1 [Source:MGI Symbol;Acc:MGI:1351338] | Grm1 | Enriched in dLGN | 1.335 | 0.00006539 | 572.7 |
| ENSMUSG00000042870 | target of myb1 trafficking protein [Source:MGI Symbol;Acc:MGI:1338026] | Tom1 | Enriched in dLGN | 1.335 | 0.005362 | 36.77 |
| ENSMUSG00000079108 | signal recognition particle 54C [Source:MGI Symbol;Acc:MGI:3714359] | Srp54c | Enriched in dLGN | 1.33 | 0.01521 | 21.13 |
| ENSMUSG00000020349 | protein phosphatase 2 (formerly 2A), catalytic subunit, alpha isoform [Source:MGI Symbol;Acc:MGI:1321159] | Ppp2ca | Enriched in dLGN | 1.329 | 4.093E-08 | 828.3 |
| ENSMUSG00000049176 | FERM and PDZ domain containing 4 [Source:MGI Symbol;Acc:MGI:3042378] | Frmpd4 | Enriched in dLGN | 1.329 | 0.0001929 | 381.5 |
| ENSMUSG00000058400 | pyroglutamylated RFamide peptide receptor [Source:MGI Symbol;Acc:MGI:2677633] | Qrfpr | Enriched in dLGN | 1.328 | 0.01456 | 24.15 |
| ENSMUSG00000021814 | annexin A7 [Source:MGI Symbol;Acc:MGI:88031] | Anxa7 | Enriched in dLGN | 1.327 | 0.0001651 | 56.22 |
| ENSMUSG00000040462 | amplified in osteosarcoma [Source:MGI Symbol;Acc:MGI:1924301] | Os9 | Enriched in dLGN | 1.324 | 2.272E-09 | 698.2 |
| ENSMUSG00000037149 | DEAD box helicase 1 [Source:MGI Symbol;Acc:MGI:2144727] | Ddx1 | Enriched in dLGN | 1.323 | 2.494E-09 | 410.1 |
| ENSMUSG00000071654 | ubiquinol-cytochrome c reductase complex assembly factor 3 [Source:MGI Symbol;Acc:MGI:2147553] | Uqcc3 | Enriched in dLGN | 1.323 | 0.000005957 | 94.54 |
| ENSMUSG00000007891 | cathepsin D [Source:MGI Symbol;Acc:MGI:88562] | Ctsd | Enriched in dLGN | 1.323 | 0.009653 | 65.28 |
| ENSMUSG00000031171 | FtsJ RNA methyltransferase homolog 1 (E. coli) [Source:MGI Symbol;Acc:MGI:1859648] | Ftsj1 | Enriched in dLGN | 1.322 | 0.004908 | 28.88 |
| ENSMUSG00000063253 | short coiled-coil protein [Source:MGI Symbol;Acc:MGI:1927654] | Scoc | Enriched in dLGN | 1.32 | 1.029E-07 | 113.5 |
| ENSMUSG00000028811 | tyrosyl-tRNA synthetase [Source:MGI Symbol;Acc:MGI:2147627] | Yars | Enriched in dLGN | 1.319 | 1.011E-21 | 315 |
| ENSMUSG00000031753 | component of oligomeric golgi complex 4 [Source:MGI Symbol;Acc:MGI:2142808] | Cog4 | Enriched in dLGN | 1.319 | 4.553E-09 | 122.4 |
| ENSMUSG00000020397 | mediator complex subunit 7 [Source:MGI Symbol;Acc:MGI:1913463] | Med7 | Enriched in dLGN | 1.318 | 0.002939 | 26.83 |
| ENSMUSG00000057315 | Rho GTPase activating protein 24 [Source:MGI Symbol;Acc:MGI:1922647] | Arhgap24 | Enriched in dLGN | 1.317 | 0.00006621 | 291.9 |
| ENSMUSG00000037772 | mitochondrial ribosomal protein L23 [Source:MGI Symbol;Acc:MGI:1196612] | Mrpl23 | Enriched in dLGN | 1.316 | 0.01235 | 20.4 |
| ENSMUSG00000039431 | myotubularin related protein 7 [Source:MGI Symbol;Acc:MGI:1891693] | Mtmr7 | Enriched in dLGN | 1.315 | 9.952E-08 | 199.1 |
| ENSMUSG00000021917 | signal peptidase complex subunit 1 homolog (S. cerevisiae) [Source:MGI Symbol;Acc:MGI:1916269] | Spcs1 | Enriched in dLGN | 1.315 | 0.0003995 | 46.82 |
| ENSMUSG00000030609 | apoptosis enhancing nuclease [Source:MGI Symbol;Acc:MGI:1915298] | Aen | Enriched in dLGN | 1.313 | 6.032E-09 | 72.83 |
| ENSMUSG00000049760 | mitochondrial contact site and cristae organizing system subunit 13 [Source:MGI Symbol;Acc:MGI:2442174] | Micos13 | Enriched in dLGN | 1.31 | 0.00001312 | 44.94 |
| ENSMUSG00000020544 | cytochrome c oxidase assembly protein 11, copper chaperone [Source:MGI Symbol;Acc:MGI:1917052] | Cox11 | Enriched in dLGN | 1.31 | 0.00004893 | 38.98 |
| ENSMUSG00000017188 | cytochrome C oxidase assembly factor 3 [Source:MGI Symbol;Acc:MGI:1098757] | Coa3 | Enriched in dLGN | 1.31 | 0.0000499 | 48.29 |
| ENSMUSG00000025287 | acyl-CoA thioesterase 9 [Source:MGI Symbol;Acc:MGI:1928939] | Acot9 | Enriched in dLGN | 1.31 | 0.01668 | 23.21 |
| ENSMUSG00000054428 | ATPase inhibitory factor 1 [Source:MGI Symbol;Acc:MGI:1196457] | Atpif1 | Enriched in dLGN | 1.309 | 0.000003872 | 829.4 |
| ENSMUSG00000029617 | CCZ1 vacuolar protein trafficking and biogenesis associated [Source:MGI Symbol;Acc:MGI:2141070] | Ccz1 | Enriched in dLGN | 1.308 | 5.951E-09 | 122.9 |
| ENSMUSG00000056486 | chimerin 1 [Source:MGI Symbol;Acc:MGI:1915674] | Chn1 | Enriched in dLGN | 1.305 | 1.185E-14 | 504.4 |
| ENSMUSG00000022295 | ATPase, H+ transporting, lysosomal V1 subunit C1 [Source:MGI Symbol;Acc:MGI:1913585] | Atp6v1c1 | Enriched in dLGN | 1.304 | 8.731E-08 | 229 |
| ENSMUSG00000028843 | SH3 domain binding glutamic acid-rich protein-like 3 [Source:MGI Symbol;Acc:MGI:1920973] | Sh3bgrl3 | Enriched in dLGN | 1.304 | 0.004279 | 64.15 |
| ENSMUSG00000006050 | steroid receptor RNA activator 1 [Source:MGI Symbol;Acc:MGI:1344414] | Sra1 | Enriched in dLGN | 1.303 | 0.00008911 | 42.72 |
| ENSMUSG00000024217 | U1 small nuclear ribonucleoprotein C [Source:MGI Symbol;Acc:MGI:109489] | Snrpc | Enriched in dLGN | 1.302 | 0.004604 | 26.15 |
| ENSMUSG00000015672 | mitochondrial ribosomal protein L32 [Source:MGI Symbol;Acc:MGI:2137226] | Mrpl32 | Enriched in dLGN | 1.302 | 0.007889 | 27.23 |
| ENSMUSG00000051256 | jagunal homolog 1 [Source:MGI Symbol;Acc:MGI:1915017] | Jagn1 | Enriched in dLGN | 1.3 | 0.0009224 | 27.81 |
| ENSMUSG00000028964 | Parkinson disease (autosomal recessive, early onset) 7 [Source:MGI Symbol;Acc:MGI:2135637] | Park7 | Enriched in dLGN | 1.299 | 2.456E-08 | 178.9 |
| ENSMUSG00000052428 | transmembrane and coiled-coil domains 1 [Source:MGI Symbol;Acc:MGI:1921173] | Tmco1 | Enriched in dLGN | 1.299 | 0.03114 | 95.8 |
| ENSMUSG00000032324 | tetraspanin 3 [Source:MGI Symbol;Acc:MGI:1928098] | Tspan3 | Enriched in dLGN | 1.298 | 2.524E-07 | 417.4 |
| ENSMUSG00000021728 | embigin [Source:MGI Symbol;Acc:MGI:95321] | Emb | Enriched in dLGN | 1.295 | 0.0007353 | 88.85 |
| ENSMUSG00000028795 | coiled coil domain containing 28B [Source:MGI Symbol;Acc:MGI:1913514] | Ccdc28b | Enriched in dLGN | 1.292 | 0.000003206 | 101 |
| ENSMUSG00000032839 | transient receptor potential cation channel, subfamily C, member 1 [Source:MGI Symbol;Acc:MGI:109528] | Trpc1 | Enriched in dLGN | 1.291 | 4.484E-09 | 140.9 |
| ENSMUSG00000053192 | myeloid/lymphoid or mixed-lineage leukemia; translocated to, 11 [Source:MGI Symbol;Acc:MGI:1929671] | Mllt11 | Enriched in dLGN | 1.29 | 9.985E-08 | 499.1 |
| ENSMUSG00000028212 | cyclin E2 [Source:MGI Symbol;Acc:MGI:1329034] | Ccne2 | Enriched in dLGN | 1.29 | 0.02425 | 12.53 |
| ENSMUSG00000029634 | ring finger protein (C3H2C3 type) 6 [Source:MGI Symbol;Acc:MGI:1921382] | Rnf6 | Enriched in dLGN | 1.289 | 9.737E-10 | 280.9 |
| ENSMUSG00000024966 | stress-induced phosphoprotein 1 [Source:MGI Symbol;Acc:MGI:109130] | Stip1 | Enriched in dLGN | 1.288 | 0.000128 | 255 |
| ENSMUSG00000024359 | heat shock protein 9 [Source:MGI Symbol;Acc:MGI:96245] | Hspa9 | Enriched in dLGN | 1.287 | 0.00003311 | 574.8 |
| ENSMUSG00000071074 | Yip1 domain family, member 3 [Source:MGI Symbol;Acc:MGI:106280] | Yipf3 | Enriched in dLGN | 1.285 | 9.153E-08 | 108.8 |
| ENSMUSG00000027637 | RAB5 interacting factor [Source:MGI Symbol;Acc:MGI:1914638] | Rab5if | Enriched in dLGN | 1.284 | 0.000000361 | 59.45 |
| ENSMUSG00000023723 | mitochondrial ribosomal protein S23 [Source:MGI Symbol;Acc:MGI:1928138] | Mrps23 | Enriched in dLGN | 1.284 | 0.000168 | 39.53 |
| ENSMUSG00000015217 | high mobility group box 3 [Source:MGI Symbol;Acc:MGI:1098219] | Hmgb3 | Enriched in dLGN | 1.283 | 0.00003602 | 141 |
| ENSMUSG00000001794 | calpain, small subunit 1 [Source:MGI Symbol;Acc:MGI:88266] | Capns1 | Enriched in dLGN | 1.282 | 0.0001683 | 150.2 |
| ENSMUSG00000030706 | mitochondrial ribosomal protein L48 [Source:MGI Symbol;Acc:MGI:1289321] | Mrpl48 | Enriched in dLGN | 1.282 | 0.0003024 | 153.1 |
| ENSMUSG00000033436 | armadillo repeat containing, X-linked 2 [Source:MGI Symbol;Acc:MGI:1914666] | Armcx2 | Enriched in dLGN | 1.28 | 1.626E-08 | 262.7 |
| ENSMUSG00000024208 | ubiquinol-cytochrome c reductase complex assembly factor 2 [Source:MGI Symbol;Acc:MGI:1914517] | Uqcc2 | Enriched in dLGN | 1.277 | 0.00005998 | 101.2 |
| ENSMUSG00000045435 | transmembrane protein 60 [Source:MGI Symbol;Acc:MGI:2673965] | Tmem60 | Enriched in dLGN | 1.277 | 0.02036 | 14.74 |
| ENSMUSG00000032046 | abhydrolase domain containing 12 [Source:MGI Symbol;Acc:MGI:1923442] | Abhd12 | Enriched in dLGN | 1.275 | 2.087E-11 | 313.9 |
| ENSMUSG00000031770 | homocysteine-inducible, endoplasmic reticulum stress-inducible, ubiquitin-like domain member 1 [Source:MGI Symbol;Acc:MGI:1927406] | Herpud1 | Enriched in dLGN | 1.275 | 0.000001694 | 93.7 |
| ENSMUSG00000020267 | histidine triad nucleotide binding protein 1 [Source:MGI Symbol;Acc:MGI:1321133] | Hint1 | Enriched in dLGN | 1.275 | 0.000002064 | 255.7 |
| ENSMUSG00000010376 | neural precursor cell expressed, developmentally down-regulated gene 8 [Source:MGI Symbol;Acc:MGI:97301] | Nedd8 | Enriched in dLGN | 1.275 | 0.00000713 | 191.2 |
| ENSMUSG00000001366 | f-box protein 9 [Source:MGI Symbol;Acc:MGI:1918788] | Fbxo9 | Enriched in dLGN | 1.273 | 1.546E-18 | 320.2 |
| ENSMUSG00000043162 | Pigy upstream reading frame [Source:MGI Symbol;Acc:MGI:1913709] | Pyurf | Enriched in dLGN | 1.273 | 0.001576 | 52.19 |
| ENSMUSG00000028138 | alcohol dehydrogenase 5 (class III), chi polypeptide [Source:MGI Symbol;Acc:MGI:87929] | Adh5 | Enriched in dLGN | 1.27 | 0.00006964 | 68.21 |
| ENSMUSG00000082229 | nucleosome assembly protein 1-like 2 [Source:MGI Symbol;Acc:MGI:106654] | Nap1l2 | Enriched in dLGN | 1.27 | 0.0002074 | 277.1 |
| ENSMUSG00000018770 | ATP synthase, H+ transporting, mitochondrial F0 complex, subunit C3 (subunit 9) [Source:MGI Symbol;Acc:MGI:2442035] | Atp5g3 | Enriched in dLGN | 1.268 | 6.514E-07 | 265.3 |
| ENSMUSG00000024194 | cutA divalent cation tolerance homolog [Source:MGI Symbol;Acc:MGI:1914925] | Cuta | Enriched in dLGN | 1.266 | 0.000002207 | 66.24 |
| ENSMUSG00000026377 | nucleolar protein interacting with the FHA domain of MKI67 [Source:MGI Symbol;Acc:MGI:1915199] | Nifk | Enriched in dLGN | 1.266 | 0.0000176 | 61.02 |
| ENSMUSG00000030588 | Yip1 interacting factor homolog B (S. cerevisiae) [Source:MGI Symbol;Acc:MGI:1924504] | Yif1b | Enriched in dLGN | 1.265 | 0.0023 | 32.44 |
| ENSMUSG00000023939 | mitochondrial ribosomal protein L14 [Source:MGI Symbol;Acc:MGI:1333864] | Mrpl14 | Enriched in dLGN | 1.265 | 0.01909 | 19.2 |
| ENSMUSG00000027206 | COP9 signalosome subunit 2 [Source:MGI Symbol;Acc:MGI:1330276] | Cops2 | Enriched in dLGN | 1.264 | 5.177E-08 | 139.2 |
| ENSMUSG00000025981 | coenzyme Q10B [Source:MGI Symbol;Acc:MGI:1915126] | Coq10b | Enriched in dLGN | 1.264 | 0.00007318 | 31.52 |
| ENSMUSG00000021947 | crystallin, lambda 1 [Source:MGI Symbol;Acc:MGI:1915881] | Cryl1 | Enriched in dLGN | 1.264 | 0.002053 | 28.77 |
| ENSMUSG00000052504 | Eph receptor A3 [Source:MGI Symbol;Acc:MGI:99612] | Epha3 | Enriched in dLGN | 1.264 | 0.004257 | 122.1 |
| ENSMUSG00000032131 | ATP binding cassette subfamily G member 4 [Source:MGI Symbol;Acc:MGI:1890594] | Abcg4 | Enriched in dLGN | 1.262 | 1.133E-10 | 245 |
| ENSMUSG00000005981 | TNF receptor-associated protein 1 [Source:MGI Symbol;Acc:MGI:1915265] | Trap1 | Enriched in dLGN | 1.261 | 0.0002402 | 166.2 |
| ENSMUSG00000021250 | FBJ osteosarcoma oncogene [Source:MGI Symbol;Acc:MGI:95574] | Fos | Enriched in dLGN | 1.259 | 0.0002755 | 57.19 |
| ENSMUSG00000054034 | transcription elongation factor A (SII)-like 5 [Source:MGI Symbol;Acc:MGI:3036236] | Tceal5 | Enriched in dLGN | 1.259 | 0.0052 | 58.79 |
| ENSMUSG00000044573 | acid phosphatase 1, soluble [Source:MGI Symbol;Acc:MGI:87881] | Acp1 | Enriched in dLGN | 1.257 | 0.001108 | 31.27 |
| ENSMUSG00000001999 | biliverdin reductase A [Source:MGI Symbol;Acc:MGI:88170] | Blvra | Enriched in dLGN | 1.256 | 0.0003259 | 31.09 |
| ENSMUSG00000030316 | TAM41 mitochondrial translocator assembly and maintenance homolog [Source:MGI Symbol;Acc:MGI:1916221] | Tamm41 | Enriched in dLGN | 1.255 | 0.004716 | 26.14 |
| ENSMUSG00000017781 | phosphatidylinositol transfer protein, alpha [Source:MGI Symbol;Acc:MGI:99887] | Pitpna | Enriched in dLGN | 1.254 | 3.911E-11 | 792.9 |
| ENSMUSG00000036309 | S-phase kinase-associated protein 1 [Source:MGI Symbol;Acc:MGI:103575] | Skp1 | Enriched in dLGN | 1.252 | 0.00001742 | 534.7 |
| ENSMUSG00000004031 | bone morphogenic protein/retinoic acid inducible neural-specific 2 [Source:MGI Symbol;Acc:MGI:2443333] | Brinp2 | Enriched in dLGN | 1.249 | 0.000004322 | 323.9 |

| | | | | | | |
|---|---|---|---|---|---|---|
| ENSMUSG00000030286 | ER membrane protein complex subunit 3 [Source:MGI Symbol;Acc:MGI:1913337] | Emc3 | Enriched in dLGN | 1.248 | 8.544E-08 | 83 |
| ENSMUSG00000032064 | DIX domain containing 1 [Source:MGI Symbol;Acc:MGI:2679721] | Dixdc1 | Enriched in dLGN | 1.245 | 2.024E-08 | 533.3 |
| ENSMUSG00000004207 | prosaposin [Source:MGI Symbol;Acc:MGI:97783] | Psap | Enriched in dLGN | 1.242 | 0.00002942 | 1320 |
| ENSMUSG00000021939 | cathepsin B [Source:MGI Symbol;Acc:MGI:88561] | Ctsb | Enriched in dLGN | 1.242 | 0.000291 | 510.6 |
| ENSMUSG00000034932 | mitochondrial ribosomal protein L54 [Source:MGI Symbol;Acc:MGI:1913297] | Mrpl54 | Enriched in dLGN | 1.242 | 0.0008324 | 34.59 |
| ENSMUSG00000021816 | protein phosphatase 3, catalytic subunit, beta isoform [Source:MGI Symbol;Acc:MGI:107163] | Ppp3cb | Enriched in dLGN | 1.241 | 4.493E-14 | 533.5 |
| ENSMUSG00000000552 | zinc finger protein 385A [Source:MGI Symbol;Acc:MGI:1352495] | Zfp385a | Enriched in dLGN | 1.241 | 4.232E-08 | 618.4 |
| ENSMUSG00000052632 | ArfGAP with SH3 domain, ankyrin repeat and PH domain 2 [Source:MGI Symbol;Acc:MGI:2685438] | Asap2 | Enriched in dLGN | 1.239 | 2.027E-11 | 463.9 |
| ENSMUSG00000029198 | GrpE-like 1, mitochondrial [Source:MGI Symbol;Acc:MGI:1334417] | Grpel1 | Enriched in dLGN | 1.239 | 0.000007906 | 89.43 |
| ENSMUSG00000055681 | coatomer protein complex, subunit epsilon [Source:MGI Symbol;Acc:MGI:1891702] | Cope | Enriched in dLGN | 1.238 | 0.0001378 | 112.7 |
| ENSMUSG00000036398 | protein phosphatase 1, regulatory inhibitor subunit 11 [Source:MGI Symbol;Acc:MGI:1923747] | Ppp1r11 | Enriched in dLGN | 1.237 | 0.000001562 | 110.1 |
| ENSMUSG00000009894 | synaptosomal-associated protein, 47 [Source:MGI Symbol;Acc:MGI:1915076] | Snap47 | Enriched in dLGN | 1.237 | 0.00007901 | 533.6 |
| ENSMUSG00000020990 | cyclin-dependent kinase-like 1 (CDC2-related kinase) [Source:MGI Symbol;Acc:MGI:1918341] | Cdkl1 | Enriched in dLGN | 1.237 | 0.0007833 | 53.67 |
| ENSMUSG00000031556 | TM2 domain containing 2 [Source:MGI Symbol;Acc:MGI:1916992] | Tm2d2 | Enriched in dLGN | 1.236 | 0.000304 | 62.81 |
| ENSMUSG00000021486 | PRELI domain containing 1 [Source:MGI Symbol;Acc:MGI:1913744] | Prelid1 | Enriched in dLGN | 1.235 | 2.675E-07 | 247.5 |
| ENSMUSG00000021494 | DEAD box helicase 41 [Source:MGI Symbol;Acc:MGI:1920185] | Ddx41 | Enriched in dLGN | 1.235 | 0.000003542 | 122.3 |
| ENSMUSG00000019796 | low density lipoprotein receptor-related protein 11 [Source:MGI Symbol;Acc:MGI:2442989] | Lrp11 | Enriched in dLGN | 1.234 | 1.284E-07 | 271 |
| ENSMUSG00000034708 | granulin [Source:MGI Symbol;Acc:MGI:95832] | Grn | Enriched in dLGN | 1.234 | 0.0001453 | 113.5 |
| ENSMUSG00000062006 | ribosomal protein L34 [Source:MGI Symbol;Acc:MGI:1915686] | Rpl34 | Enriched in dLGN | 1.23 | 0.001505 | 135.1 |
| ENSMUSG00000031791 | transmembrane protein 38A [Source:MGI Symbol;Acc:MGI:1921416] | Tmem38a | Enriched in dLGN | 1.228 | 3.901E-14 | 149.4 |
| ENSMUSG00000020720 | proteasome (prosome, macropain) 26S subunit, non-ATPase, 12 [Source:MGI Symbol;Acc:MGI:1914247] | Psmd12 | Enriched in dLGN | 1.228 | 0.00004027 | 128.2 |
| ENSMUSG00000056222 | sparc/osteonectin, cwcv and kazal-like domains proteoglycan 1 [Source:MGI Symbol;Acc:MGI:105371] | Spock1 | Enriched in dLGN | 1.228 | 0.0001568 | 1204 |
| ENSMUSG00000035232 | pyruvate dehydrogenase kinase, isoenzyme 3 [Source:MGI Symbol;Acc:MGI:2384308] | Pdk3 | Enriched in dLGN | 1.228 | 0.001665 | 64.16 |
| ENSMUSG00000024232 | BMP and activin membrane-bound inhibitor [Source:MGI Symbol;Acc:MGI:1915260] | Bambi | Enriched in dLGN | 1.228 | 0.005784 | 19.07 |
| ENSMUSG00000050199 | leucine-rich repeat-containing G protein-coupled receptor 4 [Source:MGI Symbol;Acc:MGI:1891468] | Lgr4 | Enriched in dLGN | 1.226 | 3.961E-07 | 197.4 |
| ENSMUSG00000025613 | chaperonin containing Tcp1, subunit 8 (theta) [Source:MGI Symbol;Acc:MGI:107183] | Cct8 | Enriched in dLGN | 1.225 | 0.00005393 | 259.2 |
| ENSMUSG00000048930 | transcriptional adaptor 3 [Source:MGI Symbol;Acc:MGI:1915724] | Tada3 | Enriched in dLGN | 1.224 | 5.065E-08 | 106.8 |
| ENSMUSG00000028398 | distal membrane arm assembly complex 1 [Source:MGI Symbol;Acc:MGI:1914178] | Dmac1 | Enriched in dLGN | 1.222 | 0.0004511 | 32.88 |
| ENSMUSG00000006932 | catenin (cadherin associated protein), beta 1 [Source:MGI Symbol;Acc:MGI:88276] | Ctnnb1 | Enriched in dLGN | 1.22 | 0.000009618 | 1065 |
| ENSMUSG00000036834 | phospholipase C, eta 1 [Source:MGI Symbol;Acc:MGI:2683547] | Plch1 | Enriched in dLGN | 1.218 | 9.946E-07 | 269.3 |
| ENSMUSG00000038489 | polymerase (RNA) II (DNA directed) polypeptide L [Source:MGI Symbol;Acc:MGI:1913741] | Polr2l | Enriched in dLGN | 1.218 | 0.0004021 | 28.51 |
| ENSMUSG00000036067 | solute carrier family 2 (facilitated glucose transporter), member 6 [Source:MGI Symbol;Acc:MGI:2443286] | Slc2a6 | Enriched in dLGN | 1.218 | 0.001697 | 33.47 |
| ENSMUSG00000015092 | endothelial differentiation-related factor 1 [Source:MGI Symbol;Acc:MGI:1891227] | Edf1 | Enriched in dLGN | 1.214 | 3.399E-07 | 190.3 |
| ENSMUSG00000022964 | transmembrane protein 50B [Source:MGI Symbol;Acc:MGI:1925225] | Tmem50b | Enriched in dLGN | 1.214 | 0.000001173 | 152.6 |
| ENSMUSG00000074797 | inosine triphosphatase (nucleoside triphosphate pyrophosphatase) [Source:MGI Symbol;Acc:MGI:96622] | Itpa | Enriched in dLGN | 1.212 | 0.00003121 | 70.22 |
| ENSMUSG00000037428 | VGF nerve growth factor inducible [Source:MGI Symbol;Acc:MGI:1343180] | Vgf | Enriched in dLGN | 1.212 | 0.00003362 | 1249 |
| ENSMUSG00000033578 | transmembrane protein 35A [Source:MGI Symbol;Acc:MGI:1914814] | Tmem35a | Enriched in dLGN | 1.211 | 0.004956 | 71.47 |
| ENSMUSG00000041231 | ubiquitin-like domain containing CTD phosphatase 1 [Source:MGI Symbol;Acc:MGI:1933105] | Ublcp1 | Enriched in dLGN | 1.209 | 0.0007065 | 41.53 |
| ENSMUSG00000022634 | YY1 associated factor 2 [Source:MGI Symbol;Acc:MGI:1914307] | Yaf2 | Enriched in dLGN | 1.208 | 1.232E-08 | 232.8 |
| ENSMUSG00000017221 | proteasome (prosome, macropain) 26S subunit, non-ATPase, 3 [Source:MGI Symbol;Acc:MGI:98858] | Psmd3 | Enriched in dLGN | 1.207 | 1.349E-07 | 193.7 |
| ENSMUSG00000042032 | methionine adenosyltransferase II, beta [Source:MGI Symbol;Acc:MGI:1913667] | Mat2b | Enriched in dLGN | 1.207 | 9.292E-07 | 58.43 |
| ENSMUSG00000031609 | sin3 associated polypeptide [Source:MGI Symbol;Acc:MGI:1929129] | Sap30 | Enriched in dLGN | 1.207 | 0.009974 | 26.75 |
| ENSMUSG00000033460 | armadillo repeat containing, X-linked 1 [Source:MGI Symbol;Acc:MGI:1925498] | Armcx1 | Enriched in dLGN | 1.206 | 1.923E-07 | 200.9 |
| ENSMUSG00000056185 | sorting nexin 32 [Source:MGI Symbol;Acc:MGI:2444704] | Snx32 | Enriched in dLGN | 1.206 | 0.0022 | 101.2 |
| ENSMUSG00000049792 | BCL2-associated athanogene 5 [Source:MGI Symbol;Acc:MGI:1917619] | Bag5 | Enriched in dLGN | 1.204 | 7.924E-09 | 146 |
| ENSMUSG00000017801 | MAX-like protein X [Source:MGI Symbol;Acc:MGI:108398] | Mlx | Enriched in dLGN | 1.203 | 0.005511 | 24.55 |
| ENSMUSG00000004788 | eukaryotic translation initiation factor 2B, subunit 2 beta [Source:MGI Symbol;Acc:MGI:2145118] | Eif2b2 | Enriched in dLGN | 1.202 | 0.00002313 | 73.95 |
| ENSMUSG00000061479 | small nuclear ribonucleoprotein polypeptide A [Source:MGI Symbol;Acc:MGI:1855690] | Snrpa | Enriched in dLGN | 1.202 | 0.002466 | 69.88 |
| ENSMUSG00000036503 | ring finger protein 13 [Source:MGI Symbol;Acc:MGI:1346341] | Rnf13 | Enriched in dLGN | 1.201 | 0.000365 | 102.4 |
| ENSMUSG00000064202 | spermatogenesis associated 6 like [Source:MGI Symbol;Acc:MGI:1918036] | Spata6l | Enriched in dLGN | 1.201 | 0.005792 | 24.41 |
| ENSMUSG00000025487 | proteasome (prosome, macropain) 26S subunit, non-ATPase, 13 [Source:MGI Symbol;Acc:MGI:1345192] | Psmd13 | Enriched in dLGN | 1.198 | 1.72E-08 | 179.8 |
| ENSMUSG00000009013 | dynein light chain LC8-type 1 [Source:MGI Symbol;Acc:MGI:1861457] | Dynll1 | Enriched in dLGN | 1.197 | 0.000000109 | 315.9 |
| ENSMUSG00000033047 | eukaryotic translation initiation factor 3, subunit L [Source:MGI Symbol;Acc:MGI:2386251] | Eif3l | Enriched in dLGN | 1.197 | 0.0001906 | 114.6 |
| ENSMUSG00000029458 | BRCA1 associated protein [Source:MGI Symbol;Acc:MGI:1919649] | Brap | Enriched in dLGN | 1.196 | 1.254E-10 | 203.2 |
| ENSMUSG00000024403 | ATPase, H+ transporting, lysosomal V1 subunit G2 [Source:MGI Symbol;Acc:MGI:1913487] | Atp6v1g2 | Enriched in dLGN | 1.195 | 0.000001717 | 338.6 |
| ENSMUSG00000041670 | regulating synaptic membrane exocytosis 1 [Source:MGI Symbol;Acc:MGI:2152971] | Rims1 | Enriched in dLGN | 1.195 | 0.000003516 | 1881 |
| ENSMUSG00000027263 | tubulin, gamma complex associated protein 4 [Source:MGI Symbol;Acc:MGI:1196293] | Tubgcp4 | Enriched in dLGN | 1.194 | 0.000007715 | 80.84 |
| ENSMUSG00000024829 | mitochondrial ribosomal protein L21 [Source:MGI Symbol;Acc:MGI:2660674] | Mrpl21 | Enriched in dLGN | 1.194 | 0.00003747 | 44.52 |
| ENSMUSG00000087687 | PET100 homolog [Source:MGI Symbol;Acc:MGI:3615306] | Pet100 | Enriched in dLGN | 1.191 | 0.00005187 | 59.25 |
| ENSMUSG00000024309 | prefoldin subunit 6 [Source:MGI Symbol;Acc:MGI:95908] | Pfdn6 | Enriched in dLGN | 1.188 | 0.000007491 | 72.58 |
| ENSMUSG00000031357 | synapse associated protein 1 [Source:MGI Symbol;Acc:MGI:1914293] | Syap1 | Enriched in dLGN | 1.188 | 0.0002298 | 90.22 |
| ENSMUSG00000026577 | basic leucine zipper nuclear factor 1 [Source:MGI Symbol;Acc:MGI:1201607] | Blzf1 | Enriched in dLGN | 1.185 | 0.000008461 | 70.6 |
| ENSMUSG00000068882 | small RNA binding exonuclease protection factor La [Source:MGI Symbol;Acc:MGI:98423] | Ssb | Enriched in dLGN | 1.183 | 0.00001201 | 214.7 |
| ENSMUSG00000021218 | guanosine diphosphate (GDP) dissociation inhibitor 2 [Source:MGI Symbol;Acc:MGI:99845] | Gdi2 | Enriched in dLGN | 1.182 | 0.000001277 | 530.5 |
| ENSMUSG00000019977 | Hbs1-like (S. cerevisiae) [Source:MGI Symbol;Acc:MGI:1891704] | Hbs1l | Enriched in dLGN | 1.181 | 0.000004593 | 167.3 |
| ENSMUSG00000042807 | HECT, C2 and WW domain containing E3 ubiquitin protein ligase 2 [Source:MGI Symbol;Acc:MGI:2685817] | Hecw2 | Enriched in dLGN | 1.181 | 0.0003699 | 320.7 |
| ENSMUSG00000007653 | gamma-aminobutyric acid (GABA) A receptor, subunit beta 2 [Source:MGI Symbol;Acc:MGI:95620] | Gabrb2 | Enriched in dLGN | 1.181 | 0.001594 | 322.6 |
| ENSMUSG00000078695 | CDGSH iron sulfur domain 3 [Source:MGI Symbol;Acc:MGI:101788] | Cisd3 | Enriched in dLGN | 1.181 | 0.04077 | 13.41 |
| ENSMUSG00000025236 | ADP-dependent glucokinase [Source:MGI Symbol;Acc:MGI:1919391] | Adpgk | Enriched in dLGN | 1.178 | 0.00008376 | 56.67 |
| ENSMUSG00000038762 | ATP-binding cassette, sub-family F (GCN20), member 1 [Source:MGI Symbol;Acc:MGI:1351658] | Abcf1 | Enriched in dLGN | 1.177 | 1.237E-09 | 576.3 |
| ENSMUSG00000021285 | protein phosphatase 1, regulatory subunit 13B [Source:MGI Symbol;Acc:MGI:1336199] | Ppp1r13b | Enriched in dLGN | 1.177 | 2.357E-09 | 497.8 |
| ENSMUSG00000024516 | SEC11 homolog C, signal peptidase complex subunit [Source:MGI Symbol;Acc:MGI:1913536] | Sec11c | Enriched in dLGN | 1.177 | 0.00003106 | 79.39 |
| ENSMUSG00000029767 | calumenin [Source:MGI Symbol;Acc:MGI:1097158] | Calu | Enriched in dLGN | 1.175 | 0.00004828 | 300.9 |
| ENSMUSG00000014294 | NADH:ubiquinone oxidoreductase subunit A2 [Source:MGI Symbol;Acc:MGI:1343103] | Ndufa2 | Enriched in dLGN | 1.174 | 0.000003935 | 115.8 |
| ENSMUSG00000009863 | succinate dehydrogenase complex, subunit B, iron sulfur (Ip) [Source:MGI Symbol;Acc:MGI:1914930] | Sdhb | Enriched in dLGN | 1.172 | 0.00001983 | 134.3 |
| ENSMUSG00000044067 | G protein-coupled receptor 22 [Source:MGI Symbol;Acc:MGI:1920260] | Gpr22 | Enriched in dLGN | 1.17 | 0.006055 | 30.33 |
| ENSMUSG00000021116 | eukaryotic translation initiation factor 2, subunit 1 alpha [Source:MGI Symbol;Acc:MGI:95299] | Eif2s1 | Enriched in dLGN | 1.169 | 0.0007636 | 89.93 |
| ENSMUSG00000039910 | Cbp/p300-interacting transactivator, with Glu/Asp-rich carboxy-terminal domain, 2 [Source:MGI Symbol;Acc:MGI:1306784] | Cited2 | Enriched in dLGN | 1.168 | 0.00001837 | 198.6 |
| ENSMUSG00000027905 | DEAD box helicase 20 [Source:MGI Symbol;Acc:MGI:1858415] | Ddx20 | Enriched in dLGN | 1.168 | 0.001141 | 39.64 |
| ENSMUSG00000029840 | myotrophin [Source:MGI Symbol;Acc:MGI:99445] | Mtpn | Enriched in dLGN | 1.166 | 3.993E-07 | 335.2 |
| ENSMUSG00000020697 | ligase III, DNA, ATP-dependent [Source:MGI Symbol;Acc:MGI:109152] | Lig3 | Enriched in dLGN | 1.166 | 6.332E-07 | 109.2 |
| ENSMUSG00000002767 | mitochondrial ribosomal protein L2 [Source:MGI Symbol;Acc:MGI:1351622] | Mrpl2 | Enriched in dLGN | 1.165 | 0.00002691 | 55.02 |
| ENSMUSG00000115987 | vacuolar protein sorting 28 [Source:MGI Symbol;Acc:MGI:1914164] | Vps28 | Enriched in dLGN | 1.165 | 0.0001086 | 105.7 |
| ENSMUSG00000039480 | 5'-nucleotidase domain containing 1 [Source:MGI Symbol;Acc:MGI:2442446] | Nt5dc1 | Enriched in dLGN | 1.165 | 0.011 | 84.48 |
| ENSMUSG00000051223 | basic leucine zipper and W2 domains 1 [Source:MGI Symbol;Acc:MGI:1914132] | Bzw1 | Enriched in dLGN | 1.164 | 0.00004251 | 393.6 |
| ENSMUSG00000021022 | protein phosphatase 2, regulatory subunit B'', gamma [Source:MGI Symbol;Acc:MGI:1930009] | Ppp2r3c | Enriched in dLGN | 1.164 | 0.00205 | 32.63 |
| ENSMUSG00000027742 | component of oligomeric golgi complex 6 [Source:MGI Symbol;Acc:MGI:1914792] | Cog6 | Enriched in dLGN | 1.163 | 9.456E-07 | 89.91 |
| ENSMUSG00000054423 | Ca2+-dependent secretion activator [Source:MGI Symbol;Acc:MGI:1350922] | Cadps | Enriched in dLGN | 1.162 | 0.0001621 | 1472 |
| ENSMUSG00000002015 | B cell receptor associated protein 31 [Source:MGI Symbol;Acc:MGI:1350933] | Bcap31 | Enriched in dLGN | 1.161 | 0.00001623 | 84.18 |
| ENSMUSG00000031451 | growth arrest specific 6 [Source:MGI Symbol;Acc:MGI:95660] | Gas6 | Enriched in dLGN | 1.159 | 2.608E-10 | 424.5 |
| ENSMUSG00000030417 | programmed cell death 5 [Source:MGI Symbol;Acc:MGI:1913538] | Pdcd5 | Enriched in dLGN | 1.159 | 0.0001306 | 70.53 |
| ENSMUSG00000071369 | mitogen-activated protein kinase kinase kinase 5 [Source:MGI Symbol;Acc:MGI:1346876] | Map3k5 | Enriched in dLGN | 1.153 | 0.000005863 | 236 |
| ENSMUSG00000028261 | NADH:ubiquinone oxidoreductase complex assembly factor 4 [Source:MGI Symbol;Acc:MGI:1915743] | Ndufaf4 | Enriched in dLGN | 1.151 | 0.0002498 | 70.33 |
| ENSMUSG00000018287 | sperm associated antigen 7 [Source:MGI Symbol;Acc:MGI:107380] | Spag7 | Enriched in dLGN | 1.149 | 0.000006001 | 132.5 |
| ENSMUSG00000098274 | ribosomal protein L24 [Source:MGI Symbol;Acc:MGI:1915443] | Rpl24 | Enriched in dLGN | 1.148 | 0.0008863 | 83.14 |
| ENSMUSG00000035964 | transmembrane protein 59-like [Source:MGI Symbol;Acc:MGI:1915187] | Tmem59l | Enriched in dLGN | 1.147 | 0.0002215 | 166 |
| ENSMUSG00000021996 | esterase D/formylglutathione hydrolase [Source:MGI Symbol;Acc:MGI:95421] | Esd | Enriched in dLGN | 1.145 | 0.0001276 | 87.58 |
| ENSMUSG00000024121 | ATPase, H+ transporting, lysosomal V0 subunit C [Source:MGI Symbol;Acc:MGI:88116] | Atp6v0c | Enriched in dLGN | 1.144 | 0.0001928 | 333.4 |
| ENSMUSG00000030946 | phospholysine phosphohistidine inorganic pyrophosphate phosphatase [Source:MGI Symbol;Acc:MGI:1923679] | Lhpp | Enriched in dLGN | 1.143 | 0.001341 | 34.61 |
| ENSMUSG00000002475 | abhydrolase domain containing 3 [Source:MGI Symbol;Acc:MGI:2147183] | Abhd3 | Enriched in dLGN | 1.143 | 0.008781 | 45.04 |
| ENSMUSG00000033793 | ATPase, H+ transporting, lysosomal V1 subunit H [Source:MGI Symbol;Acc:MGI:1914864] | Atp6v1h | Enriched in dLGN | 1.142 | 0.00032 | 178.8 |
| ENSMUSG00000034839 | La ribonucleoprotein 6, translational regulator [Source:MGI Symbol;Acc:MGI:1914807] | Larp6 | Enriched in dLGN | 1.14 | 0.000001023 | 87.58 |
| ENSMUSG00000027634 | N-myc downstream regulated gene 3 [Source:MGI Symbol;Acc:MGI:1352499] | Ndrg3 | Enriched in dLGN | 1.138 | 0.000001268 | 572.8 |
| ENSMUSG00000027274 | McKusick-Kaufman syndrome [Source:MGI Symbol;Acc:MGI:1891836] | Mkks | Enriched in dLGN | 1.137 | 0.02151 | 21.92 |
| ENSMUSG00000022789 | dynamin 1-like [Source:MGI Symbol;Acc:MGI:1921256] | Dnm1l | Enriched in dLGN | 1.136 | 0.00001498 | 380.5 |
| ENSMUSG00000042763 | mannosidase, endo-alpha-like [Source:MGI Symbol;Acc:MGI:2684896] | Maneal | Enriched in dLGN | 1.136 | 0.00009731 | 148.8 |
| ENSMUSG00000030079 | RuvB-like protein 1 [Source:MGI Symbol;Acc:MGI:1928760] | Ruvbl1 | Enriched in dLGN | 1.136 | 0.0008309 | 70.63 |
| ENSMUSG00000049299 | trafficking protein particle complex 1 [Source:MGI Symbol;Acc:MGI:1098727] | Trappc1 | Enriched in dLGN | 1.135 | 0.0008711 | 33.39 |
| ENSMUSG00000035049 | ribosomal RNA processing 12 homolog [Source:MGI Symbol;Acc:MGI:2147437] | Rrp12 | Enriched in dLGN | 1.133 | 0.000004075 | 124.1 |
| ENSMUSG00000033685 | uncoupling protein 2 (mitochondrial, proton carrier) [Source:MGI Symbol;Acc:MGI:109354] | Ucp2 | Enriched in dLGN | 1.133 | 0.002658 | 149.6 |
| ENSMUSG00000027195 | hydroxysteroid (17-beta) dehydrogenase 12 [Source:MGI Symbol;Acc:MGI:1926967] | Hsd17b12 | Enriched in dLGN | 1.131 | 0.00005268 | 141 |
| ENSMUSG00000010057 | NPR2 like, GATOR1 complex subunit [Source:MGI Symbol;Acc:MGI:1914482] | Nprl2 | Enriched in dLGN | 1.131 | 0.01924 | 29.57 |
| ENSMUSG00000063802 | HSPA (heat shock 70kDa) binding protein, cytoplasmic cochaperone 1 [Source:MGI Symbol;Acc:MGI:1913495] | Hspbp1 | Enriched in dLGN | 1.13 | 2.275E-07 | 128.5 |
| ENSMUSG00000030452 | non imprinted in Prader-Willi/Angelman syndrome 2 homolog (human) [Source:MGI Symbol;Acc:MGI:1913918] | Nipa2 | Enriched in dLGN | 1.13 | 0.00365 | 70.36 |
| ENSMUSG00000045538 | DEAD box helicase 28 [Source:MGI Symbol;Acc:MGI:1919236] | Ddx28 | Enriched in dLGN | 1.129 | 0.0007121 | 32.99 |
| ENSMUSG00000035790 | centrosomal protein 19 [Source:MGI Symbol;Acc:MGI:1914244] | Cep19 | Enriched in dLGN | 1.128 | 1.784E-08 | 80.5 |
| ENSMUSG00000000787 | DEAD box helicase 3, X-linked [Source:MGI Symbol;Acc:MGI:103064] | Ddx3x | Enriched in dLGN | 1.128 | 0.00001224 | 602.8 |
| ENSMUSG00000078713 | translocase of outer mitochondrial membrane 5 [Source:MGI Symbol;Acc:MGI:1915762] | Tomm5 | Enriched in dLGN | 1.127 | 0.0024 | 36.78 |
| ENSMUSG00000042182 | BEN domain containing 6 [Source:MGI Symbol;Acc:MGI:2444572] | Bend6 | Enriched in dLGN | 1.123 | 0.000002664 | 256 |
| ENSMUSG00000033319 | fem 1 homolog c [Source:MGI Symbol;Acc:MGI:2444737] | Fem1c | Enriched in dLGN | 1.123 | 0.00003409 | 163.3 |
| ENSMUSG00000025439 | chloride channel, nucleotide-sensitive, 1A [Source:MGI Symbol;Acc:MGI:109638] | Clns1a | Enriched in dLGN | 1.122 | 0.00002061 | 102.4 |
| ENSMUSG00000039989 | chromobox 4 [Source:MGI Symbol;Acc:MGI:1195985] | Cbx4 | Enriched in dLGN | 1.12 | 5.208E-11 | 266.9 |
| ENSMUSG00000021276 | cyclin-dependent kinase 2 interacting protein [Source:MGI Symbol;Acc:MGI:1914486] | Cinp | Enriched in dLGN | 1.12 | 0.00006606 | 49.08 |
| ENSMUSG00000024878 | COBW domain containing 1 [Source:MGI Symbol;Acc:MGI:2385089] | Cbwd1 | Enriched in dLGN | 1.12 | 0.01235 | 28.1 |
| ENSMUSG00000024740 | damage specific DNA binding protein 1 [Source:MGI Symbol;Acc:MGI:1202384] | Ddb1 | Enriched in dLGN | 1.119 | 0.00002894 | 974.8 |
| ENSMUSG00000027384 | NADH:ubiquinone oxidoreductase complex assembly factor 5 [Source:MGI Symbol;Acc:MGI:1916737] | Ndufaf5 | Enriched in dLGN | 1.118 | 0.0002544 | 90.19 |
| ENSMUSG00000018848 | arginyl-tRNA synthetase [Source:MGI Symbol;Acc:MGI:1914297] | Rars | Enriched in dLGN | 1.117 | 1.343E-08 | 102.9 |
| ENSMUSG00000011254 | tRNA-histidine guanylyltransferase 1-like (S. cerevisiae) [Source:MGI Symbol;Acc:MGI:1913878] | Thg1l | Enriched in dLGN | 1.116 | 0.02906 | 24.76 |

| | | | | | | |
|---|---|---|---|---|---|---|
| ENSMUSG00000025868 | HIG1 domain family, member 2A [Source:MGI Symbol;Acc:MGI:1914294] | Higd2a | Enriched in dLGN | 1.115 | 0.003835 | 47.28 |
| ENSMUSG00000021643 | small EDRK-rich factor 1 [Source:MGI Symbol;Acc:MGI:1337114] | Serf1 | Enriched in dLGN | 1.115 | 0.006452 | 40.29 |
| ENSMUSG00000026798 | coenzyme Q4 [Source:MGI Symbol;Acc:MGI:1098826] | Coq4 | Enriched in dLGN | 1.115 | 0.006792 | 24.08 |
| ENSMUSG00000032604 | glutaminyl-tRNA synthetase [Source:MGI Symbol;Acc:MGI:1915851] | Qars | Enriched in dLGN | 1.112 | 6.474E-07 | 152.8 |
| ENSMUSG00000028070 | NAD(P)HX epimerase [Source:MGI Symbol;Acc:MGI:2180167] | Naxe | Enriched in dLGN | 1.108 | 0.00003619 | 67.2 |
| ENSMUSG00000020978 | kelch domain containing 2 [Source:MGI Symbol;Acc:MGI:1916804] | Klhdc2 | Enriched in dLGN | 1.107 | 0.00001095 | 321.5 |
| ENSMUSG00000022257 | lysosomal-associated protein transmembrane 4B [Source:MGI Symbol;Acc:MGI:1890494] | Laptm4b | Enriched in dLGN | 1.107 | 0.0001086 | 165.8 |
| ENSMUSG00000021178 | protease (prosome, macropain) 26S subunit, ATPase 1 [Source:MGI Symbol;Acc:MGI:106054] | Psmc1 | Enriched in dLGN | 1.107 | 0.0001172 | 275.2 |
| ENSMUSG00000024410 | regulator of MON1-CCZ1 [Source:MGI Symbol;Acc:MGI:1916528] | Rmc1 | Enriched in dLGN | 1.106 | 0.0001577 | 76.98 |
| ENSMUSG00000028656 | CAP, adenylate cyclase-associated protein 1 (yeast) [Source:MGI Symbol;Acc:MGI:88262] | Cap1 | Enriched in dLGN | 1.106 | 0.0007309 | 243.2 |
| ENSMUSG00000087260 | late endosomal/lysosomal adaptor, MAPK and MTOR activator 5 [Source:MGI Symbol;Acc:MGI:1915826] | Lamtor5 | Enriched in dLGN | 1.105 | 0.02225 | 22.53 |
| ENSMUSG00000007036 | abhydrolase domain containing 16A [Source:MGI Symbol;Acc:MGI:99476] | Abhd16a | Enriched in dLGN | 1.104 | 7.749E-07 | 134 |
| ENSMUSG00000073471 | radial spoke 3A homolog (Chlamydomonas) [Source:MGI Symbol;Acc:MGI:1914082] | Rsph3a | Enriched in dLGN | 1.104 | 0.03247 | 41.68 |
| ENSMUSG00000062691 | CCAAT/enhancer binding protein (C/EBP), zeta, opposite strand [Source:MGI Symbol;Acc:MGI:1915804] | Cebpzos | Enriched in dLGN | 1.101 | 0.009621 | 21.83 |
| ENSMUSG00000027002 | NCK-associated protein 1 [Source:MGI Symbol;Acc:MGI:1355333] | Nckap1 | Enriched in dLGN | 1.1 | 6.989E-07 | 682.6 |
| ENSMUSG00000025531 | choroidermia (RAB escort protein 1) [Source:MGI Symbol;Acc:MGI:892979] | Chm | Enriched in dLGN | 1.1 | 0.00007837 | 90.38 |
| ENSMUSG00000025277 | abhydrolase domain containing 6 [Source:MGI Symbol;Acc:MGI:1913332] | Abhd6 | Enriched in dLGN | 1.099 | 0.0004064 | 71.87 |
| ENSMUSG00000006315 | transmembrane protein 147 [Source:MGI Symbol;Acc:MGI:1915011] | Tmem147 | Enriched in dLGN | 1.099 | 0.0007062 | 46.25 |
| ENSMUSG00000036186 | divergent protein kinase domain 1B [Source:MGI Symbol;Acc:MGI:1927576] | Dipk1b | Enriched in dLGN | 1.099 | 0.004699 | 28.18 |
| ENSMUSG00000024446 | ribonuclease P 21 subunit [Source:MGI Symbol;Acc:MGI:1914926] | Rpp21 | Enriched in dLGN | 1.099 | 0.03949 | 21.09 |
| ENSMUSG00000004070 | heme oxygenase 2 [Source:MGI Symbol;Acc:MGI:109373] | Hmox2 | Enriched in dLGN | 1.097 | 0.00032 | 96.26 |
| ENSMUSG00000020386 | secretion associated Ras related GTPase 1B [Source:MGI Symbol;Acc:MGI:1913647] | Sar1b | Enriched in dLGN | 1.097 | 0.0004945 | 55.03 |
| ENSMUSG00000031068 | glutaredoxin 3 [Source:MGI Symbol;Acc:MGI:1353653] | Glrx3 | Enriched in dLGN | 1.095 | 0.0003665 | 113.9 |
| ENSMUSG00000035517 | tudor domain containing 7 [Source:MGI Symbol;Acc:MGI:2140279] | Tdrd7 | Enriched in dLGN | 1.094 | 1.327E-07 | 468.1 |
| ENSMUSG00000019797 | mitochondrial transcription rescue factor 1 [Source:MGI Symbol;Acc:MGI:1915101] | Mtres1 | Enriched in dLGN | 1.094 | 0.00004356 | 74.57 |
| ENSMUSG00000033628 | phosphatidylinositol 3-kinase catalytic subunit type 3 [Source:MGI Symbol;Acc:MGI:2445019] | Pik3c3 | Enriched in dLGN | 1.092 | 0.0001052 | 127.1 |
| ENSMUSG00000030738 | eukaryotic translation initiation factor 3, subunit C [Source:MGI Symbol;Acc:MGI:1926966] | Eif3c | Enriched in dLGN | 1.088 | 0.00006494 | 682.4 |
| ENSMUSG00000026584 | SCY1-like 3 (S. cerevisiae) [Source:MGI Symbol;Acc:MGI:1921385] | Scyl3 | Enriched in dLGN | 1.088 | 0.001096 | 79.74 |
| ENSMUSG00000028998 | translocase of outer mitochondrial membrane 7 [Source:MGI Symbol;Acc:MGI:1913419] | Tomm7 | Enriched in dLGN | 1.087 | 0.001065 | 121.4 |
| ENSMUSG00000018882 | mitochondrial ribosomal protein L45 [Source:MGI Symbol;Acc:MGI:1914286] | Mrpl45 | Enriched in dLGN | 1.086 | 0.0000996 | 64.61 |
| ENSMUSG00000005312 | ubiquilin 1 [Source:MGI Symbol;Acc:MGI:1860276] | Ubqln1 | Enriched in dLGN | 1.082 | 2.747E-08 | 342.6 |
| ENSMUSG00000027673 | NADH:ubiquinone oxidoreductase subunit B5 [Source:MGI Symbol;Acc:MGI:1913296] | Ndufb5 | Enriched in dLGN | 1.082 | 0.00000128 | 138.6 |
| ENSMUSG00000007338 | mitochondrial ribosomal protein L49 [Source:MGI Symbol;Acc:MGI:108180] | Mrpl49 | Enriched in dLGN | 1.082 | 0.00003162 | 74.11 |
| ENSMUSG00000025208 | mitochondrial ribosomal protein L43 [Source:MGI Symbol;Acc:MGI:2137229] | Mrpl43 | Enriched in dLGN | 1.082 | 0.0001052 | 70.84 |
| ENSMUSG00000024668 | succinate dehydrogenase complex assembly factor 2 [Source:MGI Symbol;Acc:MGI:1913322] | Sdhaf2 | Enriched in dLGN | 1.079 | 0.00005209 | 57.83 |
| ENSMUSG00000021546 | heterogeneous nuclear ribonucleoprotein K [Source:MGI Symbol;Acc:MGI:99894] | Hnrnpk | Enriched in dLGN | 1.079 | 0.0003726 | 580.6 |
| ENSMUSG00000000171 | succinate dehydrogenase complex, subunit D, integral membrane protein [Source:MGI Symbol;Acc:MGI:1914175] | Sdhd | Enriched in dLGN | 1.078 | 0.000001556 | 85.24 |
| ENSMUSG00000030264 | THUMP domain containing 3 [Source:MGI Symbol;Acc:MGI:1277973] | Thumpd3 | Enriched in dLGN | 1.078 | 0.005886 | 45.23 |
| ENSMUSG00000028478 | clathrin, light polypeptide (Lca) [Source:MGI Symbol;Acc:MGI:894297] | Clta | Enriched in dLGN | 1.077 | 5.306E-10 | 866.9 |
| ENSMUSG00000030879 | mitochondrial ribosomal protein L17 [Source:MGI Symbol;Acc:MGI:1351608] | Mrpl17 | Enriched in dLGN | 1.076 | 0.0000207 | 76.99 |
| ENSMUSG00000030683 | seizure related 6 homolog like 2 [Source:MGI Symbol;Acc:MGI:2385295] | Sez6l2 | Enriched in dLGN | 1.075 | 0.00002626 | 601.7 |
| ENSMUSG00000020537 | developmentally regulated GTP binding protein 2 [Source:MGI Symbol;Acc:MGI:1342307] | Drg2 | Enriched in dLGN | 1.075 | 0.001323 | 72 |
| ENSMUSG00000041355 | signal sequence receptor, beta [Source:MGI Symbol;Acc:MGI:1913506] | Ssr2 | Enriched in dLGN | 1.073 | 0.0002305 | 72.58 |
| ENSMUSG00000001016 | interleukin enhancer binding factor 2 [Source:MGI Symbol;Acc:MGI:1915031] | Ilf2 | Enriched in dLGN | 1.072 | 0.00001546 | 131.5 |
| ENSMUSG00000030037 | mitochondrial ribosomal protein L53 [Source:MGI Symbol;Acc:MGI:1915749] | Mrpl53 | Enriched in dLGN | 1.072 | 0.003828 | 26.23 |
| ENSMUSG00000019868 | vesicle (multivesicular body) trafficking 1 [Source:MGI Symbol;Acc:MGI:1913451] | Vta1 | Enriched in dLGN | 1.072 | 0.007348 | 69.26 |
| ENSMUSG00000023460 | RAB12, member RAS oncogene family [Source:MGI Symbol;Acc:MGI:894284] | Rab12 | Enriched in dLGN | 1.071 | 1.46E-09 | 311.7 |
| ENSMUSG00000027193 | apoptosis inhibitor 5 [Source:MGI Symbol;Acc:MGI:1888993] | Api5 | Enriched in dLGN | 1.069 | 0.00007066 | 127.6 |
| ENSMUSG00000029228 | ligand of numb-protein X 1 [Source:MGI Symbol;Acc:MGI:1278335] | Lnx1 | Enriched in dLGN | 1.067 | 0.0002742 | 104.6 |
| ENSMUSG00000028790 | KH domain containing, RNA binding, signal transduction associated 1 [Source:MGI Symbol;Acc:MGI:893579] | Khdrbs1 | Enriched in dLGN | 1.066 | 0.000001346 | 341 |
| ENSMUSG00000047617 | non-homologous end joining factor [Source:MGI Symbol;Acc:MGI:2442831] | Paxx | Enriched in dLGN | 1.066 | 0.0006047 | 66.93 |
| ENSMUSG00000022024 | SGT1, suppressor of G2 allele of SKP1 (S. cerevisiae) [Source:MGI Symbol;Acc:MGI:1915205] | Sugt1 | Enriched in dLGN | 1.065 | 5.997E-07 | 199.6 |
| ENSMUSG00000024527 | AFG3-like AAA ATPase 2 [Source:MGI Symbol;Acc:MGI:1916847] | Afg3l2 | Enriched in dLGN | 1.064 | 9.687E-10 | 309.3 |
| ENSMUSG00000033379 | ATPase, H+ transporting, lysosomal V0 subunit B [Source:MGI Symbol;Acc:MGI:1890510] | Atp6v0b | Enriched in dLGN | 1.063 | 1.616E-08 | 226.1 |
| ENSMUSG00000031948 | lysyl-tRNA synthetase [Source:MGI Symbol;Acc:MGI:1934754] | Kars | Enriched in dLGN | 1.063 | 0.0000773 | 171 |
| ENSMUSG00000035297 | COP9 signalosome subunit 4 [Source:MGI Symbol;Acc:MGI:1349414] | Cops4 | Enriched in dLGN | 1.062 | 0.002418 | 86.24 |
| ENSMUSG00000049092 | G protein-coupled receptor 137C [Source:MGI Symbol;Acc:MGI:1917963] | Gpr137c | Enriched in dLGN | 1.062 | 0.002884 | 120.2 |
| ENSMUSG00000022452 | single-pass membrane protein with aspartate rich tail 1 [Source:MGI Symbol;Acc:MGI:1916279] | Smdt1 | Enriched in dLGN | 1.061 | 0.000009153 | 93.19 |
| ENSMUSG00000017707 | serine incorporator 3 [Source:MGI Symbol;Acc:MGI:1349457] | Serinc3 | Enriched in dLGN | 1.061 | 0.00001257 | 254.9 |
| ENSMUSG00000041881 | NADH:ubiquinone oxidoreductase subunit A7 [Source:MGI Symbol;Acc:MGI:1913666] | Ndufa7 | Enriched in dLGN | 1.061 | 0.0001642 | 57.87 |
| ENSMUSG00000074736 | synapse differentiation inducing 1 [Source:MGI Symbol;Acc:MGI:3702158] | Syndig1 | Enriched in dLGN | 1.06 | 0.01249 | 50.55 |
| ENSMUSG00000031993 | sorting nexin 19 [Source:MGI Symbol;Acc:MGI:1921581] | Snx19 | Enriched in dLGN | 1.059 | 1.748E-10 | 263.6 |
| ENSMUSG00000047656 | tRNA phosphotransferase 1 [Source:MGI Symbol;Acc:MGI:1333115] | Trpt1 | Enriched in dLGN | 1.059 | 0.016 | 16 |
| ENSMUSG00000035772 | mitochondrial ribosomal protein S2 [Source:MGI Symbol;Acc:MGI:2153089] | Mrps2 | Enriched in dLGN | 1.057 | 0.0001377 | 52.55 |
| ENSMUSG00000023932 | cell division cycle 5-like (S. pombe) [Source:MGI Symbol;Acc:MGI:1918952] | Cdc5l | Enriched in dLGN | 1.053 | 0.00001317 | 151.3 |
| ENSMUSG00000037260 | heparan-alpha-glucosaminide N-acetyltransferase [Source:MGI Symbol;Acc:MGI:1196297] | Hgsnat | Enriched in dLGN | 1.052 | 0.000001395 | 107.2 |
| ENSMUSG00000009630 | protein phosphatase 2 (formerly 2A), catalytic subunit, beta isoform [Source:MGI Symbol;Acc:MGI:1321161] | Ppp2cb | Enriched in dLGN | 1.048 | 1.931E-08 | 209.1 |
| ENSMUSG00000026670 | UDP-N-acetylglucosamine pyrophosphorylase 1 [Source:MGI Symbol;Acc:MGI:1334459] | Uap1 | Enriched in dLGN | 1.048 | 2.64E-08 | 143.7 |
| ENSMUSG00000027642 | ribophorin II [Source:MGI Symbol;Acc:MGI:98085] | Rpn2 | Enriched in dLGN | 1.048 | 0.000007678 | 243.8 |
| ENSMUSG00000060073 | proteasome subunit alpha 3 [Source:MGI Symbol;Acc:MGI:104883] | Psma3 | Enriched in dLGN | 1.048 | 0.00006178 | 145 |
| ENSMUSG00000028224 | nibrin [Source:MGI Symbol;Acc:MGI:1351625] | Nbn | Enriched in dLGN | 1.048 | 0.0006893 | 42.38 |
| ENSMUSG00000028932 | proteasome (prosome, macropain) 26S subunit, ATPase 2 [Source:MGI Symbol;Acc:MGI:109555] | Psmc2 | Enriched in dLGN | 1.048 | 0.004155 | 150.2 |
| ENSMUSG00000028161 | protein phosphatase 3, catalytic subunit, alpha isoform [Source:MGI Symbol;Acc:MGI:107164] | Ppp3ca | Enriched in dLGN | 1.047 | 2.119E-08 | 1200 |
| ENSMUSG00000022494 | shisa family member 9 [Source:MGI Symbol;Acc:MGI:1919805] | Shisa9 | Enriched in dLGN | 1.047 | 0.005407 | 606.7 |
| ENSMUSG00000022797 | transferrin receptor [Source:MGI Symbol;Acc:MGI:98822] | Tfrc | Enriched in dLGN | 1.046 | 0.00005225 | 218.6 |
| ENSMUSG00000028609 | mago homolog, exon junction complex core component [Source:MGI Symbol;Acc:MGI:1330312] | Magoh | Enriched in dLGN | 1.046 | 0.02627 | 35.71 |
| ENSMUSG00000001383 | zinc finger, matrin type 2 [Source:MGI Symbol;Acc:MGI:1913742] | Zmat2 | Enriched in dLGN | 1.045 | 0.000001293 | 273.6 |
| ENSMUSG00000023106 | density-regulated protein [Source:MGI Symbol;Acc:MGI:1915434] | Denr | Enriched in dLGN | 1.044 | 9.311E-07 | 114.8 |
| ENSMUSG00000003527 | ess-2 splicing factor [Source:MGI Symbol;Acc:MGI:107854] | Ess2 | Enriched in dLGN | 1.044 | 0.00001546 | 75.66 |
| ENSMUSG00000034557 | zinc finger, FYVE domain containing 9 [Source:MGI Symbol;Acc:MGI:2652838] | Zfyve9 | Enriched in dLGN | 1.044 | 0.003623 | 559.4 |
| ENSMUSG00000027341 | transmembrane protein 230 [Source:MGI Symbol;Acc:MGI:1917862] | Tmem230 | Enriched in dLGN | 1.044 | 0.01084 | 38.18 |
| ENSMUSG00000001507 | integrin alpha 3 [Source:MGI Symbol;Acc:MGI:96602] | Itga3 | Enriched in dLGN | 1.043 | 0.00002068 | 148.8 |
| ENSMUSG00000039347 | ATPase, H+ transporting, lysosomal V0 subunit E2 [Source:MGI Symbol;Acc:MGI:1923502] | Atp6v0e2 | Enriched in dLGN | 1.042 | 9.845E-07 | 554.6 |
| ENSMUSG00000025142 | alveolar soft part sarcoma chromosome region, candidate 1 (human) [Source:MGI Symbol;Acc:MGI:1916188] | Aspscr1 | Enriched in dLGN | 1.041 | 0.00002055 | 101.1 |
| ENSMUSG00000027184 | cell cycle associated protein 1 [Source:MGI Symbol;Acc:MGI:1858234] | Caprin1 | Enriched in dLGN | 1.04 | 0.000001943 | 681.9 |
| ENSMUSG00000020485 | SPT4A, DSIF elongation factor subunit [Source:MGI Symbol;Acc:MGI:107416] | Supt4a | Enriched in dLGN | 1.04 | 0.001872 | 25.99 |
| ENSMUSG00000018572 | PHD finger protein 23 [Source:MGI Symbol;Acc:MGI:1925496] | Phf23 | Enriched in dLGN | 1.039 | 0.000001665 | 112.2 |
| ENSMUSG00000041997 | tousled-like kinase 1 [Source:MGI Symbol;Acc:MGI:2441683] | Tlk1 | Enriched in dLGN | 1.038 | 0.000008574 | 318.7 |
| ENSMUSG00000033124 | autophagy related 9A [Source:MGI Symbol;Acc:MGI:2138446] | Atg9a | Enriched in dLGN | 1.037 | 2.95E-11 | 274.4 |
| ENSMUSG00000029432 | nipsnap homolog 2 [Source:MGI Symbol;Acc:MGI:1278343] | Nipsnap2 | Enriched in dLGN | 1.036 | 0.0006356 | 127.2 |
| ENSMUSG00000029104 | huntingtin [Source:MGI Symbol;Acc:MGI:96067] | Htt | Enriched in dLGN | 1.035 | 3.794E-07 | 647.6 |
| ENSMUSG00000027894 | solute carrier family 6 (neurotransmitter transporter), member 17 [Source:MGI Symbol;Acc:MGI:2442535] | Slc6a17 | Enriched in dLGN | 1.035 | 0.00002779 | 1006 |
| ENSMUSG00000063406 | transmembrane p24 trafficking protein 5 [Source:MGI Symbol;Acc:MGI:1921586] | Tmed5 | Enriched in dLGN | 1.034 | 0.004699 | 35.86 |
| ENSMUSG00000050552 | late endosomal/lysosomal adaptor, MAPK and MTOR activator 4 [Source:MGI Symbol;Acc:MGI:1913346] | Lamtor4 | Enriched in dLGN | 1.033 | 0.004655 | 48.36 |
| ENSMUSG00000015766 | epidermal growth factor receptor pathway substrate 8 [Source:MGI Symbol;Acc:MGI:104684] | Eps8 | Enriched in dLGN | 1.032 | 0.000113 | 220.6 |
| ENSMUSG00000037894 | H2A.Z variant histone 1 [Source:MGI Symbol;Acc:MGI:1888388] | H2az1 | Enriched in dLGN | 1.03 | 0.001752 | 59.57 |
| ENSMUSG00000025762 | La ribonucleoprotein 1B [Source:MGI Symbol;Acc:MGI:1914604] | Larp1b | Enriched in dLGN | 1.029 | 0.0005768 | 100 |
| ENSMUSG00000026615 | glutamyl-prolyl-tRNA synthetase [Source:MGI Symbol;Acc:MGI:97838] | Eprs | Enriched in dLGN | 1.028 | 2.023E-07 | 511 |
| ENSMUSG00000003421 | nitric oxide synthase interacting protein [Source:MGI Symbol;Acc:MGI:1913644] | Nosip | Enriched in dLGN | 1.026 | 0.00003886 | 111.7 |
| ENSMUSG00000004535 | Tax1 (human T cell leukemia virus type I) binding protein 1 [Source:MGI Symbol;Acc:MGI:1289308] | Tax1bp1 | Enriched in dLGN | 1.026 | 0.00004749 | 665.1 |
| ENSMUSG00000005362 | cereblon [Source:MGI Symbol;Acc:MGI:1913277] | Crbn | Enriched in dLGN | 1.025 | 0.002644 | 104.3 |
| ENSMUSG00000028098 | ring finger protein 115 [Source:MGI Symbol;Acc:MGI:1915095] | Rnf115 | Enriched in dLGN | 1.022 | 0.0001446 | 146.7 |
| ENSMUSG00000040896 | potassium voltage-gated channel, Shal-related family, member 3 [Source:MGI Symbol;Acc:MGI:1928743] | Kcnd3 | Enriched in dLGN | 1.022 | 0.0002476 | 579.5 |
| ENSMUSG00000068739 | seryl-aminoacyl-tRNA synthetase [Source:MGI Symbol;Acc:MGI:102809] | Sars | Enriched in dLGN | 1.02 | 1.438E-08 | 493.2 |
| ENSMUSG00000030327 | NECAP endocytosis associated 1 [Source:MGI Symbol;Acc:MGI:1914852] | Necap1 | Enriched in dLGN | 1.019 | 0.00234 | 167.4 |
| ENSMUSG00000029447 | chaperonin containing Tcp1, subunit 6a (zeta) [Source:MGI Symbol;Acc:MGI:107943] | Cct6a | Enriched in dLGN | 1.015 | 0.00005131 | 356.4 |
| ENSMUSG00000022234 | chaperonin containing Tcp1, subunit 5 (epsilon) [Source:MGI Symbol;Acc:MGI:107185] | Cct5 | Enriched in dLGN | 1.014 | 0.0005135 | 409.3 |
| ENSMUSG00000091264 | small integral membrane protein 13 [Source:MGI Symbol;Acc:MGI:2652854] | Smim13 | Enriched in dLGN | 1.013 | 0.0002833 | 346 |
| ENSMUSG00000051154 | COMM domain containing 3 [Source:MGI Symbol;Acc:MGI:88218] | Commd3 | Enriched in dLGN | 1.012 | 0.003658 | 62.85 |
| ENSMUSG00000001986 | glutamate receptor, ionotropic, AMPA3 (alpha 3) [Source:MGI Symbol;Acc:MGI:95810] | Gria3 | Enriched in dLGN | 1.011 | 0.000239 | 283.8 |
| ENSMUSG00000027792 | butyrylcholinesterase [Source:MGI Symbol;Acc:MGI:894278] | Bche | Enriched in dLGN | 1.011 | 0.04311 | 17.98 |
| ENSMUSG00000026687 | aldehyde dehydrogenase 9, subfamily A1 [Source:MGI Symbol;Acc:MGI:1861622] | Aldh9a1 | Enriched in dLGN | 1.01 | 0.0006422 | 102.9 |
| ENSMUSG00000023328 | acetylcholinesterase [Source:MGI Symbol;Acc:MGI:87876] | Ache | Enriched in dLGN | 1.009 | 0.001477 | 333.5 |
| ENSMUSG00000032171 | peptidyl-prolyl cis/trans isomerase, NIMA-interacting 1 [Source:MGI Symbol;Acc:MGI:1346036] | Pin1 | Enriched in dLGN | 1.008 | 8.459E-07 | 122 |
| ENSMUSG00000044442 | N-6 adenine-specific DNA methyltransferase 1 (putative) [Source:MGI Symbol;Acc:MGI:1915018] | N6amt1 | Enriched in dLGN | 1.008 | 0.0009846 | 39.22 |
| ENSMUSG00000041216 | clavesin 1 [Source:MGI Symbol;Acc:MGI:1921688] | Clvs1 | Enriched in dLGN | 1.008 | 0.003072 | 163.7 |
| ENSMUSG00000054162 | sparc/osteonectin, cwcv and kazal-like domains proteoglycan 3 [Source:MGI Symbol;Acc:MGI:1920152] | Spock3 | Enriched in dLGN | 1.006 | 0.003883 | 404.4 |
| ENSMUSG00000028434 | erythrocyte membrane protein band 4.1 like 4b [Source:MGI Symbol;Acc:MGI:1859149] | Epb41l4b | Enriched in dLGN | 1.005 | 0.00005755 | 333.4 |
| ENSMUSG00000029263 | phosphatidylinositol glycan anchor biosynthesis, class G [Source:MGI Symbol;Acc:MGI:3576484] | Pigg | Enriched in dLGN | 1.005 | 0.007485 | 43.15 |
| ENSMUSG00000027011 | ubiquitin-conjugating enzyme E2E 3 [Source:MGI Symbol;Acc:MGI:107412] | Ube2e3 | Enriched in dLGN | 1.003 | 0.00001907 | 305.5 |
| ENSMUSG00000027236 | eukaryotic translation initiation factor 3, subunit J1 [Source:MGI Symbol;Acc:MGI:1925905] | Eif3j1 | Enriched in dLGN | 1.002 | 0.001093 | 85.54 |
| ENSMUSG00000016526 | dual-specificity tyrosine-(Y)-phosphorylation regulated kinase 3 [Source:MGI Symbol;Acc:MGI:1330300] | Dyrk3 | Enriched in dLGN | 1 | 0.01831 | 17.86 |
| ENSMUSG00000024501 | dihydropyrimidinase-like 3 [Source:MGI Symbol;Acc:MGI:1349762] | Dpysl3 | Enriched in SCN | -1.294 | 0.00008896 | 3490 |
| ENSMUSG00000022865 | coxsackie virus and adenovirus receptor [Source:MGI Symbol;Acc:MGI:1201679] | Cxadr | Enriched in SCN | -1.688 | 0.000002913 | 403.7 |
| ENSMUSG00000025314 | protein tyrosine phosphatase, receptor type, J [Source:MGI Symbol;Acc:MGI:104574] | Ptprj | Enriched in SCN | -3.539 | 7.896E-08 | 998.9 |

**Contrast numerator: P15_KO_dLGN. Contrast denominator: P15_KO_SCN.**

| ensembl_gene_id | description | mgi_symbol | Significance | deseq_logfc | deseq_adjp | deseq_basemean |
|---|---|---|---|---|---|---|
| ENSMUSG00000064179 | troponin T1, skeletal, slow [Source:MGI Symbol;Acc:MGI:1333868] | Tnnt1 | Enriched in dLGN | 8.725 | 3.588E-38 | 308.7 |
| ENSMUSG00000022468 | endonuclease, polyU-specific [Source:MGI Symbol;Acc:MGI:97746] | Endou | Enriched in dLGN | 7.971 | 1.022E-14 | 34.93 |
| ENSMUSG00000029054 | gamma-aminobutyric acid (GABA) A receptor, subunit delta [Source:MGI Symbol;Acc:MGI:95622] | Gabrd | Enriched in dLGN | 7.833 | 1.716E-18 | 52.02 |
| ENSMUSG00000021948 | protein kinase C, delta [Source:MGI Symbol;Acc:MGI:97598] | Prkcd | Enriched in dLGN | 7.161 | 9.492E-46 | 727.9 |
| ENSMUSG00000041044 | leucine-rich repeat, immunoglobulin-like and transmembrane domains 1 [Source:MGI Symbol;Acc:MGI:2385320] | Lrit1 | Enriched in dLGN | 7.128 | 0.00004179 | 386.5 |
| ENSMUSG00000032532 | cholecystokinin [Source:MGI Symbol;Acc:MGI:88297] | Cck | Enriched in dLGN | 7.119 | 1.408E-36 | 445.5 |
| ENSMUSG00000031450 | G protein-coupled receptor kinase 1 [Source:MGI Symbol;Acc:MGI:1345146] | Grk1 | Enriched in dLGN | 7.063 | 0.000003048 | 1629 |
| ENSMUSG00000032292 | nuclear receptor subfamily 2, group E, member 3 [Source:MGI Symbol;Acc:MGI:1346317] | Nr2e3 | Enriched in dLGN | 7.001 | 0.00001856 | 617.6 |
| ENSMUSG00000029663 | guanine nucleotide binding protein (G protein), gamma transducing activity polypeptide 1 [Source:MGI Symbol;Acc:MGI:109165] | Gngt1 | Enriched in dLGN | 6.878 | 0.00004644 | 593.7 |
| ENSMUSG00000031293 | retinoschisis (X-linked, juvenile) 1 (human) [Source:MGI Symbol;Acc:MGI:1336189] | Rs1 | Enriched in dLGN | 6.712 | 0.000007641 | 1028 |
| ENSMUSG00000031142 | calcium channel, voltage-dependent, alpha 1F subunit [Source:MGI Symbol;Acc:MGI:1859639] | Cacna1f | Enriched in dLGN | 6.604 | 0.00004115 | 357 |
| ENSMUSG00000024519 | complexin 4 [Source:MGI Symbol;Acc:MGI:2685803] | Cplx4 | Enriched in dLGN | 6.316 | 0.0002202 | 325.1 |
| ENSMUSG00000039103 | nexilin [Source:MGI Symbol;Acc:MGI:1916060] | Nexn | Enriched in dLGN | 6.304 | 2.851E-23 | 125.8 |
| ENSMUSG00000046049 | retinitis pigmentosa 1 homolog like 1 [Source:MGI Symbol;Acc:MGI:2384303] | Rp1l1 | Enriched in dLGN | 6.094 | 0.00003571 | 1013 |
| ENSMUSG00000006007 | phosducin [Source:MGI Symbol;Acc:MGI:98090] | Pdc | Enriched in dLGN | 6.017 | 0.0002028 | 804 |
| ENSMUSG00000047298 | potassium channel, subfamily V, member 2 [Source:MGI Symbol;Acc:MGI:2670981] | Kcnv2 | Enriched in dLGN | 5.703 | 0.00006278 | 537.7 |
| ENSMUSG00000034829 | nucleoredoxin-like 1 [Source:MGI Symbol;Acc:MGI:1924446] | Nxnl1 | Enriched in dLGN | 5.667 | 0.000183 | 200.2 |
| ENSMUSG00000043418 | leucine-rich repeat, immunoglobulin-like and transmembrane domains 2 [Source:MGI Symbol;Acc:MGI:2444885] | Lrit2 | Enriched in dLGN | 5.662 | 0.0008016 | 108.6 |
| ENSMUSG00000039783 | kynurenine 3-monooxygenase (kynurenine 3-hydroxylase) [Source:MGI Symbol;Acc:MGI:2138151] | Kmo | Enriched in dLGN | 5.538 | 0.000001324 | 4.13 |
| ENSMUSG00000050315 | synaptopodin 2 [Source:MGI Symbol;Acc:MGI:2153070] | Synpo2 | Enriched in dLGN | 5.478 | 1.358E-59 | 581.4 |
| ENSMUSG00000029491 | phosphodiesterase 6B, cGMP, rod receptor, beta polypeptide [Source:MGI Symbol;Acc:MGI:97525] | Pde6b | Enriched in dLGN | 5.44 | 0.0001317 | 1205 |
| ENSMUSG00000091402 | retinal degeneration 3-like [Source:MGI Symbol;Acc:MGI:2675860] | Rd3l | Enriched in dLGN | 5.275 | 0.00002093 | 41.89 |
| ENSMUSG00000045776 | leucine-rich repeats and transmembrane domains 1 [Source:MGI Symbol;Acc:MGI:2442106] | Lrtm1 | Enriched in dLGN | 5.243 | 0.0009435 | 127.3 |
| ENSMUSG00000111755 | predicted gene, 47160 [Source:MGI Symbol;Acc:MGI:6095933] | Gm47160 | Enriched in dLGN | 5.242 | 0.005268 | 97.35 |
| ENSMUSG00000028024 | glutamyl aminopeptidase [Source:MGI Symbol;Acc:MGI:106645] | Enpep | Enriched in dLGN | 5.226 | 5.814E-08 | 10.7 |
| ENSMUSG00000056947 | mab-21-like 1 [Source:MGI Symbol;Acc:MGI:1333773] | Mab21l1 | Enriched in dLGN | 5.211 | 0.00005846 | 403.6 |
| ENSMUSG00000047034 | ankyrin repeat domain 33 [Source:MGI Symbol;Acc:MGI:2443398] | Ankrd33 | Enriched in dLGN | 5.194 | 0.001123 | 105.8 |
| ENSMUSG00000034837 | G protein subunit alpha transducin 1 [Source:MGI Symbol;Acc:MGI:95778] | Gnat1 | Enriched in dLGN | 5.189 | 0.000562 | 1840 |
| ENSMUSG00000056055 | S-antigen, retina and pineal gland (arrestin) [Source:MGI Symbol;Acc:MGI:98227] | Sag | Enriched in dLGN | 5.178 | 0.00001304 | 2015 |
| ENSMUSG00000031994 | a disintegrin-like and metallopeptidase (reprolysin type) with thrombospondin type 1 motif, 8 [Source:MGI Symbol;Acc:MGI:1353468] | Adamts8 | Enriched in dLGN | 5.151 | 4.062E-18 | 52.93 |
| ENSMUSG00000033044 | dehydrogenase/reductase (SDR family) member 7C [Source:MGI Symbol;Acc:MGI:1915710] | Dhrs7c | Enriched in dLGN | 5.114 | 0.0005057 | 6.079 |
| ENSMUSG00000041534 | retinol binding protein 3, interstitial [Source:MGI Symbol;Acc:MGI:97878] | Rbp3 | Enriched in dLGN | 5.075 | 0.0001836 | 3079 |
| ENSMUSG00000023979 | guanylate cyclase activator 1B [Source:MGI Symbol;Acc:MGI:1194489] | Guca1b | Enriched in dLGN | 5.035 | 0.0001638 | 220.2 |
| ENSMUSG00000019762 | iodotyrosine deiodinase [Source:MGI Symbol;Acc:MGI:1917587] | Iyd | Enriched in dLGN | 5.012 | 0.0000245 | 6.399 |
| ENSMUSG00000070683 | lactamase, beta-like 1 [Source:MGI Symbol;Acc:MGI:2448566] | Lactbl1 | Enriched in dLGN | 4.985 | 0.001438 | 222.9 |
| ENSMUSG00000024842 | calcium binding protein 4 [Source:MGI Symbol;Acc:MGI:1920910] | Cabp4 | Enriched in dLGN | 4.95 | 0.001575 | 95.11 |
| ENSMUSG00000024575 | phosphodiesterase 6A, cGMP-specific, rod, alpha [Source:MGI Symbol;Acc:MGI:97524] | Pde6a | Enriched in dLGN | 4.937 | 0.0003707 | 1301 |
| ENSMUSG00000041578 | cone-rod homeobox [Source:MGI Symbol;Acc:MGI:1194883] | Crx | Enriched in dLGN | 4.903 | 0.0004054 | 944.1 |
| ENSMUSG00000034452 | solute carrier family 24 (sodium/potassium/calcium exchanger), member 1 [Source:MGI Symbol;Acc:MGI:2384771] | Slc24a1 | Enriched in dLGN | 4.894 | 0.000341 | 1018 |
| ENSMUSG00000000617 | glutamate receptor, metabotropic 6 [Source:MGI Symbol;Acc:MGI:1351343] | Grm6 | Enriched in dLGN | 4.859 | 0.002795 | 178.8 |
| ENSMUSG00000019966 | kit ligand [Source:MGI Symbol;Acc:MGI:96974] | Kitl | Enriched in dLGN | 4.793 | 4.943E-37 | 453.1 |
| ENSMUSG00000030324 | rhodopsin [Source:MGI Symbol;Acc:MGI:97914] | Rho | Enriched in dLGN | 4.784 | 0.000217 | 5636 |
| ENSMUSG00000023978 | peripherin 2 [Source:MGI Symbol;Acc:MGI:102791] | Prph2 | Enriched in dLGN | 4.773 | 0.0006814 | 1752 |
| ENSMUSG00000030785 | cytochrome c oxidase subunit 6A2 [Source:MGI Symbol;Acc:MGI:104649] | Cox6a2 | Enriched in dLGN | 4.755 | 1.206E-08 | 11.74 |
| ENSMUSG00000024518 | retina and anterior neural fold homeobox [Source:MGI Symbol;Acc:MGI:109632] | Rax | Enriched in dLGN | 4.685 | 0.001514 | 128.9 |
| ENSMUSG00000025900 | retinitis pigmentosa 1 (human) [Source:MGI Symbol;Acc:MGI:1341105] | Rp1 | Enriched in dLGN | 4.634 | 0.00005928 | 3382 |
| ENSMUSG00000026688 | microsomal glutathione S-transferase 3 [Source:MGI Symbol;Acc:MGI:1913697] | Mgst3 | Enriched in dLGN | 4.633 | 2.05E-17 | 88.46 |
| ENSMUSG00000021123 | retinol dehydrogenase 12 [Source:MGI Symbol;Acc:MGI:1925224] | Rdh12 | Enriched in dLGN | 4.531 | 0.0001044 | 285 |
| ENSMUSG00000026525 | opsin 3 [Source:MGI Symbol;Acc:MGI:1338022] | Opn3 | Enriched in dLGN | 4.507 | 2.238E-16 | 17.62 |
| ENSMUSG00000029659 | mesenteric estrogen dependent adipogenesis [Source:MGI Symbol;Acc:MGI:1917967] | Medag | Enriched in dLGN | 4.469 | 2.801E-16 | 25.4 |
| ENSMUSG00000040554 | aryl hydrocarbon receptor-interacting protein-like 1 [Source:MGI Symbol;Acc:MGI:2148800] | Aipl1 | Enriched in dLGN | 4.455 | 0.003087 | 368.2 |
| ENSMUSG00000056043 | regulator of G-protein signalling 9 binding protein [Source:MGI Symbol;Acc:MGI:2384418] | Rgs9bp | Enriched in dLGN | 4.446 | 0.00008837 | 666 |
| ENSMUSG00000028464 | tropomyosin 2, beta [Source:MGI Symbol;Acc:MGI:98810] | Tpm2 | Enriched in dLGN | 4.416 | 9.927E-12 | 25.35 |
| ENSMUSG00000028364 | tenascin C [Source:MGI Symbol;Acc:MGI:101922] | Tnc | Enriched in dLGN | 4.386 | 2.926E-21 | 268.5 |
| ENSMUSG00000030523 | transient receptor potential cation channel, subfamily M, member 1 [Source:MGI Symbol;Acc:MGI:1330305] | Trpm1 | Enriched in dLGN | 4.376 | 0.001586 | 420.9 |
| ENSMUSG00000032776 | multiple C2 domains, transmembrane 2 [Source:MGI Symbol;Acc:MGI:2685335] | Mctp2 | Enriched in dLGN | 4.37 | 9.017E-13 | 28.35 |
| ENSMUSG00000025366 | extended synaptotagmin-like protein 1 [Source:MGI Symbol;Acc:MGI:1344426] | Esyt1 | Enriched in dLGN | 4.313 | 2.696E-21 | 159.4 |
| ENSMUSG00000018470 | potassium voltage-gated channel, shaker-related subfamily, beta member 3 [Source:MGI Symbol;Acc:MGI:1336208] | Kcnab3 | Enriched in dLGN | 4.293 | 6.989E-23 | 53.86 |
| ENSMUSG00000028873 | cell division cycle associated 8 [Source:MGI Symbol;Acc:MGI:1196274] | Cdca8 | Enriched in dLGN | 4.258 | 0.001222 | 5.365 |
| ENSMUSG00000037161 | mitochondria localized glutamic acid rich protein [Source:MGI Symbol;Acc:MGI:1914999] | Mgarp | Enriched in dLGN | 4.215 | 0.001105 | 530.5 |
| ENSMUSG00000037446 | tubby like protein 1 [Source:MGI Symbol;Acc:MGI:109571] | Tulp1 | Enriched in dLGN | 4.185 | 0.0002042 | 663.6 |
| ENSMUSG00000044375 | photoreceptor cilium actin regulator [Source:MGI Symbol;Acc:MGI:2385061] | Pcare | Enriched in dLGN | 4.131 | 0.0009867 | 962.8 |
| ENSMUSG00000045287 | reticulon 4 receptor-like 1 [Source:MGI Symbol;Acc:MGI:2661375] | Rtn4rl1 | Enriched in dLGN | 4.12 | 3.781E-46 | 303.8 |
| ENSMUSG00000038264 | sema domain, immunoglobulin domain (Ig), and GPI membrane anchor, (semaphorin) 7A [Source:MGI Symbol;Acc:MGI:1306826] | Sema7a | Enriched in dLGN | 4.111 | 6.125E-41 | 229.8 |
| ENSMUSG00000078816 | protein kinase C, gamma [Source:MGI Symbol;Acc:MGI:97597] | Prkcg | Enriched in dLGN | 4.099 | 4.082E-30 | 571.8 |
| ENSMUSG00000093865 | leucine-rich repeat, immunoglobulin-like and transmembrane domains 3 [Source:MGI Symbol;Acc:MGI:2685267] | Lrit3 | Enriched in dLGN | 4.084 | 0.01737 | 120.3 |
| ENSMUSG00000086322 | RIKEN cDNA E130218I03 gene [Source:MGI Symbol;Acc:MGI:3528958] | E130218I03Rik | Enriched in dLGN | 4.081 | 0.0001545 | 276.3 |
| ENSMUSG00000091387 | glucosaminyl (N-acetyl) transferase 4, core 2 (beta-1,6-N-acetylglucosaminyltransferase) [Source:MGI Symbol;Acc:MGI:2684919] | Gcnt4 | Enriched in dLGN | 4.066 | 9.06E-09 | 28.71 |
| ENSMUSG00000096740 | LBH domain containing 1 [Source:MGI Symbol;Acc:MGI:5516029] | Lbhd1 | Enriched in dLGN | 4.031 | 0.008123 | 104.6 |
| ENSMUSG00000040624 | pleckstrin homology domain containing, family G (with RhoGef domain) member 1 [Source:MGI Symbol;Acc:MGI:2676551] | Plekhg1 | Enriched in dLGN | 4.026 | 2.194E-46 | 465 |
| ENSMUSG00000048015 | neurogenic differentiation 4 [Source:MGI Symbol;Acc:MGI:108055] | Neurod4 | Enriched in dLGN | 3.957 | 0.003783 | 168.9 |
| ENSMUSG00000097050 | predicted gene 9918 [Source:MGI Symbol;Acc:MGI:3646315] | Gm9918 | Enriched in dLGN | 3.904 | 0.01309 | 57.93 |
| ENSMUSG00000054580 | phospholipase A2 receptor 1 [Source:MGI Symbol;Acc:MGI:102468] | Pla2r1 | Enriched in dLGN | 3.888 | 0.0004422 | 229.2 |
| ENSMUSG00000024227 | PDZ and pleckstrin homology domains 1 [Source:MGI Symbol;Acc:MGI:1916489] | Pdzph1 | Enriched in dLGN | 3.875 | 0.0001556 | 255.2 |
| ENSMUSG00000029211 | gamma-aminobutyric acid (GABA) A receptor, subunit alpha 4 [Source:MGI Symbol;Acc:MGI:95616] | Gabra4 | Enriched in dLGN | 3.847 | 1.228E-26 | 154.6 |
| ENSMUSG00000068696 | G-protein coupled receptor 88 [Source:MGI Symbol;Acc:MGI:1927653] | Gpr88 | Enriched in dLGN | 3.76 | 2.549E-29 | 66.64 |
| ENSMUSG00000040632 | neural retina leucine zipper gene [Source:MGI Symbol;Acc:MGI:102567] | Nrl | Enriched in dLGN | 3.759 | 0.001448 | 505.3 |
| ENSMUSG00000055027 | SET and MYND domain containing 1 [Source:MGI Symbol;Acc:MGI:104790] | Smyd1 | Enriched in dLGN | 3.751 | 1.222E-13 | 65.99 |
| ENSMUSG00000018821 | arginine vasopressin-induced 1 [Source:MGI Symbol;Acc:MGI:1916784] | Avpi1 | Enriched in dLGN | 3.74 | 1.843E-19 | 34.89 |
| ENSMUSG00000021359 | transcription factor AP-2, alpha [Source:MGI Symbol;Acc:MGI:104671] | Tfap2a | Enriched in dLGN | 3.732 | 0.01318 | 139.8 |
| ENSMUSG00000020953 | cochlin [Source:MGI Symbol;Acc:MGI:1278313] | Coch | Enriched in dLGN | 3.696 | 1.053E-12 | 80.94 |
| ENSMUSG00000049225 | pyruvate dehydrogenase phosphatase catalytic subunit 1 [Source:MGI Symbol;Acc:MGI:2685870] | Pdp1 | Enriched in dLGN | 3.693 | 3.818E-56 | 541.9 |
| ENSMUSG00000020733 | solute carrier family 9 (sodium/hydrogen exchanger), member 3 regulator 1 [Source:MGI Symbol;Acc:MGI:1349482] | Slc9a3r1 | Enriched in dLGN | 3.646 | 2.194E-46 | 214.9 |
| ENSMUSG00000059857 | netrin G1 [Source:MGI Symbol;Acc:MGI:1934028] | Ntng1 | Enriched in dLGN | 3.645 | 3.746E-18 | 904 |
| ENSMUSG00000027347 | RAS guanyl releasing protein 1 [Source:MGI Symbol;Acc:MGI:1314635] | Rasgrp1 | Enriched in dLGN | 3.635 | 2.419E-20 | 353.4 |
| ENSMUSG00000027530 | fatty acid binding protein 12 [Source:MGI Symbol;Acc:MGI:1922747] | Fabp12 | Enriched in dLGN | 3.623 | 0.006743 | 148.6 |
| ENSMUSG00000026235 | Eph receptor A4 [Source:MGI Symbol;Acc:MGI:98277] | Epha4 | Enriched in dLGN | 3.615 | 3.467E-37 | 500.9 |
| ENSMUSG00000030340 | sodium channel, nonvoltage-gated 1 alpha [Source:MGI Symbol;Acc:MGI:101782] | Scnn1a | Enriched in dLGN | 3.614 | 2.305E-15 | 44.62 |
| ENSMUSG00000033615 | complexin 1 [Source:MGI Symbol;Acc:MGI:104727] | Cplx1 | Enriched in dLGN | 3.612 | 3.098E-24 | 2304 |
| ENSMUSG00000005649 | calcium binding protein 5 [Source:MGI Symbol;Acc:MGI:1352746] | Cabp5 | Enriched in dLGN | 3.559 | 0.01605 | 50.22 |
| ENSMUSG00000004630 | Purkinje cell protein 2 (L7) [Source:MGI Symbol;Acc:MGI:97508] | Pcp2 | Enriched in dLGN | 3.493 | 0.03434 | 42.06 |
| ENSMUSG00000022014 | epithelial stromal interaction 1 (breast) [Source:MGI Symbol;Acc:MGI:1915168] | Epsti1 | Enriched in dLGN | 3.475 | 0.00008083 | 6.466 |
| ENSMUSG00000054196 | collagen triple helix repeat containing 1 [Source:MGI Symbol;Acc:MGI:1915838] | Cthrc1 | Enriched in dLGN | 3.473 | 1.783E-14 | 38 |
| ENSMUSG00000027716 | transient receptor potential cation channel, subfamily C, member 3 [Source:MGI Symbol;Acc:MGI:109526] | Trpc3 | Enriched in dLGN | 3.472 | 7.304E-30 | 212.4 |
| ENSMUSG00000067220 | cyclic nucleotide gated channel alpha 1 [Source:MGI Symbol;Acc:MGI:88436] | Cnga1 | Enriched in dLGN | 3.381 | 0.0003032 | 478.2 |
| ENSMUSG00000023905 | tumor necrosis factor receptor superfamily, member 12a [Source:MGI Symbol;Acc:MGI:1351484] | Tnfrsf12a | Enriched in dLGN | 3.357 | 9.454E-07 | 7.873 |
| ENSMUSG00000024727 | transient receptor potential cation channel, subfamily M, member 6 [Source:MGI Symbol;Acc:MGI:2675603] | Trpm6 | Enriched in dLGN | 3.357 | 0.00001107 | 19.86 |
| ENSMUSG00000032908 | sphingosine-1-phosphate phosphatase 2 [Source:MGI Symbol;Acc:MGI:3589109] | Sgpp2 | Enriched in dLGN | 3.348 | 1.067E-17 | 115.4 |
| ENSMUSG00000024987 | cytochrome P450, family 26, subfamily a, polypeptide 1 [Source:MGI Symbol;Acc:MGI:1096359] | Cyp26a1 | Enriched in dLGN | 3.336 | 0.04357 | 6.311 |
| ENSMUSG00000078963 | heat shock factor binding protein 1-like 1 [Source:MGI Symbol;Acc:MGI:1913505] | Hsbp1l1 | Enriched in dLGN | 3.311 | 0.0000103 | 35.03 |
| ENSMUSG00000021719 | regulator of G-protein signalling 7 binding protein [Source:MGI Symbol;Acc:MGI:106334] | Rgs7bp | Enriched in dLGN | 3.25 | 2.328E-30 | 598.8 |
| ENSMUSG00000034570 | inositol polyphosphate 5-phosphatase J [Source:MGI Symbol;Acc:MGI:2158663] | Inpp5j | Enriched in dLGN | 3.239 | 1.276E-20 | 112.1 |
| ENSMUSG00000028082 | SH3 domain protein D19 [Source:MGI Symbol;Acc:MGI:1350923] | Sh3d19 | Enriched in dLGN | 3.233 | 4.057E-20 | 657.1 |
| ENSMUSG00000024992 | phosphodiesterase 6C, cGMP specific, cone, alpha prime [Source:MGI Symbol;Acc:MGI:105956] | Pde6c | Enriched in dLGN | 3.206 | 0.01136 | 100.5 |
| ENSMUSG00000075330 | RIKEN cDNA A930003A15 gene [Source:MGI Symbol;Acc:MGI:1915412] | A930003A15Rik | Enriched in dLGN | 3.199 | 0.03493 | 40.93 |
| ENSMUSG00000040713 | cellular repressor of E1A-stimulated genes 1 [Source:MGI Symbol;Acc:MGI:1344382] | Creg1 | Enriched in dLGN | 3.18 | 2.134E-20 | 76.45 |
| ENSMUSG00000037996 | solute carrier family 24 (sodium/potassium/calcium exchanger), member 2 [Source:MGI Symbol;Acc:MGI:1923626] | Slc24a2 | Enriched in dLGN | 3.159 | 5.639E-18 | 1329 |
| ENSMUSG00000040723 | RCSD domain containing 1 [Source:MGI Symbol;Acc:MGI:2676394] | Rcsd1 | Enriched in dLGN | 3.15 | 5.231E-10 | 27.5 |
| ENSMUSG00000026609 | usherin [Source:MGI Symbol;Acc:MGI:1341292] | Ush2a | Enriched in dLGN | 3.104 | 0.0004313 | 1475 |
| ENSMUSG00000028149 | RAP1, GTP-GDP dissociation stimulator 1 [Source:MGI Symbol;Acc:MGI:2385189] | Rap1gds1 | Enriched in dLGN | 3.092 | 1.561E-48 | 847.3 |
| ENSMUSG00000039115 | integrin alpha 9 [Source:MGI Symbol;Acc:MGI:104756] | Itga9 | Enriched in dLGN | 3.079 | 0.0006117 | 1811 |
| ENSMUSG00000065968 | interferon induced transmembrane protein 7 [Source:MGI Symbol;Acc:MGI:1921732] | Ifitm7 | Enriched in dLGN | 3.039 | 3.883E-07 | 15.26 |
| ENSMUSG00000029769 | coiled-coil domain 136 [Source:MGI Symbol;Acc:MGI:1918128] | Ccdc136 | Enriched in dLGN | 2.987 | 9.95E-42 | 1034 |
| ENSMUSG00000030189 | Y box protein 3 [Source:MGI Symbol;Acc:MGI:2137670] | Ybx3 | Enriched in dLGN | 2.958 | 3.052E-17 | 527 |
| ENSMUSG00000110711 | predicted gene 45760 [Source:MGI Symbol;Acc:MGI:5804875] | Gm45760 | Enriched in dLGN | 2.944 | 0.007753 | 2.941 |
| ENSMUSG00000003541 | immediate early response 3 [Source:MGI Symbol;Acc:MGI:104814] | Ier3 | Enriched in dLGN | 2.941 | 3.256E-07 | 21.94 |
| ENSMUSG00000041460 | calcium channel, voltage-dependent, alpha 2/delta subunit 4 [Source:MGI Symbol;Acc:MGI:2442632] | Cacna2d4 | Enriched in dLGN | 2.938 | 6.07E-14 | 641.5 |
| ENSMUSG00000029632 | Ndufa4, mitochondrial complex associated [Source:MGI Symbol;Acc:MGI:107686] | Ndufa4 | Enriched in dLGN | 2.911 | 2.964E-17 | 183.2 |
| ENSMUSG00000026384 | protein tyrosine phosphatase, non-receptor type 4 [Source:MGI Symbol;Acc:MGI:1099792] | Ptpn4 | Enriched in dLGN | 2.899 | 1.229E-24 | 1013 |
| ENSMUSG00000110344 | small integral membrane protein 36 [Source:MGI Symbol;Acc:MGI:5804831] | Smim36 | Enriched in dLGN | 2.885 | 0.02304 | 107.5 |
| ENSMUSG00000053773 | retinol dehydrogenase 8 [Source:MGI Symbol;Acc:MGI:2685028] | Rdh8 | Enriched in dLGN | 2.881 | 0.02772 | 87.45 |
| ENSMUSG00000032564 | copine IV [Source:MGI Symbol;Acc:MGI:1921270] | Cpne4 | Enriched in dLGN | 2.863 | 1.554E-11 | 126.3 |
| ENSMUSG00000039629 | striatin interacting protein 2 [Source:MGI Symbol;Acc:MGI:2444363] | Strip2 | Enriched in dLGN | 2.831 | 1.307E-14 | 138 |
| ENSMUSG00000028744 | solute carrier family 66 member 1 [Source:MGI Symbol;Acc:MGI:2384837] | Slc66a1 | Enriched in dLGN | 2.825 | 0.000004436 | 195.7 |
| ENSMUSG00000027792 | butyrylcholinesterase [Source:MGI Symbol;Acc:MGI:894278] | Bche | Enriched in dLGN | 2.817 | 3.159E-07 | 17.98 |
| ENSMUSG00000031231 | cytochrome c oxidase subunit 7B [Source:MGI Symbol;Acc:MGI:1913392] | Cox7b | Enriched in dLGN | 2.8 | 7.498E-08 | 113.9 |

| | | | | | | |
|---|---|---|---|---|---|---|
| ENSMUSG00000014313 | cytochrome c oxidase subunit 6C [Source:MGI Symbol;Acc:MGI:104614] | Cox6c | Enriched in dLGN | 2.795 | 3.508E-15 | 345.4 |
| ENSMUSG00000027489 | N-terminal EF-hand calcium binding protein 3 [Source:MGI Symbol;Acc:MGI:1861721] | Necab3 | Enriched in dLGN | 2.79 | 2.107E-14 | 36.72 |
| ENSMUSG00000038128 | calcium/calmodulin-dependent protein kinase IV [Source:MGI Symbol;Acc:MGI:88258] | Camk4 | Enriched in dLGN | 2.767 | 2.699E-28 | 937 |
| ENSMUSG00000002147 | signal transducer and activator of transcription 6 [Source:MGI Symbol;Acc:MGI:103034] | Stat6 | Enriched in dLGN | 2.751 | 0.000000163 | 22.91 |
| ENSMUSG00000071866 | peptidylprolyl isomerase A [Source:MGI Symbol;Acc:MGI:97749] | Ppia | Enriched in dLGN | 2.738 | 8.279E-07 | 211.2 |
| ENSMUSG00000096764 | predicted gene 21985 [Source:MGI Symbol;Acc:MGI:5439454] | Gm21985 | Enriched in dLGN | 2.704 | 0.002551 | 6.131 |
| ENSMUSG00000061859 | PATJ, crumbs cell polarity complex component [Source:MGI Symbol;Acc:MGI:1277960] | Patj | Enriched in dLGN | 2.697 | 4.051E-12 | 609.6 |
| ENSMUSG00000029641 | RAS-like, family 11, member A [Source:MGI Symbol;Acc:MGI:1916145] | Rasl11a | Enriched in dLGN | 2.697 | 3.664E-07 | 21.43 |
| ENSMUSG00000027712 | annexin A5 [Source:MGI Symbol;Acc:MGI:106008] | Anxa5 | Enriched in dLGN | 2.693 | 1.046E-09 | 102.3 |
| ENSMUSG00000020890 | guanylate cyclase 2e [Source:MGI Symbol;Acc:MGI:105123] | Gucy2e | Enriched in dLGN | 2.689 | 0.003696 | 507.5 |
| ENSMUSG00000029610 | aminoacyl tRNA synthetase complex-interacting multifunctional protein 2 [Source:MGI Symbol;Acc:MGI:2385237] | Aimp2 | Enriched in dLGN | 2.682 | 3.561E-09 | 26.12 |
| ENSMUSG00000053930 | shisa family member 6 [Source:MGI Symbol;Acc:MGI:2685725] | Shisa6 | Enriched in dLGN | 2.677 | 8.848E-15 | 186.6 |
| ENSMUSG00000045625 | phosphatidylinositol glycan anchor biosynthesis, class Z [Source:MGI Symbol;Acc:MGI:2443822] | Pigz | Enriched in dLGN | 2.668 | 0.000001031 | 18.26 |
| ENSMUSG00000037568 | vasohibin 2 [Source:MGI Symbol;Acc:MGI:2444826] | Vash2 | Enriched in dLGN | 2.667 | 6.195E-17 | 93.06 |
| ENSMUSG00000019943 | ATPase, Ca++ transporting, plasma membrane 1 [Source:MGI Symbol;Acc:MGI:104653] | Atp2b1 | Enriched in dLGN | 2.665 | 5.052E-47 | 2706 |
| ENSMUSG00000053025 | synaptic vesicle glycoprotein 2 b [Source:MGI Symbol;Acc:MGI:1927338] | Sv2b | Enriched in dLGN | 2.653 | 3.459E-29 | 914.9 |
| ENSMUSG00000057132 | retinitis pigmentosa GTPase regulator interacting protein 1 [Source:MGI Symbol;Acc:MGI:1932134] | Rpgrip1 | Enriched in dLGN | 2.623 | 0.005268 | 1656 |
| ENSMUSG00000037492 | zinc finger, matrin type 4 [Source:MGI Symbol;Acc:MGI:2443497] | Zmat4 | Enriched in dLGN | 2.619 | 2.483E-14 | 529.7 |
| ENSMUSG00000022856 | transmembrane protein 41a [Source:MGI Symbol;Acc:MGI:1913914] | Tmem41a | Enriched in dLGN | 2.593 | 4.878E-16 | 44.73 |
| ENSMUSG00000028710 | ATP synthase mitochondrial F1 complex assembly factor 1 [Source:MGI Symbol;Acc:MGI:2180560] | Atpaf1 | Enriched in dLGN | 2.591 | 2.342E-23 | 89.62 |
| ENSMUSG00000079550 | membrane protein, palmitoylated 4 (MAGUK p55 subfamily member 4) [Source:MGI Symbol;Acc:MGI:2386681] | Mpp4 | Enriched in dLGN | 2.585 | 0.01139 | 514.2 |
| ENSMUSG00000002032 | transmembrane protein 25 [Source:MGI Symbol;Acc:MGI:1918937] | Tmem25 | Enriched in dLGN | 2.582 | 1.614E-16 | 70.33 |
| ENSMUSG00000020067 | myopalladin [Source:MGI Symbol;Acc:MGI:1916052] | Mypn | Enriched in dLGN | 2.574 | 0.0002812 | 14.16 |
| ENSMUSG00000016319 | solute carrier family 25 (mitochondrial carrier, adenine nucleotide translocator), member 5 [Source:MGI Symbol;Acc:MGI:1353496] | Slc25a5 | Enriched in dLGN | 2.562 | 2.954E-10 | 198.8 |
| ENSMUSG00000030500 | solute carrier family 17 (sodium-dependent inorganic phosphate cotransporter), member 6 [Source:MGI Symbol;Acc:MGI:2156052] | Slc17a6 | Enriched in dLGN | 2.561 | 4.978E-09 | 605.1 |
| ENSMUSG00000050541 | adrenergic receptor, alpha 1b [Source:MGI Symbol;Acc:MGI:104774] | Adra1b | Enriched in dLGN | 2.558 | 3.387E-12 | 73.77 |
| ENSMUSG00000043461 | serine palmitoyltransferase, small subunit B [Source:MGI Symbol;Acc:MGI:1913433] | Sptssb | Enriched in dLGN | 2.525 | 0.0001136 | 9.484 |
| ENSMUSG00000022180 | solute carrier family 7 (cationic amino acid transporter, y+ system), member 8 [Source:MGI Symbol;Acc:MGI:1355323] | Slc7a8 | Enriched in dLGN | 2.52 | 2.371E-18 | 249.8 |
| ENSMUSG00000029370 | Ras association (RalGDS/AF-6) domain family member 6 [Source:MGI Symbol;Acc:MGI:1920496] | Rassf6 | Enriched in dLGN | 2.518 | 0.001351 | 5.633 |
| ENSMUSG00000035000 | dipeptidylpeptidase 4 [Source:MGI Symbol;Acc:MGI:94919] | Dpp4 | Enriched in dLGN | 2.513 | 0.0002549 | 9.243 |
| ENSMUSG00000031465 | angiopoietin 2 [Source:MGI Symbol;Acc:MGI:1202890] | Angpt2 | Enriched in dLGN | 2.507 | 0.0003046 | 14.62 |
| ENSMUSG00000043629 | RIKEN cDNA 1700019D03 gene [Source:MGI Symbol;Acc:MGI:1914330] | 1700019D03Rik | Enriched in dLGN | 2.487 | 1.538E-09 | 31.11 |
| ENSMUSG00000022820 | NADH:ubiquinone oxidoreductase subunit B4 [Source:MGI Symbol;Acc:MGI:1915444] | Ndufb4 | Enriched in dLGN | 2.482 | 0.0003195 | 32.87 |
| ENSMUSG00000036578 | FXYD domain-containing ion transport regulator 7 [Source:MGI Symbol;Acc:MGI:1889006] | Fxyd7 | Enriched in dLGN | 2.481 | 1.238E-10 | 140.4 |
| ENSMUSG00000062070 | phosphoglycerate kinase 1 [Source:MGI Symbol;Acc:MGI:97555] | Pgk1 | Enriched in dLGN | 2.476 | 1.1E-15 | 300 |
| ENSMUSG00000035506 | solute carrier family 12 (potassium/chloride transporters), member 8 [Source:MGI Symbol;Acc:MGI:2443672] | Slc12a8 | Enriched in dLGN | 2.474 | 0.0002013 | 17.24 |
| ENSMUSG00000051022 | heparan sulfate (glucosamine) 3-O-sulfotransferase 1 [Source:MGI Symbol;Acc:MGI:1201606] | Hs3st1 | Enriched in dLGN | 2.471 | 9.143E-18 | 132 |
| ENSMUSG00000024064 | polypeptide N-acetylgalactosaminyltransferase 14 [Source:MGI Symbol;Acc:MGI:1918935] | Galnt14 | Enriched in dLGN | 2.464 | 4.362E-11 | 190 |
| ENSMUSG00000020907 | recoverin [Source:MGI Symbol;Acc:MGI:97883] | Rcvrn | Enriched in dLGN | 2.46 | 0.02728 | 323.5 |
| ENSMUSG00000026888 | growth factor receptor bound protein 14 [Source:MGI Symbol;Acc:MGI:1355324] | Grb14 | Enriched in dLGN | 2.455 | 2.082E-10 | 102.4 |
| ENSMUSG00000020116 | partner of NOB1 homolog [Source:MGI Symbol;Acc:MGI:1913499] | Pno1 | Enriched in dLGN | 2.44 | 1.301E-11 | 43.22 |
| ENSMUSG00000044252 | oxysterol binding protein-like 1A [Source:MGI Symbol;Acc:MGI:1927551] | Osbpl1a | Enriched in dLGN | 2.438 | 1.751E-23 | 434.2 |
| ENSMUSG00000030557 | myocyte enhancer factor 2A [Source:MGI Symbol;Acc:MGI:99532] | Mef2a | Enriched in dLGN | 2.437 | 3.624E-24 | 532.8 |
| ENSMUSG00000027674 | peroxisomal biogenesis factor 5-like [Source:MGI Symbol;Acc:MGI:1916672] | Pex5l | Enriched in dLGN | 2.416 | 7.355E-09 | 789.1 |
| ENSMUSG00000025475 | adhesion G protein-coupled receptor A1 [Source:MGI Symbol;Acc:MGI:1277167] | Adgra1 | Enriched in dLGN | 2.415 | 1.872E-25 | 360.1 |
| ENSMUSG00000025092 | heat shock protein 12A [Source:MGI Symbol;Acc:MGI:1920692] | Hspa12a | Enriched in dLGN | 2.409 | 2.015E-24 | 913 |
| ENSMUSG00000008932 | solute carrier family 1 (glutamate transporter), member 7 [Source:MGI Symbol;Acc:MGI:2444087] | Slc1a7 | Enriched in dLGN | 2.407 | 0.03806 | 185.9 |
| ENSMUSG00000029189 | sel-1 suppressor of lin-12-like 3 (C. elegans) [Source:MGI Symbol;Acc:MGI:1916941] | Sel1l3 | Enriched in dLGN | 2.406 | 3.887E-16 | 154 |
| ENSMUSG00000021803 | cadherin-related family member 1 [Source:MGI Symbol;Acc:MGI:2157782] | Cdhr1 | Enriched in dLGN | 2.394 | 0.003008 | 1092 |
| ENSMUSG00000027610 | glutathione synthetase [Source:MGI Symbol;Acc:MGI:95852] | Gss | Enriched in dLGN | 2.388 | 8.543E-12 | 48.23 |
| ENSMUSG00000018648 | dual specificity phosphatase 14 [Source:MGI Symbol;Acc:MGI:1927168] | Dusp14 | Enriched in dLGN | 2.382 | 0.0000043 | 23.93 |
| ENSMUSG00000027495 | family with sequence similarity 210, member B [Source:MGI Symbol;Acc:MGI:1914267] | Fam210b | Enriched in dLGN | 2.381 | 1.126E-10 | 55.78 |
| ENSMUSG00000040794 | C1q and tumor necrosis factor related protein 4 [Source:MGI Symbol;Acc:MGI:1914695] | C1qtnf4 | Enriched in dLGN | 2.38 | 1.983E-19 | 207.8 |
| ENSMUSG00000028367 | thioredoxin 1 [Source:MGI Symbol;Acc:MGI:98874] | Txn1 | Enriched in dLGN | 2.371 | 9.049E-13 | 92.9 |
| ENSMUSG00000027556 | carbonic anhydrase 1 [Source:MGI Symbol;Acc:MGI:88268] | Car1 | Enriched in dLGN | 2.369 | 0.00004497 | 12.27 |
| ENSMUSG00000021253 | transforming growth factor, beta 3 [Source:MGI Symbol;Acc:MGI:98727] | Tgfb3 | Enriched in dLGN | 2.366 | 9.802E-15 | 74.63 |
| ENSMUSG00000032965 | intraflagellar transport 57 [Source:MGI Symbol;Acc:MGI:1921166] | Ift57 | Enriched in dLGN | 2.358 | 1.179E-25 | 254.5 |
| ENSMUSG00000022454 | NEL-like 2 [Source:MGI Symbol;Acc:MGI:1858510] | Nell2 | Enriched in dLGN | 2.354 | 1.046E-18 | 1350 |
| ENSMUSG00000024935 | solute carrier family 1 (neuronal/epithelial high affinity glutamate transporter, system Xag), member 1 [Source:MGI Symbol;Acc:MGI:105083] | Slc1a1 | Enriched in dLGN | 2.337 | 1.973E-23 | 323.7 |
| ENSMUSG00000034825 | nuclear receptor interacting protein 3 [Source:MGI Symbol;Acc:MGI:1925843] | Nrip3 | Enriched in dLGN | 2.324 | 6.447E-11 | 285.5 |
| ENSMUSG00000038421 | Fc receptor-like A [Source:MGI Symbol;Acc:MGI:2138647] | Fcrla | Enriched in dLGN | 2.322 | 0.00134 | 28.16 |
| ENSMUSG00000031636 | PDZ and LIM domain 3 [Source:MGI Symbol;Acc:MGI:1859274] | Pdlim3 | Enriched in dLGN | 2.316 | 0.01081 | 14.05 |
| ENSMUSG00000041144 | dynein, axonemal, heavy chain 7B [Source:MGI Symbol;Acc:MGI:2684953] | Dnah7b | Enriched in dLGN | 2.281 | 0.03613 | 449.4 |
| ENSMUSG00000038552 | fibronectin type III domain containing 4 [Source:MGI Symbol;Acc:MGI:1917195] | Fndc4 | Enriched in dLGN | 2.276 | 9.053E-30 | 238.1 |
| ENSMUSG00000022900 | immunoglobulin-like domain containing receptor 1 [Source:MGI Symbol;Acc:MGI:2146574] | Ildr1 | Enriched in dLGN | 2.271 | 0.01969 | 4.707 |
| ENSMUSG00000041112 | engulfment and cell motility 1 [Source:MGI Symbol;Acc:MGI:2153044] | Elmo1 | Enriched in dLGN | 2.268 | 5.658E-12 | 598.8 |
| ENSMUSG00000062937 | methylthioadenosine phosphorylase [Source:MGI Symbol;Acc:MGI:1914152] | Mtap | Enriched in dLGN | 2.26 | 9.704E-07 | 29.38 |
| ENSMUSG00000036745 | tubulin tyrosine ligase-like family, member 7 [Source:MGI Symbol;Acc:MGI:1918142] | Ttll7 | Enriched in dLGN | 2.257 | 8.789E-14 | 648.1 |
| ENSMUSG00000032892 | RAN guanine nucleotide release factor [Source:MGI Symbol;Acc:MGI:1889073] | Rangrf | Enriched in dLGN | 2.255 | 0.002483 | 6.22 |
| ENSMUSG00000030350 | protein arginine N-methyltransferase 8 [Source:MGI Symbol;Acc:MGI:3043083] | Prmt8 | Enriched in dLGN | 2.252 | 9.492E-20 | 303.1 |
| ENSMUSG00000027533 | fatty acid binding protein 5, epidermal [Source:MGI Symbol;Acc:MGI:101790] | Fabp5 | Enriched in dLGN | 2.249 | 0.0002243 | 258.1 |
| ENSMUSG00000064288 | H4 clustered histone 12 [Source:MGI Symbol;Acc:MGI:2448439] | H4c12 | Enriched in dLGN | 2.245 | 0.0003775 | 10.87 |
| ENSMUSG00000040612 | immunoglobulin-like domain containing receptor 2 [Source:MGI Symbol;Acc:MGI:1196370] | Ildr2 | Enriched in dLGN | 2.24 | 3.431E-13 | 1307 |
| ENSMUSG00000003949 | hepatic leukemia factor [Source:MGI Symbol;Acc:MGI:96108] | Hlf | Enriched in dLGN | 2.238 | 2.85E-12 | 354.9 |
| ENSMUSG00000021520 | ubiquinol-cytochrome c reductase binding protein [Source:MGI Symbol;Acc:MGI:1914780] | Uqcrb | Enriched in dLGN | 2.237 | 5.059E-07 | 49.09 |
| ENSMUSG00000062785 | potassium voltage gated channel, Shaw-related subfamily, member 3 [Source:MGI Symbol;Acc:MGI:96669] | Kcnc3 | Enriched in dLGN | 2.236 | 2.855E-24 | 437.6 |
| ENSMUSG00000033307 | macrophage migration inhibitory factor (glycosylation-inhibiting factor) [Source:MGI Symbol;Acc:MGI:96982] | Mif | Enriched in dLGN | 2.23 | 0.000001369 | 90.39 |
| ENSMUSG00000026568 | mitochondrial pyruvate carrier 2 [Source:MGI Symbol;Acc:MGI:1917706] | Mpc2 | Enriched in dLGN | 2.228 | 2.347E-13 | 84.48 |
| ENSMUSG00000028488 | SH3-domain GRB2-like 2 [Source:MGI Symbol;Acc:MGI:700009] | Sh3gl2 | Enriched in dLGN | 2.227 | 3.146E-12 | 586.9 |
| ENSMUSG00000040430 | phosphatidylinositol transfer protein, cytoplasmic 1 [Source:MGI Symbol;Acc:MGI:1919045] | Pitpnc1 | Enriched in dLGN | 2.225 | 9.36E-25 | 764.9 |
| ENSMUSG00000033316 | polypeptide N-acetylgalactosaminyltransferase 9 [Source:MGI Symbol;Acc:MGI:2677965] | Galnt9 | Enriched in dLGN | 2.225 | 3.112E-18 | 204.6 |
| ENSMUSG00000064329 | sodium channel, voltage-gated, type I, alpha [Source:MGI Symbol;Acc:MGI:98246] | Scn1a | Enriched in dLGN | 2.217 | 3.387E-12 | 703.3 |
| ENSMUSG00000035270 | interphotoreceptor matrix proteoglycan 2 [Source:MGI Symbol;Acc:MGI:3044955] | Impg2 | Enriched in dLGN | 2.217 | 0.01514 | 754.3 |
| ENSMUSG00000032011 | thymus cell antigen 1, theta [Source:MGI Symbol;Acc:MGI:98747] | Thy1 | Enriched in dLGN | 2.215 | 1.365E-10 | 952.8 |
| ENSMUSG00000111375 | BTB (POZ) domain containing 8 [Source:MGI Symbol;Acc:MGI:3646208] | Btbd8 | Enriched in dLGN | 2.211 | 3.13E-12 | 197 |
| ENSMUSG00000069274 | H4 clustered histone 6 [Source:MGI Symbol;Acc:MGI:2448425] | H4c6 | Enriched in dLGN | 2.206 | 0.001132 | 11.96 |
| ENSMUSG00000026260 | NADH:ubiquinone oxidoreductase subunit A10 [Source:MGI Symbol;Acc:MGI:1914523] | Ndufa10 | Enriched in dLGN | 2.202 | 6.547E-11 | 263.7 |
| ENSMUSG00000075256 | ceramide kinase-like [Source:MGI Symbol;Acc:MGI:3037816] | Cerkl | Enriched in dLGN | 2.2 | 0.002896 | 144.8 |
| ENSMUSG00000021752 | potassium channel tetramerisation domain containing 6 [Source:MGI Symbol;Acc:MGI:1918643] | Kctd6 | Enriched in dLGN | 2.199 | 1.99E-12 | 62.84 |
| ENSMUSG00000024190 | dual specificity phosphatase 1 [Source:MGI Symbol;Acc:MGI:105120] | Dusp1 | Enriched in dLGN | 2.17 | 1.291E-14 | 44.73 |
| ENSMUSG00000032578 | cytokine inducible SH2-containing protein [Source:MGI Symbol;Acc:MGI:103159] | Cish | Enriched in dLGN | 2.165 | 0.00004943 | 23.3 |
| ENSMUSG00000037606 | oxysterol binding protein-like 5 [Source:MGI Symbol;Acc:MGI:1930265] | Osbpl5 | Enriched in dLGN | 2.161 | 1.518E-29 | 213.9 |
| ENSMUSG00000062542 | synaptotagmin IX [Source:MGI Symbol;Acc:MGI:1926373] | Syt9 | Enriched in dLGN | 2.161 | 1.159E-22 | 283.3 |
| ENSMUSG00000028251 | thiosulfate sulfurtransferase (rhodanese)-like domain containing 3 [Source:MGI Symbol;Acc:MGI:1924282] | Tstd3 | Enriched in dLGN | 2.16 | 2.081E-07 | 35.11 |
| ENSMUSG00000036158 | prickle planar cell polarity protein 1 [Source:MGI Symbol;Acc:MGI:1916034] | Prickle1 | Enriched in dLGN | 2.155 | 1.641E-16 | 385.1 |
| ENSMUSG00000016252 | ATP synthase, H+ transporting, mitochondrial F1 complex, epsilon subunit [Source:MGI Symbol;Acc:MGI:1855697] | Atp5e | Enriched in dLGN | 2.152 | 6.322E-09 | 153.2 |
| ENSMUSG00000027076 | translocase of inner mitochondrial membrane 10 [Source:MGI Symbol;Acc:MGI:1353429] | Timm10 | Enriched in dLGN | 2.15 | 1.073E-07 | 29.72 |
| ENSMUSG00000063531 | sema domain, immunoglobulin domain (Ig), short basic domain, secreted, (semaphorin) 3E [Source:MGI Symbol;Acc:MGI:1340034] | Sema3e | Enriched in dLGN | 2.145 | 0.000005917 | 112.6 |
| ENSMUSG00000040536 | N-terminal EF-hand calcium binding protein 1 [Source:MGI Symbol;Acc:MGI:1916602] | Necab1 | Enriched in dLGN | 2.138 | 3.237E-09 | 133 |
| ENSMUSG00000078864 | predicted gene 14322 [Source:MGI Symbol;Acc:MGI:3652185] | Gm14322 | Enriched in dLGN | 2.133 | 0.0288 | 6.289 |
| ENSMUSG00000024172 | beta galactoside alpha 2,6 sialyltransferase 2 [Source:MGI Symbol;Acc:MGI:2445190] | St6gal2 | Enriched in dLGN | 2.127 | 4.541E-11 | 150.3 |
| ENSMUSG00000045733 | shadow of prion protein [Source:MGI Symbol;Acc:MGI:3582583] | Sprn | Enriched in dLGN | 2.126 | 7.24E-22 | 239 |
| ENSMUSG00000048100 | TATA-box binding protein associated factor 13 [Source:MGI Symbol;Acc:MGI:1913500] | Taf13 | Enriched in dLGN | 2.113 | 0.000000108 | 38.39 |
| ENSMUSG00000028718 | Scl/Tal1 interrupting locus [Source:MGI Symbol;Acc:MGI:107477] | Stil | Enriched in dLGN | 2.113 | 0.01574 | 5.074 |
| ENSMUSG00000028289 | Eph receptor A7 [Source:MGI Symbol;Acc:MGI:95276] | Epha7 | Enriched in dLGN | 2.109 | 5.704E-11 | 197.6 |
| ENSMUSG00000037152 | NADH:ubiquinone oxidoreductase subunit C1 [Source:MGI Symbol;Acc:MGI:1913627] | Ndufc1 | Enriched in dLGN | 2.109 | 3.019E-07 | 49.47 |
| ENSMUSG00000000901 | matrix metallopeptidase 11 [Source:MGI Symbol;Acc:MGI:97008] | Mmp11 | Enriched in dLGN | 2.104 | 0.0006266 | 8.781 |
| ENSMUSG00000021259 | cytochrome P450, family 46, subfamily a, polypeptide 1 [Source:MGI Symbol;Acc:MGI:1341877] | Cyp46a1 | Enriched in dLGN | 2.103 | 6.986E-21 | 248.2 |
| ENSMUSG00000021290 | ATP synthase membrane subunit 6.8PL [Source:MGI Symbol;Acc:MGI:1917507] | Atp5mpl | Enriched in dLGN | 2.1 | 0.000002771 | 181.4 |
| ENSMUSG00000062352 | integrin beta 1 binding protein 1 [Source:MGI Symbol;Acc:MGI:1306802] | Itgb1bp1 | Enriched in dLGN | 2.096 | 1.535E-13 | 59.33 |
| ENSMUSG00000028773 | fatty acid binding protein 3, muscle and heart [Source:MGI Symbol;Acc:MGI:95476] | Fabp3 | Enriched in dLGN | 2.094 | 0.0001449 | 75.72 |
| ENSMUSG00000054843 | attractin like 1 [Source:MGI Symbol;Acc:MGI:2147749] | Atrnl1 | Enriched in dLGN | 2.084 | 1.109E-11 | 629.5 |
| ENSMUSG00000021591 | glutaredoxin [Source:MGI Symbol;Acc:MGI:2135625] | Glrx | Enriched in dLGN | 2.081 | 1.005E-11 | 37.82 |
| ENSMUSG00000023904 | host cell factor C1 regulator 1 (XPO1-dependent) [Source:MGI Symbol;Acc:MGI:2663619] | Hcfc1r1 | Enriched in dLGN | 2.08 | 4.832E-09 | 96.23 |
| ENSMUSG00000001056 | NHP2 ribonucleoprotein [Source:MGI Symbol;Acc:MGI:1098547] | Nhp2 | Enriched in dLGN | 2.079 | 8.196E-09 | 27.46 |
| ENSMUSG00000040373 | calcium channel, voltage-dependent, gamma subunit 5 [Source:MGI Symbol;Acc:MGI:2157946] | Cacng5 | Enriched in dLGN | 2.064 | 9.53E-15 | 203.8 |
| ENSMUSG00000020140 | leucine rich repeat containing G protein coupled receptor 5 [Source:MGI Symbol;Acc:MGI:1341817] | Lgr5 | Enriched in dLGN | 2.061 | 0.00001547 | 46.54 |
| ENSMUSG00000033998 | potassium channel, subfamily K, member 1 [Source:MGI Symbol;Acc:MGI:109322] | Kcnk1 | Enriched in dLGN | 2.059 | 4.263E-11 | 107.5 |
| ENSMUSG00000024529 | lysyl oxidase [Source:MGI Symbol;Acc:MGI:96817] | Lox | Enriched in dLGN | 2.051 | 0.002524 | 9.017 |
| ENSMUSG00000096351 | sterile alpha motif domain containing 11 [Source:MGI Symbol;Acc:MGI:2446220] | Samd11 | Enriched in dLGN | 2.046 | 0.0002974 | 401.5 |
| ENSMUSG00000045009 | proline-rich transmembrane protein 3 [Source:MGI Symbol;Acc:MGI:2444810] | Prrt3 | Enriched in dLGN | 2.027 | 5.055E-12 | 111.1 |
| ENSMUSG00000113902 | NADH:ubiquinone oxidoreductase subunit B1 [Source:MGI Symbol;Acc:MGI:3780865] | Ndufb1 | Enriched in dLGN | 2.026 | 8.254E-07 | 43.14 |
| ENSMUSG00000048332 | lipoma HMGIC fusion partner [Source:MGI Symbol;Acc:MGI:1920048] | Lhfp | Enriched in dLGN | 2.013 | 9.076E-07 | 294.9 |
| ENSMUSG00000020949 | FK506 binding protein 3 [Source:MGI Symbol;Acc:MGI:1353460] | Fkbp3 | Enriched in dLGN | 2.007 | 2.168E-08 | 160.5 |
| ENSMUSG00000032435 | dynein cytoplasmic 1 light intermediate chain 1 [Source:MGI Symbol;Acc:MGI:2135610] | Dync1li1 | Enriched in dLGN | 2.001 | 1.82E-45 | 559.8 |
| ENSMUSG00000056258 | potassium voltage-gated channel, subfamily Q, member 3 [Source:MGI Symbol;Acc:MGI:1336181] | Kcnq3 | Enriched in dLGN | 1.999 | 2.631E-10 | 1087 |
| ENSMUSG00000059409 | protein phosphatase 2, regulatory subunit B', delta [Source:MGI Symbol;Acc:MGI:2388481] | Ppp2r5d | Enriched in dLGN | 1.997 | 2.27E-21 | 362.3 |
| ENSMUSG00000044835 | ankyrin repeat domain 45 [Source:MGI Symbol;Acc:MGI:1921094] | Ankrd45 | Enriched in dLGN | 1.995 | 3.246E-10 | 83.71 |

| | | | | | | |
|---|---|---|---|---|---|---|
| ENSMUSG00000029752 | asparagine synthetase [Source:MGI Symbol;Acc:MGI:1350929] | Asns | Enriched in dLGN | 1.989 | 0.000001402 | 106.2 |
| ENSMUSG00000043091 | tubulin, alpha 1C [Source:MGI Symbol;Acc:MGI:1095409] | Tuba1c | Enriched in dLGN | 1.988 | 0.003026 | 41.77 |
| ENSMUSG00000020326 | cyclin G1 [Source:MGI Symbol;Acc:MGI:102890] | Ccng1 | Enriched in dLGN | 1.982 | 1.602E-11 | 214.5 |
| ENSMUSG00000032503 | cyclic AMP-regulated phosphoprotein, 21 [Source:MGI Symbol;Acc:MGI:107562] | Arpp21 | Enriched in dLGN | 1.977 | 0.000001038 | 594.6 |
| ENSMUSG00000041697 | cytochrome c oxidase subunit 6A1 [Source:MGI Symbol;Acc:MGI:103099] | Cox6a1 | Enriched in dLGN | 1.973 | 1.66E-09 | 168 |
| ENSMUSG00000027360 | histidine decarboxylase [Source:MGI Symbol;Acc:MGI:96062] | Hdc | Enriched in dLGN | 1.973 | 0.0000209 | 44.93 |
| ENSMUSG00000069266 | H4 clustered histone 2 [Source:MGI Symbol;Acc:MGI:2448420] | H4c2 | Enriched in dLGN | 1.972 | 0.01469 | 5.433 |
| ENSMUSG00000026496 | poly (ADP-ribose) polymerase family, member 1 [Source:MGI Symbol;Acc:MGI:1340806] | Parp1 | Enriched in dLGN | 1.967 | 7.73E-22 | 378.8 |
| ENSMUSG00000019210 | ATPase, H+ transporting, lysosomal V1 subunit E1 [Source:MGI Symbol;Acc:MGI:894326] | Atp6v1e1 | Enriched in dLGN | 1.965 | 8.203E-09 | 270.4 |
| ENSMUSG00000028795 | coiled coil domain containing 28B [Source:MGI Symbol;Acc:MGI:1913514] | Ccdc28b | Enriched in dLGN | 1.964 | 8.2E-12 | 101 |
| ENSMUSG00000078566 | BCL2/adenovirus E1B interacting protein 3 [Source:MGI Symbol;Acc:MGI:109326] | Bnip3 | Enriched in dLGN | 1.964 | 0.000001915 | 111.7 |
| ENSMUSG00000035864 | synaptotagmin I [Source:MGI Symbol;Acc:MGI:99667] | Syt1 | Enriched in dLGN | 1.963 | 3.061E-14 | 3835 |
| ENSMUSG00000042961 | EGF-like, fibronectin type III and laminin G domains [Source:MGI Symbol;Acc:MGI:2146149] | Egflam | Enriched in dLGN | 1.962 | 1.895E-07 | 370.2 |
| ENSMUSG00000029066 | mitochondrial ribosomal protein L20 [Source:MGI Symbol;Acc:MGI:2137221] | Mrpl20 | Enriched in dLGN | 1.958 | 1.465E-09 | 48.19 |
| ENSMUSG00000020321 | malate dehydrogenase 1, NAD (soluble) [Source:MGI Symbol;Acc:MGI:97051] | Mdh1 | Enriched in dLGN | 1.957 | 0.000007594 | 617.5 |
| ENSMUSG00000038717 | ATP synthase, H+ transporting, mitochondrial F0 complex, subunit G [Source:MGI Symbol;Acc:MGI:1351597] | Atp5l | Enriched in dLGN | 1.956 | 1.494E-07 | 38.82 |
| ENSMUSG00000029918 | mitochondrial ribosomal protein S33 [Source:MGI Symbol;Acc:MGI:1338046] | Mrps33 | Enriched in dLGN | 1.955 | 0.000006533 | 26.36 |
| ENSMUSG00000056201 | cofilin 1, non-muscle [Source:MGI Symbol;Acc:MGI:101757] | Cfl1 | Enriched in dLGN | 1.954 | 5.926E-10 | 762.6 |
| ENSMUSG00000029153 | OCIA domain containing 2 [Source:MGI Symbol;Acc:MGI:1916377] | Ociad2 | Enriched in dLGN | 1.954 | 1.89E-09 | 70.58 |
| ENSMUSG00000039943 | phospholipase C, beta 4 [Source:MGI Symbol;Acc:MGI:107464] | Plcb4 | Enriched in dLGN | 1.949 | 4.414E-12 | 926.4 |
| ENSMUSG00000025393 | ATP synthase, H+ transporting mitochondrial F1 complex, beta subunit [Source:MGI Symbol;Acc:MGI:107801] | Atp5b | Enriched in dLGN | 1.949 | 5.12E-09 | 1784 |
| ENSMUSG00000063931 | peptidase D [Source:MGI Symbol;Acc:MGI:97542] | Pepd | Enriched in dLGN | 1.944 | 1.895E-07 | 102.6 |
| ENSMUSG00000030718 | protein phosphatase methylesterase 1 [Source:MGI Symbol;Acc:MGI:1919840] | Ppme1 | Enriched in dLGN | 1.941 | 7.032E-12 | 576.9 |
| ENSMUSG00000021384 | sushi domain containing 3 [Source:MGI Symbol;Acc:MGI:1913579] | Susd3 | Enriched in dLGN | 1.94 | 0.03698 | 69.76 |
| ENSMUSG00000021494 | DEAD box helicase 41 [Source:MGI Symbol;Acc:MGI:1920185] | Ddx41 | Enriched in dLGN | 1.938 | 4.046E-13 | 122.3 |
| ENSMUSG00000015932 | destrin [Source:MGI Symbol;Acc:MGI:1929270] | Dstn | Enriched in dLGN | 1.935 | 3.71E-09 | 300.9 |
| ENSMUSG00000024524 | guanine nucleotide binding protein, alpha stimulating, olfactory type [Source:MGI Symbol;Acc:MGI:95774] | Gnal | Enriched in dLGN | 1.929 | 3.332E-19 | 618.2 |
| ENSMUSG00000001089 | leucine zipper protein 1 [Source:MGI Symbol;Acc:MGI:107629] | Luzp1 | Enriched in dLGN | 1.927 | 2.823E-11 | 485.1 |
| ENSMUSG00000025081 | tudor domain containing 1 [Source:MGI Symbol;Acc:MGI:1933218] | Tdrd1 | Enriched in dLGN | 1.926 | 0.02213 | 5.686 |
| ENSMUSG00000071014 | NADH:ubiquinone oxidoreductase subunit B6 [Source:MGI Symbol;Acc:MGI:2684983] | Ndufb6 | Enriched in dLGN | 1.923 | 1.334E-07 | 50.89 |
| ENSMUSG00000049422 | coiled-coil-helix-coiled-coil-helix domain containing 10 [Source:MGI Symbol;Acc:MGI:2143558] | Chchd10 | Enriched in dLGN | 1.921 | 3.881E-09 | 224.5 |
| ENSMUSG00000059040 | enolase 1B, retrotransposed [Source:MGI Symbol;Acc:MGI:3648653] | Eno1b | Enriched in dLGN | 1.921 | 0.0002277 | 124.1 |
| ENSMUSG00000018362 | karyopherin (importin) alpha 2 [Source:MGI Symbol;Acc:MGI:103561] | Kpna2 | Enriched in dLGN | 1.916 | 0.00006519 | 67.24 |
| ENSMUSG00000040111 | GRAM domain containing 1B [Source:MGI Symbol;Acc:MGI:1925037] | Gramd1b | Enriched in dLGN | 1.915 | 2.768E-17 | 1123 |
| ENSMUSG00000027131 | ER membrane protein complex subunit 4 [Source:MGI Symbol;Acc:MGI:1915282] | Emc4 | Enriched in dLGN | 1.914 | 7.543E-09 | 81.85 |
| ENSMUSG00000028271 | general transcription factor IIB [Source:MGI Symbol;Acc:MGI:2385191] | Gtf2b | Enriched in dLGN | 1.911 | 0.000002496 | 36.1 |
| ENSMUSG00000002068 | cyclin E1 [Source:MGI Symbol;Acc:MGI:88316] | Ccne1 | Enriched in dLGN | 1.909 | 0.000003255 | 29.03 |
| ENSMUSG00000041231 | ubiquitin-like domain containing CTD phosphatase 1 [Source:MGI Symbol;Acc:MGI:1933105] | Ublcp1 | Enriched in dLGN | 1.905 | 5.78E-08 | 41.53 |
| ENSMUSG00000000088 | cytochrome c oxidase subunit 5A [Source:MGI Symbol;Acc:MGI:88474] | Cox5a | Enriched in dLGN | 1.904 | 1.524E-08 | 112.1 |
| ENSMUSG00000030591 | proteasome (prosome, macropain) 26S subunit, non-ATPase, 8 [Source:MGI Symbol;Acc:MGI:1888669] | Psmd8 | Enriched in dLGN | 1.903 | 2.896E-16 | 174.8 |
| ENSMUSG00000025428 | ATP synthase, H+ transporting, mitochondrial F1 complex, alpha subunit 1 [Source:MGI Symbol;Acc:MGI:88115] | Atp5a1 | Enriched in dLGN | 1.9 | 1.428E-09 | 1689 |
| ENSMUSG00000002379 | NADH:ubiquinone oxidoreductase subunit A11 [Source:MGI Symbol;Acc:MGI:1917125] | Ndufa11 | Enriched in dLGN | 1.899 | 0.0001381 | 75.68 |
| ENSMUSG00000075702 | selenoprotein M [Source:MGI Symbol;Acc:MGI:2149786] | Selenom | Enriched in dLGN | 1.893 | 1.338E-11 | 136.2 |
| ENSMUSG00000052504 | Eph receptor A3 [Source:MGI Symbol;Acc:MGI:99612] | Epha3 | Enriched in dLGN | 1.886 | 0.00003442 | 122.1 |
| ENSMUSG00000031391 | L1 cell adhesion molecule [Source:MGI Symbol;Acc:MGI:96721] | L1cam | Enriched in dLGN | 1.879 | 0.000003494 | 803.2 |
| ENSMUSG00000017801 | MAX-like protein X [Source:MGI Symbol;Acc:MGI:108398] | Mlx | Enriched in dLGN | 1.879 | 0.00002437 | 24.55 |
| ENSMUSG00000023861 | mitochondrial pyruvate carrier 1 [Source:MGI Symbol;Acc:MGI:1915240] | Mpc1 | Enriched in dLGN | 1.877 | 1.495E-07 | 73.22 |
| ENSMUSG00000005161 | peroxiredoxin 2 [Source:MGI Symbol;Acc:MGI:109486] | Prdx2 | Enriched in dLGN | 1.874 | 1.596E-08 | 321.2 |
| ENSMUSG00000042298 | tetratricopeptide repeat domain 19 [Source:MGI Symbol;Acc:MGI:1920045] | Ttc19 | Enriched in dLGN | 1.873 | 1.497E-12 | 462.6 |
| ENSMUSG00000042660 | WD repeat domain 55 [Source:MGI Symbol;Acc:MGI:1915186] | Wdr55 | Enriched in dLGN | 1.872 | 1.142E-07 | 26.69 |
| ENSMUSG00000022376 | adenylate cyclase 8 [Source:MGI Symbol;Acc:MGI:1341110] | Adcy8 | Enriched in dLGN | 1.871 | 3.332E-08 | 297.1 |
| ENSMUSG00000070803 | Cbp/p300-interacting transactivator, with Glu/Asp-rich carboxy-terminal domain, 4 [Source:MGI Symbol;Acc:MGI:1861694] | Cited4 | Enriched in dLGN | 1.864 | 0.004667 | 9.191 |
| ENSMUSG00000042353 | Fras1 related extracellular matrix protein 3 [Source:MGI Symbol;Acc:MGI:2685641] | Frem3 | Enriched in dLGN | 1.857 | 0.000005606 | 57.69 |
| ENSMUSG00000049353 | retinal degeneration 3 [Source:MGI Symbol;Acc:MGI:1921273] | Rd3 | Enriched in dLGN | 1.857 | 0.03462 | 236.2 |
| ENSMUSG00000062825 | actin, gamma, cytoplasmic 1 [Source:MGI Symbol;Acc:MGI:87906] | Actg1 | Enriched in dLGN | 1.854 | 0.00002466 | 407 |
| ENSMUSG00000060093 | H4 clustered histone 1 [Source:MGI Symbol;Acc:MGI:2448419] | H4c1 | Enriched in dLGN | 1.85 | 0.01249 | 10.73 |
| ENSMUSG00000022269 | membrane associated ring-CH-type finger 11 [Source:MGI Symbol;Acc:MGI:3608327] | Marchf11 | Enriched in dLGN | 1.84 | 0.0006901 | 40.63 |
| ENSMUSG00000029247 | phosphoribosylaminoimidazole carboxylase, phosphoribosylaminoribosylaminoimidazole, succinocarboxamide synthetase [Source:MGI Symbol;Acc:MGI:1914304] | Paics | Enriched in dLGN | 1.836 | 0.00000063 | 114.1 |
| ENSMUSG00000010914 | pyruvate dehydrogenase complex, component X [Source:MGI Symbol;Acc:MGI:1351627] | Pdhx | Enriched in dLGN | 1.834 | 5.095E-14 | 152.7 |
| ENSMUSG00000033918 | presenilin associated, rhomboid-like [Source:MGI Symbol;Acc:MGI:1277152] | Parl | Enriched in dLGN | 1.834 | 1.771E-12 | 65.89 |
| ENSMUSG00000059734 | NADH:ubiquinone oxidoreductase core subunit S8 [Source:MGI Symbol;Acc:MGI:2385079] | Ndufs8 | Enriched in dLGN | 1.834 | 6.576E-07 | 73.63 |
| ENSMUSG00000032330 | cytochrome c oxidase subunit 7A2 [Source:MGI Symbol;Acc:MGI:1316715] | Cox7a2 | Enriched in dLGN | 1.825 | 1.168E-09 | 76.71 |
| ENSMUSG00000060402 | carbohydrate sulfotransferase 8 [Source:MGI Symbol;Acc:MGI:1916197] | Chst8 | Enriched in dLGN | 1.822 | 3.912E-07 | 67.64 |
| ENSMUSG00000006057 | ATP synthase, H+ transporting, mitochondrial F0 complex, subunit C1 (subunit 9) [Source:MGI Symbol;Acc:MGI:107653] | Atp5g1 | Enriched in dLGN | 1.82 | 0.000000234 | 76.53 |
| ENSMUSG00000028648 | NADH:ubiquinone oxidoreductase core subunit S5 [Source:MGI Symbol;Acc:MGI:1890889] | Ndufs5 | Enriched in dLGN | 1.819 | 3.003E-07 | 101.9 |
| ENSMUSG00000030103 | basic helix-loop-helix family, member e40 [Source:MGI Symbol;Acc:MGI:1097714] | Bhlhe40 | Enriched in dLGN | 1.817 | 2.509E-08 | 95.28 |
| ENSMUSG00000030303 | fatty acyl CoA reductase 2 [Source:MGI Symbol;Acc:MGI:2687035] | Far2 | Enriched in dLGN | 1.816 | 0.0001255 | 96.51 |
| ENSMUSG00000031647 | microfibrillar-associated protein 3-like [Source:MGI Symbol;Acc:MGI:1918556] | Mfap3l | Enriched in dLGN | 1.814 | 3.422E-10 | 250.9 |
| ENSMUSG00000021209 | protein phosphatase 4, regulatory subunit 4 [Source:MGI Symbol;Acc:MGI:1921771] | Ppp4r4 | Enriched in dLGN | 1.812 | 4.737E-11 | 139.4 |
| ENSMUSG00000031309 | ribosomal protein S6 kinase polypeptide 3 [Source:MGI Symbol;Acc:MGI:104557] | Rps6ka3 | Enriched in dLGN | 1.812 | 2.647E-09 | 168.6 |
| ENSMUSG00000018965 | tyrosine 3-monooxygenase/tryptophan 5-monooxygenase activation protein, eta polypeptide [Source:MGI Symbol;Acc:MGI:109194] | Ywhah | Enriched in dLGN | 1.812 | 1.142E-07 | 3239 |
| ENSMUSG00000036934 | RIKEN cDNA 4921524J17 gene [Source:MGI Symbol;Acc:MGI:1913964] | 4921524J17Rik | Enriched in dLGN | 1.806 | 0.000002845 | 42.54 |
| ENSMUSG00000029185 | family with sequence similarity 114, member A1 [Source:MGI Symbol;Acc:MGI:1915553] | Fam114a1 | Enriched in dLGN | 1.806 | 0.00004175 | 29.67 |
| ENSMUSG00000016833 | mitochondrial ribosomal protein S18C [Source:MGI Symbol;Acc:MGI:1915985] | Mrps18c | Enriched in dLGN | 1.806 | 0.00299 | 8.671 |
| ENSMUSG00000027827 | potassium voltage-gated channel, shaker-related subfamily, beta member 1 [Source:MGI Symbol;Acc:MGI:109155] | Kcnab1 | Enriched in dLGN | 1.803 | 0.000004519 | 138 |
| ENSMUSG00000005510 | NADH:ubiquinone oxidoreductase core subunit S3 [Source:MGI Symbol;Acc:MGI:1915599] | Ndufs3 | Enriched in dLGN | 1.802 | 1.05E-11 | 101.1 |
| ENSMUSG00000011752 | phosphoglycerate mutase 1 [Source:MGI Symbol;Acc:MGI:97552] | Pgam1 | Enriched in dLGN | 1.802 | 4.73E-08 | 217.3 |
| ENSMUSG00000015656 | heat shock protein 8 [Source:MGI Symbol;Acc:MGI:105384] | Hspa8 | Enriched in dLGN | 1.802 | 0.0001217 | 1562 |
| ENSMUSG00000020225 | transmembrane BAX inhibitor motif containing 4 [Source:MGI Symbol;Acc:MGI:1915462] | Tmbim4 | Enriched in dLGN | 1.797 | 0.00002272 | 25.14 |
| ENSMUSG00000034880 | mitochondrial ribosomal protein L34 [Source:MGI Symbol;Acc:MGI:2137227] | Mrpl34 | Enriched in dLGN | 1.791 | 1.739E-07 | 34.19 |
| ENSMUSG00000024841 | eukaryotic translation initiation factor 1A domain containing [Source:MGI Symbol;Acc:MGI:1917110] | Eif1ad | Enriched in dLGN | 1.786 | 3.569E-10 | 57.89 |
| ENSMUSG00000041193 | phospholipase A2, group V [Source:MGI Symbol;Acc:MGI:101899] | Pla2g5 | Enriched in dLGN | 1.785 | 0.02886 | 65.41 |
| ENSMUSG00000045427 | heterogeneous nuclear ribonucleoprotein H2 [Source:MGI Symbol;Acc:MGI:1201779] | Hnrnph2 | Enriched in dLGN | 1.777 | 3.614E-09 | 111 |
| ENSMUSG00000005233 | SPC25, NDC80 kinetochore complex component, homolog (S. cerevisiae) [Source:MGI Symbol;Acc:MGI:1913692] | Spc25 | Enriched in dLGN | 1.777 | 0.03079 | 60.05 |
| ENSMUSG00000030061 | ubiquitin-like modifier activating enzyme 3 [Source:MGI Symbol;Acc:MGI:1341217] | Uba3 | Enriched in dLGN | 1.776 | 1.092E-08 | 78.39 |
| ENSMUSG00000032279 | isocitrate dehydrogenase 3 (NAD+) alpha [Source:MGI Symbol;Acc:MGI:1915084] | Idh3a | Enriched in dLGN | 1.77 | 0.00007876 | 302 |
| ENSMUSG00000052302 | TBC1 domain family, member 30 [Source:MGI Symbol;Acc:MGI:1921944] | Tbc1d30 | Enriched in dLGN | 1.764 | 1.748E-18 | 314.2 |
| ENSMUSG00000023828 | solute carrier family 22 (organic cation transporter), member 3 [Source:MGI Symbol;Acc:MGI:1333817] | Slc22a3 | Enriched in dLGN | 1.762 | 3.857E-07 | 27.45 |
| ENSMUSG00000039809 | gamma-aminobutyric acid (GABA) B receptor, 2 [Source:MGI Symbol;Acc:MGI:2386030] | Gabbr2 | Enriched in dLGN | 1.759 | 1.245E-16 | 1664 |
| ENSMUSG00000079108 | signal recognition particle 54C [Source:MGI Symbol;Acc:MGI:3714359] | Srp54c | Enriched in dLGN | 1.758 | 0.003337 | 21.13 |
| ENSMUSG00000035372 | RIKEN cDNA 1810055G02 gene [Source:MGI Symbol;Acc:MGI:1919306] | 1810055G02Rik | Enriched in dLGN | 1.757 | 0.000006132 | 34.57 |
| ENSMUSG00000022707 | glucan (1,4-alpha-), branching enzyme 1 [Source:MGI Symbol;Acc:MGI:1921435] | Gbe1 | Enriched in dLGN | 1.756 | 0.00001534 | 124.2 |
| ENSMUSG00000035849 | keratin 222 [Source:MGI Symbol;Acc:MGI:2442728] | Krt222 | Enriched in dLGN | 1.755 | 7.048E-08 | 127 |
| ENSMUSG00000040620 | DEAH (Asp-Glu-Ala-His) box polypeptide 33 [Source:MGI Symbol;Acc:MGI:2445102] | Dhx33 | Enriched in dLGN | 1.754 | 1.087E-10 | 72.8 |
| ENSMUSG00000061482 | H4 clustered histone 4 [Source:MGI Symbol;Acc:MGI:2448423] | H4c4 | Enriched in dLGN | 1.751 | 0.00006628 | 85.46 |
| ENSMUSG00000057666 | glyceraldehyde-3-phosphate dehydrogenase [Source:MGI Symbol;Acc:MGI:95640] | Gapdh | Enriched in dLGN | 1.751 | 0.0006123 | 93.82 |
| ENSMUSG00000027108 | Obg-like ATPase 1 [Source:MGI Symbol;Acc:MGI:1914309] | Ola1 | Enriched in dLGN | 1.751 | 0.01527 | 454.9 |
| ENSMUSG00000007564 | protein phosphatase 2, regulatory subunit A, alpha [Source:MGI Symbol;Acc:MGI:1926334] | Ppp2r1a | Enriched in dLGN | 1.75 | 5.139E-08 | 634.9 |
| ENSMUSG00000025477 | inositol polyphosphate-5-phosphatase A [Source:MGI Symbol;Acc:MGI:2686961] | Inpp5a | Enriched in dLGN | 1.747 | 1.963E-11 | 299.2 |
| ENSMUSG00000020029 | nudix (nucleoside diphosphate linked moiety X)-type motif 4 [Source:MGI Symbol;Acc:MGI:1918457] | Nudt4 | Enriched in dLGN | 1.746 | 4.903E-11 | 752.5 |
| ENSMUSG00000057766 | ankyrin repeat domain 29 [Source:MGI Symbol;Acc:MGI:2687055] | Ankrd29 | Enriched in dLGN | 1.741 | 0.000000572 | 58.3 |
| ENSMUSG00000003072 | ATP synthase, H+ transporting, mitochondrial F1 complex, delta subunit [Source:MGI Symbol;Acc:MGI:1913293] | Atp5d | Enriched in dLGN | 1.738 | 1.206E-11 | 259.8 |
| ENSMUSG00000020589 | CYFIP related Rac1 interactor A [Source:MGI Symbol;Acc:MGI:1261783] | Cyria | Enriched in dLGN | 1.738 | 1.254E-11 | 283.7 |
| ENSMUSG00000036751 | cytochrome c oxidase, subunit 6B1 [Source:MGI Symbol;Acc:MGI:107460] | Cox6b1 | Enriched in dLGN | 1.726 | 1.541E-09 | 143.6 |
| ENSMUSG00000019877 | serine incorporator 1 [Source:MGI Symbol;Acc:MGI:1926228] | Serinc1 | Enriched in dLGN | 1.725 | 5.805E-07 | 662.3 |
| ENSMUSG00000031007 | ATPase, H+ transporting, lysosomal accessory protein 2 [Source:MGI Symbol;Acc:MGI:1917745] | Atp6ap2 | Enriched in dLGN | 1.722 | 1.304E-07 | 242.6 |
| ENSMUSG00000027965 | olfactomedin 3 [Source:MGI Symbol;Acc:MGI:2387329] | Olfm3 | Enriched in dLGN | 1.715 | 0.000005536 | 175.9 |
| ENSMUSG00000079480 | peptidyl-prolyl cis/trans isomerase, NIMA-interacting, 4 (parvulin) [Source:MGI Symbol;Acc:MGI:1916963] | Pin4 | Enriched in dLGN | 1.715 | 0.0002925 | 13.6 |
| ENSMUSG00000033917 | glycerophosphodiester phosphodiesterase 1 [Source:MGI Symbol;Acc:MGI:1891827] | Gde1 | Enriched in dLGN | 1.713 | 3.489E-15 | 234.5 |
| ENSMUSG00000031617 | transmembrane protein 184C [Source:MGI Symbol;Acc:MGI:2384562] | Tmem184c | Enriched in dLGN | 1.711 | 1.339E-09 | 115.9 |
| ENSMUSG00000047126 | clathrin, heavy polypeptide (Hc) [Source:MGI Symbol;Acc:MGI:2388633] | Cltc | Enriched in dLGN | 1.711 | 6.061E-09 | 1529 |
| ENSMUSG00000060429 | syntrophin, basic 1 [Source:MGI Symbol;Acc:MGI:101781] | Sntb1 | Enriched in dLGN | 1.708 | 6.838E-09 | 115.5 |
| ENSMUSG00000032366 | tropomyosin 1, alpha [Source:MGI Symbol;Acc:MGI:98809] | Tpm1 | Enriched in dLGN | 1.704 | 6.131E-29 | 654.7 |
| ENSMUSG00000022620 | arylsulfatase A [Source:MGI Symbol;Acc:MGI:88077] | Arsa | Enriched in dLGN | 1.701 | 7.187E-08 | 62.33 |
| ENSMUSG00000027603 | gamma-glutamyltransferase 7 [Source:MGI Symbol;Acc:MGI:1913385] | Ggt7 | Enriched in dLGN | 1.699 | 1.77E-10 | 194.7 |
| ENSMUSG00000058927 | predicted gene 10053 [Source:MGI Symbol;Acc:MGI:3704493] | Gm10053 | Enriched in dLGN | 1.699 | 0.002831 | 16.38 |
| ENSMUSG00000055782 | ATP-binding cassette, sub-family D (ALD), member 2 [Source:MGI Symbol;Acc:MGI:1349467] | Abcd2 | Enriched in dLGN | 1.69 | 3.059E-08 | 85.55 |
| ENSMUSG00000028410 | DnaJ heat shock protein family (Hsp40) member A1 [Source:MGI Symbol;Acc:MGI:1270129] | Dnaja1 | Enriched in dLGN | 1.69 | 0.00007336 | 251.8 |
| ENSMUSG00000038244 | microtubule associated monooxygenase, calponin and LIM domain containing 2 [Source:MGI Symbol;Acc:MGI:2444947] | Mical2 | Enriched in dLGN | 1.686 | 1.363E-07 | 255.1 |
| ENSMUSG00000031668 | eukaryotic translation initiation factor 2 alpha kinase 3 [Source:MGI Symbol;Acc:MGI:1341830] | Eif2ak3 | Enriched in dLGN | 1.68 | 4.012E-13 | 198.6 |
| ENSMUSG00000021373 | CAP, adenylate cyclase-associated protein, 2 (yeast) [Source:MGI Symbol;Acc:MGI:1914502] | Cap2 | Enriched in dLGN | 1.68 | 4.325E-09 | 147 |
| ENSMUSG00000020458 | reticulon 4 [Source:MGI Symbol;Acc:MGI:1915835] | Rtn4 | Enriched in dLGN | 1.677 | 1.525E-07 | 1440 |
| ENSMUSG00000076432 | tyrosine 3-monooxygenase/tryptophan 5-monooxygenase activation protein theta [Source:MGI Symbol;Acc:MGI:891963] | Ywhaq | Enriched in dLGN | 1.675 | 4.389E-07 | 236.1 |
| ENSMUSG00000018585 | antioxidant 1 copper chaperone [Source:MGI Symbol;Acc:MGI:1333855] | Atox1 | Enriched in dLGN | 1.673 | 0.0002334 | 25.24 |
| ENSMUSG00000032398 | small nuclear RNA activating complex, polypeptide 5 [Source:MGI Symbol;Acc:MGI:1914282] | Snapc5 | Enriched in dLGN | 1.673 | 0.0009098 | 21.39 |
| ENSMUSG00000029617 | CCZ1 vacuolar protein trafficking and biogenesis associated [Source:MGI Symbol;Acc:MGI:2141070] | Ccz1 | Enriched in dLGN | 1.669 | 2.028E-13 | 122.9 |
| ENSMUSG00000027346 | glycerophosphocholine phosphodiesterase 1 [Source:MGI Symbol;Acc:MGI:104898] | Gpcpd1 | Enriched in dLGN | 1.668 | 3.935E-10 | 161 |
| ENSMUSG00000019179 | malate dehydrogenase 2, NAD (mitochondrial) [Source:MGI Symbol;Acc:MGI:97050] | Mdh2 | Enriched in dLGN | 1.668 | 0.000002656 | 626.3 |

| | | | | | | |
|---|---|---|---|---|---|---|
| ENSMUSG00000000056 | nuclear prelamin A recognition factor [Source:MGI Symbol;Acc:MGI:1914858] | Narf | Enriched in dLGN | 1.666 | 4.238E-19 | 273.8 |
| ENSMUSG00000050830 | von Willebrand factor C domain containing 2 [Source:MGI Symbol;Acc:MGI:2442987] | Vwc2 | Enriched in dLGN | 1.665 | 0.000001557 | 127 |
| ENSMUSG00000049775 | thymosin, beta 4, X chromosome [Source:MGI Symbol;Acc:MGI:99510] | Tmsb4x | Enriched in dLGN | 1.657 | 3.592E-13 | 317.2 |
| ENSMUSG00000002102 | proteasome (prosome, macropain) 26S subunit, ATPase 3 [Source:MGI Symbol;Acc:MGI:1098754] | Psmc3 | Enriched in dLGN | 1.656 | 1.067E-09 | 304.4 |
| ENSMUSG00000034932 | mitochondrial ribosomal protein L54 [Source:MGI Symbol;Acc:MGI:1913297] | Mrpl54 | Enriched in dLGN | 1.656 | 0.000002217 | 34.59 |
| ENSMUSG00000024953 | peroxiredoxin 5 [Source:MGI Symbol;Acc:MGI:1859821] | Prdx5 | Enriched in dLGN | 1.654 | 3.716E-11 | 344.2 |
| ENSMUSG00000030695 | aldolase A, fructose-bisphosphate [Source:MGI Symbol;Acc:MGI:87994] | Aldoa | Enriched in dLGN | 1.654 | 1.137E-07 | 1713 |
| ENSMUSG00000034566 | ATP synthase, H+ transporting, mitochondrial F0 complex, subunit D [Source:MGI Symbol;Acc:MGI:1918929] | Atp5h | Enriched in dLGN | 1.65 | 3.325E-08 | 216.1 |
| ENSMUSG00000028444 | ciliary neurotrophic factor receptor [Source:MGI Symbol;Acc:MGI:99605] | Cntfr | Enriched in dLGN | 1.648 | 4.858E-10 | 219.4 |
| ENSMUSG00000024777 | protein phosphatase 2, regulatory subunit B', beta [Source:MGI Symbol;Acc:MGI:2388480] | Ppp2r5b | Enriched in dLGN | 1.646 | 1.934E-19 | 428 |
| ENSMUSG00000029363 | replication factor C (activator 1) 5 [Source:MGI Symbol;Acc:MGI:1919401] | Rfc5 | Enriched in dLGN | 1.646 | 0.002135 | 15.49 |
| ENSMUSG00000032038 | ST3 beta-galactoside alpha-2,3-sialyltransferase 4 [Source:MGI Symbol;Acc:MGI:1316743] | St3gal4 | Enriched in dLGN | 1.645 | 1.808E-07 | 102 |
| ENSMUSG00000066441 | retinol dehydrogenase 11 [Source:MGI Symbol;Acc:MGI:102581] | Rdh11 | Enriched in dLGN | 1.638 | 2.048E-11 | 82.77 |
| ENSMUSG00000000563 | ATP synthase peripheral stalk-membrane subunit b [Source:MGI Symbol;Acc:MGI:1100495] | Atp5pb | Enriched in dLGN | 1.638 | 6.419E-09 | 236 |
| ENSMUSG00000005683 | citrate synthase [Source:MGI Symbol;Acc:MGI:88529] | Cs | Enriched in dLGN | 1.637 | 2.631E-10 | 414 |
| ENSMUSG00000046798 | claudin 12 [Source:MGI Symbol;Acc:MGI:1929288] | Cldn12 | Enriched in dLGN | 1.636 | 5.089E-13 | 124.6 |
| ENSMUSG00000046442 | protein phosphatase 1E (PP2C domain containing) [Source:MGI Symbol;Acc:MGI:2444096] | Ppm1e | Enriched in dLGN | 1.636 | 2E-11 | 844.2 |
| ENSMUSG00000027203 | deoxyuridine triphosphatase [Source:MGI Symbol;Acc:MGI:1346051] | Dut | Enriched in dLGN | 1.635 | 5.145E-11 | 92.41 |
| ENSMUSG00000019849 | prolyl endopeptidase [Source:MGI Symbol;Acc:MGI:1270863] | Prep | Enriched in dLGN | 1.63 | 1.662E-08 | 207.5 |
| ENSMUSG00000036989 | tripartite motif-containing 3 [Source:MGI Symbol;Acc:MGI:1860040] | Trim3 | Enriched in dLGN | 1.628 | 6.439E-09 | 276 |
| ENSMUSG00000061518 | cytochrome c oxidase subunit 5B [Source:MGI Symbol;Acc:MGI:88475] | Cox5b | Enriched in dLGN | 1.627 | 2.849E-08 | 455.8 |
| ENSMUSG00000034891 | synuclein, beta [Source:MGI Symbol;Acc:MGI:1889011] | Sncb | Enriched in dLGN | 1.623 | 0.000000235 | 939.2 |
| ENSMUSG00000040907 | ATPase, Na+/K+ transporting, alpha 3 polypeptide [Source:MGI Symbol;Acc:MGI:88107] | Atp1a3 | Enriched in dLGN | 1.619 | 1.907E-11 | 7574 |
| ENSMUSG00000035504 | receptor accessory protein 6 [Source:MGI Symbol;Acc:MGI:1917585] | Reep6 | Enriched in dLGN | 1.618 | 0.02808 | 1054 |
| ENSMUSG00000027341 | transmembrane protein 230 [Source:MGI Symbol;Acc:MGI:1917862] | Tmem230 | Enriched in dLGN | 1.612 | 0.00001598 | 38.18 |
| ENSMUSG00000071648 | rod outer segment membrane protein 1 [Source:MGI Symbol;Acc:MGI:97998] | Rom1 | Enriched in dLGN | 1.608 | 0.02537 | 555.4 |
| ENSMUSG00000031672 | glutamatic-oxaloacetic transaminase 2, mitochondrial [Source:MGI Symbol;Acc:MGI:95792] | Got2 | Enriched in dLGN | 1.607 | 0.000001847 | 253.7 |
| ENSMUSG00000032267 | ubiquitin specific peptidase 28 [Source:MGI Symbol;Acc:MGI:2442293] | Usp28 | Enriched in dLGN | 1.606 | 9.446E-07 | 93.89 |
| ENSMUSG00000062981 | mitochondrial ribosomal protein L42 [Source:MGI Symbol;Acc:MGI:1333774] | Mrpl42 | Enriched in dLGN | 1.605 | 0.000298 | 43.37 |
| ENSMUSG00000034088 | high density lipoprotein (HDL) binding protein [Source:MGI Symbol;Acc:MGI:99256] | Hdlbp | Enriched in dLGN | 1.603 | 8.073E-15 | 727.2 |
| ENSMUSG00000026904 | solute carrier family 4, sodium bicarbonate cotransporter-like, member 10 [Source:MGI Symbol;Acc:MGI:2150150] | Slc4a10 | Enriched in dLGN | 1.603 | 9.394E-09 | 601.1 |
| ENSMUSG00000031137 | fibroblast growth factor 13 [Source:MGI Symbol;Acc:MGI:109178] | Fgf13 | Enriched in dLGN | 1.603 | 4.561E-07 | 419.6 |
| ENSMUSG00000041729 | coronin, actin binding protein, 2B [Source:MGI Symbol;Acc:MGI:2444283] | Coro2b | Enriched in dLGN | 1.598 | 1.013E-14 | 607.5 |
| ENSMUSG00000091636 | A kinase (PRKA) anchor inhibitor 1 [Source:MGI Symbol;Acc:MGI:2444600] | Akain1 | Enriched in dLGN | 1.597 | 0.002392 | 14.32 |
| ENSMUSG00000032959 | phosphatidylethanolamine binding protein 1 [Source:MGI Symbol;Acc:MGI:1344408] | Pebp1 | Enriched in dLGN | 1.596 | 0.000008762 | 164.1 |
| ENSMUSG00000021771 | voltage-dependent anion channel 2 [Source:MGI Symbol;Acc:MGI:106915] | Vdac2 | Enriched in dLGN | 1.595 | 0.00003217 | 243.1 |
| ENSMUSG00000034488 | EGF-like repeats and discoidin I-like domains 3 [Source:MGI Symbol;Acc:MGI:1329025] | Edil3 | Enriched in dLGN | 1.594 | 0.00002382 | 564.8 |
| ENSMUSG00000014243 | zinc finger SWIM-type containing 7 [Source:MGI Symbol;Acc:MGI:1916997] | Zswim7 | Enriched in dLGN | 1.593 | 0.0006256 | 26.22 |
| ENSMUSG00000008822 | acylphosphatase 1, erythrocyte (common) type [Source:MGI Symbol;Acc:MGI:1913454] | Acyp1 | Enriched in dLGN | 1.592 | 0.0002228 | 16.49 |
| ENSMUSG00000049612 | oligodendrocyte myelin glycoprotein [Source:MGI Symbol;Acc:MGI:106586] | Omg | Enriched in dLGN | 1.59 | 0.00003603 | 36.01 |
| ENSMUSG00000034171 | fatty acid amide hydrolase [Source:MGI Symbol;Acc:MGI:109609] | Faah | Enriched in dLGN | 1.589 | 1.736E-15 | 134 |
| ENSMUSG00000045624 | ESF1 nucleolar pre-rRNA processing protein homolog [Source:MGI Symbol;Acc:MGI:1913830] | Esf1 | Enriched in dLGN | 1.583 | 2.447E-08 | 125.5 |
| ENSMUSG00000059343 | aldolase 1 A, retrogene 1 [Source:MGI Symbol;Acc:MGI:2447811] | Aldoart1 | Enriched in dLGN | 1.583 | 0.0009294 | 19.45 |
| ENSMUSG00000020309 | ChaC, cation transport regulator 2 [Source:MGI Symbol;Acc:MGI:1915294] | Chac2 | Enriched in dLGN | 1.579 | 0.0007281 | 18.28 |
| ENSMUSG00000038115 | anoctamin 2 [Source:MGI Symbol;Acc:MGI:2387214] | Ano2 | Enriched in dLGN | 1.579 | 0.03198 | 281.6 |
| ENSMUSG00000020664 | dihydrolipoamide dehydrogenase [Source:MGI Symbol;Acc:MGI:107450] | Dld | Enriched in dLGN | 1.577 | 3.922E-07 | 258.4 |
| ENSMUSG00000027134 | lysophosphatidylcholine acyltransferase 4 [Source:MGI Symbol;Acc:MGI:2138993] | Lpcat4 | Enriched in dLGN | 1.575 | 2.504E-13 | 130.5 |
| ENSMUSG00000026032 | NADH:ubiquinone oxidoreductase subunit B3 [Source:MGI Symbol;Acc:MGI:1913745] | Ndufb3 | Enriched in dLGN | 1.571 | 4.937E-07 | 49.01 |
| ENSMUSG00000002728 | N(alpha)-acetyltransferase 20, NatB catalytic subunit [Source:MGI Symbol;Acc:MGI:1915127] | Naa20 | Enriched in dLGN | 1.57 | 0.00000394 | 46.04 |
| ENSMUSG00000036932 | apoptosis-inducing factor, mitochondrion-associated 1 [Source:MGI Symbol;Acc:MGI:1349419] | Aifm1 | Enriched in dLGN | 1.567 | 0.000006137 | 96.3 |
| ENSMUSG00000031996 | amyloid beta (A4) precursor-like protein 2 [Source:MGI Symbol;Acc:MGI:88047] | Aplp2 | Enriched in dLGN | 1.562 | 8.238E-08 | 2704 |
| ENSMUSG00000030423 | processing of precursor 4, ribonuclease P/MRP family, (S. cerevisiae) [Source:MGI Symbol;Acc:MGI:1913411] | Pop4 | Enriched in dLGN | 1.562 | 0.0001559 | 40.41 |
| ENSMUSG00000055409 | NEL-like 1 [Source:MGI Symbol;Acc:MGI:2443902] | Nell1 | Enriched in dLGN | 1.562 | 0.0002875 | 543.3 |
| ENSMUSG00000002280 | cytosolic iron-sulfur assembly component 3 [Source:MGI Symbol;Acc:MGI:1914813] | Ciao3 | Enriched in dLGN | 1.558 | 0.000006249 | 33.9 |
| ENSMUSG00000032459 | mitochondrial ribosomal protein S22 [Source:MGI Symbol;Acc:MGI:1928137] | Mrps22 | Enriched in dLGN | 1.558 | 0.002456 | 35.07 |
| ENSMUSG00000026527 | regulator of G protein signaling 7 [Source:MGI Symbol;Acc:MGI:1346089] | Rgs7 | Enriched in dLGN | 1.557 | 0.0003738 | 427.5 |
| ENSMUSG00000029860 | zyxin [Source:MGI Symbol;Acc:MGI:103072] | Zyx | Enriched in dLGN | 1.553 | 4.931E-09 | 118.6 |
| ENSMUSG00000063904 | dipeptidylpeptidase 3 [Source:MGI Symbol;Acc:MGI:1922471] | Dpp3 | Enriched in dLGN | 1.551 | 5.139E-11 | 154.1 |
| ENSMUSG00000023175 | basigin [Source:MGI Symbol;Acc:MGI:88208] | Bsg | Enriched in dLGN | 1.551 | 4.832E-10 | 872.5 |
| ENSMUSG00000048616 | noggin [Source:MGI Symbol;Acc:MGI:104327] | Nog | Enriched in dLGN | 1.549 | 0.00002633 | 53.91 |
| ENSMUSG00000020869 | leucine rich repeat containing 59 [Source:MGI Symbol;Acc:MGI:2138133] | Lrrc59 | Enriched in dLGN | 1.546 | 6.694E-12 | 321.9 |
| ENSMUSG00000024387 | casein kinase 2, beta polypeptide [Source:MGI Symbol;Acc:MGI:88548] | Csnk2b | Enriched in dLGN | 1.546 | 0.0001155 | 97.86 |
| ENSMUSG00000049493 | plastin 1 (I-isoform) [Source:MGI Symbol;Acc:MGI:104809] | Pls1 | Enriched in dLGN | 1.542 | 0.007935 | 15.52 |
| ENSMUSG00000023572 | cyclin D-type binding-protein 1 [Source:MGI Symbol;Acc:MGI:109595] | Ccndbp1 | Enriched in dLGN | 1.54 | 3.954E-08 | 105.3 |
| ENSMUSG00000022257 | lysosomal-associated protein transmembrane 4B [Source:MGI Symbol;Acc:MGI:1890494] | Laptm4b | Enriched in dLGN | 1.537 | 1.095E-08 | 165.8 |
| ENSMUSG00000031357 | synapse associated protein 1 [Source:MGI Symbol;Acc:MGI:1914293] | Syap1 | Enriched in dLGN | 1.535 | 0.000002534 | 90.22 |
| ENSMUSG00000025968 | NADH:ubiquinone oxidoreductase core subunit S1 [Source:MGI Symbol;Acc:MGI:2443241] | Ndufs1 | Enriched in dLGN | 1.534 | 9.251E-09 | 337.9 |
| ENSMUSG00000017167 | contactin associated protein-like 1 [Source:MGI Symbol;Acc:MGI:1858201] | Cntnap1 | Enriched in dLGN | 1.534 | 0.00001907 | 622.8 |
| ENSMUSG00000019951 | UHRF1 (ICBP90) binding protein 1-like [Source:MGI Symbol;Acc:MGI:2442888] | Uhrf1bp1l | Enriched in dLGN | 1.533 | 3.201E-21 | 276.6 |
| ENSMUSG00000044117 | bMERB domain containing 1 [Source:MGI Symbol;Acc:MGI:1914504] | Bmerb1 | Enriched in dLGN | 1.53 | 4.672E-12 | 457.3 |
| ENSMUSG00000044894 | ubiquinol-cytochrome c reductase, complex III subunit VII [Source:MGI Symbol;Acc:MGI:107807] | Uqcrq | Enriched in dLGN | 1.53 | 3.832E-07 | 236.5 |
| ENSMUSG00000021215 | neuroepithelial cell transforming gene 1 [Source:MGI Symbol;Acc:MGI:1927138] | Net1 | Enriched in dLGN | 1.529 | 0.000001233 | 50.15 |
| ENSMUSG00000025545 | citrate lyase beta like [Source:MGI Symbol;Acc:MGI:1916884] | Clybl | Enriched in dLGN | 1.528 | 0.0004144 | 35.95 |
| ENSMUSG00000031633 | solute carrier family 25 (mitochondrial carrier, adenine nucleotide translocator), member 4 [Source:MGI Symbol;Acc:MGI:1353495] | Slc25a4 | Enriched in dLGN | 1.527 | 5.414E-07 | 1489 |
| ENSMUSG00000029777 | glycyl-tRNA synthetase [Source:MGI Symbol;Acc:MGI:1309461] | Gars | Enriched in dLGN | 1.523 | 1.291E-11 | 453.4 |
| ENSMUSG00000029388 | eukaryotic translation initiation factor 2B, subunit 1 (alpha) [Source:MGI Symbol;Acc:MGI:2384802] | Eif2b1 | Enriched in dLGN | 1.519 | 0.000002373 | 40.06 |
| ENSMUSG00000015668 | PDZ domain containing 11 [Source:MGI Symbol;Acc:MGI:1919871] | Pdzd11 | Enriched in dLGN | 1.518 | 0.00003573 | 27.95 |
| ENSMUSG00000000552 | zinc finger protein 385A [Source:MGI Symbol;Acc:MGI:1352495] | Zfp385a | Enriched in dLGN | 1.511 | 2.106E-11 | 618.4 |
| ENSMUSG00000038462 | ubiquinol-cytochrome c reductase, Rieske iron-sulfur polypeptide 1 [Source:MGI Symbol;Acc:MGI:1913944] | Uqcrfs1 | Enriched in dLGN | 1.51 | 3.639E-08 | 172.4 |
| ENSMUSG00000025035 | ADP-ribosylation factor-like 3 [Source:MGI Symbol;Acc:MGI:1929699] | Arl3 | Enriched in dLGN | 1.509 | 2.445E-10 | 316.8 |
| ENSMUSG00000035958 | tyrosyl-DNA phosphodiesterase 2 [Source:MGI Symbol;Acc:MGI:1860486] | Tdp2 | Enriched in dLGN | 1.509 | 0.0001056 | 43.29 |
| ENSMUSG00000036438 | calmodulin 2 [Source:MGI Symbol;Acc:MGI:103250] | Calm2 | Enriched in dLGN | 1.509 | 0.0004691 | 1256 |
| ENSMUSG00000031176 | dynein light chain Tctex-type 3 [Source:MGI Symbol;Acc:MGI:1914367] | Dynlt3 | Enriched in dLGN | 1.507 | 0.00002026 | 112.8 |
| ENSMUSG00000079317 | trafficking protein particle complex 2 [Source:MGI Symbol;Acc:MGI:1913476] | Trappc2 | Enriched in dLGN | 1.501 | 0.0003046 | 15.83 |
| ENSMUSG00000031068 | glutaredoxin 3 [Source:MGI Symbol;Acc:MGI:1353653] | Glrx3 | Enriched in dLGN | 1.5 | 9.242E-07 | 113.9 |
| ENSMUSG00000025287 | acyl-CoA thioesterase 9 [Source:MGI Symbol;Acc:MGI:1928939] | Acot9 | Enriched in dLGN | 1.5 | 0.004603 | 23.21 |
| ENSMUSG00000002718 | chromosome segregation 1-like (S. cerevisiae) [Source:MGI Symbol;Acc:MGI:1339951] | Cse1l | Enriched in dLGN | 1.496 | 5.821E-07 | 100.5 |
| ENSMUSG00000040164 | K+ voltage-gated channel, subfamily S, 1 [Source:MGI Symbol;Acc:MGI:1197019] | Kcns1 | Enriched in dLGN | 1.495 | 0.00583 | 9.63 |
| ENSMUSG00000084786 | ubiquitin-like 5 [Source:MGI Symbol;Acc:MGI:1913427] | Ubl5 | Enriched in dLGN | 1.494 | 4.367E-09 | 89.51 |
| ENSMUSG00000034187 | N-ethylmaleimide sensitive fusion protein [Source:MGI Symbol;Acc:MGI:104560] | Nsf | Enriched in dLGN | 1.494 | 1.556E-08 | 1114 |
| ENSMUSG00000007891 | cathepsin D [Source:MGI Symbol;Acc:MGI:88562] | Ctsd | Enriched in dLGN | 1.494 | 0.003963 | 65.28 |
| ENSMUSG00000033419 | synaptosomal-associated protein 91 [Source:MGI Symbol;Acc:MGI:109132] | Snap91 | Enriched in dLGN | 1.49 | 3.061E-14 | 939.7 |
| ENSMUSG00000022587 | lymphocyte antigen 6 complex, locus E [Source:MGI Symbol;Acc:MGI:106651] | Ly6e | Enriched in dLGN | 1.488 | 0.000006563 | 95.38 |
| ENSMUSG00000083282 | cathepsin F [Source:MGI Symbol;Acc:MGI:1861434] | Ctsf | Enriched in dLGN | 1.485 | 6.798E-09 | 193.6 |
| ENSMUSG00000027350 | chromogranin B [Source:MGI Symbol;Acc:MGI:88395] | Chgb | Enriched in dLGN | 1.484 | 8.135E-08 | 1564 |
| ENSMUSG00000026188 | transmembrane protein 169 [Source:MGI Symbol;Acc:MGI:2442781] | Tmem169 | Enriched in dLGN | 1.481 | 1.324E-10 | 139.3 |
| ENSMUSG00000028837 | proteasome (prosome, macropain) subunit, beta type 2 [Source:MGI Symbol;Acc:MGI:1347045] | Psmb2 | Enriched in dLGN | 1.479 | 0.000207 | 65.51 |
| ENSMUSG00000010095 | solute carrier family 3 (activators of dibasic and neutral amino acid transport), member 2 [Source:MGI Symbol;Acc:MGI:96955] | Slc3a2 | Enriched in dLGN | 1.477 | 2.242E-10 | 248.5 |
| ENSMUSG00000032966 | FK506 binding protein 1a [Source:MGI Symbol;Acc:MGI:95541] | Fkbp1a | Enriched in dLGN | 1.475 | 8.488E-10 | 430.5 |
| ENSMUSG00000024414 | mitochondrial ribosomal protein L27 [Source:MGI Symbol;Acc:MGI:2137224] | Mrpl27 | Enriched in dLGN | 1.475 | 0.0002396 | 26.69 |
| ENSMUSG00000079426 | actin related protein 2/3 complex, subunit 4 [Source:MGI Symbol;Acc:MGI:1915339] | Arpc4 | Enriched in dLGN | 1.473 | 5.098E-08 | 89.11 |
| ENSMUSG00000028318 | polymerase (RNA) I polypeptide E [Source:MGI Symbol;Acc:MGI:1929022] | Polr1e | Enriched in dLGN | 1.47 | 0.000005534 | 28.6 |
| ENSMUSG00000024194 | cutA divalent cation tolerance homolog [Source:MGI Symbol;Acc:MGI:1914925] | Cuta | Enriched in dLGN | 1.469 | 3.17E-08 | 66.24 |
| ENSMUSG00000036972 | zinc finger protein of the cerebellum 4 [Source:MGI Symbol;Acc:MGI:107201] | Zic4 | Enriched in dLGN | 1.469 | 0.0002192 | 142.7 |
| ENSMUSG00000030037 | mitochondrial ribosomal protein L53 [Source:MGI Symbol;Acc:MGI:1915749] | Mrpl53 | Enriched in dLGN | 1.468 | 0.00002467 | 26.23 |
| ENSMUSG00000047617 | non-homologous end joining factor [Source:MGI Symbol;Acc:MGI:2442831] | Paxx | Enriched in dLGN | 1.467 | 0.000001649 | 66.93 |
| ENSMUSG00000028176 | leucine rich repeat containing 7 [Source:MGI Symbol;Acc:MGI:2676665] | Lrrc7 | Enriched in dLGN | 1.466 | 0.00007186 | 677.2 |
| ENSMUSG00000071862 | leucine rich repeat transmembrane neuronal 2 [Source:MGI Symbol;Acc:MGI:2389174] | Lrrtm2 | Enriched in dLGN | 1.465 | 0.000001409 | 78.72 |
| ENSMUSG00000026526 | fumarate hydratase 1 [Source:MGI Symbol;Acc:MGI:95530] | Fh1 | Enriched in dLGN | 1.463 | 0.0001123 | 124 |
| ENSMUSG00000051223 | basic leucine zipper and W2 domains 1 [Source:MGI Symbol;Acc:MGI:1914132] | Bzw1 | Enriched in dLGN | 1.462 | 5.832E-07 | 393.6 |
| ENSMUSG00000049176 | FERM and PDZ domain containing 4 [Source:MGI Symbol;Acc:MGI:3042378] | Frmpd4 | Enriched in dLGN | 1.462 | 0.00006362 | 381.5 |
| ENSMUSG00000031634 | UFM1-specific peptidase 2 [Source:MGI Symbol;Acc:MGI:1913679] | Ufsp2 | Enriched in dLGN | 1.461 | 0.0001122 | 89.61 |
| ENSMUSG00000073676 | heat shock protein 1 (chaperonin 10) [Source:MGI Symbol;Acc:MGI:104680] | Hspe1 | Enriched in dLGN | 1.461 | 0.0001197 | 62.31 |
| ENSMUSG00000024099 | NADH:ubiquinone oxidoreductase core subunit V2 [Source:MGI Symbol;Acc:MGI:1920150] | Ndufv2 | Enriched in dLGN | 1.459 | 1.489E-10 | 217.5 |
| ENSMUSG00000049299 | trafficking protein particle complex 1 [Source:MGI Symbol;Acc:MGI:1098727] | Trappc1 | Enriched in dLGN | 1.459 | 0.00002482 | 33.39 |
| ENSMUSG00000025917 | COP9 signalosome subunit 5 [Source:MGI Symbol;Acc:MGI:1349415] | Cops5 | Enriched in dLGN | 1.459 | 0.0001635 | 96.98 |
| ENSMUSG00000000296 | tumor protein D52-like 1 [Source:MGI Symbol;Acc:MGI:1298386] | Tpd52l1 | Enriched in dLGN | 1.458 | 0.00001515 | 61.04 |
| ENSMUSG00000019828 | glutamate receptor, metabotropic 1 [Source:MGI Symbol;Acc:MGI:1351338] | Grm1 | Enriched in dLGN | 1.456 | 0.00002628 | 572.7 |
| ENSMUSG00000030612 | mitochondrial ribosomal protein L46 [Source:MGI Symbol;Acc:MGI:1914558] | Mrpl46 | Enriched in dLGN | 1.455 | 0.00001752 | 33.73 |
| ENSMUSG00000024587 | asparaginyl-tRNA synthetase [Source:MGI Symbol;Acc:MGI:1917473] | Nars | Enriched in dLGN | 1.452 | 7.656E-08 | 419 |
| ENSMUSG00000043162 | Pigy upstream reading frame [Source:MGI Symbol;Acc:MGI:1913709] | Pyurf | Enriched in dLGN | 1.45 | 0.0001301 | 52.19 |
| ENSMUSG00000019505 | ubiquitin B [Source:MGI Symbol;Acc:MGI:98888] | Ubb | Enriched in dLGN | 1.45 | 0.000938 | 652 |
| ENSMUSG00000019796 | low density lipoprotein receptor-related protein 11 [Source:MGI Symbol;Acc:MGI:2442989] | Lrp11 | Enriched in dLGN | 1.447 | 1.062E-09 | 271 |
| ENSMUSG00000030127 | COP9 signalosome subunit 7A [Source:MGI Symbol;Acc:MGI:1349400] | Cops7a | Enriched in dLGN | 1.446 | 1.54E-09 | 167 |
| ENSMUSG00000090841 | myosin, light polypeptide 6, alkali, smooth muscle and non-muscle [Source:MGI Symbol;Acc:MGI:109318] | Myl6 | Enriched in dLGN | 1.442 | 4.951E-07 | 274.2 |
| ENSMUSG00000028609 | mago homolog, exon junction complex core component [Source:MGI Symbol;Acc:MGI:1330312] | Magoh | Enriched in dLGN | 1.442 | 0.001326 | 35.71 |
| ENSMUSG00000025410 | dynactin 2 [Source:MGI Symbol;Acc:MGI:107733] | Dctn2 | Enriched in dLGN | 1.441 | 1.508E-13 | 382.2 |
| ENSMUSG00000026585 | kinesin-associated protein 3 [Source:MGI Symbol;Acc:MGI:107566] | Kifap3 | Enriched in dLGN | 1.439 | 3.711E-07 | 663.9 |

| | | | | | | |
|---|---|---|---|---|---|---|
| ENSMUSG00000025781 | ATP synthase, H+ transporting, mitochondrial F1 complex, gamma polypeptide 1 [Source:MGI Symbol;Acc:MGI:1261437] | Atp5c1 | Enriched in dLGN | 1.438 | 1.817E-09 | 290 |
| ENSMUSG00000058400 | pyroglutamylated RFamide peptide receptor [Source:MGI Symbol;Acc:MGI:2677633] | Qrfpr | Enriched in dLGN | 1.436 | 0.00627 | 24.15 |
| ENSMUSG00000032336 | neuroplastin [Source:MGI Symbol;Acc:MGI:108077] | Nptn | Enriched in dLGN | 1.434 | 2.602E-12 | 662.1 |
| ENSMUSG00000026103 | glutaminase [Source:MGI Symbol;Acc:MGI:95752] | Gls | Enriched in dLGN | 1.434 | 2.335E-10 | 697 |
| ENSMUSG00000062526 | metallophosphoesterase 1 [Source:MGI Symbol;Acc:MGI:2661311] | Mppe1 | Enriched in dLGN | 1.433 | 0.00002063 | 32.36 |
| ENSMUSG00000056091 | ST3 beta-galactoside alpha-2,3-sialyltransferase 5 [Source:MGI Symbol;Acc:MGI:1339963] | St3gal5 | Enriched in dLGN | 1.429 | 1.611E-11 | 176.3 |
| ENSMUSG00000024044 | erythrocyte membrane protein band 4.1 like 3 [Source:MGI Symbol;Acc:MGI:103008] | Epb41l3 | Enriched in dLGN | 1.428 | 9.786E-07 | 1430 |
| ENSMUSG00000028964 | Parkinson disease (autosomal recessive, early onset) 7 [Source:MGI Symbol;Acc:MGI:2135637] | Park7 | Enriched in dLGN | 1.427 | 1.705E-09 | 178.9 |
| ENSMUSG00000022965 | interferon gamma receptor 2 [Source:MGI Symbol;Acc:MGI:107654] | Ifngr2 | Enriched in dLGN | 1.426 | 8.254E-07 | 119.8 |
| ENSMUSG00000027273 | synaptosomal-associated protein 25 [Source:MGI Symbol;Acc:MGI:98331] | Snap25 | Enriched in dLGN | 1.426 | 9.076E-07 | 1966 |
| ENSMUSG00000031609 | sin3 associated polypeptide [Source:MGI Symbol;Acc:MGI:1929129] | Sap30 | Enriched in dLGN | 1.426 | 0.001204 | 26.75 |
| ENSMUSG00000021748 | pyruvate dehydrogenase (lipoamide) beta [Source:MGI Symbol;Acc:MGI:1915513] | Pdhb | Enriched in dLGN | 1.425 | 9.236E-08 | 160.8 |
| ENSMUSG00000028690 | methylmalonic aciduria cblC type, with homocystinuria [Source:MGI Symbol;Acc:MGI:1914346] | Mmachc | Enriched in dLGN | 1.425 | 0.0003228 | 24.85 |
| ENSMUSG00000027601 | mitochondrial fission regulator 1 [Source:MGI Symbol;Acc:MGI:1914722] | Mtfr1 | Enriched in dLGN | 1.421 | 0.004934 | 98.99 |
| ENSMUSG00000002017 | family with sequence similarity 98, member A [Source:MGI Symbol;Acc:MGI:1919972] | Fam98a | Enriched in dLGN | 1.419 | 1.524E-07 | 80.81 |
| ENSMUSG00000073838 | Tu translation elongation factor, mitochondrial [Source:MGI Symbol;Acc:MGI:1923686] | Tufm | Enriched in dLGN | 1.419 | 0.0000274 | 100.2 |
| ENSMUSG00000020089 | pyrophosphatase (inorganic) 1 [Source:MGI Symbol;Acc:MGI:97831] | Ppa1 | Enriched in dLGN | 1.418 | 0.001171 | 125.6 |
| ENSMUSG00000004031 | bone morphogenic protein/retinoic acid inducible neural-specific 2 [Source:MGI Symbol;Acc:MGI:2443333] | Brinp2 | Enriched in dLGN | 1.417 | 3.716E-07 | 323.9 |
| ENSMUSG00000021548 | cyclin H [Source:MGI Symbol;Acc:MGI:1913921] | Ccnh | Enriched in dLGN | 1.416 | 0.00003242 | 69.07 |
| ENSMUSG00000027133 | NOP10 ribonucleoprotein [Source:MGI Symbol;Acc:MGI:1913431] | Nop10 | Enriched in dLGN | 1.416 | 0.0006636 | 25.83 |
| ENSMUSG00000040136 | ATP-binding cassette, sub-family C (CFTR/MRP), member 8 [Source:MGI Symbol;Acc:MGI:1352629] | Abcc8 | Enriched in dLGN | 1.415 | 0.0001605 | 86.74 |
| ENSMUSG00000017778 | cytochrome c oxidase subunit 7C [Source:MGI Symbol;Acc:MGI:103226] | Cox7c | Enriched in dLGN | 1.414 | 0.00005908 | 35.38 |
| ENSMUSG00000001763 | tetraspanin 33 [Source:MGI Symbol;Acc:MGI:1919012] | Tspan33 | Enriched in dLGN | 1.411 | 0.000001302 | 62.53 |
| ENSMUSG00000020990 | cyclin-dependent kinase-like 1 (CDC2-related kinase) [Source:MGI Symbol;Acc:MGI:1918341] | Cdkl1 | Enriched in dLGN | 1.411 | 0.0001351 | 53.67 |
| ENSMUSG00000021486 | PRELI domain containing 1 [Source:MGI Symbol;Acc:MGI:1913744] | Prelid1 | Enriched in dLGN | 1.408 | 6.493E-09 | 247.5 |
| ENSMUSG00000027698 | neutral cholesterol ester hydrolase 1 [Source:MGI Symbol;Acc:MGI:2443191] | Nceh1 | Enriched in dLGN | 1.407 | 0.000008753 | 190.1 |
| ENSMUSG00000062949 | ATPase, class VI, type 11C [Source:MGI Symbol;Acc:MGI:1859661] | Atp11c | Enriched in dLGN | 1.407 | 0.00002628 | 114.4 |
| ENSMUSG00000017412 | calcium channel, voltage-dependent, beta 4 subunit [Source:MGI Symbol;Acc:MGI:103301] | Cacnb4 | Enriched in dLGN | 1.407 | 0.0001031 | 413.3 |
| ENSMUSG00000037772 | mitochondrial ribosomal protein L23 [Source:MGI Symbol;Acc:MGI:1196612] | Mrpl23 | Enriched in dLGN | 1.406 | 0.004339 | 20.4 |
| ENSMUSG00000006932 | catenin (cadherin associated protein), beta 1 [Source:MGI Symbol;Acc:MGI:88276] | Ctnnb1 | Enriched in dLGN | 1.405 | 7.537E-07 | 1065 |
| ENSMUSG00000030647 | NADH:ubiquinone oxidoreductase subunit C2 [Source:MGI Symbol;Acc:MGI:1344370] | Ndufc2 | Enriched in dLGN | 1.405 | 0.0005425 | 77.57 |
| ENSMUSG00000000339 | RNA 3'-terminal phosphate cyclase [Source:MGI Symbol;Acc:MGI:1913618] | Rtca | Enriched in dLGN | 1.403 | 1.433E-08 | 93.2 |
| ENSMUSG00000001999 | biliverdin reductase A [Source:MGI Symbol;Acc:MGI:88170] | Blvra | Enriched in dLGN | 1.403 | 0.00002741 | 31.09 |
| ENSMUSG00000016308 | ubiquitin-conjugating enzyme E2A [Source:MGI Symbol;Acc:MGI:102959] | Ube2a | Enriched in dLGN | 1.401 | 2.378E-07 | 44.52 |
| ENSMUSG00000040767 | small nuclear ribonucleoprotein 25 (U11/U12) [Source:MGI Symbol;Acc:MGI:1925622] | Snrnp25 | Enriched in dLGN | 1.401 | 0.00002956 | 25.98 |
| ENSMUSG00000039431 | myotubularin related protein 7 [Source:MGI Symbol;Acc:MGI:1891693] | Mtmr7 | Enriched in dLGN | 1.398 | 1.805E-08 | 199.1 |
| ENSMUSG00000031149 | PRA1 domain family 2 [Source:MGI Symbol;Acc:MGI:1859607] | Praf2 | Enriched in dLGN | 1.396 | 0.02138 | 22.15 |
| ENSMUSG00000075706 | glutathione peroxidase 4 [Source:MGI Symbol;Acc:MGI:104767] | Gpx4 | Enriched in dLGN | 1.394 | 1.049E-07 | 170.9 |
| ENSMUSG00000028261 | NADH:ubiquinone oxidoreductase complex assembly factor 4 [Source:MGI Symbol;Acc:MGI:1915743] | Ndufaf4 | Enriched in dLGN | 1.394 | 0.000006917 | 70.33 |
| ENSMUSG00000000168 | dihydrolipoamide S-acetyltransferase (E2 component of pyruvate dehydrogenase complex) [Source:MGI Symbol;Acc:MGI:2385311] | Dlat | Enriched in dLGN | 1.392 | 2.683E-09 | 196.1 |
| ENSMUSG00000059278 | N(alpha)-acetyltransferase 38, NatC auxiliary subunit [Source:MGI Symbol;Acc:MGI:1925554] | Naa38 | Enriched in dLGN | 1.392 | 0.001182 | 31.83 |
| ENSMUSG00000078941 | adenylate kinase 6 [Source:MGI Symbol;Acc:MGI:5510732] | Ak6 | Enriched in dLGN | 1.39 | 0.001193 | 18.09 |
| ENSMUSG00000031701 | DnaJ heat shock protein family (Hsp40) member A2 [Source:MGI Symbol;Acc:MGI:1931882] | Dnaja2 | Enriched in dLGN | 1.389 | 0.000001102 | 401 |
| ENSMUSG00000048495 | tRNA-yW synthesizing protein 5 [Source:MGI Symbol;Acc:MGI:1915986] | Tyw5 | Enriched in dLGN | 1.389 | 0.003057 | 22.16 |
| ENSMUSG00000035517 | tudor domain containing 7 [Source:MGI Symbol;Acc:MGI:2140279] | Tdrd7 | Enriched in dLGN | 1.387 | 3.7E-11 | 468.1 |
| ENSMUSG00000019087 | ATPase, H+ transporting, lysosomal accessory protein 1 [Source:MGI Symbol;Acc:MGI:109629] | Atp6ap1 | Enriched in dLGN | 1.387 | 6.695E-07 | 276.8 |
| ENSMUSG00000026492 | transcription factor B2, mitochondrial [Source:MGI Symbol;Acc:MGI:107937] | Tfb2m | Enriched in dLGN | 1.383 | 0.0002614 | 22.6 |
| ENSMUSG00000028370 | pregnancy-associated plasma protein A [Source:MGI Symbol;Acc:MGI:97479] | Pappa | Enriched in dLGN | 1.383 | 0.00113 | 69.09 |
| ENSMUSG00000105827 | H2B clustered histone 18 [Source:MGI Symbol;Acc:MGI:2448413] | H2bc18 | Enriched in dLGN | 1.383 | 0.003865 | 42.81 |
| ENSMUSG00000049792 | BCL2-associated athanogene 5 [Source:MGI Symbol;Acc:MGI:1917619] | Bag5 | Enriched in dLGN | 1.381 | 4.136E-11 | 146 |
| ENSMUSG00000040389 | WD repeat domain 47 [Source:MGI Symbol;Acc:MGI:2139593] | Wdr47 | Enriched in dLGN | 1.38 | 1.201E-09 | 344.7 |
| ENSMUSG00000052456 | guided entry of tail-anchored proteins factor 3, ATPase [Source:MGI Symbol;Acc:MGI:1928379] | Get3 | Enriched in dLGN | 1.38 | 0.000002455 | 120.5 |
| ENSMUSG00000027282 | mitochondrial carrier 2 [Source:MGI Symbol;Acc:MGI:1929260] | Mtch2 | Enriched in dLGN | 1.379 | 6.161E-08 | 103.4 |
| ENSMUSG00000045731 | prepronociceptin [Source:MGI Symbol;Acc:MGI:105308] | Pnoc | Enriched in dLGN | 1.379 | 0.0004697 | 50.56 |
| ENSMUSG00000032249 | acidic (leucine-rich) nuclear phosphoprotein 32 family, member A [Source:MGI Symbol;Acc:MGI:108447] | Anp32a | Enriched in dLGN | 1.377 | 1.078E-14 | 803.6 |
| ENSMUSG00000023033 | sodium channel, voltage-gated, type VIII, alpha [Source:MGI Symbol;Acc:MGI:103169] | Scn8a | Enriched in dLGN | 1.377 | 1.16E-11 | 1192 |
| ENSMUSG00000021737 | proteasome (prosome, macropain) 26S subunit, non-ATPase, 6 [Source:MGI Symbol;Acc:MGI:1913663] | Psmd6 | Enriched in dLGN | 1.377 | 0.00000171 | 147.1 |
| ENSMUSG00000020349 | protein phosphatase 2 (formerly 2A), catalytic subunit, alpha isoform [Source:MGI Symbol;Acc:MGI:1321159] | Ppp2ca | Enriched in dLGN | 1.376 | 4.681E-08 | 828.3 |
| ENSMUSG00000016427 | NADH:ubiquinone oxidoreductase subunit A1 [Source:MGI Symbol;Acc:MGI:1929511] | Ndufa1 | Enriched in dLGN | 1.373 | 8.472E-07 | 84.15 |
| ENSMUSG00000037361 | splicing factor 3B, subunit 6 [Source:MGI Symbol;Acc:MGI:1913305] | Sf3b6 | Enriched in dLGN | 1.372 | 0.001311 | 35.23 |
| ENSMUSG00000027248 | protein disulfide isomerase associated 3 [Source:MGI Symbol;Acc:MGI:95834] | Pdia3 | Enriched in dLGN | 1.371 | 0.000007153 | 401 |
| ENSMUSG00000033685 | uncoupling protein 2 (mitochondrial, proton carrier) [Source:MGI Symbol;Acc:MGI:109354] | Ucp2 | Enriched in dLGN | 1.371 | 0.0003614 | 149.6 |
| ENSMUSG00000038555 | receptor accessory protein 2 [Source:MGI Symbol;Acc:MGI:2385070] | Reep2 | Enriched in dLGN | 1.37 | 1.091E-09 | 302.7 |
| ENSMUSG00000040048 | NADH:ubiquinone oxidoreductase subunit B10 [Source:MGI Symbol;Acc:MGI:1915592] | Ndufb10 | Enriched in dLGN | 1.37 | 1.798E-07 | 201.2 |
| ENSMUSG00000021114 | ATPase, H+ transporting, lysosomal V1 subunit D [Source:MGI Symbol;Acc:MGI:1921084] | Atp6v1d | Enriched in dLGN | 1.37 | 0.000001064 | 284.8 |
| ENSMUSG00000013921 | CAP-GLY domain containing linker protein 3 [Source:MGI Symbol;Acc:MGI:1923936] | Clip3 | Enriched in dLGN | 1.365 | 0.00006594 | 1062 |
| ENSMUSG00000002797 | gamma-glutamyl cyclotransferase [Source:MGI Symbol;Acc:MGI:95700] | Ggct | Enriched in dLGN | 1.364 | 0.00006137 | 25.71 |
| ENSMUSG00000039385 | cadherin 6 [Source:MGI Symbol;Acc:MGI:107435] | Cdh6 | Enriched in dLGN | 1.363 | 0.0002065 | 237 |
| ENSMUSG00000028212 | cyclin E2 [Source:MGI Symbol;Acc:MGI:1329034] | Ccne2 | Enriched in dLGN | 1.362 | 0.01442 | 12.53 |
| ENSMUSG00000029767 | calumenin [Source:MGI Symbol;Acc:MGI:1097158] | Calu | Enriched in dLGN | 1.361 | 0.000003951 | 300.9 |
| ENSMUSG00000022337 | ER membrane protein complex subunit 2 [Source:MGI Symbol;Acc:MGI:1913986] | Emc2 | Enriched in dLGN | 1.359 | 0.000000659 | 104.6 |
| ENSMUSG00000038722 | BUD31 homolog [Source:MGI Symbol;Acc:MGI:2141291] | Bud31 | Enriched in dLGN | 1.358 | 4.068E-08 | 48.97 |
| ENSMUSG00000027406 | isocitrate dehydrogenase 3 (NAD+) beta [Source:MGI Symbol;Acc:MGI:2158650] | Idh3b | Enriched in dLGN | 1.358 | 0.000002398 | 236.2 |
| ENSMUSG00000030869 | NADH:ubiquinone oxidoreductase subunit AB1 [Source:MGI Symbol;Acc:MGI:1917566] | Ndufab1 | Enriched in dLGN | 1.358 | 0.00003605 | 76.72 |
| ENSMUSG00000036181 | H1.2 linker histone, cluster member [Source:MGI Symbol;Acc:MGI:1931526] | H1f2 | Enriched in dLGN | 1.358 | 0.0002204 | 187.7 |
| ENSMUSG00000003380 | Rab acceptor 1 (prenylated) [Source:MGI Symbol;Acc:MGI:1201692] | Rabac1 | Enriched in dLGN | 1.357 | 7.452E-09 | 110.7 |
| ENSMUSG00000015217 | high mobility group box 3 [Source:MGI Symbol;Acc:MGI:1098219] | Hmgb3 | Enriched in dLGN | 1.355 | 0.0000251 | 141 |
| ENSMUSG00000041444 | Rho GTPase activating protein 32 [Source:MGI Symbol;Acc:MGI:2450166] | Arhgap32 | Enriched in dLGN | 1.349 | 3.606E-09 | 970.5 |
| ENSMUSG00000031059 | NADH:ubiquinone oxidoreductase subunit B11 [Source:MGI Symbol;Acc:MGI:1349919] | Ndufb11 | Enriched in dLGN | 1.349 | 5.157E-08 | 155.4 |
| ENSMUSG00000031918 | myotubularin related protein 2 [Source:MGI Symbol;Acc:MGI:1924366] | Mtmr2 | Enriched in dLGN | 1.349 | 6.273E-07 | 157.4 |
| ENSMUSG00000028351 | bone morphogenic protein/retinoic acid inducible neural specific 1 [Source:MGI Symbol;Acc:MGI:1928478] | Brinp1 | Enriched in dLGN | 1.349 | 0.00006956 | 586.1 |
| ENSMUSG00000019373 | COP9 signalosome subunit 3 [Source:MGI Symbol;Acc:MGI:1349409] | Cops3 | Enriched in dLGN | 1.349 | 0.0001019 | 92.31 |
| ENSMUSG00000032131 | ATP binding cassette subfamily G member 4 [Source:MGI Symbol;Acc:MGI:1890594] | Abcg4 | Enriched in dLGN | 1.348 | 1.942E-11 | 245 |
| ENSMUSG00000020115 | TANK-binding kinase 1 [Source:MGI Symbol;Acc:MGI:1929658] | Tbk1 | Enriched in dLGN | 1.345 | 0.000005985 | 121.8 |
| ENSMUSG00000052632 | ArfGAP with SH3 domain, ankyrin repeat and PH domain 2 [Source:MGI Symbol;Acc:MGI:2685438] | Asap2 | Enriched in dLGN | 1.343 | 1.355E-12 | 463.9 |
| ENSMUSG00000036834 | phospholipase C, eta 1 [Source:MGI Symbol;Acc:MGI:2683547] | Plch1 | Enriched in dLGN | 1.341 | 7.637E-08 | 269.3 |
| ENSMUSG00000022892 | amyloid beta (A4) precursor protein [Source:MGI Symbol;Acc:MGI:88059] | App | Enriched in dLGN | 1.341 | 0.00001913 | 4592 |
| ENSMUSG00000024217 | U1 small nuclear ribonucleoprotein C [Source:MGI Symbol;Acc:MGI:109489] | Snrpc | Enriched in dLGN | 1.34 | 0.00282 | 26.15 |
| ENSMUSG00000001380 | histidyl-tRNA synthetase [Source:MGI Symbol;Acc:MGI:108087] | Hars | Enriched in dLGN | 1.339 | 9.091E-10 | 256 |
| ENSMUSG00000070319 | eukaryotic translation initiation factor 3, subunit G [Source:MGI Symbol;Acc:MGI:1858258] | Eif3g | Enriched in dLGN | 1.339 | 0.0001237 | 174.2 |
| ENSMUSG00000068749 | proteasome subunit alpha 5 [Source:MGI Symbol;Acc:MGI:1347009] | Psma5 | Enriched in dLGN | 1.339 | 0.0006812 | 53.28 |
| ENSMUSG00000038208 | post-GPI attachment to proteins 3 [Source:MGI Symbol;Acc:MGI:2444461] | Pgap3 | Enriched in dLGN | 1.339 | 0.02364 | 48.45 |
| ENSMUSG00000015766 | epidermal growth factor receptor pathway substrate 8 [Source:MGI Symbol;Acc:MGI:104684] | Eps8 | Enriched in dLGN | 1.338 | 6.468E-07 | 220.6 |
| ENSMUSG00000038762 | ATP-binding cassette, sub-family F (GCN20), member 1 [Source:MGI Symbol;Acc:MGI:1351658] | Abcf1 | Enriched in dLGN | 1.337 | 1.973E-11 | 576.3 |
| ENSMUSG00000031505 | NAD(P)HX dehydratase [Source:MGI Symbol;Acc:MGI:1913353] | Naxd | Enriched in dLGN | 1.335 | 0.005382 | 43.91 |
| ENSMUSG00000037149 | DEAD box helicase 1 [Source:MGI Symbol;Acc:MGI:2144727] | Ddx1 | Enriched in dLGN | 1.334 | 4.233E-09 | 410.1 |
| ENSMUSG00000030751 | proteasome subunit alpha 1 [Source:MGI Symbol;Acc:MGI:1347005] | Psma1 | Enriched in dLGN | 1.334 | 0.0006461 | 72.84 |
| ENSMUSG00000002332 | dehydrogenase/reductase (SDR family) member 1 [Source:MGI Symbol;Acc:MGI:1196314] | Dhrs1 | Enriched in dLGN | 1.334 | 0.002363 | 49.92 |
| ENSMUSG00000030291 | mediator complex subunit 21 [Source:MGI Symbol;Acc:MGI:1347064] | Med21 | Enriched in dLGN | 1.334 | 0.003543 | 15.67 |
| ENSMUSG00000063694 | cytochrome c, somatic [Source:MGI Symbol;Acc:MGI:88578] | Cycs | Enriched in dLGN | 1.333 | 0.00007799 | 116.9 |
| ENSMUSG00000060212 | pecanex homolog 2 [Source:MGI Symbol;Acc:MGI:2445010] | Pcnx2 | Enriched in dLGN | 1.332 | 1.202E-07 | 282.9 |
| ENSMUSG00000021314 | amphiphysin [Source:MGI Symbol;Acc:MGI:103574] | Amph | Enriched in dLGN | 1.331 | 2.863E-14 | 841.6 |
| ENSMUSG00000010376 | neural precursor cell expressed, developmentally down-regulated gene 8 [Source:MGI Symbol;Acc:MGI:97301] | Nedd8 | Enriched in dLGN | 1.329 | 0.000004909 | 191.2 |
| ENSMUSG00000021024 | proteasome subunit alpha 6 [Source:MGI Symbol;Acc:MGI:1347006] | Psma6 | Enriched in dLGN | 1.329 | 0.007264 | 71.31 |
| ENSMUSG00000036748 | CUE domain containing 2 [Source:MGI Symbol;Acc:MGI:1914366] | Cuedc2 | Enriched in dLGN | 1.328 | 9.183E-07 | 145.7 |
| ENSMUSG00000074797 | inosine triphosphatase (nucleoside triphosphate pyrophosphatase) [Source:MGI Symbol;Acc:MGI:96622] | Itpa | Enriched in dLGN | 1.327 | 0.000002466 | 70.22 |
| ENSMUSG00000079508 | apolipoprotein O [Source:MGI Symbol;Acc:MGI:1915566] | Apoo | Enriched in dLGN | 1.323 | 0.02362 | 29.79 |
| ENSMUSG00000021947 | crystallin, lambda 1 [Source:MGI Symbol;Acc:MGI:1915881] | Cryl1 | Enriched in dLGN | 1.322 | 0.001099 | 28.77 |
| ENSMUSG00000043831 | LysM, putative peptidoglycan-binding, domain containing 4 [Source:MGI Symbol;Acc:MGI:1922349] | Lysmd4 | Enriched in dLGN | 1.321 | 3.194E-07 | 79.01 |
| ENSMUSG00000037072 | selenoprotein F [Source:MGI Symbol;Acc:MGI:1927947] | Selenof | Enriched in dLGN | 1.32 | 1.344E-08 | 179.7 |
| ENSMUSG00000032839 | transient receptor potential cation channel, subfamily C, member 1 [Source:MGI Symbol;Acc:MGI:109528] | Trpc1 | Enriched in dLGN | 1.316 | 2.243E-09 | 140.9 |
| ENSMUSG00000022024 | SGT1, suppressor of G2 allele of SKP1 (S. cerevisiae) [Source:MGI Symbol;Acc:MGI:1915205] | Sugt1 | Enriched in dLGN | 1.313 | 8.965E-10 | 199.6 |
| ENSMUSG00000023456 | triosephosphate isomerase 1 [Source:MGI Symbol;Acc:MGI:98797] | Tpi1 | Enriched in dLGN | 1.313 | 0.00007263 | 483.5 |
| ENSMUSG00000079941 | cytochrome c oxidase subunit 5B, pseudogene [Source:MGI Symbol;Acc:MGI:3649411] | Cox5b-ps | Enriched in dLGN | 1.313 | 0.003166 | 35.35 |
| ENSMUSG00000038690 | ATP synthase, H+ transporting, mitochondrial F0 complex, subunit F2 [Source:MGI Symbol;Acc:MGI:1927558] | Atp5j2 | Enriched in dLGN | 1.312 | 0.000003401 | 138.3 |
| ENSMUSG00000022890 | ATP synthase, H+ transporting, mitochondrial F0 complex, subunit F [Source:MGI Symbol;Acc:MGI:107777] | Atp5j | Enriched in dLGN | 1.311 | 0.00004047 | 249.2 |
| ENSMUSG00000026824 | potassium inwardly-rectifying channel, subfamily J, member 3 [Source:MGI Symbol;Acc:MGI:104742] | Kcnj3 | Enriched in dLGN | 1.31 | 0.00008792 | 206.8 |
| ENSMUSG00000019689 | formation of mitochondrial complex V assembly factor 1 [Source:MGI Symbol;Acc:MGI:1913367] | Fmc1 | Enriched in dLGN | 1.31 | 0.001762 | 21.78 |
| ENSMUSG00000021832 | proteasome (prosome, macropain) 26S subunit, ATPase, 6 [Source:MGI Symbol;Acc:MGI:1914339] | Psmc6 | Enriched in dLGN | 1.309 | 0.00003172 | 131.6 |
| ENSMUSG00000039910 | Cbp/p300-interacting transactivator, with Glu/Asp-rich carboxy-terminal domain, 2 [Source:MGI Symbol;Acc:MGI:1306784] | Cited2 | Enriched in dLGN | 1.308 | 0.000001897 | 198.6 |
| ENSMUSG00000051671 | cytochrome c oxidase assembly factor 6 [Source:MGI Symbol;Acc:MGI:1915142] | Coa6 | Enriched in dLGN | 1.306 | 0.006689 | 13.32 |
| ENSMUSG00000036850 | mitochondrial ribosomal protein L41 [Source:MGI Symbol;Acc:MGI:1333816] | Mrpl41 | Enriched in dLGN | 1.304 | 0.000002797 | 53.68 |
| ENSMUSG00000026577 | basic leucine zipper nuclear factor 1 [Source:MGI Symbol;Acc:MGI:1201607] | Blzf1 | Enriched in dLGN | 1.303 | 4.571E-07 | 70.6 |
| ENSMUSG00000055839 | elongin B [Source:MGI Symbol;Acc:MGI:1914923] | Elob | Enriched in dLGN | 1.302 | 0.00001428 | 99.23 |
| ENSMUSG00000017188 | cytochrome C oxidase assembly factor 3 [Source:MGI Symbol;Acc:MGI:1098757] | Coa3 | Enriched in dLGN | 1.302 | 0.00005192 | 48.29 |
| ENSMUSG00000057315 | Rho GTPase activating protein 24 [Source:MGI Symbol;Acc:MGI:1922647] | Arhgap24 | Enriched in dLGN | 1.302 | 0.0001324 | 291.9 |
| ENSMUSG00000022956 | ATP synthase, H+ transporting, mitochondrial F1 complex, O subunit [Source:MGI Symbol;Acc:MGI:106341] | Atp5o | Enriched in dLGN | 1.302 | 0.0002126 | 36.29 |
| ENSMUSG00000016349 | eukaryotic translation elongation factor 1 alpha 2 [Source:MGI Symbol;Acc:MGI:1096317] | Eef1a2 | Enriched in dLGN | 1.301 | 0.00007514 | 1496 |

| | | | | | | |
|---|---|---|---|---|---|---|
| ENSMUSG00000026643 | N-myristoyltransferase 2 [Source:MGI Symbol;Acc:MGI:1202298] | Nmt2 | Enriched in dLGN | 1.297 | 9.728E-12 | 203.8 |
| ENSMUSG00000037710 | CDGSH iron sulfur domain 1 [Source:MGI Symbol;Acc:MGI:1261855] | Cisd1 | Enriched in dLGN | 1.297 | 0.000007232 | 88.71 |
| ENSMUSG00000090553 | small nuclear ribonucleoprotein E [Source:MGI Symbol;Acc:MGI:98346] | Snrpe | Enriched in dLGN | 1.297 | 0.000124 | 27.79 |
| ENSMUSG00000022241 | threonyl-tRNA synthetase [Source:MGI Symbol;Acc:MGI:106314] | Tars | Enriched in dLGN | 1.295 | 3.417E-07 | 160.9 |
| ENSMUSG00000031791 | transmembrane protein 38A [Source:MGI Symbol;Acc:MGI:1921416] | Tmem38a | Enriched in dLGN | 1.284 | 2.439E-16 | 149.4 |
| ENSMUSG00000038550 | circadian associated repressor of transcription [Source:MGI Symbol;Acc:MGI:2684975] | Ciart | Enriched in dLGN | 1.281 | 0.01281 | 14.25 |
| ENSMUSG00000050730 | Rho GTPase activating protein 42 [Source:MGI Symbol;Acc:MGI:1918794] | Arhgap42 | Enriched in dLGN | 1.28 | 0.0006392 | 71.18 |
| ENSMUSG00000054423 | Ca2+-dependent secretion activator [Source:MGI Symbol;Acc:MGI:1350922] | Cadps | Enriched in dLGN | 1.279 | 0.00005952 | 1472 |
| ENSMUSG00000014294 | NADH:ubiquinone oxidoreductase subunit A2 [Source:MGI Symbol;Acc:MGI:1343103] | Ndufa2 | Enriched in dLGN | 1.276 | 8.273E-07 | 115.8 |
| ENSMUSG00000063406 | transmembrane p24 trafficking protein 5 [Source:MGI Symbol;Acc:MGI:1921586] | Tmed5 | Enriched in dLGN | 1.276 | 0.000341 | 35.86 |
| ENSMUSG00000033389 | Rho GTPase activating protein 44 [Source:MGI Symbol;Acc:MGI:2144423] | Arhgap44 | Enriched in dLGN | 1.274 | 1.079E-09 | 522.2 |
| ENSMUSG00000026456 | cytochrome b5 reductase 1 [Source:MGI Symbol;Acc:MGI:1919267] | Cyb5r1 | Enriched in dLGN | 1.272 | 0.01868 | 33.56 |
| ENSMUSG00000017781 | phosphatidylinositol transfer protein, alpha [Source:MGI Symbol;Acc:MGI:99887] | Pitpna | Enriched in dLGN | 1.271 | 9.901E-11 | 792.9 |
| ENSMUSG00000000326 | catechol-O-methyltransferase [Source:MGI Symbol;Acc:MGI:88470] | Comt | Enriched in dLGN | 1.271 | 1.478E-07 | 103.3 |
| ENSMUSG00000019861 | golgi associated PDZ and coiled-coil motif containing [Source:MGI Symbol;Acc:MGI:2149946] | Gopc | Enriched in dLGN | 1.269 | 6.13E-15 | 304.7 |
| ENSMUSG00000019814 | LTV1 ribosome biogenesis factor [Source:MGI Symbol;Acc:MGI:2447810] | Ltv1 | Enriched in dLGN | 1.269 | 0.001415 | 46.86 |
| ENSMUSG00000028447 | dynactin 3 [Source:MGI Symbol;Acc:MGI:1859251] | Dctn3 | Enriched in dLGN | 1.268 | 2.469E-08 | 123.3 |
| ENSMUSG00000021276 | cyclin-dependent kinase 2 interacting protein [Source:MGI Symbol;Acc:MGI:1914486] | Cinp | Enriched in dLGN | 1.268 | 8.351E-07 | 49.08 |
| ENSMUSG00000023460 | RAB12, member RAS oncogene family [Source:MGI Symbol;Acc:MGI:894284] | Rab12 | Enriched in dLGN | 1.266 | 1.43E-12 | 311.7 |
| ENSMUSG00000060279 | adaptor-related protein complex 2, alpha 1 subunit [Source:MGI Symbol;Acc:MGI:101921] | Ap2a1 | Enriched in dLGN | 1.265 | 4.933E-08 | 671.9 |
| ENSMUSG00000024425 | Nedd4 family interacting protein 1 [Source:MGI Symbol;Acc:MGI:1929601] | Ndfip1 | Enriched in dLGN | 1.263 | 0.000002485 | 747.3 |
| ENSMUSG00000024158 | hydroxyacyl glutathione hydrolase [Source:MGI Symbol;Acc:MGI:95745] | Hagh | Enriched in dLGN | 1.263 | 0.00009088 | 60.65 |
| ENSMUSG00000053317 | Sec61 beta subunit [Source:MGI Symbol;Acc:MGI:1913462] | Sec61b | Enriched in dLGN | 1.263 | 0.007623 | 19.39 |
| ENSMUSG00000063253 | short coiled-coil protein [Source:MGI Symbol;Acc:MGI:1927654] | Scoc | Enriched in dLGN | 1.256 | 3.572E-07 | 113.5 |
| ENSMUSG00000038279 | NOP2 nucleolar protein [Source:MGI Symbol;Acc:MGI:107891] | Nop2 | Enriched in dLGN | 1.254 | 0.000002224 | 78.7 |
| ENSMUSG00000025208 | mitochondrial ribosomal protein L43 [Source:MGI Symbol;Acc:MGI:2137229] | Mrpl43 | Enriched in dLGN | 1.254 | 0.000004805 | 70.84 |
| ENSMUSG00000028992 | nicotinamide nucleotide adenylyltransferase 1 [Source:MGI Symbol;Acc:MGI:1913704] | Nmnat1 | Enriched in dLGN | 1.252 | 0.0008909 | 20.77 |
| ENSMUSG00000039771 | polymerase (RNA) II (DNA directed) polypeptide J [Source:MGI Symbol;Acc:MGI:109582] | Polr2j | Enriched in dLGN | 1.249 | 0.003283 | 28.45 |
| ENSMUSG00000027742 | component of oligomeric golgi complex 6 [Source:MGI Symbol;Acc:MGI:1914792] | Cog6 | Enriched in dLGN | 1.248 | 2.159E-08 | 89.91 |
| ENSMUSG00000071654 | ubiquinol-cytochrome c reductase complex assembly factor 3 [Source:MGI Symbol;Acc:MGI:2147553] | Uqcc3 | Enriched in dLGN | 1.247 | 0.00001908 | 94.54 |
| ENSMUSG00000022561 | GPI anchor attachment protein 1 [Source:MGI Symbol;Acc:MGI:1202392] | Gpaa1 | Enriched in dLGN | 1.246 | 0.000004744 | 48.03 |
| ENSMUSG00000023089 | NADH:ubiquinone oxidoreductase subunit A5 [Source:MGI Symbol;Acc:MGI:1915452] | Ndufa5 | Enriched in dLGN | 1.244 | 0.00003404 | 166.1 |
| ENSMUSG00000028470 | histidine triad nucleotide binding protein 2 [Source:MGI Symbol;Acc:MGI:1916167] | Hint2 | Enriched in dLGN | 1.244 | 0.002044 | 22.68 |
| ENSMUSG00000044067 | G protein-coupled receptor 22 [Source:MGI Symbol;Acc:MGI:1920260] | Gpr22 | Enriched in dLGN | 1.241 | 0.00326 | 30.33 |
| ENSMUSG00000019927 | ubiquitin-conjugating enzyme E2D 1 [Source:MGI Symbol;Acc:MGI:2384911] | Ube2d1 | Enriched in dLGN | 1.24 | 0.0002044 | 71.7 |
| ENSMUSG00000038174 | hyccin PI4KA lipid kinase complex subunit 2 [Source:MGI Symbol;Acc:MGI:1098784] | Hycc2 | Enriched in dLGN | 1.239 | 0.0005698 | 246 |
| ENSMUSG00000015536 | molybdenum cofactor synthesis 2 [Source:MGI Symbol;Acc:MGI:1336894] | Mocs2 | Enriched in dLGN | 1.238 | 1.468E-08 | 73.92 |
| ENSMUSG00000054428 | ATPase inhibitory factor 1 [Source:MGI Symbol;Acc:MGI:1196457] | Atpif1 | Enriched in dLGN | 1.238 | 0.00002893 | 829.4 |
| ENSMUSG00000025980 | heat shock protein 1 (chaperonin) [Source:MGI Symbol;Acc:MGI:96242] | Hspd1 | Enriched in dLGN | 1.237 | 0.000004383 | 264 |
| ENSMUSG00000027602 | microtubule-associated protein 1 light chain 3 alpha [Source:MGI Symbol;Acc:MGI:1915661] | Map1lc3a | Enriched in dLGN | 1.234 | 0.00000268 | 233.2 |
| ENSMUSG00000028099 | polymerase (RNA) III (DNA directed) polypeptide C [Source:MGI Symbol;Acc:MGI:1921664] | Polr3c | Enriched in dLGN | 1.232 | 3.738E-07 | 65.81 |
| ENSMUSG00000019868 | vesicle (multivesicular body) trafficking 1 [Source:MGI Symbol;Acc:MGI:1913451] | Vta1 | Enriched in dLGN | 1.23 | 0.001853 | 69.26 |
| ENSMUSG00000022450 | NADH:ubiquinone oxidoreductase subunit A6 [Source:MGI Symbol;Acc:MGI:1914380] | Ndufa6 | Enriched in dLGN | 1.229 | 0.00000198 | 126.1 |
| ENSMUSG00000040462 | amplified in osteosarcoma [Source:MGI Symbol;Acc:MGI:1924301] | Os9 | Enriched in dLGN | 1.228 | 9.768E-08 | 698.2 |
| ENSMUSG00000027011 | ubiquitin-conjugating enzyme E2E 3 [Source:MGI Symbol;Acc:MGI:107412] | Ube2e3 | Enriched in dLGN | 1.228 | 3.069E-07 | 305.5 |
| ENSMUSG00000063882 | ubiquinol-cytochrome c reductase hinge protein [Source:MGI Symbol;Acc:MGI:1913826] | Uqcrh | Enriched in dLGN | 1.228 | 5.815E-07 | 256.5 |
| ENSMUSG00000027894 | solute carrier family 6 (neurotransmitter transporter), member 17 [Source:MGI Symbol;Acc:MGI:2442535] | Slc6a17 | Enriched in dLGN | 1.227 | 0.000001388 | 1006 |
| ENSMUSG00000026750 | proteasome (prosome, macropain) subunit, beta type 7 [Source:MGI Symbol;Acc:MGI:107637] | Psmb7 | Enriched in dLGN | 1.227 | 0.00001638 | 169.1 |
| ENSMUSG00000031167 | RNA binding motif (RNP1, RRM) protein 3 [Source:MGI Symbol;Acc:MGI:1099460] | Rbm3 | Enriched in dLGN | 1.227 | 0.001129 | 106.4 |
| ENSMUSG00000031950 | gamma-aminobutyric acid (GABA) A receptor-associated protein-like 2 [Source:MGI Symbol;Acc:MGI:1890602] | Gabarapl2 | Enriched in dLGN | 1.226 | 0.00000298 | 147.2 |
| ENSMUSG00000026664 | phytanoyl-CoA hydroxylase [Source:MGI Symbol;Acc:MGI:891978] | Phyh | Enriched in dLGN | 1.226 | 0.000006324 | 93.1 |
| ENSMUSG00000029634 | ring finger protein (C3H2C3 type) 6 [Source:MGI Symbol;Acc:MGI:1921382] | Rnf6 | Enriched in dLGN | 1.225 | 1.342E-08 | 280.9 |
| ENSMUSG00000050199 | leucine-rich repeat-containing G protein-coupled receptor 4 [Source:MGI Symbol;Acc:MGI:1891468] | Lgr4 | Enriched in dLGN | 1.225 | 6.881E-07 | 197.4 |
| ENSMUSG00000041084 | oligosaccharyltransferase complex subunit (non-catalytic) [Source:MGI Symbol;Acc:MGI:1913607] | Ostc | Enriched in dLGN | 1.224 | 0.0003823 | 62.9 |
| ENSMUSG00000024120 | leucine-rich PPR-motif containing [Source:MGI Symbol;Acc:MGI:1919666] | Lrpprc | Enriched in dLGN | 1.221 | 4.436E-10 | 294.4 |
| ENSMUSG00000030946 | phospholysine phosphohistidine inorganic pyrophosphate phosphatase [Source:MGI Symbol;Acc:MGI:1923679] | Lhpp | Enriched in dLGN | 1.221 | 0.0003133 | 34.61 |
| ENSMUSG00000021814 | annexin A7 [Source:MGI Symbol;Acc:MGI:88031] | Anxa7 | Enriched in dLGN | 1.22 | 0.0005742 | 56.22 |
| ENSMUSG00000021643 | small EDRK-rich factor 1 [Source:MGI Symbol;Acc:MGI:1337114] | Serf1 | Enriched in dLGN | 1.22 | 0.001762 | 40.29 |
| ENSMUSG00000028062 | late endosomal/lysosomal adaptor, MAPK and MTOR activator 2 [Source:MGI Symbol;Acc:MGI:1932697] | Lamtor2 | Enriched in dLGN | 1.212 | 0.0001944 | 47.8 |
| ENSMUSG00000058355 | ATP-binding cassette, sub-family E (OABP), member 1 [Source:MGI Symbol;Acc:MGI:1195458] | Abce1 | Enriched in dLGN | 1.212 | 0.0002409 | 169.9 |
| ENSMUSG00000064293 | contactin 4 [Source:MGI Symbol;Acc:MGI:1095737] | Cntn4 | Enriched in dLGN | 1.212 | 0.009818 | 402.8 |
| ENSMUSG00000024902 | mitochondrial ribosomal protein L11 [Source:MGI Symbol;Acc:MGI:2137215] | Mrpl11 | Enriched in dLGN | 1.211 | 0.00001046 | 54.62 |
| ENSMUSG00000091264 | small integral membrane protein 13 [Source:MGI Symbol;Acc:MGI:2652854] | Smim13 | Enriched in dLGN | 1.21 | 0.00002248 | 346 |
| ENSMUSG00000061474 | mitochondrial ribosomal protein S36 [Source:MGI Symbol;Acc:MGI:1913378] | Mrps36 | Enriched in dLGN | 1.21 | 0.01701 | 23.77 |
| ENSMUSG00000041763 | tripeptidyl peptidase II [Source:MGI Symbol;Acc:MGI:102724] | Tpp2 | Enriched in dLGN | 1.207 | 0.000003105 | 165.9 |
| ENSMUSG00000020736 | 5',3'-nucleotidase, cytosolic [Source:MGI Symbol;Acc:MGI:1354954] | Nt5c | Enriched in dLGN | 1.205 | 0.00001304 | 100.9 |
| ENSMUSG00000091722 | siah E3 ubiquitin protein ligase family member 3 [Source:MGI Symbol;Acc:MGI:2685758] | Siah3 | Enriched in dLGN | 1.204 | 0.00009923 | 62.95 |
| ENSMUSG00000021194 | chromogranin A [Source:MGI Symbol;Acc:MGI:88394] | Chga | Enriched in dLGN | 1.203 | 0.001184 | 812.4 |
| ENSMUSG00000020107 | anaphase promoting complex subunit 16 [Source:MGI Symbol;Acc:MGI:1289325] | Anapc16 | Enriched in dLGN | 1.203 | 0.002252 | 134.3 |
| ENSMUSG00000020171 | YEATS domain containing 4 [Source:MGI Symbol;Acc:MGI:1927224] | Yeats4 | Enriched in dLGN | 1.201 | 0.00008629 | 152.5 |
| ENSMUSG00000022789 | dynamin 1-like [Source:MGI Symbol;Acc:MGI:1921256] | Dnm1l | Enriched in dLGN | 1.2 | 0.000008899 | 380.5 |
| ENSMUSG00000004207 | prosaposin [Source:MGI Symbol;Acc:MGI:97783] | Psap | Enriched in dLGN | 1.199 | 0.0001136 | 1320 |
| ENSMUSG00000020333 | acyl-CoA synthetase long-chain family member 6 [Source:MGI Symbol;Acc:MGI:894291] | Acsl6 | Enriched in dLGN | 1.197 | 0.000000285 | 594.1 |
| ENSMUSG00000042763 | mannosidase, endo-alpha-like [Source:MGI Symbol;Acc:MGI:2684896] | Maneal | Enriched in dLGN | 1.197 | 0.00004216 | 148.8 |
| ENSMUSG00000006024 | N-ethylmaleimide sensitive fusion protein attachment protein alpha [Source:MGI Symbol;Acc:MGI:104563] | Napa | Enriched in dLGN | 1.196 | 8.848E-09 | 230.6 |
| ENSMUSG00000001630 | serine/threonine kinase 38 like [Source:MGI Symbol;Acc:MGI:1922250] | Stk38l | Enriched in dLGN | 1.195 | 0.00001432 | 71.94 |
| ENSMUSG00000032121 | transmembrane protein 218 [Source:MGI Symbol;Acc:MGI:1913529] | Tmem218 | Enriched in dLGN | 1.195 | 0.003219 | 27.24 |
| ENSMUSG00000021022 | protein phosphatase 2, regulatory subunit B'', gamma [Source:MGI Symbol;Acc:MGI:1930009] | Ppp2r3c | Enriched in dLGN | 1.194 | 0.001055 | 32.63 |
| ENSMUSG00000071528 | ATP synthase membrane subunit DAPIT [Source:MGI Symbol;Acc:MGI:1891435] | Atp5md | Enriched in dLGN | 1.193 | 0.005819 | 71.32 |
| ENSMUSG00000024537 | proteasome (prosome, macropain) assembly chaperone 2 [Source:MGI Symbol;Acc:MGI:1922901] | Psmg2 | Enriched in dLGN | 1.19 | 0.02173 | 28.6 |
| ENSMUSG00000022231 | sema domain, seven thrombospondin repeats (type 1 and type 1-like), transmembrane domain (TM) and short cytoplasmic domain, (semaphorin) 5A [Source:MGI Symbol;Acc:MGI:10 | Sema5a | Enriched in dLGN | 1.189 | 0.00006537 | 795.5 |
| ENSMUSG00000001366 | f-box protein 9 [Source:MGI Symbol;Acc:MGI:1918788] | Fbxo9 | Enriched in dLGN | 1.187 | 2.727E-16 | 320.2 |
| ENSMUSG00000026959 | glutamate receptor, ionotropic, NMDA1 (zeta 1) [Source:MGI Symbol;Acc:MGI:95819] | Grin1 | Enriched in dLGN | 1.186 | 0.000328 | 942.6 |
| ENSMUSG00000006717 | acyl-CoA thioesterase 13 [Source:MGI Symbol;Acc:MGI:1914084] | Acot13 | Enriched in dLGN | 1.181 | 0.002719 | 30.71 |
| ENSMUSG00000027963 | exostosin-like glycosyltransferase 2 [Source:MGI Symbol;Acc:MGI:1889574] | Extl2 | Enriched in dLGN | 1.179 | 0.000005122 | 151.6 |
| ENSMUSG00000018770 | ATP synthase, H+ transporting, mitochondrial F0 complex, subunit C3 (subunit 9) [Source:MGI Symbol;Acc:MGI:2442035] | Atp5g3 | Enriched in dLGN | 1.177 | 0.000009865 | 265.3 |
| ENSMUSG00000025531 | choroidermia (RAB escort protein 1) [Source:MGI Symbol;Acc:MGI:892979] | Chm | Enriched in dLGN | 1.176 | 0.0000179 | 90.38 |
| ENSMUSG00000004789 | dihydrolipoamide S-succinyltransferase (E2 component of 2-oxo-glutarate complex) [Source:MGI Symbol;Acc:MGI:1926170] | Dlst | Enriched in dLGN | 1.175 | 9.348E-11 | 409.2 |
| ENSMUSG00000028811 | tyrosyl-tRNA synthetase [Source:MGI Symbol;Acc:MGI:2147627] | Yars | Enriched in dLGN | 1.173 | 1.067E-17 | 315 |
| ENSMUSG00000056486 | chimerin 1 [Source:MGI Symbol;Acc:MGI:1915674] | Chn1 | Enriched in dLGN | 1.17 | 1.201E-11 | 504.4 |
| ENSMUSG00000052738 | succinate-CoA ligase, GDP-forming, alpha subunit [Source:MGI Symbol;Acc:MGI:1927234] | Suclg1 | Enriched in dLGN | 1.169 | 0.001074 | 138.6 |
| ENSMUSG00000078813 | leukocyte receptor cluster (LRC) member 1 [Source:MGI Symbol;Acc:MGI:1917007] | Leng1 | Enriched in dLGN | 1.168 | 0.0005103 | 41.3 |
| ENSMUSG00000005357 | solute carrier family 1 (high affinity aspartate/glutamate transporter), member 6 [Source:MGI Symbol;Acc:MGI:1096331] | Slc1a6 | Enriched in dLGN | 1.168 | 0.002127 | 52.8 |
| ENSMUSG00000058056 | palladin, cytoskeletal associated protein [Source:MGI Symbol;Acc:MGI:1919583] | Palld | Enriched in dLGN | 1.166 | 0.0003259 | 121.1 |
| ENSMUSG00000021178 | protease (prosome, macropain) 26S subunit, ATPase 1 [Source:MGI Symbol;Acc:MGI:106054] | Psmc1 | Enriched in dLGN | 1.165 | 0.00008673 | 275.2 |
| ENSMUSG00000010911 | APAF1 interacting protein [Source:MGI Symbol;Acc:MGI:1926788] | Apip | Enriched in dLGN | 1.162 | 0.01706 | 18.87 |
| ENSMUSG00000060981 | H4 clustered histone 8 [Source:MGI Symbol;Acc:MGI:2448427] | H4c8 | Enriched in dLGN | 1.162 | 0.01878 | 41.04 |
| ENSMUSG00000026646 | suppressor of variegation 3-9 2 [Source:MGI Symbol;Acc:MGI:1890396] | Suv39h2 | Enriched in dLGN | 1.161 | 0.0243 | 51.55 |
| ENSMUSG00000027170 | eukaryotic translation initiation factor 3, subunit M [Source:MGI Symbol;Acc:MGI:1351744] | Eif3m | Enriched in dLGN | 1.16 | 3.329E-07 | 71.19 |
| ENSMUSG00000055334 | snurportin 1 [Source:MGI Symbol;Acc:MGI:1913319] | Snupn | Enriched in dLGN | 1.16 | 0.001108 | 32.29 |
| ENSMUSG00000064202 | spermatogenesis associated 6 like [Source:MGI Symbol;Acc:MGI:1918036] | Spata6l | Enriched in dLGN | 1.159 | 0.005043 | 24.41 |
| ENSMUSG00000031753 | component of oligomeric golgi complex 4 [Source:MGI Symbol;Acc:MGI:2142808] | Cog4 | Enriched in dLGN | 1.158 | 2.119E-07 | 122.4 |
| ENSMUSG00000025630 | hypoxanthine guanine phosphoribosyl transferase [Source:MGI Symbol;Acc:MGI:96217] | Hprt | Enriched in dLGN | 1.156 | 6.899E-07 | 174.6 |
| ENSMUSG00000036309 | S-phase kinase-associated protein 1 [Source:MGI Symbol;Acc:MGI:103575] | Skp1 | Enriched in dLGN | 1.155 | 0.0001482 | 534.7 |
| ENSMUSG00000056185 | sorting nexin 32 [Source:MGI Symbol;Acc:MGI:2444704] | Snx32 | Enriched in dLGN | 1.154 | 0.004275 | 101.2 |
| ENSMUSG00000027293 | EH-domain containing 4 [Source:MGI Symbol;Acc:MGI:1919619] | Ehd4 | Enriched in dLGN | 1.153 | 0.0003192 | 74.05 |
| ENSMUSG00000027305 | NADH:ubiquinone oxidoreductase complex assembly factor 1 [Source:MGI Symbol;Acc:MGI:1916952] | Ndufaf1 | Enriched in dLGN | 1.152 | 0.00005503 | 32.64 |
| ENSMUSG00000021764 | NADH:ubiquinone oxidoreductase core subunit S4 [Source:MGI Symbol;Acc:MGI:1343135] | Ndufs4 | Enriched in dLGN | 1.152 | 0.00006924 | 153.5 |
| ENSMUSG00000079658 | elongin C [Source:MGI Symbol;Acc:MGI:1915173] | Eloc | Enriched in dLGN | 1.152 | 0.0001511 | 94.76 |
| ENSMUSG00000046834 | keratin 1 [Source:MGI Symbol;Acc:MGI:96698] | Krt1 | Enriched in dLGN | 1.151 | 0.009148 | 16.62 |
| ENSMUSG00000079184 | M-phase phosphoprotein 8 [Source:MGI Symbol;Acc:MGI:1922589] | Mphosph8 | Enriched in dLGN | 1.15 | 0.008481 | 685.9 |
| ENSMUSG00000017221 | proteasome (prosome, macropain) 26S subunit, non-ATPase, 3 [Source:MGI Symbol;Acc:MGI:98858] | Psmd3 | Enriched in dLGN | 1.149 | 0.000001033 | 193.7 |
| ENSMUSG00000036186 | divergent protein kinase domain 1B [Source:MGI Symbol;Acc:MGI:1927576] | Dipk1b | Enriched in dLGN | 1.149 | 0.001957 | 28.18 |
| ENSMUSG00000032551 | RIKEN cDNA 1110059G10 gene [Source:MGI Symbol;Acc:MGI:1913452] | 1110059G10Rik | Enriched in dLGN | 1.149 | 0.03243 | 11.31 |
| ENSMUSG00000041997 | tousled-like kinase 1 [Source:MGI Symbol;Acc:MGI:2441683] | Tlk1 | Enriched in dLGN | 1.146 | 0.000001354 | 318.7 |
| ENSMUSG00000020267 | histidine triad nucleotide binding protein 1 [Source:MGI Symbol;Acc:MGI:1321133] | Hint1 | Enriched in dLGN | 1.146 | 0.00003945 | 255.7 |
| ENSMUSG00000070953 | Rab9 effector protein with kelch motifs [Source:MGI Symbol;Acc:MGI:2139530] | Rabepk | Enriched in dLGN | 1.146 | 0.001258 | 41.33 |
| ENSMUSG00000041762 | G protein-coupled receptor 155 [Source:MGI Symbol;Acc:MGI:1915776] | Gpr155 | Enriched in dLGN | 1.145 | 0.00006362 | 55.46 |
| ENSMUSG00000032555 | topoisomerase (DNA) II binding protein 1 [Source:MGI Symbol;Acc:MGI:1920018] | Topbp1 | Enriched in dLGN | 1.144 | 9.446E-07 | 78.01 |
| ENSMUSG00000063873 | solute carrier family 24 (sodium/potassium/calcium exchanger), member 3 [Source:MGI Symbol;Acc:MGI:2137513] | Slc24a3 | Enriched in dLGN | 1.144 | 0.00003322 | 1240 |
| ENSMUSG00000035885 | cytochrome c oxidase subunit 8A [Source:MGI Symbol;Acc:MGI:105959] | Cox8a | Enriched in dLGN | 1.143 | 0.00002359 | 292.6 |
| ENSMUSG00000036503 | ring finger protein 13 [Source:MGI Symbol;Acc:MGI:1346341] | Rnf13 | Enriched in dLGN | 1.143 | 0.0006441 | 102.4 |
| ENSMUSG00000021775 | nuclear receptor subfamily 1, group D, member 2 [Source:MGI Symbol;Acc:MGI:2449205] | Nr1d2 | Enriched in dLGN | 1.142 | 0.00002187 | 245.4 |
| ENSMUSG00000021816 | protein phosphatase 3, catalytic subunit, beta isoform [Source:MGI Symbol;Acc:MGI:107163] | Ppp3cb | Enriched in dLGN | 1.141 | 1.149E-11 | 533.5 |
| ENSMUSG00000027502 | replication termination factor 2 [Source:MGI Symbol;Acc:MGI:1913654] | Rtf2 | Enriched in dLGN | 1.14 | 2.845E-08 | 124.2 |
| ENSMUSG00000015759 | cornichon family AMPA receptor auxiliary protein 1 [Source:MGI Symbol;Acc:MGI:1277202] | Cnih1 | Enriched in dLGN | 1.14 | 0.0001148 | 68.86 |
| ENSMUSG00000025817 | nudix (nucleoside diphosphate linked moiety X)-type motif 5 [Source:MGI Symbol;Acc:MGI:1858232] | Nudt5 | Enriched in dLGN | 1.139 | 0.0002699 | 35.8 |
| ENSMUSG00000032604 | glutaminyl-tRNA synthetase [Source:MGI Symbol;Acc:MGI:1915851] | Qars | Enriched in dLGN | 1.138 | 3.112E-07 | 152.8 |

| | | | | | | |
|---|---|---|---|---|---|---|
| ENSMUSG00000039633 | LON peptidase N-terminal domain and ring finger 1 [Source:MGI Symbol;Acc:MGI:3609241] | Lonrf1 | Enriched in dLGN | 1.138 | 0.000002524 | 125.4 |
| ENSMUSG00000028078 | doublecortin-like kinase 2 [Source:MGI Symbol;Acc:MGI:1918012] | Dclk2 | Enriched in dLGN | 1.138 | 0.00002815 | 340.9 |
| ENSMUSG00000019478 | RAB4A, member RAS oncogene family [Source:MGI Symbol;Acc:MGI:105069] | Rab4a | Enriched in dLGN | 1.137 | 2.249E-07 | 91.57 |
| ENSMUSG00000005656 | sorting nexin 6 [Source:MGI Symbol;Acc:MGI:1919433] | Snx6 | Enriched in dLGN | 1.136 | 0.00001794 | 77.18 |
| ENSMUSG00000020677 | DExD box helicase 52 [Source:MGI Symbol;Acc:MGI:1925644] | Ddx52 | Enriched in dLGN | 1.136 | 0.0003798 | 51.64 |
| ENSMUSG00000043639 | RNA binding motif protein 20 [Source:MGI Symbol;Acc:MGI:1920963] | Rbm20 | Enriched in dLGN | 1.133 | 0.0373 | 125.3 |
| ENSMUSG00000050856 | ATP synthase, H+ transporting, mitochondrial F1F0 complex, subunit E [Source:MGI Symbol;Acc:MGI:106636] | Atp5k | Enriched in dLGN | 1.132 | 0.0069 | 126.9 |
| ENSMUSG00000021606 | NADH:ubiquinone oxidoreductase core subunit S6 [Source:MGI Symbol;Acc:MGI:107932] | Ndufs6 | Enriched in dLGN | 1.131 | 0.000004931 | 70.27 |
| ENSMUSG00000032883 | acyl-CoA synthetase long-chain family member 3 [Source:MGI Symbol;Acc:MGI:1921455] | Acsl3 | Enriched in dLGN | 1.131 | 0.0000539 | 522.5 |
| ENSMUSG00000028013 | pyrophosphatase (inorganic) 2 [Source:MGI Symbol;Acc:MGI:1922026] | Ppa2 | Enriched in dLGN | 1.13 | 0.00006492 | 141.1 |
| ENSMUSG00000071662 | polymerase (RNA) II (DNA directed) polypeptide G [Source:MGI Symbol;Acc:MGI:1914960] | Polr2g | Enriched in dLGN | 1.129 | 0.0001455 | 40.09 |
| ENSMUSG00000044221 | G-rich RNA sequence binding factor 1 [Source:MGI Symbol;Acc:MGI:106479] | Grsf1 | Enriched in dLGN | 1.128 | 5.821E-07 | 242.3 |
| ENSMUSG00000042032 | methionine adenosyltransferase II, beta [Source:MGI Symbol;Acc:MGI:1913667] | Mat2b | Enriched in dLGN | 1.127 | 0.000001867 | 58.43 |
| ENSMUSG00000022094 | solute carrier family 39 (zinc transporter), member 14 [Source:MGI Symbol;Acc:MGI:2384851] | Slc39a14 | Enriched in dLGN | 1.127 | 0.000002656 | 73.63 |
| ENSMUSG00000025757 | heat shock protein 4 like [Source:MGI Symbol;Acc:MGI:107422] | Hspa4l | Enriched in dLGN | 1.127 | 0.00006092 | 545.2 |
| ENSMUSG00000030738 | eukaryotic translation initiation factor 3, subunit C [Source:MGI Symbol;Acc:MGI:1926966] | Eif3c | Enriched in dLGN | 1.127 | 0.00006844 | 682.4 |
| ENSMUSG00000005981 | TNF receptor-associated protein 1 [Source:MGI Symbol;Acc:MGI:1915265] | Trap1 | Enriched in dLGN | 1.126 | 0.001438 | 166.2 |
| ENSMUSG00000020629 | acireductone dioxygenase 1 [Source:MGI Symbol;Acc:MGI:2144929] | Adi1 | Enriched in dLGN | 1.125 | 0.0002953 | 28.33 |
| ENSMUSG00000030652 | demethyl-Q 7 [Source:MGI Symbol;Acc:MGI:107207] | Coq7 | Enriched in dLGN | 1.125 | 0.0004593 | 54.52 |
| ENSMUSG00000021807 | RNA transcription, translation and transport factor [Source:MGI Symbol;Acc:MGI:1915295] | Rtraf | Enriched in dLGN | 1.122 | 0.0002092 | 75.17 |
| ENSMUSG00000056895 | H2B clustered histone 27 [Source:MGI Symbol;Acc:MGI:1925553] | H2bc27 | Enriched in dLGN | 1.122 | 0.008101 | 37.86 |
| ENSMUSG00000042354 | guanine nucleotide binding protein-like 3 (nucleolar) [Source:MGI Symbol;Acc:MGI:1353651] | Gnl3 | Enriched in dLGN | 1.121 | 0.001883 | 45.05 |
| ENSMUSG00000011254 | tRNA-histidine guanylyltransferase 1-like (S. cerevisiae) [Source:MGI Symbol;Acc:MGI:1913878] | Thg1l | Enriched in dLGN | 1.121 | 0.02225 | 24.76 |
| ENSMUSG00000031168 | phenylalkylamine Ca2+ antagonist (emopamil) binding protein [Source:MGI Symbol;Acc:MGI:107822] | Ebp | Enriched in dLGN | 1.121 | 0.03556 | 18.75 |
| ENSMUSG00000053192 | myeloid/lymphoid or mixed-lineage leukemia; translocated to, 11 [Source:MGI Symbol;Acc:MGI:1929671] | Mllt11 | Enriched in dLGN | 1.119 | 0.000009915 | 499.1 |
| ENSMUSG00000028138 | alcohol dehydrogenase 5 (class III), chi polypeptide [Source:MGI Symbol;Acc:MGI:87929] | Adh5 | Enriched in dLGN | 1.119 | 0.0003685 | 68.21 |
| ENSMUSG00000036510 | cadherin 8 [Source:MGI Symbol;Acc:MGI:107434] | Cdh8 | Enriched in dLGN | 1.119 | 0.006048 | 272 |
| ENSMUSG00000041360 | pumilio RNA-binding family member 3 [Source:MGI Symbol;Acc:MGI:106253] | Pum3 | Enriched in dLGN | 1.118 | 0.00001559 | 123.2 |
| ENSMUSG00000074736 | synapse differentiation inducing 1 [Source:MGI Symbol;Acc:MGI:3702158] | Syndig1 | Enriched in dLGN | 1.118 | 0.009712 | 50.55 |
| ENSMUSG00000056222 | sparc/osteonectin, cwcv and kazal-like domains proteoglycan 1 [Source:MGI Symbol;Acc:MGI:105371] | Spock1 | Enriched in dLGN | 1.116 | 0.001104 | 1204 |
| ENSMUSG00000051154 | COMM domain containing 3 [Source:MGI Symbol;Acc:MGI:88218] | Commd3 | Enriched in dLGN | 1.115 | 0.001393 | 62.85 |
| ENSMUSG00000022564 | glutamate receptor, ionotropic, N-methyl D-aspartate-associated protein 1 (glutamate binding) [Source:MGI Symbol;Acc:MGI:1913418] | Grina | Enriched in dLGN | 1.114 | 0.0000101 | 437.4 |
| ENSMUSG00000033319 | fem 1 homolog c [Source:MGI Symbol;Acc:MGI:2444737] | Fem1c | Enriched in dLGN | 1.114 | 0.00005928 | 163.3 |
| ENSMUSG00000037894 | H2A.Z variant histone 1 [Source:MGI Symbol;Acc:MGI:1888388] | H2az1 | Enriched in dLGN | 1.114 | 0.0005867 | 59.57 |
| ENSMUSG00000026087 | mitochondrial ribosomal protein L30 [Source:MGI Symbol;Acc:MGI:1333820] | Mrpl30 | Enriched in dLGN | 1.113 | 0.0001627 | 45.34 |
| ENSMUSG00000020537 | developmentally regulated GTP binding protein 2 [Source:MGI Symbol;Acc:MGI:1342307] | Drg2 | Enriched in dLGN | 1.112 | 0.0009519 | 72 |
| ENSMUSG00000029036 | ATPase family, AAA domain containing 3A [Source:MGI Symbol;Acc:MGI:1919214] | Atad3a | Enriched in dLGN | 1.11 | 0.00000342 | 104.5 |
| ENSMUSG00000025762 | La ribonucleoprotein 1B [Source:MGI Symbol;Acc:MGI:1914604] | Larp1b | Enriched in dLGN | 1.108 | 0.0002995 | 100 |
| ENSMUSG00000027637 | RAB5 interacting factor [Source:MGI Symbol;Acc:MGI:1914638] | Rab5if | Enriched in dLGN | 1.106 | 0.000002027 | 59.45 |
| ENSMUSG00000029432 | nipsnap homolog 2 [Source:MGI Symbol;Acc:MGI:1278343] | Nipsnap2 | Enriched in dLGN | 1.105 | 0.0002754 | 127.2 |
| ENSMUSG00000042807 | HECT, C2 and WW domain containing E3 ubiquitin protein ligase 2 [Source:MGI Symbol;Acc:MGI:2685817] | Hecw2 | Enriched in dLGN | 1.105 | 0.001334 | 320.7 |
| ENSMUSG00000040370 | electron transfer flavoprotein regulatory factor 1 [Source:MGI Symbol;Acc:MGI:1914886] | Etfrf1 | Enriched in dLGN | 1.104 | 0.007954 | 29.52 |
| ENSMUSG00000025487 | proteasome (prosome, macropain) 26S subunit, non-ATPase, 13 [Source:MGI Symbol;Acc:MGI:1345192] | Psmd13 | Enriched in dLGN | 1.103 | 3.405E-07 | 179.8 |
| ENSMUSG00000096257 | coiled-coil glutamate-rich protein 2 [Source:MGI Symbol;Acc:MGI:3645242] | Ccer2 | Enriched in dLGN | 1.103 | 0.03376 | 12.8 |
| ENSMUSG00000024359 | heat shock protein 9 [Source:MGI Symbol;Acc:MGI:96245] | Hspa9 | Enriched in dLGN | 1.102 | 0.0007216 | 574.8 |
| ENSMUSG00000031774 | proteasome activator subunit 3 interacting protein 1 [Source:MGI Symbol;Acc:MGI:1919637] | Psme3ip1 | Enriched in dLGN | 1.101 | 4.509E-09 | 123.3 |
| ENSMUSG00000031556 | TM2 domain containing 2 [Source:MGI Symbol;Acc:MGI:1916992] | Tm2d2 | Enriched in dLGN | 1.101 | 0.001687 | 62.81 |
| ENSMUSG00000045996 | polymerase (RNA) II (DNA directed) polypeptide K [Source:MGI Symbol;Acc:MGI:102725] | Polr2k | Enriched in dLGN | 1.1 | 0.04642 | 12.72 |
| ENSMUSG00000004980 | heterogeneous nuclear ribonucleoprotein A2/B1 [Source:MGI Symbol;Acc:MGI:104819] | Hnrnpa2b1 | Enriched in dLGN | 1.099 | 2.229E-07 | 930.3 |
| ENSMUSG00000026825 | dynamin 1 [Source:MGI Symbol;Acc:MGI:107384] | Dnm1 | Enriched in dLGN | 1.099 | 0.00002279 | 2096 |
| ENSMUSG00000025940 | transmembrane protein 70 [Source:MGI Symbol;Acc:MGI:1915068] | Tmem70 | Enriched in dLGN | 1.097 | 0.00007771 | 50.14 |
| ENSMUSG00000041278 | tetratricopeptide repeat domain 1 [Source:MGI Symbol;Acc:MGI:1914077] | Ttc1 | Enriched in dLGN | 1.097 | 0.000169 | 100.1 |
| ENSMUSG00000021193 | pitrilysin metallepetidase 1 [Source:MGI Symbol;Acc:MGI:1916867] | Pitrm1 | Enriched in dLGN | 1.096 | 0.00002754 | 115.9 |
| ENSMUSG00000060904 | ADP-ribosylation factor-like 1 [Source:MGI Symbol;Acc:MGI:99436] | Arl1 | Enriched in dLGN | 1.096 | 0.0002313 | 131.5 |
| ENSMUSG00000020514 | mitochondrial ribosomal protein L22 [Source:MGI Symbol;Acc:MGI:1333794] | Mrpl22 | Enriched in dLGN | 1.096 | 0.009291 | 23.4 |
| ENSMUSG00000006315 | transmembrane protein 147 [Source:MGI Symbol;Acc:MGI:1915011] | Tmem147 | Enriched in dLGN | 1.093 | 0.0006834 | 46.25 |
| ENSMUSG00000007653 | gamma-aminobutyric acid (GABA) A receptor, subunit beta 2 [Source:MGI Symbol;Acc:MGI:95620] | Gabrb2 | Enriched in dLGN | 1.093 | 0.005272 | 322.6 |
| ENSMUSG00000048376 | coagulation factor II (thrombin) receptor [Source:MGI Symbol;Acc:MGI:101802] | F2r | Enriched in dLGN | 1.093 | 0.01561 | 37.83 |
| ENSMUSG00000039480 | 5'-nucleotidase domain containing 1 [Source:MGI Symbol;Acc:MGI:2442446] | Nt5dc1 | Enriched in dLGN | 1.093 | 0.02271 | 84.48 |
| ENSMUSG00000018882 | mitochondrial ribosomal protein L45 [Source:MGI Symbol;Acc:MGI:1914286] | Mrpl45 | Enriched in dLGN | 1.092 | 0.0000561 | 64.61 |
| ENSMUSG00000034640 | TCDD-inducible poly(ADP-ribose) polymerase [Source:MGI Symbol;Acc:MGI:2159210] | Tiparp | Enriched in dLGN | 1.09 | 0.0002497 | 38.05 |
| ENSMUSG00000033793 | ATPase, H+ transporting, lysosomal V1 subunit H [Source:MGI Symbol;Acc:MGI:1914864] | Atp6v1h | Enriched in dLGN | 1.09 | 0.0008221 | 178.8 |
| ENSMUSG00000024740 | damage specific DNA binding protein 1 [Source:MGI Symbol;Acc:MGI:1202384] | Ddb1 | Enriched in dLGN | 1.089 | 0.00009172 | 974.8 |
| ENSMUSG00000035048 | anaphase promoting complex subunit 13 [Source:MGI Symbol;Acc:MGI:1916260] | Anapc13 | Enriched in dLGN | 1.089 | 0.0004492 | 49.47 |
| ENSMUSG00000027263 | tubulin, gamma complex associated protein 4 [Source:MGI Symbol;Acc:MGI:1196293] | Tubgcp4 | Enriched in dLGN | 1.088 | 0.00004701 | 80.84 |
| ENSMUSG00000000441 | v-raf-leukemia viral oncogene 1 [Source:MGI Symbol;Acc:MGI:97847] | Raf1 | Enriched in dLGN | 1.087 | 0.00006871 | 265.3 |
| ENSMUSG00000075000 | nuclear receptor binding factor 2 [Source:MGI Symbol;Acc:MGI:1354950] | Nrbf2 | Enriched in dLGN | 1.087 | 0.0001613 | 35.46 |
| ENSMUSG00000000776 | polymerase (RNA) III (DNA directed) polypeptide D [Source:MGI Symbol;Acc:MGI:1914315] | Polr3d | Enriched in dLGN | 1.086 | 0.000001514 | 108.5 |
| ENSMUSG00000027274 | McKusick-Kaufman syndrome [Source:MGI Symbol;Acc:MGI:1891836] | Mkks | Enriched in dLGN | 1.086 | 0.01239 | 21.92 |
| ENSMUSG00000071369 | mitogen-activated protein kinase kinase kinase 5 [Source:MGI Symbol;Acc:MGI:1346876] | Map3k5 | Enriched in dLGN | 1.085 | 0.00003407 | 236 |
| ENSMUSG00000021660 | basic transcription factor 3 [Source:MGI Symbol;Acc:MGI:1202875] | Btf3 | Enriched in dLGN | 1.085 | 0.00006626 | 71.1 |
| ENSMUSG00000009863 | succinate dehydrogenase complex, subunit B, iron sulfur (Ip) [Source:MGI Symbol;Acc:MGI:1914930] | Sdhb | Enriched in dLGN | 1.085 | 0.0001103 | 134.3 |
| ENSMUSG00000030417 | programmed cell death 5 [Source:MGI Symbol;Acc:MGI:1913538] | Pdcd5 | Enriched in dLGN | 1.084 | 0.0004976 | 70.53 |
| ENSMUSG00000027615 | HPS3, biogenesis of lysosomal organelles complex 2 subunit 1 [Source:MGI Symbol;Acc:MGI:2153839] | Hps3 | Enriched in dLGN | 1.083 | 0.0002919 | 108.7 |
| ENSMUSG00000034259 | exosome component 4 [Source:MGI Symbol;Acc:MGI:1923576] | Exosc4 | Enriched in dLGN | 1.083 | 0.007604 | 28.46 |
| ENSMUSG00000027163 | COMM domain containing 9 [Source:MGI Symbol;Acc:MGI:1923751] | Commd9 | Enriched in dLGN | 1.078 | 0.0001471 | 36.63 |
| ENSMUSG00000032046 | abhydrolase domain containing 12 [Source:MGI Symbol;Acc:MGI:1923442] | Abhd12 | Enriched in dLGN | 1.077 | 3.808E-08 | 313.9 |
| ENSMUSG00000015092 | endothelial differentiation-related factor 1 [Source:MGI Symbol;Acc:MGI:1891227] | Edf1 | Enriched in dLGN | 1.077 | 0.00001335 | 190.3 |
| ENSMUSG00000029447 | chaperonin containing Tcp1, subunit 6a (zeta) [Source:MGI Symbol;Acc:MGI:107943] | Cct6a | Enriched in dLGN | 1.077 | 0.00002951 | 356.4 |
| ENSMUSG00000033845 | mitochondrial ribosomal protein L15 [Source:MGI Symbol;Acc:MGI:1351639] | Mrpl15 | Enriched in dLGN | 1.076 | 0.000009262 | 86.2 |
| ENSMUSG00000023110 | protein arginine N-methyltransferase 5 [Source:MGI Symbol;Acc:MGI:1351645] | Prmt5 | Enriched in dLGN | 1.075 | 0.00002537 | 156.1 |
| ENSMUSG00000041670 | regulating synaptic membrane exocytosis 1 [Source:MGI Symbol;Acc:MGI:2152971] | Rims1 | Enriched in dLGN | 1.075 | 0.00007148 | 1881 |
| ENSMUSG00000028656 | CAP, adenylate cyclase-associated protein 1 (yeast) [Source:MGI Symbol;Acc:MGI:88262] | Cap1 | Enriched in dLGN | 1.074 | 0.001499 | 243.2 |
| ENSMUSG00000021939 | cathepsin B [Source:MGI Symbol;Acc:MGI:88561] | Ctsb | Enriched in dLGN | 1.074 | 0.002849 | 510.6 |
| ENSMUSG00000037531 | mitochondrial ribosomal protein L47 [Source:MGI Symbol;Acc:MGI:1921850] | Mrpl47 | Enriched in dLGN | 1.074 | 0.005717 | 22.85 |
| ENSMUSG00000027206 | COP9 signalosome subunit 2 [Source:MGI Symbol;Acc:MGI:1330276] | Cops2 | Enriched in dLGN | 1.072 | 0.000005029 | 139.2 |
| ENSMUSG00000020697 | ligase III, DNA, ATP-dependent [Source:MGI Symbol;Acc:MGI:109152] | Lig3 | Enriched in dLGN | 1.071 | 0.000001504 | 109.2 |
| ENSMUSG00000047547 | clathrin, light polypeptide (Lcb) [Source:MGI Symbol;Acc:MGI:1921575] | Cltb | Enriched in dLGN | 1.07 | 4.998E-12 | 896 |
| ENSMUSG00000004798 | unc-51 like kinase 2 [Source:MGI Symbol;Acc:MGI:1352758] | Ulk2 | Enriched in dLGN | 1.069 | 3.503E-07 | 388.4 |
| ENSMUSG00000013160 | ATPase, H+ transporting, lysosomal V0 subunit D1 [Source:MGI Symbol;Acc:MGI:1201778] | Atp6v0d1 | Enriched in dLGN | 1.069 | 0.00004909 | 268.5 |
| ENSMUSG00000029657 | heat shock 105kDa/110kDa protein 1 [Source:MGI Symbol;Acc:MGI:105053] | Hsph1 | Enriched in dLGN | 1.067 | 0.000399 | 555.2 |
| ENSMUSG00000027002 | NCK-associated protein 1 [Source:MGI Symbol;Acc:MGI:1355333] | Nckap1 | Enriched in dLGN | 1.065 | 0.00000346 | 682.6 |
| ENSMUSG00000028797 | transmembrane protein 234 [Source:MGI Symbol;Acc:MGI:1924049] | Tmem234 | Enriched in dLGN | 1.063 | 0.000001089 | 162.2 |
| ENSMUSG00000053329 | glutamine amidotransferase like class 1 domain containing 3A [Source:MGI Symbol;Acc:MGI:1351861] | Gatd3a | Enriched in dLGN | 1.062 | 0.000005128 | 113.5 |
| ENSMUSG00000041020 | MAP7 domain containing 2 [Source:MGI Symbol;Acc:MGI:1917474] | Map7d2 | Enriched in dLGN | 1.062 | 0.00006973 | 736.5 |
| ENSMUSG00000026767 | MINDY lysine 48 deubiquitinase 3 [Source:MGI Symbol;Acc:MGI:1914210] | Mindy3 | Enriched in dLGN | 1.061 | 0.0009294 | 66.58 |
| ENSMUSG00000042487 | Leo1, Paf1/RNA polymerase II complex component [Source:MGI Symbol;Acc:MGI:2685031] | Leo1 | Enriched in dLGN | 1.06 | 4.594E-07 | 228.7 |
| ENSMUSG00000033781 | ankyrin repeat and SOCS box-containing 13 [Source:MGI Symbol;Acc:MGI:2145525] | Asb13 | Enriched in dLGN | 1.06 | 0.0001298 | 93.68 |
| ENSMUSG00000021196 | phosphofructokinase, platelet [Source:MGI Symbol;Acc:MGI:1891833] | Pfkp | Enriched in dLGN | 1.059 | 0.01488 | 1055 |
| ENSMUSG00000039197 | adenosine kinase [Source:MGI Symbol;Acc:MGI:87930] | Adk | Enriched in dLGN | 1.058 | 0.01644 | 98.44 |
| ENSMUSG00000026000 | LanC (bacterial lantibiotic synthetase component C)-like 1 [Source:MGI Symbol;Acc:MGI:1336997] | Lancl1 | Enriched in dLGN | 1.057 | 0.000001002 | 119.7 |
| ENSMUSG00000033460 | armadillo repeat containing, X-linked 1 [Source:MGI Symbol;Acc:MGI:1925498] | Armcx1 | Enriched in dLGN | 1.056 | 0.00001036 | 200.9 |
| ENSMUSG00000042535 | GTP binding protein 1 [Source:MGI Symbol;Acc:MGI:109443] | Gtpbp1 | Enriched in dLGN | 1.054 | 2.167E-07 | 209.5 |
| ENSMUSG00000028419 | charged multivesicular body protein 5 [Source:MGI Symbol;Acc:MGI:1924209] | Chmp5 | Enriched in dLGN | 1.054 | 0.0002447 | 102.8 |
| ENSMUSG00000022110 | succinate-Coenzyme A ligase, ADP-forming, beta subunit [Source:MGI Symbol;Acc:MGI:1306775] | Sucla2 | Enriched in dLGN | 1.054 | 0.003858 | 256.6 |
| ENSMUSG00000060882 | potassium voltage-gated channel, Shal-related family, member 2 [Source:MGI Symbol;Acc:MGI:102663] | Kcnd2 | Enriched in dLGN | 1.054 | 0.02009 | 981.4 |
| ENSMUSG00000042682 | selenoprotein K [Source:MGI Symbol;Acc:MGI:1931466] | Selenok | Enriched in dLGN | 1.053 | 0.005697 | 254.3 |
| ENSMUSG00000027534 | sorting nexin 16 [Source:MGI Symbol;Acc:MGI:1921968] | Snx16 | Enriched in dLGN | 1.052 | 0.003589 | 74.38 |
| ENSMUSG00000028790 | KH domain containing, RNA binding, signal transduction associated 1 [Source:MGI Symbol;Acc:MGI:893579] | Khdrbs1 | Enriched in dLGN | 1.049 | 0.000003952 | 341 |
| ENSMUSG00000025059 | glycerol kinase [Source:MGI Symbol;Acc:MGI:106594] | Gk | Enriched in dLGN | 1.049 | 0.003784 | 44.02 |
| ENSMUSG00000024981 | acyl-CoA synthetase long-chain family member 5 [Source:MGI Symbol;Acc:MGI:1919129] | Acsl5 | Enriched in dLGN | 1.048 | 0.000497 | 65.86 |
| ENSMUSG00000019977 | Hbs1-like (S. cerevisiae) [Source:MGI Symbol;Acc:MGI:1891704] | Hbs1l | Enriched in dLGN | 1.047 | 0.00006874 | 167.3 |
| ENSMUSG00000054321 | TATA-box binding protein associated factor 4b [Source:MGI Symbol;Acc:MGI:2152345] | Taf4b | Enriched in dLGN | 1.047 | 0.03948 | 23.21 |
| ENSMUSG00000022634 | YY1 associated factor 2 [Source:MGI Symbol;Acc:MGI:1914307] | Yaf2 | Enriched in dLGN | 1.046 | 0.000001692 | 232.8 |
| ENSMUSG00000034557 | zinc finger, FYVE domain containing 9 [Source:MGI Symbol;Acc:MGI:2652838] | Zfyve9 | Enriched in dLGN | 1.046 | 0.005189 | 559.4 |
| ENSMUSG00000015672 | mitochondrial ribosomal protein L32 [Source:MGI Symbol;Acc:MGI:2137226] | Mrpl32 | Enriched in dLGN | 1.046 | 0.01942 | 27.23 |
| ENSMUSG00000025892 | glutamate receptor, ionotropic, AMPA4 (alpha 4) [Source:MGI Symbol;Acc:MGI:95811] | Gria4 | Enriched in dLGN | 1.045 | 0.002949 | 894.8 |
| ENSMUSG00000035790 | centrosomal protein 19 [Source:MGI Symbol;Acc:MGI:1914244] | Cep19 | Enriched in dLGN | 1.044 | 3.499E-08 | 80.5 |
| ENSMUSG00000031673 | cadherin 11 [Source:MGI Symbol;Acc:MGI:99217] | Cdh11 | Enriched in dLGN | 1.044 | 0.002145 | 313.1 |
| ENSMUSG00000009630 | protein phosphatase 2 (formerly 2A), catalytic subunit, beta isoform [Source:MGI Symbol;Acc:MGI:1321161] | Ppp2cb | Enriched in dLGN | 1.043 | 4.636E-08 | 209.1 |
| ENSMUSG00000013736 | tRNA nucleotidyl transferase, CCA-adding, 1 [Source:MGI Symbol;Acc:MGI:1917297] | Trnt1 | Enriched in dLGN | 1.043 | 0.00009456 | 70.42 |
| ENSMUSG00000016831 | TOX high mobility group box family member 4 [Source:MGI Symbol;Acc:MGI:1915389] | Tox4 | Enriched in dLGN | 1.041 | 5.96E-08 | 111.7 |
| ENSMUSG00000031948 | lysyl-tRNA synthetase [Source:MGI Symbol;Acc:MGI:1934754] | Kars | Enriched in dLGN | 1.041 | 0.0001603 | 171 |
| ENSMUSG00000021218 | guanosine diphosphate (GDP) dissociation inhibitor 2 [Source:MGI Symbol;Acc:MGI:99845] | Gdi2 | Enriched in dLGN | 1.04 | 0.00004338 | 530.5 |
| ENSMUSG00000032192 | guanine nucleotide binding protein (G protein), beta 5 [Source:MGI Symbol;Acc:MGI:101848] | Gnb5 | Enriched in dLGN | 1.04 | 0.00006528 | 441.8 |
| ENSMUSG00000024487 | Yip1 domain family, member 5 [Source:MGI Symbol;Acc:MGI:1914430] | Yipf5 | Enriched in dLGN | 1.04 | 0.0007961 | 59.88 |
| ENSMUSG00000022962 | phosphoribosylglycinamide formyltransferase [Source:MGI Symbol;Acc:MGI:95654] | Gart | Enriched in dLGN | 1.039 | 0.00002121 | 92.71 |
| ENSMUSG00000022855 | SUMO/sentrin specific peptidase 2 [Source:MGI Symbol;Acc:MGI:1923076] | Senp2 | Enriched in dLGN | 1.039 | 0.00009299 | 371 |

| | | | | | | |
|---|---|---|---|---|---|---|
| ENSMUSG00000003812 | deoxyribonuclease II alpha [Source:MGI Symbol;Acc:MGI:1329019] | Dnase2a | Enriched in dLGN | 1.039 | 0.007786 | 35.72 |
| ENSMUSG00000042111 | coiled-coil domain containing 115 [Source:MGI Symbol;Acc:MGI:1916918] | Ccdc115 | Enriched in dLGN | 1.038 | 0.0012 | 46.6 |
| ENSMUSG00000022295 | ATPase, H+ transporting, lysosomal V1 subunit C1 [Source:MGI Symbol;Acc:MGI:1913585] | Atp6v1c1 | Enriched in dLGN | 1.037 | 0.00003657 | 229 |
| ENSMUSG00000021576 | programmed cell death 6 [Source:MGI Symbol;Acc:MGI:109283] | Pdcd6 | Enriched in dLGN | 1.036 | 0.009279 | 37.47 |
| ENSMUSG00000000605 | chloride channel, voltage-sensitive 4 [Source:MGI Symbol;Acc:MGI:104571] | Clcn4 | Enriched in dLGN | 1.035 | 3.267E-10 | 446.8 |
| ENSMUSG00000039953 | calsyntenin 1 [Source:MGI Symbol;Acc:MGI:1929895] | Clstn1 | Enriched in dLGN | 1.033 | 7.225E-08 | 3184 |
| ENSMUSG00000033047 | eukaryotic translation initiation factor 3, subunit L [Source:MGI Symbol;Acc:MGI:2386251] | Eif3l | Enriched in dLGN | 1.032 | 0.001563 | 114.6 |
| ENSMUSG00000067995 | general transcription factor IIF, polypeptide 2 [Source:MGI Symbol;Acc:MGI:1915955] | Gtf2f2 | Enriched in dLGN | 1.032 | 0.004724 | 71.17 |
| ENSMUSG00000041058 | WW domain containing E3 ubiquitin protein ligase 1 [Source:MGI Symbol;Acc:MGI:1861728] | Wwp1 | Enriched in dLGN | 1.031 | 0.000001723 | 207.5 |
| ENSMUSG00000039983 | coiled-coil domain containing 32 [Source:MGI Symbol;Acc:MGI:2685477] | Ccdc32 | Enriched in dLGN | 1.031 | 0.0006285 | 49.02 |
| ENSMUSG00000049092 | G protein-coupled receptor 137C [Source:MGI Symbol;Acc:MGI:1917963] | Gpr137c | Enriched in dLGN | 1.031 | 0.005024 | 120.2 |
| ENSMUSG00000029402 | small nuclear ribonucleoprotein 35 (U11/U12) [Source:MGI Symbol;Acc:MGI:1923417] | Snrnp35 | Enriched in dLGN | 1.031 | 0.01768 | 29.69 |
| ENSMUSG00000027835 | programmed cell death 10 [Source:MGI Symbol;Acc:MGI:1928396] | Pdcd10 | Enriched in dLGN | 1.03 | 0.02612 | 40.8 |
| ENSMUSG00000071180 | small integral membrane protein 15 [Source:MGI Symbol;Acc:MGI:1922866] | Smim15 | Enriched in dLGN | 1.026 | 0.001367 | 41.58 |
| ENSMUSG00000033379 | ATPase, H+ transporting, lysosomal V0 subunit B [Source:MGI Symbol;Acc:MGI:1890510] | Atp6v0b | Enriched in dLGN | 1.025 | 8.295E-08 | 226.1 |
| ENSMUSG00000026155 | small ArfGAP 1 [Source:MGI Symbol;Acc:MGI:2138261] | Smap1 | Enriched in dLGN | 1.025 | 0.00002174 | 253.2 |
| ENSMUSG00000029840 | myotrophin [Source:MGI Symbol;Acc:MGI:99445] | Mtpn | Enriched in dLGN | 1.024 | 0.00001855 | 335.2 |
| ENSMUSG00000027634 | N-myc downstream regulated gene 3 [Source:MGI Symbol;Acc:MGI:1352499] | Ndrg3 | Enriched in dLGN | 1.024 | 0.00002584 | 572.8 |
| ENSMUSG00000020435 | oxysterol binding protein 2 [Source:MGI Symbol;Acc:MGI:1921559] | Osbp2 | Enriched in dLGN | 1.023 | 0.0006106 | 628.7 |
| ENSMUSG00000022323 | reactive intermediate imine deaminase A homolog [Source:MGI Symbol;Acc:MGI:1095401] | Rida | Enriched in dLGN | 1.023 | 0.005742 | 20.62 |
| ENSMUSG00000028932 | proteasome (prosome, macropain) 26S subunit, ATPase 2 [Source:MGI Symbol;Acc:MGI:109555] | Psmc2 | Enriched in dLGN | 1.023 | 0.006642 | 150.2 |
| ENSMUSG00000025237 | poly (ADP-ribose) polymerase family, member 6 [Source:MGI Symbol;Acc:MGI:1914537] | Parp6 | Enriched in dLGN | 1.017 | 8.254E-07 | 345.3 |
| ENSMUSG00000020153 | NADH:ubiquinone oxidoreductase core subunit S7 [Source:MGI Symbol;Acc:MGI:1922656] | Ndufs7 | Enriched in dLGN | 1.017 | 0.0001418 | 124 |
| ENSMUSG00000039347 | ATPase, H+ transporting, lysosomal V0 subunit E2 [Source:MGI Symbol;Acc:MGI:1923502] | Atp6v0e2 | Enriched in dLGN | 1.016 | 0.000004451 | 554.6 |
| ENSMUSG00000030879 | mitochondrial ribosomal protein L17 [Source:MGI Symbol;Acc:MGI:1351608] | Mrpl17 | Enriched in dLGN | 1.015 | 0.00005427 | 76.99 |
| ENSMUSG00000028478 | clathrin, light polypeptide (Lca) [Source:MGI Symbol;Acc:MGI:894297] | Clta | Enriched in dLGN | 1.014 | 1.826E-08 | 866.9 |
| ENSMUSG00000049760 | mitochondrial contact site and cristae organizing system subunit 13 [Source:MGI Symbol;Acc:MGI:2442174] | Micos13 | Enriched in dLGN | 1.014 | 0.0003502 | 44.94 |
| ENSMUSG00000031451 | growth arrest specific 6 [Source:MGI Symbol;Acc:MGI:95660] | Gas6 | Enriched in dLGN | 1.012 | 7.384E-08 | 424.5 |
| ENSMUSG00000022792 | tyrosyl-tRNA synthetase 2 (mitochondrial) [Source:MGI Symbol;Acc:MGI:1917370] | Yars2 | Enriched in dLGN | 1.012 | 0.005744 | 29.46 |
| ENSMUSG00000024208 | ubiquinol-cytochrome c reductase complex assembly factor 2 [Source:MGI Symbol;Acc:MGI:1914517] | Uqcc2 | Enriched in dLGN | 1.011 | 0.001977 | 101.2 |
| ENSMUSG00000025240 | SAC1 suppressor of actin mutations 1-like (yeast) [Source:MGI Symbol;Acc:MGI:1933169] | Sacm1l | Enriched in dLGN | 1.011 | 0.002156 | 163.5 |
| ENSMUSG00000055681 | coatomer protein complex, subunit epsilon [Source:MGI Symbol;Acc:MGI:1891702] | Cope | Enriched in dLGN | 1.011 | 0.002232 | 112.7 |
| ENSMUSG00000052299 | listerin E3 ubiquitin protein ligase 1 [Source:MGI Symbol;Acc:MGI:1926163] | Ltn1 | Enriched in dLGN | 1.01 | 0.000003952 | 189.1 |
| ENSMUSG00000019782 | RWD domain containing 1 [Source:MGI Symbol;Acc:MGI:1913771] | Rwdd1 | Enriched in dLGN | 1.01 | 0.001276 | 84.05 |
| ENSMUSG00000027236 | eukaryotic translation initiation factor 3, subunit J1 [Source:MGI Symbol;Acc:MGI:1925905] | Eif3j1 | Enriched in dLGN | 1.01 | 0.001327 | 85.54 |
| ENSMUSG00000025613 | chaperonin containing Tcp1, subunit 8 (theta) [Source:MGI Symbol;Acc:MGI:107183] | Cct8 | Enriched in dLGN | 1.009 | 0.001441 | 259.2 |
| ENSMUSG00000061273 | membrane magnesium transporter 1 [Source:MGI Symbol;Acc:MGI:2384305] | Mmgt1 | Enriched in dLGN | 1.009 | 0.006795 | 82.86 |
| ENSMUSG00000028232 | transmembrane protein 68 [Source:MGI Symbol;Acc:MGI:1919348] | Tmem68 | Enriched in dLGN | 1.006 | 0.004524 | 41.85 |
| ENSMUSG00000057388 | mitochondrial ribosomal protein L18 [Source:MGI Symbol;Acc:MGI:1914931] | Mrpl18 | Enriched in dLGN | 1.005 | 0.0004133 | 67.46 |
| ENSMUSG00000054162 | sparc/osteonectin, cwcv and kazal-like domains proteoglycan 3 [Source:MGI Symbol;Acc:MGI:1920152] | Spock3 | Enriched in dLGN | 1.005 | 0.005563 | 404.4 |
| ENSMUSG00000029649 | proteasome maturation protein [Source:MGI Symbol;Acc:MGI:1913787] | Pomp | Enriched in dLGN | 1.004 | 0.0007444 | 109 |
| ENSMUSG00000004285 | ATPase, H+ transporting, lysosomal V1 subunit F [Source:MGI Symbol;Acc:MGI:1913394] | Atp6v1f | Enriched in dLGN | 1.004 | 0.01958 | 78.37 |
| ENSMUSG00000001016 | interleukin enhancer binding factor 2 [Source:MGI Symbol;Acc:MGI:1915031] | Ilf2 | Enriched in dLGN | 1.002 | 0.00004546 | 131.5 |
| ENSMUSG00000033124 | autophagy related 9A [Source:MGI Symbol;Acc:MGI:2138446] | Atg9a | Enriched in dLGN | 1 | 1.697E-10 | 274.4 |
| ENSMUSG00000061479 | small nuclear ribonucleoprotein polypeptide A [Source:MGI Symbol;Acc:MGI:1855690] | Snrpa | Enriched in dLGN | 1 | 0.01366 | 69.88 |
| ENSMUSG00000031772 | contactin associated protein-like 4 [Source:MGI Symbol;Acc:MGI:2183572] | Cntnap4 | Enriched in SCN | -1.008 | 0.007421 | 164.6 |
| ENSMUSG00000000127 | fer (fms/fps related) protein kinase [Source:MGI Symbol;Acc:MGI:105917] | Fer | Enriched in SCN | -1.013 | 0.006067 | 224 |
| ENSMUSG00000023235 | chemokine (C-C motif) ligand 25 [Source:MGI Symbol;Acc:MGI:1099448] | Ccl25 | Enriched in SCN | -1.029 | 0.03853 | 57.94 |
| ENSMUSG00000027014 | CWC22 spliceosome-associated protein [Source:MGI Symbol;Acc:MGI:2136773] | Cwc22 | Enriched in SCN | -1.035 | 0.03442 | 206.1 |
| ENSMUSG00000072623 | zinc finger protein 9 [Source:MGI Symbol;Acc:MGI:99210] | Zfp9 | Enriched in SCN | -1.102 | 0.00009753 | 48.54 |
| ENSMUSG00000034610 | terminal uridylyl transferase 4 [Source:MGI Symbol;Acc:MGI:2445126] | Tut4 | Enriched in SCN | -1.149 | 0.0000561 | 281.6 |
| ENSMUSG00000037143 | cilia and flagella associated protein 61 [Source:MGI Symbol;Acc:MGI:1926024] | Cfap61 | Enriched in SCN | -1.243 | 0.007612 | 108.6 |
| ENSMUSG00000034845 | plasmalemma vesicle associated protein [Source:MGI Symbol;Acc:MGI:1890497] | Plvap | Enriched in SCN | -1.289 | 0.01112 | 16.04 |
| ENSMUSG00000028158 | microsomal triglyceride transfer protein [Source:MGI Symbol;Acc:MGI:106926] | Mttp | Enriched in SCN | -1.36 | 0.00004479 | 48.33 |
| ENSMUSG00000033487 | fibronectin type III domain containing 3A [Source:MGI Symbol;Acc:MGI:1196463] | Fndc3a | Enriched in SCN | -1.514 | 2.587E-13 | 371.9 |
| ENSMUSG00000079410 | predicted gene 2897 [Source:MGI Symbol;Acc:MGI:3781075] | Gm2897 | Enriched in SCN | -1.884 | 0.03984 | 6.752 |
| ENSMUSG00000022803 | popeye domain containing 2 [Source:MGI Symbol;Acc:MGI:1930150] | Popdc2 | Enriched in SCN | -2.662 | 0.02136 | 4.598 |
| ENSMUSG00000040752 | myosin, heavy polypeptide 6, cardiac muscle, alpha [Source:MGI Symbol;Acc:MGI:97255] | Myh6 | Enriched in SCN | -4.532 | 1.142E-09 | 9.823 |

**Contrast numerator: P15_Het_SCN. Contrast denominator: P15_Het_retina.**

| ensembl_gene_id | description | mgi_symbol | Significance | deseq_logfc | deseq_adjp | deseq_basemean |
|---|---|---|---|---|---|---|
| ENSMUSG00000027301 | oxytocin [Source:MGI Symbol;Acc:MGI:97453] | Oxt | Enriched in SCN | 11.42 | 0.000008027 | 22.27 |
| ENSMUSG00000025314 | protein tyrosine phosphatase, receptor type, J [Source:MGI Symbol;Acc:MGI:104574] | Ptprj | Enriched in SCN | 5.488 | 1.337E-19 | 998.9 |
| ENSMUSG00000004892 | brevican [Source:MGI Symbol;Acc:MGI:1096385] | Bcan | Enriched in SCN | 4.287 | 4.501E-62 | 546.7 |
| ENSMUSG00000030854 | protein tyrosine phosphatase, non-receptor type 5 [Source:MGI Symbol;Acc:MGI:97807] | Ptpn5 | Enriched in SCN | 4.001 | 5.622E-87 | 407 |
| ENSMUSG00000042425 | FERM and PDZ domain containing 3 [Source:MGI Symbol;Acc:MGI:3646547] | Frmpd3 | Enriched in SCN | 3.303 | 4.103E-16 | 191.4 |
| ENSMUSG00000044576 | GRB2 associated regulator of MAPK1 subtype 2 [Source:MGI Symbol;Acc:MGI:2685290] | Garem2 | Enriched in SCN | 3.302 | 1.425E-28 | 62.67 |
| ENSMUSG00000022112 | glypican 5 [Source:MGI Symbol;Acc:MGI:1194894] | Gpc5 | Enriched in SCN | 3.2 | 1.792E-13 | 390.3 |
| ENSMUSG00000030889 | von Willebrand factor A domain containing 3A [Source:MGI Symbol;Acc:MGI:3041229] | Vwa3a | Enriched in SCN | 3.082 | 0.000002424 | 26.65 |
| ENSMUSG00000037428 | VGF nerve growth factor inducible [Source:MGI Symbol;Acc:MGI:1343180] | Vgf | Enriched in SCN | 2.997 | 3.263E-29 | 1249 |
| ENSMUSG00000034226 | ras homolog family member V [Source:MGI Symbol;Acc:MGI:2444227] | Rhov | Enriched in SCN | 2.947 | 9.96E-19 | 54.65 |
| ENSMUSG00000052026 | solute carrier family 6 (neurotransmitter transporter, L-proline), member 7 [Source:MGI Symbol;Acc:MGI:2147363] | Slc6a7 | Enriched in SCN | 2.918 | 9.87E-38 | 217.9 |
| ENSMUSG00000032023 | junctional cadherin complex regulator [Source:MGI Symbol;Acc:MGI:1918239] | Jhy | Enriched in SCN | 2.63 | 0.00008404 | 5.097 |
| ENSMUSG00000042216 | small G protein signaling modulator 1 [Source:MGI Symbol;Acc:MGI:107320] | Sgsm1 | Enriched in SCN | 2.594 | 1.034E-33 | 412.2 |
| ENSMUSG00000045348 | neuronal tyrosine-phosphorylated phosphoinositide 3-kinase adaptor 1 [Source:MGI Symbol;Acc:MGI:2443880] | Nyap1 | Enriched in SCN | 2.542 | 1.475E-42 | 501.1 |
| ENSMUSG00000009394 | synapsin II [Source:MGI Symbol;Acc:MGI:103020] | Syn2 | Enriched in SCN | 2.424 | 3.795E-21 | 611.4 |
| ENSMUSG00000039313 | membrane integral NOTCH2 associated receptor 1 [Source:MGI Symbol;Acc:MGI:2667167] | Minar1 | Enriched in SCN | 2.338 | 2.562E-07 | 60.24 |
| ENSMUSG00000031841 | cadherin 13 [Source:MGI Symbol;Acc:MGI:99551] | Cdh13 | Enriched in SCN | 2.258 | 1.051E-10 | 448.3 |
| ENSMUSG00000039579 | glutamate receptor ionotropic, NMDA3A [Source:MGI Symbol;Acc:MGI:1933206] | Grin3a | Enriched in SCN | 2.229 | 9.23E-18 | 314.9 |
| ENSMUSG00000079388 | RIKEN cDNA 2610042L04 gene [Source:MGI Symbol;Acc:MGI:1914305] | 2610042L04Rik | Enriched in SCN | 2.224 | 0.01464 | 4.418 |
| ENSMUSG00000027200 | sema domain, transmembrane domain (TM), and cytoplasmic domain, (semaphorin) 6D [Source:MGI Symbol;Acc:MGI:2387661] | Sema6d | Enriched in SCN | 2.199 | 9.783E-16 | 731.9 |
| ENSMUSG00000079391 | predicted gene 2974 [Source:MGI Symbol;Acc:MGI:3781152] | Gm2974 | Enriched in SCN | 2.173 | 0.007268 | 6.08 |
| ENSMUSG00000022602 | activity regulated cytoskeletal-associated protein [Source:MGI Symbol;Acc:MGI:88067] | Arc | Enriched in SCN | 2.162 | 1.933E-09 | 44.21 |
| ENSMUSG00000031860 | pre B cell leukemia homeobox 4 [Source:MGI Symbol;Acc:MGI:1931321] | Pbx4 | Enriched in SCN | 2.151 | 1.105E-13 | 37.95 |
| ENSMUSG00000023017 | acid-sensing (proton-gated) ion channel 1 [Source:MGI Symbol;Acc:MGI:1194915] | Asic1 | Enriched in SCN | 2.038 | 9.327E-22 | 275.4 |
| ENSMUSG00000045731 | prepronociceptin [Source:MGI Symbol;Acc:MGI:105308] | Pnoc | Enriched in SCN | 2.004 | 4.337E-07 | 50.56 |
| ENSMUSG00000004187 | kinesin family member C2 [Source:MGI Symbol;Acc:MGI:109187] | Kifc2 | Enriched in SCN | 1.956 | 1.102E-31 | 769 |
| ENSMUSG00000002771 | glutamate receptor, ionotropic, NMDA2D (epsilon 4) [Source:MGI Symbol;Acc:MGI:95823] | Grin2d | Enriched in SCN | 1.901 | 4.445E-19 | 466.7 |
| ENSMUSG00000066129 | kinase non-catalytic C-lobe domain (KIND) containing 1 [Source:MGI Symbol;Acc:MGI:1923734] | Kndc1 | Enriched in SCN | 1.9 | 1.124E-16 | 714.5 |
| ENSMUSG00000019785 | clavesin 2 [Source:MGI Symbol;Acc:MGI:2443223] | Clvs2 | Enriched in SCN | 1.898 | 1.38E-12 | 86.26 |
| ENSMUSG00000031963 | BMP-binding endothelial regulator [Source:MGI Symbol;Acc:MGI:1920480] | Bmper | Enriched in SCN | 1.857 | 3.636E-07 | 44.47 |
| ENSMUSG00000004558 | N-myc downstream regulated gene 2 [Source:MGI Symbol;Acc:MGI:1352498] | Ndrg2 | Enriched in SCN | 1.808 | 1.654E-14 | 450.7 |
| ENSMUSG00000042766 | tripartite motif-containing 46 [Source:MGI Symbol;Acc:MGI:2673000] | Trim46 | Enriched in SCN | 1.807 | 1.43E-22 | 200.1 |
| ENSMUSG00000028007 | sorting nexin 7 [Source:MGI Symbol;Acc:MGI:1923811] | Snx7 | Enriched in SCN | 1.787 | 1.776E-10 | 54.52 |
| ENSMUSG00000036502 | transmembrane protein 255A [Source:MGI Symbol;Acc:MGI:3045722] | Tmem255a | Enriched in SCN | 1.748 | 0.0002117 | 75.7 |
| ENSMUSG00000006711 | RIKEN cDNA D130043K22 gene [Source:MGI Symbol;Acc:MGI:3036268] | D130043K22Rik | Enriched in SCN | 1.724 | 3.379E-08 | 133.1 |
| ENSMUSG00000022861 | diacylglycerol kinase, gamma [Source:MGI Symbol;Acc:MGI:105060] | Dgkg | Enriched in SCN | 1.682 | 0.000008403 | 135 |
| ENSMUSG00000051146 | calcium/calmodulin-dependent protein kinase II inhibitor 2 [Source:MGI Symbol;Acc:MGI:1920297] | Camk2n2 | Enriched in SCN | 1.637 | 2.002E-16 | 513.1 |
| ENSMUSG00000039546 | adherens junction associated protein 1 [Source:MGI Symbol;Acc:MGI:2685419] | Ajap1 | Enriched in SCN | 1.631 | 5.36E-11 | 293.4 |
| ENSMUSG00000062184 | heparan sulfate 6-O-sulfotransferase 2 [Source:MGI Symbol;Acc:MGI:1354959] | Hs6st2 | Enriched in SCN | 1.624 | 3.73E-08 | 235.4 |
| ENSMUSG00000035835 | phospholipid phosphatase related 3 [Source:MGI Symbol;Acc:MGI:2388640] | Plppr3 | Enriched in SCN | 1.619 | 2.028E-11 | 329.9 |
| ENSMUSG00000028680 | polo like kinase 3 [Source:MGI Symbol;Acc:MGI:109604] | Plk3 | Enriched in SCN | 1.614 | 1.344E-07 | 41.32 |
| ENSMUSG00000040811 | echinoderm microtubule associated protein like 2 [Source:MGI Symbol;Acc:MGI:1919455] | Eml2 | Enriched in SCN | 1.589 | 2.55E-14 | 740.8 |
| ENSMUSG00000000627 | sema domain, immunoglobulin domain (Ig), TM domain, and short cytoplasmic domain [Source:MGI Symbol;Acc:MGI:1340055] | Sema4f | Enriched in SCN | 1.565 | 1.525E-11 | 125.4 |
| ENSMUSG00000028949 | SWI/SNF related, matrix associated, actin dependent regulator of chromatin, subfamily d, member 3 [Source:MGI Symbol;Acc:MGI:1914243] | Smarcd3 | Enriched in SCN | 1.547 | 2.801E-22 | 234.8 |
| ENSMUSG00000074736 | synapse differentiation inducing 1 [Source:MGI Symbol;Acc:MGI:3702158] | Syndig1 | Enriched in SCN | 1.536 | 0.00006582 | 50.55 |
| ENSMUSG00000001227 | sema domain, transmembrane domain (TM), and cytoplasmic domain, (semaphorin) 6B [Source:MGI Symbol;Acc:MGI:1202889] | Sema6b | Enriched in SCN | 1.501 | 1.718E-15 | 529.4 |
| ENSMUSG00000067242 | leucine-rich repeat LGI family, member 1 [Source:MGI Symbol;Acc:MGI:1861691] | Lgi1 | Enriched in SCN | 1.452 | 5.905E-07 | 72.29 |
| ENSMUSG00000055409 | NEL-like 1 [Source:MGI Symbol;Acc:MGI:2443902] | Nell1 | Enriched in SCN | 1.419 | 0.0002848 | 543.3 |
| ENSMUSG00000008153 | calsyntenin 3 [Source:MGI Symbol;Acc:MGI:2178323] | Clstn3 | Enriched in SCN | 1.414 | 2.371E-07 | 1064 |
| ENSMUSG00000052301 | double C2, alpha [Source:MGI Symbol;Acc:MGI:109446] | Doc2a | Enriched in SCN | 1.414 | 0.001709 | 56.86 |
| ENSMUSG00000036815 | dipeptidylpeptidase 10 [Source:MGI Symbol;Acc:MGI:2442409] | Dpp10 | Enriched in SCN | 1.411 | 0.0001088 | 576.4 |
| ENSMUSG00000028755 | cytidine deaminase [Source:MGI Symbol;Acc:MGI:1919519] | Cda | Enriched in SCN | 1.383 | 0.003104 | 14.43 |
| ENSMUSG00000034353 | receptor (calcitonin) activity modifying protein 1 [Source:MGI Symbol;Acc:MGI:1858418] | Ramp1 | Enriched in SCN | 1.366 | 0.0003488 | 54.71 |
| ENSMUSG00000058740 | potassium channel, subfamily T, member 1 [Source:MGI Symbol;Acc:MGI:1924627] | Kcnt1 | Enriched in SCN | 1.363 | 0.000004415 | 138.9 |
| ENSMUSG00000073433 | Rho GDP dissociation inhibitor (GDI) gamma [Source:MGI Symbol;Acc:MGI:108430] | Arhgdig | Enriched in SCN | 1.361 | 9.671E-07 | 185.1 |
| ENSMUSG00000017631 | active BCR-related gene [Source:MGI Symbol;Acc:MGI:107771] | Abr | Enriched in SCN | 1.352 | 7.222E-15 | 1239 |
| ENSMUSG00000021541 | transient receptor potential cation channel, subfamily C, member 7 [Source:MGI Symbol;Acc:MGI:1349470] | Trpc7 | Enriched in SCN | 1.346 | 0.0007411 | 62.32 |
| ENSMUSG00000020882 | calcium channel, voltage-dependent, beta 1 subunit [Source:MGI Symbol;Acc:MGI:102522] | Cacnb1 | Enriched in SCN | 1.345 | 1.191E-16 | 246.5 |
| ENSMUSG00000056413 | ArfGAP with dual PH domains 1 [Source:MGI Symbol;Acc:MGI:2442201] | Adap1 | Enriched in SCN | 1.345 | 2.013E-15 | 552.2 |
| ENSMUSG00000033061 | regulated endocrine-specific protein 18 [Source:MGI Symbol;Acc:MGI:1098222] | Resp18 | Enriched in SCN | 1.323 | 0.01084 | 74.08 |
| ENSMUSG00000075318 | sodium channel, voltage-gated, type II, alpha [Source:MGI Symbol;Acc:MGI:98248] | Scn2a | Enriched in SCN | 1.315 | 1.899E-11 | 1334 |
| ENSMUSG00000031438 | ring finger protein 128 [Source:MGI Symbol;Acc:MGI:1914139] | Rnf128 | Enriched in SCN | 1.295 | 0.01008 | 18.97 |
| ENSMUSG00000016346 | potassium voltage-gated channel, subfamily Q, member 2 [Source:MGI Symbol;Acc:MGI:1309503] | Kcnq2 | Enriched in SCN | 1.282 | 2.773E-10 | 1096 |
| ENSMUSG00000020331 | hyperpolarization-activated, cyclic nucleotide-gated K+ 2 [Source:MGI Symbol;Acc:MGI:1298210] | Hcn2 | Enriched in SCN | 1.261 | 2.537E-10 | 598.2 |
| ENSMUSG00000020297 | neuron specific gene family member 2 [Source:MGI Symbol;Acc:MGI:1202070] | Nsg2 | Enriched in SCN | 1.232 | 4.085E-13 | 922.9 |
| ENSMUSG00000004933 | megakaryocyte-associated tyrosine kinase [Source:MGI Symbol;Acc:MGI:99259] | Matk | Enriched in SCN | 1.232 | 1.363E-07 | 175.5 |
| ENSMUSG00000032503 | cyclic AMP-regulated phosphoprotein, 21 [Source:MGI Symbol;Acc:MGI:107562] | Arpp21 | Enriched in SCN | 1.203 | 0.001466 | 594.6 |
| ENSMUSG00000030592 | ryanodine receptor 1, skeletal muscle [Source:MGI Symbol;Acc:MGI:99659] | Ryr1 | Enriched in SCN | 1.186 | 0.01438 | 72.59 |
| ENSMUSG00000027220 | synaptotagmin XIII [Source:MGI Symbol;Acc:MGI:1933945] | Syt13 | Enriched in SCN | 1.166 | 2.613E-11 | 763.5 |
| ENSMUSG00000007944 | tetratricopeptide repeat domain 9B [Source:MGI Symbol;Acc:MGI:1920282] | Ttc9b | Enriched in SCN | 1.139 | 0.00001759 | 147 |
| ENSMUSG00000069911 | inhibitory synaptic factor family member 2B [Source:MGI Symbol;Acc:MGI:3643491] | Insyn2b | Enriched in SCN | 1.135 | 0.004021 | 58.95 |
| ENSMUSG00000040035 | dispatched RND transporter family member 2 [Source:MGI Symbol;Acc:MGI:2388733] | Disp2 | Enriched in SCN | 1.131 | 0.00001764 | 1280 |
| ENSMUSG00000031760 | metallothionein 3 [Source:MGI Symbol;Acc:MGI:97173] | Mt3 | Enriched in SCN | 1.131 | 0.0006641 | 131.4 |
| ENSMUSG00000022865 | coxsackie virus and adenovirus receptor [Source:MGI Symbol;Acc:MGI:1201679] | Cxadr | Enriched in SCN | 1.12 | 0.00089 | 403.7 |
| ENSMUSG00000025855 | protein kinase, cAMP dependent regulatory, type I beta [Source:MGI Symbol;Acc:MGI:97759] | Prkar1b | Enriched in SCN | 1.109 | 5.99E-08 | 871 |
| ENSMUSG00000043051 | disrupted in schizophrenia 1 [Source:MGI Symbol;Acc:MGI:2447658] | Disc1 | Enriched in SCN | 1.107 | 0.006007 | 34.75 |
| ENSMUSG00000035653 | leucine rich repeat and fibronectin type III domain containing 5 [Source:MGI Symbol;Acc:MGI:2144814] | Lrfn5 | Enriched in SCN | 1.105 | 0.001265 | 564.4 |
| ENSMUSG00000032946 | RAS, guanyl releasing protein 2 [Source:MGI Symbol;Acc:MGI:1333849] | Rasgrp2 | Enriched in SCN | 1.098 | 0.0004053 | 149.8 |
| ENSMUSG00000041362 | shootin 1 [Source:MGI Symbol;Acc:MGI:1918903] | Shtn1 | Enriched in SCN | 1.092 | 2.547E-07 | 302.5 |
| ENSMUSG00000046613 | von Willebrand factor A domain containing 5B2 [Source:MGI Symbol;Acc:MGI:2681859] | Vwa5b2 | Enriched in SCN | 1.086 | 0.0001317 | 119.4 |
| ENSMUSG00000040721 | zinc finger homeobox 2 [Source:MGI Symbol;Acc:MGI:2686934] | Zfhx2 | Enriched in SCN | 1.076 | 1.712E-08 | 608.5 |
| ENSMUSG00000029126 | neuron specific gene family member 1 [Source:MGI Symbol;Acc:MGI:109149] | Nsg1 | Enriched in SCN | 1.062 | 0.00001905 | 719.7 |
| ENSMUSG00000047344 | LanC lantibiotic synthetase component C-like 3 (bacterial) [Source:MGI Symbol;Acc:MGI:2443335] | Lancl3 | Enriched in SCN | 1.05 | 0.001469 | 26.64 |
| ENSMUSG00000020990 | cyclin-dependent kinase-like 1 (CDC2-related kinase) [Source:MGI Symbol;Acc:MGI:1918341] | Cdkl1 | Enriched in SCN | 1.029 | 0.003242 | 53.67 |
| ENSMUSG00000036995 | ArfGAP with SH3 domain, ankyrin repeat and PH domain 3 [Source:MGI Symbol;Acc:MGI:2684986] | Asap3 | Enriched in retir | -1.002 | 0.003056 | 47.68 |
| ENSMUSG00000032011 | thymus cell antigen 1, theta [Source:MGI Symbol;Acc:MGI:98747] | Thy1 | Enriched in retir | -1.063 | 0.001104 | 952.8 |
| ENSMUSG00000033917 | glycerophosphodiester phosphodiesterase 1 [Source:MGI Symbol;Acc:MGI:1891827] | Gde1 | Enriched in retir | -1.065 | 1.334E-07 | 234.5 |
| ENSMUSG00000041548 | heat shock protein 8 [Source:MGI Symbol;Acc:MGI:2135756] | Hspb8 | Enriched in retir | -1.069 | 0.02352 | 19.43 |
| ENSMUSG00000002032 | transmembrane protein 25 [Source:MGI Symbol;Acc:MGI:1918937] | Tmem25 | Enriched in retir | -1.079 | 0.0002955 | 70.33 |
| ENSMUSG00000030500 | solute carrier family 17 (sodium-dependent inorganic phosphate cotransporter), member 6 [Source:MGI Symbol;Acc:MGI:2156052] | Slc17a6 | Enriched in retir | -1.084 | 0.009563 | 605.1 |
| ENSMUSG00000022377 | ArfGAP with SH3 domain, ankyrin repeat and PH domain1 [Source:MGI Symbol;Acc:MGI:1342335] | Asap1 | Enriched in retir | -1.084 | 0.000003534 | 410.6 |
| ENSMUSG00000040813 | testis expressed gene 264 [Source:MGI Symbol;Acc:MGI:1096570] | Tex264 | Enriched in retir | -1.092 | 9.543E-10 | 124.3 |
| ENSMUSG00000033615 | complexin 1 [Source:MGI Symbol;Acc:MGI:104727] | Cplx1 | Enriched in retir | -1.125 | 0.001017 | 2304 |
| ENSMUSG00000039461 | T cell leukemia translocation altered gene [Source:MGI Symbol;Acc:MGI:1918829] | Tcta | Enriched in retir | -1.135 | 8.634E-07 | 50.88 |
| ENSMUSG00000035407 | KN motif and ankyrin repeat domains 4 [Source:MGI Symbol;Acc:MGI:3043381] | Kank4 | Enriched in retir | -1.173 | 0.007056 | 22.26 |
| ENSMUSG00000013367 | IgLON family member 5 [Source:MGI Symbol;Acc:MGI:2686277] | Iglon5 | Enriched in retir | -1.213 | 2.828E-09 | 271 |
| ENSMUSG00000046719 | neurexophilin 3 [Source:MGI Symbol;Acc:MGI:1336188] | Nxph3 | Enriched in retir | -1.225 | 0.0008099 | 27.7 |
| ENSMUSG00000062300 | nectin cell adhesion molecule 2 [Source:MGI Symbol;Acc:MGI:97822] | Nectin2 | Enriched in retir | -1.236 | 0.00004876 | 66.46 |
| ENSMUSG00000020635 | FK506 binding protein 1b [Source:MGI Symbol;Acc:MGI:1336205] | Fkbp1b | Enriched in retir | -1.24 | 0.001346 | 66.86 |
| ENSMUSG00000026688 | microsomal glutathione S-transferase 3 [Source:MGI Symbol;Acc:MGI:1913697] | Mgst3 | Enriched in retir | -1.265 | 0.01711 | 88.46 |
| ENSMUSG00000060240 | cell cycle exit and neuronal differentiation 1 [Source:MGI Symbol;Acc:MGI:1929898] | Cend1 | Enriched in retir | -1.293 | 0.001176 | 473.2 |
| ENSMUSG00000046480 | sodium channel, type IV, beta [Source:MGI Symbol;Acc:MGI:2687406] | Scn4b | Enriched in retir | -1.296 | 0.009303 | 167.6 |
| ENSMUSG00000050822 | solute carrier family 29 (nucleoside transporters), member 4 [Source:MGI Symbol;Acc:MGI:2385330] | Slc29a4 | Enriched in retir | -1.313 | 0.00003874 | 126.5 |
| ENSMUSG00000039195 | bublin coiled coil protein [Source:MGI Symbol;Acc:MGI:1920987] | Bbln | Enriched in retir | -1.333 | 0.00001676 | 50.47 |
| ENSMUSG00000020836 | coronin 6 [Source:MGI Symbol;Acc:MGI:2183448] | Coro6 | Enriched in retir | -1.336 | 0.02878 | 39.45 |
| ENSMUSG00000036023 | poly (ADP-ribose) polymerase family, member 2 [Source:MGI Symbol;Acc:MGI:1341112] | Parp2 | Enriched in retir | -1.425 | 0.0002981 | 46.73 |
| ENSMUSG00000060969 | Iroquois homeobox 1 [Source:MGI Symbol;Acc:MGI:1197515] | Irx1 | Enriched in retir | -1.433 | 0.01062 | 29.96 |
| ENSMUSG00000040557 | methyltransferase like 27 [Source:MGI Symbol;Acc:MGI:1933146] | Mettl27 | Enriched in retir | -1.45 | 0.000008655 | 18.67 |
| ENSMUSG00000018340 | annexin A6 [Source:MGI Symbol;Acc:MGI:88255] | Anxa6 | Enriched in retir | -1.452 | 0.0007421 | 254.2 |
| ENSMUSG00000014846 | tubulin polymerization-promoting protein family member 3 [Source:MGI Symbol;Acc:MGI:1915221] | Tppp3 | Enriched in retir | -1.452 | 0.000003859 | 139.8 |
| ENSMUSG00000000326 | catechol-O-methyltransferase [Source:MGI Symbol;Acc:MGI:88470] | Comt | Enriched in retir | -1.481 | 4.321E-11 | 103.3 |
| ENSMUSG00000018634 | corticotropin releasing hormone receptor 1 [Source:MGI Symbol;Acc:MGI:88498] | Crhr1 | Enriched in retir | -1.518 | 0.00009798 | 47.56 |
| ENSMUSG00000036585 | fibroblast growth factor 1 [Source:MGI Symbol;Acc:MGI:95515] | Fgf1 | Enriched in retir | -1.555 | 0.00004451 | 148.8 |
| ENSMUSG00000020810 | cytoglobin [Source:MGI Symbol;Acc:MGI:2149481] | Cygb | Enriched in retir | -1.645 | 2.95E-08 | 213.7 |
| ENSMUSG00000042115 | kelch domain containing 8A [Source:MGI Symbol;Acc:MGI:2442630] | Klhdc8a | Enriched in retir | -1.652 | 1.354E-11 | 181.2 |
| ENSMUSG00000019966 | kit ligand [Source:MGI Symbol;Acc:MGI:96974] | Kitl | Enriched in retir | -1.656 | 0.000002159 | 453.1 |
| ENSMUSG00000001333 | syncoilin [Source:MGI Symbol;Acc:MGI:1916078] | Sync | Enriched in retir | -1.694 | 0.000699 | 17.38 |
| ENSMUSG00000056596 | TMF1-regulated nuclear protein 1 [Source:MGI Symbol;Acc:MGI:1916789] | Trnp1 | Enriched in retir | -1.707 | 2.569E-10 | 640.5 |
| ENSMUSG00000040703 | cytochrome P450, family 2, subfamily s, polypeptide 1 [Source:MGI Symbol;Acc:MGI:1921384] | Cyp2s1 | Enriched in retir | -1.717 | 0.00004524 | 19.07 |
| ENSMUSG00000038457 | transmembrane protein 255B [Source:MGI Symbol;Acc:MGI:2685533] | Tmem255b | Enriched in retir | -1.778 | 2.514E-09 | 59.61 |
| ENSMUSG00000032599 | inositol hexaphosphate kinase 2 [Source:MGI Symbol;Acc:MGI:1923750] | Ip6k2 | Enriched in retir | -1.789 | 9.611E-11 | 280.5 |
| ENSMUSG00000001794 | calpain, small subunit 1 [Source:MGI Symbol;Acc:MGI:88266] | Capns1 | Enriched in retir | -1.792 | 3.063E-09 | 150.2 |
| ENSMUSG00000022055 | neurofilament, light polypeptide [Source:MGI Symbol;Acc:MGI:97313] | Nefl | Enriched in retir | -1.82 | 3.564E-09 | 7224 |
| ENSMUSG00000032908 | sphingosine-1-phosphate phosphatase 2 [Source:MGI Symbol;Acc:MGI:3589109] | Sgpp2 | Enriched in retir | -1.895 | 2.173E-07 | 115.4 |
| ENSMUSG00000025432 | advillin [Source:MGI Symbol;Acc:MGI:1333798] | Avil | Enriched in retir | -1.984 | 0.02364 | 4.817 |
| ENSMUSG00000056665 | thioesterase superfamily member 6 [Source:MGI Symbol;Acc:MGI:1925301] | Them6 | Enriched in retir | -2.019 | 7.718E-10 | 58.01 |

| | | | | | | |
|---|---|---|---|---|---|---|
| ENSMUSG00000048899 | ribosomal modification protein rimK-like family member A [Source:MGI Symbol;Acc:MGI:3040686] | Rimkla | Enriched in retir | -2.032 | 4.878E-19 | 161 |
| ENSMUSG00000021728 | embigin [Source:MGI Symbol;Acc:MGI:95321] | Emb | Enriched in retir | -2.065 | 6.156E-10 | 88.85 |
| ENSMUSG00000041147 | breast cancer 2, early onset [Source:MGI Symbol;Acc:MGI:109337] | Brca2 | Enriched in retir | -2.078 | 3.462E-10 | 53.68 |
| ENSMUSG00000038530 | regulator of G-protein signaling 4 [Source:MGI Symbol;Acc:MGI:108409] | Rgs4 | Enriched in retir | -2.09 | 0.000000063 | 578.5 |
| ENSMUSG00000028476 | reversion-inducing-cysteine-rich protein with kazal motifs [Source:MGI Symbol;Acc:MGI:1855698] | Reck | Enriched in retir | -2.098 | 0.000001055 | 71.55 |
| ENSMUSG00000002980 | basal cell adhesion molecule [Source:MGI Symbol;Acc:MGI:1929940] | Bcam | Enriched in retir | -2.136 | 2.038E-07 | 83.63 |
| ENSMUSG00000040489 | SRY (sex determining region Y)-box 30 [Source:MGI Symbol;Acc:MGI:1341157] | Sox30 | Enriched in retir | -2.175 | 0.006134 | 11.69 |
| ENSMUSG00000039158 | AT-hook transcription factor [Source:MGI Symbol;Acc:MGI:2140340] | Akna | Enriched in retir | -2.293 | 1.534E-15 | 134.6 |
| ENSMUSG00000026778 | protein kinase C, theta [Source:MGI Symbol;Acc:MGI:97601] | Prkcq | Enriched in retir | -2.435 | 6.915E-11 | 111.9 |
| ENSMUSG00000031734 | Iroquois related homeobox 3 [Source:MGI Symbol;Acc:MGI:1197522] | Irx3 | Enriched in retir | -2.454 | 0.000004301 | 16.38 |
| ENSMUSG00000020732 | RAB37, member RAS oncogene family [Source:MGI Symbol;Acc:MGI:1929945] | Rab37 | Enriched in retir | -2.544 | 0.00007396 | 50.93 |
| ENSMUSG00000030730 | ATPase, Ca++ transporting, cardiac muscle, fast twitch 1 [Source:MGI Symbol;Acc:MGI:105058] | Atp2a1 | Enriched in retir | -2.56 | 0.002667 | 39.3 |
| ENSMUSG00000030110 | ret proto-oncogene [Source:MGI Symbol;Acc:MGI:97902] | Ret | Enriched in retir | -2.586 | 7.964E-18 | 188.3 |
| ENSMUSG00000024942 | calpain 1 [Source:MGI Symbol;Acc:MGI:88263] | Capn1 | Enriched in retir | -2.592 | 0.00000003 | 59.73 |
| ENSMUSG00000096696 | zinc finger protein 960 [Source:MGI Symbol;Acc:MGI:3052731] | Zfp960 | Enriched in retir | -2.642 | 0.001081 | 4 |
| ENSMUSG00000058925 | coiled-coil domain containing 192 [Source:MGI Symbol;Acc:MGI:1922694] | Ccdc192 | Enriched in retir | -2.734 | 0.0004882 | 7.152 |
| ENSMUSG00000001504 | Iroquois homeobox 2 [Source:MGI Symbol;Acc:MGI:1197526] | Irx2 | Enriched in retir | -2.756 | 5.703E-08 | 24.13 |
| ENSMUSG00000035296 | sarcoglycan, gamma (dystrophin-associated glycoprotein) [Source:MGI Symbol;Acc:MGI:1346524] | Sgcg | Enriched in retir | -2.872 | 0.00001762 | 9.668 |
| ENSMUSG00000020908 | myosin, heavy polypeptide 3, skeletal muscle, embryonic [Source:MGI Symbol;Acc:MGI:1339709] | Myh3 | Enriched in retir | -3.009 | 0.0006168 | 16.98 |
| ENSMUSG00000035458 | troponin I, cardiac 3 [Source:MGI Symbol;Acc:MGI:98783] | Tnni3 | Enriched in retir | -3.015 | 0.0004448 | 5.883 |
| ENSMUSG00000020953 | cochlin [Source:MGI Symbol;Acc:MGI:1278313] | Coch | Enriched in retir | -3.232 | 1.046E-10 | 80.94 |
| ENSMUSG00000047746 | F-box protein 40 [Source:MGI Symbol;Acc:MGI:2443753] | Fbxo40 | Enriched in retir | -3.305 | 0.001097 | 15.08 |
| ENSMUSG00000025366 | extended synaptotagmin-like protein 1 [Source:MGI Symbol;Acc:MGI:1344426] | Esyt1 | Enriched in retir | -3.333 | 1.001E-15 | 159.4 |
| ENSMUSG00000042073 | abhydrolase domain containing 14b [Source:MGI Symbol;Acc:MGI:1923741] | Abhd14b | Enriched in retir | -3.366 | 6.278E-10 | 38.48 |
| ENSMUSG00000031737 | Iroquois homeobox 5 [Source:MGI Symbol;Acc:MGI:1859086] | Irx5 | Enriched in retir | -3.475 | 5.163E-12 | 31.09 |
| ENSMUSG00000031586 | RNA binding protein gene with multiple splicing [Source:MGI Symbol;Acc:MGI:1334446] | Rbpms | Enriched in retir | -3.543 | 8.868E-20 | 81.55 |
| ENSMUSG00000018470 | potassium voltage-gated channel, shaker-related subfamily, beta member 3 [Source:MGI Symbol;Acc:MGI:1336208] | Kcnab3 | Enriched in retir | -3.983 | 1.813E-20 | 53.86 |
| ENSMUSG00000090223 | Purkinje cell protein 4 [Source:MGI Symbol;Acc:MGI:97509] | Pcp4 | Enriched in retir | -4.019 | 2.244E-10 | 93.25 |
| ENSMUSG00000032446 | eomesodermin [Source:MGI Symbol;Acc:MGI:1201683] | Eomes | Enriched in retir | -4.083 | 5.717E-08 | 60.2 |
| ENSMUSG00000068220 | lectin, galactose binding, soluble 1 [Source:MGI Symbol;Acc:MGI:96777] | Lgals1 | Enriched in retir | -4.127 | 3.384E-10 | 30.47 |
| ENSMUSG00000005716 | parvalbumin [Source:MGI Symbol;Acc:MGI:97821] | Pvalb | Enriched in retir | -4.161 | 5.415E-09 | 26.7 |
| ENSMUSG00000032387 | RNA binding protein with multiple splicing 2 [Source:MGI Symbol;Acc:MGI:1919223] | Rbpms2 | Enriched in retir | -4.255 | 1.377E-24 | 47.72 |
| ENSMUSG00000051279 | growth differentiation factor 6 [Source:MGI Symbol;Acc:MGI:95689] | Gdf6 | Enriched in retir | -4.318 | 1.801E-07 | 11.98 |
| ENSMUSG00000002100 | myosin binding protein C, cardiac [Source:MGI Symbol;Acc:MGI:102844] | Mybpc3 | Enriched in retir | -4.972 | 0.0005616 | 4.298 |
| ENSMUSG00000079357 | predicted gene 11100 [Source:MGI Symbol;Acc:MGI:3779338] | Gm11100 | Enriched in retir | -5.261 | 0.00003531 | 4.575 |
| ENSMUSG00000069372 | cortexin 3 [Source:MGI Symbol;Acc:MGI:3642816] | Ctxn3 | Enriched in retir | -5.869 | 4.853E-14 | 38.01 |
| ENSMUSG00000023484 | peripherin [Source:MGI Symbol;Acc:MGI:97774] | Prph | Enriched in retir | -5.995 | 2.583E-13 | 64.1 |
| ENSMUSG00000029304 | secreted phosphoprotein 1 [Source:MGI Symbol;Acc:MGI:98389] | Spp1 | Enriched in retir | -6.125 | 2.257E-10 | 33.23 |
| ENSMUSG00000025389 | major intrinsic protein of lens fiber [Source:MGI Symbol;Acc:MGI:96990] | Mip | Enriched in retir | -7.441 | 0.0002181 | 36.23 |
| ENSMUSG00000031738 | Iroquois homeobox 6 [Source:MGI Symbol;Acc:MGI:1927642] | Irx6 | Enriched in retir | -8.489 | 1.391E-11 | 50.97 |
| ENSMUSG00000031688 | POU domain, class 4, transcription factor 2 [Source:MGI Symbol;Acc:MGI:102524] | Pou4f2 | Enriched in retir | -8.504 | 2.425E-11 | 45.2 |
| ENSMUSG00000042240 | crystallin, beta B2 [Source:MGI Symbol;Acc:MGI:88519] | Crybb2 | Enriched in retir | -8.841 | 2.531E-09 | 184.8 |
| ENSMUSG00000021799 | opsin 4 (melanopsin) [Source:MGI Symbol;Acc:MGI:1353425] | Opn4 | Enriched in retir | -9.15 | 3.807E-12 | 25.82 |
| ENSMUSG00000031965 | T-box 20 [Source:MGI Symbol;Acc:MGI:1888496] | Tbx20 | Enriched in retir | -9.312 | 1.595E-12 | 45.45 |

**Contrast numerator: P15_KO_SCN. Contrast denominator: P15_KO_retina.**

| ensembl_gene_id | description | mgi_symbol | Significance | deseq_logfc | deseq_adjp | deseq_basemean |
|---|---|---|---|---|---|---|
| ENSMUSG00000037727 | arginine vasopressin [Source:MGI Symbol;Acc:MGI:88121] | Avp | Enriched in SCN | 10.87 | 8.814E-07 | 32.86 |
| ENSMUSG00000040752 | myosin, heavy polypeptide 6, cardiac muscle, alpha [Source:MGI Symbol;Acc:MGI:97255] | Myh6 | Enriched in SCN | 5.353 | 2.536E-15 | 9.823 |
| ENSMUSG00000004892 | brevican [Source:MGI Symbol;Acc:MGI:1096385] | Bcan | Enriched in SCN | 4.379 | 4.244E-54 | 546.7 |
| ENSMUSG00000034226 | ras homolog family member V [Source:MGI Symbol;Acc:MGI:2444227] | Rhov | Enriched in SCN | 3.189 | 4.614E-18 | 54.65 |
| ENSMUSG00000045731 | prepronociceptin [Source:MGI Symbol;Acc:MGI:105308] | Pnoc | Enriched in SCN | 2.838 | 1.152E-11 | 50.56 |
| ENSMUSG00000022861 | diacylglycerol kinase, gamma [Source:MGI Symbol;Acc:MGI:105060] | Dgkg | Enriched in SCN | 2.745 | 4.838E-12 | 135 |
| ENSMUSG00000006711 | RIKEN cDNA D130043K22 gene [Source:MGI Symbol;Acc:MGI:3036268] | D130043K22Rik | Enriched in SCN | 2.556 | 8.997E-15 | 133.1 |
| ENSMUSG00000073406 | histocompatibility 2, blastocyst [Source:MGI Symbol;Acc:MGI:892004] | H2-Bl | Enriched in SCN | 2.554 | 0.001906 | 18.45 |
| ENSMUSG00000045348 | neuronal tyrosine-phosphorylated phosphoinositide 3-kinase adaptor 1 [Source:MGI Symbol;Acc:MGI:2443880] | Nyap1 | Enriched in SCN | 2.542 | 2.754E-36 | 501.1 |
| ENSMUSG00000032023 | junctional cadherin complex regulator [Source:MGI Symbol;Acc:MGI:1918239] | Jhy | Enriched in SCN | 2.531 | 0.0005437 | 5.097 |
| ENSMUSG00000066129 | kinase non-catalytic C-lobe domain (KIND) containing 1 [Source:MGI Symbol;Acc:MGI:1923734] | Kndc1 | Enriched in SCN | 2.377 | 2.808E-22 | 714.5 |
| ENSMUSG00000115293 | Eef1akmt4-endothelin converting enzyme 2 readthrough [Source:MGI Symbol;Acc:MGI:6121518] | Eef1ece2 | Enriched in SCN | 2.339 | 2.146E-07 | 44.17 |
| ENSMUSG00000062184 | heparan sulfate 6-O-sulfotransferase 2 [Source:MGI Symbol;Acc:MGI:1354959] | Hs6st2 | Enriched in SCN | 2.335 | 8.232E-14 | 235.4 |
| ENSMUSG00000034353 | receptor (calcitonin) activity modifying protein 1 [Source:MGI Symbol;Acc:MGI:1858418] | Ramp1 | Enriched in SCN | 2.294 | 9.814E-09 | 54.71 |
| ENSMUSG00000022840 | adenylate cyclase 5 [Source:MGI Symbol;Acc:MGI:99673] | Adcy5 | Enriched in SCN | 2.271 | 9.021E-19 | 2027 |
| ENSMUSG00000020734 | glutamate receptor, ionotropic, NMDA2C (epsilon 3) [Source:MGI Symbol;Acc:MGI:95822] | Grin2c | Enriched in SCN | 2.269 | 2.216E-08 | 127.2 |
| ENSMUSG00000031760 | metallothionein 3 [Source:MGI Symbol;Acc:MGI:97173] | Mt3 | Enriched in SCN | 2.139 | 8.074E-10 | 131.4 |
| ENSMUSG00000024873 | cornichon family AMPA receptor auxiliary protein 2 [Source:MGI Symbol;Acc:MGI:1277225] | Cnih2 | Enriched in SCN | 2.136 | 3.632E-28 | 220.1 |
| ENSMUSG00000037703 | leucine zipper, putative tumor suppressor family member 3 [Source:MGI Symbol;Acc:MGI:2656976] | Lzts3 | Enriched in SCN | 2.126 | 5.159E-19 | 572.4 |
| ENSMUSG00000039278 | proprotein convertase subtilisin/kexin type 1 inhibitor [Source:MGI Symbol;Acc:MGI:1353431] | Pcsk1n | Enriched in SCN | 2.104 | 1.327E-11 | 2810 |
| ENSMUSG00000004902 | solute carrier family 25 (mitochondrial carrier), member 18 [Source:MGI Symbol;Acc:MGI:1919053] | Slc25a18 | Enriched in SCN | 2.083 | 3.518E-07 | 68.62 |
| ENSMUSG00000096847 | transmembrane protein 151B [Source:MGI Symbol;Acc:MGI:2685169] | Tmem151b | Enriched in SCN | 2.03 | 1.615E-16 | 1096 |
| ENSMUSG00000033208 | S100 protein, beta polypeptide, neural [Source:MGI Symbol;Acc:MGI:98217] | S100b | Enriched in SCN | 2.021 | 0.000001011 | 86.39 |
| ENSMUSG00000040811 | echinoderm microtubule associated protein like 2 [Source:MGI Symbol;Acc:MGI:1919455] | Eml2 | Enriched in SCN | 1.973 | 1.35E-18 | 740.8 |
| ENSMUSG00000028782 | adhesion G protein-coupled receptor B2 [Source:MGI Symbol;Acc:MGI:2451244] | Adgrb2 | Enriched in SCN | 1.945 | 2.92E-19 | 1121 |
| ENSMUSG00000053297 | expressed sequence AI854703 [Source:MGI Symbol;Acc:MGI:2141510] | AI854703 | Enriched in SCN | 1.941 | 1.883E-15 | 90.96 |
| ENSMUSG00000004933 | megakaryocyte-associated tyrosine kinase [Source:MGI Symbol;Acc:MGI:99259] | Matk | Enriched in SCN | 1.941 | 4.853E-15 | 175.5 |
| ENSMUSG00000078794 | dishevelled-binding antagonist of beta-catenin 3 [Source:MGI Symbol;Acc:MGI:3654828] | Dact3 | Enriched in SCN | 1.928 | 4.316E-13 | 843 |
| ENSMUSG00000008496 | POU domain, class 2, transcription factor 2 [Source:MGI Symbol;Acc:MGI:101897] | Pou2f2 | Enriched in SCN | 1.874 | 5.99E-09 | 295.3 |
| ENSMUSG00000079364 | predicted gene 3558 [Source:MGI Symbol;Acc:MGI:3781735] | Gm3558 | Enriched in SCN | 1.844 | 0.008898 | 4.262 |
| ENSMUSG00000025375 | apoptosis-associated tyrosine kinase [Source:MGI Symbol;Acc:MGI:1197518] | Aatk | Enriched in SCN | 1.787 | 1.173E-15 | 1628 |
| ENSMUSG00000073433 | Rho GDP dissociation inhibitor (GDI) gamma [Source:MGI Symbol;Acc:MGI:108430] | Arhgdig | Enriched in SCN | 1.785 | 1.735E-09 | 185.1 |
| ENSMUSG00000093985 | predicted gene 10406 [Source:MGI Symbol;Acc:MGI:3711272] | Gm10406 | Enriched in SCN | 1.758 | 0.02129 | 6.055 |
| ENSMUSG00000026204 | protein tyrosine phosphatase, receptor type, N [Source:MGI Symbol;Acc:MGI:102765] | Ptprn | Enriched in SCN | 1.755 | 2.004E-08 | 802.8 |
| ENSMUSG00000040759 | CKLF-like MARVEL transmembrane domain containing 5 [Source:MGI Symbol;Acc:MGI:2447164] | Cmtm5 | Enriched in SCN | 1.75 | 0.0002621 | 15.12 |
| ENSMUSG00000041115 | IQ motif and Sec7 domain 2 [Source:MGI Symbol;Acc:MGI:3528396] | Iqsec2 | Enriched in SCN | 1.738 | 6.41E-24 | 368.7 |
| ENSMUSG00000004187 | kinesin family member C2 [Source:MGI Symbol;Acc:MGI:109187] | Kifc2 | Enriched in SCN | 1.728 | 7.198E-22 | 769 |
| ENSMUSG00000016346 | potassium voltage-gated channel, subfamily Q, member 2 [Source:MGI Symbol;Acc:MGI:1309503] | Kcnq2 | Enriched in SCN | 1.713 | 2.635E-15 | 1096 |
| ENSMUSG00000096775 | predicted gene 5796 [Source:MGI Symbol;Acc:MGI:3779522] | Gm5796 | Enriched in SCN | 1.699 | 0.03026 | 3.239 |
| ENSMUSG00000051146 | calcium/calmodulin-dependent protein kinase II inhibitor 2 [Source:MGI Symbol;Acc:MGI:1920297] | Camk2n2 | Enriched in SCN | 1.692 | 3.368E-15 | 513.1 |
| ENSMUSG00000000627 | sema domain, immunoglobulin domain (Ig), TM domain, and short cytoplasmic domain [Source:MGI Symbol;Acc:MGI:134005 | Sema4f | Enriched in SCN | 1.682 | 1.018E-11 | 125.4 |
| ENSMUSG00000002771 | glutamate receptor, ionotropic, NMDA2D (epsilon 4) [Source:MGI Symbol;Acc:MGI:95823] | Grin2d | Enriched in SCN | 1.676 | 6.27E-13 | 466.7 |
| ENSMUSG00000056413 | ArfGAP with dual PH domains 1 [Source:MGI Symbol;Acc:MGI:2442201] | Adap1 | Enriched in SCN | 1.657 | 8.085E-20 | 552.2 |
| ENSMUSG00000007944 | tetratricopeptide repeat domain 9B [Source:MGI Symbol;Acc:MGI:1920282] | Ttc9b | Enriched in SCN | 1.649 | 3.874E-09 | 147 |
| ENSMUSG00000033597 | CASK interacting protein 1 [Source:MGI Symbol;Acc:MGI:2442952] | Caskin1 | Enriched in SCN | 1.63 | 2.397E-18 | 1073 |
| ENSMUSG00000027223 | mitogen-activated protein kinase 8 interacting protein 1 [Source:MGI Symbol;Acc:MGI:1309464] | Mapk8ip1 | Enriched in SCN | 1.59 | 1.364E-13 | 1583 |
| ENSMUSG00000001227 | sema domain, transmembrane domain (TM), and cytoplasmic domain, (semaphorin) 6B [Source:MGI Symbol;Acc:MGI:120288 | Sema6b | Enriched in SCN | 1.585 | 4.846E-15 | 529.4 |
| ENSMUSG00000029120 | protein phosphatase 2, regulatory subunit B, gamma [Source:MGI Symbol;Acc:MGI:2442660] | Ppp2r2c | Enriched in SCN | 1.581 | 8.026E-18 | 1258 |
| ENSMUSG00000027220 | synaptotagmin XIII [Source:MGI Symbol;Acc:MGI:1933945] | Syt13 | Enriched in SCN | 1.581 | 2.308E-17 | 763.5 |
| ENSMUSG00000020331 | hyperpolarization-activated, cyclic nucleotide-gated K+ 2 [Source:MGI Symbol;Acc:MGI:1298210] | Hcn2 | Enriched in SCN | 1.566 | 1.904E-13 | 598.2 |
| ENSMUSG00000079396 | predicted gene 3411 [Source:MGI Symbol;Acc:MGI:3781589] | Gm3411 | Enriched in SCN | 1.546 | 0.011 | 6.451 |
| ENSMUSG00000042532 | golgi autoantigen, golgin subfamily a, 7B [Source:MGI Symbol;Acc:MGI:1918396] | Golga7b | Enriched in SCN | 1.532 | 9.281E-14 | 283.6 |
| ENSMUSG00000007594 | hyaluronan and proteoglycan link protein 4 [Source:MGI Symbol;Acc:MGI:2679531] | Hapln4 | Enriched in SCN | 1.531 | 0.002222 | 156.8 |
| ENSMUSG00000027523 | GNAS (guanine nucleotide binding protein, alpha stimulating) complex locus [Source:MGI Symbol;Acc:MGI:95777] | Gnas | Enriched in SCN | 1.522 | 3.584E-13 | 3503 |
| ENSMUSG00000044628 | ring finger protein 208 [Source:MGI Symbol;Acc:MGI:1916096] | Rnf208 | Enriched in SCN | 1.505 | 5.541E-11 | 128.2 |
| ENSMUSG00000041096 | TSPY-like 2 [Source:MGI Symbol;Acc:MGI:106244] | Tspyl2 | Enriched in SCN | 1.481 | 5.564E-10 | 347.4 |
| ENSMUSG00000039546 | adherens junction associated protein 1 [Source:MGI Symbol;Acc:MGI:2685419] | Ajap1 | Enriched in SCN | 1.469 | 5.64E-08 | 293.4 |
| ENSMUSG00000061451 | transmembrane protein 151A [Source:MGI Symbol;Acc:MGI:2147713] | Tmem151a | Enriched in SCN | 1.442 | 1.745E-09 | 350.6 |
| ENSMUSG00000022099 | dematin actin binding protein [Source:MGI Symbol;Acc:MGI:99670] | Dmtn | Enriched in SCN | 1.425 | 1.557E-12 | 452.3 |
| ENSMUSG00000029419 | apical junction component 1 [Source:MGI Symbol;Acc:MGI:2685842] | Ajm1 | Enriched in SCN | 1.389 | 2.231E-07 | 326.7 |
| ENSMUSG00000037813 | RIKEN cDNA D630003M21 gene [Source:MGI Symbol;Acc:MGI:3606579] | D630003M21Rik | Enriched in SCN | 1.381 | 0.001748 | 14.8 |
| ENSMUSG00000040867 | brain-enriched guanylate kinase-associated [Source:MGI Symbol;Acc:MGI:3044626] | Begain | Enriched in SCN | 1.347 | 0.000001398 | 445 |
| ENSMUSG00000060402 | carbohydrate sulfotransferase 8 [Source:MGI Symbol;Acc:MGI:1916197] | Chst8 | Enriched in SCN | 1.299 | 0.0004595 | 67.64 |
| ENSMUSG00000031860 | pre B cell leukemia homeobox 4 [Source:MGI Symbol;Acc:MGI:1931321] | Pbx4 | Enriched in SCN | 1.298 | 0.00003336 | 37.95 |
| ENSMUSG00000029095 | actin-binding LIM protein 2 [Source:MGI Symbol;Acc:MGI:2385758] | Ablim2 | Enriched in SCN | 1.297 | 0.00000021 | 268.3 |
| ENSMUSG00000039838 | solute carrier family 45, member 1 [Source:MGI Symbol;Acc:MGI:2653235] | Slc45a1 | Enriched in SCN | 1.272 | 2.831E-13 | 187.4 |
| ENSMUSG00000032946 | RAS, guanyl releasing protein 2 [Source:MGI Symbol;Acc:MGI:1333849] | Rasgrp2 | Enriched in SCN | 1.248 | 0.0001857 | 149.8 |
| ENSMUSG00000042388 | DLG associated protein 3 [Source:MGI Symbol;Acc:MGI:3039563] | Dlgap3 | Enriched in SCN | 1.241 | 7.184E-08 | 1641 |
| ENSMUSG00000057614 | guanine nucleotide binding protein (G protein), alpha inhibiting 1 [Source:MGI Symbol;Acc:MGI:95771] | Gnai1 | Enriched in SCN | 1.239 | 0.00003638 | 448.7 |
| ENSMUSG00000038244 | microtubule associated monooxygenase, calponin and LIM domain containing 2 [Source:MGI Symbol;Acc:MGI:2444947] | Mical2 | Enriched in SCN | 1.233 | 0.0001102 | 255.1 |
| ENSMUSG00000074652 | myosin, heavy chain 7B, cardiac muscle, beta [Source:MGI Symbol;Acc:MGI:3710243] | Myh7b | Enriched in SCN | 1.183 | 0.0004928 | 65.13 |
| ENSMUSG00000060843 | catenin (cadherin associated protein), alpha 3 [Source:MGI Symbol;Acc:MGI:2661445] | Ctnna3 | Enriched in SCN | 1.174 | 0.00001679 | 51.56 |
| ENSMUSG00000006651 | amyloid beta (A4) precursor-like protein 1 [Source:MGI Symbol;Acc:MGI:88046] | Aplp1 | Enriched in SCN | 1.134 | 0.000001415 | 2009 |
| ENSMUSG00000044024 | RELT-like 2 [Source:MGI Symbol;Acc:MGI:1918044] | Rell2 | Enriched in SCN | 1.132 | 4.978E-08 | 285.3 |
| ENSMUSG00000029875 | coiled-coil domain containing 184 [Source:MGI Symbol;Acc:MGI:2146066] | Ccdc184 | Enriched in SCN | 1.12 | 0.0003163 | 201 |
| ENSMUSG00000041263 | RUN and SH3 domain containing 1 [Source:MGI Symbol;Acc:MGI:1919546] | Rusc1 | Enriched in SCN | 1.11 | 0.0001217 | 448.9 |
| ENSMUSG00000070858 | predicted gene 1673 [Source:MGI Symbol;Acc:MGI:2686519] | Gm1673 | Enriched in SCN | 1.107 | 0.02218 | 55.45 |
| ENSMUSG00000047495 | DLG associated protein 2 [Source:MGI Symbol;Acc:MGI:2443181] | Dlgap2 | Enriched in SCN | 1.086 | 0.0191 | 201 |
| ENSMUSG00000053024 | contactin 2 [Source:MGI Symbol;Acc:MGI:104518] | Cntn2 | Enriched in SCN | 1.082 | 0.0001312 | 303.9 |
| ENSMUSG00000018865 | sulfotransferase family 4A, member 1 [Source:MGI Symbol;Acc:MGI:1888971] | Sult4a1 | Enriched in SCN | 1.051 | 0.000002527 | 795.2 |
| ENSMUSG00000031970 | dysbindin (dystrobrevin binding protein 1) domain containing 1 [Source:MGI Symbol;Acc:MGI:1919435] | Dbndd1 | Enriched in SCN | 1.049 | 0.0001061 | 82.55 |
| ENSMUSG00000055805 | formin-like 1 [Source:MGI Symbol;Acc:MGI:1888994] | Fmnl1 | Enriched in SCN | 1.041 | 7.698E-07 | 238.1 |
| ENSMUSG00000015981 | serine/threonine kinase 32C [Source:MGI Symbol;Acc:MGI:2385336] | Stk32c | Enriched in SCN | 1.005 | 0.000002163 | 234.7 |
| ENSMUSG00000030779 | retinoblastoma binding protein 6, ubiquitin ligase [Source:MGI Symbol;Acc:MGI:894835] | Rbbp6 | Enriched in retina | -1.043 | 0.00007453 | 688.1 |
| ENSMUSG00000019966 | kit ligand [Source:MGI Symbol;Acc:MGI:96974] | Kitl | Enriched in retina | -1.044 | 0.008061 | 453.1 |
| ENSMUSG00000039860 | serine/arginine repetitive matrix 3 [Source:MGI Symbol;Acc:MGI:1920309] | Srrm3 | Enriched in retina | -1.054 | 0.00001476 | 438.4 |
| ENSMUSG00000078580 | RIKEN cDNA E430018J23 gene [Source:MGI Symbol;Acc:MGI:2141981] | E430018J23Rik | Enriched in retina | -1.058 | 0.009003 | 19.55 |
| ENSMUSG00000045968 | transmembrane epididymal family member 2 [Source:MGI Symbol;Acc:MGI:1923273] | Teddm2 | Enriched in retina | -1.08 | 0.01203 | 15.55 |
| ENSMUSG00000018634 | corticotropin releasing hormone receptor 1 [Source:MGI Symbol;Acc:MGI:88498] | Crhr1 | Enriched in retina | -1.131 | 0.003879 | 47.56 |
| ENSMUSG00000050663 | TRH-degrading enzyme [Source:MGI Symbol;Acc:MGI:2384311] | Trhde | Enriched in retina | -1.192 | 0.0003887 | 309.3 |
| ENSMUSG00000036822 | topoisomerase I binding, arginine/serine-rich [Source:MGI Symbol;Acc:MGI:2146189] | Topors | Enriched in retina | -1.199 | 4.175E-09 | 180.5 |
| ENSMUSG00000048385 | scratch family zinc finger 1 [Source:MGI Symbol;Acc:MGI:2176606] | Scrt1 | Enriched in retina | -1.258 | 0.000007068 | 1666 |
| ENSMUSG00000025269 | apurinic/apyrimidinic endonuclease 2 [Source:MGI Symbol;Acc:MGI:1924872] | Apex2 | Enriched in retina | -1.26 | 0.01186 | 107 |
| ENSMUSG00000022055 | neurofilament, light polypeptide [Source:MGI Symbol;Acc:MGI:97313] | Nefl | Enriched in retina | -1.276 | 0.000172 | 7224 |
| ENSMUSG00000032418 | malic enzyme 1, NADP(+)-dependent, cytosolic [Source:MGI Symbol;Acc:MGI:97043] | Me1 | Enriched in retina | -1.276 | 0.0001188 | 251.4 |
| ENSMUSG00000026778 | protein kinase C, theta [Source:MGI Symbol;Acc:MGI:97601] | Prkcq | Enriched in retina | -1.291 | 0.001857 | 111.9 |
| ENSMUSG00000040591 | centriolar satellite-associated tubulin polyglutamylase complex regulator 1 [Source:MGI Symbol;Acc:MGI:1915079] | Cstpp1 | Enriched in retina | -1.292 | 0.00001136 | 214.8 |
| ENSMUSG00000044229 | neurexophilin and PC-esterase domain family, member 4 [Source:MGI Symbol;Acc:MGI:1924792] | Nxpe4 | Enriched in retina | -1.303 | 0.000008669 | 105.5 |
| ENSMUSG00000036995 | ArfGAP with SH3 domain, ankyrin repeat and PH domain 3 [Source:MGI Symbol;Acc:MGI:2684986] | Asap3 | Enriched in retina | -1.366 | 0.00009088 | 47.68 |
| ENSMUSG00000020994 | pinin [Source:MGI Symbol;Acc:MGI:1100514] | Pnn | Enriched in retina | -1.393 | 0.00002234 | 615.1 |
| ENSMUSG00000042115 | kelch domain containing 8A [Source:MGI Symbol;Acc:MGI:2442630] | Klhdc8a | Enriched in retina | -1.477 | 1.735E-08 | 181.2 |
| ENSMUSG00000002980 | basal cell adhesion molecule [Source:MGI Symbol;Acc:MGI:1929940] | Bcam | Enriched in retina | -1.529 | 0.0006255 | 83.63 |
| ENSMUSG00000040822 | RIKEN cDNA 1700123O20 gene [Source:MGI Symbol;Acc:MGI:1920893] | 1700123O20Rik | Enriched in retina | -1.529 | 0.000001261 | 37.7 |
| ENSMUSG00000048899 | ribosomal modification protein rimK-like family member A [Source:MGI Symbol;Acc:MGI:3040686] | Rimkla | Enriched in retina | -1.556 | 7.562E-11 | 161 |
| ENSMUSG00000039168 | death-associated protein [Source:MGI Symbol;Acc:MGI:1918190] | Dap | Enriched in retina | -1.582 | 0.0004611 | 35.43 |
| ENSMUSG00000004561 | methyltransferase like 17 [Source:MGI Symbol;Acc:MGI:1098577] | Mettl17 | Enriched in retina | -1.59 | 0.0008126 | 47.96 |
| ENSMUSG00000061462 | obscurin, cytoskeletal calmodulin and titin-interacting RhoGEF [Source:MGI Symbol;Acc:MGI:2681862] | Obscn | Enriched in retina | -1.612 | 0.002569 | 66.02 |
| ENSMUSG00000027014 | CWC22 spliceosome-associated protein [Source:MGI Symbol;Acc:MGI:2136773] | Cwc22 | Enriched in retina | -1.63 | 0.0002271 | 206.1 |
| ENSMUSG00000031257 | NADPH oxidase 1 [Source:MGI Symbol;Acc:MGI:2450016] | Nox1 | Enriched in retina | -1.731 | 0.003269 | 7.043 |
| ENSMUSG00000038457 | transmembrane protein 255B [Source:MGI Symbol;Acc:MGI:2685533] | Tmem255b | Enriched in retina | -1.749 | 5.612E-08 | 59.61 |
| ENSMUSG00000060427 | zinc finger protein 868 [Source:MGI Symbol;Acc:MGI:2142546] | Zfp868 | Enriched in retina | -1.75 | 0.000002784 | 33.49 |
| ENSMUSG00000027286 | leucine rich repeat containing 57 [Source:MGI Symbol;Acc:MGI:1913856] | Lrrc57 | Enriched in retina | -1.772 | 3.193E-08 | 34.71 |
| ENSMUSG00000028197 | collagen, type XXIV, alpha 1 [Source:MGI Symbol;Acc:MGI:1918605] | Col24a1 | Enriched in retina | -1.828 | 0.01188 | 14.18 |
| ENSMUSG00000030110 | ret proto-oncogene [Source:MGI Symbol;Acc:MGI:97902] | Ret | Enriched in retina | -1.848 | 6.991E-09 | 188.3 |
| ENSMUSG00000028248 | PNN interacting serine/arginine-rich [Source:MGI Symbol;Acc:MGI:1913875] | Pnisr | Enriched in retina | -1.868 | 0.0000171 | 892.2 |
| ENSMUSG00000040703 | cytochrome P450, family 2, subfamily s, polypeptide 1 [Source:MGI Symbol;Acc:MGI:1921384] | Cyp2s1 | Enriched in retina | -1.875 | 0.00005333 | 19.07 |
| ENSMUSG00000056596 | TMF1-regulated nuclear protein 1 [Source:MGI Symbol;Acc:MGI:1916789] | Trnp1 | Enriched in retina | -1.9 | 5.641E-11 | 640.5 |
| ENSMUSG00000072572 | solute carrier family 39 (zinc transporter), member 2 [Source:MGI Symbol;Acc:MGI:2684326] | Slc39a2 | Enriched in retina | -1.93 | 0.007388 | 19.9 |
| ENSMUSG00000023495 | poly(rC) binding protein 4 [Source:MGI Symbol;Acc:MGI:1890471] | Pcbp4 | Enriched in retina | -1.964 | 1.397E-28 | 550 |
| ENSMUSG00000029012 | origin recognition complex, subunit 5 [Source:MGI Symbol;Acc:MGI:1347044] | Orc5 | Enriched in retina | -2.036 | 0.00001425 | 47.78 |
| ENSMUSG00000029190 | DNA segment, Chr 5, ERATO Doi 579, expressed [Source:MGI Symbol;Acc:MGI:1261849] | D5Ertd579e | Enriched in retina | -2.046 | 0.00006366 | 1250 |
| ENSMUSG00000070697 | UTP3 small subunit processome component [Source:MGI Symbol;Acc:MGI:1919230] | Utp3 | Enriched in retina | -2.071 | 2.034E-11 | 195.6 |
| ENSMUSG00000031734 | Iroquois related homeobox 3 [Source:MGI Symbol;Acc:MGI:1197522] | Irx3 | Enriched in retina | -2.078 | 0.0001759 | 16.38 |
| ENSMUSG00000020953 | cochlin [Source:MGI Symbol;Acc:MGI:1278313] | Coch | Enriched in retina | -2.078 | 0.00004513 | 80.94 |
| ENSMUSG00000045275 | Leber congenital amaurosis 5-like [Source:MGI Symbol;Acc:MGI:3041157] | Lca5l | Enriched in retina | -2.099 | 0.00004008 | 37.83 |
| ENSMUSG00000060579 | fragile histidine triad gene [Source:MGI Symbol;Acc:MGI:1277947] | Fhit | Enriched in retina | -2.129 | 0.000009146 | 135.3 |

| | | | | | | |
|---|---|---|---|---|---|---|
| ENSMUSG00000045314 | sosondowah ankyrin repeat domain family member B [Source:MGI Symbol;Acc:MGI:1925338] | Sowahb | Enriched in retina | -2.181 | 0.0003262 | 9.564 |
| ENSMUSG00000046480 | sodium channel, type IV, beta [Source:MGI Symbol;Acc:MGI:2687406] | Scn4b | Enriched in retina | -2.182 | 0.0001485 | 167.6 |
| ENSMUSG00000001504 | Iroquois homeobox 2 [Source:MGI Symbol;Acc:MGI:1197526] | Irx2 | Enriched in retina | -2.207 | 0.00004067 | 24.13 |
| ENSMUSG00000043541 | cancer susceptibility candidate 1 [Source:MGI Symbol;Acc:MGI:2444480] | Casc1 | Enriched in retina | -2.208 | 0.0002844 | 13.93 |
| ENSMUSG00000046667 | RNA binding motif protein 12 B1 [Source:MGI Symbol;Acc:MGI:1919647] | Rbm12b1 | Enriched in retina | -2.235 | 0.000007149 | 35.33 |
| ENSMUSG00000021619 | autophagy related 10 [Source:MGI Symbol;Acc:MGI:1914045] | Atg10 | Enriched in retina | -2.289 | 0.00002603 | 39.83 |
| ENSMUSG00000042050 | dynein 2 intermediate chain 1 [Source:MGI Symbol;Acc:MGI:2445085] | Dync2i1 | Enriched in retina | -2.406 | 4.985E-16 | 228.8 |
| ENSMUSG00000102900 | predicted gene, 37811 [Source:MGI Symbol;Acc:MGI:5611039] | Gm37811 | Enriched in retina | -2.565 | 0.000005897 | 16.7 |
| ENSMUSG00000079418 | autophagy related 4A, cysteine peptidase [Source:MGI Symbol;Acc:MGI:2147903] | Atg4a | Enriched in retina | -2.758 | 0.005061 | 66.59 |
| ENSMUSG00000047632 | fibroblast growth factor binding protein 3 [Source:MGI Symbol;Acc:MGI:1919764] | Fgfbp3 | Enriched in retina | -2.824 | 0.00000202 | 20.4 |
| ENSMUSG00000044362 | coiled-coil domain containing 89 [Source:MGI Symbol;Acc:MGI:1917304] | Ccdc89 | Enriched in retina | -2.831 | 5.685E-07 | 17.87 |
| ENSMUSG00000031174 | retinitis pigmentosa GTPase regulator [Source:MGI Symbol;Acc:MGI:1344037] | Rpgr | Enriched in retina | -2.984 | 2.37E-09 | 194.1 |
| ENSMUSG00000025366 | extended synaptotagmin-like protein 1 [Source:MGI Symbol;Acc:MGI:1344426] | Esyt1 | Enriched in retina | -3.095 | 2.591E-12 | 159.4 |
| ENSMUSG00000024871 | double C2, gamma [Source:MGI Symbol;Acc:MGI:1926250] | Doc2g | Enriched in retina | -3.22 | 0.001351 | 6.218 |
| ENSMUSG00000096696 | zinc finger protein 960 [Source:MGI Symbol;Acc:MGI:3052731] | Zfp960 | Enriched in retina | -3.222 | 0.00003165 | 4 |
| ENSMUSG00000043727 | RIKEN cDNA F830045P16 gene [Source:MGI Symbol;Acc:MGI:3045317] | F830045P16Rik | Enriched in retina | -3.46 | 0.00004081 | 4.358 |
| ENSMUSG00000016356 | collagen, type XX, alpha 1 [Source:MGI Symbol;Acc:MGI:1920618] | Col20a1 | Enriched in retina | -3.688 | 6.097E-12 | 25.93 |
| ENSMUSG00000035314 | glycerophosphodiester phosphodiesterase domain containing 5 [Source:MGI Symbol;Acc:MGI:2686926] | Gdpd5 | Enriched in retina | -3.756 | 5.972E-24 | 328.8 |
| ENSMUSG00000025515 | mucin 2 [Source:MGI Symbol;Acc:MGI:1339364] | Muc2 | Enriched in retina | -3.778 | 0.00000209 | 78.5 |
| ENSMUSG00000045319 | proline and serine rich 2 [Source:MGI Symbol;Acc:MGI:2442238] | Proser2 | Enriched in retina | -3.788 | 2.201E-21 | 42.91 |
| ENSMUSG00000035296 | sarcoglycan, gamma (dystrophin-associated glycoprotein) [Source:MGI Symbol;Acc:MGI:1346524] | Sgcg | Enriched in retina | -3.834 | 6.891E-09 | 9.668 |
| ENSMUSG00000048337 | neuropeptide Y receptor Y4 [Source:MGI Symbol;Acc:MGI:105374] | Npy4r | Enriched in retina | -3.879 | 0.04839 | 1.332 |
| ENSMUSG00000096883 | shisa family member 8 [Source:MGI Symbol;Acc:MGI:2146080] | Shisa8 | Enriched in retina | -4.1 | 1.556E-20 | 27.13 |
| ENSMUSG00000042073 | abhydrolase domain containing 14b [Source:MGI Symbol;Acc:MGI:1923741] | Abhd14b | Enriched in retina | -4.213 | 9.922E-14 | 38.48 |
| ENSMUSG00000029601 | IQ motif containing D [Source:MGI Symbol;Acc:MGI:1922982] | Iqcd | Enriched in retina | -4.266 | 3.127E-10 | 36.37 |
| ENSMUSG00000035458 | troponin I, cardiac 3 [Source:MGI Symbol;Acc:MGI:98783] | Tnni3 | Enriched in retina | -4.428 | 0.0009313 | 5.883 |
| ENSMUSG00000040489 | SRY (sex determining region Y)-box 30 [Source:MGI Symbol;Acc:MGI:1341157] | Sox30 | Enriched in retina | -4.461 | 0.000002125 | 11.69 |
| ENSMUSG00000020732 | RAB37, member RAS oncogene family [Source:MGI Symbol;Acc:MGI:1929945] | Rab37 | Enriched in retina | -4.464 | 0.00007421 | 50.93 |
| ENSMUSG00000005716 | parvalbumin [Source:MGI Symbol;Acc:MGI:97821] | Pvalb | Enriched in retina | -4.56 | 2.142E-08 | 26.7 |
| ENSMUSG00000002100 | myosin binding protein C, cardiac [Source:MGI Symbol;Acc:MGI:102844] | Mybpc3 | Enriched in retina | -4.671 | 0.001611 | 4.298 |
| ENSMUSG00000069372 | cortexin 3 [Source:MGI Symbol;Acc:MGI:3642816] | Ctxn3 | Enriched in retina | -4.764 | 6.724E-10 | 38.01 |
| ENSMUSG00000029304 | secreted phosphoprotein 1 [Source:MGI Symbol;Acc:MGI:98389] | Spp1 | Enriched in retina | -5.527 | 4.779E-07 | 33.23 |
| ENSMUSG00000074037 | melanocortin 1 receptor [Source:MGI Symbol;Acc:MGI:99456] | Mc1r | Enriched in retina | -5.9 | 4.786E-12 | 11.83 |
| ENSMUSG00000023484 | peripherin [Source:MGI Symbol;Acc:MGI:97774] | Prph | Enriched in retina | -6.266 | 9.79E-11 | 64.1 |
| ENSMUSG00000108398 | predicted gene, 30191 [Source:MGI Symbol;Acc:MGI:5589350] | Gm30191 | Enriched in retina | -6.71 | 5.618E-07 | 4.829 |
| ENSMUSG00000073551 | serine peptidase inhibitor, Kazal type 13 [Source:MGI Symbol;Acc:MGI:3642511] | Spink13 | Enriched in retina | -6.826 | 2.397E-07 | 7.681 |
| ENSMUSG00000031688 | POU domain, class 4, transcription factor 2 [Source:MGI Symbol;Acc:MGI:102524] | Pou4f2 | Enriched in retina | -7.745 | 6.3E-10 | 45.2 |
| ENSMUSG00000031738 | Iroquois homeobox 6 [Source:MGI Symbol;Acc:MGI:1927642] | Irx6 | Enriched in retina | -7.905 | 2.171E-10 | 50.97 |
| ENSMUSG00000045493 | basic helix-loop-helix family, member e23 [Source:MGI Symbol;Acc:MGI:2153710] | Bhlhe23 | Enriched in retina | -8.543 | 2.993E-11 | 30.34 |
| ENSMUSG00000070870 | crystallin, gamma E [Source:MGI Symbol;Acc:MGI:88525] | Cryge | Enriched in retina | -8.643 | 0.0000205 | 45.26 |
| ENSMUSG00000114882 | predicted gene, 48581 [Source:MGI Symbol;Acc:MGI:6098148] | Gm48581 | Enriched in retina | -8.791 | 7.363E-10 | 15.29 |
| ENSMUSG00000051985 | immunoglobulin-like and fibronectin type III domain containing 1 [Source:MGI Symbol;Acc:MGI:3045352] | Igfn1 | Enriched in retina | -9.611 | 3.124E-12 | 21.94 |
| ENSMUSG00000042240 | crystallin, beta B2 [Source:MGI Symbol;Acc:MGI:88519] | Crybb2 | Enriched in retina | -10.39 | 1.544E-11 | 184.8 |
