## Supplementary material for "Melanopsin regulates axonal translation underlying retinohypothalamic circuit assembly": Table_S3_TRAP_seq_genotype_contrasts_significant_genes

**Contrast numerator: P8_KO_dLGN. Contrast denominator: P8_Het_dLGN.**

| ensembl_gene_id | description | mgi_symbol | Significance | deseq_logfc | deseq_adjp | deseq_basemean |
|---|---|---|---|---|---|---|
| ENSMUSG00000041144 | dynein, axonemal, heavy chain 7B [Source:MGI Symbol;Acc:MGI:2684953] | Dnah7b | Enriched in KO | 5.073 | 0.00004399 | 449.4 |
| ENSMUSG00000051864 | TBC1 domain family, member 22a [Source:MGI Symbol;Acc:MGI:1289265] | Tbc1d22a | Enriched in KO | 4.773 | 2.716E-09 | 351.4 |
| ENSMUSG00000028461 | coiled-coil domain containing 107 [Source:MGI Symbol;Acc:MGI:1913423] | Ccdc107 | Enriched in KO | 3.939 | 0.001348 | 309.5 |
| ENSMUSG00000028744 | solute carrier family 66 member 1 [Source:MGI Symbol;Acc:MGI:2384837] | Slc66a1 | Enriched in KO | 3.769 | 3.959E-08 | 195.7 |
| ENSMUSG00000039115 | integrin alpha 9 [Source:MGI Symbol;Acc:MGI:104756] | Itga9 | Enriched in KO | 3.637 | 0.001896 | 1811 |
| ENSMUSG00000020107 | anaphase promoting complex subunit 16 [Source:MGI Symbol;Acc:MGI:1289325] | Anapc16 | Enriched in KO | 3.027 | 6.285E-15 | 134.3 |
| ENSMUSG00000027104 | activating transcription factor 2 [Source:MGI Symbol;Acc:MGI:109349] | Atf2 | Enriched in KO | 2.692 | 3.965E-08 | 703.5 |
| ENSMUSG00000052372 | interleukin 1 receptor accessory protein-like 1 [Source:MGI Symbol;Acc:MGI:2687319] | Il1rapl1 | Enriched in KO | 2.147 | 0.02778 | 1897 |
| ENSMUSG00000079184 | M-phase phosphoprotein 8 [Source:MGI Symbol;Acc:MGI:1922589] | Mphosph8 | Enriched in KO | 2.006 | 0.0001123 | 685.9 |
| ENSMUSG00000026687 | aldehyde dehydrogenase 9, subfamily A1 [Source:MGI Symbol;Acc:MGI:1861622] | Aldh9a1 | Enriched in KO | 1.871 | 1.512E-08 | 102.9 |
| ENSMUSG00000005142 | mannosidase 2, alpha B1 [Source:MGI Symbol;Acc:MGI:107286] | Man2b1 | Enriched in KO | 1.517 | 0.000007308 | 85.64 |
| ENSMUSG00000020982 | nuclear export mediator factor [Source:MGI Symbol;Acc:MGI:1918305] | Nemf | Enriched in KO | 1.419 | 0.01491 | 270.7 |
| ENSMUSG00000046058 | EP300 interacting inhibitor of differentiation 2 [Source:MGI Symbol;Acc:MGI:2681174] | Eid2 | Enriched in KO | 1.404 | 0.006114 | 137.6 |
| ENSMUSG00000021156 | zinc finger, MYND domain containing 11 [Source:MGI Symbol;Acc:MGI:1913755] | Zmynd11 | Enriched in KO | 1.398 | 0.0005577 | 980.2 |
| ENSMUSG00000030706 | mitochondrial ribosomal protein L48 [Source:MGI Symbol;Acc:MGI:1289321] | Mrpl48 | Enriched in KO | 1.334 | 0.01487 | 153.1 |
| ENSMUSG00000056310 | tRNA-yW synthesizing protein 1 homolog (S. cerevisiae) [Source:MGI Symbol;Acc:MGI:214116 | Tyw1 | Enriched in KO | 1.287 | 0.00153 | 150.1 |
| ENSMUSG00000035228 | coiled-coil domain containing 106 [Source:MGI Symbol;Acc:MGI:2385900] | Ccdc106 | Enriched in KO | 1.17 | 0.00153 | 90.05 |
| ENSMUSG00000056305 | ubiquitin specific peptidase 39 [Source:MGI Symbol;Acc:MGI:107622] | Usp39 | Enriched in KO | 1.167 | 0.001879 | 113.7 |
| ENSMUSG00000052253 | zinc finger protein 622 [Source:MGI Symbol;Acc:MGI:1289282] | Zfp622 | Enriched in KO | 1.086 | 0.001348 | 157.1 |
| ENSMUSG00000020993 | trafficking protein particle complex 6B [Source:MGI Symbol;Acc:MGI:1925482] | Trappc6b | Enriched in KO | 1.072 | 0.05083 | 162.5 |
| ENSMUSG00000036087 | SLAIN motif family, member 2 [Source:MGI Symbol;Acc:MGI:1923241] | Slain2 | Enriched in KO | 1.063 | 0.00153 | 209.6 |
| ENSMUSG00000026558 | uridine-cytidine kinase 2 [Source:MGI Symbol;Acc:MGI:1931744] | Uck2 | Enriched in KO | 1.062 | 0.00153 | 349 |
| ENSMUSG00000041837 | programmed cell death 7 [Source:MGI Symbol;Acc:MGI:1859170] | Pdcd7 | Enriched in KO | 1.056 | 0.00153 | 140.5 |
| ENSMUSG00000029833 | tripartite motif-containing 24 [Source:MGI Symbol;Acc:MGI:109275] | Trim24 | Enriched in KO | 1.018 | 0.02154 | 198.7 |
| ENSMUSG00000021685 | orthopedia homeobox [Source:MGI Symbol;Acc:MGI:99835] | Otp | Enriched in Het | -11.78 | 0.03699 | 81.46 |

**Contrast numerator: P8_KO_retina. Contrast denominator: P8_Het_retina.**

| ensembl_gene_id | description | mgi_symbol | Significance | deseq_logfc | deseq_adjp | deseq_basemean |
|---|---|---|---|---|---|---|
| ENSMUSG00000072476 | predicted pseudogene 9008 [Source:MGI Symbol;Acc:MGI:3644000] | Gm9008 | KO_enriched | 8.366 | 2.311E-07 | 8.19 |
| ENSMUSG00000034883 | leucine rich repeat protein 1 [Source:MGI Symbol;Acc:MGI:1916956] | Lrr1 | KO_enriched | 5.585 | 0.0435 | 14.15 |
| ENSMUSG00000073991 | cyclic nucleotide binding domain containing 1 [Source:MGI Symbol;Acc:MGI:3650508] | Cnbd1 | KO_enriched | 2.312 | 0.03115 | 9.142 |
| ENSMUSG00000021799 | opsin 4 (melanopsin) [Source:MGI Symbol;Acc:MGI:1353425] | Opn4 | Het_enriched | -5.451 | 1.137E-20 | 25.82 |

**Contrast numerator: P8_Enriched in KO_SCN. Contrast denominator: P8_Het_SCN.**

| ensembl_gene_id | description | mgi_symbol | Significance | deseq_logfc | deseq_adjp | deseq_basemean |
|---|---|---|---|---|---|---|
| ENSMUSG00000052428 | transmembrane and coiled-coil domains 1 [Source:MGI Symbol;Acc:MGI:1921173] | Tmco1 | Enriched in KO | 3.093 | 0.00002976 | 95.8 |
| ENSMUSG00000000560 | gamma-aminobutyric acid (GABA) A receptor, subunit alpha 2 [Source:MGI Symbol;Acc:MGI:95614] | Gabra2 | Enriched in KO | 2.484 | 3.771E-07 | 394.2 |
| ENSMUSG00000029442 | WD repeat domain 66 [Source:MGI Symbol;Acc:MGI:1918495] | Wdr66 | Enriched in KO | 2.258 | 0.00002207 | 54.47 |
| ENSMUSG00000027014 | CWC22 spliceosome-associated protein [Source:MGI Symbol;Acc:MGI:2136773] | Cwc22 | Enriched in KO | 1.934 | 0.002539 | 206.1 |
| ENSMUSG00000029245 | Eph receptor A5 [Source:MGI Symbol;Acc:MGI:99654] | Epha5 | Enriched in KO | 1.722 | 0.0005972 | 636.4 |
| ENSMUSG00000025607 | coatomer protein complex, subunit gamma 2 [Source:MGI Symbol;Acc:MGI:1858683] | Copg2 | Enriched in KO | 1.568 | 0.008205 | 319.5 |
| ENSMUSG00000020107 | anaphase promoting complex subunit 16 [Source:MGI Symbol;Acc:MGI:1289325] | Anapc16 | Enriched in KO | 1.55 | 0.003244 | 134.3 |
| ENSMUSG00000042606 | HIRA interacting protein 3 [Source:MGI Symbol;Acc:MGI:2142364] | Hirip3 | Enriched in KO | 1.468 | 0.004697 | 132.7 |
| ENSMUSG00000052727 | microtubule-associated protein 1B [Source:MGI Symbol;Acc:MGI:1306778] | Map1b | Enriched in Het | -1.003 | 0.002385 | 25090 |
| ENSMUSG00000009575 | chromobox 5 [Source:MGI Symbol;Acc:MGI:109372] | Cbx5 | Enriched in Het | -1.053 | 0.02412 | 2146 |
| ENSMUSG00000021687 | secretory carrier membrane protein 1 [Source:MGI Symbol;Acc:MGI:1349480] | Scamp1 | Enriched in Het | -1.078 | 0.004055 | 570.2 |
| ENSMUSG00000021770 | sterile alpha motif domain containing 8 [Source:MGI Symbol;Acc:MGI:1914880] | Samd8 | Enriched in Het | -1.107 | 0.009049 | 676.8 |
| ENSMUSG00000021395 | spindlin 1 [Source:MGI Symbol;Acc:MGI:109242] | Spin1 | Enriched in Het | -1.126 | 0.01393 | 935.2 |
| ENSMUSG00000025134 | Aly/REF export factor [Source:MGI Symbol;Acc:MGI:1341044] | Alyref | Enriched in Het | -1.126 | 0.01698 | 194.3 |
| ENSMUSG00000022812 | glycogen synthase kinase 3 beta [Source:MGI Symbol;Acc:MGI:1861437] | Gsk3b | Enriched in Het | -1.134 | 0.02873 | 2445 |
| ENSMUSG00000054766 | SET nuclear oncogene [Source:MGI Symbol;Acc:MGI:1860267] | Set | Enriched in Het | -1.134 | 0.01069 | 841.2 |
| ENSMUSG00000031584 | glutathione reductase [Source:MGI Symbol;Acc:MGI:95804] | Gsr | Enriched in Het | -1.143 | 0.00984 | 253.9 |
| ENSMUSG00000032905 | autophagy related 12 [Source:MGI Symbol;Acc:MGI:1914776] | Atg12 | Enriched in Het | -1.151 | 0.002087 | 183.2 |
| ENSMUSG00000027983 | cytochrome P450, family 2, subfamily u, polypeptide 1 [Source:MGI Symbol;Acc:MGI:1918769] | Cyp2u1 | Enriched in Het | -1.153 | 0.003381 | 119.4 |
| ENSMUSG00000042105 | inositol polyphosphate-5-phosphatase F [Source:MGI Symbol;Acc:MGI:2141867] | Inpp5f | Enriched in Het | -1.166 | 0.003687 | 994.9 |
| ENSMUSG00000095139 | POU domain, class 3, transcription factor 2 [Source:MGI Symbol;Acc:MGI:101895] | Pou3f2 | Enriched in Het | -1.166 | 0.02412 | 124.7 |
| ENSMUSG00000024579 | prenylcysteine oxidase 1 like [Source:MGI Symbol;Acc:MGI:3606062] | Pcyox1l | Enriched in Het | -1.19 | 0.01069 | 127.8 |
| ENSMUSG00000028030 | TBC1 domain containing kinase [Source:MGI Symbol;Acc:MGI:2445052] | Tbck | Enriched in Het | -1.205 | 0.00185 | 511.5 |
| ENSMUSG00000017929 | UDP-Gal:betaGlcNAc beta 1,4-galactosyltransferase, polypeptide 5 [Source:MGI Symbol;Acc:MGI:1927169] | B4galt5 | Enriched in Het | -1.209 | 0.002539 | 566.4 |
| ENSMUSG00000032727 | MIER family member 3 [Source:MGI Symbol;Acc:MGI:2442317] | Mier3 | Enriched in Het | -1.211 | 0.0005972 | 151.2 |
| ENSMUSG00000062929 | cofilin 2, muscle [Source:MGI Symbol;Acc:MGI:101763] | Cfl2 | Enriched in Het | -1.223 | 0.01597 | 177.2 |
| ENSMUSG00000030067 | forkhead box P1 [Source:MGI Symbol;Acc:MGI:1914004] | Foxp1 | Enriched in Het | -1.23 | 0.006513 | 337.2 |
| ENSMUSG00000022180 | solute carrier family 7 (cationic amino acid transporter, y+ system), member 8 [Source:MGI Symbol;Acc:MGI:1355323] | Slc7a8 | Enriched in Het | -1.268 | 0.007505 | 249.8 |
| ENSMUSG00000032076 | cell adhesion molecule 1 [Source:MGI Symbol;Acc:MGI:1889272] | Cadm1 | Enriched in Het | -1.326 | 0.0002145 | 2724 |
| ENSMUSG00000027162 | lin-7 homolog C, crumbs cell polarity complex component [Source:MGI Symbol;Acc:MGI:1330839] | Lin7c | Enriched in Het | -1.331 | 0.001631 | 403.1 |
| ENSMUSG00000039770 | yippee like 5 [Source:MGI Symbol;Acc:MGI:1916937] | Ypel5 | Enriched in Het | -1.331 | 0.008794 | 207.1 |
| ENSMUSG00000024501 | dihydropyrimidinase-like 3 [Source:MGI Symbol;Acc:MGI:1349762] | Dpysl3 | Enriched in Het | -1.362 | 0.00424 | 3490 |
| ENSMUSG00000069662 | myristoylated alanine rich protein kinase C substrate [Source:MGI Symbol;Acc:MGI:96907] | Marcks | Enriched in Het | -1.364 | 0.02035 | 2471 |
| ENSMUSG00000029064 | guanine nucleotide binding protein (G protein), beta 1 [Source:MGI Symbol;Acc:MGI:95781] | Gnb1 | Enriched in Het | -1.368 | 0.00004537 | 5932 |
| ENSMUSG00000028381 | UDP-glucose ceramide glucosyltransferase [Source:MGI Symbol;Acc:MGI:1332243] | Ugcg | Enriched in Het | -1.414 | 0.008077 | 361.5 |
| ENSMUSG00000031226 | polysaccharide biosynthesis domain containing 1 [Source:MGI Symbol;Acc:MGI:1914933] | Pbdc1 | Enriched in Het | -1.438 | 0.00984 | 236 |
| ENSMUSG00000027879 | SEC22 homolog B, vesicle trafficking protein [Source:MGI Symbol;Acc:MGI:1338759] | Sec22b | Enriched in Het | -1.458 | 0.004237 | 253.4 |
| ENSMUSG00000032479 | microtubule-associated protein 4 [Source:MGI Symbol;Acc:MGI:97178] | Map4 | Enriched in Het | -1.467 | 0.003785 | 2983 |
| ENSMUSG00000031302 | neuroligin 3 [Source:MGI Symbol;Acc:MGI:2444609] | Nlgn3 | Enriched in Het | -1.476 | 0.004511 | 1941 |
| ENSMUSG00000039542 | neural cell adhesion molecule 1 [Source:MGI Symbol;Acc:MGI:97281] | Ncam1 | Enriched in Het | -1.546 | 0.0002145 | 5804 |
| ENSMUSG00000071359 | TATA box binding protein-like 1 [Source:MGI Symbol;Acc:MGI:1339946] | Tbpl1 | Enriched in Het | -1.551 | 0.0005322 | 195 |
| ENSMUSG00000021974 | fibroblast growth factor 9 [Source:MGI Symbol;Acc:MGI:104723] | Fgf9 | Enriched in Het | -1.587 | 0.006937 | 263.9 |
| ENSMUSG00000029580 | actin, beta [Source:MGI Symbol;Acc:MGI:87904] | Actb | Enriched in Het | -1.604 | 0.01457 | 6299 |
| ENSMUSG00000021139 | predicted gene 20498 [Source:MGI Symbol;Acc:MGI:5141963] | Gm20498 | Enriched in Het | -1.715 | 0.00984 | 272.3 |
| ENSMUSG00000108358 | predicted gene 44509 [Source:MGI Symbol;Acc:MGI:5753085] | Gm44509 | Enriched in Het | -1.726 | 0.004946 | 119 |
| ENSMUSG00000022048 | dihydropyrimidinase-like 2 [Source:MGI Symbol;Acc:MGI:1349763] | Dpysl2 | Enriched in Het | -1.731 | 0.0005972 | 6238 |
| ENSMUSG00000026434 | nuclear casein kinase and cyclin-dependent kinase substrate 1 [Source:MGI Symbol;Acc:MGI:1934811] | Nucks1 | Enriched in Het | -1.766 | 0.002385 | 1384 |
| ENSMUSG00000032262 | elongation of very long chain fatty acids (FEN1/Elo2, SUR4/Elo3, yeast)-like 4 [Source:MGI Symbol;Acc:MGI:1933331] | Elovl4 | Enriched in Het | -1.774 | 0.00006532 | 520.9 |
| ENSMUSG00000039114 | neuritin 1 [Source:MGI Symbol;Acc:MGI:1915654] | Nrn1 | Enriched in Het | -2.019 | 0.0001522 | 742.9 |
| ENSMUSG00000075324 | fidgetin [Source:MGI Symbol;Acc:MGI:1890647] | Fign | Enriched in Het | -2.149 | 0.0001635 | 251.6 |
| ENSMUSG00000064179 | troponin T1, skeletal, slow [Source:MGI Symbol;Acc:MGI:1333868] | Tnnt1 | Enriched in Het | -2.749 | 0.01741 | 308.7 |
| ENSMUSG00000070570 | solute carrier family 17 (sodium-dependent inorganic phosphate cotransporter), member 7 [Source:MGI Symbol;Acc:MGI:192021 | Slc17a7 | Enriched in Het | -3.077 | 0.00006532 | 1629 |
| ENSMUSG00000018486 | wingless-type MMTV integration site family, member 9B [Source:MGI Symbol;Acc:MGI:1197020] | Wnt9b | Enriched in Het | -4.508 | 0.00984 | 165.2 |

**Contrast numerator: P15_KO_dLGN. Contrast denominator: P15_Het_dLGN.**

| ensembl_gene_id | description | mgi_symbol | Significance | deseq_logfc | deseq_adjp | deseq_basemean |
|---|---|---|---|---|---|---|
| ENSMUSG00000034452 | solute carrier family 24 (sodium/potassium/calcium exchanger), member 1 [Source:MGI Symbol;Acc:MGI:2384871] | Slc24a1 | Enriched in KO | 6.392 | 0.001164 | 1018 |
| ENSMUSG00000044375 | photoreceptor cilium actin regulator [Source:MGI Symbol;Acc:MGI:2385061] | Pcare | Enriched in KO | 6.351 | 0.001204 | 962.8 |
| ENSMUSG00000031450 | G protein-coupled receptor kinase 1 [Source:MGI Symbol;Acc:MGI:1345146] | Grk1 | Enriched in KO | 5.877 | 0.002484 | 1629 |
| ENSMUSG00000023978 | peripherin 2 [Source:MGI Symbol;Acc:MGI:102791] | Prph2 | Enriched in KO | 5.539 | 0.006131 | 1752 |
| ENSMUSG00000110344 | small integral membrane protein 36 [Source:MGI Symbol;Acc:MGI:5804831] | Smim36 | Enriched in KO | 5.364 | 0.01548 | 107.5 |
| ENSMUSG00000041578 | cone-rod homeobox [Source:MGI Symbol;Acc:MGI:1194883] | Crx | Enriched in KO | 5.338 | 0.009016 | 944.1 |
| ENSMUSG00000034837 | G protein subunit alpha transducin 1 [Source:MGI Symbol;Acc:MGI:95778] | Gnat1 | Enriched in KO | 5.306 | 0.0164 | 1840 |
| ENSMUSG00000040632 | neural retina leucine zipper gene [Source:MGI Symbol;Acc:MGI:102567] | Nrl | Enriched in KO | 5.289 | 0.003514 | 505.3 |
| ENSMUSG00000045776 | leucine-rich repeats and transmembrane domains 1 [Source:MGI Symbol;Acc:MGI:2442106] | Lrtm1 | Enriched in KO | 5.25 | 0.02529 | 127.3 |
| ENSMUSG00000024575 | phosphodiesterase 6A, cGMP-specific, rod, alpha [Source:MGI Symbol;Acc:MGI:97524] | Pde6a | Enriched in KO | 5.23 | 0.007748 | 1301 |
| ENSMUSG00000031293 | retinoschisis (X-linked, juvenile) 1 (human) [Source:MGI Symbol;Acc:MGI:1336189] | Rs1 | Enriched in KO | 5.229 | 0.007748 | 1028 |
| ENSMUSG00000030523 | transient receptor potential cation channel, subfamily M, member 1 [Source:MGI Symbol;Acc:MGI:1330305] | Trpm1 | Enriched in KO | 5.129 | 0.01548 | 420.9 |
| ENSMUSG00000056055 | S-antigen, retina and pineal gland (arrestin) [Source:MGI Symbol;Acc:MGI:98227] | Sag | Enriched in KO | 5.083 | 0.001204 | 2015 |
| ENSMUSG00000030324 | rhodopsin [Source:MGI Symbol;Acc:MGI:97914] | Rho | Enriched in KO | 5.021 | 0.003618 | 5636 |
| ENSMUSG00000040554 | aryl hydrocarbon receptor-interacting protein-like 1 [Source:MGI Symbol;Acc:MGI:2148800] | Aipl1 | Enriched in KO | 4.966 | 0.03133 | 368.2 |
| ENSMUSG00000025900 | retinitis pigmentosa 1 (human) [Source:MGI Symbol;Acc:MGI:1341105] | Rp1 | Enriched in KO | 4.907 | 0.001204 | 3382 |
| ENSMUSG00000041044 | leucine-rich repeat, immunoglobulin-like and transmembrane domains 1 [Source:MGI Symbol;Acc:MGI:2385320] | Lrit1 | Enriched in KO | 4.897 | 0.03771 | 386.5 |
| ENSMUSG00000029663 | guanine nucleotide binding protein (G protein), gamma transducing activity polypeptide 1 [Source:MGI Symbol;Acc:MGI:10916[illegible] | Gngt1 | Enriched in KO | 4.889 | 0.04829 | 593.7 |
| ENSMUSG00000047298 | potassium channel, subfamily V, member 2 [Source:MGI Symbol;Acc:MGI:2670981] | Kcnv2 | Enriched in KO | 4.745 | 0.01834 | 537.7 |
| ENSMUSG00000029491 | phosphodiesterase 6B, cGMP, rod receptor, beta polypeptide [Source:MGI Symbol;Acc:MGI:97525] | Pde6b | Enriched in KO | 4.743 | 0.02745 | 1205 |
| ENSMUSG00000041534 | retinol binding protein 3, interstitial [Source:MGI Symbol;Acc:MGI:97878] | Rbp3 | Enriched in KO | 4.708 | 0.01556 | 3079 |
| ENSMUSG00000005649 | calcium binding protein 5 [Source:MGI Symbol;Acc:MGI:1352746] | Cabp5 | Enriched in KO | 4.488 | 0.03133 | 50.22 |
| ENSMUSG00000023979 | guanylate cyclase activator 1B [Source:MGI Symbol;Acc:MGI:1194489] | Guca1b | Enriched in KO | 4.433 | 0.02529 | 220.2 |
| ENSMUSG00000051860 | sterile alpha motif domain containing 7 [Source:MGI Symbol;Acc:MGI:1923203] | Samd7 | Enriched in KO | 4.407 | 0.04724 | 192 |
| ENSMUSG00000031789 | cyclic nucleotide gated channel beta 1 [Source:MGI Symbol;Acc:MGI:2664102] | Cngb1 | Enriched in KO | 4.388 | 0.002484 | 733.4 |
| ENSMUSG00000034829 | nucleoredoxin-like 1 [Source:MGI Symbol;Acc:MGI:1924446] | Nxnl1 | Enriched in KO | 4.353 | 0.03771 | 200.2 |
| ENSMUSG00000026609 | usherin [Source:MGI Symbol;Acc:MGI:1341292] | Ush2a | Enriched in KO | 4.184 | 0.0002212 | 1475 |
| ENSMUSG00000056947 | mab-21-like 1 [Source:MGI Symbol;Acc:MGI:1333773] | Mab21l1 | Enriched in KO | 4.154 | 0.03065 | 403.6 |
| ENSMUSG00000024227 | PDZ and pleckstrin homology domains 1 [Source:MGI Symbol;Acc:MGI:1916489] | Pdzph1 | Enriched in KO | 4.114 | 0.003514 | 255.2 |
| ENSMUSG00000020907 | recoverin [Source:MGI Symbol;Acc:MGI:97883] | Rcvrn | Enriched in KO | 4.107 | 0.03178 | 323.5 |
| ENSMUSG00000057132 | retinitis pigmentosa GTPase regulator interacting protein 1 [Source:MGI Symbol;Acc:MGI:1932134] | Rpgrip1 | Enriched in KO | 3.793 | 0.003151 | 1656 |
| ENSMUSG00000037446 | tubby like protein 1 [Source:MGI Symbol;Acc:MGI:109571] | Tulp1 | Enriched in KO | 3.782 | 0.0164 | 663.6 |
| ENSMUSG00000056043 | regulator of G-protein signalling 9 binding protein [Source:MGI Symbol;Acc:MGI:2384418] | Rgs9bp | Enriched in KO | 3.67 | 0.01749 | 666 |
| ENSMUSG00000035270 | interphotoreceptor matrix proteoglycan 2 [Source:MGI Symbol;Acc:MGI:3044955] | Impg2 | Enriched in KO | 3.556 | 0.006131 | 754.3 |
| ENSMUSG00000020890 | guanylate cyclase 2e [Source:MGI Symbol;Acc:MGI:105123] | Gucy2e | Enriched in KO | 3.493 | 0.0103 | 507.5 |
| ENSMUSG00000079550 | membrane protein, palmitoylated 4 (MAGUK p55 subfamily member 4) [Source:MGI Symbol;Acc:MGI:2386681] | Mpp4 | Enriched in KO | 3.483 | 0.03133 | 514.2 |
| ENSMUSG00000028188 | spermatogenesis associated 1 [Source:MGI Symbol;Acc:MGI:1918201] | Spata1 | Enriched in KO | 3.332 | 0.04222 | 64.99 |
| ENSMUSG00000021123 | retinol dehydrogenase 12 [Source:MGI Symbol;Acc:MGI:1925224] | Rdh12 | Enriched in KO | 3.274 | 0.04829 | 285 |
| ENSMUSG00000067220 | cyclic nucleotide gated channel alpha 1 [Source:MGI Symbol;Acc:MGI:88436] | Cnga1 | Enriched in KO | 2.982 | 0.02529 | 478.2 |
| ENSMUSG00000075410 | photoreceptor disc component [Source:MGI Symbol;Acc:MGI:3649529] | Prcd | Enriched in KO | 2.693 | 0.02529 | 168.2 |
| ENSMUSG00000021803 | cadherin-related family member 1 [Source:MGI Symbol;Acc:MGI:2157782] | Cdhr1 | Enriched in KO | 2.542 | 0.04216 | 1092 |
| ENSMUSG00000035504 | receptor accessory protein 6 [Source:MGI Symbol;Acc:MGI:1917585] | Reep6 | Enriched in KO | 2.508 | 0.0228 | 1054 |
| ENSMUSG00000038115 | anoctamin 2 [Source:MGI Symbol;Acc:MGI:2387214] | Ano2 | Enriched in KO | 2.452 | 0.03771 | 281.6 |
| ENSMUSG00000069170 | adhesion G protein-coupled receptor V1 [Source:MGI Symbol;Acc:MGI:1274784] | Adgrv1 | Enriched in KO | 2.317 | 0.005214 | 786.2 |

**Contrast numerator: P15_KO_retina. Contrast denominator: P15_Het_retina.**

| ensembl_gene_id | description | mgi_symbol | Significance | deseq_logfc | deseq_adjp | deseq_basemean |
|---|---|---|---|---|---|---|
| ENSMUSG00000037727 | arginine vasopressin [Source:MGI Symbol;Acc:MGI:88121] | Avp | Enriched in KO | 15.58 | 1.247E-08 | 32.86 |
| ENSMUSG00000063428 | D-aspartate oxidase [Source:MGI Symbol;Acc:MGI:1925528] | Ddo | Enriched in KO | 2.864 | 0.04435 | 90.03 |
| ENSMUSG00000029190 | DNA segment, Chr 5, ERATO Doi 579, expressed [Source:MGI Symbol;Acc:MGI:1261849] | D5Ertd579e | Enriched in KO | 2.457 | 0.00007177 | 1250 |
| ENSMUSG00000021619 | autophagy related 10 [Source:MGI Symbol;Acc:MGI:1914045] | Atg10 | Enriched in KO | 1.799 | 0.04435 | 39.83 |
| ENSMUSG00000060579 | fragile histidine triad gene [Source:MGI Symbol;Acc:MGI:1277947] | Fhit | Enriched in KO | 1.739 | 0.03319 | 135.3 |
| ENSMUSG00000029012 | origin recognition complex, subunit 5 [Source:MGI Symbol;Acc:MGI:1347044] | Orc5 | Enriched in KO | 1.698 | 0.02199 | 47.78 |
| ENSMUSG00000047344 | LanC lantibiotic synthetase component C-like 3 (bacterial) [Source:MGI Symbol;Acc:MGI:244333[illegible] | Lancl3 | Enriched in Het | -1.322 | 0.03319 | 26.64 |
| ENSMUSG00000056306 | serine rich and transmembrane domain containing 1 [Source:MGI Symbol;Acc:MGI:3607715] | Sertm1 | Enriched in Het | -1.456 | 0.02199 | 70.23 |
| ENSMUSG00000021799 | opsin 4 (melanopsin) [Source:MGI Symbol;Acc:MGI:1353425] | Opn4 | Enriched in Het | -9.012 | 4.424E-12 | 25.82 |
| ENSMUSG00000027301 | oxytocin [Source:MGI Symbol;Acc:MGI:97453] | Oxt | Enriched in Het | -17.41 | 2.076E-07 | 22.27 |

**Contrast numerator: P15_KO_SCN. Contrast denominator: P15_Het_SCN.**

| ensembl_gene_id | description | mgi_symbol | Significance | deseq_logfc | deseq_adjp | deseq_basemean |
|---|---|---|---|---|---|---|
| ENSMUSG00000024501 | dihydropyrimidinase-like 3 [Source:MGI Symbol;Acc:MGI:1349762] | Dpysl3 | Enriched in Het | -1.436 | 0.004222 | 3490 |
| ENSMUSG00000020431 | adenylate cyclase 1 [Source:MGI Symbol;Acc:MGI:99677] | Adcy1 | Enriched in Het | -1.523 | 0.0009236 | 5232 |
| ENSMUSG00000022865 | coxsackie virus and adenovirus receptor [Source:MGI Symbol;Acc:MGI:1201679] | Cxadr | Enriched in Het | -1.732 | 0.0009236 | 403.7 |
| ENSMUSG00000036913 | tripartite motif-containing 67 [Source:MGI Symbol;Acc:MGI:3045323] | Trim67 | Enriched in Het | -1.843 | 0.0009236 | 572.3 |
| ENSMUSG00000025314 | protein tyrosine phosphatase, receptor type, J [Source:MGI Symbol;Acc:MGI:10457 | Ptprj | Enriched in Het | -4.501 | 5.137E-09 | 998.9 |
