## Supplementary material for "Melanopsin regulates axonal translation underlying retinohypothalamic circuit assembly": Table_S4_TRAP_seq_age_contrasts_significant_genes

**Contrast numerator: P15_Het_dLGN. Contrast denominator: P8_Het_dLGN.**

| ensembl_gene_id | description | mgi_symbol | Significance | deseq_logfc | deseq_adjp | deseq_basemean |
|---|---|---|---|---|---|---|
| ENSMUSG00000021948 | protein kinase C, delta [Source:MGI Symbol;Acc:MGI:97598] | Prkcd | Enriched in P15 dLGN | 5.632 | 0.000 | 727.900 |
| ENSMUSG00000029054 | gamma-aminobutyric acid (GABA) A receptor, subunit delta [Source:MGI Symbol;Acc:MGI:95622] | Gabrd | Enriched in P15 dLGN | 4.640 | 0.000 | 52.020 |
| ENSMUSG00000019762 | iodotyrosine deiodinase [Source:MGI Symbol;Acc:MGI:1917587] | Iyd | Enriched in P15 dLGN | 4.505 | 0.000 | 6.399 |
| ENSMUSG00000027347 | RAS guanyl releasing protein 1 [Source:MGI Symbol;Acc:MGI:1314635] | Rasgrp1 | Enriched in P15 dLGN | 3.715 | 0.000 | 353.400 |
| ENSMUSG00000072476 | predicted pseudogene 9008 [Source:MGI Symbol;Acc:MGI:3644000] | Gm9008 | Enriched in P15 dLGN | 3.637 | 0.005 | 8.190 |
| ENSMUSG00000027489 | N-terminal EF-hand calcium binding protein 3 [Source:MGI Symbol;Acc:MGI:1861721] | Necab3 | Enriched in P15 dLGN | 3.407 | 0.000 | 36.720 |
| ENSMUSG00000039783 | kynurenine 3-monooxygenase (kynurenine 3-hydroxylase) [Source:MGI Symbol;Acc:MGI:2138151] | Kmo | Enriched in P15 dLGN | 3.165 | 0.001 | 4.130 |
| ENSMUSG00000000901 | matrix metallopeptidase 11 [Source:MGI Symbol;Acc:MGI:97008] | Mmp11 | Enriched in P15 dLGN | 3.116 | 0.000 | 8.781 |
| ENSMUSG00000023905 | tumor necrosis factor receptor superfamily, member 12a [Source:MGI Symbol;Acc:MGI:1351484] | Tnfrsf12a | Enriched in P15 dLGN | 3.081 | 0.000 | 7.873 |
| ENSMUSG00000041144 | dynein, axonemal, heavy chain 7B [Source:MGI Symbol;Acc:MGI:2684953] | Dnah7b | Enriched in P15 dLGN | 3.037 | 0.003 | 449.400 |
| ENSMUSG00000064179 | troponin T1, skeletal, slow [Source:MGI Symbol;Acc:MGI:1333868] | Tnnt1 | Enriched in P15 dLGN | 3.019 | 0.000 | 308.700 |
| ENSMUSG00000039115 | integrin alpha 9 [Source:MGI Symbol;Acc:MGI:104756] | Itga9 | Enriched in P15 dLGN | 2.989 | 0.001 | 1811.000 |
| ENSMUSG00000110711 | predicted gene 45760 [Source:MGI Symbol;Acc:MGI:5804875] | Gm45760 | Enriched in P15 dLGN | 2.914 | 0.008 | 2.941 |
| ENSMUSG00000043461 | serine palmitoyltransferase, small subunit B [Source:MGI Symbol;Acc:MGI:1913433] | Sptssb | Enriched in P15 dLGN | 2.852 | 0.000 | 9.484 |
| ENSMUSG00000029211 | gamma-aminobutyric acid (GABA) A receptor, subunit alpha 4 [Source:MGI Symbol;Acc:MGI:95616] | Gabra4 | Enriched in P15 dLGN | 2.851 | 0.000 | 154.600 |
| ENSMUSG00000045625 | phosphatidylinositol glycan anchor biosynthesis, class Z [Source:MGI Symbol;Acc:MGI:2443822] | Pigz | Enriched in P15 dLGN | 2.802 | 0.000 | 18.260 |
| ENSMUSG00000050541 | adrenergic receptor, alpha 1b [Source:MGI Symbol;Acc:MGI:104774] | Adra1b | Enriched in P15 dLGN | 2.776 | 0.000 | 73.770 |
| ENSMUSG00000026525 | opsin 3 [Source:MGI Symbol;Acc:MGI:1338022] | Opn3 | Enriched in P15 dLGN | 2.751 | 0.000 | 17.620 |
| ENSMUSG00000041046 | receptor (calcitonin) activity modifying protein 3 [Source:MGI Symbol;Acc:MGI:1860292] | Ramp3 | Enriched in P15 dLGN | 2.727 | 0.000 | 121.100 |
| ENSMUSG00000000884 | guanine nucleotide binding protein (G protein), beta polypeptide 1-like [Source:MGI Symbol;Acc:MGI:1338057] | Gnb1l | Enriched in P15 dLGN | 2.719 | 0.000 | 294.800 |
| ENSMUSG00000036330 | solute carrier family 18 (vesicular monoamine), member 1 [Source:MGI Symbol;Acc:MGI:106684] | Slc18a1 | Enriched in P15 dLGN | 2.706 | 0.006 | 5.605 |
| ENSMUSG00000028024 | glutamyl aminopeptidase [Source:MGI Symbol;Acc:MGI:106645] | Enpep | Enriched in P15 dLGN | 2.704 | 0.000 | 10.700 |
| ENSMUSG00000020733 | solute carrier family 9 (sodium/hydrogen exchanger), member 3 regulator 1 [Source:MGI Symbol;Acc:MGI:1349482] | Slc9a3r1 | Enriched in P15 dLGN | 2.675 | 0.000 | 214.900 |
| ENSMUSG00000060402 | carbohydrate sulfotransferase 8 [Source:MGI Symbol;Acc:MGI:1916197] | Chst8 | Enriched in P15 dLGN | 2.672 | 0.000 | 67.640 |
| ENSMUSG00000091722 | siah E3 ubiquitin protein ligase family member 3 [Source:MGI Symbol;Acc:MGI:2685758] | Siah3 | Enriched in P15 dLGN | 2.657 | 0.000 | 62.950 |
| ENSMUSG00000031778 | chemokine (C-X3-C motif) ligand 1 [Source:MGI Symbol;Acc:MGI:1097153] | Cx3cl1 | Enriched in P15 dLGN | 2.604 | 0.000 | 522.000 |
| ENSMUSG00000033615 | complexin 1 [Source:MGI Symbol;Acc:MGI:104727] | Cplx1 | Enriched in P15 dLGN | 2.595 | 0.000 | 2304.000 |
| ENSMUSG00000044017 | adhesion G protein-coupled receptor D1 [Source:MGI Symbol;Acc:MGI:3041203] | Adgrd1 | Enriched in P15 dLGN | 2.589 | 0.013 | 5.834 |
| ENSMUSG00000029641 | RAS-like, family 11, member A [Source:MGI Symbol;Acc:MGI:1916145] | Rasl11a | Enriched in P15 dLGN | 2.532 | 0.000 | 21.430 |
| ENSMUSG00000021250 | FBJ osteosarcoma oncogene [Source:MGI Symbol;Acc:MGI:95574] | Fos | Enriched in P15 dLGN | 2.519 | 0.000 | 57.190 |
| ENSMUSG00000040624 | pleckstrin homology domain containing, family G (with RhoGef domain) member 1 [Source:MGI Symbol;Acc:MGI:2676551] | Plekhg1 | Enriched in P15 dLGN | 2.509 | 0.000 | 465.000 |
| ENSMUSG00000001763 | tetraspanin 33 [Source:MGI Symbol;Acc:MGI:1919012] | Tspan33 | Enriched in P15 dLGN | 2.502 | 0.000 | 62.530 |
| ENSMUSG00000018821 | arginine vasopressin-induced 1 [Source:MGI Symbol;Acc:MGI:1916784] | Avpi1 | Enriched in P15 dLGN | 2.490 | 0.000 | 34.890 |
| ENSMUSG00000059991 | neuronal pentraxin 2 [Source:MGI Symbol;Acc:MGI:1858209] | Nptx2 | Enriched in P15 dLGN | 2.412 | 0.000 | 106.400 |
| ENSMUSG00000030785 | cytochrome c oxidase subunit 6A2 [Source:MGI Symbol;Acc:MGI:104649] | Cox6a2 | Enriched in P15 dLGN | 2.401 | 0.001 | 11.740 |
| ENSMUSG00000018470 | potassium voltage-gated channel, shaker-related subfamily, beta member 3 [Source:MGI Symbol;Acc:MGI:1336208] | Kcnab3 | Enriched in P15 dLGN | 2.394 | 0.000 | 53.860 |
| ENSMUSG00000064329 | sodium channel, voltage-gated, type I, alpha [Source:MGI Symbol;Acc:MGI:98246] | Scn1a | Enriched in P15 dLGN | 2.384 | 0.000 | 703.300 |
| ENSMUSG00000017167 | contactin associated protein-like 1 [Source:MGI Symbol;Acc:MGI:1858201] | Cntnap1 | Enriched in P15 dLGN | 2.372 | 0.000 | 622.800 |
| ENSMUSG00000022587 | lymphocyte antigen 6 complex, locus E [Source:MGI Symbol;Acc:MGI:106651] | Ly6e | Enriched in P15 dLGN | 2.365 | 0.000 | 95.380 |
| ENSMUSG00000027712 | annexin A5 [Source:MGI Symbol;Acc:MGI:106008] | Anxa5 | Enriched in P15 dLGN | 2.320 | 0.000 | 102.300 |
| ENSMUSG00000024190 | dual specificity phosphatase 1 [Source:MGI Symbol;Acc:MGI:105120] | Dusp1 | Enriched in P15 dLGN | 2.312 | 0.000 | 44.730 |
| ENSMUSG00000028710 | ATP synthase mitochondrial F1 complex assembly factor 1 [Source:MGI Symbol;Acc:MGI:2180560] | Atpaf1 | Enriched in P15 dLGN | 2.311 | 0.000 | 89.620 |
| ENSMUSG00000050315 | synaptopodin 2 [Source:MGI Symbol;Acc:MGI:2153070] | Synpo2 | Enriched in P15 dLGN | 2.289 | 0.000 | 581.400 |
| ENSMUSG00000049612 | oligodendrocyte myelin glycoprotein [Source:MGI Symbol;Acc:MGI:106586] | Omg | Enriched in P15 dLGN | 2.255 | 0.000 | 36.010 |
| ENSMUSG00000039629 | striatin interacting protein 2 [Source:MGI Symbol;Acc:MGI:2444363] | Strip2 | Enriched in P15 dLGN | 2.219 | 0.000 | 138.000 |
| ENSMUSG00000002032 | transmembrane protein 25 [Source:MGI Symbol;Acc:MGI:1918937] | Tmem25 | Enriched in P15 dLGN | 2.194 | 0.000 | 70.330 |
| ENSMUSG00000026824 | potassium inwardly-rectifying channel, subfamily J, member 3 [Source:MGI Symbol;Acc:MGI:104742] | Kcnj3 | Enriched in P15 dLGN | 2.157 | 0.000 | 206.800 |
| ENSMUSG00000022468 | endonuclease, polyU-specific [Source:MGI Symbol;Acc:MGI:97746] | Endou | Enriched in P15 dLGN | 2.146 | 0.000 | 34.930 |
| ENSMUSG00000003949 | hepatic leukemia factor [Source:MGI Symbol;Acc:MGI:96108] | Hlf | Enriched in P15 dLGN | 2.143 | 0.000 | 354.900 |
| ENSMUSG00000061859 | PATJ, crumbs cell polarity complex component [Source:MGI Symbol;Acc:MGI:1277960] | Patj | Enriched in P15 dLGN | 2.140 | 0.000 | 609.600 |
| ENSMUSG00000065968 | interferon induced transmembrane protein 7 [Source:MGI Symbol;Acc:MGI:1921732] | Ifitm7 | Enriched in P15 dLGN | 2.140 | 0.000 | 15.260 |
| ENSMUSG00000027556 | carbonic anhydrase 1 [Source:MGI Symbol;Acc:MGI:88268] | Car1 | Enriched in P15 dLGN | 2.123 | 0.000 | 12.270 |
| ENSMUSG00000010803 | gamma-aminobutyric acid (GABA) A receptor, subunit alpha 1 [Source:MGI Symbol;Acc:MGI:95613] | Gabra1 | Enriched in P15 dLGN | 2.113 | 0.000 | 231.000 |
| ENSMUSG00000031438 | ring finger protein 128 [Source:MGI Symbol;Acc:MGI:1914139] | Rnf128 | Enriched in P15 dLGN | 2.113 | 0.000 | 18.970 |
| ENSMUSG00000029659 | mesenteric estrogen dependent adipogenesis [Source:MGI Symbol;Acc:MGI:1917967] | Medag | Enriched in P15 dLGN | 2.070 | 0.000 | 25.400 |
| ENSMUSG00000025189 | cyclin M1 [Source:MGI Symbol;Acc:MGI:1891366] | Cnnm1 | Enriched in P15 dLGN | 2.069 | 0.000 | 258.600 |
| ENSMUSG00000025092 | heat shock protein 12A [Source:MGI Symbol;Acc:MGI:1920692] | Hspa12a | Enriched in P15 dLGN | 2.057 | 0.000 | 913.000 |
| ENSMUSG00000003541 | immediate early response 3 [Source:MGI Symbol;Acc:MGI:104814] | Ier3 | Enriched in P15 dLGN | 2.038 | 0.000 | 21.940 |
| ENSMUSG00000019828 | glutamate receptor, metabotropic 1 [Source:MGI Symbol;Acc:MGI:1351338] | Grm1 | Enriched in P15 dLGN | 2.029 | 0.000 | 572.700 |
| ENSMUSG00000045733 | shadow of prion protein [Source:MGI Symbol;Acc:MGI:3582583] | Sprn | Enriched in P15 dLGN | 2.016 | 0.000 | 239.000 |
| ENSMUSG00000028833 | neurochondrin [Source:MGI Symbol;Acc:MGI:1347351] | Ncdn | Enriched in P15 dLGN | 2.010 | 0.000 | 1887.000 |

| | | | | | | |
|---|---|---|---|---|---|---|
| ENSMUSG00000027894 | solute carrier family 6 (neurotransmitter transporter), member 17 [Source:MGI Symbol;Acc:MGI:2442535] | Slc6a17 | Enriched in P15 dLGN | 2.008 | 0.000 | 1006.000 |
| ENSMUSG00000032908 | sphingosine-1-phosphate phosphatase 2 [Source:MGI Symbol;Acc:MGI:3589109] | Sgpp2 | Enriched in P15 dLGN | 2.002 | 0.000 | 115.400 |
| ENSMUSG00000078963 | heat shock factor binding protein 1-like 1 [Source:MGI Symbol;Acc:MGI:1913505] | Hsbp1l1 | Enriched in P15 dLGN | 1.995 | 0.011 | 35.030 |
| ENSMUSG00000038967 | pyruvate dehydrogenase kinase, isoenzyme 2 [Source:MGI Symbol;Acc:MGI:1343087] | Pdk2 | Enriched in P15 dLGN | 1.966 | 0.000 | 136.900 |
| ENSMUSG00000078695 | CDGSH iron sulfur domain 3 [Source:MGI Symbol;Acc:MGI:101788] | Cisd3 | Enriched in P15 dLGN | 1.942 | 0.004 | 13.410 |
| ENSMUSG00000091387 | glucosaminyl (N-acetyl) transferase 4, core 2 (beta-1,6-N-acetylglucosaminyltransferase) [Source:MGI Symbol;Acc:MGI:2684919] | Gcnt4 | Enriched in P15 dLGN | 1.926 | 0.001 | 28.710 |
| ENSMUSG00000031007 | ATPase, H+ transporting, lysosomal accessory protein 2 [Source:MGI Symbol;Acc:MGI:1917745] | Atp6ap2 | Enriched in P15 dLGN | 1.877 | 0.000 | 242.600 |
| ENSMUSG00000027273 | synaptosomal-associated protein 25 [Source:MGI Symbol;Acc:MGI:98331] | Snap25 | Enriched in P15 dLGN | 1.872 | 0.000 | 1966.000 |
| ENSMUSG00000055782 | ATP-binding cassette, sub-family D (ALD), member 2 [Source:MGI Symbol;Acc:MGI:1349467] | Abcd2 | Enriched in P15 dLGN | 1.854 | 0.000 | 85.550 |
| ENSMUSG00000029769 | coiled-coil domain containing 136 [Source:MGI Symbol;Acc:MGI:1918128] | Ccdc136 | Enriched in P15 dLGN | 1.851 | 0.000 | 1034.000 |
| ENSMUSG00000039480 | 5'-nucleotidase domain containing 1 [Source:MGI Symbol;Acc:MGI:2442446] | Nt5dc1 | Enriched in P15 dLGN | 1.841 | 0.000 | 84.480 |
| ENSMUSG00000034187 | N-ethylmaleimide sensitive fusion protein [Source:MGI Symbol;Acc:MGI:104560] | Nsf | Enriched in P15 dLGN | 1.830 | 0.000 | 1114.000 |
| ENSMUSG00000031231 | cytochrome c oxidase subunit 7B [Source:MGI Symbol;Acc:MGI:1913392] | Cox7b | Enriched in P15 dLGN | 1.808 | 0.000 | 113.900 |
| ENSMUSG00000029632 | Ndufa4, mitochondrial complex associated [Source:MGI Symbol;Acc:MGI:107686] | Ndufa4 | Enriched in P15 dLGN | 1.806 | 0.000 | 183.200 |
| ENSMUSG00000030695 | aldolase A, fructose-bisphosphate [Source:MGI Symbol;Acc:MGI:87994] | Aldoa | Enriched in P15 dLGN | 1.805 | 0.000 | 1713.000 |
| ENSMUSG00000022564 | glutamate receptor, ionotropic, N-methyl D-aspartate-associated protein 1 (glutamate binding) [Source:MGI Symbol;Acc:MGI:1913418] | Grina | Enriched in P15 dLGN | 1.801 | 0.000 | 437.400 |
| ENSMUSG00000020321 | malate dehydrogenase 1, NAD (soluble) [Source:MGI Symbol;Acc:MGI:97051] | Mdh1 | Enriched in P15 dLGN | 1.798 | 0.000 | 617.500 |
| ENSMUSG00000032776 | multiple C2 domains, transmembrane 2 [Source:MGI Symbol;Acc:MGI:2685335] | Mctp2 | Enriched in P15 dLGN | 1.798 | 0.000 | 28.350 |
| ENSMUSG00000029153 | OCIA domain containing 2 [Source:MGI Symbol;Acc:MGI:1916377] | Ociad2 | Enriched in P15 dLGN | 1.790 | 0.000 | 70.580 |
| ENSMUSG00000040723 | RCSD domain containing 1 [Source:MGI Symbol;Acc:MGI:2676394] | Rcsd1 | Enriched in P15 dLGN | 1.786 | 0.000 | 27.500 |
| ENSMUSG00000025757 | heat shock protein 4 like [Source:MGI Symbol;Acc:MGI:107422] | Hspa4l | Enriched in P15 dLGN | 1.781 | 0.000 | 545.200 |
| ENSMUSG00000062070 | phosphoglycerate kinase 1 [Source:MGI Symbol;Acc:MGI:97555] | Pgk1 | Enriched in P15 dLGN | 1.781 | 0.000 | 300.000 |
| ENSMUSG00000021775 | nuclear receptor subfamily 1, group D, member 2 [Source:MGI Symbol;Acc:MGI:2449205] | Nr1d2 | Enriched in P15 dLGN | 1.757 | 0.000 | 245.400 |
| ENSMUSG00000045009 | proline-rich transmembrane protein 3 [Source:MGI Symbol;Acc:MGI:2444810] | Prrt3 | Enriched in P15 dLGN | 1.752 | 0.000 | 111.100 |
| ENSMUSG00000113149 | predicted gene, 49383 [Source:MGI Symbol;Acc:MGI:6121605] | Gm49383 | Enriched in P15 dLGN | 1.742 | 0.033 | 6.916 |
| ENSMUSG00000017417 | plexin domain containing 1 [Source:MGI Symbol;Acc:MGI:1919574] | Plxdc1 | Enriched in P15 dLGN | 1.728 | 0.000 | 77.270 |
| ENSMUSG00000078816 | protein kinase C, gamma [Source:MGI Symbol;Acc:MGI:97597] | Prkcg | Enriched in P15 dLGN | 1.725 | 0.000 | 571.800 |
| ENSMUSG00000021259 | cytochrome P450, family 46, subfamily a, polypeptide 1 [Source:MGI Symbol;Acc:MGI:1341877] | Cyp46a1 | Enriched in P15 dLGN | 1.724 | 0.000 | 248.200 |
| ENSMUSG00000062785 | potassium voltage gated channel, Shaw-related subfamily, member 3 [Source:MGI Symbol;Acc:MGI:96669] | Kcnc3 | Enriched in P15 dLGN | 1.723 | 0.000 | 437.600 |
| ENSMUSG00000032182 | Yip1 domain family, member 2 [Source:MGI Symbol;Acc:MGI:1922016] | Yipf2 | Enriched in P15 dLGN | 1.722 | 0.000 | 34.340 |
| ENSMUSG00000034825 | nuclear receptor interacting protein 3 [Source:MGI Symbol;Acc:MGI:1925843] | Nrip3 | Enriched in P15 dLGN | 1.721 | 0.000 | 285.500 |
| ENSMUSG00000004748 | mitochondrial fission process 1 [Source:MGI Symbol;Acc:MGI:1916686] | Mtfp1 | Enriched in P15 dLGN | 1.714 | 0.000 | 62.740 |
| ENSMUSG00000040612 | immunoglobulin-like domain containing receptor 2 [Source:MGI Symbol;Acc:MGI:1196370] | Ildr2 | Enriched in P15 dLGN | 1.699 | 0.000 | 1307.000 |
| ENSMUSG00000021591 | glutaredoxin [Source:MGI Symbol;Acc:MGI:2135625] | Glrx | Enriched in P15 dLGN | 1.685 | 0.000 | 37.820 |
| ENSMUSG00000035964 | transmembrane protein 59-like [Source:MGI Symbol;Acc:MGI:1915187] | Tmem59l | Enriched in P15 dLGN | 1.659 | 0.000 | 166.000 |
| ENSMUSG00000026825 | dynamin 1 [Source:MGI Symbol;Acc:MGI:107384] | Dnm1 | Enriched in P15 dLGN | 1.655 | 0.000 | 2096.000 |
| ENSMUSG00000007653 | gamma-aminobutyric acid (GABA) A receptor, subunit beta 2 [Source:MGI Symbol;Acc:MGI:95620] | Gabrb2 | Enriched in P15 dLGN | 1.649 | 0.000 | 322.600 |
| ENSMUSG00000091264 | small integral membrane protein 13 [Source:MGI Symbol;Acc:MGI:2652854] | Smim13 | Enriched in P15 dLGN | 1.635 | 0.000 | 346.000 |
| ENSMUSG00000036850 | mitochondrial ribosomal protein L41 [Source:MGI Symbol;Acc:MGI:1333816] | Mrpl41 | Enriched in P15 dLGN | 1.634 | 0.000 | 53.680 |
| ENSMUSG00000059361 | neurensin 2 [Source:MGI Symbol;Acc:MGI:2684969] | Nrsn2 | Enriched in P15 dLGN | 1.630 | 0.000 | 106.400 |
| ENSMUSG00000036186 | divergent protein kinase domain 1B [Source:MGI Symbol;Acc:MGI:1927576] | Dipk1b | Enriched in P15 dLGN | 1.619 | 0.000 | 28.180 |
| ENSMUSG00000028470 | histidine triad nucleotide binding protein 2 [Source:MGI Symbol;Acc:MGI:1916167] | Hint2 | Enriched in P15 dLGN | 1.606 | 0.000 | 22.680 |
| ENSMUSG00000040455 | ubiquitin specific petidase 45 [Source:MGI Symbol;Acc:MGI:101850] | Usp45 | Enriched in P15 dLGN | 1.606 | 0.000 | 101.700 |
| ENSMUSG00000039943 | phospholipase C, beta 4 [Source:MGI Symbol;Acc:MGI:107464] | Plcb4 | Enriched in P15 dLGN | 1.600 | 0.000 | 926.400 |
| ENSMUSG00000059343 | aldolase 1 A, retrogene 1 [Source:MGI Symbol;Acc:MGI:2447811] | Aldoart1 | Enriched in P15 dLGN | 1.597 | 0.001 | 19.450 |
| ENSMUSG00000023033 | sodium channel, voltage-gated, type VIII, alpha [Source:MGI Symbol;Acc:MGI:103169] | Scn8a | Enriched in P15 dLGN | 1.595 | 0.000 | 1192.000 |
| ENSMUSG00000068696 | G-protein coupled receptor 88 [Source:MGI Symbol;Acc:MGI:1927653] | Gpr88 | Enriched in P15 dLGN | 1.591 | 0.000 | 66.640 |
| ENSMUSG00000025981 | coenzyme Q10B [Source:MGI Symbol;Acc:MGI:1915126] | Coq10b | Enriched in P15 dLGN | 1.587 | 0.000 | 31.520 |
| ENSMUSG00000023456 | triosephosphate isomerase 1 [Source:MGI Symbol;Acc:MGI:98797] | Tpi1 | Enriched in P15 dLGN | 1.584 | 0.000 | 483.500 |
| ENSMUSG00000071369 | mitogen-activated protein kinase kinase kinase 5 [Source:MGI Symbol;Acc:MGI:1346876] | Map3k5 | Enriched in P15 dLGN | 1.583 | 0.000 | 236.000 |
| ENSMUSG00000027134 | lysophosphatidylcholine acyltransferase 4 [Source:MGI Symbol;Acc:MGI:2138993] | Lpcat4 | Enriched in P15 dLGN | 1.582 | 0.000 | 130.500 |
| ENSMUSG00000032011 | thymus cell antigen 1, theta [Source:MGI Symbol;Acc:MGI:98747] | Thy1 | Enriched in P15 dLGN | 1.580 | 0.000 | 952.800 |
| ENSMUSG00000060429 | syntrophin, basic 1 [Source:MGI Symbol;Acc:MGI:101781] | Sntb1 | Enriched in P15 dLGN | 1.575 | 0.000 | 115.500 |
| ENSMUSG00000071866 | peptidylprolyl isomerase A [Source:MGI Symbol;Acc:MGI:97749] | Ppia | Enriched in P15 dLGN | 1.574 | 0.004 | 211.200 |
| ENSMUSG00000002808 | ependymin related protein 1 (zebrafish) [Source:MGI Symbol;Acc:MGI:2145369] | Epdr1 | Enriched in P15 dLGN | 1.565 | 0.000 | 73.680 |
| ENSMUSG00000036158 | prickle planar cell polarity protein 1 [Source:MGI Symbol;Acc:MGI:1916034] | Prickle1 | Enriched in P15 dLGN | 1.561 | 0.000 | 385.100 |
| ENSMUSG00000050830 | von Willebrand factor C domain containing 2 [Source:MGI Symbol;Acc:MGI:2442987] | Vwc2 | Enriched in P15 dLGN | 1.552 | 0.000 | 127.000 |
| ENSMUSG00000046798 | claudin 12 [Source:MGI Symbol;Acc:MGI:1929288] | Cldn12 | Enriched in P15 dLGN | 1.551 | 0.000 | 124.600 |
| ENSMUSG00000016319 | solute carrier family 25 (mitochondrial carrier, adenine nucleotide translocator), member 5 [Source:MGI Symbol;Acc:MGI:1353496] | Slc25a5 | Enriched in P15 dLGN | 1.550 | 0.000 | 198.800 |
| ENSMUSG00000001986 | glutamate receptor, ionotropic, AMPA3 (alpha 3) [Source:MGI Symbol;Acc:MGI:95810] | Gria3 | Enriched in P15 dLGN | 1.548 | 0.000 | 283.800 |
| ENSMUSG00000014077 | calcineurin-like EF hand protein 1 [Source:MGI Symbol;Acc:MGI:1927185] | Chp1 | Enriched in P15 dLGN | 1.537 | 0.000 | 152.700 |
| ENSMUSG00000033917 | glycerophosphodiester phosphodiesterase 1 [Source:MGI Symbol;Acc:MGI:1891827] | Gde1 | Enriched in P15 dLGN | 1.535 | 0.000 | 234.500 |
| ENSMUSG00000034488 | EGF-like repeats and discoidin I-like domains 3 [Source:MGI Symbol;Acc:MGI:1329025] | Edil3 | Enriched in P15 dLGN | 1.520 | 0.000 | 564.800 |

| | | | | | | |
|---|---|---|---|---|---|---|
| ENSMUSG00000028271 | general transcription factor IIB [Source:MGI Symbol;Acc:MGI:2385191] | Gtf2b | Enriched in P15 dLGN | 1.515 | 0.000 | 36.100 |
| ENSMUSG00000059734 | NADH:ubiquinone oxidoreductase core subunit S8 [Source:MGI Symbol;Acc:MGI:2385079] | Ndufs8 | Enriched in P15 dLGN | 1.513 | 0.000 | 73.630 |
| ENSMUSG00000024158 | hydroxyacyl glutathione hydrolase [Source:MGI Symbol;Acc:MGI:95745] | Hagh | Enriched in P15 dLGN | 1.511 | 0.000 | 60.650 |
| ENSMUSG00000038550 | circadian associated repressor of transcription [Source:MGI Symbol;Acc:MGI:2684975] | Ciart | Enriched in P15 dLGN | 1.511 | 0.011 | 14.250 |
| ENSMUSG00000011752 | phosphoglycerate mutase 1 [Source:MGI Symbol;Acc:MGI:97552] | Pgam1 | Enriched in P15 dLGN | 1.508 | 0.000 | 217.300 |
| ENSMUSG00000026260 | NADH:ubiquinone oxidoreductase subunit A10 [Source:MGI Symbol;Acc:MGI:1914523] | Ndufa10 | Enriched in P15 dLGN | 1.508 | 0.000 | 263.700 |
| ENSMUSG00000035329 | F-box protein 33 [Source:MGI Symbol;Acc:MGI:1917861] | Fbxo33 | Enriched in P15 dLGN | 1.505 | 0.000 | 73.810 |
| ENSMUSG00000042298 | tetratricopeptide repeat domain 19 [Source:MGI Symbol;Acc:MGI:1920045] | Ttc19 | Enriched in P15 dLGN | 1.503 | 0.000 | 462.600 |
| ENSMUSG00000033998 | potassium channel, subfamily K, member 1 [Source:MGI Symbol;Acc:MGI:109322] | Kcnk1 | Enriched in P15 dLGN | 1.498 | 0.000 | 107.500 |
| ENSMUSG00000040907 | ATPase, Na+/K+ transporting, alpha 3 polypeptide [Source:MGI Symbol;Acc:MGI:88107] | Atp1a3 | Enriched in P15 dLGN | 1.497 | 0.000 | 7574.000 |
| ENSMUSG00000026688 | microsomal glutathione S-transferase 3 [Source:MGI Symbol;Acc:MGI:1913697] | Mgst3 | Enriched in P15 dLGN | 1.495 | 0.003 | 88.460 |
| ENSMUSG00000042032 | methionine adenosyltransferase II, beta [Source:MGI Symbol;Acc:MGI:1913667] | Mat2b | Enriched in P15 dLGN | 1.489 | 0.000 | 58.430 |
| ENSMUSG00000056486 | chimerin 1 [Source:MGI Symbol;Acc:MGI:1915674] | Chn1 | Enriched in P15 dLGN | 1.482 | 0.000 | 504.400 |
| ENSMUSG00000002068 | cyclin E1 [Source:MGI Symbol;Acc:MGI:88316] | Ccne1 | Enriched in P15 dLGN | 1.470 | 0.000 | 29.030 |
| ENSMUSG00000030718 | protein phosphatase methylesterase 1 [Source:MGI Symbol;Acc:MGI:1919840] | Ppme1 | Enriched in P15 dLGN | 1.468 | 0.000 | 576.900 |
| ENSMUSG00000049422 | coiled-coil-helix-coiled-coil-helix domain containing 10 [Source:MGI Symbol;Acc:MGI:2143558] | Chchd10 | Enriched in P15 dLGN | 1.468 | 0.000 | 224.500 |
| ENSMUSG00000027603 | gamma-glutamyltransferase 7 [Source:MGI Symbol;Acc:MGI:1913385] | Ggt7 | Enriched in P15 dLGN | 1.467 | 0.000 | 194.700 |
| ENSMUSG00000027274 | McKusick-Kaufman syndrome [Source:MGI Symbol;Acc:MGI:1891836] | Mkks | Enriched in P15 dLGN | 1.464 | 0.008 | 21.920 |
| ENSMUSG00000042682 | selenoprotein K [Source:MGI Symbol;Acc:MGI:1931466] | Selenok | Enriched in P15 dLGN | 1.459 | 0.000 | 254.300 |
| ENSMUSG00000014313 | cytochrome c oxidase subunit 6C [Source:MGI Symbol;Acc:MGI:104614] | Cox6c | Enriched in P15 dLGN | 1.456 | 0.000 | 345.400 |
| ENSMUSG00000005823 | G protein-coupled receptor 108 [Source:MGI Symbol;Acc:MGI:1925558] | Gpr108 | Enriched in P15 dLGN | 1.449 | 0.000 | 51.170 |
| ENSMUSG00000031176 | dynein light chain Tctex-type 3 [Source:MGI Symbol;Acc:MGI:1914367] | Dynlt3 | Enriched in P15 dLGN | 1.447 | 0.000 | 112.800 |
| ENSMUSG00000039983 | coiled-coil domain containing 32 [Source:MGI Symbol;Acc:MGI:2685477] | Ccdc32 | Enriched in P15 dLGN | 1.446 | 0.000 | 49.020 |
| ENSMUSG00000019877 | serine incorporator 1 [Source:MGI Symbol;Acc:MGI:1926228] | Serinc1 | Enriched in P15 dLGN | 1.445 | 0.000 | 662.300 |
| ENSMUSG00000022295 | ATPase, H+ transporting, lysosomal V1 subunit C1 [Source:MGI Symbol;Acc:MGI:1913585] | Atp6v1c1 | Enriched in P15 dLGN | 1.433 | 0.000 | 229.000 |
| ENSMUSG00000006057 | ATP synthase, H+ transporting, mitochondrial F0 complex, subunit C1 (subunit 9) [Source:MGI Symbol;Acc:MGI:107653] | Atp5g1 | Enriched in P15 dLGN | 1.431 | 0.000 | 76.530 |
| ENSMUSG00000037606 | oxysterol binding protein-like 5 [Source:MGI Symbol;Acc:MGI:1930265] | Osbpl5 | Enriched in P15 dLGN | 1.423 | 0.000 | 213.900 |
| ENSMUSG00000038555 | receptor accessory protein 2 [Source:MGI Symbol;Acc:MGI:2385070] | Reep2 | Enriched in P15 dLGN | 1.422 | 0.000 | 302.700 |
| ENSMUSG00000027963 | exostosin-like glycosyltransferase 2 [Source:MGI Symbol;Acc:MGI:1889574] | Extl2 | Enriched in P15 dLGN | 1.421 | 0.000 | 151.600 |
| ENSMUSG00000062542 | synaptotagmin IX [Source:MGI Symbol;Acc:MGI:1926373] | Syt9 | Enriched in P15 dLGN | 1.421 | 0.000 | 283.300 |
| ENSMUSG00000030327 | NECAP endocytosis associated 1 [Source:MGI Symbol;Acc:MGI:1914852] | Necap1 | Enriched in P15 dLGN | 1.419 | 0.000 | 167.400 |
| ENSMUSG00000018965 | tyrosine 3-monooxygenase/tryptophan 5-monooxygenase activation protein, eta polypeptide [Source:MGI Symbol;Acc:MGI:109194] | Ywhah | Enriched in P15 dLGN | 1.418 | 0.000 | 3239.000 |
| ENSMUSG00000027131 | ER membrane protein complex subunit 4 [Source:MGI Symbol;Acc:MGI:1915282] | Emc4 | Enriched in P15 dLGN | 1.418 | 0.000 | 81.850 |
| ENSMUSG00000007891 | cathepsin D [Source:MGI Symbol;Acc:MGI:88562] | Ctsd | Enriched in P15 dLGN | 1.416 | 0.006 | 65.280 |
| ENSMUSG00000020153 | NADH:ubiquinone oxidoreductase core subunit S7 [Source:MGI Symbol;Acc:MGI:1922656] | Ndufs7 | Enriched in P15 dLGN | 1.416 | 0.000 | 124.000 |
| ENSMUSG00000024194 | cutA divalent cation tolerance homolog [Source:MGI Symbol;Acc:MGI:1914925] | Cuta | Enriched in P15 dLGN | 1.416 | 0.000 | 66.240 |
| ENSMUSG00000001366 | f-box protein 9 [Source:MGI Symbol;Acc:MGI:1918788] | Fbxo9 | Enriched in P15 dLGN | 1.414 | 0.000 | 320.200 |
| ENSMUSG00000028149 | RAP1, GTP-GDP dissociation stimulator 1 [Source:MGI Symbol;Acc:MGI:2385189] | Rap1gds1 | Enriched in P15 dLGN | 1.411 | 0.000 | 847.300 |
| ENSMUSG00000027698 | neutral cholesterol ester hydrolase 1 [Source:MGI Symbol;Acc:MGI:2443191] | Nceh1 | Enriched in P15 dLGN | 1.408 | 0.000 | 190.100 |
| ENSMUSG00000028648 | NADH:ubiquinone oxidoreductase core subunit S5 [Source:MGI Symbol;Acc:MGI:1890889] | Ndufs5 | Enriched in P15 dLGN | 1.406 | 0.000 | 101.900 |
| ENSMUSG00000054162 | sparc/osteonectin, cwcv and kazal-like domains proteoglycan 3 [Source:MGI Symbol;Acc:MGI:1920152] | Spock3 | Enriched in P15 dLGN | 1.403 | 0.000 | 404.400 |
| ENSMUSG00000022890 | ATP synthase, H+ transporting, mitochondrial F0 complex, subunit F [Source:MGI Symbol;Acc:MGI:107777] | Atp5j | Enriched in P15 dLGN | 1.396 | 0.000 | 249.200 |
| ENSMUSG00000056185 | sorting nexin 32 [Source:MGI Symbol;Acc:MGI:2444704] | Snx32 | Enriched in P15 dLGN | 1.395 | 0.000 | 101.200 |
| ENSMUSG00000033389 | Rho GTPase activating protein 44 [Source:MGI Symbol;Acc:MGI:2144423] | Arhgap44 | Enriched in P15 dLGN | 1.393 | 0.000 | 522.200 |
| ENSMUSG00000026568 | mitochondrial pyruvate carrier 2 [Source:MGI Symbol;Acc:MGI:1917706] | Mpc2 | Enriched in P15 dLGN | 1.391 | 0.000 | 84.480 |
| ENSMUSG00000021728 | embigin [Source:MGI Symbol;Acc:MGI:95321] | Emb | Enriched in P15 dLGN | 1.390 | 0.000 | 88.850 |
| ENSMUSG00000049097 | ankyrin repeat domain 34A [Source:MGI Symbol;Acc:MGI:3617846] | Ankrd34a | Enriched in P15 dLGN | 1.388 | 0.000 | 84.030 |
| ENSMUSG00000006717 | acyl-CoA thioesterase 13 [Source:MGI Symbol;Acc:MGI:1914084] | Acot13 | Enriched in P15 dLGN | 1.386 | 0.001 | 30.710 |
| ENSMUSG00000029467 | ATPase, Ca++ transporting, cardiac muscle, slow twitch 2 [Source:MGI Symbol;Acc:MGI:88110] | Atp2a2 | Enriched in P15 dLGN | 1.385 | 0.000 | 2995.000 |
| ENSMUSG00000082229 | nucleosome assembly protein 1-like 2 [Source:MGI Symbol;Acc:MGI:106654] | Nap1l2 | Enriched in P15 dLGN | 1.385 | 0.000 | 277.100 |
| ENSMUSG00000028249 | syndecan binding protein [Source:MGI Symbol;Acc:MGI:1337026] | Sdcbp | Enriched in P15 dLGN | 1.383 | 0.000 | 76.000 |
| ENSMUSG00000032279 | isocitrate dehydrogenase 3 (NAD+) alpha [Source:MGI Symbol;Acc:MGI:1915084] | Idh3a | Enriched in P15 dLGN | 1.383 | 0.002 | 302.000 |
| ENSMUSG00000015668 | PDZ domain containing 11 [Source:MGI Symbol;Acc:MGI:1919871] | Pdzd11 | Enriched in P15 dLGN | 1.382 | 0.000 | 27.950 |
| ENSMUSG00000024074 | cysteine rich transmembrane BMP regulator 1 (chordin like) [Source:MGI Symbol;Acc:MGI:1354756] | Crim1 | Enriched in P15 dLGN | 1.379 | 0.000 | 661.000 |
| ENSMUSG00000031996 | amyloid beta (A4) precursor-like protein 2 [Source:MGI Symbol;Acc:MGI:88047] | Aplp2 | Enriched in P15 dLGN | 1.371 | 0.000 | 2704.000 |
| ENSMUSG00000025393 | ATP synthase, H+ transporting mitochondrial F1 complex, beta subunit [Source:MGI Symbol;Acc:MGI:107801] | Atp5b | Enriched in P15 dLGN | 1.369 | 0.000 | 1784.000 |
| ENSMUSG00000059409 | protein phosphatase 2, regulatory subunit B', delta [Source:MGI Symbol;Acc:MGI:2388481] | Ppp2r5d | Enriched in P15 dLGN | 1.366 | 0.000 | 362.300 |
| ENSMUSG00000067925 | retrotransposon Gag like 8A [Source:MGI Symbol;Acc:MGI:1913408] | Rtl8a | Enriched in P15 dLGN | 1.364 | 0.000 | 28.450 |
| ENSMUSG00000030879 | mitochondrial ribosomal protein L17 [Source:MGI Symbol;Acc:MGI:1351608] | Mrpl17 | Enriched in P15 dLGN | 1.363 | 0.000 | 76.990 |
| ENSMUSG00000035198 | tubulin, gamma 1 [Source:MGI Symbol;Acc:MGI:101834] | Tubg1 | Enriched in P15 dLGN | 1.361 | 0.001 | 105.400 |
| ENSMUSG00000053329 | glutamine amidotransferase like class 1 domain containing 3A [Source:MGI Symbol;Acc:MGI:1351861] | Gatd3a | Enriched in P15 dLGN | 1.360 | 0.000 | 113.500 |
| ENSMUSG00000027674 | peroxisomal biogenesis factor 5-like [Source:MGI Symbol;Acc:MGI:1916672] | Pex5l | Enriched in P15 dLGN | 1.359 | 0.001 | 789.100 |
| ENSMUSG00000035849 | keratin 222 [Source:MGI Symbol;Acc:MGI:2442728] | Krt222 | Enriched in P15 dLGN | 1.355 | 0.000 | 127.000 |

| | | | | | | |
|---|---|---|---|---|---|---|
| ENSMUSG00000024953 | peroxiredoxin 5 [Source:MGI Symbol;Acc:MGI:1859821] | Prdx5 | Enriched in P15 dLGN | 1.343 | 0.000 | 344.200 |
| ENSMUSG00000021576 | programmed cell death 6 [Source:MGI Symbol;Acc:MGI:109283] | Pdcd6 | Enriched in P15 dLGN | 1.341 | 0.001 | 37.470 |
| ENSMUSG00000083282 | cathepsin F [Source:MGI Symbol;Acc:MGI:1861434] | Ctsf | Enriched in P15 dLGN | 1.337 | 0.000 | 193.600 |
| ENSMUSG00000027495 | family with sequence similarity 210, member B [Source:MGI Symbol;Acc:MGI:1914267] | Fam210b | Enriched in P15 dLGN | 1.335 | 0.000 | 55.780 |
| ENSMUSG00000057766 | ankyrin repeat domain 29 [Source:MGI Symbol;Acc:MGI:2687055] | Ankrd29 | Enriched in P15 dLGN | 1.331 | 0.000 | 58.300 |
| ENSMUSG00000024414 | mitochondrial ribosomal protein L27 [Source:MGI Symbol;Acc:MGI:2137224] | Mrpl27 | Enriched in P15 dLGN | 1.320 | 0.002 | 26.690 |
| ENSMUSG00000001089 | leucine zipper protein 1 [Source:MGI Symbol;Acc:MGI:107629] | Luzp1 | Enriched in P15 dLGN | 1.316 | 0.000 | 485.100 |
| ENSMUSG00000024887 | N-acylsphingosine amidohydrolase 2 [Source:MGI Symbol;Acc:MGI:1859310] | Asah2 | Enriched in P15 dLGN | 1.312 | 0.005 | 39.790 |
| ENSMUSG00000025651 | ubiquinol-cytochrome c reductase core protein 1 [Source:MGI Symbol;Acc:MGI:107876] | Uqcrc1 | Enriched in P15 dLGN | 1.312 | 0.000 | 409.800 |
| ENSMUSG00000037152 | NADH:ubiquinone oxidoreductase subunit C1 [Source:MGI Symbol;Acc:MGI:1913627] | Ndufc1 | Enriched in P15 dLGN | 1.310 | 0.001 | 49.470 |
| ENSMUSG00000023020 | cytochrome c oxidase assembly protein 14 [Source:MGI Symbol;Acc:MGI:1913629] | Cox14 | Enriched in P15 dLGN | 1.309 | 0.000 | 73.580 |
| ENSMUSG00000063524 | enolase 1, alpha non-neuron [Source:MGI Symbol;Acc:MGI:95393] | Eno1 | Enriched in P15 dLGN | 1.308 | 0.000 | 583.300 |
| ENSMUSG00000020340 | cytoplasmic FMR1 interacting protein 2 [Source:MGI Symbol;Acc:MGI:1924134] | Cyfip2 | Enriched in P15 dLGN | 1.307 | 0.000 | 1444.000 |
| ENSMUSG00000034341 | WW domain binding protein 2 [Source:MGI Symbol;Acc:MGI:104709] | Wbp2 | Enriched in P15 dLGN | 1.297 | 0.000 | 271.900 |
| ENSMUSG00000044221 | G-rich RNA sequence binding factor 1 [Source:MGI Symbol;Acc:MGI:106479] | Grsf1 | Enriched in P15 dLGN | 1.296 | 0.000 | 242.300 |
| ENSMUSG00000031647 | microfibrillar-associated protein 3-like [Source:MGI Symbol;Acc:MGI:1918556] | Mfap3l | Enriched in P15 dLGN | 1.295 | 0.000 | 250.900 |
| ENSMUSG00000031617 | transmembrane protein 184C [Source:MGI Symbol;Acc:MGI:2384562] | Tmem184c | Enriched in P15 dLGN | 1.292 | 0.000 | 115.900 |
| ENSMUSG00000021748 | pyruvate dehydrogenase (lipoamide) beta [Source:MGI Symbol;Acc:MGI:1915513] | Pdhb | Enriched in P15 dLGN | 1.289 | 0.000 | 160.800 |
| ENSMUSG00000025428 | ATP synthase, H+ transporting, mitochondrial F1 complex, alpha subunit 1 [Source:MGI Symbol;Acc:MGI:88115] | Atp5a1 | Enriched in P15 dLGN | 1.287 | 0.000 | 1689.000 |
| ENSMUSG00000027104 | activating transcription factor 2 [Source:MGI Symbol;Acc:MGI:109349] | Atf2 | Enriched in P15 dLGN | 1.287 | 0.004 | 703.500 |
| ENSMUSG00000032324 | tetraspanin 3 [Source:MGI Symbol;Acc:MGI:1928098] | Tspan3 | Enriched in P15 dLGN | 1.287 | 0.000 | 417.400 |
| ENSMUSG00000032336 | neuroplastin [Source:MGI Symbol;Acc:MGI:108077] | Nptn | Enriched in P15 dLGN | 1.287 | 0.000 | 662.100 |
| ENSMUSG00000041697 | cytochrome c oxidase subunit 6A1 [Source:MGI Symbol;Acc:MGI:103099] | Cox6a1 | Enriched in P15 dLGN | 1.284 | 0.000 | 168.000 |
| ENSMUSG00000016349 | eukaryotic translation elongation factor 1 alpha 2 [Source:MGI Symbol;Acc:MGI:1096317] | Eef1a2 | Enriched in P15 dLGN | 1.282 | 0.000 | 1496.000 |
| ENSMUSG00000025240 | SAC1 suppressor of actin mutations 1-like (yeast) [Source:MGI Symbol;Acc:MGI:1933169] | Sacm1l | Enriched in P15 dLGN | 1.281 | 0.000 | 163.500 |
| ENSMUSG00000029634 | ring finger protein (C3H2C3 type) 6 [Source:MGI Symbol;Acc:MGI:1921382] | Rnf6 | Enriched in P15 dLGN | 1.279 | 0.000 | 280.900 |
| ENSMUSG00000020115 | TANK-binding kinase 1 [Source:MGI Symbol;Acc:MGI:1929658] | Tbk1 | Enriched in P15 dLGN | 1.275 | 0.000 | 121.800 |
| ENSMUSG00000031198 | FUN14 domain containing 2 [Source:MGI Symbol;Acc:MGI:1914641] | Fundc2 | Enriched in P15 dLGN | 1.272 | 0.000 | 67.820 |
| ENSMUSG00000039809 | gamma-aminobutyric acid (GABA) B receptor, 2 [Source:MGI Symbol;Acc:MGI:2386030] | Gabbr2 | Enriched in P15 dLGN | 1.269 | 0.000 | 1664.000 |
| ENSMUSG00000079317 | trafficking protein particle complex 2 [Source:MGI Symbol;Acc:MGI:1913476] | Trappc2 | Enriched in P15 dLGN | 1.268 | 0.009 | 15.830 |
| ENSMUSG00000027406 | isocitrate dehydrogenase 3 (NAD+) beta [Source:MGI Symbol;Acc:MGI:2158650] | Idh3b | Enriched in P15 dLGN | 1.266 | 0.000 | 236.200 |
| ENSMUSG00000048495 | tRNA-yW synthesizing protein 5 [Source:MGI Symbol;Acc:MGI:1915986] | Tyw5 | Enriched in P15 dLGN | 1.264 | 0.011 | 22.160 |
| ENSMUSG00000021373 | CAP, adenylate cyclase-associated protein, 2 (yeast) [Source:MGI Symbol;Acc:MGI:1914502] | Cap2 | Enriched in P15 dLGN | 1.258 | 0.000 | 147.000 |
| ENSMUSG00000044676 | zinc finger protein 612 [Source:MGI Symbol;Acc:MGI:2443465] | Zfp612 | Enriched in P15 dLGN | 1.258 | 0.000 | 153.100 |
| ENSMUSG00000021253 | transforming growth factor, beta 3 [Source:MGI Symbol;Acc:MGI:98727] | Tgfb3 | Enriched in P15 dLGN | 1.254 | 0.000 | 74.630 |
| ENSMUSG00000027893 | S-adenosylhomocysteine hydrolase-like 1 [Source:MGI Symbol;Acc:MGI:2385184] | Ahcyl1 | Enriched in P15 dLGN | 1.252 | 0.000 | 829.100 |
| ENSMUSG00000004207 | prosaposin [Source:MGI Symbol;Acc:MGI:97783] | Psap | Enriched in P15 dLGN | 1.251 | 0.000 | 1320.000 |
| ENSMUSG00000033316 | polypeptide N-acetylgalactosaminyltransferase 9 [Source:MGI Symbol;Acc:MGI:2677965] | Galnt9 | Enriched in P15 dLGN | 1.250 | 0.000 | 204.600 |
| ENSMUSG00000002475 | abhydrolase domain containing 3 [Source:MGI Symbol;Acc:MGI:2147183] | Abhd3 | Enriched in P15 dLGN | 1.249 | 0.005 | 45.040 |
| ENSMUSG00000031672 | glutamatic-oxaloacetic transaminase 2, mitochondrial [Source:MGI Symbol;Acc:MGI:95792] | Got2 | Enriched in P15 dLGN | 1.248 | 0.000 | 253.700 |
| ENSMUSG00000019087 | ATPase, H+ transporting, lysosomal accessory protein 1 [Source:MGI Symbol;Acc:MGI:109629] | Atp6ap1 | Enriched in P15 dLGN | 1.247 | 0.000 | 276.800 |
| ENSMUSG00000027206 | COP9 signalosome subunit 2 [Source:MGI Symbol;Acc:MGI:1330276] | Cops2 | Enriched in P15 dLGN | 1.244 | 0.000 | 139.200 |
| ENSMUSG00000045427 | heterogeneous nuclear ribonucleoprotein H2 [Source:MGI Symbol;Acc:MGI:1201779] | Hnrnph2 | Enriched in P15 dLGN | 1.242 | 0.000 | 111.000 |
| ENSMUSG00000051851 | retrotransposon Gag like 8C [Source:MGI Symbol;Acc:MGI:1920115] | Rtl8c | Enriched in P15 dLGN | 1.239 | 0.002 | 53.920 |
| ENSMUSG00000003072 | ATP synthase, H+ transporting, mitochondrial F1 complex, delta subunit [Source:MGI Symbol;Acc:MGI:1913293] | Atp5d | Enriched in P15 dLGN | 1.238 | 0.000 | 259.800 |
| ENSMUSG00000030647 | NADH:ubiquinone oxidoreductase subunit C2 [Source:MGI Symbol;Acc:MGI:1344370] | Ndufc2 | Enriched in P15 dLGN | 1.238 | 0.002 | 77.570 |
| ENSMUSG00000026032 | NADH:ubiquinone oxidoreductase subunit B3 [Source:MGI Symbol;Acc:MGI:1913745] | Ndufb3 | Enriched in P15 dLGN | 1.236 | 0.000 | 49.010 |
| ENSMUSG00000030127 | COP9 signalosome subunit 7A [Source:MGI Symbol;Acc:MGI:1349400] | Cops7a | Enriched in P15 dLGN | 1.235 | 0.000 | 167.000 |
| ENSMUSG00000033096 | adipocyte plasma membrane associated protein [Source:MGI Symbol;Acc:MGI:1919131] | Apmap | Enriched in P15 dLGN | 1.235 | 0.000 | 102.300 |
| ENSMUSG00000078566 | BCL2/adenovirus E1B interacting protein 3 [Source:MGI Symbol;Acc:MGI:109326] | Bnip3 | Enriched in P15 dLGN | 1.232 | 0.003 | 111.700 |
| ENSMUSG00000059040 | enolase 1B, retrotransposed [Source:MGI Symbol;Acc:MGI:3648653] | Eno1b | Enriched in P15 dLGN | 1.227 | 0.017 | 124.100 |
| ENSMUSG00000025868 | HIG1 domain family, member 2A [Source:MGI Symbol;Acc:MGI:1914294] | Higd2a | Enriched in P15 dLGN | 1.224 | 0.002 | 47.280 |
| ENSMUSG00000026959 | glutamate receptor, ionotropic, NMDA1 (zeta 1) [Source:MGI Symbol;Acc:MGI:95819] | Grin1 | Enriched in P15 dLGN | 1.224 | 0.000 | 942.600 |
| ENSMUSG00000032046 | abhydrolase domain containing 12 [Source:MGI Symbol;Acc:MGI:1923442] | Abhd12 | Enriched in P15 dLGN | 1.224 | 0.000 | 313.900 |
| ENSMUSG00000033124 | autophagy related 9A [Source:MGI Symbol;Acc:MGI:2138446] | Atg9a | Enriched in P15 dLGN | 1.221 | 0.000 | 274.400 |
| ENSMUSG00000052738 | succinate-CoA ligase, GDP-forming, alpha subunit [Source:MGI Symbol;Acc:MGI:1927234] | Suclg1 | Enriched in P15 dLGN | 1.219 | 0.000 | 138.600 |
| ENSMUSG00000020284 | cilia and flagella associated protein 410 [Source:MGI Symbol;Acc:MGI:1915134] | Cfap410 | Enriched in P15 dLGN | 1.218 | 0.000 | 83.690 |
| ENSMUSG00000021520 | ubiquinol-cytochrome c reductase binding protein [Source:MGI Symbol;Acc:MGI:1914780] | Uqcrb | Enriched in P15 dLGN | 1.214 | 0.006 | 49.090 |
| ENSMUSG00000032959 | phosphatidylethanolamine binding protein 1 [Source:MGI Symbol;Acc:MGI:1344408] | Pebp1 | Enriched in P15 dLGN | 1.214 | 0.000 | 164.100 |
| ENSMUSG00000031820 | BRISC and BRCA1 A complex member 1 [Source:MGI Symbol;Acc:MGI:1915501] | Babam1 | Enriched in P15 dLGN | 1.212 | 0.000 | 108.400 |
| ENSMUSG00000038486 | synaptic vesicle glycoprotein 2 a [Source:MGI Symbol;Acc:MGI:1927139] | Sv2a | Enriched in P15 dLGN | 1.212 | 0.000 | 1691.000 |
| ENSMUSG00000044252 | oxysterol binding protein-like 1A [Source:MGI Symbol;Acc:MGI:1927551] | Osbpl1a | Enriched in P15 dLGN | 1.211 | 0.000 | 434.200 |
| ENSMUSG00000000168 | dihydrolipoamide S-acetyltransferase (E2 component of pyruvate dehydrogenase complex) [Source:MGI Symbol;Acc:MGI:2385311] | Dlat | Enriched in P15 dLGN | 1.209 | 0.000 | 196.100 |

| | | | | | | |
|---|---|---|---|---|---|---|
| ENSMUSG00000029657 | heat shock 105kDa/110kDa protein 1 [Source:MGI Symbol;Acc:MGI:105053] | Hsph1 | Enriched in P15 dLGN | 1.209 | 0.000 | 555.200 |
| ENSMUSG00000034402 | potassium voltage-gated channel, subfamily H (eag-related), member 5 [Source:MGI Symbol;Acc:MGI:3584508] | Kcnh5 | Enriched in P15 dLGN | 1.208 | 0.001 | 240.300 |
| ENSMUSG00000033319 | fem 1 homolog c [Source:MGI Symbol;Acc:MGI:2444737] | Fem1c | Enriched in P15 dLGN | 1.207 | 0.000 | 163.300 |
| ENSMUSG00000028488 | SH3-domain GRB2-like 2 [Source:MGI Symbol;Acc:MGI:700009] | Sh3gl2 | Enriched in P15 dLGN | 1.204 | 0.000 | 586.900 |
| ENSMUSG00000078941 | adenylate kinase 6 [Source:MGI Symbol;Acc:MGI:5510732] | Ak6 | Enriched in P15 dLGN | 1.204 | 0.009 | 18.090 |
| ENSMUSG00000043162 | Pigy upstream reading frame [Source:MGI Symbol;Acc:MGI:1913709] | Pyurf | Enriched in P15 dLGN | 1.203 | 0.005 | 52.190 |
| ENSMUSG00000089911 | major facilitator superfamily domain containing 14A [Source:MGI Symbol;Acc:MGI:1201609] | Mfsd14a | Enriched in P15 dLGN | 1.201 | 0.000 | 53.160 |
| ENSMUSG00000026526 | fumarate hydratase 1 [Source:MGI Symbol;Acc:MGI:95530] | Fh1 | Enriched in P15 dLGN | 1.199 | 0.001 | 124.000 |
| ENSMUSG00000034891 | synuclein, beta [Source:MGI Symbol;Acc:MGI:1889011] | Sncb | Enriched in P15 dLGN | 1.197 | 0.000 | 939.200 |
| ENSMUSG00000040373 | calcium channel, voltage-dependent, gamma subunit 5 [Source:MGI Symbol;Acc:MGI:2157946] | Cacng5 | Enriched in P15 dLGN | 1.196 | 0.000 | 203.800 |
| ENSMUSG00000024044 | erythrocyte membrane protein band 4.1 like 3 [Source:MGI Symbol;Acc:MGI:103008] | Epb41l3 | Enriched in P15 dLGN | 1.194 | 0.000 | 1430.000 |
| ENSMUSG00000026154 | succinate dehydrogenase complex assembly factor 4 [Source:MGI Symbol;Acc:MGI:1915252] | Sdhaf4 | Enriched in P15 dLGN | 1.194 | 0.024 | 17.290 |
| ENSMUSG00000007338 | mitochondrial ribosomal protein L49 [Source:MGI Symbol;Acc:MGI:108180] | Mrpl49 | Enriched in P15 dLGN | 1.193 | 0.000 | 74.110 |
| ENSMUSG00000039601 | regulator of calcineurin 2 [Source:MGI Symbol;Acc:MGI:1858219] | Rcan2 | Enriched in P15 dLGN | 1.193 | 0.000 | 311.500 |
| ENSMUSG00000040713 | cellular repressor of E1A-stimulated genes 1 [Source:MGI Symbol;Acc:MGI:1344382] | Creg1 | Enriched in P15 dLGN | 1.191 | 0.000 | 76.450 |
| ENSMUSG00000071074 | Yip1 domain family, member 3 [Source:MGI Symbol;Acc:MGI:106280] | Yipf3 | Enriched in P15 dLGN | 1.191 | 0.000 | 108.800 |
| ENSMUSG00000049092 | G protein-coupled receptor 137C [Source:MGI Symbol;Acc:MGI:1917963] | Gpr137c | Enriched in P15 dLGN | 1.186 | 0.001 | 120.200 |
| ENSMUSG00000063172 | heat shock protein family B (small), member 11 [Source:MGI Symbol;Acc:MGI:1920188] | Hspb11 | Enriched in P15 dLGN | 1.186 | 0.048 | 24.210 |
| ENSMUSG00000035232 | pyruvate dehydrogenase kinase, isoenzyme 3 [Source:MGI Symbol;Acc:MGI:2384308] | Pdk3 | Enriched in P15 dLGN | 1.185 | 0.003 | 64.160 |
| ENSMUSG00000015476 | proline-rich transmembrane protein 1 [Source:MGI Symbol;Acc:MGI:1932118] | Prrt1 | Enriched in P15 dLGN | 1.183 | 0.000 | 115.500 |
| ENSMUSG00000016252 | ATP synthase, H+ transporting, mitochondrial F1 complex, epsilon subunit [Source:MGI Symbol;Acc:MGI:1855697] | Atp5e | Enriched in P15 dLGN | 1.180 | 0.001 | 153.200 |
| ENSMUSG00000034566 | ATP synthase, H+ transporting, mitochondrial F0 complex, subunit D [Source:MGI Symbol;Acc:MGI:1918929] | Atp5h | Enriched in P15 dLGN | 1.180 | 0.000 | 216.100 |
| ENSMUSG00000036934 | RIKEN cDNA 4921524J17 gene [Source:MGI Symbol;Acc:MGI:1913964] | 4921524J17I | Enriched in P15 dLGN | 1.177 | 0.002 | 42.540 |
| ENSMUSG00000021771 | voltage-dependent anion channel 2 [Source:MGI Symbol;Acc:MGI:106915] | Vdac2 | Enriched in P15 dLGN | 1.175 | 0.002 | 243.100 |
| ENSMUSG00000030849 | fibroblast growth factor receptor 2 [Source:MGI Symbol;Acc:MGI:95523] | Fgfr2 | Enriched in P15 dLGN | 1.175 | 0.037 | 937.900 |
| ENSMUSG00000032966 | FK506 binding protein 1a [Source:MGI Symbol;Acc:MGI:95541] | Fkbp1a | Enriched in P15 dLGN | 1.174 | 0.000 | 430.500 |
| ENSMUSG00000034880 | mitochondrial ribosomal protein L34 [Source:MGI Symbol;Acc:MGI:2137227] | Mrpl34 | Enriched in P15 dLGN | 1.174 | 0.000 | 34.190 |
| ENSMUSG00000062352 | integrin beta 1 binding protein 1 [Source:MGI Symbol;Acc:MGI:1306802] | Itgb1bp1 | Enriched in P15 dLGN | 1.173 | 0.000 | 59.330 |
| ENSMUSG00000026103 | glutaminase [Source:MGI Symbol;Acc:MGI:95752] | Gls | Enriched in P15 dLGN | 1.172 | 0.000 | 697.000 |
| ENSMUSG00000007564 | protein phosphatase 2, regulatory subunit A, alpha [Source:MGI Symbol;Acc:MGI:1926334] | Ppp2r1a | Enriched in P15 dLGN | 1.171 | 0.000 | 634.900 |
| ENSMUSG00000019210 | ATPase, H+ transporting, lysosomal V1 subunit E1 [Source:MGI Symbol;Acc:MGI:894326] | Atp6v1e1 | Enriched in P15 dLGN | 1.171 | 0.001 | 270.400 |
| ENSMUSG00000002379 | NADH:ubiquinone oxidoreductase subunit A11 [Source:MGI Symbol;Acc:MGI:1917125] | Ndufa11 | Enriched in P15 dLGN | 1.167 | 0.019 | 75.680 |
| ENSMUSG00000041444 | Rho GTPase activating protein 32 [Source:MGI Symbol;Acc:MGI:2450166] | Arhgap32 | Enriched in P15 dLGN | 1.165 | 0.000 | 970.500 |
| ENSMUSG00000063882 | ubiquinol-cytochrome c reductase hinge protein [Source:MGI Symbol;Acc:MGI:1913826] | Uqcrh | Enriched in P15 dLGN | 1.164 | 0.000 | 256.500 |
| ENSMUSG00000075702 | selenoprotein M [Source:MGI Symbol;Acc:MGI:2149786] | Selenom | Enriched in P15 dLGN | 1.163 | 0.000 | 136.200 |
| ENSMUSG00000031451 | growth arrest specific 6 [Source:MGI Symbol;Acc:MGI:95660] | Gas6 | Enriched in P15 dLGN | 1.162 | 0.000 | 424.500 |
| ENSMUSG00000036955 | kinesin family binding protein [Source:MGI Symbol;Acc:MGI:1919570] | Kifbp | Enriched in P15 dLGN | 1.161 | 0.000 | 286.800 |
| ENSMUSG00000000088 | cytochrome c oxidase subunit 5A [Source:MGI Symbol;Acc:MGI:88474] | Cox5a | Enriched in P15 dLGN | 1.160 | 0.000 | 112.100 |
| ENSMUSG00000019179 | malate dehydrogenase 2, NAD (mitochondrial) [Source:MGI Symbol;Acc:MGI:97050] | Mdh2 | Enriched in P15 dLGN | 1.157 | 0.001 | 626.300 |
| ENSMUSG00000020485 | SPT4A, DSIF elongation factor subunit [Source:MGI Symbol;Acc:MGI:107416] | Supt4a | Enriched in P15 dLGN | 1.157 | 0.001 | 25.990 |
| ENSMUSG00000025968 | NADH:ubiquinone oxidoreductase core subunit S1 [Source:MGI Symbol;Acc:MGI:2443241] | Ndufs1 | Enriched in P15 dLGN | 1.157 | 0.000 | 337.900 |
| ENSMUSG00000038462 | ubiquinol-cytochrome c reductase, Rieske iron-sulfur polypeptide 1 [Source:MGI Symbol;Acc:MGI:1913944] | Uqcrfs1 | Enriched in P15 dLGN | 1.157 | 0.000 | 172.400 |
| ENSMUSG00000031059 | NADH:ubiquinone oxidoreductase subunit B11 [Source:MGI Symbol;Acc:MGI:1349919] | Ndufb11 | Enriched in P15 dLGN | 1.156 | 0.000 | 155.400 |
| ENSMUSG00000029189 | sel-1 suppressor of lin-12-like 3 (C. elegans) [Source:MGI Symbol;Acc:MGI:1916941] | Sel1l3 | Enriched in P15 dLGN | 1.154 | 0.000 | 154.000 |
| ENSMUSG00000020163 | ubiquinol-cytochrome c reductase, complex III subunit XI [Source:MGI Symbol;Acc:MGI:1913844] | Uqcr11 | Enriched in P15 dLGN | 1.151 | 0.002 | 153.300 |
| ENSMUSG00000019797 | mitochondrial transcription rescue factor 1 [Source:MGI Symbol;Acc:MGI:1915101] | Mtres1 | Enriched in P15 dLGN | 1.149 | 0.000 | 74.570 |
| ENSMUSG00000021939 | cathepsin B [Source:MGI Symbol;Acc:MGI:88561] | Ctsb | Enriched in P15 dLGN | 1.148 | 0.001 | 510.600 |
| ENSMUSG00000042743 | small glutamine-rich tetratricopeptide repeat (TPR)-containing, beta [Source:MGI Symbol;Acc:MGI:2444615] | Sgtb | Enriched in P15 dLGN | 1.147 | 0.000 | 167.800 |
| ENSMUSG00000039478 | mitochondrial calcium uptake family, member 3 [Source:MGI Symbol;Acc:MGI:1925756] | Micu3 | Enriched in P15 dLGN | 1.145 | 0.000 | 313.100 |
| ENSMUSG00000020889 | nuclear receptor subfamily 1, group D, member 1 [Source:MGI Symbol;Acc:MGI:2444210] | Nr1d1 | Enriched in P15 dLGN | 1.143 | 0.000 | 187.400 |
| ENSMUSG00000025781 | ATP synthase, H+ transporting, mitochondrial F1 complex, gamma polypeptide 1 [Source:MGI Symbol;Acc:MGI:1261437] | Atp5c1 | Enriched in P15 dLGN | 1.143 | 0.000 | 290.000 |
| ENSMUSG00000031668 | eukaryotic translation initiation factor 2 alpha kinase 3 [Source:MGI Symbol;Acc:MGI:1341830] | Eif2ak3 | Enriched in P15 dLGN | 1.143 | 0.000 | 198.600 |
| ENSMUSG00000071662 | polymerase (RNA) II (DNA directed) polypeptide G [Source:MGI Symbol;Acc:MGI:1914960] | Polr2g | Enriched in P15 dLGN | 1.143 | 0.000 | 40.090 |
| ENSMUSG00000022108 | integral membrane protein 2B [Source:MGI Symbol;Acc:MGI:1309517] | Itm2b | Enriched in P15 dLGN | 1.140 | 0.000 | 723.800 |
| ENSMUSG00000025630 | hypoxanthine guanine phosphoribosyl transferase [Source:MGI Symbol;Acc:MGI:96217] | Hprt | Enriched in P15 dLGN | 1.140 | 0.000 | 174.600 |
| ENSMUSG00000021209 | protein phosphatase 4, regulatory subunit 4 [Source:MGI Symbol;Acc:MGI:1921771] | Ppp4r4 | Enriched in P15 dLGN | 1.139 | 0.000 | 139.400 |
| ENSMUSG00000024127 | prolyl endopeptidase-like [Source:MGI Symbol;Acc:MGI:2441932] | Prepl | Enriched in P15 dLGN | 1.134 | 0.000 | 815.800 |
| ENSMUSG00000071654 | ubiquinol-cytochrome c reductase complex assembly factor 3 [Source:MGI Symbol;Acc:MGI:2147553] | Uqcc3 | Enriched in P15 dLGN | 1.134 | 0.000 | 94.540 |
| ENSMUSG00000040048 | NADH:ubiquinone oxidoreductase subunit B10 [Source:MGI Symbol;Acc:MGI:1915592] | Ndufb10 | Enriched in P15 dLGN | 1.133 | 0.000 | 201.200 |
| ENSMUSG00000022257 | lysosomal-associated protein transmembrane 4B [Source:MGI Symbol;Acc:MGI:1890494] | Laptm4b | Enriched in P15 dLGN | 1.131 | 0.000 | 165.800 |
| ENSMUSG00000021711 | trafficking protein particle complex 13 [Source:MGI Symbol;Acc:MGI:1914225] | Trappc13 | Enriched in P15 dLGN | 1.130 | 0.000 | 71.190 |
| ENSMUSG00000030652 | demethyl-Q 7 [Source:MGI Symbol;Acc:MGI:107207] | Coq7 | Enriched in P15 dLGN | 1.127 | 0.000 | 54.520 |
| ENSMUSG00000006024 | N-ethylmaleimide sensitive fusion protein attachment protein alpha [Source:MGI Symbol;Acc:MGI:104563] | Napa | Enriched in P15 dLGN | 1.125 | 0.000 | 230.600 |

| | | | | | | |
|---|---|---|---|---|---|---|
| ENSMUSG00000031556 | TM2 domain containing 2 [Source:MGI Symbol;Acc:MGI:1916992] | Tm2d2 | Enriched in P15 dLGN | 1.124 | 0.001 | 62.810 |
| ENSMUSG00000021814 | annexin A7 [Source:MGI Symbol;Acc:MGI:88031] | Anxa7 | Enriched in P15 dLGN | 1.123 | 0.002 | 56.220 |
| ENSMUSG00000028528 | DnaJ heat shock protein family (Hsp40) member C6 [Source:MGI Symbol;Acc:MGI:1919935] | Dnajc6 | Enriched in P15 dLGN | 1.123 | 0.000 | 781.500 |
| ENSMUSG00000024925 | ribonuclease H2, subunit C [Source:MGI Symbol;Acc:MGI:1915459] | Rnaseh2c | Enriched in P15 dLGN | 1.122 | 0.006 | 33.010 |
| ENSMUSG00000038845 | prohibitin [Source:MGI Symbol;Acc:MGI:97572] | Phb | Enriched in P15 dLGN | 1.122 | 0.000 | 71.590 |
| ENSMUSG00000021987 | myotubularin related protein 6 [Source:MGI Symbol;Acc:MGI:2145637] | Mtmr6 | Enriched in P15 dLGN | 1.119 | 0.000 | 223.600 |
| ENSMUSG00000026687 | aldehyde dehydrogenase 9, subfamily A1 [Source:MGI Symbol;Acc:MGI:1861622] | Aldh9a1 | Enriched in P15 dLGN | 1.119 | 0.000 | 102.900 |
| ENSMUSG00000035885 | cytochrome c oxidase subunit 8A [Source:MGI Symbol;Acc:MGI:105959] | Cox8a | Enriched in P15 dLGN | 1.119 | 0.000 | 292.600 |
| ENSMUSG00000073471 | radial spoke 3A homolog (Chlamydomonas) [Source:MGI Symbol;Acc:MGI:1914082] | Rsph3a | Enriched in P15 dLGN | 1.119 | 0.030 | 41.680 |
| ENSMUSG00000039347 | ATPase, H+ transporting, lysosomal V0 subunit E2 [Source:MGI Symbol;Acc:MGI:1923502] | Atp6v0e2 | Enriched in P15 dLGN | 1.118 | 0.000 | 554.600 |
| ENSMUSG00000027088 | phosphatase, orphan 2 [Source:MGI Symbol;Acc:MGI:1920623] | Phospho2 | Enriched in P15 dLGN | 1.116 | 0.000 | 44.610 |
| ENSMUSG00000020664 | dihydrolipoamide dehydrogenase [Source:MGI Symbol;Acc:MGI:107450] | Dld | Enriched in P15 dLGN | 1.115 | 0.000 | 258.400 |
| ENSMUSG00000041112 | engulfment and cell motility 1 [Source:MGI Symbol;Acc:MGI:2153044] | Elmo1 | Enriched in P15 dLGN | 1.115 | 0.001 | 598.800 |
| ENSMUSG00000021114 | ATPase, H+ transporting, lysosomal V1 subunit D [Source:MGI Symbol;Acc:MGI:1921084] | Atp6v1d | Enriched in P15 dLGN | 1.114 | 0.000 | 284.800 |
| ENSMUSG00000031633 | solute carrier family 25 (mitochondrial carrier, adenine nucleotide translocator), member 4 [Source:MGI Symbol;Acc:MGI:1353495] | Slc25a4 | Enriched in P15 dLGN | 1.113 | 0.000 | 1489.000 |
| ENSMUSG00000029592 | ubiquitin specific peptidase 30 [Source:MGI Symbol;Acc:MGI:2140991] | Usp30 | Enriched in P15 dLGN | 1.110 | 0.000 | 129.000 |
| ENSMUSG00000067860 | zinc finger protein of the cerebellum 3 [Source:MGI Symbol;Acc:MGI:106676] | Zic3 | Enriched in P15 dLGN | 1.105 | 0.008 | 26.920 |
| ENSMUSG00000019951 | UHRF1 (ICBP90) binding protein 1-like [Source:MGI Symbol;Acc:MGI:2442888] | Uhrf1bp1l | Enriched in P15 dLGN | 1.102 | 0.000 | 276.600 |
| ENSMUSG00000036578 | FXYD domain-containing ion transport regulator 7 [Source:MGI Symbol;Acc:MGI:1889006] | Fxyd7 | Enriched in P15 dLGN | 1.102 | 0.004 | 140.400 |
| ENSMUSG00000071014 | NADH:ubiquinone oxidoreductase subunit B6 [Source:MGI Symbol;Acc:MGI:2684983] | Ndufb6 | Enriched in P15 dLGN | 1.102 | 0.002 | 50.890 |
| ENSMUSG00000060279 | adaptor-related protein complex 2, alpha 1 subunit [Source:MGI Symbol;Acc:MGI:101921] | Ap2a1 | Enriched in P15 dLGN | 1.100 | 0.000 | 671.900 |
| ENSMUSG00000027076 | translocase of inner mitochondrial membrane 10 [Source:MGI Symbol;Acc:MGI:1353429] | Timm10 | Enriched in P15 dLGN | 1.098 | 0.005 | 29.720 |
| ENSMUSG00000035790 | centrosomal protein 19 [Source:MGI Symbol;Acc:MGI:1914244] | Cep19 | Enriched in P15 dLGN | 1.097 | 0.000 | 80.500 |
| ENSMUSG00000038244 | microtubule associated monooxygenase, calponin and LIM domain containing 2 [Source:MGI Symbol;Acc:MGI:2444947] | Mical2 | Enriched in P15 dLGN | 1.093 | 0.001 | 255.100 |
| ENSMUSG00000036398 | protein phosphatase 1, regulatory inhibitor subunit 11 [Source:MGI Symbol;Acc:MGI:1923747] | Ppp1r11 | Enriched in P15 dLGN | 1.088 | 0.000 | 110.100 |
| ENSMUSG00000034640 | TCDD-inducible poly(ADP-ribose) polymerase [Source:MGI Symbol;Acc:MGI:2159210] | Tiparp | Enriched in P15 dLGN | 1.087 | 0.001 | 38.050 |
| ENSMUSG00000033918 | presenilin associated, rhomboid-like [Source:MGI Symbol;Acc:MGI:1277152] | Parl | Enriched in P15 dLGN | 1.086 | 0.000 | 65.890 |
| ENSMUSG00000001445 | mitochondrial ribosomal protein L10 [Source:MGI Symbol;Acc:MGI:1333801] | Mrpl10 | Enriched in P15 dLGN | 1.082 | 0.000 | 68.560 |
| ENSMUSG00000005510 | NADH:ubiquinone oxidoreductase core subunit S3 [Source:MGI Symbol;Acc:MGI:1915599] | Ndufs3 | Enriched in P15 dLGN | 1.082 | 0.000 | 101.100 |
| ENSMUSG00000038264 | sema domain, immunoglobulin domain (Ig), and GPI membrane anchor, (semaphorin) 7A [Source:MGI Symbol;Acc:MGI:1306826] | Sema7a | Enriched in P15 dLGN | 1.082 | 0.000 | 229.800 |
| ENSMUSG00000020029 | nudix (nucleoside diphosphate linked moiety X)-type motif 4 [Source:MGI Symbol;Acc:MGI:1918457] | Nudt4 | Enriched in P15 dLGN | 1.081 | 0.000 | 752.500 |
| ENSMUSG00000024935 | solute carrier family 1 (neuronal/epithelial high affinity glutamate transporter, system Xag), member 1 [Source:MGI Symbol;Acc:MGI:1050 | Slc1a1 | Enriched in P15 dLGN | 1.077 | 0.000 | 323.700 |
| ENSMUSG00000010914 | pyruvate dehydrogenase complex, component X [Source:MGI Symbol;Acc:MGI:1351627] | Pdhx | Enriched in P15 dLGN | 1.075 | 0.000 | 152.700 |
| ENSMUSG00000052456 | guided entry of tail-anchored proteins factor 3, ATPase [Source:MGI Symbol;Acc:MGI:1928379] | Get3 | Enriched in P15 dLGN | 1.072 | 0.000 | 120.500 |
| ENSMUSG00000020333 | acyl-CoA synthetase long-chain family member 6 [Source:MGI Symbol;Acc:MGI:894291] | Acsl6 | Enriched in P15 dLGN | 1.070 | 0.000 | 594.100 |
| ENSMUSG00000021314 | amphiphysin [Source:MGI Symbol;Acc:MGI:103574] | Amph | Enriched in P15 dLGN | 1.070 | 0.000 | 841.600 |
| ENSMUSG00000030612 | mitochondrial ribosomal protein L46 [Source:MGI Symbol;Acc:MGI:1914558] | Mrpl46 | Enriched in P15 dLGN | 1.070 | 0.002 | 33.730 |
| ENSMUSG00000047126 | clathrin, heavy polypeptide (Hc) [Source:MGI Symbol;Acc:MGI:2388633] | Cltc | Enriched in P15 dLGN | 1.069 | 0.000 | 1529.000 |
| ENSMUSG00000031753 | component of oligomeric golgi complex 4 [Source:MGI Symbol;Acc:MGI:2142808] | Cog4 | Enriched in P15 dLGN | 1.068 | 0.000 | 122.400 |
| ENSMUSG00000019505 | ubiquitin B [Source:MGI Symbol;Acc:MGI:98888] | Ubb | Enriched in P15 dLGN | 1.066 | 0.014 | 652.000 |
| ENSMUSG00000021156 | zinc finger, MYND domain containing 11 [Source:MGI Symbol;Acc:MGI:1913755] | Zmynd11 | Enriched in P15 dLGN | 1.066 | 0.000 | 980.200 |
| ENSMUSG00000031432 | phosphoribosyl pyrophosphate synthetase 1 [Source:MGI Symbol;Acc:MGI:97775] | Prps1 | Enriched in P15 dLGN | 1.064 | 0.001 | 65.650 |
| ENSMUSG00000059278 | N(alpha)-acetyltransferase 38, NatC auxiliary subunit [Source:MGI Symbol;Acc:MGI:1925554] | Naa38 | Enriched in P15 dLGN | 1.063 | 0.011 | 31.830 |
| ENSMUSG00000038880 | mitochondrial ribosomal protein S34 [Source:MGI Symbol;Acc:MGI:1930188] | Mrps34 | Enriched in P15 dLGN | 1.061 | 0.001 | 43.830 |
| ENSMUSG00000029017 | peptidase (mitochondrial processing) beta [Source:MGI Symbol;Acc:MGI:1920328] | Pmpcb | Enriched in P15 dLGN | 1.060 | 0.000 | 90.430 |
| ENSMUSG00000029047 | peroxisomal biogenesis factor 10 [Source:MGI Symbol;Acc:MGI:2684988] | Pex10 | Enriched in P15 dLGN | 1.059 | 0.012 | 29.080 |
| ENSMUSG00000024758 | reticulon 3 [Source:MGI Symbol;Acc:MGI:1339970] | Rtn3 | Enriched in P15 dLGN | 1.057 | 0.000 | 1535.000 |
| ENSMUSG00000024099 | NADH:ubiquinone oxidoreductase core subunit V2 [Source:MGI Symbol;Acc:MGI:1920150] | Ndufv2 | Enriched in P15 dLGN | 1.056 | 0.000 | 217.500 |
| ENSMUSG00000023572 | cyclin D-type binding-protein 1 [Source:MGI Symbol;Acc:MGI:109595] | Ccndbp1 | Enriched in P15 dLGN | 1.054 | 0.000 | 105.300 |
| ENSMUSG00000025964 | a disintegrin and metallopeptidase domain 23 [Source:MGI Symbol;Acc:MGI:1345162] | Adam23 | Enriched in P15 dLGN | 1.054 | 0.000 | 982.600 |
| ENSMUSG00000036067 | solute carrier family 2 (facilitated glucose transporter), member 6 [Source:MGI Symbol;Acc:MGI:2443286] | Slc2a6 | Enriched in P15 dLGN | 1.054 | 0.007 | 33.470 |
| ENSMUSG00000031993 | sorting nexin 19 [Source:MGI Symbol;Acc:MGI:1921581] | Snx19 | Enriched in P15 dLGN | 1.052 | 0.000 | 263.600 |
| ENSMUSG00000055943 | ER membrane protein complex subunit 7 [Source:MGI Symbol;Acc:MGI:1920274] | Emc7 | Enriched in P15 dLGN | 1.050 | 0.000 | 203.900 |
| ENSMUSG00000034793 | glucose 6 phosphatase, catalytic, 3 [Source:MGI Symbol;Acc:MGI:1915651] | G6pc3 | Enriched in P15 dLGN | 1.049 | 0.002 | 38.970 |
| ENSMUSG00000028447 | dynactin 3 [Source:MGI Symbol;Acc:MGI:1859251] | Dctn3 | Enriched in P15 dLGN | 1.044 | 0.000 | 123.300 |
| ENSMUSG00000031950 | gamma-aminobutyric acid (GABA) A receptor-associated protein-like 2 [Source:MGI Symbol;Acc:MGI:1890602] | Gabarapl2 | Enriched in P15 dLGN | 1.041 | 0.000 | 147.200 |
| ENSMUSG00000053025 | synaptic vesicle glycoprotein 2 b [Source:MGI Symbol;Acc:MGI:1927338] | Sv2b | Enriched in P15 dLGN | 1.040 | 0.000 | 914.900 |
| ENSMUSG00000042182 | BEN domain containing 6 [Source:MGI Symbol;Acc:MGI:2444572] | Bend6 | Enriched in P15 dLGN | 1.039 | 0.000 | 256.000 |
| ENSMUSG00000022450 | NADH:ubiquinone oxidoreductase subunit A6 [Source:MGI Symbol;Acc:MGI:1914380] | Ndufa6 | Enriched in P15 dLGN | 1.037 | 0.000 | 126.100 |
| ENSMUSG00000064202 | spermatogenesis associated 6 like [Source:MGI Symbol;Acc:MGI:1918036] | Spata6l | Enriched in P15 dLGN | 1.037 | 0.018 | 24.410 |
| ENSMUSG00000033460 | armadillo repeat containing, X-linked 1 [Source:MGI Symbol;Acc:MGI:1925498] | Armcx1 | Enriched in P15 dLGN | 1.033 | 0.000 | 200.900 |
| ENSMUSG00000028251 | thiosulfate sulfurtransferase (rhodanese)-like domain containing 3 [Source:MGI Symbol;Acc:MGI:1924282] | Tstd3 | Enriched in P15 dLGN | 1.031 | 0.012 | 35.110 |
| ENSMUSG00000048376 | coagulation factor II (thrombin) receptor [Source:MGI Symbol;Acc:MGI:101802] | F2r | Enriched in P15 dLGN | 1.031 | 0.028 | 37.830 |

| | | | | | | |
|---|---|---|---|---|---|---|
| ENSMUSG00000020544 | cytochrome c oxidase assembly protein 11, copper chaperone [Source:MGI Symbol;Acc:MGI:1917052] | Cox11 | Enriched in P15 dLGN | 1.030 | 0.001 | 38.980 |
| ENSMUSG00000034557 | zinc finger, FYVE domain containing 9 [Source:MGI Symbol;Acc:MGI:2652838] | Zfyve9 | Enriched in P15 dLGN | 1.029 | 0.005 | 559.400 |
| ENSMUSG00000075706 | glutathione peroxidase 4 [Source:MGI Symbol;Acc:MGI:104767] | Gpx4 | Enriched in P15 dLGN | 1.028 | 0.000 | 170.900 |
| ENSMUSG00000022658 | transgelin 3 [Source:MGI Symbol;Acc:MGI:1926784] | Tagln3 | Enriched in P15 dLGN | 1.025 | 0.003 | 267.200 |
| ENSMUSG00000015478 | ring finger protein 5 [Source:MGI Symbol;Acc:MGI:1860076] | Rnf5 | Enriched in P15 dLGN | 1.021 | 0.001 | 74.490 |
| ENSMUSG00000019795 | protein-L-isoaspartate (D-aspartate) O-methyltransferase 1 [Source:MGI Symbol;Acc:MGI:97502] | Pcmt1 | Enriched in P15 dLGN | 1.021 | 0.000 | 219.500 |
| ENSMUSG00000031770 | homocysteine-inducible, endoplasmic reticulum stress-inducible, ubiquitin-like domain member 1 [Source:MGI Symbol;Acc:MGI:1927406] | Herpud1 | Enriched in P15 dLGN | 1.019 | 0.000 | 93.700 |
| ENSMUSG00000041020 | MAP7 domain containing 2 [Source:MGI Symbol;Acc:MGI:1917474] | Map7d2 | Enriched in P15 dLGN | 1.018 | 0.000 | 736.500 |
| ENSMUSG00000024966 | stress-induced phosphoprotein 1 [Source:MGI Symbol;Acc:MGI:109130] | Stip1 | Enriched in P15 dLGN | 1.016 | 0.003 | 255.000 |
| ENSMUSG00000021368 | TBC1 domain family, member 7 [Source:MGI Symbol;Acc:MGI:1914296] | Tbc1d7 | Enriched in P15 dLGN | 1.015 | 0.002 | 35.710 |
| ENSMUSG00000039953 | calsyntenin 1 [Source:MGI Symbol;Acc:MGI:1929895] | Clstn1 | Enriched in P15 dLGN | 1.013 | 0.000 | 3184.000 |
| ENSMUSG00000029598 | phospholipase B domain containing 2 [Source:MGI Symbol;Acc:MGI:1919022] | Plbd2 | Enriched in P15 dLGN | 1.011 | 0.000 | 103.500 |
| ENSMUSG00000030706 | mitochondrial ribosomal protein L48 [Source:MGI Symbol;Acc:MGI:1289321] | Mrpl48 | Enriched in P15 dLGN | 1.011 | 0.005 | 153.100 |
| ENSMUSG00000022024 | SGT1, suppressor of G2 allele of SKP1 (S. cerevisiae) [Source:MGI Symbol;Acc:MGI:1915205] | Sugt1 | Enriched in P15 dLGN | 1.008 | 0.000 | 199.600 |
| ENSMUSG00000027263 | tubulin, gamma complex associated protein 4 [Source:MGI Symbol;Acc:MGI:1196293] | Tubgcp4 | Enriched in P15 dLGN | 1.008 | 0.000 | 80.840 |
| ENSMUSG00000015002 | EFR3 homolog A [Source:MGI Symbol;Acc:MGI:1923990] | Efr3a | Enriched in P15 dLGN | 1.007 | 0.000 | 259.300 |
| ENSMUSG00000003380 | Rab acceptor 1 (prenylated) [Source:MGI Symbol;Acc:MGI:1201692] | Rabac1 | Enriched in P15 dLGN | 1.004 | 0.000 | 110.700 |
| ENSMUSG00000023175 | basigin [Source:MGI Symbol;Acc:MGI:88208] | Bsg | Enriched in P15 dLGN | 1.004 | 0.000 | 872.500 |
| ENSMUSG00000039914 | coenzyme Q10A [Source:MGI Symbol;Acc:MGI:2684847] | Coq10a | Enriched in P15 dLGN | 1.004 | 0.017 | 58.280 |
| ENSMUSG00000009894 | synaptosomal-associated protein, 47 [Source:MGI Symbol;Acc:MGI:1915076] | Snap47 | Enriched in P15 dLGN | 1.002 | 0.002 | 533.600 |
| ENSMUSG00000024121 | ATPase, H+ transporting, lysosomal V0 subunit C [Source:MGI Symbol;Acc:MGI:88116] | Atp6v0c | Enriched in P15 dLGN | 1.001 | 0.001 | 333.400 |
| ENSMUSG00000045763 | brain abundant, membrane attached signal protein 1 [Source:MGI Symbol;Acc:MGI:1917600] | Basp1 | Enriched in P8 dLGN | -1.025 | 0.000 | 3099.000 |
| ENSMUSG00000042942 | growth regulation by estrogen in breast cancer-like [Source:MGI Symbol;Acc:MGI:3576497] | Greb1l | Enriched in P8 dLGN | -1.049 | 0.005 | 63.670 |
| ENSMUSG00000024172 | beta galactoside alpha 2,6 sialyltransferase 2 [Source:MGI Symbol;Acc:MGI:2445190] | St6gal2 | Enriched in P8 dLGN | -1.056 | 0.000 | 150.300 |
| ENSMUSG00000091955 | predicted pseudogene 9844 [Source:MGI Symbol;Acc:MGI:3704288] | Gm9844 | Enriched in P8 dLGN | -1.206 | 0.000 | 233.800 |
| ENSMUSG00000028007 | sorting nexin 7 [Source:MGI Symbol;Acc:MGI:1923811] | Snx7 | Enriched in P8 dLGN | -1.396 | 0.000 | 54.520 |
| ENSMUSG00000028832 | stathmin 1 [Source:MGI Symbol;Acc:MGI:96739] | Stmn1 | Enriched in P8 dLGN | -1.471 | 0.000 | 644.000 |
| ENSMUSG00000018589 | glycine receptor, alpha 2 subunit [Source:MGI Symbol;Acc:MGI:95748] | Glra2 | Enriched in P8 dLGN | -1.541 | 0.000 | 84.200 |
| ENSMUSG00000027496 | aurora kinase A [Source:MGI Symbol;Acc:MGI:894678] | Aurka | Enriched in P8 dLGN | -1.546 | 0.017 | 6.790 |
| ENSMUSG00000030137 | tubulin, alpha 8 [Source:MGI Symbol;Acc:MGI:1858225] | Tuba8 | Enriched in P8 dLGN | -1.864 | 0.000 | 18.260 |
| ENSMUSG00000079410 | predicted gene 2897 [Source:MGI Symbol;Acc:MGI:3781075] | Gm2897 | Enriched in P8 dLGN | -1.939 | 0.029 | 6.752 |
| ENSMUSG00000070883 | coiled-coil domain containing 173 [Source:MGI Symbol;Acc:MGI:1923100] | Ccdc173 | Enriched in P8 dLGN | -1.958 | 0.003 | 57.550 |
| ENSMUSG00000096740 | LBH domain containing 1 [Source:MGI Symbol;Acc:MGI:5516029] | Lbhd1 | Enriched in P8 dLGN | -3.053 | 0.027 | 104.600 |
| ENSMUSG00000025094 | solute carrier family 18 (vesicular monoamine), member 2 [Source:MGI Symbol;Acc:MGI:106677] | Slc18a2 | Enriched in P8 dLGN | -3.173 | 0.000 | 54.240 |
| ENSMUSG00000020838 | solute carrier family 6 (neurotransmitter transporter, serotonin), member 4 [Source:MGI Symbol;Acc:MGI:96285] | Slc6a4 | Enriched in P8 dLGN | -5.132 | 0.000 | 23.260 |

**Contrast numerator: P15_KO_dLGN. Contrast denominator: P8_KO_dLGN.**

| ensembl_gene_id | description | mgi_symbol | Significance | deseq_logfc | deseq_adjp | deseq_basemean |
|---|---|---|---|---|---|---|
| ENSMUSG00000034452 | solute carrier family 24 (sodium/potassium/calcium exchanger), member 1 [Source:MGI Symbol;Acc:MGI:2384871] | Slc24a1 | Enriched in P15 dLGN | 7.434 | 0.000 | 1018.000 |
| ENSMUSG00000024227 | PDZ and pleckstrin homology domains 1 [Source:MGI Symbol;Acc:MGI:1916489] | Pdzph1 | Enriched in P15 dLGN | 6.499 | 0.000 | 255.200 |
| ENSMUSG00000044375 | photoreceptor cilium actin regulator [Source:MGI Symbol;Acc:MGI:2385061] | Pcare | Enriched in P15 dLGN | 6.497 | 0.000 | 962.800 |
| ENSMUSG00000021948 | protein kinase C, delta [Source:MGI Symbol;Acc:MGI:97598] | Prkcd | Enriched in P15 dLGN | 6.430 | 0.000 | 727.900 |
| ENSMUSG00000034837 | G protein subunit alpha transducin 1 [Source:MGI Symbol;Acc:MGI:95778] | Gnat1 | Enriched in P15 dLGN | 6.374 | 0.001 | 1840.000 |
| ENSMUSG00000031450 | G protein-coupled receptor kinase 1 [Source:MGI Symbol;Acc:MGI:1345146] | Grk1 | Enriched in P15 dLGN | 5.959 | 0.001 | 1629.000 |
| ENSMUSG00000031293 | retinoschisis (X-linked, juvenile) 1 (human) [Source:MGI Symbol;Acc:MGI:1336189] | Rs1 | Enriched in P15 dLGN | 5.638 | 0.001 | 1028.000 |
| ENSMUSG00000110344 | small integral membrane protein 36 [Source:MGI Symbol;Acc:MGI:5804831] | Smim36 | Enriched in P15 dLGN | 5.534 | 0.001 | 107.500 |
| ENSMUSG00000041044 | leucine-rich repeat, immunoglobulin-like and transmembrane domains 1 [Source:MGI Symbol;Acc:MGI:2385320] | Lrit1 | Enriched in P15 dLGN | 5.511 | 0.003 | 386.500 |
| ENSMUSG00000029054 | gamma-aminobutyric acid (GABA) A receptor, subunit delta [Source:MGI Symbol;Acc:MGI:95622] | Gabrd | Enriched in P15 dLGN | 5.289 | 0.000 | 52.020 |
| ENSMUSG00000027489 | N-terminal EF-hand calcium binding protein 3 [Source:MGI Symbol;Acc:MGI:1861721] | Necab3 | Enriched in P15 dLGN | 5.220 | 0.000 | 36.720 |
| ENSMUSG00000056043 | regulator of G-protein signalling 9 binding protein [Source:MGI Symbol;Acc:MGI:2384418] | Rgs9bp | Enriched in P15 dLGN | 5.031 | 0.001 | 666.000 |
| ENSMUSG00000045776 | leucine-rich repeats and transmembrane domains 1 [Source:MGI Symbol;Acc:MGI:2442106] | Lrtm1 | Enriched in P15 dLGN | 5.014 | 0.006 | 127.300 |
| ENSMUSG00000008932 | solute carrier family 1 (glutamate transporter), member 7 [Source:MGI Symbol;Acc:MGI:2444087] | Slc1a7 | Enriched in P15 dLGN | 4.956 | 0.002 | 185.900 |
| ENSMUSG00000024519 | complexin 4 [Source:MGI Symbol;Acc:MGI:2685803] | Cplx4 | Enriched in P15 dLGN | 4.846 | 0.010 | 325.100 |
| ENSMUSG00000019762 | iodotyrosine deiodinase [Source:MGI Symbol;Acc:MGI:1917587] | Iyd | Enriched in P15 dLGN | 4.768 | 0.002 | 6.399 |
| ENSMUSG00000021123 | retinol dehydrogenase 12 [Source:MGI Symbol;Acc:MGI:1925224] | Rdh12 | Enriched in P15 dLGN | 4.678 | 0.004 | 285.000 |
| ENSMUSG00000043461 | serine palmitoyltransferase, small subunit B [Source:MGI Symbol;Acc:MGI:1913433] | Sptssb | Enriched in P15 dLGN | 4.612 | 0.002 | 9.484 |
| ENSMUSG00000024575 | phosphodiesterase 6A, cGMP-specific, rod, alpha [Source:MGI Symbol;Acc:MGI:97524] | Pde6a | Enriched in P15 dLGN | 4.611 | 0.003 | 1301.000 |
| ENSMUSG00000030324 | rhodopsin [Source:MGI Symbol;Acc:MGI:97914] | Rho | Enriched in P15 dLGN | 4.604 | 0.001 | 5636.000 |
| ENSMUSG00000070683 | lactamase, beta-like 1 [Source:MGI Symbol;Acc:MGI:2448566] | Lactbl1 | Enriched in P15 dLGN | 4.527 | 0.013 | 222.900 |
| ENSMUSG00000093865 | leucine-rich repeat, immunoglobulin-like and transmembrane domains 3 [Source:MGI Symbol;Acc:MGI:2685267] | Lrit3 | Enriched in P15 dLGN | 4.472 | 0.018 | 120.300 |
| ENSMUSG00000047298 | potassium channel, subfamily V, member 2 [Source:MGI Symbol;Acc:MGI:2670981] | Kcnv2 | Enriched in P15 dLGN | 4.445 | 0.008 | 537.700 |
| ENSMUSG00000053773 | retinol dehydrogenase 8 [Source:MGI Symbol;Acc:MGI:2685028] | Rdh8 | Enriched in P15 dLGN | 4.319 | 0.007 | 87.450 |
| ENSMUSG00000027347 | RAS guanyl releasing protein 1 [Source:MGI Symbol;Acc:MGI:1314635] | Rasgrp1 | Enriched in P15 dLGN | 4.298 | 0.000 | 353.400 |
| ENSMUSG00000031379 | pirin [Source:MGI Symbol;Acc:MGI:1916906] | Pir | Enriched in P15 dLGN | 4.214 | 0.005 | 5.860 |
| ENSMUSG00000025900 | retinitis pigmentosa 1 (human) [Source:MGI Symbol;Acc:MGI:1341105] | Rp1 | Enriched in P15 dLGN | 4.032 | 0.001 | 3382.000 |
| ENSMUSG00000111755 | predicted gene, 47160 [Source:MGI Symbol;Acc:MGI:6095933] | Gm47160 | Enriched in P15 dLGN | 4.028 | 0.043 | 97.350 |
| ENSMUSG00000035270 | interphotoreceptor matrix proteoglycan 2 [Source:MGI Symbol;Acc:MGI:3044955] | Impg2 | Enriched in P15 dLGN | 3.963 | 0.001 | 754.300 |
| ENSMUSG00000040554 | aryl hydrocarbon receptor-interacting protein-like 1 [Source:MGI Symbol;Acc:MGI:2148800] | Aipl1 | Enriched in P15 dLGN | 3.859 | 0.025 | 368.200 |
| ENSMUSG00000005649 | calcium binding protein 5 [Source:MGI Symbol;Acc:MGI:1352746] | Cabp5 | Enriched in P15 dLGN | 3.833 | 0.013 | 50.220 |
| ENSMUSG00000023978 | peripherin 2 [Source:MGI Symbol;Acc:MGI:102791] | Prph2 | Enriched in P15 dLGN | 3.817 | 0.009 | 1752.000 |
| ENSMUSG00000031142 | calcium channel, voltage-dependent, alpha 1F subunit [Source:MGI Symbol;Acc:MGI:1859639] | Cacna1f | Enriched in P15 dLGN | 3.750 | 0.038 | 357.000 |
| ENSMUSG00000046049 | retinitis pigmentosa 1 homolog like 1 [Source:MGI Symbol;Acc:MGI:2384303] | Rp1l1 | Enriched in P15 dLGN | 3.735 | 0.011 | 1013.000 |
| ENSMUSG00000030785 | cytochrome c oxidase subunit 6A2 [Source:MGI Symbol;Acc:MGI:104649] | Cox6a2 | Enriched in P15 dLGN | 3.712 | 0.000 | 11.740 |
| ENSMUSG00000043418 | leucine-rich repeat, immunoglobulin-like and transmembrane domains 2 [Source:MGI Symbol;Acc:MGI:2444885] | Lrit2 | Enriched in P15 dLGN | 3.655 | 0.045 | 108.600 |
| ENSMUSG00000056055 | S-antigen, retina and pineal gland (arrestin) [Source:MGI Symbol;Acc:MGI:98227] | Sag | Enriched in P15 dLGN | 3.632 | 0.003 | 2015.000 |
| ENSMUSG00000030523 | transient receptor potential cation channel, subfamily M, member 1 [Source:MGI Symbol;Acc:MGI:1330305] | Trpm1 | Enriched in P15 dLGN | 3.588 | 0.020 | 420.900 |
| ENSMUSG00000022820 | NADH:ubiquinone oxidoreductase subunit B4 [Source:MGI Symbol;Acc:MGI:1915444] | Ndufb4 | Enriched in P15 dLGN | 3.497 | 0.003 | 32.870 |
| ENSMUSG00000041534 | retinol binding protein 3, interstitial [Source:MGI Symbol;Acc:MGI:97878] | Rbp3 | Enriched in P15 dLGN | 3.457 | 0.013 | 3079.000 |
| ENSMUSG00000024992 | phosphodiesterase 6C, cGMP specific, cone, alpha prime [Source:MGI Symbol;Acc:MGI:105956] | Pde6c | Enriched in P15 dLGN | 3.441 | 0.029 | 100.500 |
| ENSMUSG00000037161 | mitochondria localized glutamic acid rich protein [Source:MGI Symbol;Acc:MGI:1914999] | Mgarp | Enriched in P15 dLGN | 3.348 | 0.020 | 530.500 |
| ENSMUSG00000020890 | guanylate cyclase 2e [Source:MGI Symbol;Acc:MGI:105123] | Gucy2e | Enriched in P15 dLGN | 3.337 | 0.003 | 507.500 |
| ENSMUSG00000027530 | fatty acid binding protein 12 [Source:MGI Symbol;Acc:MGI:1922747] | Fabp12 | Enriched in P15 dLGN | 3.327 | 0.025 | 148.600 |
| ENSMUSG00000031789 | cyclic nucleotide gated channel beta 1 [Source:MGI Symbol;Acc:MGI:2664102] | Cngb1 | Enriched in P15 dLGN | 3.313 | 0.004 | 733.400 |
| ENSMUSG00000069305 | H4 clustered histone 18 [Source:MGI Symbol;Acc:MGI:4843992] | H4c18 | Enriched in P15 dLGN | 3.296 | 0.040 | 2.587 |
| ENSMUSG00000041578 | cone-rod homeobox [Source:MGI Symbol;Acc:MGI:1194883] | Crx | Enriched in P15 dLGN | 3.280 | 0.022 | 944.100 |
| ENSMUSG00000006007 | phosducin [Source:MGI Symbol;Acc:MGI:98090] | Pdc | Enriched in P15 dLGN | 3.254 | 0.047 | 804.000 |
| ENSMUSG00000028024 | glutamyl aminopeptidase [Source:MGI Symbol;Acc:MGI:106645] | Enpep | Enriched in P15 dLGN | 3.251 | 0.000 | 10.700 |
| ENSMUSG00000075330 | RIKEN cDNA A930003A15 gene [Source:MGI Symbol;Acc:MGI:1915412] | A930003A15Rik | Enriched in P15 dLGN | 3.245 | 0.043 | 40.930 |
| ENSMUSG00000034829 | nucleoredoxin-like 1 [Source:MGI Symbol;Acc:MGI:1924446] | Nxnl1 | Enriched in P15 dLGN | 3.198 | 0.047 | 200.200 |
| ENSMUSG00000028212 | cyclin E2 [Source:MGI Symbol;Acc:MGI:1329034] | Ccne2 | Enriched in P15 dLGN | 3.163 | 0.015 | 12.530 |
| ENSMUSG00000029641 | RAS-like, family 11, member A [Source:MGI Symbol;Acc:MGI:1916145] | Rasl11a | Enriched in P15 dLGN | 3.158 | 0.000 | 21.430 |
| ENSMUSG00000037996 | solute carrier family 24 (sodium/potassium/calcium exchanger), member 2 [Source:MGI Symbol;Acc:MGI:1923626] | Slc24a2 | Enriched in P15 dLGN | 3.091 | 0.000 | 1329.000 |
| ENSMUSG00000023979 | guanylate cyclase activator 1B [Source:MGI Symbol;Acc:MGI:1194489] | Guca1b | Enriched in P15 dLGN | 3.084 | 0.033 | 220.200 |
| ENSMUSG00000039783 | kynurenine 3-monooxygenase (kynurenine 3-hydroxylase) [Source:MGI Symbol;Acc:MGI:2138151] | Kmo | Enriched in P15 dLGN | 3.046 | 0.006 | 4.130 |
| ENSMUSG00000029491 | phosphodiesterase 6B, cGMP, rod receptor, beta polypeptide [Source:MGI Symbol;Acc:MGI:97525] | Pde6b | Enriched in P15 dLGN | 3.029 | 0.036 | 1205.000 |
| ENSMUSG00000040624 | pleckstrin homology domain containing, family G (with RhoGef domain) member 1 [Source:MGI Symbol;Acc:MGI:2676551] | Plekhg1 | Enriched in P15 dLGN | 2.998 | 0.000 | 465.000 |
| ENSMUSG00000018470 | potassium voltage-gated channel, shaker-related subfamily, beta member 3 [Source:MGI Symbol;Acc:MGI:1336208] | Kcnab3 | Enriched in P15 dLGN | 2.981 | 0.000 | 53.860 |
| ENSMUSG00000003949 | hepatic leukemia factor [Source:MGI Symbol;Acc:MGI:96108] | Hlf | Enriched in P15 dLGN | 2.956 | 0.000 | 354.900 |
| ENSMUSG00000022468 | endonuclease, polyU-specific [Source:MGI Symbol;Acc:MGI:97746] | Endou | Enriched in P15 dLGN | 2.954 | 0.000 | 34.930 |

| | | | | | | |
|---|---|---|---|---|---|---|
| ENSMUSG00000031303 | mitogen-activated protein kinase kinase kinase 15 [Source:MGI Symbol;Acc:MGI:2448588] | Map3k15 | Enriched in P15 dLGN | 2.946 | 0.025 | 11.320 |
| ENSMUSG00000029211 | gamma-aminobutyric acid (GABA) A receptor, subunit alpha 4 [Source:MGI Symbol;Acc:MGI:95616] | Gabra4 | Enriched in P15 dLGN | 2.945 | 0.000 | 154.600 |
| ENSMUSG00000021363 | male germ cell-associated kinase [Source:MGI Symbol;Acc:MGI:96913] | Mak | Enriched in P15 dLGN | 2.925 | 0.028 | 259.300 |
| ENSMUSG00000078963 | heat shock factor binding protein 1-like 1 [Source:MGI Symbol;Acc:MGI:1913505] | Hsbp1l1 | Enriched in P15 dLGN | 2.915 | 0.008 | 35.030 |
| ENSMUSG00000031231 | cytochrome c oxidase subunit 7B [Source:MGI Symbol;Acc:MGI:1913392] | Cox7b | Enriched in P15 dLGN | 2.905 | 0.000 | 113.900 |
| ENSMUSG00000026824 | potassium inwardly-rectifying channel, subfamily J, member 3 [Source:MGI Symbol;Acc:MGI:104742] | Kcnj3 | Enriched in P15 dLGN | 2.892 | 0.000 | 206.800 |
| ENSMUSG00000056947 | mab-21-like 1 [Source:MGI Symbol;Acc:MGI:1333773] | Mab21l1 | Enriched in P15 dLGN | 2.875 | 0.030 | 403.600 |
| ENSMUSG00000067220 | cyclic nucleotide gated channel alpha 1 [Source:MGI Symbol;Acc:MGI:88436] | Cnga1 | Enriched in P15 dLGN | 2.869 | 0.005 | 478.200 |
| ENSMUSG00000031438 | ring finger protein 128 [Source:MGI Symbol;Acc:MGI:1914139] | Rnf128 | Enriched in P15 dLGN | 2.865 | 0.001 | 18.970 |
| ENSMUSG00000027610 | glutathione synthetase [Source:MGI Symbol;Acc:MGI:95852] | Gss | Enriched in P15 dLGN | 2.860 | 0.000 | 48.230 |
| ENSMUSG00000045625 | phosphatidylinositol glycan anchor biosynthesis, class Z [Source:MGI Symbol;Acc:MGI:2443822] | Pigz | Enriched in P15 dLGN | 2.845 | 0.000 | 18.260 |
| ENSMUSG00000059343 | aldolase 1 A, retrogene 1 [Source:MGI Symbol;Acc:MGI:2447811] | Aldoart1 | Enriched in P15 dLGN | 2.822 | 0.000 | 19.450 |
| ENSMUSG00000027712 | annexin A5 [Source:MGI Symbol;Acc:MGI:106008] | Anxa5 | Enriched in P15 dLGN | 2.816 | 0.000 | 102.300 |
| ENSMUSG00000021520 | ubiquinol-cytochrome c reductase binding protein [Source:MGI Symbol;Acc:MGI:1914780] | Uqcrb | Enriched in P15 dLGN | 2.812 | 0.000 | 49.090 |
| ENSMUSG00000091636 | A kinase (PRKA) anchor inhibitor 1 [Source:MGI Symbol;Acc:MGI:2444600] | Akain1 | Enriched in P15 dLGN | 2.811 | 0.008 | 14.320 |
| ENSMUSG00000022587 | lymphocyte antigen 6 complex, locus E [Source:MGI Symbol;Acc:MGI:106651] | Ly6e | Enriched in P15 dLGN | 2.795 | 0.000 | 95.380 |
| ENSMUSG00000048029 | enolase 4 [Source:MGI Symbol;Acc:MGI:2441717] | Eno4 | Enriched in P15 dLGN | 2.780 | 0.045 | 12.420 |
| ENSMUSG00000054580 | phospholipase A2 receptor 1 [Source:MGI Symbol;Acc:MGI:102468] | Pla2r1 | Enriched in P15 dLGN | 2.740 | 0.025 | 229.200 |
| ENSMUSG00000091722 | siah E3 ubiquitin protein ligase family member 3 [Source:MGI Symbol;Acc:MGI:2685758] | Siah3 | Enriched in P15 dLGN | 2.729 | 0.000 | 62.950 |
| ENSMUSG00000091402 | retinal degeneration 3-like [Source:MGI Symbol;Acc:MGI:2675860] | Rd3l | Enriched in P15 dLGN | 2.724 | 0.043 | 41.890 |
| ENSMUSG00000029632 | Ndufa4, mitochondrial complex associated [Source:MGI Symbol;Acc:MGI:107686] | Ndufa4 | Enriched in P15 dLGN | 2.706 | 0.000 | 183.200 |
| ENSMUSG00000010803 | gamma-aminobutyric acid (GABA) A receptor, subunit alpha 1 [Source:MGI Symbol;Acc:MGI:95613] | Gabra1 | Enriched in P15 dLGN | 2.704 | 0.000 | 231.000 |
| ENSMUSG00000034570 | inositol polyphosphate 5-phosphatase J [Source:MGI Symbol;Acc:MGI:2158663] | Inpp5j | Enriched in P15 dLGN | 2.698 | 0.000 | 112.100 |
| ENSMUSG00000031007 | ATPase, H+ transporting, lysosomal accessory protein 2 [Source:MGI Symbol;Acc:MGI:1917745] | Atp6ap2 | Enriched in P15 dLGN | 2.687 | 0.000 | 242.600 |
| ENSMUSG00000040723 | RCSD domain containing 1 [Source:MGI Symbol;Acc:MGI:2676394] | Rcsd1 | Enriched in P15 dLGN | 2.686 | 0.000 | 27.500 |
| ENSMUSG00000030103 | basic helix-loop-helix family, member e40 [Source:MGI Symbol;Acc:MGI:1097714] | Bhlhe40 | Enriched in P15 dLGN | 2.681 | 0.000 | 95.280 |
| ENSMUSG00000026525 | opsin 3 [Source:MGI Symbol;Acc:MGI:1338022] | Opn3 | Enriched in P15 dLGN | 2.676 | 0.000 | 17.620 |
| ENSMUSG00000055782 | ATP-binding cassette, sub-family D (ALD), member 2 [Source:MGI Symbol;Acc:MGI:1349467] | Abcd2 | Enriched in P15 dLGN | 2.647 | 0.000 | 85.550 |
| ENSMUSG00000020056 | WASH complex subunit 3 [Source:MGI Symbol;Acc:MGI:1914532] | Washc3 | Enriched in P15 dLGN | 2.638 | 0.002 | 20.630 |
| ENSMUSG00000026609 | usherin [Source:MGI Symbol;Acc:MGI:1341292] | Ush2a | Enriched in P15 dLGN | 2.621 | 0.004 | 1475.000 |
| ENSMUSG00000026384 | protein tyrosine phosphatase, non-receptor type 4 [Source:MGI Symbol;Acc:MGI:1099792] | Ptpn4 | Enriched in P15 dLGN | 2.617 | 0.000 | 1013.000 |
| ENSMUSG00000035504 | receptor accessory protein 6 [Source:MGI Symbol;Acc:MGI:1917585] | Reep6 | Enriched in P15 dLGN | 2.612 | 0.002 | 1054.000 |
| ENSMUSG00000018821 | arginine vasopressin-induced 1 [Source:MGI Symbol;Acc:MGI:1916784] | Avpi1 | Enriched in P15 dLGN | 2.594 | 0.000 | 34.890 |
| ENSMUSG00000049612 | oligodendrocyte myelin glycoprotein [Source:MGI Symbol;Acc:MGI:106586] | Omg | Enriched in P15 dLGN | 2.594 | 0.000 | 36.010 |
| ENSMUSG00000021250 | FBJ osteosarcoma oncogene [Source:MGI Symbol;Acc:MGI:95574] | Fos | Enriched in P15 dLGN | 2.590 | 0.000 | 57.190 |
| ENSMUSG00000020733 | solute carrier family 9 (sodium/hydrogen exchanger), member 3 regulator 1 [Source:MGI Symbol;Acc:MGI:1349482] | Slc9a3r1 | Enriched in P15 dLGN | 2.589 | 0.000 | 214.900 |
| ENSMUSG00000059991 | neuronal pentraxin 2 [Source:MGI Symbol;Acc:MGI:1858209] | Nptx2 | Enriched in P15 dLGN | 2.586 | 0.000 | 106.400 |
| ENSMUSG00000023905 | tumor necrosis factor receptor superfamily, member 12a [Source:MGI Symbol;Acc:MGI:1351484] | Tnfrsf12a | Enriched in P15 dLGN | 2.581 | 0.002 | 7.873 |
| ENSMUSG00000042282 | guanylate cyclase 2f [Source:MGI Symbol;Acc:MGI:105119] | Gucy2f | Enriched in P15 dLGN | 2.543 | 0.018 | 221.800 |
| ENSMUSG00000039620 | tRNA methyltransferase 9B [Source:MGI Symbol;Acc:MGI:2442328] | Trmt9b | Enriched in P15 dLGN | 2.532 | 0.001 | 24.860 |
| ENSMUSG00000039629 | striatin interacting protein 2 [Source:MGI Symbol;Acc:MGI:2444363] | Strip2 | Enriched in P15 dLGN | 2.499 | 0.000 | 138.000 |
| ENSMUSG00000064329 | sodium channel, voltage-gated, type I, alpha [Source:MGI Symbol;Acc:MGI:98246] | Scn1a | Enriched in P15 dLGN | 2.498 | 0.000 | 703.300 |
| ENSMUSG00000050541 | adrenergic receptor, alpha 1b [Source:MGI Symbol;Acc:MGI:104774] | Adra1b | Enriched in P15 dLGN | 2.490 | 0.000 | 73.770 |
| ENSMUSG00000038174 | hyccin PI4KA lipid kinase complex subunit 2 [Source:MGI Symbol;Acc:MGI:1098784] | Hycc2 | Enriched in P15 dLGN | 2.482 | 0.000 | 246.000 |
| ENSMUSG00000033615 | complexin 1 [Source:MGI Symbol;Acc:MGI:104727] | Cplx1 | Enriched in P15 dLGN | 2.475 | 0.000 | 2304.000 |
| ENSMUSG00000027273 | synaptosomal-associated protein 25 [Source:MGI Symbol;Acc:MGI:98331] | Snap25 | Enriched in P15 dLGN | 2.453 | 0.000 | 1966.000 |
| ENSMUSG00000031149 | PRA1 domain family 2 [Source:MGI Symbol;Acc:MGI:1859607] | Praf2 | Enriched in P15 dLGN | 2.452 | 0.002 | 22.150 |
| ENSMUSG00000038717 | ATP synthase, H+ transporting, mitochondrial F0 complex, subunit G [Source:MGI Symbol;Acc:MGI:1351597] | Atp5l | Enriched in P15 dLGN | 2.439 | 0.000 | 38.820 |
| ENSMUSG00000086322 | RIKEN cDNA E130218I03 gene [Source:MGI Symbol;Acc:MGI:3528958] | E130218I03Rik | Enriched in P15 dLGN | 2.435 | 0.018 | 276.300 |
| ENSMUSG00000025475 | adhesion G protein-coupled receptor A1 [Source:MGI Symbol;Acc:MGI:1277167] | Adgra1 | Enriched in P15 dLGN | 2.426 | 0.000 | 360.100 |
| ENSMUSG00000022323 | reactive intermediate imine deaminase A homolog [Source:MGI Symbol;Acc:MGI:1095401] | Rida | Enriched in P15 dLGN | 2.411 | 0.001 | 20.620 |
| ENSMUSG00000060402 | carbohydrate sulfotransferase 8 [Source:MGI Symbol;Acc:MGI:1916197] | Chst8 | Enriched in P15 dLGN | 2.411 | 0.000 | 67.640 |
| ENSMUSG00000038967 | pyruvate dehydrogenase kinase, isoenzyme 2 [Source:MGI Symbol;Acc:MGI:1343087] | Pdk2 | Enriched in P15 dLGN | 2.394 | 0.000 | 136.900 |
| ENSMUSG00000064179 | troponin T1, skeletal, slow [Source:MGI Symbol;Acc:MGI:1333868] | Tnnt1 | Enriched in P15 dLGN | 2.384 | 0.000 | 308.700 |
| ENSMUSG00000025040 | FUN14 domain containing 1 [Source:MGI Symbol;Acc:MGI:1919268] | Fundc1 | Enriched in P15 dLGN | 2.359 | 0.000 | 72.240 |
| ENSMUSG00000002032 | transmembrane protein 25 [Source:MGI Symbol;Acc:MGI:1918937] | Tmem25 | Enriched in P15 dLGN | 2.356 | 0.000 | 70.330 |
| ENSMUSG00000032776 | multiple C2 domains, transmembrane 2 [Source:MGI Symbol;Acc:MGI:2685335] | Mctp2 | Enriched in P15 dLGN | 2.354 | 0.000 | 28.350 |
| ENSMUSG00000001763 | tetraspanin 33 [Source:MGI Symbol;Acc:MGI:1919012] | Tspan33 | Enriched in P15 dLGN | 2.350 | 0.000 | 62.530 |
| ENSMUSG00000041231 | ubiquitin-like domain containing CTD phosphatase 1 [Source:MGI Symbol;Acc:MGI:1933105] | Ublcp1 | Enriched in P15 dLGN | 2.304 | 0.000 | 41.530 |
| ENSMUSG00000079508 | apolipoprotein O [Source:MGI Symbol;Acc:MGI:1915566] | Apoo | Enriched in P15 dLGN | 2.303 | 0.005 | 29.790 |
| ENSMUSG00000019828 | glutamate receptor, metabotropic 1 [Source:MGI Symbol;Acc:MGI:1351338] | Grm1 | Enriched in P15 dLGN | 2.302 | 0.000 | 572.700 |
| ENSMUSG00000043629 | RIKEN cDNA 1700019D03 gene [Source:MGI Symbol;Acc:MGI:1914330] | 1700019D03Rik | Enriched in P15 dLGN | 2.301 | 0.000 | 31.110 |
| ENSMUSG00000025092 | heat shock protein 12A [Source:MGI Symbol;Acc:MGI:1920692] | Hspa12a | Enriched in P15 dLGN | 2.291 | 0.000 | 913.000 |

| | | | | | | |
|---|---|---|---|---|---|---|
| ENSMUSG00000079550 | membrane protein, palmitoylated 4 (MAGUK p55 subfamily member 4) [Source:MGI Symbol;Acc:MGI:2386681] | Mpp4 | Enriched in P15 dLGN | 2.289 | 0.040 | 514.200 |
| ENSMUSG00000036934 | RIKEN cDNA 4921524J17 gene [Source:MGI Symbol;Acc:MGI:1913964] | 4921524J17Rik | Enriched in P15 dLGN | 2.287 | 0.000 | 42.540 |
| ENSMUSG00000031176 | dynein light chain Tctex-type 3 [Source:MGI Symbol;Acc:MGI:1914367] | Dynlt3 | Enriched in P15 dLGN | 2.271 | 0.000 | 112.800 |
| ENSMUSG00000027894 | solute carrier family 6 (neurotransmitter transporter), member 17 [Source:MGI Symbol;Acc:MGI:2442535] | Slc6a17 | Enriched in P15 dLGN | 2.252 | 0.000 | 1006.000 |
| ENSMUSG00000040455 | ubiquitin specific petidase 45 [Source:MGI Symbol;Acc:MGI:101850] | Usp45 | Enriched in P15 dLGN | 2.234 | 0.000 | 101.700 |
| ENSMUSG00000016833 | mitochondrial ribosomal protein S18C [Source:MGI Symbol;Acc:MGI:1915985] | Mrps18c | Enriched in P15 dLGN | 2.225 | 0.048 | 8.671 |
| ENSMUSG00000029659 | mesenteric estrogen dependent adipogenesis [Source:MGI Symbol;Acc:MGI:1917967] | Medag | Enriched in P15 dLGN | 2.225 | 0.000 | 25.400 |
| ENSMUSG00000021775 | nuclear receptor subfamily 1, group D, member 2 [Source:MGI Symbol;Acc:MGI:2449205] | Nr1d2 | Enriched in P15 dLGN | 2.221 | 0.000 | 245.400 |
| ENSMUSG00000032908 | sphingosine-1-phosphate phosphatase 2 [Source:MGI Symbol;Acc:MGI:3589109] | Sgpp2 | Enriched in P15 dLGN | 2.217 | 0.000 | 115.400 |
| ENSMUSG00000050315 | synaptopodin 2 [Source:MGI Symbol;Acc:MGI:2153070] | Synpo2 | Enriched in P15 dLGN | 2.216 | 0.000 | 581.400 |
| ENSMUSG00000007653 | gamma-aminobutyric acid (GABA) A receptor, subunit beta 2 [Source:MGI Symbol;Acc:MGI:95620] | Gabrb2 | Enriched in P15 dLGN | 2.205 | 0.000 | 322.600 |
| ENSMUSG00000020115 | TANK-binding kinase 1 [Source:MGI Symbol;Acc:MGI:1929658] | Tbk1 | Enriched in P15 dLGN | 2.189 | 0.000 | 121.800 |
| ENSMUSG00000034402 | potassium voltage-gated channel, subfamily H (eag-related), member 5 [Source:MGI Symbol;Acc:MGI:3584508] | Kcnh5 | Enriched in P15 dLGN | 2.168 | 0.000 | 240.300 |
| ENSMUSG00000017167 | contactin associated protein-like 1 [Source:MGI Symbol;Acc:MGI:1858201] | Cntnap1 | Enriched in P15 dLGN | 2.167 | 0.000 | 622.800 |
| ENSMUSG00000002332 | dehydrogenase/reductase (SDR family) member 1 [Source:MGI Symbol;Acc:MGI:1196314] | Dhrs1 | Enriched in P15 dLGN | 2.163 | 0.000 | 49.920 |
| ENSMUSG00000002797 | gamma-glutamyl cyclotransferase [Source:MGI Symbol;Acc:MGI:95700] | Ggct | Enriched in P15 dLGN | 2.156 | 0.000 | 25.710 |
| ENSMUSG00000034825 | nuclear receptor interacting protein 3 [Source:MGI Symbol;Acc:MGI:1925843] | Nrip3 | Enriched in P15 dLGN | 2.153 | 0.000 | 285.500 |
| ENSMUSG00000049225 | pyruvate dehydrogenase phosphatase catalytic subunit 1 [Source:MGI Symbol;Acc:MGI:2685870] | Pdp1 | Enriched in P15 dLGN | 2.150 | 0.000 | 541.900 |
| ENSMUSG00000022257 | lysosomal-associated protein transmembrane 4B [Source:MGI Symbol;Acc:MGI:1890494] | Laptm4b | Enriched in P15 dLGN | 2.130 | 0.000 | 165.800 |
| ENSMUSG00000008226 | secernin 3 [Source:MGI Symbol;Acc:MGI:1921866] | Scrn3 | Enriched in P15 dLGN | 2.104 | 0.000 | 42.230 |
| ENSMUSG00000034187 | N-ethylmaleimide sensitive fusion protein [Source:MGI Symbol;Acc:MGI:104560] | Nsf | Enriched in P15 dLGN | 2.086 | 0.000 | 1114.000 |
| ENSMUSG00000032182 | Yip1 domain family, member 2 [Source:MGI Symbol;Acc:MGI:1922016] | Yipf2 | Enriched in P15 dLGN | 2.082 | 0.000 | 34.340 |
| ENSMUSG00000002147 | signal transducer and activator of transcription 6 [Source:MGI Symbol;Acc:MGI:103034] | Stat6 | Enriched in P15 dLGN | 2.076 | 0.001 | 22.910 |
| ENSMUSG00000024190 | dual specificity phosphatase 1 [Source:MGI Symbol;Acc:MGI:105120] | Dusp1 | Enriched in P15 dLGN | 2.060 | 0.000 | 44.730 |
| ENSMUSG00000026260 | NADH:ubiquinone oxidoreductase subunit A10 [Source:MGI Symbol;Acc:MGI:1914523] | Ndufa10 | Enriched in P15 dLGN | 2.051 | 0.000 | 263.700 |
| ENSMUSG00000078974 | SEC61, gamma subunit [Source:MGI Symbol;Acc:MGI:1202066] | Sec61g | Enriched in P15 dLGN | 2.051 | 0.037 | 23.350 |
| ENSMUSG00000024072 | Yip1 domain family, member 4 [Source:MGI Symbol;Acc:MGI:1915114] | Yipf4 | Enriched in P15 dLGN | 2.039 | 0.000 | 59.480 |
| ENSMUSG00000039943 | phospholipase C, beta 4 [Source:MGI Symbol;Acc:MGI:107464] | Plcb4 | Enriched in P15 dLGN | 2.036 | 0.000 | 926.400 |
| ENSMUSG00000025757 | heat shock protein 4 like [Source:MGI Symbol;Acc:MGI:107422] | Hspa4l | Enriched in P15 dLGN | 2.016 | 0.000 | 545.200 |
| ENSMUSG00000003541 | immediate early response 3 [Source:MGI Symbol;Acc:MGI:104814] | Ier3 | Enriched in P15 dLGN | 2.008 | 0.004 | 21.940 |
| ENSMUSG00000021290 | ATP synthase membrane subunit 6.8PL [Source:MGI Symbol;Acc:MGI:1917507] | Atp5mpl | Enriched in P15 dLGN | 2.006 | 0.000 | 181.400 |
| ENSMUSG00000061859 | PATJ, crumbs cell polarity complex component [Source:MGI Symbol;Acc:MGI:1277960] | Patj | Enriched in P15 dLGN | 2.004 | 0.000 | 609.600 |
| ENSMUSG00000071862 | leucine rich repeat transmembrane neuronal 2 [Source:MGI Symbol;Acc:MGI:2389174] | Lrrtm2 | Enriched in P15 dLGN | 2.002 | 0.000 | 78.720 |
| ENSMUSG00000078816 | protein kinase C, gamma [Source:MGI Symbol;Acc:MGI:97597] | Prkcg | Enriched in P15 dLGN | 2.002 | 0.000 | 571.800 |
| ENSMUSG00000036745 | tubulin tyrosine ligase-like family, member 7 [Source:MGI Symbol;Acc:MGI:1918142] | Ttll7 | Enriched in P15 dLGN | 2.000 | 0.000 | 648.100 |
| ENSMUSG00000061474 | mitochondrial ribosomal protein S36 [Source:MGI Symbol;Acc:MGI:1913378] | Mrps36 | Enriched in P15 dLGN | 1.996 | 0.004 | 23.770 |
| ENSMUSG00000001986 | glutamate receptor, ionotropic, AMPA3 (alpha 3) [Source:MGI Symbol;Acc:MGI:95810] | Gria3 | Enriched in P15 dLGN | 1.992 | 0.000 | 283.800 |
| ENSMUSG00000027698 | neutral cholesterol ester hydrolase 1 [Source:MGI Symbol;Acc:MGI:2443191] | Nceh1 | Enriched in P15 dLGN | 1.985 | 0.000 | 190.100 |
| ENSMUSG00000022564 | glutamate receptor, ionotropic, N-methyl D-aspartate-associated protein 1 (glutamate binding) [Source:MGI Symbol;Acc:MGI:1913418] | Grina | Enriched in P15 dLGN | 1.966 | 0.000 | 437.400 |
| ENSMUSG00000032279 | isocitrate dehydrogenase 3 (NAD+) alpha [Source:MGI Symbol;Acc:MGI:1915084] | Idh3a | Enriched in P15 dLGN | 1.951 | 0.000 | 302.000 |
| ENSMUSG00000021748 | pyruvate dehydrogenase (lipoamide) beta [Source:MGI Symbol;Acc:MGI:1915513] | Pdhb | Enriched in P15 dLGN | 1.940 | 0.000 | 160.800 |
| ENSMUSG00000028710 | ATP synthase mitochondrial F1 complex assembly factor 1 [Source:MGI Symbol;Acc:MGI:2180560] | Atpaf1 | Enriched in P15 dLGN | 1.936 | 0.000 | 89.620 |
| ENSMUSG00000027963 | exostosin-like glycosyltransferase 2 [Source:MGI Symbol;Acc:MGI:1889574] | Extl2 | Enriched in P15 dLGN | 1.921 | 0.000 | 151.600 |
| ENSMUSG00000004748 | mitochondrial fission process 1 [Source:MGI Symbol;Acc:MGI:1916686] | Mtfp1 | Enriched in P15 dLGN | 1.920 | 0.000 | 62.740 |
| ENSMUSG00000036850 | mitochondrial ribosomal protein L41 [Source:MGI Symbol;Acc:MGI:1333816] | Mrpl41 | Enriched in P15 dLGN | 1.899 | 0.000 | 53.680 |
| ENSMUSG00000043518 | retinoic acid induced 2 [Source:MGI Symbol;Acc:MGI:1344378] | Rai2 | Enriched in P15 dLGN | 1.895 | 0.019 | 24.130 |
| ENSMUSG00000018648 | dual specificity phosphatase 14 [Source:MGI Symbol;Acc:MGI:1927168] | Dusp14 | Enriched in P15 dLGN | 1.892 | 0.002 | 23.930 |
| ENSMUSG00000031617 | transmembrane protein 184C [Source:MGI Symbol;Acc:MGI:2384562] | Tmem184c | Enriched in P15 dLGN | 1.881 | 0.000 | 115.900 |
| ENSMUSG00000028249 | syndecan binding protein [Source:MGI Symbol;Acc:MGI:1337026] | Sdcbp | Enriched in P15 dLGN | 1.879 | 0.000 | 76.000 |
| ENSMUSG00000030695 | aldolase A, fructose-bisphosphate [Source:MGI Symbol;Acc:MGI:87994] | Aldoa | Enriched in P15 dLGN | 1.876 | 0.000 | 1713.000 |
| ENSMUSG00000028992 | nicotinamide nucleotide adenylyltransferase 1 [Source:MGI Symbol;Acc:MGI:1913704] | Nmnat1 | Enriched in P15 dLGN | 1.870 | 0.002 | 20.770 |
| ENSMUSG00000023456 | triosephosphate isomerase 1 [Source:MGI Symbol;Acc:MGI:98797] | Tpi1 | Enriched in P15 dLGN | 1.866 | 0.000 | 483.500 |
| ENSMUSG00000028149 | RAP1, GTP-GDP dissociation stimulator 1 [Source:MGI Symbol;Acc:MGI:2385189] | Rap1gds1 | Enriched in P15 dLGN | 1.861 | 0.000 | 847.300 |
| ENSMUSG00000054162 | sparc/osteonectin, cwcv and kazal-like domains proteoglycan 3 [Source:MGI Symbol;Acc:MGI:1920152] | Spock3 | Enriched in P15 dLGN | 1.855 | 0.000 | 404.400 |
| ENSMUSG00000015672 | mitochondrial ribosomal protein L32 [Source:MGI Symbol;Acc:MGI:2137226] | Mrpl32 | Enriched in P15 dLGN | 1.850 | 0.013 | 27.230 |
| ENSMUSG00000041112 | engulfment and cell motility 1 [Source:MGI Symbol;Acc:MGI:2153044] | Elmo1 | Enriched in P15 dLGN | 1.850 | 0.000 | 598.800 |
| ENSMUSG00000021771 | voltage-dependent anion channel 2 [Source:MGI Symbol;Acc:MGI:106915] | Vdac2 | Enriched in P15 dLGN | 1.848 | 0.000 | 243.100 |
| ENSMUSG00000021719 | regulator of G-protein signalling 7 binding protein [Source:MGI Symbol;Acc:MGI:106334] | Rgs7bp | Enriched in P15 dLGN | 1.846 | 0.000 | 598.800 |
| ENSMUSG00000063882 | ubiquinol-cytochrome c reductase hinge protein [Source:MGI Symbol;Acc:MGI:1913826] | Uqcrh | Enriched in P15 dLGN | 1.842 | 0.000 | 256.500 |
| ENSMUSG00000024878 | COBW domain containing 1 [Source:MGI Symbol;Acc:MGI:2385089] | Cbwd1 | Enriched in P15 dLGN | 1.831 | 0.005 | 28.100 |
| ENSMUSG00000020321 | malate dehydrogenase 1, NAD (soluble) [Source:MGI Symbol;Acc:MGI:97051] | Mdh1 | Enriched in P15 dLGN | 1.820 | 0.000 | 617.500 |
| ENSMUSG00000027428 | retinoblastoma binding protein 9, serine hydrolase [Source:MGI Symbol;Acc:MGI:1347074] | Rbbp9 | Enriched in P15 dLGN | 1.818 | 0.001 | 45.830 |
| ENSMUSG00000091387 | glucosaminyl (N-acetyl) transferase 4, core 2 (beta-1,6-N-acetylglucosaminyltransferase) [Source:MGI Symbol;Acc:MGI:2684919] | Gcnt4 | Enriched in P15 dLGN | 1.808 | 0.006 | 28.710 |

| | | | | | | |
|---|---|---|---|---|---|---|
| ENSMUSG00000025630 | hypoxanthine guanine phosphoribosyl transferase [Source:MGI Symbol;Acc:MGI:96217] | Hprt | Enriched in P15 dLGN | 1.806 | 0.000 | 174.600 |
| ENSMUSG00000030718 | protein phosphatase methylesterase 1 [Source:MGI Symbol;Acc:MGI:1919840] | Ppme1 | Enriched in P15 dLGN | 1.805 | 0.000 | 576.900 |
| ENSMUSG00000017412 | calcium channel, voltage-dependent, beta 4 subunit [Source:MGI Symbol;Acc:MGI:103301] | Cacnb4 | Enriched in P15 dLGN | 1.804 | 0.000 | 413.300 |
| ENSMUSG00000015766 | epidermal growth factor receptor pathway substrate 8 [Source:MGI Symbol;Acc:MGI:104684] | Eps8 | Enriched in P15 dLGN | 1.799 | 0.000 | 220.600 |
| ENSMUSG00000032398 | small nuclear RNA activating complex, polypeptide 5 [Source:MGI Symbol;Acc:MGI:1914282] | Snapc5 | Enriched in P15 dLGN | 1.795 | 0.021 | 21.390 |
| ENSMUSG00000032883 | acyl-CoA synthetase long-chain family member 3 [Source:MGI Symbol;Acc:MGI:1921455] | Acsl3 | Enriched in P15 dLGN | 1.795 | 0.000 | 522.500 |
| ENSMUSG00000028690 | methylmalonic aciduria cblC type, with homocystinuria [Source:MGI Symbol;Acc:MGI:1914346] | Mmachc | Enriched in P15 dLGN | 1.789 | 0.003 | 24.850 |
| ENSMUSG00000031634 | UFM1-specific peptidase 2 [Source:MGI Symbol;Acc:MGI:1913679] | Ufsp2 | Enriched in P15 dLGN | 1.789 | 0.000 | 89.610 |
| ENSMUSG00000051671 | cytochrome c oxidase assembly factor 6 [Source:MGI Symbol;Acc:MGI:1915142] | Coa6 | Enriched in P15 dLGN | 1.789 | 0.019 | 13.320 |
| ENSMUSG00000026825 | dynamin 1 [Source:MGI Symbol;Acc:MGI:107384] | Dnm1 | Enriched in P15 dLGN | 1.783 | 0.000 | 2096.000 |
| ENSMUSG00000056486 | chimerin 1 [Source:MGI Symbol;Acc:MGI:1915674] | Chn1 | Enriched in P15 dLGN | 1.777 | 0.000 | 504.400 |
| ENSMUSG00000062785 | potassium voltage gated channel, Shaw-related subfamily, member 3 [Source:MGI Symbol;Acc:MGI:96669] | Kcnc3 | Enriched in P15 dLGN | 1.777 | 0.000 | 437.600 |
| ENSMUSG00000071866 | peptidylprolyl isomerase A [Source:MGI Symbol;Acc:MGI:97749] | Ppia | Enriched in P15 dLGN | 1.769 | 0.003 | 211.200 |
| ENSMUSG00000046798 | claudin 12 [Source:MGI Symbol;Acc:MGI:1929288] | Cldn12 | Enriched in P15 dLGN | 1.762 | 0.000 | 124.600 |
| ENSMUSG00000021711 | trafficking protein particle complex 13 [Source:MGI Symbol;Acc:MGI:1914225] | Trappc13 | Enriched in P15 dLGN | 1.753 | 0.000 | 71.190 |
| ENSMUSG00000030127 | COP9 signalosome subunit 7A [Source:MGI Symbol;Acc:MGI:1349400] | Cops7a | Enriched in P15 dLGN | 1.750 | 0.000 | 167.000 |
| ENSMUSG00000006717 | acyl-CoA thioesterase 13 [Source:MGI Symbol;Acc:MGI:1914084] | Acot13 | Enriched in P15 dLGN | 1.748 | 0.001 | 30.710 |
| ENSMUSG00000009406 | ELK1, member of ETS oncogene family [Source:MGI Symbol;Acc:MGI:101833] | Elk1 | Enriched in P15 dLGN | 1.748 | 0.000 | 79.920 |
| ENSMUSG00000028251 | thiosulfate sulfurtransferase (rhodanese)-like domain containing 3 [Source:MGI Symbol;Acc:MGI:1924282] | Tstd3 | Enriched in P15 dLGN | 1.745 | 0.000 | 35.110 |
| ENSMUSG00000096916 | zinc finger protein 850 [Source:MGI Symbol;Acc:MGI:3036281] | Zfp850 | Enriched in P15 dLGN | 1.745 | 0.043 | 22.700 |
| ENSMUSG00000029769 | coiled-coil domain containing 136 [Source:MGI Symbol;Acc:MGI:1918128] | Ccdc136 | Enriched in P15 dLGN | 1.744 | 0.000 | 1034.000 |
| ENSMUSG00000021591 | glutaredoxin [Source:MGI Symbol;Acc:MGI:2135625] | Glrx | Enriched in P15 dLGN | 1.741 | 0.000 | 37.820 |
| ENSMUSG00000057229 | distal membrane arm assembly complex 2 [Source:MGI Symbol;Acc:MGI:1913599] | Dmac2 | Enriched in P15 dLGN | 1.738 | 0.000 | 61.200 |
| ENSMUSG00000024158 | hydroxyacyl glutathione hydrolase [Source:MGI Symbol;Acc:MGI:95745] | Hagh | Enriched in P15 dLGN | 1.735 | 0.000 | 60.650 |
| ENSMUSG00000022890 | ATP synthase, H+ transporting, mitochondrial F0 complex, subunit F [Source:MGI Symbol;Acc:MGI:107777] | Atp5j | Enriched in P15 dLGN | 1.729 | 0.000 | 249.200 |
| ENSMUSG00000052738 | succinate-CoA ligase, GDP-forming, alpha subunit [Source:MGI Symbol;Acc:MGI:1927234] | Suclg1 | Enriched in P15 dLGN | 1.728 | 0.000 | 138.600 |
| ENSMUSG00000043162 | Pigy upstream reading frame [Source:MGI Symbol;Acc:MGI:1913709] | Pyurf | Enriched in P15 dLGN | 1.726 | 0.003 | 52.190 |
| ENSMUSG00000027076 | translocase of inner mitochondrial membrane 10 [Source:MGI Symbol;Acc:MGI:1353429] | Timm10 | Enriched in P15 dLGN | 1.725 | 0.000 | 29.720 |
| ENSMUSG00000025189 | cyclin M1 [Source:MGI Symbol;Acc:MGI:1891366] | Cnnm1 | Enriched in P15 dLGN | 1.716 | 0.000 | 258.600 |
| ENSMUSG00000027556 | carbonic anhydrase 1 [Source:MGI Symbol;Acc:MGI:88268] | Car1 | Enriched in P15 dLGN | 1.714 | 0.020 | 12.270 |
| ENSMUSG00000059409 | protein phosphatase 2, regulatory subunit B', delta [Source:MGI Symbol;Acc:MGI:2388481] | Ppp2r5d | Enriched in P15 dLGN | 1.714 | 0.000 | 362.300 |
| ENSMUSG00000026568 | mitochondrial pyruvate carrier 2 [Source:MGI Symbol;Acc:MGI:1917706] | Mpc2 | Enriched in P15 dLGN | 1.706 | 0.000 | 84.480 |
| ENSMUSG00000000563 | ATP synthase peripheral stalk-membrane subunit b [Source:MGI Symbol;Acc:MGI:1100495] | Atp5pb | Enriched in P15 dLGN | 1.705 | 0.000 | 236.000 |
| ENSMUSG00000002808 | ependymin related protein 1 (zebrafish) [Source:MGI Symbol;Acc:MGI:2145369] | Epdr1 | Enriched in P15 dLGN | 1.703 | 0.000 | 73.680 |
| ENSMUSG00000044676 | zinc finger protein 612 [Source:MGI Symbol;Acc:MGI:2443465] | Zfp612 | Enriched in P15 dLGN | 1.703 | 0.000 | 153.100 |
| ENSMUSG00000004285 | ATPase, H+ transporting, lysosomal V1 subunit F [Source:MGI Symbol;Acc:MGI:1913394] | Atp6v1f | Enriched in P15 dLGN | 1.690 | 0.000 | 78.370 |
| ENSMUSG00000063172 | heat shock protein family B (small), member 11 [Source:MGI Symbol;Acc:MGI:1920188] | Hspb11 | Enriched in P15 dLGN | 1.687 | 0.024 | 24.210 |
| ENSMUSG00000027531 | inositol (myo)-1(or 4)-monophosphatase 1 [Source:MGI Symbol;Acc:MGI:1933158] | Impa1 | Enriched in P15 dLGN | 1.686 | 0.000 | 75.710 |
| ENSMUSG00000032459 | mitochondrial ribosomal protein S22 [Source:MGI Symbol;Acc:MGI:1928137] | Mrps22 | Enriched in P15 dLGN | 1.675 | 0.011 | 35.070 |
| ENSMUSG00000019877 | serine incorporator 1 [Source:MGI Symbol;Acc:MGI:1926228] | Serinc1 | Enriched in P15 dLGN | 1.674 | 0.000 | 662.300 |
| ENSMUSG00000057666 | glyceraldehyde-3-phosphate dehydrogenase [Source:MGI Symbol;Acc:MGI:95640] | Gapdh | Enriched in P15 dLGN | 1.673 | 0.003 | 93.820 |
| ENSMUSG00000034488 | EGF-like repeats and discoidin I-like domains 3 [Source:MGI Symbol;Acc:MGI:1329025] | Edil3 | Enriched in P15 dLGN | 1.672 | 0.000 | 564.800 |
| ENSMUSG00000036186 | divergent protein kinase domain 1B [Source:MGI Symbol;Acc:MGI:1927576] | Dipk1b | Enriched in P15 dLGN | 1.672 | 0.000 | 28.180 |
| ENSMUSG00000005362 | cereblon [Source:MGI Symbol;Acc:MGI:1913277] | Crbn | Enriched in P15 dLGN | 1.670 | 0.000 | 104.300 |
| ENSMUSG00000029432 | nipsnap homolog 2 [Source:MGI Symbol;Acc:MGI:1278343] | Nipsnap2 | Enriched in P15 dLGN | 1.666 | 0.000 | 127.200 |
| ENSMUSG00000009207 | lunapark, ER junction formation factor [Source:MGI Symbol;Acc:MGI:1918115] | Lnpk | Enriched in P15 dLGN | 1.664 | 0.000 | 102.000 |
| ENSMUSG00000020629 | acireductone dioxygenase 1 [Source:MGI Symbol;Acc:MGI:2144929] | Adi1 | Enriched in P15 dLGN | 1.664 | 0.000 | 28.330 |
| ENSMUSG00000053930 | shisa family member 6 [Source:MGI Symbol;Acc:MGI:2685725] | Shisa6 | Enriched in P15 dLGN | 1.662 | 0.000 | 186.600 |
| ENSMUSG00000025240 | SAC1 suppressor of actin mutations 1-like (yeast) [Source:MGI Symbol;Acc:MGI:1933169] | Sacm1l | Enriched in P15 dLGN | 1.660 | 0.000 | 163.500 |
| ENSMUSG00000075256 | ceramide kinase-like [Source:MGI Symbol;Acc:MGI:3037816] | Cerkl | Enriched in P15 dLGN | 1.660 | 0.042 | 144.800 |
| ENSMUSG00000008822 | acylphosphatase 1, erythrocyte (common) type [Source:MGI Symbol;Acc:MGI:1913454] | Acyp1 | Enriched in P15 dLGN | 1.656 | 0.008 | 16.490 |
| ENSMUSG00000071648 | rod outer segment membrane protein 1 [Source:MGI Symbol;Acc:MGI:97998] | Rom1 | Enriched in P15 dLGN | 1.655 | 0.044 | 555.400 |
| ENSMUSG00000028648 | NADH:ubiquinone oxidoreductase core subunit S5 [Source:MGI Symbol;Acc:MGI:1890889] | Ndufs5 | Enriched in P15 dLGN | 1.653 | 0.000 | 101.900 |
| ENSMUSG00000048100 | TATA-box binding protein associated factor 13 [Source:MGI Symbol;Acc:MGI:1913500] | Taf13 | Enriched in P15 dLGN | 1.652 | 0.000 | 38.390 |
| ENSMUSG00000032336 | neuroplastin [Source:MGI Symbol;Acc:MGI:108077] | Nptn | Enriched in P15 dLGN | 1.645 | 0.000 | 662.100 |
| ENSMUSG00000039601 | regulator of calcineurin 2 [Source:MGI Symbol;Acc:MGI:1858219] | Rcan2 | Enriched in P15 dLGN | 1.640 | 0.000 | 311.500 |
| ENSMUSG00000031202 | RAB39B, member RAS oncogene family [Source:MGI Symbol;Acc:MGI:1915040] | Rab39b | Enriched in P15 dLGN | 1.632 | 0.000 | 57.430 |
| ENSMUSG00000078941 | adenylate kinase 6 [Source:MGI Symbol;Acc:MGI:5510732] | Ak6 | Enriched in P15 dLGN | 1.628 | 0.008 | 18.090 |
| ENSMUSG00000022295 | ATPase, H+ transporting, lysosomal V1 subunit C1 [Source:MGI Symbol;Acc:MGI:1913585] | Atp6v1c1 | Enriched in P15 dLGN | 1.627 | 0.000 | 229.000 |
| ENSMUSG00000034566 | ATP synthase, H+ transporting, mitochondrial F0 complex, subunit D [Source:MGI Symbol;Acc:MGI:1918929] | Atp5h | Enriched in P15 dLGN | 1.620 | 0.000 | 216.100 |
| ENSMUSG00000016319 | solute carrier family 25 (mitochondrial carrier, adenine nucleotide translocator), member 5 [Source:MGI Symbol;Acc:MGI:1353496] | Slc25a5 | Enriched in P15 dLGN | 1.613 | 0.000 | 198.800 |
| ENSMUSG00000083282 | cathepsin F [Source:MGI Symbol;Acc:MGI:1861434] | Ctsf | Enriched in P15 dLGN | 1.606 | 0.000 | 193.600 |
| ENSMUSG00000031996 | amyloid beta (A4) precursor-like protein 2 [Source:MGI Symbol;Acc:MGI:88047] | Aplp2 | Enriched in P15 dLGN | 1.595 | 0.000 | 2704.000 |

| | | | | | | |
|---|---|---|---|---|---|---|
| ENSMUSG00000032966 | FK506 binding protein 1a [Source:MGI Symbol;Acc:MGI:95541] | Fkbp1a | Enriched in P15 dLGN | 1.591 | 0.000 | 430.500 |
| ENSMUSG00000035790 | centrosomal protein 19 [Source:MGI Symbol;Acc:MGI:1914244] | Cep19 | Enriched in P15 dLGN | 1.584 | 0.000 | 80.500 |
| ENSMUSG00000091264 | small integral membrane protein 13 [Source:MGI Symbol;Acc:MGI:2652854] | Smim13 | Enriched in P15 dLGN | 1.584 | 0.000 | 346.000 |
| ENSMUSG00000041444 | Rho GTPase activating protein 32 [Source:MGI Symbol;Acc:MGI:2450166] | Arhgap32 | Enriched in P15 dLGN | 1.582 | 0.000 | 970.500 |
| ENSMUSG00000048376 | coagulation factor II (thrombin) receptor [Source:MGI Symbol;Acc:MGI:101802] | F2r | Enriched in P15 dLGN | 1.582 | 0.005 | 37.830 |
| ENSMUSG00000031429 | proteasome (prosome, macropain) 26S subunit, non-ATPase, 10 [Source:MGI Symbol;Acc:MGI:1858898] | Psmd10 | Enriched in P15 dLGN | 1.580 | 0.016 | 17.050 |
| ENSMUSG00000023033 | sodium channel, voltage-gated, type VIII, alpha [Source:MGI Symbol;Acc:MGI:103169] | Scn8a | Enriched in P15 dLGN | 1.575 | 0.000 | 1192.000 |
| ENSMUSG00000079317 | trafficking protein particle complex 2 [Source:MGI Symbol;Acc:MGI:1913476] | Trappc2 | Enriched in P15 dLGN | 1.571 | 0.019 | 15.830 |
| ENSMUSG00000040373 | calcium channel, voltage-dependent, gamma subunit 5 [Source:MGI Symbol;Acc:MGI:2157946] | Cacng5 | Enriched in P15 dLGN | 1.567 | 0.000 | 203.800 |
| ENSMUSG00000024966 | stress-induced phosphoprotein 1 [Source:MGI Symbol;Acc:MGI:109130] | Stip1 | Enriched in P15 dLGN | 1.564 | 0.000 | 255.000 |
| ENSMUSG00000027742 | component of oligomeric golgi complex 6 [Source:MGI Symbol;Acc:MGI:1914792] | Cog6 | Enriched in P15 dLGN | 1.564 | 0.000 | 89.910 |
| ENSMUSG00000043831 | LysM, putative peptidoglycan-binding, domain containing 4 [Source:MGI Symbol;Acc:MGI:1922349] | Lysmd4 | Enriched in P15 dLGN | 1.564 | 0.000 | 79.010 |
| ENSMUSG00000078566 | BCL2/adenovirus E1B interacting protein 3 [Source:MGI Symbol;Acc:MGI:109326] | Bnip3 | Enriched in P15 dLGN | 1.563 | 0.001 | 111.700 |
| ENSMUSG00000023175 | basigin [Source:MGI Symbol;Acc:MGI:88208] | Bsg | Enriched in P15 dLGN | 1.562 | 0.000 | 872.500 |
| ENSMUSG00000002010 | isocitrate dehydrogenase 3 (NAD+), gamma [Source:MGI Symbol;Acc:MGI:1099463] | Idh3g | Enriched in P15 dLGN | 1.561 | 0.002 | 76.060 |
| ENSMUSG00000022664 | solute carrier family 35, member A5 [Source:MGI Symbol;Acc:MGI:1921352] | Slc35a5 | Enriched in P15 dLGN | 1.556 | 0.001 | 55.760 |
| ENSMUSG00000029066 | mitochondrial ribosomal protein L20 [Source:MGI Symbol;Acc:MGI:2137221] | Mrpl20 | Enriched in P15 dLGN | 1.555 | 0.000 | 48.190 |
| ENSMUSG00000071180 | small integral membrane protein 15 [Source:MGI Symbol;Acc:MGI:1922866] | Smim15 | Enriched in P15 dLGN | 1.551 | 0.001 | 41.580 |
| ENSMUSG00000024887 | N-acylsphingosine amidohydrolase 2 [Source:MGI Symbol;Acc:MGI:1859310] | Asah2 | Enriched in P15 dLGN | 1.549 | 0.008 | 39.790 |
| ENSMUSG00000036503 | ring finger protein 13 [Source:MGI Symbol;Acc:MGI:1346341] | Rnf13 | Enriched in P15 dLGN | 1.546 | 0.000 | 102.400 |
| ENSMUSG00000028247 | coenzyme Q3 methyltransferase [Source:MGI Symbol;Acc:MGI:101813] | Coq3 | Enriched in P15 dLGN | 1.544 | 0.002 | 34.440 |
| ENSMUSG00000020340 | cytoplasmic FMR1 interacting protein 2 [Source:MGI Symbol;Acc:MGI:1924134] | Cyfip2 | Enriched in P15 dLGN | 1.542 | 0.000 | 1444.000 |
| ENSMUSG00000031647 | microfibrillar-associated protein 3-like [Source:MGI Symbol;Acc:MGI:1918556] | Mfap3l | Enriched in P15 dLGN | 1.541 | 0.000 | 250.900 |
| ENSMUSG00000020949 | FK506 binding protein 3 [Source:MGI Symbol;Acc:MGI:1353460] | Fkbp3 | Enriched in P15 dLGN | 1.539 | 0.000 | 160.500 |
| ENSMUSG00000062931 | zinc finger protein 938 [Source:MGI Symbol;Acc:MGI:3621440] | Zfp938 | Enriched in P15 dLGN | 1.539 | 0.002 | 45.710 |
| ENSMUSG00000041460 | calcium channel, voltage-dependent, alpha 2/delta subunit 4 [Source:MGI Symbol;Acc:MGI:2442632] | Cacna2d4 | Enriched in P15 dLGN | 1.536 | 0.000 | 641.500 |
| ENSMUSG00000059040 | enolase 1B, retrotransposed [Source:MGI Symbol;Acc:MGI:3648653] | Eno1b | Enriched in P15 dLGN | 1.532 | 0.006 | 124.100 |
| ENSMUSG00000030869 | NADH:ubiquinone oxidoreductase subunit AB1 [Source:MGI Symbol;Acc:MGI:1917566] | Ndufab1 | Enriched in P15 dLGN | 1.531 | 0.000 | 76.720 |
| ENSMUSG00000031059 | NADH:ubiquinone oxidoreductase subunit B11 [Source:MGI Symbol;Acc:MGI:1349919] | Ndufb11 | Enriched in P15 dLGN | 1.524 | 0.000 | 155.400 |
| ENSMUSG00000022658 | transgelin 3 [Source:MGI Symbol;Acc:MGI:1926784] | Tagln3 | Enriched in P15 dLGN | 1.522 | 0.000 | 267.200 |
| ENSMUSG00000042675 | yippee like 3 [Source:MGI Symbol;Acc:MGI:1913340] | Ypel3 | Enriched in P15 dLGN | 1.522 | 0.000 | 169.400 |
| ENSMUSG00000060429 | syntrophin, basic 1 [Source:MGI Symbol;Acc:MGI:101781] | Sntb1 | Enriched in P15 dLGN | 1.521 | 0.000 | 115.500 |
| ENSMUSG00000022024 | SGT1, suppressor of G2 allele of SKP1 (S. cerevisiae) [Source:MGI Symbol;Acc:MGI:1915205] | Sugt1 | Enriched in P15 dLGN | 1.519 | 0.000 | 199.600 |
| ENSMUSG00000040321 | zinc finger protein 770 [Source:MGI Symbol;Acc:MGI:2445100] | Zfp770 | Enriched in P15 dLGN | 1.519 | 0.001 | 64.980 |
| ENSMUSG00000029153 | OCIA domain containing 2 [Source:MGI Symbol;Acc:MGI:1916377] | Ociad2 | Enriched in P15 dLGN | 1.515 | 0.000 | 70.580 |
| ENSMUSG00000024875 | Yip1 interacting factor homolog A (S. cerevisiae) [Source:MGI Symbol;Acc:MGI:1915340] | Yif1a | Enriched in P15 dLGN | 1.511 | 0.017 | 19.910 |
| ENSMUSG00000045410 | aldo-keto reductase family 1, member E1 [Source:MGI Symbol;Acc:MGI:1914758] | Akr1e1 | Enriched in P15 dLGN | 1.509 | 0.001 | 70.940 |
| ENSMUSG00000022620 | arylsulfatase A [Source:MGI Symbol;Acc:MGI:88077] | Arsa | Enriched in P15 dLGN | 1.508 | 0.000 | 62.330 |
| ENSMUSG00000037606 | oxysterol binding protein-like 5 [Source:MGI Symbol;Acc:MGI:1930265] | Osbpl5 | Enriched in P15 dLGN | 1.508 | 0.000 | 213.900 |
| ENSMUSG00000027706 | SEC62 homolog (S. cerevisiae) [Source:MGI Symbol;Acc:MGI:1916526] | Sec62 | Enriched in P15 dLGN | 1.503 | 0.000 | 458.300 |
| ENSMUSG00000029657 | heat shock 105kDa/110kDa protein 1 [Source:MGI Symbol;Acc:MGI:105053] | Hsph1 | Enriched in P15 dLGN | 1.498 | 0.000 | 555.200 |
| ENSMUSG00000022094 | solute carrier family 39 (zinc transporter), member 14 [Source:MGI Symbol;Acc:MGI:2384851] | Slc39a14 | Enriched in P15 dLGN | 1.493 | 0.000 | 73.630 |
| ENSMUSG00000042032 | methionine adenosyltransferase II, beta [Source:MGI Symbol;Acc:MGI:1913667] | Mat2b | Enriched in P15 dLGN | 1.493 | 0.000 | 58.430 |
| ENSMUSG00000042682 | selenoprotein K [Source:MGI Symbol;Acc:MGI:1931466] | Selenok | Enriched in P15 dLGN | 1.493 | 0.000 | 254.300 |
| ENSMUSG00000025781 | ATP synthase, H+ transporting, mitochondrial F1 complex, gamma polypeptide 1 [Source:MGI Symbol;Acc:MGI:1261437] | Atp5c1 | Enriched in P15 dLGN | 1.490 | 0.000 | 290.000 |
| ENSMUSG00000020514 | mitochondrial ribosomal protein L22 [Source:MGI Symbol;Acc:MGI:1333794] | Mrpl22 | Enriched in P15 dLGN | 1.486 | 0.009 | 23.400 |
| ENSMUSG00000036158 | prickle planar cell polarity protein 1 [Source:MGI Symbol;Acc:MGI:1916034] | Prickle1 | Enriched in P15 dLGN | 1.483 | 0.000 | 385.100 |
| ENSMUSG00000026766 | methylmalonic aciduria (cobalamin deficiency) cblD type, with homocystinuria [Source:MGI Symbol;Acc:MGI:1923786] | Mmadhc | Enriched in P15 dLGN | 1.482 | 0.000 | 58.080 |
| ENSMUSG00000002015 | B cell receptor associated protein 31 [Source:MGI Symbol;Acc:MGI:1350933] | Bcap31 | Enriched in P15 dLGN | 1.479 | 0.000 | 84.180 |
| ENSMUSG00000002475 | abhydrolase domain containing 3 [Source:MGI Symbol;Acc:MGI:2147183] | Abhd3 | Enriched in P15 dLGN | 1.477 | 0.005 | 45.040 |
| ENSMUSG00000030612 | mitochondrial ribosomal protein L46 [Source:MGI Symbol;Acc:MGI:1914558] | Mrpl46 | Enriched in P15 dLGN | 1.476 | 0.001 | 33.730 |
| ENSMUSG00000001794 | calpain, small subunit 1 [Source:MGI Symbol;Acc:MGI:88266] | Capns1 | Enriched in P15 dLGN | 1.475 | 0.000 | 150.200 |
| ENSMUSG00000019916 | procollagen-proline, 2-oxoglutarate 4-dioxygenase (proline 4-hydroxylase), alpha 1 polypeptide [Source:MGI Symbol;Acc:MGI:97463] | P4ha1 | Enriched in P15 dLGN | 1.474 | 0.000 | 152.900 |
| ENSMUSG00000082229 | nucleosome assembly protein 1-like 2 [Source:MGI Symbol;Acc:MGI:106654] | Nap1l2 | Enriched in P15 dLGN | 1.472 | 0.000 | 277.100 |
| ENSMUSG00000014077 | calcineurin-like EF hand protein 1 [Source:MGI Symbol;Acc:MGI:1927185] | Chp1 | Enriched in P15 dLGN | 1.471 | 0.000 | 152.700 |
| ENSMUSG00000027495 | family with sequence similarity 210, member B [Source:MGI Symbol;Acc:MGI:1914267] | Fam210b | Enriched in P15 dLGN | 1.468 | 0.000 | 55.780 |
| ENSMUSG00000030032 | WD repeat domain 54 [Source:MGI Symbol;Acc:MGI:1922909] | Wdr54 | Enriched in P15 dLGN | 1.467 | 0.041 | 15.680 |
| ENSMUSG00000040612 | immunoglobulin-like domain containing receptor 2 [Source:MGI Symbol;Acc:MGI:1196370] | Ildr2 | Enriched in P15 dLGN | 1.465 | 0.000 | 1307.000 |
| ENSMUSG00000020333 | acyl-CoA synthetase long-chain family member 6 [Source:MGI Symbol;Acc:MGI:894291] | Acsl6 | Enriched in P15 dLGN | 1.464 | 0.000 | 594.100 |
| ENSMUSG00000021987 | myotubularin related protein 6 [Source:MGI Symbol;Acc:MGI:2145637] | Mtmr6 | Enriched in P15 dLGN | 1.464 | 0.000 | 223.600 |
| ENSMUSG00000027965 | olfactomedin 3 [Source:MGI Symbol;Acc:MGI:2387329] | Olfm3 | Enriched in P15 dLGN | 1.464 | 0.000 | 175.900 |
| ENSMUSG00000024414 | mitochondrial ribosomal protein L27 [Source:MGI Symbol;Acc:MGI:2137224] | Mrpl27 | Enriched in P15 dLGN | 1.463 | 0.006 | 26.690 |
| ENSMUSG00000044221 | G-rich RNA sequence binding factor 1 [Source:MGI Symbol;Acc:MGI:106479] | Grsf1 | Enriched in P15 dLGN | 1.463 | 0.000 | 242.300 |

| | | | | | | |
|---|---|---|---|---|---|---|
| ENSMUSG00000034640 | TCDD-inducible poly(ADP-ribose) polymerase [Source:MGI Symbol;Acc:MGI:2159210] | Tiparp | Enriched in P15 dLGN | 1.461 | 0.003 | 38.050 |
| ENSMUSG00000073676 | heat shock protein 1 (chaperonin 10) [Source:MGI Symbol;Acc:MGI:104680] | Hspe1 | Enriched in P15 dLGN | 1.461 | 0.001 | 62.310 |
| ENSMUSG00000059278 | N(alpha)-acetyltransferase 38, NatC auxiliary subunit [Source:MGI Symbol;Acc:MGI:1925554] | Naa38 | Enriched in P15 dLGN | 1.459 | 0.005 | 31.830 |
| ENSMUSG00000036932 | apoptosis-inducing factor, mitochondrion-associated 1 [Source:MGI Symbol;Acc:MGI:1349419] | Aifm1 | Enriched in P15 dLGN | 1.458 | 0.000 | 96.300 |
| ENSMUSG00000062981 | mitochondrial ribosomal protein L42 [Source:MGI Symbol;Acc:MGI:1333774] | Mrpl42 | Enriched in P15 dLGN | 1.458 | 0.014 | 43.370 |
| ENSMUSG00000015668 | PDZ domain containing 11 [Source:MGI Symbol;Acc:MGI:1919871] | Pdzd11 | Enriched in P15 dLGN | 1.456 | 0.003 | 27.950 |
| ENSMUSG00000021939 | cathepsin B [Source:MGI Symbol;Acc:MGI:88561] | Ctsb | Enriched in P15 dLGN | 1.456 | 0.000 | 510.600 |
| ENSMUSG00000022110 | succinate-Coenzyme A ligase, ADP-forming, beta subunit [Source:MGI Symbol;Acc:MGI:1306775] | Sucla2 | Enriched in P15 dLGN | 1.456 | 0.000 | 256.600 |
| ENSMUSG00000025059 | glycerol kinase [Source:MGI Symbol;Acc:MGI:106594] | Gk | Enriched in P15 dLGN | 1.455 | 0.002 | 44.020 |
| ENSMUSG00000028419 | charged multivesicular body protein 5 [Source:MGI Symbol;Acc:MGI:1924209] | Chmp5 | Enriched in P15 dLGN | 1.455 | 0.000 | 102.800 |
| ENSMUSG00000037361 | splicing factor 3B, subunit 6 [Source:MGI Symbol;Acc:MGI:1913305] | Sf3b6 | Enriched in P15 dLGN | 1.455 | 0.009 | 35.230 |
| ENSMUSG00000000326 | catechol-O-methyltransferase [Source:MGI Symbol;Acc:MGI:88470] | Comt | Enriched in P15 dLGN | 1.451 | 0.000 | 103.300 |
| ENSMUSG00000021215 | neuroepithelial cell transforming gene 1 [Source:MGI Symbol;Acc:MGI:1927138] | Net1 | Enriched in P15 dLGN | 1.451 | 0.000 | 50.150 |
| ENSMUSG00000027827 | potassium voltage-gated channel, shaker-related subfamily, beta member 1 [Source:MGI Symbol;Acc:MGI:109155] | Kcnab1 | Enriched in P15 dLGN | 1.450 | 0.000 | 138.000 |
| ENSMUSG00000040048 | NADH:ubiquinone oxidoreductase subunit B10 [Source:MGI Symbol;Acc:MGI:1915592] | Ndufb10 | Enriched in P15 dLGN | 1.445 | 0.000 | 201.200 |
| ENSMUSG00000023723 | mitochondrial ribosomal protein S23 [Source:MGI Symbol;Acc:MGI:1928138] | Mrps23 | Enriched in P15 dLGN | 1.443 | 0.001 | 39.530 |
| ENSMUSG00000031666 | RB transcriptional corepressor like 2 [Source:MGI Symbol;Acc:MGI:105085] | Rbl2 | Enriched in P15 dLGN | 1.443 | 0.000 | 128.400 |
| ENSMUSG00000004207 | prosaposin [Source:MGI Symbol;Acc:MGI:97783] | Psap | Enriched in P15 dLGN | 1.440 | 0.000 | 1320.000 |
| ENSMUSG00000060073 | proteasome subunit alpha 3 [Source:MGI Symbol;Acc:MGI:104883] | Psma3 | Enriched in P15 dLGN | 1.440 | 0.000 | 145.000 |
| ENSMUSG00000032324 | tetraspanin 3 [Source:MGI Symbol;Acc:MGI:1928098] | Tspan3 | Enriched in P15 dLGN | 1.436 | 0.000 | 417.400 |
| ENSMUSG00000000552 | zinc finger protein 385A [Source:MGI Symbol;Acc:MGI:1352495] | Zfp385a | Enriched in P15 dLGN | 1.435 | 0.000 | 618.400 |
| ENSMUSG00000027274 | McKusick-Kaufman syndrome [Source:MGI Symbol;Acc:MGI:1891836] | Mkks | Enriched in P15 dLGN | 1.435 | 0.030 | 21.920 |
| ENSMUSG00000031367 | adaptor-related protein complex 1, sigma 2 subunit [Source:MGI Symbol;Acc:MGI:1889383] | Ap1s2 | Enriched in P15 dLGN | 1.434 | 0.009 | 90.630 |
| ENSMUSG00000030614 | transmembrane protein 126B [Source:MGI Symbol;Acc:MGI:1915722] | Tmem126b | Enriched in P15 dLGN | 1.433 | 0.000 | 86.830 |
| ENSMUSG00000033319 | fem 1 homolog c [Source:MGI Symbol;Acc:MGI:2444737] | Fem1c | Enriched in P15 dLGN | 1.430 | 0.000 | 163.300 |
| ENSMUSG00000007338 | mitochondrial ribosomal protein L49 [Source:MGI Symbol;Acc:MGI:108180] | Mrpl49 | Enriched in P15 dLGN | 1.427 | 0.000 | 74.110 |
| ENSMUSG00000068696 | G-protein coupled receptor 88 [Source:MGI Symbol;Acc:MGI:1927653] | Gpr88 | Enriched in P15 dLGN | 1.426 | 0.000 | 66.640 |
| ENSMUSG00000007891 | cathepsin D [Source:MGI Symbol;Acc:MGI:88562] | Ctsd | Enriched in P15 dLGN | 1.421 | 0.015 | 65.280 |
| ENSMUSG00000108841 | FERM and PDZ domain containing 2 [Source:MGI Symbol;Acc:MGI:2685472] | Frmpd2 | Enriched in P15 dLGN | 1.421 | 0.037 | 339.600 |
| ENSMUSG00000033352 | mitogen-activated protein kinase kinase 4 [Source:MGI Symbol;Acc:MGI:1346869] | Map2k4 | Enriched in P15 dLGN | 1.420 | 0.000 | 341.700 |
| ENSMUSG00000045427 | heterogeneous nuclear ribonucleoprotein H2 [Source:MGI Symbol;Acc:MGI:1201779] | Hnrnph2 | Enriched in P15 dLGN | 1.417 | 0.000 | 111.000 |
| ENSMUSG00000034891 | synuclein, beta [Source:MGI Symbol;Acc:MGI:1889011] | Sncb | Enriched in P15 dLGN | 1.416 | 0.000 | 939.200 |
| ENSMUSG00000036257 | patatin-like phospholipase domain containing 8 [Source:MGI Symbol;Acc:MGI:1914702] | Pnpla8 | Enriched in P15 dLGN | 1.416 | 0.000 | 213.700 |
| ENSMUSG00000014313 | cytochrome c oxidase subunit 6C [Source:MGI Symbol;Acc:MGI:104614] | Cox6c | Enriched in P15 dLGN | 1.415 | 0.000 | 345.400 |
| ENSMUSG00000019087 | ATPase, H+ transporting, lysosomal accessory protein 1 [Source:MGI Symbol;Acc:MGI:109629] | Atp6ap1 | Enriched in P15 dLGN | 1.411 | 0.000 | 276.800 |
| ENSMUSG00000029918 | mitochondrial ribosomal protein S33 [Source:MGI Symbol;Acc:MGI:1338046] | Mrps33 | Enriched in P15 dLGN | 1.411 | 0.011 | 26.360 |
| ENSMUSG00000039809 | gamma-aminobutyric acid (GABA) B receptor, 2 [Source:MGI Symbol;Acc:MGI:2386030] | Gabbr2 | Enriched in P15 dLGN | 1.411 | 0.000 | 1664.000 |
| ENSMUSG00000031783 | polymerase (RNA) II (DNA directed) polypeptide C [Source:MGI Symbol;Acc:MGI:109299] | Polr2c | Enriched in P15 dLGN | 1.408 | 0.000 | 66.740 |
| ENSMUSG00000062526 | metallophosphoesterase 1 [Source:MGI Symbol;Acc:MGI:2661311] | Mppe1 | Enriched in P15 dLGN | 1.408 | 0.000 | 32.360 |
| ENSMUSG00000053641 | DENN/MADD domain containing 4A [Source:MGI Symbol;Acc:MGI:2142979] | Dennd4a | Enriched in P15 dLGN | 1.404 | 0.000 | 429.700 |
| ENSMUSG00000027263 | tubulin, gamma complex associated protein 4 [Source:MGI Symbol;Acc:MGI:1196293] | Tubgcp4 | Enriched in P15 dLGN | 1.403 | 0.000 | 80.840 |
| ENSMUSG00000025277 | abhydrolase domain containing 6 [Source:MGI Symbol;Acc:MGI:1913332] | Abhd6 | Enriched in P15 dLGN | 1.400 | 0.000 | 71.870 |
| ENSMUSG00000062070 | phosphoglycerate kinase 1 [Source:MGI Symbol;Acc:MGI:97555] | Pgk1 | Enriched in P15 dLGN | 1.399 | 0.000 | 300.000 |
| ENSMUSG00000030327 | NECAP endocytosis associated 1 [Source:MGI Symbol;Acc:MGI:1914852] | Necap1 | Enriched in P15 dLGN | 1.397 | 0.000 | 167.400 |
| ENSMUSG00000075706 | glutathione peroxidase 4 [Source:MGI Symbol;Acc:MGI:104767] | Gpx4 | Enriched in P15 dLGN | 1.394 | 0.000 | 170.900 |
| ENSMUSG00000025477 | inositol polyphosphate-5-phosphatase A [Source:MGI Symbol;Acc:MGI:2686961] | Inpp5a | Enriched in P15 dLGN | 1.393 | 0.000 | 299.200 |
| ENSMUSG00000021259 | cytochrome P450, family 46, subfamily a, polypeptide 1 [Source:MGI Symbol;Acc:MGI:1341877] | Cyp46a1 | Enriched in P15 dLGN | 1.390 | 0.000 | 248.200 |
| ENSMUSG00000039983 | coiled-coil domain containing 32 [Source:MGI Symbol;Acc:MGI:2685477] | Ccdc32 | Enriched in P15 dLGN | 1.388 | 0.000 | 49.020 |
| ENSMUSG00000000088 | cytochrome c oxidase subunit 5A [Source:MGI Symbol;Acc:MGI:88474] | Cox5a | Enriched in P15 dLGN | 1.386 | 0.000 | 112.100 |
| ENSMUSG00000049811 | family with sequence similarity 161, member A [Source:MGI Symbol;Acc:MGI:1921123] | Fam161a | Enriched in P15 dLGN | 1.386 | 0.049 | 823.600 |
| ENSMUSG00000031668 | eukaryotic translation initiation factor 2 alpha kinase 3 [Source:MGI Symbol;Acc:MGI:1341830] | Eif2ak3 | Enriched in P15 dLGN | 1.385 | 0.000 | 198.600 |
| ENSMUSG00000006057 | ATP synthase, H+ transporting, mitochondrial F0 complex, subunit C1 (subunit 9) [Source:MGI Symbol;Acc:MGI:107653] | Atp5g1 | Enriched in P15 dLGN | 1.384 | 0.000 | 76.530 |
| ENSMUSG00000031198 | FUN14 domain containing 2 [Source:MGI Symbol;Acc:MGI:1914641] | Fundc2 | Enriched in P15 dLGN | 1.384 | 0.000 | 67.820 |
| ENSMUSG00000030303 | fatty acyl CoA reductase 2 [Source:MGI Symbol;Acc:MGI:2687035] | Far2 | Enriched in P15 dLGN | 1.383 | 0.006 | 96.510 |
| ENSMUSG00000052395 | RFT1 homolog [Source:MGI Symbol;Acc:MGI:3607791] | Rft1 | Enriched in P15 dLGN | 1.382 | 0.006 | 30.510 |
| ENSMUSG00000026664 | phytanoyl-CoA hydroxylase [Source:MGI Symbol;Acc:MGI:891978] | Phyh | Enriched in P15 dLGN | 1.379 | 0.000 | 93.100 |
| ENSMUSG00000036578 | FXYD domain-containing ion transport regulator 7 [Source:MGI Symbol;Acc:MGI:1889006] | Fxyd7 | Enriched in P15 dLGN | 1.377 | 0.001 | 140.400 |
| ENSMUSG00000051978 | glutamate rich 1 [Source:MGI Symbol;Acc:MGI:3588201] | Erich1 | Enriched in P15 dLGN | 1.375 | 0.019 | 30.180 |
| ENSMUSG00000057766 | ankyrin repeat domain 29 [Source:MGI Symbol;Acc:MGI:2687055] | Ankrd29 | Enriched in P15 dLGN | 1.373 | 0.000 | 58.300 |
| ENSMUSG00000028528 | DnaJ heat shock protein family (Hsp40) member C6 [Source:MGI Symbol;Acc:MGI:1919935] | Dnajc6 | Enriched in P15 dLGN | 1.372 | 0.000 | 781.500 |
| ENSMUSG00000019210 | ATPase, H+ transporting, lysosomal V1 subunit E1 [Source:MGI Symbol;Acc:MGI:894326] | Atp6v1e1 | Enriched in P15 dLGN | 1.363 | 0.000 | 270.400 |
| ENSMUSG00000036295 | leucine rich repeat protein 3, neuronal [Source:MGI Symbol;Acc:MGI:106036] | Lrrn3 | Enriched in P15 dLGN | 1.363 | 0.001 | 123.900 |
| ENSMUSG00000042851 | zinc finger CCCH type containing 6 [Source:MGI Symbol;Acc:MGI:1926001] | Zc3h6 | Enriched in P15 dLGN | 1.362 | 0.000 | 88.340 |

| | | | | | | |
|---|---|---|---|---|---|---|
| ENSMUSG00000063888 | ribosomal protein L7-like 1 [Source:MGI Symbol;Acc:MGI:1913479] | Rpl7l1 | Enriched in P15 dLGN | 1.361 | 0.000 | 93.850 |
| ENSMUSG00000035372 | RIKEN cDNA 1810055G02 gene [Source:MGI Symbol;Acc:MGI:1919306] | 1810055G02Rik | Enriched in P15 dLGN | 1.358 | 0.006 | 34.570 |
| ENSMUSG00000027206 | COP9 signalosome subunit 2 [Source:MGI Symbol;Acc:MGI:1330276] | Cops2 | Enriched in P15 dLGN | 1.355 | 0.000 | 139.200 |
| ENSMUSG00000036438 | calmodulin 2 [Source:MGI Symbol;Acc:MGI:103250] | Calm2 | Enriched in P15 dLGN | 1.354 | 0.002 | 1256.000 |
| ENSMUSG00000025421 | haloacid dehalogenase-like hydrolase domain containing 2 [Source:MGI Symbol;Acc:MGI:1924237] | Hdhd2 | Enriched in P15 dLGN | 1.352 | 0.000 | 217.200 |
| ENSMUSG00000027346 | glycerophosphocholine phosphodiesterase 1 [Source:MGI Symbol;Acc:MGI:104898] | Gpcpd1 | Enriched in P15 dLGN | 1.352 | 0.000 | 161.000 |
| ENSMUSG00000006736 | tetraspanin 31 [Source:MGI Symbol;Acc:MGI:1914375] | Tspan31 | Enriched in P15 dLGN | 1.350 | 0.000 | 73.390 |
| ENSMUSG00000027634 | N-myc downstream regulated gene 3 [Source:MGI Symbol;Acc:MGI:1352499] | Ndrg3 | Enriched in P15 dLGN | 1.348 | 0.000 | 572.800 |
| ENSMUSG00000022139 | muscleblind like splicing factor 2 [Source:MGI Symbol;Acc:MGI:2145597] | Mbnl2 | Enriched in P15 dLGN | 1.346 | 0.000 | 483.000 |
| ENSMUSG00000032192 | guanine nucleotide binding protein (G protein), beta 5 [Source:MGI Symbol;Acc:MGI:101848] | Gnb5 | Enriched in P15 dLGN | 1.346 | 0.000 | 441.800 |
| ENSMUSG00000068749 | proteasome subunit alpha 5 [Source:MGI Symbol;Acc:MGI:1347009] | Psma5 | Enriched in P15 dLGN | 1.346 | 0.003 | 53.280 |
| ENSMUSG00000024099 | NADH:ubiquinone oxidoreductase core subunit V2 [Source:MGI Symbol;Acc:MGI:1920150] | Ndufv2 | Enriched in P15 dLGN | 1.341 | 0.000 | 217.500 |
| ENSMUSG00000032011 | thymus cell antigen 1, theta [Source:MGI Symbol;Acc:MGI:98747] | Thy1 | Enriched in P15 dLGN | 1.341 | 0.000 | 952.800 |
| ENSMUSG00000035964 | transmembrane protein 59-like [Source:MGI Symbol;Acc:MGI:1915187] | Tmem59l | Enriched in P15 dLGN | 1.340 | 0.000 | 166.000 |
| ENSMUSG00000024127 | prolyl endopeptidase-like [Source:MGI Symbol;Acc:MGI:2441932] | Prepl | Enriched in P15 dLGN | 1.338 | 0.000 | 815.800 |
| ENSMUSG00000030879 | mitochondrial ribosomal protein L17 [Source:MGI Symbol;Acc:MGI:1351608] | Mrpl17 | Enriched in P15 dLGN | 1.338 | 0.000 | 76.990 |
| ENSMUSG00000037492 | zinc finger, matrin type 4 [Source:MGI Symbol;Acc:MGI:2443497] | Zmat4 | Enriched in P15 dLGN | 1.338 | 0.000 | 529.700 |
| ENSMUSG00000045009 | proline-rich transmembrane protein 3 [Source:MGI Symbol;Acc:MGI:2444810] | Prrt3 | Enriched in P15 dLGN | 1.337 | 0.000 | 111.100 |
| ENSMUSG00000020589 | CYFIP related Rac1 interactor A [Source:MGI Symbol;Acc:MGI:1261783] | Cyria | Enriched in P15 dLGN | 1.336 | 0.000 | 283.700 |
| ENSMUSG00000020153 | NADH:ubiquinone oxidoreductase core subunit S7 [Source:MGI Symbol;Acc:MGI:1922656] | Ndufs7 | Enriched in P15 dLGN | 1.329 | 0.000 | 124.000 |
| ENSMUSG00000022307 | oxidation resistance 1 [Source:MGI Symbol;Acc:MGI:2179326] | Oxr1 | Enriched in P15 dLGN | 1.328 | 0.000 | 366.300 |
| ENSMUSG00000031095 | cullin 4B [Source:MGI Symbol;Acc:MGI:1919834] | Cul4b | Enriched in P15 dLGN | 1.325 | 0.000 | 104.200 |
| ENSMUSG00000021193 | pitrilysin metallepetidase 1 [Source:MGI Symbol;Acc:MGI:1916867] | Pitrm1 | Enriched in P15 dLGN | 1.324 | 0.000 | 115.900 |
| ENSMUSG00000027134 | lysophosphatidylcholine acyltransferase 4 [Source:MGI Symbol;Acc:MGI:2138993] | Lpcat4 | Enriched in P15 dLGN | 1.324 | 0.000 | 130.500 |
| ENSMUSG00000045733 | shadow of prion protein [Source:MGI Symbol;Acc:MGI:3582583] | Sprn | Enriched in P15 dLGN | 1.324 | 0.000 | 239.000 |
| ENSMUSG00000027406 | isocitrate dehydrogenase 3 (NAD+) beta [Source:MGI Symbol;Acc:MGI:2158650] | Idh3b | Enriched in P15 dLGN | 1.323 | 0.000 | 236.200 |
| ENSMUSG00000010914 | pyruvate dehydrogenase complex, component X [Source:MGI Symbol;Acc:MGI:1351627] | Pdhx | Enriched in P15 dLGN | 1.320 | 0.000 | 152.700 |
| ENSMUSG00000052299 | listerin E3 ubiquitin protein ligase 1 [Source:MGI Symbol;Acc:MGI:1926163] | Ltn1 | Enriched in P15 dLGN | 1.320 | 0.000 | 189.100 |
| ENSMUSG00000059734 | NADH:ubiquinone oxidoreductase core subunit S8 [Source:MGI Symbol;Acc:MGI:2385079] | Ndufs8 | Enriched in P15 dLGN | 1.320 | 0.001 | 73.630 |
| ENSMUSG00000030298 | SEC13 homolog, nuclear pore and COPII coat complex component [Source:MGI Symbol;Acc:MGI:99832] | Sec13 | Enriched in P15 dLGN | 1.316 | 0.003 | 54.450 |
| ENSMUSG00000040907 | ATPase, Na+/K+ transporting, alpha 3 polypeptide [Source:MGI Symbol;Acc:MGI:88107] | Atp1a3 | Enriched in P15 dLGN | 1.314 | 0.000 | 7574.000 |
| ENSMUSG00000029247 | phosphoribosylaminoimidazole carboxylase, phosphoribosylaminoribosylaminoimidazole, succinocarboxamide synthetase [Source:MGI Symbol;Acc:MGI:1914304] | Paics | Enriched in P15 dLGN | 1.311 | 0.001 | 114.100 |
| ENSMUSG00000031782 | coenzyme Q9 [Source:MGI Symbol;Acc:MGI:1915164] | Coq9 | Enriched in P15 dLGN | 1.311 | 0.000 | 96.750 |
| ENSMUSG00000023960 | ectonucleotide pyrophosphatase/phosphodiesterase 5 [Source:MGI Symbol;Acc:MGI:1933830] | Enpp5 | Enriched in P15 dLGN | 1.306 | 0.000 | 293.800 |
| ENSMUSG00000031672 | glutamatic-oxaloacetic transaminase 2, mitochondrial [Source:MGI Symbol;Acc:MGI:95792] | Got2 | Enriched in P15 dLGN | 1.305 | 0.000 | 253.700 |
| ENSMUSG00000023572 | cyclin D-type binding-protein 1 [Source:MGI Symbol;Acc:MGI:109595] | Ccndbp1 | Enriched in P15 dLGN | 1.304 | 0.000 | 105.300 |
| ENSMUSG00000020283 | peroxisomal biogenesis factor 13 [Source:MGI Symbol;Acc:MGI:1919379] | Pex13 | Enriched in P15 dLGN | 1.302 | 0.001 | 60.130 |
| ENSMUSG00000015656 | heat shock protein 8 [Source:MGI Symbol;Acc:MGI:105384] | Hspa8 | Enriched in P15 dLGN | 1.301 | 0.008 | 1562.000 |
| ENSMUSG00000049092 | G protein-coupled receptor 137C [Source:MGI Symbol;Acc:MGI:1917963] | Gpr137c | Enriched in P15 dLGN | 1.301 | 0.001 | 120.200 |
| ENSMUSG00000021373 | CAP, adenylate cyclase-associated protein, 2 (yeast) [Source:MGI Symbol;Acc:MGI:1914502] | Cap2 | Enriched in P15 dLGN | 1.300 | 0.000 | 147.000 |
| ENSMUSG00000033061 | regulated endocrine-specific protein 18 [Source:MGI Symbol;Acc:MGI:1098222] | Resp18 | Enriched in P15 dLGN | 1.299 | 0.050 | 74.080 |
| ENSMUSG00000038722 | BUD31 homolog [Source:MGI Symbol;Acc:MGI:2141291] | Bud31 | Enriched in P15 dLGN | 1.295 | 0.000 | 48.970 |
| ENSMUSG00000044715 | GSK3B interacting protein [Source:MGI Symbol;Acc:MGI:1914037] | Gskip | Enriched in P15 dLGN | 1.293 | 0.015 | 32.920 |
| ENSMUSG00000025971 | matrix AAA peptidase interacting protein 1 [Source:MGI Symbol;Acc:MGI:1915365] | Maip1 | Enriched in P15 dLGN | 1.291 | 0.002 | 36.680 |
| ENSMUSG00000020664 | dihydrolipoamide dehydrogenase [Source:MGI Symbol;Acc:MGI:107450] | Dld | Enriched in P15 dLGN | 1.290 | 0.000 | 258.400 |
| ENSMUSG00000021917 | signal peptidase complex subunit 1 homolog (S. cerevisiae) [Source:MGI Symbol;Acc:MGI:1916269] | Spcs1 | Enriched in P15 dLGN | 1.289 | 0.003 | 46.820 |
| ENSMUSG00000073838 | Tu translation elongation factor, mitochondrial [Source:MGI Symbol;Acc:MGI:1923686] | Tufm | Enriched in P15 dLGN | 1.289 | 0.000 | 100.200 |
| ENSMUSG00000087687 | PET100 homolog [Source:MGI Symbol;Acc:MGI:3615306] | Pet100 | Enriched in P15 dLGN | 1.289 | 0.000 | 59.250 |
| ENSMUSG00000021224 | NUMB endocytic adaptor protein [Source:MGI Symbol;Acc:MGI:107423] | Numb | Enriched in P15 dLGN | 1.288 | 0.008 | 88.320 |
| ENSMUSG00000025035 | ADP-ribosylation factor-like 3 [Source:MGI Symbol;Acc:MGI:1929699] | Arl3 | Enriched in P15 dLGN | 1.288 | 0.000 | 316.800 |
| ENSMUSG00000026709 | aspartyl-tRNA synthetase 2 (mitochondrial) [Source:MGI Symbol;Acc:MGI:2442510] | Dars2 | Enriched in P15 dLGN | 1.288 | 0.004 | 36.150 |
| ENSMUSG00000016637 | intraflagellar transport 27 [Source:MGI Symbol;Acc:MGI:1914292] | Ift27 | Enriched in P15 dLGN | 1.287 | 0.045 | 35.910 |
| ENSMUSG00000022108 | integral membrane protein 2B [Source:MGI Symbol;Acc:MGI:1309517] | Itm2b | Enriched in P15 dLGN | 1.284 | 0.000 | 723.800 |
| ENSMUSG00000056091 | ST3 beta-galactoside alpha-2,3-sialyltransferase 5 [Source:MGI Symbol;Acc:MGI:1339963] | St3gal5 | Enriched in P15 dLGN | 1.282 | 0.000 | 176.300 |
| ENSMUSG00000002767 | mitochondrial ribosomal protein L2 [Source:MGI Symbol;Acc:MGI:1351622] | Mrpl2 | Enriched in P15 dLGN | 1.281 | 0.000 | 55.020 |
| ENSMUSG00000006024 | N-ethylmaleimide sensitive fusion protein attachment protein alpha [Source:MGI Symbol;Acc:MGI:104563] | Napa | Enriched in P15 dLGN | 1.279 | 0.000 | 230.600 |
| ENSMUSG00000026688 | microsomal glutathione S-transferase 3 [Source:MGI Symbol;Acc:MGI:1913697] | Mgst3 | Enriched in P15 dLGN | 1.275 | 0.021 | 88.460 |
| ENSMUSG00000018800 | ATP-binding cassette, sub-family A (ABC1), member 5 [Source:MGI Symbol;Acc:MGI:2386607] | Abca5 | Enriched in P15 dLGN | 1.272 | 0.000 | 222.200 |
| ENSMUSG00000024194 | cutA divalent cation tolerance homolog [Source:MGI Symbol;Acc:MGI:1914925] | Cuta | Enriched in P15 dLGN | 1.272 | 0.000 | 66.240 |
| ENSMUSG00000034075 | zinc finger, DHHC domain containing 5 [Source:MGI Symbol;Acc:MGI:1923573] | Zdhhc5 | Enriched in P15 dLGN | 1.272 | 0.000 | 421.500 |
| ENSMUSG00000026959 | glutamate receptor, ionotropic, NMDA1 (zeta 1) [Source:MGI Symbol;Acc:MGI:95819] | Grin1 | Enriched in P15 dLGN | 1.271 | 0.000 | 942.600 |
| ENSMUSG00000028854 | solute carrier family 9 (sodium/hydrogen exchanger), member 1 [Source:MGI Symbol;Acc:MGI:102462] | Slc9a1 | Enriched in P15 dLGN | 1.271 | 0.000 | 175.500 |
| ENSMUSG00000036860 | mitochondrial ribosomal protein L55 [Source:MGI Symbol;Acc:MGI:1914462] | Mrpl55 | Enriched in P15 dLGN | 1.270 | 0.016 | 27.610 |

| | | | | | | |
|---|---|---|---|---|---|---|
| ENSMUSG00000024668 | succinate dehydrogenase complex assembly factor 2 [Source:MGI Symbol;Acc:MGI:1913322] | Sdhaf2 | Enriched in P15 dLGN | 1.269 | 0.000 | 57.830 |
| ENSMUSG00000019373 | COP9 signalosome subunit 3 [Source:MGI Symbol;Acc:MGI:1349409] | Cops3 | Enriched in P15 dLGN | 1.268 | 0.002 | 92.310 |
| ENSMUSG00000033917 | glycerophosphodiester phosphodiesterase 1 [Source:MGI Symbol;Acc:MGI:1891827] | Gde1 | Enriched in P15 dLGN | 1.266 | 0.000 | 234.500 |
| ENSMUSG00000000276 | diacylglycerol kinase, epsilon [Source:MGI Symbol;Acc:MGI:1889276] | Dgke | Enriched in P15 dLGN | 1.258 | 0.000 | 375.400 |
| ENSMUSG00000000168 | dihydrolipoamide S-acetyltransferase (E2 component of pyruvate dehydrogenase complex) [Source:MGI Symbol;Acc:MGI:2385311] | Dlat | Enriched in P15 dLGN | 1.251 | 0.000 | 196.100 |
| ENSMUSG00000001445 | mitochondrial ribosomal protein L10 [Source:MGI Symbol;Acc:MGI:1333801] | Mrpl10 | Enriched in P15 dLGN | 1.250 | 0.000 | 68.560 |
| ENSMUSG00000002379 | NADH:ubiquinone oxidoreductase subunit A11 [Source:MGI Symbol;Acc:MGI:1917125] | Ndufa11 | Enriched in P15 dLGN | 1.250 | 0.021 | 75.680 |
| ENSMUSG00000025545 | citrate lyase beta like [Source:MGI Symbol;Acc:MGI:1916884] | Clybl | Enriched in P15 dLGN | 1.249 | 0.014 | 35.950 |
| ENSMUSG00000031820 | BRISC and BRCA1 A complex member 1 [Source:MGI Symbol;Acc:MGI:1915501] | Babam1 | Enriched in P15 dLGN | 1.249 | 0.000 | 108.400 |
| ENSMUSG00000028902 | splicing factor 3a, subunit 3 [Source:MGI Symbol;Acc:MGI:1922312] | Sf3a3 | Enriched in P15 dLGN | 1.248 | 0.001 | 94.370 |
| ENSMUSG00000015357 | caseinolytic mitochondrial matrix peptidase chaperone subunit [Source:MGI Symbol;Acc:MGI:1346017] | Clpx | Enriched in P15 dLGN | 1.247 | 0.000 | 114.000 |
| ENSMUSG00000020225 | transmembrane BAX inhibitor motif containing 4 [Source:MGI Symbol;Acc:MGI:1915462] | Tmbim4 | Enriched in P15 dLGN | 1.245 | 0.020 | 25.140 |
| ENSMUSG00000002014 | signal sequence receptor, delta [Source:MGI Symbol;Acc:MGI:1099464] | Ssr4 | Enriched in P15 dLGN | 1.244 | 0.009 | 40.840 |
| ENSMUSG00000005823 | G protein-coupled receptor 108 [Source:MGI Symbol;Acc:MGI:1925558] | Gpr108 | Enriched in P15 dLGN | 1.243 | 0.006 | 51.170 |
| ENSMUSG00000023089 | NADH:ubiquinone oxidoreductase subunit A5 [Source:MGI Symbol;Acc:MGI:1915452] | Ndufa5 | Enriched in P15 dLGN | 1.243 | 0.000 | 166.100 |
| ENSMUSG00000063931 | peptidase D [Source:MGI Symbol;Acc:MGI:97542] | Pepd | Enriched in P15 dLGN | 1.243 | 0.002 | 102.600 |
| ENSMUSG00000090841 | myosin, light polypeptide 6, alkali, smooth muscle and non-muscle [Source:MGI Symbol;Acc:MGI:109318] | Myl6 | Enriched in P15 dLGN | 1.242 | 0.000 | 274.200 |
| ENSMUSG00000027305 | NADH:ubiquinone oxidoreductase complex assembly factor 1 [Source:MGI Symbol;Acc:MGI:1916952] | Ndufaf1 | Enriched in P15 dLGN | 1.239 | 0.002 | 32.640 |
| ENSMUSG00000015536 | molybdenum cofactor synthesis 2 [Source:MGI Symbol;Acc:MGI:1336894] | Mocs2 | Enriched in P15 dLGN | 1.238 | 0.000 | 73.920 |
| ENSMUSG00000021218 | guanosine diphosphate (GDP) dissociation inhibitor 2 [Source:MGI Symbol;Acc:MGI:99845] | Gdi2 | Enriched in P15 dLGN | 1.238 | 0.000 | 530.500 |
| ENSMUSG00000013736 | tRNA nucleotidyl transferase, CCA-adding, 1 [Source:MGI Symbol;Acc:MGI:1917297] | Trnt1 | Enriched in P15 dLGN | 1.236 | 0.000 | 70.420 |
| ENSMUSG00000050786 | coiled-coil domain containing 126 [Source:MGI Symbol;Acc:MGI:1889376] | Ccdc126 | Enriched in P15 dLGN | 1.233 | 0.006 | 193.500 |
| ENSMUSG00000053025 | synaptic vesicle glycoprotein 2 b [Source:MGI Symbol;Acc:MGI:1927338] | Sv2b | Enriched in P15 dLGN | 1.233 | 0.000 | 914.900 |
| ENSMUSG00000021494 | DEAD box helicase 41 [Source:MGI Symbol;Acc:MGI:1920185] | Ddx41 | Enriched in P15 dLGN | 1.232 | 0.000 | 122.300 |
| ENSMUSG00000021728 | embigin [Source:MGI Symbol;Acc:MGI:95321] | Emb | Enriched in P15 dLGN | 1.232 | 0.005 | 88.850 |
| ENSMUSG00000012609 | tubulin tyrosine ligase-like family, member 5 [Source:MGI Symbol;Acc:MGI:2443657] | Ttll5 | Enriched in P15 dLGN | 1.231 | 0.000 | 337.500 |
| ENSMUSG00000016349 | eukaryotic translation elongation factor 1 alpha 2 [Source:MGI Symbol;Acc:MGI:1096317] | Eef1a2 | Enriched in P15 dLGN | 1.230 | 0.000 | 1496.000 |
| ENSMUSG00000031633 | solute carrier family 25 (mitochondrial carrier, adenine nucleotide translocator), member 4 [Source:MGI Symbol;Acc:MGI:1353495] | Slc25a4 | Enriched in P15 dLGN | 1.225 | 0.000 | 1489.000 |
| ENSMUSG00000017801 | MAX-like protein X [Source:MGI Symbol;Acc:MGI:108398] | Mlx | Enriched in P15 dLGN | 1.223 | 0.032 | 24.550 |
| ENSMUSG00000027893 | S-adenosylhomocysteine hydrolase-like 1 [Source:MGI Symbol;Acc:MGI:2385184] | Ahcyl1 | Enriched in P15 dLGN | 1.222 | 0.000 | 829.100 |
| ENSMUSG00000035849 | keratin 222 [Source:MGI Symbol;Acc:MGI:2442728] | Krt222 | Enriched in P15 dLGN | 1.222 | 0.000 | 127.000 |
| ENSMUSG00000051285 | protein-L-isoaspartate (D-aspartate) O-methyltransferase domain containing 1 [Source:MGI Symbol;Acc:MGI:2441773] | Pcmtd1 | Enriched in P15 dLGN | 1.221 | 0.000 | 180.900 |
| ENSMUSG00000020650 | B cell receptor associated protein 29 [Source:MGI Symbol;Acc:MGI:101917] | Bcap29 | Enriched in P15 dLGN | 1.216 | 0.020 | 35.010 |
| ENSMUSG00000025531 | choroidermia (RAB escort protein 1) [Source:MGI Symbol;Acc:MGI:892979] | Chm | Enriched in P15 dLGN | 1.216 | 0.001 | 90.380 |
| ENSMUSG00000033849 | UDP-Gal:betaGlcNAc beta 1,3-galactosyltransferase, polypeptide 2 [Source:MGI Symbol;Acc:MGI:1349461] | B3galt2 | Enriched in P15 dLGN | 1.216 | 0.008 | 115.100 |
| ENSMUSG00000045967 | G protein-coupled receptor 158 [Source:MGI Symbol;Acc:MGI:2441697] | Gpr158 | Enriched in P15 dLGN | 1.216 | 0.000 | 404.300 |
| ENSMUSG00000026750 | proteasome (prosome, macropain) subunit, beta type 7 [Source:MGI Symbol;Acc:MGI:107637] | Psmb7 | Enriched in P15 dLGN | 1.215 | 0.000 | 169.100 |
| ENSMUSG00000027674 | peroxisomal biogenesis factor 5-like [Source:MGI Symbol;Acc:MGI:1916672] | Pex5l | Enriched in P15 dLGN | 1.213 | 0.007 | 789.100 |
| ENSMUSG00000036067 | solute carrier family 2 (facilitated glucose transporter), member 6 [Source:MGI Symbol;Acc:MGI:2443286] | Slc2a6 | Enriched in P15 dLGN | 1.212 | 0.007 | 33.470 |
| ENSMUSG00000111375 | BTB (POZ) domain containing 8 [Source:MGI Symbol;Acc:MGI:3646208] | Btbd8 | Enriched in P15 dLGN | 1.212 | 0.000 | 197.000 |
| ENSMUSG00000028364 | tenascin C [Source:MGI Symbol;Acc:MGI:101922] | Tnc | Enriched in P15 dLGN | 1.211 | 0.011 | 268.500 |
| ENSMUSG00000041729 | coronin, actin binding protein, 2B [Source:MGI Symbol;Acc:MGI:2444283] | Coro2b | Enriched in P15 dLGN | 1.210 | 0.000 | 607.500 |
| ENSMUSG00000072704 | small integral membrane protein 10 like 1 [Source:MGI Symbol;Acc:MGI:1914379] | Smim10l1 | Enriched in P15 dLGN | 1.210 | 0.002 | 297.100 |
| ENSMUSG00000020801 | mediator complex subunit 31 [Source:MGI Symbol;Acc:MGI:1914529] | Med31 | Enriched in P15 dLGN | 1.209 | 0.041 | 18.700 |
| ENSMUSG00000026037 | origin recognition complex, subunit 2 [Source:MGI Symbol;Acc:MGI:1328306] | Orc2 | Enriched in P15 dLGN | 1.209 | 0.002 | 70.880 |
| ENSMUSG00000027652 | Ral GTPase activating protein, beta subunit (non-catalytic) [Source:MGI Symbol;Acc:MGI:2444531] | Ralgapb | Enriched in P15 dLGN | 1.209 | 0.000 | 446.900 |
| ENSMUSG00000041777 | corepressor interacting with RBPJ, 1 [Source:MGI Symbol;Acc:MGI:1914185] | Cir1 | Enriched in P15 dLGN | 1.209 | 0.006 | 48.040 |
| ENSMUSG00000050830 | von Willebrand factor C domain containing 2 [Source:MGI Symbol;Acc:MGI:2442987] | Vwc2 | Enriched in P15 dLGN | 1.207 | 0.001 | 127.000 |
| ENSMUSG00000026235 | Eph receptor A4 [Source:MGI Symbol;Acc:MGI:98277] | Epha4 | Enriched in P15 dLGN | 1.205 | 0.000 | 500.900 |
| ENSMUSG00000015980 | leucine rich repeat containing 27 [Source:MGI Symbol;Acc:MGI:1923862] | Lrrc27 | Enriched in P15 dLGN | 1.204 | 0.038 | 44.920 |
| ENSMUSG00000056185 | sorting nexin 32 [Source:MGI Symbol;Acc:MGI:2444704] | Snx32 | Enriched in P15 dLGN | 1.204 | 0.005 | 101.200 |
| ENSMUSG00000040767 | small nuclear ribonucleoprotein 25 (U11/U12) [Source:MGI Symbol;Acc:MGI:1925622] | Snrnp25 | Enriched in P15 dLGN | 1.203 | 0.005 | 25.980 |
| ENSMUSG00000062352 | integrin beta 1 binding protein 1 [Source:MGI Symbol;Acc:MGI:1306802] | Itgb1bp1 | Enriched in P15 dLGN | 1.202 | 0.000 | 59.330 |
| ENSMUSG00000016427 | NADH:ubiquinone oxidoreductase subunit A1 [Source:MGI Symbol;Acc:MGI:1929511] | Ndufa1 | Enriched in P15 dLGN | 1.200 | 0.000 | 84.150 |
| ENSMUSG00000022684 | bifunctional apoptosis regulator [Source:MGI Symbol;Acc:MGI:1914368] | Bfar | Enriched in P15 dLGN | 1.199 | 0.000 | 155.000 |
| ENSMUSG00000033685 | uncoupling protein 2 (mitochondrial, proton carrier) [Source:MGI Symbol;Acc:MGI:109354] | Ucp2 | Enriched in P15 dLGN | 1.199 | 0.003 | 149.600 |
| ENSMUSG00000071014 | NADH:ubiquinone oxidoreductase subunit B6 [Source:MGI Symbol;Acc:MGI:2684983] | Ndufb6 | Enriched in P15 dLGN | 1.199 | 0.002 | 50.890 |
| ENSMUSG00000041763 | tripeptidyl peptidase II [Source:MGI Symbol;Acc:MGI:102724] | Tpp2 | Enriched in P15 dLGN | 1.197 | 0.000 | 165.900 |
| ENSMUSG00000053329 | glutamine amidotransferase like class 1 domain containing 3A [Source:MGI Symbol;Acc:MGI:1351861] | Gatd3a | Enriched in P15 dLGN | 1.196 | 0.000 | 113.500 |
| ENSMUSG00000025439 | chloride channel, nucleotide-sensitive, 1A [Source:MGI Symbol;Acc:MGI:109638] | Clns1a | Enriched in P15 dLGN | 1.195 | 0.000 | 102.400 |
| ENSMUSG00000030647 | NADH:ubiquinone oxidoreductase subunit C2 [Source:MGI Symbol;Acc:MGI:1344370] | Ndufc2 | Enriched in P15 dLGN | 1.195 | 0.007 | 77.570 |
| ENSMUSG00000025968 | NADH:ubiquinone oxidoreductase core subunit S1 [Source:MGI Symbol;Acc:MGI:2443241] | Ndufs1 | Enriched in P15 dLGN | 1.193 | 0.000 | 337.900 |
| ENSMUSG00000039197 | adenosine kinase [Source:MGI Symbol;Acc:MGI:87930] | Adk | Enriched in P15 dLGN | 1.193 | 0.011 | 98.440 |

| | | | | | | |
|---|---|---|---|---|---|---|
| ENSMUSG00000019478 | RAB4A, member RAS oncogene family [Source:MGI Symbol;Acc:MGI:105069] | Rab4a | Enriched in P15 dLGN | 1.192 | 0.000 | 91.570 |
| ENSMUSG00000021650 | pentatricopeptide repeat domain 2 [Source:MGI Symbol;Acc:MGI:1916177] | Ptcd2 | Enriched in P15 dLGN | 1.192 | 0.003 | 48.360 |
| ENSMUSG00000054452 | TLE family member 5, transcriptional modulator [Source:MGI Symbol;Acc:MGI:95806] | Tle5 | Enriched in P15 dLGN | 1.192 | 0.000 | 670.500 |
| ENSMUSG00000030057 | cellular nucleic acid binding protein [Source:MGI Symbol;Acc:MGI:88431] | Cnbp | Enriched in P15 dLGN | 1.190 | 0.000 | 267.300 |
| ENSMUSG00000059361 | neurensin 2 [Source:MGI Symbol;Acc:MGI:2684969] | Nrsn2 | Enriched in P15 dLGN | 1.188 | 0.007 | 106.400 |
| ENSMUSG00000028252 | cyclin C [Source:MGI Symbol;Acc:MGI:1858199] | Ccnc | Enriched in P15 dLGN | 1.187 | 0.002 | 88.640 |
| ENSMUSG00000015002 | EFR3 homolog A [Source:MGI Symbol;Acc:MGI:1923990] | Efr3a | Enriched in P15 dLGN | 1.186 | 0.000 | 259.300 |
| ENSMUSG00000039128 | cell division cycle 123 [Source:MGI Symbol;Acc:MGI:2138811] | Cdc123 | Enriched in P15 dLGN | 1.185 | 0.000 | 100.900 |
| ENSMUSG00000021635 | RAD17 checkpoint clamp loader component [Source:MGI Symbol;Acc:MGI:1333807] | Rad17 | Enriched in P15 dLGN | 1.181 | 0.033 | 39.840 |
| ENSMUSG00000056851 | poly(rC) binding protein 2 [Source:MGI Symbol;Acc:MGI:108202] | Pcbp2 | Enriched in P15 dLGN | 1.181 | 0.000 | 828.100 |
| ENSMUSG00000030500 | solute carrier family 17 (sodium-dependent inorganic phosphate cotransporter), member 6 [Source:MGI Symbol;Acc:MGI:2156052] | Slc17a6 | Enriched in P15 dLGN | 1.180 | 0.013 | 605.100 |
| ENSMUSG00000038555 | receptor accessory protein 2 [Source:MGI Symbol;Acc:MGI:2385070] | Reep2 | Enriched in P15 dLGN | 1.175 | 0.000 | 302.700 |
| ENSMUSG00000023861 | mitochondrial pyruvate carrier 1 [Source:MGI Symbol;Acc:MGI:1915240] | Mpc1 | Enriched in P15 dLGN | 1.171 | 0.003 | 73.220 |
| ENSMUSG00000044442 | N-6 adenine-specific DNA methyltransferase 1 (putative) [Source:MGI Symbol;Acc:MGI:1915018] | N6amt1 | Enriched in P15 dLGN | 1.168 | 0.003 | 39.220 |
| ENSMUSG00000047126 | clathrin, heavy polypeptide (Hc) [Source:MGI Symbol;Acc:MGI:2388633] | Cltc | Enriched in P15 dLGN | 1.168 | 0.000 | 1529.000 |
| ENSMUSG00000062542 | synaptotagmin IX [Source:MGI Symbol;Acc:MGI:1926373] | Syt9 | Enriched in P15 dLGN | 1.166 | 0.000 | 283.300 |
| ENSMUSG00000071662 | polymerase (RNA) II (DNA directed) polypeptide G [Source:MGI Symbol;Acc:MGI:1914960] | Polr2g | Enriched in P15 dLGN | 1.166 | 0.002 | 40.090 |
| ENSMUSG00000039431 | myotubularin related protein 7 [Source:MGI Symbol;Acc:MGI:1891693] | Mtmr7 | Enriched in P15 dLGN | 1.165 | 0.000 | 199.100 |
| ENSMUSG00000041020 | MAP7 domain containing 2 [Source:MGI Symbol;Acc:MGI:1917474] | Map7d2 | Enriched in P15 dLGN | 1.163 | 0.000 | 736.500 |
| ENSMUSG00000028020 | glycine receptor, beta subunit [Source:MGI Symbol;Acc:MGI:95751] | Glrb | Enriched in P15 dLGN | 1.161 | 0.000 | 323.700 |
| ENSMUSG00000019505 | ubiquitin B [Source:MGI Symbol;Acc:MGI:98888] | Ubb | Enriched in P15 dLGN | 1.159 | 0.011 | 652.000 |
| ENSMUSG00000022231 | sema domain, seven thrombospondin repeats (type 1 and type 1-like), transmembrane domain (TM) and short cytoplasmic domain, (semaphorin) 5A [Source:MGI Symbol;Acc:MGI:107 | Sema5a | Enriched in P15 dLGN | 1.159 | 0.000 | 795.500 |
| ENSMUSG00000022552 | SHANK-associated RH domain interacting protein [Source:MGI Symbol;Acc:MGI:1913331] | Sharpin | Enriched in P15 dLGN | 1.159 | 0.033 | 33.900 |
| ENSMUSG00000033790 | tubulin, gamma complex associated protein 5 [Source:MGI Symbol;Acc:MGI:2178836] | Tubgcp5 | Enriched in P15 dLGN | 1.159 | 0.000 | 100.600 |
| ENSMUSG00000035885 | cytochrome c oxidase subunit 8A [Source:MGI Symbol;Acc:MGI:105959] | Cox8a | Enriched in P15 dLGN | 1.158 | 0.000 | 292.600 |
| ENSMUSG00000021712 | tripartite motif-containing 23 [Source:MGI Symbol;Acc:MGI:1933161] | Trim23 | Enriched in P15 dLGN | 1.154 | 0.000 | 95.960 |
| ENSMUSG00000025393 | ATP synthase, H+ transporting mitochondrial F1 complex, beta subunit [Source:MGI Symbol;Acc:MGI:107801] | Atp5b | Enriched in P15 dLGN | 1.154 | 0.001 | 1784.000 |
| ENSMUSG00000055943 | ER membrane protein complex subunit 7 [Source:MGI Symbol;Acc:MGI:1920274] | Emc7 | Enriched in P15 dLGN | 1.153 | 0.000 | 203.900 |
| ENSMUSG00000017781 | phosphatidylinositol transfer protein, alpha [Source:MGI Symbol;Acc:MGI:99887] | Pitpna | Enriched in P15 dLGN | 1.151 | 0.000 | 792.900 |
| ENSMUSG00000021253 | transforming growth factor, beta 3 [Source:MGI Symbol;Acc:MGI:98727] | Tgfb3 | Enriched in P15 dLGN | 1.149 | 0.000 | 74.630 |
| ENSMUSG00000044252 | oxysterol binding protein-like 1A [Source:MGI Symbol;Acc:MGI:1927551] | Osbpl1a | Enriched in P15 dLGN | 1.149 | 0.000 | 434.200 |
| ENSMUSG00000057230 | AP2 associated kinase 1 [Source:MGI Symbol;Acc:MGI:1098687] | Aak1 | Enriched in P15 dLGN | 1.149 | 0.000 | 1193.000 |
| ENSMUSG00000033781 | ankyrin repeat and SOCS box-containing 13 [Source:MGI Symbol;Acc:MGI:2145525] | Asb13 | Enriched in P15 dLGN | 1.147 | 0.000 | 93.680 |
| ENSMUSG00000034341 | WW domain binding protein 2 [Source:MGI Symbol;Acc:MGI:104709] | Wbp2 | Enriched in P15 dLGN | 1.145 | 0.004 | 271.900 |
| ENSMUSG00000074736 | synapse differentiation inducing 1 [Source:MGI Symbol;Acc:MGI:3702158] | Syndig1 | Enriched in P15 dLGN | 1.145 | 0.021 | 50.550 |
| ENSMUSG00000067242 | leucine-rich repeat LGI family, member 1 [Source:MGI Symbol;Acc:MGI:1861691] | Lgi1 | Enriched in P15 dLGN | 1.142 | 0.005 | 72.290 |
| ENSMUSG00000075000 | nuclear receptor binding factor 2 [Source:MGI Symbol;Acc:MGI:1354950] | Nrbf2 | Enriched in P15 dLGN | 1.142 | 0.003 | 35.460 |
| ENSMUSG00000003721 | insulin induced gene 2 [Source:MGI Symbol;Acc:MGI:1920249] | Insig2 | Enriched in P15 dLGN | 1.140 | 0.001 | 69.600 |
| ENSMUSG00000033124 | autophagy related 9A [Source:MGI Symbol;Acc:MGI:2138446] | Atg9a | Enriched in P15 dLGN | 1.140 | 0.000 | 274.400 |
| ENSMUSG00000028618 | transmembrane protein 59 [Source:MGI Symbol;Acc:MGI:1929278] | Tmem59 | Enriched in P15 dLGN | 1.139 | 0.000 | 245.200 |
| ENSMUSG00000035772 | mitochondrial ribosomal protein S2 [Source:MGI Symbol;Acc:MGI:2153089] | Mrps2 | Enriched in P15 dLGN | 1.139 | 0.003 | 52.550 |
| ENSMUSG00000040620 | DEAH (Asp-Glu-Ala-His) box polypeptide 33 [Source:MGI Symbol;Acc:MGI:2445102] | Dhx33 | Enriched in P15 dLGN | 1.139 | 0.000 | 72.800 |
| ENSMUSG00000028464 | tropomyosin 2, beta [Source:MGI Symbol;Acc:MGI:98810] | Tpm2 | Enriched in P15 dLGN | 1.138 | 0.038 | 25.350 |
| ENSMUSG00000040713 | cellular repressor of E1A-stimulated genes 1 [Source:MGI Symbol;Acc:MGI:1344382] | Creg1 | Enriched in P15 dLGN | 1.138 | 0.001 | 76.450 |
| ENSMUSG00000028271 | general transcription factor IIB [Source:MGI Symbol;Acc:MGI:2385191] | Gtf2b | Enriched in P15 dLGN | 1.136 | 0.019 | 36.100 |
| ENSMUSG00000042743 | small glutamine-rich tetratricopeptide repeat (TPR)-containing, beta [Source:MGI Symbol;Acc:MGI:2444615] | Sgtb | Enriched in P15 dLGN | 1.133 | 0.000 | 167.800 |
| ENSMUSG00000056222 | sparc/osteonectin, cwcv and kazal-like domains proteoglycan 1 [Source:MGI Symbol;Acc:MGI:105371] | Spock1 | Enriched in P15 dLGN | 1.133 | 0.001 | 1204.000 |
| ENSMUSG00000022789 | dynamin 1-like [Source:MGI Symbol;Acc:MGI:1921256] | Dnm1l | Enriched in P15 dLGN | 1.132 | 0.000 | 380.500 |
| ENSMUSG00000027194 | tetratricopeptide repeat domain 17 [Source:MGI Symbol;Acc:MGI:1921819] | Ttc17 | Enriched in P15 dLGN | 1.132 | 0.000 | 248.100 |
| ENSMUSG00000024953 | peroxiredoxin 5 [Source:MGI Symbol;Acc:MGI:1859821] | Prdx5 | Enriched in P15 dLGN | 1.131 | 0.000 | 344.200 |
| ENSMUSG00000060279 | adaptor-related protein complex 2, alpha 1 subunit [Source:MGI Symbol;Acc:MGI:101921] | Ap2a1 | Enriched in P15 dLGN | 1.131 | 0.000 | 671.900 |
| ENSMUSG00000030061 | ubiquitin-like modifier activating enzyme 3 [Source:MGI Symbol;Acc:MGI:1341217] | Uba3 | Enriched in P15 dLGN | 1.129 | 0.001 | 78.390 |
| ENSMUSG00000056201 | cofilin 1, non-muscle [Source:MGI Symbol;Acc:MGI:101757] | Cfl1 | Enriched in P15 dLGN | 1.129 | 0.001 | 762.600 |
| ENSMUSG00000020311 | endoplasmic reticulum lectin 1 [Source:MGI Symbol;Acc:MGI:1914003] | Erlec1 | Enriched in P15 dLGN | 1.126 | 0.000 | 165.800 |
| ENSMUSG00000089911 | major facilitator superfamily domain containing 14A [Source:MGI Symbol;Acc:MGI:1201609] | Mfsd14a | Enriched in P15 dLGN | 1.126 | 0.003 | 53.160 |
| ENSMUSG00000025917 | COP9 signalosome subunit 5 [Source:MGI Symbol;Acc:MGI:1349415] | Cops5 | Enriched in P15 dLGN | 1.125 | 0.008 | 96.980 |
| ENSMUSG00000033278 | protein tyrosine phosphatase, receptor type, M [Source:MGI Symbol;Acc:MGI:102694] | Ptprm | Enriched in P15 dLGN | 1.125 | 0.002 | 370.100 |
| ENSMUSG00000033918 | presenilin associated, rhomboid-like [Source:MGI Symbol;Acc:MGI:1277152] | Parl | Enriched in P15 dLGN | 1.125 | 0.000 | 65.890 |
| ENSMUSG00000024044 | erythrocyte membrane protein band 4.1 like 3 [Source:MGI Symbol;Acc:MGI:103008] | Epb41l3 | Enriched in P15 dLGN | 1.124 | 0.000 | 1430.000 |
| ENSMUSG00000033628 | phosphatidylinositol 3-kinase catalytic subunit type 3 [Source:MGI Symbol;Acc:MGI:2445019] | Pik3c3 | Enriched in P15 dLGN | 1.121 | 0.000 | 127.100 |
| ENSMUSG00000034211 | mitochondrial ribosomal protein S17 [Source:MGI Symbol;Acc:MGI:1913508] | Mrps17 | Enriched in P15 dLGN | 1.121 | 0.005 | 54.270 |
| ENSMUSG00000021945 | zinc finger, MYM-type 2 [Source:MGI Symbol;Acc:MGI:1923257] | Zmym2 | Enriched in P15 dLGN | 1.119 | 0.000 | 289.200 |
| ENSMUSG00000021814 | annexin A7 [Source:MGI Symbol;Acc:MGI:88031] | Anxa7 | Enriched in P15 dLGN | 1.116 | 0.007 | 56.220 |

| | | | | | | |
|---|---|---|---|---|---|---|
| ENSMUSG00000026889 | RNA binding motif protein 18 [Source:MGI Symbol;Acc:MGI:1915139] | Rbm18 | Enriched in P15 dLGN | 1.116 | 0.001 | 87.370 |
| ENSMUSG00000032422 | sorting nexin 14 [Source:MGI Symbol;Acc:MGI:2155664] | Snx14 | Enriched in P15 dLGN | 1.116 | 0.002 | 77.370 |
| ENSMUSG00000021830 | thioredoxin domain containing 16 [Source:MGI Symbol;Acc:MGI:1917811] | Txndc16 | Enriched in P15 dLGN | 1.115 | 0.000 | 229.000 |
| ENSMUSG00000002068 | cyclin E1 [Source:MGI Symbol;Acc:MGI:88316] | Ccne1 | Enriched in P15 dLGN | 1.114 | 0.014 | 29.030 |
| ENSMUSG00000027131 | ER membrane protein complex subunit 4 [Source:MGI Symbol;Acc:MGI:1915282] | Emc4 | Enriched in P15 dLGN | 1.114 | 0.002 | 81.850 |
| ENSMUSG00000037152 | NADH:ubiquinone oxidoreductase subunit C1 [Source:MGI Symbol;Acc:MGI:1913627] | Ndufc1 | Enriched in P15 dLGN | 1.114 | 0.017 | 49.470 |
| ENSMUSG00000035297 | COP9 signalosome subunit 4 [Source:MGI Symbol;Acc:MGI:1349414] | Cops4 | Enriched in P15 dLGN | 1.112 | 0.007 | 86.240 |
| ENSMUSG00000008153 | calsyntenin 3 [Source:MGI Symbol;Acc:MGI:2178323] | Clstn3 | Enriched in P15 dLGN | 1.111 | 0.000 | 1064.000 |
| ENSMUSG00000030591 | proteasome (prosome, macropain) 26S subunit, non-ATPase, 8 [Source:MGI Symbol;Acc:MGI:1888669] | Psmd8 | Enriched in P15 dLGN | 1.110 | 0.000 | 174.800 |
| ENSMUSG00000040118 | calcium channel, voltage-dependent, alpha2/delta subunit 1 [Source:MGI Symbol;Acc:MGI:88295] | Cacna2d1 | Enriched in P15 dLGN | 1.108 | 0.001 | 541.200 |
| ENSMUSG00000041997 | tousled-like kinase 1 [Source:MGI Symbol;Acc:MGI:2441683] | Tlk1 | Enriched in P15 dLGN | 1.104 | 0.000 | 318.700 |
| ENSMUSG00000022722 | ADP-ribosylation factor-like 6 [Source:MGI Symbol;Acc:MGI:1927136] | Arl6 | Enriched in P15 dLGN | 1.102 | 0.020 | 147.900 |
| ENSMUSG00000019943 | ATPase, Ca++ transporting, plasma membrane 1 [Source:MGI Symbol;Acc:MGI:104653] | Atp2b1 | Enriched in P15 dLGN | 1.100 | 0.000 | 2706.000 |
| ENSMUSG00000026032 | NADH:ubiquinone oxidoreductase subunit B3 [Source:MGI Symbol;Acc:MGI:1913745] | Ndufb3 | Enriched in P15 dLGN | 1.100 | 0.003 | 49.010 |
| ENSMUSG00000022855 | SUMO/sentrin specific peptidase 2 [Source:MGI Symbol;Acc:MGI:1923076] | Senp2 | Enriched in P15 dLGN | 1.099 | 0.000 | 371.000 |
| ENSMUSG00000025868 | HIG1 domain family, member 2A [Source:MGI Symbol;Acc:MGI:1914294] | Higd2a | Enriched in P15 dLGN | 1.099 | 0.012 | 47.280 |
| ENSMUSG00000039347 | ATPase, H+ transporting, lysosomal V0 subunit E2 [Source:MGI Symbol;Acc:MGI:1923502] | Atp6v0e2 | Enriched in P15 dLGN | 1.097 | 0.000 | 554.600 |
| ENSMUSG00000025283 | spermidine/spermine N1-acetyl transferase 1 [Source:MGI Symbol;Acc:MGI:98233] | Sat1 | Enriched in P15 dLGN | 1.095 | 0.007 | 78.930 |
| ENSMUSG00000022973 | synaptojanin 1 [Source:MGI Symbol;Acc:MGI:1354961] | Synj1 | Enriched in P15 dLGN | 1.094 | 0.000 | 970.800 |
| ENSMUSG00000029776 | 3-hydroxyisobutyrate dehydrogenase [Source:MGI Symbol;Acc:MGI:1889802] | Hibadh | Enriched in P15 dLGN | 1.093 | 0.012 | 73.620 |
| ENSMUSG00000031696 | VPS35 retromer complex component [Source:MGI Symbol;Acc:MGI:1890467] | Vps35 | Enriched in P15 dLGN | 1.091 | 0.003 | 246.500 |
| ENSMUSG00000015932 | destrin [Source:MGI Symbol;Acc:MGI:1929270] | Dstn | Enriched in P15 dLGN | 1.090 | 0.002 | 300.900 |
| ENSMUSG00000028035 | DnaJ heat shock protein family (Hsp40) member B4 [Source:MGI Symbol;Acc:MGI:1914285] | Dnajb4 | Enriched in P15 dLGN | 1.088 | 0.000 | 121.400 |
| ENSMUSG00000001323 | serine racemase [Source:MGI Symbol;Acc:MGI:1351636] | Srr | Enriched in P15 dLGN | 1.086 | 0.000 | 161.100 |
| ENSMUSG00000025374 | nucleic acid binding protein 2 [Source:MGI Symbol;Acc:MGI:1917167] | Nabp2 | Enriched in P15 dLGN | 1.086 | 0.005 | 92.340 |
| ENSMUSG00000030652 | demethyl-Q 7 [Source:MGI Symbol;Acc:MGI:107207] | Coq7 | Enriched in P15 dLGN | 1.085 | 0.005 | 54.520 |
| ENSMUSG00000022201 | zinc finger RNA binding protein [Source:MGI Symbol;Acc:MGI:1341890] | Zfr | Enriched in P15 dLGN | 1.083 | 0.000 | 1105.000 |
| ENSMUSG00000024359 | heat shock protein 9 [Source:MGI Symbol;Acc:MGI:96245] | Hspa9 | Enriched in P15 dLGN | 1.083 | 0.001 | 574.800 |
| ENSMUSG00000002280 | cytosolic iron-sulfur assembly component 3 [Source:MGI Symbol;Acc:MGI:1914813] | Ciao3 | Enriched in P15 dLGN | 1.081 | 0.009 | 33.900 |
| ENSMUSG00000052456 | guided entry of tail-anchored proteins factor 3, ATPase [Source:MGI Symbol;Acc:MGI:1928379] | Get3 | Enriched in P15 dLGN | 1.081 | 0.001 | 120.500 |
| ENSMUSG00000018882 | mitochondrial ribosomal protein L45 [Source:MGI Symbol;Acc:MGI:1914286] | Mrpl45 | Enriched in P15 dLGN | 1.080 | 0.001 | 64.610 |
| ENSMUSG00000022337 | ER membrane protein complex subunit 2 [Source:MGI Symbol;Acc:MGI:1913986] | Emc2 | Enriched in P15 dLGN | 1.080 | 0.000 | 104.600 |
| ENSMUSG00000036298 | solute carrier family 2 (facilitated glucose transporter), member 13 [Source:MGI Symbol;Acc:MGI:2146030] | Slc2a13 | Enriched in P15 dLGN | 1.080 | 0.004 | 258.300 |
| ENSMUSG00000009894 | synaptosomal-associated protein, 47 [Source:MGI Symbol;Acc:MGI:1915076] | Snap47 | Enriched in P15 dLGN | 1.078 | 0.001 | 533.600 |
| ENSMUSG00000005312 | ubiquilin 1 [Source:MGI Symbol;Acc:MGI:1860276] | Ubqln1 | Enriched in P15 dLGN | 1.077 | 0.000 | 342.600 |
| ENSMUSG00000024487 | Yip1 domain family, member 5 [Source:MGI Symbol;Acc:MGI:1914430] | Yipf5 | Enriched in P15 dLGN | 1.077 | 0.007 | 59.880 |
| ENSMUSG00000036398 | protein phosphatase 1, regulatory inhibitor subunit 11 [Source:MGI Symbol;Acc:MGI:1923747] | Ppp1r11 | Enriched in P15 dLGN | 1.073 | 0.000 | 110.100 |
| ENSMUSG00000038462 | ubiquinol-cytochrome c reductase, Rieske iron-sulfur polypeptide 1 [Source:MGI Symbol;Acc:MGI:1913944] | Uqcrfs1 | Enriched in P15 dLGN | 1.073 | 0.000 | 172.400 |
| ENSMUSG00000022774 | nuclear cap binding protein subunit 2 [Source:MGI Symbol;Acc:MGI:1915342] | Ncbp2 | Enriched in P15 dLGN | 1.072 | 0.000 | 113.200 |
| ENSMUSG00000030965 | BRISC complex subunit [Source:MGI Symbol;Acc:MGI:1926116] | Abraxas2 | Enriched in P15 dLGN | 1.072 | 0.001 | 76.110 |
| ENSMUSG00000041236 | VPS41 HOPS complex subunit [Source:MGI Symbol;Acc:MGI:1929215] | Vps41 | Enriched in P15 dLGN | 1.072 | 0.000 | 365.700 |
| ENSMUSG00000029189 | sel-1 suppressor of lin-12-like 3 (C. elegans) [Source:MGI Symbol;Acc:MGI:1916941] | Sel1l3 | Enriched in P15 dLGN | 1.070 | 0.001 | 154.000 |
| ENSMUSG00000060450 | ring finger protein 14 [Source:MGI Symbol;Acc:MGI:1929668] | Rnf14 | Enriched in P15 dLGN | 1.070 | 0.000 | 686.900 |
| ENSMUSG00000034088 | high density lipoprotein (HDL) binding protein [Source:MGI Symbol;Acc:MGI:99256] | Hdlbp | Enriched in P15 dLGN | 1.069 | 0.000 | 727.200 |
| ENSMUSG00000031918 | myotubularin related protein 2 [Source:MGI Symbol;Acc:MGI:1924366] | Mtmr2 | Enriched in P15 dLGN | 1.066 | 0.000 | 157.400 |
| ENSMUSG00000027598 | itchy, E3 ubiquitin protein ligase [Source:MGI Symbol;Acc:MGI:1202301] | Itch | Enriched in P15 dLGN | 1.065 | 0.000 | 241.900 |
| ENSMUSG00000033096 | adipocyte plasma membrane associated protein [Source:MGI Symbol;Acc:MGI:1919131] | Apmap | Enriched in P15 dLGN | 1.065 | 0.000 | 102.300 |
| ENSMUSG00000027804 | peptidylprolyl isomerase D (cyclophilin D) [Source:MGI Symbol;Acc:MGI:1914988] | Ppid | Enriched in P15 dLGN | 1.063 | 0.001 | 95.780 |
| ENSMUSG00000029388 | eukaryotic translation initiation factor 2B, subunit 1 (alpha) [Source:MGI Symbol;Acc:MGI:2384802] | Eif2b1 | Enriched in P15 dLGN | 1.062 | 0.006 | 40.060 |
| ENSMUSG00000029198 | GrpE-like 1, mitochondrial [Source:MGI Symbol;Acc:MGI:1334417] | Grpel1 | Enriched in P15 dLGN | 1.060 | 0.001 | 89.430 |
| ENSMUSG00000034793 | glucose 6 phosphatase, catalytic, 3 [Source:MGI Symbol;Acc:MGI:1915651] | G6pc3 | Enriched in P15 dLGN | 1.060 | 0.009 | 38.970 |
| ENSMUSG00000034757 | transmembrane and ubiquitin-like domain containing 2 [Source:MGI Symbol;Acc:MGI:1919303] | Tmub2 | Enriched in P15 dLGN | 1.059 | 0.000 | 87.220 |
| ENSMUSG00000109865 | heat shock protein 14 [Source:MGI Symbol;Acc:MGI:1354164] | Hspa14 | Enriched in P15 dLGN | 1.059 | 0.002 | 67.240 |
| ENSMUSG00000021194 | chromogranin A [Source:MGI Symbol;Acc:MGI:88394] | Chga | Enriched in P15 dLGN | 1.056 | 0.006 | 812.400 |
| ENSMUSG00000028567 | thioredoxin domain containing 12 (endoplasmic reticulum) [Source:MGI Symbol;Acc:MGI:1913323] | Txndc12 | Enriched in P15 dLGN | 1.054 | 0.002 | 78.380 |
| ENSMUSG00000035173 | coiled-coil domain containing 186 [Source:MGI Symbol;Acc:MGI:2445022] | Ccdc186 | Enriched in P15 dLGN | 1.054 | 0.000 | 320.300 |
| ENSMUSG00000039568 | UBA-like domain containing 1 [Source:MGI Symbol;Acc:MGI:1916255] | Ubald1 | Enriched in P15 dLGN | 1.054 | 0.000 | 103.800 |
| ENSMUSG00000021209 | protein phosphatase 4, regulatory subunit 4 [Source:MGI Symbol;Acc:MGI:1921771] | Ppp4r4 | Enriched in P15 dLGN | 1.052 | 0.000 | 139.400 |
| ENSMUSG00000017707 | serine incorporator 3 [Source:MGI Symbol;Acc:MGI:1349457] | Serinc3 | Enriched in P15 dLGN | 1.044 | 0.000 | 254.900 |
| ENSMUSG00000022223 | short chain dehydrogenase/reductase family 39U, member 1 [Source:MGI Symbol;Acc:MGI:1916876] | Sdr39u1 | Enriched in P15 dLGN | 1.044 | 0.000 | 84.160 |
| ENSMUSG00000033389 | Rho GTPase activating protein 44 [Source:MGI Symbol;Acc:MGI:2144423] | Arhgap44 | Enriched in P15 dLGN | 1.041 | 0.000 | 522.200 |
| ENSMUSG00000032342 | mitochondrial tRNA translation optimization 1 [Source:MGI Symbol;Acc:MGI:1915541] | Mto1 | Enriched in P15 dLGN | 1.040 | 0.007 | 56.470 |
| ENSMUSG00000025428 | ATP synthase, H+ transporting, mitochondrial F1 complex, alpha subunit 1 [Source:MGI Symbol;Acc:MGI:88115] | Atp5a1 | Enriched in P15 dLGN | 1.039 | 0.002 | 1689.000 |

| | | | | | | |
|---|---|---|---|---|---|---|
| ENSMUSG00000031774 | proteasome activator subunit 3 interacting protein 1 [Source:MGI Symbol;Acc:MGI:1919637] | Psme3ip1 | Enriched in P15 dLGN | 1.039 | 0.000 | 123.300 |
| ENSMUSG00000037058 | polyadenylate-binding protein-interacting protein 2 [Source:MGI Symbol;Acc:MGI:1915119] | Paip2 | Enriched in P15 dLGN | 1.038 | 0.000 | 251.700 |
| ENSMUSG00000033998 | potassium channel, subfamily K, member 1 [Source:MGI Symbol;Acc:MGI:109322] | Kcnk1 | Enriched in P15 dLGN | 1.037 | 0.002 | 107.500 |
| ENSMUSG00000026643 | N-myristoyltransferase 2 [Source:MGI Symbol;Acc:MGI:1202298] | Nmt2 | Enriched in P15 dLGN | 1.036 | 0.000 | 203.800 |
| ENSMUSG00000019810 | fucosidase, alpha-L- 2, plasma [Source:MGI Symbol;Acc:MGI:1914098] | Fuca2 | Enriched in P15 dLGN | 1.035 | 0.010 | 88.280 |
| ENSMUSG00000020955 | adaptor-related protein complex AP-4, sigma 1 [Source:MGI Symbol;Acc:MGI:1337065] | Ap4s1 | Enriched in P15 dLGN | 1.034 | 0.008 | 81.030 |
| ENSMUSG00000021314 | amphiphysin [Source:MGI Symbol;Acc:MGI:103574] | Amph | Enriched in P15 dLGN | 1.033 | 0.000 | 841.600 |
| ENSMUSG00000021114 | ATPase, H+ transporting, lysosomal V1 subunit D [Source:MGI Symbol;Acc:MGI:1921084] | Atp6v1d | Enriched in P15 dLGN | 1.030 | 0.000 | 284.800 |
| ENSMUSG00000026887 | mitochondrial ribosome recycling factor [Source:MGI Symbol;Acc:MGI:1915121] | Mrrf | Enriched in P15 dLGN | 1.028 | 0.019 | 35.970 |
| ENSMUSG00000032745 | GC-rich promoter binding protein 1 [Source:MGI Symbol;Acc:MGI:1920524] | Gpbp1 | Enriched in P15 dLGN | 1.027 | 0.002 | 98.520 |
| ENSMUSG00000042298 | tetratricopeptide repeat domain 19 [Source:MGI Symbol;Acc:MGI:1920045] | Ttc19 | Enriched in P15 dLGN | 1.027 | 0.000 | 462.600 |
| ENSMUSG00000021810 | ecdysoneless cell cycle regulator [Source:MGI Symbol;Acc:MGI:1917851] | Ecd | Enriched in P15 dLGN | 1.026 | 0.002 | 102.100 |
| ENSMUSG00000027603 | gamma-glutamyltransferase 7 [Source:MGI Symbol;Acc:MGI:1913385] | Ggt7 | Enriched in P15 dLGN | 1.025 | 0.000 | 194.700 |
| ENSMUSG00000041408 | WAPL cohesin release factor [Source:MGI Symbol;Acc:MGI:2675859] | Wapl | Enriched in P15 dLGN | 1.023 | 0.002 | 427.800 |
| ENSMUSG00000024425 | Nedd4 family interacting protein 1 [Source:MGI Symbol;Acc:MGI:1929601] | Ndfip1 | Enriched in P15 dLGN | 1.021 | 0.000 | 747.300 |
| ENSMUSG00000030556 | leucine rich repeat containing 28 [Source:MGI Symbol;Acc:MGI:1915689] | Lrrc28 | Enriched in P15 dLGN | 1.021 | 0.047 | 47.460 |
| ENSMUSG00000001056 | NHP2 ribonucleoprotein [Source:MGI Symbol;Acc:MGI:1098547] | Nhp2 | Enriched in P15 dLGN | 1.019 | 0.019 | 27.460 |
| ENSMUSG00000029017 | peptidase (mitochondrial processing) beta [Source:MGI Symbol;Acc:MGI:1920328] | Pmpcb | Enriched in P15 dLGN | 1.018 | 0.001 | 90.430 |
| ENSMUSG00000019818 | CD164 antigen [Source:MGI Symbol;Acc:MGI:1859568] | Cd164 | Enriched in P15 dLGN | 1.017 | 0.042 | 102.300 |
| ENSMUSG00000031791 | transmembrane protein 38A [Source:MGI Symbol;Acc:MGI:1921416] | Tmem38a | Enriched in P15 dLGN | 1.017 | 0.000 | 149.400 |
| ENSMUSG00000038244 | microtubule associated monooxygenase, calponin and LIM domain containing 2 [Source:MGI Symbol;Acc:MGI:2444947] | Mical2 | Enriched in P15 dLGN | 1.016 | 0.003 | 255.100 |
| ENSMUSG00000021737 | proteasome (prosome, macropain) 26S subunit, non-ATPase, 6 [Source:MGI Symbol;Acc:MGI:1913663] | Psmd6 | Enriched in P15 dLGN | 1.015 | 0.001 | 147.100 |
| ENSMUSG00000025980 | heat shock protein 1 (chaperonin) [Source:MGI Symbol;Acc:MGI:96242] | Hspd1 | Enriched in P15 dLGN | 1.015 | 0.000 | 264.000 |
| ENSMUSG00000028488 | SH3-domain GRB2-like 2 [Source:MGI Symbol;Acc:MGI:700009] | Sh3gl2 | Enriched in P15 dLGN | 1.013 | 0.003 | 586.900 |
| ENSMUSG00000029238 | circadian locomotor output cycles kaput [Source:MGI Symbol;Acc:MGI:99698] | Clock | Enriched in P15 dLGN | 1.013 | 0.000 | 497.300 |
| ENSMUSG00000039953 | calsyntenin 1 [Source:MGI Symbol;Acc:MGI:1929895] | Clstn1 | Enriched in P15 dLGN | 1.013 | 0.000 | 3184.000 |
| ENSMUSG00000001366 | f-box protein 9 [Source:MGI Symbol;Acc:MGI:1918788] | Fbxo9 | Enriched in P15 dLGN | 1.012 | 0.000 | 320.200 |
| ENSMUSG00000029407 | USO1 vesicle docking factor [Source:MGI Symbol;Acc:MGI:1929095] | Uso1 | Enriched in P15 dLGN | 1.012 | 0.000 | 254.900 |
| ENSMUSG00000022471 | X-ray repair complementing defective repair in Chinese hamster cells 6 [Source:MGI Symbol;Acc:MGI:95606] | Xrcc6 | Enriched in P15 dLGN | 1.011 | 0.049 | 97.010 |
| ENSMUSG00000034708 | granulin [Source:MGI Symbol;Acc:MGI:95832] | Grn | Enriched in P15 dLGN | 1.011 | 0.007 | 113.500 |
| ENSMUSG00000002718 | chromosome segregation 1-like (S. cerevisiae) [Source:MGI Symbol;Acc:MGI:1339951] | Cse1l | Enriched in P15 dLGN | 1.010 | 0.004 | 100.500 |
| ENSMUSG00000003955 | family with sequence similarity 162, member A [Source:MGI Symbol;Acc:MGI:1917436] | Fam162a | Enriched in P15 dLGN | 1.010 | 0.003 | 60.830 |
| ENSMUSG00000020720 | proteasome (prosome, macropain) 26S subunit, non-ATPase, 12 [Source:MGI Symbol;Acc:MGI:1914247] | Psmd12 | Enriched in P15 dLGN | 1.008 | 0.002 | 128.200 |
| ENSMUSG00000022092 | protein phosphatase 3, catalytic subunit, gamma isoform [Source:MGI Symbol;Acc:MGI:107162] | Ppp3cc | Enriched in P15 dLGN | 1.008 | 0.014 | 226.200 |
| ENSMUSG00000041278 | tetratricopeptide repeat domain 1 [Source:MGI Symbol;Acc:MGI:1914077] | Ttc1 | Enriched in P15 dLGN | 1.006 | 0.002 | 100.100 |
| ENSMUSG00000026245 | phenylalanyl-tRNA synthetase, beta subunit [Source:MGI Symbol;Acc:MGI:1346035] | Farsb | Enriched in P15 dLGN | 1.005 | 0.001 | 92.510 |
| ENSMUSG00000005514 | cytochrome p450 oxidoreductase [Source:MGI Symbol;Acc:MGI:97744] | Por | Enriched in P15 dLGN | 1.001 | 0.000 | 268.000 |
| ENSMUSG00000007036 | abhydrolase domain containing 16A [Source:MGI Symbol;Acc:MGI:99476] | Abhd16a | Enriched in P15 dLGN | 1.001 | 0.000 | 134.000 |
| ENSMUSG00000025235 | Bardet-Biedl syndrome 4 (human) [Source:MGI Symbol;Acc:MGI:2143311] | Bbs4 | Enriched in P15 dLGN | 1.001 | 0.028 | 116.100 |
| ENSMUSG00000029185 | family with sequence similarity 114, member A1 [Source:MGI Symbol;Acc:MGI:1915553] | Fam114a1 | Enriched in P15 dLGN | 1.001 | 0.040 | 29.670 |
| ENSMUSG00000045624 | ESF1 nucleolar pre-rRNA processing protein homolog [Source:MGI Symbol;Acc:MGI:1913830] | Esf1 | Enriched in P15 dLGN | 1.001 | 0.001 | 125.500 |
| ENSMUSG00000027195 | hydroxysteroid (17-beta) dehydrogenase 12 [Source:MGI Symbol;Acc:MGI:1926967] | Hsd17b12 | Enriched in P15 dLGN | 1.000 | 0.001 | 141.000 |
| ENSMUSG00000024095 | heterogeneous nuclear ribonucleoprotein L-like [Source:MGI Symbol;Acc:MGI:1919942] | Hnrnpll | Enriched in P8 dLGN | -1.003 | 0.000 | 186.500 |
| ENSMUSG00000068615 | gap junction protein, delta 2 [Source:MGI Symbol;Acc:MGI:1334209] | Gjd2 | Enriched in P8 dLGN | -1.013 | 0.000 | 192.500 |
| ENSMUSG00000020481 | ankyrin repeat domain 36 [Source:MGI Symbol;Acc:MGI:1923639] | Ankrd36 | Enriched in P8 dLGN | -1.014 | 0.029 | 15.790 |
| ENSMUSG00000020086 | macroH2A.2 histone [Source:MGI Symbol;Acc:MGI:3037658] | Macroh2a2 | Enriched in P8 dLGN | -1.030 | 0.000 | 146.300 |
| ENSMUSG00000030671 | phosphodiesterase 3B, cGMP-inhibited [Source:MGI Symbol;Acc:MGI:1333863] | Pde3b | Enriched in P8 dLGN | -1.043 | 0.031 | 106.000 |
| ENSMUSG00000039913 | p21 (RAC1) activated kinase 5 [Source:MGI Symbol;Acc:MGI:1920334] | Pak5 | Enriched in P8 dLGN | -1.049 | 0.024 | 253.300 |
| ENSMUSG00000028312 | structural maintenance of chromosomes 2 [Source:MGI Symbol;Acc:MGI:106067] | Smc2 | Enriched in P8 dLGN | -1.051 | 0.008 | 42.300 |
| ENSMUSG00000079184 | M-phase phosphoprotein 8 [Source:MGI Symbol;Acc:MGI:1922589] | Mphosph8 | Enriched in P8 dLGN | -1.055 | 0.018 | 685.900 |
| ENSMUSG00000018589 | glycine receptor, alpha 2 subunit [Source:MGI Symbol;Acc:MGI:95748] | Glra2 | Enriched in P8 dLGN | -1.060 | 0.017 | 84.200 |
| ENSMUSG00000047238 | MAGE family member H1 [Source:MGI Symbol;Acc:MGI:1922875] | Mageh1 | Enriched in P8 dLGN | -1.067 | 0.001 | 63.860 |
| ENSMUSG00000031353 | retinoblastoma binding protein 7, chromatin remodeling factor [Source:MGI Symbol;Acc:MGI:1194910] | Rbbp7 | Enriched in P8 dLGN | -1.070 | 0.000 | 405.800 |
| ENSMUSG00000005357 | solute carrier family 1 (high affinity aspartate/glutamate transporter), member 6 [Source:MGI Symbol;Acc:MGI:1096331] | Slc1a6 | Enriched in P8 dLGN | -1.074 | 0.003 | 52.800 |
| ENSMUSG00000029108 | protocadherin 7 [Source:MGI Symbol;Acc:MGI:1860487] | Pcdh7 | Enriched in P8 dLGN | -1.110 | 0.000 | 1224.000 |
| ENSMUSG00000037148 | Rho GTPase activating protein 10 [Source:MGI Symbol;Acc:MGI:1925764] | Arhgap10 | Enriched in P8 dLGN | -1.121 | 0.008 | 198.100 |
| ENSMUSG00000046138 | RIKEN cDNA 9930021J03 gene [Source:MGI Symbol;Acc:MGI:2444398] | 9930021J03Rik | Enriched in P8 dLGN | -1.152 | 0.002 | 387.900 |
| ENSMUSG00000059518 | zinc finger, HIT domain containing 1 [Source:MGI Symbol;Acc:MGI:1917353] | Znhit1 | Enriched in P8 dLGN | -1.176 | 0.000 | 135.300 |
| ENSMUSG00000091337 | EP300 interacting inhibitor of differentiation 1 [Source:MGI Symbol;Acc:MGI:1889651] | Eid1 | Enriched in P8 dLGN | -1.176 | 0.000 | 1279.000 |
| ENSMUSG00000091405 | H4 clustered histone 14 [Source:MGI Symbol;Acc:MGI:2140113] | H4c14 | Enriched in P8 dLGN | -1.190 | 0.005 | 36.460 |
| ENSMUSG00000038457 | transmembrane protein 255B [Source:MGI Symbol;Acc:MGI:2685533] | Tmem255b | Enriched in P8 dLGN | -1.198 | 0.003 | 59.610 |
| ENSMUSG00000046058 | EP300 interacting inhibitor of differentiation 2 [Source:MGI Symbol;Acc:MGI:2681174] | Eid2 | Enriched in P8 dLGN | -1.207 | 0.001 | 137.600 |
| ENSMUSG00000020993 | trafficking protein particle complex 6B [Source:MGI Symbol;Acc:MGI:1925482] | Trappc6b | Enriched in P8 dLGN | -1.214 | 0.000 | 162.500 |

| | | | | | | |
|---|---|---|---|---|---|---|
| ENSMUSG00000059974 | neurotrimin [Source:MGI Symbol;Acc:MGI:2446259] | Ntm | Enriched in P8 dLGN | -1.227 | 0.000 | 2316.000 |
| ENSMUSG00000037341 | solute carrier family 9 (sodium/hydrogen exchanger), member 7 [Source:MGI Symbol;Acc:MGI:2444530] | Slc9a7 | Enriched in P8 dLGN | -1.234 | 0.000 | 237.700 |
| ENSMUSG00000008475 | actin related protein 2/3 complex, subunit 5 [Source:MGI Symbol;Acc:MGI:1915021] | Arpc5 | Enriched in P8 dLGN | -1.236 | 0.000 | 264.700 |
| ENSMUSG00000030265 | Kirsten rat sarcoma viral oncogene homolog [Source:MGI Symbol;Acc:MGI:96680] | Kras | Enriched in P8 dLGN | -1.243 | 0.000 | 342.300 |
| ENSMUSG00000020982 | nuclear export mediator factor [Source:MGI Symbol;Acc:MGI:1918305] | Nemf | Enriched in P8 dLGN | -1.257 | 0.002 | 270.700 |
| ENSMUSG00000063200 | nucleolar protein 7 [Source:MGI Symbol;Acc:MGI:1917328] | Nol7 | Enriched in P8 dLGN | -1.272 | 0.000 | 166.500 |
| ENSMUSG00000028793 | ring finger protein 19B [Source:MGI Symbol;Acc:MGI:1922484] | Rnf19b | Enriched in P8 dLGN | -1.294 | 0.000 | 239.700 |
| ENSMUSG00000002058 | unc-119 lipid binding chaperone [Source:MGI Symbol;Acc:MGI:1328357] | Unc119 | Enriched in P8 dLGN | -1.307 | 0.000 | 481.300 |
| ENSMUSG00000046791 | ribosomal oxygenase 1 [Source:MGI Symbol;Acc:MGI:1919202] | Riox1 | Enriched in P8 dLGN | -1.312 | 0.000 | 41.200 |
| ENSMUSG00000030685 | potassium channel tetramerisation domain containing 13 [Source:MGI Symbol;Acc:MGI:1923739] | Kctd13 | Enriched in P8 dLGN | -1.321 | 0.000 | 195.900 |
| ENSMUSG00000048720 | TBC1D12: TBC1 domain family, member 12 [Source:MGI Symbol;Acc:MGI:2384803] | Tbc1d12 | Enriched in P8 dLGN | -1.330 | 0.000 | 117.800 |
| ENSMUSG00000056305 | ubiquitin specific peptidase 39 [Source:MGI Symbol;Acc:MGI:107622] | Usp39 | Enriched in P8 dLGN | -1.352 | 0.000 | 113.700 |
| ENSMUSG00000024601 | isochorismatase domain containing 1 [Source:MGI Symbol;Acc:MGI:1913557] | Isoc1 | Enriched in P8 dLGN | -1.364 | 0.000 | 86.470 |
| ENSMUSG00000028007 | sorting nexin 7 [Source:MGI Symbol;Acc:MGI:1923811] | Snx7 | Enriched in P8 dLGN | -1.369 | 0.000 | 54.520 |
| ENSMUSG00000026082 | REV1, DNA directed polymerase [Source:MGI Symbol;Acc:MGI:1929074] | Rev1 | Enriched in P8 dLGN | -1.371 | 0.001 | 356.800 |
| ENSMUSG00000002908 | potassium intermediate/small conductance calcium-activated channel, subfamily N, member 1 [Source:MGI Symbol;Acc:MGI:1933993] | Kcnn1 | Enriched in P8 dLGN | -1.377 | 0.000 | 72.470 |
| ENSMUSG00000024270 | solute carrier family 39 (metal ion transporter), member 6 [Source:MGI Symbol;Acc:MGI:2147279] | Slc39a6 | Enriched in P8 dLGN | -1.378 | 0.000 | 212.000 |
| ENSMUSG00000091955 | predicted pseudogene 9844 [Source:MGI Symbol;Acc:MGI:3704288] | Gm9844 | Enriched in P8 dLGN | -1.389 | 0.000 | 233.800 |
| ENSMUSG00000028018 | glutathione S-transferase, C-terminal domain containing [Source:MGI Symbol;Acc:MGI:1914803] | Gstcd | Enriched in P8 dLGN | -1.393 | 0.001 | 51.360 |
| ENSMUSG00000074182 | zinc finger, HIT type 6 [Source:MGI Symbol;Acc:MGI:1916996] | Znhit6 | Enriched in P8 dLGN | -1.396 | 0.000 | 173.800 |
| ENSMUSG00000057858 | family with sequence similarity 204, member A [Source:MGI Symbol;Acc:MGI:1289174] | Fam204a | Enriched in P8 dLGN | -1.430 | 0.000 | 72.570 |
| ENSMUSG00000038607 | guanine nucleotide binding protein (G protein), gamma 10 [Source:MGI Symbol;Acc:MGI:1336169] | Gng10 | Enriched in P8 dLGN | -1.432 | 0.000 | 28.820 |
| ENSMUSG00000079523 | thymosin, beta 10 [Source:MGI Symbol;Acc:MGI:109146] | Tmsb10 | Enriched in P8 dLGN | -1.466 | 0.000 | 881.500 |
| ENSMUSG00000022940 | phosphatidylinositol glycan anchor biosynthesis, class P [Source:MGI Symbol;Acc:MGI:1860433] | Pigp | Enriched in P8 dLGN | -1.493 | 0.002 | 39.520 |
| ENSMUSG00000021377 | DEK proto-oncogene (DNA binding) [Source:MGI Symbol;Acc:MGI:1926209] | Dek | Enriched in P8 dLGN | -1.496 | 0.000 | 361.900 |
| ENSMUSG00000031838 | interferon gamma inducible protein 30 [Source:MGI Symbol;Acc:MGI:2137648] | Ifi30 | Enriched in P8 dLGN | -1.520 | 0.003 | 15.080 |
| ENSMUSG00000030137 | tubulin, alpha 8 [Source:MGI Symbol;Acc:MGI:1858225] | Tuba8 | Enriched in P8 dLGN | -1.522 | 0.007 | 18.260 |
| ENSMUSG00000029761 | caldesmon 1 [Source:MGI Symbol;Acc:MGI:88250] | Cald1 | Enriched in P8 dLGN | -1.526 | 0.000 | 209.000 |
| ENSMUSG00000035228 | coiled-coil domain containing 106 [Source:MGI Symbol;Acc:MGI:2385900] | Ccdc106 | Enriched in P8 dLGN | -1.541 | 0.000 | 90.050 |
| ENSMUSG00000029833 | tripartite motif-containing 24 [Source:MGI Symbol;Acc:MGI:109275] | Trim24 | Enriched in P8 dLGN | -1.547 | 0.000 | 198.700 |
| ENSMUSG00000030876 | methyltransferase like 9 [Source:MGI Symbol;Acc:MGI:1914862] | Mettl9 | Enriched in P8 dLGN | -1.550 | 0.000 | 224.200 |
| ENSMUSG00000021549 | RAS p21 protein activator 1 [Source:MGI Symbol;Acc:MGI:97860] | Rasa1 | Enriched in P8 dLGN | -1.585 | 0.000 | 293.700 |
| ENSMUSG00000039715 | dynein 2 intermediate chain 2 [Source:MGI Symbol;Acc:MGI:1919070] | Dync2i2 | Enriched in P8 dLGN | -1.591 | 0.000 | 95.480 |
| ENSMUSG00000002997 | protein kinase, cAMP dependent regulatory, type II beta [Source:MGI Symbol;Acc:MGI:97760] | Prkar2b | Enriched in P8 dLGN | -1.616 | 0.000 | 198.800 |
| ENSMUSG00000028033 | potassium voltage-gated channel, subfamily Q, member 5 [Source:MGI Symbol;Acc:MGI:1924937] | Kcnq5 | Enriched in P8 dLGN | -1.618 | 0.000 | 180.800 |
| ENSMUSG00000029814 | insulin-like growth factor 2 mRNA binding protein 3 [Source:MGI Symbol;Acc:MGI:1890359] | Igf2bp3 | Enriched in P8 dLGN | -1.643 | 0.001 | 57.120 |
| ENSMUSG00000045763 | brain abundant, membrane attached signal protein 1 [Source:MGI Symbol;Acc:MGI:1917600] | Basp1 | Enriched in P8 dLGN | -1.646 | 0.000 | 3099.000 |
| ENSMUSG00000020882 | calcium channel, voltage-dependent, beta 1 subunit [Source:MGI Symbol;Acc:MGI:102522] | Cacnb1 | Enriched in P8 dLGN | -1.708 | 0.000 | 246.500 |
| ENSMUSG00000027104 | activating transcription factor 2 [Source:MGI Symbol;Acc:MGI:109349] | Atf2 | Enriched in P8 dLGN | -1.728 | 0.000 | 703.500 |
| ENSMUSG00000036087 | SLAIN motif family, member 2 [Source:MGI Symbol;Acc:MGI:1923241] | Slain2 | Enriched in P8 dLGN | -1.755 | 0.000 | 209.600 |
| ENSMUSG00000019464 | prostaglandin E receptor 1 (subtype EP1) [Source:MGI Symbol;Acc:MGI:97793] | Ptger1 | Enriched in P8 dLGN | -1.778 | 0.003 | 8.868 |
| ENSMUSG00000006498 | polypyrimidine tract binding protein 1 [Source:MGI Symbol;Acc:MGI:97791] | Ptbp1 | Enriched in P8 dLGN | -1.800 | 0.000 | 112.300 |
| ENSMUSG00000019320 | NADPH oxidase organizer 1 [Source:MGI Symbol;Acc:MGI:1919143] | Noxo1 | Enriched in P8 dLGN | -1.816 | 0.009 | 10.810 |
| ENSMUSG00000010067 | Ras association (RalGDS/AF-6) domain family member 1 [Source:MGI Symbol;Acc:MGI:1928386] | Rassf1 | Enriched in P8 dLGN | -1.841 | 0.000 | 43.500 |
| ENSMUSG00000039166 | A kinase (PRKA) anchor protein 7 [Source:MGI Symbol;Acc:MGI:1859150] | Akap7 | Enriched in P8 dLGN | -1.883 | 0.000 | 348.800 |
| ENSMUSG00000096141 | dynein, axonemal, heavy chain 7A [Source:MGI Symbol;Acc:MGI:2685838] | Dnah7a | Enriched in P8 dLGN | -1.891 | 0.038 | 25.060 |
| ENSMUSG00000036377 | capping protein inhibiting regulator of actin [Source:MGI Symbol;Acc:MGI:2444817] | Cracd | Enriched in P8 dLGN | -1.921 | 0.000 | 1779.000 |
| ENSMUSG00000020107 | anaphase promoting complex subunit 16 [Source:MGI Symbol;Acc:MGI:1289325] | Anapc16 | Enriched in P8 dLGN | -1.955 | 0.000 | 134.300 |
| ENSMUSG00000029346 | SRR1 domain containing [Source:MGI Symbol;Acc:MGI:1917368] | Srrd | Enriched in P8 dLGN | -2.026 | 0.000 | 45.970 |
| ENSMUSG00000056310 | tRNA-yW synthesizing protein 1 homolog (S. cerevisiae) [Source:MGI Symbol;Acc:MGI:2141161] | Tyw1 | Enriched in P8 dLGN | -2.060 | 0.000 | 150.100 |
| ENSMUSG00000020814 | matrix-remodelling associated 7 [Source:MGI Symbol;Acc:MGI:1914872] | Mxra7 | Enriched in P8 dLGN | -2.077 | 0.000 | 522.100 |
| ENSMUSG00000102189 | predicted gene, 37194 [Source:MGI Symbol;Acc:MGI:5610422] | Gm37194 | Enriched in P8 dLGN | -2.168 | 0.000 | 32.700 |
| ENSMUSG00000052372 | interleukin 1 receptor accessory protein-like 1 [Source:MGI Symbol;Acc:MGI:2687319] | Il1rapl1 | Enriched in P8 dLGN | -2.249 | 0.000 | 1897.000 |
| ENSMUSG00000018169 | MFNG O-fucosylpeptide 3-beta-N-acetylglucosaminyltransferase [Source:MGI Symbol;Acc:MGI:1095404] | Mfng | Enriched in P8 dLGN | -2.384 | 0.000 | 18.920 |
| ENSMUSG00000073406 | histocompatibility 2, blastocyst [Source:MGI Symbol;Acc:MGI:892004] | H2-Bl | Enriched in P8 dLGN | -2.385 | 0.001 | 18.450 |
| ENSMUSG00000000560 | gamma-aminobutyric acid (GABA) A receptor, subunit alpha 2 [Source:MGI Symbol;Acc:MGI:95614] | Gabra2 | Enriched in P8 dLGN | -2.388 | 0.000 | 394.200 |
| ENSMUSG00000093985 | predicted gene 10406 [Source:MGI Symbol;Acc:MGI:3711272] | Gm10406 | Enriched in P8 dLGN | -2.408 | 0.011 | 6.055 |
| ENSMUSG00000021892 | SH3-domain binding protein 5 (BTK-associated) [Source:MGI Symbol;Acc:MGI:1344391] | Sh3bp5 | Enriched in P8 dLGN | -2.539 | 0.000 | 251.200 |
| ENSMUSG00000040489 | SRY (sex determining region Y)-box 30 [Source:MGI Symbol;Acc:MGI:1341157] | Sox30 | Enriched in P8 dLGN | -2.556 | 0.003 | 11.690 |
| ENSMUSG00000037013 | SS18, nBAF chromatin remodeling complex subunit [Source:MGI Symbol;Acc:MGI:107708] | Ss18 | Enriched in P8 dLGN | -2.562 | 0.000 | 158.700 |
| ENSMUSG00000041688 | angiomotin [Source:MGI Symbol;Acc:MGI:108440] | Amot | Enriched in P8 dLGN | -2.571 | 0.000 | 327.400 |
| ENSMUSG00000029155 | spermatogenesis associated 18 [Source:MGI Symbol;Acc:MGI:1920722] | Spata18 | Enriched in P8 dLGN | -2.642 | 0.018 | 10.420 |
| ENSMUSG00000041144 | dynein, axonemal, heavy chain 7B [Source:MGI Symbol;Acc:MGI:2684953] | Dnah7b | Enriched in P8 dLGN | -2.674 | 0.014 | 449.400 |

| | | | | | | |
|---|---|---|---|---|---|---|
| ENSMUSG00000028461 | coiled-coil domain containing 107 [Source:MGI Symbol;Acc:MGI:1913423] | Ccdc107 | Enriched in P8 dLGN | -2.730 | 0.005 | 309.500 |
| ENSMUSG00000050248 | EvC ciliary complex subunit 2 [Source:MGI Symbol;Acc:MGI:1915775] | Evc2 | Enriched in P8 dLGN | -2.988 | 0.000 | 56.130 |
| ENSMUSG00000027344 | fibrous sheath-interacting protein 1 [Source:MGI Symbol;Acc:MGI:1918563] | Fsip1 | Enriched in P8 dLGN | -3.021 | 0.000 | 17.620 |
| ENSMUSG00000020000 | monooxygenase, DBH-like 1 [Source:MGI Symbol;Acc:MGI:1921582] | Moxd1 | Enriched in P8 dLGN | -3.648 | 0.000 | 20.890 |
| ENSMUSG00000051864 | TBC1 domain family, member 22a [Source:MGI Symbol;Acc:MGI:1289265] | Tbc1d22a | Enriched in P8 dLGN | -4.397 | 0.000 | 351.400 |
| ENSMUSG00000034883 | leucine rich repeat protein 1 [Source:MGI Symbol;Acc:MGI:1916956] | Lrr1 | Enriched in P8 dLGN | -8.418 | 0.000 | 14.150 |

**Contrast numerator: P15_Het_retina. Contrast denominator: P8_Het_retina.**

| ensembl_gene_id | description | mgi_symbol | Significance | deseq_logfc | deseq_adjp | deseq_basemean |
|---|---|---|---|---|---|---|
| ENSMUSG00000027301 | oxytocin [Source:MGI Symbol;Acc:MGI:97453] | Oxt | Enriched in P15 retina | 15.080 | 0.000 | 22.270 |
| ENSMUSG00000072476 | predicted pseudogene 9008 [Source:MGI Symbol;Acc:MGI:3644000] | Gm9008 | Enriched in P15 retina | 6.762 | 0.000 | 8.190 |
| ENSMUSG00000030730 | ATPase, Ca++ transporting, cardiac muscle, fast twitch 1 [Source:MGI Symbol;Acc:MGI:105058] | Atp2a1 | Enriched in P15 retina | 5.852 | 0.000 | 39.300 |
| ENSMUSG00000086962 | predicted gene 12248 [Source:MGI Symbol;Acc:MGI:3651124] | Gm12248 | Enriched in P15 retina | 5.046 | 0.001 | 8.028 |
| ENSMUSG00000007594 | hyaluronan and proteoglycan link protein 4 [Source:MGI Symbol;Acc:MGI:2679531] | Hapln4 | Enriched in P15 retina | 3.966 | 0.000 | 156.800 |
| ENSMUSG00000035296 | sarcoglycan, gamma (dystrophin-associated glycoprotein) [Source:MGI Symbol;Acc:MGI:1346524] | Sgcg | Enriched in P15 retina | 3.275 | 0.000 | 9.668 |
| ENSMUSG00000033196 | myosin, heavy polypeptide 2, skeletal muscle, adult [Source:MGI Symbol;Acc:MGI:1339710] | Myh2 | Enriched in P15 retina | 3.121 | 0.006 | 4.066 |
| ENSMUSG00000050138 | potassium channel, subfamily K, member 12 [Source:MGI Symbol;Acc:MGI:2684043] | Kcnk12 | Enriched in P15 retina | 3.084 | 0.000 | 38.230 |
| ENSMUSG00000046480 | sodium channel, type IV, beta [Source:MGI Symbol;Acc:MGI:2687406] | Scn4b | Enriched in P15 retina | 3.070 | 0.000 | 167.600 |
| ENSMUSG00000005716 | parvalbumin [Source:MGI Symbol;Acc:MGI:97821] | Pvalb | Enriched in P15 retina | 3.050 | 0.000 | 26.700 |
| ENSMUSG00000042073 | abhydrolase domain containing 14b [Source:MGI Symbol;Acc:MGI:1923741] | Abhd14b | Enriched in P15 retina | 2.825 | 0.000 | 38.480 |
| ENSMUSG00000072572 | solute carrier family 39 (zinc transporter), member 2 [Source:MGI Symbol;Acc:MGI:2684326] | Slc39a2 | Enriched in P15 retina | 2.364 | 0.003 | 19.900 |
| ENSMUSG00000040489 | SRY (sex determining region Y)-box 30 [Source:MGI Symbol;Acc:MGI:1341157] | Sox30 | Enriched in P15 retina | 2.306 | 0.003 | 11.690 |
| ENSMUSG00000018470 | potassium voltage-gated channel, shaker-related subfamily, beta member 3 [Source:MGI Symbol;Acc:MGI:1336208] | Kcnab3 | Enriched in P15 retina | 2.275 | 0.000 | 53.860 |
| ENSMUSG00000030337 | vesicle-associated membrane protein 1 [Source:MGI Symbol;Acc:MGI:1313276] | Vamp1 | Enriched in P15 retina | 2.274 | 0.000 | 282.400 |
| ENSMUSG00000015354 | procollagen C-endopeptidase enhancer 2 [Source:MGI Symbol;Acc:MGI:1923727] | Pcolce2 | Enriched in P15 retina | 2.226 | 0.000 | 23.560 |
| ENSMUSG00000029304 | secreted phosphoprotein 1 [Source:MGI Symbol;Acc:MGI:98389] | Spp1 | Enriched in P15 retina | 2.210 | 0.023 | 33.230 |
| ENSMUSG00000029330 | CDP-diacylglycerol synthase 1 [Source:MGI Symbol;Acc:MGI:1921846] | Cds1 | Enriched in P15 retina | 2.188 | 0.000 | 337.700 |
| ENSMUSG00000020732 | RAB37, member RAS oncogene family [Source:MGI Symbol;Acc:MGI:1929945] | Rab37 | Enriched in P15 retina | 2.113 | 0.000 | 50.930 |
| ENSMUSG00000032908 | sphingosine-1-phosphate phosphatase 2 [Source:MGI Symbol;Acc:MGI:3589109] | Sgpp2 | Enriched in P15 retina | 2.109 | 0.000 | 115.400 |
| ENSMUSG00000061462 | obscurin, cytoskeletal calmodulin and titin-interacting RhoGEF [Source:MGI Symbol;Acc:MGI:2681862] | Obscn | Enriched in P15 retina | 2.094 | 0.000 | 66.020 |
| ENSMUSG00000020014 | cilia and flagella associated protein 54 [Source:MGI Symbol;Acc:MGI:1922208] | Cfap54 | Enriched in P15 retina | 1.955 | 0.000 | 91.990 |
| ENSMUSG00000023484 | peripherin [Source:MGI Symbol;Acc:MGI:97774] | Prph | Enriched in P15 retina | 1.902 | 0.040 | 64.100 |
| ENSMUSG00000036585 | fibroblast growth factor 1 [Source:MGI Symbol;Acc:MGI:95515] | Fgf1 | Enriched in P15 retina | 1.851 | 0.000 | 148.800 |
| ENSMUSG00000056665 | thioesterase superfamily member 6 [Source:MGI Symbol;Acc:MGI:1925301] | Them6 | Enriched in P15 retina | 1.816 | 0.000 | 58.010 |
| ENSMUSG00000051747 | titin [Source:MGI Symbol;Acc:MGI:98864] | Ttn | Enriched in P15 retina | 1.718 | 0.000 | 194.400 |
| ENSMUSG00000038457 | transmembrane protein 255B [Source:MGI Symbol;Acc:MGI:2685533] | Tmem255b | Enriched in P15 retina | 1.694 | 0.000 | 59.610 |
| ENSMUSG00000026414 | troponin T2, cardiac [Source:MGI Symbol;Acc:MGI:104597] | Tnnt2 | Enriched in P15 retina | 1.688 | 0.022 | 15.870 |
| ENSMUSG00000032599 | inositol hexaphosphate kinase 2 [Source:MGI Symbol;Acc:MGI:1923750] | Ip6k2 | Enriched in P15 retina | 1.679 | 0.000 | 280.500 |
| ENSMUSG00000035458 | troponin I, cardiac 3 [Source:MGI Symbol;Acc:MGI:98783] | Tnni3 | Enriched in P15 retina | 1.677 | 0.013 | 5.883 |
| ENSMUSG00000019194 | sodium channel, voltage-gated, type I, beta [Source:MGI Symbol;Acc:MGI:98247] | Scn1b | Enriched in P15 retina | 1.599 | 0.000 | 470.000 |
| ENSMUSG00000045968 | transmembrane epididymal family member 2 [Source:MGI Symbol;Acc:MGI:1923273] | Teddm2 | Enriched in P15 retina | 1.575 | 0.000 | 15.550 |
| ENSMUSG00000044005 | glutaminase 2 (liver, mitochondrial) [Source:MGI Symbol;Acc:MGI:2143539] | Gls2 | Enriched in P15 retina | 1.571 | 0.000 | 17.760 |
| ENSMUSG00000033208 | S100 protein, beta polypeptide, neural [Source:MGI Symbol;Acc:MGI:98217] | S100b | Enriched in P15 retina | 1.515 | 0.001 | 86.390 |
| ENSMUSG00000026672 | optineurin [Source:MGI Symbol;Acc:MGI:1918898] | Optn | Enriched in P15 retina | 1.371 | 0.000 | 212.900 |
| ENSMUSG00000026669 | minichromosome maintenance 10 replication initiation factor [Source:MGI Symbol;Acc:MGI:1917274] | Mcm10 | Enriched in P15 retina | 1.359 | 0.021 | 5.502 |
| ENSMUSG00000033595 | leucine-rich repeat LGI family, member 3 [Source:MGI Symbol;Acc:MGI:2182619] | Lgi3 | Enriched in P15 retina | 1.334 | 0.000 | 185.500 |
| ENSMUSG00000022594 | Ly6/neurotoxin 1 [Source:MGI Symbol;Acc:MGI:1345180] | Lynx1 | Enriched in P15 retina | 1.223 | 0.001 | 1133.000 |
| ENSMUSG00000001520 | nuclear receptor interacting protein 2 [Source:MGI Symbol;Acc:MGI:1891884] | Nrip2 | Enriched in P15 retina | 1.203 | 0.004 | 17.020 |
| ENSMUSG00000096696 | zinc finger protein 960 [Source:MGI Symbol;Acc:MGI:3052731] | Zfp960 | Enriched in P15 retina | 1.199 | 0.045 | 4.000 |
| ENSMUSG00000040557 | methyltransferase like 27 [Source:MGI Symbol;Acc:MGI:1933146] | Mettl27 | Enriched in P15 retina | 1.198 | 0.000 | 18.670 |
| ENSMUSG00000033705 | START domain containing 9 [Source:MGI Symbol;Acc:MGI:3045258] | Stard9 | Enriched in P15 retina | 1.181 | 0.000 | 146.400 |
| ENSMUSG00000032558 | nephronophthisis 3 (adolescent) [Source:MGI Symbol;Acc:MGI:1921275] | Nphp3 | Enriched in P15 retina | 1.161 | 0.024 | 34.650 |
| ENSMUSG00000041556 | F-box protein 2 [Source:MGI Symbol;Acc:MGI:2446216] | Fbxo2 | Enriched in P15 retina | 1.150 | 0.010 | 92.090 |
| ENSMUSG00000002032 | transmembrane protein 25 [Source:MGI Symbol;Acc:MGI:1918937] | Tmem25 | Enriched in P15 retina | 1.108 | 0.000 | 70.330 |
| ENSMUSG00000068099 | small integral membrane protein 45 [Source:MGI Symbol;Acc:MGI:1923755] | Smim45 | Enriched in P15 retina | 1.108 | 0.000 | 83.120 |
| ENSMUSG00000027014 | CWC22 spliceosome-associated protein [Source:MGI Symbol;Acc:MGI:2136773] | Cwc22 | Enriched in P15 retina | 1.054 | 0.032 | 206.100 |
| ENSMUSG00000021728 | embigin [Source:MGI Symbol;Acc:MGI:95321] | Emb | Enriched in P15 retina | 1.052 | 0.004 | 88.850 |
| ENSMUSG00000056596 | TMF1-regulated nuclear protein 1 [Source:MGI Symbol;Acc:MGI:1916789] | Trnp1 | Enriched in P15 retina | 1.002 | 0.001 | 640.500 |
| ENSMUSG00000044576 | GRB2 associated regulator of MAPK1 subtype 2 [Source:MGI Symbol;Acc:MGI:2685290] | Garem2 | Enriched in P8 retina | -1.010 | 0.006 | 62.670 |
| ENSMUSG00000021303 | guanine nucleotide binding protein (G protein), gamma 4 [Source:MGI Symbol;Acc:MGI:102703] | Gng4 | Enriched in P8 retina | -1.011 | 0.018 | 229.600 |
| ENSMUSG00000033706 | SET and MYND domain containing 5 [Source:MGI Symbol;Acc:MGI:108048] | Smyd5 | Enriched in P8 retina | -1.012 | 0.001 | 48.580 |
| ENSMUSG00000025743 | syndecan 3 [Source:MGI Symbol;Acc:MGI:1349163] | Sdc3 | Enriched in P8 retina | -1.015 | 0.000 | 1949.000 |
| ENSMUSG00000011589 | fibronectin type 3 and SPRY domain-containing protein [Source:MGI Symbol;Acc:MGI:1934858] | Fsd1 | Enriched in P8 retina | -1.019 | 0.004 | 96.920 |
| ENSMUSG00000046321 | heparan sulfate (glucosamine) 3-O-sulfotransferase 2 [Source:MGI Symbol;Acc:MGI:1333802] | Hs3st2 | Enriched in P8 retina | -1.032 | 0.005 | 41.560 |
| ENSMUSG00000041774 | YdjC homolog (bacterial) [Source:MGI Symbol;Acc:MGI:1916351] | Ydjc | Enriched in P8 retina | -1.033 | 0.000 | 46.050 |
| ENSMUSG00000061702 | transmembrane protein 91 [Source:MGI Symbol;Acc:MGI:2443589] | Tmem91 | Enriched in P8 retina | -1.049 | 0.034 | 41.880 |
| ENSMUSG00000037990 | SH3 domain containing ring finger 3 [Source:MGI Symbol;Acc:MGI:2444637] | Sh3rf3 | Enriched in P8 retina | -1.051 | 0.000 | 202.200 |
| ENSMUSG00000045763 | brain abundant, membrane attached signal protein 1 [Source:MGI Symbol;Acc:MGI:1917600] | Basp1 | Enriched in P8 retina | -1.055 | 0.000 | 3099.000 |
| ENSMUSG00000024664 | fatty acid desaturase 3 [Source:MGI Symbol;Acc:MGI:1928740] | Fads3 | Enriched in P8 retina | -1.056 | 0.000 | 75.230 |
| ENSMUSG00000034226 | ras homolog family member V [Source:MGI Symbol;Acc:MGI:2444227] | Rhov | Enriched in P8 retina | -1.060 | 0.010 | 54.650 |

| | | | | | | |
|---|---|---|---|---|---|---|
| ENSMUSG00000025889 | synuclein, alpha [Source:MGI Symbol;Acc:MGI:1277151] | Snca | Enriched in P8 retina | -1.067 | 0.006 | 118.200 |
| ENSMUSG00000033585 | necdin, MAGE family member [Source:MGI Symbol;Acc:MGI:97290] | Ndn | Enriched in P8 retina | -1.071 | 0.004 | 549.000 |
| ENSMUSG00000034037 | FYVE, RhoGEF and PH domain containing 5 [Source:MGI Symbol;Acc:MGI:2443369] | Fgd5 | Enriched in P8 retina | -1.072 | 0.007 | 83.070 |
| ENSMUSG00000032740 | coiled coil domain containing 88A [Source:MGI Symbol;Acc:MGI:1925177] | Ccdc88a | Enriched in P8 retina | -1.082 | 0.000 | 614.100 |
| ENSMUSG00000027500 | stathmin-like 2 [Source:MGI Symbol;Acc:MGI:98241] | Stmn2 | Enriched in P8 retina | -1.087 | 0.003 | 1265.000 |
| ENSMUSG00000031841 | cadherin 13 [Source:MGI Symbol;Acc:MGI:99551] | Cdh13 | Enriched in P8 retina | -1.094 | 0.009 | 448.300 |
| ENSMUSG00000038860 | GTPase activating RANGAP domain-like 3 [Source:MGI Symbol;Acc:MGI:2139309] | Garnl3 | Enriched in P8 retina | -1.094 | 0.001 | 238.500 |
| ENSMUSG00000028755 | cytidine deaminase [Source:MGI Symbol;Acc:MGI:1919519] | Cda | Enriched in P8 retina | -1.097 | 0.034 | 14.430 |
| ENSMUSG00000070644 | ethanolamine kinase 2 [Source:MGI Symbol;Acc:MGI:2443760] | Etnk2 | Enriched in P8 retina | -1.103 | 0.017 | 35.130 |
| ENSMUSG00000030376 | solute carrier family 8 (sodium/calcium exchanger), member 2 [Source:MGI Symbol;Acc:MGI:107996] | Slc8a2 | Enriched in P8 retina | -1.105 | 0.000 | 330.300 |
| ENSMUSG00000041362 | shootin 1 [Source:MGI Symbol;Acc:MGI:1918903] | Shtn1 | Enriched in P8 retina | -1.108 | 0.000 | 302.500 |
| ENSMUSG00000030303 | fatty acyl CoA reductase 2 [Source:MGI Symbol;Acc:MGI:2687035] | Far2 | Enriched in P8 retina | -1.110 | 0.025 | 96.510 |
| ENSMUSG00000038916 | SOGA family member 3 [Source:MGI Symbol;Acc:MGI:1914662] | Soga3 | Enriched in P8 retina | -1.116 | 0.000 | 604.700 |
| ENSMUSG00000029603 | deltex 1, E3 ubiquitin ligase [Source:MGI Symbol;Acc:MGI:1352744] | Dtx1 | Enriched in P8 retina | -1.122 | 0.000 | 412.900 |
| ENSMUSG00000066705 | FXYD domain-containing ion transport regulator 6 [Source:MGI Symbol;Acc:MGI:1890226] | Fxyd6 | Enriched in P8 retina | -1.130 | 0.018 | 380.300 |
| ENSMUSG00000000489 | platelet derived growth factor, B polypeptide [Source:MGI Symbol;Acc:MGI:97528] | Pdgfb | Enriched in P8 retina | -1.149 | 0.001 | 70.920 |
| ENSMUSG00000031906 | sphingomyelin phosphodiesterase 3, neutral [Source:MGI Symbol;Acc:MGI:1927578] | Smpd3 | Enriched in P8 retina | -1.154 | 0.000 | 615.600 |
| ENSMUSG00000031391 | L1 cell adhesion molecule [Source:MGI Symbol;Acc:MGI:96721] | L1cam | Enriched in P8 retina | -1.172 | 0.006 | 803.200 |
| ENSMUSG00000020230 | protein arginine N-methyltransferase 2 [Source:MGI Symbol;Acc:MGI:1316652] | Prmt2 | Enriched in P8 retina | -1.181 | 0.000 | 214.300 |
| ENSMUSG00000074793 | heat shock protein 12B [Source:MGI Symbol;Acc:MGI:1919880] | Hspa12b | Enriched in P8 retina | -1.190 | 0.002 | 37.370 |
| ENSMUSG00000030782 | transforming growth factor beta 1 induced transcript 1 [Source:MGI Symbol;Acc:MGI:102784] | Tgfb1i1 | Enriched in P8 retina | -1.193 | 0.000 | 24.560 |
| ENSMUSG00000027330 | cell division cycle 25B [Source:MGI Symbol;Acc:MGI:99701] | Cdc25b | Enriched in P8 retina | -1.201 | 0.002 | 31.550 |
| ENSMUSG00000047963 | starch binding domain 1 [Source:MGI Symbol;Acc:MGI:1261768] | Stbd1 | Enriched in P8 retina | -1.204 | 0.015 | 7.985 |
| ENSMUSG00000048070 | phosphoinositide-interacting regulator of transient receptor potential channels [Source:MGI Symbol;Acc:MGI:2443635 | Pirt | Enriched in P8 retina | -1.219 | 0.022 | 10.020 |
| ENSMUSG00000041959 | S100 calcium binding protein A10 (calpactin) [Source:MGI Symbol;Acc:MGI:1339468] | S100a10 | Enriched in P8 retina | -1.232 | 0.040 | 41.510 |
| ENSMUSG00000051652 | leucine rich repeat containing 3 [Source:MGI Symbol;Acc:MGI:2447899] | Lrrc3 | Enriched in P8 retina | -1.233 | 0.000 | 141.000 |
| ENSMUSG00000033082 | C-type lectin domain family 1, member a [Source:MGI Symbol;Acc:MGI:2444151] | Clec1a | Enriched in P8 retina | -1.250 | 0.016 | 6.168 |
| ENSMUSG00000039004 | bone morphogenetic protein 6 [Source:MGI Symbol;Acc:MGI:88182] | Bmp6 | Enriched in P8 retina | -1.265 | 0.007 | 58.770 |
| ENSMUSG00000069227 | G protein-regulated inducer of neurite outgrowth 1 [Source:MGI Symbol;Acc:MGI:1349455] | Gprin1 | Enriched in P8 retina | -1.278 | 0.000 | 480.700 |
| ENSMUSG00000022216 | proteasome (prosome, macropain) activator subunit 1 (PA28 alpha) [Source:MGI Symbol;Acc:MGI:1096367] | Psme1 | Enriched in P8 retina | -1.282 | 0.003 | 100.300 |
| ENSMUSG00000052301 | double C2, alpha [Source:MGI Symbol;Acc:MGI:109446] | Doc2a | Enriched in P8 retina | -1.295 | 0.012 | 56.860 |
| ENSMUSG00000022861 | diacylglycerol kinase, gamma [Source:MGI Symbol;Acc:MGI:105060] | Dgkg | Enriched in P8 retina | -1.307 | 0.003 | 135.000 |
| ENSMUSG00000001034 | mitogen-activated protein kinase 7 [Source:MGI Symbol;Acc:MGI:1346347] | Mapk7 | Enriched in P8 retina | -1.309 | 0.000 | 104.100 |
| ENSMUSG00000062380 | tubulin, beta 3 class III [Source:MGI Symbol;Acc:MGI:107813] | Tubb3 | Enriched in P8 retina | -1.309 | 0.026 | 1342.000 |
| ENSMUSG00000006307 | lysine (K)-specific methyltransferase 2B [Source:MGI Symbol;Acc:MGI:109565] | Kmt2b | Enriched in P8 retina | -1.325 | 0.000 | 848.200 |
| ENSMUSG00000024287 | THO complex 1 [Source:MGI Symbol;Acc:MGI:1919668] | Thoc1 | Enriched in P8 retina | -1.360 | 0.000 | 109.900 |
| ENSMUSG00000030528 | Bloom syndrome, RecQ like helicase [Source:MGI Symbol;Acc:MGI:1328362] | Blm | Enriched in P8 retina | -1.371 | 0.000 | 50.260 |
| ENSMUSG00000032946 | RAS, guanyl releasing protein 2 [Source:MGI Symbol;Acc:MGI:1333849] | Rasgrp2 | Enriched in P8 retina | -1.378 | 0.000 | 149.800 |
| ENSMUSG00000029161 | cell growth regulator with EF hand domain 1 [Source:MGI Symbol;Acc:MGI:1915817] | Cgref1 | Enriched in P8 retina | -1.385 | 0.000 | 115.300 |
| ENSMUSG00000018507 | transient receptor potential cation channel, subfamily V, member 2 [Source:MGI Symbol;Acc:MGI:1341836] | Trpv2 | Enriched in P8 retina | -1.386 | 0.000 | 57.510 |
| ENSMUSG00000034685 | family with sequence similarity 171, member A2 [Source:MGI Symbol;Acc:MGI:2448496] | Fam171a2 | Enriched in P8 retina | -1.389 | 0.000 | 403.300 |
| ENSMUSG00000030707 | coronin, actin binding protein 1A [Source:MGI Symbol;Acc:MGI:1345961] | Coro1a | Enriched in P8 retina | -1.404 | 0.001 | 102.700 |
| ENSMUSG00000036913 | tripartite motif-containing 67 [Source:MGI Symbol;Acc:MGI:3045323] | Trim67 | Enriched in P8 retina | -1.412 | 0.000 | 572.300 |
| ENSMUSG00000017897 | EYA transcriptional coactivator and phosphatase 2 [Source:MGI Symbol;Acc:MGI:109341] | Eya2 | Enriched in P8 retina | -1.413 | 0.000 | 21.410 |
| ENSMUSG00000024907 | galanin and GMAP prepropeptide [Source:MGI Symbol;Acc:MGI:95637] | Gal | Enriched in P8 retina | -1.413 | 0.008 | 30.000 |
| ENSMUSG00000021534 | RIKEN cDNA 1700001L19 gene [Source:MGI Symbol;Acc:MGI:1916565] | 1700001L19Rik | Enriched in P8 retina | -1.416 | 0.016 | 10.840 |
| ENSMUSG00000035561 | aldehyde dehydrogenase 1 family, member B1 [Source:MGI Symbol;Acc:MGI:1919785] | Aldh1b1 | Enriched in P8 retina | -1.418 | 0.004 | 22.800 |
| ENSMUSG00000022577 | lymphocyte antigen 6 complex, locus H [Source:MGI Symbol;Acc:MGI:1346030] | Ly6h | Enriched in P8 retina | -1.428 | 0.000 | 411.300 |
| ENSMUSG00000002458 | regulator of G-protein signaling 19 [Source:MGI Symbol;Acc:MGI:1915153] | Rgs19 | Enriched in P8 retina | -1.434 | 0.002 | 28.610 |
| ENSMUSG00000019066 | RAB3D, member RAS oncogene family [Source:MGI Symbol;Acc:MGI:97844] | Rab3d | Enriched in P8 retina | -1.458 | 0.000 | 46.080 |
| ENSMUSG00000025930 | musculin [Source:MGI Symbol;Acc:MGI:1333884] | Msc | Enriched in P8 retina | -1.459 | 0.031 | 4.237 |
| ENSMUSG00000102189 | predicted gene, 37194 [Source:MGI Symbol;Acc:MGI:5610422] | Gm37194 | Enriched in P8 retina | -1.480 | 0.040 | 32.700 |
| ENSMUSG00000021294 | kinesin family member 26A [Source:MGI Symbol;Acc:MGI:2447072] | Kif26a | Enriched in P8 retina | -1.482 | 0.000 | 99.430 |
| ENSMUSG00000035431 | somatostatin receptor 1 [Source:MGI Symbol;Acc:MGI:98327] | Sstr1 | Enriched in P8 retina | -1.491 | 0.029 | 18.080 |
| ENSMUSG00000005357 | solute carrier family 1 (high affinity aspartate/glutamate transporter), member 6 [Source:MGI Symbol;Acc:MGI:10963: | Slc1a6 | Enriched in P8 retina | -1.494 | 0.000 | 52.800 |
| ENSMUSG00000029503 | purinergic receptor P2X, ligand-gated ion channel, 2 [Source:MGI Symbol;Acc:MGI:2665170] | P2rx2 | Enriched in P8 retina | -1.496 | 0.033 | 6.203 |
| ENSMUSG00000047428 | delta like non-canonical Notch ligand 2 [Source:MGI Symbol;Acc:MGI:2146838] | Dlk2 | Enriched in P8 retina | -1.499 | 0.000 | 45.680 |
| ENSMUSG00000002997 | protein kinase, cAMP dependent regulatory, type II beta [Source:MGI Symbol;Acc:MGI:97760] | Prkar2b | Enriched in P8 retina | -1.500 | 0.000 | 198.800 |
| ENSMUSG00000034115 | sodium channel, voltage-gated, type XI, alpha [Source:MGI Symbol;Acc:MGI:1345149] | Scn11a | Enriched in P8 retina | -1.513 | 0.001 | 5.567 |
| ENSMUSG00000021948 | protein kinase C, delta [Source:MGI Symbol;Acc:MGI:97598] | Prkcd | Enriched in P8 retina | -1.522 | 0.005 | 727.900 |
| ENSMUSG00000001525 | tubulin, beta 5 class I [Source:MGI Symbol;Acc:MGI:107812] | Tubb5 | Enriched in P8 retina | -1.540 | 0.000 | 2392.000 |
| ENSMUSG00000033854 | potassium channel, subfamily K, member 10 [Source:MGI Symbol;Acc:MGI:1919508] | Kcnk10 | Enriched in P8 retina | -1.554 | 0.000 | 191.600 |
| ENSMUSG00000007207 | syntaxin 1A (brain) [Source:MGI Symbol;Acc:MGI:109355] | Stx1a | Enriched in P8 retina | -1.571 | 0.000 | 278.700 |

| | | | | | | |
|---|---|---|---|---|---|---|
| ENSMUSG00000028339 | collagen, type XV, alpha 1 [Source:MGI Symbol;Acc:MGI:88449] | Col15a1 | Enriched in P8 retina | -1.573 | 0.001 | 38.430 |
| ENSMUSG00000031398 | plexin A3 [Source:MGI Symbol;Acc:MGI:107683] | Plxna3 | Enriched in P8 retina | -1.576 | 0.000 | 112.900 |
| ENSMUSG00000027669 | guanine nucleotide binding protein (G protein), beta 4 [Source:MGI Symbol;Acc:MGI:104581] | Gnb4 | Enriched in P8 retina | -1.585 | 0.000 | 103.700 |
| ENSMUSG00000027217 | tetraspanin 18 [Source:MGI Symbol;Acc:MGI:1917186] | Tspan18 | Enriched in P8 retina | -1.586 | 0.000 | 78.820 |
| ENSMUSG00000028610 | DMRT-like family B with proline-rich C-terminal, 1 [Source:MGI Symbol;Acc:MGI:1927125] | Dmrtb1 | Enriched in P8 retina | -1.610 | 0.001 | 17.020 |
| ENSMUSG00000044469 | tumor necrosis factor, alpha-induced protein 8-like 1 [Source:MGI Symbol;Acc:MGI:1913693] | Tnfaip8l1 | Enriched in P8 retina | -1.612 | 0.000 | 9.788 |
| ENSMUSG00000018411 | microtubule-associated protein tau [Source:MGI Symbol;Acc:MGI:97180] | Mapt | Enriched in P8 retina | -1.628 | 0.000 | 1946.000 |
| ENSMUSG00000039976 | TBC1 domain family, member 16 [Source:MGI Symbol;Acc:MGI:2652878] | Tbc1d16 | Enriched in P8 retina | -1.657 | 0.000 | 1281.000 |
| ENSMUSG00000034993 | vesicle amine transport 1 [Source:MGI Symbol;Acc:MGI:1349450] | Vat1 | Enriched in P8 retina | -1.673 | 0.000 | 429.800 |
| ENSMUSG00000032735 | actin binding LIM protein family, member 3 [Source:MGI Symbol;Acc:MGI:2442582] | Ablim3 | Enriched in P8 retina | -1.741 | 0.000 | 445.700 |
| ENSMUSG00000030830 | integrin alpha L [Source:MGI Symbol;Acc:MGI:96606] | Itgal | Enriched in P8 retina | -1.771 | 0.002 | 6.323 |
| ENSMUSG00000058420 | synaptotagmin XVII [Source:MGI Symbol;Acc:MGI:104966] | Syt17 | Enriched in P8 retina | -1.771 | 0.000 | 49.030 |
| ENSMUSG00000063446 | phospholipid phosphatase related 1 [Source:MGI Symbol;Acc:MGI:2445015] | Plppr1 | Enriched in P8 retina | -1.806 | 0.000 | 101.600 |
| ENSMUSG00000027314 | delta like canonical Notch ligand 4 [Source:MGI Symbol;Acc:MGI:1859388] | Dll4 | Enriched in P8 retina | -1.839 | 0.000 | 11.290 |
| ENSMUSG00000029123 | serine/threonine kinase 32B [Source:MGI Symbol;Acc:MGI:1927552] | Stk32b | Enriched in P8 retina | -1.839 | 0.000 | 93.320 |
| ENSMUSG00000022044 | stathmin-like 4 [Source:MGI Symbol;Acc:MGI:1931224] | Stmn4 | Enriched in P8 retina | -1.840 | 0.000 | 638.300 |
| ENSMUSG00000025268 | MAGE family member D2 [Source:MGI Symbol;Acc:MGI:1933391] | Maged2 | Enriched in P8 retina | -1.855 | 0.000 | 189.800 |
| ENSMUSG00000035835 | phospholipid phosphatase related 3 [Source:MGI Symbol;Acc:MGI:2388640] | Plppr3 | Enriched in P8 retina | -1.896 | 0.000 | 329.900 |
| ENSMUSG00000036377 | capping protein inhibiting regulator of actin [Source:MGI Symbol;Acc:MGI:2444817] | Cracd | Enriched in P8 retina | -1.911 | 0.000 | 1779.000 |
| ENSMUSG00000006218 | family with sequence similarity 131, member C [Source:MGI Symbol;Acc:MGI:2685539] | Fam131c | Enriched in P8 retina | -1.933 | 0.000 | 42.580 |
| ENSMUSG00000037962 | refilin A [Source:MGI Symbol;Acc:MGI:1920371] | Rflna | Enriched in P8 retina | -1.949 | 0.005 | 3.272 |
| ENSMUSG00000026380 | transcription factor CP2-like 1 [Source:MGI Symbol;Acc:MGI:2444691] | Tfcp2l1 | Enriched in P8 retina | -1.970 | 0.001 | 32.750 |
| ENSMUSG00000045731 | prepronociceptin [Source:MGI Symbol;Acc:MGI:105308] | Pnoc | Enriched in P8 retina | -1.984 | 0.000 | 50.560 |
| ENSMUSG00000054555 | a disintegrin and metallopeptidase domain 12 (meltrin alpha) [Source:MGI Symbol;Acc:MGI:105378] | Adam12 | Enriched in P8 retina | -2.019 | 0.000 | 77.270 |
| ENSMUSG00000031111 | immunoglobulin superfamily, member 1 [Source:MGI Symbol;Acc:MGI:2147913] | Igsf1 | Enriched in P8 retina | -2.020 | 0.000 | 83.920 |
| ENSMUSG00000037362 | cellular communication network factor 3 [Source:MGI Symbol;Acc:MGI:109185] | Ccn3 | Enriched in P8 retina | -2.027 | 0.025 | 15.410 |
| ENSMUSG00000018012 | Rac family small GTPase 3 [Source:MGI Symbol;Acc:MGI:2180784] | Rac3 | Enriched in P8 retina | -2.060 | 0.000 | 122.200 |
| ENSMUSG00000001119 | collagen, type VI, alpha 1 [Source:MGI Symbol;Acc:MGI:88459] | Col6a1 | Enriched in P8 retina | -2.081 | 0.000 | 82.800 |
| ENSMUSG00000074923 | p21 (RAC1) activated kinase 6 [Source:MGI Symbol;Acc:MGI:2679420] | Pak6 | Enriched in P8 retina | -2.130 | 0.000 | 281.400 |
| ENSMUSG00000027306 | nucleolar and spindle associated protein 1 [Source:MGI Symbol;Acc:MGI:2675669] | Nusap1 | Enriched in P8 retina | -2.181 | 0.000 | 10.720 |
| ENSMUSG00000047261 | growth associated protein 43 [Source:MGI Symbol;Acc:MGI:95639] | Gap43 | Enriched in P8 retina | -2.287 | 0.000 | 876.700 |
| ENSMUSG00000028681 | patched 2 [Source:MGI Symbol;Acc:MGI:1095405] | Ptch2 | Enriched in P8 retina | -2.327 | 0.007 | 9.607 |
| ENSMUSG00000043165 | loricrin [Source:MGI Symbol;Acc:MGI:96816] | Lor | Enriched in P8 retina | -2.367 | 0.007 | 17.170 |
| ENSMUSG00000030854 | protein tyrosine phosphatase, non-receptor type 5 [Source:MGI Symbol;Acc:MGI:97807] | Ptpn5 | Enriched in P8 retina | -2.440 | 0.000 | 407.000 |
| ENSMUSG00000062372 | otoferlin [Source:MGI Symbol;Acc:MGI:1891247] | Otof | Enriched in P8 retina | -2.493 | 0.002 | 68.050 |
| ENSMUSG00000036856 | wingless-type MMTV integration site family, member 4 [Source:MGI Symbol;Acc:MGI:98957] | Wnt4 | Enriched in P8 retina | -2.541 | 0.000 | 80.140 |
| ENSMUSG00000023067 | cyclin-dependent kinase inhibitor 1A (P21) [Source:MGI Symbol;Acc:MGI:104556] | Cdkn1a | Enriched in P8 retina | -2.543 | 0.000 | 78.160 |
| ENSMUSG00000004098 | collagen, type V, alpha 3 [Source:MGI Symbol;Acc:MGI:1858212] | Col5a3 | Enriched in P8 retina | -2.569 | 0.000 | 67.350 |
| ENSMUSG00000029602 | RAS protein activator like 1 (GAP1 like) [Source:MGI Symbol;Acc:MGI:1330842] | Rasal1 | Enriched in P8 retina | -2.602 | 0.000 | 51.400 |
| ENSMUSG00000009356 | lactoperoxidase [Source:MGI Symbol;Acc:MGI:1923363] | Lpo | Enriched in P8 retina | -2.605 | 0.023 | 3.426 |
| ENSMUSG00000029414 | kinetochore associated 1 [Source:MGI Symbol;Acc:MGI:2673709] | Kntc1 | Enriched in P8 retina | -2.770 | 0.000 | 7.562 |
| ENSMUSG00000026875 | TNF receptor-associated factor 1 [Source:MGI Symbol;Acc:MGI:101836] | Traf1 | Enriched in P8 retina | -2.890 | 0.000 | 10.430 |
| ENSMUSG00000021070 | bradykinin receptor, beta 2 [Source:MGI Symbol;Acc:MGI:102845] | Bdkrb2 | Enriched in P8 retina | -3.200 | 0.000 | 6.429 |
| ENSMUSG00000046491 | C1q and tumor necrosis factor related protein 2 [Source:MGI Symbol;Acc:MGI:1916433] | C1qtnf2 | Enriched in P8 retina | -3.205 | 0.020 | 4.439 |
| ENSMUSG00000038112 | expressed sequence AW551984 [Source:MGI Symbol;Acc:MGI:2143322] | AW551984 | Enriched in P8 retina | -4.377 | 0.000 | 188.600 |
| ENSMUSG00000020838 | solute carrier family 6 (neurotransmitter transporter, serotonin), member 4 [Source:MGI Symbol;Acc:MGI:96285] | Slc6a4 | Enriched in P8 retina | -4.631 | 0.000 | 23.260 |
| ENSMUSG00000035355 | potassium voltage-gated channel, subfamily H (eag-related), member 4 [Source:MGI Symbol;Acc:MGI:2156184] | Kcnh4 | Enriched in P8 retina | -5.211 | 0.001 | 7.674 |

**Contrast numerator: P15_KO_retina. Contrast denominator: P8_KO_retina.**

| ensembl_gene_id | description | mgi_symbol | Significance | deseq_logfc | deseq_adjp | deseq_basemean |
|---|---|---|---|---|---|---|
| ENSMUSG00000037727 | arginine vasopressin [Source:MGI Symbol;Acc:MGI:88121] | Avp | Enriched in P15 retina | 14.590 | 0.000 | 32.860 |
| ENSMUSG00000079418 | autophagy related 4A, cysteine peptidase [Source:MGI Symbol;Acc:MGI:2147903] | Atg4a | Enriched in P15 retina | 4.825 | 0.000 | 66.590 |
| ENSMUSG00000086962 | predicted gene 12248 [Source:MGI Symbol;Acc:MGI:3651124] | Gm12248 | Enriched in P15 retina | 4.671 | 0.000 | 8.028 |
| ENSMUSG00000051985 | immunoglobulin-like and fibronectin type III domain containing 1 [Source:MGI Symbol;Acc:MGI:3045352] | Igfn1 | Enriched in P15 retina | 3.954 | 0.000 | 21.940 |
| ENSMUSG00000027401 | transglutaminase 3, E polypeptide [Source:MGI Symbol;Acc:MGI:98732] | Tgm3 | Enriched in P15 retina | 3.943 | 0.000 | 3.600 |
| ENSMUSG00000102349 | predicted gene, 37376 [Source:MGI Symbol;Acc:MGI:5610604] | Gm37376 | Enriched in P15 retina | 3.916 | 0.000 | 32.910 |
| ENSMUSG00000063428 | D-aspartate oxidase [Source:MGI Symbol;Acc:MGI:1925528] | Ddo | Enriched in P15 retina | 3.913 | 0.000 | 90.030 |
| ENSMUSG00000032356 | RAS protein-specific guanine nucleotide-releasing factor 1 [Source:MGI Symbol;Acc:MGI:99694] | Rasgrf1 | Enriched in P15 retina | 3.888 | 0.000 | 1288.000 |
| ENSMUSG00000092210 | RIKEN cDNA A930009A15 gene [Source:MGI Symbol;Acc:MGI:1925048] | A930009A15Rik | Enriched in P15 retina | 3.833 | 0.000 | 13.520 |
| ENSMUSG00000035296 | sarcoglycan, gamma (dystrophin-associated glycoprotein) [Source:MGI Symbol;Acc:MGI:1346524] | Sgcg | Enriched in P15 retina | 3.695 | 0.000 | 9.668 |
| ENSMUSG00000073551 | serine peptidase inhibitor, Kazal type 13 [Source:MGI Symbol;Acc:MGI:3642511] | Spink13 | Enriched in P15 retina | 3.489 | 0.000 | 7.681 |
| ENSMUSG00000005716 | parvalbumin [Source:MGI Symbol;Acc:MGI:97821] | Pvalb | Enriched in P15 retina | 3.410 | 0.000 | 26.700 |
| ENSMUSG00000007594 | hyaluronan and proteoglycan link protein 4 [Source:MGI Symbol;Acc:MGI:2679531] | Hapln4 | Enriched in P15 retina | 3.145 | 0.000 | 156.800 |
| ENSMUSG00000043727 | RIKEN cDNA F830045P16 gene [Source:MGI Symbol;Acc:MGI:3045317] | F830045P16Rik | Enriched in P15 retina | 3.049 | 0.000 | 4.358 |
| ENSMUSG00000114882 | predicted gene, 48581 [Source:MGI Symbol;Acc:MGI:6098148] | Gm48581 | Enriched in P15 retina | 2.815 | 0.002 | 15.290 |
| ENSMUSG00000045275 | Leber congenital amaurosis 5-like [Source:MGI Symbol;Acc:MGI:3041157] | Lca5l | Enriched in P15 retina | 2.678 | 0.000 | 37.830 |
| ENSMUSG00000029190 | DNA segment, Chr 5, ERATO Doi 579, expressed [Source:MGI Symbol;Acc:MGI:1261849] | D5Ertd579e | Enriched in P15 retina | 2.616 | 0.000 | 1250.000 |
| ENSMUSG00000060579 | fragile histidine triad gene [Source:MGI Symbol;Acc:MGI:1277947] | Fhit | Enriched in P15 retina | 2.602 | 0.000 | 135.300 |
| ENSMUSG00000044362 | coiled-coil domain containing 89 [Source:MGI Symbol;Acc:MGI:1917304] | Ccdc89 | Enriched in P15 retina | 2.582 | 0.000 | 17.870 |
| ENSMUSG00000031174 | retinitis pigmentosa GTPase regulator [Source:MGI Symbol;Acc:MGI:1344037] | Rpgr | Enriched in P15 retina | 2.535 | 0.000 | 194.100 |
| ENSMUSG00000043541 | cancer susceptibility candidate 1 [Source:MGI Symbol;Acc:MGI:2444480] | Casc1 | Enriched in P15 retina | 2.499 | 0.000 | 13.930 |
| ENSMUSG00000026414 | troponin T2, cardiac [Source:MGI Symbol;Acc:MGI:104597] | Tnnt2 | Enriched in P15 retina | 2.364 | 0.003 | 15.870 |
| ENSMUSG00000046667 | RNA binding motif protein 12 B1 [Source:MGI Symbol;Acc:MGI:1919647] | Rbm12b1 | Enriched in P15 retina | 2.305 | 0.000 | 35.330 |
| ENSMUSG00000074037 | melanocortin 1 receptor [Source:MGI Symbol;Acc:MGI:99456] | Mc1r | Enriched in P15 retina | 2.305 | 0.000 | 11.830 |
| ENSMUSG00000040490 | leucine rich repeat and fibronectin type III domain containing 2 [Source:MGI Symbol;Acc:MGI:1917780] | Lrfn2 | Enriched in P15 retina | 2.287 | 0.000 | 166.400 |
| ENSMUSG00000042073 | abhydrolase domain containing 14b [Source:MGI Symbol;Acc:MGI:1923741] | Abhd14b | Enriched in P15 retina | 2.216 | 0.000 | 38.480 |
| ENSMUSG00000029601 | IQ motif containing D [Source:MGI Symbol;Acc:MGI:1922982] | Iqcd | Enriched in P15 retina | 2.151 | 0.005 | 36.370 |
| ENSMUSG00000029012 | origin recognition complex, subunit 5 [Source:MGI Symbol;Acc:MGI:1347044] | Orc5 | Enriched in P15 retina | 2.093 | 0.000 | 47.780 |
| ENSMUSG00000072572 | solute carrier family 39 (zinc transporter), member 2 [Source:MGI Symbol;Acc:MGI:2684326] | Slc39a2 | Enriched in P15 retina | 2.065 | 0.017 | 19.900 |
| ENSMUSG00000045968 | transmembrane epididymal family member 2 [Source:MGI Symbol;Acc:MGI:1923273] | Teddm2 | Enriched in P15 retina | 2.055 | 0.000 | 15.550 |
| ENSMUSG00000080316 | sperm acrosome associated 6 [Source:MGI Symbol;Acc:MGI:1922452] | Spaca6 | Enriched in P15 retina | 2.040 | 0.001 | 101.000 |
| ENSMUSG00000016356 | collagen, type XX, alpha 1 [Source:MGI Symbol;Acc:MGI:1920618] | Col20a1 | Enriched in P15 retina | 2.026 | 0.000 | 25.930 |
| ENSMUSG00000028248 | PNN interacting serine/arginine-rich [Source:MGI Symbol;Acc:MGI:1913875] | Pnisr | Enriched in P15 retina | 2.001 | 0.000 | 892.200 |
| ENSMUSG00000031257 | NADPH oxidase 1 [Source:MGI Symbol;Acc:MGI:2450016] | Nox1 | Enriched in P15 retina | 1.993 | 0.001 | 7.043 |
| ENSMUSG00000029330 | CDP-diacylglycerol synthase 1 [Source:MGI Symbol;Acc:MGI:1921846] | Cds1 | Enriched in P15 retina | 1.929 | 0.000 | 337.700 |
| ENSMUSG00000021619 | autophagy related 10 [Source:MGI Symbol;Acc:MGI:1914045] | Atg10 | Enriched in P15 retina | 1.927 | 0.002 | 39.830 |
| ENSMUSG00000015354 | procollagen C-endopeptidase enhancer 2 [Source:MGI Symbol;Acc:MGI:1923727] | Pcolce2 | Enriched in P15 retina | 1.881 | 0.000 | 23.560 |
| ENSMUSG00000039037 | ST6 (alpha-N-acetyl-neuraminyl-2,3-beta-galactosyl-1,3)-N-acetylgalactosaminide alpha-2,6-sialyltransferase 5 [Source:MGI Symbol;Acc:MGI:13494 | St6galnac5 | Enriched in P15 retina | 1.830 | 0.000 | 194.500 |
| ENSMUSG00000072770 | proacrosin binding protein [Source:MGI Symbol;Acc:MGI:1859515] | Acrbp | Enriched in P15 retina | 1.829 | 0.012 | 25.870 |
| ENSMUSG00000096696 | zinc finger protein 960 [Source:MGI Symbol;Acc:MGI:3052731] | Zfp960 | Enriched in P15 retina | 1.756 | 0.013 | 4.000 |
| ENSMUSG00000046480 | sodium channel, type IV, beta [Source:MGI Symbol;Acc:MGI:2687406] | Scn4b | Enriched in P15 retina | 1.752 | 0.005 | 167.600 |
| ENSMUSG00000025269 | apurinic/apyrimidinic endonuclease 2 [Source:MGI Symbol;Acc:MGI:1924872] | Apex2 | Enriched in P15 retina | 1.726 | 0.003 | 107.000 |
| ENSMUSG00000020732 | RAB37, member RAS oncogene family [Source:MGI Symbol;Acc:MGI:1929945] | Rab37 | Enriched in P15 retina | 1.719 | 0.014 | 50.930 |
| ENSMUSG00000021495 | family with sequence similarity 193, member B [Source:MGI Symbol;Acc:MGI:2385851] | Fam193b | Enriched in P15 retina | 1.705 | 0.001 | 295.800 |
| ENSMUSG00000040759 | CKLF-like MARVEL transmembrane domain containing 5 [Source:MGI Symbol;Acc:MGI:2447164] | Cmtm5 | Enriched in P15 retina | 1.679 | 0.017 | 15.120 |
| ENSMUSG00000045314 | sosondowah ankyrin repeat domain family member B [Source:MGI Symbol;Acc:MGI:1925338] | Sowahb | Enriched in P15 retina | 1.597 | 0.008 | 9.564 |
| ENSMUSG00000014786 | solute carrier family 9 (sodium/hydrogen exchanger), member 5 [Source:MGI Symbol;Acc:MGI:2685542] | Slc9a5 | Enriched in P15 retina | 1.570 | 0.000 | 110.700 |
| ENSMUSG00000070697 | UTP3 small subunit processome component [Source:MGI Symbol;Acc:MGI:1919230] | Utp3 | Enriched in P15 retina | 1.514 | 0.000 | 195.600 |
| ENSMUSG00000020994 | pinin [Source:MGI Symbol;Acc:MGI:1100514] | Pnn | Enriched in P15 retina | 1.501 | 0.000 | 615.100 |
| ENSMUSG00000030337 | vesicle-associated membrane protein 1 [Source:MGI Symbol;Acc:MGI:1313276] | Vamp1 | Enriched in P15 retina | 1.484 | 0.000 | 282.400 |
| ENSMUSG00000022894 | a disintegrin-like and metallopeptidase (reprolysin type) with thrombospondin type 1 motif, 5 (aggrecanase-2) [Source:MGI Symbol;Acc:MGI:13463 | Adamts5 | Enriched in P15 retina | 1.483 | 0.000 | 56.570 |
| ENSMUSG00000020863 | LUC7-like 3 (S. cerevisiae) [Source:MGI Symbol;Acc:MGI:1914934] | Luc7l3 | Enriched in P15 retina | 1.481 | 0.000 | 973.300 |
| ENSMUSG00000035314 | glycerophosphodiester phosphodiesterase domain containing 5 [Source:MGI Symbol;Acc:MGI:2686926] | Gdpd5 | Enriched in P15 retina | 1.432 | 0.001 | 328.800 |
| ENSMUSG00000004902 | solute carrier family 25 (mitochondrial carrier), member 18 [Source:MGI Symbol;Acc:MGI:1919053] | Slc25a18 | Enriched in P15 retina | 1.417 | 0.007 | 68.620 |
| ENSMUSG00000042050 | dynein 2 intermediate chain 1 [Source:MGI Symbol;Acc:MGI:2445085] | Dync2i1 | Enriched in P15 retina | 1.416 | 0.000 | 228.800 |
| ENSMUSG00000040936 | unc-51-like kinase 4 [Source:MGI Symbol;Acc:MGI:1921622] | Ulk4 | Enriched in P15 retina | 1.415 | 0.011 | 132.600 |
| ENSMUSG00000028730 | cilia and flagella associated protein 57 [Source:MGI Symbol;Acc:MGI:2686209] | Cfap57 | Enriched in P15 retina | 1.400 | 0.008 | 18.370 |
| ENSMUSG00000047632 | fibroblast growth factor binding protein 3 [Source:MGI Symbol;Acc:MGI:1919764] | Fgfbp3 | Enriched in P15 retina | 1.372 | 0.036 | 20.400 |
| ENSMUSG00000048878 | hexamethylene bis-acetamide inducible 1 [Source:MGI Symbol;Acc:MGI:2385923] | Hexim1 | Enriched in P15 retina | 1.338 | 0.000 | 257.600 |
| ENSMUSG00000071379 | hippocalcin-like 1 [Source:MGI Symbol;Acc:MGI:1855689] | Hpcal1 | Enriched in P15 retina | 1.336 | 0.000 | 98.630 |
| ENSMUSG00000056938 | acyl-Coenzyme A binding domain containing 4 [Source:MGI Symbol;Acc:MGI:1914381] | Acbd4 | Enriched in P15 retina | 1.306 | 0.000 | 43.250 |

| | | | | | | |
|---|---|---|---|---|---|---|
| ENSMUSG00000051246 | Myb/SANT-like DNA-binding domain containing 1 [Source:MGI Symbol;Acc:MGI:2684990] | Msantd1 | Enriched in P15 retina | 1.299 | 0.021 | 15.100 |
| ENSMUSG00000045319 | proline and serine rich 2 [Source:MGI Symbol;Acc:MGI:2442238] | Proser2 | Enriched in P15 retina | 1.279 | 0.001 | 42.910 |
| ENSMUSG00000040591 | centriolar satellite-associated tubulin polyglutamylase complex regulator 1 [Source:MGI Symbol;Acc:MGI:1915079] | Cstpp1 | Enriched in P15 retina | 1.245 | 0.000 | 214.800 |
| ENSMUSG00000061462 | obscurin, cytoskeletal calmodulin and titin-interacting RhoGEF [Source:MGI Symbol;Acc:MGI:2681862] | Obscn | Enriched in P15 retina | 1.231 | 0.047 | 66.020 |
| ENSMUSG00000004561 | methyltransferase like 17 [Source:MGI Symbol;Acc:MGI:1098577] | Mettl17 | Enriched in P15 retina | 1.190 | 0.034 | 47.960 |
| ENSMUSG00000074652 | myosin, heavy chain 7B, cardiac muscle, beta [Source:MGI Symbol;Acc:MGI:3710243] | Myh7b | Enriched in P15 retina | 1.185 | 0.005 | 65.130 |
| ENSMUSG00000027286 | leucine rich repeat containing 57 [Source:MGI Symbol;Acc:MGI:1913856] | Lrrc57 | Enriched in P15 retina | 1.125 | 0.001 | 34.710 |
| ENSMUSG00000046793 | G protein-coupled receptor 61 [Source:MGI Symbol;Acc:MGI:2441719] | Gpr61 | Enriched in P15 retina | 1.108 | 0.003 | 53.440 |
| ENSMUSG00000001520 | nuclear receptor interacting protein 2 [Source:MGI Symbol;Acc:MGI:1891884] | Nrip2 | Enriched in P15 retina | 1.097 | 0.023 | 17.020 |
| ENSMUSG00000060843 | catenin (cadherin associated protein), alpha 3 [Source:MGI Symbol;Acc:MGI:2661445] | Ctnna3 | Enriched in P15 retina | 1.041 | 0.003 | 51.560 |
| ENSMUSG00000039057 | myosin XVI [Source:MGI Symbol;Acc:MGI:2685951] | Myo16 | Enriched in P8 retina | -1.000 | 0.006 | 431.200 |
| ENSMUSG00000004113 | calcium channel, voltage-dependent, N type, alpha 1B subunit [Source:MGI Symbol;Acc:MGI:88296] | Cacna1b | Enriched in P8 retina | -1.001 | 0.000 | 1028.000 |
| ENSMUSG00000050530 | family with sequence similarity 171, member A1 [Source:MGI Symbol;Acc:MGI:2442917] | Fam171a1 | Enriched in P8 retina | -1.001 | 0.000 | 240.800 |
| ENSMUSG00000048537 | pleckstrin homology like domain, family B, member 1 [Source:MGI Symbol;Acc:MGI:2143230] | Phldb1 | Enriched in P8 retina | -1.004 | 0.000 | 593.000 |
| ENSMUSG00000026156 | beta-1,3-glucuronyltransferase 2 (glucuronosyltransferase S) [Source:MGI Symbol;Acc:MGI:2389490] | B3gat2 | Enriched in P8 retina | -1.006 | 0.000 | 140.500 |
| ENSMUSG00000058420 | synaptotagmin XVII [Source:MGI Symbol;Acc:MGI:104966] | Syt17 | Enriched in P8 retina | -1.006 | 0.023 | 49.030 |
| ENSMUSG00000032024 | CXADR-like membrane protein [Source:MGI Symbol;Acc:MGI:1918816] | Clmp | Enriched in P8 retina | -1.010 | 0.001 | 151.400 |
| ENSMUSG00000061981 | flotillin 2 [Source:MGI Symbol;Acc:MGI:103309] | Flot2 | Enriched in P8 retina | -1.012 | 0.000 | 295.800 |
| ENSMUSG00000032492 | parathyroid hormone 1 receptor [Source:MGI Symbol;Acc:MGI:97801] | Pth1r | Enriched in P8 retina | -1.013 | 0.020 | 46.940 |
| ENSMUSG00000051062 | fibrillarin-like 1 [Source:MGI Symbol;Acc:MGI:3034689] | Fbll1 | Enriched in P8 retina | -1.014 | 0.011 | 42.330 |
| ENSMUSG00000024008 | copine V [Source:MGI Symbol;Acc:MGI:2385908] | Cpne5 | Enriched in P8 retina | -1.020 | 0.013 | 78.100 |
| ENSMUSG00000030844 | regulator of G-protein signalling 10 [Source:MGI Symbol;Acc:MGI:1915115] | Rgs10 | Enriched in P8 retina | -1.020 | 0.047 | 70.220 |
| ENSMUSG00000047181 | sterile alpha motif domain containing 14 [Source:MGI Symbol;Acc:MGI:2384945] | Samd14 | Enriched in P8 retina | -1.021 | 0.000 | 1407.000 |
| ENSMUSG00000021262 | Ena-vasodilator stimulated phosphoprotein [Source:MGI Symbol;Acc:MGI:1194884] | Evl | Enriched in P8 retina | -1.024 | 0.000 | 1083.000 |
| ENSMUSG00000038351 | small G protein signaling modulator 2 [Source:MGI Symbol;Acc:MGI:2144695] | Sgsm2 | Enriched in P8 retina | -1.024 | 0.000 | 381.700 |
| ENSMUSG00000031586 | RNA binding protein gene with multiple splicing [Source:MGI Symbol;Acc:MGI:1334446] | Rbpms | Enriched in P8 retina | -1.026 | 0.020 | 81.550 |
| ENSMUSG00000034994 | eukaryotic translation elongation factor 2 [Source:MGI Symbol;Acc:MGI:95288] | Eef2 | Enriched in P8 retina | -1.028 | 0.000 | 4640.000 |
| ENSMUSG00000036246 | Gem-interacting protein [Source:MGI Symbol;Acc:MGI:1926066] | Gmip | Enriched in P8 retina | -1.029 | 0.009 | 61.040 |
| ENSMUSG00000096472 | cyclin dependent kinase inhibitor 2D [Source:MGI Symbol;Acc:MGI:105387] | Cdkn2d | Enriched in P8 retina | -1.030 | 0.000 | 66.810 |
| ENSMUSG00000062184 | heparan sulfate 6-O-sulfotransferase 2 [Source:MGI Symbol;Acc:MGI:1354959] | Hs6st2 | Enriched in P8 retina | -1.036 | 0.007 | 235.400 |
| ENSMUSG00000006731 | beta-1,4-N-acetyl-galactosaminyl transferase 1 [Source:MGI Symbol;Acc:MGI:1342057] | B4galnt1 | Enriched in P8 retina | -1.037 | 0.000 | 230.200 |
| ENSMUSG00000057182 | sodium channel, voltage-gated, type III, alpha [Source:MGI Symbol;Acc:MGI:98249] | Scn3a | Enriched in P8 retina | -1.039 | 0.001 | 542.800 |
| ENSMUSG00000002771 | glutamate receptor, ionotropic, NMDA2D (epsilon 4) [Source:MGI Symbol;Acc:MGI:95823] | Grin2d | Enriched in P8 retina | -1.040 | 0.000 | 466.700 |
| ENSMUSG00000026443 | leucine rich repeat protein 2, neuronal [Source:MGI Symbol;Acc:MGI:106037] | Lrrn2 | Enriched in P8 retina | -1.041 | 0.000 | 489.200 |
| ENSMUSG00000031906 | sphingomyelin phosphodiesterase 3, neutral [Source:MGI Symbol;Acc:MGI:1927578] | Smpd3 | Enriched in P8 retina | -1.041 | 0.001 | 615.600 |
| ENSMUSG00000078249 | high mobility group AT-hook 1B [Source:MGI Symbol;Acc:MGI:96161] | Hmga1b | Enriched in P8 retina | -1.046 | 0.000 | 54.250 |
| ENSMUSG00000030352 | tetraspanin 9 [Source:MGI Symbol;Acc:MGI:1924558] | Tspan9 | Enriched in P8 retina | -1.047 | 0.000 | 538.100 |
| ENSMUSG00000024777 | protein phosphatase 2, regulatory subunit B', beta [Source:MGI Symbol;Acc:MGI:2388480] | Ppp2r5b | Enriched in P8 retina | -1.049 | 0.000 | 428.000 |
| ENSMUSG00000026778 | protein kinase C, theta [Source:MGI Symbol;Acc:MGI:97601] | Prkcq | Enriched in P8 retina | -1.049 | 0.028 | 111.900 |
| ENSMUSG00000023017 | acid-sensing (proton-gated) ion channel 1 [Source:MGI Symbol;Acc:MGI:1194915] | Asic1 | Enriched in P8 retina | -1.056 | 0.000 | 275.400 |
| ENSMUSG00000027612 | matrix metallopeptidase 24 [Source:MGI Symbol;Acc:MGI:1341867] | Mmp24 | Enriched in P8 retina | -1.056 | 0.000 | 380.900 |
| ENSMUSG00000046321 | heparan sulfate (glucosamine) 3-O-sulfotransferase 2 [Source:MGI Symbol;Acc:MGI:1333802] | Hs3st2 | Enriched in P8 retina | -1.058 | 0.010 | 41.560 |
| ENSMUSG00000003423 | PIH1 domain containing 1 [Source:MGI Symbol;Acc:MGI:1916095] | Pih1d1 | Enriched in P8 retina | -1.060 | 0.000 | 54.970 |
| ENSMUSG00000036062 | PHD finger protein 24 [Source:MGI Symbol;Acc:MGI:2140712] | Phf24 | Enriched in P8 retina | -1.061 | 0.000 | 370.000 |
| ENSMUSG00000039824 | myosin, light polypeptide 6B [Source:MGI Symbol;Acc:MGI:1917789] | Myl6b | Enriched in P8 retina | -1.061 | 0.005 | 27.880 |
| ENSMUSG00000029033 | ArfGAP with coiled-coil, ankyrin repeat and PH domains 3 [Source:MGI Symbol;Acc:MGI:2153589] | Acap3 | Enriched in P8 retina | -1.071 | 0.000 | 352.100 |
| ENSMUSG00000027330 | cell division cycle 25B [Source:MGI Symbol;Acc:MGI:99701] | Cdc25b | Enriched in P8 retina | -1.072 | 0.014 | 31.550 |
| ENSMUSG00000029223 | ubiquitin carboxy-terminal hydrolase L1 [Source:MGI Symbol;Acc:MGI:103149] | Uchl1 | Enriched in P8 retina | -1.075 | 0.009 | 1008.000 |
| ENSMUSG00000035064 | eukaryotic elongation factor-2 kinase [Source:MGI Symbol;Acc:MGI:1195261] | Eef2k | Enriched in P8 retina | -1.082 | 0.000 | 118.400 |
| ENSMUSG00000047085 | leucine rich repeat containing 4B [Source:MGI Symbol;Acc:MGI:3027390] | Lrrc4b | Enriched in P8 retina | -1.082 | 0.000 | 980.700 |
| ENSMUSG00000079003 | sterile alpha motif domain containing 1 [Source:MGI Symbol;Acc:MGI:2142433] | Samd1 | Enriched in P8 retina | -1.082 | 0.000 | 225.700 |
| ENSMUSG00000003352 | calcium channel, voltage-dependent, beta 3 subunit [Source:MGI Symbol;Acc:MGI:103307] | Cacnb3 | Enriched in P8 retina | -1.084 | 0.000 | 307.200 |
| ENSMUSG00000030600 | leucine rich repeat and fibronectin type III domain containing 1 [Source:MGI Symbol;Acc:MGI:2136810] | Lrfn1 | Enriched in P8 retina | -1.085 | 0.000 | 330.900 |
| ENSMUSG00000034390 | c-Maf inducing protein [Source:MGI Symbol;Acc:MGI:1921690] | Cmip | Enriched in P8 retina | -1.089 | 0.000 | 769.300 |
| ENSMUSG00000040563 | phospholipid phosphatase related 2 [Source:MGI Symbol;Acc:MGI:2384575] | Plppr2 | Enriched in P8 retina | -1.089 | 0.000 | 267.800 |
| ENSMUSG00000022438 | parvin, beta [Source:MGI Symbol;Acc:MGI:2153063] | Parvb | Enriched in P8 retina | -1.090 | 0.001 | 253.300 |
| ENSMUSG00000024050 | widely-interspaced zinc finger motifs [Source:MGI Symbol;Acc:MGI:1332638] | Wiz | Enriched in P8 retina | -1.091 | 0.000 | 422.100 |
| ENSMUSG00000032290 | protein tyrosine phosphatase, non-receptor type 9 [Source:MGI Symbol;Acc:MGI:1928376] | Ptpn9 | Enriched in P8 retina | -1.091 | 0.000 | 147.600 |
| ENSMUSG00000027646 | Rous sarcoma oncogene [Source:MGI Symbol;Acc:MGI:98397] | Src | Enriched in P8 retina | -1.095 | 0.000 | 473.500 |
| ENSMUSG00000030403 | vasodilator-stimulated phosphoprotein [Source:MGI Symbol;Acc:MGI:109268] | Vasp | Enriched in P8 retina | -1.099 | 0.000 | 78.390 |
| ENSMUSG00000051323 | protocadherin 19 [Source:MGI Symbol;Acc:MGI:2685563] | Pcdh19 | Enriched in P8 retina | -1.100 | 0.004 | 636.500 |
| ENSMUSG00000056306 | serine rich and transmembrane domain containing 1 [Source:MGI Symbol;Acc:MGI:3607715] | Sertm1 | Enriched in P8 retina | -1.102 | 0.016 | 70.230 |
| ENSMUSG00000003644 | ribosomal protein S6 kinase polypeptide 1 [Source:MGI Symbol;Acc:MGI:104558] | Rps6ka1 | Enriched in P8 retina | -1.104 | 0.000 | 65.070 |

| | | | | | | |
|---|---|---|---|---|---|---|
| ENSMUSG00000032297 | CUGBP, Elav-like family member 6 [Source:MGI Symbol;Acc:MGI:1923433] | Celf6 | Enriched in P8 retina | -1.104 | 0.008 | 185.400 |
| ENSMUSG00000047428 | delta like non-canonical Notch ligand 2 [Source:MGI Symbol;Acc:MGI:2146838] | Dlk2 | Enriched in P8 retina | -1.104 | 0.006 | 45.680 |
| ENSMUSG00000028458 | testis specific protein kinase 1 [Source:MGI Symbol;Acc:MGI:1201675] | Tesk1 | Enriched in P8 retina | -1.105 | 0.000 | 341.800 |
| ENSMUSG00000026321 | tumor necrosis factor receptor superfamily, member 11a, NFKB activator [Source:MGI Symbol;Acc:MGI:1314891] | Tnfrsf11a | Enriched in P8 retina | -1.106 | 0.010 | 53.110 |
| ENSMUSG00000028351 | bone morphogenic protein/retinoic acid inducible neural specific 1 [Source:MGI Symbol;Acc:MGI:1928478] | Brinp1 | Enriched in P8 retina | -1.111 | 0.004 | 586.100 |
| ENSMUSG00000027230 | cAMP responsive element binding protein 3-like 1 [Source:MGI Symbol;Acc:MGI:1347062] | Creb3l1 | Enriched in P8 retina | -1.118 | 0.034 | 27.390 |
| ENSMUSG00000042581 | thrombospondin, type I, domain containing 7B [Source:MGI Symbol;Acc:MGI:2443925] | Thsd7b | Enriched in P8 retina | -1.134 | 0.016 | 235.900 |
| ENSMUSG00000033209 | tetratricopeptide repeat domain 28 [Source:MGI Symbol;Acc:MGI:2140873] | Ttc28 | Enriched in P8 retina | -1.136 | 0.001 | 597.600 |
| ENSMUSG00000000489 | platelet derived growth factor, B polypeptide [Source:MGI Symbol;Acc:MGI:97528] | Pdgfb | Enriched in P8 retina | -1.138 | 0.003 | 70.920 |
| ENSMUSG00000028793 | ring finger protein 19B [Source:MGI Symbol;Acc:MGI:1922484] | Rnf19b | Enriched in P8 retina | -1.138 | 0.000 | 239.700 |
| ENSMUSG00000052981 | ubiquitin-conjugating enzyme E2Q family-like 1 [Source:MGI Symbol;Acc:MGI:1924230] | Ube2ql1 | Enriched in P8 retina | -1.144 | 0.000 | 310.300 |
| ENSMUSG00000039252 | leucine-rich repeat LGI family, member 2 [Source:MGI Symbol;Acc:MGI:2180196] | Lgi2 | Enriched in P8 retina | -1.149 | 0.008 | 332.300 |
| ENSMUSG00000001729 | thymoma viral proto-oncogene 1 [Source:MGI Symbol;Acc:MGI:87986] | Akt1 | Enriched in P8 retina | -1.152 | 0.000 | 701.000 |
| ENSMUSG00000006395 | hydroxypyruvate isomerase (putative) [Source:MGI Symbol;Acc:MGI:1915430] | Hyi | Enriched in P8 retina | -1.152 | 0.001 | 22.950 |
| ENSMUSG00000027394 | tubulin tyrosine ligase [Source:MGI Symbol;Acc:MGI:1916987] | Ttl | Enriched in P8 retina | -1.161 | 0.000 | 343.000 |
| ENSMUSG00000038916 | SOGA family member 3 [Source:MGI Symbol;Acc:MGI:1914662] | Soga3 | Enriched in P8 retina | -1.162 | 0.000 | 604.700 |
| ENSMUSG00000041836 | protein tyrosine phosphatase, receptor type, E [Source:MGI Symbol;Acc:MGI:97813] | Ptpre | Enriched in P8 retina | -1.168 | 0.000 | 124.900 |
| ENSMUSG00000074793 | heat shock protein 12B [Source:MGI Symbol;Acc:MGI:1919880] | Hspa12b | Enriched in P8 retina | -1.172 | 0.009 | 37.370 |
| ENSMUSG00000032020 | ubiquitin associated and SH3 domain containing, B [Source:MGI Symbol;Acc:MGI:1920078] | Ubash3b | Enriched in P8 retina | -1.173 | 0.000 | 190.100 |
| ENSMUSG00000043857 | mannoside acetylglucosaminyltransferase 5, isoenzyme B [Source:MGI Symbol;Acc:MGI:3606200] | Mgat5b | Enriched in P8 retina | -1.174 | 0.000 | 479.000 |
| ENSMUSG00000017631 | active BCR-related gene [Source:MGI Symbol;Acc:MGI:107771] | Abr | Enriched in P8 retina | -1.175 | 0.000 | 1239.000 |
| ENSMUSG00000038530 | regulator of G-protein signaling 4 [Source:MGI Symbol;Acc:MGI:108409] | Rgs4 | Enriched in P8 retina | -1.175 | 0.021 | 578.500 |
| ENSMUSG00000034839 | La ribonucleoprotein 6, translational regulator [Source:MGI Symbol;Acc:MGI:1914807] | Larp6 | Enriched in P8 retina | -1.180 | 0.000 | 87.580 |
| ENSMUSG00000020312 | SHC (Src homology 2 domain containing) transforming protein 2 [Source:MGI Symbol;Acc:MGI:106180] | Shc2 | Enriched in P8 retina | -1.182 | 0.000 | 242.800 |
| ENSMUSG00000024855 | phosphofurin acidic cluster sorting protein 1 [Source:MGI Symbol;Acc:MGI:1277113] | Pacs1 | Enriched in P8 retina | -1.188 | 0.000 | 260.200 |
| ENSMUSG00000031805 | Janus kinase 3 [Source:MGI Symbol;Acc:MGI:99928] | Jak3 | Enriched in P8 retina | -1.189 | 0.009 | 28.960 |
| ENSMUSG00000034751 | microtubule associated serine/threonine kinase family member 4 [Source:MGI Symbol;Acc:MGI:1918885] | Mast4 | Enriched in P8 retina | -1.189 | 0.000 | 628.500 |
| ENSMUSG00000033788 | dysferlin [Source:MGI Symbol;Acc:MGI:1349385] | Dysf | Enriched in P8 retina | -1.192 | 0.006 | 67.230 |
| ENSMUSG00000027200 | sema domain, transmembrane domain (TM), and cytoplasmic domain, (semaphorin) 6D [Source:MGI Symbol;Acc:MGI:2387661] | Sema6d | Enriched in P8 retina | -1.195 | 0.001 | 731.900 |
| ENSMUSG00000026888 | growth factor receptor bound protein 14 [Source:MGI Symbol;Acc:MGI:1355324] | Grb14 | Enriched in P8 retina | -1.196 | 0.007 | 102.400 |
| ENSMUSG00000041774 | YdjC homolog (bacterial) [Source:MGI Symbol;Acc:MGI:1916351] | Ydjc | Enriched in P8 retina | -1.198 | 0.000 | 46.050 |
| ENSMUSG00000018849 | WW, C2 and coiled-coil domain containing 1 [Source:MGI Symbol;Acc:MGI:2388637] | Wwc1 | Enriched in P8 retina | -1.199 | 0.000 | 625.400 |
| ENSMUSG00000058756 | thyroid hormone receptor alpha [Source:MGI Symbol;Acc:MGI:98742] | Thra | Enriched in P8 retina | -1.199 | 0.000 | 2307.000 |
| ENSMUSG00000037990 | SH3 domain containing ring finger 3 [Source:MGI Symbol;Acc:MGI:2444637] | Sh3rf3 | Enriched in P8 retina | -1.202 | 0.000 | 202.200 |
| ENSMUSG00000038860 | GTPase activating RANGAP domain-like 3 [Source:MGI Symbol;Acc:MGI:2139309] | Garnl3 | Enriched in P8 retina | -1.204 | 0.001 | 238.500 |
| ENSMUSG00000045763 | brain abundant, membrane attached signal protein 1 [Source:MGI Symbol;Acc:MGI:1917600] | Basp1 | Enriched in P8 retina | -1.204 | 0.000 | 3099.000 |
| ENSMUSG00000056258 | potassium voltage-gated channel, subfamily Q, member 3 [Source:MGI Symbol;Acc:MGI:1336181] | Kcnq3 | Enriched in P8 retina | -1.208 | 0.001 | 1087.000 |
| ENSMUSG00000045348 | neuronal tyrosine-phosphorylated phosphoinositide 3-kinase adaptor 1 [Source:MGI Symbol;Acc:MGI:2443880] | Nyap1 | Enriched in P8 retina | -1.215 | 0.000 | 501.100 |
| ENSMUSG00000038740 | multivesicular body subunit 12B [Source:MGI Symbol;Acc:MGI:1919793] | Mvb12b | Enriched in P8 retina | -1.217 | 0.000 | 759.600 |
| ENSMUSG00000063160 | numb-like [Source:MGI Symbol;Acc:MGI:894702] | Numbl | Enriched in P8 retina | -1.219 | 0.000 | 218.500 |
| ENSMUSG00000031963 | BMP-binding endothelial regulator [Source:MGI Symbol;Acc:MGI:1920480] | Bmper | Enriched in P8 retina | -1.220 | 0.009 | 44.470 |
| ENSMUSG00000033039 | microtubule associated monooxygenase, calponin and LIM domain containing -like 1 [Source:MGI Symbol;Acc:MGI:105870] | Micall1 | Enriched in P8 retina | -1.220 | 0.000 | 593.100 |
| ENSMUSG00000045045 | leucine rich repeat and fibronectin type III domain containing 4 [Source:MGI Symbol;Acc:MGI:2385612] | Lrfn4 | Enriched in P8 retina | -1.225 | 0.000 | 212.700 |
| ENSMUSG00000050272 | DS cell adhesion molecule [Source:MGI Symbol;Acc:MGI:1196281] | Dscam | Enriched in P8 retina | -1.226 | 0.001 | 1096.000 |
| ENSMUSG00000075289 | carnosine synthase 1 [Source:MGI Symbol;Acc:MGI:2147595] | Carns1 | Enriched in P8 retina | -1.227 | 0.003 | 20.470 |
| ENSMUSG00000036046 | RIKEN cDNA 5031439G07 gene [Source:MGI Symbol;Acc:MGI:2444899] | 5031439G07Rik | Enriched in P8 retina | -1.229 | 0.000 | 791.000 |
| ENSMUSG00000032946 | RAS, guanyl releasing protein 2 [Source:MGI Symbol;Acc:MGI:1333849] | Rasgrp2 | Enriched in P8 retina | -1.230 | 0.001 | 149.800 |
| ENSMUSG00000022456 | septin 3 [Source:MGI Symbol;Acc:MGI:1345148] | Septin3 | Enriched in P8 retina | -1.231 | 0.000 | 1129.000 |
| ENSMUSG00000037493 | calcium and integrin binding family member 2 [Source:MGI Symbol;Acc:MGI:1929293] | Cib2 | Enriched in P8 retina | -1.236 | 0.000 | 39.200 |
| ENSMUSG00000021294 | kinesin family member 26A [Source:MGI Symbol;Acc:MGI:2447072] | Kif26a | Enriched in P8 retina | -1.237 | 0.000 | 99.430 |
| ENSMUSG00000030411 | NOVA alternative splicing regulator 2 [Source:MGI Symbol;Acc:MGI:104296] | Nova2 | Enriched in P8 retina | -1.238 | 0.000 | 858.700 |
| ENSMUSG00000030748 | interleukin 4 receptor, alpha [Source:MGI Symbol;Acc:MGI:105367] | Il4ra | Enriched in P8 retina | -1.240 | 0.000 | 46.010 |
| ENSMUSG00000039741 | BAH domain and coiled-coil containing 1 [Source:MGI Symbol;Acc:MGI:2679272] | Bahcc1 | Enriched in P8 retina | -1.257 | 0.000 | 532.000 |
| ENSMUSG00000039004 | bone morphogenetic protein 6 [Source:MGI Symbol;Acc:MGI:88182] | Bmp6 | Enriched in P8 retina | -1.262 | 0.015 | 58.770 |
| ENSMUSG00000066705 | FXYD domain-containing ion transport regulator 6 [Source:MGI Symbol;Acc:MGI:1890226] | Fxyd6 | Enriched in P8 retina | -1.263 | 0.015 | 380.300 |
| ENSMUSG00000027500 | stathmin-like 2 [Source:MGI Symbol;Acc:MGI:98241] | Stmn2 | Enriched in P8 retina | -1.264 | 0.002 | 1265.000 |
| ENSMUSG00000034037 | FYVE, RhoGEF and PH domain containing 5 [Source:MGI Symbol;Acc:MGI:2443369] | Fgd5 | Enriched in P8 retina | -1.267 | 0.004 | 83.070 |
| ENSMUSG00000030376 | solute carrier family 8 (sodium/calcium exchanger), member 2 [Source:MGI Symbol;Acc:MGI:107996] | Slc8a2 | Enriched in P8 retina | -1.272 | 0.000 | 330.300 |
| ENSMUSG00000025743 | syndecan 3 [Source:MGI Symbol;Acc:MGI:1349163] | Sdc3 | Enriched in P8 retina | -1.280 | 0.000 | 1949.000 |
| ENSMUSG00000046287 | paraneoplastic antigen MA3 [Source:MGI Symbol;Acc:MGI:2180565] | Pnma3 | Enriched in P8 retina | -1.281 | 0.001 | 110.900 |
| ENSMUSG00000021892 | SH3-domain binding protein 5 (BTK-associated) [Source:MGI Symbol;Acc:MGI:1344391] | Sh3bp5 | Enriched in P8 retina | -1.282 | 0.000 | 251.200 |
| ENSMUSG00000047415 | G protein-coupled receptor 68 [Source:MGI Symbol;Acc:MGI:2441763] | Gpr68 | Enriched in P8 retina | -1.283 | 0.003 | 34.850 |
| ENSMUSG00000032017 | glutamate receptor, ionotropic, kainate 4 [Source:MGI Symbol;Acc:MGI:95817] | Grik4 | Enriched in P8 retina | -1.286 | 0.002 | 185.200 |

| | | | | | | |
|---|---|---|---|---|---|---|
| ENSMUSG00000035653 | leucine rich repeat and fibronectin type III domain containing 5 [Source:MGI Symbol;Acc:MGI:2144814] | Lrfn5 | Enriched in P8 retina | -1.286 | 0.003 | 564.400 |
| ENSMUSG00000011256 | a disintegrin and metallopeptidase domain 19 (meltrin beta) [Source:MGI Symbol;Acc:MGI:105377] | Adam19 | Enriched in P8 retina | -1.294 | 0.000 | 194.700 |
| ENSMUSG00000019966 | kit ligand [Source:MGI Symbol;Acc:MGI:96974] | Kitl | Enriched in P8 retina | -1.294 | 0.003 | 453.100 |
| ENSMUSG00000039579 | glutamate receptor ionotropic, NMDA3A [Source:MGI Symbol;Acc:MGI:1933206] | Grin3a | Enriched in P8 retina | -1.302 | 0.000 | 314.900 |
| ENSMUSG00000047013 | F-box protein 41 [Source:MGI Symbol;Acc:MGI:1261912] | Fbxo41 | Enriched in P8 retina | -1.305 | 0.000 | 562.600 |
| ENSMUSG00000021948 | protein kinase C, delta [Source:MGI Symbol;Acc:MGI:97598] | Prkcd | Enriched in P8 retina | -1.306 | 0.034 | 727.900 |
| ENSMUSG00000022421 | neuronal pentraxin receptor [Source:MGI Symbol;Acc:MGI:1920590] | Nptxr | Enriched in P8 retina | -1.306 | 0.001 | 1078.000 |
| ENSMUSG00000036620 | mannoside acetylglucosaminyltransferase 4, isoenzyme B [Source:MGI Symbol;Acc:MGI:2143974] | Mgat4b | Enriched in P8 retina | -1.320 | 0.000 | 283.600 |
| ENSMUSG00000022861 | diacylglycerol kinase, gamma [Source:MGI Symbol;Acc:MGI:105060] | Dgkg | Enriched in P8 retina | -1.334 | 0.007 | 135.000 |
| ENSMUSG00000020374 | RasGEF domain family, member 1C [Source:MGI Symbol;Acc:MGI:1921813] | Rasgef1c | Enriched in P8 retina | -1.335 | 0.001 | 39.330 |
| ENSMUSG00000022885 | beta galactoside alpha 2,6 sialyltransferase 1 [Source:MGI Symbol;Acc:MGI:108470] | St6gal1 | Enriched in P8 retina | -1.335 | 0.000 | 86.450 |
| ENSMUSG00000045174 | APC membrane recruitment 3 [Source:MGI Symbol;Acc:MGI:3026939] | Amer3 | Enriched in P8 retina | -1.343 | 0.000 | 95.690 |
| ENSMUSG00000001034 | mitogen-activated protein kinase 7 [Source:MGI Symbol;Acc:MGI:1346347] | Mapk7 | Enriched in P8 retina | -1.353 | 0.000 | 104.100 |
| ENSMUSG00000105867 | predicted gene 42517 [Source:MGI Symbol;Acc:MGI:5662654] | Gm42517 | Enriched in P8 retina | -1.356 | 0.001 | 89.610 |
| ENSMUSG00000046834 | keratin 1 [Source:MGI Symbol;Acc:MGI:96698] | Krt1 | Enriched in P8 retina | -1.361 | 0.019 | 16.620 |
| ENSMUSG00000034226 | ras homolog family member V [Source:MGI Symbol;Acc:MGI:2444227] | Rhov | Enriched in P8 retina | -1.362 | 0.003 | 54.650 |
| ENSMUSG00000030084 | plexin A1 [Source:MGI Symbol;Acc:MGI:107685] | Plxna1 | Enriched in P8 retina | -1.372 | 0.000 | 1259.000 |
| ENSMUSG00000073433 | Rho GDP dissociation inhibitor (GDI) gamma [Source:MGI Symbol;Acc:MGI:108430] | Arhgdig | Enriched in P8 retina | -1.376 | 0.000 | 185.100 |
| ENSMUSG00000031738 | Iroquois homeobox 6 [Source:MGI Symbol;Acc:MGI:1927642] | Irx6 | Enriched in P8 retina | -1.384 | 0.002 | 50.970 |
| ENSMUSG00000021143 | phosphofurin acidic cluster sorting protein 2 [Source:MGI Symbol;Acc:MGI:1924399] | Pacs2 | Enriched in P8 retina | -1.390 | 0.000 | 927.600 |
| ENSMUSG00000028755 | cytidine deaminase [Source:MGI Symbol;Acc:MGI:1919519] | Cda | Enriched in P8 retina | -1.390 | 0.017 | 14.430 |
| ENSMUSG00000069227 | G protein-regulated inducer of neurite outgrowth 1 [Source:MGI Symbol;Acc:MGI:1349455] | Gprin1 | Enriched in P8 retina | -1.392 | 0.000 | 480.700 |
| ENSMUSG00000021256 | vasohibin 1 [Source:MGI Symbol;Acc:MGI:2442543] | Vash1 | Enriched in P8 retina | -1.414 | 0.000 | 304.600 |
| ENSMUSG00000001333 | syncoilin [Source:MGI Symbol;Acc:MGI:1916078] | Sync | Enriched in P8 retina | -1.416 | 0.005 | 17.380 |
| ENSMUSG00000033854 | potassium channel, subfamily K, member 10 [Source:MGI Symbol;Acc:MGI:1919508] | Kcnk10 | Enriched in P8 retina | -1.416 | 0.000 | 191.600 |
| ENSMUSG00000035547 | calpain 5 [Source:MGI Symbol;Acc:MGI:1100859] | Capn5 | Enriched in P8 retina | -1.418 | 0.000 | 218.700 |
| ENSMUSG00000079523 | thymosin, beta 10 [Source:MGI Symbol;Acc:MGI:109146] | Tmsb10 | Enriched in P8 retina | -1.426 | 0.000 | 881.500 |
| ENSMUSG00000028680 | polo like kinase 3 [Source:MGI Symbol;Acc:MGI:109604] | Plk3 | Enriched in P8 retina | -1.428 | 0.000 | 41.320 |
| ENSMUSG00000030782 | transforming growth factor beta 1 induced transcript 1 [Source:MGI Symbol;Acc:MGI:102784] | Tgfb1i1 | Enriched in P8 retina | -1.438 | 0.000 | 24.560 |
| ENSMUSG00000037428 | VGF nerve growth factor inducible [Source:MGI Symbol;Acc:MGI:1343180] | Vgf | Enriched in P8 retina | -1.446 | 0.000 | 1249.000 |
| ENSMUSG00000002083 | BCL2 binding component 3 [Source:MGI Symbol;Acc:MGI:2181667] | Bbc3 | Enriched in P8 retina | -1.453 | 0.000 | 52.380 |
| ENSMUSG00000029101 | regulator of G-protein signaling 12 [Source:MGI Symbol;Acc:MGI:1918979] | Rgs12 | Enriched in P8 retina | -1.466 | 0.000 | 115.800 |
| ENSMUSG00000018507 | transient receptor potential cation channel, subfamily V, member 2 [Source:MGI Symbol;Acc:MGI:1341836] | Trpv2 | Enriched in P8 retina | -1.488 | 0.000 | 57.510 |
| ENSMUSG00000038295 | autophagy related 9B [Source:MGI Symbol;Acc:MGI:2685420] | Atg9b | Enriched in P8 retina | -1.491 | 0.040 | 13.880 |
| ENSMUSG00000031965 | T-box 20 [Source:MGI Symbol;Acc:MGI:1888496] | Tbx20 | Enriched in P8 retina | -1.499 | 0.029 | 45.450 |
| ENSMUSG00000117406 | netrin 3 [Source:MGI Symbol;Acc:MGI:1341188] | Ntn3 | Enriched in P8 retina | -1.506 | 0.001 | 52.980 |
| ENSMUSG00000036913 | tripartite motif-containing 67 [Source:MGI Symbol;Acc:MGI:3045323] | Trim67 | Enriched in P8 retina | -1.512 | 0.001 | 572.300 |
| ENSMUSG00000038094 | ATPase type 13A4 [Source:MGI Symbol;Acc:MGI:1924456] | Atp13a4 | Enriched in P8 retina | -1.515 | 0.044 | 39.550 |
| ENSMUSG00000044813 | src homology 2 domain-containing transforming protein B [Source:MGI Symbol;Acc:MGI:98294] | Shb | Enriched in P8 retina | -1.515 | 0.000 | 96.590 |
| ENSMUSG00000020230 | protein arginine N-methyltransferase 2 [Source:MGI Symbol;Acc:MGI:1316652] | Prmt2 | Enriched in P8 retina | -1.519 | 0.000 | 214.300 |
| ENSMUSG00000028039 | ephrin A3 [Source:MGI Symbol;Acc:MGI:106644] | Efna3 | Enriched in P8 retina | -1.531 | 0.000 | 71.320 |
| ENSMUSG00000046997 | splA/ryanodine receptor domain and SOCS box containing 4 [Source:MGI Symbol;Acc:MGI:2183445] | Spsb4 | Enriched in P8 retina | -1.531 | 0.000 | 36.700 |
| ENSMUSG00000044912 | synaptotagmin XVI [Source:MGI Symbol;Acc:MGI:2673872] | Syt16 | Enriched in P8 retina | -1.540 | 0.000 | 222.100 |
| ENSMUSG00000007944 | tetratricopeptide repeat domain 9B [Source:MGI Symbol;Acc:MGI:1920282] | Ttc9b | Enriched in P8 retina | -1.547 | 0.000 | 147.000 |
| ENSMUSG00000047963 | starch binding domain 1 [Source:MGI Symbol;Acc:MGI:1261768] | Stbd1 | Enriched in P8 retina | -1.560 | 0.007 | 7.985 |
| ENSMUSG00000024907 | galanin and GMAP prepropeptide [Source:MGI Symbol;Acc:MGI:95637] | Gal | Enriched in P8 retina | -1.567 | 0.012 | 30.000 |
| ENSMUSG00000049625 | TRAF-interacting protein with forkhead-associated domain, family member B [Source:MGI Symbol;Acc:MGI:2385852] | Tifab | Enriched in P8 retina | -1.578 | 0.007 | 7.994 |
| ENSMUSG00000007207 | syntaxin 1A (brain) [Source:MGI Symbol;Acc:MGI:109355] | Stx1a | Enriched in P8 retina | -1.579 | 0.000 | 278.700 |
| ENSMUSG00000066721 | zinc finger protein 575 [Source:MGI Symbol;Acc:MGI:2141921] | Zfp575 | Enriched in P8 retina | -1.590 | 0.000 | 26.990 |
| ENSMUSG00000094500 | small integral membrane protein 18 [Source:MGI Symbol;Acc:MGI:1919882] | Smim18 | Enriched in P8 retina | -1.601 | 0.028 | 4.253 |
| ENSMUSG00000031398 | plexin A3 [Source:MGI Symbol;Acc:MGI:107683] | Plxna3 | Enriched in P8 retina | -1.607 | 0.000 | 112.900 |
| ENSMUSG00000036882 | Rho GTPase activating protein 33 [Source:MGI Symbol;Acc:MGI:2673998] | Arhgap33 | Enriched in P8 retina | -1.620 | 0.000 | 502.500 |
| ENSMUSG00000018339 | glutathione peroxidase 3 [Source:MGI Symbol;Acc:MGI:105102] | Gpx3 | Enriched in P8 retina | -1.655 | 0.030 | 82.520 |
| ENSMUSG00000018411 | microtubule-associated protein tau [Source:MGI Symbol;Acc:MGI:97180] | Mapt | Enriched in P8 retina | -1.658 | 0.000 | 1946.000 |
| ENSMUSG00000026821 | ral guanine nucleotide dissociation stimulator [Source:MGI Symbol;Acc:MGI:107485] | Ralgds | Enriched in P8 retina | -1.670 | 0.000 | 543.000 |
| ENSMUSG00000043091 | tubulin, alpha 1C [Source:MGI Symbol;Acc:MGI:1095409] | Tuba1c | Enriched in P8 retina | -1.673 | 0.024 | 41.770 |
| ENSMUSG00000026220 | solute carrier family 16 (monocarboxylic acid transporters), member 14 [Source:MGI Symbol;Acc:MGI:1919031] | Slc16a14 | Enriched in P8 retina | -1.696 | 0.000 | 38.530 |
| ENSMUSG00000031659 | adenylate cyclase 7 [Source:MGI Symbol;Acc:MGI:102891] | Adcy7 | Enriched in P8 retina | -1.703 | 0.000 | 65.540 |
| ENSMUSG00000054555 | a disintegrin and metallopeptidase domain 12 (meltrin alpha) [Source:MGI Symbol;Acc:MGI:105378] | Adam12 | Enriched in P8 retina | -1.708 | 0.000 | 77.270 |
| ENSMUSG00000029603 | deltex 1, E3 ubiquitin ligase [Source:MGI Symbol;Acc:MGI:1352744] | Dtx1 | Enriched in P8 retina | -1.710 | 0.000 | 412.900 |
| ENSMUSG00000031737 | Iroquois homeobox 5 [Source:MGI Symbol;Acc:MGI:1859086] | Irx5 | Enriched in P8 retina | -1.717 | 0.000 | 31.090 |
| ENSMUSG00000019066 | RAB3D, member RAS oncogene family [Source:MGI Symbol;Acc:MGI:97844] | Rab3d | Enriched in P8 retina | -1.720 | 0.000 | 46.080 |
| ENSMUSG00000021278 | amnionless [Source:MGI Symbol;Acc:MGI:1934943] | Amn | Enriched in P8 retina | -1.728 | 0.006 | 16.050 |

| | | | | | | |
|---|---|---|---|---|---|---|
| ENSMUSG00000050271 | PEAK1 related kinase activating pseudokinase 1 [Source:MGI Symbol;Acc:MGI:1196223] | Prag1 | Enriched in P8 retina | -1.733 | 0.000 | 83.240 |
| ENSMUSG00000034685 | family with sequence similarity 171, member A2 [Source:MGI Symbol;Acc:MGI:2448496] | Fam171a2 | Enriched in P8 retina | -1.744 | 0.000 | 403.300 |
| ENSMUSG00000004098 | collagen, type V, alpha 3 [Source:MGI Symbol;Acc:MGI:1858212] | Col5a3 | Enriched in P8 retina | -1.747 | 0.001 | 67.350 |
| ENSMUSG00000002908 | potassium intermediate/small conductance calcium-activated channel, subfamily N, member 1 [Source:MGI Symbol;Acc:MGI:1933993] | Kcnn1 | Enriched in P8 retina | -1.778 | 0.000 | 72.470 |
| ENSMUSG00000018405 | mitochondrial rRNA methyltransferase 1 [Source:MGI Symbol;Acc:MGI:2443470] | Mrm1 | Enriched in P8 retina | -1.826 | 0.000 | 29.070 |
| ENSMUSG00000070644 | ethanolamine kinase 2 [Source:MGI Symbol;Acc:MGI:2443760] | Etnk2 | Enriched in P8 retina | -1.826 | 0.001 | 35.130 |
| ENSMUSG00000051652 | leucine rich repeat containing 3 [Source:MGI Symbol;Acc:MGI:2447899] | Lrrc3 | Enriched in P8 retina | -1.828 | 0.000 | 141.000 |
| ENSMUSG00000036377 | capping protein inhibiting regulator of actin [Source:MGI Symbol;Acc:MGI:2444817] | Cracd | Enriched in P8 retina | -1.858 | 0.000 | 1779.000 |
| ENSMUSG00000048070 | phosphoinositide-interacting regulator of transient receptor potential channels [Source:MGI Symbol;Acc:MGI:2443635] | Pirt | Enriched in P8 retina | -1.874 | 0.003 | 10.020 |
| ENSMUSG00000031934 | pannexin 1 [Source:MGI Symbol;Acc:MGI:1860055] | Panx1 | Enriched in P8 retina | -1.889 | 0.000 | 139.300 |
| ENSMUSG00000005357 | solute carrier family 1 (high affinity aspartate/glutamate transporter), member 6 [Source:MGI Symbol;Acc:MGI:1096331] | Slc1a6 | Enriched in P8 retina | -1.901 | 0.000 | 52.800 |
| ENSMUSG00000029581 | fascin actin-bundling protein 1 [Source:MGI Symbol;Acc:MGI:1352745] | Fscn1 | Enriched in P8 retina | -1.905 | 0.000 | 654.200 |
| ENSMUSG00000032446 | eomesodermin [Source:MGI Symbol;Acc:MGI:1201683] | Eomes | Enriched in P8 retina | -1.915 | 0.044 | 60.200 |
| ENSMUSG00000035835 | phospholipid phosphatase related 3 [Source:MGI Symbol;Acc:MGI:2388640] | Plppr3 | Enriched in P8 retina | -1.925 | 0.000 | 329.900 |
| ENSMUSG00000021303 | guanine nucleotide binding protein (G protein), gamma 4 [Source:MGI Symbol;Acc:MGI:102703] | Gng4 | Enriched in P8 retina | -1.933 | 0.000 | 229.600 |
| ENSMUSG00000110086 | small integral membrane protein 32 [Source:MGI Symbol;Acc:MGI:5791459] | Smim32 | Enriched in P8 retina | -1.934 | 0.026 | 24.370 |
| ENSMUSG00000018451 | RIKEN cDNA 6330403K07 gene [Source:MGI Symbol;Acc:MGI:1918001] | 6330403K07Rik | Enriched in P8 retina | -1.942 | 0.000 | 664.500 |
| ENSMUSG00000052301 | double C2, alpha [Source:MGI Symbol;Acc:MGI:109446] | Doc2a | Enriched in P8 retina | -1.943 | 0.001 | 56.860 |
| ENSMUSG00000055078 | gamma-aminobutyric acid (GABA) A receptor, subunit alpha 5 [Source:MGI Symbol;Acc:MGI:95617] | Gabra5 | Enriched in P8 retina | -1.949 | 0.000 | 60.370 |
| ENSMUSG00000054000 | tumor suppressor candidate 1 [Source:MGI Symbol;Acc:MGI:2684283] | Tusc1 | Enriched in P8 retina | -1.961 | 0.001 | 40.940 |
| ENSMUSG00000039976 | TBC1 domain family, member 16 [Source:MGI Symbol;Acc:MGI:2652878] | Tbc1d16 | Enriched in P8 retina | -1.965 | 0.000 | 1281.000 |
| ENSMUSG00000039313 | membrane integral NOTCH2 associated receptor 1 [Source:MGI Symbol;Acc:MGI:2667167] | Minar1 | Enriched in P8 retina | -1.975 | 0.001 | 60.240 |
| ENSMUSG00000059323 | tonsoku-like, DNA repair protein [Source:MGI Symbol;Acc:MGI:1919999] | Tonsl | Enriched in P8 retina | -2.005 | 0.000 | 30.180 |
| ENSMUSG00000034993 | vesicle amine transport 1 [Source:MGI Symbol;Acc:MGI:1349450] | Vat1 | Enriched in P8 retina | -2.057 | 0.000 | 429.800 |
| ENSMUSG00000030830 | integrin alpha L [Source:MGI Symbol;Acc:MGI:96606] | Itgal | Enriched in P8 retina | -2.067 | 0.003 | 6.323 |
| ENSMUSG00000102189 | predicted gene, 37194 [Source:MGI Symbol;Acc:MGI:5610422] | Gm37194 | Enriched in P8 retina | -2.069 | 0.010 | 32.700 |
| ENSMUSG00000022577 | lymphocyte antigen 6 complex, locus H [Source:MGI Symbol;Acc:MGI:1346030] | Ly6h | Enriched in P8 retina | -2.099 | 0.000 | 411.300 |
| ENSMUSG00000037705 | tectorin alpha [Source:MGI Symbol;Acc:MGI:109575] | Tecta | Enriched in P8 retina | -2.109 | 0.002 | 4.094 |
| ENSMUSG00000044469 | tumor necrosis factor, alpha-induced protein 8-like 1 [Source:MGI Symbol;Acc:MGI:1913693] | Tnfaip8l1 | Enriched in P8 retina | -2.111 | 0.000 | 9.788 |
| ENSMUSG00000032735 | actin binding LIM protein family, member 3 [Source:MGI Symbol;Acc:MGI:2442582] | Ablim3 | Enriched in P8 retina | -2.123 | 0.000 | 445.700 |
| ENSMUSG00000037362 | cellular communication network factor 3 [Source:MGI Symbol;Acc:MGI:109185] | Ccn3 | Enriched in P8 retina | -2.148 | 0.036 | 15.410 |
| ENSMUSG00000027669 | guanine nucleotide binding protein (G protein), beta 4 [Source:MGI Symbol;Acc:MGI:104581] | Gnb4 | Enriched in P8 retina | -2.163 | 0.000 | 103.700 |
| ENSMUSG00000034115 | sodium channel, voltage-gated, type XI, alpha [Source:MGI Symbol;Acc:MGI:1345149] | Scn11a | Enriched in P8 retina | -2.168 | 0.000 | 5.567 |
| ENSMUSG00000045731 | prepronociceptin [Source:MGI Symbol;Acc:MGI:105308] | Pnoc | Enriched in P8 retina | -2.170 | 0.000 | 50.560 |
| ENSMUSG00000035561 | aldehyde dehydrogenase 1 family, member B1 [Source:MGI Symbol;Acc:MGI:1919785] | Aldh1b1 | Enriched in P8 retina | -2.173 | 0.000 | 22.800 |
| ENSMUSG00000020090 | neuropeptide FF receptor 1 [Source:MGI Symbol;Acc:MGI:2685082] | Npffr1 | Enriched in P8 retina | -2.202 | 0.006 | 15.660 |
| ENSMUSG00000021070 | bradykinin receptor, beta 2 [Source:MGI Symbol;Acc:MGI:102845] | Bdkrb2 | Enriched in P8 retina | -2.212 | 0.000 | 6.429 |
| ENSMUSG00000043165 | loricrin [Source:MGI Symbol;Acc:MGI:96816] | Lor | Enriched in P8 retina | -2.212 | 0.015 | 17.170 |
| ENSMUSG00000030889 | von Willebrand factor A domain containing 3A [Source:MGI Symbol;Acc:MGI:3041229] | Vwa3a | Enriched in P8 retina | -2.219 | 0.019 | 26.650 |
| ENSMUSG00000015709 | aryl hydrocarbon receptor nuclear translocator 2 [Source:MGI Symbol;Acc:MGI:107188] | Arnt2 | Enriched in P8 retina | -2.220 | 0.000 | 540.300 |
| ENSMUSG00000078532 | Na+/K+ transporting ATPase interacting 1 [Source:MGI Symbol;Acc:MGI:1914399] | Nkain1 | Enriched in P8 retina | -2.223 | 0.000 | 207.800 |
| ENSMUSG00000031727 | polyamine modulated factor 1 binding protein 1 [Source:MGI Symbol;Acc:MGI:1930136] | Pmfbp1 | Enriched in P8 retina | -2.224 | 0.001 | 36.630 |
| ENSMUSG00000075334 | reprimo, TP53 dependent G2 arrest mediator candidate [Source:MGI Symbol;Acc:MGI:1915124] | Rprm | Enriched in P8 retina | -2.274 | 0.000 | 67.300 |
| ENSMUSG00000116652 | RIKEN cDNA B830017H08 gene [Source:MGI Symbol;Acc:MGI:3045365] | B830017H08Rik | Enriched in P8 retina | -2.292 | 0.006 | 15.950 |
| ENSMUSG00000029503 | purinergic receptor P2X, ligand-gated ion channel, 2 [Source:MGI Symbol;Acc:MGI:2665170] | P2rx2 | Enriched in P8 retina | -2.304 | 0.004 | 6.203 |
| ENSMUSG00000018012 | Rac family small GTPase 3 [Source:MGI Symbol;Acc:MGI:2180784] | Rac3 | Enriched in P8 retina | -2.308 | 0.000 | 122.200 |
| ENSMUSG00000090523 | glycophorin C [Source:MGI Symbol;Acc:MGI:1098566] | Gypc | Enriched in P8 retina | -2.320 | 0.002 | 5.436 |
| ENSMUSG00000032128 | roundabout guidance receptor 3 [Source:MGI Symbol;Acc:MGI:1343102] | Robo3 | Enriched in P8 retina | -2.328 | 0.002 | 9.898 |
| ENSMUSG00000044576 | GRB2 associated regulator of MAPK1 subtype 2 [Source:MGI Symbol;Acc:MGI:2685290] | Garem2 | Enriched in P8 retina | -2.365 | 0.000 | 62.670 |
| ENSMUSG00000006218 | family with sequence similarity 131, member C [Source:MGI Symbol;Acc:MGI:2685539] | Fam131c | Enriched in P8 retina | -2.449 | 0.000 | 42.580 |
| ENSMUSG00000041605 | innate immunity activator [Source:MGI Symbol;Acc:MGI:1921579] | Inava | Enriched in P8 retina | -2.451 | 0.039 | 19.290 |
| ENSMUSG00000028610 | DMRT-like family B with proline-rich C-terminal, 1 [Source:MGI Symbol;Acc:MGI:1927125] | Dmrtb1 | Enriched in P8 retina | -2.485 | 0.000 | 17.020 |
| ENSMUSG00000036198 | Rho GTPase activating protein 36 [Source:MGI Symbol;Acc:MGI:1922654] | Arhgap36 | Enriched in P8 retina | -2.583 | 0.006 | 77.620 |
| ENSMUSG00000023067 | cyclin-dependent kinase inhibitor 1A (P21) [Source:MGI Symbol;Acc:MGI:104556] | Cdkn1a | Enriched in P8 retina | -2.589 | 0.000 | 78.160 |
| ENSMUSG00000047261 | growth associated protein 43 [Source:MGI Symbol;Acc:MGI:95639] | Gap43 | Enriched in P8 retina | -2.638 | 0.000 | 876.700 |
| ENSMUSG00000029123 | serine/threonine kinase 32B [Source:MGI Symbol;Acc:MGI:1927552] | Stk32b | Enriched in P8 retina | -2.648 | 0.000 | 93.320 |
| ENSMUSG00000074923 | p21 (RAC1) activated kinase 6 [Source:MGI Symbol;Acc:MGI:2679420] | Pak6 | Enriched in P8 retina | -2.671 | 0.000 | 281.400 |
| ENSMUSG00000036856 | wingless-type MMTV integration site family, member 4 [Source:MGI Symbol;Acc:MGI:98957] | Wnt4 | Enriched in P8 retina | -2.723 | 0.000 | 80.140 |
| ENSMUSG00000030854 | protein tyrosine phosphatase, non-receptor type 5 [Source:MGI Symbol;Acc:MGI:97807] | Ptpn5 | Enriched in P8 retina | -3.033 | 0.000 | 407.000 |
| ENSMUSG00000025930 | musculin [Source:MGI Symbol;Acc:MGI:1333884] | Msc | Enriched in P8 retina | -3.116 | 0.002 | 4.237 |
| ENSMUSG00000026875 | TNF receptor-associated factor 1 [Source:MGI Symbol;Acc:MGI:101836] | Traf1 | Enriched in P8 retina | -3.231 | 0.000 | 10.430 |
| ENSMUSG00000029602 | RAS protein activator like 1 (GAP1 like) [Source:MGI Symbol;Acc:MGI:1330842] | Rasal1 | Enriched in P8 retina | -3.274 | 0.000 | 51.400 |
| ENSMUSG00000029005 | dorsal inhibitory axon guidance protein [Source:MGI Symbol;Acc:MGI:1917683] | Draxin | Enriched in P8 retina | -3.344 | 0.000 | 35.710 |

| | | | | | | |
|---|---|---|---|---|---|---|
| ENSMUSG00000020838 | solute carrier family 6 (neurotransmitter transporter, serotonin), member 4 [Source:MGI Symbol;Acc:MGI:96285] | Slc6a4 | Enriched in P8 retina | -4.394 | 0.000 | 23.260 |
| ENSMUSG00000034883 | leucine rich repeat protein 1 [Source:MGI Symbol;Acc:MGI:1916956] | Lrr1 | Enriched in P8 retina | -5.745 | 0.001 | 14.150 |

**Contrast numerator: P15_Het_SCN. Contrast denominator: P8_Het_SCN.**

| ensembl_gene_id | description | mgi_symbol | Significance | deseq_logfc | deseq_adjp | deseq_basemean |
|---|---|---|---|---|---|---|
| ENSMUSG00000025314 | protein tyrosine phosphatase, receptor type, J [Source:MGI Symbol;Acc:10457[illegible] | Ptprj | Enriched in P15 SCN | 4.696 | 0.000 | 998.900 |
| ENSMUSG00000022865 | coxsackie virus and adenovirus receptor [Source:MGI Symbol;Acc:MGI:1201679] | Cxadr | Enriched in P8 SCN | -1.275 | 0.033 | 403.700 |
| ENSMUSG00000024501 | dihydropyrimidinase-like 3 [Source:MGI Symbol;Acc:MGI:1349762] | Dpysl3 | Enriched in P8 SCN | -1.300 | 0.017 | 3490.000 |

**Contrast numerator: P15_KO_SCN. Contrast denominator: P8_KO_SCN.**

| ensembl_gene_id | description | mgi_symbol | Significance | deseq_logfc | deseq_adjp | deseq_basemean |
|---|---|---|---|---|---|---|
| ENSMUSG00000020455 | tripartite motif-containing 11 [Source:MGI Symbol;Acc:MGI:2137355] | Trim11 | Enriched in P8 SCN | -1.019 | 0.030 | 100.100 |
| ENSMUSG00000020108 | DNA-damage-inducible transcript 4 [Source:MGI Symbol;Acc:MGI:1921997] | Ddit4 | Enriched in P8 SCN | -1.021 | 0.008 | 181.800 |
| ENSMUSG00000026558 | uridine-cytidine kinase 2 [Source:MGI Symbol;Acc:MGI:1931744] | Uck2 | Enriched in P8 SCN | -1.064 | 0.000 | 349.000 |
| ENSMUSG00000042606 | HIRA interacting protein 3 [Source:MGI Symbol;Acc:MGI:2142364] | Hirip3 | Enriched in P8 SCN | -1.075 | 0.013 | 132.700 |
| ENSMUSG00000029761 | caldesmon 1 [Source:MGI Symbol;Acc:MGI:88250] | Cald1 | Enriched in P8 SCN | -1.110 | 0.000 | 209.000 |
| ENSMUSG00000029346 | SRR1 domain containing [Source:MGI Symbol;Acc:MGI:1917368] | Srrd | Enriched in P8 SCN | -1.258 | 0.002 | 45.970 |
| ENSMUSG00000036904 | frizzled class receptor 8 [Source:MGI Symbol;Acc:MGI:108460] | Fzd8 | Enriched in P8 SCN | -1.278 | 0.001 | 88.540 |
| ENSMUSG00000024937 | EH domain binding protein 1-like 1 [Source:MGI Symbol;Acc:MGI:3612340] | Ehbp1l1 | Enriched in P8 SCN | -1.351 | 0.000 | 192.900 |
| ENSMUSG00000029442 | WD repeat domain 66 [Source:MGI Symbol;Acc:MGI:1918495] | Wdr66 | Enriched in P8 SCN | -1.435 | 0.002 | 54.470 |
| ENSMUSG00000010342 | testis expressed gene 14 [Source:MGI Symbol;Acc:MGI:1933227] | Tex14 | Enriched in P8 SCN | -1.537 | 0.032 | 109.900 |
| ENSMUSG00000020107 | anaphase promoting complex subunit 16 [Source:MGI Symbol;Acc:MGI:1289325] | Anapc16 | Enriched in P8 SCN | -1.599 | 0.000 | 134.300 |
| ENSMUSG00000040952 | ribosomal protein S19 [Source:MGI Symbol;Acc:MGI:1333780] | Rps19 | Enriched in P8 SCN | -1.671 | 0.000 | 155.700 |
| ENSMUSG00000029245 | Eph receptor A5 [Source:MGI Symbol;Acc:MGI:99654] | Epha5 | Enriched in P8 SCN | -1.750 | 0.000 | 636.400 |
| ENSMUSG00000044533 | ribosomal protein S2 [Source:MGI Symbol;Acc:MGI:105110] | Rps2 | Enriched in P8 SCN | -1.824 | 0.000 | 542.000 |
| ENSMUSG00000117694 | small nucleolar RNA host gene 4 [Source:MGI Symbol;Acc:MGI:4937091] | Snhg4 | Enriched in P8 SCN | -1.968 | 0.000 | 124.900 |
| ENSMUSG00000000560 | gamma-aminobutyric acid (GABA) A receptor, subunit alpha 2 [Source:MGI Symbol;Acc:MGI:95614] | Gabra2 | Enriched in P8 SCN | -2.079 | 0.000 | 394.200 |
| ENSMUSG00000050677 | coiled-coil domain containing 96 [Source:MGI Symbol;Acc:MGI:1913967] | Ccdc96 | Enriched in P8 SCN | -2.135 | 0.002 | 66.050 |
| ENSMUSG00000051817 | SRY (sex determining region Y)-box 12 [Source:MGI Symbol;Acc:MGI:98360] | Sox12 | Enriched in P8 SCN | -2.355 | 0.000 | 619.900 |
| ENSMUSG00000052428 | transmembrane and coiled-coil domains 1 [Source:MGI Symbol;Acc:MGI:1921173] | Tmco1 | Enriched in P8 SCN | -2.481 | 0.000 | 95.800 |
| ENSMUSG00000039115 | integrin alpha 9 [Source:MGI Symbol;Acc:MGI:104756] | Itga9 | Enriched in P8 SCN | -2.526 | 0.016 | 1811.000 |
| ENSMUSG00000061462 | obscurin, cytoskeletal calmodulin and titin-interacting RhoGEF [Source:MGI Symbol;Acc:MGI:2681862 | Obscn | Enriched in P8 SCN | -2.880 | 0.000 | 66.020 |
